# Identification of RNAs bound by Hfq reveals widespread RNA partners and a sporulation regulator in the human pathogen *Clostridioides difficile*

**DOI:** 10.1101/2020.11.25.398065

**Authors:** Pierre Boudry, Emma Piattelli, Emilie Drouineau, Johann Peltier, Anaïs Boutserin, Maxence Lejars, Eliane Hajnsdorf, Marc Monot, Bruno Dupuy, Isabelle Martin-Verstraete, Daniel Gautheret, Claire Toffano-Nioche, Olga Soutourina

## Abstract

Noncoding RNAs (ncRNA) have emerged as important components of regulatory networks governing bacterial physiology and virulence. Previous deep-sequencing analysis identified a large diversity of ncRNAs in the human enteropathogen *Clostridioides (Clostridium) difficile.* Some of them are *trans*-encoded RNAs that could require the RNA chaperone protein Hfq for their action. Recent analysis suggested a pleiotropic role of Hfq in *C. difficile* with the most pronounced effect on sporulation, a key process during the infectious cycle of this pathogen. However, a global view of RNAs interacting with *C. difficile* Hfq is missing. In the present study, we performed RNA immunoprecipitation high-throughput sequencing (RIP-Seq) to identify Hfq-associated RNAs in *C. difficile*. Our work revealed a large set of Hfq-interacting mRNAs and ncRNAs, including mRNA leaders and coding regions, known and potential new ncRNAs. In addition to *trans*-encoded RNAs, new categories of Hfq ligands were found including *cis*-antisense RNAs, riboswitches and CRISPR RNAs. ncRNA-mRNA and ncRNA-ncRNA pairings were postulated through computational predictions. Investigation of one of the Hfq-associated ncRNAs, RCd1, suggests that this RNA contributes to the control of late stages of sporulation in *C. difficile*. Altogether, these data provide essential molecular basis for further studies of post-transcriptional regulatory network in this enteropathogen.

## INTRODUCTION

*Clostridioides difficile* (formerly *Clostridium difficile*) has emerged as an important human enteropathogen. This Gram-positive anaerobic spore-forming bacterium is considered as the main cause of health-care-associated diarrhea worldwide (1,2). Antibiotics exposure, age over 65 years and immune deficiency have been identified as major risk factors of *C. difficile* infection (CDI). With an increase in incidence rate, severity of infection forms, occurrence of community-acquired cases and emergence of hypervirulent strains, *C. difficile* infections are becoming a key public health concern accentuated by the general aging of population in industrial countries. Normal gut microbiota protects healthy adults from CDI, however, antibiotic treatments induce a colonic dysbiosis state that subsequently facilitates the development of this pathogen from acquired or preexisting spores in healthy carriers (2,3). In addition to two major toxins, TcdA and TcdB, causing alterations in the actin cytoskeleton of intestinal epithelial cells, several virulence and colonization factors have been identified to contribute to CDI (4–7). However, many aspects controlling the infection cycle of this emerging pathogen still remain poorly understood. Among the most challenging traits of CDI are high recurrence rates that could be associated at least in part with the ability to form highly resistant spores (6,8). Additional mechanisms contributing to the successful adaptation of this pathogen inside the host remain to be explored. A better understanding of regulatory processes governing CDI cycle and pathogenesis is urgently needed to put forward new ideas for alternative therapeutic and diagnostic strategies.

Regulatory RNAs have been identified in all living organisms as key players for major adaptive responses and are known to control virulence in major pathogens (9–13). Our recent data strongly suggest the importance of RNA-based mechanisms for the control of gene expression in *C. difficile* (14,15). By RNA-seq and differential RNA-seq we detected more than 200 regulatory RNAs including potential *trans*-encoded small regulatory RNAs (sRNA) located in intergenic regions (IGR), *cis*-antisense RNAs and riboswitches (14,15). We also detected abundant small RNAs encoded by *C. difficile* CRISPR (clustered regularly interspaced short palindromic repeats)-Cas (CRISPR associated sequences) system, that might be important for *C. difficile* survival in bacteriophage-rich gut communities and also for the control of genetic exchanges favoured within gut microbiota (15–17). We recently described type I toxin-antitoxin (TA) modules adjacent to CRISPR arrays with the first functional antisense RNAs in this pathogen (18). A total of 13 potential type I TA modules are present in the genome of *C. difficile* reference strain 630 that could also contribute to its fitness inside the host (19,20). All these studies suggest that the regulatory RNA network controlling *C. difficile* pathophysiology is in many aspects unique and deserves further special attention (14).

Most characterized sRNAs act by base-pairing with their target mRNAs. These interactions usually assisted by RNA chaperone proteins like Hfq for *trans*-encoded riboregulators lead to modulation of mRNA translation and/or stability (21–23). Hfq is an abundant bacterial RNA-binding protein closely related to the eukaryotic and archaeal Sm and Sm-like protein families forming homohexameric structures (24). Together with the recently identified ProQ, Hfq is a major RNA chaperone well-studied for its implication in the sRNA-mediated regulatory mechanisms, in particular, in Gram-negative bacteria (25,26). In these bacteria, Hfq facilitates the short and imperfect base-pairing interactions of *trans*-encoded sRNAs with their target mRNAs and thus has an important role in stress response and virulence of pathogenic species. Nevertheless, the role of Hfq in Gram-positive bacteria still remains largely questionable. The *C. difficile* genome encodes an Hfq homologue with several unique features among Gram-positive bacteria. This includes the conservation of key amino acid residues for RNA binding surfaces and the presence of a particular C-terminal region. *C. difficile* Hfq depletion has pleiotropic effects on bacterial physiology with most pronounced impact on sporulation efficiency (27,28). We recently demonstrated that *C. difficile* Hfq could be involved in the control of metabolic adaptation, biofilm formation, stress responses and sporulation (27). This is in accordance with large transcriptome changes and altered accumulation of several sRNAs upon Hfq depletion.

We have previously identified nearly 100 potential *trans*-acting regulatory RNAs in the IGR of *C. difficile* genome that could require the RNA chaperone protein Hfq for their action (15). However, a global view of Hfq-dependent regulatory networks in this species is still lacking. The development of new generation sequencing approaches provides powerful strategies for studying molecular interactions in the cells at the genome level. Here, we used a co-immunoprecipitation coupled to RNA-seq approach (RIP-seq: RNA immunoprecipitation-sequencing) to establish the first comprehensive map of the Hfq-associated RNAs in this important enteropathogen and *in silico* prediction to further associate ncRNA and target mRNA pairs. We found that *C. difficile* Hfq associates with different classes of RNAs, including mRNAs, 5’UTRs (untranslated regions), known regulatory RNAs and potential new sRNAs in IGRs. In particular, enriched RNA molecules include *trans*-encoded sRNAs but also *cis*-antisense RNAs, the toxin mRNA and antitoxin RNAs from all recently identified type I TA modules, a total of twelve CRISPR arrays and numerous riboswitches. The role of RCd1, an sRNA associated with Hfq *in vivo* and *in vitro,* has been analyzed. Our results suggest that this Hfq-associated sRNA contributes to the control of late stages of sporulation in *C. difficile*. Altogether, these data provide an essential basis for establishing links between ncRNAs and the phenotypes associated to Hfq in *C. difficile* and constitute a valuable resource for further exploration of post-transcriptional regulatory networks in this emerging enteropathogen.

## MATERIAL AND METHODS

### Plasmid and bacterial strain construction and growth conditions

*C. difficile* and *E. coli* strains and plasmids used in this study are presented in Supplementary Table S1. *C. difficile* strains were grown anaerobically (5 % H_2_, 5 % CO_2_, and 90 % N_2_) in TY (29) or Brain Heart Infusion (BHI, Difco) media in an anaerobic chamber (Jacomex). When necessary, cefoxitin (Cfx; 25 μg/ml) and thiamphenicol (Tm; 15 μg/ml) were added to *C. difficile* cultures. *E. coli* strains were grown in LB broth (30), and when needed, ampicillin (100 μg/ml) or chloramphenicol (15 μg/ml) was added to the culture medium. The non-antibiotic analog anhydrotetracycline (ATc) was used for induction of the *P*_*tet*_ promoter of pRPF185 vector derivatives in the *C. difficile* (31). Strains carrying pRPF185 derivatives were generally grown in TY medium in the presence of 250 ng.ml-1 ATc and 7.5 μg.ml-1 Tm for 7.5 h and the induction of gene expression was confirmed by qRT-PCR. The number of spores was estimated in sporulation medium (SM) (32) or BHIS medium (27) with at least three biological replicates (33). After 24 and 72 h of growth, 1 ml of culture was divided into two samples. To determine the total number of CFU, the first sample was serially diluted and plated on BHIS medium supplemented with 0.1% taurocholate. To determine the number of spores, the vegetative cells of the second sample were heat-killed by incubation for 25 min at 65 °C prior to plating on BHIS medium supplemented with 0.1% taurocholate.

All routine plasmid constructions were carried out using standard procedures (34). All primers used in this study are listed in Supplementary Table S1.

For RIP-seq experiments, the 3xFLAG-tag sequence was introduced to the C-terminal part of *hfq* gene as a translational fusion on pRFP185-derivative plasmid. The expression of 3xFLAG tagged Hfq protein was confirmed by dot blot analysis with anti-FLAG antibodies. As a positive control of CD1234 protein detection by Western blot, the 3xFLAG-tag sequence was introduced to the C-terminal part of *CD1234* gene on pRFP185-derivative plasmid.

To introduce the 3xFLAG-tag at the C-terminal part of the coding region of *CD1234* gene, directly upstream the stop codon on the *C. difficile* chromosome, we used an allelic-coupled exchange technique based on a semi-suicidal plasmid vector carrying *E. coli* cytosine deaminase *codA* gene as counter-selection marker (35) that was improved by using an inducible toxin expression from type I toxin-antitoxin module instead of *codA* as counter-selection marker (19). We used Gibson assembly to construct a plasmid for further conjugation and homologous recombination in *C. difficile*. The resulting derivative plasmid was transformed into the *E. coli* HB101 (RP4) and subsequently mated with *C. difficile* 630Δ*erm*. *C. difficile* transconjugants were selected by sub-culturing on BHI agar containing Tm (15 μg/ml) and Cfx (25 μg/ml).

### RNA extraction, quantitative real time PCR, Northern blot

Total RNA was isolated from *C. difficile* strains grown in TY or SM medium using Trizol (Sigma) or FastRNA PRO BLUE kit (MP Biomedicals) (see (36) for detailed protocol description). The cDNA synthesis by reverse transcription and quantitative real-time PCR analysis were performed as detailed in (36) using standard protocols. No template and no reverse transcriptase controls were systematically included in each experiment. In each sample, the relative expression for a gene was calculated relatively to the 16S rRNA gene or *dnaF* gene (*CD1305*) encoding DNA polymerase III. The relative change in gene expression was recorded as the ratio of normalized target concentrations (ΔΔCt) from at least three independent experiments (37). For Northern blot analysis, 5 μg of total RNA was separated on a denaturing 6% or 8% polyacrylamide gel containing 8 M urea, and transferred to Hybond-N+ membrane (Amersham) by electroblotting using the Trans-blot cell from Bio-Rad in 1x TBE buffer (89 mM Tris-base, 89 mM boric acid and 2 mM EDTA). Following UV-cross-linking of the samples to the membrane, prehybridization was carried out for 2 h at 42°C in 7 mL of prehybridization buffer ULTRAHyb (Ambion). Hybridization was performed overnight at 42°C in the same buffer in the presence of a [γ-^32^P]-labeled DNA oligonucleotide probe. Alternatively, the probe was synthesized using PCR with 5’end-labeled primer complementary to RNA sequence. After hybridization, membranes were washed twice for 5 min in 50 mL 2X SSC (300 mM sodium chloride and 30 mM sodium citrate) 0.1% sodium dodecyl sulphate (SDS) buffer and twice for 15 min in 50 mL 0.1X SSC 0.1% SDS buffer. Radioactive signal was detected with a Typhoon system (Amersham). The size of the transcripts was estimated by comparison with RNA molecular weight standards (Invitrogen). The relative intensities of the bands from Northern blot analysis via autoradiography were quantified using ImageJ software.

### RIP-seq experiment

RIP-seq experiment has been performed with four biological replicates as described previously with some modifications (38). Briefly, Hfq::3xFLAG was expressed during 4.5 h by addition of 250 ng/ml ATc inducer after 3.5 h of culture in TY medium. Cells were suspended in the lysis buffer (50 mM HEPES–KOH pH 7.5, 150 mM NaCl, 1 mM EDTA, 1% Triton X-100, 0.1% Na-deoxycholate, protease inhibitor) supplemented with 0.1% SDS and 10% glycerol and lysed using a FastPrep (2 cycles of 45 sec at 6.5 m.s^−1^). Cell debris were removed by centrifugation (16.000 g for 10 min). Total proteins were quantified with Bradford, and the concentration of protein extract was adjusted to 1 mg.ml^−1^. For 3xFLAG co-immunoprecipitation, 600 μl of magnetic beads coated with 3xFLAG antibodies (“Sigma” Anti-FLAG® M2 Magnetic Beads) were added for each sample and incubated during 1.5 h at 4°C on a rotating platform. For Hfq co-immunoprecipitation, 20 μl of serum from rabbit immunized against Hfq (27) was added and pre-immune serum serves as a negative control. Hfq was trapped with magnetic beads coated with antibodies against rabbit IgG. Using a magnet, beads were washed five times with cold lysis buffer containing additional 350 mM NaCl. Beads was suspended twice in 500 μl and 200 μl of elution buffer (50 mM Tris, 1 mM EDTA, 1% SDS, pH 8.0 supplemented with 10% of glycerol) and incubated for 15 min at 65°C. A conventional acid phenol-chloroform extraction and ethanol precipitation were performed for the preparation of nucleic acids, followed by DNase treatment. The RNA was then processed into cDNA libraries using Illumina’s TruSeq Stranded Total RNA Sample Prep Kit according to manufacturer’s instructions with an adaptation of DNA/bead ratio of 1.8 to favor small fragments and sequenced using an Illumina HiSeq 2500 sequencer over 50 cycles (50 bp single end sequencing).

### Western blot

SDS-PAGE and Western immunoblotting were carried out using standard methods in triplicate (27). Briefly, the strains were grown in SM medium for 12 h in the presence of 250 ng/ml ATc and 7.5 μg/ml Tm. 60 μg or 10 μg of whole cell lysate were loaded on the gels. The lysate of the strain overexpressing CD1234-FLAG from the plasmid served as a positive control for detection with anti-FLAG antibodies. Proteins were separated on 4-20% Bis-Tris polyacrylamide gels in Tris-Glycine SDS buffer. Immunoblotting with anti-SigA antibodies or Coomassie gel staining served as a loading control. The Hfq protein levels have been verified by Western blotting in triplicate with polyclonal anti-Hfq antibodies (27). The strains were grown in TY medium in the presence of 250 ng/ml ATc inducer and whole cell extracts were loaded on a 4-12% gradient Bis-Tris polyacrylamide gel with MES running buffer. ImageJ software was used to estimate the protein levels.

### RNA band-shift assay

Templates for the synthesis of RNA probes were obtained by PCR amplification using the Term and T7 oligonucleotides described in Supplementary Table S1. RNA synthesis, purification, concentration monitoring, 5’-labelling and Cd-HfqHis6 protein purification were performed as described in (27,28). The RNA-binding capacity of purified Hfq was evaluated by mobility-shift assay using RCd1, *spoIIID* and *CD1234*. Band-shift assays were performed as previously described (39). Briefly, RNAs were synthesized by T7 RNA polymerase and gel purified. Radiolabeled RNAs were incubated with increasing concentrations of Hfq for 10 min at 37°C in 10 μl of 100 mM NaCl/10 mM Tris-HCl (pH 8.1)/0.25% glycerol/0.5 mM DTT/1 mM EDTA/0.06% Triton X-100, and complexes were separated on polyacrylamide gels. The radioactivity level corresponding to the bound fraction of RNA was plotted versus Hfq concentrations expressed on the basis of the monomer form. The data were fitted to the Hill equation, and half-saturation values (K_1/2_) were estimated using Kaleidagraph for each RNA. When RCd1 was incubated with *spoIIID* or *CD1234,* each RNA was individually incubated in water at 70°C for 1 min and then for 5 min at room temperature. RCd1 was immediately mixed with SpoIIID or CD1234 transcripts or the equivalent amount of yeast RNA, in the presence or the absence of Hfq, in binding buffer and samples were incubated at 20°C for 20 min before loading on native polyacrylamide gels.

### *In silico* analysis description

#### RIP-seq data analysis

After adaptors trimming using AlienTrimmer (40), sequencing reads were aligned to *C. difficile* strain 630 genome (NC_009089.1 region of the GCF_000009205.2 assembly and downloaded from NCBI) with the Bowtie2 software (41) and using default parameters. RIP-seq coverage visualizations are available through the COV2HTML software (42) at : https://mmonot.eu/COV2HTML/visualisation.php?str_id=-75000032 (FLAG vs Control) https://mmonot.eu/COV2HTML/visualisation.php?str_id=-75000034 (Hfq vs PI).

#### Peak detection with MACS2

By retaining the sequencing orientation, we determined the Hfq binding regions from the strand-separated mapping data. We also excluded mapped reads on rRNA and tRNA regions because they are heavily covered and thus bias the peak-calling procedure. We applied MACS2 tool (43) thanks to the docker version (kathrinklee/macs2) with the following settings: the 4 3XFlag, and 4 control replicas for treatment and control (-t and -c options) respectively, the effective genome size (-g) at 4232334 and estimated by the percentage of bases of the genome masked by the repeatmasker tool *http://repeatmasker.org,* the --broad option adapted to our RIP-seq experiment, --nomodel option, and the --ext_size to 51 option. We doubled those MACS2 runs for wild type strain experiment but with anti-Hfq-antibodies (Hfq630E) and pre-immune serum (PI630E) as couple of treatment and control inputs respectively (same settings). The significant peaks given by MACS2 were then compared to annotation provided by the MaGe plateforme (44) (chromosome NC_009089.1 of the *Clostridium difficile* 630 genome) with the intersect tool (-wao options) of the bedtools suite (45). Results of this analysis with statistically significant enrichment of peaks are displayed in Supplementary Table S3.

#### Functional clustering with DAVID from KEGG DB

The functional categories of genes were annotated according to the KEGG database (46) based on the DAVID annotation (47) and general categories were assigned with manual curation.

#### Motifs detection with MEME

With the goal of discovering sequence motifs involved in the binding of Hfq to RNA, we applied software tools from the MEME suite (48) thanks to the docker version (ddiez/meme, v.4.11.1) on different pools of sequences corresponding to: the peaks, the 5’ region of CDS overlapped by a peak (+/− 100 bp from the translation start site), and the sRNA overlapped by a peak excluding those with multiple copies in the genome or with repeat motifs (CRISPR RNA, TA, c-di-GMP-dependent riboswitches). The fasta formatted sequences need as inputs for MEME tools were extracted from the chromosome sequence with the faidx tool of the samtools suite (49) following the region locations and their strand. The MEME background model was determined with the fasta-get-markov tool (-m 6 -dna -norc) of the MEME suite. The motif discovery was realized with the meme tool (-mod anr -dna -nmotifs 50, MEME suite) restricting the space of search with the expected motif size corresponding to an RNA-RNA interaction (-minw 4 -maxw 10) or to an Hfq-RNA interaction (-minw 6 -maxw 30).

#### Prediction of ncRNA-target mRNA interactions with IntaRNA

The predictions of ncRNA-ncRNA interaction or ncRNA-mRNA interaction have been made with the IntaRNA tool (50) thanks to the docker version (quay.io/biocontainers/intarna:2.0.4—boost1.61_1, detailled mode output option -D, reporting only one optimal interaction, -n 0). Except for the peak pool, we used the same pools of sequences as for motifs detection as entries for IntaRNA. We set the interaction energy μG cutoff to −10 kcal/mol for IntaRNA output presentation. RNAPredator (http://rna.tbi.univie.ac.at/cgi-bin/RNApredator/target_search.cgi) and RNADuplex prediction algorithms available online were also applied for RCd1 target predictions (51,52).

## RESULTS

### Co-immunoprecipitation analysis of Hfq-binding RNAs *in vivo* in *C. difficile*

In order to identify all RNAs interacting with Hfq, we performed an Hfq co-immunoprecipitation RIP-seq experiment followed by high throughput sequencing of RNAs co-purified with Hfq. For this purpose we have constructed a strain carrying the plasmid pRPF185 with the *hfq* gene in translational fusion with the 3xFLAG tag at the C-terminal end of the protein (pDIA6151) (Supplementary Table S1). This strain named CDIP303 was grown together with the control strain carrying an empty plasmid in TY broth until the late exponential growth phase (8 h of growth). The level of over-expressed FLAG-tagged Hfq was estimated to be at least 10-fold higher as compared to the endogenous Hfq level suggesting that the majority of Hfq proteins in the cell were FLAG-tagged under tested conditions (Supplementary Figure S1). We then immunoprecipitated Hfq-FLAG with an anti-FLAG antibody and purified associated RNAs. The strain carrying an empty plasmid lacking the 3xFLAG construct was used as a negative control for non-specific binding to anti-FLAG matrix. To assess the impact of 3xFLAG tag on Hfq interactions with RNAs, a control experiment was performed in parallel with the wild-type strain 630*Δerm*. From the cultures grown 8 h, we immunoprecipitated native Hfq with specific polyclonal antibodies previously used for Western blot detection of the protein (27). Pre-immune serum was used for the control sample. Generally, a lower specific signal strength was observed by this approach. The insufficient quality of polyclonal anti-Hfq antibodies led us to use 3x-FLAG tagged Hfq for this experiment. The untagged Hfq co-immunoprecipitation experiment only serves as an additional control and all further analyses were performed on replicates of Hfq-FLAG-tagged samples. The results from Hfq-untagged samples are shown in IGV visualization to illustrate the consistency between the two experiments for selected RNA partners. The results from Hfq-FLAG-tagged samples in four biological replicates were considered to establish a final list of Hfq ligands.

For each sample, between 27 and 72 million sequence reads were obtained and 97.7 to 99.1 % of reads mapped to the *C. difficile* strain 630 genome (Supplementary Table S2). All four biological replicates clustered together for Hfq-FLAG and control samples, respectively, as shown by Principal Component Analysis (Supplementary Figure S2). Sequence reads were then classified according to the strain 630 genome annotation at their mapping sites (Figure 1). For a small part of reads named ambiguous, no particular category could be assigned. In accordance with data reported for other bacterial RNA chaperones (53,54), the majority (97.5%) of sequence reads in control samples corresponded to non-specifically recovered abundant house-keeping rRNAs and tRNAs. In contrast, about 60% of sequence reads mapped to rRNAs and tRNAs regions for Hfq-FLAG samples, but a significant enrichment was observed for sRNAs, mRNA coding regions (CDS) and new IGR regions (Figure 1). Similar results were obtained for untagged wild-type strain (Supplementary Figure S3) with 64% of sequence reads from rRNAs and tRNAs regions versus 86% for control samples, and the remaining reads being specifically enriched for CDS and sRNAs.

**Figure 1.**
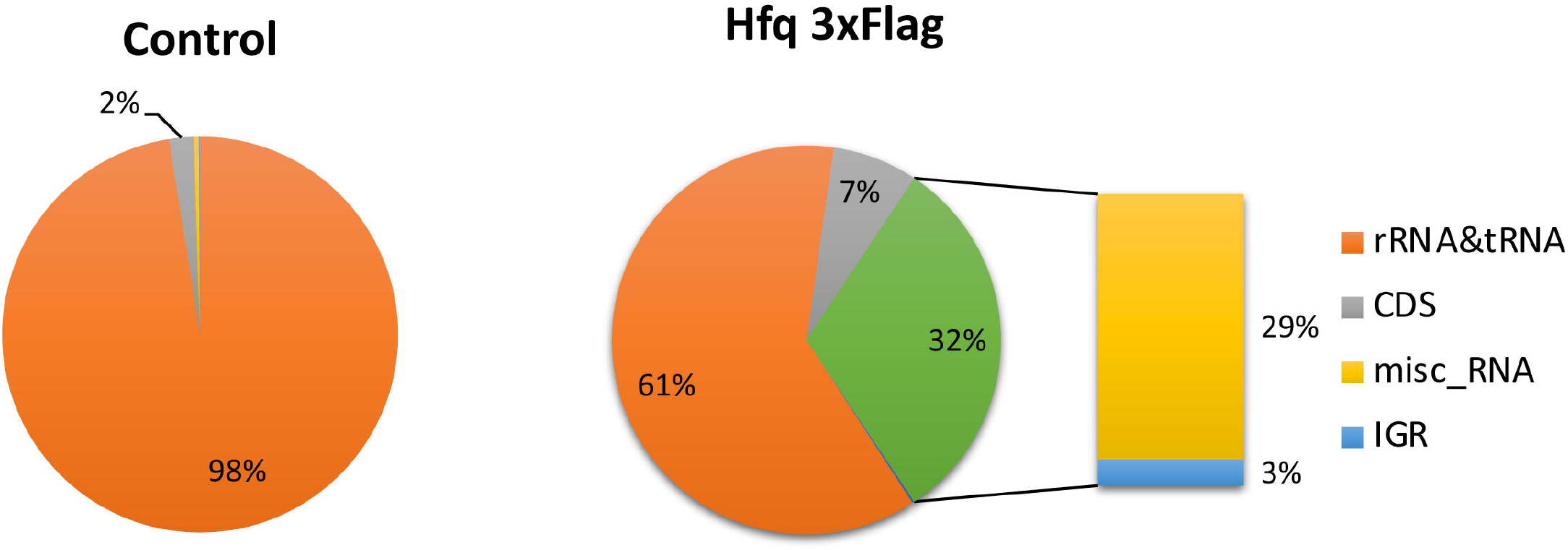
Diagram of relative proportion of different RNA species in Hfq-FLAG coIP and in control sample. All sequences that mapped to the *C. difficile* genome are represented. The rRNA and tRNA abundant housekeeping RNAs are shown in orange, the reads mapping to CDS are shown in gray. The relative proportion of other RNA species including known previously identified regulatory RNAs (named « misc_RNA ») and IGRs for Hfq-3xFLAG coIP samples is detailed in the diagram on the right. Left panel: control coIP, right panel: Hfq 3xFLAG coIP. RIP-seq experiment has been performed with four biological replicates.

### Identification of RNA peaks specifically enriched in Hfq-immunoprecipitated samples

The MACS2 program was used for the analysis of sequencing data to identify peaks of reads that were significantly enriched in the Hfq-FLAG samples as compared to a control lacking FLAG-tag in an independent manner from existing gene annotation (55,56). A similar analysis was also performed for the wild type strain experiment with anti-Hfq-antibodies and pre-immune serum as a control. Minimum peak size was set to 51 nt for MACS2 peak calling, corresponding to the size of sequencing reads. The comparative analysis of Hfq-coIP and control samples revealed that the reads corresponding to housekeeping tRNA and rRNA were overrepresented among differentially detected reads and were thus eliminated from further peak calling procedure. Peaks were then mapped to the annotated genome of *C. difficile* strain 630 available on the MaGe platform (44) and their location visualized using Integrative Genomics Viewer (IGV) program (57). Representative examples of different RNA functional groups are presented in Figure 2. This analysis identified 808 peaks overlapping with the regions of significant enrichments of reads including 634 peaks for CDS regions and 178 peaks for ncRNAs: 97 known regulatory RNAs of different functional classes and 68 potentially new regulatory RNAs or UTR regions (Supplementary Table S3). Hfq-enriched peaks were categorized into the following groups: mRNAs, sRNAs, antisense RNAs and 5’ regulatory elements. Among these peaks *trans*-encoded sRNAs (Figure 2A) but also *cis*-antisense RNAs (Figure 2B-C), the toxin mRNA and antitoxin RNAs from all 13 recently identified type I TA modules (Figure 2B), a total of twelve CRISPR arrays (Figure 2D) and numerous riboswitches (Figure 2E) were significantly enriched.

**Figure 2.**
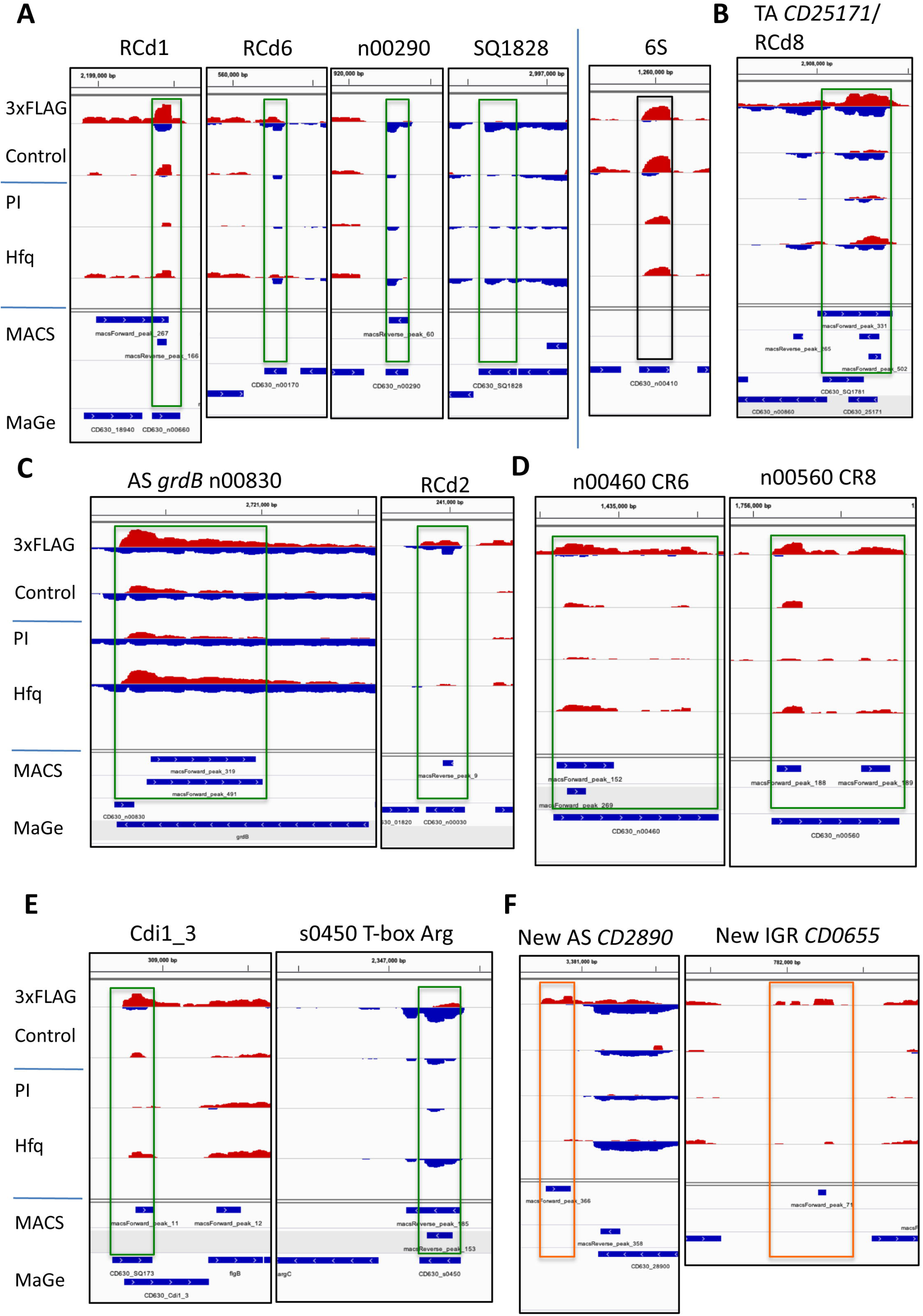
Visualization of RIP-seq data for regulatory RNAs with IGV. Representative examples from different functional groups of RNAs are presented in A) for sRNAs, B) for TA loci, C) for other antisense RNAs, D) for CRISPR RNAs, E) for riboswitches, F) for MACS peaks without annotation for new ncRNAs in IGR or new antisense RNAs. The results for “3xFLAG” Hfq-coIP sample are compared to the “Control” sample, while the “Hfq” represents Hfq-coIP sample compared to the “PI” pre-immune serum control. The genomic regions for previously identified sRNAs are presented in a green box, newly identified transcribed region are shown in orange box. 6S RNA (an Hfq-independent RNA) serves as a negative control shown in black box. The peaks identified by MACS are shown in blue together with genes from MaGe annotation at the bottom of each panel. In the IGV visualization profiles, “+” strand reads are shown in red and “-“ strand reads are shown in blue. Dynamic IGV visualization is presented with adjusted read threshold for each window to compare the Hfq-immunoprecipitated and control samples.

A hundred of enriched RIP-seq peaks were detected in antisense orientation to gene annotations in the *C. difficile* 630 genome (Supplementary Table S3). Analysis of these peaks led to the identification of 36 potential new antisense RNAs, including 22 antisense RNAs to CDS, 4 antisense RNAs to previously identified ncRNAs and 10 antisense RNAs to riboswitches responding to SAM, lysine, glycine, FMN, cobalamin and T-boxes. In 7 cases, antisense peaks corresponded to 3’ extensions of antisense RNAs previously identified by transcriptional start site (TSS) mapping and whose 3’end remained undefined. 18 peaks in antisense orientation could represent either new RNAs antisense to CDS or transcription readthroughs from adjacent genes. A total of 45 antisense peaks was associated with genomic regions covered by overlapping bi-directional transcriptional units. These included previously identified type I TA modules and other genes co-transcribed with associated antisense RNAs, as well as a number of overlapping genes in convergent orientation.

In general, RIP-seq results showed a specific increase for particular categories of RNAs for the strain bearing a FLAG tag. Over 75% of Hfq-enriched peaks mapped to a position of mRNAs and the most abundant class of regulatory RNAs enriched in Hfq-FLAG samples corresponded to antisense RNAs (Supplementary Table S3). These antisense RNAs represented more than one third of all regulatory RNA-derived peaks. We present below examples of highly enriched RNAs in Hfq immunoprecipited sample that belong to different functional classes (Figure 2).

#### sRNAs enriched in Hfq-immunoprecipitated samples

About 100 sRNAs were previously discovered in *C. difficile* that could act as potential *trans*-encoded RNA regulators in concert with Hfq protein (15). Of the Hfq-enriched peaks identified in this study, 24 corresponded to these sRNAs (Figure 2A, Supplementary Table S4). Among previously identified sRNA interacting with Hfq *in vitro* (27), we show that RCd1 and RCd6 bound Hfq with a 254- and 16-fold enrichment, respectively, although the last interaction was not statistically validated (Figure 2A). RCd1 is a ncRNA conserved in sequenced *C. difficile* strains that has been previously detected in both the historical strain 630 and the epidemic strain R20291 of 027 ribotype by Northern blot and RNA-seq (15). Further studies demonstrated that the accumulation of RCd1 is negatively affected by the depletion of Hfq and a tight interaction of RCd1 with the Hfq protein has been shown *in vitro* by RNA band-shift experiments (27). It is interesting to note that the RIP-seq experiment revealed the presence of an RNA antisense to RCd1 with a 78-fold enrichment in Hfq-FLAG-immunoprecipitated samples (Figure 2A). We detected this antisense RNA of around 100 nt by Northern blotting but the signal remained low under all conditions tested (Supplementary Figure S4). Two additional sRNAs, CD630_n00290 and SQ1828, whose accumulation was previously shown to be modulated by Hfq depletion (27) were enriched in Hfq-immunoprecipitated samples with statistical validation for CD630_n00290 (Figure 2A). Other Hfq-enriched sRNAs are presented in Supplementary Figure S5A. Interestingly, antisense RNA transcription could be detected for several of these enriched peaks, including CD630_n00440, CD630_n00930, and CD630_n00330. By contrast and in accordance with previously reported co-immunoprecipitation results with Hfq from *Salmonella* (54), cDNAs of the abundant, Hfq-independent 6S RNA were found in rather equal numbers in Hfq-immunoprecipitated and control samples (Figure 2A).

#### Antisense RNAs

Remarkably, significantly enriched peaks mapped to the regions transcribed from both sense and antisense strands forming potential RNA duplexes. Data from our RNA-seq and TSS-mapping had revealed the presence of type I TA systems in *C. difficile* with RNA antitoxin acting as antisense RNA to toxin mRNA (18–20). We further characterized some of them (18,19) and identified a total of 13 type I TA modules in the reference strain 630 including the previously unannotated TA module with CD630_n00150 antitoxin/*CD0440.1* toxin genes (20). Our RIP-seq analysis shows that the toxin and antitoxin RNAs of all these systems bind to Hfq, as is the case for CD2517.1 toxin mRNA and RCd8 antitoxin (2,650-fold enrichment) (Figure 2B) (see Supplementary Figure S5B for additional examples).

In addition to 13 antitoxin RNAs, a total of 21 previously identified antisense RNAs was detected as enriched in Hfq coIP samples. Among them we detected the antisense RNA CD630_n00830, which overlaps the *grdB* gene encoding a glycine reductase (Figure 2C). This RNA was enriched approximately 2,800-fold for 3xFLAG Hfq compared to the control strain. This peak of 544 nt, associating with a strong TSS identified in our previous work (15), is significantly enriched in both Hfq-FLAG and wild type strains and covers the 3’-coding part of *grdB* gene in antisense orientation (Figure 2C). The antisense RNA, RCd2 was also enriched in the Hfq coIP sample (Figure 2C). This RNA is transcribed in an antisense orientation to two adjacent genes: *CD0182* of unknown function and *CD0183* encoding a putative cell wall hydrolase. RCd2 overlaps the 5’-end of the *CD0183* gene. Both sense and antisense reads could be detected in this overlapping region suggesting that these RNAs co-immunoprecipitated with Hfq. Other examples of enriched antisense RNAs were CD630_n00340 overlapping the *CD0898* gene of unknown function, CD630_n00080 overlapping the *CD0249* (*fliG*) flagellar motor gene, CD630_n00910 overlapping *CD2711* gene encoding a LysR family transcriptional regulator and SQ476 overlapping *CD0740* gene encoding a putative pyridoxal phosphate-dependent aminotransferase (Supplementary Figure S5C and Table S3).

Interestingly, RIP-seq analysis revealed a peak corresponding to an antisense RNA overlapping the abundant tmRNA in *C. difficile* (Supplementary Figure S5C). Two other peaks significantly enriched in the Hfq coIP sample were detected antisense to the CD630_s0610 lysine-riboswitch sRNA and to the CD630_s0590 lysine-riboswitch sRNA (Supplementary Figure S5C). Other examples of potential new antisense RNAs co-immunoprecipitated with Hfq are presented in Supplementary Figure S5C.

#### CRISPR RNAs

Among the most abundant RNAs detected by our previous TSS mapping experiment were CRISPR RNAs (crRNAs) (15) from the adaptive immunity CRISPR-Cas system. Here, RIP-sequencing identified enriched peaks for all twelve CRISPR arrays in the *C. difficile* 630 strain. In most cases, the peak covered mainly the 5’-portion of the CRISPR arrays (e.g. CRISPR 6, CRISPR 7, CRISPR 9, CRISPR 11, CRISPR 17) corresponding to the most abundantly expressed leader-proximal part of the locus (15) (Figure 2D for CRISPR 6 and Supplementary Figure S5D for other examples). Two separate peaks were detected in both 5’- and 3’ parts of CRISPR 8 array (Figure 2D) and a longer 441-nt peak almost covering the entire CRISPR 10 array could be detected (Supplementary Figure S5D). In some cases, regulatory function has been suggested for crRNAs in addition to defensive function (58,59), it would be interesting to assess in the future this possibility for crRNA-mediated regulations of gene expression in *C. difficile*.

#### Riboswitches

Previous deep-sequencing analysis revealed that the *C. difficile* non-coding RNome is characterized by a great number (66) of 5’-RNA regulatory elements called riboswitches (15,60). In particular, 16 riboswitches responding to the signaling molecule c-di-GMP and mediating the coordinated control of motility, biofilm formation, toxin production and other community-behavior related processes have been identified in this pathogen (15,61–64). Our RIP-seq data, revealed a total of 27 riboswitches significantly enriched in the Hfq-FLAG sample, including numerous c-di-GMP riboswitches of type I and II, such as Cdi1_3 upstream of the flagellum operon (Figure 2E) and Cdi2_3 upstream of the *CD2831* gene encoding collagen-binding protein precursor (Supplementary Figure S5E). T-boxes responding to the aminoacylation level of tRNA, riboswitches responding to lysine, S-adenosyl-methionine (SAM), purine and flavine mononucleotide were also co-immunoprecipitated with Hfq (Figure 2E for Arg T-box and Supplementary Figure S5E). In most cases, the RIP-seq peak was located specifically inside the riboswitch region corresponding to the short premature-terminated transcript and the downstream mRNA region was not co-immunoprecipitated. These data raise a question on the possible implication of Hfq in riboswitch RNA-related regulatory function *in cis* but also *in trans*.

#### Potential new transcribed regions

Interestingly, 18 peaks overlapped IGR with no gene annotated in the *C. difficile* genome (Supplementary Table S3). The reannotation of TSS (65) allowed assigning 9 of these peaks to 5’UTR regions that were specifically enriched as compared to the coding part of the mRNA and one corresponded to a 3’UTR. Four others were located within IGR and might correspond to novel ncRNAs that were not identified in our previous studies, such as the region between the *CD0655* and *CD0656* genes adjacent to the pathogenicity locus (PaLoc) (Figure 2F) (see Supplementary Figure S5F for other examples). One peak was assigned to a previously identified *cis*-antisense RNA with a misannotated 3’-end. One new *cis*-antisense RNA could be also detected as well as three additional double stranded regions with potential antisense transcription.

Thus, in addition to previously identified regulatory RNAs, new potential RNAs were detected through their association with Hfq protein.

#### mRNAs

Hfq RIP-seq data showed enrichment for numerous mRNAs (634) (Supplementary Table S3) such as the mRNA of the *CD1517* (*feoB*) gene encoding an iron transporter or of the *CD1909* (*eutP*) gene encoding an ethanolamine utilization protein (Figure 3A, whose expression is also modified in our transcriptomic analysis (Supplementary Table S3, Table S5, Table S6, Figure S5F). In addition to these genes, Supplementary Figure S5G shows other examples of virulence- and host-adaptation related genes revealed by RIP-seq analysis including *CD0663 (tcdA)* and *CD0664 (tcdC)* genes encoding one of the major *C. difficile* virulence factors toxin A and a negative regulator of toxin gene expression, TcdC within the pathogenicity locus (Paloc) region, respectively, as well as the *CD2854-2851* (*dltDABC)* operon encoding the D-alanylation proteins that could provide protection against antimicrobial peptides (66,67).

**Figure 3.**
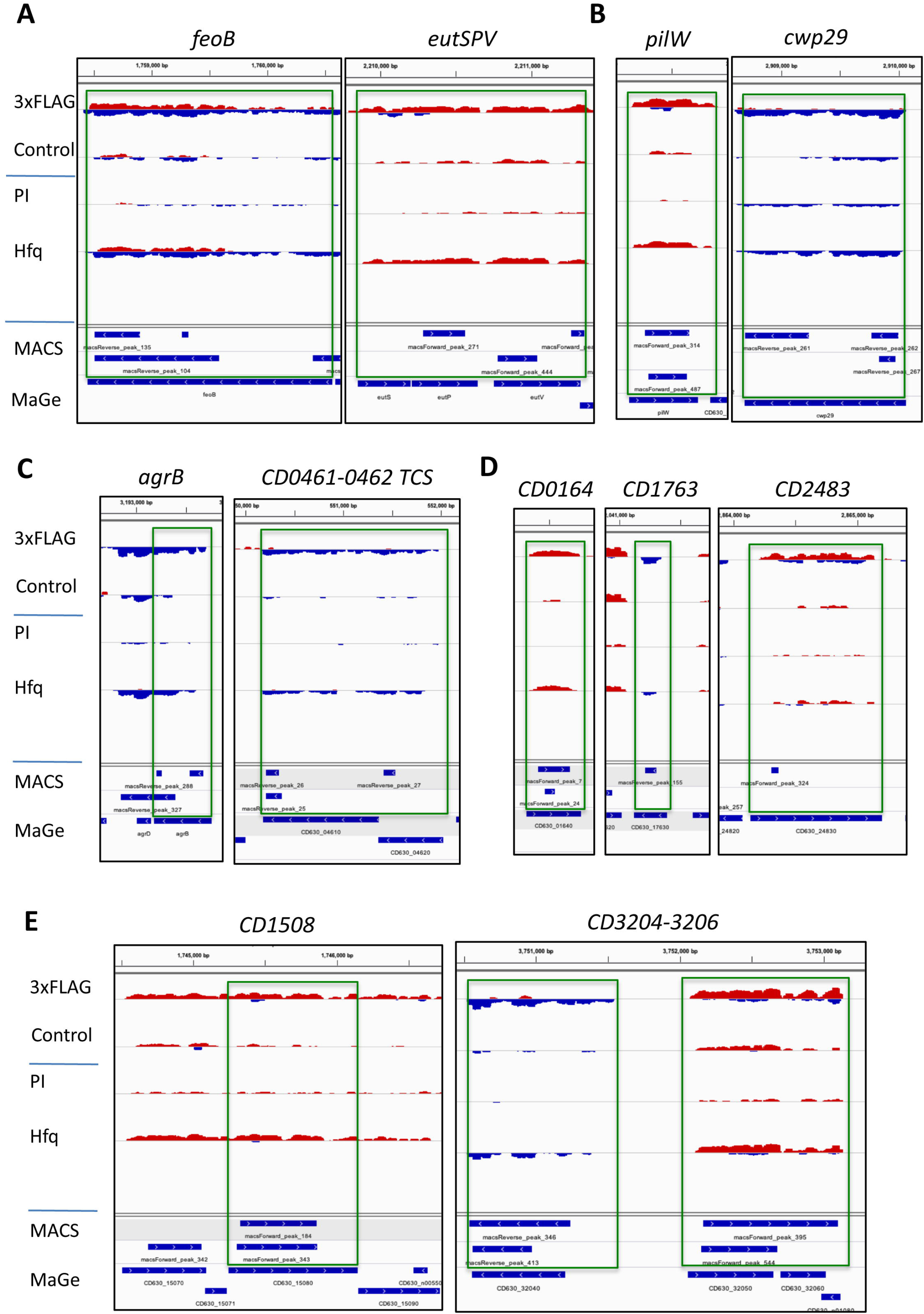
Visualization of RIP-seq data for mRNAs with IGV. Representative examples from different functional groups of mRNAs are presented in A) host-adaptation related genes, B) for type IV pili, adhesion, surface protein encoding genes, C) for regulatory and signaling pathways including quorum sensing, two-component systems and transcriptional regulatory genes, D) for membrane proteins and transporter genes, E) for metabolism genes. The IGV visualization is presented as in Figure 2.

Among genes with surface-related functions we observed a significant enrichment of *CD3504* (*pilD*)*, CD3505* (*pilT*) and *CD3511* (*pilC*) genes of *CD3513-3503* operon and *CD2305* (*pilW*) encoding type IV pili proteins, as well as genes *CD1036* (*cwp17*)*, CD1233* (*cwp26*)*, CD1803* (*cwp23*)*, CD2518* (*cwp29*) and *CD3145* (*cbpA*) (Figure 3B for *pilW* and *cwp29*, Supplementary Figure S5G for other examples).

A number of mRNAs for genes belonging to regulatory and signaling pathways were enriched in Hfq-immunoprecipitated samples including the quorum sensing gene *agrB*, the two-component systems-encoding genes *CD0461-0462* (Figure 3C) and *CD1829-1830* (*kdpDE*), the sensor histidine kinase *CD0576* (*virS*) as well as the transcriptional regulator genes *CD1076 (*LysR family)*, CD2615* (TetR family) and *CD3062* (RpiR family) (Supplementary Figure S5G). Numerous genes encoding membrane proteins and transporters were also detected such as *CD0164* (Figure 3D), *CD2483* lipoprotein, *CD0316-0318* and *CD0364-0366* ABC transporter genes (Supplementary Figure S5G). In line with the suggested pleiotropic role of the RNA chaperone protein in *C. difficile*, numerous mRNAs for metabolism genes (Figure 3E, Supplementary Figure S5G) were found as enriched in Hfq-immunoprecipitated samples, e.g. the iron-sulfur binding protein gene *CD150,* the nitroreductase gene *CD3205* (Figure 3E) and the flavodoxin/nitric oxide synthase gene *CD3121* (Supplementary Figure S5G).

Interestingly, in addition to CRISPR RNAs, the mRNAs for Cas1, Cas2 and Cas4 (CD2975-2976-2977) CRISPR-associated adaptation proteins were co-immunoprecipitated with Hfq (Supplementary Figure S5C), further suggesting the potential implication of Hfq in CRISPR-Cas-related regulatory processes.

Hfq autoregulation has previously been shown in *Escherichia coli* where Hfq represses its own translation primarily by binding to the *hfq* 5’UTR mRNA (68). Accordingly, we observed a significant enrichment of *CD1974 hfq* mRNA in the 3xFLAG Hfq-immunoprecipitated sample as compared to control with a MACS peak covering almost the entire gene being found (Supplementary Figure S5G). Intriguingly, a faint transcript in antisense orientation was also detected suggesting an additional level of regulation of *hfq* expression. In *Legionella pneumophila,* an anti-*hfq* RNA has been recently discovered as one of the central components of Hfq-dependent regulatory network governing this pathogen differentiation from nonvirulent to virulent bacteria (69). Our findings thus suggest the existence of a complex autoregulatory loop controlling *hfq* expression in *C. difficile*.

#### Validation of RIP-seq data

To validate RIP-seq enrichment we have performed qRT-PCR analysis for selected RNAs from different functional classes on Hfq-FLAG-immunoprecipitated and control samples in at least three biological replicates (Supplementary Table S7). This analysis confirmed the enrichment of 89-fold in average in Hfq-FLAG-immunoprecipitated samples as compared to control for all tested Hfq RNA partners. Among sRNAs, this validation included RCd1 that binds Hfq *in vitro* with high affinity (27), as well as CD630_n00290 and SQ1828 that were differentially expressed in *hfq*-depleted strain (27) (Figure 2A). A 34-fold enrichment for the antisense RNA RCd2 was confirmed by qRT-PCR (Supplementary Table S7, Figure 2C). Among CRISPR RNAs, the enrichment of CD630_n00460 CRISPR 6 and CD630_n00560 CRISPR 8 was also validated (Supplementary Table S7, Figure 2D). For the riboswitch class of Hfq partners, a representative Cdi1_8 c-di-GMP-responsive riboswitch was detected as highly enriched in Hfq-FLAG immunoprecipitated samples (Supplementary Figure S5E, Supplementary Table S7). Importantly, an enrichment of 15- and 30-fold was observed for the newly identified transcribed IGR *CD0655* RNA and the new antisense RNA to *CD2890*, respectively (Figure 2F). In addition to the Northern blot validation of the new antisense RNA to RCd1 revealed by RIP-seq analysis (Supplementary Figure S4), the detection of five ncRNAs enriched in RIP-seq was also validated by Northern blot (Supplementary Figure S6). We have selected pathogenesis- and colonization-related genes *tcdA, tcdC* and *pilA* among mRNA partners of Hfq for qRT-PCR analysis and confirmed their enrichment in Hfq-FLAG immunoprecipitated samples (Supplementary Figure S5G, Supplementary Table S7).

Our RIP-seq data are in complete accordance with all the previously published results from independent studies by various large-scale and targeted approaches on *C. difficile* ncRNA identification and functional characterization including antisense RNAs within TA systems, CRISPR RNAs, riboswitches and sRNAs. The majority of ncRNAs captured as Hfq ligands have been previously identified by genome-wide approaches including TSS mapping and RNA-seq (15,65) and some of them have been characterized by 5’-RACE, Northern blot, RT-PCR, *in vitro* interaction analysis and RNA stability assays (15,16,18–20,27,65).

In addition, the RIP-seq data on the enriched RNAs associated with Hfq overlapped with the results of our previous transcriptome analysis of the *C. difficile* strain depleted for Hfq under the same growth conditions (27). This analysis revealed that 53 of the Hfq-bound RNAs (40 mRNAs and 13 ncRNAs) were found differentially expressed in Hfq-depleted strain as compared to the control suggesting direct implication of Hfq-dependent post-transcriptional mechanisms in these regulations (Supplementary Table S5, Supplementary Figure S7). These genes are implicated in membrane transport, cell wall, regulation processes and sporulation including several members of the late sporulation phase sigma factor SigK regulon. Among ncRNAs, the riboswitches and CRISPR RNA classes were overrepresented.

To get a global view of functional gene categories subjected to Hfq-dependent control, we analyzed the functional categories of RIP-seq targets using clusters of orthologous groups from the KEGG databases (46). In accordance with previous phenotypic and transcriptomic analysis, this revealed three main functional categories (Supplementary Table S6, Figure 4). The most abundant category detected was composed of 199 genes encoding membrane proteins with additional 45 genes encoding membrane transporters (enrichment score 9.98). Several clusters fitted into the functional category of regulation and signaling processes including a great number of transcriptional regulators, two-component signal transduction systems and diguanylate kinase signaling proteins. A large cluster of 113 genes corresponds to genes of unknown function. Functional clusters for genes encoding metabolic proteins were also overrepresented in our analysis. These data suggest that Hfq could contribute to important regulatory pathways in *C. difficile*.

**Figure 4.**
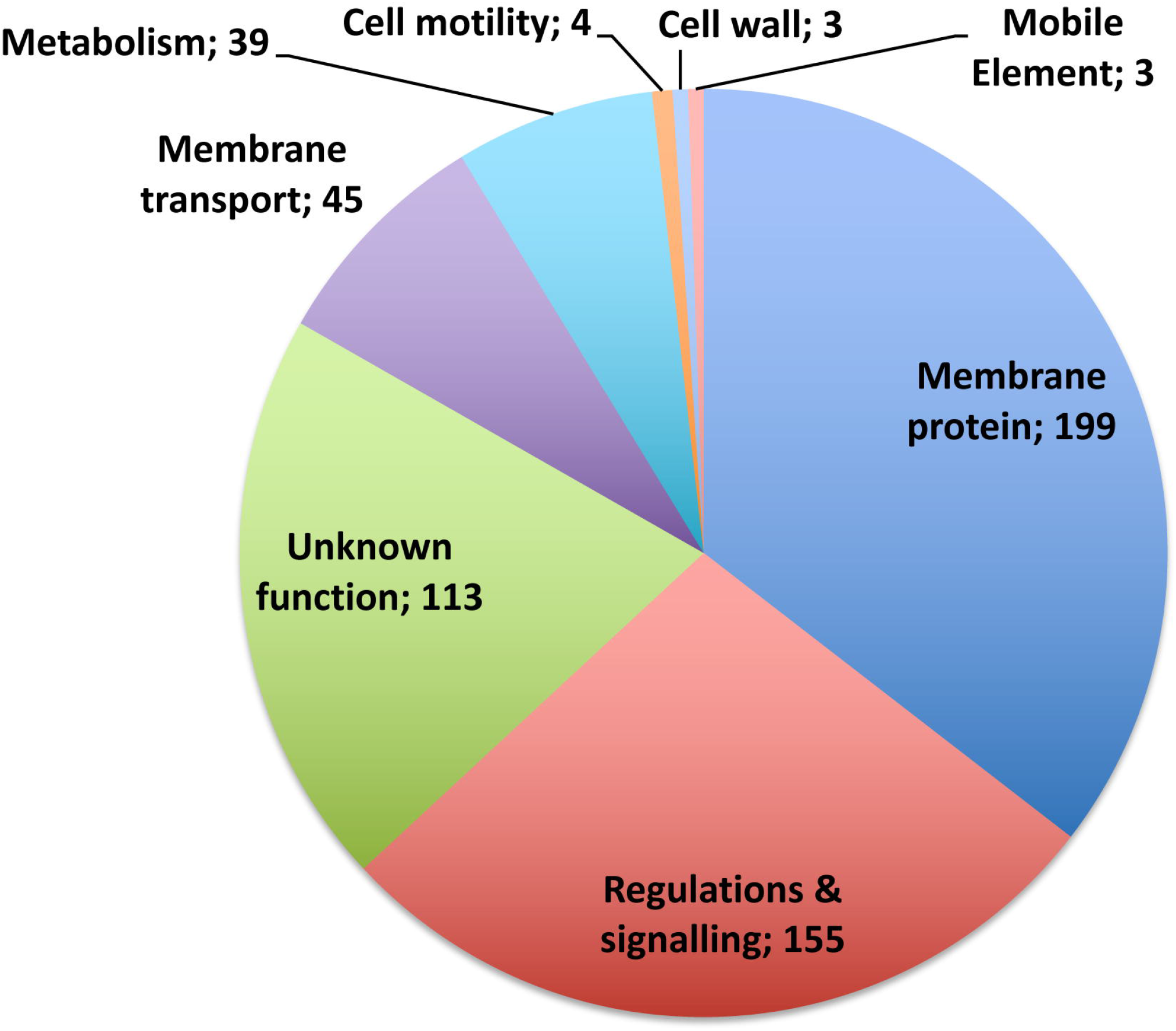
Diagram showing pathway clustering for mRNAs peaks detected by RIP-seq from KEGG database. The functional category of genes is shown with the corresponding gene number.

### Role of Hfq-bound sRNA RCd1 in sporulation control

One of the sRNAs significantly enriched in Hfq-FLAG coimmunoprecipitated samples, RCd1, has previously been identified as a potential regulator of sporulation (27). Interestingly, the increase in sporulation efficiency is also among the most pronounced effects observed for Hfq depletion and *hfq* deletion strains (27,70). To get more insight into the role of Hfq in the RCd1-dependent sporulation control, we analyzed the effect of Hfq-depletion on the sporulation efficiency in wild-type and ΔRCd1 strains. We used an antisense RNA leading to *hfq* knock-down and conditions enabling efficient Hfq-depletion (at least 5-fold) as validated in our previous work (27). Hfq-depletion in the wild-type strain led to an important increase in sporulation rate after 72 h in BHIS medium but had no impact on sporulation in the ΔRCd1 background (Figure 5A). Together with previous studies these data suggest that RCd1 could act in concert with Hfq for the negative control of sporulation. Since Hfq controls the regulon of the mother cell late sigma factor, SigK (27), we hypothesized that RCd1 also targets directly or indirectly *sigK.*

**Figure 5.**
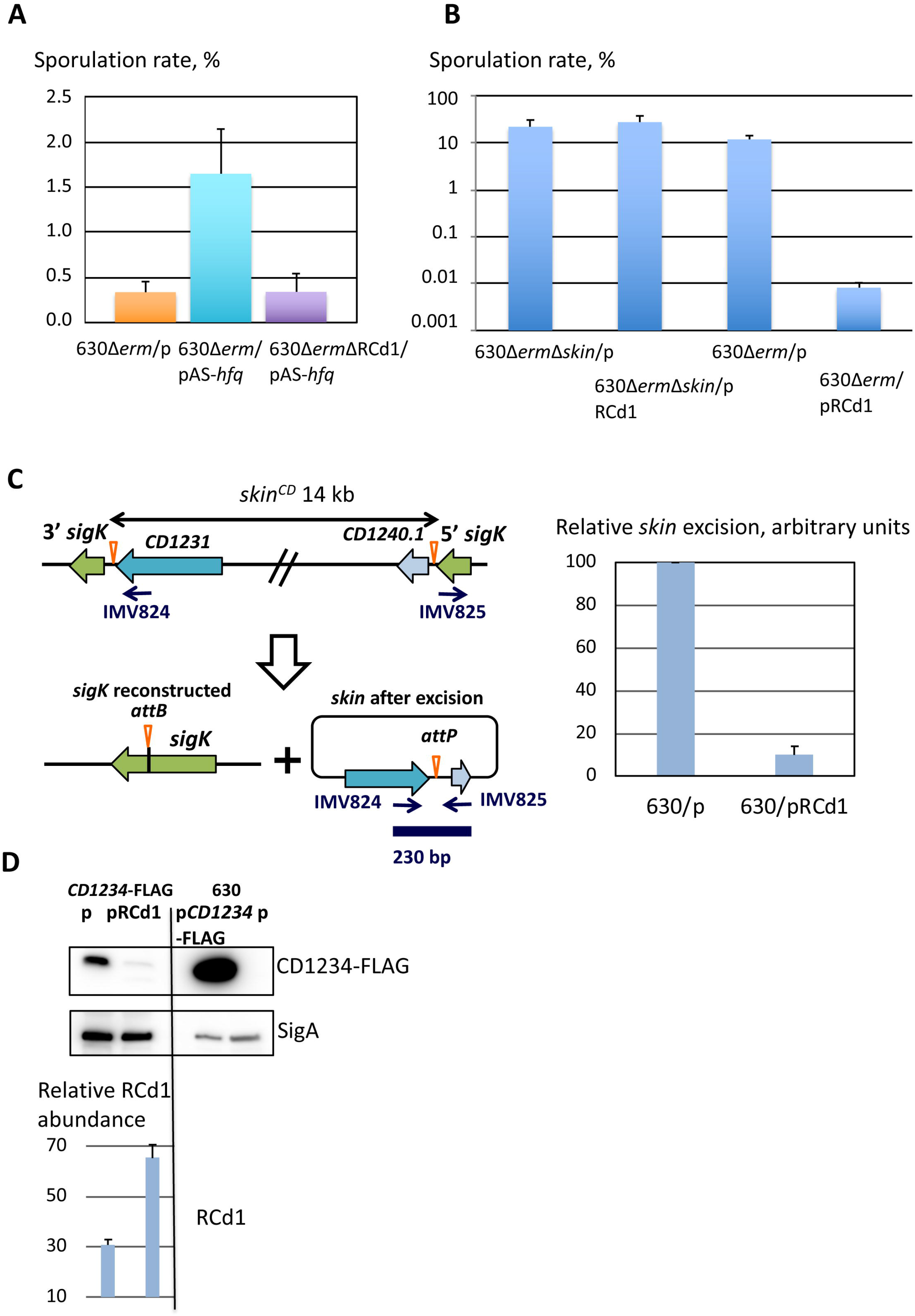
The role of RCd1 in sporulation control. **A)** Hfq depletion effect on sporulation rate was estimated in BHI-S medium after 72 h of culture in wild-type or ΔRCd1 background in strains CDIP51 (630Δ*erm*/p), CDIP53 (630Δ*erm*/pAS-*hfq*) and CDIP619 (630Δ*erm*ΔRCd1/pAS-*hfq*). The diagram shows the sporulation rate as a proportion in % of spores for the total number of CFU, mean values of results from at least three independent experiments are presented. Error bars correspond to standard deviations from three biological replicates. Statistical significance for spore formation differences between strains CDIP51 and CDIP53, and strains CDIP53 and CDIP619 was evaluated by a Welch two sample *t* test (*p*=0.03), *N*=3. **B)** Sporulation efficiency by numeration of spores formed by the strains overexpressing RCd1 and their control strains. Sporulation assays were performed in SM after 24 h of culture. Spores were selected by heat treatment at 65°C for 20 min and their number estimated for strains CDIP590 (630Δ*erm*Δ*skin*/p), CDIP592 (630Δ*erm*Δ*skin*/pRCd1), CDIP219 (630Δ*erm*/p), CDIP258 (630Δ*erm*/pRCd1). The diagram shows the sporulation rate as a proportion in % of spores for the total number of CFU, means of results from at least three independent experiments are presented. Error bars correspond to standard deviations from three biological replicates. Statistical significance for spore formation differences between strains CDIP219 and CDIP258 was evaluated by a Welch two sample *t* test (*p*=0.02), *N*=3. **C)** Diagram of the *skin* element integrated into the *sigK* gene and its episomal form shows the location of the oligonucleotides used as primers for PCR (IMV824/IMV825) to distinguish between integrated and episomal *skin* forms. The PCR results from three independent experiments for relative *skin* excision are presented. **D)** Western-blot detection of FLAG-tagged CD1234 protein in the cell extracts of *C. difficile* strains : lane 1-strain CDIP704 (CD1234-FLAG/p); lane 2-strain CDIP708 (CD1234-FLAG/pRCd1); lane 3-strain CDIP607 (630Δ*erm*/pCD1234-FLAG); lane 4-CDIP369 (630Δ*erm*/p). Immunoblotting with anti-FLAG antibodies detected a major polypeptide of ~11.5 kDa in whole cell lysates of the strain carrying *CD1234-*3xFLAG gene on the chromosome of *C. difficile* 630Δ*erm* strain but not in extracts of the wild-type strain expressing the non-tagged protein (negative control on lane 4). The representative result from three independent experiments is shown. The strains were grown in SM medium for 12 h in the presence of 250 ng/ml ATc and 7.5 μg/ml Tm. 60 μg (lanes 1-2) or 10 μg (lanes 3-4) of whole cell lysate were loaded on the gels. The lysate of the strain overexpressing CD1234-FLAG from the plasmid serves as a positive control for detection (lane 3). Proteins were separated on 4-20% Bis-Tris polyacrylamide gels in Tris-Glycine SDS buffer. Immunoblotting with anti-SigA antibodies serves as loading control (bottom panel). The relative RCd1 abundance has been estimated by qRT-PCR, the mean values with standard deviations from three independent experiments are shown.

#### Effect of RCd1 on the excision of the skin^Cd^ element

The *sigK* gene in *C. difficile* is interrupted by a 14 kb prophage-like *skin*^*Cd*^ element. The *skin* excision thus constitutes an important initial step for the σ^K^-dependent cascade activation during sporulation. Two proteins encoded within the *skin* element, CD1231, a specific recombinase, and CD1234, a recombination directionality factor, are required for *skin* excision (33). To evaluate if the sporulation control by RCd1 could be linked to the *skin* element, we tested the effect of RCd1 overexpression on sporulation efficiency in a *C. difficile* strain lacking *skin*^*Cd*^ and therefore having an intact copy of the *sigK* gene (33). We introduced the plasmid allowing the overexpression of RCd1 in the Δ*skin*^*Cd*^ strain as well as an empty plasmid used as a control. An estimated 3 to 10-fold overexpression of RCd1 was achieved in the presence of ATc inducer throughout the sporulation kinetics (Figure 6). The number of spores formed in sporulation medium (SM) after 24 h of culture at 37°C was estimated and compared to that of the 630Δ*erm* carrying the same plasmids. While we obtained an important decrease in sporulation efficiency in 630Δ*erm* strain overexpressing RCd1, similar sporulation efficiencies were observed in the Δ*skin*^*Cd*^ strain overexpressing RCd1 and carrying the control vector (Figure 5B). Thus, RCd1 loses its inhibitory effect on sporulation when the *skin*^Cd^ element is absent and the *sigK* gene is intact. This suggests that the RCd1 target(s) could be located within the *skin*^*Cd*^ element or at least be implicated in the *skin*^*Cd*^ excision. To test whether RCd1 acts on the *skin*^*Cd*^ excision, we extracted genomic DNA after 24 h of culture in SM medium for the 630Δ*erm* (pDIA6103) and 630Δ*erm* (pDIA6103-RCd1) strain. Using a pair of divergent primers at each end of the integrated sequence (33), we then quantified by qPCR the episomal form of the *skin*^*Cd*^ element representative of the excision rate in the population. After excision, these two primers converge and amplify a 230 bp fragment on the *skin*^*Cd*^ episome (Figure 5C). We found that the episomal form is 10 times more abundant in 630Δ*erm* (pDIA6103) than in 630Δ*erm* (pDIA6103-RCd1) relative to the reference *dnaF* gene encoding the DNA polymerase III (Figure 5C). This indicates that overexpression of RCd1 results in a reduction of excision rate of the *skin*^*Cd*^ element. Thus, RCd1 probably inhibits sporulation by preventing reconstitution of the intact *sigK* gene and therefore SigK production, inhibiting the late sporulation steps dependent on SigK.

**Figure 6.**
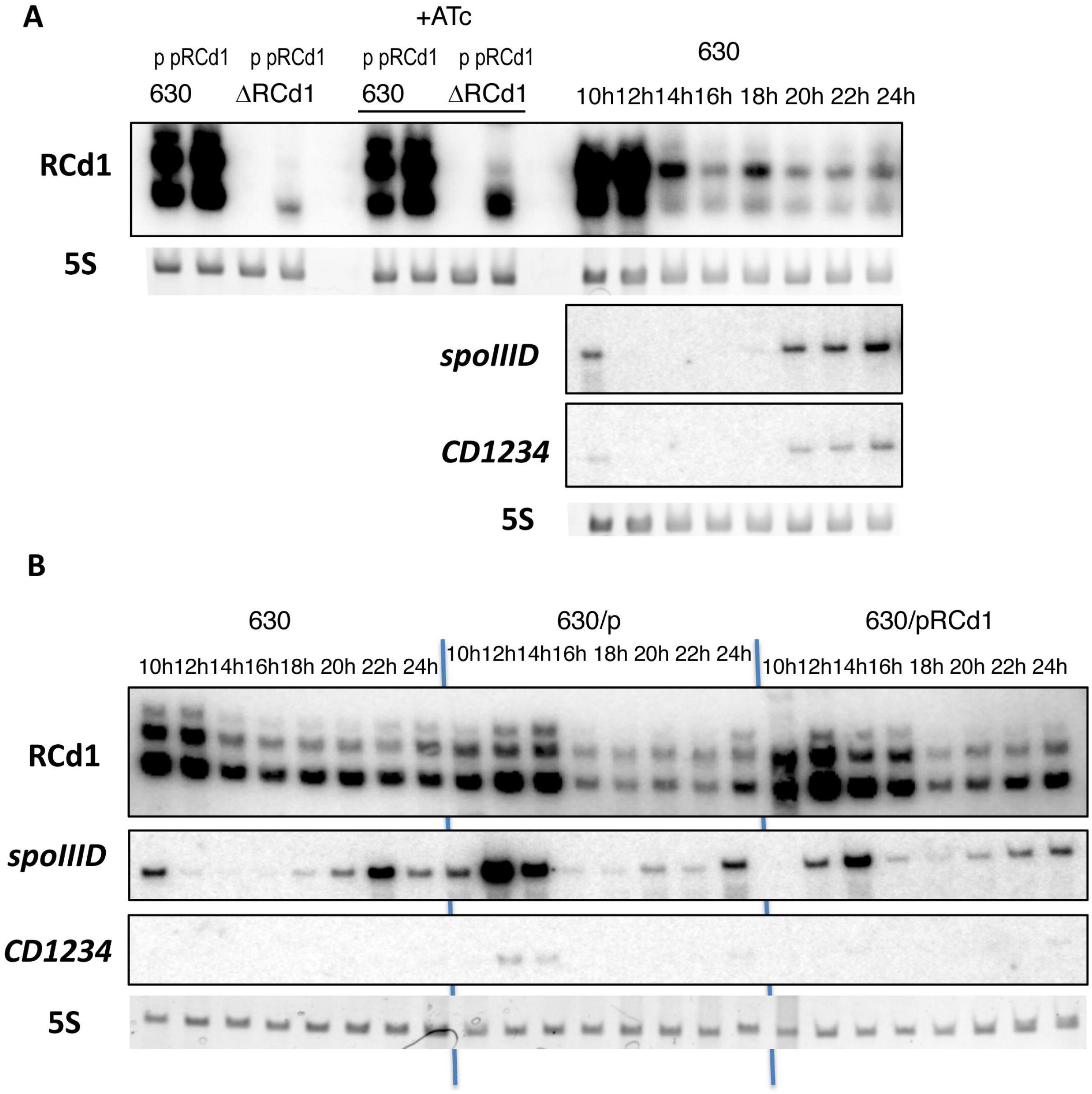
The expression profile of RCd1 and selected potential targets during sporulation. **(A)** RNA samples were extracted from 630Δ*erm* strain (630), 630Δ*erm* strain carrying an empty vector (630/p) or overexpressing RCd1 from plasmid (630/pRCd1) grown in TY medium supplemented on not with 250 ng/ml ATc (+ATc) or in SM medium for 10 h, 12 h, 14h, 16 h, 18 h, 20 h, 22 h, 24 h as indicated. The strain deleted for RCd1 (ΔRCd1) was used as a control for specific RCd1 detection. 5S RNA at the bottom serves as loading control. As indicated at the left, the blots were hybridized either with RCd1, *spoIIID* or *CD1234*-specific probes. The same 5S control panel is shown when reprobing of the same membrane was performed. **(B)** Effect of RCd1 overexpression on *spoIIID* and *CD1234* genes during sporulation was monitored in SM medium during sporulation for 10 h, 12 h, 14h, 16 h, 18 h, 20 h, 22 h, 24 h as indicated.

#### Effect of RCd1 on the abundance of potential targets

To assess the effect of RCd1 overexpression on the abundance of potential targets, we evaluated their expression by qRT-PCR analysis. To do so, the 630Δ*erm* strains carrying either a plasmid overexpressing RCd1 under the control of the inducible P_*tet*_ promoter or an empty plasmid as a control were grown in TY medium until the late exponential phase or for 12 h in SM medium. In agreement with our previous results (27), we observed that the *sigK* gene was less expressed in the strain overexpressing RCd1 compared to the control (Table 1). We also measured the impact of RCd1 overexpression on genes belonging to the SigK regulon, i.e. *CD0597* and *CD1613* genes encoding the spore coat proteins CotJB and CotA, respectively. *CD0597* and *CD1613* mRNA levels were reduced when RCd1 was overexpressed (Table 1). Thus, RCd1 is able to modulate the abundance of not only the *sigK* transcript but also its targets within the SigK regulon.

**Table 1.**
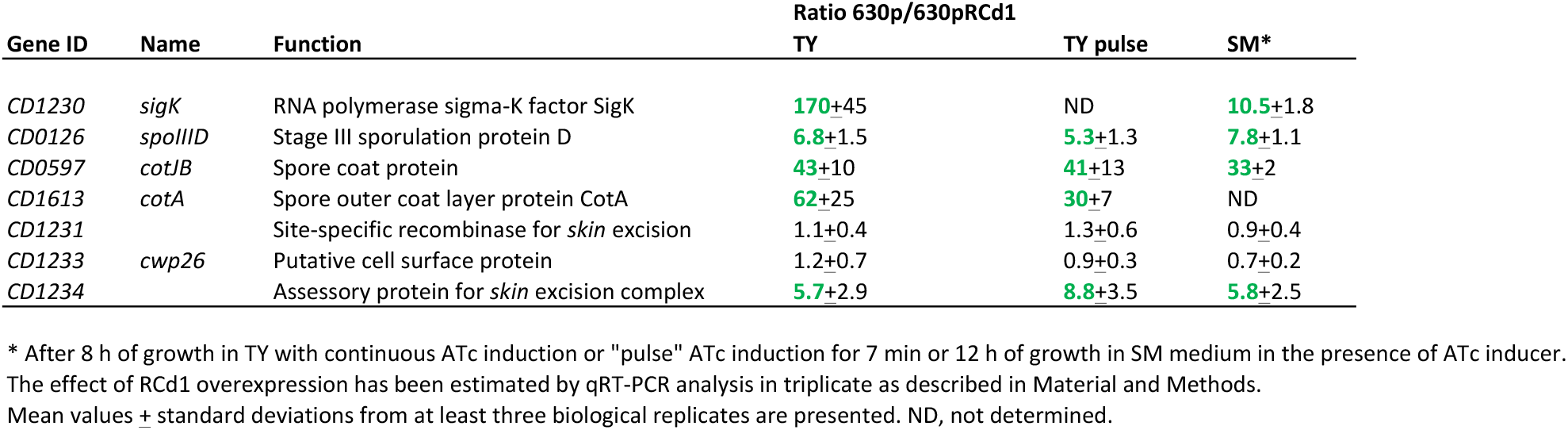
Effect of RCd1 overexpression on the abundance of potential targets. After 8 h of growth in TY medium with continuous ATc induction or “pulse” ATc induction for 7 min or 12 h of growth in SM medium in the presence of ATc inducer. The effect of RCd1 overexpression has been estimated by qRT-PCR analysis in triplicate as described in Material and Methods. Mean values ± standard deviations from at least three biological replicates are presented. ND, not determined.

As RCd1 overexpression had an impact on *skin*^*Cd*^ element excision, we further tested, as potential RCd1 targets, *spoIIID* controlling *skin*^*Cd*^ excision and *sigK* expression (33) as well as three *skin*^*Cd*^ genes. Genes *CD1231* and *CD1234* encode essential components of the *skin*^*Cd*^ excision complex (33). We included *CD1233* encoding the putative cell surface protein Cwp26 since it might be an RCd1 target as predicted by the RNAPredator algorithm. While the expression of *spoIIID* and *CD1234* decreased upon RCd1 overexpression, no effect on *CD1231* and *CD1233* expression was observed in our conditions (Table 1). We also included the *sigE* gene encoding the mother cell sigma factor involved in the transcription of *spoIIID, sigK* and *CD1234,* as well as SigE targets, i.e. *CD1192* encoding the SpoIIIAA protein*, CD1940* encoding a putative membrane protein and *CD3542 (spmA)* encoding the spore maturation protein A. After 12 h of growth in SM medium, no effect of RCd1 overexpression on the expression of *sigE* and its target was observed (data not shown) positioning *spoIIID* and *CD1234* as main regulatory components for RCd1 targeting. By short 7 min pulse induction of RCd1 overexpression, we confirmed a negative effect on expression of both genes (Table 1).

We then analyzed a kinetic of expression of RCd1 and its potential targets in SM during sporulation. We harvested culture aliquots every 2 h between 10 h and 24 h. After RNA extraction, the expression profile of RCd1 and selected potential targets was assessed by Northern blot (Figure 6A). This analysis showed that RCd1 is expressed throughout the sporulation kinetics, being most abundant during the early kinetic time points. Interestingly, as previously observed for the *sigK* gene (33), a biphasic expression pattern was detected for *spoIIID* and *CD1234* genes (Figure 6A), a first peak of expression occurring at the beginning of sporulation (10 h-14 h) and a second increased expression after 20 h-24 h of growth. We also confirmed the negative effect of RCd1 overexpression on both *spoIIID* and *CD1234* during sporulation kinetics (Figure 6B).

To further investigate the effect of RCd1 overexpression on CD1234 production for regulation of *skin*^*Cd*^ excision, we introduced a 3xFLAG tag at the C-terminal part of *CD1234* coding region on the chromosome of *C. difficile* 630Δ*erm* strain (Supplementary Table S1). Derivative strains carrying a chromosomal *CD1234*-3xFLAG copy and either an empty vector or a plasmid for RCd1 overexpression were then used for Western blot analysis. The 630Δ*erm* strain carrying an empty vector or overexpressing a CD1234-3xFLAG protein fusion from the plasmid served as negative and positive controls, respectively. Based on our expression profile analysis during the sporulation kinetics, we extracted proteins from cells grown for 12 h in SM medium. Western blot analysis with anti-FLAG tag antibodies revealed a sharp decrease in the amount of CD1234-3xFLAG protein in the strain overexpressing RCd1 (Figure 5D lane 2) compared to the strain carrying the control vector (Figure 5D lane 1). While the *CD1231* recombinase gene is constitutively expressed, the *CD1234* gene is expressed under the control of both σ^E^ and SpoIIID and is then a crucial target for the timely control of *skin* excision in the mother cell only (33). No significantly enriched peak could be detected by MACS for *CD1234* or *spoIIID* in our Hfq RIP-seq data. However, visual inspection of RIP-seq data for the *skin*^*Cd*^ region revealed a detectable signal for *CD1234* gene in both Hfq-FLAG and Hfq-immunoprecipitated samples that was absent in the control samples. An enrichment for *spoIIID* at least in Hfq-immunoprecipitated sample as compared to control was also observed (Supplementary Figure S8). RIP-seq analysis also revealed peaks enriched in Hfq-FLAG immunoprecipitated samples for *CD1230, CD1231* and *CD1233* genes within *skin* element region.

Altogether these results suggest that RCd1 could act on top of the regulatory cascade triggering the inhibition of late sporulation stage setup by affecting first the SpoIIID regulator expression and then directly or indirectly the CD1234 component of the *skin*^*Cd*^ excision complex. The decrease in *skin*^*Cd*^ excision rate would then negatively affect the native *sigK* gene recovery thus impacting the whole downstream SigK regulon.

#### Interactions of RCd1 with potential targets in vitro

Based on our results of the RCd1 overexpression effect on potential target abundance, we selected *spoIIID* and *CD1234* for further interactions analysis. To investigate these interactions *in vitro* in presence or absence of purified Hfq protein, we then setup an RNA band shift assay. To explore first the role of Hfq, we analyzed the RNA-Hfq complex formation *in vitro*. The RNA band shift assays confirmed Hfq binding to RCd1 (27) and showed that Hfq could interact individually with each of the selected mRNAs *spoIIID* and *CD1234* with similar affinity (3.3 nM and 1.8 nM, respectively, Supplementary Figure S9). Incubations of sRNA with mRNA targets revealed shifted bands corresponding to the sRNA-mRNA complex. Addition of Hfq did not allow appearance of supershifted band corresponding to sRNA-mRNA-Hfq (Figure 7). From these experiments we could conclude that RCd1 may interact with rather low affinity with the selected mRNA targets under *in vitro* conditions and that Hfq is not required for these interactions *in vitro*.

**Figure 7.**
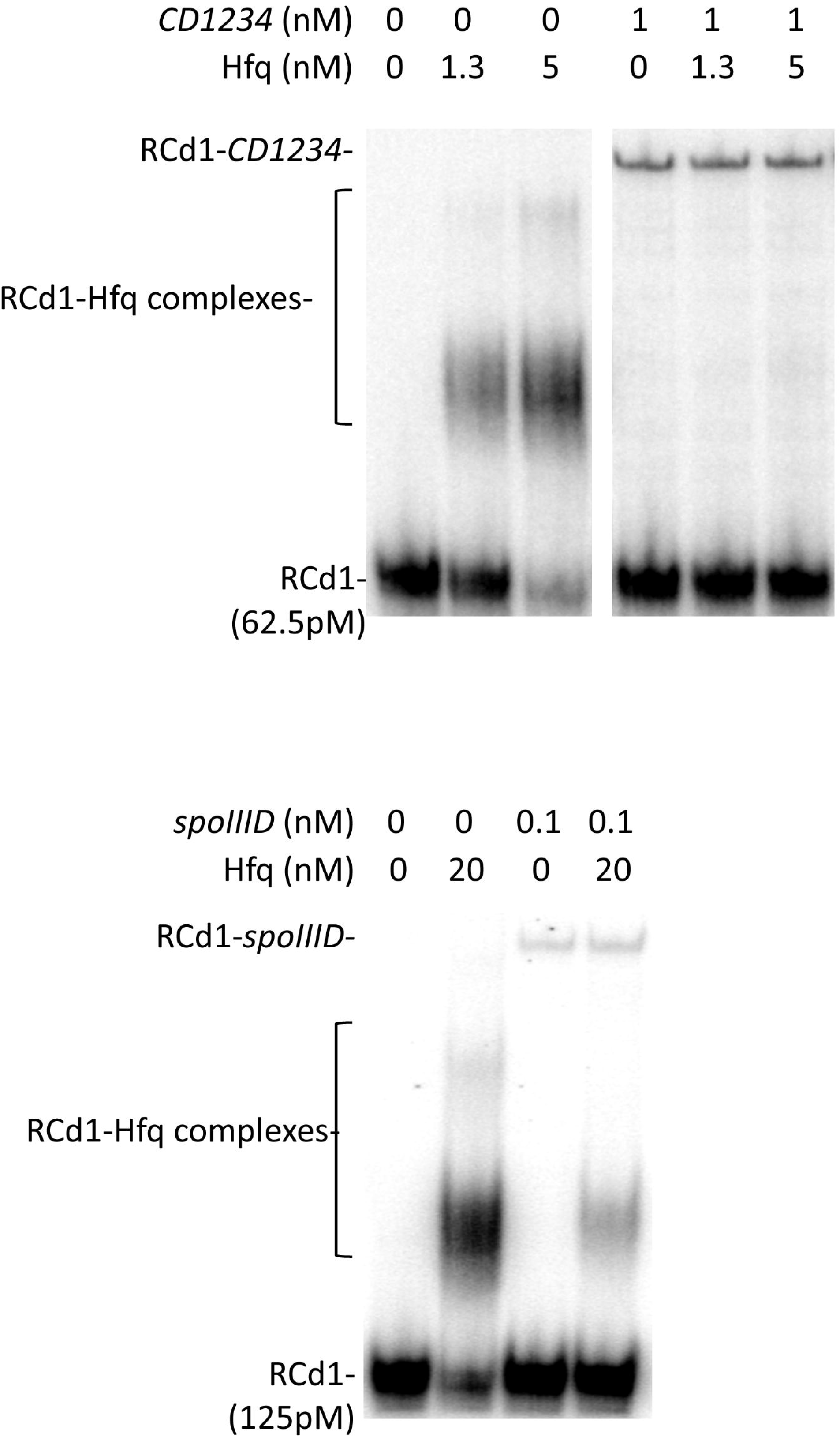
Analysis of the interaction between RCd1 and *spoIIID* or *CD1234* in the presence or the absence of Hfq. Radioactive RCd1 transcript was synthesized *in vitro* and incubated alone or together with *CD1234* or *spoIIID* transcripts with purified Hfq-His_6_ protein expressed as monomer forms. Brackets show Hfq-RCd1 complexes, the position of unbound RNA and RCd1-potential target complexes is indicated at the left.

#### Genomic analysis of RCd1 conservation and its co-occurrence with potential sporulation-related targets

The analysis of RCd1 conservation revealed the presence of close homologues only within *C. difficile* species. From 2,700 available *C. difficile* genomes, the sequences with at least 80% of nucleotide identity to RCd1 from laboratory strain 630Δ*erm* could be found in 99% of analyzed *C. difficile* strains (Supplementary Table S8). Multiple alignment of nucleotide sequences of RCd1 homologues showed extremely high level of conservation (Supplementary Figure S10). We also searched for the co-occurrence of RCd1 homologues with the important regulatory gene *spoIIID* and with the *skin*^*Cd*^ element including genes involved in the *skin*^*Cd*^ element excision, *CD1231* and *CD1234.* No clear correlation between the presence of RCd1 and the *skin* element could be observed (Supplementary Figure S11), the presence of a complete copy of the *skin* element being rather variable within *C. difficile* genomes. At 80% nucleotide sequence identity threshold, *spoIIID* homologues were found in all analyzed *C. difficile* genome sequences, while the *CD1231* and *CD1234* genes for *skin* excision complex were found in 85% of *C. difficile* genomes (Supplementary Table S8). These results suggest that RCd1 could have additional conserved functions in *C. difficile* beyond the control of *skin*^*Cd*^ excision.

## DISCUSSION

The RNA chaperone Hfq has emerged as an important player in the sRNA-based regulatory mechanisms in Gram-negative bacteria. However, its role remains less characterized in Gram-positive bacteria (71). Previous studies revealed unique features of the Hfq protein in *C. difficile* suggesting a pleiotropic role in this pathogen contrary to the majority of other studied Hfq homologues in Gram-positive bacteria (27). Transcriptomic analysis revealed that 6% of the genes are affected by Hfq depletion (27). The *C. difficile* Hfq protein was able to complement several phenotypes of the *hfq* deletion mutant in *E. coli* (28) as also demonstrated for the Hfq protein of *L. monocytogenes.* This latter bacterium is the only firmicute with at least one example of Hfq-dependent riboregulation (72,73). In the present study, the co-immunoprecipitation analysis showed that Hfq has a great number of ribonucleic ligands in *C. difficile*. During Hfq depletion, the abundance of several identified Hfq-binding RNAs transcribed from IGR was shown to be modified, suggesting that Hfq could affect their stability and contribute to their function (27). Compared to Hfq homologues from other Gram-positive bacteria, structural features of the *C. difficile* Hfq protein, such as the presence of most of the key residues within the RNA-binding sites including two of three conserved arginine residues in rim region (the outer ring on the surface of Hfq toroid structure as one of four sites for RNA contacts) and the presence of an unusual C-terminal region exceptionally rich in asparagine residues, may explain these unique characteristics. Recent results suggest that this C-terminal part is not essential but contributes to the function of the RNA chaperone (28). Extensive studies on Hfq-recognition motifs identified U-rich and A-rich or “ARN” sequences that bind to the proximal and distal face of Hfq, respectively (74). *In silico* search using MEME for potential RNA motifs recognized by *C. difficile* Hfq within the peaks significantly enriched in RIP-seq samples revealed a rather degenerated motif (Supplementary Figure S12). Presumably, the presence of conserved motifs within several classes of Hfq-associated RNAs and the AT-rich nature of *C. difficile* genome could make it difficult to precisely define an Hfq-binding motif in interacting RNA species.

To associate the ncRNAs with their potential targets in Hfq-dependent regulations in *C. difficile* we performed an *in silico* target prediction for the RNAs specifically enriched in our RIP-seq analysis using IntaRNA program (75) (Supplementary Table S9). A majority of Hfq-bound mRNAs was predicted as target for at least one Hfq-bound ncRNA, suggesting joint ncRNA-mRNA binding to Hfq. To illustrate these results, pairing predictions of several Hfq-bound ncRNAs are shown in Supplementary Figure S13.

This ncRNA-mRNA interaction prediction analysis revealed very large potential targeting capacities of crRNAs, raising the possibility of their regulatory functions in *C. difficile* as suggested in other bacteria (58). This includes the most abundant crRNAs from CRISPR 16/15 and CRISPR 3/4 arrays located in prophage regions of strain 630 chromosome (Supplementary Figure S13). No continuous coverage of entire spacer/target was observed and no functional protospacer-adjacent motif (PAM) could be associated with these potential interactions suggesting that the potential regulatory functions could be mediated by different Cas-nuclease-independent mechanisms (Supplementary Figure S13).

ncRNA-ncRNA interactions have recently been identified as additional components of RNA-mediated regulatory networks (76). Indeed, a ncRNA-sponge function has been demonstrated for several bacterial ncRNAs (77–79). Thus, in addition to mRNA targets, ncRNAs must be included as potential targets of regulatory RNAs. We thus extended our IntaRNA analysis to ncRNA identified as enriched by RIP-seq (Supplementary Table S10). This analysis revealed more than 300 potential interactions between identified ncRNAs. Intriguingly, the strongest predicted interactions involved CRISPR RNA regions including crRNA spacers targeting riboswitches, IGR sRNAs or antisense RNAs (Supplementary Figure S14). Obviously, these *in silico* predictions will require experimental validation but still suggest a complex regulatory RNA network in *C. difficile*.

Most of the genes belonging to the SigK regulon are overexpressed in the strain depleted for Hfq suggesting a link between this RNA chaperone and the sporulation control in *C. difficile* (27). An over-sporulation phenotype is observed for this depleted strain and a Δ*hfq* mutant in *C. difficile* (70). In line with these data, the entire SigK regulon, *sigK* and *spoIIID* genes are overexpressed in these conditions (27). SpoIIID, which positively controls *sigK* expression, is located upstream of SigK in the sporulation regulation cascade. The *spoIIID* gene as well as the *sigK* gene are transcribed by the sigma factor SigE (80). As no global effect of Hfq on the SigE regulon was observed (27), we can exclude SigE. Hfq rather controls the late stages of sporulation by acting either at the SpoIIID or SigK level. From *in silico* target predictions, previous phenotypic and interaction studies and present RIP-seq data, RCd1 appears among Hfq-binding ncRNAs as a good candidate to mediate this Hfq-dependent control. Indeed, RCd1 overexpression has a strong inhibitory effect on the sporulation process and on the SigK regulon expression.

As in *B. subtilis*, the *sigK* gene is interrupted by a *skin*^*Cd*^ element in most of *C. difficile* strains. The excision of this phage-like element is necessary to trigger SigK production and is carried out by the recombinase CD1231 in complex with the recombination directionality factor, CD1234 (33). *CD1231* and *CD1234* are the only two genes of the *skin*^*Cd*^ element conserved in all strains of *C. difficile*. We demonstrated in this work that deletion of the *skin* element abolished the impact of RCd1 overexpression on sporulation and that RCd1 overexpression negatively affects *skin* excision in the wild-type strain under sporulation conditions. These results strongly suggest that RCd1 act at the level of *skin* excision or upstream of this step. In the regulatory cascade of sporulation, we then propose to include an additional RNA regulatory element, RCd1, that can trigger *sigK* expression and SigK production by promoting *skin* excision. The excision of the *skin* element in the mother cell occurs at a late stage of the sporulation process and requires the specific expression of *CD1234* under the dual control of SigE and SpoIIID (33). We propose that RCd1 in concert with Hfq modulates the expression of *spoIIID* but also of *CD1234* either directly or indirectly through a negative control of *spoIIID* expression. An effect on SigE is excluded since neither Hfq-depletion nor RCd1 overexpression affected SigE targets except for *spoIIID* and *CD1234* genes. RCd1 overexpression limits the amount of CD1234 protein, an essential partner of the CD1231 recombinase leading to a decrease in the recovery of a functional *sigK* gene. A direct negative effect of RCd1 on *spoIIID* is also possible and would result, in addition to its effect on CD1234, in a decrease of *sigK* transcription affecting the entire SigK regulon and thus the sporulation efficiency. Our *in vitro* interactions studies showed that RCd1 could bind *spoIIID* and *CD1234* mRNAs. We were able to show that Hfq could interact with each RNA individually but was not required for RCd1-mRNA interactions *in vitro*. Additional missing factors or particular conditions could affect these interactions during sporulation in *C. difficile*. An Hfq-independent action of RCd1 could also be envisaged. The signals or regulatory elements affecting the RCd1 action on *sigK* network as well as the possible role of the 100-nt antisense RNA for RCd1 remain to be studied. A complex regulatory network governs SigK activation during the late stages of sporulation with SpoIIID- and SigK-dependent regulatory feedback loops and the participation of the SigE sigma factor. As for other ncRNAs, RCd1 could have a large spectrum of potential targets contributing to different facets of *C. difficile* physiology. In accordance, RIP-seq and *in silico* gene co-occurrence analysis suggests a larger role of RCd1 in regulatory processes beyond the *skin*^*Cd*^ excision control.

Several examples of bacterial RNA-based regulation of sporulation have recently been reported. In *C. perfringens*, the VirX ncRNA has been shown to negatively control the sporulation process. However, unlike RCd1 inhibiting late stages of sporulation, VirX inhibits sporulation very early, likely at the Spo0A level. This ncRNA is conserved in several clostridial species but absent in *C. difficile* or *C. tetani* (81). In *B. subtilis*, several ncRNA dependent on sporulation sigma factors have been identified, including SurA, SurC and CsfG, but their role is not yet defined (82–84). In *Bacillus anthracis*, one of these sporulation-specific sRNA, CsfG, conserved in *Bacillaceae* and some other endospore formers but absent in *C. difficile*, has been shown to bind Hfq1, one of the Hfq homologues present in *B. anthracis* (85) suggesting a possible implication of Hfq in regulatory processes during sporulation. In *B. subtilis*, sporulation was not altered in the Δ*hfq* strain (86), although CsfG was also identified among Hfq-associated sRNAs by coimmunoprecipitation analysis (87). A more specialized function connected with stationary phase physiology has recently been suggested for Hfq in *B. subtilis*. However, the Hfq-related stationary fitness phenotype could not be linked to sporulation (88). As an example of an Hfq-independent RNA contributing to sporulation control, one of the two *B. subtilis* 6S RNAs (6S-1 RNA) interacting with the housekeeping RNA polymerase in complex with SigA, was shown to be involved in the appropriate timing of the onset of endospore formation (89). Overall, the diversity of RNA-based mechanisms controlling sporulation largely remains to be explored with specific actors that could be found in different endospore formers.

In addition to classical *trans*-encoded RNAs that were expected to act in concert with Hfq, our RIP-seq analysis revealed new RNA categories of Hfq partners. *Cis*-antisense RNAs were among the most abundant classes of regulatory RNAs enriched in Hfq-immunoprecipitated samples. This RIP-seq analysis allowed us to describe multiple new regions of double-directional transcription in *C. difficile* chromosome suggesting the involvement of Hfq in these complementary RNA interactions. Emerging examples suggest the requirement of Hfq or other RNA chaperones for antisense RNA pairing with their targets (90,91). For example, Hfq facilitates the interaction of the antisense RNA RNA-OUT with its target RNA-IN in the Tn10/IS10 transposon system (91) and FinO is required for the FinP antisense RNA pairing with *traJ* mRNA in the F plasmid of *E. coli* (90).

The first functional antisense RNAs described in *C. difficile* act as antitoxins within type I TA systems (18–20). In the present study, both toxin mRNAs and antitoxin antisense RNA for all 13 type I TA modules were highly enriched in Hfq-immunoprecipitated samples. The possible role of Hfq in stabilizing type I TA interactions was suggested in our previously reported work in *C. difficile* (18). Similarly, antitoxins and toxins from type I TA modules were also enriched in Hfq coIP samples in *B. subtilis* and Hfq was able to bind antitoxins *in vitro* (87). While Hfq stabilizes SR5 antitoxin in this bacterium, it does not affect the *bsrE* toxin RNA of the type I TA module (92) and was generally not required for the function of type I TA systems (86,93). Further studies will clarify if Hfq could in some way contribute to the function of TA in *C. difficile.*

In light of our RIP-seq analysis, a potential role of Hfq in uncommon regulatory functions associated with crRNAs from CRISPR arrays and with premature terminated riboswitch transcript could be suggested. Several examples of noncanonical role of such ncRNA have been reported in other bacteria (94). However, the implication of Hfq protein has never been explored in these regulatory processes. Indeed, SAM-riboswitch-associated premature terminated transcript has been shown to act *in trans* as an independent regulatory RNA controlling the expression of *prfA* gene encoding the major virulence regulator in *L. monocytogenes* (94). The detection of multiple riboswitches from different functional groups as Hfq-binding RNAs in the present study and their large targeting potential lead to a tempting speculation about their regulatory actions through base-pairing that needs to be further explored. Similarly, a large class of regulatory RNA elements located within 5’-leader regions has been revealed among Hfq-associated RNAs in *B. subtilis* (87). These *cis*-acting regulatory RNAs respond to a variety of ligands including metabolites, tRNAs and metal ions generally through the mechanism of premature termination of transcription in many Gram-positive bacteria. Their association with Hfq opens an interesting question about the possible sRNA-like action of these abundant terminated transcripts.

The targeting of endogenous RNAs has recently been demonstrated for Cas9 protein of *Campylobacter jejuni* guided by native crRNA through imperfect complementarity (58). Other examples of CRISPR-Cas regulatory function include the down-regulation of expression of the immunostimulatory BLP lipoprotein in *Francisella novicida* by type II-B Cas9 through tracrRNA in association with another small CRISPR-Cas-associated RNA (scaRNA) by an unknown mechanism (95–97). The great potential of *C. difficile* crRNA targeting revealed in our work is in line with these findings, further suggesting a potential regulatory role for CRISPR-Cas systems in bacterial pathogens. Indeed, potential functions beyond defense and in relation with virulence control in several pathogens emerged from several recent studies (59,98–102). However, the exact mechanism implicating crRNA targeting for regulatory purposes remains the subject of debate. For example, in contrast to the previously reported regulatory potential for type I-F CRISPR-Cas in *Pseudomonas aeruginosa* (98), more recent studies did not find evidence of RNA targeting (103,104). The possible implication of Hfq RNA chaperone in crRNA interactions with potential mRNA targets implies that crRNAs could be additional components in the regulatory RNA networks and suggests an alternative mechanism for crRNA regulatory actions independent from the Cas machinery. Discontinuous base-pairing of spacer-containing regions with potential mRNA targets and the absence of functional PAM in the proximity of spacer-targeted zone are further in favor of the particular mechanism for potential crRNA regulatory functions reminiscent to the *trans*-encoded sRNA-mediated action.

In Gram-negative bacteria, the Hfq protein generally binds to AU-rich single-stranded regions upstream of the rho-independent terminator in sRNAs and could also interact with the 3′ poly(U) tail (74). The AT-rich nature of *C. difficile* genome could provide a large high-affinity interaction potential for efficient Hfq protein association with ncRNAs. Widespread post-transcriptional mechanisms of Hfq-dependent ncRNA action are linked to the mRNA RBS binding leading to the repression of translational initiation. Our RIP-seq analysis combined with *in silico* base-pairing prediction for Hfq-associated RNAs revealed a number of potential ncRNA-mRNA interactions that could fit this mechanism. Interestingly, potentially accessible short C-rich motifs could be involved in these interactions in *C. difficile*. In *Staphylococcus aureus,* another Gram-positive pathogen with a low GC-genome content, a class of ncRNAs has been identified that carries a conserved unpaired C-rich motif for targeting mRNAs RBS regions and that could repress the translation by preventing the formation of ribosomal initiation complex (105–107). Even though Hfq was not shown to be involved in these regulatory mechanisms in *S. aureus*, the presence of C-rich motifs could constitute a general hallmark for efficient mRNA targeting by ncRNAs.

## Conclusion

The present study provides the first genomic map of RNAs interacting with Hfq in *C. difficile* and paves the way for future detailed analysis of RNA-based mechanisms in this human enteropathogen. RIP-seq data analysis expands our knowledge on the great interaction potential of the Hfq RNA chaperone in a Gram-positive bacterium opening interesting perspectives for future studies and leads to the discovery of a number of new antisense RNAs and IGR ncRNA further enriching the repertoire of regulatory RNAs in *C. difficile*. Together with previous genome-wide identification of ncRNAs by TSS mapping, this global approach further confirms the importance of RNA-based mechanisms in *C. difficile*. The discovery of regulatory RNA functions still stays in its early stage in *C. difficile*. The present work provides an essential molecular basis on the repertoire of Hfq-binding RNAs and their interaction potential that will foster future studies on the post-transcriptional regulatory mechanisms to better understand the emergence and success of this important human enteropathogen.

## ACCESSION NUMBERS

RIP-Seq data have been deposited in ENA with accession no. PRJEB39335.

## SUPPLEMENTARY DATA

Supplementary Data are available online.

## ACKNOWLEDGEMENT

We are grateful to Joël Caillet for his kind help in Hfq protein purification and helpful discussions.

