## Supplementary material for "Identification of RNAs bound by Hfq reveals widespread RNA partners and a sporulation regulator in the human pathogen *Clostridioides difficile*"

**Figure S1. Detection of Hfq FLAG-tagged and untagged Hfq by Western blot.**

Immunoblotting with anti-Hfq antibody detected the native Hfq (~10 kDa) and Hfq-3xFLAG-tagged protein expressed from a plasmid (~13 kDa) in whole cell extracts in the 630/p control strain carrying an empty vector and 630/p-Hfq3xFLAG strain grown in TY medium in the presence of ATc inducer (250 ng/ml). No signal for Hfq-FLAG-tagged protein could be detected in the control strain and the native Hfq levels were similar between 630/p and 630/p-Hfq-3xFLAG strains. In 630/p-Hfq-3xFLAG strain, the Hfq-3xFLAG-tagged protein was strongly expressed, leading to the saturation of the signal. Using ImageJ software, we estimated that in 630/p-Hfq3xFLAG strain, Hfq3xFLAG levels were at least 10-fold higher compared to those of the native untagged Hfq. The quantity of loaded protein extracts was adjusted in each well to represent a final culture $OD_{600}$ of 7.5 and proteins were separated on a 4-12% gradient Bis-Tris polyacrylamide gel with MES running buffer. The experiment was done in triplicate.

**Figure S2. PCA Clustering of samples for biological replicates of RIP-seq analysis.**

Principal component analysis reveals clustering of four biological replicates F1, F2, F4, F5 for 3xFLAG-Hfq coIP shown in red and of four biological replicates of C1-C4 control samples shown in blue apart from Hfq-coIP and PI pre-immune serum control in wild-type strain.

**Figure S3. Diagram of relative proportion of different RNA species in Hfq coIP and in control PI sample in wild-type strain.**

All sequences that mapped to the *C. difficile* genome are represented. The rRNA and tRNA abundant housekeeping RNAs are shown in orange, the reads mapping to CDS are shown in gray. The relative proportion of other RNA species including known previously identified regulatory RNAs (named « misc_RNA ») and IGR is detailed in the right diagram for Hfq coIP sample. Left panel: control PI (pre-immune serum) coIP, right panel : Hfq coIP.

**Figure S4. The expression profile of an antisense RNA to RCd1.**

RNA samples were extracted from 630Δ*erm* strain (630) grown in SM medium for 10 h, 12 h, 14 h, 16 h, 18 h, 20 h, 22 h, 24 h as indicated. The RNA samples from mutant strains for sporulation regulatory genes *sigE, spo0A* and *spoIIID* were also included. 5S RNA at the bottom serves as loading control. As indicated at the left, the blots were hybridized with antisense (AS) RCd1-specific probe. The same 5S control panel is shown when reprobing of the same membrane was performed.

**Figure S5. Representative RIP-seq examples of enriched RNAs.**

Different functional groups of RNAs are presented in A) sRNAs, B) TA loci, C) other antisense RNAs, D) CRISPR RNAs, E) riboswitches, F) new potential ncRNAs, G) mRNAs. The IGV visualization is presented as in Figure S1 and Figure 2.

**Figure S6. Detection of ncRNAs by Northern blot.**

Northern blot was performed for detection of selected ncRNAs. RNA samples were extracted from 630Δ*erm* strain grown at exponential phase (**E**, 4 h of growth), late-exponential phase (**LE**, 6 h of growth), entry to stationary phase (**S**, 10 h of growth) or under nutriment starvation conditions (**St**), from R20291 strain grown at late-exponential phase (**LE**), from strains CDIP369 (**630/p**), CDIP53 strain expressing an antisense RNA for the *hfq* gene (**AS *hfq***), CDIP55 strain expressing an antisense RNA for the *rnJ* gene encoding RNase J (**AS *rnJ***) and CDIP57 strain expressing an antisense RNA for the *rny* gene encoding RNase Y (**AS *rny***). 5S RNA at the bottom serves as loading control. The arrows show the detected transcripts with their size estimated by comparison with RNA molecular weight standards.

**Figure S7. Comparison between Hfq-FLAG RIP-seq enriched peaks and differentially expressed genes identified by previously published transcriptomic analysis of *hfq*-depleted strain.**

Euler diagram representations of the number of mRNAs and ncRNAs identified as Hfq ligands by RIP-seq ("peak") and corresponding to genes differentially expressed in transcriptomic analysis ("DEG") (https://cran.r-project.org/package=eulerr). The information on up- and -down-regulated genes is presented as a separate diagram (bottom panel).

**Figure S8. Hfq RIP-seq data visualization for *C. difficile skin* region.**

An overall view of genomic region for *skin* locus is shown on the top. The IGV visualization of RIP-seq data for *CD1231, CD1233, CD1234* genes encoded inside the *skin* is presented, as well as RIP-seq data for *spoIIID* gene at the bottom panel. The IGV visualization is presented as in Figure S1. Green arrows pinpoint the *CD1234* and *spoIIID* regions.

**Figure S9. Analysis of the interaction of Hfq with RCd1, *spoIIID* and *CD1234***

5′ end-labeled RNA was mixed with increasing concentrations of Hfq ranging from (A and B) 50 pM to 100 nM or (C) from 100 pM to 50 nM, expressed on the basis of the monomer form. Half-saturation values ($K_{1/2}$) were calculated as described in Material and Methods and indicated below each band-shift experiment.

**Figure S10. Multiple alignment of nucleotide sequences of RCd1 homologues using MEGA 7 software.**

**Figure S11. Correlation between the presence of RCd1 and *skin* element in *C. difficile* strains.**

Sequence conservation of the *skin* element (Y-axis) is represented as a function of RCd1 sequence conservation (X-axis) in analyzed *C. difficile* strains.

**Figure S12. MEME motif search results**

Consensus motif for a subset of Hfq-associated ncRNA peaks excluding peaks corresponding to CRISPR, type I TA and c-di-GMP-responsible riboswitch RNAs is shown. 57 analyzed ncRNA are listed with potential motif highlighted in colour.

**Figure S13. Examples of potential ncRNA-mRNA interactions**

Each box shows the base-pairing interaction for an Hfq-bound sRNA with its predicted mRNA target. The interaction energy predicted by IntaRNA program is indicated at the top of each box. The initiation codon and RBS site are indicated in red in mRNA sequence. The spacer sequence in CRISPR RNA is depicted in blue and the trinucleotide at the position of PAM in targeted sequence is indicated in green.

**Figure S14. Examples of potential ncRNA-ncRNA interactions**

Each box shows the base-pairing interaction for an Hfq-bound sRNA with its predicted sRNA target. The interaction energy predicted by IntaRNA program is indicated at the top of each box.

**Table S1. Strains, plasmids and oligonucleotide primers used in this study**
**Table S2. Number of mapped reads per RIP-seq sample**
**Table S3. Complete list of MACS peaks with MAGe annotation**
**Table S4. List of MACS peaks with MAGe annotation for ncRNAs**
**Table S5. Transcriptome data comparison with RIP-seq**
**Table S6. Pathway clustering for mRNAs peaks detected by RIP-seq (KEGG)**
**Table S7. RIP-seq data validation by qRT-PCR**
**Table S8. RCd1 conservation in 2,700 available *C. difficile* genomes and co-occurrence of RCd1 homologues with *skin* element, and important regulatory or *skin* excision components *spoIIID*, *CD1231* and *CD1234*.**
**Table S9. IntaRNA prediction for ncRNA-mRNA interactions**
**Table S10. IntaRNA prediction for ncRNA-sRNA interactions**

Fig. S1

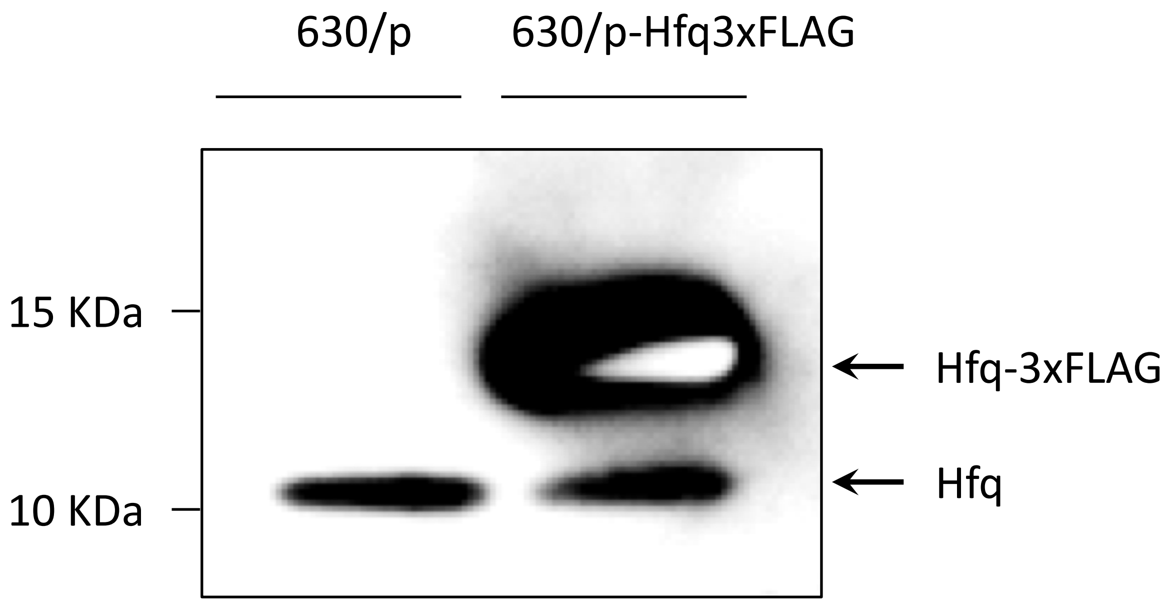

Fig. S2

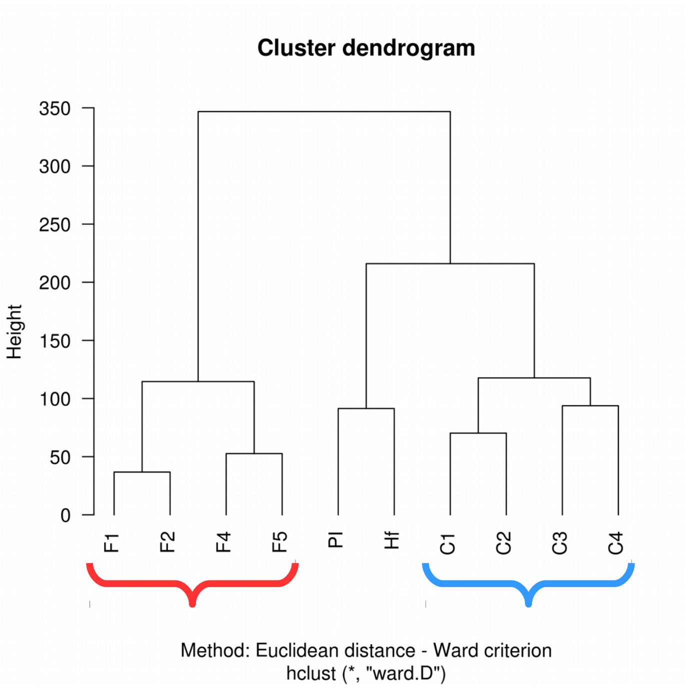
Cluster dendrogram
Height
350
300
250
200
150
100
50
0
F1
F2
F4
F5
Pl
Hf
C1
C2
C3
C4
Method: Euclidean distance - Ward criterion
hclust (*, "ward.D")

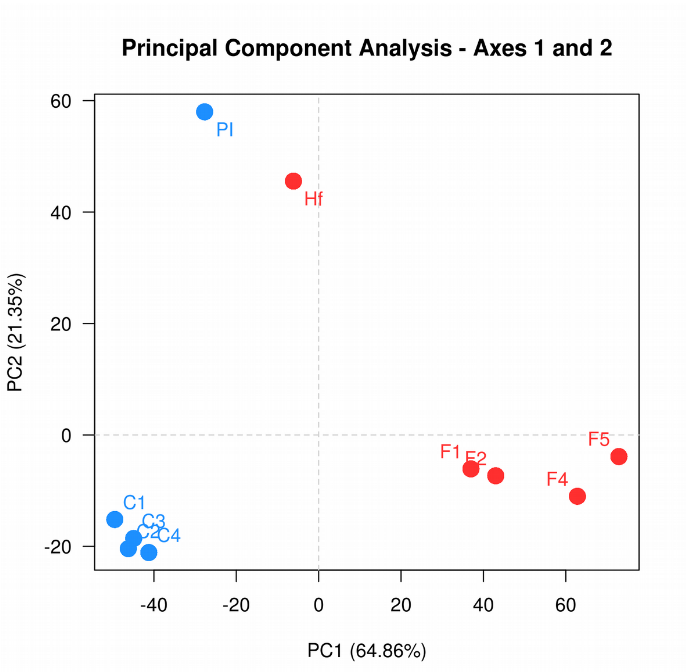
Principal Component Analysis - Axes 1 and 2
PC2 (21.35%)
PC1 (64.86%)
60
40
20
0
-20
-40
-20
0
20
40
60
Pl
Hf
F1
F2
F4
F5
C1
C3
C2
C4

Fig. S3

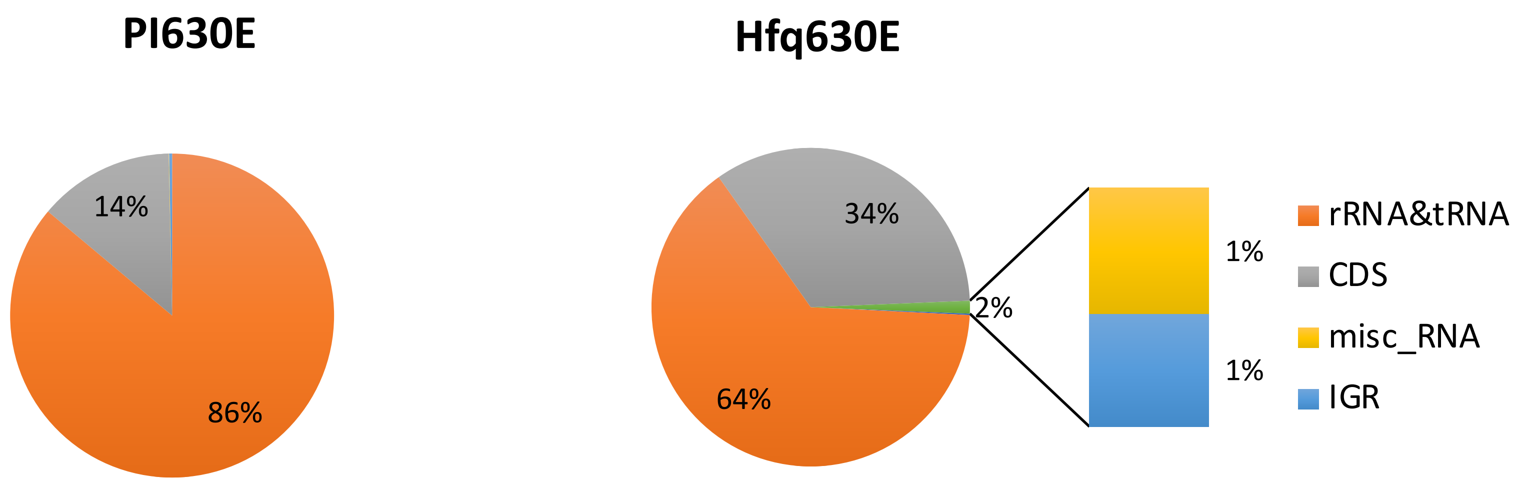

Fig. S4

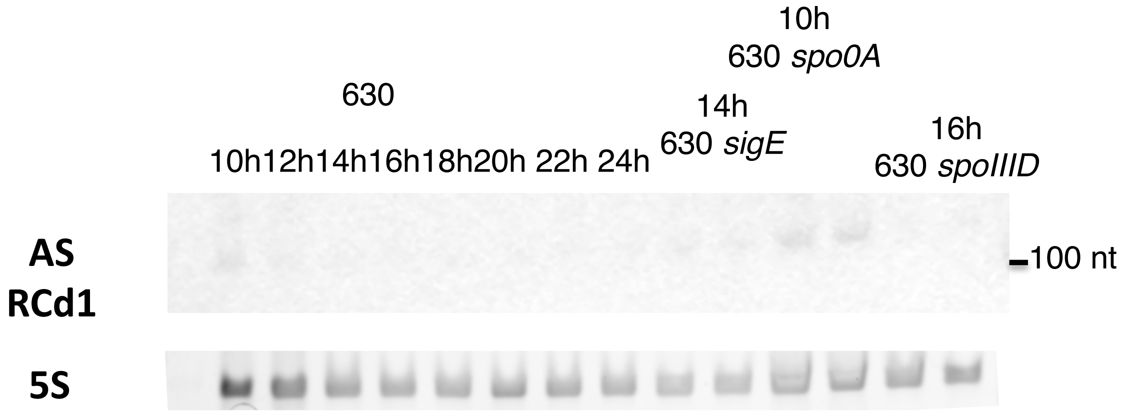
10h
630 spo0A
630
14h
630 sigE
16h
630 spoIIID
10h12h14h16h18h20h 22h 24h
AS
RCd1
100 nt
5S

Fig. S5

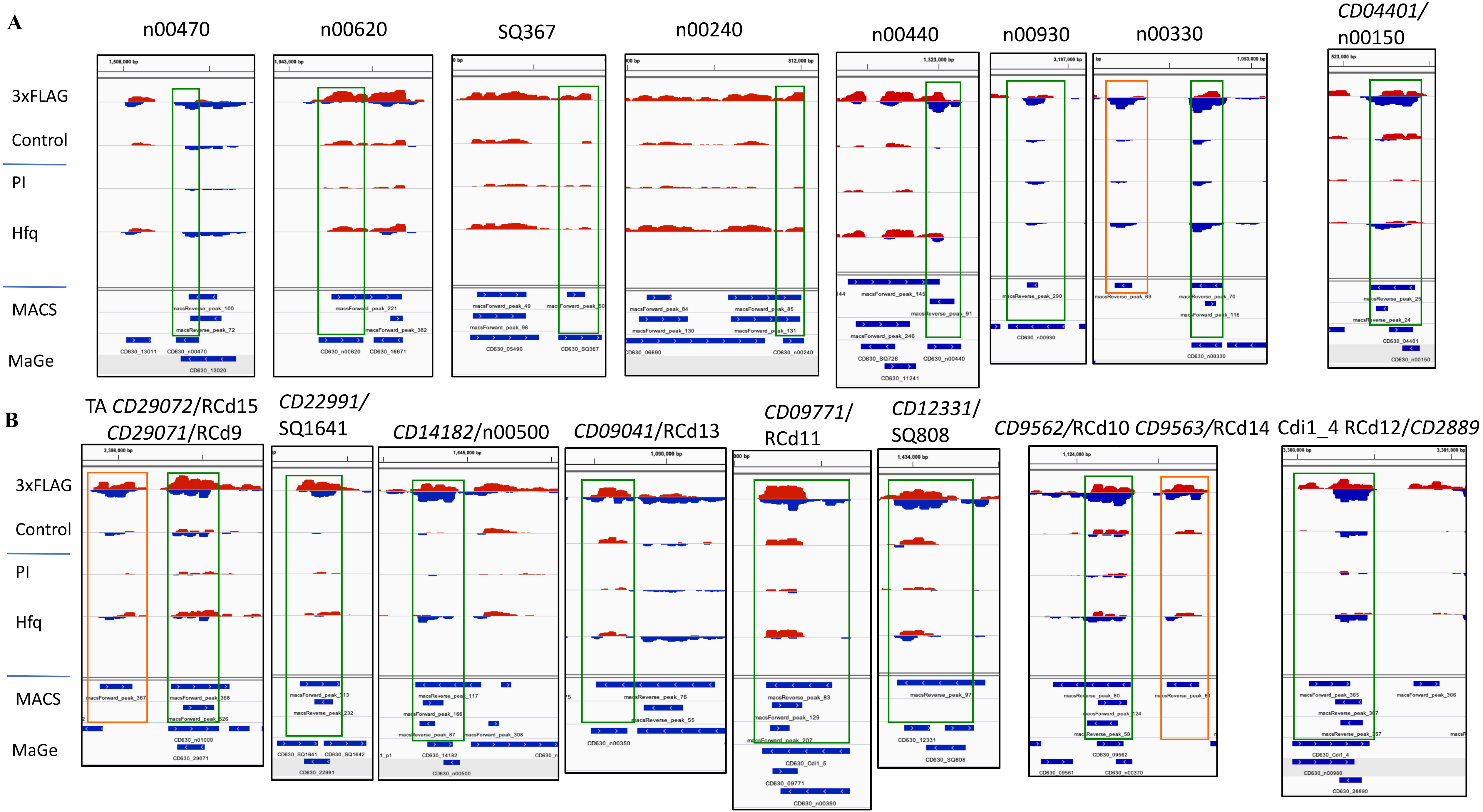
A
n00470
n00620
SQ367
n00240
n00440
n00930
n00330
CD04401/ n00150
3xFLAG
Control
PI
Hfq
MACS
MaGe
B
TA CD29072/RCd15 CD29071/RCd9
CD22991/ SQ1641
CD14182/n00500
CD09041/RCd13
CD09771/ RCd11
CD12331/ SQ808
CD9562/RCd10
CD9563/RCd14
Cdi1_4
RCd12/CD2889

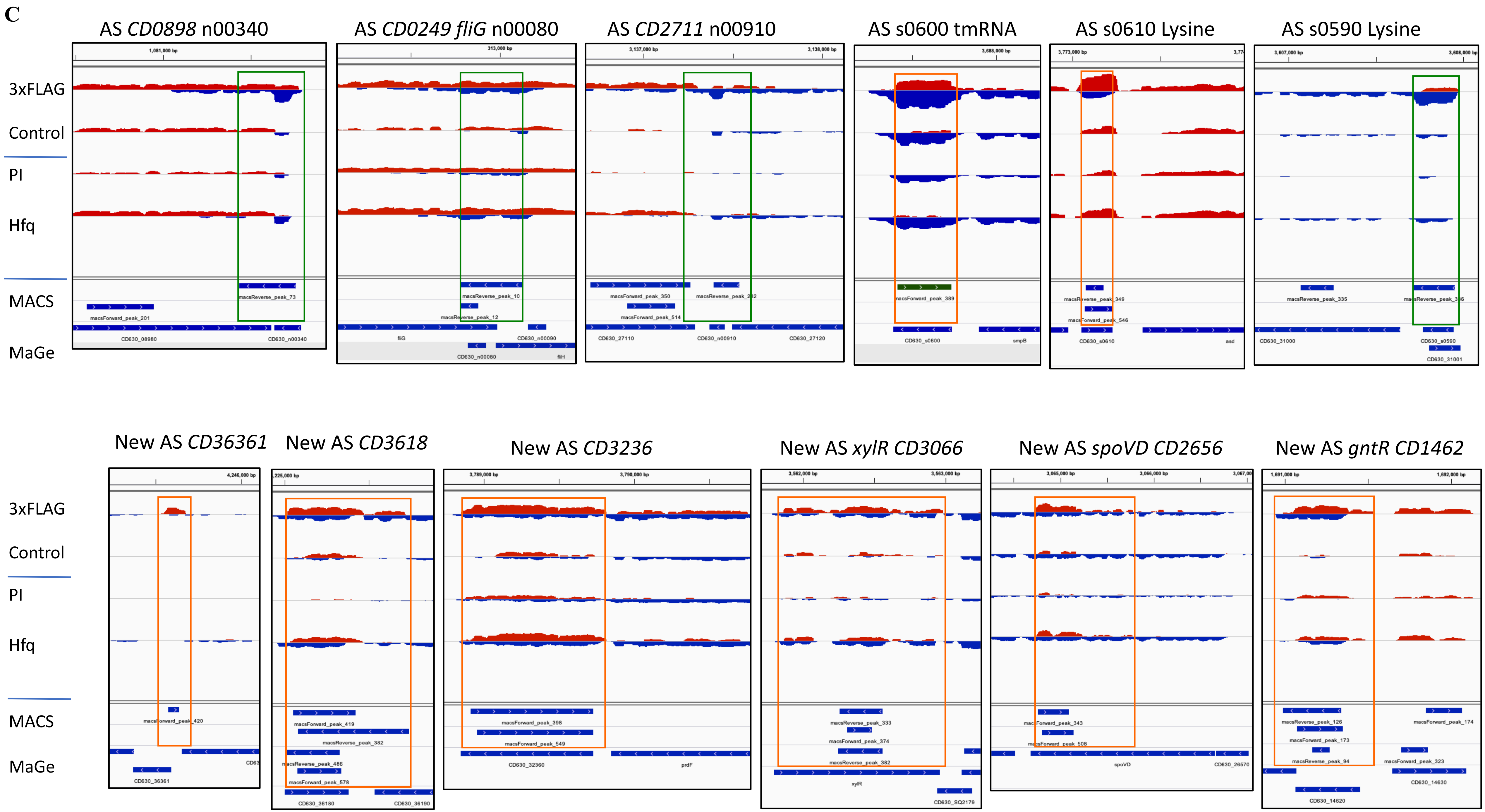
C
AS CD0898 n00340
AS CD0249 fliG n00080
AS CD2711 n00910
AS s0600 tmRNA
AS s0610 Lysine
AS s0590 Lysine
New AS CD36361
New AS CD3618
New AS CD3236
New AS xylR CD3066
New AS spoVD CD2656
New AS gntR CD1462
3xFLAG
Control
PI
Hfq
MACS
MaGe

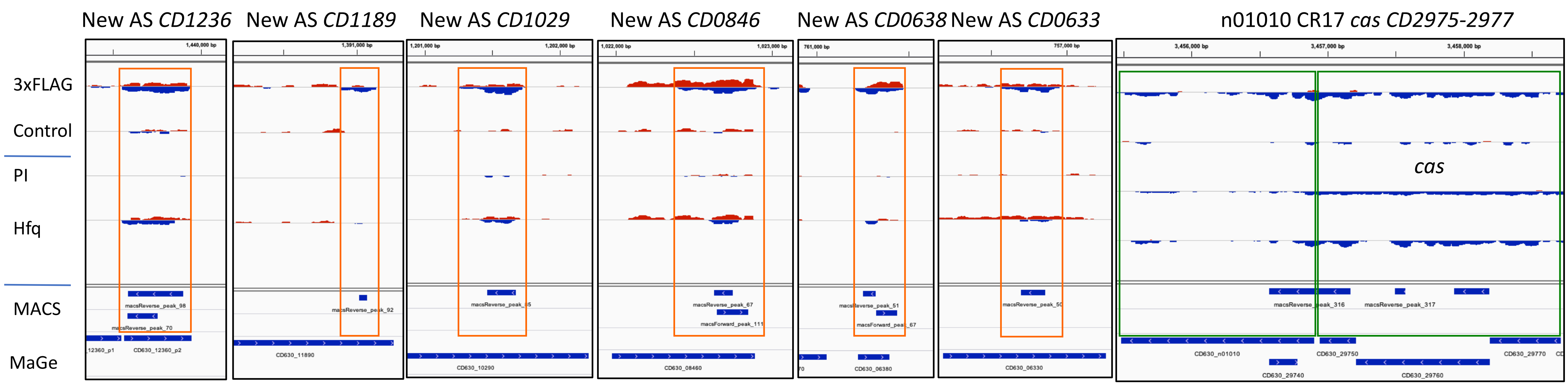
New AS CD1236
New AS CD1189
New AS CD1029
New AS CD0846
New AS CD0638
New AS CD0633
n01010 CR17 cas CD2975-2977
3xFLAG
Control
PI
Hfq
MACS
MaGe
cas

D

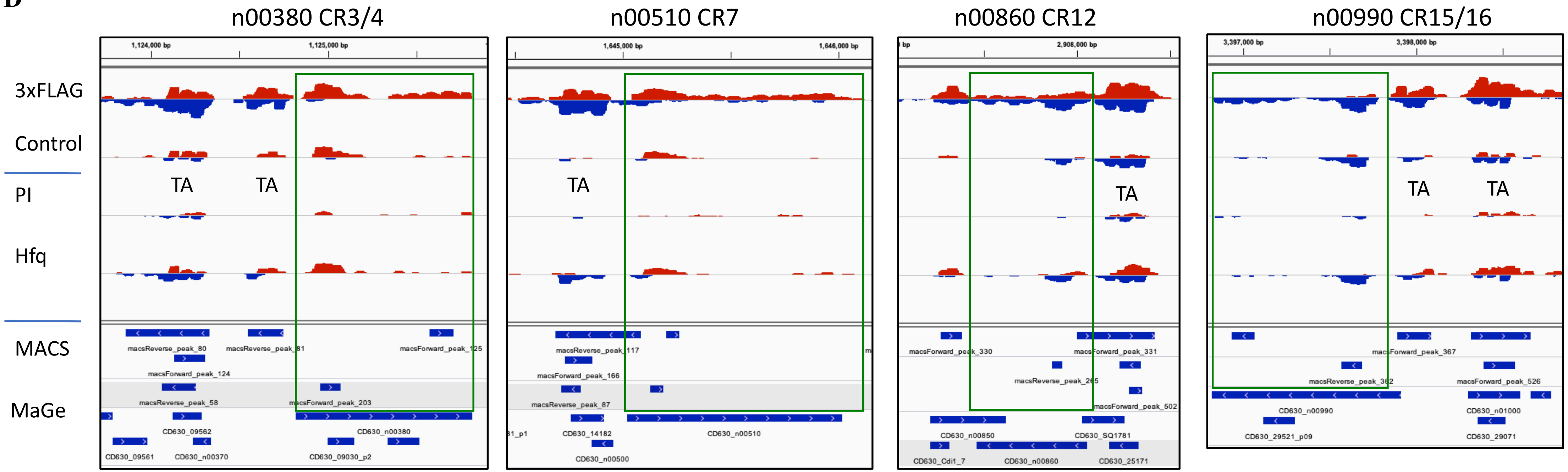
n00380 CR3/4
n00510 CR7
n00860 CR12
n00990 CR15/16
3xFLAG
Control
PI
Hfq
MACS
MaGe
TA
TA
TA
TA
TA
TA

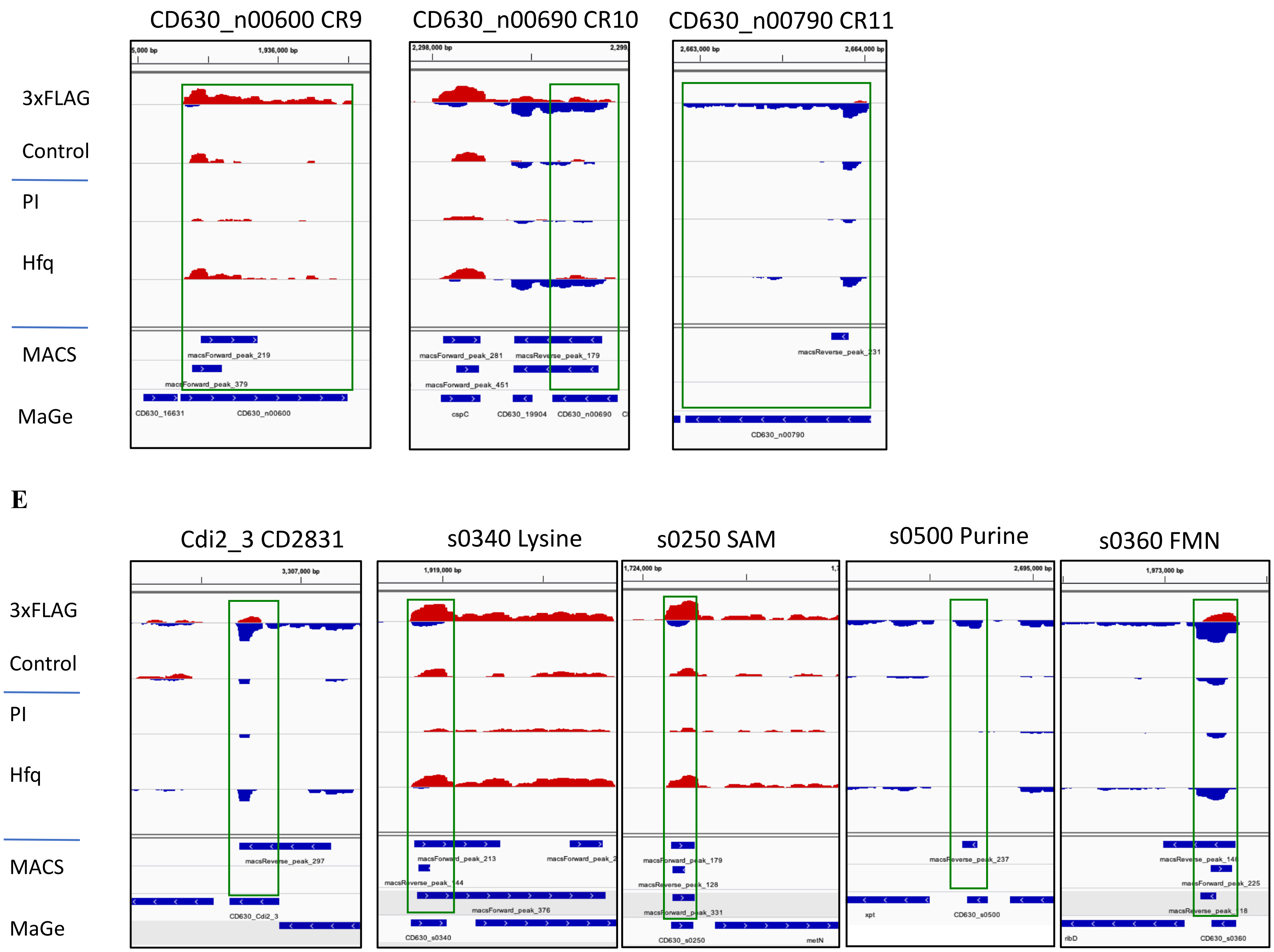
CD630_n00600 CR9
CD630_n00690 CR10
CD630_n00790 CR11
3xFLAG
Control
PI
Hfq
MACS
MaGe
E
Cdi2_3 CD2831
s0340 Lysine
s0250 SAM
s0500 Purine
s0360 FMN

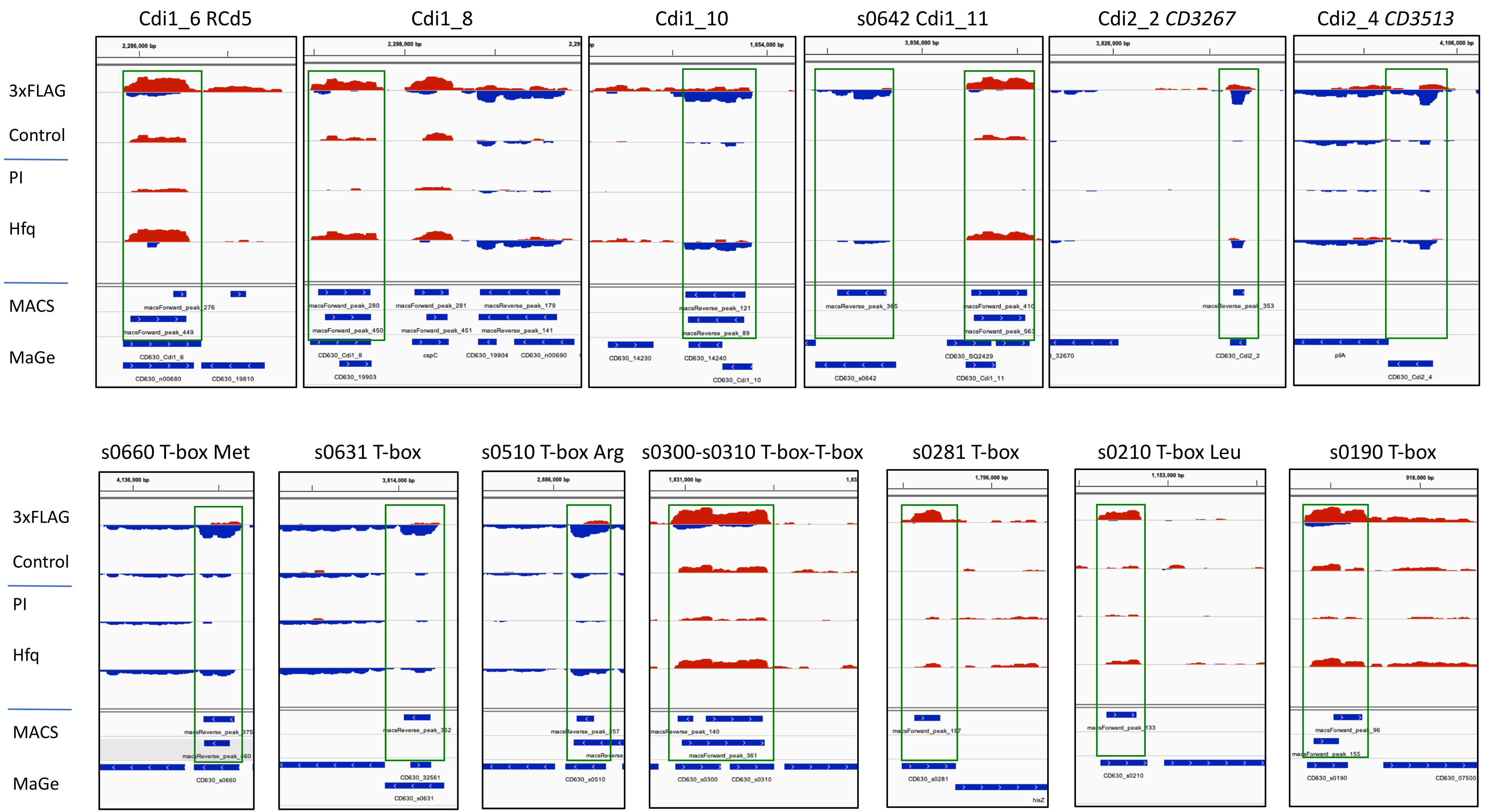

Cdi1_6 RCd5
Cdi1_8
Cdi1_10
s0642 Cdi1_11
Cdi2_2 CD3267
Cdi2_4 CD3513
3xFLAG
Control
PI
Hfq
MACS
MaGe
CD630_Cdi1_6
CD630_n00680
CD630_19810
CD630_Cdi1_8
CD630_19903
cspC
CD630_19904
CD630_n00690
CD630_14230
CD630_14240
CD630_Cdi1_10
CD630_s0642
CD630_SQ2429
CD630_Cdi1_11
CD630_Cdi2_2
pilA
CD630_Cdi2_4
s0660 T-box Met
s0631 T-box
s0510 T-box Arg
s0300-s0310 T-box-T-box
s0281 T-box
s0210 T-box Leu
s0190 T-box
CD630_s0660
CD630_32561
CD630_s0631
CD630_s0510
CD630_s0300
CD630_s0310
CD630_s0281
hisZ
CD630_s0210
CD630_s0190
CD630_07500

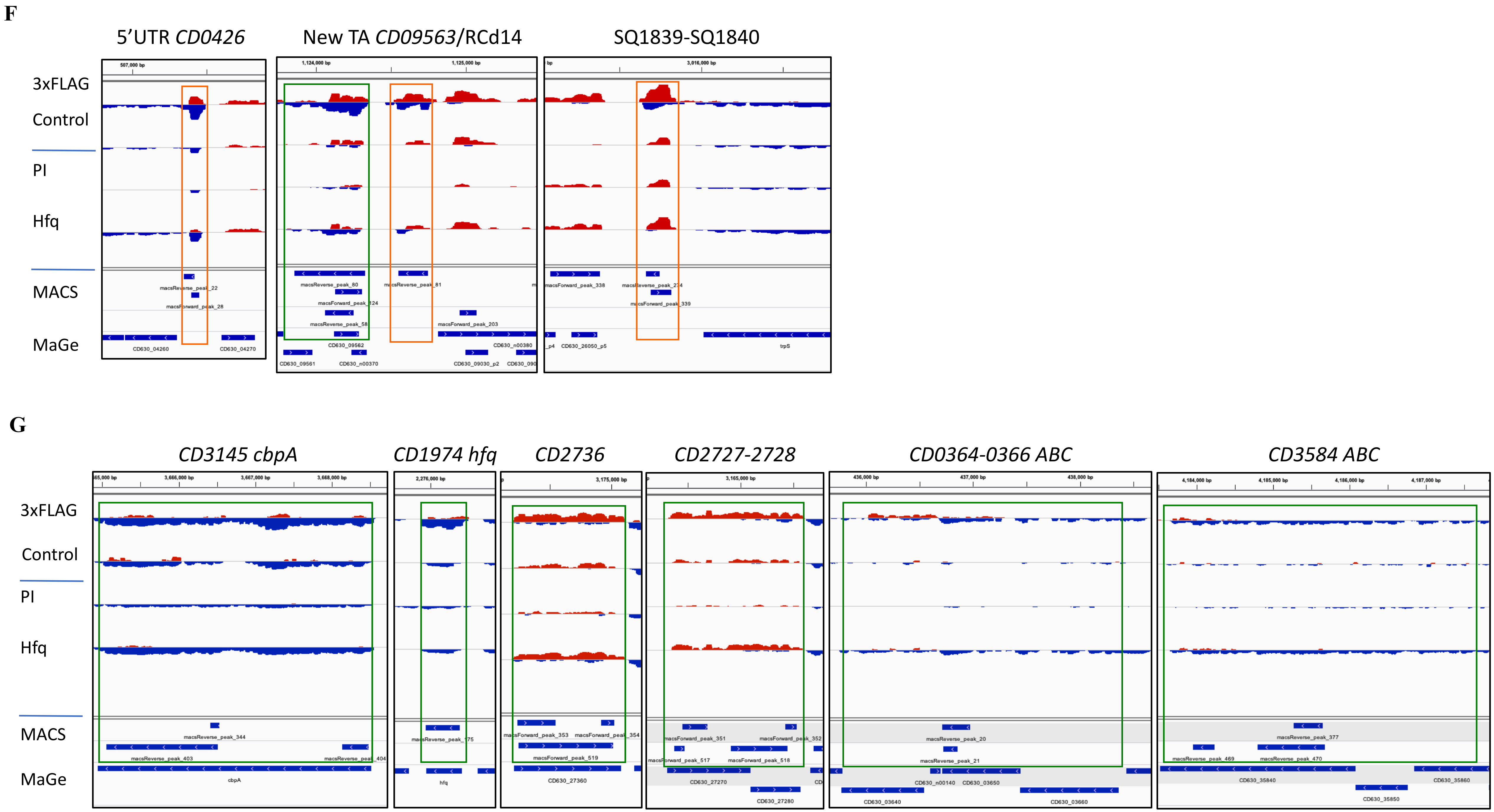

F
5'UTR CD0426
New TA CD09563/RCd14
SQ1839-SQ1840
3xFLAG
Control
PI
Hfq
MACS
MaGe
G
CD3145 cbpA
CD1974 hfq
CD2736
CD2727-2728
CD0364-0366 ABC
CD3584 ABC
3xFLAG
Control
PI
Hfq
MACS
MaGe

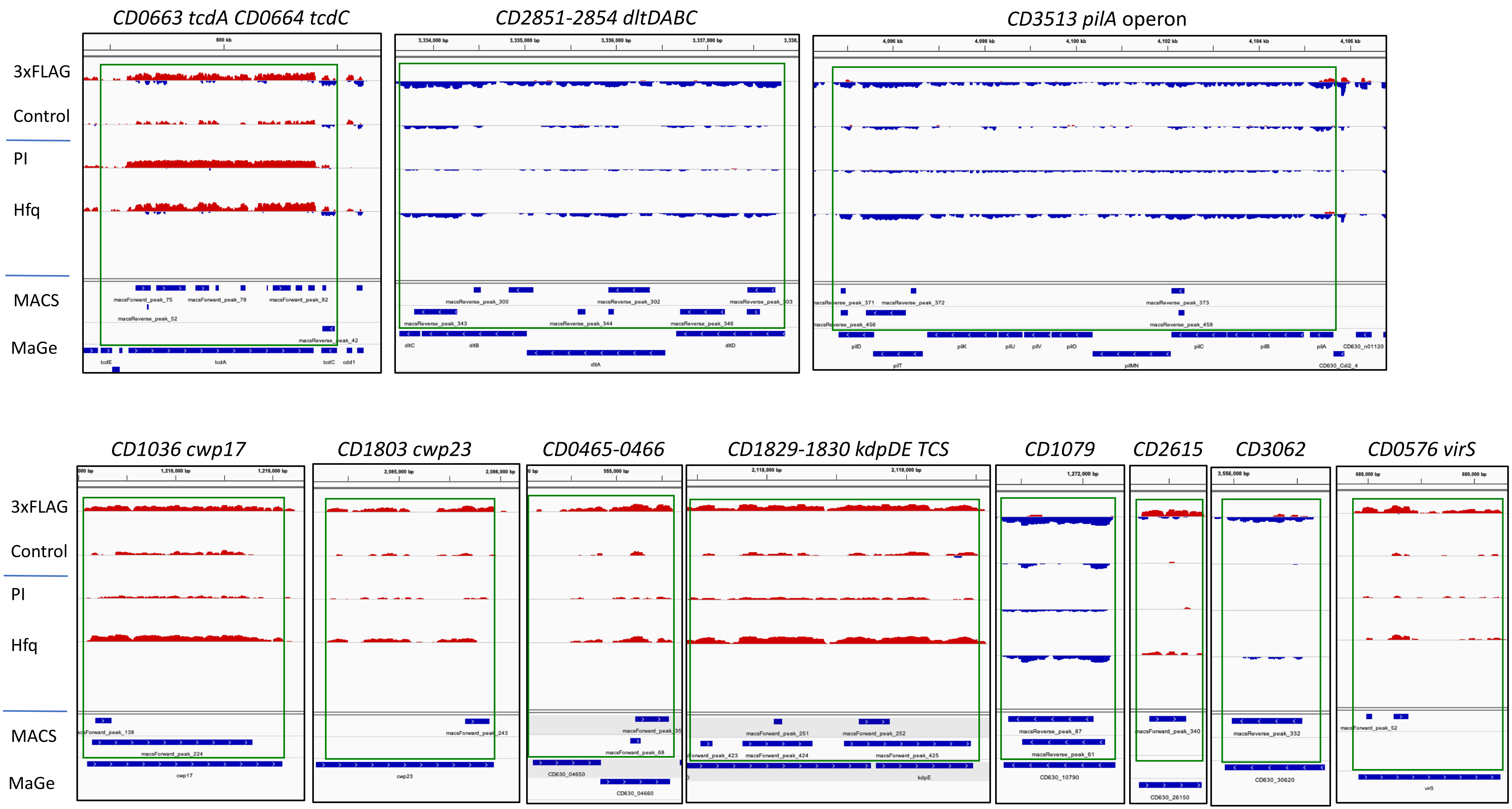

CD0663 tcdA CD0664 tcdC
CD2851-2854 dltDABC
CD3513 pilA operon
3xFLAG
Control
PI
Hfq
MACS
MaGe
CD1036 cwp17
CD1803 cwp23
CD0465-0466
CD1829-1830 kdpDE TCS
CD1079
CD2615
CD3062
CD0576 virS

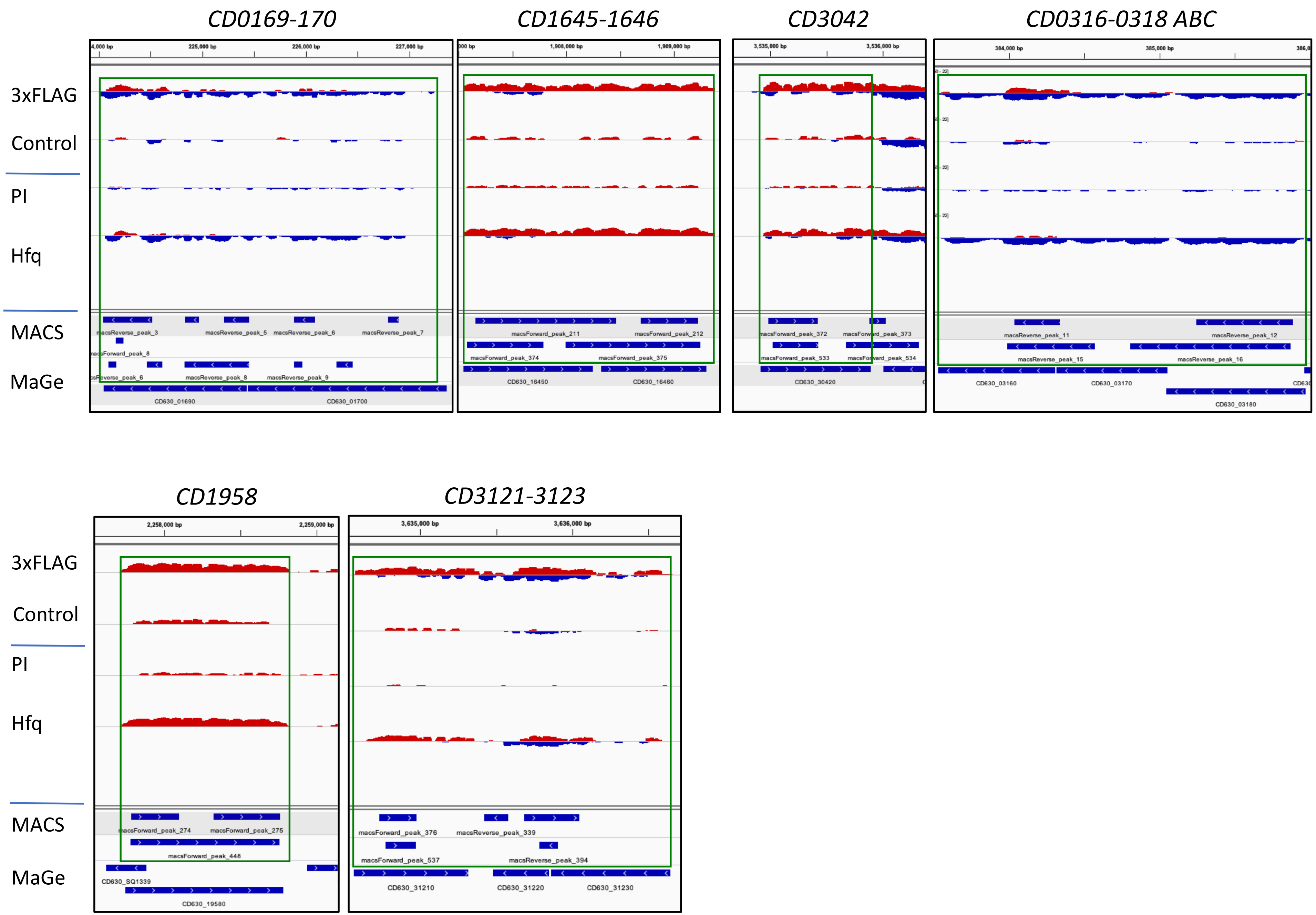

CD0169-170
CD1645-1646
CD3042
CD0316-0318 ABC
CD1958
CD3121-3123
3xFLAG
Control
PI
Hfq
MACS
MaGe

Fig. S6

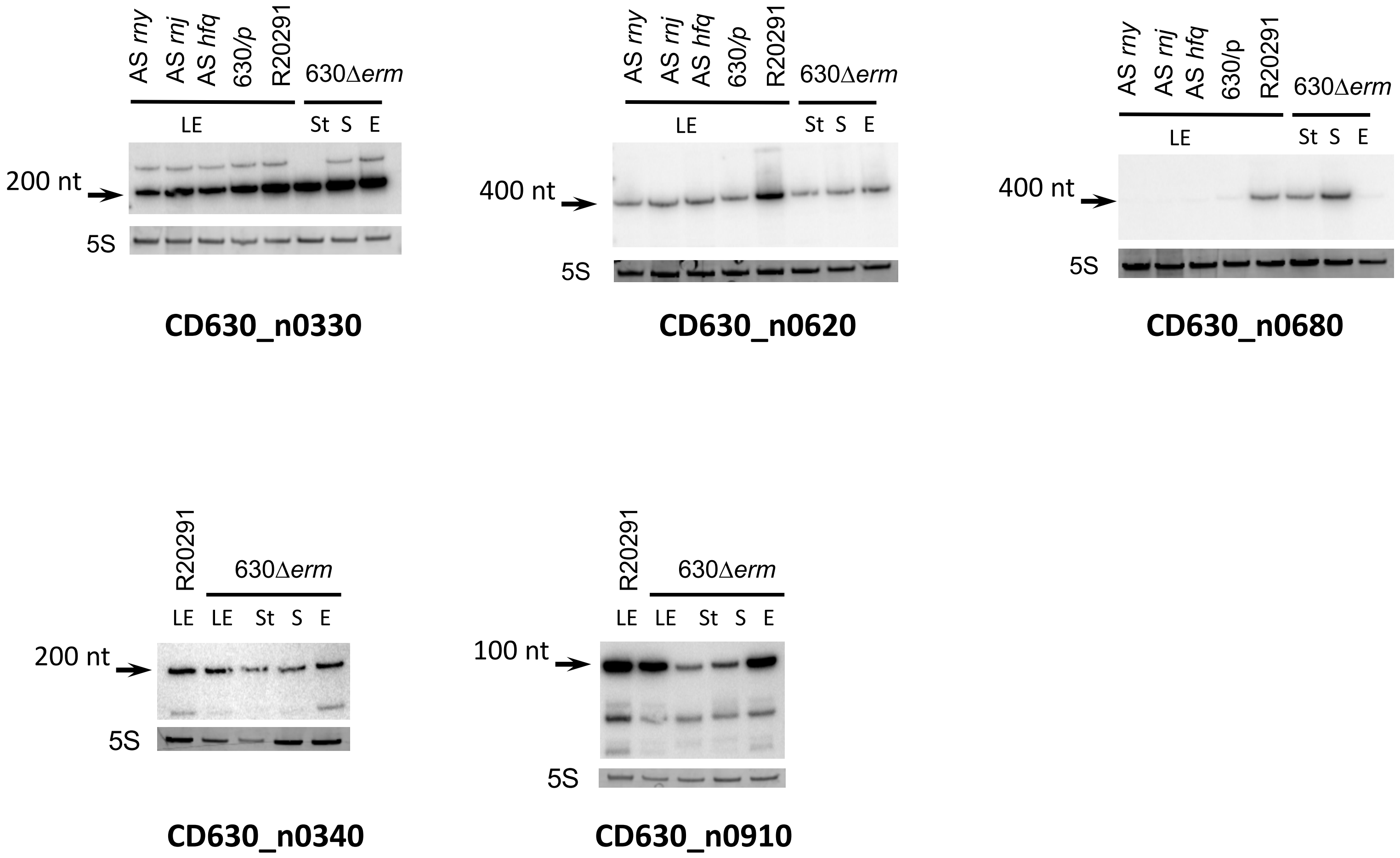

AS rny
AS rnj
AS hfq
630/p
R20291
630Δerm
LE
St S E
200 nt
5S
CD630_n0330
400 nt
CD630_n0620
CD630_n0680
R20291
LE LE St S E
CD630_n0340
100 nt
CD630_n0910

Fig. S7

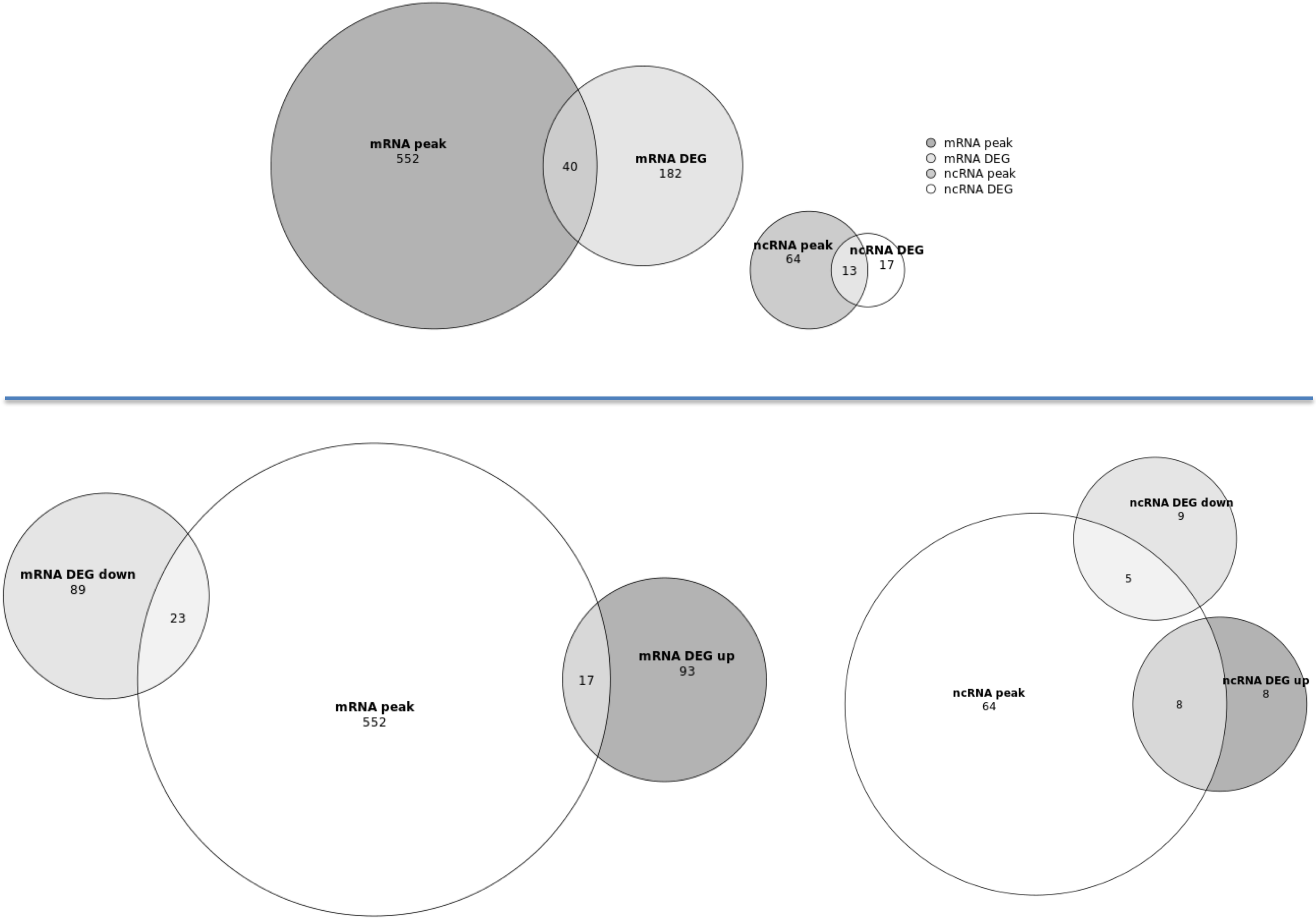
mRNA peak
552
40
mRNA DEG
182
ncRNA peak
64
13
ncRNA DEG
17
mRNA peak
mRNA DEG
ncRNA peak
ncRNA DEG
mRNA DEG down
89
23
mRNA peak
552
17
mRNA DEG up
93
ncRNA DEG down
9
5
ncRNA peak
64
8
ncRNA DEG up
8

Fig. S8

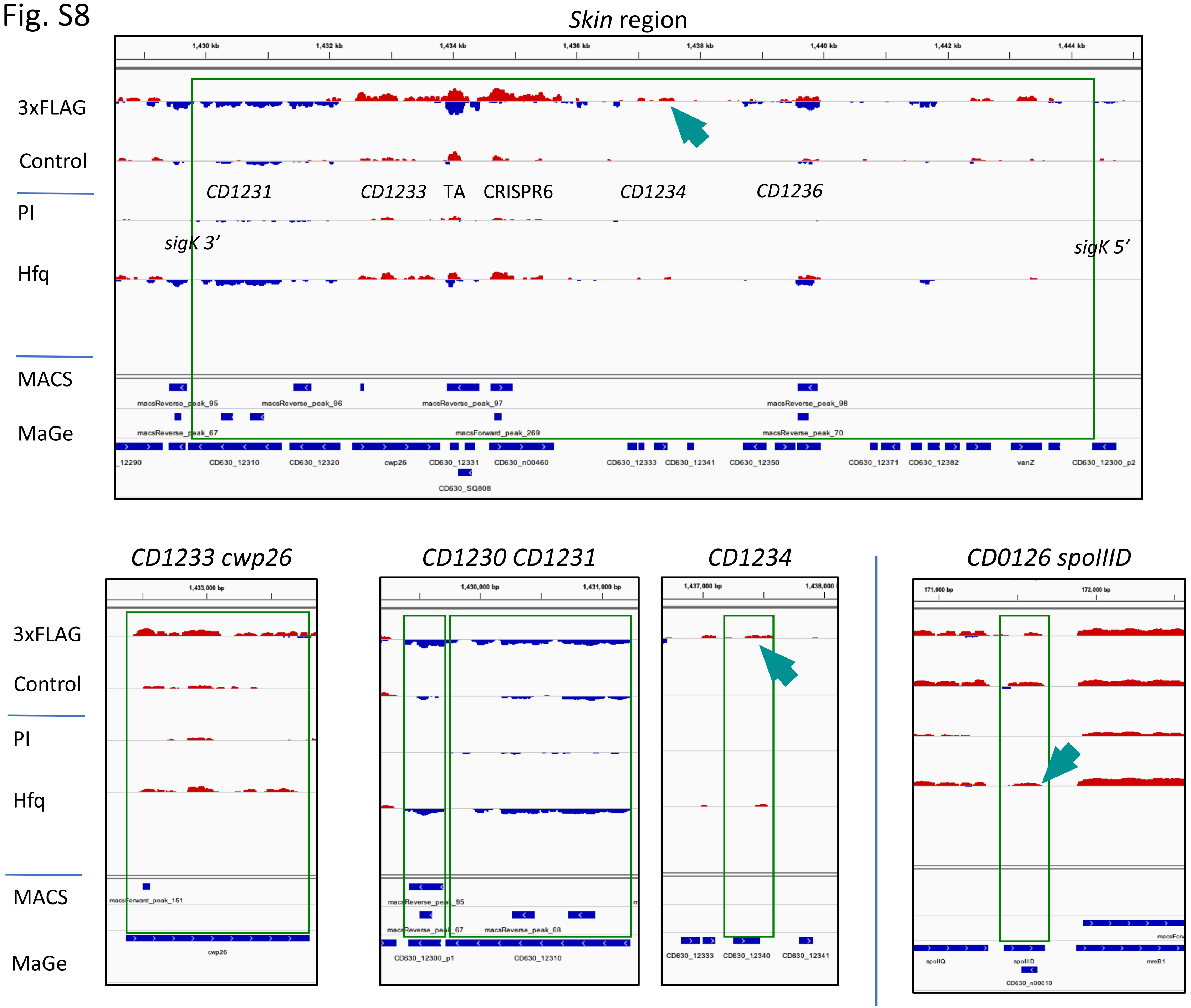

Skin region
3xFLAG
Control
PI
Hfq
MACS
MaGe
CD1231
CD1233
TA
CRISPR6
CD1234
CD1236
sigK 3'
sigK 5'
CD1233 cwp26
CD1230 CD1231
CD1234
CD0126 spoIIID

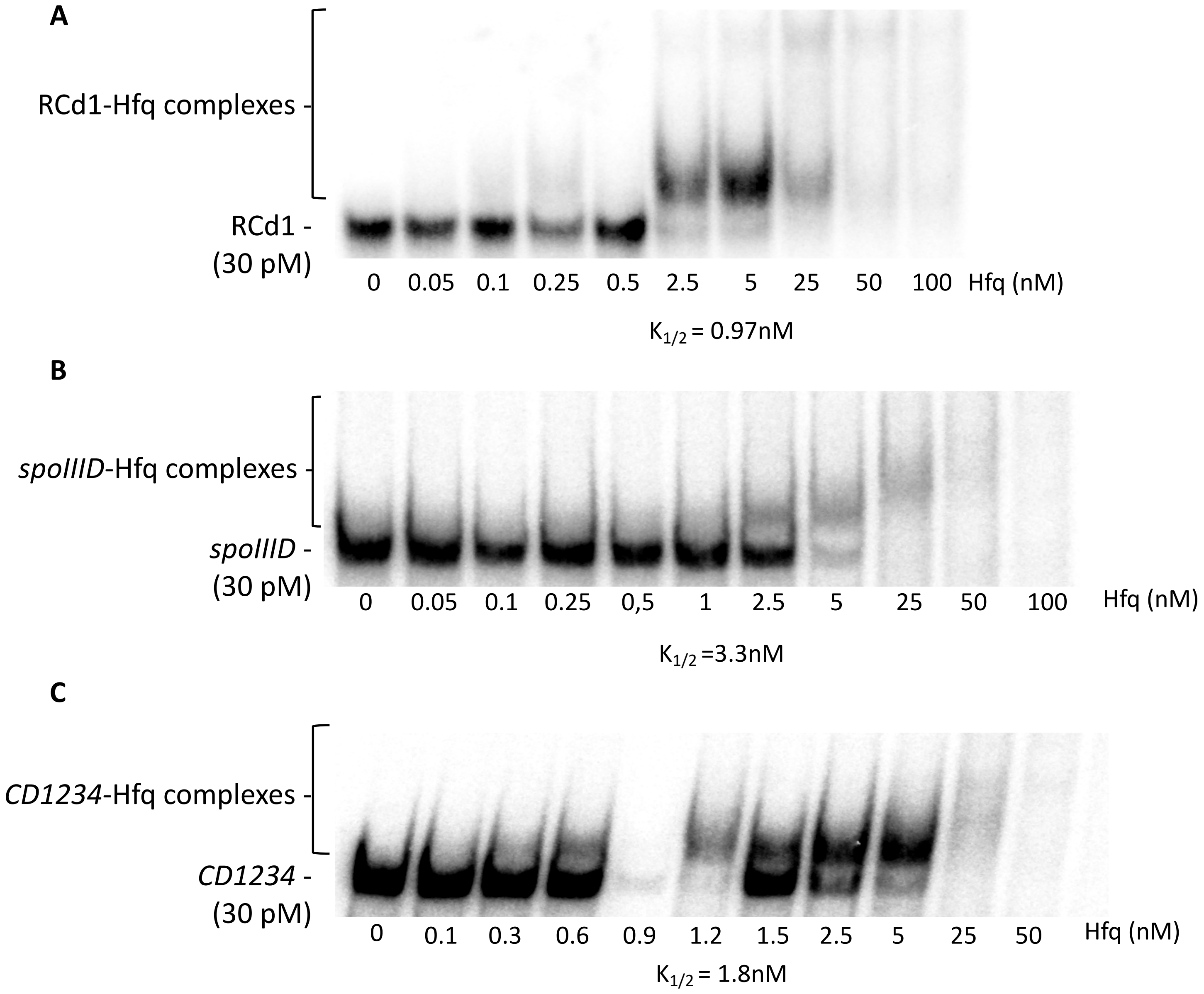
A
RCd1-Hfq complexes
RCd1
(30 pM)
0 0.05 0.1 0.25 0.5 2.5 5 25 50 100 Hfq (nM)
K1/2 = 0.97nM
B
spoIIID-Hfq complexes
spoIIID
(30 pM)
0 0.05 0.1 0.25 0,5 1 2.5 5 25 50 100 Hfq (nM)
K1/2 =3.3nM
C
CD1234-Hfq complexes
CD1234
(30 pM)
0 0.1 0.3 0.6 0.9 1.2 1.5 2.5 5 25 50 Hfq (nM)
K1/2 = 1.8nM

Fig. S9

### Fig. S10

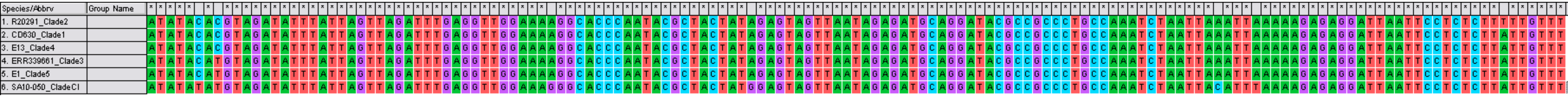

Species/Abbrv
Group Name
1. R20291_Clade2
2. CD630_Clade1
3. E13_Clade4
4. ERR339661_Clade3
5. E1_Clade5
6. SA10-050_CladeCI

Fig. S11

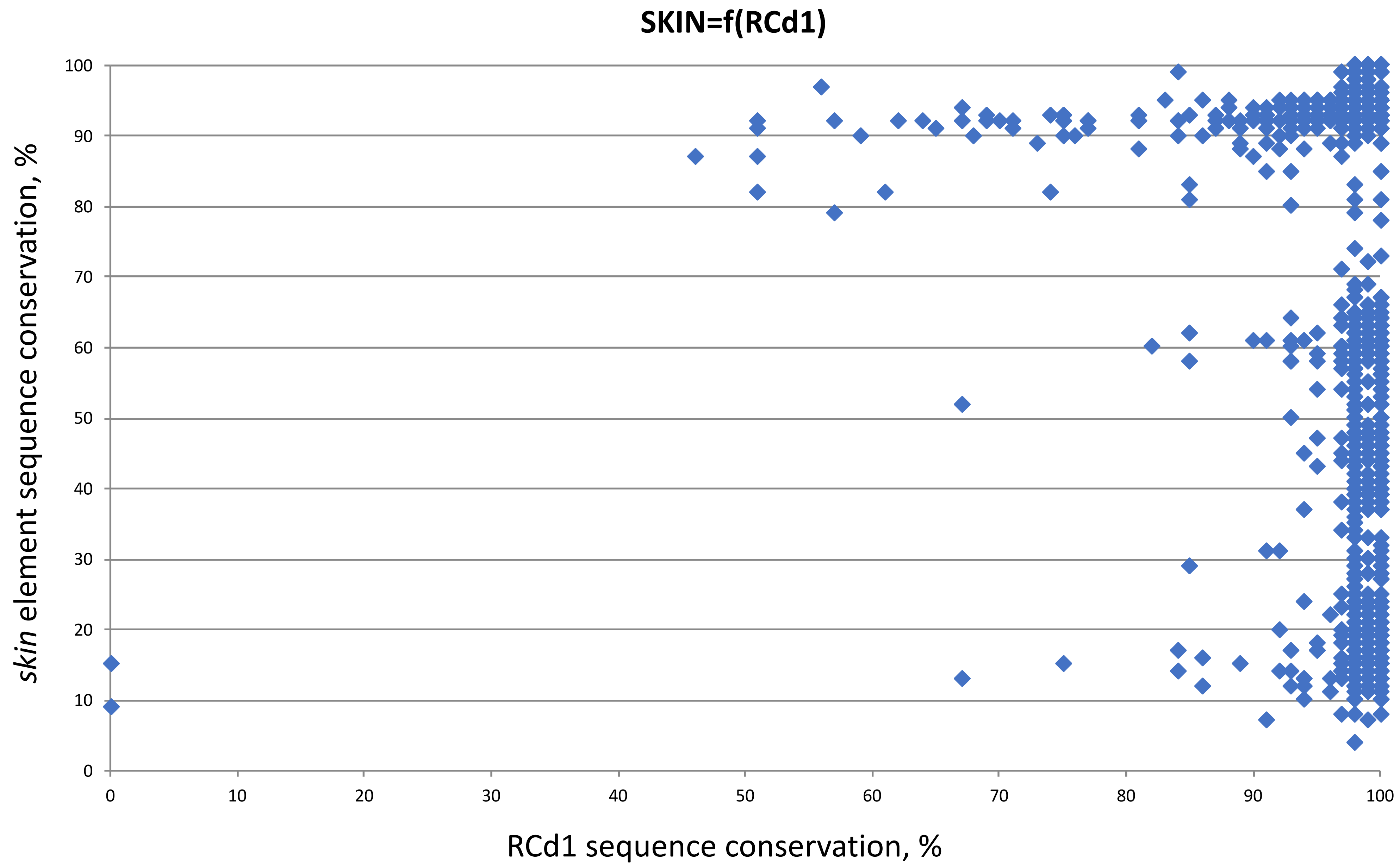

SKIN=f(RCd1)
skin element sequence conservation, %
100
90
80
70
60
50
40
30
20
10
0
0
10
20
30
40
50
60
70
80
90
100
RCd1 sequence conservation, %

### Fig. S12

***E*-value:** 2.0e-016 **Site Count:** 57 **Width:** 28

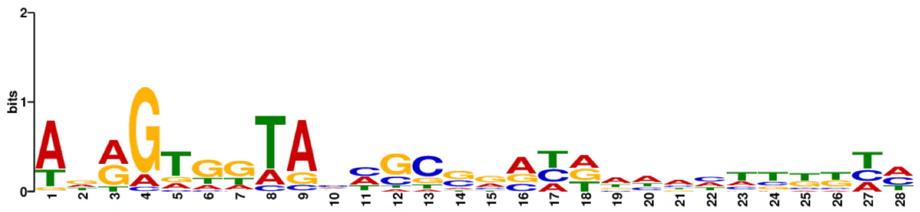

| Name | Strand | Start | *p*-value | Sites | | |
|---|---|---|---|---|---|---|
| 21. CD630_s0210 | + | 201 | 7.38e-13 | AAAGCTAAAT | AGAGTGGTACCGCGAGCAAAACCTCGTC | TCTAATTTTA |
| 33. CD630_s0450 | + | 169 | 1.06e-9 | CTTATCAATT | AGAGTGGTAACGCGGGTGTATATTCGTC | TCTTGGTCTT |
| 20. CD630_s0190 | + | 167 | 1.55e-9 | CTTATCAAAA | AGAGTGGTAACGCGGATATAATTCGTCT | CTTAGCTGTA |
| 19. CD630_s0160 | + | 194 | 4.67e-9 | TACGCCAACT | TGGGTGGAACCGCGAATTAATATCGTCC | CAAACATATT |
| 32. CD630_s0400 | + | 164 | 7.93e-9 | CTTATCAATT | AGAGTGGTAACGCGGATATATTTCGTCT | CTTGGTTTTT |
| 45. CD630_s0660 | + | 204 | 1.33e-8 | TAGATGAATT | AGGGTGGTACCGTGAATAATCTCGCCCC | TATGTTAACT |
| 16. CD630_s0010 | + | 157 | 1.33e-8 | ATAGTCAATT | AGGGTGGCAACGCGGATAAAAGATTTTC | GTCCCTTTTA |
| 34. CD630_s0480 | + | 182 | 2.58e-8 | AAAATCAATT | TGGGTGGAACCGCGGGAAACAGATTCTC | GTCCCTTTTT |
| 24. CD630_s0280 | + | 211 | 2.58e-8 | ATCTACAAAT | AAGGTGGTACCGCGGAATATAACTTTCG | TCCTTATAAG |
| 23. CD630_s0250 | + | 10 | 4.92e-8 | GTCTTATCA | AGAGTGGTGGAGGGACTGGCCCTTTGAA | ACCCGGCAAC |
| 26. CD630_s0300 | + | 192 | 1.24e-7 | AAAACTAAAT | TAGGTGGAACCGCGGGAATAATACTCTC | GTCCTTATGT |
| 22. CD630_s0220 | + | 10 | 1.24e-7 | ATCTTATCA | AGAGTGGCAGAGGGACTGGCCCTGTGAA | GCCCGGCAAC |
| 41. CD630_s0631 | + | 237 | 6.05e-7 | CTTTGGAATT | AAGGTGGTAACGCGAGCTTTTCGTCCTT | TTTAAAGAGG |
| 28. CD630_s0330 | + | 10 | 1.04e-6 | TTCTTATCC | AGAGTGGTGAAGGGACTGGCCCTATGAA | ACCCAGCAAC |
| 36. CD630_s0510 | + | 169 | 1.54e-6 | TTTACCAACT | AGAGTGGTAACACGGAATATTTCGTCTC | TTGGTCTTTT |
| 27. CD630_s0320 | + | 28 | 1.54e-6 | CCAAGAGCTG | AGAGGTATACCGACCCTTACACCTGATC | TGGATAATAC |
| 30. CD630_s0360 | + | 32 | 2.92e-6 | AAATTCCCAA | TCGGCGGTATAGCCCGCGAGCCAAGGTA | AAACTTGGTT |
| 48. CD630_SQ1005 | + | 179 | 3.31e-6 | TTGATGAACT | AAGGTGGTAACACGTAAGCAATGCTTTC | GTCCTTTTAA |
| 56. CD630_n00660 | + | 72 | 3.74e-6 | AGAGTAGTTA | ATAGAGATGCAGGATACGCCGCCCTGCC | AAATCTAATT |
| 39. CD630_s0600 | + | 140 | 3.74e-6 | CTCGTCCCTC | TTAGTCCTCCTGCCGACTAAGACTGGAC | GTCATTATGC |
| 2. CD630_n00290 | + | 1 | 4.23e-6 | | AAGGTTATCCCCCTGGTAACCCTAGTTA | GTTAATATAT |
| 40. CD630_s0610 | + | 68 | 5.39e-6 | TGATTGGAAA | AAGGATTTACAGCCGAAGAAAATATTTC | TTGCAATAAT |
| 8. CD630_n00620 | + | 210 | 1.53e-5 | ATGGGAAAGT | AAAGATGTACTCCCAATACATCTTTACT | TTAGCAGAAG |
| 17. CD630_s0050 | + | 18 | 1.71e-5 | CAAGAGAGGC | TGAGGGATAGGCCCTGTGAAGCCCAGCA | ACCACCCAAA |
| 44. CD630_s0642 | + | 209 | 2.64e-5 | ACTTTGTTAA | ATTGTGGTGTAGTGGGCTTTAATCTGCT | ACACTTTTAT |
| 37. CD630_s0590 | + | 69 | 2.64e-5 | GGCAATATAA | AAGGAGTAAGTGCCGAAGTGCTAGTGTC | CGAAGCTAGT |
| 29. CD630_s0340 | + | 2 | 4.47e-5 | T | AGAAGGGTAGAGGCGCAATAATCTATCA | GTACTAATAA |
| 18. CD630_s0080 | + | 55 | 5.49e-5 | AAAAGGTTAA | ATTCCTTTACAGCCCCCGCTACTGTGAT | GCAGACGAAA |
| 11. CD630_Cdi2_2 | + | 59 | 6.71e-5 | ATATATGGAG | TTAGTGGTGCAACCGGCTATGAATATAA | T |
| 1. CD630_Cdi2_3 | + | 186 | 7.41e-5 | ATATACTGAG | TCAGTGGTGCAACCGGCTATGAATATAA | ATTTATTTAT |
| 42. CD630_s0640 | + | 35 | 9.94e-5 | TTCCCTATCG | GCGGTAAAAGCCCGCGAGCCTTATGGCA | TAATTTGGTC |
| 25. CD630_s0281 | + | 224 | 1.20e-4 | TTATAGCCAT | ATAGGTATAAAGCCTATGTGGCTTTTTT | ATTGTTTTAA |
| 46. CD630_s0670 | + | 71 | 1.45e-4 | AACCTTTTAT | CAGGTGGTCCTGCTAACAACGATGAGAT | GACTTGAAAT |
| 52. CD630_SQ367 | + | 181 | 1.75e-4 | AGTCAAAGTA | AAAGTATTAAACCCAATTTATAATGTTA | ACATTTTTTA |
| 15. CD630_RNA_7 | + | 12 | 1.92e-4 | AACTACTCGA | AGGGGAGTAGCTTACACGAAAGTGGTAA | TTAATCGTCA |
| 14. CD630_RNA_17 | + | 37 | 2.10e-4 | TAGCACACCG | AAGGAGTAACACTCTCAGGTACTTTTAA | AAGTAGGACT |
| 31. CD630_s0370 | + | 3 | 3.00e-4 | AA | TTAGAGCTAGGGGCGCTAGAATTGGCTA | GCTGAGAGAT |
| 10. CD630_n00930 | + | 242 | 3.27e-4 | TATTATAAAT | AGGAGGGTGTCTCAAAATGAACTTTTTA | GTTTATGAGG |
| 5. CD630_n00440 | + | 47 | 3.27e-4 | CCCTAAATTT | ATAATGACAACTCTCCCTCACATCCTCA | AATTATAGTT |
| 57. CD630_n00030 | + | 104 | 3.56e-4 | TGTACCATTC | GAGGTTAAAGTGCGCATAAATATTGTTT | TATGTGTTTT |
| 53. CD630_SQ408 | + | 1 | 3.56e-4 | | TAGGTATTATAGCAGCAGGAATTGCTTT | AATTTTCTTT |
| 13. CD630_RNA_16 | + | 83 | 4.60e-4 | TAGATGAAAT | TCTCAGGTAAAGGTGACTGTACTTGGAC | G |
| 12. CD630_RNA_15 | + | 21 | 5.00e-4 | GAGACCCATT | ATGGGCGCCGAAGGGGCAAGGCTTTTAT | GCTCAATCTC |
| 9. CD630_n00910 | + | 48 | 5.43e-4 | TATAACGATA | ATTACTGAGACTCGGATGCCGACTGTCA | GTTAGGAATC |
| 6. CD630_n00470 | + | 23 | 5.89e-4 | GAAGAACAAT | ATGGTATTAATCTTGATGAAATACGTAA | GGAGCATATA |
| 43. CD630_s0641 | + | 348 | 6.39e-4 | TGTTTTTCCA | AAAGCAGAGGTCGGCATTTTACAATGCC | ATTCCTCTGC |
| 51. CD630_SQ327 | + | 141 | 6.93e-4 | ATTATTAATG | ATTGGTTTGGTCCGAGATAAAACCCACA | ACATAGTTTT |
| 3. CD630_n00330 | + | 29 | 7.50e-4 | TATATAATAA | GTACTCTTATCCACCCTAACCCTTTGAA | AATTGTGCTT |
| 54. CD630_SQ495 | + | 197 | 9.49e-4 | AATTATTTTT | AGTATAGTACCTACGCCAATAGGTGCTA | TTTCTTTTTC |
| 50. CD630_SQ1633 | + | 30 | 1.29e-3 | GCTTTATCAG | TAGGAGTACCTGCAGGTAAGACAGAATC | TCTAGTACGA |
| 7. CD630_n00590 | + | 44 | 1.49e-3 | AAATGGATAC | TGAATTTCATTGGGGATTAAAATGGGTT | GCAGGAAGAA |
| 49. CD630_SQ1339 | + | 138 | 1.85e-3 | TCTTGCAAAC | ATAGTATTTCTCTCATCAACCCTTTTCA | ATTCTATTCC |

Fig. S13

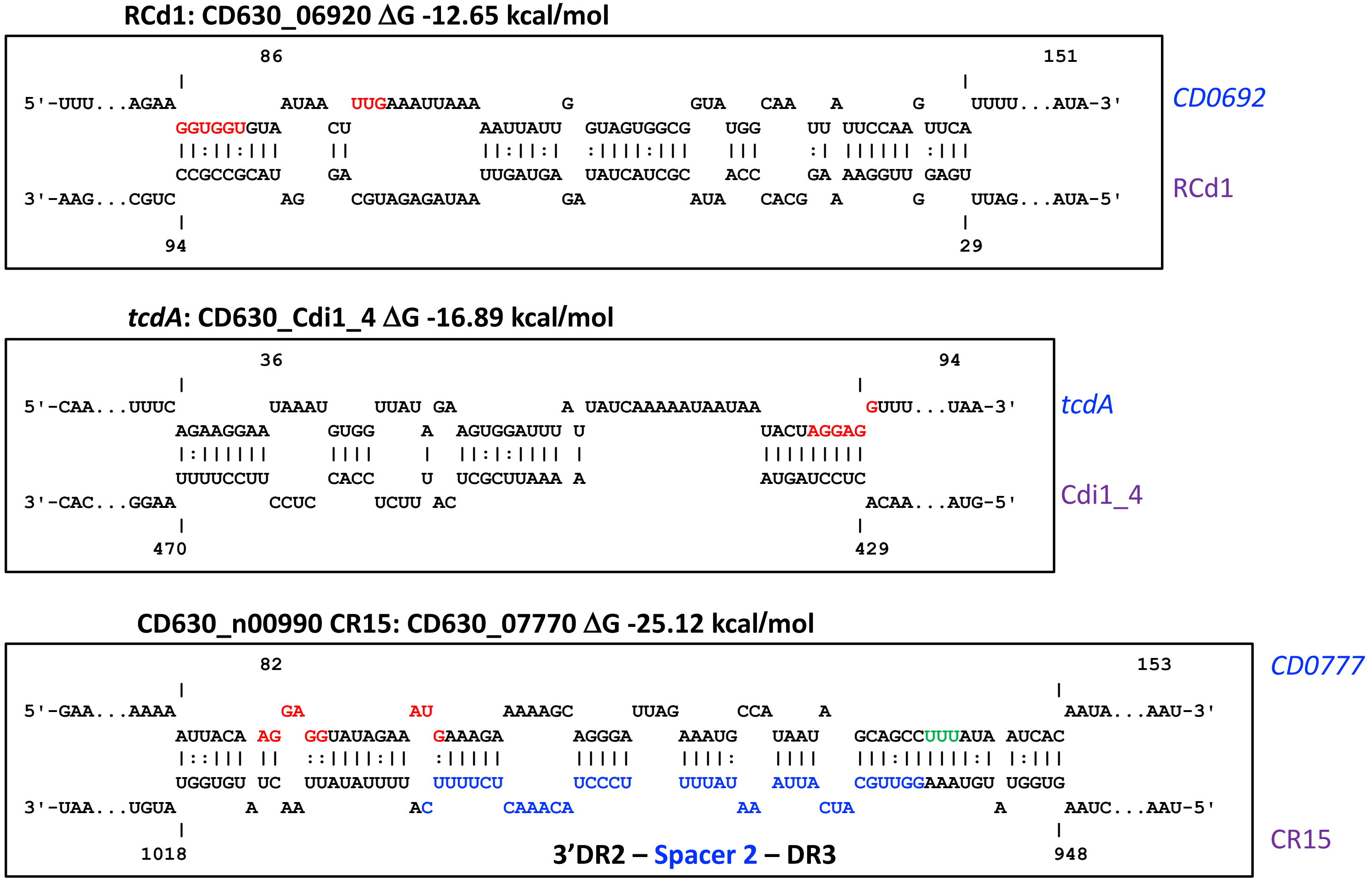
RCd1: CD630_06920 ΔG -12.65 kcal/mol
86
151
5'-UUU...AGAA AUAA UUGAAAUUAAA G GUA CAA A G UUUU...AUA-3'
GGUGGUGUA CU AAUUAUU GUAGUGGCG UGG UU UUCCAA UUCA
CCGCCGCAU GA UUGAUGA UAUCAUCGC ACC GA AAGGUU GAGU
3'-AAG...CGUC AG CGUAGAGAUAA GA AUA CACG A G UUAG...AUA-5'
94
29
CD0692
RCd1
tcdA: CD630_Cdi1_4 ΔG -16.89 kcal/mol
36
94
5'-CAA...UUUC UAAAU UUAU GA A UAUCAAAAAUAAUAA GUUU...UAA-3'
AGAAGGAA GUGG A AGUGGAUUU U UACUAGGAG
UUUUCCUU CACC U UCGCUUAAA A AUGAUCCUC
3'-CAC...GGAA CCUC UCUU AC ACAA...AUG-5'
470
429
tcdA
Cdi1_4
CD630_n00990 CR15: CD630_07770 ΔG -25.12 kcal/mol
82
153
5'-GAA...AAAA GA AU AAAAGC UUAG CCA A AAUA...AAU-3'
AUUACA AG GGUAUAGAA GAAAGA AGGGA AAAUG UAAU GCAGCCUUUAUA AUCAC
UGGUGU UC UUAUAUUUU UUUUCU UCCCU UUUAU AUUA CGUUGGAAAUGU UGGUG
3'-UAA...UGUA A AA AC CAAACA AA CUA A AAUC...AAU-5'
1018
948
3'DR2 – Spacer 2 – DR3
CD0777
CR15

**tcdA: CD630_SQ1656 ΔG -13.14 kcal/mol**

```
         58                                                                                 166
         |                                                                                   |
5'-CAA...UAUA  AG   A      AUCAAAAA                U    G  UUUA      AA              ACUCGC      A    CCAA          UAUA...UAA-3'   tcdA
             GA  UGG UUUAUU        UAAUAAUAC UAGGA GGUU UUAU UC  AUAUCUA  GAAGAGUUAAUAAA   AUAU AGC UUAGA    GAGAAAAUGAG
             ||  |:| ||||||        ||||||||| |||:| :||| |||| ||  |||||||  |||:|:||||||||   |||| |:| |||:|    ||:|||||:|:
             CU  AUC AAAUAA        AUUAUUAUG AUCUU UCAA AAUA AG  UAUAGAU  CUUUUUAAUUAUUU   UAUA UUG AAUUU    CUUUUUUAUUU
3'-UUG...ACGG  AA   A      GAUGUCUA         UU     AA   U      UUG      CGA             AAAUA    C   C    AAUAGA         UUCG...AGA-5'   SQ1656
         |                                                                                   |
        124                                                                                  11
```

**tcdA: CD630_SQ1642 ΔG -11.18 kcal/mol**

```
          9                                            60
          |                                            |
 5'-CAAAGUUU    CA      C  UU    UUUC        U      GUUAU   GUGG...UAA-3'   tcdA
            AAAAA  AGAAAAU AA  UAAA     AGA   AGGAA AAAUGUG     AGAA
            |||||  ||||||| ||  ||||     |||   ||||| ||||:||     ||||
            UUUUU  UCUUUUA UU  AUUU     UCU   UCCUU UUUAUAC     UCUU
3'-UUU...UAAC    UA          UU    UUCAU   AUA                AU      AAUA...UUG-5'   SQ1642
          |                                            |
         197                                          147
```

**tcdA: CD630_s0300 ΔG -10.24 kcal/mol**

```
         88                       111
         |                         |
5'-CAA...AAUA     A          G     AUCU...UAA-3'   tcdA
             CUAGG GGUUUUU AU UCUUUAAU
             ||||| :|||||: || :||:||||
             GAUCC UCAAAAG UA GGAGAUUA
3'-AUU...CGAG     A        U  G    AAUU...AUA-5'   s0300
         |                         |
        89                        65
```

**tcdA: CD630_n00620 ΔG -10.14 kcal/mol**

```
          137                                                          197
           |                                                            |
5'-CAA...ACUC     G   U          GAGA   GA    AAA    A          G C    AUAA-3'   tcdA
             GCAUAUA CAU AGA   CCAA  AAAU  GUAUA  CUAU  CUAAC UAAUUUA A GAAU
             |||||:| ||| |:|   ||||  ||||  :||||  ||||  ||||| ||||||| | ||||
             CGUAUGU GUA UUU   GGUU  UUUA  UAUAU  GAUA  GAUUG AUUAAAU U CUUA
3'-AAC...UUUU     G   U  AAUUAGUA  AUG   A    AAAG  CA    A         G U    GUAA...UGG-5'   n00620
           |                                                            |
          185                                                          116
```

##### *tcdC*: CD630_n00440 ΔG -15.04 kcal/mol

```
            87                   107
             |                   |
5'-AUA...UAAA    A                CUAA...UCU-3'
             AGGG GAUUGUAUUAUGUUUU
             |||| :|||:|||||||||||
             UCCC UUAAUAUAAUACAAAA
3'-UUU...UAAA    A                UUCG...AUA-5'
             |                   |
            40                   20
```

*tcdC*

n00440

##### *tcdC*: CD630_s0480 ΔG -11.00 kcal/mol

```
            51                                                                                          135
             |                                                                                          |
5'-AUA...UUUA             UUAAAAUAAAAUAUA      AAA        UG        U         AAAAAAAAUGA         A      UAAU...UCU-3'
             AGUUUUUAA CUU               UCUAAU   AGGGAGAU    UA UUA   GUUUUCU           UGGUAA CG AUUUAG
             ||||||:|| |:|               ||||||   |||||:|:    :| :||   ||||:||           |||||| || |||:||
             UCAAAAGUU GGA               AGAUUA   UCCCUUUG    GU GAU   CAAAGGA           ACCAUU GC UAAGUC
3'-CAU...UCCA         A                        A          UAAA  C   CAU       AGAG             A         UAAA...GAU-5'
             |                                                                                          |
            81                                                                                          15
```

*tcdC*

s0480

##### *agrB*: CD630_Cdi1_8 ΔG -11.63 kcal/mol

```
            41                                                                       113
             |                                                                       |
5'-UGA...AGUA     A        AAAAAAGAA         UAU          UU     AG  AAAAA            AUGC...AUU-3'
             UAUUG AUUUUCUC         UGUACUCUA   AUGUAAAAUU  UUUGG  GU     UGUUUAAAAGAU
             ||:|: ||||||||         :||||||||   ||:||||||:  |:|::  ||     |:|:|||||:||
             AUGAU UAAAAGAG         GCAUGAGAU   UAUAUUUUAG  AGAUU  CA     AUAGAUUUUUUA
3'-UAA...UGAG     G                                       U      GA  C                AUGU...AUG-5'
             |                                                                       |
           289                                                                       234
```

*agrB*

Cdi1_8

***cdtR*: CD630_n00290 ΔG -13.65 kcal/mol**

```
            81                                  114
             |                                   |
5'-AAC...GAAU       A          U                 CUUU...UGU-3'
             UUUUUAU GGAGGGUAUU   AGUGGAUAUAUUAAU
             |::|||| ::||||||||   |:|:||||||||||:
             AGGAAUA UUUCCCAUAA   UUAUCUAUAUAAUUG
3'-GAG...AUAU       C          CCC               AUUG...GAA-5'
             |                                   |
            64                                  29
```

*tcdR*

n00290

***cdtR*: CD630_s0641 ΔG -12.78 kcal/mol**

```
            3                                                                                                            106
             |                                                                                                            |
        5'-AA    CCUU                C       G   UUUUAU            UAAAAUACUUUU    U    CU        UUU   AGGAGG      A      AUAU...UGU-3'
             CUAU    AAUC    UUAUUUAU UCCUAGU UUA      GUAUU  UUCUA            GUUU UUUA  UUUGAAUU   UAU      GUAUUU GUGGAU
             ||||    ||||    |||||:|: |||||:| |||      |||||  |||||            |||| |:||  |||:||||   ||:      |||:|| ||||||
             GAUA    UUAG    AAUAAGUG AGGAUUA AAU      CAUAA  AAGAU            CAAA AGAU  AAAUUUAA   AUG      CAUGAA CACCUA
3'-UGA...UUAA    UAU     UAUU        U       G              UU                     U    AU        C     AAAA        A      AUAG...AUC-5'
             |                                                                                                            |
           124                                                                                                            38
```

*cdtR*

s0641

***cdtR*: CD630_s0190 ΔG -11.00 kcal/mol**

```
            72                                                      124
             |                                                       |
5'-AAC...UUUA     G         U   A     U      GGA        UC           GAUG...UGU-3'
             CUUUU AAUUUUUUA AGG GGGUA  UUAGU   UAUAUUAA    UUUGAUAAU
             |:|:| |||:::||| ||| |||||  ::|:|   :|||:|||    |:|||||||
             GGAGA UUAGGGAAU UCC CCCAU  GGUUA   GUAUGAUU    AGACUAUUA
3'-AUU...AAUA     G             G     UU     AUA        UCAA         GAGG...AUA-5'
             |                                                       |
           145                                                      91
```

*cdtR*

s0190

**CD630_n00600 CR9: CD630_08100 ΔG -30.61 kcal/mol**

```
            152                                            196
             |                                              |
5'-UAA...GGUC       U    GA         UAAA         A          GUAAA-3'
             GCUAAGC GAUA  AGAGGGCGC    AAGGU  GG UGGAGUAGAU
             :|||||| ||||  ||||:||||    ||||:  :| :||||:|:||
             UGAUUCG CUAU  UCUCUCGCG    UUCCG  UC GCCUCGUUUA
3'-UAA...AUUU            AA                  UA             AAUG...AUA-5'
             |                                              |
           831                                             791
```

Spacer 10

*CD0810*

n00600 CR9

**CD630_n00990 CR15: CD630_23450 ΔG -21.24 kcal/mol**

```
             56                                                                                                                                                                 201
              |                                                                                                                                                                  |
5'-GUU...UUAA         UGUCAC        U           ACU             CCA     AUAUA    AUU           UCAACUGUCU   A     A      U     AA        AAAG   U  UAU          AUC     A    -3'
             UUAUAUACU       UAAUAUG AAAUGAAAGGA            AAUA   AAUGA     AACU    UAUACUAUUU           GU AAUAU AAAACA  GACU  GGCUGCGG    UAU CA   UUCUCAA CC    CAUA CAUU
             |||:||||:       ||||||: ||||||||:||            ||||   |||||     |||:    :||||:|:||           || ||||| ||||||  :|||  :|||:|::    ||| ||   ||||||| ||    |||| ||:|
             AAUGUAUGG       AUUAUAU UUUACUUUUCU            UUAU   UUACU     UUGG    GUAUGGUGAA           CA UUAUA UUUUGU  UUGA  UCGAUGUU    AUA GU   AAGAGUU GG    GUAU GUGA
3'-UAA...GAUA         UGUAUCA                   CAAACAUCCCUU    AAA     ACG      AAAU          U            A           UC              CGAA   U  CGC       U  AAAU    G    AUCA...AAU-5'
              |                                                                                                                                                                  |
          1024                                                                                                                                                                  881
```

3'DR1 - Spacer 1 - DR2 - Spacer 2 - DR3

*CD2345*

n00990 CR15

**CD630_n00990 CR15: CD630_04341 ΔG -20.96 kcal/mol**

```
            157                                           194
             |                                              |
5'-ACC...CUAU     A      A      AG       A                  CAUUAAA-3'
             AAGAG    UGG GGAAGA     AAUG AUGCAACU   AUAUACC
             |||||    |:| ||||:|     |||| |||||||:   |:|||||
             UUCUC    AUC CCUUUU     UUAC UACGUUGG   UGUAUGG
3'-UAA...ACUU     AAAC          AUAAA            AAA        UGAA...AAU-5'
             |                                              |
           994                                            950
```

3'DR2 – Spacer 2

*CD04341*

n00990 CR15

Fig. S14

**CD630_n00600 CR9: CD630_s0450 ΔG -19.77 kcal/mol**

Spacer 10 - 5'DR11

```
           797                                     836
             |                                      |
5'-AUA...UUGC        G   UGC        AUA   C          AUUA...AAU-3'
             UCCGCUAU CCU   GCUCUCUA   UCG UUAGUUUUAU
             |||:|:|| ||:   ||||||||   ||: :|||:|||||
             AGGUGGUA GGG   CGAGAGAU   AGU GAUCGAAAUA
3'-AUU...CCGA        A   U                U          CGAU...AAA-5'
             |                                      |
            49                                      15
```

CR9

s0450

**CD630_n00290: RCd5 ΔG -16.82 kcal/mol**

```
            23                                                                                     104
             |                                                                                     |
5'-AAG...ACCC        UAA             UC  AUACCCU        G       U       A        U  GAU    U        UUAA...GAG-3'
             UAGUUAGU       UAUAUCUAU CCA       UUCAUAAG AUAUAUU UAUAUUU  UUAAAAA CG    UAG AACGGAAA
             :|||||||       ||:|||||| |||       |:|||||: ||||||: |||||::  :|||||| |:    ||: |||::|||
             GUCAAUCA       AUGUAGAUA GGU       AGGUAUUU UAUAUAG AUAUAGG  GAUUUUU GU    AUU UUGUUUUU
3'-AGU...AUAC        UAUACAC             AAAAU          G       U       GG       U  AAGU   U        CUUU...UGA-5'
             |                                                                                     |
           194                                                                                     111
```

n00290

RCd5

**RCd1: s0642 ΔG -11.49 kcal/mol**

```
            27                    48
             |                     |
5'-AUA...UAGA     G     AA          AAUA...GAA-3'
             UUUGA GUUGG  AAGGCAC CC
             |:||| :|::|  |||:||| ||
             AGACU UAGUC  UUCUGUG GG
3'-AAA...AUUG     G     CG       A  AAGU...UAA-5'
             |                     |
            39                     17
```

RCd1

s0642

**CD630_n01010 CR17: CD630_s0470 ΔG -19.84 kcal/mol**

```
            1275                                                                                          1353
             |                                                                                             |
5'-AAC...UAUC                 A    AAUA    C     CA   A                              AA                        UAAA...AAU-3'
             AACUAAGUGGUAG AUA GUAG    UUAG AAUAU  ACG GUAUU    UAGAAACUUAU  UUUAUAUU  UAAA            GCAGUAUA
             ||||||:|::||| ||| :|||    |:|| |||||  ||| |||||    |||||||||||  ::::||||  :|||            :|||||:|
             UUGAUUUAUUAUC UAU UAUC    AGUC UUAUA  UGC CAUAA    AUCUUUGAAUA  GGGUAUAA  GUUU            UGUCAUGU
          3'-             A   G                  AA          GGCC           CU        CA    UAAUAUACAAUA        UUUGUAC-5'
             |                                                                                             |
            99                                                                                             8
```

n01010 CR17

s0470

**CD630_s0400: CD630_n00080 ΔG -19.48 kcal/mol**

```
            21                                                         78
             |                                                          |
5'-AGA...CUAU    UU       A    UG   A   U  AA          UA      U      AA    ACUU...AUU-3'
             GUGU  UAUAGAG   GC  AGG UGG GG  GCUUGGCAAA    AGUA  UAGUG  AAGA
             ||||  ||||:||   :|  ||| ||: ||  |||:::||||    ||||  |||||  ||||
             CACA  AUAUUUC   UG  UCC ACU CC  CGAGUUGUUU    UCAU  AUCAC  UUCU
3'-AGA...CAAC             CAA  UG   A      GA          UUUA    UU     A     U-5'
             |                                                          |
            60                                                          2
```

s0400

n00080

**CD630_n00440: Cdi1_3 ΔG -19.35 kcal/mol**

```
            56                                  87
             |                                   |
5'-AUA...GACA    U    CACA        U    G      AUUA...UUU-3'
             ACUC CCCU    UCCUCAAA UAUA  UUGUC
             |||| |||:    |||||||| ||||  ||:||
             UGAG GGGG    AGGAGUUU AUAU  AAUAG
         3'-G    U                U    AA     AUAA...AAA-5'
             |                                   |
           496                                  468
```

n00440

Cdi1_3

**RCd1: CD630_n00790 CR11 ΔG -10.03 kcal/mol**

```
            42                                              83
             |                                              |
5'-AUA...AAAA    C         G           G    G  AA     A     GAUA...GAA-3'
             GGCA   CCAAUAC CUAC   UAUA AGUA UU  UAGAG UGCAG
             :|||   ||||||| ||||   |||| |:|| |:  ||:|: |||||
             UCGU   GGUUAUG GAUG   AUAU UUAU AG  AUUUU ACGUC
3'-UAA...UGAA    AAA            UAU    G    G  A      A     ACUC...AUA-5'
             |                                              |
           157                                             113
```

RCd1

n00790 CR11

CR11 leader

#### Table S1. Strains, plasmids and oligonucleotide primers used in this study

| **Strain** | **Genotype** | **Origin** |
|---|---|---|
| *E. coli* | | |
| DH5α | F- Φ80*lac*ZΔM15 Δ(*lacZYA-argF*) U169 *recA1 endA1 hsdR17* (rK-, mK+) *phoA supE*44 λ- *thi*-1 *gyrA*96 *relA*1 | Invitrogen |
| NEB-10 beta | Δ*(ara-leu) 7697 araD139 fhuA* Δ*lacX74 galK16 galE15 e14-* ϕ*80*d*lacZ*Δ*M15 recA1 relA1 endA1 nupG rpsL* ($Str^R$) *rph spoT1* Δ*(mrr-hsdRMS-mcrBC)* | New England Biolabs |
| *C. difficile* | | |
| 630 | Sequenced reference strain | Laboratory stock |
| 630Δ*erm* | 630Δ*erm* | Laboratory stock (1) |
| CDIP219 | 630Δ*erm* strain carrying pDIA6103 | (2) |
| CDIP258 | 630Δ*erm* strain carrying pDIA6142 | (2) |
| CDIP303 | 630Δ*erm* strain carrying pDIA6151 | This work |
| CDIP583 | 630Δ*erm* Δ$skin^{CD}$ strain | (3) |
| CDIP590 | 630Δ*erm* Δ$skin^{CD}$ strain carrying pDIA6103 | This work |
| CDIP592 | 630Δ*erm* Δ$skin^{CD}$ strain carrying pDIA6142 | This work |
| CDIP607 | 630Δ*erm* strain carrying pDIA6397 with *CD1234-3xFLAG* | This work |
| CDIP611 | 630Δ*erm* Δ*RCd1* | (4) |
| CDIP619 | 630Δ*erm* Δ*RCd1* carrying pDIA5973 plasmid | This work |
| CDIP696 | 630Δ*erm CD1234-3xFLAG*-tag | This work |
| CDIP704 | CDIP696 strain carrying pDIA6103 vector | This work |
| CDIP708 | CDIP696 strain carrying pDIA6142 plasmid | This work |
| **Plasmids** | | **Origin** |
| pRPF185 | $P_{tet}$-*gusA* $Tm^R$ | (5) |
| pMTLSC7315 | plasmid vector used for *codA* allele exchange in *C. difficile* 630 | (6) |
| p025/pMSR14 | pMTLSC7315 derivative with $P_{tet}$ *CD2517.1* toxin gene instead of *codA* for allelic exchange in *C. difficile* 630 | (7) |
| pDIA5973 | pRFF185 derivative carrying $P_{tet}$-AS-*CD1974* (*hfq*) $Tm^R$ | (2) |
| pDIA6103 | pRPF185 Δ*gus* vector derivative $Tm^R$ | (8) |
| pDIA6142 | pRFF185 derivative carrying $P_{tet}$-RCd1 $Tm^R$ | (2) |
| pDIA6151 | pRFF185 derivative carrying $P_{tet}$-ϕ*(CD1974(hfq)-3xFLAG)* $Tm^R$ | (2) |
| pDIA6397 | $P_{tet}$- *RCd1* - *CD1234-3xFLAG* $Tm^R$ | This work |
| pDIA6434 | pMTLSC7315 derivative for *CD1234-3xFLAG* gene editing | This work |

**Oligonucleotide primers**

***In vitro* interactions**

| | |
|---|---|
| T7RNA RCd1 | *TAATACGACTCACTATAGG*GATATACACGTAGATATTTATTAGTT |
| TermRCd1b | AAACAATAAGAGAGGAATTAATCCTCTC |
| T7CD1234 | *TAATACGACTCACTATAGGG*GTGATGTTATGAAAATACTACAAC |

| | | |
|---|---|---|
| TermCD1234 | TATGAGAAAGTGCTAAGTATTAGCAAC | |
| T7spoIIID | *TAATACGACTCACTATAGGG*AAAGCTATTATGTGAGGGGG | |
| TermspoIIID | TAAAAAAAGTAGGTTTTTAACCTACATTTTTTTATG | |
| **Quantitative RT-PCR/qPCR/Northern blot** | | |
| IMV824 | CCATGATTCAGATTCCCTTG | qPCR *skin* circularized |
| IMV825 | AAAAGTGTTTTGAATGGGGATT | |
| DNA polIII-*dnaF 5'* | TCCATCTATTGCAGGGTGGT | qRT-PCR or qPCR |
| DNA polIII-*dnaF 3'* | CCCAACTCTTCGCTAAGCAC | qRT-PCR or qPCR |
| LS102 | GGGCCATAGTGGTAGCAAAA | qRT-PCR 5' *CD0126-spoIIID* |
| LS103 | TGGCAAGGGATGGATTTATT | qRT-PCR 3' *CD0126-spoIIID* |
| LS155 | GAAAAACCCTTAACCCCTGA | qRT-PCR 5' *CD1230-sigK* |
| LS156 | TCATCCTGATCTTCCGTTGA | qRT-PCR 3' *CD1230-sigK* |
| LS352 | AGGCAAAAGGAAAGCATGAA | qRT-PCR 5' *CD1231* |
| LS353 | TGCAAATCACTATTCCTGCAA | qRT-PCR 3'*CD1231* |
| LS354 | AACAAGGGGTGATGTTATGAAAA | qRT-PCR 5' *CD1234* |
| LS355 | TTTTCTTCATCAGATATACGTGGA | qRT-PCR 3'*CD1234* |
| LS263 | TGCGATAAATTTACACAAGCTG | qRT-PCR 5' *CD0597* |
| LS264 | GGCCAAGGTTCAGATATCCA | qRT-PCR 3' *CD0597* |
| LS145 | GCTGCATCTACTCCATTAGCAA | qRT-PCR 5' *CD1613* |
| LS146 | GCAATCATCACAATCGCAGT | qRT-PCR 3' *CD1613* |
| PB124 | TGCTGCTAATGCAGTGAAAAA | qRT-PCR 5' *CD1233* |
| PB125 | CCCTCTGCACCCTGTTTAAT | qRT-PCR 3' *CD1233* |
| QRTBD325 | CGGAACAGATAAAGAAGGTAATGAA | *sigK* 5' *skin* |
| QRTBD326 | TCATCAAGAACATAGTTAGCCTCTG | *sigK* 3' *skin* |
| OS379 | GTTAGATTTGAGGTTGGAAAAGG | RCd1 5' |
| OS380 | GGCGTATCCTGCATCTCTATT | RCd1 3' |
| OS757 | GAATGAGGTTCTCCCTTGGA | 5' IGR *CD0655* |
| OS758 | AAAAGCTGGTGGCATCTATGA | 3' IGR *CD0655* |
| OS761 | AAAGAAATGAAAAAGCACTCTCCA | 5' AS *CD2890* |

| | | |
|---|---|---|
| OS762 | ACCCCAACGCATATTTACTGAG | 3' AS *CD2890* |
| OS664 | ATCCCCCTGGTAACCCTAGT | 5’ CD630_n00290 |
| OS665 | CCACCGAATTAATTTCCGTTAC | 3’ CD630_n00290 |
| OS668 | AATTACCCTAATTGTAATGGCAACTC | 5’ SQ1828 |
| OS669 | GGGAAAAGTTTTGTAATGACAACTATAA | 3’ SQ1828 |
| OS446 | TTTGTACCATTCGAGGTTAAAGTG | 5’ RCd2 |
| OS447 | AAAAAATATACGCCCTATAAAAGCG | 3’ RCd2 |
| AM289 | GAGAGAATTGTATAGATGTAAGTGTTG | 5’ n00460 CR6 |
| AM290 | GTGATGAATGTTCAGAAGAGGA | 3’ n00460 CR6 |
| AM175 | TGCAAATTTAAGAGAGTTGTATACG | 5’ n00560 CR8 |
| AM176 | TATCTTGAGCTGTCAATGTGAAC | 3’ n00560 CR8 |
| OS565 | AACTAAATCGGCAAAACTAGAGAAA | 5’ Cdi1_8 |
| OS566 | CTCTTTGGCAACTGGCTGAC | 3’ Cdi1_8 |
| OS234 | AGCTTTCGCTTTAGGCAGTG | 5’ *tcdA* |
| OS235 | ATGGCTGGGTTAAGGTGTTG | 3’ *tcdA* |
| OS252 | TGAAGACCATGAGGAGGTCA | 5’ *tcdC* |
| OS253 | CGTCGTCTTTCATTTTGAACC | 3’ *tcdC* |
| OS580 | CAGTAGTGGCAGTTCCAGCTT | 5’ *pilA CD3513* |
| OS581 | CCAGTTTGACCATCTGGTGT | 3’ *pilA CD3513* |
| **Cloning/strain construction** | | |
| PB9 | GAAGGCCTTTTTAGAAGTTCAATTTAATC | 5' CD1974 StuI pDIA6103 cloning |
| PB10 | GGGGATCC**TCACTA**CTTGTCATCGTCATCCTTGTAGTC GATGTCATGATCTTTATAATCACCGT CATGGTCTTTGTAGTCTCTGTTGTTATTATTATTGTTG | 3' CD1974 FLAG-stop-BamHI pDIA6103 cloning |
| OS691 | GGCTGGGACCTGAATATGTTACTTTTGTAAAGG | 5'Pro-*CD1234*-DraII cloning |
| OS692 | CTGCAGAACCAGTGTGCTGGTCACTACTTGTCATCGTC ATCCTTGTAGTCGATGTCATGATCTTTATAATCACCGT CATGGTCTTTGTAGTCACTTTCTACTGTCTCAACTTTC | 3'*CD1234*-FLAGtag-BstXI cloning |
| PB191 | gtttttgttaccctaagtttCTTACAAAGTGTTGCAAAAGAG | *CD1234*-Cter-Flag 5'homology arm |

| | | |
|---|---|---|
| PB192 | tttgtagtcACTTTCTACTGTCTCAACTTTC | |
| PB193 | gtagaaagtGACTACAAAGACCATGACGGTGATTATAAAG<br>ATCATGACATCGACTACAAGGATGACGATGACAAGTA<br>AGAGAATAGAGAAGTTGCTAATAC | *CD1234*-Cter-Flag 3'homology arm |
| PB194 | gattatcaaaaaggagtttGGAATGGAAATGAGGAAAC | |
| OS705 | CTCGAAAAGTTTCTGCTTTGGA | 5'*CD1234* outside homology arm CterFLAG strain verification |
| OS706 | GCAATAAAAATACAAAGACCAGGTC | 3'*CD1234* outside homology arm |


1. Hussain, H.A., Roberts, A.P. and Mullany, P. (2005) Generation of an erythromycin-sensitive derivative of *Clostridium difficile* strain 630 (630Deltaerm) and demonstration that the conjugative transposon Tn916DeltaE enters the genome of this strain at multiple sites. *J Med Microbiol*, **54**, 137-141.
2. Boudry, P., Gracia, C., Monot, M., Caillet, J., Saujet, L., Hajnsdorf, E., Dupuy, B., Martin-Verstraete, I. and Soutourina, O. (2014) Pleiotropic role of the RNA chaperone protein Hfq in the human pathogen *Clostridium difficile*. *Journal of bacteriology*, **196**, 3234-3248.
3. Serrano, M., Kint, N., Pereira, F.C., Saujet, L., Boudry, P., Dupuy, B., Henriques, A.O. and Martin-Verstraete, I. (2016) A Recombination Directionality Factor Controls the Cell Type-Specific Activation of sigmaK and the Fidelity of Spore Development in *Clostridium difficile*. *PLoS genetics*, **12**, e1006312.
4. Huang, L., Wang, J., Boudry, P., Wilson, T.J., Soutourina, O. and Lilley, D.M.J. (2020). Personal communication
5. Fagan, R.P. and Fairweather, N.F. (2011) *Clostridium difficile* has two parallel and essential Sec secretion systems. *The Journal of biological chemistry*, **286**, 27483-27493.
6. Cartman, S.T., Kelly, M.L., Heeg, D., Heap, J.T. and Minton, N.P. (2012) Precise manipulation of the *Clostridium difficile* chromosome reveals a lack of association between the *tcdC* genotype and toxin production. *Applied and environmental microbiology*, **78**, 4683-4690.
7. Peltier, J., Hamiot, A., Garneau, J., Boudry, P., Maikova, A., Fortier, L.C., Dupuy, B. and Soutourina, O. Type I toxin-antitoxin systems of *Clostridioides difficile* confer maintenance of mobile genetic elements. *bioRxiv 2020.03.04.976019*.
8. Soutourina, O.A., Monot, M., Boudry, P., Saujet, L., Pichon, C., Sismeiro, O., Semenova, E., Severinov, K., Le Bouguenec, C., Coppee, J.Y. *et al.* (2013) Genome-Wide Identification of Regulatory RNAs in the Human Pathogen *Clostridium difficile*. *PLoS Genet*, **9**, e1003493.

**Table S2. Number of mapped reads per RIP-seq sample**

| Strain name | Replicate number | Condition | Index | Reads number | Length, base | %Map |
|---|---|---|---|---|---|---|
| CDIP303_3xFlag | 1 | 3xFlag | 1 | 42M | 51 | 97.7% |
| CDIP303_3xFlag | 2 | 3xFlag | 3 | 28M | 51 | 98.6% |
| CDIP303_3xFlag | 3 | 3xFlag | 10 | 38M | 51 | 98.7% |
| CDIP303_3xFlag | 4 | 3xFlag | 20 | 48M | 51 | 98.8% |
| CDIP219_Control | 1 | Control | 2 | 44M | 51 | 98.1% |
| CDIP219_Control | 2 | Control | 4 | 48M | 51 | 99.1% |
| CDIP219_Control | 3 | Control | 7 | 72M | 51 | 98.5% |
| CDIP219_Control | 4 | Control | 12 | 56M | 51 | 98.8% |
| Hfq630E | 1 | Hfq | 25 | 43M | 51 | 98.4% |
| PI630E | 1 | Control | 11 | 27M | 51 | 98.8% |

**Table S3. Complete list of MACS peaks with MAGe annotation**

| source | tool | feature | start | stop | size | strand | name | -log10(pval) | overlap1 | stream (ov1) | name(ov1) | product(ov1) | overlap2 | stream (ov2) | name (ov2) | product(ov2) | overlap3 | stream (ov3) | name (ov3) | product(ov3) |
|---|---|---|---|---|---|---|---|---|---|---|---|---|---|---|---|---|---|---|---|---|
| CD630 | MACS2 | peak | 17667 | 17807 | 140 | + | macsForward_peak_1 | 4.92433 | CD630_s0010 | sense | - | T-box (Ser) | | | | | | | | |
| CD630 | MACS2 | peak | 63203 | 63280 | 77 | + | macsForward_peak_2 | 3.32169 | CD630_00390 | sense | acoL | Acetoin dehydrogenase E3 component (dihydrolipoamide dehydrogenase) | | | | | | | | |
| CD630 | MACS2 | peak | 137966 | 138495 | 529 | + | macsForward_peak_3 | 3.28290 | CD630_01080 | sense | nrdD | Anaerobic ribonucleoside triphosphate reductase | | | | | | | | |
| CD630 | MACS2 | peak | 174158 | 174267 | 109 | + | macsForward_peak_4 | 5.03432 | CD630_s0050 | sense | - | SAM | | | | | | | | |
| CD630 | MACS2 | peak | 174162 | 174227 | 65 | - | macsReverse_peak_1 | 6.79071 | CD630_s0050 | antisense | - | SAM | | | | | | | | |
| CD630 | MACS2 | peak | 206301 | 206791 | 490 | + | macsForward_peak_5 | 4.06352 | CD630_01560 | sense | - | Putative membrane protein | | | | | | | | |
| CD630 | MACS2 | peak | 207370 | 207832 | 462 | - | macsReverse_peak_2 | 6.08699 | CD630_01570_p2 | sense | - | Fragment of putative membrane protein (Part 2) | CD630_01570_p1 | sense | - | Fragment of putative membrane protein (Part 1) | | | | |
| CD630 | MACS2 | peak | 209892 | 209943 | 51 | + | macsForward_peak_6 | 3.45609 | CD630_01590 | sense | - | Putative enzyme, glyoxalase family | | | | | | | | |
| CD630 | MACS2 | peak | 217445 | 217613 | 168 | + | macsForward_peak_7 | 4.55381 | CD630_01640 | sense | - | Putative membrane protein | | | | | | | | |
| CD630 | MACS2 | peak | 224045 | 224516 | 471 | - | macsReverse_peak_3 | 4.86304 | CD630_01690 | sense | - | Putative drug/sodium antiporter, MATE family | | | | | | | | |
| CD630 | MACS2 | peak | 224167 | 224243 | 76 | + | macsForward_peak_8 | 4.40573 | CD630_01690 | antisense | - | Putative drug/sodium antiporter, MATE family | | | | | | | | |
| CD630 | MACS2 | peak | 224840 | 224968 | 128 | - | macsReverse_peak_4 | 4.67636 | CD630_01690 | sense | - | Putative drug/sodium antiporter, MATE family | | | | | | | | |
| CD630 | MACS2 | peak | 225211 | 225452 | 241 | - | macsReverse_peak_5 | 4.97234 | CD630_01690 | sense | - | Putative drug/sodium antiporter, MATE family | CD630_01700 | sense | - | ABC-type transport system, ATP-binding protein | | | | |
| CD630 | MACS2 | peak | 225890 | 226089 | 199 | - | macsReverse_peak_6 | 3.29194 | CD630_01700 | sense | - | ABC-type transport system, ATP-binding protein | | | | | | | | |
| CD630 | MACS2 | peak | 226794 | 226898 | 104 | - | macsReverse_peak_7 | 3.31800 | CD630_01700 | sense | - | ABC-type transport system, ATP-binding protein | | | | | | | | |
| CD630 | MACS2 | peak | 233846 | 233901 | 55 | - | macsReverse_peak_8 | 5.09543 | CD630_01760 | antisense | - | Putative oxidoreductase, NAD/FAD binding subunit | | | | | | | | |
| CD630 | MACS2 | peak | 240965 | 241018 | 53 | - | macsReverse_peak_9 | 3.80984 | Rcd2 | sense | | | | | | | | | | |
| CD630 | MACS2 | peak | 305369 | 305433 | 64 | + | macsForward_peak_9 | 3.52469 | CD630_02410 | sense | - | Phosphate-threonine kinase | CD630_02420 | sense | - | Phosphothreonine N-acetylglucosamine catalysis | | | | |
| CD630 | MACS2 | peak | 305961 | 306094 | 133 | + | macsForward_peak_10 | 3.71048 | CD630_02420 | sense | - | Phosphothreonine N-acetylglucosamine catalysis | CD630_02430 | sense | - | Methyltransferase | | | | |
| CD630 | MACS2 | peak | 308843 | 308955 | 112 | + | macsForward_peak_11 | 6.59405 | CD630_Cdi1_3 | sense | - | GEMM RNA motif | CD630_SQ173 | sense | - | ncRNA IGR, riboswitch c-di-GMP | | | | |
| CD630 | MACS2 | peak | 309319 | 309461 | 142 | + | macsForward_peak_12 | 3.25736 | CD630_02450 | sense | flgB | Flagellar basal-body rod protein FlgB | | | | | | | | |
| CD630 | MACS2 | peak | 312782 | 313125 | 343 | - | macsReverse_peak_10 | 4.59330 | CD630_02490 | antisense | fliG | Flagellar motor switch protein FliG | CD630_n00080 | sense | - | Antisense CDS | CD630_02500 | antisense | fliH | Flagellar assembly protein FliH |
| CD630 | MACS2 | peak | 314885 | 314937 | 52 | + | macsForward_peak_13 | 3.03666 | CD630_02510 | sense | fliI | ATP synthase subunit beta FliI | | | | | | | | |
| CD630 | MACS2 | peak | 322160 | 322289 | 129 | + | macsForward_peak_14 | 3.55524 | CD630_02620 | sense | flhB | Bifunctional flagellar biosynthesis protein FliR/FlhB | | | | | | | | |
| CD630 | MACS2 | peak | 332231 | 332297 | 66 | + | macsForward_peak_15 | 3.38938 | CD630_02720 | sense | - | Conserved hypothetical protein | CD630_n00120 | antisense | - | Antisense CDS | | | | |
| CD630 | MACS2 | peak | 334398 | 334452 | 54 | + | macsForward_peak_16 | 3.45147 | CD630_02730 | sense | htpG | Heat shock protein 90 (Heat shock protein HtpG)(High temperature protein G) | | | | | | | | |
| CD630 | MACS2 | peak | 335774 | 335999 | 225 | + | macsForward_peak_17 | 5.02277 | CD630_02740 | sense | - | Putative iron-containing alcohol dehydrogenase | | | | | | | | |
| CD630 | MACS2 | peak | 336350 | 336640 | 290 | + | macsForward_peak_18 | 4.44823 | CD630_02740 | sense | - | Putative iron-containing alcohol dehydrogenase | | | | | | | | |
| CD630 | MACS2 | peak | 340447 | 340705 | 258 | + | macsForward_peak_19 | 4.77661 | CD630_02790 | sense | - | Conserved hypothetical protein | | | | | | | | |
| CD630 | MACS2 | peak | 343199 | 343255 | 56 | + | macsForward_peak_20 | 4.89175 | CD630_02820 | antisense | - | Putative PHP hydrolase | | | | | | | | |
| CD630 | MACS2 | peak | 345668 | 345977 | 309 | + | macsForward_peak_21 | 3.57106 | CD630_02830 | sense | - | Transcription antiterminator, PTS operon regulator | | | | | | | | |
| CD630 | MACS2 | peak | 363387 | 363665 | 278 | + | macsForward_peak_22 | 3.71650 | CD630_03010 | sense | rbsA | ABC-type transport system, ribose-specific ATP-binding prote | CD630_03020 | sense | rbsC | ABC-type transport system, ribose-specific permease | | | | |
| CD630 | MACS2 | peak | 363918 | 364028 | 110 | + | macsForward_peak_23 | 3.58678 | CD630_03020 | sense | rbsC | ABC-type transport system, ribose-specific permease | | | | | | | | |
| CD630 | MACS2 | peak | 366635 | 366762 | 127 | + | macsForward_peak_24 | 3.51403 | CD630_03040 | sense | - | Conserved hypothetical protein, DUF1355 family | | | | | | | | |
| CD630 | MACS2 | peak | 384040 | 384343 | 303 | - | macsReverse_peak_11 | 3.66946 | CD630_03160 | sense | - | ABC-type transport system, permease | CD630_03170 | sense | - | ABC-type transport system, permease | | | | |
| CD630 | MACS2 | peak | 385245 | 385887 | 642 | - | macsReverse_peak_12 | 3.73138 | CD630_03180 | sense | - | ABC-type transport system, bacitracin/multidrug-family ATP-binding protein | | | | | | | | |
| CD630 | MACS2 | peak | 388326 | 388533 | 207 | + | macsForward_peak_25 | 3.36746 | CD630_03220 | sense | - | Putative ATPase | | | | | | | | |
| CD630 | MACS2 | peak | 391029 | 391210 | 181 | - | macsReverse_peak_13 | 5.05264 | CD630_s0080 | antisense | - | Cobalamin, CD630_03240 | | | | | | | | |
| CD630 | MACS2 | peak | 392744 | 393970 | 1226 | + | macsForward_peak_26 | 3.79830 | CD630_03270 | sense | cbiO | ABC-type transport system, cobalt-specific ATP-binding prote | CD630_03260 | sense | cbiQ | ABC-type transport system, cobalt-specific permease | | | | |
| CD630 | MACS2 | peak | 413318 | 413436 | 118 | - | macsReverse_peak_14 | 3.71996 | CD630_03400 | sense | - | Conserved hypothetical protein | | | | | | | | |
| CD630 | MACS2 | peak | 413650 | 414387 | 737 | - | macsReverse_peak_15 | 3.57284 | CD630_03400 | sense | - | Conserved hypothetical protein | | | | | | | | |
| CD630 | MACS2 | peak | 415311 | 415555 | 244 | - | macsReverse_peak_16 | 4.06000 | CD630_03400 | sense | - | Conserved hypothetical protein | | | | | | | | |
| CD630 | MACS2 | peak | 417422 | 417824 | 402 | - | macsReverse_peak_17 | 3.74827 | CD630_03400 | sense | - | Conserved hypothetical protein | | | | | | | | |
| CD630 | MACS2 | peak | 418199 | 418776 | 577 | - | macsReverse_peak_18 | 4.45994 | CD630_03400 | sense | - | Conserved hypothetical protein | CD630_03410 | sense | - | Conserved hypothetical protein | | | | |
| CD630 | MACS2 | peak | 419132 | 419193 | 61 | - | macsReverse_peak_19 | 4.71868 | CD630_03420 | sense | - | Conserved hypothetical protein | | | | | | | | |
| CD630 | MACS2 | peak | 428918 | 429100 | 182 | + | macsForward_peak_27 | 3.79913 | CD630_03550 | antisense | int | Integrase Tn916-like, CTn1-Orf1 | | | | | | | | |
| CD630 | MACS2 | peak | 436713 | 436975 | 262 | - | macsReverse_peak_20 | 3.93697 | CD630_03650 | sense | - | ABC-type transport system, multidrug-family permease Tn916-like, CTn1-Orf10 | | | | | | | | |
| CD630 | MACS2 | peak | 463848 | 464891 | 1043 | - | macsReverse_peak_21 | 4.15934 | CD630_03920 | sense | - | Putative radical SAM-family protein | CD630_03930 | sense | - | Putative membrane protein | | | | |
| CD630 | MACS2 | peak | 507351 | 507420 | 69 | - | macsReverse_peak_22 | 5.64455 | . | | . | NA | | | | | | | | |
| CD630 | MACS2 | peak | 507400 | 507451 | 51 | + | macsForward_peak_28 | 6.31159 | . | | . | NA | | | | | | | | |
| CD630 | MACS2 | peak | 508664 | 508897 | 233 | + | macsForward_peak_29 | 3.61457 | CD630_04280 | sense | - | Transcriptional regulator, AraC family Tn1549-like, CTn2-Orf23 | | | | | | | | |
| CD630 | MACS2 | peak | 512162 | 512386 | 224 | + | macsForward_peak_30 | 3.68342 | CD630_04320 | sense | - | ABC-type transport system, multidrug-family permease Tn1549-like, CTn2-Orf27 | | | | | | | | |
| CD630 | MACS2 | peak | 515402 | 515795 | 393 | - | macsReverse_peak_23 | 3.56926 | CD630_04340 | antisense | - | Putative drug/sodium antiporter, MATE family, Tn1549-like, CTn2-Orf29 | | | | | | | | |
| CD630 | MACS2 | peak | 516786 | 516842 | 56 | + | macsForward_peak_31 | 4.29262 | CD630_04341 | sense | - | Putative conjugative transposon protein Tn1549-like, CTn2-Orf30 | | | | | | | | |
| CD630 | MACS2 | peak | 519681 | 519820 | 139 | - | macsReverse_peak_24 | 3.10813 | CD630_04360 | antisense | - | Putative recombinase Tn1549-like, CTn2-Orf34 | | | | | | | | |
| CD630 | MACS2 | peak | 523157 | 523420 | 263 | - | macsReverse_peak_25 | 8.81823 | CD630_04401 | antisense | - | Putative phage protein | CD630_n00150 | sense | - | Antisense CDS | | | | |
| CD630 | MACS2 | peak | 524938 | 525140 | 202 | + | macsForward_peak_32 | 3.90451 | CD630_04410 | sense | - | Transcriptional regulator, sigma-54-dependent | | | | | | | | |
| CD630 | MACS2 | peak | 539473 | 539540 | 67 | + | macsForward_peak_33 | 3.33556 | CD630_04540 | sense | - | Transcriptional regulator, AraC family | | | | | | | | |
| CD630 | MACS2 | peak | 540213 | 540588 | 375 | + | macsForward_peak_34 | 3.05227 | CD630_04550 | sense | - | Putative helicase, UvrD family | | | | | | | | |
| CD630 | MACS2 | peak | 550210 | 550343 | 133 | - | macsReverse_peak_26 | 3.83025 | CD630_04610 | sense | - | Two-component sensor histidine kinase | | | | | | | | |
| CD630 | MACS2 | peak | 551418 | 551535 | 117 | - | macsReverse_peak_27 | 3.45413 | CD630_04620 | sense | - | Two-component response regulator | | | | | | | | |
| CD630 | MACS2 | peak | 555208 | 555579 | 371 | + | macsForward_peak_35 | 4.32791 | CD630_04660 | sense | - | Transcriptional regulator, RpiR family | | | | | | | | |
| CD630 | MACS2 | peak | 562266 | 562518 | 252 | - | macsReverse_peak_28 | 3.73674 | CD630_04710 | sense | blaI | Transcriptional regulator, Penicillinase repressor | | | | | | | | |
| CD630 | MACS2 | peak | 566718 | 566775 | 57 | + | macsForward_peak_36 | 3.58478 | CD630_04770 | sense | - | Conserved hypothetical protein | | | | | | | | |
| CD630 | MACS2 | peak | 570581 | 570717 | 136 | + | macsForward_peak_37 | 3.04530 | CD630_04820 | sense | spaK | Two-component sensor histidine kinase | | | | | | | | |
| CD630 | MACS2 | peak | 575235 | 575339 | 104 | - | macsReverse_peak_29 | 4.40788 | CD630_04850 | sense | - | Two-component sensor histidine kinase | | | | | | | | |
| CD630 | MACS2 | peak | 578881 | 579305 | 424 | + | macsForward_peak_38 | 4.62291 | CD630_SQ327 | sense | - | ncRNA IGR | CD630_RNA_5 | sense | - | ykkC-yxkD | CD630_04880 | sense | - | Putative small multidrug resistance SugE-like protein |
| CD630 | MACS2 | peak | 583643 | 583811 | 168 | + | macsForward_peak_39 | 4.20948 | CD630_04940 | sense | - | PTS system, mannose/fructose/sorbose IID component | | | | | | | | |
| CD630 | MACS2 | peak | 589554 | 590108 | 554 | + | macsForward_peak_40 | 4.02442 | CD630_05000 | sense | - | Putative antirestriction protein Tn5397, CTn3-Orf18 | | | | | | | | |
| CD630 | MACS2 | peak | 593130 | 593408 | 278 | + | macsForward_peak_41 | 4.03823 | CD630_05030 | sense | - | Putative membrane protein Tn5397, CTn3-Orf15 | | | | | | | | |
| CD630 | MACS2 | peak | 612995 | 613201 | 206 | - | macsReverse_peak_30 | 3.69875 | CD630_05150 | sense | - | D-alanyl-D-alanine carboxypeptidase, S11 peptidase family | | | | | | | | |
| CD630 | MACS2 | peak | 614569 | 614668 | 99 | - | macsReverse_peak_31 | 3.69285 | CD630_05160 | sense | - | Transcription antiterminator | | | | | | | | |
| CD630 | MACS2 | peak | 614935 | 615320 | 385 | - | macsReverse_peak_32 | 4.46990 | CD630_05160 | sense | - | Transcription antiterminator | | | | | | | | |
| CD630 | MACS2 | peak | 615943 | 616685 | 742 | - | macsReverse_peak_33 | 5.03573 | CD630_05160 | sense | - | Transcription antiterminator | | | | | | | | |
| CD630 | MACS2 | peak | 616895 | 616958 | 63 | - | macsReverse_peak_34 | 4.03396 | CD630_05160 | sense | - | Transcription antiterminator | | | | | | | | |
| CD630 | MACS2 | peak | 623209 | 623299 | 90 | - | macsReverse_peak_35 | 3.97048 | CD630_05211 | sense | - | Conserved hypothetical protein | | | | | | | | |
| CD630 | MACS2 | peak | 625303 | 625398 | 95 | + | macsForward_peak_42 | 3.65057 | CD630_05220 | sense | - | Putative c-di-GMP regulatory protein | | | | | | | | |
| CD630 | MACS2 | peak | 628195 | 628402 | 207 | + | macsForward_peak_43 | 3.73787 | CD630_05230 | sense | - | Putative transcriptional regulator | | | | | | | | |
| CD630 | MACS2 | peak | 632942 | 633364 | 422 | - | macsReverse_peak_36 | 3.91665 | CD630_05280 | sense | - | Putative amidohydrolase | | | | | | | | |
| CD630 | MACS2 | peak | 634294 | 634497 | 203 | - | macsReverse_peak_37 | 3.97604 | CD630_05290 | sense | - | Putative membrane protein | | | | | | | | |
| CD630 | MACS2 | peak | 634967 | 635438 | 471 | - | macsReverse_peak_38 | 4.56885 | CD630_05300 | sense | - | Putative membrane protein | | | | | | | | |
| CD630 | MACS2 | peak | 640754 | 640933 | 179 | + | macsForward_peak_44 | 3.72371 | CD630_05360 | sense | cheW | Purine-binding chemotaxis protein CheW | | | | | | | | |
| CD630 | MACS2 | peak | 641340 | 641642 | 302 | + | macsForward_peak_45 | 3.26947 | CD630_05370 | sense | - | Putative CheY-like chemosensory protein | | | | | | | | |

| | | | | | | | | | | | | | | | | | | | | |
|---|---|---|---|---|---|---|---|---|---|---|---|---|---|---|---|---|---|---|---|---|
| CD630 | MACS2 | peak | 650477 | 650715 | 238 | + | macsForward_peak_46 | 5.04761 | CD630_05450 | sense | - | Putative lipoprotein | | | | | | | | |
| CD630 | MACS2 | peak | 651395 | 651820 | 425 | + | macsForward_peak_47 | 3.98753 | CD630_05460 | sense | - | Putative membrane protein, DUF975 family | | | | | | | | |
| CD630 | MACS2 | peak | 652713 | 652775 | 62 | + | macsForward_peak_48 | 4.27854 | CD630_05480 | sense | - | Putative penicillin-binding peptidase BlaR1-like, M56 family | | | | | | | | |
| CD630 | MACS2 | peak | 654165 | 654460 | 295 | + | macsForward_peak_49 | 3.89852 | CD630_05490 | sense | - | Conserved hypothetical protein | | | | | | | | |
| CD630 | MACS2 | peak | 654703 | 654813 | 110 | + | macsForward_peak_50 | 4.08290 | CD630_SQ367 | sense | - | ncRNA IGR | | | | | | | | |
| CD630 | MACS2 | peak | 666332 | 666592 | 260 | + | macsForward_peak_51 | 3.39670 | CD630_05590 | sense | - | Transcriptional regulator, ArsR family | | | | | | | | |
| CD630 | MACS2 | peak | 683176 | 683270 | 94 | - | macsReverse_peak_39 | 3.27149 | CD630_s0160 | antisense | - | T-box (Thr) | | | | | | | | |
| CD630 | MACS2 | peak | 687984 | 688038 | 54 | + | macsForward_peak_52 | 3.64614 | CD630_05760 | sense | virS | Sensor histidine kinase | | | | | | | | |
| CD630 | MACS2 | peak | 688246 | 688382 | 136 | + | macsForward_peak_53 | 4.74085 | CD630_05760 | sense | virS | Sensor histidine kinase | | | | | | | | |
| CD630 | MACS2 | peak | 689914 | 690240 | 326 | + | macsForward_peak_54 | 5.39919 | CD630_05770 | antisense | - | Conserved hypothetical protein | | | | | | | | |
| CD630 | MACS2 | peak | 696893 | 697086 | 193 | + | macsForward_peak_55 | 3.36528 | CD630_05820 | sense | - | Putative pyruvate phosphate dikinase, PEP/pyruvate-binding | | | | | | | | |
| CD630 | MACS2 | peak | 698159 | 698289 | 130 | + | macsForward_peak_56 | 3.75661 | CD630_05820 | sense | - | Putative pyruvate phosphate dikinase, PEP/pyruvate-binding | CD630_05821 | sense | - | Conserved hypothetical protein | | | | |
| CD630 | MACS2 | peak | 698501 | 698720 | 219 | - | macsReverse_peak_40 | 4.28113 | CD630_05822 | sense | - | Fragment of putative signaling protein precursor (N-terminal region) | | | | | | | | |
| CD630 | MACS2 | peak | 699128 | 699282 | 154 | - | macsReverse_peak_41 | 3.35360 | CD630_05830 | sense | - | Putative diguanylate kinase signaling protein | | | | | | | | |
| CD630 | MACS2 | peak | 700070 | 700280 | 210 | - | macsReverse_peak_42 | 4.28779 | CD630_05840 | sense | - | Putative diguanylate kinase signaling protein | | | | | | | | |
| CD630 | MACS2 | peak | 704966 | 705546 | 580 | + | macsForward_peak_57 | 4.28339 | CD630_05870 | sense | - | Conserved hypothetical protein | CD630_05880 | sense | - | Conserved hypothetical protein | | | | |
| CD630 | MACS2 | peak | 712502 | 712627 | 125 | + | macsForward_peak_58 | 3.21934 | CD630_05900 | sense | - | Conserved hypothetical protein | | | | | | | | |
| CD630 | MACS2 | peak | 718696 | 718803 | 107 | + | macsForward_peak_59 | 4.18595 | CD630_05940 | sense | - | Conserved hypothetical protein | | | | | | | | |
| CD630 | MACS2 | peak | 719313 | 719466 | 153 | + | macsForward_peak_60 | 3.58685 | CD630_05940 | sense | - | Conserved hypothetical protein | | | | | | | | |
| CD630 | MACS2 | peak | 719844 | 719928 | 84 | + | macsForward_peak_61 | 3.31669 | CD630_05950 | sense | - | Putative nuclease, ParB-like | | | | | | | | |
| CD630 | MACS2 | peak | 723826 | 723907 | 81 | + | macsForward_peak_62 | 3.16772 | CD630_06010 | sense | ogt1 | Methylated-DNA-(protein)-cysteine S-methyltransferase 1 | | | | | | | | |
| CD630 | MACS2 | peak | 726526 | 726837 | 311 | + | macsForward_peak_63 | 3.59332 | CD630_06050 | sense | - | Fragment of conserved hypothetical protein (N-terminal region) | | | | | | | | |
| CD630 | MACS2 | peak | 727481 | 727627 | 146 | + | macsForward_peak_64 | 3.92696 | CD630_06060_p1 | sense | - | Fragment of chloride ion channel protein (part 1) | | | | | | | | |
| CD630 | MACS2 | peak | 729356 | 730057 | 701 | + | macsForward_peak_65 | 4.94050 | CD630_06060_p2 | sense | - | Fragment of chloride ion channel protein (part 2) | CD630_06060_p3 | sense | - | Fragment of chloride ion channel protein (part 3) | CD630_n00220 | sense | - | ncRNA IGR |
| CD630 | MACS2 | peak | 730360 | 730844 | 484 | - | macsReverse_peak_43 | 3.82315 | CD630_06080 | sense | - | Conserved hypothetical protein | | | | | | | | |
| CD630 | MACS2 | peak | 732355 | 732945 | 590 | - | macsReverse_peak_44 | 4.43918 | CD630_06100 | sense | - | Two-component sensor histidine kinase | | | | | | | | |
| CD630 | MACS2 | peak | 733951 | 734541 | 590 | - | macsReverse_peak_45 | 5.61380 | CD630_06110 | sense | - | Two-component response regulator | | | | | | | | |
| CD630 | MACS2 | peak | 739167 | 739290 | 123 | - | macsReverse_peak_46 | 4.12519 | CD630_06140 | sense | - | Conserved hypothetical protein | | | | | | | | |
| CD630 | MACS2 | peak | 739569 | 740604 | 1035 | - | macsReverse_peak_47 | 5.51744 | CD630_06150 | sense | - | Transcriptional regulator, TetR family | CD630_06160 | sense | - | Transcriptional regulator, MerR family | | | | |
| CD630 | MACS2 | peak | 741525 | 742062 | 537 | - | macsReverse_peak_48 | 4.95366 | CD630_06170 | sense | - | Putative membrane protein | | | | | | | | |
| CD630 | MACS2 | peak | 743177 | 743254 | 77 | - | macsReverse_peak_49 | 3.57064 | CD630_06190 | sense | - | Conserved hypothetical protein | CD630_06200 | sense | - | Conserved hypothetical protein | | | | |
| CD630 | MACS2 | peak | 756697 | 756858 | 161 | - | macsReverse_peak_50 | 4.03510 | CD630_06330 | antisense | - | Conserved hypothetical protein | | | | | | | | |
| CD630 | MACS2 | peak | 757741 | 758342 | 601 | + | macsForward_peak_66 | 4.35983 | CD630_06340 | sense | - | Conserved hypothetical protein | CD630_06350 | sense | - | Putative hydrolase | | | | |
| CD630 | MACS2 | peak | 761255 | 761321 | 66 | - | macsReverse_peak_51 | 4.44797 | CD630_06380 | antisense | - | Conserved hypothetical protein | | | | | | | | |
| CD630 | MACS2 | peak | 761326 | 761438 | 112 | + | macsForward_peak_67 | 4.32211 | CD630_06380 | sense | - | Conserved hypothetical protein | | | | | | | | |
| CD630 | MACS2 | peak | 763807 | 763862 | 55 | + | macsForward_peak_68 | 4.14404 | CD630_06410 | sense | - | Putative membrane protein | | | | | | | | |
| CD630 | MACS2 | peak | 765254 | 765407 | 153 | + | macsForward_peak_69 | 7.27863 | CD630_SQ408 | sense | - | ncRNA IGR | | | | | | | | |
| CD630 | MACS2 | peak | 779822 | 779928 | 106 | + | macsForward_peak_70 | 3.54673 | CD630_06540 | sense | - | ABC-type transport system, permease protein | | | | | | | | |
| CD630 | MACS2 | peak | 782205 | 782259 | 54 | + | macsForward_peak_71 | 3.90214 | . | | . | NA | | | | | | | | |
| CD630 | MACS2 | peak | 782989 | 783253 | 264 | + | macsForward_peak_72 | 4.02532 | CD630_06560 | sense | - | Chloride ion channel protein | | | | | | | | |
| CD630 | MACS2 | peak | 784676 | 784882 | 206 | + | macsForward_peak_73 | 2.97335 | CD630_06570 | sense | cdu2 | Na(+)/H(+) antiporter | | | | | | | | |
| CD630 | MACS2 | peak | 785310 | 785596 | 286 | + | macsForward_peak_74 | 4.46276 | CD630_06580 | sense | cdu1 | Transcriptional regulator, beta-lactams repressor | | | | | | | | |
| CD630 | MACS2 | peak | 796140 | 796813 | 673 | + | macsForward_peak_75 | 4.44902 | CD630_06630 | sense | tcdA | Toxin A | | | | | | | | |
| CD630 | MACS2 | peak | 796645 | 796721 | 76 | - | macsReverse_peak_52 | 4.21176 | CD630_06630 | antisense | tcdA | Toxin A | | | | | | | | |
| CD630 | MACS2 | peak | 797055 | 798346 | 1291 | + | macsForward_peak_76 | 4.70593 | CD630_06630 | sense | tcdA | Toxin A | | | | | | | | |
| CD630 | MACS2 | peak | 798786 | 799377 | 591 | + | macsForward_peak_77 | 4.08197 | CD630_06630 | sense | tcdA | Toxin A | | | | | | | | |
| CD630 | MACS2 | peak | 799650 | 799801 | 151 | + | macsForward_peak_78 | 4.10852 | CD630_06630 | sense | tcdA | Toxin A | | | | | | | | |
| CD630 | MACS2 | peak | 800757 | 800988 | 231 | + | macsForward_peak_79 | 4.23648 | CD630_06630 | sense | tcdA | Toxin A | | | | | | | | |
| CD630 | MACS2 | peak | 801904 | 801967 | 63 | + | macsForward_peak_80 | 3.63897 | CD630_06630 | sense | tcdA | Toxin A | | | | | | | | |
| CD630 | MACS2 | peak | 802184 | 802953 | 769 | + | macsForward_peak_81 | 3.74811 | CD630_06630 | sense | tcdA | Toxin A | | | | | | | | |
| CD630 | MACS2 | peak | 803193 | 803476 | 283 | + | macsForward_peak_82 | 3.30198 | CD630_06630 | sense | tcdA | Toxin A | | | | | | | | |
| CD630 | MACS2 | peak | 803730 | 804014 | 284 | + | macsForward_peak_83 | 4.51777 | CD630_06630 | sense | tcdA | Toxin A | | | | | | | | |
| CD630 | MACS2 | peak | 804358 | 804529 | 171 | - | macsReverse_peak_53 | 3.76101 | CD630_06640 | sense | tcdC | Negative regulator of toxin gene expression | | | | | | | | |
| CD630 | MACS2 | peak | 805860 | 806116 | 256 | - | macsReverse_peak_54 | 3.32967 | CD630_06642 | sense | - | Conserved hypothetical protein | | | | | | | | |
| CD630 | MACS2 | peak | 811114 | 811258 | 144 | + | macsForward_peak_84 | 3.61207 | CD630_06690 | sense | - | Two-component sensor histidine kinase | | | | | | | | |
| CD630 | MACS2 | peak | 811583 | 812005 | 422 | + | macsForward_peak_85 | 5.82709 | CD630_06690 | sense | - | Two-component sensor histidine kinase | CD630_n00240 | sense | - | ncRNA IGR | | | | |
| CD630 | MACS2 | peak | 831084 | 831154 | 70 | + | macsForward_peak_86 | 3.66276 | . | | . | NA | | | | | | | | |
| CD630 | MACS2 | peak | 840165 | 840546 | 381 | - | macsReverse_peak_55 | 3.13260 | CD630_06920 | sense | - | Putative metallo-hydrolase/oxidoreductase | | | | | | | | |
| CD630 | MACS2 | peak | 841749 | 841876 | 127 | + | macsForward_peak_87 | 3.27218 | CD630_06930 | antisense | - | Transcriptional regulator, MerR family | | | | | | | | |
| CD630 | MACS2 | peak | 846024 | 846141 | 117 | + | macsForward_peak_88 | 3.16317 | CD630_06960 | sense | - | Putative cation transport protein, Trk family | | | | | | | | |
| CD630 | MACS2 | peak | 847836 | 848044 | 208 | + | macsForward_peak_89 | 3.82312 | CD630_06970 | sense | - | Putative regulator of K+ conductance | | | | | | | | |
| CD630 | MACS2 | peak | 860962 | 861261 | 299 | + | macsForward_peak_90 | 3.41054 | CD630_07070 | sense | - | Putative diguanylate kinase signaling protein | | | | | | | | |
| CD630 | MACS2 | peak | 861845 | 862040 | 195 | + | macsForward_peak_91 | 3.46414 | CD630_07070 | sense | - | Putative diguanylate kinase signaling protein | | | | | | | | |
| CD630 | MACS2 | peak | 869949 | 870094 | 145 | + | macsForward_peak_92 | 4.96036 | CD630_07110 | sense | argS | Arginyl-tRNA synthetase (Arginine--tRNA ligase) (ArgRS) | | | | | | | | |
| CD630 | MACS2 | peak | 870959 | 871203 | 244 | - | macsReverse_peak_56 | 3.26177 | CD630_07120 | antisense | - | Conserved hypothetical protein | CD630_07121 | sense | - | Conserved hypothetical protein | | | | |
| CD630 | MACS2 | peak | 902630 | 903539 | 909 | - | macsReverse_peak_57 | 4.52273 | CD630_07400 | sense | - | Putative pyridoxal phosphate-dependent aminotransferase | CD630_SQ476 | antisense | - | Antisense CDS | | | | |
| CD630 | MACS2 | peak | 906407 | 906547 | 140 | - | macsReverse_peak_58 | 4.41437 | CD630_07420 | antisense | - | Putative ethanolamine transporter | | | | | | | | |
| CD630 | MACS2 | peak | 906925 | 907585 | 660 | - | macsReverse_peak_59 | 4.65568 | CD630_07430 | sense | - | Transcriptional regulator, DeoR family | | | | | | | | |
| CD630 | MACS2 | peak | 910388 | 910618 | 230 | + | macsForward_peak_93 | 3.37143 | CD630_07470 | sense | - | Putative lipoprotein | | | | | | | | |
| CD630 | MACS2 | peak | 911118 | 911241 | 123 | + | macsForward_peak_94 | 3.10955 | CD630_07470 | sense | - | Putative lipoprotein | | | | | | | | |
| CD630 | MACS2 | peak | 912318 | 912649 | 331 | + | macsForward_peak_95 | 5.18321 | CD630_07480 | sense | - | Putative diguanylate kinase signaling protein | | | | | | | | |
| CD630 | MACS2 | peak | 917517 | 917679 | 162 | + | macsForward_peak_96 | 4.98480 | CD630_s0190 | sense | - | T-box | | | | | | | | |
| CD630 | MACS2 | peak | 920250 | 920371 | 121 | - | macsReverse_peak_60 | 3.85358 | CD630_n00290 | sense | - | ncRNA IGR | | | | | | | | |
| CD630 | MACS2 | peak | 938403 | 938533 | 130 | + | macsForward_peak_97 | 3.50140 | CD630_07660 | sense | srlEb | PTS system, sorbitol-specific IIB C-terminal component (Glucitol) | | | | | | | | |
| CD630 | MACS2 | peak | 938876 | 938999 | 123 | - | macsReverse_peak_61 | 4.14905 | CD630_07670 | antisense | srlB | PTS system, sorbitol-specific IIA component (Glucitol) | CD630_SQ495 | sense | | | | | | |
| CD630 | MACS2 | peak | 939229 | 939939 | 710 | + | macsForward_peak_98 | 4.19843 | CD630_SQ495 | antisense | - | Antisense CDS | CD630_07680 | sense | srlD | Sorbitol 6-phosphate 2-dehydrogenase (Glucitol) | | | | |
| CD630 | MACS2 | peak | 947399 | 947511 | 112 | + | macsForward_peak_99 | 3.64186 | CD630_07760 | sense | - | Transcriptional regulator, LacI family | | | | | | | | |
| CD630 | MACS2 | peak | 948160 | 948233 | 73 | + | macsForward_peak_100 | 3.31158 | CD630_07770 | sense | - | Putative membrane protein | | | | | | | | |
| CD630 | MACS2 | peak | 955487 | 956332 | 845 | + | macsForward_peak_101 | 5.82750 | CD630_07830 | antisense | spoIVB | Stage IV sporulation protein SpoIVB, S55 peptidase family | | | | | | | | |
| CD630 | MACS2 | peak | 956738 | 957576 | 838 | + | macsForward_peak_102 | 4.26761 | CD630_07840 | sense | - | Putative N-acetylmuramoyl-L-alanine amidase | | | | | | | | |
| CD630 | MACS2 | peak | 960172 | 960264 | 92 | - | macsReverse_peak_62 | 3.22829 | CD630_07870 | sense | - | Putative metal-binding protein | | | | | | | | |
| CD630 | MACS2 | peak | 967587 | 967765 | 178 | - | macsReverse_peak_63 | 3.22537 | CD630_07960 | sense | - | Transcriptional regulator, LacI family | | | | | | | | |
| CD630 | MACS2 | peak | 978746 | 978828 | 82 | + | macsForward_peak_103 | 3.41204 | CD630_08050 | sense | etfA | Electron transfer flavoprotein alpha-subunit | | | | | | | | |
| CD630 | MACS2 | peak | 980667 | 981284 | 617 | - | macsReverse_peak_64 | 4.87819 | CD630_08070 | sense | - | Conserved hypothetical protein | CD630_08080 | sense | - | Conserved hypothetical protein | | | | |
| CD630 | MACS2 | peak | 981768 | 981958 | 190 | + | macsForward_peak_104 | 3.34712 | CD630_08090 | sense | - | Conserved hypothetical protein | | | | | | | | |
| CD630 | MACS2 | peak | 982163 | 982285 | 122 | + | macsForward_peak_105 | 3.57167 | CD630_08090 | sense | - | Conserved hypothetical protein | | | | | | | | |

| | | | | | | | | | | | | | | | | | | | | |
|---|---|---|---|---|---|---|---|---|---|---|---|---|---|---|---|---|---|---|---|---|
| CD630 | MACS2 | peak | 982857 | 985134 | 2277 | + | macsForward_peak_106 | 4.39181 | CD630_08110 | sense | - | Putative cyclic di-GMP regulatory protein | CD630_08100 | sense | - | Putative flavodoxin/nitric oxide synthase | | | | |
| CD630 | MACS2 | peak | 986032 | 986216 | 184 | + | macsForward_peak_107 | 4.63831 | CD630_08130 | sense | - | Putative PTS system, IIB component | | | | | | | | |
| CD630 | MACS2 | peak | 986685 | 986987 | 302 | - | macsReverse_peak_65 | 3.30159 | CD630_08140 | sense | - | Transcriptional regulator, GntR family | | | | | | | | |
| CD630 | MACS2 | peak | 999756 | 999823 | 67 | - | macsReverse_peak_66 | 3.63265 | CD630_08240 | antisense | - | ABC-type transport system, multidrug-family permease | | | | | | | | |
| CD630 | MACS2 | peak | 1013264 | 1013436 | 172 | + | macsForward_peak_108 | 3.19339 | CD630_08360 | sense | - | Putative acetyltransferase | | | | | | | | |
| CD630 | MACS2 | peak | 1013664 | 1014152 | 488 | + | macsForward_peak_109 | 3.65639 | CD630_08370 | sense | - | Nitroreductase | | | | | | | | |
| CD630 | MACS2 | peak | 1015015 | 1015404 | 389 | + | macsForward_peak_110 | 5.16000 | CD630_08390 | sense | - | Putative membrane protein | | | | | | | | |
| CD630 | MACS2 | peak | 1022632 | 1022750 | 118 | - | macsReverse_peak_67 | 4.08933 | CD630_08460 | antisense | - | Conserved hypothetical protein | | | | | | | | |
| CD630 | MACS2 | peak | 1022648 | 1022849 | 201 | + | macsForward_peak_111 | 4.15083 | CD630_08460 | sense | - | Conserved hypothetical protein | | | | | | | | |
| CD630 | MACS2 | peak | 1027293 | 1027486 | 193 | - | macsReverse_peak_68 | 5.16329 | CD630_08510 | sense | - | Putative membrane protein | | | | | | | | |
| CD630 | MACS2 | peak | 1028740 | 1028883 | 143 | + | macsForward_peak_112 | 2.98328 | CD630_08530 | sense | oppB | ABC-type transport system, oligopeptide-family permease | | | | | | | | |
| CD630 | MACS2 | peak | 1029461 | 1029687 | 226 | + | macsForward_peak_113 | 3.01695 | CD630_08540 | sense | oppC | ABC-type transport system, oligopeptide-family permease | | | | | | | | |
| CD630 | MACS2 | peak | 1043083 | 1043321 | 238 | + | macsForward_peak_114 | 3.79973 | CD630_08660 | sense | - | Conserved hypothetical protein | | | | | | | | |
| CD630 | MACS2 | peak | 1043563 | 1043773 | 210 | + | macsForward_peak_115 | 3.86467 | CD630_08660 | sense | - | Conserved hypothetical protein | | | | | | | | |
| CD630 | MACS2 | peak | 1052169 | 1052276 | 107 | - | macsReverse_peak_69 | 6.84436 | . | | . | NA | | | | | | | | |
| CD630 | MACS2 | peak | 1052643 | 1052828 | 185 | - | macsReverse_peak_70 | 6.46464 | CD630_n00330 | sense | - | ncRNA IGR | | | | | | | | |
| CD630 | MACS2 | peak | 1052720 | 1052785 | 65 | + | macsForward_peak_116 | 5.75077 | CD630_n00330 | antisense | - | ncRNA IGR | | | | | | | | |
| CD630 | MACS2 | peak | 1054774 | 1055118 | 344 | - | macsReverse_peak_71 | 3.88765 | CD630_08780 | sense | - | ABC-type transport system, sugar-specific permease | | | | | | | | |
| CD630 | MACS2 | peak | 1055331 | 1055967 | 636 | - | macsReverse_peak_72 | 7.76690 | CD630_08780 | sense | - | ABC-type transport system, sugar-specific permease | | | | | | | | |
| CD630 | MACS2 | peak | 1065385 | 1065565 | 180 | + | macsForward_peak_117 | 3.28530 | CD630_08850 | sense | glgP | Glycogen phosphorylase | | | | | | | | |
| CD630 | MACS2 | peak | 1077593 | 1077766 | 173 | + | macsForward_peak_118 | 3.72225 | CD630_08940 | sense | - | Putative iron-dependent hydrogenase | | | | | | | | |
| CD630 | MACS2 | peak | 1078215 | 1078461 | 246 | + | macsForward_peak_119 | 3.71905 | CD630_08950 | sense | - | Metallo-beta-lactamase superfamily exported protein | | | | | | | | |
| CD630 | MACS2 | peak | 1078832 | 1078918 | 86 | + | macsForward_peak_120 | 3.32619 | CD630_08950 | sense | - | Metallo-beta-lactamase superfamily exported protein | | | | | | | | |
| CD630 | MACS2 | peak | 1081434 | 1081758 | 324 | - | macsReverse_peak_73 | 5.83478 | CD630_n00340 | sense | - | Antisense CDS | CD630_08980 | antisense | - | Conserved hypothetical protein, DUF1846-family | | | | |
| CD630 | MACS2 | peak | 1084464 | 1084849 | 385 | + | macsForward_peak_121 | 4.07035 | CD630_09000 | sense | opuCA | ABC-type transport system, glycine betaine/carnitine/choline | CD630_09010 | sense | opuCC | ABC-type transport system, glycine betaine/carnitine/choline permease | | | | |
| CD630 | MACS2 | peak | 1087797 | 1087980 | 183 | + | macsForward_peak_122 | 4.21282 | CD630_09030_p1 | sense | - | Fragment of putative oxydoreductase (Part 1) | | | | | | | | |
| CD630 | MACS2 | peak | 1088211 | 1088318 | 107 | - | macsReverse_peak_74 | 4.11851 | CD630_09040 | sense | - | Putative phage integrase | | | | | | | | |
| CD630 | MACS2 | peak | 1089041 | 1089195 | 154 | - | macsReverse_peak_75 | 3.34158 | CD630_09040 | sense | - | Putative phage integrase | | | | | | | | |
| CD630 | MACS2 | peak | 1089514 | 1090345 | 831 | - | macsReverse_peak_76 | 3.96216 | CD630_n00350 | antisense | - | ncRNA IGR | CD630_09050 | sense | - | Putative phage protein | | | | |
| CD630 | MACS2 | peak | 1091758 | 1091851 | 93 | - | macsReverse_peak_77 | 4.31834 | CD630_09060 | sense | - | Putative phage DNA-binding protein | | | | | | | | |
| CD630 | MACS2 | peak | 1093642 | 1093787 | 145 | + | macsForward_peak_123 | 5.06943 | . | | . | NA | | | | | CD630_n00360 | antisense | - | Antisense CDS |
| CD630 | MACS2 | peak | 1095645 | 1096163 | 518 | - | macsReverse_peak_78 | 5.92260 | CD630_09100 | sense | - | Putative phage protein | | | | | | | | |
| CD630 | MACS2 | peak | 1111772 | 1112132 | 360 | - | macsReverse_peak_79 | 4.75651 | CD630_09401 | antisense | - | Putative phage protein | CD630_09410 | sense | - | Putative phage protein | | | | |
| CD630 | MACS2 | peak | 1123859 | 1124330 | 471 | - | macsReverse_peak_80 | 7.05302 | CD630_09561 | antisense | - | Putative phage protein | CD630_09562 | antisense | - | Putative phage protein | CD630_n00370 | sense | - | Antisense CDS |
| CD630 | MACS2 | peak | 1124131 | 1124308 | 177 | + | macsForward_peak_124 | 3.94307 | CD630_09562 | sense | - | Putative phage protein | CD630_n00370 | antisense | - | Antisense CDS | | | | |
| CD630 | MACS2 | peak | 1124551 | 1124748 | 197 | - | macsReverse_peak_81 | 5.45183 | . | | . | NA | CD630_09563 | antisense | | | SQ606, SQ2063 | antisense | | |
| CD630 | MACS2 | peak | 1125576 | 1125711 | 135 | + | macsForward_peak_125 | 4.27916 | CD630_n00380 | sense | - | ncRNA IGR | | | | | | | | |
| CD630 | MACS2 | peak | 1126073 | 1126274 | 201 | + | macsForward_peak_126 | 5.31213 | CD630_09570 | sense | - | Putative phage lipoprotein | | | | | | | | |
| CD630 | MACS2 | peak | 1139749 | 1140647 | 898 | + | macsForward_peak_127 | 4.83410 | CD630_09750 | sense | - | Putative phage protein | CD630_09760 | sense | - | Putative phage protein | | | | |
| CD630 | MACS2 | peak | 1141416 | 1141495 | 79 | + | macsForward_peak_128 | 3.22106 | CD630_09770 | sense | - | Putative phage protein | | | | | | | | |
| CD630 | MACS2 | peak | 1141753 | 1141895 | 142 | - | macsReverse_peak_82 | 3.66033 | CD630_09772 | sense | - | Putative phage protein | | | | | | | | |
| CD630 | MACS2 | peak | 1142187 | 1142564 | 377 | - | macsReverse_peak_83 | 4.83391 | CD630_Cdi1_5 | sense | - | GEMM RNA motif | CD630_09771 | antisense | - | Putative phage protein | CD630_n00390 | sense | - | Antisense CDS |
| CD630 | MACS2 | peak | 1142221 | 1142393 | 172 | + | macsForward_peak_129 | 4.95362 | CD630_09771 | antisense | - | Putative phage protein | CD630_Cdi1_5 | sense | - | GEMM RNA motif | CD630_n00390 | antisense | - | Antisense CDS |
| CD630 | MACS2 | peak | 1147792 | 1148026 | 234 | + | macsForward_peak_130 | 4.50009 | CD630_09840 | sense | - | Putative membrane protein | | | | | | | | |
| CD630 | MACS2 | peak | 1148858 | 1149204 | 346 | + | macsForward_peak_131 | 3.94885 | CD630_09850 | sense | - | Putative drug/sodium antiporter, MATE family | | | | | | | | |
| CD630 | MACS2 | peak | 1149471 | 1149527 | 56 | + | macsForward_peak_132 | 4.18679 | CD630_09850 | sense | - | Putative drug/sodium antiporter, MATE family | | | | | | | | |
| CD630 | MACS2 | peak | 1152657 | 1152825 | 168 | + | macsForward_peak_133 | 4.02122 | CD630_s0210 | sense | - | T-box (Leu) | | | | | | | | |
| CD630 | MACS2 | peak | 1166520 | 1166641 | 121 | + | macsForward_peak_134 | 3.25917 | CD630_09970 | sense | - | Two-component sensor histidine kinase | | | | | | | | |
| CD630 | MACS2 | peak | 1169554 | 1169909 | 355 | + | macsForward_peak_135 | 3.86040 | CD630_09990 | sense | - | ABC-type transport system, nitrate/sulfonate/taurine extrace | CD630_10000 | sense | - | ABC-type transport system, nitrate/sulfonate/taurine permease | | | | |
| CD630 | MACS2 | peak | 1174597 | 1174760 | 163 | + | macsForward_peak_136 | 3.93750 | CD630_10050 | sense | - | Putative NAD-dependent malic enzyme | | | | | | | | |
| CD630 | MACS2 | peak | 1175765 | 1176089 | 324 | - | macsReverse_peak_84 | 3.75625 | CD630_10070 | antisense | - | Conserved hypothetical protein | CD630_10080 | sense | - | Conserved hypothetical protein | | | | |
| CD630 | MACS2 | peak | 1194079 | 1194628 | 549 | + | macsForward_peak_137 | 4.08233 | CD630_10220 | sense | - | Putative membrane protein | | | | | | | | |
| CD630 | MACS2 | peak | 1201466 | 1201677 | 211 | - | macsReverse_peak_85 | 7.37676 | CD630_10290 | antisense | - | Putative cell wall anchored protein | | | | | | | | |
| CD630 | MACS2 | peak | 1217176 | 1217345 | 169 | + | macsForward_peak_138 | 3.43363 | CD630_10360 | sense | cwp17 | Putative N-acetylmuramoyl-L-alanine amidase, autolysin | | | | | | | | |
| CD630 | MACS2 | peak | 1243172 | 1243296 | 124 | - | macsReverse_peak_86 | 4.36817 | CD630_10490 | sense | ppnK | Inorganic polyphosphate/ATP-NAD kinase (Poly(P)/ATP NAD kinase) | | | | | | | | |
| CD630 | MACS2 | peak | 1271397 | 1272091 | 694 | - | macsReverse_peak_87 | 4.45284 | CD630_10790 | sense | - | Transcriptional regulator, LysR family | | | | | | | | |
| CD630 | MACS2 | peak | 1273181 | 1273307 | 126 | + | macsForward_peak_139 | 5.30319 | CD630_10800 | sense | - | Putative lipoprotein | | | | | | | | |
| CD630 | MACS2 | peak | 1273668 | 1273976 | 308 | + | macsForward_peak_140 | 3.26118 | CD630_10810 | sense | moeA | Molybdenum cofactor biosynthesis protein | | | | | | | | |
| CD630 | MACS2 | peak | 1275473 | 1275676 | 203 | + | macsForward_peak_141 | 4.82398 | CD630_10830 | sense | - | PTS system, lactose/cellobiose specific IIB component | | | | | | | | |
| CD630 | MACS2 | peak | 1281720 | 1281785 | 65 | + | macsForward_peak_142 | 5.05048 | CD630_10880 | sense | - | Conserved hypothetical protein | | | | | | | | |
| CD630 | MACS2 | peak | 1287723 | 1287836 | 113 | - | macsReverse_peak_88 | 3.67675 | CD630_10950 | sense | - | ABC-type transport system, multidrug-family permease Tn1549-like, CTn4-Orf30 | | | | | | | | |
| CD630 | MACS2 | peak | 1288129 | 1288208 | 79 | - | macsReverse_peak_89 | 3.08470 | CD630_10950 | sense | - | ABC-type transport system, multidrug-family permease Tn1549-like, CTn4-Orf30 | | | | | | | | |
| CD630 | MACS2 | peak | 1289031 | 1289354 | 323 | - | macsReverse_peak_90 | 3.53850 | CD630_10960 | sense | - | ABC-type transport system, multidrug-family permease Tn15 | CD630_10970 | sense | - | ABC-type transport system, multidrug-family ATP-binding protein Tn1549-like, CTn4-Orf28 | | | | |
| CD630 | MACS2 | peak | 1319427 | 1319785 | 358 | + | macsForward_peak_143 | 3.52327 | CD630_11220 | sense | - | Transcriptional regulator, RmlC-type | | | | | | | | |
| CD630 | MACS2 | peak | 1322095 | 1322159 | 64 | + | macsForward_peak_144 | 3.20274 | CD630_11240 | sense | - | Conserved hypothetical protein | | | | | | | | |
| CD630 | MACS2 | peak | 1322368 | 1323012 | 644 | + | macsForward_peak_145 | 4.85723 | CD630_11241 | antisense | - | Conserved hypothetical protein | CD630_SQ726 | sense | - | ncRNA IGR | CD630_n00440 | sense | - | ncRNA IGR |
| CD630 | MACS2 | peak | 1322939 | 1323114 | 175 | - | macsReverse_peak_91 | 8.44287 | CD630_n00440 | antisense | - | ncRNA IGR | | | | | | | | |
| CD630 | MACS2 | peak | 1364088 | 1364168 | 80 | + | macsForward_peak_146 | 3.72587 | CD630_s0220 | sense | - | SAM | | | | | | | | |
| CD630 | MACS2 | peak | 1371511 | 1372307 | 796 | + | macsForward_peak_147 | 4.80521 | CD630_11720 | sense | etfA | Electron transfer flavoprotein subunit alpha | CD630_11730 | sense | - | Putative FAD-linked oxidase | | | | |
| CD630 | MACS2 | peak | 1391016 | 1391068 | 52 | - | macsReverse_peak_92 | 4.52971 | CD630_11890 | antisense | - | Putative amino acid/polyamine transporter I | | | | | | | | |
| CD630 | MACS2 | peak | 1392215 | 1392389 | 174 | + | macsForward_peak_148 | 3.80071 | CD630_11910 | sense | fbp | Fructose-1,6-bisphosphatase | | | | | | | | |
| CD630 | MACS2 | peak | 1393906 | 1393998 | 92 | + | macsForward_peak_149 | 3.69457 | CD630_11910 | sense | fbp | Fructose-1,6-bisphosphatase | | | | | | | | |
| CD630 | MACS2 | peak | 1400555 | 1400704 | 149 | + | macsForward_peak_150 | 4.92798 | CD630_12010 | sense | nusB | N utilization substance protein B homolog (Protein NusB) | CD630_12020 | sense | gcp | O-sialoglycoprotein endopeptidase, peptidase M22 family | | | | |
| CD630 | MACS2 | peak | 1410254 | 1410604 | 350 | - | macsReverse_peak_93 | 4.36078 | CD630_12110 | sense | - | Putative N-acetyltransferase | | | | | | | | |
| CD630 | MACS2 | peak | 1410827 | 1411124 | 297 | - | macsReverse_peak_94 | 4.49926 | CD630_12120 | sense | bltD | Diamine N-acetyltransferase | | | | | | | | |
| CD630 | MACS2 | peak | 1429418 | 1429703 | 285 | - | macsReverse_peak_95 | 3.08122 | CD630_12300_p1 | sense | - | RNA polymerase sigma-K factor SigK (Part 2) | | | | | | | | |
| CD630 | MACS2 | peak | 1431431 | 1431720 | 289 | - | macsReverse_peak_96 | 3.28667 | CD630_12320 | sense | - | Putative lipoprotein | | | | | | | | |
| CD630 | MACS2 | peak | 1432503 | 1432563 | 60 | + | macsForward_peak_151 | 4.44377 | CD630_12330 | sense | cwp26 | Putative cell surface protein | | | | | | | | |
| CD630 | MACS2 | peak | 1433909 | 1434428 | 519 | - | macsReverse_peak_97 | 6.85680 | CD630_12331 | antisense | - | Fragment of phage protein | CD630_SQ808 | sense | - | Antisense 3prime UTR | CD630_12332 | antisense | - | Conserved hypothetical protein |
| CD630 | MACS2 | peak | 1434611 | 1434973 | 362 | + | macsForward_peak_152 | 5.55447 | CD630_n00460 | sense | - | ncRNA IGR | | | | | | | | |
| CD630 | MACS2 | peak | 1439587 | 1439901 | 314 | - | macsReverse_peak_98 | 4.59861 | CD630_12360_p2 | antisense | - | Fragment of putative phage protein (part 2) | | | | | | | | |
| CD630 | MACS2 | peak | 1470405 | 1470836 | 431 | - | macsReverse_peak_99 | 6.31827 | CD630_12650 | sense | - | Transcriptional regulator, CarD family | | | | | | | | |
| CD630 | MACS2 | peak | 1503439 | 1503537 | 98 | + | macsForward_peak_153 | 3.33560 | CD630_12960 | sense | scpB | Segregation and condensation protein B | | | | | | | | |
| CD630 | MACS2 | peak | 1508381 | 1508547 | 166 | - | macsReverse_peak_100 | 3.52267 | CD630_n00470 | sense | - | ncRNA IGR | CD630_13020 | sense | - | Putative MazG-like pyrophosphohydrolase | | | | |
| CD630 | MACS2 | peak | 1549853 | 1550016 | 163 | + | macsForward_peak_154 | 4.15139 | CD630_13350 | sense | - | Transcriptional regulator, MarR family | | | | | | | | |
| CD630 | MACS2 | peak | 1551208 | 1551448 | 240 | + | macsForward_peak_155 | 3.77330 | CD630_13360 | sense | - | PTS system, glucose-like IIBC component | | | | | | | | |

| | | | | | | | | | | | | | | | | | | | | |
|---|---|---|---|---|---|---|---|---|---|---|---|---|---|---|---|---|---|---|---|---|
| CD630 | MACS2 | peak | 1552688 | 1552871 | 183 | + | macsForward_peak_156 | 4.33145 | CD630_13370 | sense | - | Transcriptional regulator, RpiR family | | | | | | | | |
| CD630 | MACS2 | peak | 1553367 | 1553620 | 253 | + | macsForward_peak_157 | 4.74408 | CD630_13380 | sense | - | Putative 6-phospho-beta-glucosidase | | | | | | | | |
| CD630 | MACS2 | peak | 1556407 | 1556819 | 412 | + | macsForward_peak_158 | 3.20232 | CD630_13400 | sense | - | Putative galactose-1-phosphate uridylyltransferase | | | | | | | | |
| CD630 | MACS2 | peak | 1560856 | 1561091 | 235 | + | macsForward_peak_159 | 3.87629 | CD630_13430 | sense | glnA | Glutamine synthetase | | | | | | | | |
| CD630 | MACS2 | peak | 1569289 | 1569430 | 141 | + | macsForward_peak_160 | 3.96216 | CD630_13520 | sense | cprK | Two-component sensor histidine kinase, cationic antimicrobial peptide resistance | | | | | | | | |
| CD630 | MACS2 | peak | 1569610 | 1570248 | 638 | - | macsReverse_peak_101 | 4.09832 | CD630_13530 | sense | - | Putative phosphomethylpyrimidine kinase | | | | | | | | |
| CD630 | MACS2 | peak | 1572873 | 1572936 | 63 | + | macsForward_peak_161 | 3.38083 | CD630_13560 | sense | - | Putative CBS domain-containing protein | | | | | | | | |
| CD630 | MACS2 | peak | 1574243 | 1574325 | 82 | - | macsReverse_peak_102 | 3.80715 | CD630_13580 | sense | - | Putative membrane protein | | | | | | | | |
| CD630 | MACS2 | peak | 1574871 | 1575009 | 138 | + | macsForward_peak_162 | 5.37322 | CD630_13590 | antisense | - | Putative phage protein | CD630_n00490 | sense | | Antisense CDS | | | | |
| CD630 | MACS2 | peak | 1575171 | 1575402 | 231 | - | macsReverse_peak_103 | 4.45481 | CD630_13600 | sense | - | Transcriptional regulator, Phage-type | | | | | | | | |
| CD630 | MACS2 | peak | 1576604 | 1576736 | 132 | + | macsForward_peak_163 | 4.77627 | CD630_13620 | sense | - | Conserved hypothetical protein | | | | | | | | |
| CD630 | MACS2 | peak | 1594790 | 1595038 | 248 | - | macsReverse_peak_104 | 3.52274 | CD630_13780 | sense | - | Transcriptional regulator, Phage-type | | | | | | | | |
| CD630 | MACS2 | peak | 1596049 | 1596322 | 273 | - | macsReverse_peak_105 | 5.17374 | CD630_13790 | sense | - | Transcriptional regulator, Phage-type | | | | | | | | |
| CD630 | MACS2 | peak | 1607227 | 1607354 | 127 | - | macsReverse_peak_106 | 3.52735 | CD630_13880_p2 | sense | - | Fragment of trancriptional regulator (Part 2) | | | | | | | | |
| CD630 | MACS2 | peak | 1612357 | 1612586 | 229 | - | macsReverse_peak_107 | 3.25236 | CD630_13920 | sense | - | Putative endoribonuclease | | | | | | | | |
| CD630 | MACS2 | peak | 1613167 | 1613748 | 581 | + | macsForward_peak_164 | 4.09246 | CD630_13930 | sense | - | Putative membrane protein | | | | | | | | |
| CD630 | MACS2 | peak | 1615483 | 1615817 | 334 | - | macsReverse_peak_108 | 4.25540 | CD630_13950 | sense | - | Putative membrane protein | | | | | | | | |
| CD630 | MACS2 | peak | 1616151 | 1616408 | 257 | - | macsReverse_peak_109 | 3.30375 | CD630_13960 | sense | - | Putative amino acid amidase | | | | | | | | |
| CD630 | MACS2 | peak | 1621386 | 1621520 | 134 | - | macsReverse_peak_110 | 3.69983 | CD630_14010 | sense | - | Putative isochorismatase | | | | | | | | |
| CD630 | MACS2 | peak | 1628353 | 1628882 | 529 | - | macsReverse_peak_111 | 3.30894 | CD630_14050 | sense | - | ABC-type transport system, ATP-binding protein | | | | | | | | |
| CD630 | MACS2 | peak | 1635215 | 1635268 | 53 | - | macsReverse_peak_112 | 4.24790 | CD630_14100 | antisense | - | Putative hydrolase, HAD superfamily, subfamily IIB | | | | | | | | |
| CD630 | MACS2 | peak | 1635696 | 1635872 | 176 | - | macsReverse_peak_113 | 3.39550 | CD630_14110 | sense | - | Conserved hypothetical protein | | | | | | | | |
| CD630 | MACS2 | peak | 1638066 | 1638911 | 845 | + | macsForward_peak_165 | 4.57970 | CD630_14130 | sense | - | Putative membrane protein | | | | | | | | |
| CD630 | MACS2 | peak | 1639282 | 1639377 | 95 | - | macsReverse_peak_114 | 3.16551 | CD630_14140 | sense | - | Putative oxidoreductase, Fe-S subunit | | | | | | | | |
| CD630 | MACS2 | peak | 1640356 | 1640542 | 186 | - | macsReverse_peak_115 | 3.28251 | CD630_14150 | sense | - | Putative membrane protein | | | | | | | | |
| CD630 | MACS2 | peak | 1642388 | 1642652 | 264 | - | macsReverse_peak_116 | 3.93696 | CD630_14180 | sense | - | Putative drug/sodium antiporter, MATE family | | | | | | | | |
| CD630 | MACS2 | peak | 1644692 | 1645084 | 392 | - | macsReverse_peak_117 | 9.35359 | CD630_14182 | antisense | - | Putative phage protein | CD630_n00500 | sense | - | Antisense CDS | CD630_n00510 | antisense | - | ncRNA IGR |
| CD630 | MACS2 | peak | 1644735 | 1644859 | 124 | + | macsForward_peak_166 | 4.82360 | CD630_14182 | sense | - | Putative phage protein | CD630_n00500 | antisense | - | Antisense CDS | | | | |
| CD630 | MACS2 | peak | 1645203 | 1645262 | 59 | + | macsForward_peak_167 | 3.92762 | CD630_n00510 | sense | - | ncRNA IGR | | | | | | | | |
| CD630 | MACS2 | peak | 1646251 | 1646386 | 135 | - | macsReverse_peak_118 | 3.48168 | CD630_14190 | sense | - | Putative diguanylate kinase signaling protein | | | | | | | | |
| CD630 | MACS2 | peak | 1646699 | 1646900 | 201 | + | macsForward_peak_168 | 3.27218 | CD630_14190 | antisense | - | Putative diguanylate kinase signaling protein | | | | | | | | |
| CD630 | MACS2 | peak | 1647142 | 1647300 | 158 | - | macsReverse_peak_119 | 3.78562 | CD630_14190 | sense | - | Putative diguanylate kinase signaling protein | | | | | | | | |
| CD630 | MACS2 | peak | 1650584 | 1650704 | 120 | - | macsReverse_peak_120 | 3.82355 | CD630_14210 | sense | - | Putative diguanylate kinase signaling protein | | | | | | | | |
| CD630 | MACS2 | peak | 1653574 | 1653887 | 313 | - | macsReverse_peak_121 | 8.60766 | CD630_14240 | sense | - | Conserved hypothetical protein | CD630_Cdi1_10 | sense | - | GEMM RNA motif | | | | |
| CD630 | MACS2 | peak | 1654244 | 1654841 | 597 | - | macsReverse_peak_122 | 4.19604 | CD630_14250 | sense | - | Putative magnesium transporter | | | | | | | | |
| CD630 | MACS2 | peak | 1655183 | 1655311 | 128 | - | macsReverse_peak_123 | 3.22214 | CD630_14260_p2 | sense | - | Fragment of isochorismatase (Part 2) | | | | | | | | |
| CD630 | MACS2 | peak | 1664981 | 1665188 | 207 | + | macsForward_peak_169 | 5.10868 | CD630_14340 | sense | - | Conserved hypothetical protein | | | | | | | | |
| CD630 | MACS2 | peak | 1666809 | 1666975 | 166 | + | macsForward_peak_170 | 3.02079 | CD630_14360 | sense | - | Putative hydrolase | | | | | | | | |
| CD630 | MACS2 | peak | 1672749 | 1673086 | 337 | + | macsForward_peak_171 | 3.39433 | CD630_14410 | sense | - | Putative membrane protein | | | | | | | | |
| CD630 | MACS2 | peak | 1674075 | 1674135 | 60 | + | macsForward_peak_172 | 3.99497 | CD630_14420 | sense | - | Putative MORN repeat-containing protein | | | | | | | | |
| CD630 | MACS2 | peak | 1689357 | 1689714 | 357 | - | macsReverse_peak_124 | 5.90348 | CD630_14600 | sense | - | Putative phosphohydrolase, HD superfamily | | | | | | | | |
| CD630 | MACS2 | peak | 1690082 | 1690184 | 102 | - | macsReverse_peak_125 | 4.89428 | CD630_14610 | sense | - | Putative membrane protein | | | | | | | | |
| CD630 | MACS2 | peak | 1690989 | 1691339 | 350 | - | macsReverse_peak_126 | 4.43651 | CD630_14610 | sense | - | Putative membrane protein | CD630_14620 | sense | - | Transcriptional regulator, GntR family | | | | |
| CD630 | MACS2 | peak | 1691076 | 1691349 | 273 | + | macsForward_peak_173 | 7.81927 | CD630_14620 | antisense | - | Transcriptional regulator, GntR family | | | | | | | | |
| CD630 | MACS2 | peak | 1691851 | 1692068 | 217 | + | macsForward_peak_174 | 3.34485 | CD630_14630 | sense | - | Conserved hypothetical protein | | | | | | | | |
| CD630 | MACS2 | peak | 1702719 | 1702877 | 158 | + | macsForward_peak_175 | 3.98234 | CD630_14700 | sense | - | Putative rhodanese-like domain-containing protein | | | | | | | | |
| CD630 | MACS2 | peak | 1703212 | 1703897 | 685 | - | macsReverse_peak_127 | 3.65171 | CD630_14710 | sense | - | Putative membrane protein | | | | | | | | |
| CD630 | MACS2 | peak | 1717465 | 1717598 | 133 | + | macsForward_peak_176 | 3.27797 | CD630_14810 | sense | - | Putative tryptophan RNA-binding attenuation protein | | | | | | | | |
| CD630 | MACS2 | peak | 1717915 | 1717995 | 80 | + | macsForward_peak_177 | 3.35323 | CD630_14810 | sense | - | Putative tryptophan RNA-binding attenuation protein | | | | | | | | |
| CD630 | MACS2 | peak | 1718271 | 1718456 | 185 | + | macsForward_peak_178 | 3.45491 | CD630_14820 | sense | ssuC | ABC-type transport system, sulfonates-family permease | | | | | | | | |
| CD630 | MACS2 | peak | 1724138 | 1724253 | 115 | + | macsForward_peak_179 | 3.74616 | CD630_s0250 | sense | - | SAM, CD630_14890 | | | | | | | | |
| CD630 | MACS2 | peak | 1724145 | 1724204 | 59 | - | macsReverse_peak_128 | 4.90005 | CD630_s0250 | antisense | - | SAM, CD630_14890 | | | | | | | | |
| CD630 | MACS2 | peak | 1726654 | 1726750 | 96 | + | macsForward_peak_180 | 3.87190 | CD630_14910 | sense | metQ | ABC-type transport system, methionine-specific substrate-binding lipoprotein | | | | | | | | |
| CD630 | MACS2 | peak | 1727065 | 1727137 | 72 | + | macsForward_peak_181 | 4.35225 | CD630_14920 | sense | - | Histidine kinase, negative regulator of sporulation | | | | | | | | |
| CD630 | MACS2 | peak | 1730069 | 1730288 | 219 | + | macsForward_peak_182 | 3.31335 | CD630_14930 | sense | - | Putative 3-methyladenine DNA glycosylase | | | | | | | | |
| CD630 | MACS2 | peak | 1732484 | 1732537 | 53 | - | macsReverse_peak_129 | 3.42280 | CD630_14950 | antisense | proC | Pyrroline-5-carboxylate reductase | | | | | | | | |
| CD630 | MACS2 | peak | 1739001 | 1739435 | 434 | + | macsForward_peak_183 | 3.78619 | CD630_14990_p2 | sense | - | Putative virulence factor, MviN family protein (Disrupted IStron) | | | | | | | | |
| CD630 | MACS2 | peak | 1742883 | 1743363 | 480 | - | macsReverse_peak_130 | 3.69396 | CD630_15060 | sense | - | Putative drug/sodium antiporter, MATE family | | | | | | | | |
| CD630 | MACS2 | peak | 1745331 | 1745858 | 527 | + | macsForward_peak_184 | 4.03033 | CD630_15080 | sense | - | Putative iron-sulfur binding protein | | | | | | | | |
| CD630 | MACS2 | peak | 1749652 | 1749858 | 206 | - | macsReverse_peak_131 | 4.18664 | CD630_SQ995 | antisense | - | ncRNA IGR | CD630_15111 | sense | - | Conserved hypothetical protein | | | | |
| CD630 | MACS2 | peak | 1751362 | 1751457 | 95 | - | macsReverse_peak_132 | 3.20561 | CD630_15130 | sense | panB | Ketopantoate hydroxymethyltransferase | | | | | | | | |
| CD630 | MACS2 | peak | 1751673 | 1751843 | 170 | - | macsReverse_peak_133 | 3.40248 | CD630_15130 | sense | panB | Ketopantoate hydroxymethyltransferase | | | | | | | | |
| CD630 | MACS2 | peak | 1752748 | 1752932 | 184 | - | macsReverse_peak_134 | 5.54047 | CD630_15140 | sense | panG | Putative ketopantoate reductase | | | | | | | | |
| CD630 | MACS2 | peak | 1754138 | 1754312 | 174 | + | macsForward_peak_185 | 3.09352 | CD630_15150 | sense | - | Putative diguanylate kinase signaling protein | | | | | | | | |
| CD630 | MACS2 | peak | 1754540 | 1754732 | 192 | + | macsForward_peak_186 | 3.35564 | CD630_15150 | sense | - | Putative diguanylate kinase signaling protein | | | | | | | | |
| CD630 | MACS2 | peak | 1754949 | 1755053 | 104 | + | macsForward_peak_187 | 3.15951 | CD630_15150 | sense | - | Putative diguanylate kinase signaling protein | | | | | | | | |
| CD630 | MACS2 | peak | 1756140 | 1756276 | 136 | + | macsForward_peak_188 | 5.82053 | CD630_n00560 | sense | - | ncRNA IGR | | | | | | | | |
| CD630 | MACS2 | peak | 1756616 | 1756781 | 165 | + | macsForward_peak_189 | 4.55765 | CD630_n00560 | sense | - | ncRNA IGR | | | | | | | | |
| CD630 | MACS2 | peak | 1757497 | 1757549 | 52 | + | macsForward_peak_190 | 4.32318 | CD630_15160 | sense | - | Transcriptional regulator, MerR family | | | | | | | | |
| CD630 | MACS2 | peak | 1758507 | 1758896 | 389 | - | macsReverse_peak_135 | 4.05364 | CD630_15170 | sense | feoB | Ferrous iron transport protein B | | | | | | | | |
| CD630 | MACS2 | peak | 1759260 | 1759317 | 57 | + | macsForward_peak_191 | 3.61948 | CD630_15170 | antisense | feoB | Ferrous iron transport protein B | | | | | | | | |
| CD630 | MACS2 | peak | 1760929 | 1761290 | 361 | - | macsReverse_peak_136 | 4.43255 | CD630_SQ1002 | sense | - | ncRNA IGR | CD630_15190 | sense | - | Putative acyltransferase | | | | |
| CD630 | MACS2 | peak | 1761817 | 1762158 | 341 | + | macsForward_peak_192 | 4.00442 | CD630_15200 | sense | - | Putative membrane protein | | | | | | | | |
| CD630 | MACS2 | peak | 1762973 | 1763089 | 116 | + | macsForward_peak_193 | 3.63582 | CD630_SQ1005 | sense | - | ncRNA IGR | CD630_s0270 | sense | - | - | | | | |
| CD630 | MACS2 | peak | 1770233 | 1770580 | 347 | - | macsReverse_peak_137 | 4.66015 | CD630_15270 | sense | - | ABC-type transport system, permease | | | | | | | | |
| CD630 | MACS2 | peak | 1781500 | 1781692 | 192 | + | macsForward_peak_194 | 4.47208 | CD630_15360 | sense | - | Ferredoxin--NADP(+) reductase subunit alpha | | | | | | | | |
| CD630 | MACS2 | peak | 1793239 | 1793452 | 213 | + | macsForward_peak_195 | 5.29780 | CD630_15431 | sense | - | Conserved hypothetical protein | | | | | | | | |
| CD630 | MACS2 | peak | 1794268 | 1794904 | 636 | + | macsForward_peak_196 | 6.86296 | CD630_15460 | sense | - | Putative channel-forming hemolysin | | | | | | | | |
| CD630 | MACS2 | peak | 1795029 | 1795236 | 207 | - | macsReverse_peak_138 | 9.13442 | CD630_15461 | sense | - | Conserved hypothetical protein | | | | | | | | |
| CD630 | MACS2 | peak | 1795565 | 1795712 | 147 | + | macsForward_peak_197 | 8.30299 | CD630_s0281 | sense | - | CD630_15470 | | | | | | | | |
| CD630 | MACS2 | peak | 1803519 | 1803964 | 445 | + | macsForward_peak_198 | 3.49019 | CD630_15550 | sense | - | Putative amino acid permease | CD630_SQ1038 | sense | - | ncRNA IGR | | | | |
| CD630 | MACS2 | peak | 1810598 | 1810788 | 190 | + | macsForward_peak_199 | 5.37459 | CD630_s0280 | sense | - | - | | | | | | | | |
| CD630 | MACS2 | peak | 1823362 | 1823565 | 203 | - | macsReverse_peak_139 | 4.43222 | CD630_15740 | sense | - | Conserved hypothetical protein | | | | | | | | |
| CD630 | MACS2 | peak | 1827260 | 1827389 | 129 | + | macsForward_peak_200 | 3.45439 | CD630_15770 | sense | pgm | Alpha-phosphoglucomutase | | | | | | | | |
| CD630 | MACS2 | peak | 1830961 | 1831048 | 87 | - | macsReverse_peak_140 | 4.00174 | CD630_s0300 | antisense | - | T-box T-box | | | | | | | | |
| CD630 | MACS2 | peak | 1831121 | 1831439 | 318 | + | macsForward_peak_201 | 4.17452 | CD630_s0300 | sense | - | T-box T-box | CD630_s0310 | sense | - | - | | | | |

| | | | | | | | | | | | | | | | | | | | | |
|---|---|---|---|---|---|---|---|---|---|---|---|---|---|---|---|---|---|---|---|---|
| CD630 | MACS2 | peak | 1838227 | 1838291 | 64 | + | macsForward_peak_202 | 4.32968 | CD630_15850 | sense | - | Putative N-acetylglucosamine kinase NAGK-like | | | | | | | | |
| CD630 | MACS2 | peak | 1843247 | 1843438 | 191 | + | macsForward_peak_203 | 3.39962 | CD630_15890 | sense | - | ABC-type transport system, sugar-family extracellular solute-binding protein | | | | | | | | |
| CD630 | MACS2 | peak | 1843962 | 1844120 | 158 | + | macsForward_peak_204 | 3.17664 | CD630_15900 | sense | - | Putative membrane protein | | | | | | | | |
| CD630 | MACS2 | peak | 1851792 | 1851906 | 114 | + | macsForward_peak_205 | 4.11854 | CD630_15960 | antisense | map2 | Methionine aminopeptidase Map2 (MAP) (Peptidase M) | SQ1076 | sense | | Antisense 3'UTR | | | | |
| CD630 | MACS2 | peak | 1854581 | 1854682 | 101 | + | macsForward_peak_206 | 8.15991 | CD630_s0320 | sense | - | - | | | | | | | | |
| CD630 | MACS2 | peak | 1869500 | 1869802 | 302 | + | macsForward_peak_207 | 4.20614 | CD630_16150 | sense | - | ABC-type transport system, permease | | | | | | | | |
| CD630 | MACS2 | peak | 1870236 | 1870499 | 263 | + | macsForward_peak_208 | 3.61994 | CD630_16150 | sense | - | ABC-type transport system, permease | | | | | | | | |
| CD630 | MACS2 | peak | 1872433 | 1872625 | 192 | + | macsForward_peak_209 | 4.30055 | CD630_16160 | sense | - | Putative diguanylate kinase signaling protein | | | | | | | | |
| CD630 | MACS2 | peak | 1873098 | 1873286 | 188 | + | macsForward_peak_210 | 6.11801 | CD630_16160 | sense | - | Putative diguanylate kinase signaling protein | | | | | | | | |
| CD630 | MACS2 | peak | 1892637 | 1892935 | 298 | - | macsReverse_peak_141 | 4.11180 | CD630_16330 | sense | - | Putative serine protease | | | | | | | | |
| CD630 | MACS2 | peak | 1907159 | 1908467 | 1308 | + | macsForward_peak_211 | 4.57547 | CD630_16450 | sense | - | Putative membrane protein | CD630_16460 | sense | - | Putative peptidase, M19 family | | | | |
| CD630 | MACS2 | peak | 1908696 | 1909227 | 531 | + | macsForward_peak_212 | 4.64028 | CD630_16460 | sense | - | Putative peptidase, M19 family | | | | | | | | |
| CD630 | MACS2 | peak | 1915892 | 1916440 | 548 | - | macsReverse_peak_142 | 4.28395 | CD630_n00590 | sense | - | ncRNA IGR | CD630_16520 | sense | - | Putative tellurium resistance protein | | | | |
| CD630 | MACS2 | peak | 1916697 | 1918516 | 1819 | - | macsReverse_peak_143 | 4.37658 | CD630_s0330 | sense | - | SAM | CD630_16540 | sense | - | Putative lipoate-protein ligase | CD630_16530 | sense | - | Putative Probable D-methionine-binding lipoprotein |
| CD630 | MACS2 | peak | 1918864 | 1919290 | 426 | + | macsForward_peak_213 | 5.61064 | CD630_s0340 | sense | - | Lysine | CD630_16550 | sense | - | Putative Na+/H+ antiporter NhaC-like | | | | |
| CD630 | MACS2 | peak | 1918882 | 1918942 | 60 | - | macsReverse_peak_144 | 3.65576 | CD630_s0340 | antisense | - | Lysine | | | | | | | | |
| CD630 | MACS2 | peak | 1919635 | 1919797 | 162 | + | macsForward_peak_214 | 3.37787 | CD630_16550 | sense | - | Putative Na+/H+ antiporter NhaC-like | | | | | | | | |
| CD630 | MACS2 | peak | 1920034 | 1920870 | 836 | + | macsForward_peak_215 | 3.68720 | CD630_16550 | sense | - | Putative Na+/H+ antiporter NhaC-like | | | | | | | | |
| CD630 | MACS2 | peak | 1922433 | 1922959 | 526 | + | macsForward_peak_216 | 4.77229 | CD630_16570 | sense | gcvTPA | Bi-functional glycine dehydrogenase/aminomethyl transferase protein | | | | | | | | |
| CD630 | MACS2 | peak | 1923539 | 1923685 | 146 | + | macsForward_peak_217 | 3.83824 | CD630_16570 | sense | gcvTPA | Bi-functional glycine dehydrogenase/aminomethyl transferase protein | | | | | | | | |
| CD630 | MACS2 | peak | 1924065 | 1924319 | 254 | + | macsForward_peak_218 | 3.71071 | CD630_16570 | sense | gcvTPA | Bi-functional glycine dehydrogenase/aminomethyl transferase protein | | | | | | | | |
| CD630 | MACS2 | peak | 1935455 | 1935856 | 401 | + | macsForward_peak_219 | 5.31146 | CD630_n00600 | sense | - | ncRNA IGR | | | | | | | | |
| CD630 | MACS2 | peak | 1937906 | 1938165 | 259 | - | macsReverse_peak_145 | 3.61548 | CD630_16640 | sense | - | Transcriptional regulator, RmlC-type | | | | | | | | |
| CD630 | MACS2 | peak | 1942050 | 1942363 | 313 | + | macsForward_peak_220 | 3.33553 | CD630_16670 | sense | - | Putative maltose O-acetyltransferase | | | | | | | | |
| CD630 | MACS2 | peak | 1943259 | 1943683 | 424 | + | macsForward_peak_221 | 5.82838 | CD630_n00620 | sense | - | ncRNA IGR | CD630_16671 | antisense | - | Conserved hypothetical protein | | | | |
| CD630 | MACS2 | peak | 1951268 | 1951616 | 348 | + | macsForward_peak_222 | 5.04051 | CD630_16750 | sense | - | Transcriptional regulator, hxlR family | | | | | | | | |
| CD630 | MACS2 | peak | 1951890 | 1952028 | 138 | + | macsForward_peak_223 | 4.50056 | CD630_16760 | antisense | pcp | Pyrrolidone-carboxylate peptidase | | | | | | | | |
| CD630 | MACS2 | peak | 1965675 | 1965971 | 296 | - | macsReverse_peak_146 | 4.14665 | CD630_16940 | sense | - | Conserved hypothetical protein | | | | | | | | |
| CD630 | MACS2 | peak | 1966006 | 1966491 | 485 | + | macsForward_peak_224 | 3.99069 | CD630_16940 | antisense | - | Conserved hypothetical protein | CD630_16950 | antisense | - | Putative symporter protein | | | | |
| CD630 | MACS2 | peak | 1966939 | 1967154 | 215 | - | macsReverse_peak_147 | 4.42569 | CD630_16950 | sense | - | Putative symporter protein | | | | | | | | |
| CD630 | MACS2 | peak | 1972996 | 1973347 | 351 | - | macsReverse_peak_148 | 6.65922 | CD630_17000 | sense | ribD | Riboflavin biosynthesis protein ribD (Includes: Diaminohydrox | CD630_s0360 | sense | - | FMN | | | | |
| CD630 | MACS2 | peak | 1973228 | 1973330 | 102 | + | macsForward_peak_225 | 7.40079 | CD630_s0360 | antisense | - | FMN | | | | | | | | |
| CD630 | MACS2 | peak | 1976200 | 1976300 | 100 | + | macsForward_peak_226 | 10.21495 | CD630_s0370 | sense | - | - | | | | | | | | |
| CD630 | MACS2 | peak | 1985177 | 1985313 | 136 | + | macsForward_peak_227 | 4.25703 | CD630_17090 | sense | mobA | Molybdopterin-guanine dinucleotide biosynthesis protein A | | | | | | | | |
| CD630 | MACS2 | peak | 1987651 | 1987860 | 209 | + | macsForward_peak_228 | 3.30501 | CD630_17130 | sense | moaA | Molybdenum cofactor biosynthesis protein MoaA | | | | | | | | |
| CD630 | MACS2 | peak | 1988069 | 1988177 | 108 | + | macsForward_peak_229 | 3.83799 | CD630_17130 | sense | moaA | Molybdenum cofactor biosynthesis protein MoaA | | | | | | | | |
| CD630 | MACS2 | peak | 1989782 | 1989920 | 138 | + | macsForward_peak_230 | 6.43124 | CD630_17151 | sense | - | Conserved hypothetical protein | | | | | | | | |
| CD630 | MACS2 | peak | 1990187 | 1990408 | 221 | + | macsForward_peak_231 | 3.50228 | CD630_17160 | sense | - | Putative permease | | | | | | | | |
| CD630 | MACS2 | peak | 1991852 | 1992020 | 168 | + | macsForward_peak_232 | 4.10230 | CD630_17170 | sense | - | Conserved hypothetical protein, DUF917 family | | | | | | | | |
| CD630 | MACS2 | peak | 2001406 | 2001504 | 98 | - | macsReverse_peak_149 | 5.91544 | CD630_17281 | sense | - | Conserved hypothetical protein | | | | | | | | |
| CD630 | MACS2 | peak | 2004382 | 2004541 | 159 | + | macsForward_peak_233 | 6.25799 | CD630_17292 | sense | - | Conserved hypothetical protein | | | | | | | | |
| CD630 | MACS2 | peak | 2009614 | 2009753 | 139 | - | macsReverse_peak_150 | 3.35334 | CD630_17350 | sense | - | Putative hydrolase | | | | | | | | |
| CD630 | MACS2 | peak | 2010245 | 2010334 | 89 | - | macsReverse_peak_151 | 3.90254 | CD630_17350 | sense | - | Putative hydrolase | CD630_17360 | sense | - | Transcriptional regulator, HTH-type | | | | |
| CD630 | MACS2 | peak | 2010649 | 2010736 | 87 | - | macsReverse_peak_152 | 5.66070 | CD630_17360 | sense | - | Transcriptional regulator, HTH-type | | | | | | | | |
| CD630 | MACS2 | peak | 2012571 | 2013020 | 449 | + | macsForward_peak_234 | 5.03207 | CD630_17380 | sense | - | Two-component sensor histidine kinase | | | | | | | | |
| CD630 | MACS2 | peak | 2030332 | 2030435 | 103 | - | macsReverse_peak_153 | 5.63247 | CD630_17510 | antisense | cwp13 | Cell surface protein | | | | | | | | |
| CD630 | MACS2 | peak | 2030774 | 2030985 | 211 | - | macsReverse_peak_154 | 9.12322 | CD630_17511 | sense | - | Conserved hypothetical protein | | | | | | | | |
| CD630 | MACS2 | peak | 2034785 | 2034912 | 127 | + | macsForward_peak_235 | 3.99720 | CD630_17560 | sense | def1 | Peptide deformylase 1 (PDF 1) (Polypeptide deformylase 1) | | | | | | | | |
| CD630 | MACS2 | peak | 2036635 | 2036777 | 142 | + | macsForward_peak_236 | 8.19045 | . | | . | NA | | | | | | | | |
| CD630 | MACS2 | peak | 2041212 | 2041303 | 91 | - | macsReverse_peak_155 | 4.52408 | CD630_17630 | sense | - | Putative membrane protein | | | | | | | | |
| CD630 | MACS2 | peak | 2052419 | 2052732 | 313 | + | macsForward_peak_237 | 4.37688 | CD630_17710 | sense | - | Putative monooxygenase | | | | | | | | |
| CD630 | MACS2 | peak | 2053159 | 2053331 | 172 | + | macsForward_peak_238 | 3.22910 | CD630_17720 | sense | - | Conserved hypothetical protein | | | | | | | | |
| CD630 | MACS2 | peak | 2055318 | 2055684 | 366 | + | macsForward_peak_239 | 4.73397 | CD630_17730 | sense | - | Conserved hypothetical protein | CD630_s0390 | sense | - | T-box | | | | |
| CD630 | MACS2 | peak | 2058367 | 2058572 | 205 | + | macsForward_peak_240 | 5.06490 | CD630_17770 | sense | - | Putative arsenate reductase | | | | | | | | |
| CD630 | MACS2 | peak | 2066674 | 2066953 | 279 | + | macsForward_peak_241 | 4.51086 | CD630_s0400 | sense | - | T-box (Arg) | | | | | | | | |
| CD630 | MACS2 | peak | 2068065 | 2068293 | 228 | + | macsForward_peak_242 | 4.80300 | CD630_17850 | sense | argG | Argininosuccinate synthase (Citrulline--aspartate ligase) | | | | | | | | |
| CD630 | MACS2 | peak | 2072464 | 2072568 | 104 | - | macsReverse_peak_156 | 3.52167 | CD630_17900 | sense | - | Putative exonuclease | | | | | | | | |
| CD630 | MACS2 | peak | 2076622 | 2077032 | 410 | - | macsReverse_peak_157 | 3.71404 | CD630_17960 | sense | - | Putative nitrite and sulfite reductase subunit | | | | | | | | |
| CD630 | MACS2 | peak | 2078129 | 2078270 | 141 | - | macsReverse_peak_158 | 3.35819 | CD630_17970 | sense | - | Coenzyme A disulfide reductase (CoA-disulfide reductase) (CoADR) | | | | | | | | |
| CD630 | MACS2 | peak | 2078993 | 2079261 | 268 | - | macsReverse_peak_159 | 3.68802 | CD630_17970 | sense | - | Coenzyme A disulfide reductase (CoA-disulfide reductase) (CoADR) | | | | | | | | |
| CD630 | MACS2 | peak | 2081387 | 2081570 | 183 | - | macsReverse_peak_160 | 3.33148 | CD630_18000 | sense | terD | Tellurium resistance protein | | | | | | | | |
| CD630 | MACS2 | peak | 2082137 | 2082833 | 696 | - | macsReverse_peak_161 | 4.21967 | CD630_18010 | sense | - | Putative membrane protein | | | | | | | | |
| CD630 | MACS2 | peak | 2085655 | 2085896 | 241 | + | macsForward_peak_243 | 4.46610 | CD630_18030 | sense | cwp23 | Putative cell wall-binding protein | | | | | | | | |
| CD630 | MACS2 | peak | 2086803 | 2087019 | 216 | + | macsForward_peak_244 | 3.49166 | CD630_18040 | sense | scrR | Transcriptional regulator, LacI family | | | | | | | | |
| CD630 | MACS2 | peak | 2092621 | 2092909 | 288 | + | macsForward_peak_245 | 3.97564 | CD630_18090_p1 | sense | - | Fragment of putative permease, MFS family (Part 1) | CD630_18090_p2 | sense | - | Fragment of putative permease, MFS family (Part 2) | | | | |
| CD630 | MACS2 | peak | 2093278 | 2093611 | 333 | + | macsForward_peak_246 | 4.23961 | CD630_18090_p2 | sense | - | Fragment of putative permease, MFS family (Part 2) | | | | | | | | |
| CD630 | MACS2 | peak | 2094046 | 2094409 | 363 | + | macsForward_peak_247 | 5.71658 | CD630_18100 | sense | - | Transcriptional regulator, beta-lactams repressor | | | | | | | | |
| CD630 | MACS2 | peak | 2095843 | 2095969 | 126 | + | macsForward_peak_248 | 3.99713 | CD630_18120 | sense | - | Fragment of putative two-component response regulator (N-terminal region) | | | | | | | | |
| CD630 | MACS2 | peak | 2106156 | 2106235 | 79 | - | macsReverse_peak_162 | 4.73561 | CD630_18200_p1 | antisense | ade | Adenine deaminase (Disrupted IStron) | | | | | | | | |
| CD630 | MACS2 | peak | 2106681 | 2106816 | 135 | + | macsForward_peak_249 | 3.16670 | CD630_18220 | sense | bcp | Putative thiol peroxidase | | | | | | | | |
| CD630 | MACS2 | peak | 2107489 | 2107592 | 103 | + | macsForward_peak_250 | 3.00184 | CD630_18230 | sense | - | Conserved hypothetical protein, UPF0246 family | | | | | | | | |
| CD630 | MACS2 | peak | 2118056 | 2118114 | 58 | + | macsForward_peak_251 | 3.75086 | CD630_18290 | sense | kdpD | Two-component sensor histidine kinase KdpD | | | | | | | | |
| CD630 | MACS2 | peak | 2118662 | 2118883 | 221 | + | macsForward_peak_252 | 4.27140 | CD630_18290 | sense | kdpD | Two-component sensor histidine kinase KdpD | CD630_18300 | sense | kdpE | Two-component response regulator KdpE | | | | |
| CD630 | MACS2 | peak | 2120981 | 2121174 | 193 | + | macsForward_peak_253 | 6.86047 | CD630_18311 | sense | - | Conserved hypothetical protein | | | | | | | | |
| CD630 | MACS2 | peak | 2130231 | 2130310 | 79 | + | macsForward_peak_254 | 4.67962 | CD630_18400 | sense | - | Putative diguanylate kinase signaling protein | | | | | | | | |
| CD630 | MACS2 | peak | 2131283 | 2131423 | 140 | + | macsForward_peak_255 | 3.59739 | CD630_18410 | sense | - | Two-component sensor histidine kinase | | | | | | | | |
| CD630 | MACS2 | peak | 2155533 | 2155610 | 77 | + | macsForward_peak_256 | 4.15582 | CD630_18610 | sense | - | Putative conjugative transposon protein Tn1549-like, CTn5-Orf19 | | | | | | | | |
| CD630 | MACS2 | peak | 2167105 | 2167352 | 247 | + | macsForward_peak_257 | 3.75892 | CD630_18650 | sense | - | Putative DNA recombination protein Tn1549-like, CTn5-Orf24 | | | | | | | | |
| CD630 | MACS2 | peak | 2167699 | 2168058 | 359 | + | macsForward_peak_258 | 3.50371 | CD630_18650 | sense | - | Putative DNA recombination protein Tn1549-like, CTn5-Orf24 | CD630_18660 | sense | - | Putative conjugative transposon protein tn1549-like, CTn5-Orf25 | | | | |
| CD630 | MACS2 | peak | 2169810 | 2169916 | 106 | + | macsForward_peak_259 | 3.04315 | CD630_18680 | sense | - | Putative conjugative transposon protein Tn1549-like, CTn5-Orf27 | | | | | | | | |
| CD630 | MACS2 | peak | 2171872 | 2171929 | 57 | - | macsReverse_peak_163 | 6.94246 | . | | . | NA | | | | | | | | |
| CD630 | MACS2 | peak | 2178061 | 2178218 | 157 | + | macsForward_peak_260 | 3.93642 | CD630_18760 | sense | vncR | Two-component response regulator, Tn1549-like, CTn5-Orf36 | | | | | | | | |
| CD630 | MACS2 | peak | 2180212 | 2180339 | 127 | + | macsForward_peak_261 | 4.71613 | CD630_18780 | sense | - | Putative sigma factor Tn1549-like, CTn5-Orf38 | | | | | | | | |
| CD630 | MACS2 | peak | 2182683 | 2182801 | 118 | + | macsForward_peak_262 | 3.77660 | CD630_18781 | sense | - | Fragment of conserved hypothetical protein (N-terminal region) | | | | | | | | |
| CD630 | MACS2 | peak | 2183308 | 2183620 | 312 | + | macsForward_peak_263 | 3.32076 | CD630_18440_p2 | sense | - | Fragment of conserved hypothetical protein (Part 2) | | | | | | | | |
| CD630 | MACS2 | peak | 2183936 | 2184047 | 111 | + | macsForward_peak_264 | 4.07960 | CD630_18440_p2 | sense | - | Fragment of conserved hypothetical protein (Part 2) | | | | | | | | |

| | | | | | | | | | | | | | | | | |
|---|---|---|---|---|---|---|---|---|---|---|---|---|---|---|---|---|
| CD630 | MACS2 | peak | 2187134 | 2187231 | 97 | - | macsReverse_peak_164 | 3.30688 | CD630_18910_p2 | sense | - | Fragment of ABC-type transport system, substrate-binding protein (Part 2) | | | | |
| CD630 | MACS2 | peak | 2197336 | 2197442 | 106 | + | macsForward_peak_265 | 3.53248 | CD630_18920 | sense | - | Putative methyltransferase, type 11 family | | | | |
| CD630 | MACS2 | peak | 2197726 | 2198013 | 287 | + | macsForward_peak_266 | 4.43558 | CD630_18920 | sense | - | Putative methyltransferase, type 11 family | | | | |
| CD630 | MACS2 | peak | 2198056 | 2198484 | 428 | - | macsReverse_peak_165 | 5.32127 | CD630_18930 | sense | - | Putative oligonucleotide binding regulator | CD630_n00650 | sense | - | Antisense CDS |
| CD630 | MACS2 | peak | 2198990 | 2199467 | 477 | + | macsForward_peak_267 | 4.09988 | CD630_18940 | sense | - | Conserved hypothetical protein | Rcd1 | sense | | ncRNA IGR |
| CD630 | MACS2 | peak | 2199393 | 2199457 | 64 | - | macsReverse_peak_166 | 6.83517 | Rcd1 | antisense | | ncRNA IGR | | | | |
| CD630 | MACS2 | peak | 2199888 | 2199957 | 69 | - | macsReverse_peak_167 | 5.67942 | CD630_18950 | antisense | - | Conserved hypothetical protein | | | | |
| CD630 | MACS2 | peak | 2200090 | 2200167 | 77 | + | macsForward_peak_268 | 3.40610 | CD630_18950 | sense | - | Conserved hypothetical protein | | | | |
| CD630 | MACS2 | peak | 2200437 | 2200553 | 116 | + | macsForward_peak_269 | 4.37822 | CD630_18960 | sense | - | Conserved hypothetical protein | | | | |
| CD630 | MACS2 | peak | 2200647 | 2200958 | 311 | - | macsReverse_peak_168 | 3.57243 | CD630_18970 | sense | - | Conserved hypothetical protein | | | | |
| CD630 | MACS2 | peak | 2202760 | 2202885 | 125 | - | macsReverse_peak_169 | 4.18443 | CD630_19000 | sense | - | Conserved hypothetical protein | | | | |
| CD630 | MACS2 | peak | 2204565 | 2204664 | 99 | + | macsForward_peak_270 | 3.34240 | CD630_19030 | sense | - | Putative lipoprotein signal peptidase | | | | |
| CD630 | MACS2 | peak | 2210287 | 2210565 | 278 | + | macsForward_peak_271 | 3.82847 | CD630_19090 | sense | eutP | Ethanolamine utilization protein, GTPase family | | | | |
| CD630 | MACS2 | peak | 2211267 | 2211352 | 85 | + | macsForward_peak_272 | 4.55715 | CD630_19100 | sense | eutV | Two-component response regulator, Ethanolamine specific | CD630_19110 | sense | eutW | Two-component sensor histidine kinase, Ethanolamine specific |
| CD630 | MACS2 | peak | 2238411 | 2238542 | 131 | + | macsForward_peak_273 | 4.37084 | CD630_19401 | sense | - | Putative membrane protein | | | | |
| CD630 | MACS2 | peak | 2240648 | 2240718 | 70 | - | macsReverse_peak_170 | 3.13922 | CD630_19430 | sense | - | Conserved hypothetical protein | | | | |
| CD630 | MACS2 | peak | 2241125 | 2241339 | 214 | - | macsReverse_peak_171 | 4.53937 | CD630_19440 | sense | - | Fragment of conserved hypothetical protein (C-terminal region) | | | | |
| CD630 | MACS2 | peak | 2257786 | 2258102 | 316 | + | macsForward_peak_274 | 4.51902 | CD630_SQ1339 | antisense | - | ncRNA IGR | CD630_19580 | sense | - | Putative 4Fe-4S binding protein, putative ferredoxin, putative dehydrogenase, putative reductase |
| CD630 | MACS2 | peak | 2258327 | 2258764 | 437 | + | macsForward_peak_275 | 4.03154 | CD630_19580 | sense | - | Putative 4Fe-4S binding protein, putative ferredoxin, putative dehydrogenase, putative reductase | | | | |
| CD630 | MACS2 | peak | 2268872 | 2268974 | 102 | - | macsReverse_peak_172 | 3.20718 | CD630_19670 | sense | - | Conserved hypothetical protein | | | | |
| CD630 | MACS2 | peak | 2269965 | 2270269 | 304 | - | macsReverse_peak_173 | 3.58080 | CD630_19680 | sense | - | Putative membrane protein | | | | |
| CD630 | MACS2 | peak | 2270971 | 2271788 | 817 | - | macsReverse_peak_174 | 3.85861 | CD630_19690 | sense | - | Putative oligonucleotide binding regulator | CD630_19700 | sense | - | Putative enzyme, cyclase family |
| CD630 | MACS2 | peak | 2275965 | 2276215 | 250 | - | macsReverse_peak_175 | 5.05609 | CD630_19740 | sense | hfq | Hfq RNA chaperone | | | | |
| CD630 | MACS2 | peak | 2282546 | 2283388 | 842 | - | macsReverse_peak_176 | 3.64830 | CD630_19780 | sense | - | ABC-type transport system, ATP-binding protein | CD630_19790 | sense | - | ABC-type transport system, extracellular solute-binding protein |
| CD630 | MACS2 | peak | 2283769 | 2283895 | 126 | - | macsReverse_peak_177 | 3.79593 | CD630_19790 | sense | - | ABC-type transport system, extracellular solute-binding protein | | | | |
| CD630 | MACS2 | peak | 2284468 | 2284990 | 522 | - | macsReverse_peak_178 | 5.22035 | CD630_19800 | sense | - | ABC-type transport system, permease | CD630_19801 | sense | - | Conserved hypothetical protein, DUF77 family |
| CD630 | MACS2 | peak | 2286198 | 2286268 | 70 | + | macsForward_peak_276 | 6.52781 | CD630_n00680 | sense | - | ncRNA IGR | Rcd5 | sense | | |
| CD630 | MACS2 | peak | 2286519 | 2286607 | 88 | + | macsForward_peak_277 | 3.99438 | CD630_19810 | antisense | - | Transcriptional regulator, hxlR family | | | | |
| CD630 | MACS2 | peak | 2288265 | 2288399 | 134 | + | macsForward_peak_278 | 3.44724 | CD630_19840 | sense | - | Putative hydrolase, NUDIX family | | | | |
| CD630 | MACS2 | peak | 2289082 | 2289330 | 248 | + | macsForward_peak_279 | 6.88486 | CD630_19850 | sense | - | Putative membrane protein | | | | |
| CD630 | MACS2 | peak | 2297527 | 2297813 | 286 | + | macsForward_peak_280 | 5.79994 | CD630_Cdi1_8 | sense | - | GEMM RNA motif | CD630_19903 | sense | - | Conserved hypothetical protein |
| CD630 | MACS2 | peak | 2298058 | 2298246 | 188 | + | macsForward_peak_281 | 5.48848 | CD630_19902 | sense | cspC | Major cold shock protein CspC | | | | |
| CD630 | MACS2 | peak | 2298419 | 2298860 | 441 | - | macsReverse_peak_179 | 5.96526 | CD630_19904 | sense | - | Fragment of conserved hypothetical protein | CD630_n00690 | sense | - | ncRNA IGR |
| CD630 | MACS2 | peak | 2300303 | 2300538 | 235 | - | macsReverse_peak_180 | 5.46345 | CD630_19921 | sense | - | Conserved hypothetical protein | | | | |
| CD630 | MACS2 | peak | 2303170 | 2303236 | 66 | - | macsReverse_peak_181 | 4.24574 | CD630_19942 | sense | - | Conserved hypothetical protein | | | | |
| CD630 | MACS2 | peak | 2327638 | 2327750 | 112 | - | macsReverse_peak_182 | 4.53652 | CD630_20170 | antisense | - | Transcriptional regulator, HTH-type | | | | |
| CD630 | MACS2 | peak | 2327708 | 2327970 | 262 | + | macsForward_peak_282 | 6.90201 | CD630_20170 | sense | - | Transcriptional regulator, HTH-type | | | | |
| CD630 | MACS2 | peak | 2327984 | 2328190 | 206 | - | macsReverse_peak_183 | 6.87491 | CD630_20170 | antisense | - | Transcriptional regulator, HTH-type | CD630_20171 | sense | - | Conserved hypothetical protein |
| CD630 | MACS2 | peak | 2337894 | 2337979 | 85 | + | macsForward_peak_283 | 3.84162 | CD630_20260 | sense | - | Putative acetyltransferase | | | | |
| CD630 | MACS2 | peak | 2339976 | 2340191 | 215 | + | macsForward_peak_284 | 3.46343 | CD630_20280 | sense | - | Putative aspartate racemase RacX | | | | |
| CD630 | MACS2 | peak | 2340724 | 2340805 | 81 | - | macsReverse_peak_184 | 4.38183 | CD630_20290 | sense | - | Putative lipoprotein | CD630_n00700 | antisense | - | Antisense CDS |
| CD630 | MACS2 | peak | 2347097 | 2347397 | 300 | - | macsReverse_peak_185 | 5.67265 | CD630_s0450 | sense | - | T-box (Arg) | | | | |
| CD630 | MACS2 | peak | 2355898 | 2356092 | 194 | - | macsReverse_peak_186 | 3.23150 | CD630_20430 | antisense | - | Putative hydrolase | | | | |
| CD630 | MACS2 | peak | 2357780 | 2357980 | 200 | + | macsForward_peak_285 | 3.77388 | CD630_20440 | sense | - | Putative membrane protein | CD630_20450 | sense | - | Transcriptional regulator, HTH-type |
| CD630 | MACS2 | peak | 2358726 | 2359016 | 290 | - | macsReverse_peak_187 | 7.94034 | CD630_20460 | sense | - | Conserved hypothetical protein | | | | |
| CD630 | MACS2 | peak | 2368910 | 2369125 | 215 | - | macsReverse_peak_188 | 5.74527 | CD630_20541 | sense | - | Conserved hypothetical protein | CD630_s0460 | sense | - | Lysine |
| CD630 | MACS2 | peak | 2375625 | 2376378 | 753 | - | macsReverse_peak_189 | 4.20093 | CD630_20600 | sense | - | Putative lipid kinase | | | | |
| CD630 | MACS2 | peak | 2380292 | 2380601 | 309 | - | macsReverse_peak_190 | 4.00083 | CD630_20650 | sense | - | Transcriptional regulator, beta-lactams repressor | | | | |
| CD630 | MACS2 | peak | 2381020 | 2381125 | 105 | + | macsForward_peak_286 | 5.39321 | CD630_20660 | sense | - | Putative integrase | | | | |
| CD630 | MACS2 | peak | 2382301 | 2382556 | 255 | - | macsReverse_peak_191 | 4.42757 | CD630_20670 | sense | - | Conserved hypothetical protein | | | | |
| CD630 | MACS2 | peak | 2416363 | 2416459 | 96 | - | macsReverse_peak_192 | 4.13812 | CD630_20920 | sense | - | Transcriptional regulator, sigma-54-dependent | | | | |
| CD630 | MACS2 | peak | 2421849 | 2421910 | 61 | - | macsReverse_peak_193 | 5.65940 | CD630_20960 | sense | - | Putative DNA-binding protein | | | | |
| CD630 | MACS2 | peak | 2423513 | 2423716 | 203 | - | macsReverse_peak_194 | 8.12847 | CD630_20980 | sense | - | Conserved hypothetical protein | | | | |
| CD630 | MACS2 | peak | 2425491 | 2425835 | 344 | - | macsReverse_peak_195 | 4.45147 | CD630_20990 | sense | XdhD | Xanthine dehydrogenase subunit XdhD, molybdopterin-binding | | | | |
| CD630 | MACS2 | peak | 2427667 | 2428135 | 468 | - | macsReverse_peak_196 | 4.22212 | CD630_21020 | sense | - | Putative Na(+)/H(+) antiporter | | | | |
| CD630 | MACS2 | peak | 2428362 | 2428894 | 532 | - | macsReverse_peak_197 | 4.28666 | CD630_21020 | sense | - | Putative Na(+)/H(+) antiporter | | | | |
| CD630 | MACS2 | peak | 2430149 | 2430358 | 209 | - | macsReverse_peak_198 | 3.08733 | CD630_21030 | sense | - | ABC-type transport system, permease | | | | |
| CD630 | MACS2 | peak | 2434842 | 2434893 | 51 | - | macsReverse_peak_199 | 3.17748 | CD630_21070 | sense | - | Xanthine/uracil/thiamine/ascorbate permease family protein | | | | |
| CD630 | MACS2 | peak | 2435817 | 2436149 | 332 | - | macsReverse_peak_200 | 5.35019 | CD630_21070 | sense | - | Xanthine/uracil/thiamine/ascorbate permease family protein | CD630_s0470 | sense | - | Purine |
| CD630 | MACS2 | peak | 2436549 | 2437735 | 1186 | + | macsForward_peak_287 | 3.99742 | CD630_21080 | sense | - | Putative drug/sodium antiporter, MATE family | | | | |
| CD630 | MACS2 | peak | 2438858 | 2439250 | 392 | + | macsForward_peak_288 | 4.15803 | CD630_21090 | sense | - | Putative anion:Na+ symporter | | | | |
| CD630 | MACS2 | peak | 2440494 | 2440588 | 94 | + | macsForward_peak_289 | 3.89630 | CD630_21110 | sense | - | Transcriptional regulator, DeoR family | | | | |
| CD630 | MACS2 | peak | 2443704 | 2443870 | 166 | - | macsReverse_peak_201 | 3.33304 | CD630_21140 | sense | - | Two-component response regulator | | | | |
| CD630 | MACS2 | peak | 2451389 | 2451525 | 136 | + | macsForward_peak_290 | 4.87999 | CD630_s0480 | sense | - | T-box (Thr) | | | | |
| CD630 | MACS2 | peak | 2458858 | 2458929 | 71 | - | macsReverse_peak_202 | 3.89213 | CD630_21230 | sense | - | Putative multiprotein-complex assembly TPR repeat-containing protein | | | | |
| CD630 | MACS2 | peak | 2459564 | 2459660 | 96 | + | macsForward_peak_291 | 5.88879 | CD630_21240 | sense | - | Putative malate transporter | | | | |
| CD630 | MACS2 | peak | 2459900 | 2460374 | 474 | + | macsForward_peak_292 | 5.62627 | CD630_21240 | sense | - | Putative malate transporter | | | | |
| CD630 | MACS2 | peak | 2462344 | 2462577 | 233 | + | macsForward_peak_293 | 3.11884 | CD630_21260 | sense | - | Putative membrane protein | CD630_n00740 | antisense | - | Antisense CDS |
| CD630 | MACS2 | peak | 2462822 | 2462923 | 101 | + | macsForward_peak_294 | 4.22255 | CD630_21260 | sense | - | Putative membrane protein | | | | |
| CD630 | MACS2 | peak | 2469041 | 2469129 | 88 | + | macsForward_peak_295 | 4.10111 | CD630_21330 | sense | - | Conserved hypothetical protein | | | | |
| CD630 | MACS2 | peak | 2483582 | 2483913 | 331 | - | macsReverse_peak_203 | 3.86658 | CD630_21460 | sense | - | Conserved hypothetical protein, UPF0027 family | | | | |
| CD630 | MACS2 | peak | 2484424 | 2484769 | 345 | - | macsReverse_peak_204 | 4.83005 | CD630_21460 | sense | - | Conserved hypothetical protein, UPF0027 family | | | | |
| CD630 | MACS2 | peak | 2487179 | 2487309 | 130 | - | macsReverse_peak_205 | 3.37462 | CD630_21490 | sense | - | Putative vancomycin resistance protein, vanW family | | | | |
| CD630 | MACS2 | peak | 2492001 | 2492466 | 465 | + | macsForward_peak_296 | 3.39774 | CD630_21530 | sense | - | Conserved hypothetical protein | CD630_21540 | sense | - | Putative ATP/GTP-binding protein |
| CD630 | MACS2 | peak | 2492828 | 2493352 | 524 | + | macsForward_peak_297 | 4.33090 | CD630_21540 | sense | - | Putative ATP/GTP-binding protein | | | | |
| CD630 | MACS2 | peak | 2493641 | 2493891 | 250 | + | macsForward_peak_298 | 3.59022 | CD630_21550 | sense | bioB | Biotin synthase | | | | |
| CD630 | MACS2 | peak | 2494304 | 2494463 | 159 | + | macsForward_peak_299 | 3.30736 | CD630_21550 | sense | bioB | Biotin synthase | | | | |
| CD630 | MACS2 | peak | 2495018 | 2495300 | 282 | + | macsForward_peak_300 | 3.58230 | CD630_21560 | sense | thiH | Thiamine biosynthesis protein ThiH | | | | |
| CD630 | MACS2 | peak | 2496120 | 2496797 | 677 | + | macsForward_peak_301 | 6.48385 | CD630_21570 | sense | - | Conserved hypothetical protein | CD630_21580 | antisense | gabT | 4-aminobutyrate aminotransferase |
| CD630 | MACS2 | peak | 2496640 | 2496962 | 322 | - | macsReverse_peak_206 | 3.67067 | CD630_21580 | sense | gabT | 4-aminobutyrate aminotransferase | | | | |
| CD630 | MACS2 | peak | 2500407 | 2500524 | 117 | - | macsReverse_peak_207 | 3.16662 | CD630_21610 | sense | - | Putative membrane protein | | | | |
| CD630 | MACS2 | peak | 2500865 | 2500959 | 94 | - | macsReverse_peak_208 | 3.02649 | CD630_21620 | sense | - | Putative GCN5-related N-acetyltransferase-like protein | | | | |
| CD630 | MACS2 | peak | 2503946 | 2504024 | 78 | + | macsForward_peak_302 | 4.35818 | CD630_21640 | sense | ldh | L-lactate dehydrogenase (L-LDH) | | | | |
| CD630 | MACS2 | peak | 2510580 | 2511023 | 443 | + | macsForward_peak_303 | 3.96650 | CD630_21700 | sense | - | Putative cNMP-binding regulatory protein | | | | |
| CD630 | MACS2 | peak | 2513112 | 2513383 | 271 | - | macsReverse_peak_209 | 4.13030 | CD630_21720 | sense | - | ABC-type transport system, cystine/aminoacid-family ATP-binding protein | | | | |
| CD630 | MACS2 | peak | 2514205 | 2514714 | 509 | - | macsReverse_peak_210 | 3.45076 | CD630_21730 | sense | - | Putative peptidase, M20D family | | | | |

| | | | | | | | | | | | | | | | | | | | | |
|---|---|---|---|---|---|---|---|---|---|---|---|---|---|---|---|---|---|---|---|---|
| CD630 | MACS2 | peak | 2516200 | 2516589 | 389 | - | macsReverse_peak_211 | 3.70626 | CD630_21750 | sense | - | ABC-type transport system, cystine/aminoacid-family permease | | | | | | | | |
| CD630 | MACS2 | peak | 2517052 | 2517403 | 351 | - | macsReverse_peak_212 | 4.43846 | CD630_21760 | sense | - | ABC-type transport system, cystine/aminoacid-family permease | | | | | | | | |
| CD630 | MACS2 | peak | 2518071 | 2518439 | 368 | - | macsReverse_peak_213 | 4.02587 | CD630_21770 | sense | - | ABC-type transport system, cystine/aminoacid-family extracellular solute-binding protein | | | | | | | | |
| CD630 | MACS2 | peak | 2520944 | 2521073 | 129 | - | macsReverse_peak_214 | 4.93762 | CD630_21790 | sense | - | Putative anaerobic dehydrogenase | | | | | | | | |
| CD630 | MACS2 | peak | 2524595 | 2524806 | 211 | - | macsReverse_peak_215 | 4.73824 | CD630_21800 | antisense | - | Putative DEAD-like helicase | | | | | | | | |
| CD630 | MACS2 | peak | 2537501 | 2538246 | 745 | - | macsReverse_peak_216 | 4.66600 | CD630_21910 | sense | - | Putative phosphoesterase | | | | | | | | |
| CD630 | MACS2 | peak | 2552699 | 2552815 | 116 | + | macsForward_peak_304 | 3.29934 | CD630_22030 | sense | - | Transcriptional regulator, TetR family | | | | | | | | |
| CD630 | MACS2 | peak | 2554301 | 2555194 | 893 | - | macsReverse_peak_217 | 4.87639 | CD630_22050 | sense | - | Putative membrane protein | | | | | | | | |
| CD630 | MACS2 | peak | 2557207 | 2557580 | 373 | - | macsReverse_peak_218 | 3.94743 | CD630_22070 | sense | - | Putative flavodoxin | | | | | | | | |
| CD630 | MACS2 | peak | 2557787 | 2557839 | 52 | - | macsReverse_peak_219 | 4.09651 | CD630_22070 | sense | - | Putative flavodoxin | | | | | | | | |
| CD630 | MACS2 | peak | 2565474 | 2565873 | 399 | - | macsReverse_peak_220 | 4.21370 | CD630_22130 | sense | cynT | Carbonic anhydrase | | | | | | | | |
| CD630 | MACS2 | peak | 2568629 | 2568813 | 184 | - | macsReverse_peak_221 | 3.71929 | CD630_22170 | sense | aroD | Dehydroquinate dehydratase (3-dehydroquinate dehydratase) (Dehydroquinase) (DHQase) | | | | | | | | |
| CD630 | MACS2 | peak | 2571815 | 2572043 | 228 | + | macsForward_peak_305 | 4.18288 | CD630_22200 | sense | - | Putative cyclic nucleotides-binding protein | | | | | | | | |
| CD630 | MACS2 | peak | 2607432 | 2607529 | 97 | + | macsForward_peak_306 | 4.03688 | CD630_22500 | sense | - | Putative membrane protein | | | | | | | | |
| CD630 | MACS2 | peak | 2607863 | 2608118 | 255 | + | macsForward_peak_307 | 3.10662 | CD630_22500 | sense | - | Putative membrane protein | | | | | | | | |
| CD630 | MACS2 | peak | 2608350 | 2608676 | 326 | + | macsForward_peak_308 | 11.42763 | CD630_22510 | sense | - | Conserved hypothetical protein | | | | | | | | |
| CD630 | MACS2 | peak | 2614157 | 2614292 | 135 | - | macsReverse_peak_222 | 3.62379 | CD630_22550 | sense | - | Conserved hypothetical protein | | | | | | | | |
| CD630 | MACS2 | peak | 2626199 | 2626397 | 198 | - | macsReverse_peak_223 | 3.60496 | CD630_22660 | sense | - | Putative oxidoreductase, FAD dependent | | | | | | | | |
| CD630 | MACS2 | peak | 2626612 | 2627117 | 505 | - | macsReverse_peak_224 | 3.44541 | CD630_22660 | sense | - | Putative oxidoreductase, FAD dependent | | | | | | | | |
| CD630 | MACS2 | peak | 2632743 | 2632867 | 124 | - | macsReverse_peak_225 | 4.05623 | CD630_22710 | sense | - | Transcription antiterminator, PTS operon regulator | | | | | | | | |
| CD630 | MACS2 | peak | 2634662 | 2634855 | 193 | - | macsReverse_peak_226 | 3.81804 | CD630_22720 | sense | - | Putative protein-tyrosine phosphatase | | | | | | | | |
| CD630 | MACS2 | peak | 2637581 | 2637660 | 79 | - | macsReverse_peak_227 | 3.08733 | CD630_22740 | sense | - | Conserved hypothetical protein | | | | | | | | |
| CD630 | MACS2 | peak | 2640444 | 2640515 | 71 | - | macsReverse_peak_228 | 4.28971 | CD630_RNA_17 | antisense | - | Glycine | CD630_RNA_15 | antisense | - | Glycine | | | | |
| CD630 | MACS2 | peak | 2640559 | 2640630 | 71 | + | macsForward_peak_309 | 4.12920 | CD630_RNA_15 | sense | - | Glycine | | | | | | | | |
| CD630 | MACS2 | peak | 2652793 | 2653277 | 484 | + | macsForward_peak_310 | 3.84102 | CD630_22850 | sense | - | Transcriptional regulator, GntR family | | | | | | | | |
| CD630 | MACS2 | peak | 2656632 | 2656814 | 182 | - | macsReverse_peak_229 | 3.62455 | CD630_22880 | sense | - | Two-component sensor histidine kinase | | | | | | | | |
| CD630 | MACS2 | peak | 2657095 | 2657200 | 105 | - | macsReverse_peak_230 | 3.75591 | CD630_22890 | sense | - | Two-component response regulator | | | | | | | | |
| CD630 | MACS2 | peak | 2660762 | 2661065 | 303 | + | macsForward_peak_311 | 3.96883 | CD630_22950 | sense | - | Putative membrane protein | | | | | | | | |
| CD630 | MACS2 | peak | 2661280 | 2661357 | 77 | + | macsForward_peak_312 | 3.58866 | CD630_SQ1633 | antisense | - | Antisense 3prime UTR | CD630_22951 | antisense | - | Fragment of putative cell surface protein (C-terminal region) | | | | |
| CD630 | MACS2 | peak | 2663798 | 2663903 | 105 | - | macsReverse_peak_231 | 5.51210 | CD630_n00790 | sense | - | ncRNA IGR | | | | | | | | |
| CD630 | MACS2 | peak | 2665226 | 2665461 | 235 | + | macsForward_peak_313 | 8.65138 | CD630_SQ1641 | sense | - | Antisense CDS | CD630_22991 | antisense | - | Putative phage protein | CD630_SQ1642 | sense | - | ncRNA IGR |
| CD630 | MACS2 | peak | 2665311 | 2665419 | 108 | - | macsReverse_peak_232 | 6.03728 | CD630_SQ1641 | antisense | - | Antisense CDS | CD630_22991 | sense | - | Putative phage protein | CD630_SQ1642 | antisense | - | ncRNA IGR |
| CD630 | MACS2 | peak | 2665838 | 2666034 | 196 | - | macsReverse_peak_233 | 3.54130 | CD630_23000 | sense | - | Conserved hypothetical protein | | | | | | | | |
| CD630 | MACS2 | peak | 2669309 | 2669626 | 317 | + | macsForward_peak_314 | 4.96316 | CD630_23050 | sense | pilW | Type IV pilus protein | | | | | | | | |
| CD630 | MACS2 | peak | 2671225 | 2671566 | 341 | + | macsForward_peak_315 | 5.23027 | CD630_23070 | sense | - | Putative membrane protein | CD630_23080 | sense | - | Transcriptional regulator, HTH-type | CD630_SQ1653 | antisense | - | Antisense 5prime UTR |
| CD630 | MACS2 | peak | 2671843 | 2672139 | 296 | + | macsForward_peak_316 | 4.25622 | CD630_SQ1656 | sense | - | ncRNA IGR | CD630_Cdi1_9 | sense | - | GEMM RNA motif | CD630_23090 | sense | - | Conserved hypothetical protein |
| CD630 | MACS2 | peak | 2671845 | 2672018 | 173 | - | macsReverse_peak_234 | 3.76346 | CD630_SQ1656 | antisense | - | ncRNA IGR | CD630_Cdi1_9 | antisense | - | GEMM RNA motif | CD630_23090 | antisense | - | Conserved hypothetical protein |
| CD630 | MACS2 | peak | 2672386 | 2672539 | 153 | + | macsForward_peak_317 | 3.23587 | CD630_23100 | sense | cspD | Cold shock protein D | | | | | | | | |
| CD630 | MACS2 | peak | 2687498 | 2688101 | 603 | - | macsReverse_peak_235 | 4.25143 | CD630_23250 | sense | gatC | PTS system, galactitol-specific IIC component | | | | | | | | |
| CD630 | MACS2 | peak | 2690213 | 2690424 | 211 | - | macsReverse_peak_236 | 3.50093 | CD630_23280 | sense | - | Sigma-54 dpendent regulator, PTS fructose family | | | | | | | | |
| CD630 | MACS2 | peak | 2694659 | 2694733 | 74 | - | macsReverse_peak_237 | 3.25870 | CD630_s0500 | sense | - | Purine | | | | | | | | |
| CD630 | MACS2 | peak | 2713507 | 2713571 | 64 | - | macsReverse_peak_238 | 3.34841 | CD630_23440 | sense | - | Putative membrane protein | | | | | | | | |
| CD630 | MACS2 | peak | 2714270 | 2714482 | 212 | + | macsForward_peak_318 | 3.27218 | CD630_23450 | sense | - | Transcriptional regulator, LysR family | | | | | | | | |
| CD630 | MACS2 | peak | 2720426 | 2720970 | 544 | + | macsForward_peak_319 | 4.11757 | CD630_23510 | antisense | grdB | Glycine reductase complex component B gamma subunit (sel | CD630_n00830 | sense | - | Antisense CDS | | | | |
| CD630 | MACS2 | peak | 2725730 | 2725805 | 75 | + | macsForward_peak_320 | 3.63470 | CD630_RNA_16 | antisense | - | Glycine | | | | | | | | |
| CD630 | MACS2 | peak | 2741517 | 2741583 | 66 | - | macsReverse_peak_239 | 4.78770 | CD630_23750 | antisense | - | Conserved hypothetical protein | | | | | | | | |
| CD630 | MACS2 | peak | 2741968 | 2742275 | 307 | - | macsReverse_peak_240 | 4.54307 | CD630_23760 | sense | - | Putative membrane protein | | | | | | | | |
| CD630 | MACS2 | peak | 2742514 | 2742591 | 77 | - | macsReverse_peak_241 | 4.20865 | CD630_23760 | sense | - | Putative membrane protein | | | | | | | | |
| CD630 | MACS2 | peak | 2742956 | 2743208 | 252 | - | macsReverse_peak_242 | 3.25181 | CD630_23760 | sense | - | Putative membrane protein | CD630_23770 | sense | - | Putative hydrolase, NUDIX family | | | | |
| CD630 | MACS2 | peak | 2744002 | 2744282 | 280 | - | macsReverse_peak_243 | 3.91629 | CD630_23780 | sense | - | Putative membrane protein | | | | | | | | |
| CD630 | MACS2 | peak | 2746877 | 2747122 | 245 | - | macsReverse_peak_244 | 3.83173 | CD630_23810 | sense | iorA | Indole pyruvate ferredoxin/flavodoxin oxidoreductase | | | | | | | | |
| CD630 | MACS2 | peak | 2747598 | 2747738 | 140 | - | macsReverse_peak_245 | 3.62157 | CD630_23810 | sense | iorA | Indole pyruvate ferredoxin/flavodoxin oxidoreductase | | | | | | | | |
| CD630 | MACS2 | peak | 2748094 | 2748474 | 380 | - | macsReverse_peak_246 | 3.18419 | CD630_23810 | sense | iorA | Indole pyruvate ferredoxin/flavodoxin oxidoreductase | CD630_23820 | sense | - | Putative pyridoxal phosphate-dependent transferase | | | | |
| CD630 | MACS2 | peak | 2756316 | 2756485 | 169 | + | macsForward_peak_321 | 5.83068 | CD630_23861 | sense | - | Conserved hypothetical protein | | | | | | | | |
| CD630 | MACS2 | peak | 2758242 | 2758500 | 258 | + | macsForward_peak_322 | 4.24581 | CD630_23880 | antisense | - | Putative peptidoglycan-binding/hydrolysing protein | | | | | | | | |
| CD630 | MACS2 | peak | 2763104 | 2763200 | 96 | - | macsReverse_peak_247 | 3.05650 | CD630_23930 | sense | - | Putative glycosyl transferase, family 51 | | | | | | | | |
| CD630 | MACS2 | peak | 2774225 | 2774517 | 292 | - | macsReverse_peak_248 | 3.70192 | CD630_24050 | sense | - | Transcriptional regulator, MarR family | | | | | | | | |
| CD630 | MACS2 | peak | 2780618 | 2780709 | 91 | + | macsForward_peak_323 | 5.00641 | . | | . | NA | | | | | | | | |
| CD630 | MACS2 | peak | 2796906 | 2797084 | 178 | - | macsReverse_peak_249 | 4.18037 | CD630_24220 | sense | - | Sodium:alanine symporter | | | | | | | | |
| CD630 | MACS2 | peak | 2835265 | 2835570 | 305 | - | macsReverse_peak_250 | 3.88519 | CD630_24580 | sense | - | Transporter, Major Facilitator Superfamily (MFS) | | | | | | | | |
| CD630 | MACS2 | peak | 2836226 | 2836784 | 558 | - | macsReverse_peak_251 | 3.70401 | CD630_24580 | sense | - | Transporter, Major Facilitator Superfamily (MFS) | CD630_24590 | sense | - | Putative glucokinase, ROK family | | | | |
| CD630 | MACS2 | peak | 2838130 | 2838217 | 87 | - | macsReverse_peak_252 | 3.54497 | CD630_24591 | sense | - | Conserved hypothetical protein | | | | | | | | |
| CD630 | MACS2 | peak | 2857465 | 2857857 | 392 | - | macsReverse_peak_253 | 5.19485 | CD630_24760 | sense | - | Conserved hypothetical protein | | | | | | | | |
| CD630 | MACS2 | peak | 2859385 | 2859512 | 127 | - | macsReverse_peak_254 | 3.20798 | CD630_24770 | sense | - | Conserved hypothetical protein | | | | | | | | |
| CD630 | MACS2 | peak | 2862992 | 2863114 | 122 | - | macsReverse_peak_255 | 3.30908 | CD630_24810 | sense | ung | Uracil-DNA glycosylase (UDG) | | | | | | | | |
| CD630 | MACS2 | peak | 2864310 | 2864367 | 57 | + | macsForward_peak_324 | 4.98239 | CD630_24830 | sense | - | Putative lipoprotein | | | | | | | | |
| CD630 | MACS2 | peak | 2873964 | 2874019 | 55 | + | macsForward_peak_325 | 3.86845 | CD630_24910 | sense | pmi | Mannose-6-phosphate isomerase (Phosphomannose isomerase) (PMI) (Phosphohexomutase) | | | | | | | | |
| CD630 | MACS2 | peak | 2876898 | 2877118 | 220 | - | macsReverse_peak_256 | 3.89455 | CD630_24920 | sense | - | Two-component sensor histidine kinase, sporulation-associated spo0A | | | | | | | | |
| CD630 | MACS2 | peak | 2886131 | 2886227 | 96 | - | macsReverse_peak_257 | 3.37631 | CD630_s0510 | sense | - | T-box (Arg) | | | | | | | | |
| CD630 | MACS2 | peak | 2888136 | 2888643 | 507 | + | macsForward_peak_326 | 4.17721 | CD630_25020 | sense | - | Putative pyridoxal phosphate-dependent transferase | | | | | | | | |
| CD630 | MACS2 | peak | 2891489 | 2891549 | 60 | - | macsReverse_peak_258 | 7.42886 | CD630_RNA_7 | sense | - | yybP-ykoY | | | | | | | | |
| CD630 | MACS2 | peak | 2892158 | 2892262 | 104 | - | macsReverse_peak_259 | 3.43898 | CD630_25040 | sense | - | Putative D-alanyl-D-alanine carboxypeptidase | | | | | | | | |
| CD630 | MACS2 | peak | 2893371 | 2893894 | 523 | + | macsForward_peak_327 | 4.33624 | CD630_25050 | sense | - | Transcriptional regulator, TetR family | CD630_25060 | sense | LinCd | Transporter, Major Facilitator Superfamily (MFS) | | | | |
| CD630 | MACS2 | peak | 2894557 | 2894682 | 125 | + | macsForward_peak_328 | 3.39358 | CD630_25060 | sense | LinCd | Transporter, Major Facilitator Superfamily (MFS) | | | | | | | | |
| CD630 | MACS2 | peak | 2895196 | 2895281 | 85 | - | macsReverse_peak_260 | 3.45164 | CD630_25070 | sense | - | Conserved hypothetical protein | | | | | | | | |
| CD630 | MACS2 | peak | 2904038 | 2904405 | 367 | + | macsForward_peak_329 | 3.79339 | CD630_25150 | sense | - | Putative L-aspartate-beta-decarboxylase | | | | | | | | |
| CD630 | MACS2 | peak | 2907274 | 2907386 | 112 | + | macsForward_peak_330 | 5.53585 | CD630_n00850 | sense | - | Antisense CDS | CD630_Cdi1_7 | sense | - | GEMM RNA motif | | | | |
| CD630 | MACS2 | peak | 2908005 | 2908417 | 412 | + | macsForward_peak_331 | 6.97567 | CD630_n00860 | antisense | - | ncRNA IGR | Rcd8 | sense | | | CD630_25171 | antisense | - | Putative phage protein |
| CD630 | MACS2 | peak | 2908696 | 2909236 | 540 | - | macsReverse_peak_261 | 4.40865 | CD630_25180 | sense | cwp29 | Cell surface protein | | | | | | | | |
| CD630 | MACS2 | peak | 2909770 | 2910001 | 231 | - | macsReverse_peak_262 | 4.16259 | CD630_25180 | sense | cwp29 | Cell surface protein | | | | | | | | |
| CD630 | MACS2 | peak | 2910957 | 2911091 | 134 | + | macsForward_peak_332 | 3.19663 | CD630_25190 | sense | - | Integrase | | | | | | | | |
| CD630 | MACS2 | peak | 2911684 | 2911905 | 221 | - | macsReverse_peak_263 | 3.65158 | CD630_25200 | sense | - | Putative conjugative transposon antibiotic resistance protein | | | | | | | | |
| CD630 | MACS2 | peak | 2919119 | 2919316 | 197 | + | macsForward_peak_333 | 3.07795 | CD630_25260 | sense | glgB | 1,4-alpha-glucan branching enzyme | | | | | | | | |
| CD630 | MACS2 | peak | 2919534 | 2919671 | 137 | + | macsForward_peak_334 | 3.33867 | CD630_25260 | sense | glgB | 1,4-alpha-glucan branching enzyme | | | | | | | | |
| CD630 | MACS2 | peak | 2920792 | 2920906 | 114 | + | macsForward_peak_335 | 4.48273 | CD630_25260 | sense | glgB | 1,4-alpha-glucan branching enzyme | | | | | | | | |
| CD630 | MACS2 | peak | 2922968 | 2923023 | 55 | - | macsReverse_peak_264 | 4.49333 | CD630_25300 | sense | - | Transcriptional regulator, AraC family | | | | | | | | |

| | | | | | | | | | | | | | | | | | | | | |
|---|---|---|---|---|---|---|---|---|---|---|---|---|---|---|---|---|---|---|---|---|
| CD630 | MACS2 | peak | 2923384 | 2923616 | 232 | - | macsReverse_peak_265 | 3.21241 | CD630_25300 | sense | - | Transcriptional regulator, AraC family | | | | | | | | |
| CD630 | MACS2 | peak | 2923946 | 2924329 | 383 | - | macsReverse_peak_266 | 3.99861 | CD630_25310 | sense | - | Putative membrane protein | | | | | | | | |
| CD630 | MACS2 | peak | 2924596 | 2924838 | 242 | - | macsReverse_peak_267 | 3.31979 | CD630_25320 | sense | - | Aminotransferase, alanine--glyoxylate transaminase | | | | | | | | |
| CD630 | MACS2 | peak | 2925048 | 2925466 | 418 | - | macsReverse_peak_268 | 4.29415 | CD630_25320 | sense | - | Aminotransferase, alanine--glyoxylate transaminase | | | | | | | | |
| CD630 | MACS2 | peak | 2940729 | 2940885 | 156 | - | macsReverse_peak_269 | 4.00527 | CD630_25430 | sense | - | Putative membrane-bound 0-ACYL transferase, MBOAT family | | | | | | | | |
| CD630 | MACS2 | peak | 2963455 | 2963655 | 200 | - | macsReverse_peak_270 | 3.02685 | CD630_25610 | sense | - | Putative phosphatase | CD630_25620 | sense | - | Conserved hypothetical protein | | | | |
| CD630 | MACS2 | peak | 3001828 | 3001911 | 83 | - | macsReverse_peak_271 | 7.25310 | . | | . | NA | | | | | | | | |
| CD630 | MACS2 | peak | 3007784 | 3007871 | 87 | - | macsReverse_peak_272 | 3.53584 | CD630_26000 | sense | cstA | Carbon starvation protein, CstA | | | | | | | | |
| CD630 | MACS2 | peak | 3010752 | 3011077 | 325 | - | macsReverse_peak_273 | 4.03155 | CD630_26020 | sense | - | Two-component sensor histidine kinase | | | | | | | | |
| CD630 | MACS2 | peak | 3012301 | 3012536 | 235 | + | macsForward_peak_336 | 5.84368 | CD630_26030 | sense | cdtR | Clostridium difficile binary toxin regulatory gene, LytTR family | | | | | | | | |
| CD630 | MACS2 | peak | 3014053 | 3014378 | 325 | + | macsForward_peak_337 | 3.54815 | CD630_26050_p1 | sense | - | Fragment of ADP-ribosyltransferase CdtAB (Part 3) | | | | | | | | |
| CD630 | MACS2 | peak | 3015079 | 3015381 | 302 | + | macsForward_peak_338 | 6.99851 | CD630_26050_p4 | sense | - | Fragment of ADP-ribosyltransferase CdtAB (Part 6) | CD630_26050_p5 | sense | - | Fragment of ADP-ribosyltransferase CdtAB (Part 7) | | | | |
| CD630 | MACS2 | peak | 3015664 | 3015745 | 81 | - | macsReverse_peak_274 | 7.09360 | . | | . | NA | | | | | | | | |
| CD630 | MACS2 | peak | 3015693 | 3015816 | 123 | + | macsForward_peak_339 | 6.57474 | . | | . | NA | | | | | | | | |
| CD630 | MACS2 | peak | 3022323 | 3022661 | 338 | + | macsForward_peak_340 | 4.06302 | CD630_26150 | sense | - | Transcriptional regulator, TetR family | | | | | | | | |
| CD630 | MACS2 | peak | 3031925 | 3032242 | 317 | + | macsForward_peak_341 | 5.03379 | CD630_26250 | sense | - | Putative membrane protein | | | | | | | | |
| CD630 | MACS2 | peak | 3032803 | 3033019 | 216 | + | macsForward_peak_342 | 3.96055 | CD630_26250 | sense | - | Putative membrane protein | | | | | | | | |
| CD630 | MACS2 | peak | 3064759 | 3065092 | 333 | + | macsForward_peak_343 | 5.59616 | CD630_26560 | antisense | spoVD | Stage V sporulation protein D (Sporulation-specific penicillin-binding protein) | | | | | | | | |
| CD630 | MACS2 | peak | 3074473 | 3074751 | 278 | - | macsReverse_peak_275 | 4.61557 | CD630_26630 | sense | - | Putative signaling protein | | | | | | | | |
| CD630 | MACS2 | peak | 3082275 | 3082583 | 308 | - | macsReverse_peak_276 | 3.53956 | CD630_26680 | sense | - | Transcription antiterminator, licT family | | | | | | | | |
| CD630 | MACS2 | peak | 3084395 | 3084705 | 310 | - | macsReverse_peak_277 | 3.96230 | CD630_26691 | sense | - | Putative Na(+)/H(+) antiporter | | | | | | | | |
| CD630 | MACS2 | peak | 3085458 | 3085633 | 175 | - | macsReverse_peak_278 | 4.25570 | CD630_26700 | sense | appF | ABC-type transport system, ATP-binding protein | | | | | | | | |
| CD630 | MACS2 | peak | 3089056 | 3089111 | 55 | + | macsForward_peak_344 | 6.44103 | CD630_26730 | sense | appB | ABC-type transport system, oligopeptide-family permease protein | | | | | | | | |
| CD630 | MACS2 | peak | 3089518 | 3089977 | 459 | + | macsForward_peak_345 | 4.50438 | CD630_26730 | sense | appB | ABC-type transport system, oligopeptide-family permease pr | CD630_26740 | sense | appC | ABC-type transport system, oligopeptide-family permease protein | | | | |
| CD630 | MACS2 | peak | 3090196 | 3090539 | 343 | + | macsForward_peak_346 | 4.13830 | CD630_26740 | sense | appC | ABC-type transport system, oligopeptide-family permease protein | | | | | | | | |
| CD630 | MACS2 | peak | 3090886 | 3091560 | 674 | - | macsReverse_peak_279 | 3.60610 | CD630_26750 | sense | - | Transcriptional regulator, LysR family | | | | | | | | |
| CD630 | MACS2 | peak | 3107399 | 3107565 | 166 | - | macsReverse_peak_280 | 3.62689 | CD630_26880 | sense | sspA | Small, acid-soluble spore protein alpha | | | | | | | | |
| CD630 | MACS2 | peak | 3110802 | 3111030 | 228 | + | macsForward_peak_347 | 4.58060 | CD630_26920 | sense | - | Putative threonine-phosphate decarboxylase | | | | | | | | |
| CD630 | MACS2 | peak | 3125693 | 3126163 | 470 | + | macsForward_peak_348 | 3.51812 | CD630_27040 | sense | - | Putative permease | | | | | | | | |
| CD630 | MACS2 | peak | 3126485 | 3126703 | 218 | + | macsForward_peak_349 | 3.20020 | CD630_27040 | sense | - | Putative permease | | | | | | | | |
| CD630 | MACS2 | peak | 3128630 | 3128925 | 295 | - | macsReverse_peak_281 | 4.06049 | CD630_27060 | sense | - | Conserved hypothetical protein | | | | | | | | |
| CD630 | MACS2 | peak | 3136706 | 3137263 | 557 | + | macsForward_peak_350 | 3.78231 | CD630_27110 | sense | - | Transcriptional regulator, LysR family | | | | | | | | |
| CD630 | MACS2 | peak | 3137395 | 3137537 | 142 | - | macsReverse_peak_282 | 4.43720 | CD630_n00910 | sense | - | Antisense CDS | CD630_27120 | sense | - | Conserved hypothetical protein | | | | |
| CD630 | MACS2 | peak | 3156611 | 3156913 | 302 | - | macsReverse_peak_283 | 3.42516 | CD630_27200 | sense | - | Putative transporter | | | | | | | | |
| CD630 | MACS2 | peak | 3157783 | 3158744 | 961 | - | macsReverse_peak_284 | 4.18786 | CD630_27210 | sense | - | Two-component sensor histidine kinase | CD630_27220 | sense | - | Two-component response regulator | | | | |
| CD630 | MACS2 | peak | 3159674 | 3159815 | 141 | - | macsReverse_peak_285 | 3.29372 | CD630_27230 | sense | - | Putative membrane protein | | | | | | | | |
| CD630 | MACS2 | peak | 3160528 | 3160835 | 307 | - | macsReverse_peak_286 | 3.52462 | CD630_27230 | sense | - | Putative membrane protein | CD630_27240 | sense | - | Putative polysaccharide deacetylase | | | | |
| CD630 | MACS2 | peak | 3164455 | 3164689 | 234 | + | macsForward_peak_351 | 3.50892 | CD630_27270 | sense | - | Putative membrane protein | | | | | | | | |
| CD630 | MACS2 | peak | 3165426 | 3165535 | 109 | + | macsForward_peak_352 | 3.24738 | CD630_27280 | sense | - | Putative membrane protein | | | | | | | | |
| CD630 | MACS2 | peak | 3166680 | 3166781 | 101 | - | macsReverse_peak_287 | 4.01622 | CD630_27310 | sense | - | Putative membrane protein | | | | | | | | |
| CD630 | MACS2 | peak | 3174237 | 3174549 | 312 | + | macsForward_peak_353 | 3.79570 | CD630_27360 | sense | - | Transcriptional regulator, LysR family | | | | | | | | |
| CD630 | MACS2 | peak | 3174921 | 3175027 | 106 | + | macsForward_peak_354 | 3.75375 | CD630_27360 | sense | - | Transcriptional regulator, LysR family | | | | | | | | |
| CD630 | MACS2 | peak | 3193205 | 3193257 | 52 | - | macsReverse_peak_288 | 3.15785 | CD630_27500 | sense | agrB | Accessory gene regulator | | | | | | | | |
| CD630 | MACS2 | peak | 3193539 | 3193680 | 141 | - | macsReverse_peak_289 | 3.92598 | CD630_27500 | sense | agrB | Accessory gene regulator | | | | | | | | |
| CD630 | MACS2 | peak | 3195959 | 3196394 | 435 | + | macsForward_peak_355 | 4.27693 | CD630_27520 | sense | - | Conserved hypothetical protein | | | | | | | | |
| CD630 | MACS2 | peak | 3196739 | 3196812 | 73 | - | macsReverse_peak_290 | 5.79130 | CD630_n00930 | sense | - | ncRNA IGR | | | | | | | | |
| CD630 | MACS2 | peak | 3204656 | 3205099 | 443 | - | macsReverse_peak_291 | 3.46216 | CD630_27570 | sense | - | Putative acyl transferase/phospholipase | | | | | | | | |
| CD630 | MACS2 | peak | 3208424 | 3208558 | 134 | - | macsReverse_peak_292 | 3.48384 | CD630_27600 | sense | - | Putative signaling protein | | | | | | | | |
| CD630 | MACS2 | peak | 3209637 | 3209750 | 113 | - | macsReverse_peak_293 | 3.96465 | CD630_27610 | sense | - | Putative N-acetylmuramoyl-L-alanine amidase | | | | | | | | |
| CD630 | MACS2 | peak | 3211700 | 3211841 | 141 | + | macsForward_peak_356 | 3.54199 | CD630_27630 | sense | - | Putative lipoprotein | | | | | | | | |
| CD630 | MACS2 | peak | 3228110 | 3228214 | 104 | - | macsReverse_peak_294 | 4.08588 | CD630_27770 | sense | - | Putative membrane protein | | | | | | | | |
| CD630 | MACS2 | peak | 3285851 | 3286030 | 179 | + | macsForward_peak_357 | 4.18447 | CD630_28150 | sense | - | Conserved hypothetical protein | | | | | | | | |
| CD630 | MACS2 | peak | 3290645 | 3290922 | 277 | - | macsReverse_peak_295 | 3.91299 | CD630_28181 | sense | - | Fragment of dihydrofolate reductase family protein | | | | | | | | |
| CD630 | MACS2 | peak | 3298865 | 3299164 | 299 | - | macsReverse_peak_296 | 3.51613 | CD630_28260 | sense | - | Putative aldolase | | | | | | | | |
| CD630 | MACS2 | peak | 3300239 | 3300951 | 712 | + | macsForward_peak_358 | 4.49556 | CD630_28280 | sense | - | Putative pyridoxal phosphate-dependent transferase | | | | | | | | |
| CD630 | MACS2 | peak | 3301186 | 3301288 | 102 | + | macsForward_peak_359 | 4.59833 | CD630_28280 | sense | - | Putative pyridoxal phosphate-dependent transferase | | | | | | | | |
| CD630 | MACS2 | peak | 3301681 | 3302536 | 855 | + | macsForward_peak_360 | 5.54192 | CD630_28290 | sense | - | Putative membrane protein | | | | | | | | |
| CD630 | MACS2 | peak | 3306694 | 3307153 | 459 | - | macsReverse_peak_297 | 3.99184 | CD630_Cdi2_3 | sense | - | c-di-GMP-II | CD630_28320 | sense | - | Putative cation efflux protein | | | | |
| CD630 | MACS2 | peak | 3308890 | 3309067 | 177 | - | macsReverse_peak_298 | 3.29122 | CD630_28330 | sense | - | Putative calcium-transporting ATPase | | | | | | | | |
| CD630 | MACS2 | peak | 3320836 | 3320959 | 123 | - | macsReverse_peak_299 | 3.45858 | CD630_28400 | sense | serS1 | Seryl-tRNA synthetase 1 (Seryl-tRNA(Ser/Sec) synthetase 1) (Serine--tRNA ligase 1) (SerRS 1) | | | | | | | | |
| CD630 | MACS2 | peak | 3334444 | 3334522 | 78 | - | macsReverse_peak_300 | 3.08671 | CD630_28520 | sense | dltB | D-alanyl transferase DltB, MBOAT family | | | | | | | | |
| CD630 | MACS2 | peak | 3334832 | 3335101 | 269 | - | macsReverse_peak_301 | 3.81485 | CD630_28520 | sense | dltB | D-alanyl transferase DltB, MBOAT family | CD630_28530 | sense | dltA | D-alanine--poly(phosphoribitol) ligase subunit 1 | | | | |
| CD630 | MACS2 | peak | 3335920 | 3336371 | 451 | - | macsReverse_peak_302 | 3.85429 | CD630_28530 | sense | dltA | D-alanine--poly(phosphoribitol) ligase subunit 1 | | | | | | | | |
| CD630 | MACS2 | peak | 3337439 | 3337746 | 307 | - | macsReverse_peak_303 | 4.40777 | CD630_28540 | sense | dltD | D-alanine transferase | | | | | | | | |
| CD630 | MACS2 | peak | 3338191 | 3338530 | 339 | - | macsReverse_peak_304 | 3.51196 | CD630_28550 | sense | - | Putative NAD(P)-binding oxidoreductase | | | | | | | | |
| CD630 | MACS2 | peak | 3339390 | 3339742 | 352 | - | macsReverse_peak_305 | 3.96803 | CD630_28560 | sense | - | Putative membrane protein | | | | | | | | |
| CD630 | MACS2 | peak | 3341811 | 3341933 | 122 | - | macsReverse_peak_306 | 4.67130 | CD630_28580 | sense | - | Putative membrane protein | | | | | | | | |
| CD630 | MACS2 | peak | 3357684 | 3358377 | 693 | + | macsForward_peak_361 | 4.49207 | CD630_28700 | sense | kdgT1 | 2-keto-3-deoxygluconate permease 1 | | | | | | | | |
| CD630 | MACS2 | peak | 3358863 | 3359026 | 163 | + | macsForward_peak_362 | 4.26165 | CD630_28720 | sense | uxaA | Fragment of C-terminal D-galactate dehydratase/Altronate h | CD630_28710 | sense | uxaAprim | Fragment of N-terminal carbohydrate hydrolase, SAF domain | | | | |
| CD630 | MACS2 | peak | 3359649 | 3359947 | 298 | + | macsForward_peak_363 | 3.58606 | CD630_28720 | sense | uxaA | Fragment of C-terminal D-galactate dehydratase/Altronate hydrolase | | | | | | | | |
| CD630 | MACS2 | peak | 3376194 | 3376419 | 225 | + | macsForward_peak_364 | 3.41991 | CD630_28870 | sense | - | Putative diguanylate kinase signaling protein | | | | | | | | |
| CD630 | MACS2 | peak | 3380080 | 3380348 | 268 | + | macsForward_peak_365 | 5.31318 | CD630_n00980 | sense | - | Antisense CDS | CD630_Cdi1_4 | sense | - | GEMM RNA motif | CD630_28890 | antisense | - | Putative phage protein |
| CD630 | MACS2 | peak | 3380250 | 3380421 | 171 | - | macsReverse_peak_307 | 7.52469 | CD630_n00980 | antisense | - | Antisense CDS | CD630_Cdi1_4 | antisense | - | GEMM RNA motif | CD630_28890 | sense | - | Putative phage protein |
| CD630 | MACS2 | peak | 3380757 | 3380927 | 170 | + | macsForward_peak_366 | 4.12438 | . | | . | NA | | | | | | | | |
| CD630 | MACS2 | peak | 3382212 | 3382724 | 512 | - | macsReverse_peak_308 | 4.51664 | CD630_28910 | sense | - | Putative phage protein | CD630_28920 | sense | - | Putative phage protein | | | | |
| CD630 | MACS2 | peak | 3396473 | 3396567 | 94 | - | macsReverse_peak_309 | 3.89477 | CD630_29070 | sense | - | Putative phage lipoprotein | | | | | | | | |
| CD630 | MACS2 | peak | 3396937 | 3397064 | 127 | - | macsReverse_peak_310 | 3.72051 | CD630_n00990 | sense | - | ncRNA IGR | | | | | | | | |
| CD630 | MACS2 | peak | 3397893 | 3398087 | 194 | + | macsForward_peak_367 | 5.45665 | CD630_n00990 | antisense | - | ncRNA IGR | | | | | CD630_29072 | antisense | | |
| CD630 | MACS2 | peak | 3398317 | 3398663 | 346 | + | macsForward_peak_368 | 8.96925 | Rcd9 | sense | | Antisense CDS | CD630_29071 | antisense | - | Putative phage protein | | | | |
| CD630 | MACS2 | peak | 3410514 | 3410870 | 356 | + | macsForward_peak_369 | 4.61901 | CD630_29250 | sense | - | Putative phage protein | CD630_29521_p07 | antisense | - | Fragment of ABC-type transport system, ATP-binding protein (Part 7) | | | | |
| CD630 | MACS2 | peak | 3412624 | 3413086 | 462 | - | macsReverse_peak_311 | 3.70174 | CD630_29270 | sense | - | Transcriptional regulator, Phage-type | | | | | | | | |
| CD630 | MACS2 | peak | 3426605 | 3426911 | 306 | + | macsForward_peak_370 | 3.70644 | CD630_29500 | sense | - | Transcriptional regulator, HTH-type | | | | | | | | |
| CD630 | MACS2 | peak | 3442721 | 3442849 | 128 | - | macsReverse_peak_312 | 3.67128 | CD630_29650 | sense | - | Putative signaling protein | | | | | | | | |
| CD630 | MACS2 | peak | 3446562 | 3446767 | 205 | - | macsReverse_peak_313 | 3.06485 | CD630_29660 | sense | adhE | Aldehyde-alcohol dehydrogenase | | | | | | | | |
| CD630 | MACS2 | peak | 3448748 | 3449176 | 428 | - | macsReverse_peak_314 | 4.62758 | CD630_29670 | sense | dpaB | Dipicolinate synthase subunit B | | | | | | | | |
| CD630 | MACS2 | peak | 3450136 | 3450257 | 121 | - | macsReverse_peak_315 | 4.65364 | CD630_29680 | sense | dpaA | Dipicolinate synthase subunit A | | | | | | | | |

| | | | | | | | | | | | | | | | | | | | | |
|---|---|---|---|---|---|---|---|---|---|---|---|---|---|---|---|---|---|---|---|---|
| CD630 | MACS2 | peak | 3456571 | 3457171 | 600 | - | macsReverse_peak_316 | 4.59146 | CD630_n01010 | sense | - | ncRNA IGR | CD630_29740 | antisense | - | Conserved hypothetical protein | CD630_29750 | sense | - | Putative CRISPR-associated Cas2 family protein |
| CD630 | MACS2 | peak | 3457495 | 3457573 | 78 | - | macsReverse_peak_317 | 3.51074 | CD630_29760 | sense | - | Putative CRISPR-associated Cas1 family protein | | | | | | | | |
| CD630 | MACS2 | peak | 3457931 | 3458190 | 259 | - | macsReverse_peak_318 | 3.39713 | CD630_29760 | sense | - | Putative CRISPR-associated Cas1 family protein | | | | | | | | |
| CD630 | MACS2 | peak | 3466591 | 3466752 | 161 | - | macsReverse_peak_319 | 4.48825 | CD630_29840 | sense | - | ABC-type transport system, permease | | | | | | | | |
| CD630 | MACS2 | peak | 3480497 | 3481132 | 635 | + | macsForward_peak_371 | 4.51904 | CD630_29960 | sense | - | Conserved hypothetical protein | | | | | | | | |
| CD630 | MACS2 | peak | 3496733 | 3497052 | 319 | - | macsReverse_peak_320 | 4.13234 | CD630_30100 | sense | - | Putative cytochrome C assembly protein | | | | | | | | |
| CD630 | MACS2 | peak | 3497884 | 3498012 | 128 | - | macsReverse_peak_321 | 3.31041 | CD630_30110 | sense | - | Transcriptional regulator, HxlR family DUF24 | | | | | | | | |
| CD630 | MACS2 | peak | 3498582 | 3498672 | 90 | - | macsReverse_peak_322 | 3.15521 | CD630_30120 | sense | - | Putative alpha-mannosidase | | | | | | | | |
| CD630 | MACS2 | peak | 3498932 | 3499296 | 364 | - | macsReverse_peak_323 | 3.49980 | CD630_30120 | sense | - | Putative alpha-mannosidase | | | | | | | | |
| CD630 | MACS2 | peak | 3500184 | 3500314 | 130 | - | macsReverse_peak_324 | 4.04859 | CD630_30120 | sense | - | Putative alpha-mannosidase | | | | | | | | |
| CD630 | MACS2 | peak | 3500662 | 3500784 | 122 | - | macsReverse_peak_325 | 3.12380 | CD630_30120 | sense | - | Putative alpha-mannosidase | | | | | | | | |
| CD630 | MACS2 | peak | 3502315 | 3502449 | 134 | - | macsReverse_peak_326 | 3.28343 | CD630_30130 | sense | - | PTS system, mannose-specfic IIC component | CD630_30140 | sense | - | PTS system, mannose-specfic IIB component | | | | |
| CD630 | MACS2 | peak | 3507621 | 3507806 | 185 | - | macsReverse_peak_327 | 3.27394 | CD630_30190 | sense | - | Putative transporter | | | | | | | | |
| CD630 | MACS2 | peak | 3517968 | 3518194 | 226 | - | macsReverse_peak_328 | 4.46345 | CD630_30290 | sense | malY | Bifunctional protein: cystathionine beta-lyase / repressor | | | | | | | | |
| CD630 | MACS2 | peak | 3520119 | 3520512 | 393 | - | macsReverse_peak_329 | 4.56059 | CD630_30310 | sense | - | Transcription antiterminator, PTS operon regulator | | | | | | | | |
| CD630 | MACS2 | peak | 3526616 | 3526948 | 332 | - | macsReverse_peak_330 | 4.26509 | CD630_30360 | sense | - | Transporter, Major Facilitator Superfamily (MFS) | | | | | | | | |
| CD630 | MACS2 | peak | 3534988 | 3535424 | 436 | + | macsForward_peak_372 | 5.27539 | CD630_30420 | sense | - | Putative membrane protein | | | | | | | | |
| CD630 | MACS2 | peak | 3535884 | 3536027 | 143 | + | macsForward_peak_373 | 4.04338 | CD630_30420 | sense | - | Putative membrane protein | CD630_30430 | antisense | - | Conserved hypothetical protein | | | | |
| CD630 | MACS2 | peak | 3546922 | 3547539 | 617 | - | macsReverse_peak_331 | 3.63804 | CD630_30540 | sense | - | Two-component response regulator | | | | | | | | |
| CD630 | MACS2 | peak | 3555992 | 3556542 | 550 | - | macsReverse_peak_332 | 4.15443 | CD630_30620 | sense | - | Transcriptional regulator, RpiR family | | | | | | | | |
| CD630 | MACS2 | peak | 3562259 | 3562560 | 301 | - | macsReverse_peak_333 | 3.41240 | CD630_30660 | antisense | xylR | Transcriptional regulator, Xylose repressor | CD630_SQ2179 | sense | | | | | | |
| CD630 | MACS2 | peak | 3562312 | 3562484 | 172 | + | macsForward_peak_374 | 3.95671 | CD630_30660 | sense | xylR | Transcriptional regulator, Xylose repressor | | | | | | | | |
| CD630 | MACS2 | peak | 3569463 | 3569707 | 244 | + | macsForward_peak_375 | 3.13236 | CD630_30730 | sense | - | Putative membrane protein | | | | | | | | |
| CD630 | MACS2 | peak | 3581623 | 3581826 | 203 | - | macsReverse_peak_334 | 3.72836 | CD630_30840 | sense | garK | Glycerate kinase | | | | | | | | |
| CD630 | MACS2 | peak | 3607070 | 3607259 | 189 | - | macsReverse_peak_335 | 3.70946 | CD630_31000 | sense | - | Putative C4-dicarboxylate anaerobic carrier, DcuC family | | | | | | | | |
| CD630 | MACS2 | peak | 3607716 | 3607953 | 237 | - | macsReverse_peak_336 | 8.13716 | CD630_s0590 | sense | - | Lysine | CD630_31001 | antisense | - | Fragment of artefact | | | | |
| CD630 | MACS2 | peak | 3608237 | 3608330 | 93 | - | macsReverse_peak_337 | 4.96305 | CD630_31010 | sense | - | PTS system, glucose-like IIBC component | | | | | | | | |
| CD630 | MACS2 | peak | 3631128 | 3631193 | 65 | - | macsReverse_peak_338 | 6.84967 | CD630_s0591 | sense | - | CD630_31170 | | | | | | | | |
| CD630 | MACS2 | peak | 3634735 | 3634977 | 242 | + | macsForward_peak_376 | 4.17693 | CD630_31210 | sense | - | Putative flavodoxin/nitric oxide synthase | | | | | | | | |
| CD630 | MACS2 | peak | 3635424 | 3635580 | 156 | - | macsReverse_peak_339 | 3.73779 | CD630_31220 | sense | - | Transcriptional regulator, ArsR family | | | | | | | | |
| CD630 | MACS2 | peak | 3635687 | 3636047 | 360 | + | macsForward_peak_377 | 4.01108 | CD630_31220 | antisense | - | Transcriptional regulator, ArsR family | CD630_31230 | antisense | - | Putative nitroreductase | | | | |
| CD630 | MACS2 | peak | 3647311 | 3647661 | 350 | - | macsReverse_peak_340 | 4.50315 | CD630_31300 | sense | bglA | 6-phospho-beta-glucosidase | | | | | | | | |
| CD630 | MACS2 | peak | 3655352 | 3655419 | 67 | - | macsReverse_peak_341 | 4.86402 | CD630_31360 | sense | bglA | 6-phospho-beta-glucosidase | | | | | | | | |
| CD630 | MACS2 | peak | 3659601 | 3659656 | 55 | - | macsReverse_peak_342 | 5.17071 | . | | . | NA | | | | | | | | |
| CD630 | MACS2 | peak | 3659940 | 3660193 | 253 | - | macsReverse_peak_343 | 3.49137 | CD630_31381 | sense | - | Fragment of putative phage transcriptional regulator (C-term | CD630_31382 | sense | - | Fragment of conserved hypothetical protein (Central region) | | | | |
| CD630 | MACS2 | peak | 3666416 | 3666531 | 115 | - | macsReverse_peak_344 | 3.03040 | CD630_31450 | sense | cbpA | Surface-exposed adhesin | | | | | | | | |
| CD630 | MACS2 | peak | 3669949 | 3670006 | 57 | + | macsForward_peak_378 | 4.41476 | CD630_31461 | sense | - | Fragment of putative transposase, IS4 family (C-terminal region) | | | | | | | | |
| CD630 | MACS2 | peak | 3672550 | 3672989 | 439 | + | macsForward_peak_379 | 4.88897 | CD630_31480 | sense | - | Hypothetical protein | | | | | | | | |
| CD630 | MACS2 | peak | 3674264 | 3674325 | 61 | + | macsForward_peak_380 | 3.61126 | CD630_31480 | sense | - | Hypothetical protein | | | | | | | | |
| CD630 | MACS2 | peak | 3674820 | 3675853 | 1033 | + | macsForward_peak_381 | 3.51267 | CD630_31480 | sense | - | Hypothetical protein | CD630_31490 | sense | - | DNA helicase | | | | |
| CD630 | MACS2 | peak | 3676158 | 3676686 | 528 | + | macsForward_peak_382 | 4.08593 | CD630_31490 | sense | - | DNA helicase | | | | | | | | |
| CD630 | MACS2 | peak | 3676899 | 3677179 | 280 | + | macsForward_peak_383 | 3.37847 | CD630_31490 | sense | - | DNA helicase | | | | | | | | |
| CD630 | MACS2 | peak | 3677797 | 3678077 | 280 | + | macsForward_peak_384 | 3.80651 | CD630_31490 | sense | - | DNA helicase | | | | | | | | |
| CD630 | MACS2 | peak | 3678516 | 3678944 | 428 | + | macsForward_peak_385 | 4.33960 | CD630_31500 | sense | - | Conserved hypothetical protein | | | | | | | | |
| CD630 | MACS2 | peak | 3679182 | 3679686 | 504 | + | macsForward_peak_386 | 3.66435 | CD630_31500 | sense | - | Conserved hypothetical protein | | | | | | | | |
| CD630 | MACS2 | peak | 3680089 | 3680331 | 242 | + | macsForward_peak_387 | 4.90207 | CD630_31500 | sense | - | Conserved hypothetical protein | | | | | | | | |
| CD630 | MACS2 | peak | 3682591 | 3682846 | 255 | + | macsForward_peak_388 | 3.49100 | CD630_31530 | sense | - | Transcriptional regulator, Phage-type | | | | | | | | |
| CD630 | MACS2 | peak | 3687415 | 3687733 | 318 | + | macsForward_peak_389 | 11.25295 | CD630_s0600 | antisense | - | tmRNA | | | | | | | | |
| CD630 | MACS2 | peak | 3689471 | 3689705 | 234 | + | macsForward_peak_390 | 3.51756 | CD630_31580 | sense | - | Transcriptional regulator, TRAP family | | | | | | | | |
| CD630 | MACS2 | peak | 3702050 | 3702239 | 189 | + | macsForward_peak_391 | 3.71193 | CD630_31690 | sense | - | Putative malate transporter | | | | | | | | |
| CD630 | MACS2 | peak | 3731537 | 3731706 | 169 | + | macsForward_peak_392 | 3.97522 | CD630_31880 | sense | - | Conserved hypothetical protein | CD630_n01060 | sense | - | Antisense CDS | | | | |
| CD630 | MACS2 | peak | 3733655 | 3733916 | 261 | - | macsReverse_peak_345 | 3.48314 | CD630_31910 | sense | - | Putative oxidoreductase | | | | | | | | |
| CD630 | MACS2 | peak | 3741795 | 3742010 | 215 | + | macsForward_peak_393 | 4.93757 | CD630_31960 | sense | - | Putative penicillin-binding protein | | | | | | | | |
| CD630 | MACS2 | peak | 3742877 | 3743043 | 166 | + | macsForward_peak_394 | 3.24511 | CD630_31970 | sense | - | Transcriptional regulator, MerR family | | | | | | | | |
| CD630 | MACS2 | peak | 3750540 | 3751240 | 700 | - | macsReverse_peak_346 | 4.93516 | CD630_32040 | sense | - | Putative membrane protein | | | | | | | | |
| CD630 | MACS2 | peak | 3752164 | 3753101 | 937 | + | macsForward_peak_395 | 5.73797 | CD630_32050 | sense | - | Putative nitroreductase | CD630_32060 | sense | - | Putative membrane protein | CD630_n01080 | antisense | - | Antisense CDS |
| CD630 | MACS2 | peak | 3754026 | 3754200 | 174 | - | macsReverse_peak_347 | 3.51964 | CD630_32070 | sense | - | Putative multi antimicrobial extrusion protein, MatE family | | | | | | | | |
| CD630 | MACS2 | peak | 3757092 | 3757177 | 85 | + | macsForward_peak_396 | 4.45291 | CD630_32110 | sense | - | Conserved hypothetical protein | | | | | | | | |
| CD630 | MACS2 | peak | 3762706 | 3762768 | 62 | - | macsReverse_peak_348 | 3.57422 | . | | . | NA | | | | | | | | |
| CD630 | MACS2 | peak | 3773076 | 3773176 | 100 | - | macsReverse_peak_349 | 5.84172 | CD630_s0610 | antisense | - | Lysine | | | | | | | | |
| CD630 | MACS2 | peak | 3779121 | 3779466 | 345 | + | macsForward_peak_397 | 3.27218 | CD630_32290 | sense | dapB2 | Dihydrodipicolinate reductase 2 (DHPR 2) | | | | | | | | |
| CD630 | MACS2 | peak | 3788915 | 3789729 | 814 | + | macsForward_peak_398 | 4.47959 | CD630_32360 | antisense | - | Putative membrane protein | | | | | | | | |
| CD630 | MACS2 | peak | 3801097 | 3801335 | 238 | - | macsReverse_peak_350 | 3.76949 | . | | . | NA | | | | | | | | |
| CD630 | MACS2 | peak | 3804771 | 3805235 | 464 | + | macsForward_peak_399 | 6.17288 | CD630_32500 | sense | - | Conserved hypothetical protein | | | | | | | | |
| CD630 | MACS2 | peak | 3806306 | 3806517 | 211 | + | macsForward_peak_400 | 4.06098 | CD630_32520 | sense | - | Conserved hypothetical protein | | | | | | | | |
| CD630 | MACS2 | peak | 3806645 | 3806832 | 187 | - | macsReverse_peak_351 | 3.85634 | CD630_32530 | sense | folC | Folylpolyglutamate synthase | | | | | | | | |
| CD630 | MACS2 | peak | 3814035 | 3814189 | 154 | - | macsReverse_peak_352 | 4.95025 | CD630_s0631 | sense | - | T-box, CD630_32560 | CD630_32561 | antisense | - | Conserved hypothetical protein | | | | |
| CD630 | MACS2 | peak | 3826624 | 3826684 | 60 | - | macsReverse_peak_353 | 6.72203 | CD630_n01090 | sense | | | CD630_Cdi2_2 | sense | - | c-di-GMP-II | | | | |
| CD630 | MACS2 | peak | 3829054 | 3829143 | 89 | - | macsReverse_peak_354 | 7.32676 | CD630_32690 | sense | - | Putative oligoendopeptidase F, M3B family | | | | | | | | |
| CD630 | MACS2 | peak | 3830181 | 3831358 | 1177 | - | macsReverse_peak_355 | 4.74276 | CD630_32700 | sense | - | Putative magnesium transport ATPase, MgtC/SapB family | CD630_32710 | sense | - | Conserved hypothetical protein | CD630_32720 | sense | - | Putative membrane protein |
| CD630 | MACS2 | peak | 3833638 | 3833721 | 83 | + | macsForward_peak_401 | 4.52257 | CD630_32740 | sense | feoB | Ferrous iron transport protein B | | | | | | | | |
| CD630 | MACS2 | peak | 3852208 | 3852547 | 339 | - | macsReverse_peak_356 | 3.41276 | CD630_32880 | sense | - | Conserved hypothetical protein | CD630_32890 | sense | - | Conserved hypothetical protein | | | | |
| CD630 | MACS2 | peak | 3861679 | 3863304 | 1625 | + | macsForward_peak_402 | 5.24925 | CD630_s0640 | sense | - | FMN | CD630_32990 | sense | - | Transporter, Major Facilitator Superfamily (MFS) | | | | |
| CD630 | MACS2 | peak | 3866856 | 3867709 | 853 | + | macsForward_peak_403 | 5.27255 | CD630_33020 | sense | chrA | Chromate transport protein ChrA | CD630_33020 | sense | chrA | Chromate transport protein ChrA | | | | |
| CD630 | MACS2 | peak | 3885176 | 3885433 | 257 | - | macsReverse_peak_357 | 4.13899 | CD630_33190 | sense | - | Putative membrane protein | | | | | | | | |
| CD630 | MACS2 | peak | 3894294 | 3894595 | 301 | + | macsForward_peak_404 | 3.79149 | CD630_33300 | sense | - | Transcriptional regulator, HTH-type Tn916-like, CTn6-Orf6 | CD630_33310 | antisense | - | Putative conjugative transposon protein Tn916-like, CTn6-Orf7 | | | | |
| CD630 | MACS2 | peak | 3906191 | 3906255 | 64 | - | macsReverse_peak_358 | 3.22095 | CD630_33420 | sense | - | Putative conjugative transposon protein Tn916-like, CTn6-Orf19 | | | | | | | | |
| CD630 | MACS2 | peak | 3911606 | 3911959 | 353 | + | macsForward_peak_405 | 3.67613 | CD630_33480 | sense | - | Hypothetical protein | | | | | | | | |
| CD630 | MACS2 | peak | 3920112 | 3920763 | 651 | + | macsForward_peak_406 | 4.03147 | CD630_33560 | sense | - | Transcriptional regulator, hxlR family | CD630_33570 | antisense | - | Putative protease/amidase | | | | |
| CD630 | MACS2 | peak | 3920168 | 3921107 | 939 | - | macsReverse_peak_359 | 3.95778 | CD630_33560 | antisense | - | Transcriptional regulator, hxlR family | CD630_33570 | sense | - | Putative protease/amidase | CD630_33580 | sense | - | Fragment of C-terminal transcriptional regulator, AsnC/Lrp family |
| CD630 | MACS2 | peak | 3921333 | 3921396 | 63 | - | macsReverse_peak_360 | 3.26349 | CD630_33580 | sense | - | Fragment of C-terminal transcriptional regulator, AsnC/Lrp family | | | | | | | | |
| CD630 | MACS2 | peak | 3922102 | 3922277 | 175 | + | macsForward_peak_407 | 6.68432 | CD630_s0641 | sense | - | CD630_33590 | CD630_33581 | antisense | - | Conserved hypothetical protein | | | | |
| CD630 | MACS2 | peak | 3923637 | 3923759 | 122 | - | macsReverse_peak_361 | 3.93606 | CD630_33590 | antisense | - | ABC-type transport system, ATP-binding protein | | | | | | | | |
| CD630 | MACS2 | peak | 3924240 | 3924607 | 367 | - | macsReverse_peak_362 | 3.73196 | CD630_33600 | sense | - | Two-component response regulator | CD630_SQ2424 | antisense | - | Antisense CDS | | | | |
| CD630 | MACS2 | peak | 3925870 | 3926384 | 514 | - | macsReverse_peak_363 | 3.79322 | CD630_33610 | sense | - | Putative drug/sodium antiporter, MATE family | | | | | | | | |
| CD630 | MACS2 | peak | 3929833 | 3929909 | 76 | - | macsReverse_peak_364 | 4.88427 | . | | . | NA | | | | | | | | |

| | | | | | | | | | | | | | | | | | | | | |
|---|---|---|---|---|---|---|---|---|---|---|---|---|---|---|---|---|---|---|---|---|
| CD630 | MACS2 | peak | 3930508 | 3930589 | 81 | + | macsForward_peak_408 | 4.80565 | CD630_33650 | sense | - | Putative diguanylate kinase signaling protein | | | | | | | | |
| CD630 | MACS2 | peak | 3930834 | 3931383 | 549 | + | macsForward_peak_409 | 3.97291 | CD630_33650 | sense | - | Putative diguanylate kinase signaling protein | | | | | | | | |
| CD630 | MACS2 | peak | 3935555 | 3935815 | 260 | - | macsReverse_peak_365 | 6.00600 | CD630_s0642 | sense | - | CD630_33681 | | | | | | | | |
| CD630 | MACS2 | peak | 3936262 | 3936555 | 293 | + | macsForward_peak_410 | 5.77720 | CD630_SQ2429 | sense | - | ncRNA IGR | CD630_Cdi1_11 | sense | - | GEMM RNA motif | CD630_33682 | sense | - | Conserved hypothetical protein |
| CD630 | MACS2 | peak | 3981099 | 3981244 | 145 | - | macsReverse_peak_366 | 3.05766 | CD630_34030 | sense | - | Putative formate/nitrite transporter | | | | | | | | |
| CD630 | MACS2 | peak | 3982479 | 3983475 | 996 | + | macsForward_peak_411 | 4.46177 | CD630_34040 | sense | cls | Cardiolipin synthetase (Cardiolipin synthase) (CL synthase) | | | | | | | | |
| CD630 | MACS2 | peak | 3983981 | 3984227 | 246 | + | macsForward_peak_412 | 4.03863 | CD630_34050 | sense | - | Putative iron-only hydrogenase,electron-transferring subunit HymA-like | | | | | | | | |
| CD630 | MACS2 | peak | 3989488 | 3989614 | 126 | - | macsReverse_peak_367 | 3.20224 | CD630_34080 | sense | - | Putative DNA mismatch repair ATPase MutS | | | | | | | | |
| CD630 | MACS2 | peak | 4025902 | 4025984 | 82 | - | macsReverse_peak_368 | 3.13524 | CD630_34350 | sense | cobQ | Cobyric acid synthase CobQ | | | | | | | | |
| CD630 | MACS2 | peak | 4027816 | 4028101 | 285 | - | macsReverse_peak_369 | 3.15854 | CD630_34360 | sense | cobC | Cobalamin biosynthesis phosphoglycerate mutase CobC | | | | | | | | |
| CD630 | MACS2 | peak | 4028550 | 4028808 | 258 | - | macsReverse_peak_370 | 3.67481 | CD630_34370 | sense | cobS | Cobalamin synthase CobS (Cobalamin-5-phosphate synthase) | | | | | | | | |
| CD630 | MACS2 | peak | 4045046 | 4045248 | 202 | + | macsForward_peak_413 | 3.20416 | CD630_34520 | sense | - | Transcriptional regulator, GntR family | | | | | | | | |
| CD630 | MACS2 | peak | 4045567 | 4046184 | 617 | + | macsForward_peak_414 | 3.83016 | CD630_34520 | sense | - | Transcriptional regulator, GntR family | CD630_34530 | sense | - | N-acetylglucosamine-6-phosphate deacetylase | | | | |
| CD630 | MACS2 | peak | 4046522 | 4046945 | 423 | + | macsForward_peak_415 | 3.56778 | CD630_34530 | sense | - | N-acetylglucosamine-6-phosphate deacetylase | | | | | | | | |
| CD630 | MACS2 | peak | 4050222 | 4050669 | 447 | + | macsForward_peak_416 | 4.02604 | CD630_34560 | sense | - | 5-formyltetrahydrofolate cyclo-ligase | | | | | | | | |
| CD630 | MACS2 | peak | 4094860 | 4094971 | 111 | - | macsReverse_peak_371 | 4.54123 | CD630_35040 | sense | pilD | Type IV prepilin peptidase, A24A family | | | | | | | | |
| CD630 | MACS2 | peak | 4096400 | 4096514 | 114 | - | macsReverse_peak_372 | 3.24639 | CD630_35050 | sense | pilT | Type IV pilus twitching motility protein | | | | | | | | |
| CD630 | MACS2 | peak | 4102100 | 4102382 | 282 | - | macsReverse_peak_373 | 3.54230 | CD630_35110 | sense | pilC | Type IV pilus secretion protein | | | | | | | | |
| CD630 | MACS2 | peak | 4120254 | 4120928 | 674 | - | macsReverse_peak_374 | 5.77277 | CD630_35260 | sense | - | ABC-type transport system, iron-family permease | | | | | | | | |
| CD630 | MACS2 | peak | 4136414 | 4136594 | 180 | - | macsReverse_peak_375 | 4.92199 | CD630_s0660 | sense | - | T-box (Met) | | | | | | | | |
| CD630 | MACS2 | peak | 4138045 | 4138314 | 269 | + | macsForward_peak_417 | 4.28798 | CD630_35430 | sense | - | Putative nicotinate phosphribosyltransferase | | | | | | | | |
| CD630 | MACS2 | peak | 4178829 | 4179302 | 473 | - | macsReverse_peak_376 | 3.50107 | CD630_35780 | sense | - | Resolvase | | | | | | | | |
| CD630 | MACS2 | peak | 4179670 | 4179814 | 144 | + | macsForward_peak_418 | 6.97665 | CD630_35790 | sense | - | Conserved hypothetical protein | | | | | | | | |
| CD630 | MACS2 | peak | 4185273 | 4185652 | 379 | - | macsReverse_peak_377 | 3.14170 | CD630_35840 | sense | - | ABC-type transport system, permease | | | | | | | | |
| CD630 | MACS2 | peak | 4206354 | 4206645 | 291 | - | macsReverse_peak_378 | 4.56362 | CD630_s0670 | sense | - | CD630_35980 | CD630_35990 | sense | rolA | Two-component sensor histidine kinase | | | | |
| CD630 | MACS2 | peak | 4207759 | 4207810 | 51 | - | macsReverse_peak_379 | 3.60270 | CD630_36000 | sense | rolB | Two-component response regulator | | | | | | | | |
| CD630 | MACS2 | peak | 4211779 | 4211921 | 142 | - | macsReverse_peak_380 | 3.52009 | CD630_36040 | sense | - | Conserved hypothetical protein | | | | | | | | |
| CD630 | MACS2 | peak | 4223877 | 4223928 | 51 | - | macsReverse_peak_381 | 3.72380 | CD630_36170 | sense | - | Conserved hypothetical protein | | | | | | | | |
| CD630 | MACS2 | peak | 4225051 | 4225424 | 373 | + | macsForward_peak_419 | 4.83428 | CD630_36180 | sense | - | Conserved hypothetical protein | | | | | | | | |
| CD630 | MACS2 | peak | 4225081 | 4225742 | 661 | - | macsReverse_peak_382 | 4.63449 | CD630_36180 | antisense | - | Conserved hypothetical protein | CD630_36190 | sense | - | Putative acyl-CoA N-acyltransferase | | | | |
| CD630 | MACS2 | peak | 4231977 | 4232711 | 734 | - | macsReverse_peak_383 | 4.10826 | CD630_36250 | sense | - | Putative signaling protein | | | | | | | | |
| CD630 | MACS2 | peak | 4232968 | 4234315 | 1347 | - | macsReverse_peak_384 | 4.13224 | CD630_36250 | sense | - | Putative signaling protein | | | | | | | | |
| CD630 | MACS2 | peak | 4245575 | 4245640 | 65 | + | macsForward_peak_420 | 5.08247 | CD630_36361 | antisense | - | Conserved hypothetical protein | | | | | | | | |
| CD630 | MACS2 | peak | 4264134 | 4264337 | 203 | + | macsForward_peak_421 | 3.56078 | CD630_36530 | sense | dnaD | DNA replication protein DnaD/DnaB | CD630_36540 | sense | dnaC | DNA replication protein DnaC | | | | |
| CD630 | MACS2 | peak | 4264744 | 4265189 | 445 | + | macsForward_peak_422 | 6.30796 | CD630_36540 | sense | dnaC | DNA replication protein DnaC | | | | | | | | |
| CD630 | MACS2 | peak | 4278957 | 4279155 | 198 | + | macsForward_peak_423 | 3.53761 | CD630_36690 | sense | - | Putative lipoprotein | | | | | | | | |
| CD630 | MACS2 | peak | 4288746 | 4288831 | 85 | - | macsReverse_peak_385 | 3.75946 | CD630_36781 | sense | - | Conserved hypothetical protein | | | | | | | | |

**Table S4. List of MACS peaks with MAGe annotation for ncRNAs**

| name | start | stop | size | strand | -log10(pval) | overlap1 | stream (ov1) | name(ov1) | product(ov1) | overlap2 | stream (ov2) | name (ov2) | product(ov2) | overlap3 | stream (ov3) | name (ov3) | product(ov3) | comment |
|---|---|---|---|---|---|---|---|---|---|---|---|---|---|---|---|---|---|---|
| macsForward_peak_1 | 17667 | 17807 | 140 | + | 4.92433 | CD630_s0010 | sense | - | T-box (Ser) | | | | | | | | | T-box (Ser) |
| macsForward_peak_4 | 174158 | 174267 | 109 | + | 5.03432 | CD630_s0050 | sense | - | SAM | | | | | | | | | SAM riboswitch |
| macsReverse_peak_1 | 174162 | 174227 | 65 | - | 6.79071 | CD630_s0050 | antisense | - | SAM | | | | | | | | | New antisense for SAM riboswitch |
| macsForward_peak_8 | 224167 | 224243 | 76 | + | 4.40573 | CD630_01690 | antisense | - | Putative drug/sodium antiporter, MATE family | | | | | | | | | New antisense CDS or CD630_01680 readthrough |
| macsReverse_peak_8 | 233846 | 233901 | 55 | - | 5.09543 | CD630_01760 | antisense | - | Putative oxidoreductase, NAD/FAD binding subunit | | | | | | | | | New antisense CDS or CD630_01770 readthrough |
| macsReverse_peak_9 | 240965 | 241018 | 53 | - | 3.80984 | Rcd2 | sense | | Antisense CDS | | | | | | | | | Antisense CDS |
| macsForward_peak_11 | 308843 | 308955 | 112 | + | 6.59405 | CD630_Cdi1_3 | sense | - | GEMM RNA motif | CD630_SQ173 | sense | - | ncRNA IGR, riboswitch c-di-GMP | | | | | GEMM RNA motif |
| macsReverse_peak_10 | 312782 | 313125 | 343 | - | 4.59330 | CD630_02490 | antisense | fliG | Flagellar motor switch protein FliG | CD630_n00080 | sense | - | Antisense CDS | CD630_02500 | antisense | fliH | Flagellar assembly protein FliH | Antisense CDS |
| macsReverse_peak_15 | 332231 | 332297 | 66 | + | 3.38938 | CD630_02720 | sense | - | Conserved hypothetical protein | CD630_n00120 | antisense | - | Antisense CDS | | | | | Antisense CDS |
| macsForward_peak_20 | 343199 | 343255 | 56 | + | 4.89175 | CD630_02820 | antisense | - | Putative PHP hydrolase | | | | | | | | | New antisense CDS |
| macsReverse_peak_13 | 391029 | 391210 | 181 | - | 5.05264 | CD630_s0080 | antisense | - | Cobalamin, CD630_03240 | | | | | | | | | New antisense for cobalamin riboswitch |
| macsForward_peak_27 | 428918 | 429100 | 182 | + | 3.79913 | CD630_03550 | antisense | int | Integrase Tn916-like, CTn1-Orf1 | | | | | | | | | New antisense CDS or CD630_03540_p1 readthrough |
| macsReverse_peak_22 | 507351 | 507420 | 69 | - | 5.64455 | . | | . | NA | | | | | | | | | 5'UTR CD630_04260 |
| macsForward_peak_28 | 507400 | 507451 | 51 | + | 6.31159 | . | | . | NA | | | | | | | | | 5'UTR CD630_04270 |
| macsReverse_peak_23 | 515402 | 515795 | 393 | - | 3.56926 | CD630_04340 | antisense | - | Putative drug/sodium antiporter, MATE family, Tn1549-like, CTn2-Orf29 | | | | | | | | | New antisense CDS |
| macsReverse_peak_24 | 519681 | 519820 | 139 | - | 3.10813 | CD630_04360 | antisense | - | Putative recombinase Tn1549-like, CTn2-Orf34 | | | | | | | | | New antisense CDS or CD630_04370 readthrough |
| macsReverse_peak_25 | 523157 | 523420 | 263 | - | 8.81823 | CD630_04401 | antisense | - | Putative phage protein | CD630_n00150 | sense | - | Antisense CDS | | | | | Antisense CDS, TA |
| macsReverse_peak_38 | 578881 | 579305 | 424 | + | 4.62291 | CD630_SQ327 | sense | - | ncRNA IGR | CD630_RNA_5 | sense | - | ykkC-yxkD | CD630_04880 | sense | - | Putative small multidrug resistan | ykkC-yxkD |
| macsForward_peak_50 | 654703 | 654813 | 110 | + | 4.08290 | CD630_SQ367 | sense | - | ncRNA IGR | | | | | | | | | ncRNA IGR |
| macsReverse_peak_39 | 683176 | 683270 | 94 | - | 3.27149 | CD630_s0160 | antisense | - | T-box (Thr) | | | | | | | | | New antisense for T-box |
| macsReverse_peak_54 | 689914 | 690240 | 326 | + | 5.39919 | CD630_05770 | antisense | - | Conserved hypothetical protein | | | | | | | | | New antisense CDS |
| macsForward_peak_65 | 729356 | 730057 | 701 | + | 4.94050 | CD630_06060_p2 | sense | - | Fragment of chloride ion channel protein (part 2) | CD630_06060_p3 | sense | - | Fragment of chloride ion channel protein (part 3) | CD630_n00220 | sense | - | ncRNA IGR | ncRNA IGR |
| macsReverse_peak_50 | 756697 | 756858 | 161 | - | 4.03510 | CD630_06330 | antisense | - | Conserved hypothetical protein | | | | | | | | | New antisense CDS |
| macsReverse_peak_51 | 761255 | 761321 | 66 | - | 4.44797 | CD630_06380 | antisense | - | Conserved hypothetical protein | | | | | | | | | New antisense CDS |
| macsForward_peak_69 | 765254 | 765407 | 153 | + | 7.27863 | CD630_SQ408 | sense | - | ncRNA IGR | | | | | | | | | ncRNA IGR |
| macsForward_peak_71 | 782205 | 782259 | 54 | + | 3.90214 | . | | . | NA | | | | | | | | | New ncRNA |
| macsReverse_peak_52 | 796645 | 796721 | 76 | - | 4.21176 | CD630_06630 | antisense | tcdA | Toxin A | | | | | | | | | New antisense CDS |
| macsForward_peak_85 | 811583 | 812005 | 422 | + | 5.82709 | CD630_06690 | sense | - | Two-component sensor histidine kinase | CD630_n00240 | sense | - | ncRNA IGR | | | | | ncRNA IGR |
| macsForward_peak_86 | 831084 | 831154 | 70 | + | 3.66276 | . | | . | NA | | | | | | | | | 5'UTR CD630_06850 |
| macsForward_peak_87 | 841749 | 841876 | 127 | + | 3.27218 | CD630_06930 | antisense | - | Transcriptional regulator, MerR family | | | | | | | | | New antisense CDS |
| macsReverse_peak_57 | 902630 | 903539 | 909 | - | 4.52273 | CD630_07400 | sense | - | Putative pyridoxal phosphate-dependent aminotransferase | CD630_SQ476 | antisense | - | Antisense CDS | | | | | Antisense CDS |
| macsReverse_peak_58 | 906407 | 906547 | 140 | - | 4.41437 | CD630_07420 | antisense | - | Putative ethanolamine transporter | | | | | | | | | New antisense CDS or CD630_07430 readthrough |
| macsForward_peak_96 | 917517 | 917679 | 162 | + | 4.98480 | CD630_s0190 | sense | - | T-box | | | | | | | | | T-box |
| macsReverse_peak_60 | 920250 | 920371 | 121 | - | 3.85358 | CD630_n00290 | sense | - | ncRNA IGR | | | | | | | | | ncRNA IGR |
| macsReverse_peak_61 | 938876 | 938999 | 123 | - | 4.14905 | CD630_07670 | antisense | srlB | PTS system, sorbitol-specific IIA component (Glucitol) | CD630_SQ495 | sense | | | | | | | Antisense CDS |
| macsForward_peak_98 | 939229 | 939939 | 710 | + | 4.19843 | CD630_SQ495 | antisense | - | Antisense CDS | CD630_07680 | sense | srlD | Sorbitol 6-phosphate 2-dehydrogenase (Glucitol) | | | | | Antisense CDS |
| macsForward_peak_101 | 955487 | 956332 | 845 | + | 5.82750 | CD630_07830 | antisense | spoIVB | Stage IV sporulation protein SpoIVB, S55 peptidase family | | | | | | | | | New antisense CDS or CD630_07820 readthrough |
| macsReverse_peak_66 | 999756 | 999823 | 67 | - | 3.63265 | CD630_08240 | antisense | - | ABC-type transport system, multidrug-family permease | | | | | | | | | New antisense CDS or CD360_SQ523 3'-end |
| macsReverse_peak_67 | 1022632 | 1022750 | 118 | - | 4.08933 | CD630_08460 | antisense | - | Conserved hypothetical protein | | | | | | | | | New antisense CDS |
| macsReverse_peak_69 | 1052169 | 1052276 | 107 | - | 6.84436 | . | | . | NA | | | | | | | | | 5'UTR CD63_08750 |
| macsReverse_peak_70 | 1052643 | 1052828 | 185 | - | 6.46464 | CD630_n00330 | sense | - | ncRNA IGR | | | | | | | | | ncRNA IGR |
| macsForward_peak_116 | 1052720 | 1052785 | 65 | + | 5.75077 | CD630_n00330 | antisense | - | ncRNA IGR | | | | | | | | | New antisense for ncRNA IGR |
| macsReverse_peak_73 | 1081434 | 1081758 | 324 | - | 5.83478 | CD630_n00340 | sense | - | Antisense CDS | CD630_08980 | antisense | - | Conserved hypothetical protein, DUF1846-family | | | | | Antisense CDS |
| macsReverse_peak_76 | 1089514 | 1090345 | 831 | - | 3.96216 | CD630_n00350 | antisense | - | ncRNA IGR | CD630_09050 | sense | - | Putative phage protein | | | | | ncRNA IGR, TA |
| macsForward_peak_123 | 1093642 | 1093787 | 145 | + | 5.06943 | . | | . | NA | | | | | CD630_n00360 | antisense | - | Antisense CDS | Antisense CDS |
| macsReverse_peak_80 | 1123859 | 1124330 | 471 | - | 7.05302 | CD630_09561 | antisense | - | Putative phage protein | CD630_09562 | antisense | - | Putative phage protein | CD630_n00370 | sense | - | Antisense CDS | TA |
| macsForward_peak_124 | 1124131 | 1124308 | 177 | + | 3.94307 | CD630_09562 | sense | - | Putative phage protein | CD630_n00370 | antisense | - | Antisense CDS | | | | | TA |
| macsReverse_peak_81 | 1124551 | 1124748 | 197 | - | 5.45183 | . | | . | NA | CD630_09563 | antisense | | | SQ606, SQ2063 | antisense | | | ncRNA IGR TA |
| macsForward_peak_125 | 1125576 | 1125711 | 135 | + | 4.27916 | CD630_n00380 | sense | - | ncRNA IGR | | | | | | | | | CRISPR |
| macsReverse_peak_83 | 1142187 | 1142564 | 377 | - | 4.83391 | CD630_Cdi1_5 | sense | - | GEMM RNA motif | CD630_09771 | antisense | - | Putative phage protein | CD630_n00390 | sense | - | Antisense CDS | TA |
| macsForward_peak_129 | 1142221 | 1142393 | 172 | + | 4.95362 | CD630_09771 | antisense | - | Putative phage protein | CD630_Cdi1_5 | sense | - | GEMM RNA motif | CD630_n00390 | antisense | - | Antisense CDS | TA |
| macsForward_peak_133 | 1152657 | 1152825 | 168 | + | 4.02122 | CD630_s0210 | sense | - | T-box (Leu) | | | | | | | | | T-box |
| macsReverse_peak_85 | 1201466 | 1201677 | 211 | - | 7.37676 | CD630_10290 | antisense | - | Putative cell wall anchored protein | | | | | | | | | New antisense CDS |
| macsForward_peak_145 | 1322368 | 1323012 | 644 | + | 4.85723 | CD630_11241 | antisense | - | Conserved hypothetical protein | CD630_SQ726 | sense | - | ncRNA IGR | CD630_n00440 | sense | - | ncRNA IGR | ncRNA IGR |
| macsReverse_peak_91 | 1322939 | 1323114 | 175 | - | 8.44287 | CD630_n00440 | antisense | - | ncRNA IGR | | | | | | | | | New antisense for ncRNA IGR |
| macsForward_peak_146 | 1364088 | 1364168 | 80 | + | 3.72587 | CD630_s0220 | sense | - | SAM | | | | | | | | | SAM riboswitch |
| macsReverse_peak_92 | 1391016 | 1391068 | 52 | - | 4.52971 | CD630_11890 | antisense | - | Putative amino acid/polyamine transporter I | | | | | | | | | New antisense CDS |
| macsReverse_peak_97 | 1433909 | 1434428 | 519 | - | 6.85680 | CD630_12331 | antisense | - | Fragment of phage protein | CD630_SQ808 | sense | - | Antisense 3prime UTR | CD630_12332 | antisense | - | Conserved hypothetical protein | TA |
| macsForward_peak_152 | 1434611 | 1434973 | 362 | + | 5.55447 | CD630_n00460 | sense | - | ncRNA IGR | | | | | | | | | CRISPR |
| macsReverse_peak_98 | 1439587 | 1439901 | 314 | - | 4.59861 | CD630_12360_p2 | antisense | - | Fragment of putative phage protein (part 2) | | | | | | | | | New antisense CDS |
| macsReverse_peak_100 | 1508381 | 1508547 | 166 | - | 3.52267 | CD630_n00470 | sense | - | ncRNA IGR | CD630_13020 | sense | - | Putative MazG-like pyrophosphohydrolase | | | | | ncRNA IGR |
| macsForward_peak_162 | 1574871 | 1575009 | 138 | + | 5.37322 | CD630_13590 | antisense | - | Putative phage protein | CD630_n00490 | sense | | Antisense CDS | | | | | Antisense CDS 3'end undefined |
| macsReverse_peak_112 | 1635215 | 1635268 | 53 | - | 4.24790 | CD630_14100 | antisense | - | Putative hydrolase, HAD superfamily, subfamily IIB | | | | | | | | | New antisense CDS or CD630_14110 readthrough |
| macsReverse_peak_117 | 1644692 | 1645084 | 392 | - | 9.35359 | CD630_14182 | antisense | - | Putative phage protein | CD630_n00500 | sense | - | Antisense CDS | CD630_n00510 | antisense | - | ncRNA IGR | TA |
| macsForward_peak_166 | 1644735 | 1644859 | 124 | + | 4.82360 | CD630_14182 | sense | - | Putative phage protein | CD630_n00500 | antisense | - | Antisense CDS | | | | | TA |
| macsForward_peak_167 | 1645203 | 1645262 | 59 | + | 3.92762 | CD630_n00510 | sense | - | ncRNA IGR | | | | | | | | | CRISPR |
| macsForward_peak_168 | 1646699 | 1646900 | 201 | + | 3.27218 | CD630_14190 | antisense | - | Putative diguanylate kinase signaling protein | | | | | | | | | New antisense CDS or CD630_n00510 readthrough |
| macsReverse_peak_121 | 1653574 | 1653887 | 313 | - | 8.60766 | CD630_14240 | sense | - | Conserved hypothetical protein | CD630_Cdi1_10 | sense | - | GEMM RNA motif | | | | | GEMM RNA motif |
| macsForward_peak_173 | 1691076 | 1691349 | 273 | + | 7.81927 | CD630_14620 | antisense | - | Transcriptional regulator, GntR family | | | | | | | | | New antisense CDS |
| macsForward_peak_179 | 1724138 | 1724253 | 115 | + | 3.74616 | CD630_s0250 | sense | - | SAM, CD630_14890 | | | | | | | | | SAM riboswitch |
| macsReverse_peak_128 | 1724145 | 1724204 | 59 | - | 4.90005 | CD630_s0250 | antisense | - | SAM, CD630_14890 | | | | | | | | | New antisense for SAM riboswitch |
| macsReverse_peak_129 | 1732484 | 1732537 | 53 | - | 3.42280 | CD630_14950 | antisense | proC | Pyrroline-5-carboxylate reductase | | | | | | | | | New antisense CDS or CD630_14960 readthrough |
| macsReverse_peak_131 | 1749652 | 1749858 | 206 | - | 4.18664 | CD630_SQ995 | antisense | - | ncRNA IGR | CD630_15111 | sense | - | Conserved hypothetical protein | | | | | ncRNA IGR |
| macsForward_peak_188 | 1756140 | 1756276 | 136 | + | 5.82053 | CD630_n00560 | sense | - | ncRNA IGR | | | | | | | | | CRISPR |
| macsForward_peak_189 | 1756616 | 1756781 | 165 | + | 4.55765 | CD630_n00560 | sense | - | ncRNA IGR | | | | | | | | | CRISPR |
| macsForward_peak_191 | 1759260 | 1759317 | 57 | + | 3.61948 | CD630_15170 | antisense | feoB | Ferrous iron transport protein B | | | | | | | | | New antisense CDS or CD630_15160 readthrough |
| macsReverse_peak_136 | 1760929 | 1761290 | 361 | - | 4.43255 | CD630_SQ1002 | sense | - | ncRNA IGR | CD630_15190 | sense | - | Putative acyltransferase | | | | | ncRNA IGR |
| macsForward_peak_193 | 1762973 | 1763089 | 116 | + | 3.63582 | CD630_SQ1005 | sense | - | ncRNA IGR | CD630_s0270 | sense | - | - | | | | | ncRNA IGR |
| macsForward_peak_197 | 1795565 | 1795712 | 147 | + | 8.30299 | CD630_s0281 | sense | - | CD630_15470 | | | | | | | | | |
| macsForward_peak_198 | 1803519 | 1803964 | 445 | + | 3.49019 | CD630_15550 | sense | - | Putative amino acid permease | CD630_SQ1038 | sense | - | ncRNA IGR | | | | | ncRNA IGR |
| macsForward_peak_199 | 1810598 | 1810788 | 190 | + | 5.37459 | CD630_s0280 | sense | - | - | | | | | | | | | |

| | | | | | | | | | | | | | | | | | | |
|---|---|---|---|---|---|---|---|---|---|---|---|---|---|---|---|---|---|---|
| macsReverse_peak_140 | 1830961 | 1831048 | 87 | - | 4.00174 | CD630_s0300 | antisense | - | T-box T-box | | | | | | | | | New antisense for T-box |
| macsForward_peak_201 | 1831121 | 1831439 | 318 | + | 4.17452 | CD630_s0300 | sense | - | T-box T-box | CD630_s0310 | sense | - | - | | | | | T-box; T-box |
| macsForward_peak_205 | 1851792 | 1851906 | 114 | + | 4.11854 | CD630_15960 | antisense | map2 | Methionine aminopeptidase Map2 (MAP) (Peptidase M) | SQ1076 | sense | | Antisense 3'UTR | | | | | 3' end undefined |
| macsForward_peak_206 | 1854581 | 1854682 | 101 | + | 8.15991 | CD630_s0320 | sense | - | - | | | | | | | | | |
| macsReverse_peak_142 | 1915892 | 1916440 | 548 | - | 4.28395 | CD630_n00590 | sense | - | ncRNA IGR | CD630_16520 | sense | - | Putative tellurium resistance protein | | | | | ncRNA IGR |
| macsReverse_peak_143 | 1916697 | 1918516 | 1819 | - | 4.37658 | CD630_s0330 | sense | - | SAM | CD630_16540 | sense | - | Putative lipoate-protein ligase | CD630_16530 | sense | - | Putative Probable D-methionine-l | SAM riboswitch |
| macsForward_peak_213 | 1918864 | 1919290 | 426 | + | 5.61064 | CD630_s0340 | sense | - | Lysine | CD630_16550 | sense | - | Putative Na+/H+ antiporter NhaC-like | | | | | Lysine riboswitch |
| macsReverse_peak_144 | 1918882 | 1918942 | 60 | - | 3.65576 | CD630_s0340 | antisense | - | Lysine | | | | | | | | | New antisense for Lysine riboswitch |
| macsForward_peak_219 | 1935455 | 1935856 | 401 | + | 5.31146 | CD630_n00600 | sense | - | ncRNA IGR | | | | | | | | | CRISPR |
| macsForward_peak_221 | 1943259 | 1943683 | 424 | + | 5.82838 | CD630_n00620 | sense | - | ncRNA IGR | CD630_16671 | antisense | - | Conserved hypothetical protein | | | | | ncRNA IGR |
| macsForward_peak_223 | 1951890 | 1952028 | 138 | + | 4.50056 | CD630_16760 | antisense | pcp | Pyrrolidone-carboxylate peptidase | | | | | | | | | New antisense CDS or CD630_1675 readthrough |
| macsForward_peak_224 | 1966006 | 1966491 | 485 | + | 3.99069 | CD630_16940 | antisense | - | Conserved hypothetical protein | CD630_16950 | antisense | - | Putative symporter protein | | | | | New antisense CDS or CD630_1692_1693 readthrough |
| macsReverse_peak_148 | 1972996 | 1973347 | 351 | - | 6.65922 | CD630_17000 | sense | ribD | Riboflavin biosynthesis protein ribD (Includes: Diaminohydrox | CD630_s0360 | sense | - | FMN | | | | | FMN riboswitch |
| macsForward_peak_225 | 1973228 | 1973330 | 102 | + | 7.40079 | CD630_s0360 | antisense | - | FMN | | | | | | | | | New antisense for FMN riboswitch |
| macsForward_peak_226 | 1976200 | 1976300 | 100 | + | 10.21495 | CD630_s0370 | sense | - | - | | | | | | | | | |
| macsReverse_peak_153 | 2030332 | 2030435 | 103 | - | 5.63247 | CD630_17510 | antisense | cwp13 | Cell surface protein | | | | | | | | | New antisense CDS or CD630_17511 readthrough |
| macsForward_peak_236 | 2036635 | 2036777 | 142 | + | 8.19045 | . | | . | NA | | | | | | | | | 5'UTR CD630_17590 |
| macsForward_peak_239 | 2055318 | 2055684 | 366 | + | 4.73397 | CD630_17730 | sense | - | Conserved hypothetical protein | CD630_s0390 | sense | - | T-box | | | | | T-box |
| macsForward_peak_241 | 2066674 | 2066953 | 279 | + | 4.51086 | CD630_s0400 | sense | - | T-box (Arg) | | | | | | | | | T-box (Arg) |
| macsReverse_peak_162 | 2106156 | 2106235 | 79 | - | 4.73561 | CD630_18200_p1 | antisense | ade | Adenine deaminase (Disrupted IStron) | | | | | | | | | New antisense CDS |
| macsReverse_peak_163 | 2171872 | 2171929 | 57 | - | 6.94246 | . | | . | NA | | | | | | | | | 5'UTR CD630_18700 |
| macsReverse_peak_165 | 2198056 | 2198484 | 428 | - | 5.32127 | CD630_18930 | sense | - | Putative oligonucleotide binding regulator | CD630_n00650 | sense | - | Antisense CDS | | | | | Antisense CDS |
| macsReverse_peak_267 | 2198990 | 2199467 | 477 | + | 4.09988 | CD630_18940 | sense | - | Conserved hypothetical protein | Rcd1 | sense | | ncRNA IGR | | | | | ncRNA IGR |
| macsReverse_peak_166 | 2199393 | 2199457 | 64 | - | 6.83517 | Rcd1 | antisense | | ncRNA IGR | | | | | | | | | New antisense for ncRNA IGR |
| macsReverse_peak_167 | 2199888 | 2199957 | 69 | - | 5.67942 | CD630_18950 | antisense | - | Conserved hypothetical protein | | | | | | | | | New antisense CDS |
| macsReverse_peak_274 | 2257786 | 2258102 | 316 | + | 4.51902 | CD630_SQ1339 | antisense | - | ncRNA IGR | CD630_19580 | sense | - | Putative 4Fe-4S binding protein, putative ferredoxin, putative dehydrogenase, putative reductase | | | | | ncRNA IGR |
| macsReverse_peak_276 | 2286198 | 2286268 | 70 | + | 6.52781 | CD630_n00680 | sense | - | ncRNA IGR | Rcd5 | sense | | ncRNA IGR | | | | | GEMM RNA motif |
| macsReverse_peak_277 | 2286519 | 2286607 | 88 | + | 3.99438 | CD630_19810 | antisense | - | Transcriptional regulator, hxlR family | | | | | | | | | RCd5 3' end |
| macsReverse_peak_280 | 2297527 | 2297813 | 286 | + | 5.79994 | CD630_Cdi1_8 | sense | - | GEMM RNA motif | CD630_19903 | sense | - | Conserved hypothetical protein | | | | | GEMM RNA motif |
| macsReverse_peak_179 | 2298419 | 2298860 | 441 | - | 5.96526 | CD630_19904 | sense | - | Fragment of conserved hypothetical protein | CD630_n00690 | sense | - | ncRNA IGR | | | | | CRISPR |
| macsReverse_peak_182 | 2327638 | 2327750 | 112 | - | 4.53652 | CD630_20170 | antisense | - | Transcriptional regulator, HTH-type | | | | | | | | | Double stranded region |
| macsReverse_peak_184 | 2340724 | 2340805 | 81 | - | 4.38183 | CD630_20290 | sense | - | Putative lipoprotein | CD630_n00700 | antisense | - | Antisense CDS | | | | | Antisense CDS |
| macsReverse_peak_185 | 2347097 | 2347397 | 300 | - | 5.67265 | CD630_s0450 | sense | - | T-box (Arg) | | | | | | | | | T-box (Arg) |
| macsReverse_peak_186 | 2355898 | 2356092 | 194 | - | 3.23150 | CD630_20430 | antisense | - | Putative hydrolase | | | | | | | | | New antisense CDS |
| macsReverse_peak_188 | 2368910 | 2369125 | 215 | - | 5.74527 | CD630_20541 | sense | - | Conserved hypothetical protein | CD630_s0460 | sense | - | Lysine | | | | | Lysine riboswitch |
| macsReverse_peak_200 | 2435817 | 2436149 | 332 | - | 5.35019 | CD630_21070 | sense | - | Xanthine/uracil/thiamine/ascorbate permease family proteir | CD630_s0470 | sense | - | Purine | | | | | Purine riboswitch |
| macsForward_peak_290 | 2451389 | 2451525 | 136 | + | 4.87999 | CD630_s0480 | sense | - | T-box (Thr) | | | | | | | | | T-box (Thr) |
| macsForward_peak_293 | 2462344 | 2462577 | 233 | + | 3.11884 | CD630_21260 | sense | - | Putative membrane protein | CD630_n00740 | antisense | - | Antisense CDS | | | | | Antisense CDS |
| macsReverse_peak_215 | 2524595 | 2524806 | 211 | - | 4.73824 | CD630_21800 | antisense | - | Putative DEAD-like helicase | | | | | | | | | New antisense CDS or CD630_21810 readthrough |
| macsReverse_peak_228 | 2640444 | 2640515 | 71 | - | 4.28971 | CD630_RNA_17 | antisense | - | Glycine | CD630_RNA_15 | antisense | - | Glycine | | | | | New antisense for glycine riboswitch |
| macsForward_peak_312 | 2661280 | 2661357 | 77 | + | 3.58866 | CD630_SQ1633 | antisense | - | Antisense 3prime UTR | CD630_22951 | antisense | - | Fragment of putative cell surface protein (C-terminal region) | | | | | Antisense 3'UTR |
| macsReverse_peak_231 | 2663798 | 2663903 | 105 | - | 5.51210 | CD630_n00790 | sense | - | ncRNA IGR | | | | | | | | | CRISPR |
| macsForward_peak_313 | 2665226 | 2665461 | 235 | + | 8.65138 | CD630_SQ1641 | sense | - | Antisense CDS | CD630_22991 | antisense | - | Putative phage protein | CD630_SQ1642 | sense | - | ncRNA IGR | TA |
| macsReverse_peak_232 | 2665311 | 2665419 | 108 | - | 6.03728 | CD630_SQ1641 | antisense | - | Antisense CDS | CD630_22991 | sense | - | Putative phage protein | CD630_SQ1642 | antisense | - | ncRNA IGR | TA |
| macsForward_peak_315 | 2671225 | 2671566 | 341 | + | 5.23027 | CD630_23070 | sense | - | Putative membrane protein | CD630_23080 | sense | - | Transcriptional regulator, HTH-type | CD630_SQ1653 | antisense | - | Antisense 5prime UTR | Antisense 5'UTR |
| macsForward_peak_316 | 2671843 | 2672139 | 296 | + | 4.25622 | CD630_SQ1656 | sense | - | ncRNA IGR | CD630_Cdi1_9 | sense | - | GEMM RNA motif | CD630_23090 | sense | - | Conserved hypothetical protein | GEMM RNA motif |
| macsReverse_peak_234 | 2671845 | 2672018 | 173 | - | 3.76346 | CD630_SQ1656 | antisense | - | ncRNA IGR | CD630_Cdi1_9 | antisense | - | GEMM RNA motif | CD630_23090 | antisense | - | Conserved hypothetical protein | New antisense CDS |
| macsReverse_peak_237 | 2694659 | 2694733 | 74 | - | 3.25870 | CD630_s0500 | sense | - | Purine | | | | | | | | | Purine riboswitch |
| macsForward_peak_319 | 2720426 | 2720970 | 544 | + | 4.11757 | CD630_23510 | antisense | grdB | Glycine reductase complex component B gamma subunit (sel | CD630_n00830 | sense | - | Antisense CDS | | | | | Antisense CDS |
| macsForward_peak_320 | 2725730 | 2725805 | 75 | + | 3.63470 | CD630_RNA_16 | antisense | - | Glycine | | | | | | | | | New antisense for glycine riboswitch |
| macsReverse_peak_239 | 2741517 | 2741583 | 66 | - | 4.78770 | CD630_23750 | antisense | - | Conserved hypothetical protein | | | | | | | | | New antisense CDS |
| macsForward_peak_322 | 2758242 | 2758500 | 258 | + | 4.24581 | CD630_23880 | antisense | - | Putative peptidoglycan-binding/hydrolysing protein | | | | | | | | | New antisense CDS or readthrough CD630_23870 |
| macsForward_peak_323 | 2780618 | 2780709 | 91 | + | 5.00641 | . | | . | NA | | | | | | | | | 3'UTR CD630_24080 |
| macsReverse_peak_257 | 2886131 | 2886227 | 96 | - | 3.37631 | CD630_s0510 | sense | - | T-box (Arg) | | | | | | | | | T-box (Arg) |
| macsReverse_peak_258 | 2891489 | 2891549 | 60 | - | 7.42886 | CD630_RNA_7 | sense | - | yybP-ykoY | | | | | | | | | yybP-ykoY |
| macsForward_peak_330 | 2907274 | 2907386 | 112 | + | 5.53585 | CD630_n00850 | sense | - | Antisense CDS | CD630_Cdi1_7 | sense | - | GEMM RNA motif | | | | | GEMM RNA motif |
| macsForward_peak_331 | 2908005 | 2908417 | 412 | + | 6.97567 | CD630_n00860 | antisense | - | ncRNA IGR | Rcd8 | sense | | Antisense CDS | CD630_25171 | antisense | - | Putative phage protein | TA |
| macsReverse_peak_271 | 3001828 | 3001911 | 83 | - | 7.25310 | . | | . | NA | | | | | | | | | 5'UTR CD630_25930 |
| macsReverse_peak_274 | 3015664 | 3015745 | 81 | - | 7.09360 | . | | . | NA | | | | | | | | | 3'UTR CD630_26100 |
| macsForward_peak_339 | 3015693 | 3015816 | 123 | + | 6.57474 | . | | . | NA | | | | | | | | | ncRNA IGR |
| macsForward_peak_343 | 3064759 | 3065092 | 333 | + | 5.59616 | CD630_26560 | antisense | spoVD | Stage V sporulation protein D (Sporulation-specific penicillin-binding protein) | | | | | | | | | New antisense CDS |
| macsReverse_peak_282 | 3137395 | 3137537 | 142 | - | 4.43720 | CD630_n00910 | sense | - | Antisense CDS | CD630_27120 | sense | - | Conserved hypothetical protein | | | | | Antisense CDS |
| macsReverse_peak_290 | 3196739 | 3196812 | 73 | - | 5.79130 | CD630_n00930 | sense | - | ncRNA IGR | | | | | | | | | ncRNA IGR |
| macsReverse_peak_297 | 3306694 | 3307153 | 459 | - | 3.99184 | CD630_Cdi2_3 | sense | - | c-di-GMP-II | CD630_28320 | sense | - | Putative cation efflux protein | | | | | c-di-GMP-II |
| macsForward_peak_365 | 3380080 | 3380348 | 268 | + | 5.31318 | CD630_n00980 | sense | - | Antisense CDS | CD630_Cdi1_4 | sense | - | GEMM RNA motif | CD630_28890 | antisense | - | Putative phage protein | GEMM RNA motif |
| macsReverse_peak_307 | 3380250 | 3380421 | 171 | - | 7.52469 | CD630_n00980 | antisense | - | Antisense CDS | CD630_Cdi1_4 | antisense | - | GEMM RNA motif | CD630_28890 | sense | - | Putative phage protein | TA |
| macsForward_peak_366 | 3380757 | 3380927 | 170 | + | 4.12438 | . | | . | NA | | | | | | | | | New antisense CDS |
| macsReverse_peak_310 | 3396937 | 3397064 | 127 | - | 3.72051 | CD630_n00990 | sense | - | ncRNA IGR | | | | | | | | | CRISPR |
| macsForward_peak_367 | 3397893 | 3398087 | 194 | + | 5.45665 | CD630_n00990 | antisense | - | ncRNA IGR | | | | | CD630_29072 | antisense | | | TA |
| macsForward_peak_368 | 3398317 | 3398663 | 346 | + | 8.96925 | Rcd9 | sense | | Antisense CDS | CD630_29071 | antisense | - | Putative phage protein | | | | | TA |
| macsReverse_peak_316 | 3456571 | 3457171 | 600 | - | 4.59146 | CD630_n01010 | sense | - | ncRNA IGR | CD630_29740 | antisense | - | Conserved hypothetical protein | CD630_29750 | sense | - | Putative CRISPR-associated Cas2 | CRISPR |
| macsReverse_peak_333 | 3562259 | 3562560 | 301 | - | 3.41240 | CD630_30660 | antisense | xylR | Transcriptional regulator, Xylose repressor | CD630_SQ2179 | sense | | | | | | | 3'-end undefined |
| macsReverse_peak_336 | 3607716 | 3607953 | 237 | - | 8.13716 | CD630_s0590 | sense | - | Lysine | CD630_31001 | antisense | - | Fragment of artefact | | | | | Lysine riboswitch |
| macsReverse_peak_338 | 3631128 | 3631193 | 65 | - | 6.84967 | CD630_s0591 | sense | - | CD630_31170 | | | | | | | | | |
| macsForward_peak_377 | 3635687 | 3636047 | 360 | + | 4.01108 | CD630_31220 | antisense | - | Transcriptional regulator, ArsR family | CD630_31230 | antisense | - | Putative nitroreductase | | | | | New antisense CDS or CD630_31210 readthrough |
| macsReverse_peak_342 | 3659601 | 3659656 | 55 | - | 5.17071 | . | | . | NA | | | | | | | | | 5'UTR CD630_31380 |
| macsForward_peak_389 | 3687415 | 3687733 | 318 | + | 11.25295 | CD630_s0600 | antisense | - | tmRNA | | | | | | | | | New antisense for tmRNA |
| macsForward_peak_392 | 3731537 | 3731706 | 169 | + | 3.97522 | CD630_31880 | sense | - | Conserved hypothetical protein | CD630_n01060 | sense | - | Antisense CDS | | | | | Antisense CDS |
| macsForward_peak_395 | 3752164 | 3753101 | 937 | + | 5.73797 | CD630_32050 | sense | - | Putative nitroreductase | CD630_32060 | sense | - | Putative membrane protein | CD630_n01080 | antisense | - | Antisense CDS | Antisense CDS |
| macsReverse_peak_348 | 3762706 | 3762768 | 62 | - | 3.57422 | . | | . | NA | | | | | | | | | 5'UTR CD630_32160 |
| macsReverse_peak_349 | 3773076 | 3773176 | 100 | - | 5.84172 | CD630_s0610 | antisense | - | Lysine | | | | | | | | | New antisense for lysine riboswitch |
| macsForward_peak_398 | 3788915 | 3789729 | 814 | + | 4.47959 | CD630_32360 | antisense | - | Putative membrane protein | | | | | | | | | New antisense CDS |
| macsReverse_peak_350 | 3801097 | 3801335 | 238 | - | 3.76949 | . | | . | NA | | | | | | | | | ncRNA promoter upstream Cdi2_1 |
| macsReverse_peak_352 | 3814035 | 3814189 | 154 | - | 4.95025 | CD630_s0631 | sense | - | T-box, CD630_32560 | CD630_32561 | antisense | - | Conserved hypothetical protein | | | | | T-box |

| | | | | | | | | | | | | | | | | | | |
|---|---|---|---|---|---|---|---|---|---|---|---|---|---|---|---|---|---|---|
| macsReverse_peak_353 | 3826624 | 3826684 | 60 | - | 6.72203 | CD630_n01090 | sense | | | CD630_Cdi2_2 | sense | - | c-di-GMP-II | | | | | c-di-GMP-II |
| macsForward_peak_402 | 3861679 | 3863304 | 1625 | + | 5.24925 | CD630_s0640 | sense | - | FMN | CD630_32990 | sense | - | Transporter, Major Facilitator Superfamily (MFS) | | | | | FMN riboswitch |
| macsReverse_peak_359 | 3920168 | 3921107 | 939 | - | 3.95778 | CD630_33560 | antisense | - | Transcriptional regulator, hxlR family | CD630_33570 | sense | - | Putative protease/amidase | CD630_33580 | sense | - | Fragment of C-terminal transcrip | New antisense CDS or CD630_33570 readthrough |
| macsForward_peak_407 | 3922102 | 3922277 | 175 | + | 6.68432 | CD630_s0641 | sense | - | CD630_33590 | CD630_33581 | antisense | - | Conserved hypothetical protein | | | | | uup |
| macsReverse_peak_361 | 3923637 | 3923759 | 122 | - | 3.93606 | CD630_33590 | antisense | - | ABC-type transport system, ATP-binding protein | | | | | | | | | New antisense CDS |
| macsReverse_peak_362 | 3924240 | 3924607 | 367 | - | 3.73196 | CD630_33600 | sense | - | Two-component response regulator | CD630_SQ2424 | antisense | - | Antisense CDS | | | | | Antisense CDS |
| macsReverse_peak_364 | 3929833 | 3929909 | 76 | - | 4.88427 | . | | . | NA | | | | | | | | | 5'UTR CD630_33640 |
| macsReverse_peak_365 | 3935555 | 3935815 | 260 | - | 6.00600 | CD630_s0642 | sense | - | CD630_33681 | | | | | | | | | |
| macsForward_peak_410 | 3936262 | 3936555 | 293 | + | 5.77720 | CD630_SQ2429 | sense | - | ncRNA IGR | CD630_Cdi1_11 | sense | - | GEMM RNA motif | CD630_33682 | sense | - | Conserved hypothetical protein | GEMM RNA motif |
| macsReverse_peak_375 | 4136414 | 4136594 | 180 | - | 4.92199 | CD630_s0660 | sense | - | T-box (Met) | | | | | | | | | T-box (Met) |
| macsReverse_peak_378 | 4206354 | 4206645 | 291 | - | 4.56362 | CD630_s0670 | sense | - | CD630_35980 | CD630_35990 | sense | rolA | Two-component sensor histidine kinase | | | | | |
| macsReverse_peak_382 | 4225081 | 4225742 | 661 | - | 4.63449 | CD630_36180 | antisense | - | Conserved hypothetical protein | CD630_36190 | sense | - | Putative acyl-CoA N-acyltransferase | | | | | double stranded region |
| macsForward_peak_420 | 4245575 | 4245640 | 65 | + | 5.08247 | CD630_36361 | antisense | - | Conserved hypothetical protein | | | | | | | | | New antisense CDS |

**Table S5. Transcriptome data comparison with RIP-seq**

| RIP-seq peak name | Fold_enrichment | -log10 (Pval) | Gene name | 630pAS-hfq/630p expression ratio micro-array* | Product | Category | Alternative gene name | Comment |
|---|---|---|---|---|---|---|---|---|
| macsForward_peak_5 | 3.10785 | 4.06352 | CD630_01560 | 2.54 | Putative membrane protein | Cell Wall | | |
| macsForward_peak_26 | 2.28523 | 3.79830 | CD630_03260 | 0.45 | ABC-type transport system, cobalt-specific permease | Membrane Transport | *cbiQ* | |
| macsForward_peak_26 | 2.28523 | 3.79830 | CD630_03270 | 0.47 | ABC-type transport system, cobalt-specific ATP-binding protein | Membrane Transport | *cbiO* | |
| macsReverse_peak_28 | 3.09086 | 3.73674 | CD630_04710 | 2.01 | Transcriptional regulator, Penicillinase repressor | Regulations | *blaI* | |
| macsForward_peak_49 | 2.64967 | 3.89852 | CD630_05490 | 2.63 | Conserved hypothetical protein | Unknown function | | |
| macsForward_peak_55 | 2.60589 | 3.36528 | CD630_05820 | 2.16 | Putative pyruvate phosphate dikinase, PEP/pyruvate-binding | Carbon Metabolism | | |
| macsForward_peak_56 | 2.74078 | 3.75661 | CD630_05820 | 2.16 | Putative pyruvate phosphate dikinase, PEP/pyruvate-binding | | | |
| macsReverse_peak_64 | 2.93320 | 4.87819 | CD630_08070 | 0.37 | Conserved hypothetical protein | Unknown function | | |
| macsReverse_peak_64 | 2.93320 | 4.87819 | CD630_08080 | 0.4 | Conserved hypothetical protein | Unknown function | | |
| macsForward_peak_112 | 1.92522 | 2.98328 | CD630_08530 | 0.39 | ABC-type transport system, oligopeptide-family permease | Membrane Transport | *oppB* | |
| macsReverse_peak_93 | 3.25684 | 4.36078 | CD630_12110 | 0.46 | Putative N-acetyltransferase | Cell Factor and Growth | | |
| macsReverse_peak_101 | 2.96604 | 4.09832 | CD630_13530 | 0.49 | Putative phosphomethylpyrimidine kinase | Nucleic Acid Metabolism | | |
| macsReverse_peak_135 | 3.13620 | 4.05364 | CD630_15170 | 3.2 | Ferrous iron transport protein B | Membrane Transport | *feoB* | SigK regulon |
| macsForward_peak_195 | 3.15854 | 5.29780 | CD630_15431 | 0.23 | Conserved hypothetical protein | Unknown function | | |
| macsForward_peak_204 | 2.01793 | 3.17664 | CD630_15900 | 2.18 | Putative membrane protein | Cell Wall | | |
| macsForward_peak_209 | 3.24371 | 4.30055 | CD630_16160 | 0.42 | Putative diguanylate kinase signaling protein | Regulations | | |
| macsForward_peak_210 | 3.91509 | 6.11801 | CD630_16160 | 0.42 | Putative diguanylate kinase signaling protein | | | |
| macsReverse_peak_142 | 2.55917 | 4.28395 | CD630_16520 | 0.31 | Putative tellurium resistance protein | Stress | | |
| macsReverse_peak_143 | 3.26114 | 4.37658 | CD630_16540 | 2.09 | Putative lipoate-protein ligase | Cell Wall | | |
| macsForward_peak_253 | 3.79465 | 6.86047 | CD630_18311 | 0.43 | Conserved hypothetical protein | Unknown function | | |
| macsReverse_peak_168 | 2.67116 | 3.57243 | CD630_18970 | 2.62 | Conserved hypothetical protein | Unknown function | | SigK regulon |
| macsForward_peak_271 | 2.99513 | 3.82847 | CD630_19090 | 2.06 | Ethanolamine utilization protein, GTPase family | Carbon Metabolism | *eutP* | |
| macsReverse_peak_170 | 2.06645 | 3.13922 | CD630_19430 | 0.38 | Conserved hypothetical protein | Unknown function | | |
| macsReverse_peak_194 | 4.48342 | 8.12847 | CD630_20980 | 0.42 | Conserved hypothetical protein | Unknown function | | |
| macsReverse_peak_240 | 2.94676 | 4.54307 | CD630_23760 | 0.5 | Putative membrane protein | Cell Wall | | |
| macsReverse_peak_241 | 2.73772 | 4.20865 | CD630_23760 | 0.5 | Putative membrane protein | | | |
| macsReverse_peak_242 | 2.47316 | 3.25181 | CD630_23760 | 0.5 | Putative membrane protein | | | |
| macsReverse_peak_272 | 2.25104 | 3.53584 | CD630_26000 | 3.81 | Carbon starvation protein, CstA | Carbon Metabolism | *cstA* | |
| macsForward_peak_344 | 3.88390 | 6.44103 | CD630_26730 | 0.46 | ABC-type transport system, oligopeptide-family permease protein | Membrane Transport | *appB* | |
| macsForward_peak_345 | 3.12352 | 4.50438 | CD630_26730 | 0.46 | ABC-type transport system, oligopeptide-family permease protein | | | |
| macsReverse_peak_280 | 2.51982 | 3.62689 | CD630_26880 | 36.89 | Small, acid-soluble spore protein alpha | Sporulation | *sspA* | |
| macsReverse_peak_283 | 2.32558 | 3.42516 | CD630_27200 | 2.2 | Putative transporter | Membrane Transport | | SigK regulon |
| macsReverse_peak_300 | 2.50890 | 3.08671 | CD630_28520 | 0.47 | D-alanyl transferase DltB, MBOAT family | Cell Wall | *dltB* | |
| macsReverse_peak_301 | 3.09816 | 3.81485 | CD630_28520 | 0.47 | D-alanyl transferase DltB, MBOAT family | | | |
| macsForward_peak_364 | 2.81772 | 3.41991 | CD630_28870 | 2.08 | Putative diguanylate kinase signaling protein | Regulations | | |
| macsForward_peak_364 | 2.81772 | 3.41991 | CD630_28870 | 2.08 | Putative diguanylate kinase signaling protein | | | |
| macsReverse_peak_311 | 2.45690 | 3.70174 | CD630_29270 | 2.03 | Transcriptional regulator, Phage-type | Regulations | | |
| macsReverse_peak_314 | 3.43492 | 4.62758 | CD630_29670 | 2.67 | Dipicolinate synthase subunit B | Sporulation | *spoVFB* | SigK regulon |
| macsReverse_peak_326 | 2.44957 | 3.28343 | CD630_30130 | 0.27 | PTS system, mannose-specfic IIC component | Membrane Transport | | |
| macsReverse_peak_326 | 2.44957 | 3.28343 | CD630_30140 | 0.38 | PTS system, mannose-specfic IIB component | Membrane Transport | | |
| macsReverse_peak_328 | 3.22288 | 4.46345 | CD630_30290 | 0.29 | Bifunctional protein: cystathionine beta-lyase / repressor | Membrane Transport | *malY* | |
| macsReverse_peak_329 | 3.04962 | 4.56059 | CD630_30310 | 0.15 | Transcription antiterminator, PTS operon regulator | Membrane Transport | | |
| macsForward_peak_375 | 2.02620 | 3.13236 | CD630_30730 | 0.08 | Putative membrane protein | Cell Wall | | |
| macsReverse_peak_341 | 3.24092 | 4.86402 | CD630_31360 | 0.35 | 6-phospho-beta-glucosidase | Membrane Transport | *bglA* | |
| macsForward_peak_385 | 2.90670 | 4.33960 | CD630_31500 | 0.48 | Conserved hypothetical protein | Unknown function | | |
| macsForward_peak_386 | 2.57350 | 3.66435 | CD630_31500 | 0.48 | Conserved hypothetical protein | | | |
| macsForward_peak_387 | 2.77981 | 4.90207 | CD630_31500 | 0.48 | Conserved hypothetical protein | | | |
| macsReverse_peak_60 | 3.15093 | 3.85358 | CD630_n00290 | 9.08 | ncRNA IGR | | | |
| macsForward_peak_152 | 3.46743 | 5.55447 | CD630_n00460 | 0.43 | ncRNA IGR | CRISPR 6 | | |
| macsForward_peak_188 | 3.75094 | 5.82053 | CD630_n00560 | 0.75 | ncRNA IGR | CRISPR 8 | | |
| macsForward_peak_189 | 3.33940 | 4.55765 | CD630_n00560 | 0.75 | ncRNA IGR | | | |
| macsForward_peak_221 | 3.17161 | 5.82838 | CD630_n00620 | 1.52 | ncRNA IGR | | | |
| macsReverse_peak_179 | 3.73919 | 5.96526 | CD630_n00690 | 0.55 | ncRNA IGR | CRISPR 10 | | |
| macsForward_peak_38 | 3.11295 | 4.62291 | CD630_RNA_5 | 0.53 | ykkC-yxkD | | | |
| macsForward_peak_133 | 2.89201 | 4.02122 | CD630_s0210 | 1.85 | T-box (Leu) | | | |
| macsForward_peak_201 | 2.41417 | 4.17452 | CD630_s0300 | 1.61 | T-box T-box | | | |
| macsReverse_peak_185 | 3.43732 | 5.67265 | CD630_s0450 | 1.55 | T-box (Arg) | | | |
| macsReverse_peak_188 | 3.10318 | 5.74527 | CD630_s0460 | 2.87 | Lysine | | | |
| macsReverse_peak_338 | 4.28465 | 6.84967 | CD630_s0591 | 0.40 | CD630_31170 | | | |
| macsReverse_peak_378 | 2.59678 | 4.56362 | CD630_s0670 | 1.85 | CD630_35980 | | | |
| macsForward_peak_11 | 3.56775 | 6.59405 | CD630_SQ173 | 2.01 | ncRNA IGR, riboswitch c-di-GMP | | | |
| macsReverse_peak_95 | 2.72052 | 3.08122 | CD630_12300_p2 | 4.12 | RNA polymerase sigma-K factor SigK (Part 2) | Sporulation | *sigK* | SigK regulon |
| macsReverse_peak_164 | 2.72993 | 3.30688 | CD630_18910_p2 | 3.28 | Fragment of ABC-type transport system, substrate-binding protein | Membrane Transport | | |

* Transcriptome data from Boudry et al. J. Bacteriol. 2014. 196: 3234-3248.

**Table S6. Pathway clustering for mRNAs peaks detected by RIP-seq from KEGG**

| Cluster | Enrichment_Score | Classification | Gene_number | Category | Total gene_number per category |
|---|---|---|---|---|---|
| 28 | 0.018586963841872393 | Cell mobility, Flagellar assembly | 4 | Cell motility | 4 |
| 17 | 0.35461545126523714 | Cell wall hydrolase/autolysin, catalytic | 3 | Cell wall | 3 |
| 1 | 9.983922294624588 | Membrane protein | 199 | Membrane protein | 199 |
| 26 | 0.032132329533603496 | ABC transporter | 27 | Membrane transport | 45 |
| 4 | 1.9137264497182604 | Drug transmembrane transporter activity | 7 | Membrane transport | |
| 27 | 0.02336918665624426 | Phosphotransferase system | 6 | Membrane transport | |
| 15 | 0.4285048522166124 | Major facilitator superfamily | 5 | Membrane transport | |
| 3 | 2.464358789115692 | N-acetyltransferase activity | 10 | Metabolism | 39 |
| 12 | 0.7347958176596812 | Aminotransferase | 7 | Metabolism | |
| 14 | 0.4448749282704527 | Amino acids biosynthesis | 6 | Metabolism | |
| 24 | 0.0963974490591501 | Iron-sulfur cluster | 6 | Metabolism | |
| 22 | 0.21885504802629352 | Sugar isomerase | 4 | Metabolism | |
| 11 | 0.8093382747290931 | Lipid metabolism | 3 | Metabolism | |
| 25 | 0.06713471006721808 | Radical SAM family protein | 3 | Metabolism | |
| 20 | 0.2854292113006537 | Integrase/recombinase | 3 | Mobile Element | 3 |
| 6 | 1.368410723905778 | DNA-transcription regulation | 40 | Regulations & signaling | 155 |
| 23 | 0.17540864018495575 | Transcription regulator HTH, MerR | 23 | Regulations & signaling | |
| 5 | 1.6356471184030128 | Signal transduction histidine kinase-related protein | 17 | Regulations & signaling | |
| 2 | 4.3638636752239 | Diguanylate kinase signaling protein | 15 | Regulations & signaling | |
| 7 | 1.2792201034015007 | DNA binding protein | 15 | Regulations & signaling | |
| 10 | 0.8400776125092168 | Signal transduction response regulator | 13 | Regulations & signaling | |
| 18 | 0.30581699608824675 | Transcriptional regulator | 6 | Regulations & signaling | |
| 8 | 1.2189357219735004 | PAS-associated protein | 5 | Regulations & signaling | |
| 9 | 1.0424263968301977 | PAS fold-protein | 5 | Regulations & signaling | |
| 19 | 0.29221482242346075 | Transcription antiterminator | 5 | Regulations & signaling | |
| 13 | 0.45282455932546456 | RmlC-like domain protein | 4 | Regulations & signaling | |
| 16 | 0.3902258513508365 | Beta-lactams repressor | 4 | Regulations & signaling | |
| 21 | 0.23175952730041166 | Tetratricopeptide repeat-domain protein | 3 | Regulations & signaling | |
| 0 | NA | Hypothetical protein | 113 | Unknown function | 113 |

**Table S7. RIP-seq data validation by qRT-PCR**

| | Control | | 3x FLAG | | **Fold Change** |
|---|---|---|---|---|---|
| | Mean | SD | Mean | SD | |
| RCd1 | 1.36 | 0.31 | 100.91 | 36.46 | **74.03** |
| n00290 | 0.63 | 0.21 | 18.79 | 11.84 | **30.05** |
| SQ1828 | 0.58 | 0.18 | 13.78 | 1.91 | **23.83** |
| RCd2 | 0.40 | 0.08 | 13.74 | 5.26 | **33.93** |
| n00460 | 0.32 | 0.06 | 19.36 | 12.98 | **60.75** |
| n00560 | 0.12 | 0.04 | 1.79 | 0.84 | **14.54** |
| Cdi1_8 | 1.83 | 0.99 | 1496.84 | 538.39 | **819.96** |
| New IGR *CD0655* | 0.22 | 0.07 | 3.37 | 1.70 | **15.60** |
| New AS *CD2890* | 0.20 | 0.07 | 6.41 | 5.80 | **31.89** |
| *tcdA* | 0.43 | 0.18 | 13.16 | 9.38 | **30.73** |
| *tcdC* | 0.39 | 0.20 | 1.30 | 0.30 | **3.38** |
| *pilA* | 0.20 | 0.06 | 1.05 | 0.66 | **5.29** |

The data was normalized with the *dnaF* gene encoding DNA III polymerase. The mean values and standard deviations “SD” from at least three biological replicates are presented. Fold changes represent the ratio between the mean value observed for the Hfq3xFLAG-immunoprecipitated sample and the mean value for the control sample.

**Table S8. RCd1 conservation in 2,700 available *C. difficile* genomes and co-occurrence of RCd1 homologues with *skin* element, and important regulatory or skin excision components spoIIID, CD1231 and CD1234.**

| order | Unik ID | OPEN | Ribotype | ST | Length | RCd1 | *skin*-CD630 | SpoIIID | CD1231 | CD1234 |
|---|---|---|---|---|---|---|---|---|---|---|
| 1269 | ERR340084 | 1 | | 18 | 4217020 | 0 | 9 | 100 | 6 | 0 |
| 1285 | ERR340100 | 1 | | 2 | 4176768 | 0 | 15 | 99 | 6 | 0 |
| 2629 | ERR045748 | 1 | | 1 | 3814792 | 46 | 87 | 97 | 99 | 78 |
| 30 | ERR025349 | 1 | 027 | NF | 4158304 | 51 | 91 | 99 | 99 | 73 |
| 57 | ERR026373 | 1 | 027 | 1 | 4130260 | 51 | 92 | 99 | 99 | 95 |
| 297 | ERR030343 | 1 | 027 | 371 | 4029272 | 51 | 87 | 100 | 99 | 98 |
| 310 | ERR030356 | 1 | 027 | 1 | 3900514 | 51 | 82 | 99 | 99 | 73 |
| 275 | ERR030321 | 1 | | NF | 4248873 | 56 | 97 | 99 | 99 | 99 |
| 39 | ERR026355 | 1 | 027 | 1 | 4124498 | 57 | 92 | 99 | 99 | 99 |
| 323 | ERR030369 | 1 | 027 | 1 | 3963985 | 57 | 79 | 99 | 99 | 85 |
| 35 | ERR026351 | 1 | | NF | 4186497 | 59 | 90 | 97 | 99 | 73 |
| 2626 | ERR022124 | 1 | | 1 | 3819633 | 61 | 82 | 98 | 99 | 83 |
| 318 | ERR030364 | 1 | 027 | 1 | 4106256 | 62 | 92 | 99 | 99 | 98 |
| 70 | ERR026386 | 1 | 027 | 1 | 4139876 | 64 | 92 | 99 | 100 | 98 |
| 2637 | ERR045756 | 1 | 027 | 1 | 4114828 | 65 | 91 | 98 | 99 | 100 |
| 212 | ERR029519 | 1 | | 1 | 4391492 | 67 | 94 | 100 | 99 | 100 |
| 1265 | ERR340080 | 1 | | 53 | 4104840 | 67 | 92 | 99 | 99 | 99 |
| 1290 | ERR340105 | 1 | | 2 | 4161995 | 67 | 13 | 99 | 0 | 45 |
| 2126 | ERR232381 | 1 | | 3 | 4364844 | 67 | 52 | 99 | 99 | 95 |
| 2187 | ERR251752 | 1 | | 8 | 4051704 | 68 | 90 | 99 | 99 | 99 |
| 199 | ERR029506 | 1 | | 1 | 4217662 | 69 | 93 | 99 | 99 | 95 |
| 2099 | ERR142590 | 1 | | NF | 4494464 | 69 | 92 | 99 | 96 | 94 |
| 1248 | ERR340063 | 1 | | 8 | 4086874 | 70 | 92 | 99 | 100 | 99 |
| 96 | ERR026412 | 1 | 027 | 1 | 4138797 | 71 | 92 | 99 | 99 | 99 |
| 2153 | ERR251718 | 1 | | 2 | 4246736 | 71 | 91 | 100 | 100 | 100 |
| 1263 | ERR340078 | 1 | | 1 | 4132877 | 73 | 89 | 100 | 100 | 99 |
| 271 | ERR030317 | 1 | 027 | 1 | 4038041 | 74 | 82 | 99 | 99 | 86 |
| 1274 | ERR340089 | 1 | | 52 | 4199715 | 74 | 93 | 100 | 99 | 100 |
| 31 | ERR025351 | 1 | 027 | 1 | 4145725 | 75 | 93 | 99 | 100 | 99 |
| 109 | ERR026425 | 1 | 027 | 1 | 4149353 | 75 | 92 | 100 | 100 | 99 |
| 169 | ERR028729 | 1 | 027 | 1 | 4145278 | 75 | 93 | 100 | 100 | 100 |
| 186 | ERR029493 | 1 | | 1 | 4147771 | 75 | 93 | 100 | 99 | 100 |
| 1264 | ERR340079 | 1 | | 2 | 4131270 | 75 | 90 | 99 | 99 | 100 |
| 1273 | ERR340088 | 1 | | 2 | 4371170 | 75 | 92 | 100 | 99 | 99 |
| 1281 | ERR340096 | 1 | | 43 | 4143125 | 75 | 15 | 100 | 0 | 0 |
| 1556 | ERR126293 | 1 | | NF | 4270049 | 76 | 90 | 100 | 100 | 99 |
| 157 | ERR026878 | 1 | 027 | 1 | 4152114 | 77 | 92 | 100 | 99 | 78 |
| 2609 | ERR008623 | 1 | | 1 | 4131344 | 77 | 91 | 100 | 100 | 99 |
| 2631 | ERR045750 | 1 | 027 | 1 | 4245458 | 77 | 92 | 99 | 100 | 94 |
| 314 | ERR030360 | 1 | | 1 | 4090749 | 81 | 92 | 94 | 99 | 92 |
| 364 | ERR031686 | 1 | | 1 | 4178342 | 81 | 92 | 99 | 99 | 99 |
| 1268 | ERR340083 | 1 | | 8 | 4326715 | 81 | 93 | 99 | 99 | 100 |
| 2628 | ERR045747 | 1 | 027 | 1 | 3963590 | 81 | 88 | 100 | 99 | 90 |
| 1890 | ERR142381 | 1 | | NF | 4455250 | 82 | 60 | 99 | 100 | 99 |
| 2286 | ERR251860 | 1 | | 42 | 4231362 | 83 | 95 | 99 | 99 | 99 |
| 44 | ERR026360 | 1 | 027 | 1 | 4103904 | 84 | 90 | 100 | 99 | 98 |
| 351 | ERR031673 | 1 | | 1 | 4157280 | 84 | 92 | 99 | 99 | 99 |
| 1472 | ERR340287 | 1 | | 54 | 4540335 | 84 | 99 | 93 | 99 | 98 |
| 1817 | ERR142308 | 1 | | NF | 4462810 | 84 | 17 | 99 | 0 | 0 |
| 1916 | ERR142407 | 1 | | 44 | 4411760 | 84 | 14 | 99 | 0 | 0 |
| 2606 | ERR008620 | 1 | | 1 | 4225616 | 84 | 92 | 100 | 99 | 100 |
| 87 | ERR026403 | 1 | | 1 | 4129450 | 85 | 93 | 94 | 99 | 98 |
| 1398 | ERR340213 | 1 | | 17 | 4272885 | 85 | 58 | 99 | 99 | 94 |
| 1795 | ERR142286 | 1 | | NF | 4331063 | 85 | 62 | 99 | 100 | 92 |
| 2484 | ERR272243 | 1 | | 44 | 4261650 | 85 | 29 | 100 | 25 | 0 |
| 2624 | ERR022122 | 1 | | 1 | 4002633 | 85 | 81 | 99 | 99 | 94 |
| 2625 | ERR022123 | 1 | | 1 | 3923061 | 85 | 83 | 100 | 100 | 85 |
| 1282 | ERR340097 | 1 | | 2 | 4189317 | 86 | 90 | 100 | 99 | 99 |
| 1283 | ERR340098 | 1 | | 2 | 4221400 | 86 | 12 | 100 | 0 | 0 |
| 1359 | ERR340174 | 1 | | 8 | 4330048 | 86 | 95 | 100 | 99 | 99 |
| 1794 | ERR142285 | 1 | | NF | 4389469 | 86 | 16 | 100 | 0 | 0 |
| 143 | ERR026864 | 1 | 027 | 1 | 4147271 | 87 | 93 | 99 | 100 | 100 |
| 1557 | ERR126294 | 1 | 038 | 48 | 4169551 | 87 | 91 | 100 | 99 | 99 |
| 2049 | ERR142540 | 1 | | NF | 4256455 | 87 | 92 | 99 | 99 | 94 |
| 1169 | ERR339983 | 1 | | 8 | 4210660 | 88 | 95 | 100 | 100 | 100 |
| 1249 | ERR340064 | 1 | | 8 | 4153469 | 88 | 92 | 99 | 99 | 99 |
| 1799 | ERR142290 | 1 | | NF | 4445366 | 88 | 94 | 99 | 97 | 94 |
| 2636 | ERR045755 | 1 | 027 | 1 | 4107280 | 88 | 92 | 100 | 99 | 99 |
| 249 | ERR030295 | 1 | | 1 | 4091818 | 89 | 92 | 87 | 99 | 98 |
| 1255 | ERR340070 | 1 | | 3 | 4135170 | 89 | 15 | 99 | 0 | 0 |
| 1286 | ERR340101 | 1 | | 2 | 4178827 | 89 | 89 | 99 | 100 | 99 |
| 1288 | ERR340103 | 1 | | 2 | 4093839 | 89 | 91 | 100 | 99 | 99 |
| 2649 | ERR111464 | 1 | 027 | 1 | 3944288 | 89 | 88 | 100 | 99 | 99 |
| 2650 | ERR111465 | 1 | 027 | 1 | 4043026 | 89 | 91 | 99 | 99 | 92 |
| 23 | ERR025342 | 1 | | 1 | 4152839 | 90 | 93 | 100 | 99 | 92 |
| 91 | ERR026407 | 1 | 027 | 1 | 4079190 | 90 | 93 | 100 | 99 | 100 |
| 1278 | ERR340093 | 1 | | 46 | 4257773 | 90 | 94 | 100 | 99 | 100 |
| 2161 | ERR251726 | 1 | | NF | 4405157 | 90 | 61 | 99 | 100 | 98 |
| 2616 | ERR008630 | 1 | | 1 | 3983257 | 90 | 87 | 98 | 99 | 97 |
| 2632 | ERR045751 | 1 | 027 | 1 | 4508704 | 90 | 92 | 100 | 99 | 99 |
| 327 | ERR030373 | 1 | | 1 | 4221865 | 91 | 93 | 100 | 99 | 88 |
| 390 | ERR034220 | 1 | | 1 | 4171536 | 91 | 93 | 99 | 99 | 100 |
| 719 | ERR339531 | 1 | | 2 | 4113917 | 91 | 92 | 99 | 99 | 99 |
| 1257 | ERR340072 | 1 | | 2 | 4147318 | 91 | 85 | 99 | 100 | 98 |
| 1400 | ERR340215 | 1 | | 55 | 4086319 | 91 | 94 | 99 | 99 | 100 |
| 1526 | ERR126263 | 1 | | 3 | 4117712 | 91 | 7 | 98 | 0 | 0 |
| 2033 | ERR142524 | 1 | | 2 | 4113080 | 91 | 92 | 99 | 99 | 99 |
| 2144 | ERR232399 | 1 | | 178 | 4075709 | 91 | 31 | 100 | 80 | 0 |
| 2385 | ERR272144 | 1 | | 3 | 4192551 | 91 | 61 | 100 | 99 | 99 |
| 2601 | ERR008615 | 1 | | 371 | 4167238 | 91 | 92 | 100 | 99 | 98 |
| 2639 | ERR045758 | 1 | | 371 | 4083163 | 91 | 91 | 95 | 99 | 98 |
| 2643 | ERR111458 | 1 | 027 | 197 | 4030241 | 91 | 89 | 100 | 99 | 95 |
| 123 | ERR026842 | 1 | | 353 | 4477550 | 92 | 94 | 94 | 99 | 97 |
| 695 | ERR339507 | 1 | | NF | 4375274 | 92 | 95 | 100 | 99 | 99 |
| 1262 | ERR340077 | 1 | | 1 | 4138072 | 92 | 90 | 99 | 99 | 99 |
| 1910 | ERR142401 | 1 | | 44 | 4388272 | 92 | 20 | 100 | 13 | 0 |
| 2146 | ERR232401 | 1 | | 18 | 4283392 | 92 | 31 | 99 | 77 | 91 |
| 2310 | ERR272069 | 1 | | 58 | 4050846 | 92 | 14 | 99 | 0 | 0 |
| 2608 | ERR008622 | 1 | | 1 | 4234235 | 92 | 92 | 97 | 100 | 90 |
| 2634 | ERR045753 | 1 | 027 | 1 | 3938350 | 92 | 88 | 95 | 100 | 85 |
| 83 | ERR026399 | 1 | 027 | 1 | 4141026 | 93 | 93 | 99 | 100 | 99 |
| 100 | ERR026416 | 1 | | 1 | 4091732 | 93 | 92 | 93 | 99 | 98 |
| 133 | ERR026853 | 1 | 027 | 6 | 4290718 | 93 | 50 | 99 | 64 | 0 |
| 147 | ERR026868 | 1 | | 1 | 4107653 | 93 | 92 | 97 | 99 | 99 |
| 236 | ERR030282 | 1 | | 1 | 4101015 | 93 | 93 | 100 | 99 | 99 |
| 262 | ERR030308 | 1 | | 1 | 4097724 | 93 | 93 | 99 | 99 | 99 |
| 808 | ERR339620 | 1 | | 45 | 4468702 | 93 | 94 | 99 | 99 | 99 |
| 1253 | ERR340068 | 1 | | 15 | 4213050 | 93 | 92 | 99 | 99 | 100 |
| 1270 | ERR340085 | 1 | | 2 | 4231138 | 93 | 91 | 100 | 100 | 100 |
| 1294 | ERR340109 | 1 | | 2 | 4235891 | 93 | 90 | 100 | 99 | 99 |
| 1302 | ERR340117 | 1 | | 9 | 4322037 | 93 | 17 | 100 | 0 | 0 |
| 1767 | ERR142258 | 1 | | NF | 4437585 | 93 | 94 | 100 | 99 | 98 |
| 1801 | ERR142292 | 1 | | NF | 4570839 | 93 | 61 | 99 | 99 | 85 |
| 1843 | ERR142334 | 1 | | 6 | 4583939 | 93 | 14 | 100 | 0 | 0 |
| 1917 | ERR142408 | 1 | | NF | 4090795 | 93 | 95 | 99 | 99 | 99 |
| 1922 | ERR142413 | 1 | | NF | 4368395 | 93 | 58 | 99 | 100 | 97 |
| 1986 | ERR142477 | 1 | | NF | 4304132 | 93 | 58 | 99 | 99 | 87 |
| 2062 | ERR142553 | 1 | | 271 | 4342604 | 93 | 12 | 99 | 0 | 0 |
| 2063 | ERR142554 | 1 | | 1 | 4124778 | 93 | 93 | 98 | 96 | 99 |
| 2090 | ERR142581 | 1 | | 8 | 4237722 | 93 | 94 | 99 | 99 | 99 |
| 2105 | ERR142596 | 1 | | 49 | 4171765 | 93 | 92 | 99 | 99 | 95 |
| 2116 | ERR142607 | 1 | | 55 | 4091068 | 93 | 94 | 96 | 99 | 100 |
| 2189 | ERR251754 | 1 | | 2 | 4192325 | 93 | 92 | 100 | 99 | 100 |
| 2219 | ERR251784 | 1 | | 6 | 4356178 | 93 | 64 | 100 | 100 | 99 |
| 2479 | ERR272238 | 1 | | NF | 4362661 | 93 | 60 | 99 | 99 | 89 |
| 2623 | ERR022121 | 1 | | 1 | 3840594 | 93 | 80 | 99 | 98 | 79 |
| 2640 | ERR111454 | 1 | 027 | NF | 3911567 | 93 | 85 | 86 | 99 | 96 |
| 132 | ERR026852 | 1 | 027 | 1 | 4160625 | 94 | 93 | 99 | 99 | 97 |
| 151 | ERR026872 | 1 | 027 | 1 | 4153179 | 94 | 93 | 100 | 99 | 100 |
| 477 | ERR178542 | 1 | 017 | 37 | 4321907 | 94 | 94 | 100 | 99 | 99 |
| 736 | ERR339548 | 1 | | NF | 4152407 | 94 | 13 | 99 | 13 | 0 |
| 774 | ERR339586 | 1 | | 9 | 4348440 | 94 | 10 | 99 | 0 | 0 |
| 932 | ERR339744 | 1 | | 5 | 4277857 | 94 | 37 | 99 | 97 | 94 |
| 1250 | ERR340065 | 1 | | 8 | 4428113 | 94 | 95 | 100 | 99 | 99 |
| 1271 | ERR340086 | 1 | | 2 | 4186807 | 94 | 13 | 100 | 0 | 0 |
| 1284 | ERR340099 | 1 | | 2 | 4128146 | 94 | 88 | 100 | 99 | 99 |
| 1455 | ERR340270 | 1 | | 14 | 4335674 | 94 | 92 | 99 | 99 | 100 |
| 1760 | ERR142251 | 1 | | NF | 4198864 | 94 | 95 | 100 | 100 | 99 |
| 1806 | ERR142297 | 1 | | NF | 4638422 | 94 | 61 | 99 | 100 | 93 |
| 1936 | ERR142427 | 1 | | 42 | 4139861 | 94 | 95 | 100 | 100 | 98 |
| 1961 | ERR142452 | 1 | | NF | 4555295 | 94 | 61 | 100 | 99 | 94 |
| 2042 | ERR142533 | 1 | | NF | 4234446 | 94 | 12 | 100 | 0 | 45 |
| 2048 | ERR142539 | 1 | | NF | 4145721 | 94 | 12 | 99 | 13 | 0 |
| 2054 | ERR142545 | 1 | | 11 | 4029998 | 94 | 45 | 99 | 99 | 99 |
| 2094 | ERR142585 | 1 | | NF | 4353928 | 94 | 91 | 98 | 99 | 100 |
| 2095 | ERR142586 | 1 | | NF | 4462671 | 94 | 93 | 100 | 99 | 99 |
| 2173 | ERR251738 | 1 | | 8 | 4326391 | 94 | 95 | 100 | 100 | 100 |
| 2429 | ERR272188 | 1 | | 6 | 4332814 | 94 | 24 | 100 | 32 | 0 |
| 2602 | ERR008616 | 1 | | 1 | 4300403 | 94 | 93 | 100 | 100 | 97 |
| 2610 | ERR008624 | 1 | | 1 | 4509751 | 94 | 93 | 100 | 100 | 99 |
| 27 | ERR025346 | 1 | 027 | 1 | 4127502 | 95 | 93 | 99 | 100 | 99 |
| 74 | ERR026390 | 1 | | 1 | 4122954 | 95 | 92 | 99 | 99 | 99 |
| 139 | ERR026859 | 1 | 027 | 1 | 4139331 | 95 | 92 | 99 | 100 | 100 |
| 173 | ERR029480 | 1 | | 1 | 4143196 | 95 | 92 | 99 | 99 | 99 |
| 258 | ERR030304 | 1 | 027 | 1 | 4156817 | 95 | 92 | 100 | 99 | 99 |
| 992 | ERR339804 | 1 | | NF | 4579389 | 95 | 92 | 99 | 100 | 100 |
| 1261 | ERR340076 | 1 | | 1 | 4138766 | 95 | 92 | 100 | 99 | 100 |
| 1289 | ERR340104 | 1 | | 2 | 4196326 | 95 | 18 | 100 | 0 | 0 |
| 1293 | ERR340108 | 1 | | 51 | 4188342 | 95 | 95 | 99 | 99 | 100 |
| 1347 | ERR340162 | 1 | | 1 | 4441139 | 95 | 94 | 99 | 100 | 99 |
| 1410 | ERR340225 | 1 | | 8 | 4337142 | 95 | 95 | 100 | 99 | 99 |
| 1464 | ERR340279 | 1 | | 114 | 4323784 | 95 | 17 | 100 | 0 | 0 |
| 1558 | ERR126295 | 1 | | 2 | 4375612 | 95 | 91 | 100 | 100 | 99 |
| 1710 | ERR142201 | 1 | | 3 | 4406630 | 95 | 58 | 100 | 100 | 99 |
| 1724 | ERR142215 | 1 | | NF | 4156638 | 95 | 92 | 100 | 99 | 99 |
| 1765 | ERR142256 | 1 | | NF | 4436147 | 95 | 95 | 99 | 99 | 100 |
| 1770 | ERR142261 | 1 | | 1 | 4409194 | 95 | 95 | 100 | 100 | 99 |
| 1797 | ERR142288 | 1 | | 44 | 4369293 | 95 | 62 | 100 | 100 | 98 |
| 1879 | ERR142370 | 1 | | 11 | 4168101 | 95 | 47 | 99 | 99 | 99 |
| 1901 | ERR142392 | 1 | | 42 | 4207271 | 95 | 95 | 99 | 99 | 99 |
| 2002 | ERR142493 | 1 | | NF | 4309725 | 95 | 94 | 99 | 99 | 100 |
| 2081 | ERR142572 | 1 | | 46 | 4410462 | 95 | 94 | 99 | 99 | 99 |
| 2218 | ERR251783 | 1 | | 76 | 4138899 | 95 | 17 | 100 | 6 | 0 |
| 2227 | ERR251792 | 1 | | 1 | 4187791 | 95 | 95 | 99 | 99 | 99 |
| 2298 | ERR272057 | 1 | | 3 | 4399687 | 95 | 54 | 99 | 100 | 99 |
| 2358 | ERR272117 | 1 | | 3 | 4293036 | 95 | 59 | 99 | 99 | 99 |
| 2604 | ERR008618 | 1 | | 1 | 4330146 | 95 | 93 | 100 | 100 | 99 |
| 2605 | ERR008619 | 1 | | 1 | 4119115 | 95 | 93 | 100 | 100 | 100 |
| 2613 | ERR008627 | 1 | | 1 | 4244738 | 95 | 93 | 100 | 100 | 99 |
| 2617 | ERR008631 | 1 | | 1 | 4136379 | 95 | 92 | 99 | 99 | 99 |
| 2621 | ERR008635 | 1 | | 1 | 4181413 | 95 | 92 | 99 | 99 | 99 |
| 2635 | ERR045754 | 1 | 027 | 1 | 4133877 | 95 | 92 | 98 | 99 | 93 |
| 2638 | ERR045757 | 1 | 027 | 1 | 4199743 | 95 | 92 | 99 | 99 | 98 |
| 2646 | ERR111461 | 1 | 027 | 1 | 4179817 | 95 | 92 | 99 | 99 | 99 |
| 2647 | ERR111462 | 1 | 027 | 1 | 4180314 | 95 | 92 | 99 | 99 | 98 |

| | % Souche (80%) |
|---|---|
| **SpoIIID** | **100%** |
| **CD1231** | **86%** |
| **CD1234** | **85%** |
| **RCd1** | **99%** |

| | | | | | | | | | | |
|---|---|---|---|---|---|---|---|---|---|---|
| 2648 | ERR111463 | 1 | 027 | 1 | 4155508 | 95 | 92 | 98 | 99 | 98 |
| 2699 | ERR605029 | 1 | 078 | 11 | 3975505 | 95 | 43 | 99 | 99 | 99 |
| 302 | ERR030348 | 1 | 027 | 1 | 4135494 | 96 | 93 | 97 | 100 | 99 |
| 355 | ERR031677 | 1 | 027 | 1 | 4142836 | 96 | 93 | 100 | 100 | 100 |
| 775 | ERR339587 | 1 | | NF | 4291943 | 96 | 92 | 99 | 99 | 99 |
| 818 | ERR339630 | 1 | | 7 | 4106701 | 96 | 22 | 100 | 19 | 0 |
| 825 | ERR339637 | 1 | | 9 | 4310655 | 96 | 13 | 99 | 0 | 0 |
| 1229 | ERR340044 | 1 | | 8 | 4116357 | 96 | 92 | 99 | 99 | 98 |
| 1292 | ERR340107 | 1 | | 2 | 4185509 | 96 | 89 | 99 | 99 | 99 |
| 1390 | ERR340205 | 1 | | 48 | 4301452 | 96 | 95 | 100 | 99 | 100 |
| 1725 | ERR142216 | 1 | | 1 | 4138305 | 96 | 93 | 100 | 99 | 100 |
| 1790 | ERR142281 | 1 | | 35 | 4348398 | 96 | 11 | 99 | 0 | 0 |
| 1919 | ERR142410 | 1 | | NF | 4140086 | 96 | 95 | 99 | 99 | 99 |
| 2028 | ERR142519 | 1 | | NF | 4407005 | 96 | 94 | 100 | 100 | 91 |
| 2061 | ERR142552 | 1 | | 1 | 4157525 | 96 | 94 | 99 | 99 | 99 |
| 2085 | ERR142576 | 1 | | 1 | 4129211 | 96 | 93 | 99 | 98 | 99 |
| 2645 | ERR111460 | 1 | 027 | 1 | 4203698 | 96 | 92 | 98 | 99 | 98 |
| 46 | ERR026362 | 1 | 027 | 1 | 4133785 | 97 | 93 | 99 | 100 | 99 |
| 48 | ERR026364 | 1 | | 1 | 4147856 | 97 | 91 | 98 | 99 | 72 |
| 104 | ERR026420 | 1 | 027 | 1 | 4117135 | 97 | 93 | 100 | 100 | 99 |
| 112 | ERR026829 | 1 | | 1 | 4108148 | 97 | 92 | 96 | 99 | 97 |
| 144 | ERR026865 | 1 | 027 | 1 | 4133016 | 97 | 93 | 99 | 100 | 100 |
| 164 | ERR028724 | 1 | 027 | 1 | 4133371 | 97 | 93 | 100 | 99 | 99 |
| 190 | ERR029497 | 1 | 027 | 1 | 4139798 | 97 | 93 | 100 | 99 | 99 |
| 208 | ERR029515 | 1 | 027 | 1 | 4107745 | 97 | 93 | 100 | 100 | 100 |
| 234 | ERR029542 | 1 | 027 | 1 | 4117193 | 97 | 93 | 99 | 99 | 99 |
| 240 | ERR030286 | 1 | 027 | 1 | 4134433 | 97 | 93 | 100 | 100 | 99 |
| 305 | ERR030351 | 1 | 027 | 1 | 4118293 | 97 | 89 | 100 | 99 | 97 |
| 508 | ERR178573 | 1 | 017 | 37 | 4236519 | 97 | 94 | 99 | 99 | 99 |
| 682 | ERR339494 | 1 | | 8 | 4312546 | 97 | 95 | 100 | 99 | 100 |
| 714 | ERR339526 | 1 | | 1 | 4182497 | 97 | 92 | 99 | 99 | 98 |
| 731 | ERR339543 | 1 | | 14 | 4241133 | 97 | 92 | 94 | 99 | 99 |
| 735 | ERR339547 | 1 | | 2 | 4143396 | 97 | 15 | 99 | 0 | 54 |
| 785 | ERR339597 | 1 | | 9 | 4617905 | 97 | 54 | 100 | 57 | 64 |
| 787 | ERR339599 | 1 | | 9 | 4253633 | 97 | 14 | 99 | 11 | 0 |
| 850 | ERR339662 | 1 | | 5 | 4221819 | 97 | 44 | 99 | 99 | 99 |
| 978 | ERR339790 | 1 | | 1 | 4207427 | 97 | 94 | 99 | 99 | 100 |
| 1084 | ERR339897 | 1 | | 1 | 4145000 | 97 | 93 | 100 | 99 | 100 |
| 1109 | ERR339922 | 1 | | 1 | 4158936 | 97 | 93 | 99 | 99 | 100 |
| 1153 | ERR339967 | 1 | | 2 | 4178836 | 97 | 92 | 100 | 100 | 100 |
| 1200 | ERR340014 | 1 | | 2 | 4129098 | 97 | 92 | 100 | 100 | 100 |
| 1235 | ERR340050 | 1 | | 2 | 4144869 | 97 | 18 | 99 | 13 | 0 |
| 1252 | ERR340067 | 1 | | 15 | 4263860 | 97 | 94 | 99 | 100 | 99 |
| 1276 | ERR340091 | 1 | | 71 | 4486818 | 97 | 93 | 99 | 100 | 99 |
| 1279 | ERR340094 | 1 | | 2 | 4191589 | 97 | 92 | 100 | 100 | 99 |
| 1280 | ERR340095 | 1 | | 2 | 4172221 | 97 | 91 | 100 | 100 | 99 |
| 1295 | ERR340110 | 1 | | 53 | 4115805 | 97 | 92 | 99 | 99 | 99 |
| 1300 | ERR340115 | 1 | | 2 | 4158839 | 97 | 92 | 100 | 99 | 100 |
| 1325 | ERR340140 | 1 | | 6 | 4384101 | 97 | 25 | 100 | 42 | 0 |
| 1330 | ERR340145 | 1 | | 13 | 4396779 | 97 | 92 | 100 | 100 | 99 |
| 1333 | ERR340148 | 1 | | 8 | 4287963 | 97 | 95 | 99 | 99 | 99 |
| 1338 | ERR340153 | 1 | | 5 | 4359270 | 97 | 38 | 99 | 99 | 99 |
| 1346 | ERR340161 | 1 | | 2 | 4164615 | 97 | 93 | 99 | 99 | 99 |
| 1355 | ERR340170 | 1 | | 49 | 4154417 | 97 | 92 | 99 | 100 | 99 |
| 1360 | ERR340175 | 1 | | 1 | 4175930 | 97 | 94 | 99 | 99 | 99 |
| 1361 | ERR340176 | 1 | | 99 | 3997345 | 97 | 99 | 99 | 99 | 99 |
| 1364 | ERR340179 | 1 | | 7 | 4259751 | 97 | 15 | 99 | 0 | 0 |
| 1385 | ERR340200 | 1 | | 34 | 4225928 | 97 | 64 | 99 | 100 | 99 |
| 1388 | ERR340203 | 1 | | 37 | 4306528 | 97 | 94 | 100 | 99 | 98 |
| 1422 | ERR340237 | 1 | | 3 | 4155640 | 97 | 23 | 100 | 21 | 0 |
| 1432 | ERR340247 | 1 | | 49 | 4227419 | 97 | 92 | 99 | 99 | 100 |
| 1444 | ERR340259 | 1 | | 1 | 4350567 | 97 | 94 | 99 | 100 | 100 |
| 1466 | ERR340281 | 1 | | 13 | 4332191 | 97 | 92 | 100 | 100 | 100 |
| 1530 | ERR126267 | 1 | | 59 | 4190473 | 97 | 94 | 100 | 99 | 99 |
| 1571 | ERR142062 | 1 | | 1 | 4174102 | 97 | 95 | 99 | 99 | 99 |
| 1618 | ERR142109 | 1 | | 1 | 4128144 | 97 | 94 | 100 | 99 | 100 |
| 1621 | ERR142112 | 1 | | 1 | 4130639 | 97 | 94 | 99 | 99 | 100 |
| 1622 | ERR142113 | 1 | | 17 | 4245972 | 97 | 63 | 99 | 100 | 94 |
| 1628 | ERR142119 | 1 | | 1 | 4135155 | 97 | 96 | 99 | 100 | 100 |
| 1660 | ERR142151 | 1 | | 2 | 4123953 | 97 | 93 | 99 | 100 | 100 |
| 1728 | ERR142219 | 1 | | 37 | 4229122 | 97 | 94 | 99 | 99 | 100 |
| 1737 | ERR142228 | 1 | | 37 | 4258274 | 97 | 94 | 100 | 99 | 99 |
| 1772 | ERR142263 | 1 | | 1 | 4372170 | 97 | 94 | 99 | 100 | 98 |
| 1775 | ERR142266 | 1 | | 1 | 4353045 | 97 | 94 | 100 | 99 | 99 |
| 1780 | ERR142271 | 1 | | 1 | 4403814 | 97 | 94 | 100 | 100 | 100 |
| 1809 | ERR142300 | 1 | | 44 | 4366445 | 97 | 63 | 99 | 99 | 99 |
| 1867 | ERR142358 | 1 | | NF | 4262731 | 97 | 13 | 99 | 0 | 0 |
| 1868 | ERR142359 | 1 | | 63 | 4257769 | 97 | 99 | 99 | 100 | 100 |
| 1869 | ERR142360 | 1 | | 17 | 4473192 | 97 | 66 | 99 | 99 | 99 |
| 1877 | ERR142368 | 1 | | 37 | 4338363 | 97 | 94 | 99 | 99 | 99 |
| 1878 | ERR142369 | 1 | | 14 | 4541560 | 97 | 92 | 99 | 99 | 99 |
| 1896 | ERR142387 | 1 | | NF | 4177207 | 97 | 95 | 99 | 99 | 99 |
| 1905 | ERR142396 | 1 | | NF | 4113061 | 97 | 95 | 99 | 99 | 99 |
| 1907 | ERR142398 | 1 | | 11 | 4024209 | 97 | 45 | 99 | 99 | 100 |
| 1908 | ERR142399 | 1 | | NF | 4354961 | 97 | 95 | 100 | 99 | 99 |
| 1915 | ERR142406 | 1 | | NF | 4315775 | 97 | 94 | 99 | 99 | 99 |
| 1918 | ERR142409 | 1 | | NF | 4356783 | 97 | 94 | 99 | 99 | 99 |
| 1927 | ERR142418 | 1 | | NF | 4369439 | 97 | 13 | 99 | 0 | 0 |
| 1933 | ERR142424 | 1 | | NF | 4087644 | 97 | 95 | 99 | 99 | 99 |
| 1940 | ERR142431 | 1 | | 42 | 4212903 | 97 | 95 | 100 | 100 | 99 |
| 1947 | ERR142438 | 1 | | NF | 4279845 | 97 | 95 | 98 | 100 | 99 |
| 1959 | ERR142450 | 1 | | NF | 4515637 | 97 | 64 | 100 | 99 | 92 |
| 1960 | ERR142451 | 1 | | 42 | 4166669 | 97 | 95 | 99 | 99 | 99 |
| 1970 | ERR142461 | 1 | | 6 | 4231025 | 97 | 15 | 99 | 0 | 0 |
| 1979 | ERR142470 | 1 | | 6 | 4376312 | 97 | 60 | 99 | 99 | 97 |
| 1981 | ERR142472 | 1 | | 11 | 4003164 | 97 | 47 | 99 | 99 | 100 |
| 1992 | ERR142483 | 1 | | 1 | 4475208 | 97 | 94 | 99 | 99 | 100 |
| 1995 | ERR142486 | 1 | | 44 | 4529674 | 97 | 58 | 99 | 99 | 91 |
| 2006 | ERR142497 | 1 | | NF | 4356859 | 97 | 60 | 99 | 100 | 96 |
| 2012 | ERR142503 | 1 | | 44 | 4521906 | 97 | 59 | 99 | 100 | 98 |
| 2014 | ERR142505 | 1 | | 144 | 4185432 | 97 | 57 | 100 | 99 | 87 |
| 2026 | ERR142517 | 1 | | 1 | 4145611 | 97 | 91 | 98 | 99 | 98 |
| 2030 | ERR142521 | 1 | | 1 | 4147328 | 97 | 93 | 99 | 99 | 99 |
| 2031 | ERR142522 | 1 | | 1 | 4151219 | 97 | 93 | 99 | 99 | 100 |
| 2059 | ERR142550 | 1 | | 271 | 4403580 | 97 | 14 | 99 | 13 | 0 |
| 2060 | ERR142551 | 1 | | 1 | 4146662 | 97 | 93 | 99 | 99 | 99 |
| 2073 | ERR142564 | 1 | | 1 | 4143083 | 97 | 93 | 100 | 99 | 100 |
| 2075 | ERR142566 | 1 | | 46 | 4467412 | 97 | 94 | 100 | 99 | 99 |
| 2076 | ERR142567 | 1 | | 1 | 4149346 | 97 | 91 | 99 | 100 | 90 |
| 2079 | ERR142570 | 1 | | 8 | 4277919 | 97 | 95 | 99 | 100 | 99 |
| 2082 | ERR142573 | 1 | | 8 | 4230489 | 97 | 95 | 98 | 99 | 96 |
| 2083 | ERR142574 | 1 | | 1 | 4152296 | 97 | 94 | 99 | 99 | 99 |
| 2084 | ERR142575 | 1 | | 1 | 4168117 | 97 | 93 | 94 | 99 | 99 |
| 2089 | ERR142580 | 1 | | 1 | 4173408 | 97 | 93 | 99 | 100 | 99 |
| 2103 | ERR142594 | 1 | | 49 | 4446676 | 97 | 92 | 99 | 99 | 99 |
| 2104 | ERR142595 | 1 | | NF | 4201678 | 97 | 91 | 100 | 99 | 99 |
| 2121 | ERR232376 | 1 | | 7 | 4476437 | 97 | 19 | 100 | 0 | 0 |
| 2127 | ERR232382 | 1 | | NF | 4204876 | 97 | 87 | 100 | 99 | 89 |
| 2147 | ERR232402 | 1 | | 181 | 4092735 | 97 | 20 | 99 | 67 | 0 |
| 2149 | ERR251714 | 1 | | 14 | 4407184 | 97 | 92 | 100 | 100 | 100 |
| 2246 | ERR251811 | 1 | | 2 | 4904572 | 97 | 95 | 99 | 100 | 100 |
| 2293 | ERR251874 | 1 | | 1 | 4237811 | 97 | 8 | 99 | 0 | 0 |
| 2294 | ERR251876 | 1 | | 8 | 4287839 | 97 | 95 | 100 | 99 | 100 |
| 2311 | ERR272070 | 1 | | 58 | 4176717 | 97 | 13 | 99 | 0 | 0 |
| 2357 | ERR272116 | 1 | | 3 | 4245495 | 97 | 60 | 99 | 100 | 95 |
| 2366 | ERR272125 | 1 | | 3 | 4222344 | 97 | 60 | 99 | 100 | 100 |
| 2376 | ERR272135 | 1 | | 3 | 4361085 | 97 | 58 | 99 | 100 | 93 |
| 2401 | ERR272160 | 1 | | NF | 4246281 | 97 | 14 | 100 | 0 | 0 |
| 2407 | ERR272166 | 1 | | 6 | 4437455 | 97 | 34 | 100 | 19 | 0 |
| 2411 | ERR272170 | 1 | | 6 | 4513509 | 97 | 20 | 99 | 13 | 0 |
| 2460 | ERR272219 | 1 | | 7 | 4233630 | 97 | 19 | 100 | 13 | 0 |
| 2494 | ERR272253 | 1 | | 44 | 4479359 | 97 | 71 | 100 | 99 | 86 |
| 2497 | ERR272256 | 1 | | 44 | 4410027 | 97 | 94 | 99 | 100 | 99 |
| 2503 | ERR272262 | 1 | | 44 | 4633492 | 97 | 16 | 99 | 13 | 0 |
| 2534 | ERR467560 | 1 | | 1 | 4178203 | 97 | 93 | 100 | 100 | 99 |
| 2548 | ERR467574 | 1 | | 8 | 4190488 | 97 | 95 | 100 | 100 | 99 |
| 2600 | ERR008614 | 1 | | 1 | 4126976 | 97 | 92 | 100 | 99 | 98 |
| 2611 | ERR008625 | 1 | | 1 | 4164066 | 97 | 91 | 100 | 100 | 96 |
| 2656 | E12 | 1 | 106 | 42 | 4044621 | 97 | 97 | 100 | 100 | 99 |
| 1 | ERR024642 | 1 | 027 | 1 | 4155525 | 98 | 93 | 99 | 99 | 99 |
| 5 | ERR024647 | 1 | 027 | 1 | 4167095 | 98 | 94 | 99 | 99 | 100 |
| 13 | ERR024656 | 1 | 027 | 1 | 4134915 | 98 | 94 | 100 | 100 | 99 |
| 18 | ERR024662 | 1 | 027 | 1 | 4818638 | 98 | 93 | 100 | 99 | 99 |
| 19 | ERR024663 | 1 | 027 | 1 | 4138042 | 98 | 92 | 99 | 99 | 99 |
| 20 | ERR024664 | 1 | 027 | 1 | 4133722 | 98 | 93 | 100 | 99 | 99 |
| 21 | ERR024665 | 1 | 027 | 1 | 4141337 | 98 | 94 | 99 | 100 | 99 |
| 22 | ERR024666 | 1 | 027 | 1 | 4142369 | 98 | 94 | 99 | 99 | 99 |
| 24 | ERR025343 | 1 | 027 | 1 | 4227412 | 98 | 93 | 100 | 100 | 100 |
| 25 | ERR025344 | 1 | 027 | 1 | 4130185 | 98 | 93 | 100 | 99 | 100 |
| 28 | ERR025347 | 1 | 027 | 1 | 4131322 | 98 | 93 | 100 | 99 | 98 |
| 29 | ERR025348 | 1 | 027 | 1 | 4137774 | 98 | 93 | 99 | 99 | 99 |
| 37 | ERR026353 | 1 | 027 | 1 | 4569806 | 98 | 95 | 100 | 99 | 100 |
| 41 | ERR026357 | 1 | 027 | 1 | 4765510 | 98 | 93 | 99 | 99 | 99 |
| 43 | ERR026359 | 1 | 027 | 1 | 4130394 | 98 | 93 | 99 | 99 | 99 |
| 45 | ERR026361 | 1 | 027 | 1 | 4135762 | 98 | 93 | 100 | 99 | 99 |
| 51 | ERR026367 | 1 | 027 | 1 | 4132791 | 98 | 93 | 100 | 100 | 99 |
| 52 | ERR026368 | 1 | 027 | 1 | 4136247 | 98 | 93 | 100 | 99 | 98 |
| 58 | ERR026374 | 1 | 027 | 1 | 4225593 | 98 | 93 | 99 | 99 | 99 |
| 61 | ERR026377 | 1 | | 1 | 4144493 | 98 | 92 | 99 | 99 | 99 |
| 64 | ERR026380 | 1 | 027 | 1 | 4138427 | 98 | 93 | 100 | 100 | 99 |
| 69 | ERR026385 | 1 | 027 | 5 | 4422806 | 98 | 46 | 99 | 99 | 81 |
| 71 | ERR026387 | 1 | 027 | 1 | 4131698 | 98 | 93 | 100 | 99 | 99 |
| 72 | ERR026388 | 1 | 027 | 1 | 4134638 | 98 | 93 | 100 | 100 | 99 |
| 75 | ERR026391 | 1 | 027 | 1 | 4139277 | 98 | 93 | 99 | 99 | 100 |
| 81 | ERR026397 | 1 | 027 | 1 | 4136217 | 98 | 93 | 100 | 100 | 100 |
| 90 | ERR026406 | 1 | 027 | 1 | 4124130 | 98 | 93 | 99 | 99 | 99 |
| 92 | ERR026408 | 1 | 027 | 1 | 4138735 | 98 | 93 | 99 | 100 | 99 |
| 94 | ERR026410 | 1 | 027 | 1 | 4128458 | 98 | 93 | 99 | 100 | 100 |
| 95 | ERR026411 | 1 | 027 | 1 | 4136473 | 98 | 93 | 99 | 99 | 100 |
| 97 | ERR026413 | 1 | 027 | 1 | 4135627 | 98 | 93 | 100 | 99 | 100 |
| 98 | ERR026414 | 1 | 027 | 1 | 4131527 | 98 | 93 | 100 | 100 | 99 |
| 105 | ERR026421 | 1 | 027 | 1 | 4137605 | 98 | 93 | 99 | 100 | 99 |
| 110 | ERR026426 | 1 | 027 | 1 | 4141871 | 98 | 93 | 99 | 99 | 100 |
| 111 | ERR026427 | 1 | 027 | 1 | 4138959 | 98 | 93 | 99 | 99 | 100 |
| 113 | ERR026832 | 1 | 027 | 1 | 4135081 | 98 | 93 | 100 | 100 | 98 |
| 115 | ERR026834 | 1 | 027 | 1 | 4132455 | 98 | 93 | 100 | 99 | 100 |
| 117 | ERR026836 | 1 | 027 | 1 | 4135344 | 98 | 93 | 100 | 99 | 100 |
| 119 | ERR026838 | 1 | 027 | 1 | 4160033 | 98 | 93 | 100 | 100 | 100 |
| 120 | ERR026839 | 1 | 027 | 1 | 4132890 | 98 | 93 | 100 | 100 | 100 |
| 121 | ERR026840 | 1 | 027 | 1 | 4131545 | 98 | 93 | 99 | 99 | 99 |
| 122 | ERR026841 | 1 | 027 | 1 | 4073855 | 98 | 93 | 99 | 99 | 100 |
| 125 | ERR026844 | 1 | 027 | 1 | 4131330 | 98 | 93 | 99 | 99 | 100 |
| 130 | ERR026850 | 1 | 027 | 1 | 4134383 | 98 | 93 | 100 | 99 | 99 |
| 135 | ERR026855 | 1 | | 1 | 4110805 | 98 | 92 | 98 | 99 | 99 |
| 137 | ERR026857 | 1 | 027 | 1 | 4142545 | 98 | 93 | 100 | 99 | 99 |
| 138 | ERR026858 | 1 | 027 | 1 | 4131901 | 98 | 93 | 99 | 99 | 100 |
| 140 | ERR026860 | 1 | 027 | 1 | 4145235 | 98 | 93 | 100 | 99 | 100 |

| | | | | | | | | | | |
|---|---|---|---|---|---|---|---|---|---|---|
| 149 | ERR026870 | 1 | 027 | 1 | 4139709 | 98 | 93 | 99 | 99 | 99 |
| 152 | ERR026873 | 1 | 027 | 1 | 4136209 | 98 | 93 | 99 | 99 | 100 |
| 155 | ERR026876 | 1 | 027 | 1 | 4166539 | 98 | 93 | 100 | 100 | 100 |
| 156 | ERR026877 | 1 | 027 | 1 | 4159013 | 98 | 93 | 100 | 99 | 100 |
| 158 | ERR026879 | 1 | 027 | 1 | 4146269 | 98 | 93 | 99 | 100 | 99 |
| 161 | ERR028721 | 1 | 027 | 1 | 4140739 | 98 | 93 | 100 | 100 | 99 |
| 167 | ERR028727 | 1 | 027 | 1 | 4133799 | 98 | 93 | 100 | 100 | 100 |
| 171 | ERR028731 | 1 | 027 | 1 | 4134062 | 98 | 93 | 100 | 99 | 100 |
| 172 | ERR028732 | 1 | 027 | 1 | 4136497 | 98 | 93 | 99 | 99 | 99 |
| 174 | ERR029481 | 1 | 027 | 1 | 4135968 | 98 | 93 | 100 | 100 | 99 |
| 175 | ERR029482 | 1 | 027 | 1 | 4129714 | 98 | 93 | 99 | 100 | 99 |
| 179 | ERR029486 | 1 | 027 | 1 | 4158548 | 98 | 93 | 100 | 99 | 99 |
| 180 | ERR029487 | 1 | 027 | 1 | 4135260 | 98 | 93 | 100 | 100 | 100 |
| 182 | ERR029489 | 1 | 027 | 1 | 4153039 | 98 | 92 | 99 | 99 | 100 |
| 193 | ERR029500 | 1 | 027 | 1 | 4134868 | 98 | 93 | 100 | 99 | 100 |
| 195 | ERR029502 | 1 | 027 | 1 | 4199552 | 98 | 93 | 99 | 99 | 98 |
| 198 | ERR029505 | 1 | 027 | 1 | 4138309 | 98 | 93 | 100 | 100 | 100 |
| 203 | ERR029510 | 1 | 027 | 1 | 4092821 | 98 | 93 | 99 | 99 | 99 |
| 205 | ERR029512 | 1 | 027 | 1 | 4192916 | 98 | 93 | 100 | 100 | 100 |
| 213 | ERR029520 | 1 | 027 | 1 | 4091759 | 98 | 93 | 100 | 100 | 99 |
| 215 | ERR029522 | 1 | 27 | 123 | 4082471 | 98 | 95 | 99 | 99 | 99 |
| 216 | ERR029523 | 1 | 027 | 1 | 4165300 | 98 | 93 | 100 | 100 | 100 |
| 220 | ERR029527 | 1 | 027 | 1 | 4259392 | 98 | 93 | 100 | 99 | 99 |
| 221 | ERR029528 | 1 | 027 | 1 | 4280828 | 98 | 92 | 99 | 100 | 99 |
| 222 | ERR029529 | 1 | 027 | 1 | 4092442 | 98 | 93 | 100 | 99 | 100 |
| 223 | ERR029530 | 1 | 027 | 47 | 4071306 | 98 | 13 | 100 | 0 | 0 |
| 228 | ERR029536 | 1 | 027 | 1 | 4136949 | 98 | 93 | 99 | 100 | 99 |
| 233 | ERR029541 | 1 | 027 | 1 | 4101541 | 98 | 93 | 100 | 100 | 100 |
| 235 | ERR029543 | 1 | 027 | 1 | 4144755 | 98 | 93 | 99 | 99 | 100 |
| 237 | ERR030283 | 1 | 027 | 1 | 4134546 | 98 | 93 | 100 | 100 | 100 |
| 239 | ERR030285 | 1 | 027 | 1 | 4134025 | 98 | 93 | 99 | 99 | 99 |
| 242 | ERR030288 | 1 | 027 | 1 | 4131540 | 98 | 93 | 100 | 100 | 100 |
| 243 | ERR030289 | 1 | 027 | 1 | 4158157 | 98 | 93 | 100 | 99 | 98 |
| 245 | ERR030291 | 1 | 027 | 1 | 4158835 | 98 | 93 | 100 | 99 | 100 |
| 246 | ERR030292 | 1 | 027 | 1 | 4132754 | 98 | 93 | 100 | 99 | 99 |
| 250 | ERR030296 | 1 | 027 | 1 | 4134145 | 98 | 93 | 99 | 99 | 99 |
| 251 | ERR030297 | 1 | 027 | 1 | 4132525 | 98 | 93 | 99 | 100 | 99 |
| 253 | ERR030299 | 1 | 027 | 1 | 4140726 | 98 | 93 | 99 | 99 | 100 |
| 257 | ERR030303 | 1 | 027 | 1 | 4155637 | 98 | 93 | 99 | 100 | 100 |
| 259 | ERR030305 | 1 | 027 | 1 | 4135003 | 98 | 93 | 100 | 99 | 100 |
| 260 | ERR030306 | 1 | 027 | 1 | 4134728 | 98 | 93 | 100 | 99 | 100 |
| 265 | ERR030311 | 1 | 027 | 1 | 4135705 | 98 | 93 | 100 | 100 | 99 |
| 274 | ERR030320 | 1 | 027 | 1 | 4136770 | 98 | 93 | 99 | 99 | 99 |
| 276 | ERR030322 | 1 | 027 | 1 | 4142678 | 98 | 93 | 99 | 99 | 100 |
| 280 | ERR030326 | 1 | 027 | 1 | 4132600 | 98 | 93 | 100 | 99 | 99 |
| 284 | ERR030330 | 1 | 027 | 1 | 4145988 | 98 | 91 | 100 | 99 | 98 |
| 285 | ERR030331 | 1 | 027 | 1 | 4162921 | 98 | 93 | 100 | 100 | 100 |
| 286 | ERR030332 | 1 | 027 | 54 | 4268303 | 98 | 99 | 100 | 99 | 99 |
| 288 | ERR030334 | 1 | | 1 | 4250607 | 98 | 92 | 100 | 99 | 92 |
| 289 | ERR030335 | 1 | 027 | 1 | 4136885 | 98 | 93 | 100 | 99 | 100 |
| 292 | ERR030338 | 1 | 027 | 1 | 4128030 | 98 | 90 | 100 | 99 | 98 |
| 295 | ERR030341 | 1 | 027 | 1 | 4122682 | 98 | 93 | 99 | 100 | 99 |
| 296 | ERR030342 | 1 | 027 | 1 | 4155444 | 98 | 93 | 99 | 100 | 99 |
| 299 | ERR030345 | 1 | 027 | 1 | 4085819 | 98 | 93 | 100 | 99 | 100 |
| 311 | ERR030357 | 1 | 027 | 1 | 4132941 | 98 | 93 | 100 | 100 | 99 |
| 313 | ERR030359 | 1 | 027 | 1 | 4076317 | 98 | 93 | 100 | 99 | 99 |
| 315 | ERR030361 | 1 | 027 | 1 | 4140434 | 98 | 93 | 100 | 100 | 99 |
| 316 | ERR030362 | 1 | 027 | 1 | 4135306 | 98 | 93 | 100 | 100 | 99 |
| 317 | ERR030363 | 1 | 027 | 1 | 4136094 | 98 | 93 | 99 | 100 | 99 |
| 319 | ERR030365 | 1 | 027 | 1 | 4135537 | 98 | 93 | 100 | 100 | 100 |
| 320 | ERR030366 | 1 | 027 | 1 | 4134883 | 98 | 93 | 99 | 100 | 100 |
| 324 | ERR030370 | 1 | 027 | 1 | 4133757 | 98 | 93 | 100 | 100 | 100 |
| 325 | ERR030371 | 1 | 027 | 1 | 4135207 | 98 | 93 | 100 | 99 | 100 |
| 326 | ERR030372 | 1 | 027 | 1 | 4136458 | 98 | 93 | 99 | 99 | 100 |
| 330 | ERR030376 | 1 | 027 | 1 | 4131578 | 98 | 93 | 100 | 99 | 99 |
| 331 | ERR030377 | 1 | 027 | 1 | 4145489 | 98 | 93 | 99 | 100 | 100 |
| 334 | ERR030380 | 1 | 027 | 1 | 4148822 | 98 | 93 | 100 | 100 | 100 |
| 338 | ERR030386 | 1 | | 1 | 4140732 | 98 | 92 | 99 | 99 | 98 |
| 340 | ERR030388 | 1 | 027 | 1 | 4136121 | 98 | 93 | 100 | 100 | 100 |
| 342 | ERR030390 | 1 | 027 | 1 | 4182894 | 98 | 93 | 99 | 100 | 99 |
| 345 | ERR030393 | 1 | 027 | 1 | 4130920 | 98 | 93 | 100 | 100 | 100 |
| 346 | ERR030394 | 1 | 027 | 1 | 4132923 | 98 | 93 | 99 | 100 | 100 |
| 347 | ERR030395 | 1 | 027 | 1 | 4149141 | 98 | 92 | 99 | 99 | 100 |
| 350 | ERR030398 | 1 | 027 | 1 | 4130571 | 98 | 93 | 100 | 99 | 100 |
| 356 | ERR031678 | 1 | 027 | 1 | 4135917 | 98 | 93 | 100 | 100 | 99 |
| 357 | ERR031679 | 1 | 027 | 1 | 4135120 | 98 | 93 | 100 | 100 | 100 |
| 360 | ERR031682 | 1 | 027 | 1 | 4187166 | 98 | 93 | 99 | 99 | 100 |
| 361 | ERR031683 | 1 | 027 | 1 | 4143785 | 98 | 93 | 100 | 99 | 100 |
| 362 | ERR031684 | 1 | 027 | 1 | 4144741 | 98 | 93 | 100 | 99 | 100 |
| 369 | ERR031691 | 1 | 027 | 1 | 4138900 | 98 | 93 | 99 | 100 | 99 |
| 371 | ERR031693 | 1 | 027 | 1 | 4130392 | 98 | 93 | 99 | 100 | 100 |
| 373 | ERR031695 | 1 | 027 | 1 | 4165088 | 98 | 91 | 100 | 100 | 99 |
| 374 | ERR031696 | 1 | 027 | 1 | 4076101 | 98 | 93 | 100 | 100 | 100 |
| 376 | ERR031698 | 1 | 027 | 1 | 4135854 | 98 | 93 | 100 | 99 | 99 |
| 380 | ERR031702 | 1 | 027 | 1 | 4229077 | 98 | 93 | 100 | 100 | 99 |
| 385 | ERR031707 | 1 | 027 | 1 | 4159122 | 98 | 93 | 99 | 99 | 100 |
| 387 | ERR031709 | 1 | 027 | 1 | 4132300 | 98 | 93 | 100 | 99 | 100 |
| 388 | ERR031710 | 1 | 027 | 1 | 4127157 | 98 | 93 | 99 | 99 | 99 |
| 395 | ERR034225 | 1 | 027 | 1 | 4133224 | 98 | 93 | 100 | 100 | 100 |
| 398 | ERR034228 | 1 | 027 | 1 | 4137779 | 98 | 94 | 100 | 100 | 99 |
| 409 | ERR044811 | 1 | 027 | NF | 4090262 | 98 | 95 | 100 | 100 | 100 |
| 412 | ERR044814 | 1 | 027 | 1 | 4295804 | 98 | 93 | 100 | 100 | 100 |
| 445 | ERR178510 | 1 | 017 | 8 | 4441393 | 98 | 90 | 100 | 99 | 99 |
| 446 | ERR178511 | 1 | 017 | 37 | 4253629 | 98 | 94 | 99 | 99 | 100 |
| 447 | ERR178512 | 1 | 017 | 37 | 4264090 | 98 | 94 | 99 | 100 | 99 |
| 449 | ERR178514 | 1 | 017 | 37 | 4269795 | 98 | 94 | 99 | 99 | 100 |
| 452 | ERR178517 | 1 | 017 | 37 | 4255872 | 98 | 94 | 100 | 99 | 100 |
| 456 | ERR178521 | 1 | 017 | 37 | 4262050 | 98 | 94 | 99 | 99 | 99 |
| 459 | ERR178524 | 1 | 017 | 37 | 4467461 | 98 | 94 | 99 | 99 | 99 |
| 460 | ERR178525 | 1 | 017 | 37 | 4371645 | 98 | 94 | 100 | 99 | 100 |
| 463 | ERR178528 | 1 | 017 | 37 | 4129508 | 98 | 94 | 100 | 99 | 100 |
| 466 | ERR178531 | 1 | 017 | 37 | 4440386 | 98 | 94 | 100 | 100 | 99 |
| 469 | ERR178534 | 1 | 017 | 37 | 4291537 | 98 | 94 | 99 | 99 | 99 |
| 471 | ERR178536 | 1 | 017 | 37 | 4251943 | 98 | 94 | 99 | 99 | 99 |
| 472 | ERR178537 | 1 | 017 | 37 | 4241159 | 98 | 94 | 100 | 100 | 99 |
| 474 | ERR178539 | 1 | 017 | 37 | 4270414 | 98 | 94 | 99 | 99 | 99 |
| 476 | ERR178541 | 1 | 017 | 37 | 4280513 | 98 | 94 | 99 | 99 | 99 |
| 483 | ERR178548 | 1 | 017 | 37 | 4244679 | 98 | 94 | 99 | 99 | 99 |
| 486 | ERR178551 | 1 | 017 | 37 | 4238700 | 98 | 94 | 98 | 99 | 99 |
| 489 | ERR178554 | 1 | 017 | 37 | 4228055 | 98 | 94 | 100 | 99 | 100 |
| 490 | ERR178555 | 1 | 017 | 37 | 4219874 | 98 | 94 | 100 | 99 | 99 |
| 494 | ERR178559 | 1 | 017 | 37 | 4214886 | 98 | 94 | 100 | 99 | 99 |
| 499 | ERR178564 | 1 | 017 | 37 | 4181031 | 98 | 93 | 100 | 99 | 99 |
| 500 | ERR178565 | 1 | 017 | 37 | 4180892 | 98 | 94 | 99 | 99 | 99 |
| 506 | ERR178571 | 1 | 017 | 37 | 4238757 | 98 | 94 | 99 | 99 | 99 |
| 507 | ERR178572 | 1 | 017 | 37 | 4238458 | 98 | 94 | 99 | 99 | 100 |
| 509 | ERR178574 | 1 | 017 | 37 | 4238595 | 98 | 93 | 99 | 99 | 99 |
| 511 | ERR178576 | 1 | 017 | 37 | 4242068 | 98 | 94 | 99 | 99 | 97 |
| 513 | ERR178579 | 1 | 017 | 37 | 4240159 | 98 | 94 | 100 | 99 | 99 |
| 517 | ERR178583 | 1 | 017 | 37 | 4239274 | 98 | 94 | 99 | 99 | 98 |
| 518 | ERR178584 | 1 | 017 | 37 | 4237841 | 98 | 94 | 98 | 99 | 99 |
| 519 | ERR178585 | 1 | 017 | 37 | 4242763 | 98 | 94 | 99 | 100 | 99 |
| 520 | ERR178586 | 1 | 017 | 37 | 4239591 | 98 | 94 | 100 | 99 | 100 |
| 521 | ERR178587 | 1 | 017 | 37 | 4238026 | 98 | 94 | 100 | 99 | 99 |
| 522 | ERR178588 | 1 | 017 | 37 | 4243314 | 98 | 94 | 99 | 99 | 99 |
| 524 | ERR178590 | 1 | 017 | 37 | 4246241 | 98 | 94 | 100 | 99 | 99 |
| 531 | ERR178597 | 1 | 017 | 37 | 4276648 | 98 | 94 | 100 | 99 | 100 |
| 533 | ERR178599 | 1 | 017 | 37 | 4209638 | 98 | 94 | 100 | 100 | 100 |
| 535 | ERR178601 | 1 | 017 | 37 | 4237363 | 98 | 94 | 100 | 99 | 98 |
| 538 | ERR178604 | 1 | 017 | 37 | 4259087 | 98 | 94 | 99 | 99 | 99 |
| 557 | ERR340608 | 1 | 017 | 37 | 4224750 | 98 | 94 | 100 | 99 | 100 |
| 561 | ERR340616 | 1 | 017 | 37 | 4252100 | 98 | 94 | 99 | 99 | 100 |
| 567 | ERR340624 | 1 | 017 | 8 | 4451359 | 98 | 95 | 99 | 99 | 99 |
| 568 | ERR340626 | 1 | 017 | 37 | 4191479 | 98 | 94 | 100 | 100 | 99 |
| 574 | ERR340636 | 1 | 017 | 37 | 4302392 | 98 | 94 | 100 | 99 | 100 |
| 577 | ERR340639 | 1 | 017 | 37 | 4454040 | 98 | 94 | 100 | 100 | 99 |
| 578 | ERR340640 | 1 | 017 | 37 | 4422736 | 98 | 94 | 100 | 100 | 100 |
| 587 | ERR340659 | 1 | | 37 | 4350552 | 98 | 94 | 100 | 99 | 100 |
| 600 | ERR387373 | 1 | 017 | 37 | 4307899 | 98 | 94 | 100 | 99 | 99 |
| 610 | ERR387387 | 1 | | 37 | 4227728 | 98 | 94 | 100 | 100 | 99 |
| 614 | ERR387391 | 1 | 017 | 37 | 4210832 | 98 | 93 | 99 | 99 | 99 |
| 621 | ERR387402 | 1 | | 37 | 4253201 | 98 | 94 | 99 | 100 | 100 |
| 623 | ERR387405 | 1 | 017 | 37 | 4241586 | 98 | 13 | 99 | 6 | 0 |
| 641 | ERR387426 | 1 | | 37 | 4319678 | 98 | 94 | 100 | 99 | 99 |
| 642 | ERR387427 | 1 | | 37 | 4306808 | 98 | 94 | 100 | 99 | 99 |
| 643 | ERR387428 | 1 | 017 | 37 | 4253666 | 98 | 94 | 99 | 99 | 99 |
| 648 | ERR387435 | 1 | | 37 | 4301218 | 98 | 94 | 100 | 99 | 99 |
| 667 | ERR485013 | 1 | | 37 | 4254450 | 98 | 94 | 100 | 100 | 100 |
| 668 | ERR485014 | 1 | | 37 | 4240402 | 98 | 94 | 100 | 100 | 100 |
| 680 | ERR339492 | 1 | | 2 | 4295377 | 98 | 94 | 100 | 100 | 99 |
| 684 | ERR339496 | 1 | | 1 | 4187940 | 98 | 93 | 100 | 99 | 100 |
| 686 | ERR339498 | 1 | | 17 | 4236282 | 98 | 61 | 100 | 100 | 100 |
| 694 | ERR339506 | 1 | | NF | 4378453 | 98 | 94 | 100 | 100 | 99 |
| 696 | ERR339508 | 1 | | NF | 4406500 | 98 | 24 | 100 | 13 | 0 |
| 698 | ERR339510 | 1 | | 11 | 4030001 | 98 | 46 | 100 | 99 | 99 |
| 699 | ERR339511 | 1 | | 44 | 4507733 | 98 | 59 | 99 | 100 | 98 |
| 701 | ERR339513 | 1 | | 44 | 4416302 | 98 | 59 | 99 | 99 | 87 |
| 702 | ERR339514 | 1 | | 11 | 4060393 | 98 | 46 | 99 | 99 | 100 |
| 703 | ERR339515 | 1 | | NF | 4407927 | 98 | 65 | 99 | 100 | 83 |
| 707 | ERR339519 | 1 | | NF | 4179443 | 98 | 95 | 99 | 99 | 99 |
| 709 | ERR339521 | 1 | | 1 | 4225287 | 98 | 93 | 99 | 100 | 99 |
| 710 | ERR339522 | 1 | | 42 | 4126860 | 98 | 95 | 99 | 99 | 99 |
| 715 | ERR339527 | 1 | | 1 | 4159803 | 98 | 93 | 100 | 99 | 100 |
| 716 | ERR339528 | 1 | | 1 | 4163968 | 98 | 93 | 100 | 99 | 99 |
| 720 | ERR339532 | 1 | | 2 | 4140334 | 98 | 26 | 100 | 13 | 0 |
| 725 | ERR339537 | 1 | | 2 | 4322301 | 98 | 14 | 99 | 10 | 0 |
| 729 | ERR339541 | 1 | | 3 | 4271418 | 98 | 62 | 99 | 99 | 99 |
| 730 | ERR339542 | 1 | | 36 | 4188821 | 98 | 94 | 99 | 100 | 100 |
| 734 | ERR339546 | 1 | | 36 | 4186284 | 98 | 94 | 99 | 100 | 99 |
| 737 | ERR339549 | 1 | | NF | 4237617 | 98 | 92 | 99 | 99 | 99 |
| 738 | ERR339550 | 1 | | NF | 4118453 | 98 | 94 | 99 | 100 | 99 |
| 739 | ERR339551 | 1 | | 66 | 4305333 | 98 | 94 | 99 | 100 | 98 |
| 740 | ERR339552 | 1 | | 45 | 4260251 | 98 | 94 | 100 | 99 | 100 |
| 741 | ERR339553 | 1 | | 66 | 4291159 | 98 | 94 | 99 | 100 | 98 |
| 743 | ERR339555 | 1 | | 11 | 4042981 | 98 | 48 | 99 | 99 | 99 |
| 746 | ERR339558 | 1 | | 9 | 4343540 | 98 | 20 | 99 | 0 | 0 |
| 749 | ERR339561 | 1 | | 14 | 4258600 | 98 | 93 | 99 | 100 | 99 |
| 750 | ERR339562 | 1 | | 16 | 4418944 | 98 | 94 | 100 | 99 | 100 |
| 752 | ERR339564 | 1 | | 13 | 4319393 | 98 | 92 | 100 | 99 | 99 |
| 754 | ERR339566 | 1 | | 9 | 4336326 | 98 | 39 | 99 | 99 | 95 |
| 756 | ERR339568 | 1 | | NF | 4345938 | 98 | 94 | 99 | 99 | 100 |
| 758 | ERR339570 | 1 | | 9 | 4262770 | 98 | 12 | 100 | 0 | 0 |
| 759 | ERR339571 | 1 | | 14 | 4177263 | 98 | 93 | 100 | 99 | 100 |
| 760 | ERR339572 | 1 | | 36 | 4273282 | 98 | 94 | 100 | 100 | 99 |
| 762 | ERR339574 | 1 | | 36 | 4435505 | 98 | 94 | 99 | 100 | 99 |
| 763 | ERR339575 | 1 | | 9 | 4283109 | 98 | 28 | 99 | 16 | 45 |
| 764 | ERR339576 | 1 | | 7 | 4135707 | 98 | 17 | 100 | 0 | 0 |

| | | | | | | | | | |
|---|---|---|---|---|---|---|---|---|---|
| 765 | ERR339577 | 1 | NF | 4304263 | 98 | 18 | 100 | 0 | 45 |
| 767 | ERR339579 | 1 | 14 | 4156754 | 98 | 16 | 99 | 0 | 0 |
| 768 | ERR339580 | 1 | NF | 4406483 | 98 | 57 | 99 | 75 | 0 |
| 770 | ERR339582 | 1 | NF | 4279350 | 98 | 31 | 100 | 25 | 0 |
| 771 | ERR339583 | 1 | 14 | 4190409 | 98 | 93 | 100 | 99 | 100 |
| 781 | ERR339593 | 1 | 33 | 4794917 | 98 | 94 | 99 | 99 | 99 |
| 782 | ERR339594 | 1 | NF | 4179185 | 98 | 92 | 100 | 99 | 99 |
| 786 | ERR339598 | 1 | NF | 4215954 | 98 | 92 | 100 | 99 | 99 |
| 790 | ERR339602 | 1 | 13 | 4276416 | 98 | 92 | 100 | 99 | 99 |
| 791 | ERR339603 | 1 | 33 | 4261645 | 98 | 94 | 99 | 99 | 99 |
| 793 | ERR339605 | 1 | 14 | 4166090 | 98 | 92 | 99 | 99 | 99 |
| 795 | ERR339607 | 1 | 7 | 4115712 | 98 | 19 | 99 | 9 | 45 |
| 801 | ERR339613 | 1 | 36 | 4187602 | 98 | 94 | 100 | 99 | 100 |
| 803 | ERR339615 | 1 | 7 | 4188244 | 98 | 15 | 99 | 0 | 0 |
| 805 | ERR339617 | 1 | 45 | 4428310 | 98 | 94 | 99 | 99 | 99 |
| 810 | ERR339622 | 1 | 36 | 4188400 | 98 | 94 | 99 | 100 | 100 |
| 811 | ERR339623 | 1 | 13 | 4199670 | 98 | 92 | 99 | 100 | 99 |
| 813 | ERR339625 | 1 | 37 | 4249286 | 98 | 94 | 99 | 99 | 100 |
| 816 | ERR339628 | 1 | NF | 4230325 | 98 | 92 | 99 | 99 | 99 |
| 817 | ERR339629 | 1 | 14 | 4216617 | 98 | 94 | 100 | 99 | 99 |
| 819 | ERR339631 | 1 | 7 | 4415019 | 98 | 21 | 99 | 26 | 0 |
| 821 | ERR339633 | 1 | 36 | 4140430 | 98 | 94 | 100 | 99 | 99 |
| 827 | ERR339639 | 1 | 13 | 4281564 | 98 | 92 | 99 | 100 | 100 |
| 829 | ERR339641 | 1 | NF | 4393317 | 98 | 92 | 99 | 100 | 100 |
| 832 | ERR339644 | 1 | 14 | 4185772 | 98 | 92 | 99 | 99 | 99 |
| 834 | ERR339646 | 1 | 271 | 4208375 | 98 | 64 | 100 | 99 | 98 |
| 842 | ERR339654 | 1 | 5 | 4190366 | 98 | 42 | 99 | 99 | 99 |
| 843 | ERR339655 | 1 | 5 | 4317001 | 98 | 40 | 100 | 99 | 98 |
| 845 | ERR339657 | 1 | 3 | 4321248 | 98 | 18 | 99 | 34 | 0 |
| 851 | ERR339663 | 1 | 36 | 4378940 | 98 | 94 | 99 | 99 | 100 |
| 860 | ERR339672 | 1 | 17 | 4282347 | 98 | 62 | 100 | 100 | 95 |
| 862 | ERR339674 | 1 | 1 | 4169422 | 98 | 94 | 100 | 99 | 99 |
| 864 | ERR339676 | 1 | 1 | 4159116 | 98 | 94 | 100 | 99 | 100 |
| 878 | ERR339690 | 1 | 8 | 4070706 | 98 | 95 | 100 | 100 | 100 |
| 879 | ERR339691 | 1 | 8 | 4123961 | 98 | 95 | 100 | 99 | 99 |
| 880 | ERR339692 | 1 | 8 | 4296773 | 98 | 54 | 99 | 100 | 99 |
| 881 | ERR339693 | 1 | 8 | 4241319 | 98 | 95 | 100 | 99 | 100 |
| 885 | ERR339697 | 1 | 8 | 4075827 | 98 | 95 | 100 | 100 | 99 |
| 886 | ERR339698 | 1 | 8 | 4242790 | 98 | 95 | 100 | 99 | 99 |
| 888 | ERR339700 | 1 | 8 | 4286274 | 98 | 52 | 99 | 99 | 87 |
| 890 | ERR339702 | 1 | 1 | 4162065 | 98 | 93 | 100 | 99 | 99 |
| 892 | ERR339704 | 1 | 8 | 4314271 | 98 | 95 | 100 | 99 | 100 |
| 893 | ERR339705 | 1 | 8 | 4418754 | 98 | 91 | 100 | 99 | 96 |
| 894 | ERR339706 | 1 | 8 | 4146289 | 98 | 95 | 100 | 100 | 99 |
| 895 | ERR339707 | 1 | 8 | 4297488 | 98 | 95 | 99 | 100 | 100 |
| 900 | ERR339712 | 1 | 8 | 4183816 | 98 | 95 | 99 | 99 | 99 |
| 901 | ERR339713 | 1 | 1 | 4157557 | 98 | 93 | 100 | 99 | 100 |
| 902 | ERR339714 | 1 | 1 | 4166941 | 98 | 93 | 99 | 99 | 100 |
| 903 | ERR339715 | 1 | 1 | 4216632 | 98 | 93 | 100 | 99 | 98 |
| 904 | ERR339716 | 1 | 8 | 4086646 | 98 | 95 | 99 | 100 | 100 |
| 908 | ERR339720 | 1 | 8 | 4363675 | 98 | 95 | 100 | 100 | 99 |
| 909 | ERR339721 | 1 | 8 | 4154118 | 98 | 95 | 99 | 100 | 100 |
| 919 | ERR339731 | 1 | 5 | 4300435 | 98 | 40 | 99 | 99 | 98 |
| 923 | ERR339735 | 1 | 5 | 4278387 | 98 | 40 | 100 | 99 | 99 |
| 930 | ERR339742 | 1 | 5 | 4277825 | 98 | 39 | 100 | 98 | 99 |
| 933 | ERR339745 | 1 | 5 | 4197842 | 98 | 40 | 100 | 100 | 92 |
| 936 | ERR339748 | 1 | 6 | 4360445 | 98 | 60 | 99 | 100 | 99 |
| 938 | ERR339750 | 1 | 1 | 4154974 | 98 | 93 | 100 | 99 | 100 |
| 940 | ERR339752 | 1 | 1 | 4149738 | 98 | 93 | 100 | 100 | 99 |
| 943 | ERR339755 | 1 | 1 | 4155473 | 98 | 93 | 100 | 99 | 99 |
| 944 | ERR339756 | 1 | 1 | 4158693 | 98 | 93 | 100 | 100 | 100 |
| 947 | ERR339759 | 1 | 1 | 4150201 | 98 | 93 | 100 | 99 | 100 |
| 948 | ERR339760 | 1 | 1 | 4142635 | 98 | 93 | 99 | 99 | 99 |
| 953 | ERR339765 | 1 | 1 | 4160566 | 98 | 94 | 99 | 99 | 100 |
| 954 | ERR339766 | 1 | 3 | 4341770 | 98 | 29 | 100 | 20 | 0 |
| 963 | ERR339775 | 1 | 18 | 4409013 | 98 | 69 | 99 | 73 | 89 |
| 964 | ERR339776 | 1 | 1 | 4180541 | 98 | 94 | 100 | 99 | 100 |
| 969 | ERR339781 | 1 | 1 | 4145711 | 98 | 93 | 100 | 99 | 99 |
| 976 | ERR339788 | 1 | 46 | 4340518 | 98 | 94 | 100 | 99 | 99 |
| 979 | ERR339791 | 1 | 1 | 4199085 | 98 | 94 | 99 | 100 | 100 |
| 983 | ERR339795 | 1 | 1 | 4187481 | 98 | 94 | 99 | 99 | 99 |
| 988 | ERR339800 | 1 | 1 | 4230014 | 98 | 93 | 100 | 99 | 100 |
| 990 | ERR339802 | 1 | 49 | 4745063 | 98 | 92 | 100 | 99 | 100 |
| 993 | ERR339805 | 1 | NF | 4496993 | 98 | 94 | 100 | 99 | 99 |
| 996 | ERR339808 | 1 | 55 | 4074340 | 98 | 94 | 100 | 99 | 100 |
| 997 | ERR339809 | 1 | 45 | 4509290 | 98 | 94 | 100 | 99 | 99 |
| 1000 | ERR339812 | 1 | NF | 4541577 | 98 | 94 | 100 | 99 | 99 |
| 1001 | ERR339813 | 1 | 49 | 4424293 | 98 | 92 | 99 | 99 | 100 |
| 1005 | ERR339817 | 1 | 49 | 4294030 | 98 | 94 | 100 | 100 | 99 |
| 1007 | ERR339819 | 1 | 45 | 4449263 | 98 | 94 | 100 | 99 | 99 |
| 1010 | ERR339822 | 1 | 37 | 4076616 | 98 | 94 | 99 | 100 | 99 |
| 1019 | ERR339831 | 1 | 1 | 4179430 | 98 | 93 | 100 | 99 | 100 |
| 1020 | ERR339832 | 1 | 1 | 4144531 | 98 | 93 | 100 | 99 | 100 |
| 1023 | ERR339835 | 1 | 1 | 4147450 | 98 | 93 | 100 | 100 | 99 |
| 1025 | ERR339837 | 1 | 1 | 4147661 | 98 | 93 | 99 | 100 | 99 |
| 1029 | ERR339841 | 1 | 1 | 4179826 | 98 | 93 | 99 | 100 | 99 |
| 1032 | ERR339844 | 1 | 1 | 4152257 | 98 | 93 | 99 | 99 | 100 |
| 1034 | ERR339846 | 1 | 1 | 4166139 | 98 | 93 | 100 | 100 | 100 |
| 1036 | ERR339848 | 1 | 1 | 4154379 | 98 | 94 | 100 | 99 | 100 |
| 1042 | ERR339854 | 1 | 1 | 4141727 | 98 | 93 | 99 | 99 | 100 |
| 1045 | ERR339857 | 1 | 1 | 4149370 | 98 | 93 | 100 | 99 | 99 |
| 1052 | ERR339864 | 1 | 1 | 4088940 | 98 | 93 | 100 | 99 | 99 |
| 1053 | ERR339866 | 1 | 1 | 4146325 | 98 | 94 | 99 | 100 | 100 |
| 1056 | ERR339869 | 1 | 1 | 4152155 | 98 | 93 | 100 | 99 | 100 |
| 1058 | ERR339871 | 1 | 1 | 4145827 | 98 | 94 | 99 | 99 | 100 |
| 1059 | ERR339872 | 1 | 1 | 4144412 | 98 | 93 | 99 | 100 | 100 |
| 1060 | ERR339873 | 1 | 1 | 4155992 | 98 | 93 | 100 | 99 | 100 |
| 1064 | ERR339877 | 1 | 1 | 4178444 | 98 | 93 | 99 | 99 | 99 |
| 1071 | ERR339884 | 1 | 1 | 4147252 | 98 | 93 | 100 | 99 | 99 |
| 1072 | ERR339885 | 1 | 1 | 4151064 | 98 | 93 | 99 | 99 | 99 |
| 1076 | ERR339889 | 1 | 1 | 4156151 | 98 | 93 | 99 | 99 | 100 |
| 1078 | ERR339891 | 1 | 1 | 4142624 | 98 | 93 | 99 | 99 | 100 |
| 1079 | ERR339892 | 1 | 1 | 4143645 | 98 | 93 | 100 | 99 | 99 |
| 1082 | ERR339895 | 1 | 1 | 4148671 | 98 | 94 | 100 | 99 | 100 |
| 1089 | ERR339902 | 1 | 1 | 4143808 | 98 | 93 | 100 | 99 | 99 |
| 1090 | ERR339903 | 1 | 1 | 4151299 | 98 | 94 | 100 | 100 | 100 |
| 1093 | ERR339906 | 1 | 1 | 4153139 | 98 | 93 | 99 | 99 | 100 |
| 1105 | ERR339918 | 1 | 1 | 4152796 | 98 | 94 | 99 | 99 | 99 |
| 1113 | ERR339926 | 1 | 1 | 4137932 | 98 | 94 | 100 | 99 | 100 |
| 1118 | ERR339931 | 1 | 1 | 4149963 | 98 | 93 | 99 | 99 | 100 |
| 1120 | ERR339933 | 1 | 1 | 4170554 | 98 | 93 | 100 | 99 | 99 |
| 1121 | ERR339934 | 1 | 1 | 4171858 | 98 | 94 | 100 | 99 | 100 |
| 1123 | ERR339936 | 1 | 1 | 4149782 | 98 | 93 | 100 | 100 | 99 |
| 1126 | ERR339939 | 1 | 1 | 4155265 | 98 | 94 | 99 | 99 | 99 |
| 1127 | ERR339940 | 1 | 1 | 4148549 | 98 | 93 | 100 | 100 | 100 |
| 1128 | ERR339942 | 1 | 1 | 4143200 | 98 | 94 | 99 | 99 | 99 |
| 1131 | ERR339945 | 1 | 1 | 4145799 | 98 | 93 | 99 | 100 | 100 |
| 1137 | ERR339951 | 1 | 1 | 4147866 | 98 | 93 | 99 | 100 | 100 |
| 1140 | ERR339954 | 1 | 1 | 4147243 | 98 | 93 | 100 | 99 | 99 |
| 1141 | ERR339955 | 1 | 1 | 4159244 | 98 | 94 | 99 | 100 | 100 |
| 1142 | ERR339956 | 1 | 1 | 4630543 | 98 | 95 | 100 | 99 | 100 |
| 1144 | ERR339958 | 1 | 72 | 4351479 | 98 | 65 | 99 | 99 | 96 |
| 1146 | ERR339960 | 1 | 21 | 4132989 | 98 | 94 | 100 | 100 | 100 |
| 1149 | ERR339963 | 1 | 67 | 4313209 | 98 | 94 | 99 | 99 | 100 |
| 1151 | ERR339965 | 1 | 4 | 4047343 | 98 | 94 | 99 | 99 | 99 |
| 1154 | ERR339968 | 1 | 2 | 4194734 | 98 | 20 | 100 | 26 | 0 |
| 1155 | ERR339969 | 1 | 2 | 4120046 | 98 | 92 | 100 | 99 | 99 |
| 1159 | ERR339973 | 1 | 8 | 4062088 | 98 | 56 | 99 | 99 | 99 |
| 1161 | ERR339975 | 1 | 65 | 4150868 | 98 | 95 | 100 | 99 | 99 |
| 1164 | ERR339978 | 1 | 2 | 4194538 | 98 | 28 | 100 | 40 | 45 |
| 1170 | ERR339984 | 1 | 8 | 4089493 | 98 | 95 | 99 | 99 | 100 |
| 1172 | ERR339986 | 1 | 2 | 4224579 | 98 | 95 | 99 | 99 | 99 |
| 1175 | ERR339989 | 1 | 66 | 4270972 | 98 | 94 | 100 | 99 | 99 |
| 1177 | ERR339991 | 1 | 42 | 4127461 | 98 | 95 | 100 | 99 | 100 |
| 1178 | ERR339992 | 1 | 42 | 4130209 | 98 | 95 | 100 | 100 | 99 |
| 1180 | ERR339994 | 1 | 44 | 4622474 | 98 | 67 | 100 | 99 | 98 |
| 1184 | ERR339998 | 1 | 2 | 4150863 | 98 | 92 | 100 | 100 | 100 |
| 1187 | ERR340001 | 1 | 2 | 4111971 | 98 | 93 | 99 | 99 | 100 |
| 1189 | ERR340003 | 1 | 75 | 4231377 | 98 | 62 | 100 | 99 | 99 |
| 1190 | ERR340004 | 1 | 2 | 4132787 | 98 | 92 | 100 | 99 | 99 |
| 1193 | ERR340007 | 1 | 2 | 4195130 | 98 | 20 | 99 | 13 | 0 |
| 1197 | ERR340011 | 1 | 2 | 4135790 | 98 | 92 | 100 | 99 | 99 |
| 1198 | ERR340012 | 1 | 2 | 4310356 | 98 | 94 | 100 | 100 | 100 |
| 1205 | ERR340019 | 1 | 8 | 4163120 | 98 | 52 | 99 | 99 | 93 |
| 1208 | ERR340023 | 1 | 2 | 4208550 | 98 | 20 | 99 | 0 | 0 |
| 1211 | ERR340026 | 1 | 78 | 4322636 | 98 | 94 | 99 | 99 | 99 |
| 1212 | ERR340027 | 1 | 68 | 4133114 | 98 | 17 | 100 | 13 | 0 |
| 1213 | ERR340028 | 1 | 2 | 4213526 | 98 | 14 | 99 | 13 | 0 |
| 1214 | ERR340029 | 1 | 9 | 4331731 | 98 | 60 | 100 | 99 | 93 |
| 1216 | ERR340031 | 1 | 91 | 4353181 | 98 | 33 | 100 | 26 | 0 |
| 1224 | ERR340039 | 1 | 8 | 4251817 | 98 | 95 | 100 | 100 | 100 |
| 1227 | ERR340042 | 1 | 8 | 4781695 | 98 | 95 | 99 | 99 | 99 |
| 1236 | ERR340051 | 1 | 8 | 4308526 | 98 | 95 | 99 | 99 | 99 |
| 1241 | ERR340056 | 1 | 8 | 4284551 | 98 | 49 | 100 | 99 | 94 |
| 1247 | ERR340062 | 1 | 8 | 4229529 | 98 | 95 | 100 | 99 | 99 |
| 1251 | ERR340066 | 1 | 23 | 4275423 | 98 | 92 | 100 | 99 | 99 |
| 1259 | ERR340074 | 1 | 17 | 4454749 | 98 | 60 | 99 | 99 | 97 |
| 1260 | ERR340075 | 1 | 1 | 4377570 | 98 | 94 | 100 | 100 | 99 |
| 1272 | ERR340087 | 1 | 24 | 4358265 | 98 | 16 | 99 | 0 | 0 |
| 1275 | ERR340090 | 1 | 46 | 4377047 | 98 | 94 | 99 | 99 | 100 |
| 1277 | ERR340092 | 1 | 7 | 4172454 | 98 | 62 | 100 | 99 | 99 |
| 1291 | ERR340106 | 1 | 2 | 4170634 | 98 | 91 | 100 | 100 | 100 |
| 1297 | ERR340112 | 1 | 2 | 4236148 | 98 | 14 | 100 | 0 | 0 |
| 1301 | ERR340116 | 1 | 6 | 4427217 | 98 | 65 | 99 | 100 | 99 |
| 1303 | ERR340118 | 1 | 44 | 4501386 | 98 | 63 | 99 | 100 | 95 |
| 1306 | ERR340121 | 1 | 46 | 4353613 | 98 | 94 | 99 | 99 | 99 |
| 1310 | ERR340125 | 1 | 16 | 4496031 | 98 | 12 | 99 | 13 | 0 |
| 1311 | ERR340126 | 1 | 248 | 4095584 | 98 | 64 | 100 | 100 | 94 |
| 1312 | ERR340127 | 1 | 1 | 4152655 | 98 | 94 | 99 | 99 | 100 |
| 1314 | ERR340129 | 1 | 44 | 4400051 | 98 | 28 | 99 | 32 | 45 |
| 1318 | ERR340133 | 1 | 35 | 4322752 | 98 | 23 | 100 | 0 | 0 |
| 1319 | ERR340134 | 1 | 248 | 4098623 | 98 | 64 | 99 | 100 | 95 |
| 1320 | ERR340135 | 1 | 13 | 4543281 | 98 | 98 | 100 | 100 | 100 |
| 1323 | ERR340138 | 1 | 9 | 4340089 | 98 | 27 | 100 | 21 | 0 |
| 1326 | ERR340141 | 1 | 8 | 4233324 | 98 | 95 | 99 | 99 | 99 |
| 1327 | ERR340142 | 1 | 1 | 4176635 | 98 | 94 | 99 | 99 | 99 |
| 1328 | ERR340143 | 1 | 6 | 4381546 | 98 | 21 | 99 | 22 | 0 |
| 1329 | ERR340144 | 1 | 11 | 4040975 | 98 | 51 | 99 | 99 | 99 |
| 1332 | ERR340147 | 1 | 1 | 4140719 | 98 | 94 | 99 | 100 | 99 |
| 1334 | ERR340149 | 1 | 14 | 4175858 | 98 | 92 | 99 | 100 | 99 |
| 1335 | ERR340150 | 1 | 8 | 4064415 | 98 | 49 | 100 | 100 | 92 |
| 1336 | ERR340151 | 1 | 26 | 4259244 | 98 | 95 | 100 | 99 | 100 |
| 1337 | ERR340152 | 1 | 11 | 4021903 | 98 | 47 | 99 | 99 | 99 |
| 1339 | ERR340154 | 1 | 8 | 4274264 | 98 | 95 | 99 | 99 | 99 |
| 1340 | ERR340155 | 1 | 2 | 4138393 | 98 | 93 | 100 | 100 | 100 |
| 1341 | ERR340156 | 1 | 9 | 4356355 | 98 | 18 | 100 | 0 | 0 |
| 1343 | ERR340158 | 1 | 8 | 4308596 | 98 | 95 | 100 | 99 | 99 |

| | | | | | | | | | | |
|---|---|---|---|---|---|---|---|---|---|---|
| 1345 | ERR340160 | 1 | | 8 | 4289095 | 98 | 95 | 100 | 99 | 100 |
| 1348 | ERR340163 | 1 | | 51 | 4143136 | 98 | 17 | 100 | 0 | 45 |
| 1350 | ERR340165 | 1 | | 13 | 4141461 | 98 | 92 | 99 | 99 | 100 |
| 1351 | ERR340166 | 1 | | 8 | 4209055 | 98 | 95 | 99 | 99 | 99 |
| 1352 | ERR340167 | 1 | | 8 | 4105411 | 98 | 63 | 99 | 99 | 94 |
| 1353 | ERR340168 | 1 | | 11 | 4112630 | 98 | 51 | 99 | 100 | 100 |
| 1356 | ERR340171 | 1 | | 5 | 4247037 | 98 | 49 | 99 | 99 | 92 |
| 1358 | ERR340173 | 1 | | 36 | 4117462 | 98 | 18 | 99 | 19 | 66 |
| 1362 | ERR340177 | 1 | | 11 | 3962633 | 98 | 53 | 99 | 100 | 99 |
| 1365 | ERR340180 | 1 | | 5 | 4303457 | 98 | 46 | 100 | 100 | 98 |
| 1375 | ERR340190 | 1 | | 16 | 4543394 | 98 | 95 | 99 | 99 | 100 |
| 1377 | ERR340192 | 1 | | 51 | 4113685 | 98 | 24 | 99 | 35 | 0 |
| 1379 | ERR340194 | 1 | | 1 | 4428116 | 98 | 94 | 100 | 99 | 100 |
| 1380 | ERR340195 | 1 | | 92 | 4138706 | 98 | 94 | 100 | 99 | 99 |
| 1381 | ERR340196 | 1 | | 45 | 4349204 | 98 | 94 | 100 | 99 | 100 |
| 1392 | ERR340207 | 1 | | 49 | 4166642 | 98 | 92 | 100 | 100 | 99 |
| 1393 | ERR340208 | 1 | | 13 | 4259758 | 98 | 92 | 100 | 100 | 99 |
| 1395 | ERR340210 | 1 | | 8 | 4229497 | 98 | 95 | 99 | 99 | 98 |
| 1396 | ERR340211 | 1 | | 1 | 4367542 | 98 | 94 | 99 | 100 | 99 |
| 1397 | ERR340212 | 1 | | 49 | 4271251 | 98 | 92 | 100 | 99 | 100 |
| 1399 | ERR340214 | 1 | | 49 | 4268411 | 98 | 92 | 99 | 100 | 100 |
| 1402 | ERR340217 | 1 | | 2 | 4188054 | 98 | 92 | 99 | 99 | 99 |
| 1403 | ERR340218 | 1 | | 44 | 4268099 | 98 | 19 | 100 | 0 | 0 |
| 1412 | ERR340227 | 1 | | 53 | 4159181 | 98 | 93 | 99 | 99 | 99 |
| 1414 | ERR340229 | 1 | | 8 | 4399151 | 98 | 47 | 99 | 99 | 88 |
| 1415 | ERR340230 | 1 | | 2 | 4155135 | 98 | 16 | 100 | 0 | 45 |
| 1417 | ERR340232 | 1 | | 49 | 4174696 | 98 | 92 | 100 | 99 | 100 |
| 1423 | ERR340238 | 1 | | 11 | 3964412 | 98 | 49 | 100 | 99 | 99 |
| 1424 | ERR340239 | 1 | | 8 | 4301993 | 98 | 95 | 99 | 99 | 100 |
| 1427 | ERR340242 | 1 | | 13 | 4372231 | 98 | 93 | 99 | 99 | 100 |
| 1428 | ERR340243 | 1 | | 37 | 4239353 | 98 | 94 | 99 | 100 | 99 |
| 1430 | ERR340245 | 1 | | 37 | 4304095 | 98 | 93 | 100 | 99 | 99 |
| 1431 | ERR340246 | 1 | | 17 | 4518923 | 98 | 95 | 99 | 100 | 99 |
| 1434 | ERR340249 | 1 | | 7 | 4180269 | 98 | 12 | 100 | 0 | 0 |
| 1436 | ERR340251 | 1 | | 98 | 4258311 | 98 | 60 | 99 | 99 | 99 |
| 1441 | ERR340256 | 1 | | 8 | 4202940 | 98 | 95 | 100 | 99 | 99 |
| 1442 | ERR340257 | 1 | | 92 | 4283898 | 98 | 94 | 99 | 100 | 99 |
| 1445 | ERR340260 | 1 | | 8 | 4473427 | 98 | 53 | 100 | 100 | 99 |
| 1447 | ERR340262 | 1 | | 49 | 4357086 | 98 | 92 | 99 | 99 | 98 |
| 1448 | ERR340263 | 1 | | 13 | 4270207 | 98 | 92 | 99 | 99 | 99 |
| 1449 | ERR340264 | 1 | | 2 | 4149456 | 98 | 13 | 99 | 0 | 0 |
| 1450 | ERR340265 | 1 | | 112 | 4282982 | 98 | 20 | 99 | 13 | 0 |
| 1451 | ERR340266 | 1 | | 35 | 4357586 | 98 | 17 | 99 | 0 | 0 |
| 1453 | ERR340268 | 1 | | 1 | 4351195 | 98 | 94 | 99 | 99 | 99 |
| 1454 | ERR340269 | 1 | | 2 | 4143393 | 98 | 23 | 99 | 33 | 0 |
| 1456 | ERR340271 | 1 | | 14 | 4170788 | 98 | 92 | 100 | 99 | 99 |
| 1459 | ERR340274 | 1 | | 37 | 4238785 | 98 | 94 | 100 | 99 | 99 |
| 1461 | ERR340276 | 1 | | 16 | 4563447 | 98 | 10 | 99 | 0 | 0 |
| 1462 | ERR340277 | 1 | | 1 | 4372947 | 98 | 94 | 100 | 99 | 100 |
| 1467 | ERR340282 | 1 | | 11 | 4032569 | 98 | 60 | 100 | 99 | 100 |
| 1469 | ERR340284 | 1 | | 17 | 4316519 | 98 | 68 | 100 | 99 | 92 |
| 1479 | ERR340294 | 1 | | 45 | 4698355 | 98 | 94 | 100 | 99 | 100 |
| 1480 | ERR340295 | 1 | | 17 | 4187862 | 98 | 63 | 99 | 100 | 94 |
| 1483 | ERR340298 | 1 | | 16 | 4502943 | 98 | 95 | 99 | 100 | 99 |
| 1487 | ERR340302 | 1 | | 1 | 4145239 | 98 | 93 | 99 | 99 | 100 |
| 1488 | ERR340303 | 1 | | 1 | 4137333 | 98 | 94 | 99 | 99 | 99 |
| 1489 | ERR340304 | 1 | | 1 | 4142195 | 98 | 93 | 99 | 100 | 100 |
| 1490 | ERR340305 | 1 | | 1 | 4142710 | 98 | 94 | 99 | 99 | 100 |
| 1492 | ERR340307 | 1 | | 2 | 4161414 | 98 | 92 | 100 | 100 | 99 |
| 1493 | ERR340308 | 1 | | 2 | 4248847 | 98 | 92 | 100 | 99 | 99 |
| 1496 | ERR340311 | 1 | | 2 | 4137778 | 98 | 22 | 100 | 13 | 0 |
| 1502 | ERR340317 | 1 | | 3 | 4258983 | 98 | 63 | 99 | 99 | 99 |
| 1503 | ERR340318 | 1 | | 11 | 5186708 | 98 | 53 | 99 | 100 | 100 |
| 1506 | ERR126243 | 1 | | 4 | 4062529 | 98 | 97 | 99 | 100 | 99 |
| 1507 | ERR126244 | 1 | | 45 | 4359689 | 98 | 94 | 99 | 99 | 99 |
| 1510 | ERR126247 | 1 | 015 | 1 | 4204169 | 98 | 17 | 99 | 13 | 0 |
| 1523 | ERR126260 | 1 | 054 | 43 | 4219640 | 98 | 22 | 99 | 7 | 0 |
| 1532 | ERR126269 | 1 | | 41 | 4251230 | 98 | 94 | 100 | 100 | 99 |
| 1533 | ERR126270 | 1 | 078 | 11 | 4040669 | 98 | 48 | 100 | 99 | 99 |
| 1534 | ERR126271 | 1 | 193 | 11 | 4064114 | 98 | 51 | 99 | 100 | 100 |
| 1541 | ERR126278 | 1 | | NF | 4401870 | 98 | 99 | 99 | 100 | 99 |
| 1545 | ERR126282 | 1 | | 36 | 4221732 | 98 | 94 | 99 | 99 | 99 |
| 1546 | ERR126283 | 1 | | 66 | 4287777 | 98 | 94 | 99 | 99 | 99 |
| 1547 | ERR126284 | 1 | 029 | 16 | 4398429 | 98 | 95 | 100 | 99 | 100 |
| 1550 | ERR126287 | 1 | | 15 | 4215605 | 98 | 94 | 100 | 99 | 99 |
| 1554 | ERR126291 | 1 | | 23 | 4326404 | 98 | 94 | 99 | 99 | 100 |
| 1563 | ERR142054 | 1 | | 2 | 4190913 | 98 | 92 | 99 | 100 | 100 |
| 1567 | ERR142058 | 1 | | 44 | 4199528 | 98 | 96 | 100 | 100 | 100 |
| 1568 | ERR142059 | 1 | | 1 | 4153828 | 98 | 95 | 100 | 99 | 99 |
| 1569 | ERR142060 | 1 | | 44 | 4397638 | 98 | 63 | 100 | 99 | 99 |
| 1573 | ERR142064 | 1 | | 1 | 4140019 | 98 | 94 | 100 | 100 | 99 |
| 1575 | ERR142066 | 1 | | 42 | 4155468 | 98 | 97 | 100 | 100 | 100 |
| 1576 | ERR142067 | 1 | | 42 | 4141529 | 98 | 96 | 99 | 100 | 100 |
| 1578 | ERR142069 | 1 | | 1 | 4134579 | 98 | 93 | 100 | 99 | 100 |
| 1579 | ERR142070 | 1 | | 58 | 4222212 | 98 | 62 | 100 | 99 | 94 |
| 1582 | ERR142073 | 1 | | 1 | 4175510 | 98 | 93 | 99 | 100 | 100 |
| 1583 | ERR142074 | 1 | | 3 | 4373426 | 98 | 64 | 99 | 100 | 100 |
| 1586 | ERR142077 | 1 | | 44 | 4483779 | 98 | 74 | 99 | 100 | 98 |
| 1597 | ERR142088 | 1 | | 1 | 4199622 | 98 | 93 | 99 | 99 | 100 |
| 1598 | ERR142089 | 1 | | 6 | 4435371 | 98 | 15 | 99 | 0 | 0 |
| 1601 | ERR142092 | 1 | | 1 | 4157161 | 98 | 93 | 100 | 100 | 100 |
| 1603 | ERR142094 | 1 | | 57 | 4255773 | 98 | 16 | 99 | 0 | 0 |
| 1608 | ERR142099 | 1 | | 1 | 4198849 | 98 | 92 | 100 | 99 | 99 |
| 1611 | ERR142102 | 1 | | 1 | 4194320 | 98 | 93 | 100 | 99 | 100 |
| 1614 | ERR142105 | 1 | | NF | 4262283 | 98 | 64 | 100 | 99 | 98 |
| 1616 | ERR142107 | 1 | | 1 | 4166354 | 98 | 95 | 100 | 99 | 100 |
| 1617 | ERR142108 | 1 | | 1 | 4132912 | 98 | 95 | 100 | 99 | 99 |
| 1632 | ERR142123 | 1 | | 6 | 4326015 | 98 | 67 | 99 | 99 | 94 |
| 1636 | ERR142127 | 1 | | 1 | 4141796 | 98 | 94 | 100 | 100 | 100 |
| 1640 | ERR142131 | 1 | | 58 | 4223264 | 98 | 64 | 100 | 100 | 98 |
| 1641 | ERR142132 | 1 | | 8 | 4428461 | 98 | 95 | 100 | 100 | 100 |
| 1646 | ERR142137 | 1 | | 1 | 4212832 | 98 | 92 | 99 | 100 | 100 |
| 1674 | ERR142165 | 1 | | 1 | 4155301 | 98 | 93 | 100 | 99 | 99 |
| 1675 | ERR142166 | 1 | | 17 | 4467340 | 98 | 63 | 100 | 100 | 100 |
| 1679 | ERR142170 | 1 | | 17 | 4246707 | 98 | 59 | 100 | 100 | 98 |
| 1680 | ERR142171 | 1 | | 17 | 4211795 | 98 | 60 | 100 | 100 | 99 |
| 1686 | ERR142177 | 1 | | 1 | 4157475 | 98 | 93 | 100 | 100 | 100 |
| 1688 | ERR142179 | 1 | | 17 | 4270841 | 98 | 60 | 100 | 99 | 100 |
| 1691 | ERR142182 | 1 | | 54 | 4530031 | 98 | 100 | 100 | 100 | 100 |
| 1701 | ERR142192 | 1 | | 58 | 4294057 | 98 | 60 | 99 | 99 | 99 |
| 1702 | ERR142193 | 1 | | 58 | 4095238 | 98 | 10 | 100 | 0 | 0 |
| 1709 | ERR142200 | 1 | | 14 | 4322404 | 98 | 92 | 100 | 99 | 100 |
| 1712 | ERR142203 | 1 | | 44 | 4358262 | 98 | 65 | 99 | 100 | 91 |
| 1713 | ERR142204 | 1 | | 1 | 4147375 | 98 | 93 | 99 | 100 | 100 |
| 1715 | ERR142206 | 1 | | 8 | 4279366 | 98 | 95 | 99 | 100 | 99 |
| 1717 | ERR142208 | 1 | | NF | 4087827 | 98 | 95 | 99 | 99 | 99 |
| 1718 | ERR142209 | 1 | | NF | 4150109 | 98 | 92 | 100 | 99 | 99 |
| 1719 | ERR142210 | 1 | | 1 | 4147799 | 98 | 93 | 99 | 100 | 99 |
| 1720 | ERR142211 | 1 | | NF | 4188155 | 98 | 92 | 100 | 100 | 100 |
| 1723 | ERR142214 | 1 | | 2 | 4156026 | 98 | 92 | 100 | 99 | 99 |
| 1726 | ERR142217 | 1 | | 37 | 4229030 | 98 | 94 | 100 | 99 | 99 |
| 1727 | ERR142218 | 1 | | 375 | 4181181 | 98 | 94 | 100 | 100 | 99 |
| 1729 | ERR142220 | 1 | | 37 | 4266404 | 98 | 94 | 99 | 100 | 100 |
| 1730 | ERR142221 | 1 | | 37 | 4225321 | 98 | 94 | 99 | 99 | 99 |
| 1731 | ERR142222 | 1 | | 37 | 4250581 | 98 | 94 | 99 | 100 | 100 |
| 1733 | ERR142224 | 1 | | NF | 4185925 | 98 | 95 | 99 | 99 | 99 |
| 1735 | ERR142226 | 1 | | 37 | 4246649 | 98 | 94 | 99 | 100 | 99 |
| 1738 | ERR142229 | 1 | | 1 | 4143001 | 98 | 93 | 100 | 100 | 100 |
| 1739 | ERR142230 | 1 | | 5 | 4294824 | 98 | 40 | 99 | 99 | 98 |
| 1741 | ERR142232 | 1 | | 5 | 4222460 | 98 | 39 | 99 | 99 | 94 |
| 1742 | ERR142233 | 1 | | NF | 4300139 | 98 | 36 | 100 | 99 | 92 |
| 1743 | ERR142234 | 1 | | NF | 4539326 | 98 | 43 | 100 | 99 | 98 |
| 1744 | ERR142235 | 1 | | 22 | 4440465 | 98 | 41 | 100 | 99 | 99 |
| 1745 | ERR142236 | 1 | | NF | 4258915 | 98 | 38 | 100 | 99 | 98 |
| 1747 | ERR142238 | 1 | | 11 | 4137008 | 98 | 47 | 99 | 99 | 100 |
| 1748 | ERR142239 | 1 | | 11 | 3966916 | 98 | 48 | 99 | 99 | 100 |
| 1751 | ERR142242 | 1 | | 11 | 4091879 | 98 | 47 | 99 | 99 | 99 |
| 1753 | ERR142244 | 1 | | 11 | 3990992 | 98 | 47 | 99 | 99 | 100 |
| 1755 | ERR142246 | 1 | | NF | 4133516 | 98 | 95 | 99 | 99 | 100 |
| 1757 | ERR142248 | 1 | | 11 | 4020458 | 98 | 47 | 99 | 100 | 100 |
| 1758 | ERR142249 | 1 | | 1 | 4144552 | 98 | 94 | 100 | 99 | 99 |
| 1759 | ERR142250 | 1 | | NF | 4267936 | 98 | 63 | 100 | 99 | 96 |
| 1761 | ERR142252 | 1 | | NF | 4381050 | 98 | 94 | 99 | 99 | 99 |
| 1762 | ERR142253 | 1 | | 1 | 4470683 | 98 | 81 | 99 | 92 | 51 |
| 1763 | ERR142254 | 1 | | NF | 4397882 | 98 | 94 | 99 | 99 | 99 |
| 1766 | ERR142257 | 1 | | 1 | 4385292 | 98 | 94 | 99 | 100 | 99 |
| 1768 | ERR142259 | 1 | | NF | 4480802 | 98 | 94 | 100 | 100 | 99 |
| 1769 | ERR142260 | 1 | | NF | 4381928 | 98 | 94 | 100 | 100 | 99 |
| 1773 | ERR142264 | 1 | | NF | 4371762 | 98 | 94 | 99 | 100 | 100 |
| 1774 | ERR142265 | 1 | | NF | 4370542 | 98 | 94 | 100 | 99 | 99 |
| 1776 | ERR142267 | 1 | | NF | 4377432 | 98 | 94 | 100 | 99 | 100 |
| 1777 | ERR142268 | 1 | | NF | 4382316 | 98 | 12 | 100 | 0 | 0 |
| 1779 | ERR142270 | 1 | | 1 | 4394001 | 98 | 94 | 97 | 99 | 99 |
| 1782 | ERR142273 | 1 | | NF | 4437079 | 98 | 94 | 99 | 99 | 99 |
| 1784 | ERR142275 | 1 | | 35 | 4290464 | 98 | 10 | 99 | 0 | 0 |
| 1785 | ERR142276 | 1 | | 35 | 4420876 | 98 | 12 | 100 | 0 | 0 |
| 1786 | ERR142277 | 1 | | NF | 4313578 | 98 | 13 | 99 | 0 | 0 |
| 1787 | ERR142278 | 1 | | NF | 4277293 | 98 | 13 | 99 | 0 | 0 |
| 1789 | ERR142280 | 1 | | 35 | 4334186 | 98 | 13 | 100 | 0 | 0 |
| 1791 | ERR142282 | 1 | | NF | 4523083 | 98 | 61 | 99 | 100 | 91 |
| 1792 | ERR142283 | 1 | | NF | 4493821 | 98 | 19 | 100 | 13 | 0 |
| 1793 | ERR142284 | 1 | | 44 | 4351813 | 98 | 94 | 99 | 99 | 99 |
| 1796 | ERR142287 | 1 | | NF | 4373854 | 98 | 18 | 99 | 0 | 0 |
| 1800 | ERR142291 | 1 | | 44 | 4345150 | 98 | 61 | 99 | 100 | 98 |
| 1802 | ERR142293 | 1 | | NF | 4489995 | 98 | 94 | 99 | 99 | 99 |
| 1805 | ERR142296 | 1 | | NF | 4389566 | 98 | 61 | 100 | 99 | 98 |
| 1807 | ERR142298 | 1 | | NF | 4405437 | 98 | 60 | 100 | 99 | 98 |
| 1808 | ERR142299 | 1 | | NF | 4341903 | 98 | 63 | 99 | 99 | 90 |
| 1811 | ERR142302 | 1 | | NF | 4643383 | 98 | 62 | 99 | 99 | 94 |
| 1812 | ERR142303 | 1 | | NF | 4948102 | 98 | 96 | 100 | 100 | 99 |
| 1814 | ERR142305 | 1 | | NF | 4568417 | 98 | 95 | 99 | 100 | 99 |
| 1815 | ERR142306 | 1 | | 44 | 4501669 | 98 | 62 | 99 | 100 | 97 |
| 1828 | ERR142319 | 1 | | NF | 4150588 | 98 | 58 | 100 | 99 | 81 |
| 1832 | ERR142323 | 1 | | 63 | 4173438 | 98 | 99 | 100 | 100 | 100 |
| 1836 | ERR142327 | 1 | | 63 | 4019005 | 98 | 99 | 100 | 100 | 99 |
| 1841 | ERR142332 | 1 | | 6 | 4220811 | 98 | 17 | 100 | 13 | 0 |
| 1844 | ERR142335 | 1 | | 6 | 4391708 | 98 | 14 | 100 | 0 | 0 |
| 1845 | ERR142336 | 1 | | 6 | 4623048 | 98 | 17 | 99 | 10 | 0 |
| 1846 | ERR142337 | 1 | | 6 | 4610973 | 98 | 17 | 100 | 11 | 0 |
| 1847 | ERR142338 | 1 | | 67 | 4191410 | 98 | 94 | 100 | 99 | 99 |
| 1848 | ERR142339 | 1 | | NF | 4372642 | 98 | 19 | 100 | 0 | 0 |
| 1849 | ERR142340 | 1 | | 6 | 4302481 | 98 | 18 | 100 | 0 | 0 |
| 1852 | ERR142343 | 1 | | 6 | 4333265 | 98 | 25 | 100 | 24 | 62 |
| 1853 | ERR142344 | 1 | | 6 | 4380264 | 98 | 15 | 100 | 0 | 0 |
| 1858 | ERR142349 | 1 | | 6 | 4337463 | 98 | 20 | 100 | 0 | 0 |
| 1859 | ERR142350 | 1 | | 6 | 4602502 | 98 | 34 | 99 | 34 | 0 |
| 1863 | ERR142354 | 1 | | 6 | 4362417 | 98 | 64 | 99 | 99 | 99 |

| | | | | | | | | | | |
|---|---|---|---|---|---|---|---|---|---|---|
| 1864 | ERR142355 | 1 | | 6 | 4329029 | 98 | 23 | 100 | 28 | 0 |
| 1870 | ERR142361 | 1 | | 37 | 4198618 | 98 | 94 | 100 | 100 | 99 |
| 1871 | ERR142362 | 1 | 023 | 25 | 4220062 | 98 | 39 | 100 | 99 | 99 |
| 1872 | ERR142363 | 1 | | 12 | 4255033 | 98 | 94 | 99 | 99 | 99 |
| 1873 | ERR142364 | 1 | | 12 | 4145827 | 98 | 94 | 99 | 99 | 99 |
| 1875 | ERR142366 | 1 | | 44 | 4356113 | 98 | 17 | 99 | 0 | 0 |
| 1876 | ERR142367 | 1 | | NF | 4373693 | 98 | 18 | 99 | 0 | 0 |
| 1880 | ERR142371 | 1 | | NF | 4363124 | 98 | 20 | 99 | 17 | 0 |
| 1882 | ERR142373 | 1 | | 1 | 4169149 | 98 | 93 | 100 | 99 | 99 |
| 1883 | ERR142374 | 1 | | 1 | 4393902 | 98 | 95 | 99 | 99 | 98 |
| 1884 | ERR142375 | 1 | | 11 | 4103645 | 98 | 45 | 99 | 99 | 99 |
| 1887 | ERR142378 | 1 | | NF | 4645865 | 98 | 16 | 100 | 29 | 61 |
| 1888 | ERR142379 | 1 | | 6 | 4476151 | 98 | 16 | 99 | 6 | 0 |
| 1889 | ERR142380 | 1 | | NF | 4421766 | 98 | 16 | 99 | 0 | 0 |
| 1891 | ERR142382 | 1 | | NF | 4099097 | 98 | 95 | 98 | 100 | 99 |
| 1892 | ERR142383 | 1 | | NF | 4089978 | 98 | 95 | 99 | 100 | 99 |
| 1893 | ERR142384 | 1 | | 11 | 3977528 | 98 | 48 | 99 | 99 | 98 |
| 1894 | ERR142385 | 1 | | 42 | 4136695 | 98 | 95 | 99 | 99 | 99 |
| 1898 | ERR142389 | 1 | | 44 | 4324470 | 98 | 16 | 100 | 0 | 0 |
| 1900 | ERR142391 | 1 | | NF | 4370974 | 98 | 94 | 99 | 99 | 95 |
| 1902 | ERR142393 | 1 | | NF | 4509960 | 98 | 57 | 100 | 100 | 97 |
| 1904 | ERR142395 | 1 | | NF | 4179834 | 98 | 95 | 99 | 100 | 100 |
| 1906 | ERR142397 | 1 | | 42 | 4180582 | 98 | 95 | 100 | 99 | 99 |
| 1911 | ERR142402 | 1 | | NF | 4246954 | 98 | 94 | 99 | 99 | 99 |
| 1912 | ERR142403 | 1 | | NF | 4182208 | 98 | 94 | 99 | 99 | 99 |
| 1920 | ERR142411 | 1 | | 35 | 4308553 | 98 | 15 | 99 | 0 | 0 |
| 1926 | ERR142417 | 1 | | 1 | 4358366 | 98 | 14 | 99 | 6 | 0 |
| 1928 | ERR142419 | 1 | | NF | 4499056 | 98 | 95 | 99 | 99 | 98 |
| 1929 | ERR142420 | 1 | | 1 | 4222830 | 98 | 94 | 99 | 99 | 100 |
| 1930 | ERR142421 | 1 | | NF | 4144879 | 98 | 95 | 99 | 99 | 99 |
| 1931 | ERR142422 | 1 | | NF | 4069027 | 98 | 95 | 99 | 99 | 99 |
| 1934 | ERR142425 | 1 | | 1 | 4156445 | 98 | 93 | 99 | 99 | 93 |
| 1935 | ERR142426 | 1 | | 11 | 3989838 | 98 | 48 | 99 | 100 | 100 |
| 1937 | ERR142428 | 1 | | 6 | 4338785 | 98 | 20 | 100 | 11 | 0 |
| 1938 | ERR142429 | 1 | | 1 | 4349621 | 98 | 10 | 99 | 0 | 0 |
| 1941 | ERR142432 | 1 | | 42 | 4122949 | 98 | 95 | 99 | 99 | 99 |
| 1943 | ERR142434 | 1 | | 6 | 4332436 | 98 | 57 | 99 | 100 | 98 |
| 1944 | ERR142435 | 1 | | 11 | 4017705 | 98 | 50 | 100 | 99 | 99 |
| 1945 | ERR142436 | 1 | | 42 | 4182943 | 98 | 95 | 99 | 99 | 100 |
| 1946 | ERR142437 | 1 | | 42 | 4168131 | 98 | 95 | 100 | 99 | 100 |
| 1951 | ERR142442 | 1 | | NF | 4203458 | 98 | 94 | 100 | 99 | 99 |
| 1952 | ERR142443 | 1 | | NF | 4155823 | 98 | 95 | 99 | 99 | 99 |
| 1953 | ERR142444 | 1 | | NF | 4239462 | 98 | 60 | 99 | 100 | 98 |
| 1954 | ERR142445 | 1 | | NF | 4195855 | 98 | 94 | 99 | 99 | 100 |
| 1956 | ERR142447 | 1 | | NF | 4413800 | 98 | 35 | 98 | 48 | 90 |
| 1957 | ERR142448 | 1 | | 17 | 4228171 | 98 | 59 | 99 | 99 | 99 |
| 1958 | ERR142449 | 1 | | 6 | 4441890 | 98 | 14 | 99 | 0 | 0 |
| 1962 | ERR142453 | 1 | | 11 | 4083867 | 98 | 45 | 99 | 99 | 100 |
| 1963 | ERR142454 | 1 | | 6 | 4483701 | 98 | 15 | 98 | 9 | 0 |
| 1964 | ERR142455 | 1 | | 6 | 4396662 | 98 | 13 | 99 | 0 | 0 |
| 1965 | ERR142456 | 1 | 005 | 6 | 4500873 | 98 | 17 | 99 | 11 | 0 |
| 1967 | ERR142458 | 1 | | 6 | 4487084 | 98 | 17 | 95 | 0 | 0 |
| 1968 | ERR142459 | 1 | | 6 | 4248231 | 98 | 15 | 99 | 0 | 0 |
| 1969 | ERR142460 | 1 | | 6 | 4463967 | 98 | 62 | 99 | 99 | 88 |
| 1971 | ERR142462 | 1 | | 17 | 4233116 | 98 | 58 | 99 | 100 | 99 |
| 1973 | ERR142464 | 1 | | 6 | 4310525 | 98 | 65 | 100 | 99 | 91 |
| 1974 | ERR142465 | 1 | | 42 | 4075314 | 98 | 95 | 99 | 99 | 99 |
| 1975 | ERR142466 | 1 | | 6 | 4418267 | 98 | 62 | 99 | 99 | 93 |
| 1977 | ERR142468 | 1 | | 11 | 4133080 | 98 | 13 | 99 | 0 | 0 |
| 1978 | ERR142469 | 1 | | 6 | 4323361 | 98 | 59 | 99 | 99 | 98 |
| 1982 | ERR142473 | 1 | | 11 | 3996978 | 98 | 47 | 100 | 98 | 100 |
| 1983 | ERR142474 | 1 | | 6 | 4272303 | 98 | 62 | 99 | 99 | 99 |
| 1985 | ERR142476 | 1 | | NF | 4375473 | 98 | 61 | 98 | 99 | 100 |
| 1987 | ERR142478 | 1 | | NF | 4471975 | 98 | 29 | 99 | 47 | 60 |
| 1989 | ERR142480 | 1 | | 11 | 4138355 | 98 | 44 | 100 | 99 | 99 |
| 1990 | ERR142481 | 1 | | NF | 4162921 | 98 | 95 | 100 | 99 | 100 |
| 1991 | ERR142482 | 1 | | 11 | 4113283 | 98 | 45 | 100 | 99 | 99 |
| 1994 | ERR142485 | 1 | | 11 | 4026007 | 98 | 46 | 99 | 99 | 99 |
| 1997 | ERR142488 | 1 | | 6 | 4185475 | 98 | 13 | 100 | 0 | 0 |
| 1998 | ERR142489 | 1 | | 11 | 4032437 | 98 | 47 | 100 | 99 | 99 |
| 1999 | ERR142490 | 1 | | NF | 4341614 | 98 | 61 | 99 | 100 | 99 |
| 2000 | ERR142491 | 1 | | 44 | 4482118 | 98 | 62 | 100 | 99 | 94 |
| 2008 | ERR142499 | 1 | | NF | 4376697 | 98 | 62 | 100 | 99 | 98 |
| 2009 | ERR142500 | 1 | | NF | 4382801 | 98 | 63 | 100 | 99 | 92 |
| 2010 | ERR142501 | 1 | | 42 | 4091106 | 98 | 95 | 99 | 99 | 99 |
| 2011 | ERR142502 | 1 | | 11 | 4092115 | 98 | 46 | 100 | 99 | 99 |
| 2013 | ERR142504 | 1 | | 44 | 4399287 | 98 | 58 | 100 | 99 | 92 |
| 2015 | ERR142506 | 1 | | 11 | 4338946 | 98 | 57 | 95 | 99 | 96 |
| 2016 | ERR142507 | 1 | | 1 | 4153874 | 98 | 93 | 98 | 99 | 98 |
| 2017 | ERR142508 | 1 | | 1 | 4197980 | 98 | 93 | 100 | 99 | 98 |
| 2018 | ERR142509 | 1 | | NF | 4167882 | 98 | 94 | 95 | 99 | 98 |
| 2020 | ERR142511 | 1 | | 1 | 4178944 | 98 | 92 | 99 | 100 | 99 |
| 2021 | ERR142512 | 1 | | 3 | 4264258 | 98 | 57 | 99 | 99 | 89 |
| 2022 | ERR142513 | 1 | | NF | 4197183 | 98 | 95 | 99 | 99 | 99 |
| 2023 | ERR142514 | 1 | | 1 | 4127945 | 98 | 92 | 99 | 99 | 99 |
| 2025 | ERR142516 | 1 | | 1 | 4168553 | 98 | 93 | 99 | 99 | 99 |
| 2027 | ERR142518 | 1 | | NF | 4150776 | 98 | 94 | 98 | 99 | 99 |
| 2029 | ERR142520 | 1 | | 1 | 4143671 | 98 | 93 | 100 | 99 | 100 |
| 2032 | ERR142523 | 1 | | 1 | 4188966 | 98 | 93 | 99 | 99 | 99 |
| 2034 | ERR142525 | 1 | | NF | 4165576 | 98 | 16 | 99 | 0 | 0 |
| 2039 | ERR142530 | 1 | | 2 | 4334014 | 98 | 13 | 99 | 0 | 0 |
| 2043 | ERR142534 | 1 | | 271 | 4355932 | 98 | 60 | 100 | 100 | 99 |
| 2044 | ERR142535 | 1 | | 271 | 4254848 | 98 | 59 | 99 | 99 | 91 |
| 2045 | ERR142536 | 1 | | 14 | 4336799 | 98 | 93 | 99 | 99 | 100 |
| 2047 | ERR142538 | 1 | | 36 | 4224105 | 98 | 94 | 99 | 100 | 97 |
| 2050 | ERR142541 | 1 | | 66 | 4316126 | 98 | 94 | 99 | 100 | 97 |
| 2051 | ERR142542 | 1 | | NF | 4267487 | 98 | 94 | 99 | 99 | 99 |
| 2052 | ERR142543 | 1 | | 66 | 4355024 | 98 | 94 | 99 | 99 | 98 |
| 2055 | ERR142546 | 1 | | 11 | 4024772 | 98 | 48 | 99 | 99 | 99 |
| 2057 | ERR142548 | 1 | | 375 | 4171362 | 98 | 94 | 99 | 99 | 99 |
| 2058 | ERR142549 | 1 | | 1 | 4158357 | 98 | 94 | 100 | 99 | 100 |
| 2065 | ERR142556 | 1 | | 1 | 4155947 | 98 | 93 | 99 | 99 | 98 |
| 2067 | ERR142558 | 1 | | 18 | 4266540 | 98 | 8 | 98 | 0 | 0 |
| 2070 | ERR142561 | 1 | | 1 | 4159736 | 98 | 93 | 99 | 100 | 100 |
| 2071 | ERR142562 | 1 | | 1 | 4149000 | 98 | 92 | 95 | 99 | 98 |
| 2072 | ERR142563 | 1 | | 18 | 4266466 | 98 | 34 | 99 | 42 | 80 |
| 2074 | ERR142565 | 1 | | 1 | 4144658 | 98 | 93 | 99 | 99 | 99 |
| 2080 | ERR142571 | 1 | | 1 | 4154913 | 98 | 93 | 100 | 99 | 99 |
| 2086 | ERR142577 | 1 | | 1 | 4130919 | 98 | 93 | 99 | 99 | 99 |
| 2091 | ERR142582 | 1 | | 1 | 4101490 | 98 | 92 | 100 | 99 | 96 |
| 2093 | ERR142584 | 1 | | 49 | 4490874 | 98 | 92 | 99 | 99 | 99 |
| 2098 | ERR142589 | 1 | | 55 | 4035290 | 98 | 94 | 99 | 99 | 100 |
| 2102 | ERR142593 | 1 | | NF | 4521284 | 98 | 94 | 99 | 99 | 99 |
| 2106 | ERR142597 | 1 | | NF | 4354326 | 98 | 92 | 100 | 99 | 99 |
| 2107 | ERR142598 | 1 | | 55 | 4045487 | 98 | 94 | 99 | 99 | 93 |
| 2109 | ERR142600 | 1 | | 37 | 4149900 | 98 | 93 | 99 | 99 | 99 |
| 2110 | ERR142601 | 1 | | 49 | 4276088 | 98 | 92 | 100 | 99 | 98 |
| 2111 | ERR142602 | 1 | | 55 | 4150271 | 98 | 95 | 99 | 99 | 97 |
| 2112 | ERR142603 | 1 | | 45 | 4413577 | 98 | 94 | 100 | 99 | 98 |
| 2113 | ERR142604 | 1 | | NF | 4239570 | 98 | 10 | 99 | 0 | 0 |
| 2115 | ERR142606 | 1 | | 37 | 4056729 | 98 | 93 | 99 | 100 | 99 |
| 2117 | ERR142608 | 1 | | 45 | 4420237 | 98 | 10 | 100 | 0 | 0 |
| 2118 | ERR142609 | 1 | | 55 | 4125053 | 98 | 94 | 99 | 99 | 99 |
| 2122 | ERR232377 | 1 | | 7 | 4085190 | 98 | 27 | 100 | 13 | 45 |
| 2124 | ERR232379 | 1 | 067 | 27 | 4366050 | 98 | 95 | 100 | 100 | 100 |
| 2125 | ERR232380 | 1 | | 29 | 4221923 | 98 | 94 | 100 | 99 | 99 |
| 2129 | ERR232384 | 1 | | 4 | 4193557 | 98 | 93 | 100 | 99 | 100 |
| 2130 | ERR232385 | 1 | 012 | 54 | 4398961 | 98 | 100 | 100 | 99 | 100 |
| 2131 | ERR232386 | 1 | | 6 | 4184427 | 98 | 59 | 100 | 99 | 99 |
| 2132 | ERR232387 | 1 | 036 | 62 | 4089105 | 98 | 4 | 99 | 0 | 0 |
| 2133 | ERR232388 | 1 | | 13 | 4142484 | 98 | 98 | 100 | 100 | 100 |
| 2134 | ERR232389 | 1 | | 17 | 4309632 | 98 | 95 | 100 | 99 | 100 |
| 2136 | ERR232391 | 1 | | 122 | 4106478 | 98 | 95 | 100 | 99 | 100 |
| 2140 | ERR232395 | 1 | | 167 | 4138758 | 98 | 50 | 100 | 100 | 99 |
| 2141 | ERR232396 | 1 | 239 | 168 | 4090253 | 98 | 55 | 100 | 99 | 99 |
| 2142 | ERR232397 | 1 | | 17 | 4117314 | 98 | 22 | 100 | 0 | 0 |
| 2145 | ERR232400 | 1 | | 179 | 4477056 | 98 | 39 | 100 | 79 | 0 |
| 2148 | ERR251713 | 1 | | 8 | 4299569 | 98 | 30 | 100 | 33 | 0 |
| 2152 | ERR251717 | 1 | | 2 | 4325035 | 98 | 23 | 99 | 38 | 0 |
| 2154 | ERR251719 | 1 | | 1 | 4147647 | 98 | 94 | 99 | 100 | 99 |
| 2155 | ERR251720 | 1 | | 55 | 4045244 | 98 | 94 | 100 | 99 | 99 |
| 2157 | ERR251722 | 1 | | 26 | 4368989 | 98 | 91 | 100 | 99 | 100 |
| 2159 | ERR251724 | 1 | | 2 | 4284881 | 98 | 92 | 99 | 99 | 99 |
| 2164 | ERR251729 | 1 | | 1 | 4171114 | 98 | 94 | 99 | 99 | 99 |
| 2168 | ERR251733 | 1 | | NF | 4250453 | 98 | 92 | 100 | 99 | 99 |
| 2171 | ERR251736 | 1 | | 2 | 4149415 | 98 | 92 | 100 | 100 | 100 |
| 2172 | ERR251737 | 1 | | 5 | 4238942 | 98 | 92 | 99 | 99 | 99 |
| 2176 | ERR251741 | 1 | | 58 | 4295840 | 98 | 62 | 99 | 99 | 99 |
| 2177 | ERR251742 | 1 | | 1 | 4150918 | 98 | 93 | 99 | 100 | 99 |
| 2178 | ERR251743 | 1 | | 42 | 4179553 | 98 | 95 | 100 | 100 | 100 |
| 2181 | ERR251746 | 1 | | 51 | 4309851 | 98 | 31 | 100 | 6 | 0 |
| 2182 | ERR251747 | 1 | | 1 | 4208222 | 98 | 93 | 99 | 99 | 100 |
| 2183 | ERR251748 | 1 | | NF | 4217321 | 98 | 93 | 99 | 99 | 99 |
| 2184 | ERR251749 | 1 | | NF | 4187179 | 98 | 92 | 99 | 100 | 100 |
| 2188 | ERR251753 | 1 | | 5 | 4232435 | 98 | 39 | 99 | 99 | 94 |
| 2195 | ERR251760 | 1 | | 9 | 4409162 | 98 | 12 | 99 | 0 | 0 |
| 2198 | ERR251763 | 1 | | 14 | 4179523 | 98 | 92 | 100 | 99 | 99 |
| 2199 | ERR251764 | 1 | | 44 | 4532000 | 98 | 94 | 99 | 100 | 99 |
| 2201 | ERR251766 | 1 | | 2 | 4148520 | 98 | 18 | 100 | 0 | 0 |
| 2207 | ERR251772 | 1 | | 7 | 4085890 | 98 | 20 | 100 | 0 | 0 |
| 2208 | ERR251773 | 1 | | 8 | 4081289 | 98 | 95 | 99 | 99 | 99 |
| 2211 | ERR251776 | 1 | | 1 | 4155775 | 98 | 93 | 100 | 99 | 100 |
| 2216 | ERR251781 | 1 | | 14 | 4212855 | 98 | 92 | 99 | 99 | 99 |
| 2220 | ERR251785 | 1 | | 1 | 4165427 | 98 | 93 | 100 | 99 | 100 |
| 2228 | ERR251793 | 1 | | NF | 4903393 | 98 | 95 | 99 | 99 | 100 |
| 2231 | ERR251796 | 1 | | 1 | 5257406 | 98 | 94 | 100 | 100 | 99 |
| 2234 | ERR251799 | 1 | | NF | 4901677 | 98 | 92 | 99 | 99 | 99 |
| 2236 | ERR251801 | 1 | | NF | 4620124 | 98 | 95 | 99 | 99 | 100 |
| 2237 | ERR251802 | 1 | | 44 | 4783119 | 98 | 95 | 100 | 100 | 100 |
| 2238 | ERR251803 | 1 | | 66 | 4595995 | 98 | 95 | 100 | 100 | 99 |
| 2240 | ERR251805 | 1 | | 1 | 4143105 | 98 | 93 | 100 | 99 | 99 |
| 2247 | ERR251812 | 1 | | 67 | 5026877 | 98 | 95 | 99 | 99 | 100 |
| 2248 | ERR251813 | 1 | | 44 | 4921733 | 98 | 93 | 100 | 99 | 99 |
| 2251 | ERR251816 | 1 | | 2 | 4917087 | 98 | 93 | 100 | 99 | 99 |
| 2254 | ERR251819 | 1 | | 13 | 4539262 | 98 | 95 | 99 | 99 | 99 |
| 2255 | ERR251820 | 1 | | 44 | 4616797 | 98 | 95 | 99 | 99 | 99 |
| 2256 | ERR251821 | 1 | | 49 | 4471254 | 98 | 95 | 100 | 100 | 99 |
| 2263 | ERR251828 | 1 | | 46 | 4359345 | 98 | 94 | 100 | 99 | 99 |
| 2265 | ERR251830 | 1 | | 14 | 4549473 | 98 | 95 | 99 | 99 | 100 |
| 2267 | ERR251832 | 1 | | 1 | 4871446 | 98 | 94 | 100 | 99 | 100 |
| 2274 | ERR251839 | 1 | | 8 | 4418048 | 98 | 93 | 100 | 99 | 100 |
| 2278 | ERR251843 | 1 | | 8 | 4332680 | 98 | 95 | 99 | 100 | 100 |
| 2281 | ERR251846 | 1 | | 46 | 4347184 | 98 | 94 | 99 | 99 | 100 |
| 2283 | ERR251857 | 1 | | 1 | 4139728 | 98 | 93 | 100 | 100 | 100 |
| 2284 | ERR251858 | 1 | | 35 | 4283784 | 98 | 15 | 100 | 0 | 0 |
| 2285 | ERR251859 | 1 | | 6 | 4419666 | 98 | 17 | 100 | 0 | 0 |
| 2287 | ERR251864 | 1 | | 1 | 4135834 | 98 | 94 | 99 | 99 | 100 |

| | | | | | | | | | | |
|---|---|---|---|---|---|---|---|---|---|---|
| 2300 | ERR272059 | 1 | | 37 | 4160253 | 98 | 93 | 100 | 99 | 99 |
| 2302 | ERR272061 | 1 | | 39 | 4252581 | 98 | 94 | 100 | 100 | 100 |
| 2304 | ERR272063 | 1 | | 39 | 4111326 | 98 | 94 | 99 | 99 | 100 |
| 2309 | ERR272068 | 1 | | 58 | 4383703 | 98 | 12 | 99 | 0 | 0 |
| 2313 | ERR272072 | 1 | | 58 | 4202998 | 98 | 16 | 100 | 13 | 0 |
| 2314 | ERR272073 | 1 | | 54 | 4572691 | 98 | 99 | 100 | 99 | 99 |
| 2317 | ERR272076 | 1 | | 54 | 4191943 | 98 | 99 | 100 | 99 | 99 |
| 2318 | ERR272077 | 1 | | 54 | 4176895 | 98 | 99 | 99 | 99 | 100 |
| 2320 | ERR272079 | 1 | | 54 | 4240076 | 98 | 99 | 99 | 100 | 99 |
| 2321 | ERR272080 | 1 | | 54 | 4239020 | 98 | 99 | 100 | 100 | 99 |
| 2325 | ERR272084 | 1 | | 3 | 4118733 | 98 | 12 | 99 | 0 | 0 |
| 2327 | ERR272086 | 1 | | 3 | 4404571 | 98 | 59 | 99 | 99 | 99 |
| 2333 | ERR272092 | 1 | | 271 | 4250941 | 98 | 60 | 99 | 100 | 99 |
| 2337 | ERR272096 | 1 | | 3 | 4190757 | 98 | 60 | 99 | 100 | 96 |
| 2338 | ERR272097 | 1 | | 3 | 4221935 | 98 | 62 | 99 | 100 | 98 |
| 2341 | ERR272100 | 1 | | 271 | 4233553 | 98 | 61 | 99 | 100 | 100 |
| 2343 | ERR272102 | 1 | | 3 | 4233882 | 98 | 62 | 99 | 99 | 96 |
| 2345 | ERR272104 | 1 | | 3 | 4167725 | 98 | 61 | 99 | 99 | 99 |
| 2346 | ERR272105 | 1 | | 3 | 4232342 | 98 | 62 | 99 | 99 | 99 |
| 2349 | ERR272108 | 1 | | 3 | 4430519 | 98 | 60 | 100 | 99 | 93 |
| 2350 | ERR272109 | 1 | | 271 | 4330165 | 98 | 61 | 99 | 100 | 96 |
| 2351 | ERR272110 | 1 | | 271 | 4211946 | 98 | 62 | 100 | 100 | 98 |
| 2355 | ERR272114 | 1 | | 3 | 4203870 | 98 | 58 | 99 | 100 | 99 |
| 2356 | ERR272115 | 1 | | 3 | 4252977 | 98 | 59 | 100 | 100 | 98 |
| 2362 | ERR272121 | 1 | | 3 | 4227565 | 98 | 58 | 100 | 99 | 96 |
| 2367 | ERR272126 | 1 | | 3 | 4375716 | 98 | 59 | 100 | 100 | 94 |
| 2368 | ERR272127 | 1 | | 3 | 4405916 | 98 | 59 | 99 | 99 | 99 |
| 2370 | ERR272129 | 1 | | 3 | 4364709 | 98 | 59 | 100 | 100 | 98 |
| 2371 | ERR272130 | 1 | | 3 | 4315081 | 98 | 59 | 100 | 100 | 99 |
| 2372 | ERR272131 | 1 | | 3 | 4459834 | 98 | 62 | 100 | 100 | 99 |
| 2377 | ERR272136 | 1 | | 3 | 4255889 | 98 | 61 | 100 | 99 | 99 |
| 2378 | ERR272137 | 1 | | 3 | 4362483 | 98 | 60 | 99 | 100 | 94 |
| 2380 | ERR272139 | 1 | | 3 | 4359424 | 98 | 64 | 99 | 99 | 95 |
| 2383 | ERR272142 | 1 | | 3 | 4192393 | 98 | 59 | 100 | 100 | 99 |
| 2384 | ERR272143 | 1 | | 3 | 4215131 | 98 | 18 | 100 | 0 | 0 |
| 2386 | ERR272145 | 1 | | 3 | 4178478 | 98 | 60 | 99 | 99 | 96 |
| 2387 | ERR272146 | 1 | | 3 | 4147837 | 98 | 25 | 99 | 13 | 0 |
| 2389 | ERR272148 | 1 | | 3 | 4213980 | 98 | 60 | 99 | 100 | 99 |
| 2391 | ERR272150 | 1 | | 3 | 4448455 | 98 | 24 | 100 | 19 | 0 |
| 2393 | ERR272152 | 1 | | 6 | 4329682 | 98 | 16 | 99 | 0 | 0 |
| 2396 | ERR272155 | 1 | | 6 | 4279600 | 98 | 18 | 100 | 0 | 0 |
| 2399 | ERR272158 | 1 | | 6 | 4432344 | 98 | 65 | 100 | 99 | 97 |
| 2400 | ERR272159 | 1 | | NF | 4370033 | 98 | 59 | 100 | 99 | 99 |
| 2403 | ERR272162 | 1 | | NF | 4332484 | 98 | 16 | 99 | 6 | 0 |
| 2405 | ERR272164 | 1 | | 6 | 4299532 | 98 | 17 | 100 | 0 | 0 |
| 2406 | ERR272165 | 1 | | 6 | 4336370 | 98 | 13 | 99 | 0 | 0 |
| 2408 | ERR272167 | 1 | | 6 | 4535031 | 98 | 22 | 100 | 14 | 0 |
| 2409 | ERR272168 | 1 | | 6 | 4512186 | 98 | 15 | 99 | 0 | 0 |
| 2412 | ERR272171 | 1 | | 6 | 4330560 | 98 | 38 | 99 | 55 | 45 |
| 2417 | ERR272176 | 1 | | 6 | 4331159 | 98 | 21 | 99 | 15 | 0 |
| 2420 | ERR272179 | 1 | | 6 | 4414095 | 98 | 16 | 100 | 0 | 0 |
| 2424 | ERR272183 | 1 | | 6 | 4698826 | 98 | 19 | 99 | 0 | 0 |
| 2425 | ERR272184 | 1 | | 6 | 4330813 | 98 | 19 | 99 | 32 | 0 |
| 2426 | ERR272185 | 1 | | 6 | 4217779 | 98 | 29 | 99 | 37 | 0 |
| 2432 | ERR272191 | 1 | | 6 | 4381279 | 98 | 22 | 99 | 0 | 45 |
| 2436 | ERR272195 | 1 | | NF | 4267383 | 98 | 14 | 100 | 0 | 0 |
| 2437 | ERR272196 | 1 | | 7 | 4132181 | 98 | 25 | 99 | 31 | 0 |
| 2438 | ERR272197 | 1 | | NF | 4594032 | 98 | 17 | 100 | 8 | 0 |
| 2440 | ERR272199 | 1 | | 7 | 4401328 | 98 | 14 | 100 | 0 | 0 |
| 2441 | ERR272200 | 1 | | 7 | 4302016 | 98 | 11 | 99 | 0 | 0 |
| 2442 | ERR272201 | 1 | | 7 | 4429130 | 98 | 20 | 100 | 26 | 0 |
| 2444 | ERR272203 | 1 | | 7 | 4190457 | 98 | 20 | 100 | 0 | 0 |
| 2446 | ERR272205 | 1 | | NF | 4412594 | 98 | 18 | 99 | 13 | 0 |
| 2447 | ERR272206 | 1 | | NF | 4206184 | 98 | 17 | 100 | 0 | 0 |
| 2448 | ERR272207 | 1 | | 7 | 4269784 | 98 | 25 | 99 | 22 | 0 |
| 2452 | ERR272211 | 1 | | 7 | 4130400 | 98 | 37 | 99 | 41 | 0 |
| 2455 | ERR272214 | 1 | | 7 | 4249104 | 98 | 20 | 99 | 26 | 0 |
| 2457 | ERR272216 | 1 | | 7 | 4307410 | 98 | 16 | 100 | 19 | 0 |
| 2458 | ERR272217 | 1 | | 7 | 4717758 | 98 | 93 | 100 | 98 | 98 |
| 2459 | ERR272218 | 1 | | 7 | 4307421 | 98 | 21 | 100 | 0 | 0 |
| 2461 | ERR272220 | 1 | | 7 | 4357298 | 98 | 33 | 100 | 26 | 0 |
| 2465 | ERR272224 | 1 | | 7 | 4259166 | 98 | 19 | 100 | 23 | 0 |
| 2467 | ERR272226 | 1 | | 7 | 4394586 | 98 | 17 | 99 | 6 | 0 |
| 2468 | ERR272227 | 1 | | 7 | 4486465 | 98 | 16 | 100 | 0 | 0 |
| 2469 | ERR272228 | 1 | | 7 | 4234033 | 98 | 27 | 100 | 32 | 0 |
| 2472 | ERR272231 | 1 | | 7 | 4285900 | 98 | 34 | 100 | 31 | 0 |
| 2474 | ERR272233 | 1 | | 7 | 4357846 | 98 | 19 | 100 | 0 | 0 |
| 2476 | ERR272235 | 1 | | 7 | 4655961 | 98 | 25 | 100 | 23 | 0 |
| 2478 | ERR272237 | 1 | | 7 | 4189820 | 98 | 17 | 99 | 13 | 0 |
| 2480 | ERR272239 | 1 | | NF | 4404443 | 98 | 60 | 100 | 99 | 99 |
| 2481 | ERR272240 | 1 | | NF | 4518084 | 98 | 57 | 99 | 99 | 98 |
| 2482 | ERR272241 | 1 | | 44 | 4411853 | 98 | 59 | 99 | 99 | 81 |
| 2485 | ERR272244 | 1 | | 44 | 4493150 | 98 | 94 | 100 | 99 | 99 |
| 2489 | ERR272248 | 1 | | 44 | 4363973 | 98 | 64 | 100 | 100 | 99 |
| 2490 | ERR272249 | 1 | | 44 | 4328012 | 98 | 63 | 100 | 100 | 94 |
| 2491 | ERR272250 | 1 | | 44 | 4438710 | 98 | 27 | 99 | 19 | 0 |
| 2495 | ERR272254 | 1 | | 44 | 4438121 | 98 | 95 | 99 | 99 | 99 |
| 2496 | ERR272255 | 1 | | NF | 4488458 | 98 | 63 | 99 | 99 | 89 |
| 2498 | ERR272257 | 1 | | 44 | 4340537 | 98 | 95 | 100 | 99 | 100 |
| 2499 | ERR272258 | 1 | | 44 | 4650581 | 98 | 94 | 100 | 99 | 99 |
| 2500 | ERR272259 | 1 | | 44 | 4427178 | 98 | 95 | 100 | 100 | 100 |
| 2504 | ERR467530 | 1 | | 54 | 4568939 | 98 | 100 | 100 | 100 | 100 |
| 2507 | ERR467533 | 1 | | 2 | 4198240 | 98 | 20 | 99 | 7 | 0 |
| 2508 | ERR467534 | 1 | | 54 | 4659829 | 98 | 99 | 99 | 99 | 99 |
| 2509 | ERR467535 | 1 | | | 4356353 | 98 | 92 | 99 | 99 | 100 |
| 2511 | ERR467537 | 1 | | 58 | 4156796 | 98 | 16 | 100 | 0 | 0 |
| 2512 | ERR467538 | 1 | | 54 | 4599372 | 98 | 99 | 100 | 99 | 100 |
| 2513 | ERR467539 | 1 | | 54 | 4599129 | 98 | 99 | 99 | 99 | 100 |
| 2514 | ERR467540 | 1 | | 58 | 4155931 | 98 | 25 | 99 | 23 | 45 |
| 2515 | ERR467541 | 1 | | 54 | 4572694 | 98 | 99 | 100 | 99 | 99 |
| 2519 | ERR467545 | 1 | | 54 | 4606805 | 98 | 99 | 99 | 100 | 100 |
| 2520 | ERR467546 | 1 | | 54 | 4571408 | 98 | 99 | 100 | 100 | 99 |
| 2521 | ERR467547 | 1 | | 54 | 4606545 | 98 | 99 | 99 | 99 | 99 |
| 2528 | ERR467554 | 1 | | 1 | 4194645 | 98 | 93 | 100 | 99 | 100 |
| 2529 | ERR467555 | 1 | | 54 | 4574780 | 98 | 99 | 99 | 99 | 99 |
| 2530 | ERR467556 | 1 | | 1 | 4199943 | 98 | 93 | 99 | 100 | 99 |
| 2537 | ERR467563 | 1 | | 3 | 4247180 | 98 | 14 | 99 | 0 | 0 |
| 2539 | ERR467565 | 1 | | 54 | 4606104 | 98 | 100 | 100 | 99 | 100 |
| 2540 | ERR467566 | 1 | | 1 | 4247706 | 98 | 93 | 100 | 100 | 99 |
| 2542 | ERR467568 | 1 | | 54 | 4597691 | 98 | 99 | 100 | 100 | 99 |
| 2543 | ERR467569 | 1 | | 54 | 4572797 | 98 | 100 | 100 | 99 | 99 |
| 2547 | ERR467573 | 1 | | 8 | 4195149 | 98 | 95 | 100 | 100 | 99 |
| 2550 | ERR467576 | 1 | | 1 | 4193632 | 98 | 93 | 99 | 100 | 99 |
| 2551 | ERR467577 | 1 | | 1 | 4199153 | 98 | 93 | 100 | 100 | 99 |
| 2552 | ERR467578 | 1 | | 54 | 4602601 | 98 | 99 | 99 | 99 | 99 |
| 2553 | ERR467579 | 1 | | 2 | 4199731 | 98 | 26 | 100 | 13 | 0 |
| 2554 | ERR467580 | 1 | | 8 | 4159535 | 98 | 95 | 99 | 100 | 99 |
| 2555 | ERR467581 | 1 | | 54 | 4605295 | 98 | 100 | 99 | 100 | 99 |
| 2556 | ERR467582 | 1 | | 67 | 4113907 | 98 | 95 | 100 | 100 | 99 |
| 2557 | ERR467583 | 1 | | 1 | 4194323 | 98 | 93 | 99 | 100 | 99 |
| 2558 | ERR467584 | 1 | | 1 | 4196176 | 98 | 93 | 99 | 99 | 99 |
| 2559 | ERR467585 | 1 | | 8 | 4093298 | 98 | 95 | 99 | 100 | 99 |
| 2560 | ERR467586 | 1 | | 54 | 4549929 | 98 | 98 | 100 | 99 | 100 |
| 2561 | ERR467587 | 1 | | 54 | 4546536 | 98 | 98 | 99 | 100 | 98 |
| 2571 | ERR467597 | 1 | | 2 | 4248803 | 98 | 12 | 99 | 0 | 0 |
| 2573 | ERR467599 | 1 | | 37 | 4259460 | 98 | 93 | 100 | 99 | 99 |
| 2574 | ERR467600 | 1 | | - | 4206474 | 98 | 15 | 99 | 13 | 0 |
| 2576 | ERR467602 | 1 | | 54 | 4586248 | 98 | 16 | 100 | 0 | 0 |
| 2579 | ERR467605 | 1 | | 37 | 4266063 | 98 | 95 | 99 | 100 | 100 |
| 2580 | ERR467606 | 1 | | 8 | 4078790 | 98 | 95 | 100 | 100 | 100 |
| 2583 | ERR467609 | 1 | | 37 | 4267726 | 98 | 95 | 99 | 99 | 100 |
| 2586 | ERR467612 | 1 | | 3 | 4150067 | 98 | 61 | 99 | 99 | 93 |
| 2589 | ERR467615 | 1 | | 53 | 4078517 | 98 | 92 | 100 | 99 | 99 |
| 2591 | ERR467617 | 1 | | 37 | 4291347 | 98 | 93 | 99 | 99 | 99 |
| 2593 | ERR467619 | 1 | | 37 | 4256719 | 98 | 94 | 100 | 99 | 100 |
| 2594 | ERR467620 | 1 | | 11 | 3931184 | 98 | 45 | 100 | 100 | 99 |
| 2595 | ERR467621 | 1 | | 37 | 4266872 | 98 | 94 | 99 | 99 | 100 |
| 2597 | ERR467623 | 1 | | 154 | 4328723 | 98 | 94 | 99 | 99 | 99 |
| 2607 | ERR008621 | 1 | | 1 | 4235369 | 98 | 91 | 99 | 99 | 97 |
| 2615 | ERR008629 | 1 | | 1 | 4214078 | 98 | 91 | 99 | 99 | 99 |
| 2618 | ERR008632 | 1 | | 1 | 4193378 | 98 | 93 | 99 | 99 | 98 |
| 2620 | ERR008634 | 1 | | 1 | 3975706 | 98 | 90 | 98 | 99 | 99 |
| 2627 | ERR022125 | 1 | 027 | 1 | 3635136 | 98 | 79 | 99 | 100 | 55 |
| 2630 | ERR045749 | 1 | | 1 | 4360695 | 98 | 92 | 99 | 100 | 98 |
| 2633 | ERR045752 | 1 | 027 | 1 | 4025598 | 98 | 89 | 97 | 99 | 93 |
| 2641 | ERR111455 | 1 | 027 | 1 | 4191520 | 98 | 83 | 98 | 98 | 98 |
| 2671 | T15 | 1 | 005 | 6 | 4413466 | 98 | 25 | 100 | 19 | 0 |
| 2683 | ERR604986 | 1 | 050 | 16 | 4507621 | 98 | 95 | 99 | 100 | 99 |
| 2685 | ERR604991 | 1 | 056 | 58 | 4377961 | 98 | 18 | 99 | 20 | 0 |
| 2686 | ERR604995 | 1 | 017 | 37 | 4372867 | 98 | 91 | 99 | 99 | 99 |
| 2688 | ERR605000 | 1 | 002 | 8 | 4225258 | 98 | 95 | 99 | 100 | 96 |
| 2689 | ERR605004 | 1 | 078 | 11 | 4026499 | 98 | 45 | 99 | 100 | 99 |
| 2690 | ERR605006 | 1 | 050 | 16 | 4539762 | 98 | 95 | 99 | 99 | 99 |
| 2691 | ERR605007 | 1 | 070 | 55 | 4156486 | 98 | 94 | 100 | 99 | 99 |
| 2692 | ERR605009 | 1 | 020 | 2 | 4064306 | 98 | 93 | 99 | 100 | 99 |
| 2695 | ERR605022 | 1 | 023 | 5 | 4429249 | 98 | 41 | 100 | 100 | 99 |
| 2696 | ERR605023 | 1 | 015 | 44 | 4954442 | 98 | 60 | 100 | 99 | 90 |
| 2697 | ERR605025 | 1 | 078 | 11 | 4086486 | 98 | 49 | 100 | 100 | 99 |
| 2 | ERR024643 | 1 | 027 | 1 | 4135521 | 99 | 94 | 100 | 99 | 100 |
| 4 | ERR024646 | 1 | 027 | 1 | 4147119 | 99 | 94 | 100 | 99 | 100 |
| 6 | ERR024648 | 1 | 027 | 1 | 4140044 | 99 | 94 | 99 | 100 | 99 |
| 10 | ERR024652 | 1 | 027 | 1 | 4146164 | 99 | 94 | 100 | 99 | 100 |
| 11 | ERR024653 | 1 | 027 | 1 | 4134758 | 99 | 93 | 100 | 99 | 100 |
| 15 | ERR024659 | 1 | 027 | 1 | 4136984 | 99 | 94 | 100 | 100 | 100 |
| 16 | ERR024660 | 1 | 027 | 1 | 4141194 | 99 | 93 | 99 | 99 | 100 |
| 33 | ERR025353 | 1 | 027 | 1 | 4134130 | 99 | 93 | 100 | 100 | 99 |
| 34 | ERR025354 | 1 | 027 | 1 | 4135105 | 99 | 93 | 100 | 100 | 99 |
| 47 | ERR026363 | 1 | 027 | 1 | 4128661 | 99 | 93 | 100 | 100 | 99 |
| 50 | ERR026366 | 1 | 027 | 1 | 4132531 | 99 | 93 | 99 | 100 | 100 |
| 55 | ERR026371 | 1 | 027 | 1 | 4137376 | 99 | 93 | 100 | 99 | 99 |
| 60 | ERR026376 | 1 | 027 | 1 | 4135905 | 99 | 93 | 100 | 99 | 99 |
| 65 | ERR026381 | 1 | 027 | 1 | 4139789 | 99 | 93 | 100 | 99 | 100 |
| 66 | ERR026382 | 1 | 027 | 1 | 4136648 | 99 | 93 | 100 | 99 | 100 |
| 67 | ERR026383 | 1 | 027 | 1 | 4133818 | 99 | 93 | 98 | 99 | 99 |
| 73 | ERR026389 | 1 | 027 | 1 | 4135635 | 99 | 93 | 100 | 100 | 100 |
| 78 | ERR026394 | 1 | 027 | 1 | 4143884 | 99 | 93 | 100 | 99 | 100 |
| 79 | ERR026395 | 1 | 027 | 1 | 4130288 | 99 | 93 | 100 | 99 | 99 |
| 84 | ERR026400 | 1 | 027 | 1 | 4139289 | 99 | 93 | 100 | 100 | 100 |
| 93 | ERR026409 | 1 | 027 | 1 | 4133613 | 99 | 93 | 100 | 99 | 100 |
| 101 | ERR026417 | 1 | 027 | 1 | 4138685 | 99 | 93 | 99 | 99 | 99 |
| 102 | ERR026418 | 1 | 027 | 1 | 4186613 | 99 | 93 | 99 | 100 | 100 |
| 108 | ERR026424 | 1 | 027 | 1 | 4151435 | 99 | 93 | 100 | 99 | 99 |
| 114 | ERR026833 | 1 | 027 | 1 | 4133956 | 99 | 93 | 99 | 99 | 99 |
| 126 | ERR026845 | 1 | 027 | 1 | 4132812 | 99 | 93 | 100 | 99 | 99 |
| 128 | ERR026847 | 1 | 027 | 1 | 4137719 | 99 | 93 | 100 | 100 | 100 |
| 131 | ERR026851 | 1 | 027 | 18 | 4279518 | 99 | 30 | 100 | 53 | 0 |
| 141 | ERR026862 | 1 | 027 | 1 | 4145979 | 99 | 93 | 99 | 99 | 99 |

| | | | | | | | | | | |
|---|---|---|---|---|---|---|---|---|---|---|
| 148 | ERR026869 | 1 | 027 | 1 | 4130996 | 99 | 93 | 100 | 100 | 100 |
| 165 | ERR028725 | 1 | 027 | 1 | 4130499 | 99 | 93 | 99 | 99 | 100 |
| 168 | ERR028728 | 1 | 027 | 1 | 4134436 | 99 | 93 | 100 | 100 | 100 |
| 181 | ERR029488 | 1 | 027 | 1 | 4138307 | 99 | 93 | 100 | 99 | 99 |
| 184 | ERR029491 | 1 | 027 | 1 | 4130714 | 99 | 93 | 99 | 100 | 100 |
| 185 | ERR029492 | 1 | 027 | 1 | 4130974 | 99 | 93 | 99 | 100 | 99 |
| 187 | ERR029494 | 1 | 027 | 1 | 4147174 | 99 | 93 | 99 | 99 | 99 |
| 207 | ERR029514 | 1 | 027 | 1 | 4178689 | 99 | 93 | 100 | 99 | 99 |
| 211 | ERR029518 | 1 | 027 | 1 | 4157760 | 99 | 94 | 100 | 99 | 100 |
| 217 | ERR029524 | 1 | 027 | 1 | 4095445 | 99 | 93 | 99 | 100 | 99 |
| 219 | ERR029526 | 1 | 027 | 1 | 4096961 | 99 | 93 | 100 | 99 | 99 |
| 230 | ERR029538 | 1 | 027 | 1 | 4145401 | 99 | 93 | 99 | 99 | 99 |
| 241 | ERR030287 | 1 | 027 | 1 | 4137655 | 99 | 93 | 99 | 99 | 100 |
| 252 | ERR030298 | 1 | 027 | 1 | 4134249 | 99 | 93 | 100 | 99 | 100 |
| 263 | ERR030309 | 1 | 027 | 1 | 4144396 | 99 | 93 | 99 | 100 | 100 |
| 266 | ERR030312 | 1 | 027 | 1 | 4142948 | 99 | 92 | 100 | 100 | 99 |
| 269 | ERR030315 | 1 | 027 | 1 | 4133248 | 99 | 93 | 99 | 99 | 100 |
| 270 | ERR030316 | 1 | 027 | 1 | 4135013 | 99 | 93 | 99 | 100 | 99 |
| 272 | ERR030318 | 1 | 027 | 1 | 4132417 | 99 | 93 | 100 | 99 | 99 |
| 273 | ERR030319 | 1 | 027 | 1 | 4082560 | 99 | 93 | 100 | 100 | 100 |
| 281 | ERR030327 | 1 | 027 | 1 | 4135364 | 99 | 93 | 99 | 100 | 100 |
| 290 | ERR030336 | 1 | 027 | 1 | 4134166 | 99 | 93 | 100 | 100 | 99 |
| 293 | ERR030339 | 1 | 027 | 1 | 4256097 | 99 | 93 | 99 | 99 | 99 |
| 294 | ERR030340 | 1 | 027 | 1 | 4086551 | 99 | 93 | 100 | 99 | 100 |
| 301 | ERR030347 | 1 | | NF | 4097041 | 99 | 92 | 95 | 99 | 97 |
| 303 | ERR030349 | 1 | 027 | 1 | 4134846 | 99 | 93 | 100 | 100 | 98 |
| 304 | ERR030350 | 1 | 027 | 1 | 4134163 | 99 | 93 | 100 | 100 | 99 |
| 309 | ERR030355 | 1 | 027 | 1 | 4134976 | 99 | 93 | 100 | 99 | 100 |
| 321 | ERR030367 | 1 | 27 | 1 | 4133929 | 99 | 93 | 99 | 99 | 100 |
| 329 | ERR030375 | 1 | 027 | 1 | 4134584 | 99 | 93 | 100 | 100 | 100 |
| 333 | ERR030379 | 1 | 027 | 1 | 4146056 | 99 | 93 | 100 | 99 | 99 |
| 336 | ERR030384 | 1 | 027 | 1 | 4647539 | 99 | 95 | 100 | 99 | 100 |
| 343 | ERR030391 | 1 | 027 | 1 | 4132251 | 99 | 93 | 100 | 99 | 100 |
| 348 | ERR030396 | 1 | 027 | 1 | 4136671 | 99 | 93 | 99 | 99 | 99 |
| 368 | ERR031690 | 1 | 027 | 1 | 4203540 | 99 | 93 | 100 | 99 | 100 |
| 370 | ERR031692 | 1 | 027 | 1 | 4135689 | 99 | 93 | 100 | 99 | 100 |
| 372 | ERR031694 | 1 | 027 | 1 | 4135501 | 99 | 93 | 100 | 99 | 99 |
| 377 | ERR031699 | 1 | | 1 | 4100842 | 99 | 92 | 94 | 99 | 98 |
| 379 | ERR031701 | 1 | 027 | 1 | 4132826 | 99 | 93 | 100 | 100 | 99 |
| 394 | ERR034224 | 1 | 027 | 1 | 4130057 | 99 | 94 | 100 | 100 | 100 |
| 396 | ERR034226 | 1 | 027 | 1 | 4143070 | 99 | 94 | 100 | 100 | 100 |
| 401 | ERR044803 | 1 | 027 | 1 | 4135081 | 99 | 93 | 100 | 100 | 100 |
| 403 | ERR044805 | 1 | 027 | 1 | 4130984 | 99 | 93 | 100 | 100 | 100 |
| 405 | ERR044807 | 1 | 027 | 1 | 4136027 | 99 | 94 | 100 | 100 | 100 |
| 408 | ERR044810 | 1 | 027 | 67 | 4097395 | 99 | 94 | 100 | 99 | 99 |
| 416 | ERR044818 | 1 | 027 | 1 | 4244774 | 99 | 94 | 100 | 100 | 100 |
| 427 | ERR044829 | 1 | 027 | 1 | 4153404 | 99 | 93 | 100 | 100 | 99 |
| 429 | ERR044831 | 1 | 027 | 1 | 4210416 | 99 | 94 | 100 | 100 | 99 |
| 441 | ERR044844 | 1 | 027 | 1 | 4126272 | 99 | 93 | 100 | 99 | 100 |
| 442 | ERR044845 | 1 | 027 | 1 | 4126735 | 99 | 93 | 100 | 99 | 100 |
| 443 | ERR044846 | 1 | 027 | 1 | 4129438 | 99 | 93 | 100 | 100 | 100 |
| 448 | ERR178513 | 1 | 017 | 37 | 4179967 | 99 | 94 | 100 | 99 | 99 |
| 451 | ERR178516 | 1 | 017 | 37 | 4224548 | 99 | 94 | 100 | 99 | 100 |
| 457 | ERR178522 | 1 | 017 | 37 | 4258390 | 99 | 94 | 100 | 100 | 98 |
| 458 | ERR178523 | 1 | 017 | 37 | 4267040 | 99 | 94 | 100 | 99 | 100 |
| 465 | ERR178530 | 1 | 017 | 37 | 4191399 | 99 | 94 | 100 | 99 | 99 |
| 470 | ERR178535 | 1 | 017 | 37 | 4165273 | 99 | 94 | 100 | 99 | 99 |
| 478 | ERR178543 | 1 | 017 | 37 | 4322578 | 99 | 94 | 99 | 99 | 99 |
| 480 | ERR178545 | 1 | 017 | 37 | 4363065 | 99 | 94 | 100 | 99 | 99 |
| 481 | ERR178546 | 1 | 017 | 37 | 4200117 | 99 | 94 | 99 | 99 | 99 |
| 484 | ERR178549 | 1 | 017 | 37 | 4292957 | 99 | 94 | 98 | 99 | 99 |
| 493 | ERR178558 | 1 | 017 | 86 | 4143844 | 99 | 94 | 99 | 99 | 100 |
| 502 | ERR178567 | 1 | 017 | 37 | 4239159 | 99 | 94 | 99 | 100 | 99 |
| 503 | ERR178568 | 1 | 017 | 37 | 4237912 | 99 | 94 | 99 | 99 | 99 |
| 505 | ERR178570 | 1 | 017 | 37 | 4238396 | 99 | 94 | 100 | 99 | 99 |
| 516 | ERR178582 | 1 | 017 | 37 | 4209939 | 99 | 94 | 99 | 99 | 99 |
| 527 | ERR178593 | 1 | 017 | 37 | 4249148 | 99 | 94 | 100 | 99 | 99 |
| 528 | ERR178594 | 1 | 017 | 37 | 4273998 | 99 | 94 | 99 | 99 | 99 |
| 534 | ERR178600 | 1 | 017 | 37 | 4209881 | 99 | 94 | 99 | 100 | 99 |
| 539 | ERR178605 | 1 | 017 | 37 | 4277032 | 99 | 94 | 100 | 100 | 99 |
| 545 | ERR340585 | 1 | | 375 | 4238952 | 99 | 94 | 100 | 99 | 100 |
| 546 | ERR340588 | 1 | | 37 | 4360850 | 99 | 94 | 100 | 100 | 100 |
| 556 | ERR340606 | 1 | 017 | 37 | 4187438 | 99 | 94 | 100 | 99 | 100 |
| 558 | ERR340612 | 1 | 017 | 37 | 4227719 | 99 | 94 | 100 | 100 | 99 |
| 562 | ERR340617 | 1 | 017 | 37 | 4253605 | 99 | 94 | 99 | 100 | 99 |
| 573 | ERR340635 | 1 | 017 | 37 | 4235383 | 99 | 94 | 100 | 100 | 100 |
| 580 | ERR340644 | 1 | 017 | 37 | 4376709 | 99 | 94 | 100 | 99 | 100 |
| 581 | ERR340645 | 1 | | 37 | 4237601 | 99 | 94 | 100 | 100 | 99 |
| 582 | ERR340648 | 1 | | 37 | 4216895 | 99 | 94 | 100 | 99 | 99 |
| 594 | ERR387365 | 1 | 017 | 37 | 4276776 | 99 | 94 | 99 | 100 | 100 |
| 609 | ERR387386 | 1 | | 37 | 4232987 | 99 | 94 | 100 | 99 | 99 |
| 612 | ERR387389 | 1 | 017 | 37 | 4225850 | 99 | 94 | 100 | 100 | 99 |
| 617 | ERR387397 | 1 | 017 | 37 | 4200802 | 99 | 94 | 100 | 99 | 99 |
| 618 | ERR387398 | 1 | 017 | 37 | 4193593 | 99 | 94 | 100 | 99 | 100 |
| 634 | ERR387416 | 1 | | 37 | 4234706 | 99 | 94 | 100 | 99 | 99 |
| 636 | ERR387418 | 1 | | 37 | 4330886 | 99 | 94 | 100 | 99 | 99 |
| 645 | ERR387430 | 1 | 017 | 37 | 4210918 | 99 | 94 | 100 | 99 | 100 |
| 656 | ERR387444 | 1 | 017 | 37 | 4289995 | 99 | 94 | 100 | 99 | 100 |
| 662 | ERR387451 | 1 | | 37 | 4266892 | 99 | 94 | 100 | 99 | 100 |
| 685 | ERR339497 | 1 | | 2 | 4089178 | 99 | 92 | 100 | 100 | 100 |
| 693 | ERR339505 | 1 | | 1 | 4388659 | 99 | 94 | 99 | 100 | 99 |
| 700 | ERR339512 | 1 | | 11 | 4914165 | 99 | 61 | 99 | 99 | 100 |
| 706 | ERR339518 | 1 | | 1 | 4299690 | 99 | 94 | 100 | 99 | 99 |
| 711 | ERR339523 | 1 | | 1 | 4140516 | 99 | 93 | 99 | 99 | 99 |
| 712 | ERR339524 | 1 | | 271 | 4224913 | 99 | 61 | 99 | 99 | 99 |
| 713 | ERR339525 | 1 | | 1 | 4138917 | 99 | 93 | 100 | 100 | 99 |
| 717 | ERR339529 | 1 | | 1 | 4141993 | 99 | 93 | 99 | 100 | 100 |
| 722 | ERR339534 | 1 | | NF | 4070455 | 99 | 94 | 99 | 99 | 96 |
| 726 | ERR339538 | 1 | | 1 | 4140816 | 99 | 93 | 99 | 99 | 99 |
| 727 | ERR339539 | 1 | | 1 | 4151940 | 99 | 93 | 99 | 99 | 100 |
| 728 | ERR339540 | 1 | | 2 | 4234882 | 99 | 15 | 100 | 0 | 45 |
| 742 | ERR339554 | 1 | | 2 | 4116871 | 99 | 15 | 100 | 9 | 0 |
| 744 | ERR339556 | 1 | | NF | 4364962 | 99 | 15 | 100 | 0 | 0 |
| 772 | ERR339584 | 1 | | 14 | 4909754 | 99 | 93 | 100 | 100 | 100 |
| 776 | ERR339588 | 1 | | 9 | 4277536 | 99 | 21 | 99 | 52 | 0 |
| 777 | ERR339589 | 1 | | 14 | 4197140 | 99 | 92 | 100 | 100 | 100 |
| 778 | ERR339590 | 1 | | 13 | 4235059 | 99 | 92 | 100 | 100 | 100 |
| 783 | ERR339595 | 1 | | 16 | 4566616 | 99 | 18 | 99 | 0 | 0 |
| 788 | ERR339600 | 1 | | 7 | 4158564 | 99 | 15 | 99 | 0 | 0 |
| 797 | ERR339609 | 1 | | NF | 4330922 | 99 | 13 | 99 | 0 | 0 |
| 800 | ERR339612 | 1 | | 36 | 4358547 | 99 | 94 | 100 | 99 | 99 |
| 812 | ERR339624 | 1 | | 36 | 4169043 | 99 | 94 | 99 | 99 | 99 |
| 815 | ERR339627 | 1 | | 55 | 4209049 | 99 | 94 | 100 | 100 | 99 |
| 822 | ERR339634 | 1 | | 36 | 4226199 | 99 | 94 | 100 | 100 | 99 |
| 828 | ERR339640 | 1 | | 13 | 4279711 | 99 | 92 | 100 | 99 | 99 |
| 833 | ERR339645 | 1 | | 36 | 4273132 | 99 | 94 | 99 | 99 | 100 |
| 836 | ERR339648 | 1 | | 5 | 4315241 | 99 | 39 | 99 | 99 | 99 |
| 844 | ERR339656 | 1 | | 5 | 4200814 | 99 | 39 | 99 | 99 | 99 |
| 849 | ERR339661 | 1 | | 5 | 4323632 | 99 | 41 | 99 | 99 | 98 |
| 856 | ERR339668 | 1 | | NF | 4131508 | 99 | 95 | 99 | 99 | 99 |
| 861 | ERR339673 | 1 | | 8 | 4245288 | 99 | 95 | 100 | 100 | 100 |
| 863 | ERR339675 | 1 | | 8 | 4198985 | 99 | 17 | 99 | 19 | 0 |
| 872 | ERR339684 | 1 | | 1 | 4154096 | 99 | 93 | 99 | 100 | 100 |
| 873 | ERR339685 | 1 | | 8 | 4528695 | 99 | 95 | 100 | 99 | 100 |
| 874 | ERR339686 | 1 | | 8 | 4297596 | 99 | 95 | 100 | 100 | 99 |
| 875 | ERR339687 | 1 | | 8 | 4284622 | 99 | 95 | 100 | 100 | 100 |
| 876 | ERR339688 | 1 | | 1 | 4149135 | 99 | 94 | 99 | 99 | 100 |
| 882 | ERR339694 | 1 | | 8 | 4289937 | 99 | 95 | 100 | 99 | 99 |
| 887 | ERR339699 | 1 | | 1 | 4157589 | 99 | 93 | 99 | 99 | 99 |
| 889 | ERR339701 | 1 | | 8 | 4352428 | 99 | 95 | 99 | 99 | 100 |
| 896 | ERR339708 | 1 | | 8 | 4274811 | 99 | 95 | 99 | 100 | 99 |
| 898 | ERR339710 | 1 | | 8 | 4243255 | 99 | 95 | 100 | 99 | 99 |
| 899 | ERR339711 | 1 | | 1 | 4160372 | 99 | 94 | 99 | 99 | 99 |
| 906 | ERR339718 | 1 | | 8 | 4183045 | 99 | 95 | 99 | 100 | 99 |
| 910 | ERR339722 | 1 | | 8 | 4252503 | 99 | 95 | 100 | 99 | 99 |
| 916 | ERR339728 | 1 | | 5 | 4381680 | 99 | 38 | 99 | 99 | 93 |
| 922 | ERR339734 | 1 | | 5 | 4298118 | 99 | 37 | 100 | 99 | 93 |
| 934 | ERR339746 | 1 | | 5 | 4291956 | 99 | 40 | 99 | 99 | 91 |
| 937 | ERR339749 | 1 | | 1 | 4153443 | 99 | 93 | 99 | 99 | 100 |
| 951 | ERR339763 | 1 | | 1 | 4165853 | 99 | 94 | 100 | 99 | 100 |
| 956 | ERR339768 | 1 | | 1 | 4161901 | 99 | 94 | 100 | 99 | 99 |
| 957 | ERR339769 | 1 | | 1 | 4273425 | 99 | 93 | 100 | 99 | 99 |
| 958 | ERR339770 | 1 | | 1 | 4181275 | 99 | 94 | 100 | 99 | 99 |
| 962 | ERR339774 | 1 | | 1 | 4147577 | 99 | 94 | 99 | 99 | 100 |
| 965 | ERR339777 | 1 | | 1 | 4149674 | 99 | 94 | 100 | 99 | 99 |
| 970 | ERR339782 | 1 | | 1 | 4152391 | 99 | 94 | 99 | 100 | 99 |
| 971 | ERR339783 | 1 | | 1 | 4145818 | 99 | 94 | 99 | 100 | 99 |
| 972 | ERR339784 | 1 | | 1 | 4154387 | 99 | 94 | 99 | 99 | 100 |
| 973 | ERR339785 | 1 | | 8 | 4282934 | 99 | 95 | 99 | 100 | 99 |
| 974 | ERR339786 | 1 | | 1 | 4160225 | 99 | 94 | 100 | 99 | 100 |
| 975 | ERR339787 | 1 | | 1 | 4181034 | 99 | 94 | 100 | 99 | 99 |
| 994 | ERR339806 | 1 | | 55 | 4478001 | 99 | 94 | 99 | 100 | 99 |
| 1008 | ERR339820 | 1 | | 49 | 4236766 | 99 | 20 | 99 | 7 | 45 |
| 1011 | ERR339823 | 1 | | 55 | 4078018 | 99 | 94 | 99 | 99 | 100 |
| 1015 | ERR339827 | 1 | | 1 | 4209821 | 99 | 94 | 99 | 99 | 100 |
| 1021 | ERR339833 | 1 | | 1 | 4182987 | 99 | 93 | 99 | 99 | 99 |
| 1028 | ERR339840 | 1 | | 1 | 4145777 | 99 | 93 | 100 | 100 | 99 |
| 1031 | ERR339843 | 1 | | 1 | 4238388 | 99 | 93 | 100 | 99 | 99 |
| 1037 | ERR339849 | 1 | | 1 | 4156881 | 99 | 93 | 99 | 99 | 99 |
| 1041 | ERR339853 | 1 | | 1 | 4148042 | 99 | 93 | 99 | 99 | 100 |
| 1049 | ERR339861 | 1 | | 1 | 4210408 | 99 | 94 | 99 | 100 | 100 |
| 1051 | ERR339863 | 1 | | 1 | 4143679 | 99 | 93 | 99 | 100 | 99 |
| 1054 | ERR339867 | 1 | | 1 | 4157696 | 99 | 93 | 99 | 99 | 100 |
| 1057 | ERR339870 | 1 | | 1 | 4158418 | 99 | 94 | 100 | 99 | 100 |
| 1061 | ERR339874 | 1 | | 1 | 4147237 | 99 | 94 | 99 | 99 | 100 |
| 1062 | ERR339875 | 1 | | 1 | 4157936 | 99 | 93 | 100 | 99 | 100 |
| 1063 | ERR339876 | 1 | | 1 | 4138676 | 99 | 94 | 99 | 99 | 98 |
| 1073 | ERR339886 | 1 | | 1 | 4157044 | 99 | 93 | 99 | 99 | 99 |
| 1075 | ERR339888 | 1 | | 1 | 4144504 | 99 | 93 | 100 | 99 | 99 |
| 1080 | ERR339893 | 1 | | 1 | 4147614 | 99 | 93 | 99 | 99 | 100 |
| 1085 | ERR339898 | 1 | | 1 | 4144205 | 99 | 94 | 99 | 99 | 100 |
| 1086 | ERR339899 | 1 | | 1 | 4092540 | 99 | 93 | 100 | 99 | 100 |
| 1092 | ERR339905 | 1 | | 1 | 4147932 | 99 | 93 | 100 | 99 | 99 |
| 1094 | ERR339907 | 1 | | 1 | 4142156 | 99 | 93 | 99 | 100 | 99 |
| 1095 | ERR339908 | 1 | | 1 | 4140757 | 99 | 93 | 100 | 100 | 100 |
| 1097 | ERR339910 | 1 | | 1 | 4155036 | 99 | 94 | 99 | 99 | 99 |
| 1107 | ERR339920 | 1 | | 1 | 4215741 | 99 | 93 | 100 | 99 | 99 |
| 1110 | ERR339923 | 1 | | 1 | 4145805 | 99 | 93 | 99 | 100 | 99 |
| 1115 | ERR339928 | 1 | | 1 | 4158476 | 99 | 94 | 100 | 99 | 100 |
| 1133 | ERR339947 | 1 | | 1 | 4240974 | 99 | 94 | 100 | 99 | 98 |
| 1134 | ERR339948 | 1 | | 1 | 4143754 | 99 | 94 | 99 | 99 | 100 |
| 1135 | ERR339949 | 1 | | 1 | 4144330 | 99 | 94 | 100 | 100 | 100 |
| 1136 | ERR339950 | 1 | | 1 | 4144741 | 99 | 93 | 100 | 100 | 100 |
| 1138 | ERR339952 | 1 | | 1 | 4145564 | 99 | 93 | 99 | 100 | 99 |
| 1148 | ERR339962 | 1 | | 8 | 4124284 | 99 | 69 | 100 | 100 | 99 |
| 1150 | ERR339964 | 1 | | 53 | 4082647 | 99 | 94 | 100 | 99 | 99 |
| 1156 | ERR339970 | 1 | | 65 | 4202405 | 99 | 94 | 100 | 99 | 99 |

| | | | | | | | | | | |
|---|---|---|---|---|---|---|---|---|---|---|
| 1163 | ERR339977 | 1 | | 8 | 4241055 | 99 | 95 | 100 | 100 | 100 |
| 1166 | ERR339980 | 1 | | 2 | 4127687 | 99 | 28 | 99 | 43 | 0 |
| 1171 | ERR339985 | 1 | | 2 | 4145143 | 99 | 94 | 100 | 99 | 100 |
| 1179 | ERR339993 | 1 | | 42 | 4133151 | 99 | 95 | 100 | 100 | 100 |
| 1181 | ERR339995 | 1 | | 42 | 4138834 | 99 | 95 | 99 | 99 | 99 |
| 1182 | ERR339996 | 1 | | 44 | 4999674 | 99 | 94 | 100 | 99 | 91 |
| 1183 | ERR339997 | 1 | | 44 | 4858244 | 99 | 94 | 99 | 99 | 99 |
| 1186 | ERR340000 | 1 | | 2 | 4135761 | 99 | 19 | 100 | 13 | 0 |
| 1188 | ERR340002 | 1 | | 73 | 4383977 | 99 | 93 | 99 | 100 | 99 |
| 1192 | ERR340006 | 1 | | 2 | 4212069 | 99 | 20 | 100 | 0 | 0 |
| 1203 | ERR340017 | 1 | | 2 | 4186415 | 99 | 25 | 100 | 13 | 0 |
| 1209 | ERR340024 | 1 | | 8 | 4138809 | 99 | 95 | 99 | 99 | 99 |
| 1218 | ERR340033 | 1 | | 8 | 4251412 | 99 | 95 | 100 | 99 | 98 |
| 1219 | ERR340034 | 1 | | 8 | 4229718 | 99 | 95 | 99 | 99 | 99 |
| 1223 | ERR340038 | 1 | | 92 | 4158977 | 99 | 94 | 99 | 99 | 99 |
| 1230 | ERR340045 | 1 | | 2 | 4160149 | 99 | 30 | 100 | 47 | 45 |
| 1231 | ERR340046 | 1 | | 72 | 4351353 | 99 | 64 | 100 | 99 | 82 |
| 1232 | ERR340047 | 1 | | 2 | 4097534 | 99 | 92 | 100 | 99 | 99 |
| 1239 | ERR340054 | 1 | | 2 | 4135737 | 99 | 92 | 100 | 100 | 100 |
| 1240 | ERR340055 | 1 | | 2 | 4133091 | 99 | 92 | 100 | 99 | 100 |
| 1243 | ERR340058 | 1 | | 8 | 4246101 | 99 | 95 | 99 | 100 | 100 |
| 1245 | ERR340060 | 1 | | 2 | 4139425 | 99 | 92 | 100 | 99 | 100 |
| 1254 | ERR340069 | 1 | | 15 | 4231082 | 99 | 92 | 100 | 99 | 99 |
| 1256 | ERR340071 | 1 | | 1 | 4252549 | 99 | 93 | 99 | 100 | 99 |
| 1258 | ERR340073 | 1 | | 45 | 4500076 | 99 | 92 | 100 | 99 | 100 |
| 1266 | ERR340081 | 1 | | 56 | 4122929 | 99 | 59 | 100 | 100 | 88 |
| 1267 | ERR340082 | 1 | | 74 | 4099420 | 99 | 91 | 99 | 99 | 99 |
| 1298 | ERR340113 | 1 | | 1 | 4162943 | 99 | 94 | 100 | 99 | 100 |
| 1299 | ERR340114 | 1 | | 97 | 4227581 | 99 | 94 | 99 | 99 | 99 |
| 1324 | ERR340139 | 1 | | 44 | 4437408 | 99 | 22 | 99 | 0 | 0 |
| 1331 | ERR340146 | 1 | | 13 | 4146580 | 99 | 93 | 100 | 100 | 100 |
| 1363 | ERR340178 | 1 | | 1 | 4176298 | 99 | 94 | 100 | 99 | 100 |
| 1382 | ERR340197 | 1 | | 14 | 5180153 | 99 | 92 | 100 | 99 | 100 |
| 1387 | ERR340202 | 1 | | 11 | 3967380 | 99 | 52 | 100 | 99 | 99 |
| 1391 | ERR340206 | 1 | | 44 | 4458327 | 99 | 66 | 100 | 99 | 93 |
| 1401 | ERR340216 | 1 | | 92 | 4284626 | 99 | 94 | 99 | 99 | 99 |
| 1404 | ERR340219 | 1 | | 5 | 4198266 | 99 | 39 | 100 | 99 | 86 |
| 1411 | ERR340226 | 1 | | 8 | 4090395 | 99 | 95 | 99 | 99 | 99 |
| 1420 | ERR340235 | 1 | | 1 | 4371680 | 99 | 94 | 100 | 100 | 100 |
| 1438 | ERR340253 | 1 | | 2 | 4144827 | 99 | 13 | 100 | 0 | 0 |
| 1439 | ERR340254 | 1 | | 8 | 4344006 | 99 | 45 | 99 | 99 | 83 |
| 1443 | ERR340258 | 1 | | 2 | 4141346 | 99 | 92 | 100 | 100 | 100 |
| 1446 | ERR340261 | 1 | | 16 | 4612802 | 99 | 95 | 99 | 100 | 100 |
| 1458 | ERR340273 | 1 | | 37 | 4241037 | 99 | 94 | 99 | 99 | 100 |
| 1465 | ERR340280 | 1 | | 1 | 4345429 | 99 | 33 | 99 | 6 | 0 |
| 1471 | ERR340286 | 1 | | 35 | 4239996 | 99 | 19 | 100 | 22 | 45 |
| 1473 | ERR340288 | 1 | | 1 | 4145232 | 99 | 93 | 99 | 99 | 100 |
| 1475 | ERR340290 | 1 | | 1 | 4134147 | 99 | 94 | 100 | 99 | 100 |
| 1476 | ERR340291 | 1 | | 8 | 4280429 | 99 | 95 | 99 | 100 | 100 |
| 1477 | ERR340292 | 1 | | 11 | 4004789 | 99 | 72 | 99 | 99 | 100 |
| 1482 | ERR340297 | 1 | | 5 | 4372474 | 99 | 39 | 99 | 99 | 93 |
| 1495 | ERR340310 | 1 | | 34 | 4094858 | 99 | 64 | 99 | 100 | 99 |
| 1498 | ERR340313 | 1 | | 2 | 4179677 | 99 | 19 | 99 | 13 | 0 |
| 1499 | ERR340314 | 1 | | 31 | 4229666 | 99 | 94 | 99 | 99 | 99 |
| 1505 | ERR126242 | 1 | | 6 | 4426590 | 99 | 23 | 99 | 6 | 0 |
| 1509 | ERR126246 | 1 | 027 | 1 | 4175801 | 99 | 93 | 100 | 100 | 100 |
| 1512 | ERR126249 | 1 | 002 | 8 | 4293970 | 99 | 95 | 99 | 100 | 99 |
| 1517 | ERR126254 | 1 | 284 | 35 | 4253805 | 99 | 25 | 100 | 20 | 0 |
| 1535 | ERR126272 | 1 | | 32 | 4321058 | 99 | 94 | 100 | 100 | 100 |
| 1538 | ERR126275 | 1 | 058 | 96 | 4186624 | 99 | 39 | 100 | 99 | 98 |
| 1539 | ERR126276 | 1 | 075 | 95 | 4126720 | 99 | 94 | 100 | 100 | 100 |
| 1542 | ERR126279 | 1 | 053 | 63 | 4027736 | 99 | 99 | 99 | 99 | 100 |
| 1548 | ERR126285 | 1 | 011 | 36 | 4189173 | 99 | 94 | 100 | 100 | 100 |
| 1552 | ERR126289 | 1 | 375 | 47 | 4284323 | 99 | 94 | 100 | 99 | 100 |
| 1553 | ERR126290 | 1 | | 3 | 4108247 | 99 | 49 | 99 | 100 | 94 |
| 1555 | ERR126292 | 1 | 014 | 49 | 4299911 | 99 | 92 | 100 | 100 | 99 |
| 1560 | ERR126297 | 1 | | 16 | 4608082 | 99 | 94 | 100 | 99 | 95 |
| 1562 | ERR142053 | 1 | | 2 | 4215148 | 99 | 95 | 100 | 100 | 100 |
| 1564 | ERR142055 | 1 | | 2 | 4334356 | 99 | 17 | 99 | 6 | 0 |
| 1565 | ERR142056 | 1 | | 6 | 4206285 | 99 | 65 | 100 | 99 | 85 |
| 1566 | ERR142057 | 1 | | 42 | 4151521 | 99 | 97 | 99 | 100 | 99 |
| 1570 | ERR142061 | 1 | | 1 | 4139664 | 99 | 94 | 99 | 100 | 99 |
| 1574 | ERR142065 | 1 | | 42 | 4188748 | 99 | 96 | 99 | 100 | 99 |
| 1580 | ERR142071 | 1 | | 1 | 4147591 | 99 | 93 | 100 | 100 | 99 |
| 1581 | ERR142072 | 1 | | 1 | 4144880 | 99 | 93 | 99 | 99 | 99 |
| 1585 | ERR142076 | 1 | | 42 | 4259034 | 99 | 95 | 99 | 100 | 100 |
| 1587 | ERR142078 | 1 | | 1 | 4150849 | 99 | 93 | 99 | 100 | 100 |
| 1592 | ERR142083 | 1 | | 1 | 4134791 | 99 | 93 | 100 | 100 | 100 |
| 1602 | ERR142093 | 1 | | 2 | 4182206 | 99 | 92 | 100 | 100 | 100 |
| 1612 | ERR142103 | 1 | | 1 | 4135843 | 99 | 93 | 99 | 100 | 99 |
| 1619 | ERR142110 | 1 | | 1 | 4136615 | 99 | 94 | 100 | 100 | 100 |
| 1623 | ERR142114 | 1 | | 8 | 4303646 | 99 | 95 | 99 | 99 | 99 |
| 1625 | ERR142116 | 1 | | 17 | 4221547 | 99 | 62 | 99 | 100 | 100 |
| 1626 | ERR142117 | 1 | | 14 | 4169920 | 99 | 93 | 100 | 100 | 100 |
| 1630 | ERR142121 | 1 | | 1 | 4149811 | 99 | 94 | 100 | 99 | 99 |
| 1635 | ERR142126 | 1 | | 1 | 4199868 | 99 | 94 | 100 | 99 | 100 |
| 1644 | ERR142135 | 1 | | 77 | 4310015 | 99 | 95 | 100 | 100 | 100 |
| 1651 | ERR142142 | 1 | | 1 | 4137178 | 99 | 93 | 100 | 100 | 100 |
| 1652 | ERR142143 | 1 | | 1 | 4202379 | 99 | 93 | 100 | 99 | 98 |
| 1653 | ERR142144 | 1 | | 1 | 4139593 | 99 | 93 | 100 | 99 | 99 |
| 1657 | ERR142148 | 1 | | 1 | 4145921 | 99 | 93 | 100 | 100 | 99 |
| 1659 | ERR142150 | 1 | | 1 | 4185190 | 99 | 93 | 99 | 100 | 100 |
| 1664 | ERR142155 | 1 | | 6 | 4265360 | 99 | 17 | 99 | 0 | 0 |
| 1667 | ERR142158 | 1 | | 58 | 4266360 | 99 | 13 | 100 | 0 | 0 |
| 1668 | ERR142159 | 1 | | 12 | 4101421 | 99 | 94 | 100 | 100 | 100 |
| 1670 | ERR142161 | 1 | | 12 | 4138801 | 99 | 94 | 100 | 100 | 100 |
| 1672 | ERR142163 | 1 | | 12 | 4052640 | 99 | 94 | 99 | 100 | 100 |
| 1676 | ERR142167 | 1 | | 17 | 4237517 | 99 | 59 | 100 | 99 | 100 |
| 1678 | ERR142169 | 1 | | 17 | 4286919 | 99 | 61 | 100 | 99 | 99 |
| 1684 | ERR142175 | 1 | | 13 | 4188743 | 99 | 92 | 100 | 100 | 100 |
| 1687 | ERR142178 | 1 | | 1 | 4192613 | 99 | 94 | 100 | 100 | 100 |
| 1694 | ERR142185 | 1 | | 54 | 4256327 | 99 | 99 | 100 | 100 | 99 |
| 1698 | ERR142189 | 1 | | 58 | 4278117 | 99 | 60 | 100 | 100 | 99 |
| 1700 | ERR142191 | 1 | | 58 | 4105934 | 99 | 12 | 99 | 0 | 0 |
| 1703 | ERR142194 | 1 | | 37 | 4251979 | 99 | 95 | 99 | 99 | 100 |
| 1706 | ERR142197 | 1 | | 37 | 4228519 | 99 | 94 | 99 | 99 | 100 |
| 1711 | ERR142202 | 1 | | 54 | 4381581 | 99 | 100 | 99 | 99 | 99 |
| 1732 | ERR142223 | 1 | | 37 | 4310109 | 99 | 94 | 100 | 99 | 99 |
| 1734 | ERR142225 | 1 | | 37 | 4278573 | 99 | 94 | 100 | 100 | 99 |
| 1736 | ERR142227 | 1 | | 11 | 4135684 | 99 | 52 | 99 | 99 | 100 |
| 1749 | ERR142240 | 1 | | 11 | 4073747 | 99 | 47 | 99 | 99 | 99 |
| 1754 | ERR142245 | 1 | | NF | 4147620 | 99 | 92 | 99 | 100 | 100 |
| 1756 | ERR142247 | 1 | | 5 | 4286377 | 99 | 42 | 100 | 99 | 100 |
| 1771 | ERR142262 | 1 | | NF | 4409654 | 99 | 95 | 100 | 100 | 99 |
| 1781 | ERR142272 | 1 | | NF | 4405104 | 99 | 94 | 99 | 99 | 99 |
| 1798 | ERR142289 | 1 | | NF | 4338432 | 99 | 61 | 99 | 99 | 97 |
| 1816 | ERR142307 | 1 | | NF | 4652766 | 99 | 61 | 100 | 99 | 93 |
| 1819 | ERR142310 | 1 | | 17 | 4214215 | 99 | 61 | 100 | 100 | 99 |
| 1823 | ERR142314 | 1 | | 1 | 4335853 | 99 | 94 | 99 | 100 | 100 |
| 1824 | ERR142315 | 1 | | 1 | 4205465 | 99 | 21 | 100 | 24 | 0 |
| 1825 | ERR142316 | 1 | | NF | 4388142 | 99 | 30 | 100 | 14 | 0 |
| 1831 | ERR142322 | 1 | | 58 | 4202142 | 99 | 17 | 100 | 26 | 0 |
| 1833 | ERR142324 | 1 | | 63 | 4025238 | 99 | 99 | 99 | 99 | 100 |
| 1834 | ERR142325 | 1 | | 63 | 4172600 | 99 | 99 | 100 | 100 | 100 |
| 1839 | ERR142330 | 1 | | 6 | 4325804 | 99 | 15 | 100 | 0 | 0 |
| 1840 | ERR142331 | 1 | | NF | 4641016 | 99 | 15 | 100 | 0 | 0 |
| 1860 | ERR142351 | 1 | | 6 | 4386076 | 99 | 17 | 100 | 0 | 0 |
| 1866 | ERR142357 | 1 | | 6 | 4388539 | 99 | 63 | 99 | 99 | 99 |
| 1874 | ERR142365 | 1 | | 44 | 4354320 | 99 | 60 | 98 | 99 | 89 |
| 1895 | ERR142386 | 1 | | NF | 4154709 | 99 | 95 | 99 | 100 | 98 |
| 1897 | ERR142388 | 1 | | 35 | 4251156 | 99 | 7 | 99 | 9 | 0 |
| 1903 | ERR142394 | 1 | | 11 | 4056570 | 99 | 48 | 99 | 100 | 100 |
| 1913 | ERR142404 | 1 | | NF | 4439077 | 99 | 94 | 99 | 99 | 98 |
| 1921 | ERR142412 | 1 | | NF | 4134361 | 99 | 95 | 98 | 99 | 99 |
| 1924 | ERR142415 | 1 | | 11 | 3992315 | 99 | 47 | 99 | 99 | 99 |
| 1932 | ERR142423 | 1 | | NF | 4138983 | 99 | 95 | 100 | 99 | 99 |
| 1942 | ERR142433 | 1 | | 42 | 4042012 | 99 | 95 | 100 | 99 | 99 |
| 1948 | ERR142439 | 1 | | NF | 4135958 | 99 | 95 | 99 | 99 | 99 |
| 1949 | ERR142440 | 1 | | 42 | 4228574 | 99 | 95 | 99 | 100 | 99 |
| 1950 | ERR142441 | 1 | | NF | 4287782 | 99 | 95 | 99 | 99 | 89 |
| 1966 | ERR142457 | 1 | | 6 | 4343054 | 99 | 18 | 99 | 13 | 0 |
| 1972 | ERR142463 | 1 | | 11 | 4011657 | 99 | 46 | 99 | 99 | 100 |
| 1976 | ERR142467 | 1 | | NF | 4631224 | 99 | 61 | 99 | 99 | 96 |
| 1980 | ERR142471 | 1 | | 6 | 4236745 | 99 | 62 | 100 | 99 | 93 |
| 1984 | ERR142475 | 1 | | 44 | 4347257 | 99 | 94 | 99 | 99 | 98 |
| 1993 | ERR142484 | 1 | | 11 | 4036207 | 99 | 44 | 99 | 99 | 99 |
| 1996 | ERR142487 | 1 | | NF | 4310716 | 99 | 14 | 99 | 0 | 0 |
| 2003 | ERR142494 | 1 | | 1 | 4362693 | 99 | 94 | 100 | 99 | 100 |
| 2024 | ERR142515 | 1 | | 3 | 4225267 | 99 | 58 | 99 | 99 | 100 |
| 2035 | ERR142526 | 1 | | NF | 4148478 | 99 | 92 | 99 | 99 | 98 |
| 2036 | ERR142527 | 1 | | NF | 4071313 | 99 | 94 | 99 | 99 | 96 |
| 2040 | ERR142531 | 1 | | 1 | 4160276 | 99 | 93 | 99 | 99 | 99 |
| 2041 | ERR142532 | 1 | | 1 | 4189467 | 99 | 92 | 99 | 99 | 99 |
| 2053 | ERR142544 | 1 | | 2 | 4115155 | 99 | 15 | 100 | 9 | 0 |
| 2056 | ERR142547 | 1 | | NF | 4287653 | 99 | 38 | 99 | 99 | 91 |
| 2064 | ERR142555 | 1 | | 1 | 4152112 | 99 | 93 | 100 | 99 | 99 |
| 2066 | ERR142557 | 1 | | 1 | 4145891 | 99 | 93 | 100 | 99 | 99 |
| 2077 | ERR142568 | 1 | | 1 | 4145750 | 99 | 92 | 99 | 99 | 80 |
| 2078 | ERR142569 | 1 | | 1 | 4139639 | 99 | 93 | 99 | 99 | 99 |
| 2092 | ERR142583 | 1 | | 2 | 4686958 | 99 | 14 | 99 | 0 | 45 |
| 2096 | ERR142587 | 1 | | 55 | 4183949 | 99 | 94 | 99 | 99 | 99 |
| 2114 | ERR142605 | 1 | | 55 | 4101535 | 99 | 94 | 98 | 99 | 99 |
| 2119 | ERR142610 | 1 | | 37 | 4193310 | 99 | 94 | 99 | 99 | 99 |
| 2123 | ERR232378 | 1 | 039 | 26 | 4302249 | 99 | 95 | 100 | 99 | 100 |
| 2137 | ERR232392 | 1 | | 152 | 4156043 | 99 | 95 | 100 | 99 | 99 |
| 2138 | ERR232393 | 1 | | 162 | 4165704 | 99 | 41 | 100 | 99 | 94 |
| 2143 | ERR232398 | 1 | | 177 | 4249273 | 99 | 40 | 99 | 76 | 0 |
| 2156 | ERR251721 | 1 | | 49 | 4236452 | 99 | 92 | 100 | 99 | 100 |
| 2165 | ERR251730 | 1 | | 1 | 4157806 | 99 | 94 | 99 | 100 | 100 |
| 2166 | ERR251731 | 1 | | 8 | 4282732 | 99 | 95 | 99 | 99 | 100 |
| 2175 | ERR251740 | 1 | | 1 | 4159812 | 99 | 93 | 100 | 99 | 100 |
| 2179 | ERR251744 | 1 | | 1 | 4156387 | 99 | 93 | 100 | 99 | 99 |
| 2186 | ERR251751 | 1 | | 1 | 4182487 | 99 | 94 | 99 | 99 | 100 |
| 2204 | ERR251769 | 1 | | 8 | 4267058 | 99 | 95 | 99 | 99 | 99 |
| 2205 | ERR251770 | 1 | | 1 | 4144619 | 99 | 93 | 100 | 99 | 100 |
| 2206 | ERR251771 | 1 | | 2 | 4135947 | 99 | 14 | 99 | 0 | 0 |
| 2213 | ERR251778 | 1 | | NF | 4171501 | 99 | 15 | 100 | 0 | 0 |
| 2215 | ERR251780 | 1 | | NF | 4257271 | 99 | 93 | 100 | 99 | 100 |
| 2217 | ERR251782 | 1 | | 2 | 4160525 | 99 | 24 | 100 | 17 | 0 |
| 2225 | ERR251790 | 1 | | 1 | 4121008 | 99 | 94 | 100 | 99 | 100 |
| 2229 | ERR251794 | 1 | | 67 | 4893559 | 99 | 95 | 100 | 99 | 99 |
| 2232 | ERR251797 | 1 | | 353 | 5141710 | 99 | 93 | 100 | 99 | 94 |
| 2233 | ERR251798 | 1 | | 1 | 4908156 | 99 | 93 | 99 | 99 | 100 |
| 2235 | ERR251800 | 1 | | NF | 4728873 | 99 | 95 | 99 | 100 | 99 |
| 2249 | ERR251814 | 1 | | 16 | 5448254 | 99 | 94 | 100 | 100 | 100 |
| 2250 | ERR251815 | 1 | | 44 | 5144407 | 99 | 93 | 100 | 99 | 98 |
| 2252 | ERR251817 | 1 | | 33 | 4894843 | 99 | 93 | 100 | 99 | 100 |

| | | | | | | | | | | |
|---|---|---|---|---|---|---|---|---|---|---|
| 2253 | ERR251818 | 1 | | 44 | 4591049 | 99 | 95 | 100 | 99 | 100 |
| 2257 | ERR251822 | 1 | | 54 | 4631384 | 99 | 100 | 100 | 99 | 100 |
| 2258 | ERR251823 | 1 | | 1 | 4144685 | 99 | 94 | 100 | 100 | 99 |
| 2259 | ERR251824 | 1 | | 3 | 4310895 | 99 | 62 | 100 | 99 | 93 |
| 2262 | ERR251827 | 1 | | 14 | 4369195 | 99 | 92 | 99 | 100 | 99 |
| 2266 | ERR251831 | 1 | | 1 | 4733475 | 99 | 93 | 100 | 99 | 100 |
| 2272 | ERR251837 | 1 | | 8 | 4496514 | 99 | 95 | 100 | 99 | 99 |
| 2280 | ERR251845 | 1 | | 14 | 4397928 | 99 | 92 | 100 | 100 | 99 |
| 2282 | ERR251847 | 1 | | 3 | 5049312 | 99 | 55 | 96 | 99 | 89 |
| 2290 | ERR251868 | 1 | | NF | 4276286 | 99 | 46 | 91 | 98 | 90 |
| 2291 | ERR251872 | 1 | | 44 | 4403322 | 99 | 62 | 100 | 99 | 92 |
| 2295 | ERR251877 | 1 | | 3 | 4280328 | 99 | 20 | 100 | 13 | 0 |
| 2306 | ERR272065 | 1 | | 58 | 4248208 | 99 | 15 | 99 | 11 | 0 |
| 2316 | ERR272075 | 1 | | 54 | 5073336 | 99 | 98 | 100 | 100 | 99 |
| 2323 | ERR272082 | 1 | | 271 | 4174509 | 99 | 61 | 99 | 100 | 98 |
| 2324 | ERR272083 | 1 | | 3 | 4311515 | 99 | 18 | 100 | 0 | 45 |
| 2326 | ERR272085 | 1 | | 271 | 4300458 | 99 | 24 | 100 | 30 | 0 |
| 2329 | ERR272088 | 1 | | 3 | 4271798 | 99 | 58 | 99 | 100 | 89 |
| 2330 | ERR272089 | 1 | | 3 | 4195493 | 99 | 60 | 100 | 100 | 99 |
| 2332 | ERR272091 | 1 | | 271 | 4233451 | 99 | 59 | 99 | 100 | 94 |
| 2335 | ERR272094 | 1 | | 271 | 4196087 | 99 | 60 | 100 | 100 | 99 |
| 2342 | ERR272101 | 1 | | 3 | 4846724 | 99 | 60 | 99 | 100 | 93 |
| 2344 | ERR272103 | 1 | | 271 | 4385981 | 99 | 60 | 100 | 99 | 99 |
| 2347 | ERR272106 | 1 | | 3 | 4231716 | 99 | 59 | 100 | 99 | 96 |
| 2348 | ERR272107 | 1 | | 271 | 4166232 | 99 | 14 | 99 | 27 | 0 |
| 2353 | ERR272112 | 1 | | 3 | 4189371 | 99 | 59 | 100 | 99 | 99 |
| 2363 | ERR272122 | 1 | | 3 | 4070501 | 99 | 60 | 100 | 100 | 99 |
| 2365 | ERR272124 | 1 | | 3 | 4337668 | 99 | 15 | 100 | 13 | 0 |
| 2374 | ERR272133 | 1 | | 3 | 4235662 | 99 | 64 | 100 | 100 | 94 |
| 2379 | ERR272138 | 1 | | 3 | 4258830 | 99 | 62 | 100 | 100 | 94 |
| 2388 | ERR272147 | 1 | | 3 | 4259852 | 99 | 59 | 100 | 99 | 99 |
| 2394 | ERR272153 | 1 | | 6 | 4212040 | 99 | 66 | 99 | 100 | 99 |
| 2395 | ERR272154 | 1 | | 6 | 4443790 | 99 | 15 | 100 | 0 | 0 |
| 2402 | ERR272161 | 1 | | 6 | 4445301 | 99 | 15 | 99 | 0 | 0 |
| 2410 | ERR272169 | 1 | | 6 | 4568740 | 99 | 62 | 99 | 99 | 96 |
| 2413 | ERR272172 | 1 | | 6 | 4283065 | 99 | 61 | 99 | 100 | 99 |
| 2415 | ERR272174 | 1 | | 6 | 4441419 | 99 | 64 | 99 | 100 | 94 |
| 2427 | ERR272186 | 1 | | 6 | 4324481 | 99 | 59 | 100 | 99 | 93 |
| 2428 | ERR272187 | 1 | | 6 | 4329806 | 99 | 18 | 100 | 36 | 0 |
| 2433 | ERR272192 | 1 | | 7 | 4387144 | 99 | 18 | 100 | 32 | 0 |
| 2434 | ERR272193 | 1 | | NF | 4114850 | 99 | 14 | 99 | 13 | 0 |
| 2443 | ERR272202 | 1 | | 7 | 4255424 | 99 | 16 | 100 | 0 | 0 |
| 2449 | ERR272208 | 1 | | 7 | 4151332 | 99 | 20 | 99 | 0 | 0 |
| 2454 | ERR272213 | 1 | | 7 | 4322148 | 99 | 24 | 99 | 15 | 0 |
| 2464 | ERR272223 | 1 | | 7 | 4222825 | 99 | 22 | 100 | 13 | 0 |
| 2466 | ERR272225 | 1 | | 7 | 4222224 | 99 | 19 | 99 | 0 | 0 |
| 2483 | ERR272242 | 1 | | NF | 4502564 | 99 | 60 | 100 | 99 | 99 |
| 2487 | ERR272246 | 1 | | 44 | 4327951 | 99 | 64 | 100 | 99 | 99 |
| 2488 | ERR272247 | 1 | | 44 | 4460316 | 99 | 28 | 100 | 0 | 0 |
| 2492 | ERR272251 | 1 | | 44 | 4356126 | 99 | 66 | 99 | 100 | 87 |
| 2501 | ERR272260 | 1 | | 44 | 4447813 | 99 | 94 | 99 | 100 | 99 |
| 2502 | ERR272261 | 1 | | 44 | 4456261 | 99 | 94 | 99 | 99 | 100 |
| 2505 | ERR467531 | 1 | | NF | 4674931 | 99 | 55 | 99 | 92 | 94 |
| 2506 | ERR467532 | 1 | | 41 | 4259982 | 99 | 94 | 100 | 99 | 100 |
| 2517 | ERR467543 | 1 | | 54 | 4612233 | 99 | 100 | 99 | 100 | 99 |
| 2518 | ERR467544 | 1 | | 54 | 4606462 | 99 | 100 | 99 | 99 | 99 |
| 2522 | ERR467548 | 1 | | 8 | 4307316 | 99 | 95 | 99 | 100 | 99 |
| 2525 | ERR467551 | 1 | | 1 | 4245295 | 99 | 94 | 99 | 99 | 99 |
| 2531 | ERR467557 | 1 | | 1 | 4198244 | 99 | 93 | 100 | 100 | 99 |
| 2533 | ERR467559 | 1 | | 54 | 4612847 | 99 | 99 | 99 | 99 | 99 |
| 2538 | ERR467564 | 1 | | 1 | 4193813 | 99 | 93 | 100 | 99 | 100 |
| 2546 | ERR467572 | 1 | | 58 | 4219409 | 99 | 65 | 99 | 100 | 93 |
| 2566 | ERR467592 | 1 | | 54 | 4600296 | 99 | 100 | 100 | 99 | 100 |
| 2568 | ERR467594 | 1 | | 54 | 4598810 | 99 | 100 | 100 | 100 | 100 |
| 2569 | ERR467595 | 1 | | 54 | 4600647 | 99 | 99 | 100 | 100 | 99 |
| 2570 | ERR467596 | 1 | | 54 | 4600803 | 99 | 99 | 99 | 99 | 99 |
| 2575 | ERR467601 | 1 | | 54 | 4571238 | 99 | 99 | 100 | 99 | 99 |
| 2578 | ERR467604 | 1 | | 8 | 4292187 | 99 | 95 | 100 | 99 | 100 |
| 2581 | ERR467607 | 1 | | 54 | 4607000 | 99 | 99 | 99 | 99 | 100 |
| 2584 | ERR467610 | 1 | | 54 | 4595066 | 99 | 100 | 99 | 99 | 99 |
| 2585 | ERR467611 | 1 | | 54 | 4666383 | 99 | 99 | 100 | 100 | 99 |
| 2596 | ERR467622 | 1 | | 37 | 4264152 | 99 | 93 | 99 | 100 | 100 |
| 2598 | ERR467624 | 1 | | 37 | 4272496 | 99 | 93 | 99 | 99 | 99 |
| 2599 | ERR467625 | 1 | | 2 | 4188703 | 99 | 16 | 99 | 0 | 0 |
| 2603 | ERR008617 | 1 | | 1 | 4216349 | 99 | 93 | 100 | 99 | 98 |
| 2614 | ERR008628 | 1 | | 1 | 4186855 | 99 | 92 | 100 | 99 | 99 |
| 2619 | ERR008633 | 1 | | 1 | 4155854 | 99 | 92 | 100 | 100 | 98 |
| 2642 | ERR111457 | 1 | 027 | 1 | 4098710 | 99 | 90 | 100 | 99 | 99 |
| 2644 | ERR111459 | 1 | 027 | 1 | 4124822 | 99 | 91 | 99 | 99 | 89 |
| 2657 | E13 | 1 | 017 | 37 | 4172340 | 99 | 95 | 100 | 100 | 100 |
| 2678 | T3 | 1 | 012 | 54 | 4087983 | 99 | 100 | 100 | 100 | 100 |
| 2684 | ERR604989 | 1 | 003 | 12 | 5207782 | 99 | 91 | 100 | 100 | 66 |
| 2687 | ERR604998 | 1 | 351 | 53 | 4203772 | 99 | 94 | 99 | 100 | 99 |
| 2693 | ERR605015 | 1 | 003 | 12 | 4118842 | 99 | 95 | 100 | 99 | 99 |
| 2694 | ERR605018 | 1 | 131 | 122 | 4250877 | 99 | 96 | 100 | 100 | 99 |
| 2698 | ERR605027 | 1 | 020 | 2 | 4187574 | 99 | 12 | 100 | 0 | 0 |
| 2700 | ERR605030 | 1 | 020 | 2 | 4035871 | 99 | 11 | 99 | 0 | 0 |
| 3 | ERR024644 | 1 | 027 | 49 | 4365167 | 100 | 92 | 100 | 100 | 99 |
| 7 | ERR024649 | 1 | 027 | 1 | 4172262 | 100 | 94 | 100 | 99 | 99 |
| 8 | ERR024650 | 1 | 027 | 1 | 4171736 | 100 | 94 | 100 | 99 | 99 |
| 9 | ERR024651 | 1 | 027 | 1 | 4167415 | 100 | 93 | 100 | 99 | 100 |
| 12 | ERR024655 | 1 | 027 | 1 | 4139798 | 100 | 94 | 99 | 99 | 100 |
| 14 | ERR024657 | 1 | 027 | 1 | 4132647 | 100 | 94 | 99 | 99 | 100 |
| 17 | ERR024661 | 1 | 027 | 1 | 4169258 | 100 | 93 | 100 | 99 | 99 |
| 26 | ERR025345 | 1 | 027 | 1 | 4139359 | 100 | 93 | 100 | 99 | 100 |
| 32 | ERR025352 | 1 | 027 | 1 | 4129805 | 100 | 93 | 100 | 99 | 100 |
| 36 | ERR026352 | 1 | 027 | 1 | 4080839 | 100 | 93 | 100 | 100 | 100 |
| 38 | ERR026354 | 1 | 027 | 1 | 4130380 | 100 | 93 | 100 | 99 | 99 |
| 40 | ERR026356 | 1 | 027 | 1 | 4133421 | 100 | 93 | 99 | 99 | 99 |
| 42 | ERR026358 | 1 | 027 | 1 | 4129794 | 100 | 93 | 99 | 99 | 99 |
| 49 | ERR026365 | 1 | 027 | 1 | 4137018 | 100 | 93 | 100 | 99 | 99 |
| 53 | ERR026369 | 1 | 027 | 1 | 4133638 | 100 | 93 | 99 | 100 | 99 |
| 54 | ERR026370 | 1 | 027 | 1 | 4134014 | 100 | 93 | 100 | 100 | 99 |
| 56 | ERR026372 | 1 | 027 | 1 | 4139920 | 100 | 93 | 100 | 99 | 99 |
| 59 | ERR026375 | 1 | 027 | 1 | 4175585 | 100 | 93 | 99 | 100 | 100 |
| 62 | ERR026378 | 1 | 027 | 1 | 4127748 | 100 | 93 | 100 | 100 | 100 |
| 63 | ERR026379 | 1 | 027 | 1 | 4130136 | 100 | 93 | 99 | 100 | 99 |
| 68 | ERR026384 | 1 | 027 | 1 | 4138520 | 100 | 93 | 99 | 99 | 100 |
| 76 | ERR026392 | 1 | 027 | 1 | 4131494 | 100 | 93 | 100 | 100 | 99 |
| 77 | ERR026393 | 1 | 027 | 1 | 4138010 | 100 | 93 | 99 | 100 | 99 |
| 80 | ERR026396 | 1 | 027 | 1 | 4141951 | 100 | 93 | 100 | 99 | 99 |
| 82 | ERR026398 | 1 | 027 | 1 | 4135469 | 100 | 93 | 100 | 99 | 99 |
| 85 | ERR026401 | 1 | 027 | 1 | 4138883 | 100 | 93 | 100 | 100 | 100 |
| 86 | ERR026402 | 1 | 027 | 1 | 4076629 | 100 | 93 | 100 | 99 | 99 |
| 88 | ERR026404 | 1 | 027 | 1 | 4133937 | 100 | 93 | 100 | 99 | 99 |
| 89 | ERR026405 | 1 | 027 | 1 | 4131293 | 100 | 93 | 99 | 99 | 99 |
| 99 | ERR026415 | 1 | 027 | 1 | 4136311 | 100 | 93 | 100 | 99 | 99 |
| 103 | ERR026419 | 1 | 027 | 1 | 4134698 | 100 | 93 | 99 | 99 | 99 |
| 106 | ERR026422 | 1 | 027 | 1 | 4135752 | 100 | 93 | 100 | 100 | 100 |
| 107 | ERR026423 | 1 | 027 | 1 | 4131729 | 100 | 93 | 100 | 100 | 99 |
| 116 | ERR026835 | 1 | 027 | 1 | 4135419 | 100 | 93 | 99 | 100 | 99 |
| 118 | ERR026837 | 1 | 027 | 1 | 4147116 | 100 | 93 | 99 | 100 | 99 |
| 124 | ERR026843 | 1 | 027 | 1 | 4156826 | 100 | 93 | 100 | 100 | 100 |
| 127 | ERR026846 | 1 | 027 | 1 | 4370707 | 100 | 94 | 100 | 100 | 99 |
| 129 | ERR026848 | 1 | 027 | 1 | 4132540 | 100 | 93 | 99 | 100 | 100 |
| 134 | ERR026854 | 1 | 027 | NF | 4135983 | 100 | 93 | 99 | 100 | 98 |
| 136 | ERR026856 | 1 | 027 | 1 | 4146678 | 100 | 93 | 100 | 100 | 100 |
| 142 | ERR026863 | 1 | 027 | 1 | 4138133 | 100 | 93 | 100 | 100 | 100 |
| 145 | ERR026866 | 1 | 027 | 1 | 4136226 | 100 | 93 | 100 | 100 | 100 |
| 146 | ERR026867 | 1 | 027 | 1 | 4143149 | 100 | 93 | 100 | 99 | 100 |
| 150 | ERR026871 | 1 | 027 | 1 | 4150003 | 100 | 93 | 100 | 100 | 99 |
| 153 | ERR026874 | 1 | 027 | 1 | 4136598 | 100 | 93 | 99 | 99 | 99 |
| 154 | ERR026875 | 1 | 027 | 1 | 4143786 | 100 | 93 | 100 | 100 | 99 |
| 159 | ERR026880 | 1 | 027 | 1 | 4137127 | 100 | 93 | 99 | 100 | 99 |
| 160 | ERR028720 | 1 | | 1 | 4108918 | 100 | 93 | 98 | 99 | 98 |
| 162 | ERR028722 | 1 | 27 | 1 | 4143519 | 100 | 93 | 99 | 99 | 100 |
| 163 | ERR028723 | 1 | 027 | 1 | 4082605 | 100 | 93 | 100 | 99 | 100 |
| 166 | ERR028726 | 1 | 027 | 1 | 4134085 | 100 | 93 | 99 | 100 | 99 |
| 170 | ERR028730 | 1 | 027 | 1 | 4136823 | 100 | 93 | 100 | 100 | 99 |
| 176 | ERR029483 | 1 | 027 | 1 | 4147374 | 100 | 93 | 100 | 99 | 100 |
| 177 | ERR029484 | 1 | 027 | 1 | 4127492 | 100 | 93 | 100 | 99 | 99 |
| 178 | ERR029485 | 1 | 027 | 1 | 4131765 | 100 | 93 | 99 | 99 | 100 |
| 183 | ERR029490 | 1 | 027 | 1 | 4139757 | 100 | 93 | 99 | 99 | 99 |
| 188 | ERR029495 | 1 | 027 | 1 | 4148096 | 100 | 93 | 100 | 99 | 100 |
| 189 | ERR029496 | 1 | 027 | 1 | 4144615 | 100 | 93 | 100 | 100 | 99 |
| 191 | ERR029498 | 1 | 027 | 1 | 4145246 | 100 | 93 | 99 | 100 | 99 |
| 192 | ERR029499 | 1 | 027 | 1 | 4135010 | 100 | 93 | 100 | 100 | 99 |
| 194 | ERR029501 | 1 | 027 | 1 | 4136568 | 100 | 93 | 100 | 99 | 100 |
| 196 | ERR029503 | 1 | 027 | 1 | 4136329 | 100 | 93 | 99 | 99 | 100 |
| 197 | ERR029504 | 1 | 027 | 1 | 4195444 | 100 | 93 | 99 | 100 | 100 |
| 200 | ERR029507 | 1 | 027 | 1 | 4174147 | 100 | 93 | 100 | 99 | 100 |
| 201 | ERR029508 | 1 | 027 | 1 | 4078949 | 100 | 93 | 99 | 99 | 99 |
| 202 | ERR029509 | 1 | 027 | 1 | 4146254 | 100 | 93 | 100 | 100 | 99 |
| 204 | ERR029511 | 1 | 027 | 1 | 4172320 | 100 | 93 | 100 | 99 | 99 |
| 206 | ERR029513 | 1 | 027 | 1 | 4133034 | 100 | 93 | 99 | 100 | 99 |
| 209 | ERR029516 | 1 | 027 | 1 | 4144424 | 100 | 93 | 100 | 99 | 99 |
| 210 | ERR029517 | 1 | 027 | 1 | 4157292 | 100 | 93 | 100 | 99 | 99 |
| 214 | ERR029521 | 1 | 027 | 61 | 4152181 | 100 | 94 | 100 | 100 | 99 |
| 218 | ERR029525 | 1 | 027 | 1 | 4097590 | 100 | 93 | 100 | 99 | 100 |
| 224 | ERR029531 | 1 | 027 | 1 | 4140073 | 100 | 93 | 100 | 99 | 99 |
| 225 | ERR029532 | 1 | | 1 | 4143578 | 100 | 93 | 99 | 99 | 99 |
| 226 | ERR029534 | 1 | 027 | 1 | 4146724 | 100 | 93 | 100 | 99 | 99 |
| 227 | ERR029535 | 1 | 027 | 1 | 4136717 | 100 | 93 | 100 | 100 | 99 |
| 229 | ERR029537 | 1 | 027 | 1 | 4427165 | 100 | 93 | 100 | 100 | 100 |
| 231 | ERR029539 | 1 | 027 | 1 | 4173003 | 100 | 93 | 100 | 100 | 99 |
| 232 | ERR029540 | 1 | 027 | 1 | 4137556 | 100 | 93 | 100 | 100 | 99 |
| 238 | ERR030284 | 1 | 027 | 1 | 4133822 | 100 | 93 | 100 | 100 | 100 |
| 244 | ERR030290 | 1 | 027 | 1 | 4132726 | 100 | 93 | 99 | 99 | 99 |
| 247 | ERR030293 | 1 | 027 | 1 | 4134552 | 100 | 93 | 99 | 100 | 100 |
| 248 | ERR030294 | 1 | 027 | 1 | 4140365 | 100 | 93 | 100 | 100 | 99 |
| 254 | ERR030300 | 1 | 027 | 1 | 4133460 | 100 | 93 | 99 | 100 | 100 |
| 255 | ERR030301 | 1 | 027 | 1 | 4135249 | 100 | 93 | 100 | 100 | 99 |
| 256 | ERR030302 | 1 | 027 | 1 | 4134803 | 100 | 93 | 99 | 100 | 100 |
| 261 | ERR030307 | 1 | 027 | 1 | 4130345 | 100 | 93 | 99 | 99 | 100 |
| 264 | ERR030310 | 1 | 027 | 1 | 4133174 | 100 | 93 | 99 | 99 | 100 |
| 267 | ERR030313 | 1 | 027 | 1 | 4138294 | 100 | 93 | 100 | 99 | 100 |
| 268 | ERR030314 | 1 | 027 | 1 | 4135298 | 100 | 93 | 100 | 100 | 100 |
| 277 | ERR030323 | 1 | 027 | 1 | 4071551 | 100 | 93 | 100 | 100 | 100 |
| 278 | ERR030324 | 1 | 027 | 1 | 4255370 | 100 | 93 | 100 | 99 | 99 |
| 279 | ERR030325 | 1 | 027 | 1 | 4126228 | 100 | 93 | 100 | 99 | 100 |
| 282 | ERR030328 | 1 | 027 | 1 | 4137106 | 100 | 93 | 100 | 100 | 99 |
| 283 | ERR030329 | 1 | 027 | 1 | 4132151 | 100 | 93 | 99 | 99 | 99 |
| 287 | ERR030333 | 1 | 027 | 1 | 4166265 | 100 | 93 | 99 | 99 | 99 |
| 291 | ERR030337 | 1 | 027 | 1 | 4292614 | 100 | 93 | 99 | 99 | 100 |
| 298 | ERR030344 | 1 | 027 | 1 | 4205184 | 100 | 93 | 99 | 99 | 99 |
| 300 | ERR030346 | 1 | 027 | 1 | 4146538 | 100 | 93 | 99 | 99 | 99 |
| 306 | ERR030352 | 1 | 027 | 1 | 4137141 | 100 | 93 | 99 | 100 | 99 |

| | | | | | | | | | | |
|---|---|---|---|---|---|---|---|---|---|---|
| 307 | ERR030353 | 1 | 027 | 1 | 4134144 | 100 | 93 | 100 | 100 | 99 |
| 308 | ERR030354 | 1 | 027 | 1 | 4137370 | 100 | 93 | 100 | 99 | 100 |
| 312 | ERR030358 | 1 | 027 | 1 | 4132960 | 100 | 93 | 99 | 100 | 99 |
| 322 | ERR030368 | 1 | 027 | 1 | 4082589 | 100 | 93 | 99 | 100 | 100 |
| 328 | ERR030374 | 1 | 027 | 1 | 4140350 | 100 | 93 | 100 | 100 | 100 |
| 332 | ERR030378 | 1 | 027 | 1 | 4139917 | 100 | 93 | 100 | 99 | 100 |
| 335 | ERR030381 | 1 | 027 | 1 | 4999683 | 100 | 93 | 99 | 100 | 99 |
| 337 | ERR030385 | 1 | 027 | 1 | 4078480 | 100 | 93 | 100 | 100 | 99 |
| 339 | ERR030387 | 1 | 027 | 1 | 4136968 | 100 | 93 | 100 | 99 | 100 |
| 341 | ERR030389 | 1 | 027 | 1 | 4135067 | 100 | 93 | 100 | 99 | 100 |
| 344 | ERR030392 | 1 | 027 | 1 | 4134064 | 100 | 93 | 100 | 100 | 100 |
| 349 | ERR030397 | 1 | 027 | 1 | 4077605 | 100 | 93 | 100 | 99 | 100 |
| 352 | ERR031674 | 1 | 027 | 1 | 4158897 | 100 | 93 | 100 | 99 | 99 |
| 353 | ERR031675 | 1 | 027 | 1 | 4191291 | 100 | 93 | 100 | 99 | 99 |
| 354 | ERR031676 | 1 | 027 | 1 | 4142516 | 100 | 93 | 99 | 99 | 100 |
| 358 | ERR031680 | 1 | 027 | 1 | 4136475 | 100 | 93 | 100 | 99 | 100 |
| 359 | ERR031681 | 1 | 027 | 1 | 4140699 | 100 | 93 | 100 | 99 | 100 |
| 363 | ERR031685 | 1 | 027 | 1 | 4146601 | 100 | 93 | 100 | 100 | 100 |
| 365 | ERR031687 | 1 | 027 | 1 | 4340904 | 100 | 93 | 100 | 99 | 100 |
| 366 | ERR031688 | 1 | 027 | 371 | 4167258 | 100 | 93 | 100 | 100 | 100 |
| 367 | ERR031689 | 1 | 027 | 1 | 4184988 | 100 | 93 | 99 | 99 | 100 |
| 375 | ERR031697 | 1 | 027 | 1 | 4141152 | 100 | 93 | 99 | 99 | 100 |
| 378 | ERR031700 | 1 | 027 | 1 | 4137740 | 100 | 93 | 100 | 99 | 100 |
| 381 | ERR031703 | 1 | 027 | 1 | 4141602 | 100 | 93 | 100 | 99 | 100 |
| 382 | ERR031704 | 1 | 027 | 1 | 4131191 | 100 | 93 | 99 | 99 | 99 |
| 383 | ERR031705 | 1 | 027 | 1 | 4163413 | 100 | 93 | 100 | 100 | 100 |
| 384 | ERR031706 | 1 | 027 | 1 | 4158134 | 100 | 93 | 100 | 100 | 100 |
| 386 | ERR031708 | 1 | 027 | 1 | 4168058 | 100 | 93 | 99 | 100 | 96 |
| 389 | ERR031711 | 1 | 027 | 1 | 4080095 | 100 | 93 | 100 | 100 | 99 |
| 391 | ERR034221 | 1 | 027 | 371 | 4125934 | 100 | 94 | 100 | 100 | 99 |
| 392 | ERR034222 | 1 | 027 | 1 | 4135167 | 100 | 94 | 100 | 99 | 100 |
| 393 | ERR034223 | 1 | 027 | 1 | 4137541 | 100 | 94 | 100 | 100 | 100 |
| 397 | ERR034227 | 1 | 027 | 1 | 4134548 | 100 | 93 | 100 | 100 | 100 |
| 399 | ERR044801 | 1 | 027 | 1 | 4179012 | 100 | 93 | 100 | 100 | 100 |
| 400 | ERR044802 | 1 | 027 | 1 | 4129950 | 100 | 27 | 100 | 24 | 34 |
| 402 | ERR044804 | 1 | 027 | 1 | 4151456 | 100 | 93 | 100 | 100 | 100 |
| 404 | ERR044806 | 1 | 027 | 1 | 4158511 | 100 | 33 | 100 | 26 | 0 |
| 406 | ERR044808 | 1 | 027 | 1 | 4130167 | 100 | 93 | 100 | 99 | 99 |
| 407 | ERR044809 | 1 | 027 | 1 | 4133138 | 100 | 93 | 100 | 100 | 99 |
| 410 | ERR044812 | 1 | 027 | 1 | 4132040 | 100 | 93 | 100 | 100 | 100 |
| 411 | ERR044813 | 1 | 027 | 1 | 4161597 | 100 | 93 | 100 | 100 | 100 |
| 413 | ERR044815 | 1 | 027 | 1 | 4166613 | 100 | 93 | 100 | 100 | 100 |
| 414 | ERR044816 | 1 | 027 | 1 | 4072734 | 100 | 93 | 100 | 100 | 100 |
| 415 | ERR044817 | 1 | 027 | 1 | 4213446 | 100 | 32 | 100 | 48 | 63 |
| 417 | ERR044819 | 1 | 027 | 1 | 4137335 | 100 | 93 | 100 | 100 | 100 |
| 418 | ERR044820 | 1 | 027 | 1 | 4131708 | 100 | 93 | 100 | 100 | 100 |
| 419 | ERR044821 | 1 | 027 | 1 | 4136128 | 100 | 93 | 99 | 100 | 100 |
| 420 | ERR044822 | 1 | 027 | 1 | 4134769 | 100 | 93 | 100 | 99 | 99 |
| 421 | ERR044823 | 1 | 027 | 1 | 4131687 | 100 | 93 | 100 | 100 | 100 |
| 422 | ERR044824 | 1 | 027 | 1 | 4133148 | 100 | 93 | 100 | 100 | 100 |
| 423 | ERR044825 | 1 | 027 | 1 | 4183010 | 100 | 93 | 100 | 100 | 100 |
| 424 | ERR044826 | 1 | 027 | 1 | 4158186 | 100 | 94 | 100 | 100 | 99 |
| 425 | ERR044827 | 1 | 027 | 1 | 4136246 | 100 | 94 | 100 | 100 | 100 |
| 426 | ERR044828 | 1 | 027 | 1 | 4130173 | 100 | 93 | 100 | 100 | 100 |
| 428 | ERR044830 | 1 | 027 | 1 | 4132607 | 100 | 94 | 100 | 99 | 100 |
| 430 | ERR044832 | 1 | 027 | 1 | 4127446 | 100 | 94 | 100 | 100 | 100 |
| 431 | ERR044833 | 1 | 027 | 1 | 4156904 | 100 | 94 | 100 | 100 | 100 |
| 432 | ERR044834 | 1 | 027 | 1 | 4128360 | 100 | 93 | 100 | 100 | 100 |
| 433 | ERR044835 | 1 | 027 | 1 | 4131621 | 100 | 94 | 100 | 100 | 100 |
| 434 | ERR044837 | 1 | 027 | 1 | 4119858 | 100 | 50 | 100 | 53 | 0 |
| 435 | ERR044838 | 1 | 027 | 1 | 4127971 | 100 | 94 | 100 | 100 | 100 |
| 436 | ERR044839 | 1 | 027 | 1 | 4130413 | 100 | 94 | 100 | 100 | 100 |
| 437 | ERR044840 | 1 | 027 | 1 | 4126610 | 100 | 94 | 100 | 100 | 100 |
| 438 | ERR044841 | 1 | 027 | 1 | 4132357 | 100 | 94 | 100 | 100 | 100 |
| 439 | ERR044842 | 1 | 027 | 1 | 4136055 | 100 | 94 | 100 | 100 | 100 |
| 440 | ERR044843 | 1 | 027 | 1 | 4134219 | 100 | 94 | 100 | 100 | 100 |
| 444 | ERR044848 | 1 | 027 | 1 | 4132182 | 100 | 94 | 100 | 100 | 100 |
| 450 | ERR178515 | 1 | 017 | 37 | 4260418 | 100 | 94 | 99 | 100 | 99 |
| 453 | ERR178518 | 1 | 017 | 37 | 4261384 | 100 | 94 | 100 | 99 | 100 |
| 454 | ERR178519 | 1 | 017 | 37 | 4267093 | 100 | 94 | 100 | 100 | 99 |
| 455 | ERR178520 | 1 | 017 | 37 | 4262147 | 100 | 94 | 99 | 100 | 100 |
| 461 | ERR178526 | 1 | 017 | 37 | 4287865 | 100 | 94 | 100 | 99 | 99 |
| 462 | ERR178527 | 1 | 017 | 37 | 4279329 | 100 | 94 | 99 | 99 | 99 |
| 464 | ERR178529 | 1 | 017 | 37 | 4266060 | 100 | 94 | 100 | 100 | 99 |
| 467 | ERR178532 | 1 | 017 | 37 | 4296224 | 100 | 94 | 100 | 99 | 100 |
| 468 | ERR178533 | 1 | 017 | 37 | 4227574 | 100 | 94 | 98 | 99 | 100 |
| 473 | ERR178538 | 1 | 017 | 37 | 4237199 | 100 | 94 | 100 | 100 | 99 |
| 475 | ERR178540 | 1 | 017 | 37 | 4276001 | 100 | 94 | 99 | 99 | 100 |
| 479 | ERR178544 | 1 | 017 | 37 | 4315080 | 100 | 94 | 100 | 99 | 99 |
| 482 | ERR178547 | 1 | 017 | 37 | 4251171 | 100 | 94 | 100 | 99 | 100 |
| 485 | ERR178550 | 1 | 017 | 37 | 4251068 | 100 | 94 | 100 | 99 | 100 |
| 487 | ERR178552 | 1 | 017 | 37 | 4190081 | 100 | 94 | 99 | 99 | 99 |
| 488 | ERR178553 | 1 | 017 | 37 | 4228599 | 100 | 94 | 100 | 99 | 100 |
| 491 | ERR178556 | 1 | 017 | 37 | 4174558 | 100 | 94 | 99 | 99 | 100 |
| 492 | ERR178557 | 1 | 017 | 37 | 4179492 | 100 | 94 | 99 | 100 | 99 |
| 495 | ERR178560 | 1 | 017 | 37 | 4212963 | 100 | 94 | 100 | 99 | 100 |
| 496 | ERR178561 | 1 | 017 | 37 | 4213300 | 100 | 94 | 99 | 99 | 99 |
| 497 | ERR178562 | 1 | 017 | 37 | 4239044 | 100 | 94 | 100 | 99 | 99 |
| 498 | ERR178563 | 1 | 017 | 37 | 4237830 | 100 | 94 | 100 | 99 | 99 |
| 501 | ERR178566 | 1 | 017 | 37 | 4240122 | 100 | 94 | 99 | 100 | 100 |
| 504 | ERR178569 | 1 | 017 | 37 | 4241610 | 100 | 94 | 100 | 99 | 100 |
| 510 | ERR178575 | 1 | 017 | 37 | 4244468 | 100 | 94 | 100 | 99 | 100 |
| 512 | ERR178578 | 1 | 017 | 37 | 4239145 | 100 | 94 | 100 | 100 | 100 |
| 514 | ERR178580 | 1 | 017 | 37 | 4210966 | 100 | 94 | 100 | 99 | 100 |
| 515 | ERR178581 | 1 | 017 | 37 | 4209826 | 100 | 94 | 99 | 99 | 100 |
| 523 | ERR178589 | 1 | 017 | 37 | 4282092 | 100 | 94 | 99 | 99 | 99 |
| 525 | ERR178591 | 1 | 017 | 37 | 4187469 | 100 | 94 | 100 | 99 | 100 |
| 526 | ERR178592 | 1 | 017 | 37 | 4201157 | 100 | 94 | 100 | 99 | 99 |
| 529 | ERR178595 | 1 | 017 | 37 | 4313129 | 100 | 94 | 99 | 100 | 99 |
| 530 | ERR178596 | 1 | 017 | 37 | 4243267 | 100 | 94 | 99 | 99 | 99 |
| 532 | ERR178598 | 1 | 017 | 37 | 4196851 | 100 | 94 | 100 | 99 | 99 |
| 536 | ERR178602 | 1 | 017 | 11 | 3999724 | 100 | 50 | 100 | 99 | 99 |
| 537 | ERR178603 | 1 | 017 | 37 | 4208077 | 100 | 94 | 99 | 99 | 99 |
| 540 | ERR340575 | 1 | | 37 | 4299669 | 100 | 94 | 100 | 100 | 100 |
| 541 | ERR340576 | 1 | | 37 | 4265068 | 100 | 94 | 100 | 99 | 99 |
| 542 | ERR340580 | 1 | | 37 | 4249078 | 100 | 94 | 100 | 99 | 100 |
| 543 | ERR340582 | 1 | | 37 | 4241167 | 100 | 94 | 100 | 100 | 100 |
| 544 | ERR340583 | 1 | | 37 | 4379577 | 100 | 94 | 100 | 99 | 99 |
| 547 | ERR340592 | 1 | 017 | 37 | 4197429 | 100 | 94 | 100 | 99 | 100 |
| 548 | ERR340594 | 1 | 017 | 37 | 4178404 | 100 | 94 | 99 | 99 | 100 |
| 549 | ERR340595 | 1 | 017 | 37 | 4277862 | 100 | 94 | 100 | 100 | 99 |
| 550 | ERR340597 | 1 | 017 | 37 | 4243641 | 100 | 94 | 99 | 100 | 100 |
| 551 | ERR340599 | 1 | 017 | 37 | 4235858 | 100 | 94 | 100 | 100 | 100 |
| 552 | ERR340600 | 1 | 017 | 37 | 4189295 | 100 | 94 | 100 | 100 | 99 |
| 553 | ERR340603 | 1 | 017 | 37 | 4238325 | 100 | 94 | 100 | 100 | 99 |
| 554 | ERR340604 | 1 | 017 | 37 | 4241453 | 100 | 94 | 99 | 99 | 100 |
| 555 | ERR340605 | 1 | 017 | 37 | 4255509 | 100 | 94 | 100 | 99 | 99 |
| 559 | ERR340613 | 1 | 017 | 37 | 4180115 | 100 | 94 | 100 | 100 | 100 |
| 560 | ERR340614 | 1 | 017 | 37 | 4204809 | 100 | 94 | 99 | 99 | 99 |
| 563 | ERR340618 | 1 | 017 | 37 | 4296440 | 100 | 94 | 100 | 99 | 99 |
| 564 | ERR340620 | 1 | 017 | 37 | 4273518 | 100 | 94 | 100 | 100 | 100 |
| 565 | ERR340621 | 1 | 017 | 37 | 4400777 | 100 | 94 | 100 | 99 | 100 |
| 566 | ERR340623 | 1 | 017 | 37 | 4297782 | 100 | 94 | 100 | 99 | 100 |
| 569 | ERR340629 | 1 | 017 | 2 | 4368300 | 100 | 94 | 100 | 99 | 100 |
| 570 | ERR340632 | 1 | 017 | 37 | 4239152 | 100 | 94 | 100 | 99 | 100 |
| 571 | ERR340633 | 1 | 017 | 37 | 4236790 | 100 | 94 | 100 | 99 | 100 |
| 572 | ERR340634 | 1 | 017 | 37 | 4410602 | 100 | 94 | 100 | 100 | 100 |
| 575 | ERR340637 | 1 | 017 | 37 | 4375553 | 100 | 94 | 100 | 100 | 100 |
| 576 | ERR340638 | 1 | 017 | 37 | 4374331 | 100 | 94 | 99 | 100 | 100 |
| 579 | ERR340641 | 1 | 017 | 37 | 4239912 | 100 | 25 | 100 | 13 | 0 |
| 583 | ERR340649 | 1 | | 37 | 4399675 | 100 | 94 | 100 | 99 | 100 |
| 584 | ERR340650 | 1 | | 37 | 4320308 | 100 | 94 | 100 | 100 | 99 |
| 585 | ERR340656 | 1 | | 37 | 4265155 | 100 | 94 | 100 | 99 | 100 |
| 586 | ERR340657 | 1 | | 37 | 4290469 | 100 | 94 | 100 | 100 | 100 |
| 588 | ERR340660 | 1 | | 37 | 4271416 | 100 | 94 | 100 | 99 | 100 |
| 589 | ERR340661 | 1 | | 37 | 4276528 | 100 | 94 | 100 | 100 | 99 |
| 590 | ERR340662 | 1 | | 37 | 4260413 | 100 | 94 | 99 | 100 | 100 |
| 591 | ERR351986 | 1 | 017 | 37 | 4785893 | 100 | 14 | 99 | 18 | 0 |
| 592 | ERR351988 | 1 | 017 | 37 | 4229615 | 100 | 94 | 100 | 100 | 99 |
| 593 | ERR387364 | 1 | | 37 | 4248858 | 100 | 94 | 100 | 100 | 99 |
| 595 | ERR387366 | 1 | 017 | 37 | 4245717 | 100 | 94 | 100 | 99 | 100 |
| 596 | ERR387368 | 1 | | 37 | 4237269 | 100 | 94 | 100 | 100 | 100 |
| 597 | ERR387369 | 1 | | 37 | 4547417 | 100 | 94 | 100 | 99 | 100 |
| 598 | ERR387370 | 1 | | 37 | 4282134 | 100 | 94 | 100 | 99 | 100 |
| 599 | ERR387371 | 1 | | 37 | 4290040 | 100 | 94 | 100 | 100 | 100 |
| 601 | ERR387374 | 1 | 017 | 37 | 4246900 | 100 | 94 | 99 | 100 | 99 |
| 602 | ERR387378 | 1 | | 37 | 5101419 | 100 | 94 | 100 | 99 | 100 |
| 603 | ERR387379 | 1 | | 37 | 4248337 | 100 | 94 | 100 | 99 | 99 |
| 604 | ERR387380 | 1 | | 37 | 4250000 | 100 | 94 | 99 | 99 | 100 |
| 605 | ERR387381 | 1 | 017 | 37 | 4249386 | 100 | 94 | 99 | 99 | 100 |
| 606 | ERR387382 | 1 | 017 | 37 | 4429475 | 100 | 94 | 100 | 100 | 99 |
| 607 | ERR387383 | 1 | | 37 | 4248479 | 100 | 94 | 100 | 100 | 99 |
| 608 | ERR387384 | 1 | | 37 | 4232981 | 100 | 94 | 99 | 99 | 99 |
| 611 | ERR387388 | 1 | | 37 | 4254840 | 100 | 94 | 100 | 99 | 100 |
| 613 | ERR387390 | 1 | 017 | 37 | 4232079 | 100 | 94 | 100 | 99 | 99 |
| 615 | ERR387395 | 1 | | 37 | 4308538 | 100 | 94 | 100 | 100 | 100 |
| 616 | ERR387396 | 1 | | 37 | 4260269 | 100 | 94 | 100 | 100 | 99 |
| 619 | ERR387400 | 1 | | 37 | 4236505 | 100 | 94 | 100 | 100 | 99 |
| 620 | ERR387401 | 1 | | 37 | 4212316 | 100 | 94 | 100 | 99 | 99 |
| 622 | ERR387404 | 1 | | 37 | 4201408 | 100 | 94 | 99 | 100 | 99 |
| 624 | ERR387406 | 1 | 017 | 37 | 4193422 | 100 | 94 | 100 | 99 | 100 |
| 625 | ERR387407 | 1 | | 37 | 4250349 | 100 | 94 | 100 | 99 | 100 |
| 626 | ERR387408 | 1 | | 37 | 4237541 | 100 | 94 | 99 | 99 | 100 |
| 627 | ERR387409 | 1 | | 37 | 4251080 | 100 | 94 | 100 | 99 | 99 |
| 628 | ERR387410 | 1 | | 37 | 4192289 | 100 | 94 | 100 | 100 | 100 |
| 629 | ERR387411 | 1 | | 37 | 4316430 | 100 | 94 | 100 | 99 | 99 |
| 630 | ERR387412 | 1 | 017 | 37 | 4208269 | 100 | 94 | 100 | 99 | 99 |
| 631 | ERR387413 | 1 | 017 | 37 | 4250791 | 100 | 94 | 100 | 99 | 100 |
| 632 | ERR387414 | 1 | 017 | 37 | 4277750 | 100 | 94 | 100 | 99 | 100 |
| 633 | ERR387415 | 1 | | 37 | 4233357 | 100 | 94 | 100 | 99 | 100 |
| 635 | ERR387417 | 1 | | 37 | 4290412 | 100 | 94 | 100 | 99 | 100 |
| 637 | ERR387419 | 1 | | 37 | 4319548 | 100 | 94 | 99 | 99 | 100 |
| 638 | ERR387420 | 1 | 017 | 37 | 4319690 | 100 | 94 | 100 | 99 | 99 |
| 639 | ERR387424 | 1 | | 37 | 4236505 | 100 | 94 | 99 | 100 | 100 |
| 640 | ERR387425 | 1 | | 37 | 5173918 | 100 | 94 | 100 | 99 | 100 |
| 644 | ERR387429 | 1 | 017 | 37 | 4199212 | 100 | 94 | 100 | 99 | 100 |
| 646 | ERR387431 | 1 | | 37 | 4234562 | 100 | 94 | 100 | 100 | 100 |
| 647 | ERR387432 | 1 | | 37 | 4232345 | 100 | 94 | 100 | 99 | 99 |
| 649 | ERR387436 | 1 | 017 | 37 | 4183279 | 100 | 14 | 100 | 0 | 0 |
| 650 | ERR387437 | 1 | 017 | 37 | 4240497 | 100 | 94 | 100 | 99 | 100 |
| 651 | ERR387438 | 1 | 017 | 37 | 4348112 | 100 | 94 | 100 | 100 | 100 |
| 652 | ERR387440 | 1 | | 37 | 4234438 | 100 | 94 | 100 | 99 | 99 |
| 653 | ERR387441 | 1 | | 37 | 4277426 | 100 | 94 | 100 | 99 | 99 |
| 654 | ERR387442 | 1 | | 37 | 4240421 | 100 | 94 | 100 | 99 | 99 |
| 655 | ERR387443 | 1 | | 37 | 4191566 | 100 | 94 | 100 | 100 | 99 |
| 657 | ERR387445 | 1 | 017 | 37 | 4192637 | 100 | 94 | 100 | 100 | 100 |
| 658 | ERR387447 | 1 | | 37 | 4247523 | 100 | 94 | 99 | 100 | 99 |

| | | | | | | | | | | |
|---|---|---|---|---|---|---|---|---|---|---|
| 659 | ERR387448 | 1 | | 37 | 4234942 | 100 | 94 | 100 | 100 | 100 |
| 660 | ERR387449 | 1 | | 37 | 4347630 | 100 | 94 | 100 | 99 | 100 |
| 661 | ERR387450 | 1 | | 37 | 4333337 | 100 | 94 | 100 | 100 | 99 |
| 663 | ERR387452 | 1 | 017 | 37 | 4317427 | 100 | 30 | 99 | 13 | 0 |
| 664 | ERR387455 | 1 | | 37 | 4236134 | 100 | 94 | 99 | 99 | 99 |
| 665 | ERR387457 | 1 | | | 4181200 | 100 | 94 | 100 | 99 | 99 |
| 666 | ERR387458 | 1 | | 37 | 4376296 | 100 | 94 | 99 | 100 | 99 |
| 669 | ERR788959 | 1 | | 37 | 3872058 | 100 | 94 | 100 | 99 | 99 |
| 670 | NAP08 | 1 | 078 | 11 | 4033133 | 100 | 45 | 100 | 95 | 100 |
| 671 | QCD-23m63 | 1 | 078 | NF | 3942185 | 100 | 45 | 100 | 95 | 100 |
| 672 | QCD-32g58 | 1 | 027 | 1 | 3995934 | 100 | 91 | 100 | 100 | 100 |
| 673 | QCD-37x79 | 1 | 027 | 1 | 4334388 | 100 | 91 | 100 | 100 | 100 |
| 674 | QCD-63q42 | 1 | 001 | 3 | 4446437 | 100 | 57 | 100 | 100 | 98 |
| 675 | QCD-66c26 | 1 | 027 | 1 | 4129150 | 100 | 91 | 100 | 100 | 100 |
| 676 | QCD-76w55 | 1 | 027 | 1 | 4399195 | 100 | 91 | 100 | 100 | 100 |
| 677 | QCD-97b34 | 1 | 027 | 1 | 4065010 | 100 | 91 | 100 | 100 | 100 |
| 678 | R20291 | 1 | 027 | 1 | 4191339 | 100 | 91 | 100 | 100 | 100 |
| 679 | VPI10463 | 1 | 087 | 46 | 4210279 | 100 | 94 | 100 | 100 | 100 |
| 681 | ERR339493 | 1 | | 1 | 4191943 | 100 | 93 | 100 | 100 | 100 |
| 683 | ERR339495 | 1 | | 1 | 4136583 | 100 | 95 | 100 | 100 | 99 |
| 687 | ERR339499 | 1 | | 17 | 4209582 | 100 | 60 | 99 | 100 | 99 |
| 688 | ERR339500 | 1 | | 17 | 4245577 | 100 | 62 | 100 | 100 | 99 |
| 689 | ERR339501 | 1 | | 2 | 4195065 | 100 | 92 | 100 | 100 | 100 |
| 690 | ERR339502 | 1 | | 1 | 4161154 | 100 | 93 | 100 | 99 | 100 |
| 691 | ERR339503 | 1 | | 12 | 4110855 | 100 | 94 | 100 | 99 | 99 |
| 692 | ERR339504 | 1 | | 1 | 4371813 | 100 | 94 | 99 | 100 | 100 |
| 697 | ERR339509 | 1 | | 44 | 4403978 | 100 | 62 | 99 | 100 | 99 |
| 704 | ERR339516 | 1 | | 12 | 4184494 | 100 | 94 | 99 | 99 | 99 |
| 705 | ERR339517 | 1 | | 1 | 4175269 | 100 | 93 | 99 | 99 | 99 |
| 708 | ERR339520 | 1 | | 1 | 4159307 | 100 | 93 | 100 | 99 | 98 |
| 718 | ERR339530 | 1 | | 1 | 4153667 | 100 | 93 | 99 | 99 | 99 |
| 721 | ERR339533 | 1 | | NF | 4122064 | 100 | 92 | 99 | 99 | 99 |
| 723 | ERR339535 | 1 | | 1 | 4156315 | 100 | 93 | 100 | 99 | 100 |
| 724 | ERR339536 | 1 | | 2 | 4198405 | 100 | 22 | 99 | 46 | 0 |
| 732 | ERR339544 | 1 | | 45 | 4243994 | 100 | 94 | 99 | 99 | 100 |
| 733 | ERR339545 | 1 | | 45 | 4241239 | 100 | 94 | 99 | 99 | 100 |
| 745 | ERR339557 | 1 | | NF | 4519035 | 100 | 92 | 99 | 100 | 100 |
| 747 | ERR339559 | 1 | | 9 | 4245320 | 100 | 13 | 99 | 0 | 0 |
| 748 | ERR339560 | 1 | | 33 | 4405517 | 100 | 18 | 99 | 13 | 0 |
| 751 | ERR339563 | 1 | | NF | 4333604 | 100 | 93 | 99 | 99 | 100 |
| 753 | ERR339565 | 1 | | 33 | 4339489 | 100 | 93 | 100 | 99 | 99 |
| 755 | ERR339567 | 1 | | 14 | 4226502 | 100 | 92 | 99 | 99 | 99 |
| 757 | ERR339569 | 1 | | 33 | 4301467 | 100 | 94 | 99 | 99 | 99 |
| 761 | ERR339573 | 1 | | NF | 4295381 | 100 | 14 | 100 | 0 | 0 |
| 766 | ERR339578 | 1 | | NF | 4195674 | 100 | 16 | 100 | 36 | 0 |
| 769 | ERR339581 | 1 | | 9 | 4307879 | 100 | 15 | 100 | 22 | 0 |
| 773 | ERR339585 | 1 | | 16 | 4490536 | 100 | 10 | 99 | 0 | 0 |
| 779 | ERR339591 | 1 | | 9 | 4358240 | 100 | 14 | 99 | 0 | 0 |
| 780 | ERR339592 | 1 | | 14 | 4266922 | 100 | 92 | 99 | 99 | 99 |
| 784 | ERR339596 | 1 | | 49 | 4324211 | 100 | 92 | 99 | 100 | 99 |
| 789 | ERR339601 | 1 | | 16 | 4546484 | 100 | 94 | 99 | 99 | 99 |
| 792 | ERR339604 | 1 | | 13 | 4285704 | 100 | 92 | 100 | 100 | 99 |
| 794 | ERR339606 | 1 | | 33 | 4298553 | 100 | 93 | 99 | 99 | 99 |
| 796 | ERR339608 | 1 | | NF | 4198987 | 100 | 93 | 100 | 100 | 98 |
| 798 | ERR339610 | 1 | | NF | 4246515 | 100 | 94 | 99 | 100 | 100 |
| 799 | ERR339611 | 1 | | 13 | 4241774 | 100 | 94 | 100 | 100 | 100 |
| 802 | ERR339614 | 1 | | 16 | 4535092 | 100 | 14 | 100 | 0 | 0 |
| 804 | ERR339616 | 1 | | 55 | 4095986 | 100 | 94 | 99 | 99 | 100 |
| 806 | ERR339618 | 1 | | 55 | 4170240 | 100 | 95 | 99 | 99 | 100 |
| 807 | ERR339619 | 1 | | 13 | 4356448 | 100 | 92 | 100 | 100 | 99 |
| 809 | ERR339621 | 1 | | 16 | 4371755 | 100 | 94 | 100 | 100 | 99 |
| 814 | ERR339626 | 1 | | 55 | 4099643 | 100 | 94 | 99 | 99 | 99 |
| 820 | ERR339632 | 1 | | 14 | 4272059 | 100 | 92 | 99 | 100 | 100 |
| 823 | ERR339635 | 1 | | 13 | 4259696 | 100 | 93 | 99 | 99 | 100 |
| 824 | ERR339636 | 1 | | 13 | 4292475 | 100 | 93 | 100 | 99 | 99 |
| 826 | ERR339638 | 1 | | 33 | 4143053 | 100 | 20 | 100 | 24 | 0 |
| 830 | ERR339642 | 1 | | 37 | 4217824 | 100 | 94 | 100 | 99 | 100 |
| 831 | ERR339643 | 1 | | 14 | 4201494 | 100 | 92 | 100 | 99 | 99 |
| 835 | ERR339647 | 1 | | 5 | 4198854 | 100 | 41 | 99 | 99 | 99 |
| 837 | ERR339649 | 1 | | 5 | 4353378 | 100 | 46 | 99 | 99 | 99 |
| 838 | ERR339650 | 1 | | 5 | 4523041 | 100 | 52 | 99 | 99 | 94 |
| 839 | ERR339651 | 1 | | 5 | 4109670 | 100 | 49 | 99 | 99 | 93 |
| 840 | ERR339652 | 1 | | 14 | 4257604 | 100 | 92 | 100 | 99 | 99 |
| 841 | ERR339653 | 1 | | 5 | 4167206 | 100 | 44 | 99 | 99 | 95 |
| 846 | ERR339658 | 1 | | 5 | 4214531 | 100 | 39 | 99 | 99 | 94 |
| 847 | ERR339659 | 1 | | 5 | 4215616 | 100 | 54 | 100 | 99 | 96 |
| 848 | ERR339660 | 1 | | 5 | 4384579 | 100 | 45 | 100 | 99 | 92 |
| 852 | ERR339664 | 1 | | 5 | 4206466 | 100 | 40 | 100 | 99 | 99 |
| 853 | ERR339665 | 1 | | 5 | 4207261 | 100 | 43 | 99 | 99 | 91 |
| 854 | ERR339666 | 1 | | 271 | 4211486 | 100 | 59 | 100 | 100 | 99 |
| 855 | ERR339667 | 1 | | 5 | 4083132 | 100 | 44 | 100 | 99 | 99 |
| 857 | ERR339669 | 1 | | 42 | 4159569 | 100 | 95 | 99 | 99 | 100 |
| 858 | ERR339670 | 1 | | 1 | 4148507 | 100 | 94 | 100 | 99 | 100 |
| 859 | ERR339671 | 1 | | 17 | 4232930 | 100 | 62 | 99 | 99 | 98 |
| 865 | ERR339677 | 1 | | 8 | 4226247 | 100 | 95 | 100 | 100 | 99 |
| 866 | ERR339678 | 1 | | 8 | 4281589 | 100 | 95 | 100 | 100 | 99 |
| 867 | ERR339679 | 1 | | 8 | 4233504 | 100 | 17 | 99 | 13 | 0 |
| 868 | ERR339680 | 1 | | 1 | 4168420 | 100 | 94 | 100 | 99 | 99 |
| 869 | ERR339681 | 1 | | 8 | 4133545 | 100 | 95 | 100 | 100 | 99 |
| 870 | ERR339682 | 1 | | 8 | 4270358 | 100 | 95 | 100 | 100 | 100 |
| 871 | ERR339683 | 1 | | 8 | 4186359 | 100 | 95 | 99 | 99 | 100 |
| 877 | ERR339689 | 1 | | 8 | 4294887 | 100 | 95 | 99 | 99 | 99 |
| 883 | ERR339695 | 1 | | 8 | 4314839 | 100 | 95 | 100 | 100 | 99 |
| 884 | ERR339696 | 1 | | 8 | 4320688 | 100 | 95 | 100 | 100 | 99 |
| 891 | ERR339703 | 1 | | 8 | 4064376 | 100 | 19 | 100 | 0 | 0 |
| 897 | ERR339709 | 1 | | 1 | 4180140 | 100 | 94 | 99 | 99 | 99 |
| 905 | ERR339717 | 1 | | 8 | 4280256 | 100 | 95 | 99 | 99 | 99 |
| 907 | ERR339719 | 1 | | 8 | 4314514 | 100 | 95 | 99 | 99 | 100 |
| 911 | ERR339723 | 1 | | 8 | 4304879 | 100 | 95 | 100 | 99 | 100 |
| 912 | ERR339724 | 1 | | 8 | 4121855 | 100 | 95 | 100 | 99 | 99 |
| 913 | ERR339725 | 1 | | 5 | 4293404 | 100 | 41 | 100 | 99 | 99 |
| 914 | ERR339726 | 1 | | 5 | 4161966 | 100 | 41 | 99 | 99 | 95 |
| 915 | ERR339727 | 1 | | 5 | 4190941 | 100 | 38 | 99 | 99 | 99 |
| 917 | ERR339729 | 1 | | 5 | 4244347 | 100 | 38 | 99 | 99 | 100 |
| 918 | ERR339730 | 1 | | 5 | 4339783 | 100 | 39 | 99 | 99 | 91 |
| 920 | ERR339732 | 1 | | 5 | 4168002 | 100 | 39 | 99 | 99 | 97 |
| 921 | ERR339733 | 1 | | 5 | 4856161 | 100 | 56 | 100 | 100 | 99 |
| 924 | ERR339736 | 1 | | 5 | 4337415 | 100 | 40 | 99 | 99 | 97 |
| 925 | ERR339737 | 1 | | 5 | 4022062 | 100 | 45 | 99 | 99 | 94 |
| 926 | ERR339738 | 1 | | 5 | 4248656 | 100 | 38 | 99 | 99 | 92 |
| 927 | ERR339739 | 1 | | 5 | 4340893 | 100 | 39 | 100 | 99 | 99 |
| 928 | ERR339740 | 1 | | 5 | 4322280 | 100 | 41 | 99 | 99 | 99 |
| 929 | ERR339741 | 1 | | 5 | 4243729 | 100 | 40 | 100 | 99 | 98 |
| 931 | ERR339743 | 1 | | 5 | 4451588 | 100 | 37 | 100 | 98 | 93 |
| 935 | ERR339747 | 1 | | 5 | 4236650 | 100 | 41 | 99 | 97 | 95 |
| 939 | ERR339751 | 1 | | 1 | 4144491 | 100 | 93 | 99 | 99 | 99 |
| 941 | ERR339753 | 1 | | 1 | 4163892 | 100 | 94 | 100 | 99 | 100 |
| 942 | ERR339754 | 1 | | 1 | 4149274 | 100 | 93 | 99 | 99 | 99 |
| 945 | ERR339757 | 1 | | 1 | 4142070 | 100 | 94 | 100 | 100 | 100 |
| 946 | ERR339758 | 1 | | 1 | 4148462 | 100 | 93 | 99 | 100 | 99 |
| 949 | ERR339761 | 1 | | 15 | 4172996 | 100 | 16 | 100 | 13 | 0 |
| 950 | ERR339762 | 1 | | 1 | 4143839 | 100 | 93 | 99 | 99 | 100 |
| 952 | ERR339764 | 1 | | 1 | 4157047 | 100 | 94 | 100 | 99 | 100 |
| 955 | ERR339767 | 1 | | 1 | 4210395 | 100 | 94 | 99 | 99 | 99 |
| 959 | ERR339771 | 1 | | 18 | 4237157 | 100 | 14 | 99 | 13 | 0 |
| 960 | ERR339772 | 1 | | 1 | 4164419 | 100 | 94 | 99 | 99 | 100 |
| 961 | ERR339773 | 1 | | 1 | 4158279 | 100 | 94 | 99 | 100 | 100 |
| 966 | ERR339778 | 1 | | 46 | 4962599 | 100 | 94 | 100 | 99 | 99 |
| 967 | ERR339779 | 1 | | 1 | 4180493 | 100 | 93 | 99 | 100 | 99 |
| 968 | ERR339780 | 1 | | 1 | 4149663 | 100 | 94 | 100 | 99 | 99 |
| 977 | ERR339789 | 1 | | 8 | 4337167 | 100 | 95 | 99 | 99 | 100 |
| 980 | ERR339792 | 1 | | 1 | 4142019 | 100 | 94 | 99 | 99 | 100 |
| 981 | ERR339793 | 1 | | 1 | 4145691 | 100 | 93 | 99 | 100 | 100 |
| 982 | ERR339794 | 1 | | 1 | 4152334 | 100 | 94 | 100 | 99 | 100 |
| 984 | ERR339796 | 1 | | 8 | 4317005 | 100 | 95 | 99 | 99 | 100 |
| 985 | ERR339797 | 1 | | 1 | 4154220 | 100 | 93 | 100 | 99 | 99 |
| 986 | ERR339798 | 1 | | 1 | 4180702 | 100 | 94 | 100 | 100 | 99 |
| 987 | ERR339799 | 1 | | 8 | 4221022 | 100 | 95 | 100 | 99 | 99 |
| 989 | ERR339801 | 1 | | 2 | 5212921 | 100 | 15 | 100 | 6 | 45 |
| 991 | ERR339803 | 1 | | NF | 4395005 | 100 | 92 | 99 | 99 | 100 |
| 995 | ERR339807 | 1 | | 45 | 4585654 | 100 | 94 | 100 | 100 | 100 |
| 998 | ERR339810 | 1 | | 55 | 4041597 | 100 | 94 | 100 | 99 | 99 |
| 999 | ERR339811 | 1 | | 55 | 4048356 | 100 | 94 | 99 | 99 | 100 |
| 1002 | ERR339814 | 1 | | 49 | 4176314 | 100 | 93 | 99 | 99 | 99 |
| 1003 | ERR339815 | 1 | | NF | 4216931 | 100 | 92 | 100 | 99 | 100 |
| 1004 | ERR339816 | 1 | | 55 | 4051919 | 100 | 94 | 100 | 100 | 100 |
| 1006 | ERR339818 | 1 | | 55 | 4132558 | 100 | 95 | 99 | 99 | 100 |
| 1009 | ERR339821 | 1 | | 55 | 4078339 | 100 | 95 | 99 | 99 | 99 |
| 1012 | ERR339824 | 1 | | 55 | 4121729 | 100 | 95 | 99 | 100 | 99 |
| 1013 | ERR339825 | 1 | | 37 | 4200711 | 100 | 94 | 99 | 99 | 99 |
| 1014 | ERR339826 | 1 | | 1 | 4145683 | 100 | 94 | 100 | 99 | 100 |
| 1016 | ERR339828 | 1 | | 1 | 4150240 | 100 | 93 | 100 | 100 | 99 |
| 1017 | ERR339829 | 1 | | 1 | 4232248 | 100 | 93 | 100 | 99 | 99 |
| 1018 | ERR339830 | 1 | | 1 | 4121068 | 100 | 94 | 100 | 99 | 100 |
| 1022 | ERR339834 | 1 | | 1 | 4145042 | 100 | 94 | 100 | 99 | 100 |
| 1024 | ERR339836 | 1 | | 1 | 4157283 | 100 | 94 | 100 | 99 | 100 |
| 1026 | ERR339838 | 1 | | 1 | 4212200 | 100 | 93 | 100 | 100 | 100 |
| 1027 | ERR339839 | 1 | | 1 | 4157769 | 100 | 93 | 99 | 99 | 100 |
| 1030 | ERR339842 | 1 | | 1 | 4146387 | 100 | 94 | 100 | 100 | 99 |
| 1033 | ERR339845 | 1 | | 1 | 4163033 | 100 | 94 | 99 | 99 | 99 |
| 1035 | ERR339847 | 1 | | 1 | 4166805 | 100 | 94 | 100 | 99 | 100 |
| 1038 | ERR339850 | 1 | | 1 | 4143175 | 100 | 93 | 100 | 100 | 99 |
| 1039 | ERR339851 | 1 | | 1 | 4324950 | 100 | 94 | 100 | 100 | 98 |
| 1040 | ERR339852 | 1 | | 1 | 4145953 | 100 | 93 | 100 | 99 | 99 |
| 1043 | ERR339855 | 1 | | 1 | 4147598 | 100 | 93 | 100 | 99 | 99 |
| 1044 | ERR339856 | 1 | | 1 | 4142110 | 100 | 93 | 100 | 99 | 100 |
| 1046 | ERR339858 | 1 | | 1 | 4141461 | 100 | 93 | 100 | 99 | 100 |
| 1047 | ERR339859 | 1 | | 1 | 4188580 | 100 | 93 | 99 | 99 | 100 |
| 1048 | ERR339860 | 1 | | 1 | 4152511 | 100 | 93 | 100 | 99 | 100 |
| 1050 | ERR339862 | 1 | | 1 | 4146795 | 100 | 94 | 99 | 99 | 99 |
| 1055 | ERR339868 | 1 | | 1 | 4185844 | 100 | 94 | 100 | 99 | 100 |
| 1065 | ERR339878 | 1 | | 1 | 4161407 | 100 | 93 | 99 | 99 | 100 |
| 1066 | ERR339879 | 1 | | 1 | 4146660 | 100 | 93 | 100 | 99 | 100 |
| 1067 | ERR339880 | 1 | | 1 | 4163924 | 100 | 93 | 99 | 100 | 100 |
| 1068 | ERR339881 | 1 | | 1 | 4145365 | 100 | 94 | 99 | 99 | 99 |
| 1069 | ERR339882 | 1 | | 1 | 4154893 | 100 | 93 | 99 | 100 | 99 |
| 1070 | ERR339883 | 1 | | 1 | 4143472 | 100 | 94 | 100 | 100 | 99 |
| 1074 | ERR339887 | 1 | | 1 | 4244124 | 100 | 93 | 100 | 99 | 99 |
| 1077 | ERR339890 | 1 | | 1 | 4146109 | 100 | 94 | 100 | 99 | 100 |
| 1081 | ERR339894 | 1 | | 1 | 4155883 | 100 | 93 | 99 | 99 | 100 |
| 1083 | ERR339896 | 1 | | 1 | 4146671 | 100 | 93 | 99 | 99 | 100 |
| 1087 | ERR339900 | 1 | | 1 | 4187051 | 100 | 94 | 99 | 99 | 100 |
| 1088 | ERR339901 | 1 | | 1 | 4143677 | 100 | 95 | 100 | 100 | 99 |
| 1091 | ERR339904 | 1 | | 1 | 4156885 | 100 | 94 | 100 | 99 | 100 |
| 1096 | ERR339909 | 1 | | 1 | 4143330 | 100 | 93 | 99 | 99 | 100 |
| 1098 | ERR339911 | 1 | | 1 | 4147733 | 100 | 93 | 100 | 99 | 100 |
| 1099 | ERR339912 | 1 | | 1 | 4150270 | 100 | 93 | 100 | 99 | 99 |

| | | | | | | | | | | |
|---|---|---|---|---|---|---|---|---|---|---|
| 1100 | ERR339913 | 1 | | 1 | 4148634 | 100 | 93 | 100 | 99 | 99 |
| 1101 | ERR339914 | 1 | | 1 | 4148636 | 100 | 94 | 100 | 100 | 100 |
| 1102 | ERR339915 | 1 | | 1 | 4149567 | 100 | 94 | 100 | 99 | 100 |
| 1103 | ERR339916 | 1 | | 1 | 4143428 | 100 | 93 | 99 | 99 | 99 |
| 1104 | ERR339917 | 1 | | 1 | 4143017 | 100 | 93 | 99 | 99 | 99 |
| 1106 | ERR339919 | 1 | | 1 | 4144052 | 100 | 93 | 100 | 99 | 100 |
| 1108 | ERR339921 | 1 | | 1 | 4149765 | 100 | 94 | 100 | 100 | 99 |
| 1111 | ERR339924 | 1 | | 1 | 4148467 | 100 | 93 | 100 | 100 | 99 |
| 1112 | ERR339925 | 1 | | 1 | 4166406 | 100 | 94 | 99 | 99 | 99 |
| 1114 | ERR339927 | 1 | | 1 | 4145139 | 100 | 93 | 100 | 99 | 100 |
| 1116 | ERR339929 | 1 | | 1 | 4139614 | 100 | 93 | 100 | 99 | 99 |
| 1117 | ERR339930 | 1 | | 1 | 4158688 | 100 | 93 | 100 | 100 | 100 |
| 1119 | ERR339932 | 1 | | 1 | 4158771 | 100 | 93 | 100 | 99 | 100 |
| 1122 | ERR339935 | 1 | | 1 | 4144464 | 100 | 94 | 100 | 99 | 99 |
| 1124 | ERR339937 | 1 | | 1 | 4158892 | 100 | 94 | 99 | 100 | 100 |
| 1125 | ERR339938 | 1 | | 1 | 4157185 | 100 | 93 | 99 | 99 | 99 |
| 1129 | ERR339943 | 1 | | 1 | 4151187 | 100 | 93 | 100 | 100 | 99 |
| 1130 | ERR339944 | 1 | | 1 | 4146647 | 100 | 93 | 100 | 99 | 99 |
| 1132 | ERR339946 | 1 | | 1 | 4332389 | 100 | 94 | 99 | 99 | 100 |
| 1139 | ERR339953 | 1 | | 1 | 4152133 | 100 | 94 | 99 | 99 | 99 |
| 1143 | ERR339957 | 1 | | 1 | 4161610 | 100 | 94 | 100 | 99 | 100 |
| 1145 | ERR339959 | 1 | | NF | 4338019 | 100 | 94 | 100 | 99 | 100 |
| 1147 | ERR339961 | 1 | | 8 | 4175781 | 100 | 95 | 100 | 99 | 100 |
| 1152 | ERR339966 | 1 | | 2 | 4172083 | 100 | 92 | 100 | 99 | 99 |
| 1157 | ERR339971 | 1 | | 8 | 4236298 | 100 | 95 | 99 | 99 | 99 |
| 1158 | ERR339972 | 1 | | 8 | 4080175 | 100 | 95 | 99 | 99 | 99 |
| 1160 | ERR339974 | 1 | | 2 | 4190937 | 100 | 18 | 100 | 0 | 0 |
| 1162 | ERR339976 | 1 | | 4 | 4098201 | 100 | 95 | 100 | 100 | 100 |
| 1165 | ERR339979 | 1 | | 59 | 4204614 | 100 | 95 | 100 | 99 | 100 |
| 1167 | ERR339981 | 1 | | 8 | 4220798 | 100 | 95 | 100 | 99 | 100 |
| 1168 | ERR339982 | 1 | | 2 | 4142129 | 100 | 22 | 100 | 41 | 0 |
| 1173 | ERR339987 | 1 | | 8 | 4183618 | 100 | 95 | 100 | 100 | 99 |
| 1174 | ERR339988 | 1 | | 8 | 4338750 | 100 | 95 | 100 | 99 | 99 |
| 1176 | ERR339990 | 1 | | 43 | 4213944 | 100 | 27 | 100 | 40 | 0 |
| 1185 | ERR339999 | 1 | | 2 | 4212163 | 100 | 15 | 99 | 0 | 0 |
| 1191 | ERR340005 | 1 | | 2 | 4276952 | 100 | 63 | 100 | 84 | 45 |
| 1194 | ERR340008 | 1 | | 2 | 4136655 | 100 | 17 | 99 | 13 | 0 |
| 1195 | ERR340009 | 1 | | 2 | 4131084 | 100 | 92 | 100 | 100 | 99 |
| 1196 | ERR340010 | 1 | | 2 | 4148518 | 100 | 15 | 99 | 0 | 0 |
| 1199 | ERR340013 | 1 | | 43 | 4219037 | 100 | 27 | 100 | 13 | 0 |
| 1201 | ERR340015 | 1 | | 75 | 4352043 | 100 | 20 | 100 | 0 | 0 |
| 1202 | ERR340016 | 1 | | 18 | 4426618 | 100 | 16 | 100 | 0 | 0 |
| 1204 | ERR340018 | 1 | | 2 | 4194777 | 100 | 20 | 99 | 14 | 0 |
| 1206 | ERR340021 | 1 | | 53 | 4068733 | 100 | 93 | 100 | 99 | 99 |
| 1207 | ERR340022 | 1 | | 2 | 4158761 | 100 | 92 | 100 | 99 | 99 |
| 1210 | ERR340025 | 1 | | 56 | 4046661 | 100 | 10 | 99 | 0 | 0 |
| 1215 | ERR340030 | 1 | | 8 | 4305628 | 100 | 91 | 100 | 99 | 100 |
| 1217 | ERR340032 | 1 | | 53 | 4079185 | 100 | 93 | 100 | 99 | 100 |
| 1220 | ERR340035 | 1 | | 8 | 4267009 | 100 | 95 | 99 | 99 | 100 |
| 1221 | ERR340036 | 1 | | 8 | 4059394 | 100 | 95 | 99 | 99 | 99 |
| 1222 | ERR340037 | 1 | | 8 | 4253863 | 100 | 95 | 99 | 100 | 99 |
| 1225 | ERR340040 | 1 | | 2 | 4215964 | 100 | 43 | 99 | 50 | 0 |
| 1226 | ERR340041 | 1 | | 18 | 4212977 | 100 | 17 | 99 | 0 | 0 |
| 1228 | ERR340043 | 1 | | 43 | 4446018 | 100 | 17 | 100 | 0 | 0 |
| 1233 | ERR340048 | 1 | | 8 | 4066487 | 100 | 92 | 99 | 99 | 99 |
| 1234 | ERR340049 | 1 | | 2 | 4175676 | 100 | 94 | 100 | 100 | 99 |
| 1237 | ERR340052 | 1 | | 8 | 4310372 | 100 | 95 | 100 | 99 | 100 |
| 1238 | ERR340053 | 1 | | 8 | 4224487 | 100 | 95 | 100 | 100 | 99 |
| 1242 | ERR340057 | 1 | | 2 | 4203375 | 100 | 27 | 99 | 29 | 0 |
| 1244 | ERR340059 | 1 | | 2 | 4167088 | 100 | 93 | 100 | 99 | 100 |
| 1246 | ERR340061 | 1 | | 8 | 4271765 | 100 | 95 | 99 | 100 | 100 |
| 1287 | ERR340102 | 1 | | 19 | 4125626 | 100 | 8 | 100 | 0 | 0 |
| 1296 | ERR340111 | 1 | | 56 | 4095855 | 100 | 12 | 100 | 0 | 0 |
| 1304 | ERR340119 | 1 | | 37 | 4173309 | 100 | 94 | 100 | 99 | 100 |
| 1305 | ERR340120 | 1 | | 26 | 4286887 | 100 | 95 | 99 | 100 | 100 |
| 1307 | ERR340122 | 1 | | 5 | 4224504 | 100 | 44 | 99 | 99 | 99 |
| 1308 | ERR340123 | 1 | | 1 | 4177449 | 100 | 95 | 100 | 99 | 99 |
| 1309 | ERR340124 | 1 | | 36 | 4391772 | 100 | 94 | 98 | 99 | 99 |
| 1313 | ERR340128 | 1 | | 9 | 5024736 | 100 | 29 | 100 | 53 | 0 |
| 1315 | ERR340130 | 1 | | 5 | 4292400 | 100 | 46 | 100 | 99 | 99 |
| 1316 | ERR340131 | 1 | | 1 | 4140712 | 100 | 94 | 100 | 100 | 100 |
| 1317 | ERR340132 | 1 | | 5 | 4236767 | 100 | 39 | 100 | 99 | 91 |
| 1321 | ERR340136 | 1 | | 35 | 4640186 | 100 | 92 | 100 | 99 | 96 |
| 1322 | ERR340137 | 1 | | 3 | 4363206 | 100 | 61 | 100 | 100 | 90 |
| 1342 | ERR340157 | 1 | | 2 | 4137870 | 100 | 17 | 100 | 13 | 0 |
| 1344 | ERR340159 | 1 | | 11 | 4052976 | 100 | 55 | 100 | 100 | 100 |
| 1349 | ERR340164 | 1 | | 5 | 4318409 | 100 | 50 | 99 | 99 | 100 |
| 1354 | ERR340169 | 1 | | 8 | 4248028 | 100 | 95 | 99 | 100 | 100 |
| 1357 | ERR340172 | 1 | | 42 | 4100893 | 100 | 95 | 100 | 99 | 100 |
| 1366 | ERR340181 | 1 | | 1 | 4141885 | 100 | 94 | 100 | 99 | 99 |
| 1367 | ERR340182 | 1 | | 44 | 4354796 | 100 | 67 | 99 | 99 | 87 |
| 1368 | ERR340183 | 1 | | 8 | 4245971 | 100 | 95 | 100 | 100 | 100 |
| 1369 | ERR340184 | 1 | | 17 | 4327773 | 100 | 61 | 99 | 99 | 99 |
| 1370 | ERR340185 | 1 | | 5 | 4328602 | 100 | 54 | 100 | 99 | 99 |
| 1371 | ERR340186 | 1 | | 49 | 4331826 | 100 | 92 | 100 | 99 | 99 |
| 1372 | ERR340187 | 1 | | 14 | 4220313 | 100 | 92 | 100 | 99 | 100 |
| 1373 | ERR340188 | 1 | | 16 | 4522471 | 100 | 95 | 99 | 100 | 99 |
| 1374 | ERR340189 | 1 | | 26 | 4645526 | 100 | 95 | 100 | 99 | 99 |
| 1376 | ERR340191 | 1 | | 5 | 4324197 | 100 | 53 | 99 | 99 | 94 |
| 1378 | ERR340193 | 1 | | 18 | 4276882 | 100 | 21 | 99 | 52 | 0 |
| 1383 | ERR340198 | 1 | | 8 | 4233942 | 100 | 95 | 99 | 100 | 99 |
| 1384 | ERR340199 | 1 | | 11 | 3905600 | 100 | 56 | 99 | 99 | 99 |
| 1386 | ERR340201 | 1 | | 8 | 4369841 | 100 | 95 | 100 | 100 | 100 |
| 1389 | ERR340204 | 1 | | 56 | 4050365 | 100 | 16 | 100 | 0 | 0 |
| 1394 | ERR340209 | 1 | | 11 | 3966844 | 100 | 47 | 100 | 99 | 100 |
| 1405 | ERR340220 | 1 | | 5 | 4249856 | 100 | 41 | 99 | 99 | 93 |
| 1406 | ERR340221 | 1 | | 11 | 4012027 | 100 | 46 | 99 | 99 | 100 |
| 1407 | ERR340222 | 1 | | 5 | 4117166 | 100 | 41 | 100 | 99 | 97 |
| 1408 | ERR340223 | 1 | | 11 | 4017280 | 100 | 48 | 100 | 100 | 100 |
| 1409 | ERR340224 | 1 | | 8 | 4303661 | 100 | 95 | 100 | 99 | 100 |
| 1413 | ERR340228 | 1 | | 5 | 4287008 | 100 | 39 | 100 | 99 | 99 |
| 1416 | ERR340231 | 1 | | 113 | 4386514 | 100 | 18 | 100 | 13 | 0 |
| 1418 | ERR340233 | 1 | | 35 | 4297698 | 100 | 16 | 99 | 0 | 0 |
| 1419 | ERR340234 | 1 | | 2 | 4164202 | 100 | 93 | 100 | 99 | 99 |
| 1421 | ERR340236 | 1 | | 11 | 4051055 | 100 | 46 | 100 | 99 | 99 |
| 1425 | ERR340240 | 1 | | 8 | 4279487 | 100 | 95 | 99 | 99 | 99 |
| 1426 | ERR340241 | 1 | | 2 | 4146163 | 100 | 92 | 100 | 100 | 99 |
| 1429 | ERR340244 | 1 | | 11 | 3965079 | 100 | 50 | 100 | 99 | 99 |
| 1433 | ERR340248 | 1 | | 111 | 4393894 | 100 | 13 | 100 | 0 | 0 |
| 1435 | ERR340250 | 1 | | 1 | 4357162 | 100 | 94 | 99 | 99 | 100 |
| 1437 | ERR340252 | 1 | | 17 | 4381870 | 100 | 95 | 100 | 99 | 99 |
| 1440 | ERR340255 | 1 | | 26 | 4269419 | 100 | 95 | 100 | 99 | 100 |
| 1452 | ERR340267 | 1 | | 8 | 4119079 | 100 | 95 | 100 | 100 | 100 |
| 1457 | ERR340272 | 1 | | 14 | 4301802 | 100 | 92 | 99 | 100 | 100 |
| 1460 | ERR340275 | 1 | | 42 | 4071511 | 100 | 95 | 99 | 99 | 100 |
| 1463 | ERR340278 | 1 | | 5 | 4249102 | 100 | 37 | 100 | 99 | 93 |
| 1468 | ERR340283 | 1 | | 13 | 4270831 | 100 | 92 | 99 | 99 | 100 |
| 1470 | ERR340285 | 1 | | 18 | 4249862 | 100 | 17 | 100 | 0 | 0 |
| 1474 | ERR340289 | 1 | | 49 | 4387752 | 100 | 92 | 99 | 100 | 100 |
| 1478 | ERR340293 | 1 | | 1 | 4362556 | 100 | 94 | 100 | 99 | 99 |
| 1481 | ERR340296 | 1 | | 16 | 4491164 | 100 | 95 | 99 | 100 | 100 |
| 1484 | ERR340299 | 1 | | 42 | 4081241 | 100 | 95 | 100 | 100 | 100 |
| 1485 | ERR340300 | 1 | | 3 | 4203895 | 100 | 63 | 100 | 100 | 99 |
| 1486 | ERR340301 | 1 | | 15 | 4180167 | 100 | 94 | 99 | 100 | 99 |
| 1491 | ERR340306 | 1 | | 2 | 4060926 | 100 | 92 | 99 | 100 | 99 |
| 1494 | ERR340309 | 1 | | 2 | 4172310 | 100 | 42 | 99 | 71 | 45 |
| 1497 | ERR340312 | 1 | | 2 | 4112319 | 100 | 21 | 100 | 19 | 0 |
| 1500 | ERR340315 | 1 | | 2 | 4211256 | 100 | 93 | 99 | 99 | 100 |
| 1501 | ERR340316 | 1 | | 48 | 4172177 | 100 | 94 | 100 | 99 | 100 |
| 1504 | ERR126241 | 1 | | 44 | 4572355 | 100 | 24 | 100 | 13 | 0 |
| 1508 | ERR126245 | 1 | 017 | 37 | 4173054 | 100 | 94 | 100 | 100 | 100 |
| 1511 | ERR126248 | 1 | | 2 | 4052720 | 100 | 92 | 100 | 99 | 100 |
| 1513 | ERR126250 | 1 | 018 | 17 | 4236341 | 100 | 64 | 100 | 100 | 99 |
| 1514 | ERR126251 | 1 | 129 | 13 | 4306949 | 100 | 94 | 100 | 99 | 100 |
| 1515 | ERR126252 | 1 | 014 | 14 | 4193646 | 100 | 93 | 100 | 100 | 100 |
| 1516 | ERR126253 | 1 | 106 | 42 | 4138193 | 100 | 95 | 100 | 99 | 99 |
| 1518 | ERR126255 | 1 | | 3 | 4239231 | 100 | 63 | 100 | 100 | 98 |
| 1519 | ERR126256 | 1 | 011 | 77 | 4243949 | 100 | 94 | 99 | 100 | 100 |
| 1520 | ERR126257 | 1 | 106 | 41 | 4267557 | 100 | 95 | 100 | 100 | 100 |
| 1521 | ERR126258 | 1 | 015 | 44 | 4521335 | 100 | 94 | 100 | 99 | 100 |
| 1522 | ERR126259 | 1 | 056 | 58 | 4146757 | 100 | 16 | 99 | 13 | 0 |
| 1524 | ERR126261 | 1 | | 54 | 4272012 | 100 | 100 | 99 | 100 | 100 |
| 1525 | ERR126262 | 1 | | 54 | 4422900 | 100 | 99 | 99 | 99 | 100 |
| 1527 | ERR126264 | 1 | | 7 | 4139724 | 100 | 14 | 100 | 0 | 0 |
| 1528 | ERR126265 | 1 | | 12 | 4147710 | 100 | 94 | 100 | 100 | 99 |
| 1529 | ERR126266 | 1 | | 5 | 4249563 | 100 | 37 | 100 | 98 | 99 |
| 1531 | ERR126268 | 1 | 023 | 22 | 4252826 | 100 | 39 | 100 | 99 | 90 |
| 1536 | ERR126273 | 1 | 321 | 41 | 4214840 | 100 | 95 | 99 | 100 | 100 |
| 1537 | ERR126274 | 1 | | 11 | 4033972 | 100 | 13 | 100 | 0 | 0 |
| 1540 | ERR126277 | 1 | 066 | 11 | 4114491 | 100 | 59 | 100 | 100 | 99 |
| 1543 | ERR126280 | 1 | | 11 | 5306781 | 100 | 92 | 99 | 100 | 99 |
| 1544 | ERR126281 | 1 | 019 | 67 | 4142092 | 100 | 94 | 100 | 99 | 100 |
| 1549 | ERR126286 | 1 | 069 | 5 | 4048009 | 100 | 43 | 100 | 99 | 99 |
| 1551 | ERR126288 | 1 | 454 | 45 | 4519853 | 100 | 18 | 100 | 26 | 45 |
| 1559 | ERR126296 | 1 | | 16 | 4559109 | 100 | 29 | 99 | 53 | 62 |
| 1561 | ERR142052 | 1 | | 2 | 4361618 | 100 | 19 | 100 | 10 | 0 |
| 1572 | ERR142063 | 1 | | 42 | 4170453 | 100 | 95 | 100 | 100 | 99 |
| 1577 | ERR142068 | 1 | | 42 | 4172134 | 100 | 97 | 100 | 100 | 100 |
| 1584 | ERR142075 | 1 | | 3 | 4387527 | 100 | 62 | 100 | 100 | 99 |
| 1588 | ERR142079 | 1 | | 1 | 4135289 | 100 | 92 | 100 | 99 | 99 |
| 1589 | ERR142080 | 1 | | 1 | 4135660 | 100 | 93 | 99 | 100 | 99 |
| 1590 | ERR142081 | 1 | | 1 | 4147299 | 100 | 93 | 100 | 99 | 100 |
| 1591 | ERR142082 | 1 | | 1 | 4231280 | 100 | 93 | 99 | 100 | 99 |
| 1593 | ERR142084 | 1 | | 1 | 4142715 | 100 | 93 | 100 | 100 | 100 |
| 1594 | ERR142085 | 1 | | 5 | 4332653 | 100 | 40 | 100 | 100 | 97 |
| 1595 | ERR142086 | 1 | | 1 | 4204041 | 100 | 92 | 99 | 100 | 99 |
| 1596 | ERR142087 | 1 | | 1 | 4261196 | 100 | 95 | 100 | 100 | 100 |
| 1599 | ERR142090 | 1 | | 89 | 4375665 | 100 | 30 | 99 | 17 | 23 |
| 1600 | ERR142091 | 1 | | 1 | 4218771 | 100 | 93 | 99 | 100 | 100 |
| 1604 | ERR142095 | 1 | | 1 | 4189405 | 100 | 92 | 99 | 99 | 99 |
| 1605 | ERR142096 | 1 | | 1 | 4205635 | 100 | 93 | 99 | 100 | 100 |
| 1606 | ERR142097 | 1 | | 1 | 4167228 | 100 | 92 | 100 | 99 | 98 |
| 1607 | ERR142098 | 1 | | 1 | 4198160 | 100 | 93 | 100 | 99 | 99 |
| 1609 | ERR142100 | 1 | | 1 | 4147413 | 100 | 93 | 99 | 99 | 99 |
| 1610 | ERR142101 | 1 | | 1 | 4164736 | 100 | 93 | 99 | 99 | 100 |
| 1613 | ERR142104 | 1 | | 17 | 4248530 | 100 | 63 | 100 | 100 | 99 |
| 1615 | ERR142106 | 1 | | NF | 4267725 | 100 | 18 | 99 | 12 | 0 |
| 1620 | ERR142111 | 1 | | 1 | 4132606 | 100 | 94 | 100 | 99 | 99 |
| 1624 | ERR142115 | 1 | | 17 | 4225580 | 100 | 63 | 100 | 99 | 100 |
| 1627 | ERR142118 | 1 | | 1 | 4133112 | 100 | 95 | 99 | 100 | 100 |
| 1629 | ERR142120 | 1 | | 1 | 4145190 | 100 | 95 | 100 | 99 | 100 |
| 1631 | ERR142122 | 1 | | 1 | 4154178 | 100 | 97 | 100 | 100 | 99 |
| 1633 | ERR142124 | 1 | | 57 | 4150306 | 100 | 23 | 99 | 13 | 0 |
| 1634 | ERR142125 | 1 | | 14 | 4166923 | 100 | 94 | 99 | 100 | 99 |
| 1637 | ERR142128 | 1 | | 1 | 4140044 | 100 | 94 | 100 | 100 | 100 |
| 1638 | ERR142129 | 1 | | 1 | 4137208 | 100 | 95 | 100 | 99 | 100 |
| 1639 | ERR142130 | 1 | | 58 | 4229596 | 100 | 62 | 100 | 100 | 99 |

| | | | | | | | | | | |
|---|---|---|---|---|---|---|---|---|---|---|
| 1642 | ERR142133 | 1 | | 16 | 4385080 | 100 | 96 | 100 | 100 | 100 |
| 1643 | ERR142134 | 1 | | 6 | 4347563 | 100 | 19 | 100 | 10 | 0 |
| 1645 | ERR142136 | 1 | | 1 | 4194979 | 100 | 94 | 100 | 99 | 100 |
| 1647 | ERR142138 | 1 | | 1 | 4241455 | 100 | 93 | 99 | 99 | 100 |
| 1648 | ERR142139 | 1 | | 1 | 4136478 | 100 | 94 | 100 | 100 | 100 |
| 1649 | ERR142140 | 1 | | 1 | 4136940 | 100 | 93 | 99 | 99 | 98 |
| 1650 | ERR142141 | 1 | | 1 | 4145018 | 100 | 93 | 99 | 99 | 100 |
| 1654 | ERR142145 | 1 | | 1 | 4139068 | 100 | 94 | 99 | 100 | 100 |
| 1655 | ERR142146 | 1 | | 1 | 4166773 | 100 | 92 | 100 | 99 | 99 |
| 1656 | ERR142147 | 1 | | 1 | 4136756 | 100 | 94 | 100 | 100 | 99 |
| 1658 | ERR142149 | 1 | | 1 | 4422230 | 100 | 95 | 99 | 100 | 99 |
| 1661 | ERR142152 | 1 | | 17 | 4189441 | 100 | 62 | 100 | 100 | 99 |
| 1662 | ERR142153 | 1 | | 17 | 4201169 | 100 | 61 | 100 | 100 | 94 |
| 1663 | ERR142154 | 1 | | 1 | 4147494 | 100 | 93 | 100 | 100 | 99 |
| 1665 | ERR142156 | 1 | | 14 | 4174770 | 100 | 93 | 100 | 99 | 99 |
| 1666 | ERR142157 | 1 | | 2 | 4070583 | 100 | 92 | 100 | 100 | 100 |
| 1669 | ERR142160 | 1 | | 12 | 4144707 | 100 | 94 | 99 | 99 | 100 |
| 1671 | ERR142162 | 1 | | 12 | 4186402 | 100 | 94 | 100 | 100 | 100 |
| 1673 | ERR142164 | 1 | | 12 | 4137927 | 100 | 94 | 100 | 100 | 99 |
| 1677 | ERR142168 | 1 | | 17 | 4252437 | 100 | 61 | 100 | 99 | 99 |
| 1681 | ERR142172 | 1 | | 17 | 4255188 | 100 | 60 | 100 | 100 | 97 |
| 1682 | ERR142173 | 1 | | 17 | 4448825 | 100 | 61 | 100 | 99 | 99 |
| 1683 | ERR142174 | 1 | | 2 | 4226110 | 100 | 18 | 100 | 27 | 0 |
| 1685 | ERR142176 | 1 | | 42 | 4086238 | 100 | 95 | 100 | 100 | 100 |
| 1689 | ERR142180 | 1 | | 3 | 4135052 | 100 | 61 | 100 | 100 | 97 |
| 1690 | ERR142181 | 1 | | 54 | 4546565 | 100 | 100 | 100 | 99 | 100 |
| 1692 | ERR142183 | 1 | | 54 | 4298597 | 100 | 100 | 99 | 99 | 100 |
| 1693 | ERR142184 | 1 | | 54 | 4234422 | 100 | 99 | 100 | 100 | 100 |
| 1695 | ERR142186 | 1 | | NF | 4438711 | 100 | 99 | 99 | 99 | 99 |
| 1696 | ERR142187 | 1 | | 54 | 4280454 | 100 | 99 | 99 | 100 | 100 |
| 1697 | ERR142188 | 1 | | 58 | 4172717 | 100 | 14 | 100 | 0 | 0 |
| 1699 | ERR142190 | 1 | | 58 | 4162458 | 100 | 17 | 100 | 8 | 0 |
| 1704 | ERR142195 | 1 | | 375 | 4256717 | 100 | 93 | 100 | 100 | 100 |
| 1705 | ERR142196 | 1 | | 375 | 4239577 | 100 | 94 | 99 | 99 | 100 |
| 1707 | ERR142198 | 1 | | 17 | 4284108 | 100 | 60 | 100 | 99 | 100 |
| 1708 | ERR142199 | 1 | | 44 | 4538605 | 100 | 94 | 99 | 100 | 100 |
| 1714 | ERR142205 | 1 | | NF | 4370624 | 100 | 92 | 100 | 99 | 99 |
| 1716 | ERR142207 | 1 | | 6 | 4374639 | 100 | 13 | 99 | 0 | 0 |
| 1721 | ERR142212 | 1 | | 2 | 4180110 | 100 | 15 | 100 | 0 | 0 |
| 1722 | ERR142213 | 1 | | NF | 4204233 | 100 | 92 | 100 | 99 | 99 |
| 1740 | ERR142231 | 1 | | NF | 4351289 | 100 | 38 | 100 | 99 | 100 |
| 1746 | ERR142237 | 1 | | 22 | 4331216 | 100 | 37 | 99 | 99 | 89 |
| 1750 | ERR142241 | 1 | | 11 | 3989911 | 100 | 47 | 100 | 99 | 99 |
| 1752 | ERR142243 | 1 | | 11 | 3989394 | 100 | 48 | 100 | 100 | 99 |
| 1764 | ERR142255 | 1 | | 1 | 4367385 | 100 | 95 | 100 | 100 | 99 |
| 1778 | ERR142269 | 1 | | NF | 4407317 | 100 | 94 | 99 | 99 | 99 |
| 1783 | ERR142274 | 1 | | 1 | 4369679 | 100 | 94 | 99 | 99 | 100 |
| 1788 | ERR142279 | 1 | | NF | 4298506 | 100 | 12 | 99 | 0 | 0 |
| 1803 | ERR142294 | 1 | | NF | 4581465 | 100 | 60 | 99 | 100 | 94 |
| 1804 | ERR142295 | 1 | | NF | 4340184 | 100 | 62 | 99 | 99 | 99 |
| 1810 | ERR142301 | 1 | | 44 | 4202907 | 100 | 94 | 99 | 100 | 99 |
| 1813 | ERR142304 | 1 | | 1 | 4388488 | 100 | 95 | 99 | 99 | 99 |
| 1818 | ERR142309 | 1 | | 1 | 4162345 | 100 | 92 | 100 | 100 | 100 |
| 1820 | ERR142311 | 1 | | 5 | 4280198 | 100 | 41 | 100 | 99 | 99 |
| 1821 | ERR142312 | 1 | | NF | 4299205 | 100 | 94 | 99 | 100 | 100 |
| 1822 | ERR142313 | 1 | | 1 | 4155728 | 100 | 94 | 100 | 100 | 100 |
| 1826 | ERR142317 | 1 | | 41 | 4170062 | 100 | 94 | 100 | 100 | 99 |
| 1827 | ERR142318 | 1 | | 58 | 4677097 | 100 | 89 | 100 | 100 | 99 |
| 1829 | ERR142320 | 1 | | 58 | 4177219 | 100 | 63 | 99 | 99 | 99 |
| 1830 | ERR142321 | 1 | | 58 | 4166420 | 100 | 18 | 100 | 22 | 0 |
| 1835 | ERR142326 | 1 | | 63 | 4182737 | 100 | 99 | 99 | 99 | 99 |
| 1837 | ERR142328 | 1 | | 58 | 4212403 | 100 | 66 | 100 | 100 | 99 |
| 1838 | ERR142329 | 1 | | 6 | 4501092 | 100 | 67 | 100 | 99 | 100 |
| 1842 | ERR142333 | 1 | | 6 | 4425197 | 100 | 16 | 100 | 0 | 0 |
| 1850 | ERR142341 | 1 | | 6 | 4384896 | 100 | 15 | 99 | 0 | 0 |
| 1851 | ERR142342 | 1 | | 6 | 4402790 | 100 | 64 | 99 | 99 | 99 |
| 1854 | ERR142345 | 1 | | NF | 4279200 | 100 | 64 | 100 | 99 | 99 |
| 1855 | ERR142346 | 1 | | 6 | 4365719 | 100 | 64 | 100 | 100 | 99 |
| 1856 | ERR142347 | 1 | | 6 | 4520908 | 100 | 24 | 99 | 16 | 0 |
| 1857 | ERR142348 | 1 | | 41 | 4283090 | 100 | 95 | 100 | 100 | 100 |
| 1861 | ERR142352 | 1 | | 6 | 4331407 | 100 | 18 | 100 | 13 | 0 |
| 1862 | ERR142353 | 1 | | 6 | 4572169 | 100 | 16 | 99 | 0 | 0 |
| 1865 | ERR142356 | 1 | | 6 | 4302008 | 100 | 23 | 100 | 26 | 45 |
| 1881 | ERR142372 | 1 | | NF | 4570263 | 100 | 60 | 99 | 100 | 89 |
| 1885 | ERR142376 | 1 | | NF | 4135139 | 100 | 95 | 99 | 100 | 99 |
| 1886 | ERR142377 | 1 | | 44 | 4386266 | 100 | 94 | 99 | 99 | 99 |
| 1899 | ERR142390 | 1 | | 42 | 4191497 | 100 | 95 | 99 | 99 | 92 |
| 1909 | ERR142400 | 1 | | NF | 4387524 | 100 | 60 | 99 | 99 | 98 |
| 1914 | ERR142405 | 1 | | 44 | 4400214 | 100 | 61 | 99 | 99 | 99 |
| 1923 | ERR142414 | 1 | | 42 | 4177018 | 100 | 95 | 99 | 99 | 99 |
| 1925 | ERR142416 | 1 | | NF | 4681248 | 100 | 58 | 99 | 99 | 93 |
| 1939 | ERR142430 | 1 | | NF | 4377517 | 100 | 61 | 100 | 100 | 99 |
| 1955 | ERR142446 | 1 | | NF | 4241924 | 100 | 56 | 99 | 99 | 99 |
| 1988 | ERR142479 | 1 | | 6 | 4312260 | 100 | 19 | 99 | 0 | 0 |
| 2001 | ERR142492 | 1 | | 11 | 3972507 | 100 | 46 | 100 | 99 | 100 |
| 2004 | ERR142495 | 1 | | NF | 4532365 | 100 | 94 | 100 | 99 | 99 |
| 2005 | ERR142496 | 1 | | NF | 4201680 | 100 | 61 | 100 | 99 | 97 |
| 2007 | ERR142498 | 1 | | 44 | 4371496 | 100 | 60 | 99 | 99 | 96 |
| 2019 | ERR142510 | 1 | | 1 | 4130572 | 100 | 91 | 100 | 99 | 98 |
| 2037 | ERR142528 | 1 | | 1 | 4150587 | 100 | 92 | 99 | 99 | 99 |
| 2038 | ERR142529 | 1 | | 2 | 4195886 | 100 | 20 | 99 | 37 | 0 |
| 2046 | ERR142537 | 1 | | 45 | 4244118 | 100 | 94 | 99 | 99 | 100 |
| 2068 | ERR142559 | 1 | | 1 | 4151192 | 100 | 93 | 99 | 99 | 100 |
| 2069 | ERR142560 | 1 | | 1 | 4157512 | 100 | 93 | 99 | 99 | 98 |
| 2087 | ERR142578 | 1 | | 8 | 4304973 | 100 | 94 | 99 | 99 | 100 |
| 2088 | ERR142579 | 1 | | 1 | 4158056 | 100 | 93 | 99 | 99 | 99 |
| 2097 | ERR142588 | 1 | | 45 | 4580201 | 100 | 94 | 99 | 100 | 100 |
| 2100 | ERR142591 | 1 | | 55 | 4021267 | 100 | 94 | 99 | 99 | 99 |
| 2101 | ERR142592 | 1 | | 55 | 4040810 | 100 | 94 | 98 | 99 | 99 |
| 2108 | ERR142599 | 1 | | 55 | 4053300 | 100 | 94 | 100 | 100 | 100 |
| 2120 | ERR232375 | 1 | 023 | 5 | 4242373 | 100 | 39 | 100 | 99 | 95 |
| 2128 | ERR232383 | 1 | | 39 | 4212504 | 100 | 94 | 100 | 99 | 99 |
| 2135 | ERR232390 | 1 | | 238 | 4214716 | 100 | 95 | 99 | 100 | 99 |
| 2139 | ERR232394 | 1 | | 164 | 4025963 | 100 | 63 | 100 | 99 | 100 |
| 2150 | ERR251715 | 1 | | 1 | 4145965 | 100 | 93 | 100 | 99 | 99 |
| 2151 | ERR251716 | 1 | | 8 | 4292907 | 100 | 95 | 99 | 99 | 99 |
| 2158 | ERR251723 | 1 | | 49 | 4357939 | 100 | 92 | 99 | 99 | 100 |
| 2160 | ERR251725 | 1 | | 1 | 4160038 | 100 | 93 | 100 | 99 | 100 |
| 2162 | ERR251727 | 1 | | NF | 4348419 | 100 | 92 | 100 | 99 | 99 |
| 2163 | ERR251728 | 1 | | 3 | 4238660 | 100 | 58 | 99 | 99 | 100 |
| 2167 | ERR251732 | 1 | | 1 | 4155989 | 100 | 93 | 100 | 99 | 100 |
| 2169 | ERR251734 | 1 | | 1 | 4198317 | 100 | 94 | 100 | 99 | 99 |
| 2170 | ERR251735 | 1 | | 2 | 4165904 | 100 | 92 | 99 | 100 | 99 |
| 2174 | ERR251739 | 1 | | 8 | 4306395 | 100 | 95 | 100 | 100 | 100 |
| 2180 | ERR251745 | 1 | | 3 | 4231547 | 100 | 60 | 100 | 100 | 99 |
| 2185 | ERR251750 | 1 | | 8 | 4053097 | 100 | 52 | 100 | 100 | 93 |
| 2190 | ERR251755 | 1 | | 3 | 4363472 | 100 | 59 | 100 | 99 | 98 |
| 2191 | ERR251756 | 1 | | 1 | 4149216 | 100 | 93 | 100 | 99 | 100 |
| 2192 | ERR251757 | 1 | | 1 | 4152083 | 100 | 93 | 100 | 99 | 100 |
| 2193 | ERR251758 | 1 | | 1 | 4151346 | 100 | 93 | 99 | 99 | 100 |
| 2194 | ERR251759 | 1 | | 13 | 4273159 | 100 | 92 | 100 | 99 | 99 |
| 2196 | ERR251761 | 1 | | 72 | 4351285 | 100 | 67 | 99 | 99 | 92 |
| 2197 | ERR251762 | 1 | | 37 | 4249498 | 100 | 94 | 99 | 100 | 99 |
| 2200 | ERR251765 | 1 | | 33 | 4356138 | 100 | 93 | 100 | 99 | 99 |
| 2202 | ERR251767 | 1 | | 1 | 4172801 | 100 | 93 | 99 | 99 | 99 |
| 2203 | ERR251768 | 1 | | 1 | 4151624 | 100 | 93 | 99 | 99 | 99 |
| 2209 | ERR251774 | 1 | | 5 | 4212809 | 100 | 39 | 100 | 98 | 97 |
| 2210 | ERR251775 | 1 | | 1 | 4156639 | 100 | 93 | 100 | 99 | 99 |
| 2212 | ERR251777 | 1 | | 8 | 4228578 | 100 | 53 | 99 | 100 | 93 |
| 2214 | ERR251779 | 1 | | 8 | 4236667 | 100 | 95 | 99 | 99 | 99 |
| 2221 | ERR251786 | 1 | | 5 | 4294457 | 100 | 38 | 100 | 99 | 99 |
| 2222 | ERR251787 | 1 | | 271 | 4228764 | 100 | 60 | 99 | 99 | 89 |
| 2223 | ERR251788 | 1 | | 53 | 4076028 | 100 | 93 | 99 | 100 | 100 |
| 2224 | ERR251789 | 1 | | 22 | 4676415 | 100 | 73 | 99 | 99 | 91 |
| 2226 | ERR251791 | 1 | | 1 | 4089961 | 100 | 93 | 100 | 99 | 100 |
| 2230 | ERR251795 | 1 | | NF | 5031932 | 100 | 93 | 99 | 100 | 99 |
| 2239 | ERR251804 | 1 | | NF | 4605471 | 100 | 99 | 100 | 100 | 99 |
| 2241 | ERR251806 | 1 | | 3 | 4415139 | 100 | 81 | 99 | 99 | 98 |
| 2242 | ERR251807 | 1 | | 8 | 4345357 | 100 | 95 | 99 | 99 | 100 |
| 2243 | ERR251808 | 1 | | 5 | 4371709 | 100 | 44 | 100 | 99 | 99 |
| 2244 | ERR251809 | 1 | | 14 | 4370821 | 100 | 92 | 100 | 99 | 99 |
| 2245 | ERR251810 | 1 | | 46 | 4352585 | 100 | 94 | 99 | 99 | 99 |
| 2260 | ERR251825 | 1 | | 8 | 4347464 | 100 | 95 | 100 | 99 | 99 |
| 2261 | ERR251826 | 1 | | 5 | 4375857 | 100 | 47 | 99 | 99 | 93 |
| 2264 | ERR251829 | 1 | | 1 | 4518864 | 100 | 95 | 99 | 99 | 100 |
| 2268 | ERR251833 | 1 | | 1 | 4851731 | 100 | 94 | 100 | 99 | 100 |
| 2269 | ERR251834 | 1 | | 1 | 4507631 | 100 | 94 | 99 | 100 | 100 |
| 2270 | ERR251835 | 1 | | 34 | 4634983 | 100 | 91 | 99 | 99 | 99 |
| 2271 | ERR251836 | 1 | | 14 | 4546691 | 100 | 95 | 100 | 99 | 99 |
| 2273 | ERR251838 | 1 | | 14 | 4558143 | 100 | 95 | 100 | 100 | 100 |
| 2275 | ERR251840 | 1 | | 2 | 4415984 | 100 | 99 | 100 | 99 | 99 |
| 2276 | ERR251841 | 1 | | 1 | 4142500 | 100 | 93 | 100 | 99 | 99 |
| 2277 | ERR251842 | 1 | | 3 | 4307621 | 100 | 63 | 100 | 99 | 95 |
| 2279 | ERR251844 | 1 | | 5 | 4354942 | 100 | 37 | 99 | 99 | 93 |
| 2288 | ERR251865 | 1 | | 2 | 4095668 | 100 | 13 | 100 | 0 | 0 |
| 2289 | ERR251866 | 1 | | 1 | 4320987 | 100 | 94 | 99 | 99 | 98 |
| 2292 | ERR251873 | 1 | | 6 | 4239131 | 100 | 19 | 99 | 13 | 0 |
| 2296 | ERR266409 | 1 | | 58 | 4106651 | 100 | 13 | 100 | 0 | 0 |
| 2297 | ERR266410 | 1 | | 58 | 4227281 | 100 | 65 | 100 | 99 | 99 |
| 2299 | ERR272058 | 1 | | 37 | 4305105 | 100 | 94 | 99 | 99 | 99 |
| 2301 | ERR272060 | 1 | | 37 | 4252438 | 100 | 94 | 99 | 99 | 99 |
| 2303 | ERR272062 | 1 | | 39 | 4162831 | 100 | 94 | 100 | 100 | 99 |
| 2305 | ERR272064 | 1 | | 39 | 4227864 | 100 | 94 | 99 | 99 | 99 |
| 2307 | ERR272066 | 1 | | 58 | 4100033 | 100 | 12 | 100 | 0 | 0 |
| 2308 | ERR272067 | 1 | | 58 | 4408919 | 100 | 14 | 100 | 8 | 0 |
| 2312 | ERR272071 | 1 | | 58 | 4118938 | 100 | 13 | 100 | 0 | 0 |
| 2315 | ERR272074 | 1 | | 54 | 4282061 | 100 | 100 | 99 | 99 | 99 |
| 2319 | ERR272078 | 1 | | 54 | 4453392 | 100 | 100 | 100 | 99 | 100 |
| 2322 | ERR272081 | 1 | | 54 | 4245145 | 100 | 100 | 100 | 99 | 100 |
| 2328 | ERR272087 | 1 | | 3 | 4244668 | 100 | 59 | 100 | 99 | 99 |
| 2331 | ERR272090 | 1 | | 3 | 4293245 | 100 | 41 | 100 | 47 | 45 |
| 2334 | ERR272093 | 1 | | 3 | 4194582 | 100 | 60 | 100 | 100 | 99 |
| 2336 | ERR272095 | 1 | | 271 | 4179673 | 100 | 61 | 99 | 99 | 99 |
| 2339 | ERR272098 | 1 | | 3 | 4244807 | 100 | 63 | 99 | 99 | 99 |
| 2340 | ERR272099 | 1 | | 271 | 4168318 | 100 | 59 | 100 | 99 | 94 |
| 2352 | ERR272111 | 1 | | 271 | 4214784 | 100 | 62 | 100 | 100 | 99 |
| 2354 | ERR272113 | 1 | | 3 | 4178049 | 100 | 59 | 100 | 100 | 99 |
| 2359 | ERR272118 | 1 | | 3 | 4275263 | 100 | 11 | 100 | 0 | 0 |
| 2360 | ERR272119 | 1 | | 3 | 4144350 | 100 | 15 | 100 | 34 | 0 |
| 2361 | ERR272120 | 1 | | 3 | 4209112 | 100 | 58 | 100 | 100 | 99 |
| 2364 | ERR272123 | 1 | | 271 | 4335449 | 100 | 19 | 99 | 26 | 0 |
| 2369 | ERR272128 | 1 | | 3 | 4367664 | 100 | 58 | 99 | 100 | 93 |
| 2373 | ERR272132 | 1 | | 3 | 4393545 | 100 | 28 | 100 | 26 | 0 |
| 2375 | ERR272134 | 1 | | 3 | 4443362 | 100 | 58 | 99 | 99 | 84 |
| 2381 | ERR272140 | 1 | | 3 | 4258707 | 100 | 61 | 100 | 99 | 94 |
| 2382 | ERR272141 | 1 | | 3 | 4364721 | 100 | 61 | 100 | 79 | 67 |
| 2390 | ERR272149 | 1 | | 3 | 4122830 | 100 | 61 | 99 | 99 | 99 |

| | | | | | | | | | | |
|---|---|---|---|---|---|---|---|---|---|---|
| 2392 | ERR272151 | 1 | | 3 | 4441764 | 100 | 21 | 100 | 0 | 0 |
| 2397 | ERR272156 | 1 | | 6 | 4392938 | 100 | 23 | 99 | 29 | 0 |
| 2398 | ERR272157 | 1 | | 6 | 4688390 | 100 | 92 | 100 | 100 | 96 |
| 2404 | ERR272163 | 1 | | 6 | 4249052 | 100 | 60 | 99 | 100 | 95 |
| 2414 | ERR272173 | 1 | | 6 | 4337680 | 100 | 63 | 100 | 99 | 98 |
| 2416 | ERR272175 | 1 | | 6 | 4313886 | 100 | 63 | 100 | 100 | 90 |
| 2418 | ERR272177 | 1 | | 6 | 4253442 | 100 | 21 | 100 | 13 | 45 |
| 2419 | ERR272178 | 1 | | 6 | 4387692 | 100 | 64 | 99 | 99 | 91 |
| 2421 | ERR272180 | 1 | | 6 | 4362020 | 100 | 24 | 100 | 0 | 0 |
| 2422 | ERR272181 | 1 | | 6 | 4360170 | 100 | 24 | 100 | 0 | 0 |
| 2423 | ERR272182 | 1 | | NF | 4351746 | 100 | 15 | 100 | 13 | 0 |
| 2430 | ERR272189 | 1 | | 6 | 4288240 | 100 | 66 | 100 | 100 | 99 |
| 2431 | ERR272190 | 1 | | 6 | 4291693 | 100 | 64 | 100 | 100 | 91 |
| 2435 | ERR272194 | 1 | | 7 | 4435051 | 100 | 23 | 99 | 37 | 45 |
| 2439 | ERR272198 | 1 | | 7 | 4183639 | 100 | 21 | 100 | 26 | 0 |
| 2445 | ERR272204 | 1 | | NF | 4084354 | 100 | 15 | 99 | 0 | 45 |
| 2450 | ERR272209 | 1 | | 7 | 4172393 | 100 | 18 | 100 | 26 | 0 |
| 2451 | ERR272210 | 1 | | 7 | 4118433 | 100 | 13 | 99 | 0 | 0 |
| 2453 | ERR272212 | 1 | | 7 | 4087016 | 100 | 21 | 100 | 13 | 0 |
| 2456 | ERR272215 | 1 | | 7 | 4121374 | 100 | 19 | 100 | 23 | 0 |
| 2462 | ERR272221 | 1 | | 7 | 4202371 | 100 | 28 | 100 | 22 | 0 |
| 2463 | ERR272222 | 1 | | 7 | 4429828 | 100 | 17 | 100 | 6 | 0 |
| 2470 | ERR272229 | 1 | | 7 | 4546544 | 100 | 78 | 99 | 79 | 82 |
| 2471 | ERR272230 | 1 | | 7 | 4191336 | 100 | 31 | 99 | 57 | 53 |
| 2473 | ERR272232 | 1 | | 7 | 4216618 | 100 | 18 | 100 | 0 | 0 |
| 2475 | ERR272234 | 1 | | 7 | 4353627 | 100 | 23 | 99 | 6 | 0 |
| 2477 | ERR272236 | 1 | | 7 | 4226455 | 100 | 15 | 99 | 0 | 0 |
| 2486 | ERR272245 | 1 | | 44 | 4356788 | 100 | 63 | 99 | 100 | 94 |
| 2493 | ERR272252 | 1 | | 44 | 4417649 | 100 | 66 | 99 | 99 | 91 |
| 2510 | ERR467536 | 1 | | 54 | 4603659 | 100 | 100 | 100 | 99 | 100 |
| 2516 | ERR467542 | 1 | | 54 | 4604252 | 100 | 100 | 99 | 100 | 99 |
| 2523 | ERR467549 | 1 | | 1 | 4250321 | 100 | 93 | 100 | 99 | 99 |
| 2524 | ERR467550 | 1 | | 54 | 4556331 | 100 | 100 | 99 | 100 | 100 |
| 2526 | ERR467552 | 1 | | 54 | 4604742 | 100 | 100 | 100 | 100 | 99 |
| 2527 | ERR467553 | 1 | | 1 | 4198596 | 100 | 93 | 100 | 100 | 100 |
| 2532 | ERR467558 | 1 | | 1 | 4190867 | 100 | 93 | 100 | 99 | 100 |
| 2535 | ERR467561 | 1 | | 1 | 4195634 | 100 | 93 | 100 | 99 | 100 |
| 2536 | ERR467562 | 1 | | 1 | 4163551 | 100 | 93 | 99 | 100 | 99 |
| 2541 | ERR467567 | 1 | | 37 | 4268026 | 100 | 93 | 99 | 99 | 100 |
| 2544 | ERR467570 | 1 | | 14 | 4200440 | 100 | 92 | 100 | 100 | 99 |
| 2545 | ERR467571 | 1 | | 14 | 4423202 | 100 | 92 | 100 | 99 | 99 |
| 2549 | ERR467575 | 1 | | | 4253465 | 100 | 92 | 99 | 100 | 100 |
| 2562 | ERR467588 | 1 | | 54 | 4663795 | 100 | 100 | 100 | 99 | 100 |
| 2563 | ERR467589 | 1 | | 1 | 4152390 | 100 | 93 | 99 | 99 | 99 |
| 2564 | ERR467590 | 1 | | 1 | 4194184 | 100 | 92 | 100 | 99 | 99 |
| 2565 | ERR467591 | 1 | | 54 | 4677775 | 100 | 100 | 100 | 99 | 99 |
| 2567 | ERR467593 | 1 | | 1 | 4203095 | 100 | 93 | 100 | 99 | 100 |
| 2572 | ERR467598 | 1 | | - | 4192055 | 100 | 18 | 100 | 0 | 0 |
| 2577 | ERR467603 | 1 | | 8 | 4358888 | 100 | 95 | 100 | 100 | 99 |
| 2582 | ERR467608 | 1 | | 2 | 4189192 | 100 | 15 | 100 | 0 | 0 |
| 2587 | ERR467613 | 1 | | 15 | 4075504 | 100 | 94 | 100 | 100 | 100 |
| 2588 | ERR467614 | 1 | | 37 | 4267851 | 100 | 94 | 98 | 99 | 99 |
| 2590 | ERR467616 | 1 | | - | 4064196 | 100 | 44 | 99 | 98 | 90 |
| 2592 | ERR467618 | 1 | | 2 | 4139634 | 100 | 17 | 100 | 0 | 0 |
| 2612 | ERR008626 | 1 | | 1 | 4196796 | 100 | 92 | 99 | 99 | 99 |
| 2622 | ERR008636 | 1 | | 1 | 4197644 | 100 | 92 | 99 | 100 | 99 |
| 2651 | MID11-7032[illegible] | 1 | 010 | 15 | 4492700 | 100 | 94 | 100 | 100 | 100 |
| 2652 | MID12-7032[illegible] | 1 | 010 | 15 | 4157479 | 100 | 94 | 100 | 100 | 100 |
| 2653 | MID13-7032[illegible] | 1 | 010 | 15 | 4156582 | 100 | 94 | 100 | 100 | 100 |
| 2654 | E1 | 1 | 126 | 11 | 3966825 | 100 | 47 | 100 | 97 | 100 |
| 2655 | E10 | 1 | 033 | 11 | 3865634 | 100 | 17 | 100 | 0 | 0 |
| 2658 | E14 | 1 | 014 | 2 | 4163067 | 100 | 94 | 100 | 100 | 100 |
| 2659 | E15 | 1 | 075 | 95 | 4370091 | 100 | 10 | 100 | 0 | 0 |
| 2660 | E16 | 1 | 015 | 7 | 4263783 | 100 | 94 | 100 | 99 | 99 |
| 2661 | E19 | 1 | 577 | 62 | 4016285 | 100 | 17 | 100 | 4 | 0 |
| 2662 | E23 | 1 | 001 | 3 | 4041692 | 100 | 85 | 100 | 100 | 89 |
| 2663 | E24 | 1 | 020 | 2 | 4016330 | 100 | 97 | 100 | 100 | 100 |
| 2664 | E25 | 1 | 005 | 6 | 4148697 | 100 | 61 | 100 | 100 | 89 |
| 2665 | E28 | 1 | 012 | 54 | 4011167 | 100 | 99 | 100 | 100 | 100 |
| 2666 | E7 | 1 | 053 | 63 | 4251106 | 100 | 99 | 100 | 100 | 100 |
| 2667 | E9 | 1 | 009 | 3 | 4193506 | 100 | 15 | 100 | 2 | 16 |
| 2668 | T10 | 1 | 165 | 3 | 4088109 | 100 | 60 | 100 | 100 | 89 |
| 2669 | T11 | 1 | 075 | 95 | 4226426 | 100 | 12 | 100 | 4 | 0 |
| 2670 | T14 | 1 | 156 | 42 | 4037952 | 100 | 97 | 99 | 100 | 100 |
| 2672 | T17 | 1 | 025 | 49 | 4033051 | 100 | 97 | 100 | 100 | 99 |
| 2673 | T19 | 1 | 057 | 55 | 3980802 | 100 | 97 | 100 | 100 | 100 |
| 2674 | T20 | 1 | 078 | 11 | 3836875 | 100 | 47 | 100 | 97 | 100 |
| 2675 | T22 | 1 | | 6 | 4116425 | 100 | 17 | 100 | 4 | 0 |
| 2676 | T22bis | 1 | | 8 | 4227904 | 100 | 95 | 100 | 100 | 100 |
| 2677 | T23 | 1 | 019 | 67 | 4108039 | 100 | 97 | 100 | 100 | 100 |
| 2679 | T42 | 1 | 020 | 2 | 4137327 | 100 | 96 | 100 | 95 | 94 |
| 2680 | T5 | 1 | 079 | 11 | 3983922 | 100 | 47 | 100 | 99 | 100 |
| 2681 | T6 | 1 | 095 | 13 | 4345136 | 100 | 94 | 100 | 100 | 100 |
| 2682 | T61 | 1 | 015 | 7 | 4079543 | 100 | 89 | 100 | 98 | 58 |

**Table S9. IntaRNA prediction for ncRNA-mRNA interactions**

Strand "+"

| | | | | | RIPseq log p-value | |
|---|---|---|---|---|---|---|
| **CDS** | **ncRNA** | **interaction CDS** | **interaction ncRNA** | **interaction energy (kcal/mol)** | **cds logpVal** | **ncRNA logpVal** |
| CD630_05460 | CD630_Cdi1_11 | 51 – 113 | 93 – 151 | -15.7611 | 3.98753 | 5.77720 |
| CD630_21700 | CD630_Cdi1_11 | 22 – 41 | 125 – 149 | -15.289 | 3.96650 | 5.77720 |
| CD630_06080 | CD630_Cdi1_11 | 1 – 57 | 97 – 151 | -14.7334 | 3.82315 | 5.77720 |
| CD630_19401 | CD630_Cdi1_11 | 50 – 107 | 98 – 157 | -14.6303 | 4.37084 | 5.77720 |
| CD630_31461 | CD630_Cdi1_11 | 141 – 190 | 97 – 145 | -14.155 | 4.41476 | 5.77720 |
| CD630_17281 | CD630_Cdi1_11 | 50 – 93 | 114 – 158 | -13.9786 | 5.91544 | 5.77720 |
| CD630_17511 | CD630_Cdi1_11 | 51 – 109 | 96 – 150 | -13.6149 | 9.12322 | 5.77720 |
| CD630_20450 | CD630_Cdi1_11 | 113 – 165 | 93 – 148 | -13.5235 | 3.77388 | 5.77720 |
| CD630_03420 | CD630_Cdi1_11 | 138 – 177 | 111 – 151 | -13.026 | 4.71868 | 5.77720 |
| CD630_16540 | CD630_Cdi1_11 | 161 – 184 | 122 – 145 | -12.9793 | 4.37658 | 5.77720 |
| feoB | CD630_Cdi1_11 | 134 – 197 | 88 – 145 | -12.4775 | 4.05364 | 5.77720 |
| CD630_08080 | CD630_Cdi1_11 | 31 – 51 | 124 – 146 | -12.4487 | 4.87819 | 5.77720 |
| CD630_21570 | CD630_Cdi1_11 | 157 – 190 | 110 – 143 | -12.087 | 6.48385 | 5.77720 |
| CD630_23070 | CD630_Cdi1_11 | 51 – 186 | 17 – 131 | -12.0419 | 5.23027 | 5.77720 |
| CD630_06170 | CD630_Cdi1_11 | 117 – 154 | 105 – 146 | -12.0109 | 4.95366 | 5.77720 |
| CD630_21260 | CD630_Cdi1_11 | 140 – 152 | 44 – 56 | -11.8608 | 4.22255 | 5.77720 |
| glgP | CD630_Cdi1_11 | 75 – 129 | 97 – 158 | -11.8245 | 3.28530 | 5.77720 |
| CD630_24830 | CD630_Cdi1_11 | 1 – 28 | 104 – 145 | -11.7711 | 4.98239 | 5.77720 |
| CD630_13020 | CD630_Cdi1_11 | 85 – 123 | 107 – 145 | -11.5882 | 3.52267 | 5.77720 |
| CD630_14700 | CD630_Cdi1_11 | 125 – 159 | 111 – 146 | -11.3814 | 3.98234 | 5.77720 |
| CD630_18960 | CD630_Cdi1_11 | 2 – 33 | 110 – 141 | -11.3448 | 4.37822 | 5.77720 |
| CD630_14600 | CD630_Cdi1_11 | 137 – 156 | 130 – 149 | -11.336 | 5.90348 | 5.77720 |
| CD630_22200 | CD630_Cdi1_11 | 143 – 173 | 115 – 146 | -11.3014 | 4.18288 | 5.77720 |
| CD630_05830 | CD630_Cdi1_11 | 144 – 178 | 125 – 157 | -11.2328 | 3.35360 | 5.77720 |
| CD630_20600 | CD630_Cdi1_11 | 150 – 178 | 124 – 152 | -11.1527 | 4.20093 | 5.77720 |
| ppnK | CD630_Cdi1_11 | 142 – 162 | 133 – 148 | -11.1419 | 4.36817 | 5.77720 |
| CD630_03040 | CD630_Cdi1_11 | 51 – 96 | 111 – 159 | -11.0959 | 3.51403 | 5.77720 |
| sspA | CD630_Cdi1_11 | 50 – 109 | 96 – 156 | -11.0512 | 3.62689 | 5.77720 |
| CD630_13790 | CD630_Cdi1_11 | 139 – 196 | 93 – 145 | -10.8687 | 5.17374 | 5.77720 |
| CD630_22550 | CD630_Cdi1_11 | 26 – 51 | 125 – 150 | -10.7309 | 3.62379 | 5.77720 |
| CD630_22030 | CD630_Cdi1_11 | 138 – 200 | 93 – 156 | -10.7224 | 3.29934 | 5.77720 |
| kdpD | CD630_Cdi1_11 | 152 – 201 | 94 – 140 | -10.6711 | 4.27140 | 5.77720 |
| ldh | CD630_Cdi1_11 | 157 – 171 | 9 – 23 | -10.6229 | 4.35818 | 5.77720 |
| CD630_02420 | CD630_Cdi1_11 | 9 – 50 | 48 – 94 | -10.5298 | 3.71048 | 5.77720 |
| CD630_34050 | CD630_Cdi1_11 | 8 – 20 | 4 – 16 | -10.4905 | 4.03863 | 5.77720 |
| CD630_09060 | CD630_Cdi1_11 | 40 – 93 | 112 – 155 | -10.4761 | 4.31834 | 5.77720 |
| CD630_04620 | CD630_Cdi1_11 | 157 – 169 | 135 – 147 | -10.3911 | 3.45413 | 5.77720 |
| CD630_15360 | CD630_Cdi1_11 | 24 – 37 | 42 – 55 | -10.3438 | 4.47208 | 5.77720 |
| CD630_13370 | CD630_Cdi1_11 | 156 – 182 | 123 – 149 | -10.3027 | 4.33145 | 5.77720 |
| CD630_20670 | CD630_Cdi1_11 | 179 – 197 | 78 – 95 | -10.2721 | 4.42757 | 5.77720 |
| CD630_25190 | CD630_Cdi1_11 | 174 – 181 | 93 – 100 | -10.247 | 3.19663 | 5.77720 |
| CD630_31530 | CD630_Cdi1_11 | 118 – 168 | 90 – 143 | -10.2313 | 3.49100 | 5.77720 |
| CD630_20170 | CD630_Cdi1_11 | 2 – 23 | 141 – 159 | -10.1416 | 6.90201 | 5.77720 |
| pilD | CD630_Cdi1_11 | 3 – 71 | 73 – 138 | -10.0901 | 4.54123 | 5.77720 |
| CD630_30310 | CD630_Cdi1_11 | 142 – 154 | 134 – 146 | -10.0889 | 4.56059 | 5.77720 |
| CD630_10880 | CD630_Cdi1_11 | 5 – 25 | 137 – 158 | -10.0436 | 5.05048 | 5.77720 |
| CD630_13620 | CD630_Cdi1_11 | 133 – 164 | 124 – 155 | -9.9987 | 4.77627 | 5.77720 |
| CD630_34030 | CD630_Cdi1_11 | 42 – 66 | 134 – 157 | -9.85881 | 3.05766 | 5.77720 |
| CD630_19904 | CD630_Cdi1_11 | 23 – 36 | 134 – 148 | -9.73264 | 5.96526 | 5.77720 |
| appC | CD630_Cdi1_11 | 49 – 72 | 137 – 157 | -9.62671 | 4.50438 | 5.77720 |
| CD630_05160 | CD630_Cdi1_11 | 151 – 188 | 113 – 149 | -9.59154 | 5.03573 | 5.77720 |
| glnA | CD630_Cdi1_11 | 162 – 172 | 134 – 144 | -9.51916 | 3.87629 | 5.77720 |
| CD630_15850 | CD630_Cdi1_3 | 16 – 50 | 291 – 323 | -17.1219 | 4.32968 | 6.59405 |
| CD630_27120 | CD630_Cdi1_3 | 128 – 179 | 324 – 374 | -16.7799 | 4.43720 | 6.59405 |
| CD630_08080 | CD630_Cdi1_3 | 26 – 36 | 486 – 496 | -16.5146 | 4.87819 | 6.59405 |
| CD630_03410 | CD630_Cdi1_3 | 154 – 165 | 485 – 496 | -15.3605 | 4.45994 | 6.59405 |
| CD630_10220 | CD630_Cdi1_3 | 113 – 173 | 205 – 264 | -15.225 | 4.08233 | 6.59405 |
| CD630_04410 | CD630_Cdi1_3 | 60 – 102 | 70 – 124 | -15.1374 | 3.90451 | 6.59405 |
| CD630_17970 | CD630_Cdi1_3 | 108 – 143 | 424 – 458 | -15.1221 | 3.68802 | 6.59405 |
| cdtR | CD630_Cdi1_3 | 29 – 51 | 184 – 206 | -14.9666 | 5.84368 | 6.59405 |
| CD630_09570 | CD630_Cdi1_3 | 5 – 49 | 304 – 347 | -14.58 | 5.31213 | 6.59405 |
| CD630_29070 | CD630_Cdi1_3 | 5 – 49 | 304 – 347 | -14.58 | 3.89477 | 6.59405 |
| CD630_31490 | CD630_Cdi1_3 | 36 – 96 | 440 – 494 | -14.3537 | 4.08593 | 6.59405 |
| CD630_31910 | CD630_Cdi1_3 | 135 – 167 | 427 – 461 | -14.2766 | 3.48314 | 6.59405 |
| CD630_23070 | CD630_Cdi1_3 | 172 – 187 | 76 – 90 | -14.1207 | 5.23027 | 6.59405 |
| CD630_20280 | CD630_Cdi1_3 | 117 – 137 | 72 – 91 | -13.966 | 3.46343 | 6.59405 |
| CD630_09100 | CD630_Cdi1_3 | 93 – 108 | 475 – 491 | -13.9607 | 5.92260 | 6.59405 |
| CD630_22200 | CD630_Cdi1_3 | 79 – 140 | 309 – 364 | -13.9411 | 4.18288 | 6.59405 |
| CD630_26150 | CD630_Cdi1_3 | 158 – 199 | 425 – 450 | -13.7703 | 4.06302 | 6.59405 |
| CD630_06170 | CD630_Cdi1_3 | 115 – 150 | 463 – 493 | -13.6846 | 4.95366 | 6.59405 |
| CD630_05830 | CD630_Cdi1_3 | 10 – 29 | 142 – 161 | -13.6693 | 3.35360 | 6.59405 |
| CD630_19921 | CD630_Cdi1_3 | 24 – 59 | 433 – 473 | -13.6185 | 5.46345 | 6.59405 |
| cbiQ | CD630_Cdi1_3 | 101 – 170 | 146 – 225 | -13.4038 | 3.79830 | 6.59405 |
| CD630_19942 | CD630_Cdi1_3 | 114 – 130 | 317 – 333 | -13.2269 | 4.24574 | 6.59405 |
| CD630_16160 | CD630_Cdi1_3 | 32 – 52 | 241 – 262 | -13.154 | 6.11801 | 6.59405 |
| CD630_16940 | CD630_Cdi1_3 | 87 – 128 | 169 – 214 | -13.0926 | 4.14665 | 6.59405 |
| cwp17 | CD630_Cdi1_3 | 131 – 150 | 84 – 101 | -12.9854 | 3.43363 | 6.59405 |
| CD630_32050 | CD630_Cdi1_3 | 129 – 150 | 311 – 333 | -12.9806 | 5.73797 | 6.59405 |
| CD630_08950 | CD630_Cdi1_3 | 15 – 71 | 422 – 496 | -12.9384 | 3.71905 | 6.59405 |
| CD630_08370 | CD630_Cdi1_3 | 126 – 198 | 266 – 333 | -12.8658 | 3.65639 | 6.59405 |
| CD630_14620 | CD630_Cdi1_3 | 18 – 58 | 191 – 231 | -12.7694 | 4.43651 | 6.59405 |
| CD630_23780 | CD630_Cdi1_3 | 62 – 101 | 167 – 217 | -12.6744 | 3.91629 | 6.59405 |
| CD630_07400 | CD630_Cdi1_3 | 159 – 170 | 485 – 496 | -12.6443 | 4.52273 | 6.59405 |
| CD630_17160 | CD630_Cdi1_3 | 86 – 111 | 309 – 336 | -12.5804 | 3.50228 | 6.59405 |
| CD630_21620 | CD630_Cdi1_3 | 137 – 185 | 289 – 336 | -12.5378 | 3.02649 | 6.59405 |
| CD630_22510 | CD630_Cdi1_3 | 111 – 144 | 459 – 497 | -12.5346 | 11.42763 | 6.59405 |
| srlD | CD630_Cdi1_3 | 172 – 198 | 426 – 454 | -12.5008 | 4.19843 | 6.59405 |
| CD630_15200 | CD630_Cdi1_3 | 13 – 62 | 425 – 471 | -12.4931 | 4.00442 | 6.59405 |
| CD630_12650 | CD630_Cdi1_3 | 100 – 123 | 203 – 226 | -12.4104 | 6.31827 | 6.59405 |
| CD630_26680 | CD630_Cdi1_3 | 147 – 173 | 300 – 325 | -12.3746 | 3.53956 | 6.59405 |
| CD630_21720 | CD630_Cdi1_3 | 180 – 193 | 251 – 264 | -12.3666 | 4.13030 | 6.59405 |
| CD630_23861 | CD630_Cdi1_3 | 89 – 144 | 148 – 214 | -12.3222 | 5.83068 | 6.59405 |
| tcdC | CD630_Cdi1_3 | 116 – 166 | 317 – 375 | -12.3116 | 3.76101 | 6.59405 |
| CD630_24591 | CD630_Cdi1_3 | 39 – 93 | 440 – 493 | -12.2873 | 3.54497 | 6.59405 |
| CD630_21260 | CD630_Cdi1_3 | 138 – 152 | 37 – 51 | -12.2721 | 4.22255 | 6.59405 |
| CD630_34530 | CD630_Cdi1_3 | 9 – 24 | 250 – 265 | -12.133 | 3.83016 | 6.59405 |
| CD630_20260 | CD630_Cdi1_3 | 15 – 89 | 141 – 212 | -12.1201 | 3.84162 | 6.59405 |
| CD630_12320 | CD630_Cdi1_3 | 42 – 54 | 204 – 216 | -12.0022 | 3.28667 | 6.59405 |
| CD630_18311 | CD630_Cdi1_3 | 160 – 185 | 79 – 103 | -11.9947 | 6.86047 | 6.59405 |
| CD630_25190 | CD630_Cdi1_3 | 174 – 187 | 75 – 87 | -11.8902 | 3.19663 | 6.59405 |
| CD630_13560 | CD630_Cdi1_3 | 127 – 147 | 432 – 451 | -11.8827 | 3.38083 | 6.59405 |
| CD630_28181 | CD630_Cdi1_3 | 66 – 90 | 71 – 100 | -11.7935 | 3.91299 | 6.59405 |
| CD630_33420 | CD630_Cdi1_3 | 16 – 47 | 240 – 265 | -11.7776 | 3.22095 | 6.59405 |
| nrdD | CD630_Cdi1_3 | 69 – 108 | 77 – 114 | -11.7656 | 3.28290 | 6.59405 |
| CD630_14130 | CD630_Cdi1_3 | 167 – 196 | 442 – 472 | -11.7157 | 4.57970 | 6.59405 |
| CD630_25200 | CD630_Cdi1_3 | 50 – 101 | 426 – 469 | -11.6326 | 3.65158 | 6.59405 |
| CD630_18910 | CD630_Cdi1_3 | 25 – 71 | 464 – 497 | -11.6066 | 3.30688 | 6.59405 |
| CD630_09040 | CD630_Cdi1_3 | 163 – 185 | 143 – 164 | -11.5851 | 4.11851 | 6.59405 |
| CD630_32520 | CD630_Cdi1_3 | 151 – 167 | 317 – 334 | -11.569 | 4.06098 | 6.59405 |
| CD630_06540 | CD630_Cdi1_3 | 7 – 44 | 301 – 338 | -11.5158 | 3.54673 | 6.59405 |
| CD630_28870 | CD630_Cdi1_3 | 173 – 193 | 471 – 490 | -11.5148 | 3.41991 | 6.59405 |
| CD630_21110 | CD630_Cdi1_3 | 77 – 90 | 424 – 437 | -11.4883 | 3.89630 | 6.59405 |
| dapB2 | CD630_Cdi1_3 | 117 – 200 | 256 – 338 | -11.4492 | 3.27218 | 6.59405 |
| CD630_27310 | CD630_Cdi1_3 | 70 – 101 | 312 – 342 | -11.4331 | 4.01622 | 6.59405 |
| CD630_24770 | CD630_Cdi1_3 | 26 – 57 | 425 – 455 | -11.425 | 3.20798 | 6.59405 |
| CD630_35260 | CD630_Cdi1_3 | 116 – 180 | 392 – 454 | -11.4213 | 5.77277 | 6.59405 |
| CD630_21540 | CD630_Cdi1_3 | 127 – 156 | 85 – 117 | -11.418 | 4.33090 | 6.59405 |
| CD630_06060 | CD630_Cdi1_3 | 166 – 185 | 78 – 101 | -11.4127 | 4.94050 | 6.59405 |
| CD630_23440 | CD630_Cdi1_3 | 68 – 109 | 326 – 364 | -11.3984 | 3.34841 | 6.59405 |
| CD630_19801 | CD630_Cdi1_3 | 102 – 119 | 72 – 90 | -11.3124 | 5.22035 | 6.59405 |
| CD630_32070 | CD630_Cdi1_3 | 29 – 56 | 437 – 465 | -11.2999 | 3.51964 | 6.59405 |
| cdu2 | CD630_Cdi1_3 | 49 – 78 | 342 – 374 | -11.2733 | 2.97335 | 6.59405 |
| bglA | CD630_Cdi1_3 | 90 – 126 | 299 – 333 | -11.2211 | 4.50315 | 6.59405 |
| CD630_21140 | CD630_Cdi1_3 | 15 – 51 | 240 – 273 | -11.1894 | 3.33304 | 6.59405 |
| CD630_05030 | CD630_Cdi1_3 | 129 – 170 | 69 – 100 | -11.1887 | 4.03823 | 6.59405 |
| ribD | CD630_Cdi1_3 | 126 – 157 | 180 – 211 | -11.1831 | 6.65922 | 6.59405 |
| CD630_26691 | CD630_Cdi1_3 | 67 – 104 | 432 – 470 | -11.161 | 3.96230 | 6.59405 |
| CD630_18781 | CD630_Cdi1_3 | 76 – 109 | 425 – 455 | -11.1567 | 3.77660 | 6.59405 |
| bltD | CD630_Cdi1_3 | 133 – 151 | 70 – 87 | -11.1021 | 4.49926 | 6.59405 |
| CD630_24050 | CD630_Cdi1_3 | 111 – 140 | 90 – 112 | -11.0767 | 3.70192 | 6.59405 |
| CD630_32890 | CD630_Cdi1_3 | 62 – 110 | 325 – 379 | -11.0709 | 3.41276 | 6.59405 |
| CD630_08100 | CD630_Cdi1_3 | 22 – 65 | 62 – 98 | -11.0461 | 4.39181 | 6.59405 |
| CD630_28280 | CD630_Cdi1_3 | 22 – 67 | 252 – 293 | -11.0306 | 4.59833 | 6.59405 |
| CD630_16150 | CD630_Cdi1_3 | 156 – 185 | 214 – 246 | -11.0278 | 4.20614 | 6.59405 |
| CD630_17710 | CD630_Cdi1_3 | 82 – 107 | 297 – 322 | -11.0047 | 4.37688 | 6.59405 |
| garK | CD630_Cdi1_3 | 75 – 141 | 270 – 333 | -10.9876 | 3.72836 | 6.59405 |
| CD630_18230 | CD630_Cdi1_3 | 108 – 187 | 252 – 337 | -10.9808 | 3.00184 | 6.59405 |
| CD630_15890 | CD630_Cdi1_3 | 142 – 170 | 199 – 224 | -10.9263 | 3.39962 | 6.59405 |
| CD630_33580 | CD630_Cdi1_3 | 12 – 38 | 176 – 199 | -10.9169 | 3.95778 | 6.59405 |
| CD630_08460 | CD630_Cdi1_3 | 119 – 198 | 171 – 262 | -10.9155 | 4.15083 | 6.59405 |
| CD630_14420 | CD630_Cdi1_3 | 115 – 162 | 173 – 211 | -10.8896 | 3.99497 | 6.59405 |
| def1 | CD630_Cdi1_3 | 38 – 62 | 171 – 196 | -10.8849 | 3.99720 | 6.59405 |
| CD630_21760 | CD630_Cdi1_3 | 39 – 71 | 79 – 102 | -10.8548 | 4.43846 | 6.59405 |
| CD630_03400 | CD630_Cdi1_3 | 46 – 75 | 470 – 496 | -10.8196 | 4.45994 | 6.59405 |
| glgB | CD630_Cdi1_3 | 158 – 201 | 313 – 354 | -10.8042 | 4.48273 | 6.59405 |
| CD630_15160 | CD630_Cdi1_3 | 46 – 75 | 427 – 455 | -10.8031 | 4.32318 | 6.59405 |
| terD | CD630_Cdi1_3 | 98 – 125 | 420 – 449 | -10.758 | 3.33148 | 6.59405 |
| ppnK | CD630_Cdi1_3 | 134 – 160 | 310 – 338 | -10.7369 | 4.36817 | 6.59405 |
| feoB | CD630_Cdi1_3 | 185 – 197 | 75 – 88 | -10.7349 | 4.05364 | 6.59405 |
| CD630_17170 | CD630_Cdi1_3 | 188 – 200 | 78 – 89 | -10.7336 | 4.10230 | 6.59405 |
| ogt1 | CD630_Cdi1_3 | 49 – 103 | 299 – 356 | -10.7271 | 3.16772 | 6.59405 |
| CD630_19850 | CD630_Cdi1_3 | 110 – 127 | 196 – 213 | -10.7264 | 6.88486 | 6.59405 |
| CD630_18610 | CD630_Cdi1_3 | 20 – 35 | 323 – 336 | -10.7232 | 4.15582 | 6.59405 |
| CD630_27210 | CD630_Cdi1_3 | 2 – 36 | 350 – 379 | -10.707 | 4.18786 | 6.59405 |
| CD630_15360 | CD630_Cdi1_3 | 117 – 156 | 331 – 361 | -10.6831 | 4.47208 | 6.59405 |
| CD630_34560 | CD630_Cdi1_3 | 15 – 63 | 395 – 440 | -10.6555 | 4.02604 | 6.59405 |
| CD630_30420 | CD630_Cdi1_3 | 32 – 43 | 486 – 497 | -10.6441 | 5.27539 | 6.59405 |
| CD630_19903 | CD630_Cdi1_3 | 54 – 145 | 183 – 274 | -10.6386 | 5.79994 | 6.59405 |
| cynT | CD630_Cdi1_3 | 32 – 88 | 140 – 199 | -10.6314 | 4.21370 | 6.59405 |
| CD630_21090 | CD630_Cdi1_3 | 158 – 169 | 485 – 495 | -10.6289 | 4.15803 | 6.59405 |
| CD630_04320 | CD630_Cdi1_3 | 8 – 39 | 425 – 455 | -10.6128 | 3.68342 | 6.59405 |
| CD630_06100 | CD630_Cdi1_3 | 106 – 142 | 468 – 497 | -10.6107 | 4.43918 | 6.59405 |
| CD630_32720 | CD630_Cdi1_3 | 46 – 127 | 413 – 479 | -10.5992 | 4.74276 | 6.59405 |
| CD630_25610 | CD630_Cdi1_3 | 182 – 200 | 63 – 81 | -10.5955 | 3.02685 | 6.59405 |
| thiH | CD630_Cdi1_3 | 175 – 194 | 75 – 89 | -10.5455 | 3.58230 | 6.59405 |
| CD630_21490 | CD630_Cdi1_3 | 85 – 110 | 425 – 449 | -10.534 | 3.37462 | 6.59405 |
| CD630_18660 | CD630_Cdi1_3 | 9 – 37 | 417 – 446 | -10.5253 | 3.50371 | 6.59405 |
| CD630_30110 | CD630_Cdi1_3 | 161 – 191 | 424 – 454 | -10.5153 | 3.31041 | 6.59405 |
| kdpD | CD630_Cdi1_3 | 194 – 201 | 81 – 88 | -10.4983 | 4.27140 | 6.59405 |
| uxaA | CD630_Cdi1_3 | 8 – 22 | 73 – 88 | -10.4979 | 4.26165 | 6.59405 |
| CD630_07121 | CD630_Cdi1_3 | 5 – 12 | 486 – 493 | -10.4962 | 3.26177 | 6.59405 |
| CD630_05211 | CD630_Cdi1_3 | 126 – 157 | 180 – 222 | -10.4576 | 3.97048 | 6.59405 |
| CD630_27270 | CD630_Cdi1_3 | 149 – 176 | 240 – 273 | -10.4515 | 3.50892 | 6.59405 |
| CD630_28550 | CD630_Cdi1_3 | 74 – 115 | 206 – 263 | -10.3929 | 3.51196 | 6.59405 |
| CD630_13400 | CD630_Cdi1_3 | 131 – 156 | 426 – 447 | -10.3857 | 3.20232 | 6.59405 |
| CD630_01640 | CD630_Cdi1_3 | 140 – 159 | 236 – 259 | -10.3751 | 4.55381 | 6.59405 |
| CD630_32060 | CD630_Cdi1_3 | 31 – 48 | 240 – 256 | -10.3603 | 5.73797 | 6.59405 |
| CD630_28320 | CD630_Cdi1_3 | 185 – 200 | 72 – 88 | -10.3335 | 3.99184 | 6.59405 |
| CD630_08130 | CD630_Cdi1_3 | 176 – 194 | 66 – 82 | -10.3126 | 4.63831 | 6.59405 |
| CD630_19790 | CD630_Cdi1_3 | 41 – 68 | 423 – 453 | -10.2873 | 3.79593 | 6.59405 |
| oppC | CD630_Cdi1_3 | 131 – 176 | 311 – 358 | -10.2228 | 3.01695 | 6.59405 |
| fliI | CD630_Cdi1_3 | 32 – 79 | 309 – 355 | -10.218 | 3.03666 | 6.59405 |
| CD630_28150 | CD630_Cdi1_3 | 26 – 54 | 148 – 177 | -10.1835 | 4.18447 | 6.59405 |
| cspD | CD630_Cdi1_3 | 95 – 131 | 301 – 341 | -10.1826 | 3.23587 | 6.59405 |
| CD630_21240 | CD630_Cdi1_3 | 52 – 85 | 60 – 99 | -10.1702 | 5.88879 | 6.59405 |
| CD630_23930 | CD630_Cdi1_3 | 141 – 169 | 239 – 264 | -10.1515 | 3.05650 | 6.59405 |
| CD630_16540 | CD630_Cdi1_3 | 172 – 188 | 324 – 340 | -10.1274 | 4.37658 | 6.59405 |
| CD630_25040 | CD630_Cdi1_3 | 9 – 16 | 437 – 444 | -10.1253 | 3.43898 | 6.59405 |
| CD630_11241 | CD630_Cdi1_3 | 166 – 181 | 437 – 451 | -10.1233 | 4.85723 | 6.59405 |
| CD630_22070 | CD630_Cdi1_3 | 6 – 53 | 272 – 324 | -10.1207 | 4.09651 | 6.59405 |
| chrA | CD630_Cdi1_3 | 154 – 167 | 79 – 93 | -10.1171 | 5.27255 | 6.59405 |
| pilT | CD630_Cdi1_3 | 79 – 106 | 296 – 323 | -10.1123 | 3.24639 | 6.59405 |
| CD630_36690 | CD630_Cdi1_3 | 89 – 135 | 288 – 333 | -10.112 | 3.53761 | 6.59405 |
| CD630_03650 | CD630_Cdi1_3 | 151 – 163 | 87 – 99 | -10.1051 | 3.93697 | 6.59405 |
| CD630_13920 | CD630_Cdi1_3 | 34 – 48 | 118 – 132 | -10.0952 | 3.25236 | 6.59405 |
| CD630_14240 | CD630_Cdi1_3 | 54 – 93 | 239 – 275 | -10.0939 | 8.60766 | 6.59405 |
| CD630_06060 | CD630_Cdi1_3 | 107 – 145 | 203 – 246 | -10.0762 | 4.94050 | 6.59405 |
| CD630_13350 | CD630_Cdi1_3 | 44 – 56 | 173 – 185 | -10.0626 | 4.15139 | 6.59405 |
| pgm | CD630_Cdi1_3 | 145 – 186 | 310 – 340 | -10.0593 | 3.45439 | 6.59405 |
| dnaC | CD630_Cdi1_3 | 177 – 199 | 108 – 131 | -10.0526 | 6.30796 | 6.59405 |
| CD630_04341 | CD630_Cdi1_3 | 130 – 146 | 317 – 333 | -10.0339 | 4.29262 | 6.59405 |
| CD630_08390 | CD630_Cdi1_3 | 85 – 104 | 240 – 257 | -10.0236 | 5.16000 | 6.59405 |
| CD630_30100 | CD630_Cdi1_3 | 175 – 186 | 6 – 17 | -9.9904 | 4.13234 | 6.59405 |
| rbsC | CD630_Cdi1_3 | 34 – 62 | 77 – 100 | -9.93763 | 3.71650 | 6.59405 |
| CD630_27240 | CD630_Cdi1_3 | 5 – 45 | 301 – 336 | -9.93073 | 3.52462 | 6.59405 |
| CD630_31010 | CD630_Cdi1_3 | 136 – 155 | 429 – 447 | -9.89332 | 4.96305 | 6.59405 |
| cbiO | CD630_Cdi1_3 | 2 – 24 | 418 – 444 | -9.88752 | 3.79830 | 6.59405 |
| CD630_08510 | CD630_Cdi1_3 | 46 – 85 | 423 – 461 | -9.88666 | 5.16329 | 6.59405 |
| CD630_09760 | CD630_Cdi1_3 | 86 – 145 | 297 – 362 | -9.84131 | 4.83410 | 6.59405 |
| CD630_28910 | CD630_Cdi1_3 | 86 – 145 | 297 – 362 | -9.84131 | 4.51664 | 6.59405 |
| cobC | CD630_Cdi1_3 | 13 – 33 | 244 – 265 | -9.83639 | 3.15854 | 6.59405 |
| CD630_25620 | CD630_Cdi1_3 | 118 – 140 | 311 – 333 | -9.83547 | 3.02685 | 6.59405 |
| CD630_04940 | CD630_Cdi1_3 | 110 – 128 | 435 – 453 | -9.83226 | 4.20948 | 6.59405 |
| CD630_14700 | CD630_Cdi1_3 | 138 – 151 | 442 – 455 | -9.82992 | 3.98234 | 6.59405 |
| CD630_05450 | CD630_Cdi1_3 | 4 – 14 | 205 – 215 | -9.8194 | 5.04761 | 6.59405 |
| CD630_21230 | CD630_Cdi1_3 | 177 – 188 | 479 – 490 | -9.79391 | 3.89213 | 6.59405 |
| iorA | CD630_Cdi1_3 | 138 – 151 | 321 – 335 | -9.79011 | 3.83173 | 6.59405 |
| CD630_30620 | CD630_Cdi1_3 | 56 – 103 | 66 – 117 | -9.75651 | 4.15443 | 6.59405 |
| CD630_35840 | CD630_Cdi1_3 | 116 – 126 | 269 – 279 | -9.74885 | 3.14170 | 6.59405 |
| CD630_33560 | CD630_Cdi1_3 | 90 – 119 | 289 – 333 | -9.73717 | 4.03147 | 6.59405 |
| CD630_03420 | CD630_Cdi1_3 | 66 – 87 | 393 – 414 | -9.72803 | 4.71868 | 6.59405 |
| CD630_29250 | CD630_Cdi1_3 | 78 – 95 | 172 – 189 | -9.7113 | 4.61901 | 6.59405 |
| CD630_09410 | CD630_Cdi1_3 | 78 – 95 | 172 – 189 | -9.7113 | 4.75651 | 6.59405 |
| CD630_14150 | CD630_Cdi1_3 | 135 – 156 | 434 – 456 | -9.70771 | 3.28251 | 6.59405 |
| CD630_36250 | CD630_Cdi1_3 | 155 – 196 | 295 – 336 | -9.6988 | 4.13224 | 6.59405 |
| CD630_17292 | CD630_Cdi1_3 | 164 – 188 | 308 – 333 | -9.69239 | 6.25799 | 6.59405 |
| CD630_07470 | CD630_Cdi1_3 | 13 – 52 | 328 – 368 | -9.68586 | 3.37143 | 6.59405 |
| kdpE | CD630_Cdi1_3 | 119 – 135 | 443 – 459 | -9.64719 | 4.27140 | 6.59405 |
| CD630_01570 | CD630_Cdi1_3 | 7 – 23 | 330 – 346 | -9.64714 | 6.08699 | 6.59405 |
| CD630_03220 | CD630_Cdi1_3 | 31 – 44 | 442 – 455 | -9.63329 | 3.36746 | 6.59405 |
| CD630_22880 | CD630_Cdi1_3 | 104 – 123 | 72 – 89 | -9.60569 | 3.62455 | 6.59405 |
| CD630_07960 | CD630_Cdi1_3 | 90 – 108 | 84 – 101 | -9.58705 | 3.22537 | 6.59405 |
| acoL | CD630_Cdi1_3 | 131 – 145 | 320 – 333 | -9.55696 | 3.32169 | 6.59405 |
| CD630_05230 | CD630_Cdi1_3 | 3 – 59 | 314 – 359 | -9.55609 | 3.73787 | 6.59405 |
| CD630_23760 | CD630_Cdi1_3 | 56 – 69 | 129 – 142 | -9.54437 | 4.54307 | 6.59405 |
| CD630_08660 | CD630_Cdi1_3 | 157 – 172 | 385 – 400 | -9.53419 | 3.86467 | 6.59405 |
| CD630_09771 | CD630_Cdi1_4 | 52 – 201 | 350 – 499 | -191.788 | 4.95362 | 5.31318 |
| CD630_28890 | CD630_Cdi1_4 | 52 – 201 | 350 – 499 | -191.788 | 7.52469 | 5.31318 |
| CD630_17170 | CD630_Cdi1_4 | 158 – 177 | 447 – 466 | -19.3106 | 4.10230 | 5.31318 |
| srlD | CD630_Cdi1_4 | 125 – 149 | 441 – 465 | -18.9315 | 4.19843 | 5.31318 |
| cheW | CD630_Cdi1_4 | 85 – 105 | 446 – 467 | -18.7067 | 3.72371 | 5.31318 |
| CD630_30140 | CD630_Cdi1_4 | 84 – 110 | 442 – 465 | -18.609 | 3.28343 | 5.31318 |
| CD630_14410 | CD630_Cdi1_4 | 89 – 109 | 442 – 464 | -18.5063 | 3.39433 | 5.31318 |
| CD630_13920 | CD630_Cdi1_4 | 119 – 139 | 445 – 465 | -18.3788 | 3.25236 | 5.31318 |
| CD630_21490 | CD630_Cdi1_4 | 90 – 175 | 409 – 461 | -18.1678 | 3.37462 | 5.31318 |
| CD630_10220 | CD630_Cdi1_4 | 87 – 157 | 394 – 463 | -17.7394 | 4.08233 | 5.31318 |
| CD630_21610 | CD630_Cdi1_4 | 88 – 123 | 425 – 463 | -17.3975 | 3.16662 | 5.31318 |
| CD630_23780 | CD630_Cdi1_4 | 90 – 184 | 269 – 360 | -17.3747 | 3.91629 | 5.31318 |
| metQ | CD630_Cdi1_4 | 85 – 200 | 356 – 470 | -17.3663 | 3.87190 | 5.31318 |
| CD630_19680 | CD630_Cdi1_4 | 88 – 136 | 397 – 461 | -17.1587 | 3.58080 | 5.31318 |
| tcdA | CD630_Cdi1_4 | 36 – 94 | 429 – 470 | -16.8929 | 4.70593 | 5.31318 |
| CD630_06970 | CD630_Cdi1_4 | 84 – 128 | 427 – 467 | -16.8761 | 3.82312 | 5.31318 |
| CD630_35260 | CD630_Cdi1_4 | 90 – 104 | 447 – 461 | -16.7628 | 5.77277 | 5.31318 |
| CD630_27570 | CD630_Cdi1_4 | 102 – 140 | 429 – 471 | -16.657 | 3.46216 | 5.31318 |
| CD630_09570 | CD630_Cdi1_4 | 21 – 52 | 433 – 464 | -16.5741 | 5.31213 | 5.31318 |
| CD630_29070 | CD630_Cdi1_4 | 21 – 52 | 433 – 464 | -16.5741 | 3.89477 | 5.31318 |
| CD630_20450 | CD630_Cdi1_4 | 111 – 165 | 92 – 149 | -16.5042 | 3.77388 | 5.31318 |
| CD630_18120 | CD630_Cdi1_4 | 78 – 109 | 442 – 471 | -16.4526 | 3.99713 | 5.31318 |
| CD630_21570 | CD630_Cdi1_4 | 151 – 186 | 117 – 148 | -16.4301 | 6.48385 | 5.31318 |
| CD630_09990 | CD630_Cdi1_4 | 90 – 174 | 379 – 463 | -16.3863 | 3.86040 | 5.31318 |
| CD630_09562 | CD630_Cdi1_4 | 87 – 195 | 365 – 463 | -16.3732 | 3.94307 | 5.31318 |
| CD630_11241 | CD630_Cdi1_4 | 164 – 193 | 185 – 213 | -16.3643 | 4.85723 | 5.31318 |
| CD630_14700 | CD630_Cdi1_4 | 76 – 157 | 113 – 207 | -16.3357 | 3.98234 | 5.31318 |
| kdpE | CD630_Cdi1_4 | 91 – 106 | 445 – 462 | -16.3097 | 4.27140 | 5.31318 |
| CD630_04341 | CD630_Cdi1_4 | 159 – 188 | 433 – 461 | -16.2838 | 4.29262 | 5.31318 |
| CD630_26050 | CD630_Cdi1_4 | 169 – 179 | 450 – 460 | -16.1864 | 6.99851 | 5.31318 |
| xdhA | CD630_Cdi1_4 | 47 – 101 | 112 – 164 | -16.07 | 4.45147 | 5.31318 |
| CD630_26250 | CD630_Cdi1_4 | 86 – 157 | 384 – 464 | -16.0424 | 5.03379 | 5.31318 |
| CD630_06140 | CD630_Cdi1_4 | 86 – 123 | 428 – 460 | -16.0109 | 4.12519 | 5.31318 |
| CD630_05211 | CD630_Cdi1_4 | 112 – 153 | 219 – 267 | -16.0082 | 3.97048 | 5.31318 |
| CD630_17281 | CD630_Cdi1_4 | 50 – 94 | 113 – 158 | -15.9585 | 5.91544 | 5.31318 |
| CD630_18910 | CD630_Cdi1_4 | 118 – 138 | 435 – 460 | -15.9172 | 3.30688 | 5.31318 |
| oppC | CD630_Cdi1_4 | 143 – 193 | 106 – 157 | -15.8425 | 3.01695 | 5.31318 |
| CD630_32710 | CD630_Cdi1_4 | 89 – 185 | 380 – 458 | -15.718 | 4.74276 | 5.31318 |
| cbpA | CD630_Cdi1_4 | 84 – 95 | 458 – 469 | -15.6735 | 3.03040 | 5.31318 |
| CD630_18680 | CD630_Cdi1_4 | 86 – 111 | 432 – 458 | -15.5711 | 3.04315 | 5.31318 |
| cbiQ | CD630_Cdi1_4 | 1 – 89 | 378 – 458 | -15.5485 | 3.79830 | 5.31318 |
| CD630_19780 | CD630_Cdi1_4 | 14 – 32 | 443 – 461 | -15.5028 | 3.64830 | 5.31318 |
| terD | CD630_Cdi1_4 | 87 – 141 | 418 – 464 | -15.495 | 3.33148 | 5.31318 |
| CD630_04550 | CD630_Cdi1_4 | 89 – 137 | 308 – 360 | -15.471 | 3.05227 | 5.31318 |
| CD630_06060 | CD630_Cdi1_4 | 29 – 39 | 451 – 461 | -15.4466 | 4.94050 | 5.31318 |
| CD630_24590 | CD630_Cdi1_4 | 123 – 190 | 426 – 471 | -15.4071 | 3.70401 | 5.31318 |
| CD630_13960 | CD630_Cdi1_4 | 117 – 175 | 190 – 253 | -15.3818 | 3.30375 | 5.31318 |
| CD630_31530 | CD630_Cdi1_4 | 107 – 146 | 113 – 151 | -15.3302 | 3.49100 | 5.31318 |
| CD630_05830 | CD630_Cdi1_4 | 47 – 94 | 109 – 156 | -15.3056 | 3.35360 | 5.31318 |
| CD630_06080 | CD630_Cdi1_4 | 156 – 182 | 184 – 212 | -15.2631 | 3.82315 | 5.31318 |
| bglA | CD630_Cdi1_4 | 155 – 190 | 445 – 465 | -15.2409 | 4.86402 | 5.31318 |
| CD630_15360 | CD630_Cdi1_4 | 105 – 176 | 276 – 339 | -15.2203 | 4.47208 | 5.31318 |
| rbsC | CD630_Cdi1_4 | 34 – 63 | 442 – 466 | -15.1728 | 3.71650 | 5.31318 |
| CD630_07840 | CD630_Cdi1_4 | 109 – 163 | 112 – 160 | -15.1266 | 4.26761 | 5.31318 |
| CD630_14250 | CD630_Cdi1_4 | 83 – 104 | 447 – 465 | -15.1266 | 4.19604 | 5.31318 |
| CD630_05821 | CD630_Cdi1_4 | 80 – 111 | 333 – 364 | -15.1231 | 3.75661 | 5.31318 |
| CD630_19690 | CD630_Cdi1_4 | 109 – 126 | 443 – 456 | -15.0682 | 3.85861 | 5.31318 |
| CD630_02790 | CD630_Cdi1_4 | 127 – 170 | 328 – 363 | -15.0512 | 4.77661 | 5.31318 |
| CD630_10950 | CD630_Cdi1_4 | 85 – 130 | 436 – 466 | -15.024 | 3.67675 | 5.31318 |
| CD630_32700 | CD630_Cdi1_4 | 180 – 194 | 199 – 213 | -14.9589 | 4.74276 | 5.31318 |
| CD630_18410 | CD630_Cdi1_4 | 85 – 104 | 443 – 463 | -14.9138 | 3.59739 | 5.31318 |
| CD630_21020 | CD630_Cdi1_4 | 85 – 108 | 445 – 466 | -14.8727 | 4.28666 | 5.31318 |
| CD630_28181 | CD630_Cdi1_4 | 28 – 79 | 292 – 349 | -14.8648 | 3.91299 | 5.31318 |
| CD630_26750 | CD630_Cdi1_4 | 85 – 104 | 447 – 467 | -14.7497 | 3.60610 | 5.31318 |
| CD630_14920 | CD630_Cdi1_4 | 95 – 113 | 447 – 465 | -14.6638 | 4.35225 | 5.31318 |
| CD630_14420 | CD630_Cdi1_4 | 75 – 97 | 445 – 467 | -14.6189 | 3.99497 | 5.31318 |
| ldh | CD630_Cdi1_4 | 39 – 97 | 350 – 415 | -14.5657 | 4.35818 | 5.31318 |
| CD630_07480 | CD630_Cdi1_4 | 87 – 106 | 445 – 468 | -14.5358 | 5.18321 | 5.31318 |
| CD630_10050 | CD630_Cdi1_4 | 152 – 198 | 430 – 470 | -14.4302 | 3.93750 | 5.31318 |
| CD630_31000 | CD630_Cdi1_4 | 97 – 126 | 439 – 470 | -14.3885 | 3.70946 | 5.31318 |
| CD630_21620 | CD630_Cdi1_4 | 4 – 38 | 117 – 148 | -14.3735 | 3.02649 | 5.31318 |
| cprK | CD630_Cdi1_4 | 84 – 112 | 426 – 457 | -14.343 | 3.96216 | 5.31318 |
| CD630_02720 | CD630_Cdi1_4 | 1 – 50 | 310 – 354 | -14.3415 | 3.38938 | 5.31318 |
| CD630_28550 | CD630_Cdi1_4 | 78 – 109 | 428 – 458 | -14.3095 | 3.51196 | 5.31318 |
| CD630_29250 | CD630_Cdi1_4 | 87 – 106 | 445 – 464 | -14.2892 | 4.61901 | 5.31318 |
| CD630_09410 | CD630_Cdi1_4 | 87 – 106 | 445 – 464 | -14.2892 | 4.75651 | 5.31318 |
| CD630_05480 | CD630_Cdi1_4 | 118 – 181 | 387 – 454 | -14.2885 | 4.27854 | 5.31318 |
| CD630_16550 | CD630_Cdi1_4 | 79 – 108 | 442 – 472 | -14.2719 | 5.61064 | 5.31318 |
| CD630_16150 | CD630_Cdi1_4 | 83 – 105 | 446 – 467 | -14.2455 | 4.20614 | 5.31318 |
| CD630_19580 | CD630_Cdi1_4 | 52 – 65 | 429 – 442 | -14.1506 | 4.51902 | 5.31318 |
| CD630_14630 | CD630_Cdi1_4 | 139 – 176 | 429 – 460 | -14.0856 | 3.34485 | 5.31318 |
| pilD | CD630_Cdi1_4 | 3 – 31 | 111 – 138 | -14.0685 | 4.54123 | 5.31318 |
| CD630_20290 | CD630_Cdi1_4 | 80 – 180 | 379 – 472 | -14.0337 | 4.38183 | 5.31318 |
| CD630_16670 | CD630_Cdi1_4 | 122 – 137 | 349 – 365 | -14.0304 | 3.33553 | 5.31318 |
| CD630_28150 | CD630_Cdi1_4 | 25 – 74 | 311 – 360 | -14.011 | 4.18447 | 5.31318 |
| gabT | CD630_Cdi1_4 | 90 – 114 | 443 – 461 | -14.0108 | 3.67067 | 5.31318 |
| CD630_21260 | CD630_Cdi1_4 | 138 – 152 | 44 – 58 | -13.9821 | 4.22255 | 5.31318 |
| CD630_01640 | CD630_Cdi1_4 | 78 – 104 | 447 – 474 | -13.9626 | 4.55381 | 5.31318 |
| tcdC | CD630_Cdi1_4 | 112 – 158 | 281 – 321 | -13.9354 | 3.76101 | 5.31318 |
| CD630_30540 | CD630_Cdi1_4 | 116 – 136 | 306 – 326 | -13.9036 | 3.63804 | 5.31318 |
| CD630_19790 | CD630_Cdi1_4 | 85 – 125 | 181 – 216 | -13.8585 | 3.79593 | 5.31318 |
| CD630_35430 | CD630_Cdi1_4 | 142 – 182 | 121 – 157 | -13.8354 | 4.28798 | 5.31318 |
| CD630_33480 | CD630_Cdi1_4 | 86 – 106 | 445 – 464 | -13.8037 | 3.67613 | 5.31318 |
| CD630_04280 | CD630_Cdi1_4 | 82 – 109 | 442 – 464 | -13.7851 | 3.61457 | 5.31318 |
| CD630_14150 | CD630_Cdi1_4 | 36 – 184 | 283 – 414 | -13.7812 | 3.28251 | 5.31318 |
| CD630_17292 | CD630_Cdi1_4 | 82 – 105 | 430 – 456 | -13.7807 | 6.25799 | 5.31318 |
| CD630_17160 | CD630_Cdi1_4 | 88 – 105 | 442 – 457 | -13.7797 | 3.50228 | 5.31318 |
| CD630_36250 | CD630_Cdi1_4 | 170 – 199 | 186 – 213 | -13.7561 | 4.13224 | 5.31318 |
| CD630_21460 | CD630_Cdi1_4 | 72 – 79 | 356 – 363 | -13.752 | 4.83005 | 5.31318 |
| serS1 | CD630_Cdi1_4 | 164 – 201 | 326 – 363 | -13.7452 | 3.45858 | 5.31318 |
| CD630_05290 | CD630_Cdi1_4 | 19 – 63 | 113 – 161 | -13.7324 | 3.97604 | 5.31318 |
| CD630_26680 | CD630_Cdi1_4 | 75 – 104 | 194 – 223 | -13.7243 | 3.53956 | 5.31318 |
| CD630_15080 | CD630_Cdi1_4 | 85 – 104 | 447 – 466 | -13.6735 | 4.03033 | 5.31318 |
| CD630_04540 | CD630_Cdi1_4 | 82 – 93 | 460 – 471 | -13.6728 | 3.33556 | 5.31318 |
| CD630_28320 | CD630_Cdi1_4 | 78 – 93 | 457 – 471 | -13.6719 | 3.99184 | 5.31318 |
| CD630_23440 | CD630_Cdi1_4 | 2 – 17 | 449 – 463 | -13.635 | 3.34841 | 5.31318 |
| CD630_27240 | CD630_Cdi1_4 | 2 – 11 | 445 – 454 | -13.6339 | 3.52462 | 5.31318 |
| flgB | CD630_Cdi1_4 | 89 – 101 | 444 – 456 | -13.5754 | 3.25736 | 5.31318 |
| acoL | CD630_Cdi1_4 | 104 – 143 | 347 – 386 | -13.5731 | 3.32169 | 5.31318 |
| cdu2 | CD630_Cdi1_4 | 82 – 139 | 427 – 470 | -13.5493 | 2.97335 | 5.31318 |
| CD630_21750 | CD630_Cdi1_4 | 59 – 106 | 277 – 330 | -13.5103 | 3.70626 | 5.31318 |
| CD630_11240 | CD630_Cdi1_4 | 83 – 108 | 443 – 466 | -13.4824 | 3.20274 | 5.31318 |
| etfA | CD630_Cdi1_4 | 88 – 157 | 300 – 361 | -13.456 | 3.41204 | 5.31318 |
| CD630_22890 | CD630_Cdi1_4 | 31 – 48 | 442 – 458 | -13.4509 | 3.75591 | 5.31318 |
| CD630_19000 | CD630_Cdi1_4 | 83 – 116 | 184 – 216 | -13.4366 | 4.18443 | 5.31318 |
| dapB2 | CD630_Cdi1_4 | 130 – 162 | 426 – 457 | -13.4155 | 3.27218 | 5.31318 |
| feoB | CD630_Cdi1_4 | 131 – 197 | 87 – 148 | -13.4125 | 4.05364 | 5.31318 |
| CD630_14190 | CD630_Cdi1_4 | 82 – 106 | 445 – 470 | -13.2987 | 3.78562 | 5.31318 |
| CD630_06340 | CD630_Cdi1_4 | 90 – 135 | 278 – 321 | -13.276 | 4.35983 | 5.31318 |
| CD630_03420 | CD630_Cdi1_4 | 138 – 175 | 113 – 154 | -13.2503 | 4.71868 | 5.31318 |
| CD630_14130 | CD630_Cdi1_4 | 87 – 179 | 377 – 470 | -13.1931 | 4.57970 | 5.31318 |
| CD630_13360 | CD630_Cdi1_4 | 89 – 139 | 430 – 462 | -13.1889 | 3.77330 | 5.31318 |
| cspD | CD630_Cdi1_4 | 160 – 184 | 434 – 462 | -13.1851 | 3.23587 | 5.31318 |
| CD630_16450 | CD630_Cdi1_4 | 88 – 112 | 438 – 462 | -13.1762 | 4.57547 | 5.31318 |
| CD630_24770 | CD630_Cdi1_4 | 87 – 100 | 349 – 363 | -13.1497 | 3.20798 | 5.31318 |
| CD630_15850 | CD630_Cdi1_4 | 12 – 39 | 445 – 470 | -13.1241 | 4.32968 | 5.31318 |
| CD630_30310 | CD630_Cdi1_4 | 70 – 93 | 429 – 451 | -13.1214 | 4.56059 | 5.31318 |
| CD630_28560 | CD630_Cdi1_4 | 64 – 92 | 429 – 455 | -13.0952 | 3.96803 | 5.31318 |
| CD630_05490 | CD630_Cdi1_4 | 155 – 184 | 430 – 456 | -13.0871 | 3.89852 | 5.31318 |
| CD630_22200 | CD630_Cdi1_4 | 141 – 173 | 115 – 148 | -13.0054 | 4.18288 | 5.31318 |
| CD630_31490 | CD630_Cdi1_4 | 90 – 122 | 429 – 463 | -13.0007 | 4.08593 | 5.31318 |
| CD630_23000 | CD630_Cdi1_4 | 22 – 33 | 429 – 440 | -12.9994 | 3.54130 | 5.31318 |
| CD630_27610 | CD630_Cdi1_4 | 5 – 52 | 278 – 323 | -12.952 | 3.96465 | 5.31318 |
| CD630_21770 | CD630_Cdi1_4 | 87 – 196 | 111 – 212 | -12.9463 | 4.02587 | 5.31318 |
| CD630_08360 | CD630_Cdi1_4 | 161 – 176 | 443 – 458 | -12.9089 | 3.19339 | 5.31318 |
| eutV | CD630_Cdi1_4 | 90 – 109 | 450 – 468 | -12.8995 | 4.55715 | 5.31318 |

| | | | | | | |
|---|---|---|---|---|---|---|
| scrR | CD630_Cdi1_4 | 134 -- 148 | 351 -- 365 | -12.8559 | 3.49166 | 5.31318 |
| bcp | CD630_Cdi1_4 | 44 -- 97 | 109 -- 164 | -12.8175 | 3.16670 | 5.31318 |
| CD630_05280 | CD630_Cdi1_4 | 153 -- 186 | 180 -- 213 | -12.7883 | 3.91665 | 5.31318 |
| CD630_33300 | CD630_Cdi1_4 | 87 -- 105 | 444 -- 462 | -12.7506 | 3.79149 | 5.31318 |
| sspA | CD630_Cdi1_4 | 53 -- 109 | 95 -- 156 | -12.7503 | 3.62689 | 5.31318 |
| thiH | CD630_Cdi1_4 | 169 -- 196 | 271 -- 295 | -12.7222 | 3.58230 | 5.31318 |
| CD630_25190 | CD630_Cdi1_4 | 91 -- 139 | 111 -- 164 | -12.7137 | 3.19663 | 5.31318 |
| CD630_05590 | CD630_Cdi1_4 | 90 -- 135 | 427 -- 467 | -12.7063 | 3.39670 | 5.31318 |
| oppB | CD630_Cdi1_4 | 88 -- 99 | 453 -- 463 | -12.6972 | 2.98328 | 5.31318 |
| CD630_01590 | CD630_Cdi1_4 | 167 -- 201 | 184 -- 218 | -12.6501 | 3.45609 | 5.31318 |
| CD630_34530 | CD630_Cdi1_4 | 2 -- 50 | 115 -- 165 | -12.6264 | 3.83016 | 5.31318 |
| CD630_11220 | CD630_Cdi1_4 | 136 -- 160 | 445 -- 468 | -12.6171 | 3.52327 | 5.31318 |
| CD630_17960 | CD630_Cdi1_4 | 126 -- 147 | 452 -- 470 | -12.6142 | 3.71404 | 5.31318 |
| CD630_20170 | CD630_Cdi1_4 | 123 -- 164 | 113 -- 149 | -12.5997 | 6.90201 | 5.31318 |
| CD630_18960 | CD630_Cdi1_4 | 135 -- 153 | 143 -- 161 | -12.5983 | 4.37822 | 5.31318 |
| agrB | CD630_Cdi1_4 | 17 -- 61 | 378 -- 418 | -12.568 | 3.92598 | 5.31318 |
| CD630_03220 | CD630_Cdi1_4 | 29 -- 70 | 428 -- 459 | -12.556 | 3.36746 | 5.31318 |
| CD630_06200 | CD630_Cdi1_4 | 118 -- 150 | 111 -- 151 | -12.5113 | 3.57064 | 5.31318 |
| CD630_02420 | CD630_Cdi1_4 | 18 -- 40 | 446 -- 466 | -12.4979 | 3.71048 | 5.31318 |
| CD630_03040 | CD630_Cdi1_4 | 51 -- 95 | 112 -- 159 | -12.4872 | 3.51403 | 5.31318 |
| nusB | CD630_Cdi1_4 | 88 -- 157 | 288 -- 360 | -12.4755 | 4.92798 | 5.31318 |
| gcvTPA | CD630_Cdi1_4 | 87 -- 111 | 440 -- 464 | -12.4456 | 4.77229 | 5.31318 |
| CD630_13780 | CD630_Cdi1_4 | 130 -- 176 | 181 -- 216 | -12.4444 | 3.52274 | 5.31318 |
| CD630_31910 | CD630_Cdi1_4 | 84 -- 121 | 430 -- 464 | -12.4371 | 3.48314 | 5.31318 |
| CD630_23280 | CD630_Cdi1_4 | 99 -- 134 | 429 -- 468 | -12.4266 | 3.50093 | 5.31318 |
| CD630_29270 | CD630_Cdi1_4 | 26 -- 57 | 452 -- 478 | -12.3483 | 3.70174 | 5.31318 |
| ung | CD630_Cdi1_4 | 1 -- 38 | 109 -- 148 | -12.3454 | 3.30908 | 5.31318 |
| CD630_08100 | CD630_Cdi1_4 | 180 -- 198 | 445 -- 460 | -12.3438 | 4.39181 | 5.31318 |
| CD630_27110 | CD630_Cdi1_4 | 133 -- 178 | 106 -- 151 | -12.3348 | 3.78231 | 5.31318 |
| CD630_16160 | CD630_Cdi1_4 | 87 -- 111 | 439 -- 466 | -12.3077 | 6.11801 | 5.31318 |
| CD630_30100 | CD630_Cdi1_4 | 54 -- 80 | 182 -- 209 | -12.2928 | 4.13234 | 5.31318 |
| CD630_14810 | CD630_Cdi1_4 | 144 -- 165 | 435 -- 456 | -12.2108 | 3.35323 | 5.31318 |
| CD630_28290 | CD630_Cdi1_4 | 86 -- 94 | 456 -- 464 | -12.2034 | 5.54192 | 5.31318 |
| CD630_22050 | CD630_Cdi1_4 | 82 -- 104 | 447 -- 471 | -12.1779 | 4.87639 | 5.31318 |
| CD630_14260 | CD630_Cdi1_4 | 96 -- 109 | 445 -- 458 | -12.1745 | 3.22214 | 5.31318 |
| bglA | CD630_Cdi1_4 | 86 -- 112 | 430 -- 465 | -12.1745 | 4.50315 | 5.31318 |
| CD630_18950 | CD630_Cdi1_4 | 180 -- 194 | 86 -- 99 | -12.1735 | 3.40610 | 5.31318 |
| CD630_11730 | CD630_Cdi1_4 | 88 -- 129 | 310 -- 361 | -12.0796 | 4.80521 | 5.31318 |
| CD630_22550 | CD630_Cdi1_4 | 84 -- 104 | 447 -- 471 | -12.072 | 3.62379 | 5.31318 |
| CD630_32050 | CD630_Cdi1_4 | 176 -- 199 | 191 -- 213 | -12.0678 | 5.73797 | 5.31318 |
| CD630_15550 | CD630_Cdi1_4 | 88 -- 105 | 446 -- 464 | -12.0643 | 3.49019 | 5.31318 |
| CD630_09030 | CD630_Cdi1_4 | 148 -- 197 | 314 -- 364 | -12.0354 | 4.21282 | 5.31318 |
| CD630_14360 | CD630_Cdi1_4 | 86 -- 106 | 445 -- 464 | -12.0294 | 3.02079 | 5.31318 |
| CD630_31580 | CD630_Cdi1_4 | 87 -- 108 | 439 -- 459 | -12.0195 | 3.51756 | 5.31318 |
| CD630_15740 | CD630_Cdi1_4 | 48 -- 93 | 429 -- 457 | -12.0141 | 4.43222 | 5.31318 |
| CD630_23820 | CD630_Cdi1_4 | 88 -- 106 | 345 -- 361 | -11.9908 | 3.18419 | 5.31318 |
| CD630_21090 | CD630_Cdi1_4 | 131 -- 188 | 112 -- 161 | -11.9649 | 4.15803 | 5.31318 |
| CD630_21540 | CD630_Cdi1_4 | 77 -- 106 | 445 -- 470 | -11.919 | 4.33090 | 5.31318 |
| CD630_17730 | CD630_Cdi1_4 | 87 -- 109 | 442 -- 460 | -11.9061 | 4.73397 | 5.31318 |
| def1 | CD630_Cdi1_4 | 86 -- 179 | 305 -- 362 | -11.8806 | 3.99720 | 5.31318 |
| CD630_20171 | CD630_Cdi1_4 | 151 -- 179 | 427 -- 454 | -11.8669 | 6.87491 | 5.31318 |
| CD630_15270 | CD630_Cdi1_4 | 89 -- 111 | 440 -- 462 | -11.8478 | 4.66015 | 5.31318 |
| CD630_32520 | CD630_Cdi1_4 | 138 -- 164 | 291 -- 321 | -11.8117 | 4.06098 | 5.31318 |
| CD630_28260 | CD630_Cdi1_4 | 66 -- 104 | 426 -- 469 | -11.7917 | 3.51613 | 5.31318 |
| CD630_21910 | CD630_Cdi1_4 | 85 -- 96 | 460 -- 471 | -11.7881 | 4.66600 | 5.31318 |
| CD630_20460 | CD630_Cdi1_4 | 98 -- 112 | 448 -- 461 | -11.7875 | 7.94034 | 5.31318 |
| CD630_18650 | CD630_Cdi1_4 | 167 -- 191 | 297 -- 320 | -11.7812 | 3.75892 | 5.31318 |
| srlEb | CD630_Cdi1_4 | 79 -- 177 | 379 -- 468 | -11.7758 | 3.50140 | 5.31318 |
| CD630_33600 | CD630_Cdi1_4 | 139 -- 189 | 272 -- 325 | -11.7698 | 3.73196 | 5.31318 |
| CD630_29650 | CD630_Cdi1_4 | 87 -- 115 | 449 -- 468 | -11.7602 | 3.67128 | 5.31318 |
| CD630_07870 | CD630_Cdi1_4 | 90 -- 109 | 442 -- 463 | -11.7549 | 3.22829 | 5.31318 |
| CD630_28580 | CD630_Cdi1_4 | 48 -- 64 | 344 -- 363 | -11.7196 | 4.67130 | 5.31318 |
| ogt1 | CD630_Cdi1_4 | 63 -- 93 | 429 -- 453 | -11.7016 | 3.16772 | 5.31318 |
| dnaC | CD630_Cdi1_4 | 145 -- 175 | 185 -- 213 | -11.6399 | 6.30796 | 5.31318 |
| CD630_15900 | CD630_Cdi1_4 | 112 -- 127 | 378 -- 393 | -11.6288 | 3.17664 | 5.31318 |
| pgm | CD630_Cdi1_4 | 145 -- 176 | 184 -- 217 | -11.6275 | 3.45439 | 5.31318 |
| vncR | CD630_Cdi1_4 | 138 -- 157 | 190 -- 213 | -11.6078 | 3.93642 | 5.31318 |
| CD630_31010 | CD630_Cdi1_4 | 41 -- 98 | 109 -- 164 | -11.6009 | 4.96305 | 5.31318 |
| CD630_32890 | CD630_Cdi1_4 | 85 -- 118 | 430 -- 465 | -11.5913 | 3.41276 | 5.31318 |
| ribD | CD630_Cdi1_4 | 85 -- 109 | 438 -- 463 | -11.5819 | 6.65922 | 5.31318 |
| CD630_09772 | CD630_Cdi1_4 | 39 -- 96 | 111 -- 165 | -11.5683 | 3.66033 | 5.31318 |
| CD630_18311 | CD630_Cdi1_4 | 88 -- 113 | 181 -- 210 | -11.5657 | 6.86047 | 5.31318 |
| CD630_23760 | CD630_Cdi1_4 | 82 -- 105 | 193 -- 216 | -11.5604 | 4.54307 | 5.31318 |
| CD630_14340 | CD630_Cdi1_4 | 98 -- 111 | 440 -- 453 | -11.5516 | 5.10868 | 5.31318 |
| CD630_05870 | CD630_Cdi1_4 | 87 -- 106 | 445 -- 464 | -11.5509 | 4.28339 | 5.31318 |
| cspC | CD630_Cdi1_4 | 160 -- 184 | 434 -- 462 | -11.5479 | 5.48848 | 5.31318 |
| CD630_20260 | CD630_Cdi1_4 | 86 -- 114 | 183 -- 212 | -11.5295 | 3.84162 | 5.31318 |
| CD630_17710 | CD630_Cdi1_4 | 87 -- 108 | 443 -- 464 | -11.5276 | 4.37688 | 5.31318 |
| CD630_31461 | CD630_Cdi1_4 | 6 -- 25 | 139 -- 156 | -11.5276 | 4.41476 | 5.31318 |
| CD630_09040 | CD630_Cdi1_4 | 91 -- 139 | 111 -- 164 | -11.5059 | 4.11851 | 5.31318 |
| CD630_23770 | CD630_Cdi1_4 | 2 -- 31 | 313 -- 341 | -11.502 | 3.25181 | 5.31318 |
| CD630_18610 | CD630_Cdi1_4 | 85 -- 106 | 445 -- 464 | -11.4999 | 4.15582 | 5.31318 |
| LinCd | CD630_Cdi1_4 | 2 -- 94 | 308 -- 407 | -11.4882 | 4.33624 | 5.31318 |
| cwp29 | CD630_Cdi1_4 | 29 -- 43 | 450 -- 464 | -11.462 | 4.40865 | 5.31318 |
| CD630_20670 | CD630_Cdi1_4 | 74 -- 112 | 184 -- 226 | -11.4298 | 4.42757 | 5.31318 |
| CD630_15431 | CD630_Cdi1_4 | 125 -- 185 | 109 -- 164 | -11.4213 | 5.29780 | 5.31318 |
| CD630_34030 | CD630_Cdi1_4 | 42 -- 66 | 134 -- 161 | -11.4158 | 3.05766 | 5.31318 |
| CD630_22030 | CD630_Cdi1_4 | 7 -- 23 | 51 -- 67 | -11.402 | 3.29934 | 5.31318 |
| CD630_33560 | CD630_Cdi1_4 | 57 -- 94 | 113 -- 149 | -11.3923 | 4.03147 | 5.31318 |
| CD630_10000 | CD630_Cdi1_4 | 52 -- 88 | 429 -- 455 | -11.3689 | 3.86040 | 5.31318 |
| CD630_22880 | CD630_Cdi1_4 | 175 -- 199 | 132 -- 157 | -11.3654 | 3.62455 | 5.31318 |
| CD630_20660 | CD630_Cdi1_4 | 134 -- 167 | 115 -- 157 | -11.3333 | 5.39321 | 5.31318 |
| CD630_36040 | CD630_Cdi1_4 | 64 -- 125 | 427 -- 468 | -11.3289 | 3.52009 | 5.31318 |
| glgB | CD630_Cdi1_4 | 87 -- 112 | 442 -- 459 | -11.3237 | 4.48273 | 5.31318 |
| CD630_29960 | CD630_Cdi1_4 | 135 -- 201 | 72 -- 145 | -11.316 | 4.51904 | 5.31318 |
| CD630_12110 | CD630_Cdi1_4 | 53 -- 61 | 456 -- 464 | -11.3134 | 4.36078 | 5.31318 |
| CD630_32690 | CD630_Cdi1_4 | 3 -- 34 | 433 -- 465 | -11.2935 | 7.32676 | 5.31318 |
| cobC | CD630_Cdi1_4 | 164 -- 198 | 111 -- 148 | -11.2858 | 3.15854 | 5.31318 |
| CD630_22850 | CD630_Cdi1_4 | 167 -- 201 | 112 -- 156 | -11.2854 | 3.84102 | 5.31318 |
| CD630_17970 | CD630_Cdi1_4 | 164 -- 174 | 443 -- 453 | -11.2792 | 3.68802 | 5.31318 |
| CD630_24580 | CD630_Cdi1_4 | 85 -- 141 | 410 -- 462 | -11.2602 | 3.88519 | 5.31318 |
| uxaA | CD630_Cdi1_4 | 72 -- 91 | 429 -- 458 | -11.2559 | 4.26165 | 5.31318 |
| ssuC | CD630_Cdi1_4 | 61 -- 95 | 428 -- 470 | -11.2517 | 3.45491 | 5.31318 |
| CD630_18090 | CD630_Cdi1_4 | 10 -- 32 | 191 -- 212 | -11.2492 | 4.23961 | 5.31318 |
| CD630_16950 | CD630_Cdi1_4 | 185 -- 196 | 88 -- 100 | -11.2437 | 4.42569 | 5.31318 |
| CD630_33580 | CD630_Cdi1_4 | 91 -- 99 | 456 -- 464 | -11.2064 | 3.95778 | 5.31318 |
| CD630_10960 | CD630_Cdi1_4 | 87 -- 131 | 430 -- 463 | -11.1677 | 3.53850 | 5.31318 |
| CD630_10880 | CD630_Cdi1_4 | 86 -- 105 | 430 -- 457 | -11.1546 | 5.05048 | 5.31318 |
| glnA | CD630_Cdi1_4 | 151 -- 187 | 115 -- 154 | -11.1194 | 3.87629 | 5.31318 |
| CD630_02740 | CD630_Cdi1_4 | 88 -- 104 | 447 -- 463 | -11.1065 | 5.02277 | 5.31318 |
| CD630_13400 | CD630_Cdi1_4 | 67 -- 118 | 112 -- 157 | -11.1024 | 3.20232 | 5.31318 |
| CD630_15150 | CD630_Cdi1_4 | 8 -- 52 | 110 -- 163 | -11.0672 | 3.35564 | 5.31318 |
| CD630_09970 | CD630_Cdi1_4 | 3 -- 39 | 110 -- 147 | -11.0498 | 3.25917 | 5.31318 |
| CD630_14710 | CD630_Cdi1_4 | 87 -- 97 | 452 -- 463 | -11.0441 | 3.65171 | 5.31318 |
| spoVFB | CD630_Cdi1_4 | 83 -- 129 | 428 -- 471 | -11.0393 | 4.62758 | 5.31318 |
| CD630_27200 | CD630_Cdi1_4 | 1 -- 36 | 278 -- 302 | -11.0259 | 3.42516 | 5.31318 |
| CD630_12320 | CD630_Cdi1_4 | 37 -- 80 | 315 -- 355 | -11.0037 | 3.28667 | 5.31318 |
| glgP | CD630_Cdi1_4 | 71 -- 115 | 427 -- 463 | -10.9776 | 3.28530 | 5.31318 |
| CD630_16460 | CD630_Cdi1_4 | 15 -- 86 | 172 -- 251 | -10.9693 | 4.64028 | 5.31318 |
| CD630_04610 | CD630_Cdi1_4 | 48 -- 71 | 115 -- 146 | -10.9688 | 3.83025 | 5.31318 |
| CD630_17151 | CD630_Cdi1_4 | 166 -- 197 | 428 -- 454 | -10.9381 | 6.43124 | 5.31318 |
| adhE | CD630_Cdi1_4 | 35 -- 82 | 315 -- 361 | -10.9332 | 3.06485 | 5.31318 |
| opuCA | CD630_Cdi1_4 | 87 -- 112 | 182 -- 213 | -10.9324 | 4.07035 | 5.31318 |
| CD630_21080 | CD630_Cdi1_4 | 89 -- 105 | 442 -- 457 | -10.8993 | 3.99742 | 5.31318 |
| CD630_13880 | CD630_Cdi1_4 | 41 -- 55 | 446 -- 460 | -10.8978 | 3.52735 | 5.31318 |
| CD630_21070 | CD630_Cdi1_4 | 185 -- 196 | 351 -- 362 | -10.8869 | 5.35019 | 5.31318 |
| CD630_34080 | CD630_Cdi1_4 | 83 -- 106 | 430 -- 454 | -10.8776 | 3.20224 | 5.31318 |
| CD630_31220 | CD630_Cdi1_4 | 85 -- 105 | 430 -- 454 | -10.8748 | 3.73779 | 5.31318 |
| CD630_30130 | CD630_Cdi1_4 | 1 -- 28 | 182 -- 217 | -10.8571 | 3.28343 | 5.31318 |
| CD630_06150 | CD630_Cdi1_4 | 99 -- 128 | 110 -- 131 | -10.8386 | 5.51744 | 5.31318 |
| CD630_13350 | CD630_Cdi1_4 | 163 -- 181 | 134 -- 149 | -10.838 | 4.15139 | 5.31318 |
| appF | CD630_Cdi1_4 | 84 -- 129 | 428 -- 470 | -10.8236 | 4.25570 | 5.31318 |
| CD630_36781 | CD630_Cdi1_4 | 20 -- 38 | 347 -- 364 | -10.8226 | 3.75946 | 5.31318 |
| folC | CD630_Cdi1_4 | 87 -- 125 | 428 -- 465 | -10.8174 | 3.85634 | 5.31318 |
| CD630_24760 | CD630_Cdi1_4 | 98 -- 143 | 111 -- 161 | -10.808 | 5.19485 | 5.31318 |
| cls | CD630_Cdi1_4 | 141 -- 173 | 384 -- 416 | -10.8076 | 4.46177 | 5.31318 |
| CD630_05820 | CD630_Cdi1_4 | 87 -- 93 | 456 -- 462 | -10.8038 | 3.75661 | 5.31318 |
| CD630_04660 | CD630_Cdi1_4 | 91 -- 105 | 442 -- 456 | -10.8002 | 4.32791 | 5.31318 |
| CD630_01690 | CD630_Cdi1_4 | 14 -- 45 | 120 -- 151 | -10.7811 | 4.97234 | 5.31318 |
| uxaA%27 | CD630_Cdi1_4 | 86 -- 108 | 445 -- 464 | -10.7742 | 4.26165 | 5.31318 |
| CD630_34560 | CD630_Cdi1_4 | 84 -- 115 | 439 -- 468 | -10.7479 | 4.02604 | 5.31318 |
| CD630_19401 | CD630_Cdi1_4 | 35 -- 82 | 125 -- 172 | -10.7356 | 4.37084 | 5.31318 |
| CD630_35780 | CD630_Cdi1_4 | 16 -- 67 | 278 -- 337 | -10.6864 | 3.50107 | 5.31318 |
| CD630_25620 | CD630_Cdi1_4 | 88 -- 107 | 444 -- 464 | -10.6782 | 3.02685 | 5.31318 |
| CD630_06920 | CD630_Cdi1_4 | 83 -- 125 | 427 -- 465 | -10.6675 | 3.13260 | 5.31318 |
| CD630_02430 | CD630_Cdi1_4 | 39 -- 84 | 117 -- 145 | -10.6616 | 3.71048 | 5.31318 |
| garK | CD630_Cdi1_4 | 123 -- 178 | 429 -- 477 | -10.6569 | 3.72836 | 5.31318 |
| CD630_09060 | CD630_Cdi1_4 | 2 -- 23 | 379 -- 400 | -10.6229 | 4.31834 | 5.31318 |
| CD630_25040 | CD630_Cdi1_4 | 144 -- 177 | 26 -- 61 | -10.607 | 3.43898 | 5.31318 |
| CD630_09100 | CD630_Cdi1_4 | 2 -- 50 | 27 -- 68 | -10.6054 | 5.92260 | 5.31318 |
| CD630_16530 | CD630_Cdi1_4 | 87 -- 166 | 394 -- 467 | -10.5713 | 4.37658 | 5.31318 |
| CD630_01560 | CD630_Cdi1_4 | 13 -- 61 | 175 -- 218 | -10.5705 | 4.06352 | 5.31318 |
| CD630_04880 | CD630_Cdi1_4 | 125 -- 152 | 343 -- 362 | -10.5632 | 4.62291 | 5.31318 |
| CD630_33420 | CD630_Cdi1_4 | 88 -- 105 | 442 -- 458 | -10.5481 | 3.22095 | 5.31318 |
| blaI | CD630_Cdi1_4 | 87 -- 104 | 447 -- 465 | -10.5428 | 3.73674 | 5.31318 |
| CD630_09760 | CD630_Cdi1_4 | 90 -- 125 | 180 -- 213 | -10.5355 | 4.83410 | 5.31318 |
| CD630_28910 | CD630_Cdi1_4 | 90 -- 125 | 180 -- 213 | -10.5355 | 4.51664 | 5.31318 |
| CD630_30120 | CD630_Cdi1_4 | 43 -- 100 | 111 -- 164 | -10.5294 | 4.04859 | 5.31318 |
| CD630_25610 | CD630_Cdi1_4 | 44 -- 94 | 279 -- 320 | -10.5293 | 3.02685 | 5.31318 |
| CD630_08940 | CD630_Cdi1_4 | 75 -- 91 | 74 -- 90 | -10.5078 | 3.72225 | 5.31318 |
| CD630_27310 | CD630_Cdi1_4 | 76 -- 94 | 451 -- 468 | -10.5054 | 4.01622 | 5.31318 |
| CD630_17360 | CD630_Cdi1_4 | 85 -- 122 | 429 -- 464 | -10.4998 | 5.66070 | 5.31318 |
| CD630_02830 | CD630_Cdi1_4 | 182 -- 198 | 300 -- 317 | -10.4833 | 3.57106 | 5.31318 |
| CD630_09850 | CD630_Cdi1_4 | 87 -- 104 | 447 -- 464 | -10.4805 | 4.18679 | 5.31318 |
| CD630_21760 | CD630_Cdi1_4 | 64 -- 74 | 89 -- 98 | -10.4599 | 4.43846 | 5.31318 |
| CD630_18010 | CD630_Cdi1_4 | 154 -- 186 | 132 -- 163 | -10.4453 | 4.21967 | 5.31318 |
| CD630_03930 | CD630_Cdi1_4 | 87 -- 113 | 447 -- 468 | -10.4291 | 4.15934 | 5.31318 |
| CD630_19030 | CD630_Cdi1_4 | 105 -- 113 | 456 -- 464 | -10.4229 | 3.34240 | 5.31318 |
| CD630_14050 | CD630_Cdi1_4 | 186 -- 201 | 198 -- 213 | -10.4163 | 3.30894 | 5.31318 |
| CD630_08090 | CD630_Cdi1_4 | 143 -- 174 | 109 -- 139 | -10.4078 | 3.57167 | 5.31318 |
| CD630_06690 | CD630_Cdi1_4 | 86 -- 96 | 350 -- 360 | -10.3871 | 5.82709 | 5.31318 |
| CD630_27270 | CD630_Cdi1_4 | 89 -- 118 | 442 -- 462 | -10.3495 | 3.50892 | 5.31318 |
| CD630_08390 | CD630_Cdi1_4 | 81 -- 101 | 430 -- 450 | -10.3304 | 5.16000 | 5.31318 |
| CD630_31880 | CD630_Cdi1_4 | 15 -- 34 | 22 -- 42 | -10.3237 | 3.97522 | 5.31318 |
| CD630_06190 | CD630_Cdi1_4 | 141 -- 151 | 48 -- 58 | -10.2851 | 3.57064 | 5.31318 |
| CD630_18920 | CD630_Cdi1_4 | 170 -- 194 | 445 -- 466 | -10.2733 | 4.43558 | 5.31318 |
| CD630_33190 | CD630_Cdi1_4 | 115 -- 126 | 184 -- 195 | -10.2718 | 4.13899 | 5.31318 |
| opuCC | CD630_Cdi1_4 | 95 -- 144 | 115 -- 164 | -10.2683 | 4.07035 | 5.31318 |
| CD630_27220 | CD630_Cdi1_4 | 86 -- 112 | 446 -- 470 | -10.2665 | 4.18786 | 5.31318 |
| feoB | CD630_Cdi1_4 | 85 -- 124 | 430 -- 469 | -10.2627 | 4.52257 | 5.31318 |
| CD630_06350 | CD630_Cdi1_4 | 138 -- 167 | 184 -- 213 | -10.2618 | 4.35983 | 5.31318 |
| CD630_08510 | CD630_Cdi1_4 | 42 -- 76 | 178 -- 215 | -10.2448 | 5.16329 | 5.31318 |
| cwp26 | CD630_Cdi1_4 | 131 -- 161 | 142 -- 172 | -10.2308 | 4.44377 | 5.31318 |
| CD630_13380 | CD630_Cdi1_4 | 177 -- 186 | 443 -- 452 | -10.2292 | 4.74408 | 5.31318 |
| CD630_19801 | CD630_Cdi1_4 | 88 -- 98 | 350 -- 360 | -10.2168 | 5.22035 | 5.31318 |
| CD630_10830 | CD630_Cdi1_4 | 86 -- 95 | 352 -- 362 | -10.1989 | 4.82398 | 5.31318 |
| CD630_05450 | CD630_Cdi1_4 | 85 -- 92 | 458 -- 465 | -10.1949 | 5.04761 | 5.31318 |
| CD630_35840 | CD630_Cdi1_4 | 80 -- 92 | 459 -- 471 | -10.1883 | 3.14170 | 5.31318 |
| CD630_27120 | CD630_Cdi1_4 | 116 -- 161 | 273 -- 321 | -10.1805 | 4.43720 | 5.31318 |
| cdu1 | CD630_Cdi1_4 | 87 -- 96 | 352 -- 362 | -10.178 | 4.46276 | 5.31318 |
| CD630_18400 | CD630_Cdi1_4 | 121 -- 158 | 112 -- 149 | -10.1631 | 4.67962 | 5.31318 |
| cdtR | CD630_Cdi1_4 | 76 -- 92 | 429 -- 445 | -10.1512 | 5.84368 | 5.31318 |
| CD630_22720 | CD630_Cdi1_4 | 122 -- 158 | 333 -- 363 | -10.1446 | 3.81804 | 5.31318 |
| CD630_01700 | CD630_Cdi1_4 | 85 -- 98 | 449 -- 464 | -10.1374 | 4.97234 | 5.31318 |
| CD630_32040 | CD630_Cdi1_4 | 24 -- 42 | 257 -- 281 | -10.1189 | 4.93516 | 5.31318 |
| iorA | CD630_Cdi1_4 | 57 -- 98 | 438 -- 460 | -10.1111 | 3.83173 | 5.31318 |
| malY | CD630_Cdi1_4 | 77 -- 121 | 316 -- 364 | -10.098 | 4.46345 | 5.31318 |
| CD630_06060 | CD630_Cdi1_4 | 41 -- 59 | 349 -- 362 | -10.0858 | 4.94050 | 5.31318 |
| pilW | CD630_Cdi1_4 | 88 -- 118 | 430 -- 459 | -10.0702 | 4.96316 | 5.31318 |
| pilC | CD630_Cdi1_4 | 129 -- 139 | 460 -- 470 | -10.0584 | 3.54230 | 5.31318 |
| CD630_36190 | CD630_Cdi1_4 | 73 -- 128 | 447 -- 472 | -10.0462 | 4.63449 | 5.31318 |
| etfA | CD630_Cdi1_4 | 124 -- 163 | 179 -- 212 | -10.0345 | 4.80521 | 5.31318 |
| CD630_28280 | CD630_Cdi1_4 | 97 -- 105 | 425 -- 433 | -10.0258 | 4.59833 | 5.31318 |
| CD630_06050 | CD630_Cdi1_4 | 74 -- 90 | 446 -- 460 | -10.0205 | 3.59332 | 5.31318 |
| argG | CD630_Cdi1_4 | 87 -- 134 | 429 -- 466 | -9.99621 | 4.80300 | 5.31318 |
| CD630_13600 | CD630_Cdi1_4 | 129 -- 176 | 444 -- 470 | -9.9804 | 4.45481 | 5.31318 |
| CD630_06060 | CD630_Cdi1_4 | 87 -- 94 | 458 -- 465 | -9.94986 | 4.94050 | 5.31318 |
| flhB | CD630_Cdi1_4 | 18 -- 33 | 395 -- 412 | -9.93724 | 3.55524 | 5.31318 |
| CD630_15060 | CD630_Cdi1_4 | 173 -- 191 | 227 -- 244 | -9.91105 | 3.69396 | 5.31318 |
| CD630_36180 | CD630_Cdi1_4 | 176 -- 183 | 458 -- 465 | -9.91002 | 4.83428 | 5.31318 |
| CD630_33650 | CD630_Cdi1_4 | 151 -- 180 | 379 -- 410 | -9.90236 | 4.80565 | 5.31318 |
| CD630_25070 | CD630_Cdi1_4 | 24 -- 42 | 436 -- 453 | -9.89825 | 3.45164 | 5.31318 |
| CD630_30730 | CD630_Cdi1_4 | 186 -- 201 | 382 -- 398 | -9.89082 | 3.13236 | 5.31318 |
| cynT | CD630_Cdi1_4 | 146 -- 169 | 108 -- 127 | -9.88715 | 4.21370 | 5.31318 |
| CD630_15890 | CD630_Cdi1_4 | 146 -- 166 | 295 -- 316 | -9.88669 | 3.39962 | 5.31318 |
| CD630_25320 | CD630_Cdi1_4 | 144 -- 173 | 184 -- 213 | -9.88459 | 4.29415 | 5.31318 |
| CD630_29760 | CD630_Cdi1_4 | 86 -- 93 | 458 -- 465 | -9.86919 | 3.51074 | 5.31318 |
| CD630_06100 | CD630_Cdi1_4 | 65 -- 74 | 381 -- 390 | -9.869 | 4.43918 | 5.31318 |
| CD630_03650 | CD630_Cdi1_4 | 85 -- 94 | 427 -- 436 | -9.86074 | 3.93697 | 5.31318 |
| CD630_18781 | CD630_Cdi1_4 | 109 -- 125 | 196 -- 212 | -9.84787 | 3.77660 | 5.31318 |
| CD630_05950 | CD630_Cdi1_4 | 34 -- 69 | 180 -- 212 | -9.84437 | 3.31669 | 5.31318 |
| CD630_05150 | CD630_Cdi1_4 | 87 -- 109 | 430 -- 457 | -9.83147 | 3.69875 | 5.31318 |
| CD630_05230 | CD630_Cdi1_4 | 2 -- 72 | 376 -- 467 | -9.82876 | 3.73787 | 5.31318 |
| aroD | CD630_Cdi1_4 | 137 -- 176 | 430 -- 462 | -9.80768 | 3.71929 | 5.31318 |
| CD630_27280 | CD630_Cdi1_4 | 122 -- 141 | 382 -- 401 | -9.80759 | 3.24738 | 5.31318 |
| CD630_24920 | CD630_Cdi1_4 | 89 -- 150 | 112 -- 164 | -9.75926 | 3.89455 | 5.31318 |
| CD630_21030 | CD630_Cdi1_4 | 57 -- 107 | 114 -- 163 | -9.75456 | 3.08733 | 5.31318 |
| CD630_22510 | CD630_Cdi1_4 | 114 -- 127 | 225 -- 238 | -9.74758 | 11.42763 | 5.31318 |
| CD630_20980 | CD630_Cdi1_4 | 101 -- 110 | 87 -- 96 | -9.71741 | 8.12847 | 5.31318 |
| CD630_09840 | CD630_Cdi1_4 | 162 -- 177 | 44 -- 58 | -9.69133 | 4.50009 | 5.31318 |
| xylR | CD630_Cdi1_4 | 126 -- 139 | 442 -- 455 | -9.67919 | 3.95671 | 5.31318 |
| hfq | CD630_Cdi1_4 | 153 -- 190 | 292 -- 321 | -9.6355 | 5.05609 | 5.31318 |
| CD630_19840 | CD630_Cdi1_4 | 90 -- 96 | 457 -- 463 | -9.63057 | 3.44724 | 5.31318 |
| CD630_14600 | CD630_Cdi1_4 | 65 -- 101 | 126 -- 163 | -9.62582 | 5.90348 | 5.31318 |
| CD630_19850 | CD630_Cdi1_4 | 88 -- 115 | 430 -- 462 | -9.6181 | 6.88486 | 5.31318 |
| CD630_06560 | CD630_Cdi1_4 | 137 -- 187 | 397 -- 459 | -9.5996 | 4.02532 | 5.31318 |
| CD630_18660 | CD630_Cdi1_4 | 83 -- 107 | 444 -- 469 | -9.59753 | 3.50371 | 5.31318 |
| CD630_22070 | CD630_Cdi1_4 | 42 -- 53 | 445 -- 457 | -9.59743 | 4.09651 | 5.31318 |
| CD630_22710 | CD630_Cdi1_4 | 84 -- 115 | 180 -- 213 | -9.59639 | 4.05623 | 5.31318 |
| CD630_32880 | CD630_Cdi1_4 | 137 -- 190 | 278 -- 320 | -9.59553 | 3.41276 | 5.31318 |
| gatC | CD630_Cdi1_4 | 86 -- 123 | 426 -- 464 | -9.58961 | 4.25143 | 5.31318 |
| CD630_24591 | CD630_Cdi1_4 | 88 -- 96 | 352 -- 360 | -9.58466 | 3.54497 | 5.31318 |
| CD630_34520 | CD630_Cdi1_4 | 77 -- 113 | 114 -- 146 | -9.57564 | 3.83016 | 5.31318 |
| dltB | CD630_Cdi1_4 | 7 -- 25 | 380 -- 398 | -9.55275 | 3.81485 | 5.31318 |
| panG | CD630_Cdi1_4 | 47 -- 53 | 456 -- 462 | -9.54192 | 5.54047 | 5.31318 |
| CD630_32990 | CD630_Cdi1_4 | 35 -- 98 | 188 -- 261 | -9.54162 | 5.24925 | 5.31318 |
| htpG | CD630_Cdi1_4 | 86 -- 121 | 428 -- 464 | -9.52978 | 3.45147 | 5.31318 |
| CD630_23070 | CD630_Cdi1_4 | 5 -- 16 | 30 -- 41 | -9.52112 | 5.23027 | 5.31318 |
| CD630_08130 | CD630_Cdi1_4 | 73 -- 103 | 180 -- 208 | -9.517 | 4.63831 | 5.31318 |
| pilT | CD630_Cdi1_4 | 193 -- 199 | 454 -- 460 | -9.5062 | 3.24639 | 5.31318 |
| CD630_19942 | CD630_Cdi1_4 | 186 -- 197 | 443 -- 453 | -9.50382 | 4.24574 | 5.31318 |
| CD630_15460 | CD630_Cdi1_4 | 88 -- 97 | 352 -- 362 | -9.50274 | 6.86296 | 5.31318 |
| CD630_29750 | CD630_Cdi1_4 | 65 -- 95 | 455 -- 471 | -9.50018 | 4.59146 | 5.31318 |
| CD630_21260 | CD630_Cdi1_7 | 138 -- 152 | 43 -- 57 | -12.381 | 4.22255 | 5.53585 |
| ldh | CD630_Cdi1_7 | 157 -- 171 | 8 -- 22 | -10.8856 | 4.35818 | 5.53585 |
| CD630_25190 | CD630_Cdi1_7 | 174 -- 181 | 93 -- 100 | -10.8499 | 3.19663 | 5.53585 |
| CD630_15360 | CD630_Cdi1_7 | 24 -- 37 | 41 -- 54 | -10.5792 | 4.47208 | 5.53585 |
| CD630_34050 | CD630_Cdi1_7 | 8 -- 20 | 3 -- 15 | -10.3528 | 4.03863 | 5.53585 |
| CD630_20670 | CD630_Cdi1_7 | 179 -- 199 | 76 -- 95 | -10.2297 | 4.42757 | 5.53585 |
| CD630_32890 | CD630_Cdi1_7 | 89 -- 113 | 56 -- 79 | -10.1203 | 3.41276 | 5.53585 |
| CD630_22200 | CD630_Cdi1_8 | 102 -- 173 | 117 -- 188 | -19.3729 | 4.18288 | 5.79994 |
| feoB | CD630_Cdi1_8 | 134 -- 197 | 89 -- 147 | -19.2594 | 4.05364 | 5.79994 |
| CD630_06140 | CD630_Cdi1_8 | 16 -- 93 | 114 -- 190 | -19.0622 | 4.12519 | 5.79994 |
| CD630_31530 | CD630_Cdi1_8 | 11 -- 31 | 181 -- 201 | -17.103 | 3.49100 | 5.79994 |
| CD630_05160 | CD630_Cdi1_8 | 180 -- 196 | 283 -- 300 | -16.6083 | 5.03573 | 5.79994 |
| CD630_08080 | CD630_Cdi1_8 | 3 -- 51 | 126 -- 176 | -16.235 | 4.87819 | 5.79994 |
| CD630_20450 | CD630_Cdi1_8 | 113 -- 165 | 94 -- 150 | -16.218 | 3.77388 | 5.79994 |
| CD630_31461 | CD630_Cdi1_8 | 141 -- 190 | 98 -- 147 | -15.8877 | 4.41476 | 5.79994 |
| CD630_14600 | CD630_Cdi1_8 | 137 -- 201 | 99 -- 151 | -15.4078 | 5.90348 | 5.79994 |
| CD630_15270 | CD630_Cdi1_8 | 92 -- 158 | 240 -- 305 | -15.3185 | 4.66015 | 5.79994 |
| adhE | CD630_Cdi1_8 | 1 -- 60 | 79 -- 144 | -15.0643 | 3.06485 | 5.79994 |
| flhB | CD630_Cdi1_8 | 97 -- 166 | 94 -- 164 | -14.8852 | 3.55524 | 5.79994 |
| CD630_15431 | CD630_Cdi1_8 | 139 -- 192 | 255 -- 307 | -14.7025 | 5.29780 | 5.79994 |
| CD630_27270 | CD630_Cdi1_8 | 57 -- 175 | 89 -- 212 | -14.6733 | 3.50892 | 5.79994 |
| CD630_26920 | CD630_Cdi1_8 | 40 -- 111 | 121 -- 190 | -14.3976 | 4.58060 | 5.79994 |
| CD630_06170 | CD630_Cdi1_8 | 117 -- 154 | 104 -- 148 | -14.1991 | 4.95366 | 5.79994 |
| CD630_21700 | CD630_Cdi1_8 | 22 -- 41 | 127 -- 151 | -14.1833 | 3.96650 | 5.79994 |
| CD630_13350 | CD630_Cdi1_8 | 33 -- 181 | 136 -- 281 | -14.0509 | 4.15139 | 5.79994 |
| CD630_05290 | CD630_Cdi1_8 | 1 -- 48 | 278 -- 324 | -13.703 | 3.97604 | 5.79994 |
| CD630_02790 | CD630_Cdi1_8 | 14 -- 52 | 255 -- 290 | -13.5977 | 4.77661 | 5.79994 |
| CD630_27770 | CD630_Cdi1_8 | 26 -- 39 | 292 -- 305 | -13.5613 | 4.08588 | 5.79994 |
| CD630_27110 | CD630_Cdi1_8 | 21 -- 138 | 94 -- 199 | -13.4649 | 3.78231 | 5.79994 |
| opuCC | CD630_Cdi1_8 | 123 -- 194 | 240 -- 321 | -13.2649 | 4.07035 | 5.79994 |
| CD630_32710 | CD630_Cdi1_8 | 136 -- 191 | 168 -- 211 | -13.1447 | 4.74276 | 5.79994 |
| CD630_21620 | CD630_Cdi1_8 | 120 -- 167 | 113 -- 164 | -13.1133 | 3.02649 | 5.79994 |
| ppnK | CD630_Cdi1_8 | 88 -- 162 | 135 -- 206 | -13.0987 | 4.36817 | 5.79994 |
| CD630_29500 | CD630_Cdi1_8 | 40 -- 60 | 180 -- 199 | -13.0694 | 3.70644 | 5.79994 |
| CD630_06100 | CD630_Cdi1_8 | 121 -- 155 | 256 -- 292 | -13.0436 | 4.43918 | 5.79994 |
| cblQ | CD630_Cdi1_8 | 6 -- 77 | 61 -- 128 | -13.0296 | 3.79830 | 5.79994 |
| CD630_07960 | CD630_Cdi1_8 | 186 -- 201 | 283 -- 298 | -12.9698 | 3.22537 | 5.79994 |
| CD630_19401 | CD630_Cdi1_8 | 26 -- 107 | 99 -- 181 | -12.9423 | 4.37084 | 5.79994 |
| CD630_34530 | CD630_Cdi1_8 | 4 -- 56 | 110 -- 166 | -12.6779 | 3.83016 | 5.79994 |
| CD630_23450 | CD630_Cdi1_8 | 90 -- 157 | 249 -- 321 | -12.6466 | 3.27218 | 5.79994 |
| CD630_08140 | CD630_Cdi1_8 | 5 -- 124 | 35 -- 147 | -12.5013 | 3.30159 | 5.79994 |
| pilD | CD630_Cdi1_8 | 3 -- 29 | 113 -- 140 | -12.4686 | 4.54123 | 5.79994 |
| CD630_13020 | CD630_Cdi1_8 | 113 -- 166 | 280 -- 325 | -12.4034 | 3.52267 | 5.79994 |
| CD630_21260 | CD630_Cdi1_8 | 138 -- 163 | 31 -- 58 | -12.3422 | 4.22255 | 5.79994 |
| CD630_16540 | CD630_Cdi1_8 | 161 -- 184 | 124 -- 147 | -12.3014 | 4.37658 | 5.79994 |
| CD630_28181 | CD630_Cdi1_8 | 14 -- 76 | 94 -- 145 | -12.2578 | 3.91299 | 5.79994 |
| CD630_31970 | CD630_Cdi1_8 | 44 -- 82 | 277 -- 315 | -12.1272 | 3.24511 | 5.79994 |
| CD630_05460 | CD630_Cdi1_8 | 12 -- 113 | 94 -- 203 | -12.0838 | 3.98753 | 5.79994 |
| CD630_20670 | CD630_Cdi1_8 | 179 -- 199 | 77 -- 96 | -12.0721 | 4.42757 | 5.79994 |
| CD630_18910 | CD630_Cdi1_8 | 178 -- 200 | 184 -- 206 | -12.0614 | 3.30688 | 5.79994 |
| CD630_23770 | CD630_Cdi1_8 | 28 -- 77 | 171 -- 208 | -11.9758 | 3.25181 | 5.79994 |
| CD630_16160 | CD630_Cdi1_8 | 51 -- 68 | 315 -- 332 | -11.9704 | 6.11801 | 5.79994 |
| CD630_05480 | CD630_Cdi1_8 | 169 -- 191 | 231 -- 252 | -11.8726 | 4.27854 | 5.79994 |
| CD630_22660 | CD630_Cdi1_8 | 133 -- 201 | 85 -- 146 | -11.8662 | 3.60496 | 5.79994 |
| CD630_13620 | CD630_Cdi1_8 | 186 -- 200 | 288 -- 304 | -11.8588 | 4.77627 | 5.79994 |
| CD630_31220 | CD630_Cdi1_8 | 129 -- 190 | 84 -- 142 | -11.8335 | 3.73779 | 5.79994 |
| CD630_27220 | CD630_Cdi1_8 | 93 -- 116 | 290 -- 309 | -11.8331 | 4.18786 | 5.79994 |
| tcdC | CD630_Cdi1_8 | 86 -- 109 | 277 -- 300 | -11.8047 | 3.76101 | 5.79994 |
| CD630_16670 | CD630_Cdi1_8 | 2 -- 43 | 153 -- 190 | -11.7978 | 3.33553 | 5.79994 |
| agrB | CD630_Cdi1_8 | 41 -- 113 | 234 -- 289 | -11.6312 | 3.92598 | 5.79994 |
| CD630_29960 | CD630_Cdi1_8 | 109 -- 146 | 136 -- 173 | -11.6082 | 4.51904 | 5.79994 |
| CD630_18660 | CD630_Cdi1_8 | 136 -- 192 | 250 -- 306 | -11.6023 | 3.50371 | 5.79994 |
| kdpD | CD630_Cdi1_8 | 152 -- 201 | 95 -- 142 | -11.5694 | 4.27140 | 5.79994 |
| CD630_22880 | CD630_Cdi1_8 | 173 -- 199 | 290 -- 315 | -11.4798 | 3.62455 | 5.79994 |
| CD630_16530 | CD630_Cdi1_8 | 117 -- 157 | 246 -- 289 | -11.445 | 4.37658 | 5.79994 |
| CD630_25020 | CD630_Cdi1_8 | 22 -- 55 | 250 -- 290 | -11.4341 | 4.17721 | 5.79994 |
| CD630_07770 | CD630_Cdi1_8 | 42 -- 79 | 229 -- 264 | -11.4187 | 3.31158 | 5.79994 |
| CD630_21720 | CD630_Cdi1_8 | 35 -- 177 | 82 -- 215 | -11.411 | 4.13030 | 5.79994 |
| CD630_23070 | CD630_Cdi1_8 | 165 -- 187 | 90 -- 110 | -11.4084 | 5.23027 | 5.79994 |
| panB | CD630_Cdi1_8 | 120 -- 181 | 82 -- 147 | -11.3943 | 3.40248 | 5.79994 |
| CD630_13960 | CD630_Cdi1_8 | 49 -- 73 | 254 -- 280 | -11.3712 | 3.30375 | 5.79994 |
| CD630_04620 | CD630_Cdi1_8 | 10 -- 33 | 259 -- 283 | -11.3621 | 3.45413 | 5.79994 |
| oppC | CD630_Cdi1_8 | 164 -- 193 | 108 -- 139 | -11.2741 | 3.01695 | 5.79994 |
| CD630_32060 | CD630_Cdi1_8 | 161 -- 189 | 290 -- 325 | -11.1671 | 5.73797 | 5.79994 |
| CD630_20460 | CD630_Cdi1_8 | 94 -- 182 | 136 -- 208 | -11.1279 | 7.94034 | 5.79994 |
| CD630_30420 | CD630_Cdi1_8 | 152 -- 172 | 304 -- 324 | -11.1215 | 5.27539 | 5.79994 |
| CD630_21570 | CD630_Cdi1_8 | 157 -- 186 | 119 -- 145 | -11.1048 | 6.48385 | 5.79994 |
| CD630_09100 | CD630_Cdi1_8 | 17 -- 66 | 231 -- 292 | -11.1042 | 5.92260 | 5.79994 |
| CD630_28870 | CD630_Cdi1_8 | 180 -- 197 | 177 -- 194 | -11.08 | 3.41991 | 5.79994 |
| CD630_08950 | CD630_Cdi1_8 | 156 -- 191 | 285 -- 322 | -11.0662 | 3.71905 | 5.79994 |
| CD630_25190 | CD630_Cdi1_8 | 174 -- 181 | 94 -- 101 | -11.0578 | 3.19663 | 5.79994 |
| CD630_08510 | CD630_Cdi1_8 | 21 -- 79 | 94 -- 141 | -11.0298 | 5.16329 | 5.79994 |
| CD630_10080 | CD630_Cdi1_8 | 1 -- 50 | 282 -- 334 | -11.011 | 3.75625 | 5.79994 |
| CD630_17511 | CD630_Cdi1_8 | 103 -- 115 | 283 -- 294 | -11.0107 | 9.12322 | 5.79994 |
| CD630_04610 | CD630_Cdi1_8 | 46 -- 76 | 113 -- 150 | -10.9941 | 3.83025 | 5.79994 |
| CD630_10950 | CD630_Cdi1_8 | 176 -- 192 | 270 -- 285 | -10.9937 | 3.67675 | 5.79994 |
| CD630_22850 | CD630_Cdi1_8 | 123 -- 186 | 94 -- 153 | -10.9638 | 3.84102 | 5.79994 |
| CD630_08780 | CD630_Cdi1_8 | 127 -- 174 | 82 -- 125 | -10.8746 | 7.76690 | 5.79994 |
| ldh | CD630_Cdi1_8 | 157 -- 171 | 9 -- 23 | -10.8471 | 4.35818 | 5.79994 |
| CD630_19942 | CD630_Cdi1_8 | 90 -- 141 | 94 -- 142 | -10.8324 | 4.24574 | 5.79994 |
| CD630_14700 | CD630_Cdi1_8 | 111 -- 157 | 115 -- 162 | -10.8207 | 3.98234 | 5.79994 |
| CD630_15900 | CD630_Cdi1_8 | 153 -- 187 | 284 -- 319 | -10.7947 | 3.17664 | 5.79994 |
| CD630_17770 | CD630_Cdi1_8 | 109 -- 137 | 285 -- 312 | -10.7487 | 5.06490 | 5.79994 |
| CD630_05830 | CD630_Cdi1_8 | 86 -- 144 | 29 -- 96 | -10.7355 | 3.35360 | 5.79994 |
| CD630_21750 | CD630_Cdi1_8 | 146 -- 169 | 246 -- 264 | -10.7307 | 3.70626 | 5.79994 |
| CD630_32890 | CD630_Cdi1_8 | 89 -- 113 | 57 -- 80 | -10.7277 | 3.41276 | 5.79994 |
| etfA | CD630_Cdi1_8 | 77 -- 93 | 293 -- 309 | -10.7228 | 4.80521 | 5.79994 |
| CD630_06060 | CD630_Cdi1_8 | 41 -- 178 | 48 -- 194 | -10.7161 | 4.94050 | 5.79994 |
| CD630_28560 | CD630_Cdi1_8 | 117 -- 132 | 277 -- 292 | -10.7073 | 3.96803 | 5.79994 |
| CD630_27120 | CD630_Cdi1_8 | 153 -- 174 | 77 -- 96 | -10.6818 | 4.43720 | 5.79994 |
| CD630_33480 | CD630_Cdi1_8 | 66 -- 142 | 87 -- 147 | -10.6815 | 3.67613 | 5.79994 |
| fliI | CD630_Cdi1_8 | 82 -- 106 | 287 -- 312 | -10.6641 | 3.03666 | 5.79994 |
| CD630_21760 | CD630_Cdi1_8 | 132 -- 158 | 161 -- 189 | -10.653 | 4.43846 | 5.79994 |
| CD630_20660 | CD630_Cdi1_8 | 106 -- 167 | 117 -- 188 | -10.6333 | 5.39321 | 5.79994 |
| CD630_05950 | CD630_Cdi1_8 | 81 -- 153 | 234 -- 301 | -10.5935 | 3.31669 | 5.79994 |
| CD630_02720 | CD630_Cdi1_8 | 10 -- 26 | 292 -- 308 | -10.5507 | 3.38938 | 5.79994 |
| CD630_14250 | CD630_Cdi1_8 | 28 -- 40 | 281 -- 293 | -10.5453 | 4.19604 | 5.79994 |
| CD630_28320 | CD630_Cdi1_8 | 160 -- 195 | 42 -- 79 | -10.5376 | 3.99184 | 5.79994 |
| CD630_25050 | CD630_Cdi1_8 | 6 -- 51 | 134 -- 182 | -10.506 | 4.33624 | 5.79994 |
| CD630_19904 | CD630_Cdi1_8 | 133 -- 175 | 91 -- 143 | -10.5026 | 5.96526 | 5.79994 |
| metQ | CD630_Cdi1_8 | 119 -- 143 | 295 -- 323 | -10.4876 | 3.87190 | 5.79994 |
| CD630_05880 | CD630_Cdi1_8 | 93 -- 108 | 291 -- 305 | -10.4837 | 4.28339 | 5.79994 |
| CD630_34050 | CD630_Cdi1_8 | 8 -- 20 | 4 -- 16 | -10.4822 | 4.03863 | 5.79994 |
| CD630_06080 | CD630_Cdi1_8 | 1 -- 28 | 125 -- 153 | -10.3755 | 3.82315 | 5.79994 |
| CD630_07870 | CD630_Cdi1_8 | 162 -- 193 | 241 -- 269 | -10.3568 | 3.22829 | 5.79994 |
| CD630_26050 | CD630_Cdi1_8 | 96 -- 123 | 287 -- 312 | -10.3114 | 6.99851 | 5.79994 |
| CD630_27280 | CD630_Cdi1_8 | 140 -- 192 | 249 -- 305 | -10.2452 | 3.24738 | 5.79994 |
| CD630_05300 | CD630_Cdi1_8 | 5 -- 46 | 137 -- 185 | -10.2374 | 4.56885 | 5.79994 |
| CD630_21110 | CD630_Cdi1_8 | 120 -- 158 | 126 -- 169 | -10.1917 | 3.89630 | 5.79994 |
| CD630_13790 | CD630_Cdi1_8 | 139 -- 196 | 94 -- 147 | -10.1866 | 5.17374 | 5.79994 |
| malY | CD630_Cdi1_8 | 8 -- 26 | 190 -- 208 | -10.1684 | 4.46345 | 5.79994 |
| CD630_18680 | CD630_Cdi1_8 | 66 -- 154 | 111 -- 206 | -10.1605 | 3.04315 | 5.79994 |

| | | | | | | |
|---|---|---|---|---|---|---|
| acoL | CD630_Cdi1_8 | 44↔105 | 107↔164 | -10.1415 | 3.32169 | 5.79994 |
| CD630_13920 | CD630_Cdi1_8 | 173↔191 | 285↔303 | -10.1397 | 3.25236 | 5.79994 |
| sspA | CD630_Cdi1_8 | 54↔109 | 97↔154 | -10.0885 | 3.62689 | 5.79994 |
| appC | CD630_Cdi1_8 | 44↔66 | 313↔332 | -10.0669 | 4.50438 | 5.79994 |
| CD630_23760 | CD630_Cdi1_8 | 56↔69 | 193↔206 | -10.0626 | 4.54307 | 5.79994 |
| CD630_27570 | CD630_Cdi1_8 | 1↔22 | 165↔189 | -10.0302 | 3.46216 | 5.79994 |
| glgP | CD630_Cdi1_8 | 81↔129 | 98↔141 | -10.0167 | 3.28530 | 5.79994 |
| CD630_30310 | CD630_Cdi1_8 | 142↔167 | 122↔148 | -10.005 | 4.56059 | 5.79994 |
| cbpA | CD630_Cdi1_8 | 76↔132 | 109↔158 | -9.9959 | 3.03040 | 5.79994 |
| bioB | CD630_Cdi1_8 | 129↔148 | 281↔301 | -9.98996 | 3.59022 | 5.79994 |
| CD630_25150 | CD630_Cdi1_8 | 28↔106 | 126↔202 | -9.97928 | 3.79339 | 5.79994 |
| CD630_16640 | CD630_Cdi1_8 | 65↔180 | 91↔188 | -9.94887 | 3.61548 | 5.79994 |
| rbsC | CD630_Cdi1_8 | 156↔174 | 236↔253 | -9.9462 | 3.71650 | 5.79994 |
| CD630_36781 | CD630_Cdi1_8 | 140↔195 | 116↔168 | -9.93009 | 3.75946 | 5.79994 |
| CD630_13580 | CD630_Cdi1_8 | 142↔158 | 101↔117 | -9.92958 | 3.80715 | 5.79994 |
| CD630_07760 | CD630_Cdi1_8 | 181↔193 | 235↔247 | -9.91466 | 3.64186 | 5.79994 |
| CD630_02830 | CD630_Cdi1_8 | 140↔187 | 94↔150 | -9.90785 | 3.57106 | 5.79994 |
| chrA | CD630_Cdi1_8 | 69↔133 | 135↔196 | -9.88948 | 5.27255 | 5.79994 |
| CD630_33600 | CD630_Cdi1_8 | 41↔61 | 283↔300 | -9.88408 | 3.73196 | 5.79994 |
| CD630_23780 | CD630_Cdi1_8 | 40↔72 | 240↔277 | -9.88311 | 3.91629 | 5.79994 |
| CD630_03420 | CD630_Cdi1_8 | 138↔175 | 115↔153 | -9.86708 | 4.71868 | 5.79994 |
| CD630_17170 | CD630_Cdi1_8 | 136↔155 | 186↔205 | -9.86416 | 4.10230 | 5.79994 |
| CD630_24050 | CD630_Cdi1_8 | 117↔160 | 269↔310 | -9.86284 | 3.70192 | 5.79994 |
| uxaA | CD630_Cdi1_8 | 8↔23 | 86↔102 | -9.77418 | 4.26165 | 5.79994 |
| etfA | CD630_Cdi1_8 | 22↔66 | 123↔166 | -9.73629 | 3.41204 | 5.79994 |
| CD630_32990 | CD630_Cdi1_8 | 53↔97 | 242↔287 | -9.72725 | 5.24925 | 5.79994 |
| CD630_05900 | CD630_Cdi1_8 | 4↔51 | 285↔331 | -9.70492 | 3.21934 | 5.79994 |
| CD630_33580 | CD630_Cdi1_8 | 189↔201 | 313↔325 | -9.69396 | 3.95778 | 5.79994 |
| CD630_20600 | CD630_Cdi1_8 | 150↔178 | 126↔154 | -9.68651 | 4.20093 | 5.79994 |
| mobA | CD630_Cdi1_8 | 119↔192 | 95↔164 | -9.67932 | 4.25703 | 5.79994 |
| CD630_16460 | CD630_Cdi1_8 | 130↔148 | 77↔96 | -9.67437 | 4.64028 | 5.79994 |
| CD630_27240 | CD630_Cdi1_8 | 6↔20 | 191↔208 | -9.67008 | 3.52462 | 5.79994 |
| CD630_04940 | CD630_Cdi1_8 | 77↔135 | 136↔198 | -9.66198 | 4.20948 | 5.79994 |
| glnA | CD630_Cdi1_8 | 149↔172 | 136↔158 | -9.65053 | 3.87629 | 5.79994 |
| CD630_13880 | CD630_Cdi1_8 | 80↔151 | 109↔187 | -9.62064 | 3.52735 | 5.79994 |
| CD630_22550 | CD630_Cdi1_8 | 26↔51 | 127↔152 | -9.6052 | 3.62379 | 5.79994 |
| CD630_02740 | CD630_Cdi1_8 | 36↔93 | 118↔198 | -9.59753 | 5.02277 | 5.79994 |
| CD630_15460 | CD630_Cdi1_8 | 168↔184 | 97↔112 | -9.59721 | 6.86296 | 5.79994 |
| CD630_19840 | CD630_Cdi1_8 | 175↔196 | 160↔181 | -9.58962 | 3.44724 | 5.79994 |
| CD630_18610 | CD630_Cdi1_8 | 151↔182 | 234↔255 | -9.58697 | 4.15582 | 5.79994 |
| CD630_15360 | CD630_Cdi1_8 | 24↔37 | 42↔55 | -9.55423 | 4.47208 | 5.79994 |
| CD630_16550 | CD630_Cdi1_8 | 187↔199 | 283↔295 | -9.53274 | 5.61064 | 5.79994 |
| CD630_33190 | CD630_Cdi1_8 | 126↔138 | 283↔295 | -9.53144 | 4.13899 | 5.79994 |
| CD630_36250 | CD630_Cdi1_8 | 55↔94 | 224↔262 | -9.51619 | 4.13224 | 5.79994 |
| CD630_10220 | CD630_Cdi1_8 | 122↔159 | 4↔37 | -9.50979 | 4.08233 | 5.79994 |
| CD630_12650 | CD630_Cdi1_8 | 120↔188 | 126↔191 | -9.50877 | 6.31827 | 5.79994 |
| CD630_06060 | CD630_Cdi1_8 | 82↔106 | 77↔102 | -9.50782 | 4.94050 | 5.79994 |
| CD630_20450 | CD630_Cdi1_9 | 110↔165 | 86↔148 | -15.4708 | 3.77388 | 4.25622 |
| CD630_14700 | CD630_Cdi1_9 | 125↔156 | 107↔140 | -13.3538 | 3.98234 | 4.25622 |
| CD630_16540 | CD630_Cdi1_9 | 161↔183 | 117↔139 | -12.8948 | 4.37658 | 4.25622 |
| feoB | CD630_Cdi1_9 | 134↔197 | 81↔139 | -12.719 | 4.05364 | 4.25622 |
| CD630_14600 | CD630_Cdi1_9 | 137↔166 | 116↔143 | -12.564 | 5.90348 | 4.25622 |
| CD630_02420 | CD630_Cdi1_9 | 9↔50 | 48↔87 | -12.3377 | 3.71048 | 4.25622 |
| CD630_36690 | CD630_Cdi1_9 | 90↔118 | 103↔134 | -12.0666 | 3.53761 | 4.25622 |
| CD630_28181 | CD630_Cdi1_9 | 14↔82 | 86↔137 | -11.9829 | 3.91299 | 4.25622 |
| CD630_17170 | CD630_Cdi1_9 | 90↔175 | 58↔134 | -11.8719 | 4.10230 | 4.25622 |
| CD630_21260 | CD630_Cdi1_9 | 138↔152 | 44↔58 | -11.7578 | 4.22255 | 4.25622 |
| CD630_15360 | CD630_Cdi1_9 | 34↔70 | 116↔150 | -11.6766 | 4.47208 | 4.25622 |
| etfA | CD630_Cdi1_9 | 164↔182 | 103↔121 | -11.5755 | 3.41204 | 4.25622 |
| cdtR | CD630_Cdi1_9 | 16↔41 | 121↔147 | -11.4897 | 5.84368 | 4.25622 |
| CD630_21700 | CD630_Cdi1_9 | 22↔38 | 122↔143 | -11.4206 | 3.96650 | 4.25622 |
| CD630_17511 | CD630_Cdi1_9 | 48↔109 | 89↔147 | -11.3679 | 9.12322 | 4.25622 |
| CD630_30310 | CD630_Cdi1_9 | 134↔154 | 128↔148 | -11.1331 | 4.56059 | 4.25622 |
| CD630_05940 | CD630_Cdi1_9 | 125↔156 | 124↔151 | -10.9187 | 4.18595 | 4.25622 |
| CD630_15160 | CD630_Cdi1_9 | 186↔201 | 133↔148 | -10.8492 | 4.32318 | 4.25622 |
| CD630_16550 | CD630_Cdi1_9 | 90↔124 | 101↔134 | -10.829 | 5.61064 | 4.25622 |
| CD630_23280 | CD630_Cdi1_9 | 134↔152 | 103↔121 | -10.8248 | 3.50093 | 4.25622 |
| dnaC | CD630_Cdi1_9 | 37↔82 | 105↔139 | -10.6434 | 6.30796 | 4.25622 |
| CD630_10050 | CD630_Cdi1_9 | 140↔163 | 103↔124 | -10.5993 | 3.93750 | 4.25622 |
| ldh | CD630_Cdi1_9 | 157↔171 | 9↔23 | -10.5742 | 4.35818 | 4.25622 |
| CD630_25040 | CD630_Cdi1_9 | 12↔58 | 99↔142 | -10.5294 | 3.43898 | 4.25622 |
| CD630_34050 | CD630_Cdi1_9 | 8↔20 | 4↔16 | -10.5005 | 4.03863 | 4.25622 |
| CD630_13020 | CD630_Cdi1_9 | 85↔106 | 118↔139 | -10.477 | 3.52267 | 4.25622 |
| CD630_22660 | CD630_Cdi1_9 | 132↔201 | 77↔142 | -10.4585 | 3.60496 | 4.25622 |
| CD630_21760 | CD630_Cdi1_9 | 52↔87 | 76↔108 | -10.3613 | 4.43846 | 4.25622 |
| appC | CD630_Cdi1_9 | 39↔58 | 72↔87 | -10.2699 | 4.50438 | 4.25622 |
| CD630_13600 | CD630_Cdi1_9 | 133↔182 | 55↔95 | -10.0602 | 4.45481 | 4.25622 |
| CD630_18230 | CD630_Cdi1_9 | 108↔140 | 114↔152 | -10.0591 | 3.00184 | 4.25622 |
| CD630_33610 | CD630_Cdi1_9 | 106↔187 | 71↔152 | -10.0436 | 3.79322 | 4.25622 |
| sspA | CD630_Cdi1_9 | 79↔109 | 89↔118 | -10.0222 | 3.62689 | 4.25622 |
| CD630_27280 | CD630_Cdi1_9 | 146↔189 | 48↔94 | -9.99022 | 3.24738 | 4.25622 |
| CD630_14130 | CD630_Cdi1_9 | 173↔201 | 82↔110 | -9.9543 | 4.57970 | 4.25622 |
| CD630_25190 | CD630_Cdi1_9 | 174↔181 | 86↔93 | -9.93953 | 3.19663 | 4.25622 |
| CD630_10880 | CD630_Cdi1_9 | 138↔197 | 73↔142 | -9.91369 | 5.05048 | 4.25622 |
| ppnK | CD630_Cdi1_9 | 142↔162 | 127↔142 | -9.9074 | 4.36817 | 4.25622 |
| CD630_23070 | CD630_Cdi1_9 | 158↔187 | 82↔107 | -9.89656 | 5.23027 | 4.25622 |
| CD630_13620 | CD630_Cdi1_9 | 140↔164 | 118↔142 | -9.7816 | 4.77627 | 4.25622 |
| CD630_36781 | CD630_Cdi1_9 | 118↔131 | 135↔148 | -9.77094 | 3.75946 | 4.25622 |
| CD630_21770 | CD630_Cdi1_9 | 188↔200 | 109↔121 | -9.71414 | 4.02587 | 4.25622 |
| CD630_21020 | CD630_Cdi1_9 | 70↔107 | 117↔152 | -9.63261 | 4.28666 | 4.25622 |
| CD630_18610 | CD630_Cdi1_9 | 17↔38 | 50↔69 | -9.61793 | 4.15582 | 4.25622 |
| CD630_31530 | CD630_Cdi1_9 | 118↔168 | 83↔137 | -9.59339 | 3.49100 | 4.25622 |
| CD630_05280 | CD630_Cdi1_9 | 121↔165 | 101↔142 | -9.53221 | 3.91665 | 4.25622 |
| CD630_27570 | CD630_n00220 | 5↔22 | 2↔18 | -13.7069 | 3.46216 | 4.94050 |
| CD630_19904 | CD630_n00220 | 2↔33 | 1↔35 | -12.5507 | 5.96526 | 4.94050 |
| CD630_34520 | CD630_n00220 | 28↔55 | 5↔32 | -11.9518 | 3.83016 | 4.94050 |
| CD630_31690 | CD630_n00220 | 99↔139 | 4↔45 | -11.9225 | 3.71193 | 4.94050 |
| appC | CD630_n00220 | 156↔200 | 39↔80 | -11.6615 | 4.50438 | 4.94050 |
| CD630_27280 | CD630_n00220 | 187↔198 | 5↔15 | -11.4701 | 3.24738 | 4.94050 |
| CD630_03410 | CD630_n00220 | 46↔70 | 1↔25 | -11.4298 | 4.45994 | 4.94050 |
| CD630_09050 | CD630_n00220 | 168↔193 | 56↔85 | -10.9159 | 3.96216 | 4.94050 |
| CD630_24830 | CD630_n00220 | 138↔185 | 1↔56 | -10.9024 | 4.98239 | 4.94050 |
| CD630_21090 | CD630_n00220 | 127↔201 | 11↔85 | -10.6156 | 4.15803 | 4.94050 |
| CD630_09100 | CD630_n00220 | 51↔96 | 4↔45 | -10.318 | 5.92260 | 4.94050 |
| CD630_13370 | CD630_n00220 | 157↔188 | 12↔48 | -10.255 | 4.33145 | 4.94050 |
| CD630_18410 | CD630_n00220 | 139↔175 | 5↔45 | -10.1812 | 3.59739 | 4.94050 |
| CD630_24220 | CD630_n00220 | 134↔186 | 6↔59 | -10.1199 | 4.18037 | 4.94050 |
| CD630_30420 | CD630_n00220 | 132↔174 | 46↔89 | -9.92901 | 5.27539 | 4.94050 |
| dnaD | CD630_n00220 | 21↔50 | 32↔60 | -9.87884 | 3.56078 | 4.94050 |
| CD630_15190 | CD630_n00220 | 119↔159 | 1↔42 | -9.85751 | 4.43255 | 4.94050 |
| CD630_17720 | CD630_n00220 | 1↔38 | 23↔55 | -9.79332 | 3.22910 | 4.94050 |
| CD630_08660 | CD630_n00220 | 165↔178 | 3↔16 | -9.79205 | 3.86467 | 4.94050 |
| CD630_24591 | CD630_n00220 | 18↔51 | 1↔26 | -9.66699 | 3.54497 | 4.94050 |
| CD630_20280 | CD630_n00220 | 118↔193 | 2↔79 | -9.55558 | 3.46343 | 4.94050 |
| thiH | CD630_n00240 | 177↔201 | 5↔32 | -16.1397 | 3.58230 | 5.82709 |
| cls | CD630_n00240 | 68↔99 | 1↔33 | -14.4902 | 4.46177 | 5.82709 |
| vncR | CD630_n00240 | 124↔145 | 4↔27 | -14.0574 | 3.93642 | 5.82709 |
| CD630_16150 | CD630_n00240 | 71↔90 | 5↔25 | -13.8943 | 4.20614 | 5.82709 |
| CD630_18940 | CD630_n00240 | 145↔164 | 3↔23 | -13.388 | 4.09988 | 5.82709 |
| CD630_11730 | CD630_n00240 | 136↔160 | 4↔29 | -12.7584 | 4.80521 | 5.82709 |
| cobQ | CD630_n00240 | 69↔94 | 5↔30 | -12.7407 | 3.13524 | 5.82709 |
| CD630_20600 | CD630_n00240 | 17↔73 | 75↔117 | -12.6757 | 4.20093 | 5.82709 |
| dltB | CD630_n00240 | 3↔31 | 94↔121 | -12.4204 | 3.81485 | 5.82709 |
| CD630_20650 | CD630_n00240 | 18↔47 | 6↔32 | -12.1818 | 4.00083 | 5.82709 |
| CD630_14250 | CD630_n00240 | 167↔193 | 3↔31 | -12.1644 | 4.19604 | 5.82709 |
| cynT | CD630_n00240 | 106↔128 | 3↔23 | -12.1109 | 4.21370 | 5.82709 |
| CD630_33682 | CD630_n00240 | 19↔90 | 3↔78 | -12.1094 | 5.77720 | 5.82709 |
| CD630_06080 | CD630_n00240 | 8↔32 | 4↔31 | -12.0751 | 3.82315 | 5.82709 |
| CD630_21770 | CD630_n00240 | 73↔95 | 3↔26 | -11.9298 | 4.02587 | 5.82709 |
| CD630_28150 | CD630_n00240 | 58↔78 | 10↔27 | -11.8716 | 4.18447 | 5.82709 |
| rbsA | CD630_n00240 | 6↔24 | 7↔33 | -11.6382 | 3.71650 | 5.82709 |
| CD630_03220 | CD630_n00240 | 23↔44 | 4↔24 | -11.625 | 3.36746 | 5.82709 |
| CD630_04341 | CD630_n00240 | 59↔80 | 5↔26 | -11.595 | 4.29262 | 5.82709 |
| CD630_04280 | CD630_n00240 | 63↔108 | 68↔118 | -11.5917 | 3.61457 | 5.82709 |
| cdu2 | CD630_n00240 | 38↔70 | 4↔26 | -11.5467 | 2.97335 | 5.82709 |
| CD630_21620 | CD630_n00240 | 147↔164 | 6↔27 | -11.5156 | 3.02649 | 5.82709 |
| serS1 | CD630_n00240 | 103↔120 | 3↔20 | -11.508 | 3.45858 | 5.82709 |
| CD630_06110 | CD630_n00240 | 76↔93 | 8↔25 | -11.3705 | 5.61380 | 5.82709 |
| appF | CD630_n00240 | 94↔120 | 1↔26 | -11.3071 | 4.25570 | 5.82709 |
| CD630_10950 | CD630_n00240 | 70↔99 | 5↔31 | -11.2537 | 3.67675 | 5.82709 |
| CD630_31490 | CD630_n00240 | 132↔151 | 4↔24 | -11.1339 | 4.08593 | 5.82709 |
| CD630_08080 | CD630_n00240 | 170↔186 | 100↔118 | -11.0917 | 4.87819 | 5.82709 |
| CD630_20450 | CD630_n00240 | 40↔69 | 4↔31 | -11.0874 | 3.77388 | 5.82709 |
| CD630_28181 | CD630_n00240 | 39↔52 | 3↔16 | -10.9446 | 3.91299 | 5.82709 |
| CD630_14240 | CD630_n00240 | 19↔43 | 55↔78 | -10.9324 | 8.60766 | 5.82709 |
| CD630_27230 | CD630_n00240 | 107↔125 | 7↔26 | -10.9228 | 3.52462 | 5.82709 |
| CD630_29250 | CD630_n00240 | 180↔200 | 12↔31 | -10.8616 | 4.61901 | 5.82709 |
| CD630_09410 | CD630_n00240 | 180↔200 | 12↔31 | -10.8616 | 4.75651 | 5.82709 |
| CD630_11220 | CD630_n00240 | 41↔58 | 9↔27 | -10.832 | 3.52327 | 5.82709 |
| CD630_21730 | CD630_n00240 | 174↔188 | 102↔117 | -10.8145 | 3.45076 | 5.82709 |
| CD630_27310 | CD630_n00240 | 82↔95 | 3↔16 | -10.7276 | 4.01622 | 5.82709 |
| def1 | CD630_n00240 | 130↔154 | 3↔27 | -10.6929 | 3.99720 | 5.82709 |
| CD630_10880 | CD630_n00240 | 145↔159 | 4↔17 | -10.6292 | 5.05048 | 5.82709 |
| xdhA | CD630_n00240 | 93↔121 | 4↔26 | -10.6027 | 4.45147 | 5.82709 |
| CD630_09990 | CD630_n00240 | 68↔101 | 5↔33 | -10.5993 | 3.86040 | 5.82709 |
| CD630_06540 | CD630_n00240 | 35↔64 | 5↔32 | -10.5126 | 3.54673 | 5.82709 |
| hfq | CD630_n00240 | 146↔157 | 4↔15 | -10.4904 | 5.05609 | 5.82709 |
| kdpE | CD630_n00240 | 122↔149 | 4↔28 | -10.4843 | 4.27140 | 5.82709 |
| tcdC | CD630_n00240 | 131↔151 | 4↔29 | -10.4509 | 3.76101 | 5.82709 |
| CD630_13600 | CD630_n00240 | 121↔139 | 1↔23 | -10.3982 | 4.45481 | 5.82709 |
| CD630_23440 | CD630_n00240 | 1↔12 | 5↔16 | -10.3955 | 3.34841 | 5.82709 |
| CD630_21760 | CD630_n00240 | 62↔80 | 5↔23 | -10.3913 | 4.43846 | 5.82709 |
| feoB | CD630_n00240 | 35↔44 | 7↔16 | -10.3643 | 4.05364 | 5.82709 |
| CD630_05880 | CD630_n00240 | 127↔157 | 4↔32 | -10.3596 | 4.28339 | 5.82709 |
| cbiO | CD630_n00240 | 8↔38 | 5↔32 | -10.3566 | 3.79830 | 5.82709 |
| dapB2 | CD630_n00240 | 123↔146 | 6↔31 | -10.3422 | 3.27218 | 5.82709 |
| CD630_02830 | CD630_n00240 | 174↔201 | 2↔31 | -10.3164 | 3.57106 | 5.82709 |
| CD630_19580 | CD630_n00240 | 74↔95 | 7↔28 | -10.3015 | 4.51902 | 5.82709 |
| dnaC | CD630_n00240 | 126↔151 | 5↔32 | -10.2641 | 6.30796 | 5.82709 |
| mobA | CD630_n00240 | 4↔28 | 4↔27 | -10.2291 | 4.25703 | 5.82709 |
| CD630_09040 | CD630_n00240 | 116↔141 | 7↔31 | -10.1398 | 4.11851 | 5.82709 |
| CD630_34030 | CD630_n00240 | 49↔73 | 7↔33 | -10.139 | 3.05766 | 5.82709 |
| CD630_33600 | CD630_n00240 | 135↔151 | 3↔18 | -10.1273 | 3.73196 | 5.82709 |
| nusB | CD630_n00240 | 155↔196 | 76↔120 | -10.1158 | 4.92798 | 5.82709 |
| CD630_05870 | CD630_n00240 | 164↔196 | 2↔29 | -10.1117 | 4.28339 | 5.82709 |
| CD630_24920 | CD630_n00240 | 122↔150 | 5↔25 | -10.0583 | 3.89455 | 5.82709 |
| CD630_16460 | CD630_n00240 | 13↔38 | 7↔32 | -10.0556 | 4.64028 | 5.82709 |
| pilC | CD630_n00240 | 171↔187 | 15↔31 | -10.0543 | 3.54230 | 5.82709 |
| CD630_24591 | CD630_n00240 | 62↔93 | 3↔32 | -10.0518 | 3.54497 | 5.82709 |
| CD630_17170 | CD630_n00240 | 80↔96 | 7↔23 | -10.0148 | 4.10230 | 5.82709 |
| CD630_27570 | CD630_n00240 | 149↔159 | 50↔60 | -10.012 | 3.46216 | 5.82709 |
| CD630_22200 | CD630_n00240 | 69↔95 | 5↔29 | -10.006 | 4.18288 | 5.82709 |
| moaA | CD630_n00240 | 1↔21 | 4↔25 | -10.0053 | 3.83799 | 5.82709 |
| CD630_17630 | CD630_n00240 | 167↔178 | 5↔16 | -9.96649 | 4.52408 | 5.82709 |
| CD630_23070 | CD630_n00240 | 5↔30 | 3↔32 | -9.93834 | 5.23027 | 5.82709 |
| CD630_03170 | CD630_n00240 | 70↔96 | 4↔32 | -9.93554 | 3.66946 | 5.82709 |
| CD630_31500 | CD630_n00240 | 14↔58 | 78↔119 | -9.93203 | 4.90207 | 5.82709 |
| CD630_30130 | CD630_n00240 | 2↔16 | 4↔19 | -9.89948 | 3.28343 | 5.82709 |
| iorA | CD630_n00240 | 73↔144 | 46↔116 | -9.88742 | 3.83173 | 5.82709 |
| CD630_08090 | CD630_n00240 | 147↔173 | 3↔31 | -9.87753 | 3.57167 | 5.82709 |
| etfA | CD630_n00240 | 125↔152 | 1↔26 | -9.86867 | 4.80521 | 5.82709 |
| CD630_06410 | CD630_n00240 | 4↔32 | 83↔111 | -9.86779 | 4.14404 | 5.82709 |
| CD630_19921 | CD630_n00240 | 187↔198 | 74↔85 | -9.819 | 5.46345 | 5.82709 |
| CD630_10800 | CD630_n00240 | 69↔90 | 5↔32 | -9.80118 | 5.30319 | 5.82709 |
| CD630_12320 | CD630_n00240 | 39↔61 | 12↔31 | -9.79207 | 3.28667 | 5.82709 |
| CD630_18410 | CD630_n00240 | 155↔168 | 107↔120 | -9.77097 | 3.59739 | 5.82709 |
| CD630_32710 | CD630_n00240 | 131↔158 | 5↔26 | -9.76921 | 4.74276 | 5.82709 |
| CD630_16670 | CD630_n00240 | 116↔133 | 7↔23 | -9.75683 | 3.33553 | 5.82709 |
| CD630_17360 | CD630_n00240 | 16↔31 | 106↔121 | -9.75135 | 5.66070 | 5.82709 |
| CD630_17292 | CD630_n00240 | 110↔131 | 4↔28 | -9.70711 | 6.25799 | 5.82709 |
| gabT | CD630_n00240 | 59↔73 | 103↔117 | -9.69629 | 3.67067 | 5.82709 |
| kdgT1 | CD630_n00240 | 178↔190 | 106↔118 | -9.68411 | 4.49207 | 5.82709 |
| CD630_19903 | CD630_n00240 | 19↔90 | 3↔77 | -9.65651 | 5.79994 | 5.82709 |
| CD630_04620 | CD630_n00240 | 152↔171 | 4↔23 | -9.65515 | 3.45413 | 5.82709 |
| CD630_31461 | CD630_n00240 | 82↔175 | 2↔82 | -9.65123 | 4.41476 | 5.82709 |
| CD630_13560 | CD630_n00240 | 15↔25 | 5↔15 | -9.59164 | 3.38083 | 5.82709 |
| CD630_07840 | CD630_n00240 | 83↔94 | 4↔15 | -9.5761 | 4.26761 | 5.82709 |
| CD630_06350 | CD630_n00240 | 18↔36 | 13↔31 | -9.55256 | 4.35983 | 5.82709 |
| CD630_21460 | CD630_n00240 | 72↔81 | 7↔16 | -9.53615 | 4.83005 | 5.82709 |
| CD630_06560 | CD630_n00240 | 83↔115 | 2↔31 | -9.53578 | 4.02532 | 5.82709 |
| CD630_25320 | CD630_n00240 | 152↔172 | 4↔25 | -9.52658 | 4.29415 | 5.82709 |
| CD630_06340 | CD630_n00240 | 76↔94 | 4↔24 | -9.51658 | 4.35983 | 5.82709 |
| CD630_13350 | CD630_n00240 | 23↔42 | 101↔120 | -9.51332 | 4.15139 | 5.82709 |
| CD630_16670 | CD630_n00380 | 106↔157 | 307↔344 | -19.9028 | 3.33553 | 4.27916 |
| CD630_07760 | CD630_n00380 | 73↔109 | 871↔915 | -19.3754 | 3.64186 | 4.27916 |
| nusB | CD630_n00380 | 123↔186 | 839↔907 | -19.0833 | 4.92798 | 4.27916 |
| CD630_21620 | CD630_n00380 | 50↔189 | 872↔992 | -18.9529 | 3.02649 | 4.27916 |
| CD630_31000 | CD630_n00380 | 101↔152 | 859↔897 | -18.9114 | 3.70946 | 4.27916 |
| terD | CD630_n00380 | 1↔65 | 759↔822 | -18.8389 | 3.33148 | 4.27916 |
| CD630_18610 | CD630_n00380 | 14↔155 | 789↔913 | -18.7996 | 4.15582 | 4.27916 |
| moeA | CD630_n00380 | 49↔123 | 859↔933 | -18.7461 | 3.26118 | 4.27916 |
| CD630_24590 | CD630_n00380 | 2↔144 | 826↔974 | -18.6448 | 3.70401 | 4.27916 |
| CD630_19942 | CD630_n00380 | 83↔145 | 828↔900 | -18.6053 | 4.24574 | 4.27916 |
| gcvTPA | CD630_n00380 | 37↔120 | 303↔375 | -18.5996 | 4.77229 | 4.27916 |
| CD630_03170 | CD630_n00380 | 119↔192 | 755↔820 | -18.5954 | 3.66946 | 4.27916 |
| CD630_32710 | CD630_n00380 | 124↔167 | 294↔352 | -18.4994 | 4.74276 | 4.27916 |
| CD630_25620 | CD630_n00380 | 89↔201 | 788↔895 | -18.4939 | 3.02685 | 4.27916 |
| CD630_15080 | CD630_n00380 | 22↔111 | 299↔387 | -18.3041 | 4.03033 | 4.27916 |
| CD630_02430 | CD630_n00380 | 67↔141 | 311↔380 | -18.1958 | 3.71048 | 4.27916 |
| CD630_10880 | CD630_n00380 | 148↔200 | 286↔345 | -18.0472 | 5.05048 | 4.27916 |
| CD630_19401 | CD630_n00380 | 21↔64 | 875↔916 | -17.9061 | 4.37084 | 4.27916 |
| CD630_16450 | CD630_n00380 | 20↔110 | 302↔391 | -17.5987 | 4.57547 | 4.27916 |
| CD630_34030 | CD630_n00380 | 23↔78 | 885↔935 | -17.53 | 3.05766 | 4.27916 |
| CD630_22070 | CD630_n00380 | 85↔170 | 370↔445 | -17.4329 | 4.09651 | 4.27916 |
| srlEb | CD630_n00380 | 65↔97 | 305↔339 | -17.4218 | 3.50140 | 4.27916 |
| CD630_13370 | CD630_n00380 | 103↔168 | 155↔210 | -17.4026 | 4.33145 | 4.27916 |
| CD630_31490 | CD630_n00380 | 59↔125 | 312↔385 | -17.3589 | 4.08593 | 4.27916 |
| appF | CD630_n00380 | 52↔122 | 296↔383 | -17.3484 | 4.25570 | 4.27916 |
| CD630_03040 | CD630_n00380 | 84↔127 | 889↔924 | -17.0506 | 3.51403 | 4.27916 |
| CD630_28181 | CD630_n00380 | 4↔76 | 280↔350 | -16.9484 | 3.91299 | 4.27916 |
| CD630_17170 | CD630_n00380 | 115↔150 | 872↔907 | -16.9198 | 4.10230 | 4.27916 |
| garK | CD630_n00380 | 174↔191 | 301↔319 | -16.8243 | 3.72836 | 4.27916 |
| CD630_08080 | CD630_n00380 | 2↔65 | 627↔691 | -16.785 | 4.87819 | 4.27916 |
| CD630_11241 | CD630_n00380 | 158↔195 | 323↔355 | -16.7205 | 4.85723 | 4.27916 |
| CD630_22500 | CD630_n00380 | 115↔150 | 41↔74 | -16.6901 | 4.03688 | 4.27916 |
| CD630_14110 | CD630_n00380 | 5↔41 | 157↔197 | -16.6892 | 3.39550 | 4.27916 |
| CD630_20171 | CD630_n00380 | 71↔154 | 316↔410 | -16.6464 | 6.87491 | 4.27916 |
| CD630_04850 | CD630_n00380 | 165↔198 | 642↔676 | -16.5754 | 4.40788 | 4.27916 |
| CD630_20450 | CD630_n00380 | 110↔165 | 833↔897 | -16.4224 | 3.77388 | 4.27916 |
| opuCC | CD630_n00380 | 102↔198 | 575↔699 | -16.3896 | 4.07035 | 4.27916 |
| spaK | CD630_n00380 | 72↔110 | 871↔915 | -16.3061 | 3.04530 | 4.27916 |
| CD630_25610 | CD630_n00380 | 111↔153 | 875↔917 | -16.2571 | 3.02685 | 4.27916 |
| CD630_08940 | CD630_n00380 | 45↔123 | 871↔933 | -16.2351 | 3.72225 | 4.27916 |
| CD630_19700 | CD630_n00380 | 48↔95 | 435↔477 | -16.2311 | 3.85861 | 4.27916 |
| CD630_27280 | CD630_n00380 | 136↔197 | 666↔723 | -16.1861 | 3.24738 | 4.27916 |
| CD630_21260 | CD630_n00380 | 121↔143 | 658↔682 | -16.1852 | 4.22255 | 4.27916 |
| CD630_34520 | CD630_n00380 | 88↔161 | 835↔893 | -16.096 | 3.83016 | 4.27916 |
| ung | CD630_n00380 | 100↔141 | 305↔356 | -16.0365 | 3.30908 | 4.27916 |
| CD630_19690 | CD630_n00380 | 29↔130 | 297↔393 | -16.0333 | 3.85861 | 4.27916 |
| CD630_15900 | CD630_n00380 | 140↔190 | 628↔679 | -15.9338 | 3.17664 | 4.27916 |
| CD630_06060 | CD630_n00380 | 129↔144 | 738↔753 | -15.9265 | 4.94050 | 4.27916 |
| glgB | CD630_n00380 | 100↔172 | 844↔918 | -15.8817 | 4.48273 | 4.27916 |
| CD630_19790 | CD630_n00380 | 21↔157 | 371↔507 | -15.8416 | 3.79593 | 4.27916 |
| CD630_29500 | CD630_n00380 | 47↔139 | 295↔394 | -15.7942 | 3.70644 | 4.27916 |
| CD630_21570 | CD630_n00380 | 39↔108 | 305↔365 | -15.6988 | 6.48385 | 4.27916 |
| CD630_04410 | CD630_n00380 | 142↔201 | 828↔893 | -15.6522 | 3.90451 | 4.27916 |
| CD630_36690 | CD630_n00380 | 140↔197 | 176↔240 | -15.6462 | 3.53761 | 4.27916 |
| CD630_29650 | CD630_n00380 | 34↔129 | 293↔393 | -15.6092 | 3.67128 | 4.27916 |
| oppB | CD630_n00380 | 46↔102 | 186↔242 | -15.6075 | 2.98328 | 4.27916 |
| etfA | CD630_n00380 | 146↔183 | 434↔474 | -15.5282 | 3.41204 | 4.27916 |
| cbiO | CD630_n00380 | 91↔107 | 875↔897 | -15.4871 | 3.79830 | 4.27916 |
| oppC | CD630_n00380 | 106↔156 | 872↔929 | -15.4297 | 3.01695 | 4.27916 |
| CD630_23000 | CD630_n00380 | 1↔53 | 451↔503 | -15.4268 | 3.54130 | 4.27916 |
| CD630_26630 | CD630_n00380 | 58↔193 | 65↔184 | -15.4139 | 4.61557 | 4.27916 |
| CD630_19921 | CD630_n00380 | 145↔198 | 859↔919 | -15.3643 | 5.46345 | 4.27916 |
| CD630_10970 | CD630_n00380 | 3↔146 | 739↔880 | -15.3411 | 3.53850 | 4.27916 |
| CD630_14920 | CD630_n00380 | 87↔107 | 891↔913 | -15.2906 | 4.35225 | 4.27916 |
| CD630_30730 | CD630_n00380 | 8↔36 | 165↔190 | -15.2852 | 3.13236 | 4.27916 |
| CD630_21090 | CD630_n00380 | 111↔183 | 757↔814 | -15.2555 | 4.15803 | 4.27916 |
| CD630_32690 | CD630_n00380 | 122↔159 | 874↔915 | -15.2403 | 7.32676 | 4.27916 |
| CD630_05820 | CD630_n00380 | 65↔93 | 312↔347 | -15.175 | 3.75661 | 4.27916 |
| CD630_13960 | CD630_n00380 | 130↔156 | 441↔465 | -15.1683 | 3.30375 | 4.27916 |
| CD630_18970 | CD630_n00380 | 86↔158 | 859↔922 | -15.1481 | 3.57243 | 4.27916 |
| CD630_08140 | CD630_n00380 | 6↔55 | 771↔812 | -15.1188 | 3.30159 | 4.27916 |
| CD630_18100 | CD630_n00380 | 11↔148 | 824↔968 | -15.0943 | 5.71658 | 4.27916 |
| CD630_18930 | CD630_n00380 | 13↔53 | 674↔713 | -15.0032 | 5.32127 | 4.27916 |
| CD630_06340 | CD630_n00380 | 68↔109 | 874↔917 | -14.9575 | 4.35983 | 4.27916 |
| CD630_17511 | CD630_n00380 | 3↔60 | 159↔215 | -14.9488 | 9.12322 | 4.27916 |
| CD630_19850 | CD630_n00380 | 90↔115 | 441↔465 | -14.918 | 6.88486 | 4.27916 |
| CD630_28280 | CD630_n00380 | 21↔116 | 618↔721 | -14.9116 | 4.59833 | 4.27916 |
| CD630_28550 | CD630_n00380 | 68↔106 | 876↔920 | -14.8849 | 3.51196 | 4.27916 |
| CD630_09760 | CD630_n00380 | 111↔144 | 875↔915 | -14.7982 | 4.83410 | 4.27916 |
| CD630_28910 | CD630_n00380 | 111↔144 | 875↔915 | -14.7982 | 4.51664 | 4.27916 |
| CD630_06540 | CD630_n00380 | 127↔195 | 24↔94 | -14.7776 | 3.54673 | 4.27916 |
| CD630_32990 | CD630_n00380 | 22↔54 | 694↔728 | -14.7589 | 5.24925 | 4.27916 |
| cobQ | CD630_n00380 | 88↔110 | 888↔908 | -14.7101 | 3.13524 | 4.27916 |
| CD630_08100 | CD630_n00380 | 87↔135 | 834↔895 | -14.7014 | 4.39181 | 4.27916 |
| rolB | CD630_n00380 | 79↔144 | 399↔453 | -14.6257 | 3.60270 | 4.27916 |
| CD630_04610 | CD630_n00380 | 143↔201 | 663↔726 | -14.5914 | 3.83025 | 4.27916 |
| CD630_19030 | CD630_n00380 | 109↔198 | 637↔720 | -14.5726 | 3.34240 | 4.27916 |
| CD630_05950 | CD630_n00380 | 29↔63 | 305↔344 | -14.5698 | 3.31669 | 4.27916 |
| chrA | CD630_n00380 | 1↔52 | 758↔812 | -14.5431 | 5.27255 | 4.27916 |
| CD630_27570 | CD630_n00380 | 84↔194 | 807↔926 | -14.5067 | 3.46216 | 4.27916 |
| CD630_06160 | CD630_n00380 | 164↔199 | 165↔201 | -14.487 | 5.51744 | 4.27916 |
| CD630_21760 | CD630_n00380 | 173↔199 | 157↔185 | -14.4617 | 4.43846 | 4.27916 |
| CD630_01590 | CD630_n00380 | 2↔28 | 655↔677 | -14.4573 | 3.45609 | 4.27916 |
| CD630_27110 | CD630_n00380 | 174↔199 | 154↔181 | -14.442 | 3.78231 | 4.27916 |
| CD630_28290 | CD630_n00380 | 25↔74 | 760↔802 | -14.4328 | 5.54192 | 4.27916 |
| CD630_21730 | CD630_n00380 | 166↔193 | 149↔176 | -14.4144 | 3.45076 | 4.27916 |
| CD630_15360 | CD630_n00380 | 36↔106 | 303↔370 | -14.3487 | 4.47208 | 4.27916 |
| CD630_06080 | CD630_n00380 | 45↔104 | 285↔346 | -14.3418 | 3.82315 | 4.27916 |
| CD630_28150 | CD630_n00380 | 21↔53 | 309↔347 | -14.3192 | 4.18447 | 4.27916 |
| CD630_16330 | CD630_n00380 | 149↔200 | 158↔210 | -14.3084 | 4.11180 | 4.27916 |
| CD630_32700 | CD630_n00380 | 102↔119 | 437↔454 | -14.2673 | 4.74276 | 4.27916 |
| CD630_21240 | CD630_n00380 | 121↔152 | 699↔725 | -14.2657 | 5.88879 | 4.27916 |
| CD630_15200 | CD630_n00380 | 8↔138 | 594↔711 | -14.2517 | 4.00442 | 4.27916 |
| CD630_06970 | CD630_n00380 | 83↔151 | 829↔896 | -14.239 | 3.82312 | 4.27916 |
| kdpD | CD630_n00380 | 141↔199 | 871↔913 | -14.2351 | 4.27140 | 4.27916 |
| moaA | CD630_n00380 | 51↔120 | 439↔487 | -14.2338 | 3.83799 | 4.27916 |
| CD630_21140 | CD630_n00380 | 128↔195 | 816↔882 | -14.1954 | 3.33304 | 4.27916 |
| fbp | CD630_n00380 | 172↔189 | 709↔729 | -14.1864 | 3.80071 | 4.27916 |
| CD630_14360 | CD630_n00380 | 40↔116 | 865↔929 | -14.1674 | 3.02079 | 4.27916 |
| CD630_27610 | CD630_n00380 | 11↔29 | 373↔391 | -14.0759 | 3.96465 | 4.27916 |
| CD630_01700 | CD630_n00380 | 1↔50 | 157↔206 | -14.0537 | 4.97234 | 4.27916 |
| pilC | CD630_n00380 | 11↔110 | 302↔382 | -14.0361 | 3.54230 | 4.27916 |
| CD630_22550 | CD630_n00380 | 125↔196 | 149↔238 | -14.0346 | 3.62379 | 4.27916 |
| CD630_31010 | CD630_n00380 | 133↔158 | 856↔882 | -14.0279 | 4.96305 | 4.27916 |
| CD630_20960 | CD630_n00380 | 50↔111 | 303↔357 | -14.0124 | 5.65940 | 4.27916 |
| CD630_17710 | CD630_n00380 | 86↔109 | 872↔898 | -14.0018 | 4.37688 | 4.27916 |
| dltB | CD630_n00380 | 1↔34 | 72↔110 | -13.976 | 3.81485 | 4.27916 |
| eutW | CD630_n00380 | 129↔144 | 163↔178 | -13.9663 | 4.55715 | 4.27916 |
| CD630_17360 | CD630_n00380 | 89↔123 | 314↔339 | -13.964 | 5.66070 | 4.27916 |
| CD630_26920 | CD630_n00380 | 33↔57 | 156↔180 | -13.9528 | 4.58060 | 4.27916 |
| CD630_13880 | CD630_n00380 | 111↔158 | 140↔181 | -13.9478 | 3.52735 | 4.27916 |
| CD630_19840 | CD630_n00380 | 133↔178 | 580↔630 | -13.9389 | 3.44724 | 4.27916 |
| CD630_05000 | CD630_n00380 | 110↔159 | 833↔878 | -13.9297 | 4.02442 | 4.27916 |
| cdu2 | CD630_n00380 | 36↔162 | 304↔416 | -13.9047 | 2.97335 | 4.27916 |
| CD630_22740 | CD630_n00380 | 62↔119 | 372↔423 | -13.9012 | 3.08733 | 4.27916 |
| CD630_09772 | CD630_n00380 | 69↔92 | 902↔925 | -13.8969 | 3.66033 | 4.27916 |
| CD630_32520 | CD630_n00380 | 166↔201 | 886↔919 | -13.8898 | 4.06098 | 4.27916 |
| CD630_30120 | CD630_n00380 | 70↔134 | 858↔918 | -13.8806 | 4.04859 | 4.27916 |
| cobC | CD630_n00380 | 165↔195 | 901↔929 | -13.8566 | 3.15854 | 4.27916 |
| CD630_06050 | CD630_n00380 | 89↔166 | 295↔382 | -13.8301 | 3.59332 | 4.27916 |
| CD630_05450 | CD630_n00380 | 83↔103 | 879↔899 | -13.822 | 5.04761 | 4.27916 |
| CD630_35260 | CD630_n00380 | 171↔188 | 436↔454 | -13.7314 | 5.77277 | 4.27916 |
| CD630_13950 | CD630_n00380 | 149↔201 | 660↔710 | -13.7019 | 4.25540 | 4.27916 |
| bcp | CD630_n00380 | 89↔151 | 241↔309 | -13.701 | 3.16670 | 4.27916 |
| uxaA%27 | CD630_n00380 | 88↔160 | 801↔893 | -13.6809 | 4.26165 | 4.27916 |
| CD630_14630 | CD630_n00380 | 55↔134 | 863↔933 | -13.6768 | 3.34485 | 4.27916 |
| CD630_05460 | CD630_n00380 | 51↔93 | 900↔940 | -13.6761 | 3.98753 | 4.27916 |
| CD630_22030 | CD630_n00380 | 156↔198 | 834↔892 | -13.6365 | 3.29934 | 4.27916 |
| CD630_14930 | CD630_n00380 | 2↔43 | 609↔641 | -13.6042 | 3.31335 | 4.27916 |
| CD630_13530 | CD630_n00380 | 5↔31 | 676↔701 | -13.6027 | 4.09832 | 4.27916 |
| CD630_16150 | CD630_n00380 | 5↔75 | 380↔452 | -13.5995 | 4.20614 | 4.27916 |
| cbpA | CD630_n00380 | 75↔107 | 872↔913 | -13.5839 | 3.03040 | 4.27916 |
| xdhA | CD630_n00380 | 99↔150 | 859↔926 | -13.5696 | 4.45147 | 4.27916 |
| CD630_05821 | CD630_n00380 | 176↔199 | 792↔815 | -13.5593 | 3.75661 | 4.27916 |
| CD630_14130 | CD630_n00380 | 89↔138 | 872↔922 | -13.5105 | 4.57970 | 4.27916 |
| CD630_23820 | CD630_n00380 | 44↔109 | 873↔926 | -13.4666 | 3.18419 | 4.27916 |
| CD630_23780 | CD630_n00380 | 5↔54 | 623↔672 | -13.4381 | 3.91629 | 4.27916 |
| CD630_32040 | CD630_n00380 | 22↔41 | 298↔318 | -13.4371 | 4.93516 | 4.27916 |
| CD630_25320 | CD630_n00380 | 142↔153 | 885↔896 | -13.4154 | 4.29415 | 4.27916 |
| appC | CD630_n00380 | 43↔76 | 196↔231 | -13.4061 | 4.50438 | 4.27916 |
| CD630_20980 | CD630_n00380 | 78↔152 | 312↔410 | -13.3979 | 8.12847 | 4.27916 |
| CD630_27230 | CD630_n00380 | 6↔26 | 858↔879 | -13.3958 | 3.52462 | 4.27916 |
| dapB2 | CD630_n00380 | 136↔160 | 885↔911 | -13.3876 | 3.27218 | 4.27916 |
| CD630_13350 | CD630_n00380 | 158↔199 | 224↔265 | -13.3843 | 4.15139 | 4.27916 |
| CD630_27210 | CD630_n00380 | 118↔150 | 485↔518 | -13.3746 | 4.18786 | 4.27916 |
| CD630_10950 | CD630_n00380 | 30↔106 | 305↔380 | -13.3552 | 3.67675 | 4.27916 |
| CD630_24580 | CD630_n00380 | 11↔46 | 140↔177 | -13.3238 | 3.88519 | 4.27916 |
| scpB | CD630_n00380 | 92↔151 | 862↔918 | -13.3052 | 3.33560 | 4.27916 |
| def1 | CD630_n00380 | 86↔188 | 838↔897 | -13.3035 | 3.99720 | 4.27916 |
| CD630_31210 | CD630_n00380 | 3↔69 | 619↔678 | -13.2859 | 4.17693 | 4.27916 |
| CD630_14260 | CD630_n00380 | 22↔68 | 137↔170 | -13.2774 | 3.22214 | 4.27916 |
| CD630_27630 | CD630_n00380 | 1↔73 | 604↔675 | -13.2625 | 3.54199 | 4.27916 |

| | | | | | | |
|---|---|---|---|---|---|---|
| CD630_23080 | CD630_n00380 | 1 -- 56 | 162 -- 214 | -13.2518 | 5.23027 | 4.27916 |
| argG | CD630_n00380 | 70 -- 107 | 305 -- 336 | -13.2492 | 4.80300 | 4.27916 |
| cwp29 | CD630_n00380 | 89 -- 137 | 848 -- 895 | -13.2463 | 4.40865 | 4.27916 |
| CD630_21610 | CD630_n00380 | 156 -- 171 | 684 -- 699 | -13.137 | 3.16662 | 4.27916 |
| CD630_06380 | CD630_n00380 | 147 -- 201 | 643 -- 696 | -13.1264 | 4.32211 | 4.27916 |
| CD630_27270 | CD630_n00380 | 158 -- 174 | 877 -- 897 | -13.1192 | 3.50892 | 4.27916 |
| CD630_25300 | CD630_n00380 | 73 -- 154 | 135 -- 211 | -13.1104 | 4.49333 | 4.27916 |
| CD630_16460 | CD630_n00380 | 91 -- 160 | 827 -- 893 | -13.0767 | 4.64028 | 4.27916 |
| CD630_23930 | CD630_n00380 | 71 -- 200 | 789 -- 933 | -13.0615 | 3.05650 | 4.27916 |
| CD630_13780 | CD630_n00380 | 85 -- 130 | 296 -- 345 | -13.0479 | 3.52274 | 4.27916 |
| CD630_07840 | CD630_n00380 | 114 -- 154 | 432 -- 471 | -13.0162 | 4.26761 | 4.27916 |
| dnaD | CD630_n00380 | 114 -- 172 | 163 -- 211 | -13.0132 | 3.56078 | 4.27916 |
| CD630_34530 | CD630_n00380 | 65 -- 121 | 874 -- 931 | -12.9866 | 3.83016 | 4.27916 |
| CD630_21770 | CD630_n00380 | 52 -- 112 | 870 -- 933 | -12.8947 | 4.02587 | 4.27916 |
| CD630_30130 | CD630_n00380 | 31 -- 76 | 874 -- 922 | -12.8796 | 3.28343 | 4.27916 |
| CD630_15850 | CD630_n00380 | 121 -- 197 | 853 -- 912 | -12.8389 | 4.32968 | 4.27916 |
| CD630_24760 | CD630_n00380 | 55 -- 161 | 299 -- 407 | -12.8313 | 5.19485 | 4.27916 |
| CD630_10830 | CD630_n00380 | 66 -- 104 | 878 -- 916 | -12.8219 | 4.82398 | 4.27916 |
| CD630_13600 | CD630_n00380 | 35 -- 165 | 305 -- 433 | -12.8055 | 4.45481 | 4.27916 |
| CD630_21700 | CD630_n00380 | 121 -- 139 | 155 -- 174 | -12.8023 | 3.96650 | 4.27916 |
| CD630_19430 | CD630_n00380 | 88 -- 116 | 866 -- 895 | -12.7372 | 3.13922 | 4.27916 |
| malY | CD630_n00380 | 4 -- 18 | 371 -- 384 | -12.7149 | 4.46345 | 4.27916 |
| CD630_16750 | CD630_n00380 | 4 -- 36 | 631 -- 666 | -12.7133 | 5.04051 | 4.27916 |
| CD630_18650 | CD630_n00380 | 29 -- 98 | 312 -- 389 | -12.7091 | 3.75892 | 4.27916 |
| CD630_31580 | CD630_n00380 | 85 -- 114 | 873 -- 897 | -12.702 | 3.51756 | 4.27916 |
| CD630_30140 | CD630_n00380 | 15 -- 62 | 838 -- 882 | -12.6995 | 3.28343 | 4.27916 |
| srlD | CD630_n00380 | 88 -- 104 | 878 -- 895 | -12.6558 | 4.19843 | 4.27916 |
| CD630_05230 | CD630_n00380 | 127 -- 169 | 195 -- 244 | -12.6443 | 3.73787 | 4.27916 |
| eutV | CD630_n00380 | 102 -- 135 | 871 -- 896 | -12.6313 | 4.55715 | 4.27916 |
| CD630_06060 | CD630_n00380 | 172 -- 193 | 867 -- 896 | -12.5974 | 4.94050 | 4.27916 |
| ssuC | CD630_n00380 | 47 -- 83 | 885 -- 918 | -12.5856 | 3.45491 | 4.27916 |
| CD630_06100 | CD630_n00380 | 137 -- 201 | 660 -- 701 | -12.5806 | 4.43918 | 4.27916 |
| CD630_09750 | CD630_n00380 | 89 -- 128 | 887 -- 921 | -12.5647 | 4.83410 | 4.27916 |
| CD630_28920 | CD630_n00380 | 89 -- 128 | 887 -- 921 | -12.5647 | 4.51664 | 4.27916 |
| CD630_25050 | CD630_n00380 | 1 -- 67 | 615 -- 676 | -12.5075 | 4.33624 | 4.27916 |
| CD630_32050 | CD630_n00380 | 132 -- 177 | 727 -- 765 | -12.4965 | 5.73797 | 4.27916 |
| CD630_13020 | CD630_n00380 | 117 -- 168 | 230 -- 260 | -12.4818 | 3.52267 | 4.27916 |
| CD630_09770 | CD630_n00380 | 155 -- 181 | 23 -- 50 | -12.472 | 3.22106 | 4.27916 |
| CD630_25310 | CD630_n00380 | 22 -- 31 | 441 -- 450 | -12.4706 | 3.99861 | 4.27916 |
| CD630_25020 | CD630_n00380 | 12 -- 53 | 644 -- 686 | -12.4192 | 4.17721 | 4.27916 |
| CD630_04550 | CD630_n00380 | 94 -- 136 | 178 -- 216 | -12.4017 | 3.05227 | 4.27916 |
| CD630_18090 | CD630_n00380 | 81 -- 168 | 337 -- 450 | -12.3592 | 4.23961 | 4.27916 |
| CD630_19000 | CD630_n00380 | 54 -- 136 | 304 -- 373 | -12.3513 | 4.18443 | 4.27916 |
| CD630_05830 | CD630_n00380 | 134 -- 183 | 668 -- 722 | -12.3496 | 3.35360 | 4.27916 |
| CD630_04940 | CD630_n00380 | 89 -- 100 | 884 -- 895 | -12.3351 | 4.20948 | 4.27916 |
| CD630_05490 | CD630_n00380 | 137 -- 178 | 295 -- 332 | -12.322 | 3.89852 | 4.27916 |
| bglA | CD630_n00380 | 151 -- 201 | 875 -- 922 | -12.3197 | 4.86402 | 4.27916 |
| CD630_06170 | CD630_n00380 | 102 -- 168 | 549 -- 629 | -12.3126 | 4.95366 | 4.27916 |
| CD630_04280 | CD630_n00380 | 156 -- 169 | 316 -- 328 | -12.3011 | 3.61457 | 4.27916 |
| CD630_19680 | CD630_n00380 | 39 -- 97 | 335 -- 395 | -12.3011 | 3.58080 | 4.27916 |
| argS | CD630_n00380 | 176 -- 195 | 257 -- 274 | -12.2762 | 4.96036 | 4.27916 |
| CD630_05822 | CD630_n00380 | 172 -- 201 | 802 -- 834 | -12.2754 | 4.28113 | 4.27916 |
| pgm | CD630_n00380 | 16 -- 43 | 293 -- 316 | -12.264 | 3.45439 | 4.27916 |
| pilT | CD630_n00380 | 102 -- 180 | 599 -- 677 | -12.2523 | 3.24639 | 4.27916 |
| CD630_07870 | CD630_n00380 | 159 -- 200 | 758 -- 809 | -12.2502 | 3.22829 | 4.27916 |
| CD630_23770 | CD630_n00380 | 59 -- 201 | 281 -- 422 | -12.2349 | 3.25181 | 4.27916 |
| CD630_31500 | CD630_n00380 | 135 -- 168 | 805 -- 843 | -12.2327 | 4.90207 | 4.27916 |
| CD630_06110 | CD630_n00380 | 170 -- 186 | 892 -- 908 | -12.2267 | 5.61380 | 4.27916 |
| CD630_33420 | CD630_n00380 | 142 -- 159 | 706 -- 723 | -12.1676 | 3.22095 | 4.27916 |
| panG | CD630_n00380 | 161 -- 199 | 743 -- 776 | -12.1658 | 5.54047 | 4.27916 |
| CD630_32070 | CD630_n00380 | 158 -- 179 | 710 -- 729 | -12.1466 | 3.51964 | 4.27916 |
| CD630_15550 | CD630_n00380 | 85 -- 139 | 860 -- 911 | -12.1381 | 3.49019 | 4.27916 |
| CD630_05880 | CD630_n00380 | 104 -- 174 | 872 -- 955 | -12.138 | 4.28339 | 4.27916 |
| CD630_20170 | CD630_n00380 | 99 -- 119 | 312 -- 332 | -12.1366 | 6.90201 | 4.27916 |
| CD630_22050 | CD630_n00380 | 134 -- 191 | 759 -- 820 | -12.1354 | 4.87639 | 4.27916 |
| CD630_09040 | CD630_n00380 | 23 -- 53 | 835 -- 860 | -12.1326 | 4.11851 | 4.27916 |
| CD630_30420 | CD630_n00380 | 137 -- 175 | 658 -- 696 | -12.0895 | 5.27539 | 4.27916 |
| CD630_19580 | CD630_n00380 | 63 -- 95 | 888 -- 922 | -12.0892 | 4.51902 | 4.27916 |
| kdgT1 | CD630_n00380 | 67 -- 111 | 869 -- 931 | -12.0629 | 4.49207 | 4.27916 |
| CD630_18010 | CD630_n00380 | 175 -- 201 | 672 -- 697 | -12.0301 | 4.21967 | 4.27916 |
| CD630_05870 | CD630_n00380 | 158 -- 174 | 875 -- 891 | -12.0106 | 4.28339 | 4.27916 |
| CD630_10080 | CD630_n00380 | 9 -- 51 | 617 -- 665 | -12.0073 | 3.75625 | 4.27916 |
| CD630_16520 | CD630_n00380 | 148 -- 193 | 837 -- 893 | -11.9742 | 4.28395 | 4.27916 |
| CD630_13560 | CD630_n00380 | 131 -- 179 | 834 -- 882 | -11.9709 | 3.38083 | 4.27916 |
| hfq | CD630_n00380 | 150 -- 197 | 840 -- 893 | -11.962 | 5.05609 | 4.27916 |
| CD630_08130 | CD630_n00380 | 13 -- 96 | 297 -- 381 | -11.9518 | 4.63831 | 4.27916 |
| CD630_17292 | CD630_n00380 | 93 -- 149 | 873 -- 918 | -11.9503 | 6.25799 | 4.27916 |
| CD630_23861 | CD630_n00380 | 17 -- 41 | 159 -- 178 | -11.9495 | 5.83068 | 4.27916 |
| CD630_14710 | CD630_n00380 | 60 -- 106 | 734 -- 782 | -11.9468 | 3.65171 | 4.27916 |
| CD630_15060 | CD630_n00380 | 170 -- 196 | 166 -- 186 | -11.9458 | 3.69396 | 4.27916 |
| CD630_23440 | CD630_n00380 | 53 -- 80 | 875 -- 913 | -11.9383 | 3.34841 | 4.27916 |
| CD630_27310 | CD630_n00380 | 78 -- 161 | 828 -- 907 | -11.9122 | 4.01622 | 4.27916 |
| cspC | CD630_n00380 | 160 -- 177 | 875 -- 893 | -11.9041 | 5.48848 | 4.27916 |
| CD630_32720 | CD630_n00380 | 113 -- 151 | 430 -- 476 | -11.903 | 4.74276 | 4.27916 |
| CD630_12320 | CD630_n00380 | 50 -- 105 | 877 -- 926 | -11.9018 | 3.28667 | 4.27916 |
| CD630_10050 | CD630_n00380 | 154 -- 163 | 886 -- 895 | -11.8978 | 3.93750 | 4.27916 |
| CD630_22880 | CD630_n00380 | 39 -- 64 | 155 -- 179 | -11.8845 | 3.62455 | 4.27916 |
| CD630_10000 | CD630_n00380 | 82 -- 113 | 727 -- 755 | -11.8843 | 3.86040 | 4.27916 |
| CD630_31220 | CD630_n00380 | 165 -- 195 | 235 -- 261 | -11.8649 | 3.73779 | 4.27916 |
| CD630_14240 | CD630_n00380 | 70 -- 133 | 859 -- 910 | -11.8608 | 8.60766 | 4.27916 |
| CD630_09840 | CD630_n00380 | 105 -- 130 | 159 -- 183 | -11.827 | 4.50009 | 4.27916 |
| iorA | CD630_n00380 | 2 -- 22 | 726 -- 746 | -11.8259 | 3.83173 | 4.27916 |
| CD630_03160 | CD630_n00380 | 67 -- 80 | 439 -- 452 | -11.8188 | 3.66946 | 4.27916 |
| uxaA | CD630_n00380 | 74 -- 98 | 301 -- 318 | -11.796 | 4.26165 | 4.27916 |
| CD630_02740 | CD630_n00380 | 140 -- 162 | 875 -- 894 | -11.7888 | 5.02277 | 4.27916 |
| CD630_08370 | CD630_n00380 | 112 -- 155 | 306 -- 347 | -11.7868 | 3.65639 | 4.27916 |
| CD630_18960 | CD630_n00380 | 11 -- 55 | 886 -- 917 | -11.775 | 4.37822 | 4.27916 |
| CD630_28320 | CD630_n00380 | 87 -- 132 | 267 -- 319 | -11.7745 | 3.99184 | 4.27916 |
| CD630_18410 | CD630_n00380 | 123 -- 172 | 158 -- 203 | -11.7644 | 3.59739 | 4.27916 |
| CD630_16550 | CD630_n00380 | 80 -- 107 | 305 -- 345 | -11.7541 | 5.61064 | 4.27916 |
| CD630_20670 | CD630_n00380 | 151 -- 159 | 890 -- 898 | -11.7506 | 4.42757 | 4.27916 |
| CD630_10960 | CD630_n00380 | 11 -- 40 | 306 -- 334 | -11.7336 | 3.53850 | 4.27916 |
| CD630_28260 | CD630_n00380 | 39 -- 108 | 309 -- 382 | -11.7211 | 3.51613 | 4.27916 |
| CD630_19780 | CD630_n00380 | 2 -- 18 | 377 -- 393 | -11.7195 | 3.64830 | 4.27916 |
| CD630_24050 | CD630_n00380 | 5 -- 52 | 618 -- 674 | -11.7158 | 3.70192 | 4.27916 |
| CD630_14810 | CD630_n00380 | 54 -- 65 | 314 -- 325 | -11.7024 | 3.35323 | 4.27916 |
| CD630_31461 | CD630_n00380 | 141 -- 201 | 161 -- 206 | -11.5869 | 4.41476 | 4.27916 |
| thiH | CD630_n00380 | 68 -- 99 | 895 -- 927 | -11.573 | 3.58230 | 4.27916 |
| CD630_02420 | CD630_n00380 | 3 -- 35 | 304 -- 339 | -11.5637 | 3.71048 | 4.27916 |
| CD630_30540 | CD630_n00380 | 123 -- 156 | 293 -- 319 | -11.5598 | 3.63804 | 4.27916 |
| CD630_21490 | CD630_n00380 | 149 -- 169 | 702 -- 722 | -11.551 | 3.37462 | 4.27916 |
| cwp26 | CD630_n00380 | 70 -- 88 | 878 -- 896 | -11.5349 | 4.44377 | 4.27916 |
| CD630_31690 | CD630_n00380 | 112 -- 160 | 163 -- 209 | -11.5343 | 3.71193 | 4.27916 |
| CD630_27200 | CD630_n00380 | 122 -- 137 | 223 -- 238 | -11.5262 | 3.42516 | 4.27916 |
| CD630_08070 | CD630_n00380 | 86 -- 125 | 863 -- 894 | -11.5261 | 4.87819 | 4.27916 |
| CD630_16160 | CD630_n00380 | 66 -- 111 | 870 -- 915 | -11.5237 | 6.11801 | 4.27916 |
| htpG | CD630_n00380 | 46 -- 93 | 901 -- 933 | -11.5019 | 3.45147 | 4.27916 |
| cdu1 | CD630_n00380 | 3 -- 35 | 443 -- 476 | -11.5003 | 4.46276 | 4.27916 |
| CD630_13790 | CD630_n00380 | 78 -- 118 | 888 -- 915 | -11.4958 | 5.17374 | 4.27916 |
| CD630_28870 | CD630_n00380 | 32 -- 73 | 463 -- 505 | -11.4836 | 3.41991 | 4.27916 |
| CD630_20600 | CD630_n00380 | 79 -- 124 | 862 -- 915 | -11.4664 | 4.20093 | 4.27916 |
| CD630_08460 | CD630_n00380 | 148 -- 166 | 433 -- 454 | -11.4482 | 4.15083 | 4.27916 |
| CD630_12110 | CD630_n00380 | 167 -- 191 | 646 -- 669 | -11.4476 | 4.36078 | 4.27916 |
| CD630_19904 | CD630_n00380 | 154 -- 200 | 227 -- 277 | -11.4443 | 5.96526 | 4.27916 |
| CD630_32060 | CD630_n00380 | 161 -- 189 | 602 -- 632 | -11.4404 | 5.73797 | 4.27916 |
| CD630_03220 | CD630_n00380 | 86 -- 135 | 303 -- 355 | -11.4121 | 3.36746 | 4.27916 |
| metQ | CD630_n00380 | 88 -- 104 | 878 -- 895 | -11.4097 | 3.87190 | 4.27916 |
| CD630_23280 | CD630_n00380 | 71 -- 140 | 872 -- 933 | -11.4051 | 3.50093 | 4.27916 |
| CD630_01690 | CD630_n00380 | 151 -- 171 | 761 -- 780 | -11.3978 | 4.97234 | 4.27916 |
| panB | CD630_n00380 | 153 -- 187 | 149 -- 181 | -11.3821 | 3.40248 | 4.27916 |
| CD630_04880 | CD630_n00380 | 122 -- 160 | 794 -- 834 | -11.3812 | 4.62291 | 4.27916 |
| CD630_29760 | CD630_n00380 | 134 -- 150 | 902 -- 918 | -11.3698 | 3.51074 | 4.27916 |
| CD630_26150 | CD630_n00380 | 19 -- 83 | 623 -- 698 | -11.3253 | 4.06302 | 4.27916 |
| CD630_36250 | CD630_n00380 | 53 -- 152 | 828 -- 926 | -11.3107 | 4.13224 | 4.27916 |
| CD630_05900 | CD630_n00380 | 116 -- 174 | 777 -- 821 | -11.3058 | 3.21934 | 4.27916 |
| CD630_13920 | CD630_n00380 | 25 -- 49 | 156 -- 179 | -11.2962 | 3.25236 | 4.27916 |
| CD630_18781 | CD630_n00380 | 139 -- 159 | 165 -- 183 | -11.2957 | 3.77660 | 4.27916 |
| CD630_07430 | CD630_n00380 | 3 -- 22 | 167 -- 186 | -11.2936 | 4.65568 | 4.27916 |
| dpaA | CD630_n00380 | 40 -- 54 | 141 -- 156 | -11.2918 | 4.65364 | 4.27916 |
| cspD | CD630_n00380 | 121 -- 178 | 874 -- 924 | -11.2913 | 3.23587 | 4.27916 |
| CD630_20460 | CD630_n00380 | 77 -- 100 | 901 -- 924 | -11.2905 | 7.94034 | 4.27916 |
| CD630_13400 | CD630_n00380 | 149 -- 180 | 796 -- 834 | -11.2647 | 3.20232 | 4.27916 |
| CD630_02790 | CD630_n00380 | 71 -- 134 | 315 -- 375 | -11.2629 | 4.77661 | 4.27916 |
| CD630_25040 | CD630_n00380 | 1 -- 53 | 837 -- 887 | -11.2618 | 3.43898 | 4.27916 |
| CD630_15740 | CD630_n00380 | 71 -- 97 | 303 -- 330 | -11.2574 | 4.43222 | 4.27916 |
| CD630_17281 | CD630_n00380 | 74 -- 151 | 189 -- 263 | -11.2397 | 5.91544 | 4.27916 |
| CD630_29250 | CD630_n00380 | 81 -- 135 | 272 -- 331 | -11.2373 | 4.61901 | 4.27916 |
| CD630_09410 | CD630_n00380 | 81 -- 135 | 272 -- 331 | -11.2373 | 4.75651 | 4.27916 |
| CD630_02830 | CD630_n00380 | 155 -- 174 | 735 -- 755 | -11.2338 | 3.57106 | 4.27916 |
| CD630_18311 | CD630_n00380 | 96 -- 181 | 792 -- 865 | -11.2285 | 6.86047 | 4.27916 |
| CD630_32880 | CD630_n00380 | 106 -- 196 | 144 -- 226 | -11.2145 | 3.41276 | 4.27916 |
| cls | CD630_n00380 | 168 -- 201 | 586 -- 624 | -11.1989 | 4.46177 | 4.27916 |
| CD630_27060 | CD630_n00380 | 102 -- 161 | 837 -- 896 | -11.1905 | 4.06049 | 4.27916 |
| CD630_08110 | CD630_n00380 | 106 -- 116 | 370 -- 380 | -11.1853 | 4.39181 | 4.27916 |
| CD630_04660 | CD630_n00380 | 73 -- 100 | 902 -- 929 | -11.1763 | 4.32791 | 4.27916 |
| CD630_26680 | CD630_n00380 | 70 -- 98 | 308 -- 339 | -11.161 | 3.53956 | 4.27916 |
| CD630_03930 | CD630_n00380 | 138 -- 166 | 650 -- 677 | -11.1564 | 4.15934 | 4.27916 |
| CD630_18920 | CD630_n00380 | 116 -- 189 | 305 -- 394 | -11.1294 | 4.43558 | 4.27916 |
| CD630_34050 | CD630_n00380 | 80 -- 151 | 829 -- 907 | -11.097 | 4.03863 | 4.27916 |
| CD630_27240 | CD630_n00380 | 147 -- 170 | 164 -- 186 | -11.0936 | 3.52462 | 4.27916 |
| CD630_14610 | CD630_n00380 | 152 -- 183 | 595 -- 627 | -11.0541 | 4.89428 | 4.27916 |
| mobA | CD630_n00380 | 149 -- 195 | 439 -- 476 | -11.0516 | 4.25703 | 4.27916 |
| CD630_25150 | CD630_n00380 | 99 -- 157 | 598 -- 674 | -11.0372 | 3.79339 | 4.27916 |
| CD630_26050 | CD630_n00380 | 68 -- 135 | 852 -- 918 | -11.0341 | 6.99851 | 4.27916 |
| CD630_06150 | CD630_n00380 | 144 -- 177 | 155 -- 186 | -11.0258 | 5.51744 | 4.27916 |
| CD630_26050 | CD630_n00380 | 171 -- 188 | 878 -- 897 | -10.998 | 6.99851 | 4.27916 |
| CD630_33580 | CD630_n00380 | 174 -- 201 | 622 -- 659 | -10.9928 | 3.95778 | 4.27916 |
| CD630_21070 | CD630_n00380 | 87 -- 98 | 309 -- 320 | -10.9704 | 5.35019 | 4.27916 |
| CD630_33600 | CD630_n00380 | 142 -- 174 | 303 -- 328 | -10.9635 | 3.73196 | 4.27916 |
| CD630_23760 | CD630_n00380 | 151 -- 200 | 215 -- 263 | -10.9193 | 4.54307 | 4.27916 |
| pilW | CD630_n00380 | 90 -- 104 | 878 -- 895 | -10.9144 | 4.96316 | 4.27916 |
| CD630_08360 | CD630_n00380 | 30 -- 66 | 312 -- 344 | -10.911 | 3.19339 | 4.27916 |
| CD630_33570 | CD630_n00380 | 132 -- 168 | 701 -- 739 | -10.8859 | 3.95778 | 4.27916 |
| pilD | CD630_n00380 | 35 -- 77 | 827 -- 867 | -10.8826 | 4.54123 | 4.27916 |
| CD630_07121 | CD630_n00380 | 44 -- 77 | 154 -- 184 | -10.8798 | 3.26177 | 4.27916 |
| CD630_08390 | CD630_n00380 | 87 -- 116 | 428 -- 454 | -10.8707 | 5.16000 | 4.27916 |
| CD630_24591 | CD630_n00380 | 59 -- 107 | 890 -- 933 | -10.8667 | 3.54497 | 4.27916 |
| CD630_09030 | CD630_n00380 | 94 -- 136 | 886 -- 937 | -10.8541 | 4.21282 | 4.27916 |
| vncR | CD630_n00380 | 126 -- 178 | 861 -- 915 | -10.8432 | 3.93642 | 4.27916 |
| CD630_31960 | CD630_n00380 | 125 -- 146 | 793 -- 812 | -10.8368 | 4.93757 | 4.27916 |
| CD630_21110 | CD630_n00380 | 13 -- 32 | 79 -- 100 | -10.8286 | 3.89630 | 4.27916 |
| CD630_05370 | CD630_n00380 | 72 -- 107 | 888 -- 918 | -10.8275 | 3.26947 | 4.27916 |
| CD630_06350 | CD630_n00380 | 139 -- 157 | 878 -- 895 | -10.8248 | 4.35983 | 4.27916 |
| CD630_17160 | CD630_n00380 | 85 -- 108 | 888 -- 912 | -10.8198 | 3.50228 | 4.27916 |
| CD630_05840 | CD630_n00380 | 52 -- 138 | 594 -- 690 | -10.7518 | 4.28779 | 4.27916 |
| CD630_14340 | CD630_n00380 | 51 -- 92 | 877 -- 920 | -10.7258 | 5.10868 | 4.27916 |
| CD630_11220 | CD630_n00380 | 138 -- 168 | 824 -- 856 | -10.6826 | 3.52327 | 4.27916 |
| CD630_15111 | CD630_n00380 | 20 -- 45 | 905 -- 932 | -10.6619 | 4.18664 | 4.27916 |
| CD630_20650 | CD630_n00380 | 26 -- 122 | 859 -- 981 | -10.6583 | 4.00083 | 4.27916 |
| CD630_02720 | CD630_n00380 | 138 -- 176 | 767 -- 801 | -10.6499 | 3.38938 | 4.27916 |
| cdtR | CD630_n00380 | 4 -- 73 | 659 -- 725 | -10.6481 | 5.84368 | 4.27916 |
| CD630_13380 | CD630_n00380 | 20 -- 84 | 840 -- 891 | -10.6468 | 4.74408 | 4.27916 |
| CD630_08660 | CD630_n00380 | 52 -- 81 | 838 -- 868 | -10.6422 | 3.86467 | 4.27916 |
| CD630_23070 | CD630_n00380 | 176 -- 196 | 862 -- 882 | -10.6344 | 5.23027 | 4.27916 |
| rbsC | CD630_n00380 | 160 -- 190 | 871 -- 912 | -10.6316 | 3.71650 | 4.27916 |
| CD630_05220 | CD630_n00380 | 160 -- 197 | 888 -- 934 | -10.5906 | 3.65057 | 4.27916 |
| CD630_22720 | CD630_n00380 | 92 -- 147 | 873 -- 918 | -10.582 | 3.81804 | 4.27916 |
| CD630_24920 | CD630_n00380 | 86 -- 96 | 434 -- 444 | -10.5695 | 3.89455 | 4.27916 |
| feoB | CD630_n00380 | 144 -- 181 | 895 -- 926 | -10.5691 | 4.05364 | 4.27916 |
| CD630_22991 | CD630_n00380 | 190 -- 200 | 250 -- 261 | -10.5634 | 6.03728 | 4.27916 |
| CD630_33480 | CD630_n00380 | 15 -- 33 | 169 -- 185 | -10.5633 | 3.67613 | 4.27916 |
| CD630_03650 | CD630_n00380 | 177 -- 197 | 657 -- 687 | -10.5234 | 3.93697 | 4.27916 |
| CD630_01640 | CD630_n00380 | 18 -- 56 | 694 -- 732 | -10.515 | 4.55381 | 4.27916 |
| cheW | CD630_n00380 | 71 -- 105 | 304 -- 338 | -10.5144 | 3.72371 | 4.27916 |
| CD630_27770 | CD630_n00380 | 87 -- 111 | 869 -- 895 | -10.5077 | 4.08588 | 4.27916 |
| CD630_26750 | CD630_n00380 | 166 -- 193 | 250 -- 281 | -10.4821 | 3.60610 | 4.27916 |
| CD630_05290 | CD630_n00380 | 6 -- 39 | 206 -- 236 | -10.4817 | 3.97604 | 4.27916 |
| CD630_24830 | CD630_n00380 | 21 -- 39 | 666 -- 685 | -10.4765 | 4.98239 | 4.27916 |
| CD630_27520 | CD630_n00380 | 111 -- 147 | 888 -- 926 | -10.4764 | 4.27693 | 4.27916 |
| CD630_36190 | CD630_n00380 | 76 -- 104 | 878 -- 913 | -10.4681 | 4.63449 | 4.27916 |
| CD630_27040 | CD630_n00380 | 37 -- 69 | 159 -- 187 | -10.4565 | 3.51812 | 4.27916 |
| CD630_07480 | CD630_n00380 | 5 -- 48 | 639 -- 684 | -10.4513 | 5.18321 | 4.27916 |
| sspA | CD630_n00380 | 87 -- 161 | 805 -- 893 | -10.4434 | 3.62689 | 4.27916 |
| CD630_19440 | CD630_n00380 | 82 -- 124 | 733 -- 785 | -10.4257 | 4.53937 | 4.27916 |
| CD630_25200 | CD630_n00380 | 168 -- 198 | 664 -- 702 | -10.4251 | 3.65158 | 4.27916 |
| CD630_09850 | CD630_n00380 | 1 -- 10 | 886 -- 895 | -10.4184 | 4.18679 | 4.27916 |
| CD630_04770 | CD630_n00380 | 85 -- 105 | 325 -- 345 | -10.415 | 3.58478 | 4.27916 |
| CD630_16530 | CD630_n00380 | 112 -- 137 | 24 -- 48 | -10.4137 | 4.37658 | 4.27916 |
| CD630_36170 | CD630_n00380 | 148 -- 165 | 871 -- 892 | -10.4018 | 3.72380 | 4.27916 |
| CD630_29840 | CD630_n00380 | 129 -- 166 | 457 -- 496 | -10.3991 | 4.48825 | 4.27916 |
| CD630_09570 | CD630_n00380 | 16 -- 26 | 901 -- 911 | -10.3924 | 5.31213 | 4.27916 |
| CD630_29070 | CD630_n00380 | 16 -- 26 | 901 -- 911 | -10.3924 | 3.89477 | 4.27916 |
| tcdC | CD630_n00380 | 114 -- 151 | 316 -- 344 | -10.3616 | 3.76101 | 4.27916 |
| CD630_06140 | CD630_n00380 | 87 -- 103 | 879 -- 895 | -10.3588 | 4.12519 | 4.27916 |
| CD630_33560 | CD630_n00380 | 91 -- 128 | 163 -- 210 | -10.3573 | 4.03147 | 4.27916 |
| CD630_34560 | CD630_n00380 | 85 -- 104 | 878 -- 895 | -10.3304 | 4.02604 | 4.27916 |
| CD630_21910 | CD630_n00380 | 106 -- 142 | 154 -- 186 | -10.3289 | 4.66600 | 4.27916 |
| CD630_11240 | CD630_n00380 | 2 -- 39 | 141 -- 176 | -10.3286 | 3.20274 | 4.27916 |
| CD630_05590 | CD630_n00380 | 65 -- 114 | 299 -- 344 | -10.3019 | 3.39670 | 4.27916 |
| rbsA | CD630_n00380 | 45 -- 74 | 322 -- 354 | -10.2959 | 3.71650 | 4.27916 |
| serS1 | CD630_n00380 | 151 -- 165 | 891 -- 911 | -10.279 | 3.45858 | 4.27916 |
| gcp | CD630_n00380 | 87 -- 145 | 830 -- 893 | -10.2563 | 4.92798 | 4.27916 |
| CD630_18910 | CD630_n00380 | 166 -- 192 | 885 -- 911 | -10.2546 | 3.30688 | 4.27916 |
| CD630_06560 | CD630_n00380 | 120 -- 167 | 198 -- 237 | -10.2531 | 4.02532 | 4.27916 |
| CD630_05150 | CD630_n00380 | 79 -- 112 | 888 -- 917 | -10.2518 | 3.69875 | 4.27916 |
| CD630_17970 | CD630_n00380 | 50 -- 123 | 377 -- 442 | -10.2461 | 3.68802 | 4.27916 |
| CD630_14420 | CD630_n00380 | 90 -- 147 | 325 -- 382 | -10.2409 | 3.99497 | 4.27916 |
| CD630_14700 | CD630_n00380 | 138 -- 155 | 902 -- 919 | -10.23 | 3.98234 | 4.27916 |
| CD630_25190 | CD630_n00380 | 128 -- 141 | 888 -- 907 | -10.2298 | 3.19663 | 4.27916 |
| CD630_20290 | CD630_n00380 | 119 -- 175 | 141 -- 208 | -10.2154 | 4.38183 | 4.27916 |
| CD630_19801 | CD630_n00380 | 125 -- 177 | 875 -- 931 | -10.2134 | 5.22035 | 4.27916 |
| CD630_18940 | CD630_n00380 | 89 -- 126 | 733 -- 786 | -10.2083 | 4.09988 | 4.27916 |
| CD630_03420 | CD630_n00380 | 81 -- 136 | 873 -- 929 | -10.2026 | 4.71868 | 4.27916 |
| spoVFB | CD630_n00380 | 179 -- 200 | 315 -- 339 | -10.1828 | 4.62758 | 4.27916 |
| CD630_26020 | CD630_n00380 | 105 -- 124 | 64 -- 81 | -10.1789 | 4.03155 | 4.27916 |
| chrA | CD630_n00380 | 133 -- 154 | 376 -- 392 | -10.1732 | 5.27255 | 4.27916 |
| CD630_03920 | CD630_n00380 | 151 -- 168 | 481 -- 497 | -10.1596 | 4.15934 | 4.27916 |
| CD630_13360 | CD630_n00380 | 165 -- 189 | 161 -- 184 | -10.1483 | 3.77330 | 4.27916 |
| CD630_18680 | CD630_n00380 | 41 -- 153 | 270 -- 405 | -10.1179 | 3.04315 | 4.27916 |
| CD630_26250 | CD630_n00380 | 139 -- 178 | 203 -- 243 | -10.1156 | 5.03379 | 4.27916 |
| flgB | CD630_n00380 | 77 -- 185 | 780 -- 918 | -10.0825 | 3.25736 | 4.27916 |
| CD630_29960 | CD630_n00380 | 1 -- 15 | 789 -- 803 | -10.08 | 4.51904 | 4.27916 |
| CD630_31970 | CD630_n00380 | 159 -- 184 | 652 -- 675 | -10.0785 | 3.24511 | 4.27916 |
| CD630_20920 | CD630_n00380 | 15 -- 74 | 727 -- 784 | -10.0641 | 4.13812 | 4.27916 |
| CD630_27220 | CD630_n00380 | 40 -- 118 | 290 -- 371 | -10.0624 | 4.18786 | 4.27916 |
| CD630_14150 | CD630_n00380 | 27 -- 71 | 763 -- 808 | -10.0354 | 3.28251 | 4.27916 |
| appB | CD630_n00380 | 140 -- 173 | 628 -- 677 | -10.0105 | 6.44103 | 4.27916 |
| ldh | CD630_n00380 | 180 -- 191 | 895 -- 906 | -10.0094 | 4.35818 | 4.27916 |
| CD630_29270 | CD630_n00380 | 103 -- 175 | 825 -- 893 | -9.99597 | 3.70174 | 4.27916 |
| CD630_06060 | CD630_n00380 | 169 -- 186 | 162 -- 179 | -9.98449 | 4.94050 | 4.27916 |
| CD630_03180 | CD630_n00380 | 88 -- 121 | 307 -- 339 | -9.97258 | 3.73138 | 4.27916 |
| dltD | CD630_n00380 | 127 -- 177 | 156 -- 208 | -9.9497 | 4.40777 | 4.27916 |
| glgP | CD630_n00380 | 178 -- 199 | 155 -- 178 | -9.93762 | 3.28530 | 4.27916 |
| CD630_21020 | CD630_n00380 | 12 -- 28 | 114 -- 130 | -9.92715 | 4.28666 | 4.27916 |
| CD630_26691 | CD630_n00380 | 1 -- 42 | 848 -- 892 | -9.92325 | 3.96230 | 4.27916 |
| CD630_14250 | CD630_n00380 | 10 -- 46 | 222 -- 255 | -9.91303 | 4.19604 | 4.27916 |
| cobS | CD630_n00380 | 157 -- 173 | 876 -- 898 | -9.90017 | 3.67481 | 4.27916 |
| agrB | CD630_n00380 | 128 -- 146 | 149 -- 168 | -9.89921 | 3.92598 | 4.27916 |
| CD630_22660 | CD630_n00380 | 9 -- 40 | 634 -- 666 | -9.8971 | 3.60496 | 4.27916 |
| eutP | CD630_n00380 | 79 -- 126 | 889 -- 929 | -9.89618 | 3.82847 | 4.27916 |
| CD630_21790 | CD630_n00380 | 142 -- 176 | 856 -- 893 | -9.89193 | 4.93762 | 4.27916 |
| CD630_05160 | CD630_n00380 | 4 -- 13 | 577 -- 586 | -9.89042 | 5.03573 | 4.27916 |
| bglA | CD630_n00380 | 78 -- 115 | 889 -- 916 | -9.85771 | 4.50315 | 4.27916 |
| CD630_03400 | CD630_n00380 | 28 -- 51 | 651 -- 675 | -9.85019 | 4.45994 | 4.27916 |
| LinCd | CD630_n00380 | 1 -- 36 | 162 -- 186 | -9.83909 | 4.33624 | 4.27916 |
| CD630_17630 | CD630_n00380 | 8 -- 40 | 644 -- 678 | -9.82905 | 4.52408 | 4.27916 |
| CD630_08780 | CD630_n00380 | 161 -- 189 | 152 -- 180 | -9.80724 | 7.76690 | 4.27916 |
| CD630_10220 | CD630_n00380 | 85 -- 174 | 366 -- 446 | -9.79135 | 4.08233 | 4.27916 |
| CD630_09970 | CD630_n00380 | 167 -- 190 | 152 -- 176 | -9.77934 | 3.25917 | 4.27916 |
| CD630_06920 | CD630_n00380 | 40 -- 65 | 697 -- 724 | -9.7575 | 3.13260 | 4.27916 |
| CD630_14190 | CD630_n00380 | 89 -- 171 | 839 -- 895 | -9.75617 | 3.78562 | 4.27916 |
| CD630_19903 | CD630_n00380 | 170 -- 199 | 850 -- 877 | -9.74477 | 5.79994 | 4.27916 |
| CD630_30310 | CD630_n00380 | 48 -- 104 | 878 -- 933 | -9.73818 | 4.56059 | 4.27916 |
| etfA | CD630_n00380 | 75 -- 96 | 886 -- 906 | -9.72673 | 4.80521 | 4.27916 |
| CD630_31480 | CD630_n00380 | 9 -- 22 | 617 -- 630 | -9.7266 | 4.88897 | 4.27916 |
| adhE | CD630_n00380 | 160 -- 194 | 168 -- 216 | -9.7265 | 3.06485 | 4.27916 |
| CD630_05211 | CD630_n00380 | 89 -- 104 | 878 -- 894 | -9.72504 | 3.97048 | 4.27916 |
| CD630_14210 | CD630_n00380 | 4 -- 39 | 163 -- 198 | -9.71386 | 3.82355 | 4.27916 |
| CD630_36781 | CD630_n00380 | 140 -- 166 | 651 -- 677 | -9.70613 | 3.75946 | 4.27916 |
| CD630_30110 | CD630_n00380 | 84 -- 109 | 872 -- 908 | -9.68506 | 3.31041 | 4.27916 |
| CD630_31382 | CD630_n00380 | 1 -- 20 | 432 -- 450 | -9.67634 | 3.49137 | 4.27916 |
| CD630_33682 | CD630_n00380 | 70 -- 112 | 877 -- 910 | -9.66633 | 5.77720 | 4.27916 |
| CD630_13620 | CD630_n00380 | 32 -- 71 | 741 -- 783 | -9.64396 | 4.77627 | 4.27916 |
| CD630_22710 | CD630_n00380 | 9 -- 49 | 400 -- 440 | -9.62957 | 4.05623 | 4.27916 |
| CD630_21540 | CD630_n00380 | 31 -- 45 | 162 -- 177 | -9.62739 | 4.33090 | 4.27916 |
| CD630_21750 | CD630_n00380 | 49 -- 92 | 889 -- 933 | -9.6051 | 3.70626 | 4.27916 |
| CD630_34080 | CD630_n00380 | 22 -- 78 | 762 -- 822 | -9.58773 | 3.20224 | 4.27916 |
| CD630_17380 | CD630_n00380 | 184 -- 196 | 430 -- 442 | -9.5845 | 5.03207 | 4.27916 |
| CD630_32500 | CD630_n00380 | 53 -- 101 | 882 -- 923 | -9.55775 | 6.17288 | 4.27916 |
| CD630_21460 | CD630_n00380 | 180 -- 201 | 900 -- 922 | -9.51771 | 4.83005 | 4.27916 |
| CD630_17151 | CD630_n00380 | 132 -- 191 | 727 -- 782 | -9.51133 | 6.43124 | 4.27916 |
| CD630_22850 | CD630_n00380 | 16 -- 57 | 629 -- 680 | -9.50565 | 3.84102 | 4.27916 |
| CD630_14010 | CD630_n00380 | 40 -- 86 | 633 -- 678 | -9.50385 | 3.69983 | 4.27916 |
| CD630_07070 | CD630_n00380 | 32 -- 47 | 793 -- 807 | -9.50138 | 3.46414 | 4.27916 |
| CD630_28181 | CD630_n00440 | 146 -- 164 | 60 -- 80 | -17.9114 | 3.91299 | 4.85723 |
| adhE | CD630_n00440 | 37 -- 82 | 201 -- 240 | -17.6765 | 3.06485 | 4.85723 |
| CD630_19790 | CD630_n00440 | 48 -- 94 | 59 -- 102 | -17.5994 | 3.79593 | 4.85723 |
| CD630_19401 | CD630_n00440 | 42 -- 66 | 14 -- 41 | -17.4395 | 4.37084 | 4.85723 |
| metQ | CD630_n00440 | 87 -- 96 | 56 -- 65 | -17.024 | 3.87190 | 4.85723 |
| CD630_17151 | CD630_n00440 | 168 -- 201 | 197 -- 229 | -16.8014 | 6.43124 | 4.85723 |
| etfA | CD630_n00440 | 62 -- 95 | 56 -- 78 | -16.6023 | 4.80521 | 4.85723 |
| CD630_04940 | CD630_n00440 | 82 -- 93 | 58 -- 69 | -16.4117 | 4.20948 | 4.85723 |
| CD630_06080 | CD630_n00440 | 126 -- 197 | 166 -- 236 | -16.3368 | 3.82315 | 4.85723 |
| CD630_15850 | CD630_n00440 | 122 -- 133 | 58 -- 70 | -16.0101 | 4.32968 | 4.85723 |
| CD630_31910 | CD630_n00440 | 57 -- 99 | 196 -- 236 | -15.6565 | 3.48314 | 4.85723 |
| folC | CD630_n00440 | 47 -- 95 | 55 -- 98 | -15.4451 | 3.85634 | 4.85723 |
| CD630_18311 | CD630_n00440 | 104 -- 151 | 187 -- 240 | -15.4338 | 6.86047 | 4.85723 |
| CD630_13930 | CD630_n00440 | 91 -- 136 | 201 -- 240 | -15.3326 | 4.09246 | 4.85723 |
| flgB | CD630_n00440 | 72 -- 100 | 56 -- 87 | -15.1963 | 3.25736 | 4.85723 |
| tcdC | CD630_n00440 | 87 -- 107 | 20 -- 40 | -15.043 | 3.76101 | 4.85723 |
| CD630_23930 | CD630_n00440 | 141 -- 195 | 185 -- 236 | -14.8142 | 3.05650 | 4.85723 |
| bcp | CD630_n00440 | 42 -- 64 | 12 -- 41 | -14.6965 | 3.16670 | 4.85723 |
| cwp29 | CD630_n00440 | 126 -- 157 | 9 -- 41 | -14.624 | 4.40865 | 4.85723 |
| CD630_31580 | CD630_n00440 | 171 -- 191 | 59 -- 78 | -14.5091 | 3.51756 | 4.85723 |
| CD630_17170 | CD630_n00440 | 87 -- 120 | 202 -- 239 | -14.3882 | 4.10230 | 4.85723 |
| CD630_15111 | CD630_n00440 | 85 -- 110 | 17 -- 43 | -14.2481 | 4.18664 | 4.85723 |
| CD630_10800 | CD630_n00440 | 41 -- 57 | 71 -- 87 | -14.2255 | 5.30319 | 4.85723 |
| moaA | CD630_n00440 | 86 -- 131 | 195 -- 240 | -14.1729 | 3.83799 | 4.85723 |
| CD630_03410 | CD630_n00440 | 30 -- 75 | 185 -- 235 | -14.1577 | 4.45994 | 4.85723 |
| CD630_32720 | CD630_n00440 | 61 -- 99 | 194 -- 235 | -14.1205 | 4.74276 | 4.85723 |
| dapB2 | CD630_n00440 | 108 -- 148 | 55 -- 102 | -14.1136 | 3.27218 | 4.85723 |
| CD630_21490 | CD630_n00440 | 52 -- 81 | 12 -- 39 | -14.0203 | 3.37462 | 4.85723 |
| CD630_22720 | CD630_n00440 | 136 -- 163 | 60 -- 74 | -13.9617 | 3.81804 | 4.85723 |
| cbpA | CD630_n00440 | 85 -- 96 | 58 -- 72 | -13.9517 | 3.03040 | 4.85723 |
| CD630_32520 | CD630_n00440 | 157 -- 198 | 58 -- 91 | -13.873 | 4.06098 | 4.85723 |
| argG | CD630_n00440 | 56 -- 92 | 62 -- 96 | -13.7621 | 4.80300 | 4.85723 |
| CD630_06170 | CD630_n00440 | 34 -- 72 | 194 -- 233 | -13.706 | 4.95366 | 4.85723 |
| CD630_10220 | CD630_n00440 | 91 -- 138 | 12 -- 40 | -13.7022 | 4.08233 | 4.85723 |
| CD630_12320 | CD630_n00440 | 87 -- 103 | 24 -- 40 | -13.6394 | 3.28667 | 4.85723 |
| def1 | CD630_n00440 | 87 -- 115 | 13 -- 41 | -13.5855 | 3.99720 | 4.85723 |
| CD630_06970 | CD630_n00440 | 100 -- 133 | 58 -- 90 | -13.4746 | 3.82312 | 4.85723 |
| CD630_32690 | CD630_n00440 | 60 -- 91 | 204 -- 236 | -13.3385 | 7.32676 | 4.85723 |
| CD630_35840 | CD630_n00440 | 86 -- 119 | 204 -- 240 | -13.3334 | 3.14170 | 4.85723 |
| CD630_26750 | CD630_n00440 | 63 -- 94 | 59 -- 93 | -13.2891 | 3.60610 | 4.85723 |
| CD630_21570 | CD630_n00440 | 78 -- 94 | 58 -- 78 | -13.2136 | 6.48385 | 4.85723 |
| CD630_33300 | CD630_n00440 | 116 -- 142 | 61 -- 87 | -13.2105 | 3.79149 | 4.85723 |
| CD630_13880 | CD630_n00440 | 41 -- 77 | 202 -- 238 | -13.1498 | 3.52735 | 4.85723 |
| CD630_35260 | CD630_n00440 | 89 -- 124 | 202 -- 240 | -13.1449 | 5.77277 | 4.85723 |
| CD630_01590 | CD630_n00440 | 92 -- 133 | 197 -- 240 | -13.1419 | 3.45609 | 4.85723 |
| CD630_17900 | CD630_n00440 | 96 -- 105 | 65 -- 74 | -13.1203 | 3.52167 | 4.85723 |
| CD630_04341 | CD630_n00440 | 38 -- 71 | 60 -- 95 | -13.0571 | 4.29262 | 4.85723 |
| nusB | CD630_n00440 | 136 -- 184 | 202 -- 239 | -13.0542 | 4.92798 | 4.85723 |
| CD630_16550 | CD630_n00440 | 84 -- 95 | 58 -- 72 | -13.0405 | 5.61064 | 4.85723 |
| CD630_08390 | CD630_n00440 | 82 -- 113 | 11 -- 41 | -13.015 | 5.16000 | 4.85723 |
| CD630_08090 | CD630_n00440 | 172 -- 186 | 12 -- 26 | -13.0081 | 3.57167 | 4.85723 |
| CD630_20170 | CD630_n00440 | 112 -- 148 | 196 -- 238 | -12.9913 | 6.90201 | 4.85723 |
| CD630_16150 | CD630_n00440 | 51 -- 66 | 66 -- 80 | -12.8733 | 4.20614 | 4.85723 |
| CD630_07400 | CD630_n00440 | 19 -- 50 | 185 -- 230 | -12.8579 | 4.52273 | 4.85723 |
| CD630_28280 | CD630_n00440 | 62 -- 97 | 203 -- 237 | -12.8515 | 4.59833 | 4.85723 |
| CD630_25050 | CD630_n00440 | 86 -- 137 | 185 -- 240 | -12.8457 | 4.33624 | 4.85723 |
| CD630_31382 | CD630_n00440 | 7 -- 14 | 66 -- 73 | -12.8284 | 3.49137 | 4.85723 |
| CD630_09570 | CD630_n00440 | 80 -- 123 | 7 -- 45 | -12.8277 | 5.31213 | 4.85723 |
| CD630_29070 | CD630_n00440 | 80 -- 123 | 7 -- 45 | -12.8277 | 3.89477 | 4.85723 |
| CD630_14700 | CD630_n00440 | 65 -- 100 | 207 -- 238 | -12.8193 | 3.98234 | 4.85723 |
| CD630_03220 | CD630_n00440 | 29 -- 73 | 197 -- 237 | -12.7786 | 3.36746 | 4.85723 |
| mobA | CD630_n00440 | 170 -- 193 | 58 -- 86 | -12.726 | 4.25703 | 4.85723 |
| ldh | CD630_n00440 | 48 -- 92 | 56 -- 91 | -12.5691 | 4.35818 | 4.85723 |
| CD630_25430 | CD630_n00440 | 151 -- 174 | 15 -- 38 | -12.5103 | 4.00527 | 4.85723 |
| CD630_17710 | CD630_n00440 | 86 -- 94 | 58 -- 66 | -12.472 | 4.37688 | 4.85723 |
| iorA | CD630_n00440 | 62 -- 93 | 60 -- 84 | -12.4631 | 3.83173 | 4.85723 |
| CD630_32070 | CD630_n00440 | 39 -- 74 | 202 -- 235 | -12.4527 | 3.51964 | 4.85723 |
| ung | CD630_n00440 | 90 -- 126 | 60 -- 91 | -12.4307 | 3.30908 | 4.85723 |
| CD630_16940 | CD630_n00440 | 87 -- 95 | 65 -- 73 | -12.4032 | 4.14665 | 4.85723 |
| CD630_18680 | CD630_n00440 | 91 -- 124 | 194 -- 229 | -12.297 | 3.04315 | 4.85723 |
| CD630_10970 | CD630_n00440 | 183 -- 199 | 58 -- 69 | -12.2776 | 3.53850 | 4.85723 |

| | | | | | | |
|---|---|---|---|---|---|---|
| CD630_32040 | CD630_n00440 | 18 -- 43 | 19 -- 44 | -12.2733 | 4.93516 | 4.85723 |
| CD630_13960 | CD630_n00440 | 89 -- 103 | 24 -- 40 | -12.2678 | 3.30375 | 4.85723 |
| CD630_05370 | CD630_n00440 | 74 -- 102 | 1 -- 32 | -12.2215 | 3.26947 | 4.85723 |
| CD630_20980 | CD630_n00440 | 88 -- 109 | 12 -- 40 | -12.181 | 8.12847 | 4.85723 |
| CD630_09040 | CD630_n00440 | 170 -- 197 | 202 -- 229 | -12.1706 | 4.11851 | 4.85723 |
| etfA | CD630_n00440 | 1 -- 33 | 196 -- 224 | -12.1253 | 3.41204 | 4.85723 |
| CD630_18960 | CD630_n00440 | 46 -- 67 | 217 -- 239 | -12.11 | 4.37822 | 4.85723 |
| cprK | CD630_n00440 | 25 -- 45 | 215 -- 234 | -12.0915 | 3.96216 | 4.85723 |
| garK | CD630_n00440 | 10 -- 36 | 215 -- 240 | -12.0596 | 3.72836 | 4.85723 |
| CD630_11241 | CD630_n00440 | 172 -- 182 | 65 -- 74 | -11.9808 | 4.85723 | 4.85723 |
| CD630_19690 | CD630_n00440 | 147 -- 189 | 183 -- 229 | -11.9224 | 3.85861 | 4.85723 |
| CD630_20670 | CD630_n00440 | 84 -- 127 | 2 -- 43 | -11.8731 | 4.42757 | 4.85723 |
| CD630_26680 | CD630_n00440 | 178 -- 193 | 61 -- 72 | -11.806 | 3.53956 | 4.85723 |
| CD630_32050 | CD630_n00440 | 45 -- 86 | 196 -- 234 | -11.781 | 5.73797 | 4.85723 |
| bglA | CD630_n00440 | 155 -- 198 | 205 -- 240 | -11.7653 | 4.86402 | 4.85723 |
| CD630_23861 | CD630_n00440 | 88 -- 94 | 59 -- 65 | -11.7626 | 5.83068 | 4.85723 |
| CD630_10950 | CD630_n00440 | 89 -- 129 | 192 -- 239 | -11.7286 | 3.67675 | 4.85723 |
| ssuC | CD630_n00440 | 64 -- 98 | 9 -- 40 | -11.704 | 3.45491 | 4.85723 |
| CD630_22710 | CD630_n00440 | 123 -- 143 | 203 -- 224 | -11.6839 | 4.05623 | 4.85723 |
| vncR | CD630_n00440 | 122 -- 145 | 60 -- 90 | -11.6793 | 3.93642 | 4.85723 |
| CD630_19904 | CD630_n00440 | 161 -- 175 | 13 -- 40 | -11.6188 | 5.96526 | 4.85723 |
| CD630_05450 | CD630_n00440 | 84 -- 92 | 58 -- 66 | -11.615 | 5.04761 | 4.85723 |
| CD630_33682 | CD630_n00440 | 85 -- 110 | 17 -- 40 | -11.5886 | 5.77720 | 4.85723 |
| CD630_19800 | CD630_n00440 | 40 -- 52 | 59 -- 71 | -11.5317 | 5.22035 | 4.85723 |
| CD630_06140 | CD630_n00440 | 85 -- 92 | 57 -- 64 | -11.524 | 4.12519 | 4.85723 |
| CD630_30130 | CD630_n00440 | 4 -- 10 | 60 -- 66 | -11.4991 | 3.28343 | 4.85723 |
| CD630_04610 | CD630_n00440 | 21 -- 43 | 197 -- 221 | -11.4981 | 3.83025 | 4.85723 |
| CD630_05230 | CD630_n00440 | 2 -- 38 | 203 -- 238 | -11.4696 | 3.73787 | 4.85723 |
| glnA | CD630_n00440 | 119 -- 156 | 197 -- 236 | -11.4547 | 3.87629 | 4.85723 |
| CD630_19700 | CD630_n00440 | 11 -- 47 | 203 -- 236 | -11.4087 | 3.85861 | 4.85723 |
| CD630_22660 | CD630_n00440 | 154 -- 182 | 210 -- 238 | -11.3195 | 3.60496 | 4.85723 |
| CD630_14240 | CD630_n00440 | 85 -- 110 | 17 -- 40 | -11.3047 | 8.60766 | 4.85723 |
| CD630_20290 | CD630_n00440 | 84 -- 90 | 59 -- 65 | -11.2774 | 4.38183 | 4.85723 |
| CD630_05870 | CD630_n00440 | 166 -- 197 | 54 -- 91 | -11.2267 | 4.28339 | 4.85723 |
| CD630_15550 | CD630_n00440 | 17 -- 33 | 1 -- 19 | -11.2243 | 3.49019 | 4.85723 |
| CD630_25320 | CD630_n00440 | 143 -- 174 | 202 -- 239 | -11.1817 | 4.29415 | 4.85723 |
| CD630_08070 | CD630_n00440 | 159 -- 168 | 69 -- 78 | -11.1615 | 4.87819 | 4.85723 |
| CD630_30310 | CD630_n00440 | 15 -- 49 | 196 -- 229 | -11.1527 | 4.56059 | 4.85723 |
| sspA | CD630_n00440 | 88 -- 120 | 206 -- 240 | -11.141 | 3.62689 | 4.85723 |
| CD630_29960 | CD630_n00440 | 88 -- 124 | 203 -- 240 | -11.0954 | 4.51904 | 4.85723 |
| cdu2 | CD630_n00440 | 89 -- 131 | 3 -- 41 | -11.0952 | 2.97335 | 4.85723 |
| CD630_34030 | CD630_n00440 | 160 -- 186 | 204 -- 230 | -11.0902 | 3.05766 | 4.85723 |
| CD630_08940 | CD630_n00440 | 75 -- 93 | 58 -- 70 | -11.0633 | 3.72225 | 4.85723 |
| eutW | CD630_n00440 | 9 -- 16 | 67 -- 74 | -11.0298 | 4.55715 | 4.85723 |
| CD630_32700 | CD630_n00440 | 112 -- 138 | 213 -- 238 | -11.0098 | 4.74276 | 4.85723 |
| CD630_03040 | CD630_n00440 | 87 -- 142 | 201 -- 240 | -10.9853 | 3.51403 | 4.85723 |
| CD630_17720 | CD630_n00440 | 4 -- 39 | 1 -- 43 | -10.9772 | 3.22910 | 4.85723 |
| CD630_09760 | CD630_n00440 | 73 -- 105 | 60 -- 88 | -10.9398 | 4.83410 | 4.85723 |
| CD630_28910 | CD630_n00440 | 73 -- 105 | 60 -- 88 | -10.9398 | 4.51664 | 4.85723 |
| CD630_19840 | CD630_n00440 | 91 -- 111 | 216 -- 240 | -10.9283 | 3.44724 | 4.85723 |
| CD630_31480 | CD630_n00440 | 157 -- 177 | 205 -- 225 | -10.9084 | 4.88897 | 4.85723 |
| CD630_10080 | CD630_n00440 | 85 -- 126 | 206 -- 240 | -10.9075 | 3.75625 | 4.85723 |
| CD630_28260 | CD630_n00440 | 72 -- 100 | 62 -- 93 | -10.9009 | 3.51613 | 4.85723 |
| cheW | CD630_n00440 | 10 -- 55 | 199 -- 238 | -10.8716 | 3.72371 | 4.85723 |
| CD630_23000 | CD630_n00440 | 93 -- 101 | 64 -- 72 | -10.8681 | 3.54130 | 4.85723 |
| CD630_17360 | CD630_n00440 | 111 -- 140 | 10 -- 47 | -10.8633 | 5.66070 | 4.85723 |
| CD630_13380 | CD630_n00440 | 119 -- 141 | 63 -- 86 | -10.8133 | 4.74408 | 4.85723 |
| CD630_16750 | CD630_n00440 | 173 -- 191 | 220 -- 237 | -10.7892 | 5.04051 | 4.85723 |
| CD630_04880 | CD630_n00440 | 140 -- 155 | 10 -- 26 | -10.7885 | 4.62291 | 4.85723 |
| CD630_21140 | CD630_n00440 | 43 -- 79 | 196 -- 231 | -10.7872 | 3.33304 | 4.85723 |
| gcvTPA | CD630_n00440 | 87 -- 96 | 55 -- 64 | -10.7691 | 4.77229 | 4.85723 |
| CD630_25040 | CD630_n00440 | 23 -- 73 | 1 -- 42 | -10.7435 | 3.43898 | 4.85723 |
| cwp26 | CD630_n00440 | 126 -- 167 | 1 -- 41 | -10.7207 | 4.44377 | 4.85723 |
| CD630_05950 | CD630_n00440 | 24 -- 44 | 197 -- 219 | -10.6863 | 3.31669 | 4.85723 |
| CD630_10050 | CD630_n00440 | 152 -- 190 | 197 -- 240 | -10.6715 | 3.93750 | 4.85723 |
| CD630_08460 | CD630_n00440 | 91 -- 140 | 200 -- 240 | -10.6316 | 4.15083 | 4.85723 |
| CD630_21720 | CD630_n00440 | 152 -- 176 | 213 -- 240 | -10.6094 | 4.13030 | 4.85723 |
| CD630_27310 | CD630_n00440 | 86 -- 104 | 223 -- 240 | -10.6089 | 4.01622 | 4.85723 |
| CD630_31210 | CD630_n00440 | 84 -- 93 | 58 -- 67 | -10.5751 | 4.17693 | 4.85723 |
| panG | CD630_n00440 | 174 -- 194 | 203 -- 224 | -10.5734 | 5.54047 | 4.85723 |
| opuCA | CD630_n00440 | 87 -- 110 | 58 -- 73 | -10.5549 | 4.07035 | 4.85723 |
| CD630_14010 | CD630_n00440 | 170 -- 186 | 71 -- 87 | -10.5523 | 3.69983 | 4.85723 |
| CD630_14360 | CD630_n00440 | 194 -- 200 | 58 -- 64 | -10.5488 | 3.02079 | 4.85723 |
| CD630_23090 | CD630_n00440 | 81 -- 110 | 17 -- 43 | -10.5109 | 4.25622 | 4.85723 |
| CD630_06200 | CD630_n00440 | 89 -- 112 | 17 -- 40 | -10.5052 | 3.57064 | 4.85723 |
| CD630_15890 | CD630_n00440 | 149 -- 167 | 11 -- 31 | -10.4934 | 3.39962 | 4.85723 |
| CD630_32060 | CD630_n00440 | 25 -- 45 | 58 -- 72 | -10.478 | 5.73797 | 4.85723 |
| CD630_33580 | CD630_n00440 | 176 -- 187 | 213 -- 224 | -10.4691 | 3.95778 | 4.85723 |
| CD630_22070 | CD630_n00440 | 89 -- 125 | 2 -- 40 | -10.446 | 4.09651 | 4.85723 |
| glgB | CD630_n00440 | 190 -- 200 | 64 -- 74 | -10.4098 | 4.48273 | 4.85723 |
| argS | CD630_n00440 | 74 -- 114 | 198 -- 238 | -10.3731 | 4.96036 | 4.85723 |
| CD630_02740 | CD630_n00440 | 158 -- 165 | 63 -- 70 | -10.3548 | 5.02277 | 4.85723 |
| bglA | CD630_n00440 | 77 -- 96 | 13 -- 33 | -10.3333 | 4.50315 | 4.85723 |
| CD630_06060 | CD630_n00440 | 77 -- 102 | 214 -- 236 | -10.2966 | 4.94050 | 4.85723 |
| CD630_32710 | CD630_n00440 | 91 -- 129 | 197 -- 237 | -10.2867 | 4.74276 | 4.85723 |
| CD630_36250 | CD630_n00440 | 130 -- 157 | 201 -- 233 | -10.2662 | 4.13224 | 4.85723 |
| CD630_19000 | CD630_n00440 | 64 -- 94 | 58 -- 98 | -10.2482 | 4.18443 | 4.85723 |
| CD630_21540 | CD630_n00440 | 85 -- 108 | 212 -- 239 | -10.24 | 4.33090 | 4.85723 |
| CD630_30140 | CD630_n00440 | 84 -- 122 | 206 -- 240 | -10.2386 | 3.28343 | 4.85723 |
| CD630_16460 | CD630_n00440 | 129 -- 163 | 202 -- 238 | -10.2167 | 4.64028 | 4.85723 |
| CD630_06560 | CD630_n00440 | 53 -- 79 | 212 -- 240 | -10.2056 | 4.02532 | 4.85723 |
| CD630_27200 | CD630_n00440 | 1 -- 14 | 66 -- 79 | -10.1882 | 3.42516 | 4.85723 |
| xdhA | CD630_n00440 | 3 -- 31 | 201 -- 231 | -10.1533 | 4.45147 | 4.85723 |
| opuCC | CD630_n00440 | 96 -- 121 | 6 -- 38 | -10.1505 | 4.07035 | 4.85723 |
| CD630_07121 | CD630_n00440 | 79 -- 86 | 62 -- 69 | -10.1503 | 3.26177 | 4.85723 |
| CD630_22850 | CD630_n00440 | 84 -- 94 | 60 -- 69 | -10.1393 | 3.84102 | 4.85723 |
| CD630_25620 | CD630_n00440 | 128 -- 159 | 208 -- 237 | -10.1314 | 3.02685 | 4.85723 |
| CD630_20171 | CD630_n00440 | 94 -- 132 | 202 -- 237 | -10.1254 | 6.87491 | 4.85723 |
| CD630_06110 | CD630_n00440 | 175 -- 186 | 62 -- 71 | -10.1169 | 5.61380 | 4.85723 |
| CD630_02420 | CD630_n00440 | 154 -- 183 | 205 -- 238 | -10.1028 | 3.71048 | 4.85723 |
| CD630_15200 | CD630_n00440 | 34 -- 78 | 200 -- 240 | -10.0997 | 4.00442 | 4.85723 |
| CD630_05490 | CD630_n00440 | 6 -- 30 | 64 -- 94 | -10.0982 | 3.89852 | 4.85723 |
| CD630_11730 | CD630_n00440 | 89 -- 124 | 202 -- 239 | -10.0982 | 4.80521 | 4.85723 |
| CD630_08130 | CD630_n00440 | 86 -- 109 | 215 -- 240 | -10.0975 | 4.63831 | 4.85723 |
| CD630_29840 | CD630_n00440 | 31 -- 57 | 6 -- 40 | -10.094 | 4.48825 | 4.85723 |
| CD630_31461 | CD630_n00440 | 91 -- 99 | 64 -- 72 | -10.0906 | 4.41476 | 4.85723 |
| CD630_16450 | CD630_n00440 | 88 -- 104 | 223 -- 240 | -10.0772 | 4.57547 | 4.85723 |
| CD630_06540 | CD630_n00440 | 92 -- 119 | 14 -- 40 | -10.0454 | 3.54673 | 4.85723 |
| CD630_21910 | CD630_n00440 | 91 -- 117 | 216 -- 240 | -10.0144 | 4.66600 | 4.85723 |
| CD630_14130 | CD630_n00440 | 142 -- 158 | 14 -- 40 | -10.0106 | 4.57970 | 4.85723 |
| CD630_29270 | CD630_n00440 | 3 -- 24 | 196 -- 221 | -9.98032 | 3.70174 | 4.85723 |
| cwp17 | CD630_n00440 | 89 -- 128 | 206 -- 239 | -9.9671 | 3.43363 | 4.85723 |
| CD630_08370 | CD630_n00440 | 144 -- 153 | 64 -- 72 | -9.96611 | 3.65639 | 4.85723 |
| CD630_02430 | CD630_n00440 | 24 -- 30 | 60 -- 66 | -9.96108 | 3.71048 | 4.85723 |
| CD630_07960 | CD630_n00440 | 76 -- 94 | 68 -- 86 | -9.95786 | 3.22537 | 4.85723 |
| kdpE | CD630_n00440 | 128 -- 134 | 67 -- 73 | -9.94646 | 4.27140 | 4.85723 |
| CD630_05900 | CD630_n00440 | 29 -- 56 | 202 -- 240 | -9.94089 | 3.21934 | 4.85723 |
| CD630_17770 | CD630_n00440 | 62 -- 110 | 196 -- 238 | -9.93824 | 5.06490 | 4.85723 |
| CD630_23820 | CD630_n00440 | 90 -- 121 | 12 -- 40 | -9.93476 | 3.18419 | 4.85723 |
| CD630_05840 | CD630_n00440 | 90 -- 97 | 66 -- 73 | -9.90482 | 4.28779 | 4.85723 |
| CD630_22030 | CD630_n00440 | 82 -- 127 | 202 -- 240 | -9.89821 | 3.29934 | 4.85723 |
| CD630_18090 | CD630_n00440 | 17 -- 31 | 66 -- 80 | -9.898 | 4.23961 | 4.85723 |
| CD630_21770 | CD630_n00440 | 172 -- 201 | 201 -- 228 | -9.87151 | 4.02587 | 4.85723 |
| CD630_06920 | CD630_n00440 | 83 -- 108 | 213 -- 240 | -9.86453 | 3.13260 | 4.85723 |
| eutV | CD630_n00440 | 176 -- 196 | 215 -- 237 | -9.86424 | 4.55715 | 4.85723 |
| fliI | CD630_n00440 | 89 -- 137 | 204 -- 237 | -9.86118 | 3.03666 | 4.85723 |
| CD630_04320 | CD630_n00440 | 2 -- 44 | 199 -- 240 | -9.85159 | 3.68342 | 4.85723 |
| CD630_33190 | CD630_n00440 | 54 -- 89 | 201 -- 235 | -9.84955 | 4.13899 | 4.85723 |
| CD630_15060 | CD630_n00440 | 2 -- 25 | 2 -- 25 | -9.83906 | 3.69396 | 4.85723 |
| CD630_06050 | CD630_n00440 | 88 -- 107 | 220 -- 239 | -9.8247 | 3.59332 | 4.85723 |
| CD630_25610 | CD630_n00440 | 125 -- 137 | 58 -- 72 | -9.81962 | 3.02685 | 4.85723 |
| CD630_19780 | CD630_n00440 | 64 -- 87 | 215 -- 240 | -9.78722 | 3.64830 | 4.85723 |
| CD630_14190 | CD630_n00440 | 116 -- 162 | 185 -- 229 | -9.75295 | 3.78562 | 4.85723 |
| CD630_27520 | CD630_n00440 | 113 -- 147 | 58 -- 92 | -9.75218 | 4.27693 | 4.85723 |
| nrdD | CD630_n00440 | 56 -- 93 | 200 -- 239 | -9.74905 | 3.28290 | 4.85723 |
| CD630_15740 | CD630_n00440 | 49 -- 78 | 208 -- 237 | -9.73773 | 4.43222 | 4.85723 |
| dnaD | CD630_n00440 | 51 -- 74 | 201 -- 223 | -9.73623 | 3.56078 | 4.85723 |
| CD630_25300 | CD630_n00440 | 88 -- 97 | 10 -- 19 | -9.7323 | 4.49333 | 4.85723 |
| CD630_27240 | CD630_n00440 | 5 -- 31 | 197 -- 224 | -9.72254 | 3.52462 | 4.85723 |
| CD630_12650 | CD630_n00440 | 112 -- 143 | 59 -- 91 | -9.71718 | 6.31827 | 4.85723 |
| CD630_16950 | CD630_n00440 | 166 -- 187 | 207 -- 240 | -9.70976 | 4.42569 | 4.85723 |
| CD630_18400 | CD630_n00440 | 136 -- 157 | 201 -- 221 | -9.70788 | 4.67962 | 4.85723 |
| CD630_30620 | CD630_n00440 | 68 -- 92 | 68 -- 89 | -9.70671 | 4.15443 | 4.85723 |
| CD630_14410 | CD630_n00440 | 83 -- 95 | 58 -- 70 | -9.6834 | 3.39433 | 4.85723 |
| CD630_21750 | CD630_n00440 | 79 -- 87 | 13 -- 21 | -9.68315 | 3.70626 | 4.85723 |
| ppnK | CD630_n00440 | 89 -- 139 | 196 -- 240 | -9.67996 | 4.36817 | 4.85723 |
| CD630_28320 | CD630_n00440 | 180 -- 193 | 12 -- 29 | -9.67876 | 3.99184 | 4.85723 |
| malY | CD630_n00440 | 79 -- 127 | 196 -- 240 | -9.67561 | 4.46345 | 4.85723 |
| CD630_10830 | CD630_n00440 | 175 -- 197 | 215 -- 237 | -9.67544 | 4.82398 | 4.85723 |
| cstA | CD630_n00440 | 89 -- 95 | 67 -- 73 | -9.6652 | 3.53584 | 4.85723 |
| cobQ | CD630_n00440 | 163 -- 187 | 216 -- 238 | -9.66018 | 3.13524 | 4.85723 |
| CD630_28870 | CD630_n00440 | 28 -- 78 | 55 -- 93 | -9.64414 | 3.41991 | 4.85723 |
| sigK | CD630_n00440 | 78 -- 106 | 211 -- 240 | -9.6428 | 3.08122 | 4.85723 |
| cynT | CD630_n00440 | 113 -- 164 | 196 -- 240 | -9.61289 | 4.21370 | 4.85723 |
| CD630_18970 | CD630_n00440 | 71 -- 100 | 202 -- 232 | -9.59804 | 3.57243 | 4.85723 |
| oppB | CD630_n00440 | 88 -- 96 | 54 -- 62 | -9.59161 | 2.98328 | 4.85723 |
| chrA | CD630_n00440 | 22 -- 58 | 1 -- 36 | -9.58544 | 5.27255 | 4.85723 |
| CD630_23440 | CD630_n00440 | 1 -- 7 | 58 -- 64 | -9.58177 | 3.34841 | 4.85723 |
| CD630_29500 | CD630_n00440 | 131 -- 158 | 196 -- 229 | -9.5747 | 3.70644 | 4.85723 |
| CD630_22200 | CD630_n00440 | 79 -- 95 | 60 -- 78 | -9.56254 | 4.18288 | 4.85723 |
| CD630_03920 | CD630_n00440 | 61 -- 95 | 60 -- 100 | -9.55987 | 4.15934 | 4.85723 |
| CD630_14420 | CD630_n00440 | 144 -- 178 | 197 -- 230 | -9.52394 | 3.99497 | 4.85723 |
| CD630_13950 | CD630_n00440 | 133 -- 155 | 1 -- 22 | -9.51537 | 4.25540 | 4.85723 |
| CD630_15150 | CD630_n00440 | 1 -- 26 | 14 -- 38 | -9.51216 | 3.35564 | 4.85723 |
| CD630_32520 | CD630_n00460 | 154 -- 201 | 211 -- 249 | -19.5709 | 4.06098 | 5.55447 |
| CD630_04341 | CD630_n00460 | 26 -- 143 | 203 -- 351 | -19.4872 | 4.29262 | 5.55447 |
| CD630_23820 | CD630_n00460 | 52 -- 108 | 727 -- 783 | -18.553 | 3.18419 | 5.55447 |
| malY | CD630_n00460 | 57 -- 103 | 331 -- 373 | -17.9833 | 4.46345 | 5.55447 |
| CD630_09760 | CD630_n00460 | 105 -- 144 | 1001 -- 1038 | -17.9078 | 4.83410 | 5.55447 |
| CD630_28910 | CD630_n00460 | 105 -- 144 | 1001 -- 1038 | -17.9078 | 4.51664 | 5.55447 |
| CD630_28260 | CD630_n00460 | 70 -- 104 | 210 -- 245 | -17.826 | 3.51613 | 5.55447 |
| CD630_08090 | CD630_n00460 | 150 -- 181 | 218 -- 247 | -17.685 | 3.57167 | 5.55447 |
| argS | CD630_n00460 | 64 -- 94 | 210 -- 245 | -17.6789 | 4.96036 | 5.55447 |
| CD630_21620 | CD630_n00460 | 137 -- 185 | 402 -- 443 | -17.4036 | 3.02649 | 5.55447 |
| CD630_16460 | CD630_n00460 | 91 -- 163 | 291 -- 355 | -17.2299 | 4.64028 | 5.55447 |
| CD630_27110 | CD630_n00460 | 119 -- 185 | 537 -- 609 | -16.9612 | 3.78231 | 5.55447 |
| CD630_36040 | CD630_n00460 | 82 -- 93 | 210 -- 221 | -16.7867 | 3.52009 | 5.55447 |
| bglA | CD630_n00460 | 68 -- 106 | 203 -- 243 | -16.5931 | 4.50315 | 5.55447 |
| CD630_14250 | CD630_n00460 | 81 -- 97 | 205 -- 222 | -16.5794 | 4.19604 | 5.55447 |
| CD630_33420 | CD630_n00460 | 2 -- 31 | 398 -- 425 | -16.1802 | 3.22095 | 5.55447 |
| folC | CD630_n00460 | 46 -- 134 | 779 -- 852 | -16.0449 | 3.85634 | 5.55447 |
| CD630_24590 | CD630_n00460 | 8 -- 129 | 351 -- 489 | -16.0161 | 3.70401 | 5.55447 |
| rbsC | CD630_n00460 | 31 -- 53 | 340 -- 367 | -15.8615 | 3.71650 | 5.55447 |
| CD630_22510 | CD630_n00460 | 109 -- 143 | 113 -- 146 | -15.8515 | 11.42763 | 5.55447 |
| srlEb | CD630_n00460 | 59 -- 94 | 213 -- 245 | -15.8102 | 3.50140 | 5.55447 |
| eutV | CD630_n00460 | 99 -- 181 | 347 -- 432 | -15.6867 | 4.55715 | 5.55447 |
| appF | CD630_n00460 | 49 -- 192 | 407 -- 553 | -15.6608 | 4.25570 | 5.55447 |
| opuCC | CD630_n00460 | 88 -- 195 | 132 -- 240 | -15.4969 | 4.07035 | 5.55447 |
| kdpE | CD630_n00460 | 185 -- 201 | 347 -- 363 | -15.2583 | 4.27140 | 5.55447 |
| moeA | CD630_n00460 | 14 -- 74 | 117 -- 167 | -15.2469 | 3.26118 | 5.55447 |
| cdu2 | CD630_n00460 | 39 -- 68 | 215 -- 242 | -15.1082 | 2.97335 | 5.55447 |
| CD630_06060 | CD630_n00460 | 174 -- 193 | 221 -- 240 | -15.0971 | 4.94050 | 5.55447 |
| CD630_10880 | CD630_n00460 | 147 -- 182 | 208 -- 242 | -15.0346 | 5.05048 | 5.55447 |
| CD630_10220 | CD630_n00460 | 81 -- 94 | 214 -- 227 | -15.0215 | 4.08233 | 5.55447 |
| CD630_27240 | CD630_n00460 | 137 -- 172 | 649 -- 682 | -14.9426 | 3.52462 | 5.55447 |
| CD630_20460 | CD630_n00460 | 69 -- 112 | 206 -- 245 | -14.9085 | 7.94034 | 5.55447 |
| CD630_23930 | CD630_n00460 | 134 -- 158 | 1006 -- 1031 | -14.8985 | 3.05650 | 5.55447 |
| appC | CD630_n00460 | 35 -- 76 | 159 -- 203 | -14.7984 | 4.50438 | 5.55447 |
| CD630_17151 | CD630_n00460 | 164 -- 200 | 710 -- 746 | -14.7754 | 6.43124 | 5.55447 |
| bglA | CD630_n00460 | 162 -- 201 | 1001 -- 1034 | -14.7554 | 4.86402 | 5.55447 |
| CD630_15850 | CD630_n00460 | 137 -- 197 | 838 -- 898 | -14.7137 | 4.32968 | 5.55447 |
| CD630_19904 | CD630_n00460 | 9 -- 33 | 118 -- 141 | -14.7108 | 5.96526 | 5.55447 |
| metQ | CD630_n00460 | 144 -- 181 | 212 -- 250 | -14.6977 | 3.87190 | 5.55447 |
| cbpA | CD630_n00460 | 75 -- 95 | 286 -- 306 | -14.5902 | 3.03040 | 5.55447 |
| CD630_32890 | CD630_n00460 | 60 -- 92 | 214 -- 246 | -14.5374 | 3.41276 | 5.55447 |
| CD630_03400 | CD630_n00460 | 109 -- 192 | 101 -- 194 | -14.5282 | 4.45994 | 5.55447 |
| CD630_26750 | CD630_n00460 | 78 -- 95 | 285 -- 303 | -14.5211 | 3.60610 | 5.55447 |
| pilT | CD630_n00460 | 128 -- 144 | 821 -- 837 | -14.5005 | 3.24639 | 5.55447 |
| CD630_02720 | CD630_n00460 | 117 -- 173 | 860 -- 911 | -14.457 | 3.38938 | 5.55447 |
| CD630_19670 | CD630_n00460 | 73 -- 90 | 290 -- 308 | -14.3671 | 3.20718 | 5.55447 |
| CD630_12650 | CD630_n00460 | 103 -- 144 | 210 -- 244 | -14.3644 | 6.31827 | 5.55447 |
| CD630_20260 | CD630_n00460 | 110 -- 141 | 410 -- 438 | -14.3211 | 3.84162 | 5.55447 |
| adhE | CD630_n00460 | 138 -- 200 | 857 -- 911 | -14.3143 | 3.06485 | 5.55447 |
| CD630_10830 | CD630_n00460 | 163 -- 183 | 1004 -- 1024 | -14.2899 | 4.82398 | 5.55447 |
| CD630_33570 | CD630_n00460 | 141 -- 168 | 595 -- 625 | -14.1777 | 3.95778 | 5.55447 |
| scpB | CD630_n00460 | 122 -- 160 | 104 -- 140 | -14.1649 | 3.33560 | 5.55447 |
| CD630_25300 | CD630_n00460 | 62 -- 108 | 998 -- 1043 | -14.1272 | 4.49333 | 5.55447 |
| CD630_13880 | CD630_n00460 | 103 -- 158 | 103 -- 146 | -14.0965 | 3.52735 | 5.55447 |
| CD630_14182 | CD630_n00460 | 118 -- 141 | 595 -- 618 | -14.0672 | 4.82360 | 5.55447 |
| mobA | CD630_n00460 | 4 -- 39 | 204 -- 233 | -14.0501 | 4.25703 | 5.55447 |
| CD630_31490 | CD630_n00460 | 106 -- 122 | 100 -- 116 | -14.0461 | 4.08593 | 5.55447 |
| CD630_13560 | CD630_n00460 | 168 -- 194 | 21 -- 47 | -14.0289 | 3.38083 | 5.55447 |
| CD630_05370 | CD630_n00460 | 80 -- 106 | 203 -- 227 | -14.0227 | 3.26947 | 5.55447 |
| terD | CD630_n00460 | 44 -- 187 | 351 -- 471 | -14.0063 | 3.33148 | 5.55447 |
| def1 | CD630_n00460 | 95 -- 134 | 408 -- 443 | -13.9886 | 3.99720 | 5.55447 |
| CD630_05840 | CD630_n00460 | 83 -- 171 | 128 -- 224 | -13.9837 | 4.28779 | 5.55447 |
| CD630_21760 | CD630_n00460 | 168 -- 199 | 120 -- 147 | -13.9837 | 4.43846 | 5.55447 |
| moaA | CD630_n00460 | 81 -- 111 | 401 -- 427 | -13.9699 | 3.83799 | 5.55447 |
| CD630_14630 | CD630_n00460 | 116 -- 146 | 408 -- 440 | -13.9667 | 3.34485 | 5.55447 |
| CD630_14130 | CD630_n00460 | 119 -- 198 | 684 -- 754 | -13.9487 | 4.57970 | 5.55447 |
| oppB | CD630_n00460 | 120 -- 181 | 977 -- 1019 | -13.8968 | 2.98328 | 5.55447 |
| CD630_21020 | CD630_n00460 | 86 -- 120 | 186 -- 221 | -13.8753 | 4.28666 | 5.55447 |
| CD630_04320 | CD630_n00460 | 10 -- 70 | 325 -- 375 | -13.8611 | 3.68342 | 5.55447 |
| CD630_25040 | CD630_n00460 | 11 -- 62 | 203 -- 249 | -13.8424 | 3.43898 | 5.55447 |
| etfA | CD630_n00460 | 64 -- 103 | 1002 -- 1047 | -13.7759 | 3.41204 | 5.55447 |
| sspA | CD630_n00460 | 68 -- 143 | 100 -- 167 | -13.7738 | 3.62689 | 5.55447 |
| CD630_19690 | CD630_n00460 | 67 -- 129 | 724 -- 769 | -13.7507 | 3.85861 | 5.55447 |
| CD630_13380 | CD630_n00460 | 26 -- 81 | 828 -- 870 | -13.7424 | 4.74408 | 5.55447 |
| CD630_33600 | CD630_n00460 | 134 -- 188 | 599 -- 645 | -13.7339 | 3.73196 | 5.55447 |
| CD630_05030 | CD630_n00460 | 132 -- 161 | 731 -- 759 | -13.7327 | 4.03823 | 5.55447 |
| CD630_08140 | CD630_n00460 | 85 -- 121 | 185 -- 227 | -13.711 | 3.30159 | 5.55447 |
| CD630_04850 | CD630_n00460 | 157 -- 188 | 249 -- 277 | -13.7061 | 4.40788 | 5.55447 |
| CD630_20170 | CD630_n00460 | 96 -- 193 | 820 -- 919 | -13.6929 | 6.90201 | 5.55447 |
| CD630_25050 | CD630_n00460 | 42 -- 82 | 815 -- 862 | -13.67 | 4.33624 | 5.55447 |
| gcvTPA | CD630_n00460 | 56 -- 105 | 204 -- 249 | -13.6553 | 4.77229 | 5.55447 |
| CD630_28320 | CD630_n00460 | 82 -- 99 | 546 -- 560 | -13.6478 | 3.99184 | 5.55447 |
| CD630_07470 | CD630_n00460 | 22 -- 56 | 1005 -- 1049 | -13.6091 | 3.37143 | 5.55447 |
| CD630_21570 | CD630_n00460 | 34 -- 176 | 118 -- 266 | -13.5819 | 6.48385 | 5.55447 |
| CD630_21490 | CD630_n00460 | 75 -- 112 | 216 -- 249 | -13.495 | 3.37462 | 5.55447 |
| CD630_19942 | CD630_n00460 | 87 -- 127 | 203 -- 236 | -13.4918 | 4.24574 | 5.55447 |
| CD630_05870 | CD630_n00460 | 168 -- 197 | 211 -- 242 | -13.4794 | 4.28339 | 5.55447 |
| CD630_32700 | CD630_n00460 | 117 -- 181 | 689 -- 748 | -13.4676 | 4.74276 | 5.55447 |
| CD630_13020 | CD630_n00460 | 146 -- 171 | 128 -- 148 | -13.4328 | 3.52267 | 5.55447 |
| CD630_14260 | CD630_n00460 | 13 -- 67 | 799 -- 864 | -13.428 | 3.22214 | 5.55447 |
| CD630_30140 | CD630_n00460 | 69 -- 106 | 203 -- 245 | -13.3697 | 3.28343 | 5.55447 |
| CD630_21030 | CD630_n00460 | 169 -- 195 | 757 -- 784 | -13.366 | 3.08733 | 5.55447 |
| CD630_27610 | CD630_n00460 | 12 -- 33 | 698 -- 718 | -13.3616 | 3.96465 | 5.55447 |
| CD630_06170 | CD630_n00460 | 101 -- 154 | 122 -- 192 | -13.3425 | 4.95366 | 5.55447 |
| CD630_05000 | CD630_n00460 | 140 -- 170 | 403 -- 433 | -13.3302 | 4.02442 | 5.55447 |
| cstA | CD630_n00460 | 64 -- 114 | 196 -- 243 | -13.3227 | 3.53584 | 5.55447 |
| CD630_08360 | CD630_n00460 | 148 -- 176 | 413 -- 441 | -13.2809 | 3.19339 | 5.55447 |
| CD630_05950 | CD630_n00460 | 95 -- 106 | 203 -- 214 | -13.278 | 3.31669 | 5.55447 |
| etfA | CD630_n00460 | 11 -- 37 | 1002 -- 1029 | -13.2714 | 4.80521 | 5.55447 |
| bcp | CD630_n00460 | 61 -- 107 | 404 -- 444 | -13.2711 | 3.16670 | 5.55447 |
| CD630_21140 | CD630_n00460 | 128 -- 181 | 347 -- 405 | -13.2674 | 3.33304 | 5.55447 |
| CD630_20980 | CD630_n00460 | 118 -- 149 | 215 -- 246 | -13.2478 | 8.12847 | 5.55447 |
| CD630_02830 | CD630_n00460 | 62 -- 104 | 205 -- 245 | -13.2389 | 3.57106 | 5.55447 |
| CD630_18090 | CD630_n00460 | 117 -- 157 | 528 -- 565 | -13.2221 | 4.23961 | 5.55447 |
| pilD | CD630_n00460 | 35 -- 88 | 411 -- 458 | -13.2127 | 4.54123 | 5.55447 |
| CD630_15360 | CD630_n00460 | 114 -- 139 | 949 -- 974 | -13.212 | 4.47208 | 5.55447 |
| CD630_20920 | CD630_n00460 | 89 -- 175 | 361 -- 447 | -13.1755 | 4.13812 | 5.55447 |
| CD630_11241 | CD630_n00460 | 162 -- 190 | 328 -- 357 | -13.1702 | 4.85723 | 5.55447 |
| CD630_01570 | CD630_n00460 | 128 -- 148 | 129 -- 148 | -13.1562 | 6.08699 | 5.55447 |
| CD630_18930 | CD630_n00460 | 111 -- 153 | 387 -- 436 | -13.068 | 5.32127 | 5.55447 |
| CD630_36180 | CD630_n00460 | 173 -- 184 | 213 -- 224 | -13.0633 | 4.83428 | 5.55447 |
| rolB | CD630_n00460 | 108 -- 132 | 101 -- 133 | -13.0599 | 3.60270 | 5.55447 |
| CD630_05490 | CD630_n00460 | 5 -- 37 | 401 -- 435 | -13.0383 | 3.89852 | 5.55447 |
| CD630_19801 | CD630_n00460 | 157 -- 180 | 356 -- 378 | -13.0332 | 5.22035 | 5.55447 |
| dpaA | CD630_n00460 | 26 -- 54 | 104 -- 133 | -13.032 | 4.65364 | 5.55447 |
| CD630_19430 | CD630_n00460 | 175 -- 198 | 554 -- 577 | -13.0025 | 3.13922 | 5.55447 |
| CD630_19440 | CD630_n00460 | 56 -- 105 | 112 -- 150 | -12.9849 | 4.53937 | 5.55447 |
| CD630_34520 | CD630_n00460 | 76 -- 162 | 825 -- 914 | -12.9711 | 3.83016 | 5.55447 |
| CD630_32880 | CD630_n00460 | 142 -- 188 | 473 -- 525 | -12.9661 | 3.41276 | 5.55447 |
| nrdD | CD630_n00460 | 13 -- 60 | 292 -- 355 | -12.9154 | 3.28290 | 5.55447 |
| CD630_31210 | CD630_n00460 | 147 -- 181 | 402 -- 440 | -12.9138 | 4.17693 | 5.55447 |
| CD630_07400 | CD630_n00460 | 36 -- 67 | 118 -- 150 | -12.8601 | 4.52273 | 5.55447 |
| xdhA | CD630_n00460 | 89 -- 121 | 213 -- 242 | -12.8271 | 4.45147 | 5.55447 |
| CD630_21700 | CD630_n00460 | 167 -- 198 | 396 -- 430 | -12.7857 | 3.96650 | 5.55447 |
| CD630_19780 | CD630_n00460 | 1 -- 31 | 202 -- 232 | -12.7748 | 3.64830 | 5.55447 |
| CD630_12320 | CD630_n00460 | 45 -- 75 | 410 -- 440 | -12.7704 | 3.28667 | 5.55447 |
| CD630_08130 | CD630_n00460 | 51 -- 89 | 202 -- 245 | -12.7666 | 4.63831 | 5.55447 |
| CD630_19680 | CD630_n00460 | 137 -- 154 | 350 -- 367 | -12.7086 | 3.58080 | 5.55447 |
| CD630_20450 | CD630_n00460 | 47 -- 78 | 409 -- 440 | -12.6947 | 3.77388 | 5.55447 |
| CD630_28181 | CD630_n00460 | 48 -- 77 | 402 -- 430 | -12.6903 | 3.91299 | 5.55447 |
| glgB | CD630_n00460 | 163 -- 201 | 204 -- 246 | -12.6537 | 4.48273 | 5.55447 |
| CD630_32990 | CD630_n00460 | 55 -- 104 | 101 -- 145 | -12.652 | 5.24925 | 5.55447 |
| CD630_14920 | CD630_n00460 | 32 -- 62 | 401 -- 430 | -12.6513 | 4.35225 | 5.55447 |
| CD630_22890 | CD630_n00460 | 8 -- 50 | 202 -- 249 | -12.6399 | 3.75591 | 5.55447 |
| CD630_06340 | CD630_n00460 | 14 -- 43 | 1002 -- 1031 | -12.6285 | 4.35983 | 5.55447 |
| CD630_28150 | CD630_n00460 | 60 -- 78 | 420 -- 440 | -12.6195 | 4.18447 | 5.55447 |
| pgm | CD630_n00460 | 16 -- 37 | 1003 -- 1024 | -12.564 | 3.45439 | 5.55447 |
| CD630_34530 | CD630_n00460 | 35 -- 120 | 330 -- 425 | -12.5417 | 3.83016 | 5.55447 |
| CD630_14990 | CD630_n00460 | 1 -- 49 | 333 -- 378 | -12.5416 | 3.78619 | 5.55447 |
| CD630_14210 | CD630_n00460 | 41 -- 73 | 118 -- 146 | -12.5349 | 3.82355 | 5.55447 |
| eutW | CD630_n00460 | 103 -- 195 | 796 -- 896 | -12.5343 | 4.55715 | 5.55447 |
| dltD | CD630_n00460 | 37 -- 106 | 203 -- 261 | -12.5239 | 4.40777 | 5.55447 |
| dltB | CD630_n00460 | 121 -- 147 | 648 -- 674 | -12.5205 | 3.81485 | 5.55447 |
| CD630_20670 | CD630_n00460 | 72 -- 92 | 291 -- 311 | -12.5174 | 4.42757 | 5.55447 |
| CD630_14700 | CD630_n00460 | 130 -- 159 | 210 -- 241 | -12.5094 | 3.98234 | 5.55447 |
| chrA | CD630_n00460 | 10 -- 47 | 1010 -- 1049 | -12.5092 | 5.27255 | 5.55447 |
| CD630_15060 | CD630_n00460 | 170 -- 194 | 129 -- 149 | -12.4863 | 3.69396 | 5.55447 |
| CD630_24920 | CD630_n00460 | 119 -- 150 | 410 -- 443 | -12.4809 | 3.89455 | 5.55447 |
| CD630_10000 | CD630_n00460 | 77 -- 90 | 350 -- 364 | -12.4788 | 3.86040 | 5.55447 |
| CD630_28550 | CD630_n00460 | 77 -- 109 | 275 -- 311 | -12.4715 | 3.51196 | 5.55447 |
| CD630_04540 | CD630_n00460 | 62 -- 107 | 395 -- 445 | -12.4425 | 3.33556 | 5.55447 |
| CD630_15461 | CD630_n00460 | 151 -- 184 | 989 -- 1022 | -12.396 | 9.13442 | 5.55447 |
| CD630_28870 | CD630_n00460 | 60 -- 160 | 126 -- 217 | -12.3889 | 3.41991 | 5.55447 |
| opuCA | CD630_n00460 | 56 -- 104 | 205 -- 246 | -12.3816 | 4.07035 | 5.55447 |
| CD630_13960 | CD630_n00460 | 27 -- 71 | 29 -- 75 | -12.3777 | 3.30375 | 5.55447 |
| CD630_14420 | CD630_n00460 | 147 -- 178 | 413 -- 443 | -12.3667 | 3.99497 | 5.55447 |
| CD630_24760 | CD630_n00460 | 79 -- 118 | 329 -- 369 | -12.3393 | 5.19485 | 5.55447 |
| CD630_33650 | CD630_n00460 | 121 -- 138 | 947 -- 964 | -12.3335 | 4.80565 | 5.55447 |
| CD630_19790 | CD630_n00460 | 49 -- 105 | 204 -- 249 | -12.3311 | 3.79593 | 5.55447 |
| CD630_32720 | CD630_n00460 | 37 -- 95 | 348 -- 402 | -12.3311 | 4.74276 | 5.55447 |
| CD630_31000 | CD630_n00460 | 43 -- 57 | 128 -- 142 | -12.3283 | 3.70946 | 5.55447 |
| nusB | CD630_n00460 | 136 -- 194 | 691 -- 745 | -12.3057 | 4.92798 | 5.55447 |
| CD630_06080 | CD630_n00460 | 15 -- 37 | 1004 -- 1029 | -12.2953 | 3.82315 | 5.55447 |
| CD630_36170 | CD630_n00460 | 3 -- 37 | 410 -- 434 | -12.256 | 3.72380 | 5.55447 |
| CD630_14810 | CD630_n00460 | 144 -- 155 | 284 -- 296 | -12.223 | 3.35323 | 5.55447 |
| CD630_16670 | CD630_n00460 | 112 -- 139 | 721 -- 746 | -12.2033 | 3.33553 | 5.55447 |
| CD630_10050 | CD630_n00460 | 156 -- 178 | 604 -- 624 | -12.1752 | 3.93750 | 5.55447 |
| fliI | CD630_n00460 | 42 -- 94 | 204 -- 250 | -12.1697 | 3.03666 | 5.55447 |
| panB | CD630_n00460 | 21 -- 163 | 819 -- 967 | -12.1647 | 3.40248 | 5.55447 |
| CD630_17170 | CD630_n00460 | 39 -- 171 | 286 -- 432 | -12.1446 | 4.10230 | 5.55447 |
| CD630_03650 | CD630_n00460 | 152 -- 193 | 32 -- 72 | -12.1405 | 3.93697 | 5.55447 |
| CD630_05290 | CD630_n00460 | 166 -- 200 | 402 -- 441 | -12.1281 | 3.97604 | 5.55447 |
| acoL | CD630_n00460 | 175 -- 196 | 338 -- 356 | -12.1155 | 3.32169 | 5.55447 |
| CD630_21770 | CD630_n00460 | 2 -- 13 | 3 -- 14 | -12.0304 | 4.02587 | 5.55447 |
| dapB2 | CD630_n00460 | 170 -- 198 | 398 -- 427 | -12.0053 | 3.27218 | 5.55447 |
| CD630_23080 | CD630_n00460 | 38 -- 107 | 867 -- 920 | -11.9804 | 5.23027 | 5.55447 |
| CD630_08110 | CD630_n00460 | 88 -- 101 | 545 -- 558 | -11.9773 | 4.39181 | 5.55447 |
| CD630_08100 | CD630_n00460 | 143 -- 176 | 208 -- 240 | -11.9593 | 4.39181 | 5.55447 |
| CD630_31880 | CD630_n00460 | 144 -- 191 | 389 -- 439 | -11.9355 | 3.97522 | 5.55447 |
| CD630_05150 | CD630_n00460 | 121 -- 138 | 135 -- 152 | -11.9294 | 3.69875 | 5.55447 |
| CD630_14410 | CD630_n00460 | 97 -- 166 | 558 -- 624 | -11.9293 | 3.39433 | 5.55447 |
| CD630_22950 | CD630_n00460 | 115 -- 155 | 408 -- 447 | -11.9115 | 3.96883 | 5.55447 |
| CD630_06140 | CD630_n00460 | 86 -- 93 | 289 -- 296 | -11.9003 | 4.12519 | 5.55447 |
| CD630_33560 | CD630_n00460 | 70 -- 124 | 844 -- 892 | -11.8884 | 4.03147 | 5.55447 |
| CD630_09100 | CD630_n00460 | 40 -- 89 | 843 -- 894 | -11.8853 | 5.92260 | 5.55447 |
| CD630_27770 | CD630_n00460 | 143 -- 180 | 512 -- 546 | -11.8836 | 4.08588 | 5.55447 |
| cdtR | CD630_n00460 | 4 -- 40 | 31 -- 73 | -11.8799 | 5.84368 | 5.55447 |
| CD630_22740 | CD630_n00460 | 55 -- 89 | 201 -- 246 | -11.8618 | 3.08733 | 5.55447 |
| CD630_35260 | CD630_n00460 | 58 -- 104 | 205 -- 245 | -11.8478 | 5.77277 | 5.55447 |
| CD630_22660 | CD630_n00460 | 17 -- 55 | 166 -- 204 | -11.8389 | 3.60496 | 5.55447 |
| iorA | CD630_n00460 | 2 -- 23 | 217 -- 240 | -11.8371 | 3.83173 | 5.55447 |
| CD630_29270 | CD630_n00460 | 40 -- 59 | 408 -- 427 | -11.8244 | 3.70174 | 5.55447 |
| CD630_09030 | CD630_n00460 | 110 -- 159 | 202 -- 248 | -11.8203 | 4.21282 | 5.55447 |
| CD630_19580 | CD630_n00460 | 185 -- 201 | 1011 -- 1027 | -11.8129 | 4.51902 | 5.55447 |
| CD630_17160 | CD630_n00460 | 89 -- 107 | 202 -- 220 | -11.8082 | 3.50228 | 5.55447 |
| CD630_18960 | CD630_n00460 | 35 -- 87 | 397 -- 438 | -11.7962 | 4.37822 | 5.55447 |
| CD630_24580 | CD630_n00460 | 28 -- 64 | 192 -- 225 | -11.794 | 3.88519 | 5.55447 |
| serS1 | CD630_n00460 | 132 -- 165 | 214 -- 250 | -11.7908 | 3.45858 | 5.55447 |
| spaK | CD630_n00460 | 130 -- 170 | 642 -- 675 | -11.744 | 3.04530 | 5.55447 |
| CD630_23000 | CD630_n00460 | 92 -- 123 | 210 -- 241 | -11.7309 | 3.54130 | 5.55447 |
| pilC | CD630_n00460 | 42 -- 68 | 413 -- 438 | -11.6823 | 3.54230 | 5.55447 |
| CD630_13600 | CD630_n00460 | 88 -- 106 | 396 -- 415 | -11.6624 | 4.45481 | 5.55447 |
| srlD | CD630_n00460 | 101 -- 133 | 213 -- 242 | -11.6377 | 4.19843 | 5.55447 |
| CD630_18311 | CD630_n00460 | 88 -- 107 | 202 -- 218 | -11.6353 | 6.86047 | 5.55447 |
| CD630_25620 | CD630_n00460 | 85 -- 116 | 85 -- 123 | -11.6345 | 3.02685 | 5.55447 |
| CD630_23760 | CD630_n00460 | 45 -- 112 | 685 -- 747 | -11.6296 | 4.54307 | 5.55447 |
| CD630_01560 | CD630_n00460 | 134 -- 200 | 186 -- 245 | -11.6199 | 4.06352 | 5.55447 |
| CD630_08460 | CD630_n00460 | 19 -- 45 | 1004 -- 1034 | -11.5893 | 4.15083 | 5.55447 |
| CD630_02790 | CD630_n00460 | 86 -- 163 | 698 -- 754 | -11.5859 | 4.77661 | 5.55447 |
| CD630_19030 | CD630_n00460 | 143 -- 182 | 112 -- 151 | -11.5646 | 3.34240 | 5.55447 |
| CD630_03420 | CD630_n00460 | 147 -- 160 | 943 -- 956 | -11.5614 | 4.71868 | 5.55447 |

| | | | | | | |
|---|---|---|---|---|---|---|
| kdpD | CD630_n00460 | 59 – 93 | 214 – 247 | -11.54 | 4.27140 | 5.55447 |
| CD630_06690 | CD630_n00460 | 15 – 49 | 400 – 435 | -11.5183 | 5.82709 | 5.55447 |
| CD630_18610 | CD630_n00460 | 111 – 160 | 61 – 114 | -11.5129 | 4.15582 | 5.55447 |
| CD630_17960 | CD630_n00460 | 18 – 64 | 329 – 364 | -11.5118 | 3.71404 | 5.55447 |
| cprK | CD630_n00460 | 142 – 167 | 21 – 47 | -11.5017 | 3.96216 | 5.55447 |
| CD630_10080 | CD630_n00460 | 21 – 41 | 422 – 444 | -11.5002 | 3.75625 | 5.55447 |
| CD630_06110 | CD630_n00460 | 54 – 105 | 478 – 526 | -11.498 | 5.61380 | 5.55447 |
| CD630_31580 | CD630_n00460 | 85 – 92 | 291 – 298 | -11.4948 | 3.51756 | 5.55447 |
| CD630_16530 | CD630_n00460 | 169 – 200 | 405 – 439 | -11.4881 | 4.37658 | 5.55447 |
| CD630_05211 | CD630_n00460 | 125 – 149 | 863 – 887 | -11.4855 | 3.97048 | 5.55447 |
| CD630_04620 | CD630_n00460 | 21 – 79 | 823 – 888 | -11.468 | 3.45413 | 5.55447 |
| CD630_18010 | CD630_n00460 | 181 – 200 | 668 – 688 | -11.4594 | 4.21967 | 5.55447 |
| CD630_08080 | CD630_n00460 | 1 – 21 | 651 – 669 | -11.4352 | 4.87819 | 5.55447 |
| CD630_31970 | CD630_n00460 | 67 – 83 | 133 – 148 | -11.4253 | 3.24511 | 5.55447 |
| glgP | CD630_n00460 | 1 – 34 | 399 – 427 | -11.3939 | 3.28530 | 5.55447 |
| CD630_21730 | CD630_n00460 | 135 – 171 | 1003 – 1029 | -11.3688 | 3.45076 | 5.55447 |
| CD630_31381 | CD630_n00460 | 87 – 127 | 37 – 72 | -11.362 | 3.49137 | 5.55447 |
| CD630_30120 | CD630_n00460 | 79 – 151 | 628 – 707 | -11.3416 | 4.04859 | 5.55447 |
| CD630_06060 | CD630_n00460 | 94 – 115 | 1006 – 1028 | -11.329 | 4.94050 | 5.55447 |
| CD630_32070 | CD630_n00460 | 38 – 83 | 202 – 247 | -11.324 | 3.51964 | 5.55447 |
| cobQ | CD630_n00460 | 101 – 108 | 291 – 298 | -11.3109 | 3.13524 | 5.55447 |
| CD630_18910 | CD630_n00460 | 12 – 118 | 814 – 929 | -11.3081 | 3.30688 | 5.55447 |
| ssuC | CD630_n00460 | 59 – 75 | 731 – 747 | -11.295 | 3.45491 | 5.55447 |
| CD630_06920 | CD630_n00460 | 47 – 93 | 283 – 333 | -11.291 | 3.13260 | 5.55447 |
| CD630_14620 | CD630_n00460 | 166 – 190 | 415 – 439 | -11.2896 | 4.43651 | 5.55447 |
| CD630_32690 | CD630_n00460 | 68 – 182 | 203 – 311 | -11.285 | 7.32676 | 5.55447 |
| CD630_27280 | CD630_n00460 | 6 – 65 | 30 – 78 | -11.284 | 3.24738 | 5.55447 |
| CD630_32050 | CD630_n00460 | 42 – 69 | 1003 – 1030 | -11.2812 | 5.73797 | 5.55447 |
| CD630_04880 | CD630_n00460 | 100 – 145 | 988 – 1032 | -11.279 | 4.62291 | 5.55447 |
| CD630_36250 | CD630_n00460 | 50 – 107 | 202 – 240 | -11.24 | 4.13224 | 5.55447 |
| CD630_15200 | CD630_n00460 | 59 – 112 | 384 – 438 | -11.2129 | 4.00442 | 5.55447 |
| CD630_36781 | CD630_n00460 | 24 – 52 | 681 – 711 | -11.2038 | 3.75946 | 5.55447 |
| CD630_30420 | CD630_n00460 | 145 – 199 | 808 – 867 | -11.1973 | 5.27539 | 5.55447 |
| CD630_07121 | CD630_n00460 | 6 – 59 | 126 – 171 | -11.1952 | 3.26177 | 5.55447 |
| uxaA%27 | CD630_n00460 | 123 – 151 | 998 – 1031 | -11.1855 | 4.26165 | 5.55447 |
| CD630_03220 | CD630_n00460 | 49 – 91 | 1003 – 1047 | -11.1702 | 3.36746 | 5.55447 |
| CD630_10950 | CD630_n00460 | 135 – 156 | 877 – 899 | -11.1512 | 3.67675 | 5.55447 |
| CD630_18120 | CD630_n00460 | 89 – 107 | 202 – 220 | -11.1272 | 3.99713 | 5.55447 |
| CD630_31382 | CD630_n00460 | 5 – 28 | 203 – 219 | -11.1115 | 3.49137 | 5.55447 |
| CD630_30110 | CD630_n00460 | 100 – 149 | 851 – 903 | -11.1058 | 3.31041 | 5.55447 |
| CD630_17970 | CD630_n00460 | 119 – 173 | 225 – 277 | -11.1046 | 3.68802 | 5.55447 |
| CD630_30540 | CD630_n00460 | 120 – 175 | 292 – 357 | -11.0998 | 3.63804 | 5.55447 |
| CD630_21790 | CD630_n00460 | 132 – 146 | 550 – 564 | -11.0874 | 4.93762 | 5.55447 |
| CD630_20171 | CD630_n00460 | 86 – 166 | 218 – 297 | -11.0802 | 6.87491 | 5.55447 |
| CD630_21090 | CD630_n00460 | 109 – 148 | 40 – 73 | -11.0799 | 4.15803 | 5.55447 |
| CD630_13780 | CD630_n00460 | 161 – 184 | 347 – 372 | -11.0328 | 3.52274 | 5.55447 |
| CD630_21110 | CD630_n00460 | 143 – 163 | 725 – 746 | -11.017 | 3.89630 | 5.55447 |
| CD630_22880 | CD630_n00460 | 37 – 48 | 292 – 303 | -11.011 | 3.62455 | 5.55447 |
| eutP | CD630_n00460 | 107 – 123 | 294 – 311 | -11.002 | 3.82847 | 5.55447 |
| CD630_23861 | CD630_n00460 | 88 – 94 | 290 – 296 | -10.9999 | 5.83068 | 5.55447 |
| CD630_22991 | CD630_n00460 | 187 – 196 | 665 – 674 | -10.9716 | 6.03728 | 5.55447 |
| CD630_27120 | CD630_n00460 | 139 – 155 | 215 – 233 | -10.9477 | 4.43720 | 5.55447 |
| bltD | CD630_n00460 | 110 – 145 | 409 – 443 | -10.9423 | 4.49926 | 5.55447 |
| CD630_20600 | CD630_n00460 | 4 – 73 | 845 – 912 | -10.9403 | 4.20093 | 5.55447 |
| CD630_01640 | CD630_n00460 | 79 – 105 | 409 – 434 | -10.937 | 4.55381 | 5.55447 |
| CD630_01690 | CD630_n00460 | 81 – 176 | 646 – 749 | -10.9213 | 4.97234 | 5.55447 |
| CD630_27210 | CD630_n00460 | 68 – 109 | 271 – 310 | -10.9199 | 4.18786 | 5.55447 |
| CD630_06150 | CD630_n00460 | 88 – 108 | 201 – 219 | -10.9119 | 5.51744 | 5.55447 |
| ppnK | CD630_n00460 | 124 – 177 | 872 – 919 | -10.9019 | 4.36817 | 5.55447 |
| CD630_27570 | CD630_n00460 | 97 – 169 | 298 – 379 | -10.8538 | 3.46216 | 5.55447 |
| CD630_10970 | CD630_n00460 | 48 – 83 | 324 – 367 | -10.8361 | 3.53850 | 5.55447 |
| CD630_04610 | CD630_n00460 | 13 – 51 | 779 – 818 | -10.8285 | 3.83025 | 5.55447 |
| CD630_14110 | CD630_n00460 | 7 – 49 | 21 – 61 | -10.8143 | 3.39550 | 5.55447 |
| feoB | CD630_n00460 | 84 – 97 | 208 – 222 | -10.8103 | 4.52257 | 5.55447 |
| CD630_30730 | CD630_n00460 | 8 – 52 | 120 – 153 | -10.8101 | 3.13236 | 5.55447 |
| CD630_23770 | CD630_n00460 | 175 – 185 | 346 – 356 | -10.796 | 3.25181 | 5.55447 |
| CD630_05830 | CD630_n00460 | 158 – 179 | 117 – 138 | -10.7764 | 3.35360 | 5.55447 |
| CD630_28280 | CD630_n00460 | 80 – 118 | 997 – 1034 | -10.7748 | 4.59833 | 5.55447 |
| CD630_18680 | CD630_n00460 | 34 – 75 | 1007 – 1045 | -10.7706 | 3.04315 | 5.55447 |
| CD630_31960 | CD630_n00460 | 32 – 82 | 23 – 74 | -10.7645 | 4.93757 | 5.55447 |
| CD630_14240 | CD630_n00460 | 75 – 120 | 204 – 249 | -10.7583 | 8.60766 | 5.55447 |
| flgB | CD630_n00460 | 152 – 195 | 822 – 863 | -10.7477 | 3.25736 | 5.55447 |
| CD630_18940 | CD630_n00460 | 46 – 108 | 779 – 827 | -10.7442 | 4.09988 | 5.55447 |
| CD630_07760 | CD630_n00460 | 73 – 105 | 409 – 440 | -10.7425 | 3.64186 | 5.55447 |
| cwp17 | CD630_n00460 | 186 – 196 | 203 – 213 | -10.7408 | 3.43363 | 5.55447 |
| CD630_07070 | CD630_n00460 | 24 – 47 | 649 – 672 | -10.7278 | 3.46414 | 5.55447 |
| CD630_27060 | CD630_n00460 | 96 – 133 | 203 – 242 | -10.7139 | 4.06049 | 5.55447 |
| CD630_05450 | CD630_n00460 | 20 – 61 | 401 – 440 | -10.696 | 5.04761 | 5.55447 |
| CD630_02420 | CD630_n00460 | 127 – 148 | 548 – 570 | -10.6957 | 3.71048 | 5.55447 |
| CD630_04940 | CD630_n00460 | 19 – 67 | 94 – 147 | -10.6856 | 4.20948 | 5.55447 |
| CD630_15080 | CD630_n00460 | 92 – 169 | 137 – 221 | -10.6828 | 4.03033 | 5.55447 |
| CD630_34030 | CD630_n00460 | 174 – 200 | 119 – 150 | -10.6754 | 3.05766 | 5.55447 |
| rolA | CD630_n00460 | 99 – 179 | 609 – 711 | -10.6646 | 4.56362 | 5.55447 |
| CD630_17292 | CD630_n00460 | 143 – 191 | 324 – 378 | -10.6559 | 6.25799 | 5.55447 |
| CD630_16940 | CD630_n00460 | 85 – 105 | 204 – 219 | -10.6522 | 4.14665 | 5.55447 |
| blaI | CD630_n00460 | 44 – 148 | 203 – 350 | -10.6496 | 3.73674 | 5.55447 |
| CD630_27630 | CD630_n00460 | 32 – 64 | 104 – 145 | -10.6459 | 3.54199 | 5.55447 |
| CD630_05880 | CD630_n00460 | 128 – 190 | 394 – 435 | -10.6407 | 4.28339 | 5.55447 |
| appB | CD630_n00460 | 94 – 174 | 972 – 1052 | -10.6198 | 6.44103 | 5.55447 |
| CD630_29750 | CD630_n00460 | 71 – 94 | 109 – 133 | -10.6179 | 4.59146 | 5.55447 |
| CD630_26050 | CD630_n00460 | 42 – 60 | 258 – 278 | -10.6049 | 6.99851 | 5.55447 |
| CD630_05230 | CD630_n00460 | 30 – 55 | 1003 – 1028 | -10.5958 | 3.73787 | 5.55447 |
| CD630_08940 | CD630_n00460 | 55 – 95 | 212 – 250 | -10.5946 | 3.72225 | 5.55447 |
| CD630_26150 | CD630_n00460 | 130 – 185 | 37 – 82 | -10.5787 | 4.06302 | 5.55447 |
| CD630_14930 | CD630_n00460 | 9 – 25 | 516 – 532 | -10.5617 | 3.31335 | 5.55447 |
| CD630_32710 | CD630_n00460 | 139 – 157 | 215 – 241 | -10.5612 | 4.74276 | 5.55447 |
| CD630_27520 | CD630_n00460 | 134 – 158 | 607 – 632 | -10.5484 | 4.27693 | 5.55447 |
| CD630_27230 | CD630_n00460 | 108 – 150 | 853 – 900 | -10.5469 | 3.52462 | 5.55447 |
| CD630_09990 | CD630_n00460 | 120 – 164 | 209 – 246 | -10.5398 | 3.86040 | 5.55447 |
| CD630_05900 | CD630_n00460 | 23 – 63 | 21 – 57 | -10.5362 | 3.21934 | 5.55447 |
| CD630_02740 | CD630_n00460 | 79 – 94 | 350 – 367 | -10.5286 | 5.02277 | 5.55447 |
| CD630_04410 | CD630_n00460 | 84 – 91 | 558 – 565 | -10.5234 | 3.90451 | 5.55447 |
| kdgT1 | CD630_n00460 | 31 – 73 | 801 – 860 | -10.5174 | 4.49207 | 5.55447 |
| CD630_06190 | CD630_n00460 | 71 – 117 | 324 – 371 | -10.4977 | 3.57064 | 5.55447 |
| CD630_24050 | CD630_n00460 | 84 – 95 | 208 – 220 | -10.4975 | 3.70192 | 5.55447 |
| CD630_25310 | CD630_n00460 | 96 – 113 | 1005 – 1026 | -10.4929 | 3.99861 | 5.55447 |
| hfq | CD630_n00460 | 146 – 155 | 549 – 558 | -10.4852 | 5.05609 | 5.55447 |
| cobS | CD630_n00460 | 57 – 106 | 203 – 250 | -10.4801 | 3.67481 | 5.55447 |
| cheW | CD630_n00460 | 39 – 95 | 288 – 354 | -10.48 | 3.72371 | 5.55447 |
| CD630_31500 | CD630_n00460 | 92 – 121 | 409 – 438 | -10.4707 | 4.90207 | 5.55447 |
| CD630_19700 | CD630_n00460 | 85 – 102 | 339 – 357 | -10.4697 | 3.85861 | 5.55447 |
| CD630_23450 | CD630_n00460 | 69 – 126 | 842 – 895 | -10.4684 | 3.27218 | 5.55447 |
| vncR | CD630_n00460 | 126 – 170 | 599 – 639 | -10.4531 | 3.93642 | 5.55447 |
| CD630_24220 | CD630_n00460 | 32 – 54 | 684 – 706 | -10.4399 | 4.18037 | 5.55447 |
| CD630_03040 | CD630_n00460 | 179 – 198 | 870 – 889 | -10.4199 | 3.51403 | 5.55447 |
| CD630_18410 | CD630_n00460 | 132 – 168 | 121 – 157 | -10.4053 | 3.59739 | 5.55447 |
| CD630_06560 | CD630_n00460 | 3 – 45 | 408 – 436 | -10.3792 | 4.02532 | 5.55447 |
| CD630_24770 | CD630_n00460 | 158 – 201 | 324 – 369 | -10.3638 | 3.20798 | 5.55447 |
| CD630_15190 | CD630_n00460 | 80 – 93 | 350 – 364 | -10.3636 | 4.43255 | 5.55447 |
| CD630_06060 | CD630_n00460 | 90 – 165 | 184 – 257 | -10.3416 | 4.94050 | 5.55447 |
| CD630_02430 | CD630_n00460 | 54 – 110 | 399 – 440 | -10.3254 | 3.71048 | 5.55447 |
| CD630_16330 | CD630_n00460 | 145 – 197 | 820 – 875 | -10.3142 | 4.11180 | 5.55447 |
| CD630_05821 | CD630_n00460 | 69 – 87 | 350 – 367 | -10.2858 | 3.75661 | 5.55447 |
| CD630_33682 | CD630_n00460 | 69 – 89 | 291 – 309 | -10.2791 | 5.77720 | 5.55447 |
| CD630_18100 | CD630_n00460 | 80 – 122 | 186 – 225 | -10.2752 | 5.71658 | 5.55447 |
| CD630_17360 | CD630_n00460 | 8 – 37 | 516 – 545 | -10.2583 | 5.66070 | 5.55447 |
| ogt1 | CD630_n00460 | 6 – 35 | 520 – 556 | -10.2263 | 3.16772 | 5.55447 |
| CD630_30620 | CD630_n00460 | 1 – 35 | 820 – 856 | -10.2249 | 4.15443 | 5.55447 |
| CD630_19903 | CD630_n00460 | 64 – 89 | 291 – 314 | -10.2222 | 5.79994 | 5.55447 |
| CD630_07770 | CD630_n00460 | 47 – 94 | 1003 – 1045 | -10.2179 | 3.31158 | 5.55447 |
| CD630_29250 | CD630_n00460 | 167 – 197 | 477 – 509 | -10.2137 | 4.61901 | 5.55447 |
| CD630_09410 | CD630_n00460 | 167 – 197 | 477 – 509 | -10.2137 | 4.75651 | 5.55447 |
| gabT | CD630_n00460 | 9 – 60 | 844 – 895 | -10.2096 | 3.67067 | 5.55447 |
| CD630_32040 | CD630_n00460 | 137 – 161 | 1002 – 1024 | -10.2021 | 4.93516 | 5.55447 |
| CD630_24591 | CD630_n00460 | 7 – 46 | 661 – 704 | -10.1793 | 3.54497 | 5.55447 |
| CD630_04770 | CD630_n00460 | 81 – 99 | 545 – 565 | -10.1665 | 3.58478 | 5.55447 |
| CD630_20960 | CD630_n00460 | 156 – 183 | 216 – 249 | -10.1609 | 5.65940 | 5.55447 |
| CD630_27310 | CD630_n00460 | 80 – 118 | 392 – 438 | -10.1552 | 4.01622 | 5.55447 |
| CD630_29960 | CD630_n00460 | 98 – 198 | 340 – 445 | -10.1496 | 4.51904 | 5.55447 |
| CD630_35780 | CD630_n00460 | 29 – 49 | 1010 – 1031 | -10.1299 | 3.50107 | 5.55447 |
| CD630_15160 | CD630_n00460 | 106 – 128 | 1002 – 1029 | -10.1297 | 4.32318 | 5.55447 |
| CD630_19800 | CD630_n00460 | 63 – 90 | 409 – 436 | -10.1199 | 5.22035 | 5.55447 |
| CD630_13620 | CD630_n00460 | 113 – 196 | 557 – 635 | -10.119 | 4.77627 | 5.55447 |
| CD630_18970 | CD630_n00460 | 123 – 174 | 401 – 447 | -10.1171 | 3.57243 | 5.55447 |
| CD630_19401 | CD630_n00460 | 17 – 78 | 29 – 81 | -10.1166 | 4.37084 | 5.55447 |
| CD630_28290 | CD630_n00460 | 49 – 74 | 650 – 677 | -10.0957 | 5.54192 | 5.55447 |
| CD630_32500 | CD630_n00460 | 57 – 89 | 1004 – 1036 | -10.0772 | 6.17288 | 5.55447 |
| cdu1 | CD630_n00460 | 16 – 28 | 205 – 216 | -10.0723 | 4.46276 | 5.55447 |
| CD630_14050 | CD630_n00460 | 160 – 200 | 733 – 767 | -10.0677 | 3.30894 | 5.55447 |
| flhB | CD630_n00460 | 15 – 29 | 208 – 221 | -10.0663 | 3.55524 | 5.55447 |
| CD630_08390 | CD630_n00460 | 79 – 179 | 404 – 501 | -10.0558 | 5.16000 | 5.55447 |
| CD630_07870 | CD630_n00460 | 156 – 191 | 866 – 900 | -10.0531 | 3.22829 | 5.55447 |
| cspC | CD630_n00460 | 108 – 143 | 212 – 242 | -10.0524 | 5.48848 | 5.55447 |
| CD630_03920 | CD630_n00460 | 132 – 168 | 837 – 870 | -10.0497 | 4.15934 | 5.55447 |
| CD630_20290 | CD630_n00460 | 47 – 84 | 42 – 79 | -10.0402 | 4.38183 | 5.55447 |
| aroD | CD630_n00460 | 91 – 106 | 203 – 218 | -10.0389 | 3.71929 | 5.55447 |
| CD630_17730 | CD630_n00460 | 169 – 197 | 182 – 210 | -10.0362 | 4.73397 | 5.55447 |
| CD630_09050 | CD630_n00460 | 1 – 49 | 716 – 762 | -10.0214 | 3.96216 | 5.55447 |
| uxaA | CD630_n00460 | 72 – 98 | 208 – 234 | -10.0083 | 4.26165 | 5.55447 |
| agrB | CD630_n00460 | 111 – 131 | 476 – 497 | -9.99261 | 3.92598 | 5.55447 |
| CD630_25610 | CD630_n00460 | 98 – 140 | 1001 – 1043 | -9.99109 | 3.02685 | 5.55447 |
| CD630_22550 | CD630_n00460 | 154 – 181 | 516 – 544 | -9.98681 | 3.62379 | 5.55447 |
| htpG | CD630_n00460 | 31 – 71 | 302 – 341 | -9.96154 | 3.45147 | 5.55447 |
| CD630_16150 | CD630_n00460 | 139 – 196 | 647 – 707 | -9.94603 | 4.20614 | 5.55447 |
| CD630_27200 | CD630_n00460 | 26 – 42 | 745 – 762 | -9.94143 | 3.42516 | 5.55447 |
| CD630_21540 | CD630_n00460 | 37 – 90 | 1002 – 1049 | -9.93778 | 4.33090 | 5.55447 |
| CD630_05590 | CD630_n00460 | 67 – 97 | 214 – 245 | -9.93059 | 3.39670 | 5.55447 |
| ribD | CD630_n00460 | 88 – 111 | 598 – 618 | -9.90216 | 6.65922 | 5.55447 |
| thiH | CD630_n00460 | 69 – 144 | 359 – 432 | -9.89725 | 3.58230 | 5.55447 |
| CD630_21910 | CD630_n00460 | 71 – 97 | 1003 – 1030 | -9.89264 | 4.66600 | 5.55447 |
| CD630_35840 | CD630_n00460 | 68 – 126 | 38 – 101 | -9.87909 | 3.14170 | 5.55447 |
| CD630_27040 | CD630_n00460 | 107 – 127 | 128 – 148 | -9.86635 | 3.51812 | 5.55447 |
| CD630_31220 | CD630_n00460 | 26 – 74 | 727 – 782 | -9.86263 | 3.73779 | 5.55447 |
| CD630_05300 | CD630_n00460 | 163 – 185 | 419 – 442 | -9.85812 | 4.56885 | 5.55447 |
| CD630_19000 | CD630_n00460 | 145 – 174 | 1004 – 1030 | -9.85358 | 4.18443 | 5.55447 |
| CD630_22050 | CD630_n00460 | 139 – 191 | 844 – 896 | -9.84939 | 4.87639 | 5.55447 |
| CD630_28560 | CD630_n00460 | 69 – 93 | 351 – 373 | -9.84621 | 3.96803 | 5.55447 |
| CD630_14360 | CD630_n00460 | 46 – 103 | 211 – 250 | -9.83681 | 3.02079 | 5.55447 |
| rbsA | CD630_n00460 | 2 – 23 | 874 – 896 | -9.82397 | 3.71650 | 5.55447 |
| CD630_35790 | CD630_n00460 | 152 – 158 | 290 – 296 | -9.81767 | 6.97665 | 5.55447 |
| CD630_25070 | CD630_n00460 | 125 – 152 | 103 – 131 | -9.80318 | 3.45164 | 5.55447 |
| feoB | CD630_n00460 | 143 – 153 | 487 – 497 | -9.78996 | 4.05364 | 5.55447 |
| CD630_20650 | CD630_n00460 | 46 – 122 | 649 – 731 | -9.75837 | 4.00083 | 5.55447 |
| CD630_27600 | CD630_n00460 | 88 – 119 | 731 – 755 | -9.75687 | 3.48384 | 5.55447 |
| CD630_30310 | CD630_n00460 | 60 – 105 | 204 – 244 | -9.75597 | 4.56059 | 5.55447 |
| CD630_08370 | CD630_n00460 | 146 – 153 | 404 – 411 | -9.74683 | 3.65639 | 5.55447 |
| CD630_29650 | CD630_n00460 | 156 – 184 | 129 – 153 | -9.74436 | 3.67128 | 5.55447 |
| CD630_06410 | CD630_n00460 | 119 – 177 | 103 – 155 | -9.72631 | 4.14404 | 5.55447 |
| CD630_11240 | CD630_n00460 | 69 – 88 | 291 – 311 | -9.72473 | 3.20274 | 5.55447 |
| CD630_09770 | CD630_n00460 | 6 – 50 | 142 – 183 | -9.72133 | 3.22106 | 5.55447 |
| CD630_25200 | CD630_n00460 | 158 – 173 | 335 – 351 | -9.71843 | 3.65158 | 5.55447 |
| CD630_06960 | CD630_n00460 | 99 – 124 | 947 – 973 | -9.707 | 3.16317 | 5.55447 |
| CD630_23090 | CD630_n00460 | 166 – 199 | 409 – 445 | -9.69582 | 4.25622 | 5.55447 |
| CD630_15111 | CD630_n00460 | 2 – 42 | 98 – 140 | -9.69373 | 4.18664 | 5.55447 |
| CD630_17900 | CD630_n00460 | 177 – 198 | 404 – 425 | -9.66528 | 3.52167 | 5.55447 |
| CD630_09060 | CD630_n00460 | 100 – 145 | 685 – 735 | -9.65699 | 4.31834 | 5.55447 |
| CD630_08780 | CD630_n00460 | 170 – 200 | 113 – 143 | -9.65425 | 7.76690 | 5.55447 |
| spoVFB | CD630_n00460 | 164 – 200 | 100 – 139 | -9.64169 | 4.62758 | 5.55447 |
| CD630_05820 | CD630_n00460 | 41 – 82 | 204 – 248 | -9.63933 | 3.75661 | 5.55447 |
| CD630_29840 | CD630_n00460 | 104 – 116 | 1022 – 1034 | -9.63123 | 4.48825 | 5.55447 |
| CD630_14180 | CD630_n00460 | 112 – 161 | 846 – 892 | -9.62913 | 3.93696 | 5.55447 |
| tcdA | CD630_n00460 | 165 – 180 | 561 – 576 | -9.62627 | 4.70593 | 5.55447 |
| CD630_22030 | CD630_n00460 | 176 – 200 | 751 – 776 | -9.6194 | 3.29934 | 5.55447 |
| CD630_21260 | CD630_n00460 | 117 – 147 | 113 – 146 | -9.61677 | 4.22255 | 5.55447 |
| CD630_30190 | CD630_n00460 | 168 – 194 | 103 – 143 | -9.61308 | 3.27394 | 5.55447 |
| CD630_13930 | CD630_n00460 | 88 – 111 | 726 – 746 | -9.58758 | 4.09246 | 5.55447 |
| CD630_15550 | CD630_n00460 | 185 – 197 | 668 – 680 | -9.58157 | 3.49019 | 5.55447 |
| CD630_21720 | CD630_n00460 | 155 – 175 | 615 – 632 | -9.57524 | 4.13030 | 5.55447 |
| CD630_16540 | CD630_n00460 | 168 – 190 | 679 – 702 | -9.55898 | 4.37658 | 5.55447 |
| dnaC | CD630_n00460 | 131 – 162 | 404 – 435 | -9.55466 | 6.30796 | 5.55447 |
| CD630_19921 | CD630_n00460 | 75 – 101 | 1006 – 1033 | -9.55365 | 5.46345 | 5.55447 |
| oppC | CD630_n00460 | 66 – 79 | 160 – 173 | -9.52689 | 3.01695 | 5.55447 |
| CD630_21750 | CD630_n00460 | 82 – 119 | 698 – 737 | -9.50645 | 3.70626 | 5.55447 |
| CD630_23280 | CD630_n00460 | 104 – 121 | 339 – 356 | -9.50092 | 3.50093 | 5.55447 |
| CD630_04850 | CD630_n00510 | 26 – 170 | 784 – 932 | -19.2112 | 4.40788 | 3.92762 |
| CD630_06060 | CD630_n00510 | 147 – 186 | 123 – 157 | -18.6374 | 4.94050 | 3.92762 |
| CD630_14260 | CD630_n00510 | 22 – 70 | 213 – 261 | -17.2109 | 3.22214 | 3.92762 |
| CD630_06540 | CD630_n00510 | 42 – 120 | 844 – 907 | -17.0865 | 3.54673 | 3.92762 |
| CD630_05870 | CD630_n00510 | 168 – 197 | 748 – 776 | -16.8411 | 4.28339 | 3.92762 |
| CD630_24580 | CD630_n00510 | 11 – 46 | 104 – 139 | -16.321 | 3.88519 | 3.92762 |
| CD630_18610 | CD630_n00510 | 74 – 155 | 848 – 925 | -16.2198 | 4.15582 | 3.92762 |
| CD630_09100 | CD630_n00510 | 21 – 95 | 332 – 404 | -15.8848 | 5.92260 | 3.92762 |
| appC | CD630_n00510 | 43 – 76 | 160 – 195 | -15.5352 | 4.50438 | 3.92762 |
| CD630_06080 | CD630_n00510 | 163 – 196 | 746 – 775 | -15.3398 | 3.82315 | 3.92762 |
| bglA | CD630_n00510 | 68 – 94 | 484 – 511 | -15.2736 | 4.50315 | 3.92762 |
| CD630_08460 | CD630_n00510 | 8 – 47 | 212 – 246 | -15.19 | 4.15083 | 3.92762 |
| CD630_27240 | CD630_n00510 | 134 – 172 | 786 – 826 | -14.9982 | 3.52462 | 3.92762 |
| CD630_06690 | CD630_n00510 | 21 – 68 | 732 – 768 | -14.8911 | 5.82709 | 3.92762 |
| CD630_13560 | CD630_n00510 | 73 – 104 | 89 – 122 | -14.8904 | 3.38083 | 3.92762 |
| folC | CD630_n00510 | 46 – 134 | 520 – 595 | -14.6915 | 3.85634 | 3.92762 |
| pilD | CD630_n00510 | 2 – 63 | 501 – 565 | -14.3763 | 4.54123 | 3.92762 |
| CD630_34050 | CD630_n00510 | 133 – 177 | 619 – 678 | -14.3737 | 4.03863 | 3.92762 |
| CD630_25150 | CD630_n00510 | 150 – 201 | 10 – 43 | -14.2859 | 3.79339 | 3.92762 |
| CD630_24920 | CD630_n00510 | 88 – 149 | 485 – 549 | -14.2416 | 3.89455 | 3.92762 |
| CD630_10220 | CD630_n00510 | 144 – 188 | 227 – 269 | -14.2181 | 4.08233 | 3.92762 |
| CD630_31490 | CD630_n00510 | 102 – 122 | 101 – 122 | -14.1701 | 4.08593 | 3.92762 |
| CD630_30100 | CD630_n00510 | 24 – 64 | 756 – 796 | -14.1589 | 4.13234 | 3.92762 |
| CD630_28550 | CD630_n00510 | 68 – 110 | 735 – 781 | -14.0389 | 3.51196 | 3.92762 |
| CD630_18230 | CD630_n00510 | 79 – 166 | 207 – 307 | -13.9999 | 3.00184 | 3.92762 |
| bglA | CD630_n00510 | 158 – 201 | 734 – 783 | -13.8788 | 4.86402 | 3.92762 |
| glgB | CD630_n00510 | 136 – 191 | 555 – 609 | -13.8738 | 4.48273 | 3.92762 |
| dnaD | CD630_n00510 | 5 – 27 | 721 – 743 | -13.8567 | 3.56078 | 3.92762 |
| CD630_31000 | CD630_n00510 | 84 – 152 | 720 – 779 | -13.8549 | 3.70946 | 3.92762 |
| CD630_32520 | CD630_n00510 | 123 – 196 | 546 – 610 | -13.7922 | 4.06098 | 3.92762 |
| opuCC | CD630_n00510 | 98 – 135 | 927 – 962 | -13.5965 | 4.07035 | 3.92762 |
| CD630_21750 | CD630_n00510 | 83 – 126 | 829 – 881 | -13.454 | 3.70626 | 3.92762 |
| CD630_19942 | CD630_n00510 | 89 – 134 | 229 – 278 | -13.3756 | 4.24574 | 3.92762 |
| CD630_04610 | CD630_n00510 | 147 – 184 | 6 – 41 | -13.3511 | 3.83025 | 3.92762 |
| CD630_11220 | CD630_n00510 | 138 – 167 | 749 – 783 | -13.3479 | 3.52327 | 3.92762 |
| CD630_08110 | CD630_n00510 | 77 – 172 | 801 – 904 | -13.2846 | 4.39181 | 3.92762 |
| CD630_32040 | CD630_n00510 | 166 – 192 | 943 – 972 | -13.2794 | 4.93516 | 3.92762 |
| appF | CD630_n00510 | 43 – 86 | 172 – 216 | -13.2268 | 4.25570 | 3.92762 |
| CD630_03920 | CD630_n00510 | 127 – 168 | 646 – 690 | -13.2196 | 4.15934 | 3.92762 |
| CD630_21620 | CD630_n00510 | 137 – 184 | 468 – 513 | -13.2071 | 3.02649 | 3.92762 |
| fliI | CD630_n00510 | 82 – 129 | 529 – 591 | -13.1066 | 3.03666 | 3.92762 |
| CD630_15360 | CD630_n00510 | 114 – 156 | 213 – 251 | -13.0757 | 4.47208 | 3.92762 |
| CD630_04320 | CD630_n00510 | 78 – 108 | 734 – 763 | -12.9831 | 3.68342 | 3.92762 |
| CD630_28290 | CD630_n00510 | 65 – 129 | 451 – 511 | -12.9777 | 5.54192 | 3.92762 |
| CD630_08370 | CD630_n00510 | 185 – 196 | 630 – 641 | -12.9244 | 3.65639 | 3.92762 |
| CD630_19850 | CD630_n00510 | 106 – 119 | 869 – 882 | -12.8639 | 6.88486 | 3.92762 |
| CD630_22030 | CD630_n00510 | 176 – 199 | 695 – 717 | -12.8339 | 3.29934 | 3.92762 |
| CD630_14110 | CD630_n00510 | 1 – 29 | 318 – 350 | -12.7563 | 3.39550 | 3.92762 |
| CD630_13620 | CD630_n00510 | 158 – 199 | 123 – 160 | -12.7402 | 4.77627 | 3.92762 |
| CD630_15900 | CD630_n00510 | 147 – 189 | 357 – 405 | -12.5765 | 3.17664 | 3.92762 |
| garK | CD630_n00510 | 1 – 66 | 538 – 602 | -12.5415 | 3.72836 | 3.92762 |
| CD630_27770 | CD630_n00510 | 154 – 180 | 848 – 873 | -12.5066 | 4.08588 | 3.92762 |
| CD630_02420 | CD630_n00510 | 168 – 186 | 734 – 755 | -12.4213 | 3.71048 | 3.92762 |
| dpaA | CD630_n00510 | 40 – 54 | 105 – 120 | -12.3924 | 4.65364 | 3.92762 |
| CD630_03410 | CD630_n00510 | 53 – 76 | 323 – 346 | -12.3309 | 4.45994 | 3.92762 |
| CD630_22880 | CD630_n00510 | 39 – 85 | 851 – 899 | -12.2902 | 3.62455 | 3.92762 |
| CD630_15850 | CD630_n00510 | 156 – 197 | 846 – 883 | -12.2258 | 4.32968 | 3.92762 |
| CD630_16530 | CD630_n00510 | 150 – 168 | 266 – 286 | -12.2154 | 4.37658 | 3.92762 |
| CD630_32060 | CD630_n00510 | 38 – 60 | 224 – 246 | -12.2058 | 5.73797 | 3.92762 |
| CD630_05211 | CD630_n00510 | 127 – 170 | 833 – 882 | -12.2009 | 3.97048 | 3.92762 |
| CD630_02790 | CD630_n00510 | 170 – 195 | 114 – 140 | -12.1742 | 4.77661 | 3.92762 |
| CD630_09850 | CD630_n00510 | 45 – 99 | 199 – 245 | -12.1606 | 4.18679 | 3.92762 |
| CD630_04540 | CD630_n00510 | 103 – 142 | 5 – 43 | -12.1078 | 3.33556 | 3.92762 |
| CD630_27280 | CD630_n00510 | 96 – 154 | 633 – 686 | -12.0801 | 3.24738 | 3.92762 |
| CD630_17350 | CD630_n00510 | 51 – 93 | 212 – 251 | -12.0778 | 3.35334 | 3.92762 |
| CD630_21090 | CD630_n00510 | 156 – 189 | 936 – 966 | -12.0697 | 4.15803 | 3.92762 |
| CD630_13400 | CD630_n00510 | 132 – 153 | 587 – 610 | -12.0511 | 3.20232 | 3.92762 |
| CD630_31910 | CD630_n00510 | 132 – 184 | 569 – 636 | -12.0333 | 3.48314 | 3.92762 |
| CD630_22850 | CD630_n00510 | 172 – 190 | 892 – 909 | -12.0301 | 3.84102 | 3.92762 |
| CD630_14710 | CD630_n00510 | 5 – 102 | 832 – 944 | -12.0225 | 3.65171 | 3.92762 |
| CD630_13960 | CD630_n00510 | 1 – 42 | 644 – 690 | -11.9875 | 3.30375 | 3.92762 |
| CD630_27610 | CD630_n00510 | 15 – 34 | 236 – 255 | -11.9775 | 3.96465 | 3.92762 |
| CD630_18930 | CD630_n00510 | 2 – 80 | 301 – 378 | -11.937 | 5.32127 | 3.92762 |
| htpG | CD630_n00510 | 23 – 56 | 651 – 682 | -11.9081 | 3.45147 | 3.92762 |
| hfq | CD630_n00510 | 164 – 199 | 298 – 333 | -11.8982 | 5.05609 | 3.92762 |
| spoVFB | CD630_n00510 | 14 – 24 | 814 – 824 | -11.895 | 4.62758 | 3.92762 |
| CD630_21140 | CD630_n00510 | 126 – 173 | 226 – 274 | -11.8547 | 3.33304 | 3.92762 |
| scpB | CD630_n00510 | 134 – 160 | 105 – 127 | -11.8374 | 3.33560 | 3.92762 |
| CD630_25020 | CD630_n00510 | 127 – 158 | 875 – 906 | -11.8061 | 4.17721 | 3.92762 |
| CD630_17970 | CD630_n00510 | 155 – 177 | 752 – 776 | -11.7978 | 3.68802 | 3.92762 |
| CD630_24590 | CD630_n00510 | 186 – 201 | 231 – 246 | -11.7483 | 3.70401 | 3.92762 |
| CD630_19801 | CD630_n00510 | 113 – 132 | 652 – 674 | -11.62 | 5.22035 | 3.92762 |
| CD630_33600 | CD630_n00510 | 42 – 67 | 6 – 31 | -11.5936 | 3.73196 | 3.92762 |
| CD630_13950 | CD630_n00510 | 11 – 115 | 299 – 404 | -11.5917 | 4.25540 | 3.92762 |
| CD630_09772 | CD630_n00510 | 16 – 106 | 519 – 607 | -11.5692 | 3.66033 | 3.92762 |
| CD630_33190 | CD630_n00510 | 37 – 111 | 867 – 943 | -11.5633 | 4.13899 | 3.92762 |
| CD630_21260 | CD630_n00510 | 121 – 170 | 787 – 833 | -11.5576 | 4.22255 | 3.92762 |
| CD630_05830 | CD630_n00510 | 117 – 164 | 785 – 824 | -11.5511 | 3.35360 | 3.92762 |
| CD630_26050 | CD630_n00510 | 49 – 67 | 227 – 245 | -11.5217 | 6.99851 | 3.92762 |
| CD630_20670 | CD630_n00510 | 72 – 110 | 271 – 312 | -11.5073 | 4.42757 | 3.92762 |
| CD630_17900 | CD630_n00510 | 99 – 195 | 810 – 887 | -11.4874 | 3.52167 | 3.92762 |
| CD630_16950 | CD630_n00510 | 182 – 201 | 95 – 115 | -11.479 | 4.42569 | 3.92762 |
| CD630_32880 | CD630_n00510 | 142 – 180 | 560 – 596 | -11.471 | 3.41276 | 3.92762 |
| CD630_33420 | CD630_n00510 | 141 – 169 | 847 – 880 | -11.4684 | 3.22095 | 3.92762 |
| CD630_30120 | CD630_n00510 | 141 – 181 | 447 – 490 | -11.4657 | 4.04859 | 3.92762 |
| srlEb | CD630_n00510 | 59 – 94 | 213 – 246 | -11.4235 | 3.50140 | 3.92762 |
| thiH | CD630_n00510 | 183 – 201 | 757 – 776 | -11.4184 | 3.58230 | 3.92762 |
| nusB | CD630_n00510 | 130 – 197 | 558 – 630 | -11.4039 | 4.92798 | 3.92762 |
| CD630_23780 | CD630_n00510 | 45 – 104 | 338 – 397 | -11.3786 | 3.91629 | 3.92762 |
| CD630_05230 | CD630_n00510 | 38 – 59 | 752 – 768 | -11.3095 | 3.73787 | 3.92762 |
| CD630_22950 | CD630_n00510 | 3 – 27 | 17 – 42 | -11.3092 | 3.96883 | 3.92762 |
| virS | CD630_n00510 | 115 – 140 | 11 – 41 | -11.2851 | 4.74085 | 3.92762 |
| CD630_18970 | CD630_n00510 | 123 – 144 | 226 – 250 | -11.2777 | 3.57243 | 3.92762 |
| nrdD | CD630_n00510 | 58 – 92 | 484 – 517 | -11.2775 | 3.28290 | 3.92762 |
| CD630_13380 | CD630_n00510 | 40 – 106 | 546 – 598 | -11.2727 | 4.74408 | 3.92762 |
| opuCA | CD630_n00510 | 5 – 38 | 840 – 870 | -11.2671 | 4.07035 | 3.92762 |
| CD630_32110 | CD630_n00510 | 133 – 171 | 211 – 250 | -11.2548 | 4.45291 | 3.92762 |
| CD630_31381 | CD630_n00510 | 2 – 27 | 453 – 483 | -11.2456 | 3.49137 | 3.92762 |
| xylR | CD630_n00510 | 167 – 182 | 803 – 817 | -11.2404 | 3.95671 | 3.92762 |
| CD630_16460 | CD630_n00510 | 95 – 147 | 300 – 352 | -11.2247 | 4.64028 | 3.92762 |
| CD630_27200 | CD630_n00510 | 122 – 140 | 449 – 467 | -11.2223 | 3.42516 | 3.92762 |
| terD | CD630_n00510 | 182 – 194 | 752 – 764 | -11.2143 | 3.33148 | 3.92762 |
| CD630_08950 | CD630_n00510 | 167 – 201 | 8 – 40 | -11.1899 | 3.71905 | 3.92762 |
| CD630_05300 | CD630_n00510 | 108 – 159 | 812 – 847 | -11.1795 | 4.56885 | 3.92762 |
| gcp | CD630_n00510 | 20 – 57 | 531 – 577 | -11.1286 | 4.92798 | 3.92762 |
| CD630_17710 | CD630_n00510 | 16 – 81 | 193 – 248 | -11.1253 | 4.37688 | 3.92762 |
| CD630_03400 | CD630_n00510 | 109 – 140 | 161 – 195 | -11.1179 | 4.45994 | 3.92762 |
| CD630_22720 | CD630_n00510 | 140 – 164 | 756 – 779 | -11.1129 | 3.81804 | 3.92762 |
| CD630_06960 | CD630_n00510 | 147 – 182 | 11 – 43 | -11.1117 | 3.16317 | 3.92762 |
| CD630_18960 | CD630_n00510 | 155 – 192 | 930 – 964 | -11.1013 | 4.37822 | 3.92762 |
| CD630_09990 | CD630_n00510 | 143 – 201 | 365 – 419 | -11.0938 | 3.86040 | 3.92762 |
| CD630_31690 | CD630_n00510 | 116 – 162 | 315 – 365 | -11.0851 | 3.71193 | 3.92762 |
| CD630_20171 | CD630_n00510 | 76 – 156 | 212 – 304 | -11.0811 | 6.87491 | 3.92762 |
| CD630_14130 | CD630_n00510 | 99 – 201 | 667 – 764 | -11.0532 | 4.57970 | 3.92762 |
| blaI | CD630_n00510 | 26 – 46 | 665 – 687 | -11.0405 | 3.73674 | 3.92762 |
| CD630_34030 | CD630_n00510 | 166 – 198 | 4 – 31 | -11.0194 | 3.05766 | 3.92762 |
| CD630_23450 | CD630_n00510 | 36 – 47 | 553 – 564 | -11.004 | 3.27218 | 3.92762 |
| panB | CD630_n00510 | 108 – 179 | 851 – 923 | -10.9995 | 3.40248 | 3.92762 |
| CD630_25300 | CD630_n00510 | 129 – 142 | 261 – 274 | -10.9964 | 4.49333 | 3.92762 |
| CD630_13780 | CD630_n00510 | 17 – 58 | 659 – 696 | -10.9811 | 3.52274 | 3.92762 |
| CD630_14930 | CD630_n00510 | 5 – 25 | 385 – 411 | -10.9777 | 3.31335 | 3.92762 |
| oppB | CD630_n00510 | 87 – 179 | 813 – 896 | -10.969 | 2.98328 | 3.92762 |
| CD630_05490 | CD630_n00510 | 90 – 103 | 881 – 894 | -10.947 | 3.89852 | 3.92762 |
| CD630_06380 | CD630_n00510 | 147 – 189 | 786 – 822 | -10.9159 | 4.32211 | 3.92762 |
| CD630_08080 | CD630_n00510 | 2 – 21 | 321 – 342 | -10.9032 | 4.87819 | 3.92762 |
| CD630_05840 | CD630_n00510 | 124 – 161 | 357 – 392 | -10.8635 | 4.28779 | 3.92762 |
| spaK | CD630_n00510 | 174 – 200 | 123 – 148 | -10.8572 | 3.04530 | 3.92762 |
| bioB | CD630_n00510 | 115 – 149 | 934 – 961 | -10.8297 | 3.59022 | 3.92762 |
| CD630_21770 | CD630_n00510 | 1 – 13 | 4 – 16 | -10.8107 | 4.02587 | 3.92762 |
| CD630_07400 | CD630_n00510 | 168 – 187 | 385 – 404 | -10.7992 | 4.52273 | 3.92762 |
| CD630_06340 | CD630_n00510 | 104 – 190 | 886 – 976 | -10.7934 | 4.35983 | 3.92762 |
| CD630_33570 | CD630_n00510 | 147 – 174 | 755 – 781 | -10.7898 | 3.95778 | 3.92762 |
| CD630_20600 | CD630_n00510 | 36 – 56 | 118 – 140 | -10.7721 | 4.20093 | 3.92762 |
| CD630_01700 | CD630_n00510 | 81 – 93 | 481 – 494 | -10.7678 | 4.97234 | 3.92762 |
| CD630_09970 | CD630_n00510 | 167 – 183 | 122 – 138 | -10.762 | 3.25917 | 3.92762 |
| CD630_10830 | CD630_n00510 | 173 – 200 | 226 – 252 | -10.7507 | 4.82398 | 3.92762 |
| CD630_04620 | CD630_n00510 | 20 – 67 | 907 – 953 | -10.7439 | 3.45413 | 3.92762 |
| CD630_06050 | CD630_n00510 | 111 – 166 | 227 – 291 | -10.7432 | 3.59332 | 3.92762 |
| CD630_31880 | CD630_n00510 | 2 – 23 | 211 – 233 | -10.7241 | 3.97522 | 3.92762 |
| CD630_03180 | CD630_n00510 | 42 – 50 | 601 – 609 | -10.7162 | 3.73138 | 3.92762 |
| CD630_23820 | CD630_n00510 | 83 – 107 | 749 – 772 | -10.7058 | 3.18419 | 3.92762 |
| CD630_25620 | CD630_n00510 | 91 – 107 | 99 – 115 | -10.6974 | 3.02685 | 3.92762 |
| CD630_21760 | CD630_n00510 | 125 – 157 | 236 – 270 | -10.6958 | 4.43846 | 3.92762 |
| CD630_15890 | CD630_n00510 | 121 – 169 | 882 – 943 | -10.6669 | 3.39962 | 3.92762 |
| dltB | CD630_n00510 | 135 – 177 | 215 – 264 | -10.6267 | 3.81485 | 3.92762 |
| CD630_34520 | CD630_n00510 | 115 – 133 | 496 – 514 | -10.6262 | 3.83016 | 3.92762 |
| CD630_36781 | CD630_n00510 | 45 – 91 | 351 – 399 | -10.6165 | 3.75946 | 3.92762 |
| CD630_21110 | CD630_n00510 | 44 – 114 | 890 – 960 | -10.6147 | 3.89630 | 3.92762 |
| CD630_30420 | CD630_n00510 | 56 – 201 | 228 – 358 | -10.5919 | 5.27539 | 3.92762 |
| CD630_34530 | CD630_n00510 | 1 – 71 | 520 – 577 | -10.5915 | 3.83016 | 3.92762 |
| CD630_25190 | CD630_n00510 | 70 – 100 | 477 – 511 | -10.5683 | 3.19663 | 3.92762 |

| | | | | | | |
|---|---|---|---|---|---|---|
| CD630_14600 | CD630_n00510 | 66 -- 81 | 451 -- 466 | -10.5644 | 5.90348 | 3.92762 |
| CD630_18910 | CD630_n00510 | 14 -- 83 | 768 -- 833 | -10.5627 | 3.30688 | 3.92762 |
| CD630_31530 | CD630_n00510 | 7 -- 30 | 124 -- 148 | -10.55 | 3.49100 | 3.92762 |
| CD630_16450 | CD630_n00510 | 3 -- 30 | 932 -- 960 | -10.5089 | 4.57547 | 3.92762 |
| CD630_18940 | CD630_n00510 | 171 -- 185 | 552 -- 566 | -10.5052 | 4.09988 | 3.92762 |
| CD630_14210 | CD630_n00510 | 14 -- 28 | 792 -- 806 | -10.4964 | 3.82355 | 3.92762 |
| CD630_25050 | CD630_n00510 | 101 -- 133 | 4 -- 33 | -10.4892 | 4.33624 | 3.92762 |
| CD630_19700 | CD630_n00510 | 151 -- 168 | 810 -- 825 | -10.485 | 3.85861 | 3.92762 |
| sigK | CD630_n00510 | 9 -- 114 | 734 -- 840 | -10.4761 | 3.08122 | 3.92762 |
| CD630_21540 | CD630_n00510 | 57 -- 82 | 484 -- 508 | -10.4598 | 4.33090 | 3.92762 |
| CD630_29750 | CD630_n00510 | 25 -- 36 | 267 -- 278 | -10.4591 | 4.59146 | 3.92762 |
| rolA | CD630_n00510 | 152 -- 195 | 139 -- 179 | -10.4553 | 4.56362 | 3.92762 |
| CD630_05000 | CD630_n00510 | 124 -- 174 | 868 -- 922 | -10.4368 | 4.02442 | 3.92762 |
| CD630_14610 | CD630_n00510 | 173 -- 184 | 31 -- 42 | -10.4291 | 4.89428 | 3.92762 |
| CD630_19430 | CD630_n00510 | 175 -- 197 | 226 -- 247 | -10.378 | 3.13922 | 3.92762 |
| CD630_16640 | CD630_n00510 | 92 -- 107 | 99 -- 115 | -10.3553 | 3.61548 | 3.92762 |
| CD630_25610 | CD630_n00510 | 12 -- 41 | 557 -- 585 | -10.3514 | 3.02685 | 3.92762 |
| CD630_13530 | CD630_n00510 | 160 -- 187 | 934 -- 961 | -10.3493 | 4.09832 | 3.92762 |
| CD630_15060 | CD630_n00510 | 1 -- 22 | 359 -- 380 | -10.3432 | 3.69396 | 3.92762 |
| CD630_25310 | CD630_n00510 | 21 -- 33 | 871 -- 883 | -10.341 | 3.99861 | 3.92762 |
| CD630_15200 | CD630_n00510 | 73 -- 112 | 519 -- 564 | -10.3279 | 4.00442 | 3.92762 |
| CD630_10050 | CD630_n00510 | 57 -- 94 | 213 -- 246 | -10.3174 | 3.93750 | 3.92762 |
| CD630_23000 | CD630_n00510 | 90 -- 114 | 667 -- 692 | -10.3064 | 3.54130 | 3.92762 |
| CD630_35260 | CD630_n00510 | 58 -- 105 | 606 -- 646 | -10.2706 | 5.77277 | 3.92762 |
| CD630_28260 | CD630_n00510 | 70 -- 105 | 753 -- 779 | -10.2583 | 3.51613 | 3.92762 |
| CD630_03930 | CD630_n00510 | 135 -- 171 | 786 -- 823 | -10.2569 | 4.15934 | 3.92762 |
| CD630_21030 | CD630_n00510 | 169 -- 199 | 418 -- 457 | -10.2189 | 3.08733 | 3.92762 |
| CD630_14620 | CD630_n00510 | 103 -- 146 | 656 -- 696 | -10.2142 | 4.43651 | 3.92762 |
| CD630_15740 | CD630_n00510 | 7 -- 92 | 873 -- 958 | -10.209 | 4.43222 | 3.92762 |
| CD630_07960 | CD630_n00510 | 1 -- 55 | 530 -- 578 | -10.1851 | 3.22537 | 3.92762 |
| CD630_26150 | CD630_n00510 | 25 -- 56 | 16 -- 43 | -10.1805 | 4.06302 | 3.92762 |
| iorA | CD630_n00510 | 55 -- 69 | 102 -- 117 | -10.1722 | 3.83173 | 3.92762 |
| CD630_13790 | CD630_n00510 | 157 -- 174 | 550 -- 568 | -10.1573 | 5.17374 | 3.92762 |
| CD630_13020 | CD630_n00510 | 123 -- 150 | 699 -- 725 | -10.1452 | 3.52267 | 3.92762 |
| CD630_14340 | CD630_n00510 | 33 -- 108 | 205 -- 270 | -10.1398 | 5.10868 | 3.92762 |
| CD630_03040 | CD630_n00510 | 10 -- 34 | 484 -- 506 | -10.135 | 3.51403 | 3.92762 |
| CD630_17720 | CD630_n00510 | 3 -- 43 | 793 -- 830 | -10.1294 | 3.22910 | 3.92762 |
| oppC | CD630_n00510 | 66 -- 80 | 160 -- 174 | -10.1174 | 3.01695 | 3.92762 |
| fbp | CD630_n00510 | 159 -- 186 | 223 -- 250 | -10.1114 | 3.80071 | 3.92762 |
| CD630_28330 | CD630_n00510 | 14 -- 30 | 53 -- 69 | -10.1048 | 3.29122 | 3.92762 |
| vncR | CD630_n00510 | 127 -- 175 | 734 -- 780 | -10.0958 | 3.93642 | 3.92762 |
| CD630_29960 | CD630_n00510 | 47 -- 67 | 653 -- 673 | -10.0838 | 4.51904 | 3.92762 |
| CD630_15111 | CD630_n00510 | 20 -- 55 | 228 -- 260 | -10.0531 | 4.18664 | 3.92762 |
| CD630_04940 | CD630_n00510 | 163 -- 174 | 269 -- 280 | -10.0524 | 4.20948 | 3.92762 |
| CD630_23090 | CD630_n00510 | 10 -- 32 | 880 -- 903 | -10.0487 | 4.25622 | 3.92762 |
| CD630_29840 | CD630_n00510 | 123 -- 194 | 13 -- 69 | -10.0299 | 4.48825 | 3.92762 |
| CD630_06410 | CD630_n00510 | 152 -- 181 | 934 -- 971 | -10.0201 | 4.14404 | 3.92762 |
| CD630_16750 | CD630_n00510 | 177 -- 193 | 230 -- 246 | -10.0126 | 5.04051 | 3.92762 |
| chrA | CD630_n00510 | 68 -- 82 | 727 -- 741 | -10.0118 | 5.27255 | 3.92762 |
| CD630_21080 | CD630_n00510 | 159 -- 179 | 805 -- 825 | -10.0077 | 3.99742 | 3.92762 |
| CD630_26750 | CD630_n00510 | 104 -- 167 | 187 -- 246 | -10.0063 | 3.60610 | 3.92762 |
| CD630_06100 | CD630_n00510 | 135 -- 195 | 817 -- 869 | -9.98549 | 4.43918 | 3.92762 |
| CD630_17770 | CD630_n00510 | 122 -- 149 | 6 -- 30 | -9.97893 | 5.06490 | 3.92762 |
| CD630_14010 | CD630_n00510 | 32 -- 51 | 942 -- 961 | -9.95225 | 3.69983 | 3.92762 |
| CD630_27110 | CD630_n00510 | 73 -- 118 | 849 -- 905 | -9.9515 | 3.78231 | 3.92762 |
| CD630_17511 | CD630_n00510 | 4 -- 59 | 218 -- 268 | -9.9294 | 9.12322 | 3.92762 |
| CD630_32890 | CD630_n00510 | 101 -- 157 | 699 -- 755 | -9.92761 | 3.41276 | 3.92762 |
| CD630_23861 | CD630_n00510 | 7 -- 38 | 517 -- 548 | -9.9131 | 5.83068 | 3.92762 |
| CD630_04550 | CD630_n00510 | 181 -- 199 | 331 -- 352 | -9.89939 | 3.05227 | 3.92762 |
| CD630_13360 | CD630_n00510 | 176 -- 195 | 212 -- 229 | -9.89206 | 3.77330 | 3.92762 |
| CD630_32710 | CD630_n00510 | 119 -- 169 | 723 -- 774 | -9.88092 | 4.74276 | 3.92762 |
| CD630_32500 | CD630_n00510 | 28 -- 64 | 373 -- 408 | -9.86911 | 6.17288 | 3.92762 |
| CD630_02720 | CD630_n00510 | 124 -- 172 | 631 -- 675 | -9.82248 | 3.38938 | 3.92762 |
| CD630_08660 | CD630_n00510 | 48 -- 86 | 887 -- 937 | -9.80357 | 3.86467 | 3.92762 |
| CD630_24760 | CD630_n00510 | 100 -- 122 | 875 -- 897 | -9.80353 | 5.19485 | 3.92762 |
| CD630_36690 | CD630_n00510 | 24 -- 50 | 16 -- 40 | -9.78582 | 3.53761 | 3.92762 |
| eutV | CD630_n00510 | 113 -- 173 | 749 -- 792 | -9.77766 | 4.55715 | 3.92762 |
| uxaA%27 | CD630_n00510 | 119 -- 170 | 48 -- 96 | -9.77205 | 4.26165 | 3.92762 |
| CD630_21230 | CD630_n00510 | 91 -- 104 | 102 -- 115 | -9.76513 | 3.89213 | 3.92762 |
| CD630_24050 | CD630_n00510 | 151 -- 174 | 538 -- 561 | -9.76174 | 3.70192 | 3.92762 |
| CD630_14150 | CD630_n00510 | 47 -- 168 | 350 -- 465 | -9.74565 | 3.28251 | 3.92762 |
| CD630_19780 | CD630_n00510 | 96 -- 114 | 136 -- 156 | -9.74465 | 3.64830 | 3.92762 |
| CD630_11240 | CD630_n00510 | 88 -- 107 | 99 -- 115 | -9.72821 | 3.20274 | 3.92762 |
| CD630_11730 | CD630_n00510 | 147 -- 198 | 431 -- 476 | -9.72326 | 4.80521 | 3.92762 |
| CD630_09750 | CD630_n00510 | 153 -- 200 | 249 -- 297 | -9.70604 | 4.83410 | 3.92762 |
| CD630_28920 | CD630_n00510 | 153 -- 200 | 249 -- 297 | -9.70604 | 4.51664 | 3.92762 |
| CD630_34080 | CD630_n00510 | 148 -- 160 | 629 -- 641 | -9.68949 | 3.20224 | 3.92762 |
| CD630_20650 | CD630_n00510 | 78 -- 122 | 452 -- 494 | -9.68455 | 4.00083 | 3.92762 |
| CD630_08070 | CD630_n00510 | 184 -- 200 | 528 -- 544 | -9.68445 | 4.87819 | 3.92762 |
| CD630_27310 | CD630_n00510 | 25 -- 80 | 213 -- 256 | -9.66242 | 4.01622 | 3.92762 |
| CD630_32070 | CD630_n00510 | 38 -- 74 | 873 -- 912 | -9.66158 | 3.51964 | 3.92762 |
| CD630_21070 | CD630_n00510 | 178 -- 188 | 226 -- 236 | -9.65759 | 5.35019 | 3.92762 |
| CD630_23770 | CD630_n00510 | 1 -- 20 | 227 -- 246 | -9.64446 | 3.25181 | 3.92762 |
| CD630_25320 | CD630_n00510 | 90 -- 130 | 873 -- 914 | -9.63967 | 4.29415 | 3.92762 |
| CD630_04660 | CD630_n00510 | 162 -- 200 | 929 -- 965 | -9.63792 | 4.32791 | 3.92762 |
| CD630_22740 | CD630_n00510 | 88 -- 114 | 748 -- 776 | -9.63422 | 3.08733 | 3.92762 |
| CD630_22500 | CD630_n00510 | 124 -- 143 | 11 -- 28 | -9.62523 | 4.03688 | 3.92762 |
| CD630_26020 | CD630_n00510 | 125 -- 156 | 222 -- 248 | -9.60668 | 4.03155 | 3.92762 |
| CD630_16520 | CD630_n00510 | 1 -- 69 | 327 -- 387 | -9.59456 | 4.28395 | 3.92762 |
| CD630_17281 | CD630_n00510 | 5 -- 38 | 851 -- 883 | -9.58768 | 5.91544 | 3.92762 |
| CD630_20260 | CD630_n00510 | 7 -- 72 | 391 -- 458 | -9.58683 | 3.84162 | 3.92762 |
| CD630_09770 | CD630_n00510 | 6 -- 35 | 872 -- 914 | -9.57736 | 3.22106 | 3.92762 |
| dapB2 | CD630_n00510 | 186 -- 196 | 276 -- 286 | -9.56194 | 3.27218 | 3.92762 |
| CD630_26250 | CD630_n00510 | 144 -- 175 | 321 -- 355 | -9.55909 | 5.03379 | 3.92762 |
| CD630_08100 | CD630_n00510 | 126 -- 134 | 483 -- 491 | -9.54446 | 4.39181 | 3.92762 |
| rbsC | CD630_n00510 | 8 -- 44 | 737 -- 784 | -9.54221 | 3.71650 | 3.92762 |
| cdtR | CD630_n00510 | 95 -- 149 | 397 -- 454 | -9.51888 | 5.84368 | 3.92762 |
| etfA | CD630_n00560 | 49 -- 94 | 658 -- 698 | -18.9596 | 4.80521 | 5.82053 |
| CD630_19680 | CD630_n00560 | 5 -- 152 | 97 -- 235 | -18.5052 | 3.58080 | 5.82053 |
| dapB2 | CD630_n00560 | 149 -- 200 | 445 -- 496 | -17.9233 | 3.27218 | 5.82053 |
| thiH | CD630_n00560 | 176 -- 196 | 526 -- 546 | -17.1505 | 3.58230 | 5.82053 |
| CD630_18910 | CD630_n00560 | 163 -- 197 | 442 -- 476 | -16.8306 | 3.30688 | 5.82053 |
| CD630_10880 | CD630_n00560 | 147 -- 182 | 671 -- 703 | -16.6436 | 5.05048 | 5.82053 |
| CD630_15850 | CD630_n00560 | 96 -- 128 | 674 -- 708 | -16.099 | 4.32968 | 5.82053 |
| CD630_11240 | CD630_n00560 | 148 -- 168 | 670 -- 690 | -16.0602 | 3.20274 | 5.82053 |
| bglA | CD630_n00560 | 75 -- 117 | 462 -- 492 | -15.6368 | 4.50315 | 5.82053 |
| CD630_04660 | CD630_n00560 | 62 -- 96 | 461 -- 493 | -15.596 | 4.32791 | 5.82053 |
| CD630_22880 | CD630_n00560 | 74 -- 116 | 657 -- 705 | -15.5288 | 3.62455 | 5.82053 |
| CD630_27210 | CD630_n00560 | 29 -- 50 | 462 -- 483 | -15.3983 | 4.18786 | 5.82053 |
| cdu2 | CD630_n00560 | 65 -- 89 | 460 -- 482 | -15.2855 | 2.97335 | 5.82053 |
| malY | CD630_n00560 | 158 -- 191 | 456 -- 487 | -15.2016 | 4.46345 | 5.82053 |
| CD630_13350 | CD630_n00560 | 85 -- 200 | 541 -- 667 | -15.1749 | 4.15139 | 5.82053 |
| CD630_03920 | CD630_n00560 | 151 -- 201 | 673 -- 722 | -15.1349 | 4.15934 | 5.82053 |
| CD630_32690 | CD630_n00560 | 47 -- 186 | 462 -- 588 | -14.9924 | 7.32676 | 5.82053 |
| CD630_25190 | CD630_n00560 | 9 -- 62 | 663 -- 707 | -14.9918 | 3.19663 | 5.82053 |
| CD630_28260 | CD630_n00560 | 37 -- 85 | 427 -- 478 | -14.7079 | 3.51613 | 5.82053 |
| CD630_31910 | CD630_n00560 | 115 -- 193 | 159 -- 230 | -14.6319 | 3.48314 | 5.82053 |
| CD630_13620 | CD630_n00560 | 142 -- 199 | 122 -- 170 | -14.4699 | 4.77627 | 5.82053 |
| CD630_19790 | CD630_n00560 | 49 -- 104 | 659 -- 710 | -14.382 | 3.79593 | 5.82053 |
| serS1 | CD630_n00560 | 141 -- 160 | 464 -- 483 | -14.3457 | 3.45858 | 5.82053 |
| CD630_32880 | CD630_n00560 | 91 -- 196 | 107 -- 203 | -14.1622 | 3.41276 | 5.82053 |
| CD630_27060 | CD630_n00560 | 14 -- 152 | 71 -- 211 | -14.1458 | 4.06049 | 5.82053 |
| flgB | CD630_n00560 | 81 -- 194 | 164 -- 289 | -14.0787 | 3.25736 | 5.82053 |
| CD630_19903 | CD630_n00560 | 166 -- 201 | 332 -- 364 | -14.0617 | 5.79994 | 5.82053 |
| CD630_32710 | CD630_n00560 | 132 -- 172 | 448 -- 486 | -13.9181 | 4.74276 | 5.82053 |
| CD630_03170 | CD630_n00560 | 2 -- 16 | 689 -- 703 | -13.8901 | 3.66946 | 5.82053 |
| CD630_07470 | CD630_n00560 | 9 -- 66 | 141 -- 217 | -13.8402 | 3.37143 | 5.82053 |
| CD630_06060 | CD630_n00560 | 72 -- 94 | 670 -- 692 | -13.8136 | 4.94050 | 5.82053 |
| CD630_08370 | CD630_n00560 | 122 -- 139 | 465 -- 482 | -13.6832 | 3.65639 | 5.82053 |
| folC | CD630_n00560 | 110 -- 124 | 467 -- 481 | -13.6418 | 3.85634 | 5.82053 |
| CD630_27600 | CD630_n00560 | 92 -- 131 | 455 -- 494 | -13.5879 | 3.48384 | 5.82053 |
| CD630_29270 | CD630_n00560 | 33 -- 57 | 462 -- 483 | -13.3627 | 3.70174 | 5.82053 |
| CD630_27570 | CD630_n00560 | 97 -- 128 | 674 -- 707 | -13.2844 | 3.46216 | 5.82053 |
| iorA | CD630_n00560 | 2 -- 23 | 680 -- 701 | -13.2828 | 3.83173 | 5.82053 |
| CD630_04341 | CD630_n00560 | 156 -- 192 | 446 -- 483 | -13.2651 | 4.29262 | 5.82053 |
| CD630_15461 | CD630_n00560 | 46 -- 189 | 271 -- 415 | -13.2316 | 9.13442 | 5.82053 |
| ung | CD630_n00560 | 100 -- 156 | 462 -- 514 | -13.1864 | 3.30908 | 5.82053 |
| CD630_21770 | CD630_n00560 | 143 -- 162 | 537 -- 558 | -13.1675 | 4.02587 | 5.82053 |
| fbp | CD630_n00560 | 159 -- 195 | 280 -- 314 | -13.1287 | 3.80071 | 5.82053 |
| CD630_10050 | CD630_n00560 | 160 -- 201 | 676 -- 710 | -13.1239 | 3.93750 | 5.82053 |
| CD630_01700 | CD630_n00560 | 1 -- 45 | 123 -- 169 | -13.0994 | 4.97234 | 5.82053 |
| CD630_22550 | CD630_n00560 | 28 -- 60 | 137 -- 171 | -13.0638 | 3.62379 | 5.82053 |
| CD630_07400 | CD630_n00560 | 36 -- 72 | 115 -- 150 | -13.0412 | 4.52273 | 5.82053 |
| eutW | CD630_n00560 | 10 -- 20 | 673 -- 683 | -13.0231 | 4.55715 | 5.82053 |
| CD630_17151 | CD630_n00560 | 155 -- 191 | 455 -- 491 | -13.0149 | 6.43124 | 5.82053 |
| CD630_19904 | CD630_n00560 | 148 -- 195 | 648 -- 696 | -13.0094 | 5.96526 | 5.82053 |
| CD630_05230 | CD630_n00560 | 6 -- 22 | 665 -- 682 | -12.8745 | 3.73787 | 5.82053 |
| cspD | CD630_n00560 | 116 -- 155 | 322 -- 373 | -12.8642 | 3.23587 | 5.82053 |
| CD630_21610 | CD630_n00560 | 52 -- 88 | 406 -- 441 | -12.8004 | 3.16662 | 5.82053 |
| CD630_28181 | CD630_n00560 | 29 -- 77 | 439 -- 477 | -12.7233 | 3.91299 | 5.82053 |
| CD630_18680 | CD630_n00560 | 135 -- 193 | 460 -- 513 | -12.7108 | 3.04315 | 5.82053 |
| CD630_04850 | CD630_n00560 | 155 -- 189 | 119 -- 148 | -12.7091 | 4.40788 | 5.82053 |
| CD630_32050 | CD630_n00560 | 153 -- 195 | 670 -- 701 | -12.7071 | 5.73797 | 5.82053 |
| cbpA | CD630_n00560 | 181 -- 201 | 25 -- 45 | -12.6994 | 3.03040 | 5.82053 |
| appC | CD630_n00560 | 63 -- 76 | 159 -- 172 | -12.6422 | 4.50438 | 5.82053 |
| panB | CD630_n00560 | 129 -- 180 | 123 -- 171 | -12.6188 | 3.40248 | 5.82053 |
| def1 | CD630_n00560 | 76 -- 190 | 461 -- 558 | -12.6183 | 3.99720 | 5.82053 |
| CD630_08390 | CD630_n00560 | 79 -- 113 | 668 -- 698 | -12.5211 | 5.16000 | 5.82053 |
| argS | CD630_n00560 | 12 -- 61 | 673 -- 703 | -12.499 | 4.96036 | 5.82053 |
| CD630_32700 | CD630_n00560 | 169 -- 184 | 461 -- 477 | -12.4506 | 4.74276 | 5.82053 |
| CD630_21260 | CD630_n00560 | 131 -- 189 | 19 -- 72 | -12.4391 | 4.22255 | 5.82053 |
| mobA | CD630_n00560 | 121 -- 153 | 673 -- 705 | -12.4386 | 4.25703 | 5.82053 |
| CD630_03220 | CD630_n00560 | 158 -- 171 | 462 -- 475 | -12.4284 | 3.36746 | 5.82053 |
| kdpE | CD630_n00560 | 164 -- 199 | 532 -- 558 | -12.4075 | 4.27140 | 5.82053 |
| CD630_20960 | CD630_n00560 | 170 -- 185 | 267 -- 283 | -12.4061 | 5.65940 | 5.82053 |
| CD630_32070 | CD630_n00560 | 31 -- 75 | 461 -- 496 | -12.3985 | 3.51964 | 5.82053 |
| CD630_32520 | CD630_n00560 | 154 -- 198 | 657 -- 710 | -12.3953 | 4.06098 | 5.82053 |
| CD630_18100 | CD630_n00560 | 120 -- 146 | 673 -- 697 | -12.3809 | 5.71658 | 5.82053 |
| moaA | CD630_n00560 | 10 -- 25 | 671 -- 686 | -12.3439 | 3.83799 | 5.82053 |
| spoVFB | CD630_n00560 | 154 -- 201 | 673 -- 708 | -12.2402 | 4.62758 | 5.82053 |
| CD630_21750 | CD630_n00560 | 108 -- 199 | 146 -- 247 | -12.2285 | 3.70626 | 5.82053 |
| CD630_23090 | CD630_n00560 | 166 -- 201 | 332 -- 364 | -12.2207 | 4.25622 | 5.82053 |
| CD630_06340 | CD630_n00560 | 74 -- 106 | 525 -- 556 | -12.2045 | 4.35983 | 5.82053 |
| CD630_14360 | CD630_n00560 | 59 -- 116 | 266 -- 316 | -12.192 | 3.02079 | 5.82053 |
| CD630_24590 | CD630_n00560 | 131 -- 160 | 84 -- 139 | -12.1651 | 3.70401 | 5.82053 |
| CD630_19700 | CD630_n00560 | 69 -- 106 | 674 -- 712 | -12.1507 | 3.85861 | 5.82053 |
| CD630_27270 | CD630_n00560 | 141 -- 166 | 476 -- 503 | -12.1436 | 3.50892 | 5.82053 |
| CD630_17960 | CD630_n00560 | 182 -- 198 | 124 -- 139 | -12.1398 | 3.71404 | 5.82053 |
| tcdA | CD630_n00560 | 143 -- 160 | 461 -- 476 | -12.1218 | 4.70593 | 5.82053 |
| CD630_24580 | CD630_n00560 | 10 -- 54 | 100 -- 141 | -12.1158 | 3.88519 | 5.82053 |
| CD630_10000 | CD630_n00560 | 99 -- 107 | 673 -- 681 | -12.0955 | 3.86040 | 5.82053 |
| CD630_21570 | CD630_n00560 | 39 -- 65 | 694 -- 722 | -12.0637 | 6.48385 | 5.82053 |
| CD630_13530 | CD630_n00560 | 175 -- 186 | 393 -- 404 | -12.0463 | 4.09832 | 5.82053 |
| CD630_17170 | CD630_n00560 | 64 -- 109 | 654 -- 706 | -12.0427 | 4.10230 | 5.82053 |
| CD630_05150 | CD630_n00560 | 110 -- 169 | 112 -- 166 | -12.0125 | 3.69875 | 5.82053 |
| bioB | CD630_n00560 | 114 -- 148 | 393 -- 419 | -12.0069 | 3.59022 | 5.82053 |
| CD630_27230 | CD630_n00560 | 93 -- 150 | 661 -- 714 | -12.0019 | 3.52462 | 5.82053 |
| CD630_21240 | CD630_n00560 | 84 -- 113 | 395 -- 422 | -11.9854 | 5.88879 | 5.82053 |
| CD630_22740 | CD630_n00560 | 79 -- 109 | 522 -- 559 | -11.9735 | 3.08733 | 5.82053 |
| CD630_14010 | CD630_n00560 | 29 -- 90 | 282 -- 359 | -11.9244 | 3.69983 | 5.82053 |
| feoB | CD630_n00560 | 127 -- 176 | 125 -- 169 | -11.9141 | 4.52257 | 5.82053 |
| CD630_21090 | CD630_n00560 | 161 -- 181 | 110 -- 132 | -11.9135 | 4.15803 | 5.82053 |
| CD630_01570 | CD630_n00560 | 157 -- 197 | 179 -- 212 | -11.9102 | 6.08699 | 5.82053 |
| CD630_02420 | CD630_n00560 | 9 -- 95 | 461 -- 534 | -11.8816 | 3.71048 | 5.82053 |
| glgB | CD630_n00560 | 3 -- 24 | 211 -- 232 | -11.8604 | 4.48273 | 5.82053 |
| CD630_22850 | CD630_n00560 | 172 -- 198 | 679 -- 705 | -11.8339 | 3.84102 | 5.82053 |
| CD630_02790 | CD630_n00560 | 90 -- 115 | 467 -- 486 | -11.8021 | 4.77661 | 5.82053 |
| CD630_14630 | CD630_n00560 | 109 -- 167 | 659 -- 712 | -11.795 | 3.34485 | 5.82053 |
| CD630_13960 | CD630_n00560 | 121 -- 160 | 675 -- 707 | -11.7887 | 3.30375 | 5.82053 |
| CD630_25320 | CD630_n00560 | 127 -- 163 | 395 -- 437 | -11.7753 | 4.29415 | 5.82053 |
| cdtR | CD630_n00560 | 19 -- 43 | 576 -- 600 | -11.7657 | 5.84368 | 5.82053 |
| CD630_21760 | CD630_n00560 | 138 -- 164 | 466 -- 492 | -11.7437 | 4.43846 | 5.82053 |
| dnaC | CD630_n00560 | 122 -- 156 | 676 -- 701 | -11.7363 | 6.30796 | 5.82053 |
| CD630_07770 | CD630_n00560 | 106 -- 125 | 461 -- 482 | -11.7359 | 3.31158 | 5.82053 |
| CD630_13880 | CD630_n00560 | 63 -- 79 | 464 -- 482 | -11.6977 | 3.52735 | 5.82053 |
| oppC | CD630_n00560 | 67 -- 80 | 159 -- 172 | -11.6905 | 3.01695 | 5.82053 |
| CD630_06170 | CD630_n00560 | 8 -- 35 | 322 -- 350 | -11.6763 | 4.95366 | 5.82053 |
| rbsC | CD630_n00560 | 32 -- 52 | 674 -- 692 | -11.6631 | 3.71650 | 5.82053 |
| CD630_10960 | CD630_n00560 | 39 -- 60 | 474 -- 494 | -11.6569 | 3.53850 | 5.82053 |
| CD630_17292 | CD630_n00560 | 143 -- 173 | 674 -- 706 | -11.6568 | 6.25799 | 5.82053 |
| CD630_29960 | CD630_n00560 | 175 -- 201 | 459 -- 485 | -11.654 | 4.51904 | 5.82053 |
| CD630_20660 | CD630_n00560 | 6 -- 31 | 124 -- 147 | -11.6378 | 5.39321 | 5.82053 |
| CD630_29650 | CD630_n00560 | 163 -- 200 | 626 -- 666 | -11.6192 | 3.67128 | 5.82053 |
| CD630_31490 | CD630_n00560 | 102 -- 116 | 106 -- 121 | -11.6148 | 4.08593 | 5.82053 |
| CD630_08080 | CD630_n00560 | 30 -- 69 | 123 -- 171 | -11.6051 | 4.87819 | 5.82053 |
| terD | CD630_n00560 | 21 -- 33 | 127 -- 139 | -11.603 | 3.33148 | 5.82053 |
| etfA | CD630_n00560 | 158 -- 180 | 463 -- 484 | -11.5495 | 3.41204 | 5.82053 |
| CD630_15431 | CD630_n00560 | 166 -- 190 | 668 -- 693 | -11.5312 | 5.29780 | 5.82053 |
| adhE | CD630_n00560 | 164 -- 196 | 444 -- 481 | -11.4981 | 3.06485 | 5.82053 |
| CD630_20980 | CD630_n00560 | 67 -- 109 | 669 -- 704 | -11.4361 | 8.12847 | 5.82053 |
| CD630_19030 | CD630_n00560 | 3 -- 76 | 141 -- 202 | -11.4304 | 3.34240 | 5.82053 |
| CD630_33600 | CD630_n00560 | 147 -- 174 | 673 -- 701 | -11.4228 | 3.73196 | 5.82053 |
| CD630_14260 | CD630_n00560 | 129 -- 148 | 466 -- 486 | -11.4166 | 3.22214 | 5.82053 |
| CD630_06060 | CD630_n00560 | 22 -- 48 | 106 -- 135 | -11.4068 | 4.94050 | 5.82053 |
| CD630_31970 | CD630_n00560 | 111 -- 124 | 617 -- 630 | -11.4022 | 3.24511 | 5.82053 |
| CD630_14110 | CD630_n00560 | 5 -- 37 | 123 -- 160 | -11.39 | 3.39550 | 5.82053 |
| CD630_26050 | CD630_n00560 | 103 -- 154 | 425 -- 476 | -11.3387 | 6.99851 | 5.82053 |
| CD630_02830 | CD630_n00560 | 185 -- 199 | 462 -- 476 | -11.3243 | 3.57106 | 5.82053 |
| CD630_06540 | CD630_n00560 | 45 -- 117 | 215 -- 283 | -11.3215 | 3.54673 | 5.82053 |
| CD630_26750 | CD630_n00560 | 104 -- 149 | 668 -- 706 | -11.2951 | 3.60610 | 5.82053 |
| pilC | CD630_n00560 | 177 -- 186 | 670 -- 679 | -11.2644 | 3.54230 | 5.82053 |
| CD630_06100 | CD630_n00560 | 63 -- 188 | 38 -- 184 | -11.2472 | 4.43918 | 5.82053 |
| CD630_18610 | CD630_n00560 | 21 -- 33 | 666 -- 679 | -11.2404 | 4.15582 | 5.82053 |
| CD630_24920 | CD630_n00560 | 120 -- 153 | 454 -- 486 | -11.2304 | 3.89455 | 5.82053 |
| CD630_09770 | CD630_n00560 | 1 -- 27 | 556 -- 585 | -11.2285 | 3.22106 | 5.82053 |
| cobC | CD630_n00560 | 175 -- 197 | 473 -- 495 | -11.2161 | 3.15854 | 5.82053 |
| gatC | CD630_n00560 | 14 -- 157 | 98 -- 242 | -11.2151 | 4.25143 | 5.82053 |
| CD630_22070 | CD630_n00560 | 8 -- 48 | 465 -- 502 | -11.2144 | 4.09651 | 5.82053 |
| CD630_03040 | CD630_n00560 | 102 -- 128 | 460 -- 483 | -11.2105 | 3.51403 | 5.82053 |
| CD630_14700 | CD630_n00560 | 172 -- 190 | 675 -- 693 | -11.2071 | 3.98234 | 5.82053 |
| CD630_19840 | CD630_n00560 | 165 -- 195 | 444 -- 483 | -11.2007 | 3.44724 | 5.82053 |
| CD630_27520 | CD630_n00560 | 113 -- 150 | 673 -- 712 | -11.1729 | 4.27693 | 5.82053 |
| CD630_05280 | CD630_n00560 | 10 -- 50 | 101 -- 143 | -11.1573 | 3.91665 | 5.82053 |
| xdhA | CD630_n00560 | 41 -- 57 | 534 -- 547 | -11.1527 | 4.45147 | 5.82053 |
| CD630_24760 | CD630_n00560 | 97 -- 129 | 462 -- 490 | -11.1026 | 5.19485 | 5.82053 |
| CD630_17770 | CD630_n00560 | 129 -- 147 | 474 -- 493 | -11.0897 | 5.06490 | 5.82053 |
| CD630_14990 | CD630_n00560 | 1 -- 36 | 675 -- 706 | -11.0745 | 3.78619 | 5.82053 |
| CD630_06970 | CD630_n00560 | 90 -- 107 | 673 -- 690 | -11.072 | 3.82312 | 5.82053 |
| CD630_27240 | CD630_n00560 | 3 -- 28 | 459 -- 482 | -11.0612 | 3.52462 | 5.82053 |
| bcp | CD630_n00560 | 93 -- 135 | 158 -- 203 | -11.0516 | 3.16670 | 5.82053 |
| spaK | CD630_n00560 | 78 -- 96 | 671 -- 693 | -11.0508 | 3.04530 | 5.82053 |
| CD630_18120 | CD630_n00560 | 50 -- 99 | 675 -- 708 | -11.0416 | 3.99713 | 5.82053 |
| bglA | CD630_n00560 | 166 -- 186 | 460 -- 483 | -11.0202 | 4.86402 | 5.82053 |
| CD630_05821 | CD630_n00560 | 68 -- 100 | 79 -- 113 | -10.9943 | 3.75661 | 5.82053 |
| CD630_20450 | CD630_n00560 | 37 -- 71 | 676 -- 701 | -10.9849 | 3.77388 | 5.82053 |
| CD630_16150 | CD630_n00560 | 14 -- 61 | 657 -- 701 | -10.9834 | 4.20614 | 5.82053 |
| CD630_03410 | CD630_n00560 | 103 -- 132 | 527 -- 558 | -10.9762 | 4.45994 | 5.82053 |
| CD630_15360 | CD630_n00560 | 113 -- 132 | 297 -- 316 | -10.9739 | 4.47208 | 5.82053 |
| cobQ | CD630_n00560 | 130 -- 153 | 472 -- 494 | -10.9625 | 3.13524 | 5.82053 |
| CD630_21330 | CD630_n00560 | 8 -- 52 | 671 -- 713 | -10.9534 | 4.10111 | 5.82053 |
| CD630_31210 | CD630_n00560 | 147 -- 178 | 671 -- 703 | -10.9406 | 4.17693 | 5.82053 |
| CD630_18960 | CD630_n00560 | 22 -- 74 | 441 -- 482 | -10.9373 | 4.37822 | 5.82053 |
| CD630_14610 | CD630_n00560 | 33 -- 54 | 458 -- 477 | -10.9188 | 4.89428 | 5.82053 |
| kdgT1 | CD630_n00560 | 173 -- 201 | 119 -- 149 | -10.8985 | 4.49207 | 5.82053 |
| CD630_15080 | CD630_n00560 | 48 -- 73 | 280 -- 306 | -10.8965 | 4.03033 | 5.82053 |
| CD630_08090 | CD630_n00560 | 150 -- 186 | 669 -- 708 | -10.8918 | 3.57167 | 5.82053 |
| CD630_06060 | CD630_n00560 | 95 -- 119 | 94 -- 115 | -10.8478 | 4.94050 | 5.82053 |
| CD630_21490 | CD630_n00560 | 191 -- 201 | 521 -- 531 | -10.8478 | 3.37462 | 5.82053 |
| CD630_13560 | CD630_n00560 | 1 -- 28 | 675 -- 701 | -10.8303 | 3.38083 | 5.82053 |
| CD630_08100 | CD630_n00560 | 170 -- 190 | 526 -- 559 | -10.8186 | 4.39181 | 5.82053 |
| CD630_08130 | CD630_n00560 | 74 -- 117 | 197 -- 235 | -10.8056 | 4.63831 | 5.82053 |
| dltB | CD630_n00560 | 25 -- 59 | 21 -- 58 | -10.7818 | 3.81485 | 5.82053 |
| CD630_30360 | CD630_n00560 | 158 -- 184 | 101 -- 131 | -10.7802 | 4.26509 | 5.82053 |
| CD630_21700 | CD630_n00560 | 25 -- 49 | 151 -- 170 | -10.7756 | 3.96650 | 5.82053 |
| CD630_19401 | CD630_n00560 | 16 -- 84 | 23 -- 82 | -10.7671 | 4.37084 | 5.82053 |
| CD630_14600 | CD630_n00560 | 15 -- 37 | 535 -- 559 | -10.7663 | 5.90348 | 5.82053 |
| glnA | CD630_n00560 | 132 -- 159 | 679 -- 703 | -10.7553 | 3.87629 | 5.82053 |
| sspA | CD630_n00560 | 64 -- 94 | 671 -- 699 | -10.7457 | 3.62689 | 5.82053 |
| CD630_16450 | CD630_n00560 | 81 -- 106 | 674 -- 703 | -10.7409 | 4.57547 | 5.82053 |
| CD630_10220 | CD630_n00560 | 119 -- 166 | 181 -- 228 | -10.7324 | 4.08233 | 5.82053 |
| CD630_16330 | CD630_n00560 | 149 -- 200 | 121 -- 173 | -10.7316 | 4.11180 | 5.82053 |
| CD630_18930 | CD630_n00560 | 147 -- 164 | 531 -- 546 | -10.7156 | 5.32127 | 5.82053 |
| CD630_09570 | CD630_n00560 | 136 -- 191 | 657 -- 713 | -10.706 | 5.31213 | 5.82053 |
| CD630_29070 | CD630_n00560 | 136 -- 191 | 657 -- 713 | -10.706 | 3.89477 | 5.82053 |
| CD630_05880 | CD630_n00560 | 65 -- 109 | 671 -- 708 | -10.7043 | 4.28339 | 5.82053 |
| CD630_33560 | CD630_n00560 | 147 -- 178 | 693 -- 723 | -10.689 | 4.03147 | 5.82053 |
| CD630_29750 | CD630_n00560 | 71 -- 94 | 109 -- 133 | -10.6884 | 4.59146 | 5.82053 |
| CD630_25040 | CD630_n00560 | 142 -- 188 | 473 -- 514 | -10.6827 | 3.43898 | 5.82053 |
| hfq | CD630_n00560 | 3 -- 13 | 125 -- 135 | -10.6737 | 5.05609 | 5.82053 |
| srlD | CD630_n00560 | 102 -- 134 | 461 -- 483 | -10.6726 | 4.19843 | 5.82053 |
| pilT | CD630_n00560 | 128 -- 150 | 618 -- 638 | -10.6697 | 3.24639 | 5.82053 |
| CD630_23780 | CD630_n00560 | 38 -- 107 | 527 -- 585 | -10.6649 | 3.91629 | 5.82053 |
| CD630_34030 | CD630_n00560 | 171 -- 189 | 41 -- 58 | -10.6581 | 3.05766 | 5.82053 |
| CD630_05300 | CD630_n00560 | 148 -- 179 | 533 -- 565 | -10.6542 | 4.56885 | 5.82053 |
| CD630_03420 | CD630_n00560 | 174 -- 194 | 464 -- 486 | -10.647 | 4.71868 | 5.82053 |
| CD630_02430 | CD630_n00560 | 64 -- 131 | 411 -- 481 | -10.5919 | 3.71048 | 5.82053 |
| CD630_17160 | CD630_n00560 | 90 -- 112 | 461 -- 478 | -10.588 | 3.50228 | 5.82053 |
| moeA | CD630_n00560 | 102 -- 119 | 677 -- 693 | -10.5879 | 3.26118 | 5.82053 |
| CD630_21620 | CD630_n00560 | 66 -- 119 | 157 -- 212 | -10.5861 | 3.02649 | 5.82053 |
| CD630_30540 | CD630_n00560 | 182 -- 198 | 676 -- 692 | -10.5843 | 3.63804 | 5.82053 |
| uxaA%27 | CD630_n00560 | 138 -- 188 | 86 -- 144 | -10.5522 | 4.26165 | 5.82053 |
| CD630_06080 | CD630_n00560 | 158 -- 193 | 668 -- 701 | -10.5072 | 3.82315 | 5.82053 |
| CD630_29250 | CD630_n00560 | 11 -- 43 | 335 -- 365 | -10.5005 | 4.61901 | 5.82053 |
| CD630_09410 | CD630_n00560 | 11 -- 43 | 335 -- 365 | -10.5005 | 4.75651 | 5.82053 |
| CD630_30140 | CD630_n00560 | 174 -- 194 | 462 -- 483 | -10.4734 | 3.28343 | 5.82053 |
| CD630_13790 | CD630_n00560 | 165 -- 197 | 395 -- 423 | -10.4562 | 5.17374 | 5.82053 |
| CD630_14810 | CD630_n00560 | 121 -- 150 | 463 -- 501 | -10.4458 | 3.35323 | 5.82053 |
| CD630_03180 | CD630_n00560 | 176 -- 193 | 474 -- 492 | -10.4449 | 3.73138 | 5.82053 |
| CD630_30730 | CD630_n00560 | 8 -- 35 | 567 -- 599 | -10.43 | 3.13236 | 5.82053 |
| CD630_05870 | CD630_n00560 | 135 -- 144 | 274 -- 283 | -10.4294 | 4.28339 | 5.82053 |
| CD630_13780 | CD630_n00560 | 144 -- 192 | 125 -- 174 | -10.4272 | 3.52274 | 5.82053 |
| CD630_27610 | CD630_n00560 | 15 -- 35 | 695 -- 715 | -10.4191 | 3.96465 | 5.82053 |
| CD630_24050 | CD630_n00560 | 35 -- 57 | 337 -- 361 | -10.4135 | 3.70192 | 5.82053 |
| CD630_21460 | CD630_n00560 | 134 -- 172 | 656 -- 693 | -10.4118 | 4.83005 | 5.82053 |
| CD630_22200 | CD630_n00560 | 141 -- 173 | 462 -- 494 | -10.384 | 4.18288 | 5.82053 |
| CD630_02740 | CD630_n00560 | 157 -- 199 | 470 -- 508 | -10.3389 | 5.02277 | 5.82053 |
| CD630_01690 | CD630_n00560 | 11 -- 22 | 665 -- 677 | -10.2987 | 4.97234 | 5.82053 |
| CD630_23770 | CD630_n00560 | 120 -- 133 | 526 -- 539 | -10.2738 | 3.25181 | 5.82053 |
| CD630_14250 | CD630_n00560 | 177 -- 198 | 157 -- 179 | -10.2687 | 4.19604 | 5.82053 |
| CD630_33570 | CD630_n00560 | 153 -- 175 | 676 -- 697 | -10.2629 | 3.95778 | 5.82053 |
| cheW | CD630_n00560 | 12 -- 49 | 674 -- 703 | -10.2553 | 3.72371 | 5.82053 |
| CD630_12110 | CD630_n00560 | 133 -- 150 | 465 -- 482 | -10.2457 | 4.36078 | 5.82053 |
| CD630_10830 | CD630_n00560 | 164 -- 184 | 461 -- 484 | -10.2363 | 4.82398 | 5.82053 |
| CD630_36250 | CD630_n00560 | 50 -- 77 | 671 -- 701 | -10.234 | 4.13224 | 5.82053 |
| CD630_07070 | CD630_n00560 | 29 -- 53 | 124 -- 148 | -10.2304 | 3.46414 | 5.82053 |
| CD630_18010 | CD630_n00560 | 6 -- 22 | 332 -- 348 | -10.2263 | 4.21967 | 5.82053 |
| CD630_31580 | CD630_n00560 | 129 -- 153 | 463 -- 483 | -10.2079 | 3.51756 | 5.82053 |
| CD630_36170 | CD630_n00560 | 4 -- 13 | 536 -- 545 | -10.2062 | 3.72380 | 5.82053 |
| srlEb | CD630_n00560 | 59 -- 96 | 674 -- 706 | -10.1948 | 3.50140 | 5.82053 |
| CD630_17710 | CD630_n00560 | 161 -- 188 | 457 -- 484 | -10.1875 | 4.37688 | 5.82053 |
| CD630_13920 | CD630_n00560 | 132 -- 157 | 84 -- 109 | -10.1771 | 3.25236 | 5.82053 |
| CD630_08360 | CD630_n00560 | 112 -- 146 | 465 -- 494 | -10.1576 | 3.19339 | 5.82053 |
| CD630_35790 | CD630_n00560 | 97 -- 129 | 468 -- 508 | -10.1561 | 6.97665 | 5.82053 |
| CD630_31530 | CD630_n00560 | 22 -- 44 | 262 -- 284 | -10.1503 | 3.49100 | 5.82053 |
| CD630_11730 | CD630_n00560 | 142 -- 159 | 679 -- 700 | -10.1408 | 4.80521 | 5.82053 |
| CD630_28870 | CD630_n00560 | 141 -- 160 | 126 -- 150 | -10.1316 | 3.41991 | 5.82053 |
| CD630_07121 | CD630_n00560 | 148 -- 176 | 117 -- 149 | -10.1148 | 3.26177 | 5.82053 |
| fliI | CD630_n00560 | 41 -- 87 | 672 -- 712 | -10.1059 | 3.03666 | 5.82053 |
| CD630_05490 | CD630_n00560 | 7 -- 35 | 529 -- 557 | -10.1003 | 3.89852 | 5.82053 |
| CD630_26050 | CD630_n00560 | 34 -- 53 | 671 -- 690 | -10.08 | 6.99851 | 5.82053 |
| CD630_25310 | CD630_n00560 | 23 -- 40 | 461 -- 475 | -10.0683 | 3.99861 | 5.82053 |
| CD630_16460 | CD630_n00560 | 135 -- 168 | 462 -- 486 | -10.0664 | 4.64028 | 5.82053 |
| CD630_22720 | CD630_n00560 | 151 -- 166 | 470 -- 485 | -10.0637 | 3.81804 | 5.82053 |
| CD630_23820 | CD630_n00560 | 180 -- 194 | 463 -- 477 | -10.0542 | 3.18419 | 5.82053 |
| CD630_31381 | CD630_n00560 | 59 -- 104 | 330 -- 386 | -10.0444 | 3.49137 | 5.82053 |
| CD630_01560 | CD630_n00560 | 155 -- 194 | 27 -- 68 | -10.026 | 4.06352 | 5.82053 |
| CD630_20600 | CD630_n00560 | 33 -- 60 | 674 -- 701 | -10.0175 | 4.20093 | 5.82053 |
| CD630_13380 | CD630_n00560 | 171 -- 190 | 468 -- 487 | -10.0108 | 4.74408 | 5.82053 |
| CD630_32110 | CD630_n00560 | 81 -- 112 | 473 -- 507 | -10.0049 | 4.45291 | 5.82053 |
| LinCd | CD630_n00560 | 17 -- 28 | 461 -- 472 | -9.98209 | 4.33624 | 5.82053 |
| CD630_15890 | CD630_n00560 | 138 -- 167 | 668 -- 697 | -9.95473 | 3.39962 | 5.82053 |
| CD630_08510 | CD630_n00560 | 29 -- 80 | 395 -- 443 | -9.94987 | 5.16329 | 5.82053 |
| eutP | CD630_n00560 | 1 -- 27 | 470 -- 500 | -9.92942 | 3.82847 | 5.82053 |
| cprK | CD630_n00560 | 79 -- 97 | 537 -- 556 | -9.92893 | 3.96216 | 5.82053 |
| CD630_18230 | CD630_n00560 | 121 -- 159 | 462 -- 494 | -9.92574 | 3.00184 | 5.82053 |
| CD630_23440 | CD630_n00560 | 185 -- 199 | 470 -- 484 | -9.92549 | 3.34841 | 5.82053 |
| CD630_34530 | CD630_n00560 | 1 -- 16 | 686 -- 703 | -9.9249 | 3.83016 | 5.82053 |
| panG | CD630_n00560 | 167 -- 178 | 221 -- 232 | -9.91578 | 5.54047 | 5.82053 |
| CD630_22030 | CD630_n00560 | 113 -- 128 | 466 -- 482 | -9.91195 | 3.29934 | 5.82053 |
| CD630_32890 | CD630_n00560 | 31 -- 165 | 46 -- 174 | -9.90067 | 3.41276 | 5.82053 |
| kdpD | CD630_n00560 | 134 -- 149 | 674 -- 690 | -9.89496 | 4.27140 | 5.82053 |
| CD630_14140 | CD630_n00560 | 50 -- 73 | 62 -- 85 | -9.88238 | 3.16551 | 5.82053 |
| CD630_24770 | CD630_n00560 | 23 -- 78 | 112 -- 168 | -9.8801 | 3.20798 | 5.82053 |
| dpaA | CD630_n00560 | 83 -- 93 | 474 -- 484 | -9.87118 | 4.65364 | 5.82053 |
| CD630_16950 | CD630_n00560 | 187 -- 197 | 272 -- 282 | -9.86989 | 4.42569 | 5.82053 |
| CD630_25300 | CD630_n00560 | 87 -- 98 | 666 -- 677 | -9.86899 | 4.49333 | 5.82053 |
| CD630_31220 | CD630_n00560 | 137 -- 160 | 19 -- 44 | -9.86099 | 3.73779 | 5.82053 |
| CD630_07760 | CD630_n00560 | 73 -- 105 | 669 -- 703 | -9.85926 | 3.64186 | 5.82053 |
| rolB | CD630_n00560 | 171 -- 200 | 674 -- 712 | -9.85911 | 3.60270 | 5.82053 |
| CD630_19921 | CD630_n00560 | 116 -- 130 | 468 -- 482 | -9.85711 | 5.46345 | 5.82053 |
| CD630_05950 | CD630_n00560 | 65 -- 92 | 459 -- 488 | -9.8532 | 3.31669 | 5.82053 |
| CD630_06200 | CD630_n00560 | 71 -- 92 | 617 -- 638 | -9.81826 | 3.57064 | 5.82053 |
| CD630_04410 | CD630_n00560 | 132 -- 162 | 669 -- 703 | -9.79597 | 3.90451 | 5.82053 |
| CD630_09970 | CD630_n00560 | 9 -- 29 | 462 -- 483 | -9.79487 | 3.25917 | 5.82053 |
| CD630_07840 | CD630_n00560 | 87 -- 108 | 464 -- 482 | -9.78967 | 4.26761 | 5.82053 |

| | | | | | | |
|---|---|---|---|---|---|---|
| CD630_19690 | CD630_n00560 | 13 -- 42 | 526 -- 558 | -9.78859 | 3.85861 | 5.82053 |
| CD630_18781 | CD630_n00560 | 15 -- 46 | 665 -- 710 | -9.77694 | 3.77660 | 5.82053 |
| CD630_06350 | CD630_n00560 | 64 -- 104 | 462 -- 498 | -9.74726 | 4.35983 | 5.82053 |
| CD630_11241 | CD630_n00560 | 38 -- 52 | 695 -- 710 | -9.747 | 4.85723 | 5.82053 |
| CD630_06140 | CD630_n00560 | 35 -- 58 | 66 -- 90 | -9.74274 | 4.12519 | 5.82053 |
| CD630_13580 | CD630_n00560 | 133 -- 154 | 547 -- 568 | -9.74262 | 3.80715 | 5.82053 |
| CD630_13400 | CD630_n00560 | 94 -- 179 | 161 -- 248 | -9.74205 | 3.20232 | 5.82053 |
| CD630_28290 | CD630_n00560 | 16 -- 104 | 282 -- 360 | -9.74038 | 5.54192 | 5.82053 |
| CD630_04320 | CD630_n00560 | 79 -- 121 | 472 -- 513 | -9.73162 | 3.68342 | 5.82053 |
| CD630_14130 | CD630_n00560 | 120 -- 168 | 524 -- 559 | -9.72236 | 4.57970 | 5.82053 |
| CD630_18970 | CD630_n00560 | 10 -- 46 | 462 -- 507 | -9.71957 | 3.57243 | 5.82053 |
| CD630_06050 | CD630_n00560 | 49 -- 144 | 625 -- 703 | -9.71898 | 3.59332 | 5.82053 |
| CD630_08070 | CD630_n00560 | 106 -- 134 | 127 -- 168 | -9.71403 | 4.87819 | 5.82053 |
| CD630_26250 | CD630_n00560 | 172 -- 192 | 474 -- 494 | -9.71259 | 5.03379 | 5.82053 |
| CD630_14180 | CD630_n00560 | 169 -- 188 | 268 -- 288 | -9.70831 | 3.93696 | 5.82053 |
| CD630_31382 | CD630_n00560 | 42 -- 76 | 467 -- 498 | -9.69565 | 3.49137 | 5.82053 |
| opuCA | CD630_n00560 | 26 -- 92 | 656 -- 712 | -9.69285 | 4.07035 | 5.82053 |
| cwp17 | CD630_n00560 | 4 -- 27 | 337 -- 360 | -9.66456 | 3.43363 | 5.82053 |
| CD630_07430 | CD630_n00560 | 57 -- 67 | 37 -- 47 | -9.66158 | 4.65568 | 5.82053 |
| CD630_08460 | CD630_n00560 | 174 -- 199 | 675 -- 706 | -9.65816 | 4.15083 | 5.82053 |
| CD630_32720 | CD630_n00560 | 47 -- 95 | 671 -- 722 | -9.6427 | 4.74276 | 5.82053 |
| CD630_27280 | CD630_n00560 | 124 -- 162 | 521 -- 558 | -9.64168 | 3.24738 | 5.82053 |
| CD630_04620 | CD630_n00560 | 106 -- 131 | 674 -- 702 | -9.64037 | 3.45413 | 5.82053 |
| dnaD | CD630_n00560 | 109 -- 129 | 462 -- 483 | -9.63699 | 3.56078 | 5.82053 |
| CD630_15060 | CD630_n00560 | 43 -- 92 | 462 -- 517 | -9.6351 | 3.69396 | 5.82053 |
| ldh | CD630_n00560 | 144 -- 189 | 457 -- 499 | -9.63286 | 4.35818 | 5.82053 |
| CD630_23000 | CD630_n00560 | 149 -- 183 | 673 -- 701 | -9.63116 | 3.54130 | 5.82053 |
| gabT | CD630_n00560 | 98 -- 136 | 82 -- 119 | -9.62259 | 3.67067 | 5.82053 |
| CD630_04940 | CD630_n00560 | 93 -- 123 | 673 -- 701 | -9.62019 | 4.20948 | 5.82053 |
| CD630_04550 | CD630_n00560 | 104 -- 170 | 106 -- 170 | -9.61363 | 3.05227 | 5.82053 |
| CD630_04770 | CD630_n00560 | 79 -- 99 | 457 -- 477 | -9.60956 | 3.58478 | 5.82053 |
| CD630_16550 | CD630_n00560 | 72 -- 102 | 650 -- 697 | -9.60478 | 5.61064 | 5.82053 |
| CD630_15160 | CD630_n00560 | 89 -- 135 | 113 -- 179 | -9.60177 | 4.32318 | 5.82053 |
| scpB | CD630_n00560 | 141 -- 157 | 535 -- 551 | -9.59528 | 3.33560 | 5.82053 |
| CD630_06560 | CD630_n00560 | 90 -- 139 | 378 -- 422 | -9.59254 | 4.02532 | 5.82053 |
| CD630_05940 | CD630_n00560 | 64 -- 86 | 338 -- 366 | -9.58711 | 4.18595 | 5.82053 |
| CD630_07960 | CD630_n00560 | 185 -- 198 | 532 -- 545 | -9.56677 | 3.22537 | 5.82053 |
| CD630_36781 | CD630_n00560 | 165 -- 196 | 466 -- 493 | -9.56365 | 3.75946 | 5.82053 |
| CD630_33610 | CD630_n00560 | 11 -- 34 | 103 -- 133 | -9.56246 | 3.79322 | 5.82053 |
| cstA | CD630_n00560 | 163 -- 182 | 467 -- 485 | -9.56169 | 3.53584 | 5.82053 |
| CD630_31000 | CD630_n00560 | 80 -- 125 | 668 -- 703 | -9.54098 | 3.70946 | 5.82053 |
| CD630_10970 | CD630_n00560 | 131 -- 153 | 125 -- 146 | -9.53681 | 3.53850 | 5.82053 |
| CD630_14340 | CD630_n00560 | 51 -- 71 | 534 -- 561 | -9.53609 | 5.10868 | 5.82053 |
| CD630_07480 | CD630_n00560 | 167 -- 190 | 24 -- 50 | -9.53286 | 5.18321 | 5.82053 |
| CD630_09760 | CD630_n00560 | 120 -- 154 | 473 -- 507 | -9.53275 | 4.83410 | 5.82053 |
| CD630_28910 | CD630_n00560 | 120 -- 154 | 473 -- 507 | -9.53275 | 4.51664 | 5.82053 |
| CD630_30310 | CD630_n00560 | 154 -- 194 | 522 -- 560 | -9.5219 | 4.56059 | 5.82053 |
| gcvTPA | CD630_n00560 | 37 -- 44 | 539 -- 546 | -9.51141 | 4.77229 | 5.82053 |
| CD630_08110 | CD630_n00560 | 110 -- 146 | 415 -- 451 | -9.50503 | 4.39181 | 5.82053 |
| CD630_20541 | CD630_n00560 | 41 -- 55 | 533 -- 546 | -9.50363 | 5.74527 | 5.82053 |
| CD630_13350 | CD630_n00600 | 165 -- 201 | 517 -- 557 | -19.6351 | 4.15139 | 5.31146 |
| CD630_24590 | CD630_n00600 | 52 -- 200 | 363 -- 508 | -19.2754 | 3.70401 | 5.31146 |
| cheW | CD630_n00600 | 4 -- 80 | 506 -- 572 | -18.7533 | 3.72371 | 5.31146 |
| etfA | CD630_n00600 | 125 -- 173 | 792 -- 839 | -18.6597 | 3.41204 | 5.31146 |
| CD630_21620 | CD630_n00600 | 50 -- 189 | 731 -- 847 | -18.5956 | 3.02649 | 5.31146 |
| CD630_32880 | CD630_n00600 | 134 -- 201 | 537 -- 601 | -18.3725 | 3.41276 | 5.31146 |
| nusB | CD630_n00600 | 54 -- 170 | 1004 -- 1129 | -18.2441 | 4.92798 | 5.31146 |
| CD630_28580 | CD630_n00600 | 124 -- 177 | 917 -- 961 | -17.9883 | 4.67130 | 5.31146 |
| CD630_18940 | CD630_n00600 | 87 -- 163 | 542 -- 605 | -17.8314 | 4.09988 | 5.31146 |
| pilT | CD630_n00600 | 121 -- 188 | 747 -- 814 | -17.7012 | 3.24639 | 5.31146 |
| cspD | CD630_n00600 | 116 -- 155 | 193 -- 235 | -17.6408 | 3.23587 | 5.31146 |
| CD630_24220 | CD630_n00600 | 74 -- 178 | 268 -- 380 | -17.6032 | 4.18037 | 5.31146 |
| CD630_21570 | CD630_n00600 | 123 -- 162 | 797 -- 836 | -17.5618 | 6.48385 | 5.31146 |
| CD630_32690 | CD630_n00600 | 68 -- 130 | 530 -- 576 | -17.4364 | 7.32676 | 5.31146 |
| srlEb | CD630_n00600 | 44 -- 93 | 735 -- 799 | -17.355 | 3.50140 | 5.31146 |
| CD630_31000 | CD630_n00600 | 106 -- 153 | 517 -- 551 | -17.2763 | 3.70946 | 5.31146 |
| CD630_27520 | CD630_n00600 | 86 -- 181 | 669 -- 750 | -17.2748 | 4.27693 | 5.31146 |
| panB | CD630_n00600 | 100 -- 187 | 114 -- 201 | -17.134 | 3.40248 | 5.31146 |
| CD630_04280 | CD630_n00600 | 87 -- 112 | 539 -- 566 | -17.033 | 3.61457 | 5.31146 |
| flgB | CD630_n00600 | 97 -- 201 | 486 -- 604 | -16.9754 | 3.25736 | 5.31146 |
| CD630_27280 | CD630_n00600 | 150 -- 179 | 931 -- 959 | -16.8096 | 3.24738 | 5.31146 |
| CD630_19904 | CD630_n00600 | 130 -- 180 | 793 -- 838 | -16.6736 | 5.96526 | 5.31146 |
| CD630_32520 | CD630_n00600 | 126 -- 153 | 665 -- 691 | -16.6406 | 4.06098 | 5.31146 |
| CD630_07400 | CD630_n00600 | 36 -- 67 | 120 -- 152 | -16.5961 | 4.52273 | 5.31146 |
| CD630_19921 | CD630_n00600 | 60 -- 199 | 779 -- 924 | -16.5579 | 5.46345 | 5.31146 |
| CD630_15360 | CD630_n00600 | 79 -- 157 | 988 -- 1080 | -16.5334 | 4.47208 | 5.31146 |
| etfA | CD630_n00600 | 63 -- 127 | 759 -- 821 | -16.5233 | 4.80521 | 5.31146 |
| eutV | CD630_n00600 | 87 -- 172 | 734 -- 818 | -16.5176 | 4.55715 | 5.31146 |
| CD630_06060 | CD630_n00600 | 70 -- 179 | 468 -- 575 | -16.4929 | 4.94050 | 5.31146 |
| CD630_31880 | CD630_n00600 | 130 -- 180 | 515 -- 569 | -16.2563 | 3.97522 | 5.31146 |
| mobA | CD630_n00600 | 132 -- 200 | 790 -- 836 | -16.2138 | 4.25703 | 5.31146 |
| CD630_15850 | CD630_n00600 | 161 -- 197 | 1035 -- 1078 | -16.1518 | 4.32968 | 5.31146 |
| scpB | CD630_n00600 | 120 -- 201 | 501 -- 576 | -16.1288 | 3.33560 | 5.31146 |
| CD630_33570 | CD630_n00600 | 98 -- 175 | 565 -- 641 | -16.1108 | 3.95778 | 5.31146 |
| CD630_24770 | CD630_n00600 | 168 -- 186 | 544 -- 564 | -16.0814 | 3.20798 | 5.31146 |
| CD630_02430 | CD630_n00600 | 68 -- 126 | 407 -- 446 | -16.0453 | 3.71048 | 5.31146 |
| bglA | CD630_n00600 | 22 -- 108 | 668 -- 748 | -16.0315 | 4.50315 | 5.31146 |
| CD630_27120 | CD630_n00600 | 5 -- 48 | 599 -- 643 | -15.9632 | 4.43720 | 5.31146 |
| CD630_24580 | CD630_n00600 | 11 -- 46 | 105 -- 140 | -15.9042 | 3.88519 | 5.31146 |
| CD630_08090 | CD630_n00600 | 150 -- 194 | 530 -- 578 | -15.8019 | 3.57167 | 5.31146 |
| CD630_13780 | CD630_n00600 | 156 -- 196 | 538 -- 576 | -15.7658 | 3.52274 | 5.31146 |
| CD630_14260 | CD630_n00600 | 22 -- 70 | 547 -- 591 | -15.7122 | 3.22214 | 5.31146 |
| cprK | CD630_n00600 | 87 -- 149 | 532 -- 571 | -15.7019 | 3.96216 | 5.31146 |
| CD630_09840 | CD630_n00600 | 90 -- 156 | 539 -- 618 | -15.6523 | 4.50009 | 5.31146 |
| xdhA | CD630_n00600 | 73 -- 160 | 539 -- 615 | -15.6288 | 4.45147 | 5.31146 |
| CD630_23770 | CD630_n00600 | 102 -- 150 | 556 -- 606 | -15.6219 | 3.25181 | 5.31146 |
| def1 | CD630_n00600 | 106 -- 179 | 508 -- 574 | -15.5664 | 3.99720 | 5.31146 |
| CD630_02740 | CD630_n00600 | 144 -- 174 | 538 -- 571 | -15.5561 | 5.02277 | 5.31146 |
| bglA | CD630_n00600 | 48 -- 70 | 536 -- 563 | -15.5293 | 4.86402 | 5.31146 |
| CD630_32720 | CD630_n00600 | 61 -- 197 | 248 -- 381 | -15.5092 | 4.74276 | 5.31146 |
| argS | CD630_n00600 | 75 -- 108 | 413 -- 440 | -15.5059 | 4.96036 | 5.31146 |
| CD630_27570 | CD630_n00600 | 152 -- 200 | 793 -- 843 | -15.4935 | 3.46216 | 5.31146 |
| CD630_19801 | CD630_n00600 | 160 -- 200 | 530 -- 575 | -15.4857 | 5.22035 | 5.31146 |
| CD630_25190 | CD630_n00600 | 164 -- 201 | 536 -- 568 | -15.4852 | 3.19663 | 5.31146 |
| CD630_28290 | CD630_n00600 | 172 -- 198 | 58 -- 82 | -15.4823 | 5.54192 | 5.31146 |
| rbsC | CD630_n00600 | 157 -- 189 | 795 -- 832 | -15.4301 | 3.71650 | 5.31146 |
| malY | CD630_n00600 | 57 -- 102 | 726 -- 768 | -15.3853 | 4.46345 | 5.31146 |
| CD630_10880 | CD630_n00600 | 147 -- 200 | 522 -- 573 | -15.3701 | 5.05048 | 5.31146 |
| CD630_25320 | CD630_n00600 | 142 -- 171 | 792 -- 819 | -15.3672 | 4.29415 | 5.31146 |
| CD630_32890 | CD630_n00600 | 79 -- 118 | 200 -- 236 | -15.2947 | 3.41276 | 5.31146 |
| CD630_24920 | CD630_n00600 | 22 -- 129 | 304 -- 445 | -15.2564 | 3.89455 | 5.31146 |
| CD630_09040 | CD630_n00600 | 164 -- 201 | 536 -- 568 | -15.2533 | 4.11851 | 5.31146 |
| CD630_11730 | CD630_n00600 | 140 -- 184 | 403 -- 442 | -15.2076 | 4.80521 | 5.31146 |
| CD630_17380 | CD630_n00600 | 168 -- 197 | 665 -- 693 | -15.2012 | 5.03207 | 5.31146 |
| CD630_09770 | CD630_n00600 | 6 -- 31 | 360 -- 384 | -15.1216 | 3.22106 | 5.31146 |
| CD630_19000 | CD630_n00600 | 164 -- 188 | 800 -- 824 | -15.0697 | 4.18443 | 5.31146 |
| CD630_16150 | CD630_n00600 | 19 -- 115 | 347 -- 456 | -15.0696 | 4.20614 | 5.31146 |
| CD630_13950 | CD630_n00600 | 25 -- 76 | 325 -- 371 | -15.0638 | 4.25540 | 5.31146 |
| CD630_10050 | CD630_n00600 | 157 -- 181 | 804 -- 834 | -15.0509 | 3.93750 | 5.31146 |
| CD630_17970 | CD630_n00600 | 59 -- 180 | 483 -- 631 | -15.0252 | 3.68802 | 5.31146 |
| CD630_19790 | CD630_n00600 | 49 -- 100 | 730 -- 777 | -14.9909 | 3.79593 | 5.31146 |
| CD630_08390 | CD630_n00600 | 18 -- 115 | 351 -- 439 | -14.96 | 5.16000 | 5.31146 |
| CD630_36781 | CD630_n00600 | 143 -- 169 | 930 -- 961 | -14.9331 | 3.75946 | 5.31146 |
| CD630_19840 | CD630_n00600 | 154 -- 195 | 710 -- 755 | -14.929 | 3.44724 | 5.31146 |
| CD630_11220 | CD630_n00600 | 36 -- 92 | 746 -- 818 | -14.9237 | 3.52327 | 5.31146 |
| CD630_09760 | CD630_n00600 | 170 -- 191 | 818 -- 839 | -14.9217 | 4.83410 | 5.31146 |
| CD630_28910 | CD630_n00600 | 170 -- 191 | 818 -- 839 | -14.9217 | 4.51664 | 5.31146 |
| CD630_22890 | CD630_n00600 | 3 -- 47 | 1053 -- 1111 | -14.9062 | 3.75591 | 5.31146 |
| CD630_07770 | CD630_n00600 | 69 -- 161 | 482 -- 598 | -14.8854 | 3.31158 | 5.31146 |
| cobQ | CD630_n00600 | 89 -- 147 | 486 -- 551 | -14.8564 | 3.13524 | 5.31146 |
| flhB | CD630_n00600 | 159 -- 197 | 12 -- 54 | -14.8105 | 3.55524 | 5.31146 |
| CD630_14610 | CD630_n00600 | 136 -- 177 | 168 -- 214 | -14.8019 | 4.89428 | 5.31146 |
| CD630_31382 | CD630_n00600 | 98 -- 185 | 675 -- 744 | -14.7897 | 3.49137 | 5.31146 |
| glgB | CD630_n00600 | 91 -- 147 | 500 -- 551 | -14.7236 | 4.48273 | 5.31146 |
| CD630_33300 | CD630_n00600 | 89 -- 180 | 286 -- 384 | -14.6634 | 3.79149 | 5.31146 |
| CD630_18100 | CD630_n00600 | 88 -- 123 | 779 -- 816 | -14.6382 | 5.71658 | 5.31146 |
| CD630_14110 | CD630_n00600 | 149 -- 200 | 517 -- 568 | -14.58 | 3.39550 | 5.31146 |
| CD630_33600 | CD630_n00600 | 8 -- 33 | 352 -- 385 | -14.5274 | 3.73196 | 5.31146 |
| cdtR | CD630_n00600 | 177 -- 195 | 214 -- 233 | -14.5071 | 5.84368 | 5.31146 |
| appC | CD630_n00600 | 43 -- 76 | 161 -- 196 | -14.4976 | 4.50438 | 5.31146 |
| CD630_07480 | CD630_n00600 | 69 -- 100 | 797 -- 827 | -14.4965 | 5.18321 | 5.31146 |
| metQ | CD630_n00600 | 149 -- 197 | 1037 -- 1081 | -14.4897 | 3.87190 | 5.31146 |
| CD630_10830 | CD630_n00600 | 92 -- 199 | 517 -- 605 | -14.4861 | 4.82398 | 5.31146 |
| CD630_25610 | CD630_n00600 | 69 -- 122 | 311 -- 365 | -14.4838 | 3.02685 | 5.31146 |
| CD630_10970 | CD630_n00600 | 25 -- 153 | 127 -- 273 | -14.4738 | 3.53850 | 5.31146 |
| CD630_07760 | CD630_n00600 | 10 -- 53 | 513 -- 562 | -14.4608 | 3.64186 | 5.31146 |
| CD630_36250 | CD630_n00600 | 50 -- 90 | 529 -- 571 | -14.4059 | 4.13224 | 5.31146 |
| CD630_13560 | CD630_n00600 | 76 -- 102 | 93 -- 120 | -14.3928 | 3.38083 | 5.31146 |
| blaI | CD630_n00600 | 79 -- 132 | 564 -- 621 | -14.3815 | 3.73674 | 5.31146 |
| spaK | CD630_n00600 | 174 -- 200 | 124 -- 149 | -14.3106 | 3.04530 | 5.31146 |
| CD630_31910 | CD630_n00600 | 39 -- 64 | 213 -- 238 | -14.2584 | 3.48314 | 5.31146 |
| CD630_25620 | CD630_n00600 | 85 -- 117 | 86 -- 125 | -14.2522 | 3.02685 | 5.31146 |
| CD630_04320 | CD630_n00600 | 80 -- 157 | 517 -- 606 | -14.1935 | 3.68342 | 5.31146 |
| CD630_18120 | CD630_n00600 | 3 -- 34 | 134 -- 167 | -14.1503 | 3.99713 | 5.31146 |
| CD630_18400 | CD630_n00600 | 108 -- 124 | 800 -- 818 | -14.1223 | 4.67962 | 5.31146 |
| CD630_05950 | CD630_n00600 | 64 -- 109 | 599 -- 641 | -14.0874 | 3.31669 | 5.31146 |
| CD630_35260 | CD630_n00600 | 19 -- 109 | 85 -- 170 | -14.072 | 5.77277 | 5.31146 |
| CD630_08460 | CD630_n00600 | 95 -- 168 | 214 -- 280 | -14.0474 | 4.15083 | 5.31146 |
| CD630_35840 | CD630_n00600 | 58 -- 91 | 538 -- 567 | -14.0439 | 3.14170 | 5.31146 |
| CD630_12650 | CD630_n00600 | 93 -- 118 | 478 -- 503 | -14.0386 | 6.31827 | 5.31146 |
| iorA | CD630_n00600 | 86 -- 163 | 534 -- 606 | -14.0348 | 3.83173 | 5.31146 |
| gcvTPA | CD630_n00600 | 139 -- 198 | 543 -- 615 | -14.0079 | 4.77229 | 5.31146 |
| CD630_14190 | CD630_n00600 | 133 -- 172 | 759 -- 797 | -13.9935 | 3.78562 | 5.31146 |
| CD630_24760 | CD630_n00600 | 63 -- 196 | 321 -- 448 | -13.9757 | 5.19485 | 5.31146 |
| cwp29 | CD630_n00600 | 81 -- 137 | 507 -- 575 | -13.9498 | 4.40865 | 5.31146 |
| CD630_14250 | CD630_n00600 | 80 -- 198 | 1062 -- 1163 | -13.9467 | 4.19604 | 5.31146 |
| CD630_20171 | CD630_n00600 | 71 -- 105 | 797 -- 834 | -13.8667 | 6.87491 | 5.31146 |
| rolB | CD630_n00600 | 82 -- 133 | 1119 -- 1162 | -13.8576 | 3.60270 | 5.31146 |
| CD630_01700 | CD630_n00600 | 1 -- 45 | 125 -- 171 | -13.8194 | 4.97234 | 5.31146 |
| CD630_05480 | CD630_n00600 | 165 -- 183 | 665 -- 682 | -13.8105 | 4.27854 | 5.31146 |
| cspC | CD630_n00600 | 79 -- 161 | 184 -- 274 | -13.8002 | 5.48848 | 5.31146 |
| CD630_31490 | CD630_n00600 | 38 -- 122 | 102 -- 173 | -13.7932 | 4.08593 | 5.31146 |
| CD630_16460 | CD630_n00600 | 118 -- 141 | 129 -- 151 | -13.7797 | 4.64028 | 5.31146 |
| garK | CD630_n00600 | 152 -- 196 | 531 -- 571 | -13.7729 | 3.72836 | 5.31146 |
| acoL | CD630_n00600 | 92 -- 152 | 744 -- 818 | -13.7421 | 3.32169 | 5.31146 |
| CD630_05880 | CD630_n00600 | 89 -- 199 | 338 -- 428 | -13.7179 | 4.28339 | 5.31146 |
| CD630_30140 | CD630_n00600 | 70 -- 108 | 539 -- 584 | -13.6628 | 3.28343 | 5.31146 |
| CD630_02420 | CD630_n00600 | 175 -- 198 | 555 -- 579 | -13.6504 | 3.71048 | 5.31146 |
| CD630_13620 | CD630_n00600 | 185 -- 196 | 805 -- 816 | -13.6373 | 4.77627 | 5.31146 |
| cls | CD630_n00600 | 144 -- 173 | 122 -- 149 | -13.6165 | 4.46177 | 5.31146 |
| CD630_20460 | CD630_n00600 | 15 -- 100 | 677 -- 756 | -13.6054 | 7.94034 | 5.31146 |
| CD630_22950 | CD630_n00600 | 124 -- 136 | 351 -- 363 | -13.5328 | 3.96883 | 5.31146 |
| CD630_19680 | CD630_n00600 | 88 -- 142 | 560 -- 624 | -13.5262 | 3.58080 | 5.31146 |
| cobC | CD630_n00600 | 168 -- 197 | 675 -- 707 | -13.5181 | 3.15854 | 5.31146 |
| CD630_28550 | CD630_n00600 | 77 -- 110 | 410 -- 445 | -13.4963 | 3.51196 | 5.31146 |
| CD630_24591 | CD630_n00600 | 85 -- 108 | 546 -- 571 | -13.4945 | 3.54497 | 5.31146 |
| CD630_06150 | CD630_n00600 | 87 -- 142 | 714 -- 752 | -13.4416 | 5.51744 | 5.31146 |
| serS1 | CD630_n00600 | 115 -- 169 | 792 -- 834 | -13.4037 | 3.45858 | 5.31146 |
| CD630_08130 | CD630_n00600 | 31 -- 143 | 537 -- 624 | -13.4016 | 4.63831 | 5.31146 |
| CD630_31220 | CD630_n00600 | 65 -- 83 | 38 -- 55 | -13.3933 | 3.73779 | 5.31146 |
| CD630_17960 | CD630_n00600 | 82 -- 150 | 346 -- 425 | -13.3711 | 3.71404 | 5.31146 |
| dapB2 | CD630_n00600 | 88 -- 186 | 663 -- 753 | -13.358 | 3.27218 | 5.31146 |
| ppnK | CD630_n00600 | 81 -- 103 | 472 -- 495 | -13.356 | 4.36817 | 5.31146 |
| CD630_06100 | CD630_n00600 | 144 -- 201 | 33 -- 89 | -13.3494 | 4.43918 | 5.31146 |
| CD630_32700 | CD630_n00600 | 154 -- 201 | 525 -- 569 | -13.3273 | 4.74276 | 5.31146 |
| CD630_06560 | CD630_n00600 | 109 -- 179 | 480 -- 543 | -13.3137 | 4.02532 | 5.31146 |
| CD630_15200 | CD630_n00600 | 4 -- 135 | 1019 -- 1162 | -13.3125 | 4.00442 | 5.31146 |
| CD630_21090 | CD630_n00600 | 146 -- 174 | 121 -- 145 | -13.2863 | 4.15803 | 5.31146 |
| CD630_34530 | CD630_n00600 | 1 -- 72 | 320 -- 377 | -13.2673 | 3.83016 | 5.31146 |
| CD630_18970 | CD630_n00600 | 117 -- 161 | 318 -- 368 | -13.2235 | 3.57243 | 5.31146 |
| CD630_28181 | CD630_n00600 | 38 -- 77 | 508 -- 550 | -13.2205 | 3.91299 | 5.31146 |
| CD630_19690 | CD630_n00600 | 40 -- 131 | 727 -- 816 | -13.2096 | 3.85861 | 5.31146 |
| CD630_33190 | CD630_n00600 | 56 -- 93 | 224 -- 256 | -13.1742 | 4.13899 | 5.31146 |
| CD630_34520 | CD630_n00600 | 185 -- 200 | 335 -- 353 | -13.173 | 3.83016 | 5.31146 |
| CD630_08370 | CD630_n00600 | 17 -- 29 | 743 -- 755 | -13.1569 | 3.65639 | 5.31146 |
| CD630_20670 | CD630_n00600 | 55 -- 200 | 1056 -- 1172 | -13.1012 | 4.42757 | 5.31146 |
| CD630_13380 | CD630_n00600 | 116 -- 142 | 419 -- 445 | -13.096 | 4.74408 | 5.31146 |
| CD630_31580 | CD630_n00600 | 122 -- 164 | 1057 -- 1095 | -13.0774 | 3.51756 | 5.31146 |
| CD630_22550 | CD630_n00600 | 28 -- 72 | 128 -- 173 | -13.0754 | 3.62379 | 5.31146 |
| CD630_10000 | CD630_n00600 | 98 -- 151 | 524 -- 567 | -13.0688 | 3.86040 | 5.31146 |
| CD630_31461 | CD630_n00600 | 112 -- 191 | 730 -- 819 | -12.9806 | 4.41476 | 5.31146 |
| CD630_21330 | CD630_n00600 | 140 -- 167 | 1136 -- 1164 | -12.934 | 4.10111 | 5.31146 |
| CD630_05280 | CD630_n00600 | 42 -- 92 | 746 -- 813 | -12.9027 | 3.91665 | 5.31146 |
| CD630_23780 | CD630_n00600 | 39 -- 107 | 87 -- 152 | -12.8589 | 3.91629 | 5.31146 |
| CD630_21140 | CD630_n00600 | 147 -- 178 | 481 -- 515 | -12.8587 | 3.33304 | 5.31146 |
| CD630_22740 | CD630_n00600 | 67 -- 87 | 670 -- 691 | -12.8375 | 3.08733 | 5.31146 |
| CD630_06540 | CD630_n00600 | 62 -- 120 | 510 -- 566 | -12.8082 | 3.54673 | 5.31146 |
| spoVFB | CD630_n00600 | 175 -- 201 | 795 -- 821 | -12.7916 | 4.62758 | 5.31146 |
| CD630_28260 | CD630_n00600 | 39 -- 78 | 503 -- 550 | -12.7634 | 3.51613 | 5.31146 |
| CD630_28320 | CD630_n00600 | 161 -- 197 | 793 -- 824 | -12.7248 | 3.99184 | 5.31146 |
| CD630_26750 | CD630_n00600 | 104 -- 152 | 660 -- 708 | -12.7011 | 3.60610 | 5.31146 |
| dltD | CD630_n00600 | 88 -- 119 | 520 -- 551 | -12.6692 | 4.40777 | 5.31146 |
| cdu2 | CD630_n00600 | 39 -- 74 | 671 -- 705 | -12.6557 | 2.97335 | 5.31146 |
| appF | CD630_n00600 | 49 -- 122 | 525 -- 604 | -12.6498 | 4.25570 | 5.31146 |
| CD630_23440 | CD630_n00600 | 182 -- 196 | 794 -- 809 | -12.6375 | 3.34841 | 5.31146 |
| CD630_27230 | CD630_n00600 | 112 -- 142 | 513 -- 548 | -12.6075 | 3.52462 | 5.31146 |
| CD630_06170 | CD630_n00600 | 65 -- 113 | 580 -- 637 | -12.5894 | 4.95366 | 5.31146 |
| CD630_27770 | CD630_n00600 | 50 -- 79 | 349 -- 378 | -12.5861 | 4.08588 | 5.31146 |
| CD630_05870 | CD630_n00600 | 27 -- 59 | 518 -- 551 | -12.5332 | 4.28339 | 5.31146 |
| pilD | CD630_n00600 | 23 -- 81 | 227 -- 273 | -12.4945 | 4.54123 | 5.31146 |
| CD630_25310 | CD630_n00600 | 84 -- 144 | 513 -- 570 | -12.4765 | 3.99861 | 5.31146 |
| CD630_07430 | CD630_n00600 | 83 -- 154 | 1003 -- 1070 | -12.462 | 4.65568 | 5.31146 |
| CD630_14620 | CD630_n00600 | 183 -- 194 | 540 -- 551 | -12.4601 | 4.43651 | 5.31146 |
| CD630_10950 | CD630_n00600 | 168 -- 201 | 47 -- 83 | -12.4592 | 3.67675 | 5.31146 |
| CD630_32060 | CD630_n00600 | 20 -- 61 | 793 -- 832 | -12.4552 | 5.73797 | 5.31146 |
| CD630_21540 | CD630_n00600 | 14 -- 94 | 672 -- 745 | -12.4512 | 4.33090 | 5.31146 |
| CD630_28150 | CD630_n00600 | 151 -- 201 | 335 -- 370 | -12.4504 | 4.18447 | 5.31146 |
| CD630_21020 | CD630_n00600 | 40 -- 95 | 670 -- 719 | -12.4363 | 4.28666 | 5.31146 |
| CD630_14700 | CD630_n00600 | 21 -- 59 | 863 -- 903 | -12.4356 | 3.98234 | 5.31146 |
| CD630_21700 | CD630_n00600 | 37 -- 60 | 1119 -- 1141 | -12.4279 | 3.96650 | 5.31146 |
| CD630_17151 | CD630_n00600 | 1 -- 54 | 495 -- 544 | -12.3821 | 6.43124 | 5.31146 |
| nrdD | CD630_n00600 | 51 -- 105 | 745 -- 807 | -12.3758 | 3.28290 | 5.31146 |
| CD630_33420 | CD630_n00600 | 7 -- 42 | 1120 -- 1161 | -12.3143 | 3.22095 | 5.31146 |
| dnaC | CD630_n00600 | 129 -- 160 | 664 -- 697 | -12.3111 | 6.30796 | 5.31146 |
| pmi | CD630_n00600 | 176 -- 200 | 544 -- 574 | -12.2897 | 3.86845 | 5.31146 |
| CD630_15080 | CD630_n00600 | 48 -- 95 | 530 -- 572 | -12.2877 | 4.03033 | 5.31146 |
| CD630_07121 | CD630_n00600 | 26 -- 42 | 680 -- 695 | -12.283 | 3.26177 | 5.31146 |
| CD630_06920 | CD630_n00600 | 102 -- 135 | 800 -- 835 | -12.2599 | 3.13260 | 5.31146 |
| CD630_21730 | CD630_n00600 | 155 -- 198 | 727 -- 768 | -12.2506 | 3.45076 | 5.31146 |
| CD630_21460 | CD630_n00600 | 44 -- 55 | 1061 -- 1072 | -12.2411 | 4.83005 | 5.31146 |
| CD630_14810 | CD630_n00600 | 112 -- 153 | 798 -- 825 | -12.2291 | 3.35323 | 5.31146 |
| CD630_03420 | CD630_n00600 | 83 -- 196 | 528 -- 610 | -12.2205 | 4.71868 | 5.31146 |
| pgm | CD630_n00600 | 112 -- 159 | 191 -- 244 | -12.202 | 3.45439 | 5.31146 |
| CD630_21230 | CD630_n00600 | 48 -- 67 | 677 -- 696 | -12.192 | 3.89213 | 5.31146 |
| cdu1 | CD630_n00600 | 84 -- 148 | 500 -- 562 | -12.1895 | 4.46276 | 5.31146 |
| CD630_21750 | CD630_n00600 | 49 -- 109 | 125 -- 196 | -12.1632 | 3.70626 | 5.31146 |
| CD630_02790 | CD630_n00600 | 87 -- 164 | 763 -- 816 | -12.1366 | 4.77661 | 5.31146 |
| CD630_17170 | CD630_n00600 | 64 -- 169 | 538 -- 641 | -12.1347 | 4.10230 | 5.31146 |
| CD630_25040 | CD630_n00600 | 12 -- 47 | 670 -- 698 | -12.1054 | 3.43898 | 5.31146 |
| terD | CD630_n00600 | 157 -- 195 | 596 -- 639 | -12.1014 | 3.33148 | 5.31146 |
| CD630_04341 | CD630_n00600 | 158 -- 192 | 515 -- 550 | -12.0876 | 4.29262 | 5.31146 |
| CD630_05490 | CD630_n00600 | 112 -- 139 | 480 -- 506 | -12.0816 | 3.89852 | 5.31146 |
| CD630_06080 | CD630_n00600 | 102 -- 167 | 533 -- 606 | -12.0811 | 3.82315 | 5.31146 |
| CD630_29840 | CD630_n00600 | 25 -- 52 | 730 -- 756 | -12.08 | 4.48825 | 5.31146 |
| CD630_14240 | CD630_n00600 | 75 -- 117 | 669 -- 712 | -12.0464 | 8.60766 | 5.31146 |
| CD630_27270 | CD630_n00600 | 144 -- 173 | 103 -- 131 | -12.0422 | 3.50892 | 5.31146 |
| CD630_09990 | CD630_n00600 | 78 -- 110 | 529 -- 560 | -12.0335 | 3.86040 | 5.31146 |
| CD630_30310 | CD630_n00600 | 183 -- 198 | 341 -- 357 | -12.0275 | 4.56059 | 5.31146 |
| CD630_10800 | CD630_n00600 | 89 -- 114 | 531 -- 551 | -12.001 | 5.30319 | 5.31146 |
| CD630_30360 | CD630_n00600 | 158 -- 169 | 122 -- 133 | -12.0003 | 4.26509 | 5.31146 |
| CD630_27060 | CD630_n00600 | 69 -- 119 | 333 -- 380 | -11.9891 | 4.06049 | 5.31146 |
| cstA | CD630_n00600 | 85 -- 97 | 671 -- 683 | -11.9832 | 3.53584 | 5.31146 |
| CD630_29960 | CD630_n00600 | 131 -- 201 | 528 -- 614 | -11.9702 | 4.51904 | 5.31146 |
| CD630_05030 | CD630_n00600 | 79 -- 90 | 547 -- 558 | -11.9626 | 4.03823 | 5.31146 |
| CD630_27610 | CD630_n00600 | 19 -- 42 | 1150 -- 1171 | -11.9597 | 3.96465 | 5.31146 |
| CD630_06050 | CD630_n00600 | 46 -- 193 | 538 -- 681 | -11.9185 | 3.59332 | 5.31146 |
| CD630_15740 | CD630_n00600 | 90 -- 146 | 320 -- 366 | -11.9151 | 4.43222 | 5.31146 |
| uxaA%27 | CD630_n00600 | 87 -- 112 | 730 -- 751 | -11.905 | 4.26165 | 5.31146 |
| CD630_14182 | CD630_n00600 | 116 -- 143 | 781 -- 810 | -11.8992 | 4.82360 | 5.31146 |
| CD630_21070 | CD630_n00600 | 176 -- 192 | 359 -- 375 | -11.893 | 5.35019 | 5.31146 |
| CD630_22500 | CD630_n00600 | 117 -- 156 | 1 -- 37 | -11.8923 | 4.03688 | 5.31146 |
| CD630_34030 | CD630_n00600 | 23 -- 58 | 296 -- 331 | -11.8771 | 3.05766 | 5.31146 |
| moaA | CD630_n00600 | 26 -- 122 | 530 -- 615 | -11.8697 | 3.83799 | 5.31146 |
| CD630_18650 | CD630_n00600 | 143 -- 157 | 261 -- 275 | -11.8549 | 3.75892 | 5.31146 |
| CD630_05821 | CD630_n00600 | 61 -- 90 | 414 -- 442 | -11.8445 | 3.75661 | 5.31146 |
| CD630_09970 | CD630_n00600 | 166 -- 177 | 426 -- 438 | -11.8441 | 3.25917 | 5.31146 |
| CD630_18440 | CD630_n00600 | 168 -- 183 | 351 -- 365 | -11.8331 | 4.07960 | 5.31146 |
| bcp | CD630_n00600 | 58 -- 187 | 478 -- 584 | -11.8188 | 3.16670 | 5.31146 |
| CD630_04610 | CD630_n00600 | 21 -- 46 | 531 -- 561 | -11.8172 | 3.83025 | 5.31146 |
| CD630_15060 | CD630_n00600 | 42 -- 76 | 1074 -- 1111 | -11.8066 | 3.69396 | 5.31146 |
| CD630_10960 | CD630_n00600 | 11 -- 30 | 863 -- 882 | -11.8002 | 3.53850 | 5.31146 |
| cobS | CD630_n00600 | 85 -- 174 | 349 -- 430 | -11.7935 | 3.67481 | 5.31146 |
| cwp26 | CD630_n00600 | 56 -- 89 | 1056 -- 1087 | -11.79 | 4.44377 | 5.31146 |
| bltD | CD630_n00600 | 192 -- 200 | 415 -- 423 | -11.7879 | 4.49926 | 5.31146 |
| CD630_33480 | CD630_n00600 | 57 -- 98 | 529 -- 564 | -11.7814 | 3.67613 | 5.31146 |
| CD630_30110 | CD630_n00600 | 140 -- 190 | 1037 -- 1095 | -11.7766 | 3.31041 | 5.31146 |
| CD630_23280 | CD630_n00600 | 113 -- 169 | 520 -- 571 | -11.7735 | 3.50093 | 5.31146 |
| CD630_10080 | CD630_n00600 | 43 -- 102 | 662 -- 720 | -11.7661 | 3.75625 | 5.31146 |
| opuCC | CD630_n00600 | 169 -- 199 | 34 -- 60 | -11.7595 | 4.07035 | 5.31146 |
| CD630_25300 | CD630_n00600 | 138 -- 161 | 859 -- 880 | -11.7475 | 4.49333 | 5.31146 |
| CD630_06060 | CD630_n00600 | 100 -- 130 | 405 -- 435 | -11.7419 | 4.94050 | 5.31146 |
| CD630_27210 | CD630_n00600 | 31 -- 85 | 493 -- 550 | -11.7362 | 4.18786 | 5.31146 |
| gatC | CD630_n00600 | 73 -- 147 | 304 -- 378 | -11.7305 | 4.25143 | 5.31146 |
| CD630_23820 | CD630_n00600 | 27 -- 158 | 752 -- 867 | -11.7281 | 3.18419 | 5.31146 |
| CD630_20450 | CD630_n00600 | 61 -- 108 | 731 -- 770 | -11.7124 | 3.77388 | 5.31146 |
| CD630_08070 | CD630_n00600 | 157 -- 187 | 521 -- 550 | -11.7043 | 4.87819 | 5.31146 |
| CD630_15461 | CD630_n00600 | 169 -- 178 | 271 -- 280 | -11.7008 | 9.13442 | 5.31146 |
| adhE | CD630_n00600 | 165 -- 196 | 513 -- 544 | -11.6956 | 3.06485 | 5.31146 |
| CD630_03400 | CD630_n00600 | 50 -- 144 | 157 -- 240 | -11.6786 | 4.45994 | 5.31146 |
| CD630_19430 | CD630_n00600 | 74 -- 101 | 344 -- 377 | -11.6748 | 3.13922 | 5.31146 |
| CD630_23080 | CD630_n00600 | 157 -- 197 | 400 -- 440 | -11.6508 | 5.23027 | 5.31146 |
| oppC | CD630_n00600 | 59 -- 188 | 303 -- 442 | -11.6359 | 3.01695 | 5.31146 |
| CD630_20170 | CD630_n00600 | 98 -- 122 | 671 -- 694 | -11.6323 | 6.90201 | 5.31146 |
| rolA | CD630_n00600 | 1 -- 50 | 356 -- 404 | -11.6197 | 4.56362 | 5.31146 |
| CD630_19780 | CD630_n00600 | 168 -- 187 | 1119 -- 1139 | -11.6182 | 3.64830 | 5.31146 |
| CD630_14360 | CD630_n00600 | 59 -- 103 | 600 -- 647 | -11.606 | 3.02079 | 5.31146 |
| CD630_22880 | CD630_n00600 | 52 -- 83 | 25 -- 56 | -11.5861 | 3.62455 | 5.31146 |
| CD630_08360 | CD630_n00600 | 4 -- 38 | 801 -- 832 | -11.5779 | 3.19339 | 5.31146 |
| CD630_07070 | CD630_n00600 | 29 -- 52 | 182 -- 205 | -11.5742 | 3.46414 | 5.31146 |
| CD630_05230 | CD630_n00600 | 184 -- 200 | 360 -- 376 | -11.5738 | 3.73787 | 5.31146 |
| CD630_32990 | CD630_n00600 | 178 -- 199 | 62 -- 82 | -11.5564 | 5.24925 | 5.31146 |
| CD630_29270 | CD630_n00600 | 2 -- 16 | 361 -- 375 | -11.5521 | 3.70174 | 5.31146 |
| CD630_06350 | CD630_n00600 | 65 -- 103 | 811 -- 842 | -11.5486 | 4.35983 | 5.31146 |
| CD630_08940 | CD630_n00600 | 45 -- 125 | 727 -- 788 | -11.5292 | 3.72225 | 5.31146 |
| CD630_17360 | CD630_n00600 | 5 -- 40 | 185 -- 210 | -11.5239 | 5.66070 | 5.31146 |
| CD630_01590 | CD630_n00600 | 77 -- 109 | 667 -- 695 | -11.5158 | 3.45609 | 5.31146 |
| CD630_36190 | CD630_n00600 | 77 -- 91 | 813 -- 828 | -11.5129 | 4.63449 | 5.31146 |
| ung | CD630_n00600 | 114 -- 164 | 520 -- 562 | -11.4759 | 3.30908 | 5.31146 |
| CD630_06340 | CD630_n00600 | 36 -- 72 | 515 -- 558 | -11.4721 | 4.35983 | 5.31146 |
| CD630_13400 | CD630_n00600 | 153 -- 163 | 1067 -- 1077 | -11.471 | 3.20232 | 5.31146 |
| CD630_14630 | CD630_n00600 | 107 -- 178 | 337 -- 388 | -11.4323 | 3.34485 | 5.31146 |
| fliI | CD630_n00600 | 90 -- 186 | 1055 -- 1163 | -11.4253 | 3.03666 | 5.31146 |
| CD630_31970 | CD630_n00600 | 107 -- 135 | 117 -- 145 | -11.4172 | 3.24511 | 5.31146 |
| CD630_27200 | CD630_n00600 | 1 -- 50 | 666 -- 713 | -11.4043 | 3.42516 | 5.31146 |
| CD630_16160 | CD630_n00600 | 29 -- 63 | 328 -- 365 | -11.3986 | 6.11801 | 5.31146 |
| CD630_22850 | CD630_n00600 | 177 -- 191 | 546 -- 561 | -11.3907 | 3.84102 | 5.31146 |
| kdpE | CD630_n00600 | 25 -- 36 | 283 -- 294 | -11.3836 | 4.27140 | 5.31146 |
| dpaA | CD630_n00600 | 21 -- 54 | 106 -- 140 | -11.368 | 4.65364 | 5.31146 |
| pilC | CD630_n00600 | 4 -- 33 | 527 -- 557 | -11.3652 | 3.54230 | 5.31146 |
| CD630_29750 | CD630_n00600 | 84 -- 95 | 798 -- 809 | -11.3556 | 4.59146 | 5.31146 |
| CD630_20260 | CD630_n00600 | 84 -- 143 | 559 -- 618 | -11.3431 | 3.84162 | 5.31146 |
| CD630_05150 | CD630_n00600 | 81 -- 109 | 530 -- 564 | -11.3263 | 3.69875 | 5.31146 |
| CD630_20660 | CD630_n00600 | 77 -- 99 | 1065 -- 1095 | -11.3217 | 5.39321 | 5.31146 |
| CD630_07840 | CD630_n00600 | 81 -- 98 | 792 -- 806 | -11.3069 | 4.26761 | 5.31146 |
| CD630_14130 | CD630_n00600 | 61 -- 196 | 596 -- 739 | -11.2902 | 4.57970 | 5.31146 |
| CD630_15550 | CD630_n00600 | 173 -- 181 | 422 -- 430 | -11.2471 | 3.49019 | 5.31146 |
| CD630_05370 | CD630_n00600 | 79 -- 106 | 670 -- 688 | -11.2449 | 3.26947 | 5.31146 |
| CD630_06960 | CD630_n00600 | 134 -- 146 | 556 -- 568 | -11.2402 | 3.16317 | 5.31146 |
| CD630_26691 | CD630_n00600 | 1 -- 69 | 1084 -- 1145 | -11.2319 | 3.96230 | 5.31146 |
| CD630_01690 | CD630_n00600 | 174 -- 198 | 59 -- 83 | -11.1847 | 4.97234 | 5.31146 |
| CD630_19440 | CD630_n00600 | 92 -- 106 | 52 -- 66 | -11.1847 | 4.53937 | 5.31146 |
| CD630_20980 | CD630_n00600 | 85 -- 116 | 727 -- 753 | -11.1731 | 8.12847 | 5.31146 |
| eutW | CD630_n00600 | 1 -- 38 | 595 -- 636 | -11.171 | 4.55715 | 5.31146 |
| CD630_01570 | CD630_n00600 | 132 -- 155 | 25 -- 47 | -11.1464 | 6.08699 | 5.31146 |
| CD630_20541 | CD630_n00600 | 39 -- 58 | 795 -- 815 | -11.1066 | 5.74527 | 5.31146 |
| CD630_05590 | CD630_n00600 | 67 -- 124 | 399 -- 445 | -11.0965 | 3.39670 | 5.31146 |
| CD630_06970 | CD630_n00600 | 26 -- 62 | 677 -- 707 | -11.0952 | 3.82312 | 5.31146 |
| CD630_18410 | CD630_n00600 | 123 -- 168 | 123 -- 168 | -11.07 | 3.59739 | 5.31146 |
| CD630_17900 | CD630_n00600 | 98 -- 159 | 1106 -- 1156 | -11.0626 | 3.52167 | 5.31146 |
| CD630_05460 | CD630_n00600 | 77 -- 114 | 804 -- 837 | -11.0433 | 3.98753 | 5.31146 |
| CD630_26630 | CD630_n00600 | 65 -- 113 | 350 -- 394 | -11.0343 | 4.61557 | 5.31146 |
| CD630_14410 | CD630_n00600 | 97 -- 117 | 532 -- 550 | -11.0224 | 3.39433 | 5.31146 |
| CD630_08140 | CD630_n00600 | 84 -- 106 | 670 -- 688 | -11.0107 | 3.30159 | 5.31146 |
| CD630_29650 | CD630_n00600 | 119 -- 144 | 123 -- 148 | -11.0083 | 3.67128 | 5.31146 |
| dnaD | CD630_n00600 | 7 -- 27 | 519 -- 538 | -11.0036 | 3.56078 | 5.31146 |
| CD630_04540 | CD630_n00600 | 62 -- 92 | 677 -- 710 | -10.9793 | 3.33556 | 5.31146 |
| CD630_10220 | CD630_n00600 | 80 -- 103 | 666 -- 688 | -10.9528 | 4.08233 | 5.31146 |
| CD630_19580 | CD630_n00600 | 70 -- 102 | 518 -- 550 | -10.9517 | 4.51902 | 5.31146 |
| CD630_18090 | CD630_n00600 | 49 -- 69 | 1119 -- 1142 | -10.9169 | 4.23961 | 5.31146 |
| CD630_13020 | CD630_n00600 | 120 -- 173 | 28 -- 64 | -10.9023 | 3.52267 | 5.31146 |
| CD630_26050 | CD630_n00600 | 155 -- 193 | 324 -- 364 | -10.8927 | 6.99851 | 5.31146 |
| CD630_11241 | CD630_n00600 | 161 -- 186 | 93 -- 116 | -10.8813 | 4.85723 | 5.31146 |
| CD630_05900 | CD630_n00600 | 19 -- 63 | 23 -- 54 | -10.8722 | 3.21934 | 5.31146 |
| CD630_25070 | CD630_n00600 | 150 -- 160 | 1058 -- 1068 | -10.8555 | 3.45164 | 5.31146 |
| CD630_22710 | CD630_n00600 | 104 -- 148 | 1121 -- 1170 | -10.852 | 4.05623 | 5.31146 |

| | | | | | | |
|---|---|---|---|---|---|---|
| CD630_18680 | CD630_n00600 | 97 – 146 | 526 – 568 | -10.8491 | 3.04315 | 5.31146 |
| gcp | CD630_n00600 | 20 – 75 | 524 – 573 | -10.8487 | 4.92798 | 5.31146 |
| CD630_17630 | CD630_n00600 | 23 – 50 | 120 – 147 | -10.848 | 4.52408 | 5.31146 |
| CD630_06060 | CD630_n00600 | 35 – 96 | 517 – 574 | -10.8323 | 4.94050 | 5.31146 |
| CD630_23930 | CD630_n00600 | 129 – 142 | 757 – 770 | -10.8303 | 3.05650 | 5.31146 |
| CD630_03170 | CD630_n00600 | 2 – 36 | 996 – 1031 | -10.816 | 3.66946 | 5.31146 |
| CD630_04940 | CD630_n00600 | 131 – 187 | 327 – 375 | -10.801 | 4.20948 | 5.31146 |
| CD630_15431 | CD630_n00600 | 139 – 179 | 539 – 578 | -10.7886 | 5.29780 | 5.31146 |
| CD630_26680 | CD630_n00600 | 1 – 8 | 421 – 428 | -10.7877 | 3.53956 | 5.31146 |
| CD630_05820 | CD630_n00600 | 190 – 197 | 421 – 428 | -10.7568 | 3.75661 | 5.31146 |
| CD630_13880 | CD630_n00600 | 105 – 158 | 105 – 149 | -10.7322 | 3.52735 | 5.31146 |
| CD630_20920 | CD630_n00600 | 89 – 136 | 797 – 843 | -10.7201 | 4.13812 | 5.31146 |
| srlD | CD630_n00600 | 88 – 146 | 522 – 566 | -10.7191 | 4.19843 | 5.31146 |
| CD630_34050 | CD630_n00600 | 4 – 21 | 930 – 947 | -10.7141 | 4.03863 | 5.31146 |
| CD630_05160 | CD630_n00600 | 79 – 109 | 678 – 716 | -10.7114 | 5.03573 | 5.31146 |
| CD630_12110 | CD630_n00600 | 161 – 174 | 625 – 638 | -10.6987 | 4.36078 | 5.31146 |
| CD630_06190 | CD630_n00600 | 37 – 95 | 755 – 814 | -10.6984 | 3.57064 | 5.31146 |
| CD630_08110 | CD630_n00600 | 64 – 130 | 498 – 576 | -10.6726 | 4.39181 | 5.31146 |
| fbp | CD630_n00600 | 143 – 184 | 119 – 163 | -10.6723 | 3.80071 | 5.31146 |
| CD630_36180 | CD630_n00600 | 5 – 57 | 1006 – 1051 | -10.6513 | 4.83428 | 5.31146 |
| CD630_18610 | CD630_n00600 | 108 – 150 | 389 – 426 | -10.6457 | 4.15582 | 5.31146 |
| CD630_06690 | CD630_n00600 | 88 – 106 | 199 – 215 | -10.6441 | 5.82709 | 5.31146 |
| CD630_21260 | CD630_n00600 | 117 – 165 | 38 – 88 | -10.6391 | 4.22255 | 5.31146 |
| vncR | CD630_n00600 | 74 – 94 | 796 – 814 | -10.6309 | 3.93642 | 5.31146 |
| CD630_16940 | CD630_n00600 | 128 – 181 | 825 – 872 | -10.6162 | 4.14665 | 5.31146 |
| CD630_20600 | CD630_n00600 | 36 – 56 | 119 – 139 | -10.6157 | 4.20093 | 5.31146 |
| CD630_22720 | CD630_n00600 | 120 – 157 | 539 – 567 | -10.6135 | 3.81804 | 5.31146 |
| CD630_22660 | CD630_n00600 | 161 – 201 | 1063 – 1100 | -10.6068 | 3.60496 | 5.31146 |
| CD630_08780 | CD630_n00600 | 148 – 197 | 1005 – 1048 | -10.595 | 7.76690 | 5.31146 |
| kdgT1 | CD630_n00600 | 180 – 199 | 125 – 149 | -10.5946 | 4.49207 | 5.31146 |
| CD630_16750 | CD630_n00600 | 87 – 157 | 751 – 818 | -10.5782 | 5.04051 | 5.31146 |
| CD630_21720 | CD630_n00600 | 164 – 188 | 793 – 820 | -10.5728 | 4.13030 | 5.31146 |
| CD630_15160 | CD630_n00600 | 123 – 153 | 401 – 426 | -10.5714 | 4.32318 | 5.31146 |
| CD630_13920 | CD630_n00600 | 98 – 132 | 540 – 576 | -10.569 | 3.25236 | 5.31146 |
| CD630_18780 | CD630_n00600 | 147 – 170 | 126 – 150 | -10.5681 | 4.71613 | 5.31146 |
| CD630_15111 | CD630_n00600 | 20 – 60 | 1011 – 1048 | -10.5583 | 4.18664 | 5.31146 |
| CD630_14600 | CD630_n00600 | 2 – 103 | 215 – 306 | -10.5463 | 5.90348 | 5.31146 |
| CD630_28330 | CD630_n00600 | 39 – 68 | 484 – 512 | -10.5337 | 3.29122 | 5.31146 |
| CD630_06140 | CD630_n00600 | 35 – 77 | 53 – 92 | -10.5281 | 4.12519 | 5.31146 |
| CD630_05290 | CD630_n00600 | 86 – 94 | 810 – 818 | -10.5279 | 3.97604 | 5.31146 |
| tcdC | CD630_n00600 | 123 – 155 | 337 – 366 | -10.5138 | 3.76101 | 5.31146 |
| CD630_04880 | CD630_n00600 | 68 – 143 | 674 – 753 | -10.4755 | 4.62291 | 5.31146 |
| CD630_06642 | CD630_n00600 | 106 – 178 | 37 – 112 | -10.474 | 3.32967 | 5.31146 |
| CD630_31960 | CD630_n00600 | 107 – 129 | 1059 – 1078 | -10.4685 | 4.93757 | 5.31146 |
| CD630_19850 | CD630_n00600 | 6 – 65 | 621 – 684 | -10.4658 | 6.88486 | 5.31146 |
| CD630_05830 | CD630_n00600 | 20 – 61 | 916 – 959 | -10.4607 | 3.35360 | 5.31146 |
| CD630_14140 | CD630_n00600 | 51 – 144 | 75 – 155 | -10.4405 | 3.16551 | 5.31146 |
| CD630_01570 | CD630_n00600 | 146 – 159 | 555 – 568 | -10.44 | 6.08699 | 5.31146 |
| CD630_18230 | CD630_n00600 | 85 – 190 | 320 – 430 | -10.4354 | 3.00184 | 5.31146 |
| CD630_23000 | CD630_n00600 | 147 – 190 | 788 – 819 | -10.4213 | 3.54130 | 5.31146 |
| moeA | CD630_n00600 | 49 – 124 | 320 – 395 | -10.4128 | 3.26118 | 5.31146 |
| CD630_07470 | CD630_n00600 | 10 – 64 | 350 – 407 | -10.3829 | 3.37143 | 5.31146 |
| CD630_31530 | CD630_n00600 | 137 – 172 | 118 – 169 | -10.37 | 3.49100 | 5.31146 |
| CD630_23070 | CD630_n00600 | 167 – 192 | 38 – 66 | -10.3652 | 5.23027 | 5.31146 |
| CD630_07870 | CD630_n00600 | 156 – 185 | 285 – 311 | -10.3589 | 3.22829 | 5.31146 |
| CD630_08510 | CD630_n00600 | 126 – 147 | 126 – 149 | -10.3456 | 5.16329 | 5.31146 |
| CD630_07960 | CD630_n00600 | 66 – 106 | 322 – 366 | -10.3288 | 3.22537 | 5.31146 |
| CD630_16670 | CD630_n00600 | 116 – 143 | 347 – 375 | -10.3243 | 3.33553 | 5.31146 |
| CD630_06110 | CD630_n00600 | 167 – 187 | 795 – 816 | -10.296 | 5.61380 | 5.31146 |
| CD630_13960 | CD630_n00600 | 121 – 161 | 670 – 709 | -10.2873 | 3.30375 | 5.31146 |
| thiH | CD630_n00600 | 47 – 98 | 552 – 605 | -10.2802 | 3.58230 | 5.31146 |
| CD630_17730 | CD630_n00600 | 64 – 106 | 595 – 628 | -10.2733 | 4.73397 | 5.31146 |
| CD630_19670 | CD630_n00600 | 72 – 96 | 533 – 565 | -10.2699 | 3.20718 | 5.31146 |
| glnA | CD630_n00600 | 28 – 49 | 590 – 611 | -10.2667 | 3.87629 | 5.31146 |
| CD630_09030 | CD630_n00600 | 79 – 149 | 1001 – 1068 | -10.2502 | 4.21282 | 5.31146 |
| CD630_25020 | CD630_n00600 | 119 – 150 | 536 – 571 | -10.25 | 4.17721 | 5.31146 |
| CD630_17281 | CD630_n00600 | 3 – 37 | 321 – 354 | -10.2455 | 5.91544 | 5.31146 |
| CD630_13930 | CD630_n00600 | 90 – 110 | 733 – 751 | -10.2439 | 4.09246 | 5.31146 |
| CD630_06380 | CD630_n00600 | 1 – 22 | 666 – 690 | -10.2335 | 4.32211 | 5.31146 |
| CD630_05211 | CD630_n00600 | 84 – 188 | 480 – 578 | -10.2198 | 3.97048 | 5.31146 |
| CD630_17292 | CD630_n00600 | 143 – 190 | 608 – 641 | -10.2038 | 6.25799 | 5.31146 |
| CD630_14990 | CD630_n00600 | 94 – 112 | 545 – 562 | -10.2025 | 3.78619 | 5.31146 |
| CD630_32110 | CD630_n00600 | 98 – 188 | 120 – 195 | -10.1977 | 4.45291 | 5.31146 |
| CD630_31480 | CD630_n00600 | 9 – 22 | 185 – 198 | -10.1913 | 4.88897 | 5.31146 |
| CD630_18910 | CD630_n00600 | 25 – 86 | 1051 – 1116 | -10.1884 | 3.30688 | 5.31146 |
| CD630_30620 | CD630_n00600 | 9 – 38 | 753 – 784 | -10.1628 | 4.15443 | 5.31146 |
| CD630_17160 | CD630_n00600 | 85 – 114 | 540 – 570 | -10.1538 | 3.50228 | 5.31146 |
| CD630_12320 | CD630_n00600 | 38 – 76 | 213 – 241 | -10.1481 | 3.28667 | 5.31146 |
| cynT | CD630_n00600 | 106 – 126 | 816 – 834 | -10.1427 | 4.21370 | 5.31146 |
| CD630_05840 | CD630_n00600 | 79 – 115 | 1038 – 1069 | -10.1372 | 4.28779 | 5.31146 |
| ogt1 | CD630_n00600 | 30 – 45 | 476 – 492 | -10.1137 | 3.16772 | 5.31146 |
| CD630_27310 | CD630_n00600 | 45 – 66 | 804 – 816 | -10.1081 | 4.01622 | 5.31146 |
| CD630_21030 | CD630_n00600 | 28 – 74 | 97 – 153 | -10.1061 | 3.08733 | 5.31146 |
| CD630_14420 | CD630_n00600 | 81 – 96 | 226 – 241 | -10.1049 | 3.99497 | 5.31146 |
| aroD | CD630_n00600 | 136 – 169 | 397 – 431 | -10.0792 | 3.71929 | 5.31146 |
| panG | CD630_n00600 | 11 – 71 | 159 – 218 | -10.0617 | 5.54047 | 5.31146 |
| CD630_17720 | CD630_n00600 | 145 – 172 | 122 – 150 | -10.0406 | 3.22910 | 5.31146 |
| CD630_14930 | CD630_n00600 | 17 – 40 | 356 – 379 | -10.0322 | 3.31335 | 5.31146 |
| CD630_15900 | CD630_n00600 | 144 – 153 | 286 – 295 | -10.0313 | 3.17664 | 5.31146 |
| CD630_04660 | CD630_n00600 | 30 – 42 | 674 – 685 | -10.0199 | 4.32791 | 5.31146 |
| CD630_09050 | CD630_n00600 | 110 – 137 | 525 – 549 | -10.0187 | 3.96216 | 5.31146 |
| CD630_08080 | CD630_n00600 | 157 – 180 | 187 – 211 | -10.0176 | 4.87819 | 5.31146 |
| CD630_14920 | CD630_n00600 | 101 – 116 | 594 – 609 | -10.0093 | 4.35225 | 5.31146 |
| CD630_30100 | CD630_n00600 | 117 – 155 | 123 – 155 | -9.99338 | 4.13234 | 5.31146 |
| CD630_13600 | CD630_n00600 | 120 – 134 | 539 – 551 | -9.96655 | 4.45481 | 5.31146 |
| CD630_09570 | CD630_n00600 | 22 – 31 | 742 – 751 | -9.93465 | 5.31213 | 5.31146 |
| CD630_29070 | CD630_n00600 | 22 – 31 | 742 – 751 | -9.93465 | 3.89477 | 5.31146 |
| CD630_27240 | CD630_n00600 | 8 – 94 | 673 – 749 | -9.92402 | 3.52462 | 5.31146 |
| ssuC | CD630_n00600 | 47 – 79 | 801 – 824 | -9.92324 | 3.45491 | 5.31146 |
| CD630_26050 | CD630_n00600 | 44 – 83 | 779 – 818 | -9.91856 | 6.99851 | 5.31146 |
| ldh | CD630_n00600 | 129 – 167 | 918 – 959 | -9.89373 | 4.35818 | 5.31146 |
| CD630_28890 | CD630_n00600 | 33 – 52 | 1054 – 1071 | -9.89204 | 7.52469 | 5.31146 |
| CD630_33610 | CD630_n00600 | 131 – 171 | 714 – 767 | -9.88764 | 3.79322 | 5.31146 |
| CD630_03410 | CD630_n00600 | 87 – 112 | 787 – 812 | -9.88602 | 4.45994 | 5.31146 |
| glgP | CD630_n00600 | 30 – 62 | 530 – 561 | -9.88336 | 3.28530 | 5.31146 |
| CD630_09100 | CD630_n00600 | 5 – 51 | 38 – 81 | -9.85314 | 5.92260 | 5.31146 |
| dltA | CD630_n00600 | 119 – 150 | 1063 – 1097 | -9.85149 | 3.85429 | 5.31146 |
| CD630_21790 | CD630_n00600 | 143 – 161 | 1124 – 1145 | -9.84964 | 4.93762 | 5.31146 |
| virS | CD630_n00600 | 144 – 190 | 194 – 239 | -9.84305 | 4.74085 | 5.31146 |
| xylR | CD630_n00600 | 160 – 195 | 800 – 837 | -9.82387 | 3.95671 | 5.31146 |
| CD630_36170 | CD630_n00600 | 10 – 74 | 318 – 377 | -9.80841 | 3.72380 | 5.31146 |
| CD630_16450 | CD630_n00600 | 88 – 109 | 734 – 750 | -9.77663 | 4.57547 | 5.31146 |
| hfq | CD630_n00600 | 66 – 78 | 854 – 866 | -9.77464 | 5.05609 | 5.31146 |
| CD630_30420 | CD630_n00600 | 4 – 45 | 121 – 178 | -9.76044 | 5.27539 | 5.31146 |
| CD630_26150 | CD630_n00600 | 180 – 195 | 933 – 949 | -9.72864 | 4.06302 | 5.31146 |
| CD630_25050 | CD630_n00600 | 181 – 201 | 743 – 762 | -9.72662 | 4.33624 | 5.31146 |
| CD630_22991 | CD630_n00600 | 6 – 45 | 1054 – 1088 | -9.6983 | 6.03728 | 5.31146 |
| CD630_27630 | CD630_n00600 | 41 – 80 | 1148 – 1184 | -9.69507 | 3.54199 | 5.31146 |
| CD630_06200 | CD630_n00600 | 89 – 98 | 741 – 750 | -9.68762 | 3.57064 | 5.31146 |
| CD630_15190 | CD630_n00600 | 146 – 194 | 540 – 574 | -9.68623 | 4.43255 | 5.31146 |
| chrA | CD630_n00600 | 146 – 167 | 347 – 366 | -9.6695 | 5.27255 | 5.31146 |
| CD630_36690 | CD630_n00600 | 54 – 87 | 1056 – 1093 | -9.6367 | 3.53761 | 5.31146 |
| CD630_17350 | CD630_n00600 | 181 – 197 | 621 – 636 | -9.63304 | 3.35334 | 5.31146 |
| CD630_21760 | CD630_n00600 | 27 – 69 | 543 – 576 | -9.62455 | 4.43846 | 5.31146 |
| CD630_03920 | CD630_n00600 | 26 – 34 | 800 – 808 | -9.61427 | 4.15934 | 5.31146 |
| CD630_31690 | CD630_n00600 | 112 – 134 | 19 – 45 | -9.61224 | 3.71193 | 5.31146 |
| ribD | CD630_n00600 | 83 – 92 | 744 – 753 | -9.6103 | 6.65922 | 5.31146 |
| CD630_28280 | CD630_n00600 | 97 – 116 | 1039 – 1060 | -9.58815 | 4.59833 | 5.31146 |
| uxaA | CD630_n00600 | 110 – 125 | 1056 – 1074 | -9.58632 | 4.26165 | 5.31146 |
| CD630_14180 | CD630_n00600 | 179 – 189 | 426 – 437 | -9.58626 | 3.93696 | 5.31146 |
| CD630_21490 | CD630_n00600 | 183 – 201 | 195 – 222 | -9.5829 | 3.37462 | 5.31146 |
| CD630_10790 | CD630_n00600 | 163 – 170 | 360 – 367 | -9.58286 | 4.45284 | 5.31146 |
| cbiO | CD630_n00600 | 89 – 118 | 1136 – 1167 | -9.55082 | 3.79830 | 5.31146 |
| CD630_30730 | CD630_n00600 | 136 – 172 | 1062 – 1107 | -9.54469 | 3.13236 | 5.31146 |
| CD630_21080 | CD630_n00600 | 85 – 113 | 344 – 371 | -9.52495 | 3.99742 | 5.31146 |
| CD630_30120 | CD630_n00600 | 6 – 75 | 135 – 198 | -9.5112 | 4.04859 | 5.31146 |
| CD630_27040 | CD630_n00620 | 174 – 199 | 68 – 94 | -17.9694 | 3.51812 | 5.82838 |
| CD630_14110 | CD630_n00620 | 37 – 177 | 43 – 180 | -17.5443 | 3.39550 | 5.82838 |
| CD630_09100 | CD630_n00620 | 1 – 51 | 46 – 98 | -15.7057 | 5.92260 | 5.82838 |
| sspA | CD630_n00620 | 131 – 161 | 70 – 96 | -15.5022 | 3.62689 | 5.82838 |
| CD630_27310 | CD630_n00620 | 151 – 194 | 135 – 183 | -14.7464 | 4.01622 | 5.82838 |
| eutW | CD630_n00620 | 10 – 28 | 176 – 195 | -14.6369 | 4.55715 | 5.82838 |
| CD630_04850 | CD630_n00620 | 132 – 201 | 154 – 213 | -14.6074 | 4.40788 | 5.82838 |
| CD630_14620 | CD630_n00620 | 8 – 152 | 31 – 177 | -14.5164 | 4.43651 | 5.82838 |
| CD630_22510 | CD630_n00620 | 151 – 201 | 138 – 182 | -14.2384 | 11.42763 | 5.82838 |
| CD630_28290 | CD630_n00620 | 28 – 90 | 117 – 181 | -14.2113 | 5.54192 | 5.82838 |
| CD630_09060 | CD630_n00620 | 52 – 86 | 153 – 188 | -13.8886 | 4.31834 | 5.82838 |
| oppB | CD630_n00620 | 128 – 177 | 53 – 98 | -13.8044 | 2.98328 | 5.82838 |
| CD630_28260 | CD630_n00620 | 142 – 192 | 70 – 120 | -13.5439 | 3.51613 | 5.82838 |
| CD630_25020 | CD630_n00620 | 15 – 36 | 49 – 72 | -13.3608 | 4.17721 | 5.82838 |
| feoB | CD630_n00620 | 8 – 75 | 140 – 204 | -13.323 | 4.52257 | 5.82838 |
| CD630_17292 | CD630_n00620 | 76 – 200 | 88 – 201 | -13.3033 | 6.25799 | 5.82838 |
| CD630_06060 | CD630_n00620 | 36 – 74 | 146 – 181 | -13.2176 | 4.94050 | 5.82838 |
| CD630_27280 | CD630_n00620 | 51 – 196 | 49 – 195 | -13.0916 | 3.24738 | 5.82838 |
| CD630_16530 | CD630_n00620 | 32 – 86 | 51 – 102 | -12.9452 | 4.37658 | 5.82838 |
| etfA | CD630_n00620 | 82 – 134 | 151 – 203 | -12.9374 | 3.41204 | 5.82838 |
| CD630_05950 | CD630_n00620 | 84 – 140 | 156 – 198 | -12.9188 | 3.31669 | 5.82838 |
| CD630_36690 | CD630_n00620 | 11 – 34 | 55 – 77 | -12.7599 | 3.53761 | 5.82838 |
| CD630_23080 | CD630_n00620 | 176 – 198 | 175 – 198 | -12.5227 | 5.23027 | 5.82838 |
| CD630_36170 | CD630_n00620 | 147 – 194 | 152 – 196 | -12.4277 | 3.72380 | 5.82838 |
| CD630_18930 | CD630_n00620 | 135 – 175 | 160 – 197 | -12.4024 | 5.32127 | 5.82838 |
| CD630_31690 | CD630_n00620 | 79 – 172 | 109 – 198 | -12.3065 | 3.71193 | 5.82838 |
| CD630_06170 | CD630_n00620 | 102 – 160 | 51 – 97 | -12.2702 | 4.95366 | 5.82838 |
| CD630_05830 | CD630_n00620 | 123 – 167 | 38 – 98 | -12.2639 | 3.35360 | 5.82838 |
| cls | CD630_n00620 | 143 – 190 | 68 – 96 | -11.9979 | 4.46177 | 5.82838 |
| CD630_27520 | CD630_n00620 | 115 – 154 | 180 – 220 | -11.9462 | 4.27693 | 5.82838 |
| CD630_18781 | CD630_n00620 | 109 – 159 | 152 – 196 | -11.945 | 3.77660 | 5.82838 |
| CD630_32890 | CD630_n00620 | 30 – 165 | 62 – 189 | -11.9247 | 3.41276 | 5.82838 |
| CD630_07870 | CD630_n00620 | 94 – 196 | 95 – 196 | -11.9145 | 3.22829 | 5.82838 |
| cwp17 | CD630_n00620 | 128 – 192 | 62 – 123 | -11.833 | 3.43363 | 5.82838 |
| CD630_19440 | CD630_n00620 | 144 – 184 | 61 – 98 | -11.7368 | 4.53937 | 5.82838 |
| CD630_18920 | CD630_n00620 | 6 – 26 | 54 – 74 | -11.6787 | 4.43558 | 5.82838 |
| CD630_23760 | CD630_n00620 | 54 – 76 | 163 – 189 | -11.6629 | 4.54307 | 5.82838 |
| CD630_19030 | CD630_n00620 | 155 – 173 | 1 – 19 | -11.6618 | 3.34240 | 5.82838 |
| cbiQ | CD630_n00620 | 122 – 160 | 153 – 185 | -11.6395 | 3.79830 | 5.82838 |
| CD630_19700 | CD630_n00620 | 7 – 48 | 152 – 199 | -11.5695 | 3.85861 | 5.82838 |
| CD630_36781 | CD630_n00620 | 5 – 57 | 52 – 97 | -11.557 | 3.75946 | 5.82838 |
| cobS | CD630_n00620 | 4 – 21 | 31 – 48 | -11.5254 | 3.67481 | 5.82838 |
| CD630_06380 | CD630_n00620 | 117 – 201 | 88 – 181 | -11.5012 | 4.32211 | 5.82838 |
| CD630_17710 | CD630_n00620 | 165 – 196 | 152 – 190 | -11.4768 | 4.37688 | 5.82838 |
| CD630_25300 | CD630_n00620 | 58 – 93 | 189 – 222 | -11.4662 | 4.49333 | 5.82838 |
| CD630_06060 | CD630_n00620 | 149 – 189 | 45 – 96 | -11.3863 | 4.94050 | 5.82838 |
| CD630_06100 | CD630_n00620 | 110 – 163 | 49 – 102 | -11.3694 | 4.43918 | 5.82838 |
| CD630_06960 | CD630_n00620 | 114 – 179 | 63 – 125 | -11.2535 | 3.16317 | 5.82838 |
| uxaA%27 | CD630_n00620 | 133 – 162 | 64 – 95 | -11.1955 | 4.26165 | 5.82838 |
| CD630_03650 | CD630_n00620 | 125 – 183 | 46 – 101 | -11.175 | 3.93697 | 5.82838 |
| folC | CD630_n00620 | 28 – 87 | 154 – 198 | -11.1586 | 3.85634 | 5.82838 |
| CD630_31490 | CD630_n00620 | 32 – 71 | 65 – 99 | -11.145 | 4.08593 | 5.82838 |
| CD630_14710 | CD630_n00620 | 4 – 54 | 163 – 215 | -11.0268 | 3.65171 | 5.82838 |
| CD630_14810 | CD630_n00620 | 77 – 165 | 65 – 168 | -10.9789 | 3.35323 | 5.82838 |
| moeA | CD630_n00620 | 104 – 127 | 171 – 198 | -10.8769 | 3.26118 | 5.82838 |
| CD630_07840 | CD630_n00620 | 22 – 82 | 131 – 186 | -10.8087 | 4.26761 | 5.82838 |
| CD630_32710 | CD630_n00620 | 146 – 181 | 160 – 201 | -10.7663 | 4.74276 | 5.82838 |
| CD630_15900 | CD630_n00620 | 147 – 189 | 60 – 101 | -10.7633 | 3.17664 | 5.82838 |
| CD630_08460 | CD630_n00620 | 24 – 52 | 70 – 98 | -10.758 | 4.15083 | 5.82838 |
| dpaA | CD630_n00620 | 179 – 196 | 87 – 105 | -10.753 | 4.65364 | 5.82838 |
| CD630_28280 | CD630_n00620 | 42 – 89 | 41 – 99 | -10.7474 | 4.59833 | 5.82838 |
| CD630_16330 | CD630_n00620 | 147 – 200 | 49 – 98 | -10.7261 | 4.11180 | 5.82838 |
| CD630_04320 | CD630_n00620 | 65 – 113 | 164 – 213 | -10.7114 | 3.68342 | 5.82838 |
| CD630_10970 | CD630_n00620 | 89 – 129 | 153 – 196 | -10.6896 | 3.53850 | 5.82838 |
| CD630_21770 | CD630_n00620 | 1 – 18 | 82 – 98 | -10.657 | 4.02587 | 5.82838 |
| CD630_33480 | CD630_n00620 | 141 – 178 | 61 – 97 | -10.6234 | 3.67613 | 5.82838 |
| CD630_24770 | CD630_n00620 | 49 – 191 | 58 – 184 | -10.5969 | 3.20798 | 5.82838 |
| CD630_07121 | CD630_n00620 | 7 – 57 | 154 – 204 | -10.5228 | 3.26177 | 5.82838 |
| CD630_18910 | CD630_n00620 | 32 – 40 | 10 – 18 | -10.5209 | 3.30688 | 5.82838 |
| cdtR | CD630_n00620 | 5 – 54 | 43 – 93 | -10.5181 | 5.84368 | 5.82838 |
| hfq | CD630_n00620 | 150 – 200 | 150 – 196 | -10.4415 | 5.05609 | 5.82838 |
| kdgT1 | CD630_n00620 | 4 – 45 | 144 – 175 | -10.4102 | 4.49207 | 5.82838 |
| CD630_18940 | CD630_n00620 | 57 – 172 | 78 – 182 | -10.3923 | 4.09988 | 5.82838 |
| CD630_22070 | CD630_n00620 | 45 – 193 | 49 – 196 | -10.3626 | 4.09651 | 5.82838 |
| CD630_21620 | CD630_n00620 | 100 – 188 | 120 – 199 | -10.3571 | 3.02649 | 5.82838 |
| CD630_05490 | CD630_n00620 | 155 – 201 | 156 – 196 | -10.3466 | 3.89852 | 5.82838 |
| acoL | CD630_n00620 | 107 – 145 | 171 – 198 | -10.3237 | 3.32169 | 5.82838 |
| CD630_31381 | CD630_n00620 | 33 – 92 | 159 – 204 | -10.2255 | 3.49137 | 5.82838 |
| CD630_12110 | CD630_n00620 | 101 – 122 | 65 – 82 | -10.1906 | 4.36078 | 5.82838 |
| bltD | CD630_n00620 | 188 – 201 | 177 – 189 | -10.1831 | 4.49926 | 5.82838 |
| CD630_13350 | CD630_n00620 | 166 – 199 | 176 – 204 | -10.1712 | 4.15139 | 5.82838 |
| tcdA | CD630_n00620 | 137 – 197 | 116 – 185 | -10.1431 | 4.70593 | 5.82838 |
| CD630_33650 | CD630_n00620 | 86 – 122 | 153 – 191 | -10.1382 | 4.80565 | 5.82838 |
| CD630_26150 | CD630_n00620 | 137 – 146 | 89 – 99 | -10.0983 | 4.06302 | 5.82838 |
| CD630_18010 | CD630_n00620 | 186 – 200 | 88 – 102 | -10.0833 | 4.21967 | 5.82838 |
| CD630_31970 | CD630_n00620 | 175 – 200 | 65 – 89 | -10.0673 | 3.24511 | 5.82838 |
| CD630_09772 | CD630_n00620 | 5 – 29 | 71 – 97 | -10.0286 | 3.66033 | 5.82838 |
| CD630_18410 | CD630_n00620 | 125 – 171 | 48 – 94 | -10.0067 | 3.59739 | 5.82838 |
| CD630_22050 | CD630_n00620 | 137 – 199 | 32 – 96 | -10.0038 | 4.87639 | 5.82838 |
| CD630_15150 | CD630_n00620 | 83 – 152 | 145 – 199 | -9.94619 | 3.35564 | 5.82838 |
| CD630_06140 | CD630_n00620 | 117 – 181 | 126 – 190 | -9.94493 | 4.12519 | 5.82838 |
| CD630_32500 | CD630_n00620 | 7 – 32 | 149 – 174 | -9.91815 | 6.17288 | 5.82838 |
| CD630_07480 | CD630_n00620 | 158 – 165 | 12 – 19 | -9.90751 | 5.18321 | 5.82838 |
| bcp | CD630_n00620 | 109 – 145 | 170 – 199 | -9.87279 | 3.16670 | 5.82838 |
| CD630_06540 | CD630_n00620 | 12 – 57 | 154 – 195 | -9.83535 | 3.54673 | 5.82838 |
| CD630_17900 | CD630_n00620 | 87 – 115 | 69 – 95 | -9.80859 | 3.52167 | 5.82838 |
| CD630_27110 | CD630_n00620 | 6 – 20 | 36 – 51 | -9.80708 | 3.78231 | 5.82838 |
| CD630_25040 | CD630_n00620 | 78 – 102 | 180 – 206 | -9.79901 | 3.43898 | 5.82838 |
| cbpA | CD630_n00620 | 147 – 192 | 156 – 205 | -9.78901 | 3.03040 | 5.82838 |
| CD630_15160 | CD630_n00620 | 129 – 177 | 44 – 94 | -9.77121 | 4.32318 | 5.82838 |
| CD630_24220 | CD630_n00620 | 52 – 76 | 42 – 66 | -9.76516 | 4.18037 | 5.82838 |
| CD630_14130 | CD630_n00620 | 134 – 183 | 49 – 96 | -9.74376 | 4.57970 | 5.82838 |
| CD630_28870 | CD630_n00620 | 172 – 186 | 8 – 22 | -9.74141 | 3.41991 | 5.82838 |
| CD630_05880 | CD630_n00620 | 130 – 143 | 186 – 199 | -9.70177 | 4.28339 | 5.82838 |
| appC | CD630_n00620 | 85 – 182 | 28 – 121 | -9.67764 | 4.50438 | 5.82838 |
| CD630_14250 | CD630_n00620 | 137 – 198 | 70 – 123 | -9.66292 | 4.19604 | 5.82838 |
| CD630_26630 | CD630_n00620 | 8 – 68 | 148 – 199 | -9.65602 | 4.61557 | 5.82838 |
| CD630_18970 | CD630_n00620 | 116 – 141 | 174 – 201 | -9.64001 | 3.57243 | 5.82838 |
| CD630_10220 | CD630_n00620 | 3 – 20 | 166 – 183 | -9.62936 | 4.08233 | 5.82838 |
| CD630_12650 | CD630_n00620 | 96 – 134 | 159 – 186 | -9.62234 | 6.31827 | 5.82838 |
| CD630_20280 | CD630_n00620 | 16 – 68 | 53 – 95 | -9.62123 | 3.46343 | 5.82838 |
| CD630_09050 | CD630_n00620 | 179 – 187 | 10 – 18 | -9.60144 | 3.96216 | 5.82838 |
| CD630_25310 | CD630_n00620 | 136 – 168 | 52 – 86 | -9.59921 | 3.99861 | 5.82838 |
| CD630_08140 | CD630_n00620 | 2 – 25 | 53 – 81 | -9.59607 | 3.30159 | 5.82838 |
| CD630_28550 | CD630_n00620 | 159 – 180 | 167 – 190 | -9.57102 | 3.51196 | 5.82838 |
| dltB | CD630_n00620 | 153 – 193 | 116 – 165 | -9.55442 | 3.81485 | 5.82838 |
| CD630_17720 | CD630_n00620 | 4 – 33 | 46 – 77 | -9.54045 | 3.22910 | 5.82838 |
| CD630_22030 | CD630_n00620 | 9 – 36 | 74 – 101 | -9.52563 | 3.29934 | 5.82838 |
| garK | CD630_n00620 | 14 – 67 | 143 – 196 | -9.5159 | 3.72836 | 5.82838 |
| CD630_16160 | CD630_n00620 | 178 – 201 | 44 – 65 | -9.50909 | 6.11801 | 5.82838 |
| CD630_22200 | CD630_n00680 | 102 – 173 | 109 – 179 | -19.6136 | 4.18288 | 6.52781 |
| CD630_21770 | CD630_n00680 | 74 – 161 | 283 – 366 | -19.0043 | 4.02587 | 6.52781 |
| feoB | CD630_n00680 | 135 – 197 | 82 – 138 | -17.3971 | 4.05364 | 6.52781 |
| CD630_09100 | CD630_n00680 | 3 – 85 | 213 – 285 | -17.1181 | 5.92260 | 6.52781 |
| CD630_05870 | CD630_n00680 | 160 – 196 | 349 – 393 | -16.6381 | 4.28339 | 6.52781 |
| CD630_20450 | CD630_n00680 | 118 – 165 | 87 – 137 | -16.4579 | 3.77388 | 6.52781 |
| CD630_06140 | CD630_n00680 | 16 – 92 | 107 – 181 | -16.0356 | 4.12519 | 6.52781 |
| CD630_05280 | CD630_n00680 | 8 – 58 | 246 – 302 | -15.7554 | 3.91665 | 6.52781 |
| CD630_32710 | CD630_n00680 | 110 – 152 | 351 – 385 | -15.6948 | 4.74276 | 6.52781 |
| CD630_05160 | CD630_n00680 | 165 – 196 | 274 – 306 | -15.5527 | 5.03573 | 6.52781 |
| CD630_19580 | CD630_n00680 | 68 – 95 | 339 – 366 | -15.0753 | 4.51902 | 6.52781 |
| CD630_21090 | CD630_n00680 | 146 – 171 | 266 – 293 | -15.0565 | 4.15803 | 6.52781 |
| thiH | CD630_n00680 | 171 – 200 | 351 – 375 | -14.9995 | 3.58230 | 6.52781 |
| CD630_31461 | CD630_n00680 | 38 – 80 | 247 – 295 | -14.9731 | 4.41476 | 6.52781 |
| CD630_21260 | CD630_n00680 | 138 – 163 | 32 – 59 | -14.6657 | 4.22255 | 6.52781 |
| CD630_21620 | CD630_n00680 | 11 – 38 | 111 – 137 | -14.5028 | 3.02649 | 6.52781 |
| CD630_21570 | CD630_n00680 | 157 – 186 | 111 – 137 | -14.4523 | 6.48385 | 6.52781 |
| CD630_10220 | CD630_n00680 | 113 – 171 | 181 – 237 | -14.4178 | 4.08233 | 6.52781 |
| CD630_08080 | CD630_n00680 | 3 – 56 | 114 – 167 | -14.3304 | 4.87819 | 6.52781 |
| CD630_32720 | CD630_n00680 | 64 – 94 | 349 – 378 | -14.0183 | 4.74276 | 6.52781 |
| metQ | CD630_n00680 | 82 – 143 | 286 – 360 | -14.0083 | 3.87190 | 6.52781 |
| kdpD | CD630_n00680 | 136 – 199 | 314 – 381 | -13.9794 | 4.27140 | 6.52781 |
| cobQ | CD630_n00680 | 59 – 106 | 351 – 399 | -13.9315 | 3.13524 | 6.52781 |
| CD630_27570 | CD630_n00680 | 108 – 138 | 352 – 397 | -13.8438 | 3.46216 | 6.52781 |
| glnA | CD630_n00680 | 163 – 187 | 109 – 137 | -13.5433 | 3.87629 | 6.52781 |
| etfA | CD630_n00680 | 9 – 53 | 354 – 386 | -13.5055 | 3.41204 | 6.52781 |
| CD630_05290 | CD630_n00680 | 1 – 48 | 269 – 315 | -13.4702 | 3.97604 | 6.52781 |
| CD630_23280 | CD630_n00680 | 87 – 155 | 334 – 397 | -13.2807 | 3.50093 | 6.52781 |
| folC | CD630_n00680 | 113 – 160 | 247 – 296 | -13.2383 | 3.85634 | 6.52781 |
| CD630_18311 | CD630_n00680 | 71 – 108 | 352 – 386 | -13.1447 | 6.86047 | 6.52781 |
| CD630_12650 | CD630_n00680 | 97 – 133 | 168 – 205 | -13.0995 | 6.31827 | 6.52781 |
| CD630_36690 | CD630_n00680 | 121 – 151 | 157 – 192 | -13.001 | 3.53761 | 6.52781 |
| CD630_02790 | CD630_n00680 | 82 – 165 | 286 – 359 | -12.9345 | 4.77661 | 6.52781 |
| CD630_04850 | CD630_n00680 | 142 – 196 | 258 – 312 | -12.8956 | 4.40788 | 6.52781 |
| malY | CD630_n00680 | 48 – 83 | 352 – 391 | -12.8537 | 4.46345 | 6.52781 |
| pilD | CD630_n00680 | 3 – 25 | 109 – 132 | -12.7992 | 4.54123 | 6.52781 |
| CD630_28320 | CD630_n00680 | 84 – 193 | 87 – 195 | -12.7801 | 3.99184 | 6.52781 |
| CD630_01560 | CD630_n00680 | 79 – 147 | 111 – 194 | -12.772 | 4.06352 | 6.52781 |
| CD630_14250 | CD630_n00680 | 163 – 201 | 356 – 394 | -12.7282 | 4.19604 | 6.52781 |
| CD630_14110 | CD630_n00680 | 17 – 57 | 144 – 179 | -12.702 | 3.39550 | 6.52781 |
| CD630_20280 | CD630_n00680 | 55 – 132 | 84 – 171 | -12.6511 | 3.46343 | 6.52781 |
| CD630_20460 | CD630_n00680 | 46 – 102 | 350 – 399 | -12.6394 | 7.94034 | 6.52781 |
| CD630_17151 | CD630_n00680 | 145 – 196 | 110 – 151 | -12.587 | 6.43124 | 6.52781 |
| CD630_30140 | CD630_n00680 | 66 – 112 | 106 – 149 | -12.5142 | 3.28343 | 6.52781 |
| CD630_15460 | CD630_n00680 | 146 – 180 | 50 – 79 | -12.4712 | 6.86296 | 6.52781 |
| rolB | CD630_n00680 | 112 – 137 | 352 – 393 | -12.4052 | 3.60270 | 6.52781 |
| CD630_19700 | CD630_n00680 | 17 – 81 | 334 – 394 | -12.3081 | 3.85861 | 6.52781 |
| CD630_31530 | CD630_n00680 | 118 – 144 | 109 – 137 | -12.2866 | 3.49100 | 6.52781 |
| CD630_27770 | CD630_n00680 | 78 – 102 | 333 – 364 | -12.2531 | 4.08588 | 6.52781 |
| CD630_17511 | CD630_n00680 | 11 – 34 | 54 – 79 | -12.2384 | 9.12322 | 6.52781 |
| cspD | CD630_n00680 | 91 – 156 | 318 – 393 | -12.2212 | 3.23587 | 6.52781 |
| CD630_10970 | CD630_n00680 | 175 – 194 | 110 – 131 | -12.1561 | 3.53850 | 6.52781 |
| CD630_21330 | CD630_n00680 | 1 – 97 | 106 – 178 | -12.1444 | 4.10111 | 6.52781 |
| def1 | CD630_n00680 | 165 – 188 | 353 – 375 | -12.1415 | 3.99720 | 6.52781 |
| CD630_07121 | CD630_n00680 | 24 – 54 | 157 – 190 | -12.1395 | 3.26177 | 6.52781 |
| CD630_18610 | CD630_n00680 | 185 – 200 | 263 – 278 | -12.1103 | 4.15582 | 6.52781 |
| bcp | CD630_n00680 | 77 – 91 | 109 – 123 | -12.1063 | 3.16670 | 6.52781 |
| rolA | CD630_n00680 | 23 – 59 | 354 – 394 | -12.0938 | 4.56362 | 6.52781 |
| CD630_05830 | CD630_n00680 | 112 – 144 | 30 – 71 | -11.9755 | 3.35360 | 6.52781 |
| CD630_14930 | CD630_n00680 | 55 – 140 | 325 – 395 | -11.9345 | 3.31335 | 6.52781 |
| CD630_13020 | CD630_n00680 | 115 – 162 | 278 – 323 | -11.8999 | 3.52267 | 6.52781 |
| iorA | CD630_n00680 | 53 – 92 | 351 – 397 | -11.8479 | 3.83173 | 6.52781 |
| CD630_21110 | CD630_n00680 | 119 – 154 | 126 – 163 | -11.8217 | 3.89630 | 6.52781 |
| CD630_33600 | CD630_n00680 | 139 – 171 | 354 – 386 | -11.7919 | 3.73196 | 6.52781 |
| sspA | CD630_n00680 | 64 – 109 | 90 – 130 | -11.786 | 3.62689 | 6.52781 |
| CD630_25020 | CD630_n00680 | 10 – 66 | 120 – 173 | -11.7794 | 4.17721 | 6.52781 |
| bglA | CD630_n00680 | 88 – 112 | 351 – 375 | -11.7486 | 4.50315 | 6.52781 |
| gcp | CD630_n00680 | 10 – 38 | 213 – 237 | -11.6922 | 4.92798 | 6.52781 |
| CD630_03650 | CD630_n00680 | 150 – 181 | 31 – 62 | -11.5862 | 3.93697 | 6.52781 |
| CD630_14130 | CD630_n00680 | 92 – 122 | 350 – 382 | -11.5678 | 4.57970 | 6.52781 |
| cspC | CD630_n00680 | 91 – 141 | 352 – 393 | -11.5626 | 5.48848 | 6.52781 |
| CD630_35430 | CD630_n00680 | 163 – 182 | 115 – 134 | -11.56 | 4.28798 | 6.52781 |
| CD630_21910 | CD630_n00680 | 76 – 95 | 351 – 369 | -11.5063 | 4.66600 | 6.52781 |
| CD630_36781 | CD630_n00680 | 145 – 198 | 115 – 174 | -11.4974 | 3.75946 | 6.52781 |
| CD630_17170 | CD630_n00680 | 123 – 147 | 213 – 239 | -11.4772 | 4.10230 | 6.52781 |
| CD630_13530 | CD630_n00680 | 156 – 194 | 281 – 321 | -11.4729 | 4.09832 | 6.52781 |
| CD630_15160 | CD630_n00680 | 157 – 198 | 266 – 293 | -11.3906 | 4.32318 | 6.52781 |
| CD630_27360 | CD630_n00680 | 58 – 77 | 68 – 87 | -11.3772 | 3.79570 | 6.52781 |
| CD630_06060 | CD630_n00680 | 13 – 44 | 220 – 266 | -11.3604 | 4.94050 | 6.52781 |
| CD630_24590 | CD630_n00680 | 52 – 148 | 284 – 377 | -11.3134 | 3.70401 | 6.52781 |
| CD630_22050 | CD630_n00680 | 170 – 194 | 299 – 325 | -11.3042 | 4.87639 | 6.52781 |
| srlD | CD630_n00680 | 84 – 106 | 334 – 358 | -11.2482 | 4.19843 | 6.52781 |
| CD630_08360 | CD630_n00680 | 22 – 63 | 353 – 385 | -11.2295 | 3.19339 | 6.52781 |
| glgP | CD630_n00680 | 81 – 129 | 91 – 133 | -11.2222 | 3.28530 | 6.52781 |
| CD630_06080 | CD630_n00680 | 157 – 186 | 352 – 382 | -11.2152 | 3.82315 | 6.52781 |
| adhE | CD630_n00680 | 54 – 95 | 104 – 139 | -11.1858 | 3.06485 | 6.52781 |
| CD630_31880 | CD630_n00680 | 2 – 14 | 215 – 227 | -11.1772 | 3.97522 | 6.52781 |
| CD630_21020 | CD630_n00680 | 21 – 80 | 193 – 248 | -11.1668 | 4.28666 | 6.52781 |
| CD630_31210 | CD630_n00680 | 139 – 178 | 205 – 240 | -11.158 | 4.17693 | 6.52781 |
| CD630_16330 | CD630_n00680 | 154 – 198 | 124 – 162 | -11.1498 | 4.11180 | 6.52781 |
| CD630_04610 | CD630_n00680 | 18 – 45 | 354 – 377 | -11.128 | 3.83025 | 6.52781 |
| CD630_30310 | CD630_n00680 | 142 – 167 | 114 – 140 | -11.1123 | 4.56059 | 6.52781 |
| CD630_04341 | CD630_n00680 | 48 – 80 | 352 – 387 | -11.1022 | 4.29262 | 6.52781 |
| dltB | CD630_n00680 | 6 – 30 | 124 – 148 | -11.0654 | 3.81485 | 6.52781 |
| CD630_21700 | CD630_n00680 | 20 – 41 | 119 – 145 | -11.0577 | 3.96650 | 6.52781 |
| CD630_21540 | CD630_n00680 | 47 – 70 | 215 – 238 | -11.0441 | 4.33090 | 6.52781 |
| CD630_08140 | CD630_n00680 | 9 – 124 | 36 – 143 | -11.0416 | 3.30159 | 6.52781 |
| cbpA | CD630_n00680 | 163 – 184 | 291 – 312 | -10.97 | 3.03040 | 6.52781 |
| CD630_32070 | CD630_n00680 | 50 – 79 | 104 – 134 | -10.8565 | 3.51964 | 6.52781 |
| CD630_20670 | CD630_n00680 | 179 – 194 | 74 – 89 | -10.8089 | 4.42757 | 6.52781 |
| CD630_25040 | CD630_n00680 | 144 – 177 | 27 – 62 | -10.8006 | 3.43898 | 6.52781 |
| CD630_27230 | CD630_n00680 | 90 – 121 | 353 – 375 | -10.7536 | 3.52462 | 6.52781 |
| CD630_06340 | CD630_n00680 | 4 – 45 | 216 – 258 | -10.7532 | 4.35983 | 6.52781 |
| CD630_06560 | CD630_n00680 | 4 – 19 | 290 – 306 | -10.7344 | 4.02532 | 6.52781 |
| CD630_10050 | CD630_n00680 | 176 – 198 | 354 – 373 | -10.6691 | 3.93750 | 6.52781 |
| CD630_29500 | CD630_n00680 | 43 – 60 | 171 – 187 | -10.6636 | 3.70644 | 6.52781 |
| CD630_22950 | CD630_n00680 | 3 – 25 | 245 – 269 | -10.6522 | 3.96883 | 6.52781 |
| CD630_31960 | CD630_n00680 | 31 – 52 | 371 – 393 | -10.6362 | 4.93757 | 6.52781 |
| CD630_08370 | CD630_n00680 | 2 – 26 | 110 – 135 | -10.6261 | 3.65639 | 6.52781 |
| CD630_24220 | CD630_n00680 | 126 – 169 | 211 – 255 | -10.5927 | 4.18037 | 6.52781 |
| CD630_33570 | CD630_n00680 | 150 – 162 | 227 – 239 | -10.5897 | 3.95778 | 6.52781 |
| CD630_23080 | CD630_n00680 | 1 – 88 | 257 – 347 | -10.5636 | 5.23027 | 6.52781 |
| CD630_10830 | CD630_n00680 | 61 – 93 | 352 – 383 | -10.4823 | 4.82398 | 6.52781 |
| CD630_32050 | CD630_n00680 | 153 – 184 | 104 – 132 | -10.4261 | 5.73797 | 6.52781 |
| xdhA | CD630_n00680 | 3 – 148 | 102 – 231 | -10.4071 | 4.45147 | 6.52781 |
| CD630_18120 | CD630_n00680 | 6 – 62 | 115 – 164 | -10.3962 | 3.99713 | 6.52781 |
| CD630_25050 | CD630_n00680 | 153 – 164 | 346 – 357 | -10.3816 | 4.33624 | 6.52781 |
| CD630_27270 | CD630_n00680 | 93 – 105 | 227 – 239 | -10.3748 | 3.50892 | 6.52781 |
| CD630_33420 | CD630_n00680 | 26 – 76 | 324 – 386 | -10.3563 | 3.22095 | 6.52781 |

| | | | | | | |
|---|---|---|---|---|---|---|
| CD630_11240 | CD630_n00680 | 102 -- 130 | 63 -- 90 | -10.3343 | 3.20274 | 6.52781 |
| CD630_34520 | CD630_n00680 | 22 -- 55 | 138 -- 178 | -10.2832 | 3.83016 | 6.52781 |
| CD630_28150 | CD630_n00680 | 153 -- 201 | 347 -- 399 | -10.2711 | 4.18447 | 6.52781 |
| CD630_26750 | CD630_n00680 | 22 -- 61 | 111 -- 149 | -10.2397 | 3.60610 | 6.52781 |
| oppC | CD630_n00680 | 164 -- 182 | 111 -- 131 | -10.2387 | 3.01695 | 6.52781 |
| CD630_31490 | CD630_n00680 | 101 -- 129 | 360 -- 384 | -10.2306 | 4.08593 | 6.52781 |
| CD630_22660 | CD630_n00680 | 134 -- 201 | 78 -- 137 | -10.2274 | 3.60496 | 6.52781 |
| CD630_15111 | CD630_n00680 | 37 -- 177 | 166 -- 297 | -10.2144 | 4.18664 | 6.52781 |
| rbsC | CD630_n00680 | 34 -- 51 | 335 -- 352 | -10.1944 | 3.71650 | 6.52781 |
| CD630_04550 | CD630_n00680 | 95 -- 131 | 212 -- 247 | -10.1874 | 3.05227 | 6.52781 |
| CD630_06170 | CD630_n00680 | 117 -- 136 | 113 -- 140 | -10.1807 | 4.95366 | 6.52781 |
| CD630_19670 | CD630_n00680 | 72 -- 87 | 351 -- 367 | -10.1769 | 3.20718 | 6.52781 |
| pgm | CD630_n00680 | 118 -- 140 | 349 -- 379 | -10.1699 | 3.45439 | 6.52781 |
| CD630_06190 | CD630_n00680 | 141 -- 151 | 49 -- 59 | -10.166 | 3.57064 | 6.52781 |
| CD630_15360 | CD630_n00680 | 20 -- 44 | 229 -- 253 | -10.1579 | 4.47208 | 6.52781 |
| CD630_25190 | CD630_n00680 | 174 -- 181 | 87 -- 94 | -10.1564 | 3.19663 | 6.52781 |
| CD630_17770 | CD630_n00680 | 76 -- 104 | 336 -- 364 | -10.1444 | 5.06490 | 6.52781 |
| CD630_20260 | CD630_n00680 | 7 -- 43 | 265 -- 295 | -10.1427 | 3.84162 | 6.52781 |
| CD630_06200 | CD630_n00680 | 44 -- 66 | 286 -- 313 | -10.1422 | 3.57064 | 6.52781 |
| CD630_17730 | CD630_n00680 | 76 -- 97 | 104 -- 124 | -10.1393 | 4.73397 | 6.52781 |
| CD630_27120 | CD630_n00680 | 3 -- 36 | 226 -- 262 | -10.1229 | 4.43720 | 6.52781 |
| CD630_18960 | CD630_n00680 | 4 -- 28 | 109 -- 134 | -10.1036 | 4.37822 | 6.52781 |
| dapB2 | CD630_n00680 | 114 -- 191 | 348 -- 398 | -10.0949 | 3.27218 | 6.52781 |
| CD630_34050 | CD630_n00680 | 8 -- 20 | 5 -- 17 | -10.0935 | 4.03863 | 6.52781 |
| CD630_27610 | CD630_n00680 | 15 -- 28 | 234 -- 248 | -10.0861 | 3.96465 | 6.52781 |
| cobC | CD630_n00680 | 169 -- 195 | 351 -- 378 | -10.0802 | 3.15854 | 6.52781 |
| CD630_08660 | CD630_n00680 | 148 -- 186 | 213 -- 251 | -10.0721 | 3.86467 | 6.52781 |
| CD630_07960 | CD630_n00680 | 180 -- 201 | 274 -- 295 | -10.0633 | 3.22537 | 6.52781 |
| CD630_11241 | CD630_n00680 | 159 -- 184 | 235 -- 259 | -10.0618 | 4.85723 | 6.52781 |
| CD630_20541 | CD630_n00680 | 89 -- 113 | 350 -- 374 | -10.0614 | 5.74527 | 6.52781 |
| CD630_23070 | CD630_n00680 | 172 -- 187 | 83 -- 97 | -10.0583 | 5.23027 | 6.52781 |
| fliI | CD630_n00680 | 51 -- 107 | 197 -- 245 | -10.0497 | 3.03666 | 6.52781 |
| CD630_01570 | CD630_n00680 | 11 -- 64 | 324 -- 391 | -10.0486 | 6.08699 | 6.52781 |
| CD630_05950 | CD630_n00680 | 19 -- 50 | 112 -- 142 | -10.0295 | 3.31669 | 6.52781 |
| CD630_25070 | CD630_n00680 | 106 -- 131 | 111 -- 135 | -10.0238 | 3.45164 | 6.52781 |
| CD630_19904 | CD630_n00680 | 25 -- 37 | 127 -- 141 | -10.0162 | 5.96526 | 6.52781 |
| CD630_30100 | CD630_n00680 | 176 -- 186 | 14 -- 24 | -9.9901 | 4.13234 | 6.52781 |
| CD630_13580 | CD630_n00680 | 88 -- 110 | 372 -- 393 | -9.98481 | 3.80715 | 6.52781 |
| opuCC | CD630_n00680 | 184 -- 197 | 169 -- 183 | -9.9252 | 4.07035 | 6.52781 |
| CD630_02740 | CD630_n00680 | 88 -- 150 | 286 -- 355 | -9.92302 | 5.02277 | 6.52781 |
| CD630_30360 | CD630_n00680 | 88 -- 181 | 287 -- 393 | -9.90997 | 4.26509 | 6.52781 |
| bglA | CD630_n00680 | 151 -- 183 | 352 -- 393 | -9.90197 | 4.86402 | 6.52781 |
| CD630_14610 | CD630_n00680 | 146 -- 177 | 258 -- 298 | -9.89087 | 4.89428 | 6.52781 |
| CD630_22500 | CD630_n00680 | 149 -- 195 | 95 -- 148 | -9.88528 | 4.03688 | 6.52781 |
| CD630_08950 | CD630_n00680 | 156 -- 191 | 276 -- 313 | -9.87247 | 3.71905 | 6.52781 |
| CD630_05220 | CD630_n00680 | 67 -- 79 | 335 -- 347 | -9.85476 | 3.65057 | 6.52781 |
| CD630_32690 | CD630_n00680 | 139 -- 173 | 352 -- 377 | -9.84272 | 7.32676 | 6.52781 |
| CD630_06970 | CD630_n00680 | 110 -- 152 | 211 -- 256 | -9.83701 | 3.82312 | 6.52781 |
| CD630_07870 | CD630_n00680 | 184 -- 199 | 158 -- 174 | -9.82911 | 3.22829 | 6.52781 |
| CD630_28181 | CD630_n00680 | 14 -- 76 | 87 -- 137 | -9.81231 | 3.91299 | 6.52781 |
| CD630_19790 | CD630_n00680 | 53 -- 92 | 351 -- 380 | -9.80759 | 3.79593 | 6.52781 |
| ssuC | CD630_n00680 | 54 -- 68 | 350 -- 364 | -9.79755 | 3.45491 | 6.52781 |
| CD630_19401 | CD630_n00680 | 26 -- 107 | 92 -- 172 | -9.77057 | 4.37084 | 6.52781 |
| CD630_06410 | CD630_n00680 | 86 -- 130 | 247 -- 292 | -9.76527 | 4.14404 | 6.52781 |
| CD630_09970 | CD630_n00680 | 8 -- 29 | 113 -- 136 | -9.76199 | 3.25917 | 6.52781 |
| CD630_14630 | CD630_n00680 | 79 -- 91 | 352 -- 364 | -9.75547 | 3.34485 | 6.52781 |
| cwp26 | CD630_n00680 | 124 -- 161 | 304 -- 340 | -9.74978 | 4.44377 | 6.52781 |
| CD630_29960 | CD630_n00680 | 137 -- 168 | 114 -- 137 | -9.7354 | 4.51904 | 6.52781 |
| garK | CD630_n00680 | 15 -- 38 | 353 -- 375 | -9.72713 | 3.72836 | 6.52781 |
| dpaA | CD630_n00680 | 163 -- 188 | 344 -- 364 | -9.71701 | 4.65364 | 6.52781 |
| CD630_04410 | CD630_n00680 | 126 -- 160 | 351 -- 386 | -9.71385 | 3.90451 | 6.52781 |
| CD630_10000 | CD630_n00680 | 136 -- 166 | 104 -- 135 | -9.71302 | 3.86040 | 6.52781 |
| CD630_18230 | CD630_n00680 | 2 -- 75 | 271 -- 343 | -9.68857 | 3.00184 | 6.52781 |
| CD630_14990 | CD630_n00680 | 120 -- 151 | 220 -- 266 | -9.68788 | 3.78619 | 6.52781 |
| CD630_22720 | CD630_n00680 | 122 -- 162 | 351 -- 385 | -9.67863 | 3.81804 | 6.52781 |
| CD630_30730 | CD630_n00680 | 118 -- 130 | 265 -- 277 | -9.67136 | 3.13236 | 6.52781 |
| CD630_06380 | CD630_n00680 | 135 -- 151 | 345 -- 360 | -9.66554 | 4.32211 | 6.52781 |
| CD630_32890 | CD630_n00680 | 188 -- 200 | 189 -- 201 | -9.64358 | 3.41276 | 6.52781 |
| CD630_22550 | CD630_n00680 | 141 -- 182 | 277 -- 327 | -9.61724 | 3.62379 | 6.52781 |
| CD630_14700 | CD630_n00680 | 130 -- 155 | 352 -- 376 | -9.60212 | 3.98234 | 6.52781 |
| CD630_26150 | CD630_n00680 | 42 -- 90 | 117 -- 164 | -9.60062 | 4.06302 | 6.52781 |
| CD630_24830 | CD630_n00680 | 139 -- 171 | 257 -- 285 | -9.60049 | 4.98239 | 6.52781 |
| CD630_33650 | CD630_n00680 | 88 -- 111 | 336 -- 357 | -9.5988 | 4.80565 | 6.52781 |
| appC | CD630_n00680 | 39 -- 58 | 73 -- 88 | -9.58936 | 4.50438 | 6.52781 |
| CD630_21730 | CD630_n00680 | 131 -- 166 | 109 -- 136 | -9.56137 | 3.45076 | 6.52781 |
| cheW | CD630_n00680 | 11 -- 49 | 353 -- 385 | -9.51992 | 3.72371 | 6.52781 |
| CD630_15431 | CD630_n00680 | 139 -- 153 | 284 -- 298 | -9.51448 | 5.29780 | 6.52781 |
| CD630_22200 | CD630_n00850 | 102 -- 173 | 116 -- 187 | -19.2433 | 4.18288 | 5.53585 |
| CD630_24220 | CD630_n00850 | 126 -- 182 | 334 -- 394 | -17.4624 | 4.18037 | 5.53585 |
| CD630_08080 | CD630_n00850 | 3 -- 51 | 125 -- 175 | -16.985 | 4.87819 | 5.53585 |
| CD630_24590 | CD630_n00850 | 157 -- 192 | 363 -- 401 | -16.6993 | 3.70401 | 5.53585 |
| CD630_06340 | CD630_n00850 | 78 -- 137 | 293 -- 385 | -16.3658 | 4.35983 | 5.53585 |
| CD630_31530 | CD630_n00850 | 118 -- 168 | 90 -- 144 | -16.2882 | 3.49100 | 5.53585 |
| feoB | CD630_n00850 | 134 -- 197 | 88 -- 146 | -16.1585 | 4.05364 | 5.53585 |
| eutW | CD630_n00850 | 39 -- 110 | 207 -- 268 | -16.1363 | 4.55715 | 5.53585 |
| CD630_20450 | CD630_n00850 | 118 -- 165 | 93 -- 145 | -15.8031 | 3.77388 | 5.53585 |
| CD630_05830 | CD630_n00850 | 125 -- 191 | 127 -- 203 | -15.631 | 3.35360 | 5.53585 |
| mobA | CD630_n00850 | 136 -- 195 | 348 -- 400 | -15.2964 | 4.25703 | 5.53585 |
| CD630_06100 | CD630_n00850 | 105 -- 166 | 128 -- 186 | -15.0588 | 4.43918 | 5.53585 |
| CD630_06140 | CD630_n00850 | 16 -- 94 | 112 -- 189 | -15.024 | 4.12519 | 5.53585 |
| CD630_21610 | CD630_n00850 | 19 -- 56 | 325 -- 360 | -14.7553 | 3.16662 | 5.53585 |
| CD630_05460 | CD630_n00850 | 12 -- 113 | 93 -- 202 | -14.239 | 3.98753 | 5.53585 |
| CD630_05870 | CD630_n00850 | 172 -- 198 | 369 -- 400 | -14.1352 | 4.28339 | 5.53585 |
| CD630_21700 | CD630_n00850 | 22 -- 41 | 126 -- 150 | -14.0567 | 3.96650 | 5.53585 |
| CD630_05160 | CD630_n00850 | 125 -- 188 | 114 -- 173 | -14.0297 | 5.03573 | 5.53585 |
| CD630_13790 | CD630_n00850 | 162 -- 200 | 367 -- 400 | -13.8874 | 5.17374 | 5.53585 |
| moaA | CD630_n00850 | 87 -- 119 | 322 -- 359 | -13.4377 | 3.83799 | 5.53585 |
| CD630_04880 | CD630_n00850 | 90 -- 144 | 352 -- 392 | -13.0793 | 4.62291 | 5.53585 |
| CD630_09760 | CD630_n00850 | 117 -- 147 | 323 -- 352 | -13.079 | 4.83410 | 5.53585 |
| CD630_28910 | CD630_n00850 | 117 -- 147 | 323 -- 352 | -13.079 | 4.51664 | 5.53585 |
| CD630_14600 | CD630_n00850 | 55 -- 90 | 323 -- 354 | -13.076 | 5.90348 | 5.53585 |
| CD630_23780 | CD630_n00850 | 80 -- 122 | 363 -- 404 | -12.9986 | 3.91629 | 5.53585 |
| CD630_17281 | CD630_n00850 | 40 -- 94 | 114 -- 172 | -12.9478 | 5.91544 | 5.53585 |
| CD630_31381 | CD630_n00850 | 57 -- 112 | 349 -- 400 | -12.89 | 3.49137 | 5.53585 |
| CD630_10220 | CD630_n00850 | 64 -- 123 | 322 -- 391 | -12.8547 | 4.08233 | 5.53585 |
| CD630_25190 | CD630_n00850 | 174 -- 191 | 84 -- 100 | -12.8442 | 3.19663 | 5.53585 |
| CD630_26920 | CD630_n00850 | 40 -- 111 | 120 -- 189 | -12.8348 | 4.58060 | 5.53585 |
| CD630_27270 | CD630_n00850 | 57 -- 175 | 88 -- 211 | -12.8249 | 3.50892 | 5.53585 |
| CD630_19680 | CD630_n00850 | 77 -- 150 | 342 -- 403 | -12.7952 | 3.58080 | 5.53585 |
| CD630_09750 | CD630_n00850 | 12 -- 43 | 382 -- 405 | -12.7135 | 4.83410 | 5.53585 |
| CD630_28920 | CD630_n00850 | 12 -- 43 | 382 -- 405 | -12.7135 | 4.51664 | 5.53585 |
| kdpD | CD630_n00850 | 152 -- 201 | 94 -- 141 | -12.5256 | 4.27140 | 5.53585 |
| CD630_29500 | CD630_n00850 | 40 -- 60 | 179 -- 198 | -12.4452 | 3.70644 | 5.53585 |
| CD630_02420 | CD630_n00850 | 9 -- 37 | 372 -- 399 | -12.4391 | 3.71048 | 5.53585 |
| CD630_20960 | CD630_n00850 | 35 -- 160 | 272 -- 386 | -12.432 | 5.65940 | 5.53585 |
| CD630_18910 | CD630_n00850 | 178 -- 200 | 183 -- 205 | -12.3679 | 3.30688 | 5.53585 |
| CD630_21720 | CD630_n00850 | 35 -- 177 | 81 -- 214 | -12.3677 | 4.13030 | 5.53585 |
| CD630_20670 | CD630_n00850 | 179 -- 199 | 76 -- 95 | -12.3356 | 4.42757 | 5.53585 |
| CD630_34530 | CD630_n00850 | 4 -- 50 | 116 -- 165 | -12.3328 | 3.83016 | 5.53585 |
| CD630_13960 | CD630_n00850 | 64 -- 117 | 237 -- 295 | -12.3321 | 3.30375 | 5.53585 |
| CD630_05590 | CD630_n00850 | 68 -- 116 | 350 -- 396 | -12.2453 | 3.39670 | 5.53585 |
| CD630_14130 | CD630_n00850 | 60 -- 199 | 213 -- 351 | -12.1421 | 4.57970 | 5.53585 |
| CD630_14340 | CD630_n00850 | 62 -- 103 | 213 -- 259 | -12.1247 | 5.10868 | 5.53585 |
| CD630_09970 | CD630_n00850 | 164 -- 192 | 363 -- 402 | -12.1077 | 3.25917 | 5.53585 |
| CD630_24760 | CD630_n00850 | 95 -- 105 | 390 -- 400 | -12.0906 | 5.19485 | 5.53585 |
| CD630_14700 | CD630_n00850 | 111 -- 157 | 114 -- 161 | -12.039 | 3.98234 | 5.53585 |
| cspD | CD630_n00850 | 108 -- 128 | 375 -- 394 | -12.0332 | 3.23587 | 5.53585 |
| CD630_23070 | CD630_n00850 | 172 -- 187 | 89 -- 103 | -12.0148 | 5.23027 | 5.53585 |
| CD630_16670 | CD630_n00850 | 2 -- 114 | 76 -- 189 | -11.9717 | 3.33553 | 5.53585 |
| rbsC | CD630_n00850 | 32 -- 77 | 349 -- 387 | -11.8642 | 3.71650 | 5.53585 |
| CD630_20460 | CD630_n00850 | 94 -- 185 | 111 -- 207 | -11.8181 | 7.94034 | 5.53585 |
| CD630_29960 | CD630_n00850 | 109 -- 146 | 135 -- 172 | -11.7198 | 4.51904 | 5.53585 |
| acoL | CD630_n00850 | 39 -- 61 | 379 -- 401 | -11.6894 | 3.32169 | 5.53585 |
| CD630_23440 | CD630_n00850 | 146 -- 188 | 353 -- 398 | -11.6445 | 3.34841 | 5.53585 |
| CD630_23770 | CD630_n00850 | 28 -- 77 | 170 -- 207 | -11.6287 | 3.25181 | 5.53585 |
| CD630_27200 | CD630_n00850 | 5 -- 14 | 386 -- 395 | -11.5654 | 3.42516 | 5.53585 |
| CD630_21260 | CD630_n00850 | 138 -- 152 | 43 -- 57 | -11.5234 | 4.22255 | 5.53585 |
| CD630_16540 | CD630_n00850 | 133 -- 184 | 123 -- 173 | -11.5073 | 4.37658 | 5.53585 |
| srlD | CD630_n00850 | 124 -- 148 | 369 -- 393 | -11.4642 | 4.19843 | 5.53585 |
| glnA | CD630_n00850 | 81 -- 127 | 348 -- 400 | -11.3946 | 3.87629 | 5.53585 |
| CD630_30110 | CD630_n00850 | 180 -- 192 | 389 -- 401 | -11.3797 | 3.31041 | 5.53585 |
| CD630_31461 | CD630_n00850 | 141 -- 190 | 97 -- 146 | -11.3404 | 4.41476 | 5.53585 |
| CD630_12650 | CD630_n00850 | 112 -- 138 | 234 -- 258 | -11.3067 | 6.31827 | 5.53585 |
| pilT | CD630_n00850 | 83 -- 97 | 382 -- 396 | -11.2551 | 3.24639 | 5.53585 |
| CD630_29650 | CD630_n00850 | 58 -- 134 | 329 -- 405 | -11.2412 | 3.67128 | 5.53585 |
| CD630_33570 | CD630_n00850 | 161 -- 174 | 387 -- 400 | -11.2195 | 3.95778 | 5.53585 |
| adhE | CD630_n00850 | 1 -- 63 | 76 -- 143 | -11.1639 | 3.06485 | 5.53585 |
| CD630_31220 | CD630_n00850 | 52 -- 87 | 76 -- 124 | -11.1523 | 3.73779 | 5.53585 |
| CD630_06170 | CD630_n00850 | 26 -- 115 | 84 -- 169 | -11.1221 | 4.95366 | 5.53585 |
| CD630_20171 | CD630_n00850 | 72 -- 162 | 246 -- 391 | -11.0386 | 6.87491 | 5.53585 |
| cspC | CD630_n00850 | 108 -- 128 | 375 -- 394 | -10.9599 | 5.48848 | 5.53585 |
| CD630_27120 | CD630_n00850 | 153 -- 174 | 76 -- 95 | -10.9542 | 4.43720 | 5.53585 |
| CD630_22950 | CD630_n00850 | 19 -- 46 | 202 -- 226 | -10.9135 | 3.96883 | 5.53585 |
| CD630_04610 | CD630_n00850 | 25 -- 71 | 116 -- 171 | -10.8961 | 3.83025 | 5.53585 |
| dapB2 | CD630_n00850 | 121 -- 162 | 353 -- 401 | -10.8946 | 3.27218 | 5.53585 |
| CD630_08780 | CD630_n00850 | 141 -- 174 | 81 -- 111 | -10.8896 | 7.76690 | 5.53585 |
| CD630_10050 | CD630_n00850 | 27 -- 45 | 374 -- 391 | -10.8276 | 3.93750 | 5.53585 |
| CD630_06080 | CD630_n00850 | 169 -- 193 | 366 -- 388 | -10.8091 | 3.82315 | 5.53585 |
| CD630_14630 | CD630_n00850 | 144 -- 180 | 212 -- 259 | -10.7997 | 3.34485 | 5.53585 |
| CD630_17970 | CD630_n00850 | 85 -- 127 | 366 -- 393 | -10.784 | 3.68802 | 5.53585 |
| ldh | CD630_n00850 | 157 -- 171 | 8 -- 22 | -10.7095 | 4.35818 | 5.53585 |
| CD630_32520 | CD630_n00850 | 155 -- 198 | 356 -- 403 | -10.7007 | 4.06098 | 5.53585 |
| etfA | CD630_n00850 | 139 -- 159 | 372 -- 401 | -10.6976 | 3.41204 | 5.53585 |
| CD630_27230 | CD630_n00850 | 9 -- 26 | 289 -- 307 | -10.6665 | 3.52462 | 5.53585 |
| CD630_08390 | CD630_n00850 | 69 -- 100 | 373 -- 403 | -10.6179 | 5.16000 | 5.53585 |
| CD630_36781 | CD630_n00850 | 146 -- 178 | 222 -- 253 | -10.5916 | 3.75946 | 5.53585 |
| pilD | CD630_n00850 | 8 -- 31 | 112 -- 133 | -10.5915 | 4.54123 | 5.53585 |
| CD630_07070 | CD630_n00850 | 78 -- 90 | 390 -- 402 | -10.5368 | 3.46414 | 5.53585 |
| CD630_34050 | CD630_n00850 | 8 -- 20 | 3 -- 15 | -10.5329 | 4.03863 | 5.53585 |
| CD630_15900 | CD630_n00850 | 155 -- 171 | 388 -- 405 | -10.5319 | 3.17664 | 5.53585 |
| CD630_23080 | CD630_n00850 | 5 -- 14 | 383 -- 392 | -10.5091 | 5.23027 | 5.53585 |
| CD630_32710 | CD630_n00850 | 136 -- 191 | 167 -- 210 | -10.5031 | 4.74276 | 5.53585 |
| CD630_08100 | CD630_n00850 | 91 -- 121 | 219 -- 257 | -10.48 | 4.39181 | 5.53585 |
| CD630_14110 | CD630_n00850 | 82 -- 105 | 377 -- 400 | -10.4754 | 3.39550 | 5.53585 |
| CD630_21760 | CD630_n00850 | 132 -- 158 | 160 -- 188 | -10.3977 | 4.43846 | 5.53585 |
| CD630_25610 | CD630_n00850 | 83 -- 129 | 92 -- 134 | -10.3969 | 3.02685 | 5.53585 |
| CD630_06200 | CD630_n00850 | 94 -- 125 | 370 -- 401 | -10.3716 | 3.57064 | 5.53585 |
| CD630_02740 | CD630_n00850 | 152 -- 181 | 237 -- 268 | -10.3693 | 5.02277 | 5.53585 |
| CD630_04540 | CD630_n00850 | 38 -- 130 | 83 -- 170 | -10.3647 | 3.33556 | 5.53585 |
| CD630_19942 | CD630_n00850 | 90 -- 141 | 93 -- 141 | -10.3528 | 4.24574 | 5.53585 |
| CD630_19401 | CD630_n00850 | 26 -- 107 | 98 -- 180 | -10.3328 | 4.37084 | 5.53585 |
| spaK | CD630_n00850 | 76 -- 96 | 370 -- 390 | -10.2985 | 3.04530 | 5.53585 |
| CD630_06050 | CD630_n00850 | 190 -- 201 | 389 -- 400 | -10.2723 | 3.59332 | 5.53585 |
| CD630_21620 | CD630_n00850 | 120 -- 166 | 113 -- 163 | -10.2633 | 3.02649 | 5.53585 |
| appC | CD630_n00850 | 152 -- 190 | 259 -- 296 | -10.2295 | 4.50438 | 5.53585 |
| CD630_19801 | CD630_n00850 | 102 -- 120 | 84 -- 103 | -10.1828 | 5.22035 | 5.53585 |
| CD630_05280 | CD630_n00850 | 171 -- 179 | 368 -- 376 | -10.1735 | 3.91665 | 5.53585 |
| kdgT1 | CD630_n00850 | 3 -- 48 | 93 -- 143 | -10.173 | 4.49207 | 5.53585 |
| CD630_23760 | CD630_n00850 | 56 -- 69 | 192 -- 205 | -10.1716 | 4.54307 | 5.53585 |
| uxaA | CD630_n00850 | 8 -- 23 | 85 -- 101 | -10.1529 | 4.26165 | 5.53585 |
| CD630_21770 | CD630_n00850 | 177 -- 196 | 112 -- 130 | -10.124 | 4.02587 | 5.53585 |
| CD630_16640 | CD630_n00850 | 162 -- 201 | 347 -- 379 | -10.1059 | 3.61548 | 5.53585 |
| CD630_14190 | CD630_n00850 | 121 -- 162 | 267 -- 308 | -10.0896 | 3.78562 | 5.53585 |
| malY | CD630_n00850 | 156 -- 176 | 379 -- 399 | -10.0366 | 4.46345 | 5.53585 |
| CD630_28320 | CD630_n00850 | 171 -- 200 | 35 -- 68 | -10.0023 | 3.99184 | 5.53585 |
| nusB | CD630_n00850 | 59 -- 117 | 343 -- 403 | -9.99058 | 4.92798 | 5.53585 |
| CD630_25150 | CD630_n00850 | 42 -- 107 | 124 -- 187 | -9.98928 | 3.79339 | 5.53585 |
| CD630_30140 | CD630_n00850 | 7 -- 32 | 382 -- 400 | -9.98881 | 3.28343 | 5.53585 |
| CD630_17720 | CD630_n00850 | 127 -- 160 | 219 -- 251 | -9.98814 | 3.22910 | 5.53585 |
| bglA | CD630_n00850 | 152 -- 201 | 300 -- 354 | -9.98737 | 4.86402 | 5.53585 |
| dltB | CD630_n00850 | 17 -- 38 | 223 -- 242 | -9.94651 | 3.81485 | 5.53585 |
| CD630_16460 | CD630_n00850 | 130 -- 148 | 76 -- 95 | -9.93194 | 4.64028 | 5.53585 |
| CD630_31580 | CD630_n00850 | 85 -- 104 | 374 -- 394 | -9.9127 | 3.51756 | 5.53585 |
| cbiQ | CD630_n00850 | 6 -- 77 | 60 -- 127 | -9.91226 | 3.79830 | 5.53585 |
| CD630_13370 | CD630_n00850 | 7 -- 21 | 275 -- 289 | -9.88777 | 4.33145 | 5.53585 |
| CD630_20170 | CD630_n00850 | 1 -- 24 | 142 -- 161 | -9.87568 | 6.90201 | 5.53585 |
| CD630_26680 | CD630_n00850 | 170 -- 195 | 371 -- 405 | -9.85083 | 3.53956 | 5.53585 |
| CD630_32720 | CD630_n00850 | 140 -- 169 | 348 -- 375 | -9.84928 | 4.74276 | 5.53585 |
| CD630_09030 | CD630_n00850 | 146 -- 201 | 209 -- 268 | -9.84842 | 4.21282 | 5.53585 |
| CD630_06060 | CD630_n00850 | 82 -- 106 | 76 -- 101 | -9.83029 | 4.94050 | 5.53585 |
| CD630_04770 | CD630_n00850 | 66 -- 102 | 361 -- 397 | -9.82075 | 3.58478 | 5.53585 |
| CD630_25310 | CD630_n00850 | 70 -- 131 | 228 -- 296 | -9.81746 | 3.99861 | 5.53585 |
| CD630_06642 | CD630_n00850 | 144 -- 171 | 91 -- 122 | -9.80717 | 3.32967 | 5.53585 |
| CD630_30620 | CD630_n00850 | 7 -- 27 | 275 -- 296 | -9.80383 | 4.15443 | 5.53585 |
| CD630_11240 | CD630_n00850 | 77 -- 115 | 367 -- 400 | -9.7993 | 3.20274 | 5.53585 |
| CD630_27570 | CD630_n00850 | 1 -- 19 | 167 -- 188 | -9.79667 | 3.46216 | 5.53585 |
| CD630_03420 | CD630_n00850 | 136 -- 175 | 114 -- 157 | -9.7837 | 4.71868 | 5.53585 |
| CD630_15460 | CD630_n00850 | 168 -- 184 | 96 -- 111 | -9.78158 | 6.86296 | 5.53585 |
| CD630_04620 | CD630_n00850 | 157 -- 169 | 136 -- 148 | -9.76782 | 3.45413 | 5.53585 |
| bcp | CD630_n00850 | 77 -- 100 | 107 -- 130 | -9.74392 | 3.16670 | 5.53585 |
| CD630_21090 | CD630_n00850 | 131 -- 198 | 131 -- 195 | -9.72394 | 4.15803 | 5.53585 |
| CD630_15550 | CD630_n00850 | 101 -- 137 | 353 -- 394 | -9.72173 | 3.49019 | 5.53585 |
| CD630_32880 | CD630_n00850 | 104 -- 117 | 388 -- 400 | -9.70464 | 3.41276 | 5.53585 |
| eutV | CD630_n00850 | 135 -- 167 | 264 -- 295 | -9.69505 | 4.55715 | 5.53585 |
| CD630_15360 | CD630_n00850 | 24 -- 37 | 41 -- 54 | -9.68834 | 4.47208 | 5.53585 |
| CD630_32890 | CD630_n00850 | 89 -- 113 | 56 -- 79 | -9.67521 | 3.41276 | 5.53585 |
| CD630_20600 | CD630_n00850 | 150 -- 178 | 125 -- 153 | -9.66826 | 4.20093 | 5.53585 |
| CD630_14360 | CD630_n00850 | 94 -- 103 | 387 -- 396 | -9.66774 | 3.02079 | 5.53585 |
| CD630_32700 | CD630_n00850 | 53 -- 159 | 275 -- 403 | -9.65675 | 4.74276 | 5.53585 |
| CD630_15431 | CD630_n00850 | 9 -- 22 | 380 -- 394 | -9.61405 | 5.29780 | 5.53585 |
| CD630_09040 | CD630_n00850 | 116 -- 137 | 115 -- 136 | -9.61124 | 4.11851 | 5.53585 |
| glgP | CD630_n00850 | 81 -- 129 | 97 -- 140 | -9.61012 | 3.28530 | 5.53585 |
| CD630_22030 | CD630_n00850 | 29 -- 44 | 283 -- 298 | -9.60793 | 3.29934 | 5.53585 |
| CD630_25320 | CD630_n00850 | 91 -- 110 | 76 -- 94 | -9.57822 | 4.29415 | 5.53585 |
| CD630_24920 | CD630_n00850 | 4 -- 29 | 155 -- 181 | -9.56387 | 3.89455 | 5.53585 |
| argS | CD630_n00850 | 65 -- 97 | 97 -- 130 | -9.54053 | 4.96036 | 5.53585 |
| CD630_10950 | CD630_n00850 | 49 -- 65 | 88 -- 109 | -9.52477 | 3.67675 | 5.53585 |
| CD630_20450 | CD630_n00980 | 111 -- 165 | 92 -- 149 | -16.5042 | 3.77388 | 5.31318 |
| CD630_21570 | CD630_n00980 | 151 -- 186 | 117 -- 148 | -16.4301 | 6.48385 | 5.31318 |
| CD630_11241 | CD630_n00980 | 164 -- 193 | 185 -- 213 | -16.3643 | 4.85723 | 5.31318 |
| CD630_14700 | CD630_n00980 | 76 -- 157 | 113 -- 207 | -16.3357 | 3.98234 | 5.31318 |
| xdhA | CD630_n00980 | 47 -- 101 | 112 -- 164 | -16.07 | 4.45147 | 5.31318 |
| CD630_05211 | CD630_n00980 | 112 -- 153 | 219 -- 267 | -16.0082 | 3.97048 | 5.31318 |
| CD630_17281 | CD630_n00980 | 50 -- 94 | 113 -- 158 | -15.9585 | 5.91544 | 5.31318 |
| CD630_23780 | CD630_n00980 | 90 -- 184 | 269 -- 360 | -15.8506 | 3.91629 | 5.31318 |
| oppC | CD630_n00980 | 143 -- 193 | 106 -- 157 | -15.8425 | 3.01695 | 5.31318 |
| CD630_04550 | CD630_n00980 | 89 -- 137 | 308 -- 360 | -15.8011 | 3.05227 | 5.31318 |
| CD630_02790 | CD630_n00980 | 127 -- 170 | 328 -- 363 | -15.5904 | 4.77661 | 5.31318 |
| CD630_05821 | CD630_n00980 | 79 -- 101 | 347 -- 365 | -15.405 | 3.75661 | 5.31318 |
| CD630_13960 | CD630_n00980 | 117 -- 175 | 190 -- 253 | -15.3818 | 3.30375 | 5.31318 |
| CD630_15360 | CD630_n00980 | 105 -- 176 | 276 -- 339 | -15.3685 | 4.47208 | 5.31318 |
| CD630_31530 | CD630_n00980 | 107 -- 146 | 113 -- 151 | -15.3302 | 3.49100 | 5.31318 |
| CD630_05830 | CD630_n00980 | 47 -- 94 | 109 -- 156 | -15.3056 | 3.35360 | 5.31318 |
| CD630_06080 | CD630_n00980 | 156 -- 182 | 184 -- 212 | -15.2631 | 3.82315 | 5.31318 |
| CD630_07840 | CD630_n00980 | 109 -- 163 | 112 -- 160 | -15.1266 | 4.26761 | 5.31318 |
| CD630_32700 | CD630_n00980 | 180 -- 194 | 199 -- 213 | -14.9589 | 4.74276 | 5.31318 |
| CD630_16670 | CD630_n00980 | 122 -- 137 | 349 -- 365 | -14.7821 | 3.33553 | 5.31318 |
| CD630_02720 | CD630_n00980 | 1 -- 50 | 310 -- 354 | -14.6912 | 3.38938 | 5.31318 |
| acoL | CD630_n00980 | 104 -- 143 | 347 -- 386 | -14.4345 | 3.32169 | 5.31318 |
| CD630_21460 | CD630_n00980 | 72 -- 79 | 356 -- 363 | -14.3745 | 4.83005 | 5.31318 |
| CD630_21620 | CD630_n00980 | 4 -- 38 | 117 -- 148 | -14.3735 | 3.02649 | 5.31318 |
| CD630_32710 | CD630_n00980 | 150 -- 188 | 181 -- 213 | -14.3133 | 4.74276 | 5.31318 |
| serS1 | CD630_n00980 | 164 -- 201 | 326 -- 363 | -14.308 | 3.45858 | 5.31318 |
| CD630_28150 | CD630_n00980 | 25 -- 74 | 311 -- 360 | -14.2811 | 4.18447 | 5.31318 |
| tcdC | CD630_n00980 | 112 -- 158 | 281 -- 321 | -14.0718 | 3.76101 | 5.31318 |
| pilD | CD630_n00980 | 3 -- 31 | 111 -- 138 | -14.0685 | 4.54123 | 5.31318 |
| mobA | CD630_n00980 | 181 -- 193 | 353 -- 366 | -14.0053 | 4.25703 | 5.31318 |
| CD630_21260 | CD630_n00980 | 138 -- 152 | 44 -- 58 | -13.9821 | 4.22255 | 5.31318 |
| CD630_21490 | CD630_n00980 | 127 -- 193 | 270 -- 336 | -13.9241 | 3.37462 | 5.31318 |
| CD630_30540 | CD630_n00980 | 116 -- 136 | 306 -- 326 | -13.9187 | 3.63804 | 5.31318 |
| etfA | CD630_n00980 | 88 -- 157 | 300 -- 361 | -13.8758 | 3.41204 | 5.31318 |
| CD630_19790 | CD630_n00980 | 85 -- 125 | 181 -- 216 | -13.8585 | 3.79593 | 5.31318 |
| CD630_14150 | CD630_n00980 | 145 -- 184 | 283 -- 321 | -13.8542 | 3.28251 | 5.31318 |
| CD630_35430 | CD630_n00980 | 142 -- 182 | 121 -- 157 | -13.8354 | 4.28798 | 5.31318 |
| CD630_24770 | CD630_n00980 | 87 -- 100 | 349 -- 363 | -13.7608 | 3.20798 | 5.31318 |
| CD630_36250 | CD630_n00980 | 170 -- 199 | 186 -- 213 | -13.7561 | 4.13224 | 5.31318 |
| CD630_05290 | CD630_n00980 | 19 -- 63 | 113 -- 161 | -13.7324 | 3.97604 | 5.31318 |
| CD630_26680 | CD630_n00980 | 75 -- 104 | 194 -- 223 | -13.7243 | 3.53956 | 5.31318 |
| scrR | CD630_n00980 | 134 -- 148 | 351 -- 365 | -13.6116 | 3.49166 | 5.31318 |
| CD630_21750 | CD630_n00980 | 59 -- 106 | 277 -- 330 | -13.5264 | 3.70626 | 5.31318 |
| CD630_28550 | CD630_n00980 | 90 -- 104 | 347 -- 360 | -13.4751 | 3.51196 | 5.31318 |
| CD630_19000 | CD630_n00980 | 83 -- 116 | 184 -- 216 | -13.4366 | 4.18443 | 5.31318 |
| feoB | CD630_n00980 | 131 -- 197 | 87 -- 148 | -13.4125 | 4.05364 | 5.31318 |
| CD630_03420 | CD630_n00980 | 138 -- 175 | 113 -- 154 | -13.2503 | 4.71868 | 5.31318 |
| cbpA | CD630_n00980 | 87 -- 125 | 328 -- 363 | -13.1894 | 3.03040 | 5.31318 |
| cdu2 | CD630_n00980 | 66 -- 105 | 178 -- 213 | -13.0697 | 2.97335 | 5.31318 |
| CD630_29250 | CD630_n00980 | 102 -- 159 | 113 -- 165 | -13.0495 | 4.61901 | 5.31318 |
| CD630_09410 | CD630_n00980 | 102 -- 159 | 113 -- 165 | -13.0495 | 4.75651 | 5.31318 |
| CD630_06340 | CD630_n00980 | 90 -- 135 | 278 -- 321 | -13.0336 | 4.35983 | 5.31318 |
| CD630_22200 | CD630_n00980 | 141 -- 173 | 115 -- 148 | -13.0054 | 4.18288 | 5.31318 |
| CD630_21770 | CD630_n00980 | 87 -- 196 | 111 -- 212 | -12.9463 | 4.02587 | 5.31318 |
| bcp | CD630_n00980 | 44 -- 97 | 109 -- 164 | -12.8175 | 3.16670 | 5.31318 |
| CD630_05280 | CD630_n00980 | 153 -- 186 | 180 -- 213 | -12.7883 | 3.91665 | 5.31318 |
| CD630_27610 | CD630_n00980 | 5 -- 52 | 278 -- 323 | -12.7803 | 3.96465 | 5.31318 |
| CD630_14130 | CD630_n00980 | 140 -- 194 | 314 -- 362 | -12.762 | 4.57970 | 5.31318 |
| CD630_28181 | CD630_n00980 | 27 -- 79 | 292 -- 350 | -12.7581 | 3.91299 | 5.31318 |
| sspA | CD630_n00980 | 53 -- 109 | 95 -- 156 | -12.7503 | 3.62689 | 5.31318 |
| CD630_25190 | CD630_n00980 | 91 -- 139 | 111 -- 164 | -12.7137 | 3.19663 | 5.31318 |
| CD630_01590 | CD630_n00980 | 167 -- 201 | 184 -- 218 | -12.6501 | 3.45609 | 5.31318 |
| CD630_23440 | CD630_n00980 | 62 -- 84 | 341 -- 360 | -12.6493 | 3.34841 | 5.31318 |
| CD630_30140 | CD630_n00980 | 70 -- 110 | 114 -- 158 | -12.629 | 3.28343 | 5.31318 |
| CD630_34530 | CD630_n00980 | 2 -- 50 | 115 -- 165 | -12.6264 | 3.83016 | 5.31318 |
| CD630_20170 | CD630_n00980 | 123 -- 164 | 113 -- 149 | -12.5997 | 6.90201 | 5.31318 |
| CD630_18960 | CD630_n00980 | 135 -- 153 | 143 -- 161 | -12.5983 | 4.37822 | 5.31318 |
| CD630_06200 | CD630_n00980 | 118 -- 150 | 111 -- 151 | -12.5113 | 3.57064 | 5.31318 |
| CD630_03040 | CD630_n00980 | 51 -- 95 | 112 -- 159 | -12.4872 | 3.51403 | 5.31318 |
| CD630_13780 | CD630_n00980 | 130 -- 176 | 181 -- 216 | -12.4444 | 3.52274 | 5.31318 |
| thiH | CD630_n00980 | 169 -- 196 | 271 -- 295 | -12.4363 | 3.58230 | 5.31318 |
| CD630_09030 | CD630_n00980 | 148 -- 197 | 314 -- 364 | -12.3966 | 4.21282 | 5.31318 |
| CD630_11730 | CD630_n00980 | 88 -- 129 | 310 -- 361 | -12.3686 | 4.80521 | 5.31318 |
| ung | CD630_n00980 | 1 -- 38 | 109 -- 148 | -12.3454 | 3.30908 | 5.31318 |
| CD630_27110 | CD630_n00980 | 133 -- 178 | 106 -- 151 | -12.3348 | 3.78231 | 5.31318 |
| CD630_22991 | CD630_n00980 | 178 -- 196 | 364 -- 382 | -12.319 | 6.03728 | 5.31318 |
| CD630_30100 | CD630_n00980 | 54 -- 80 | 182 -- 209 | -12.2928 | 4.13234 | 5.31318 |
| CD630_15900 | CD630_n00980 | 112 -- 127 | 378 -- 393 | -12.2313 | 3.17664 | 5.31318 |
| def1 | CD630_n00980 | 86 -- 179 | 305 -- 362 | -12.2079 | 3.99720 | 5.31318 |
| CD630_18950 | CD630_n00980 | 180 -- 194 | 86 -- 99 | -12.1735 | 3.40610 | 5.31318 |
| ldh | CD630_n00980 | 85 -- 97 | 350 -- 362 | -12.1414 | 4.35818 | 5.31318 |
| CD630_32050 | CD630_n00980 | 176 -- 199 | 191 -- 213 | -12.0678 | 5.73797 | 5.31318 |
| CD630_33480 | CD630_n00980 | 89 -- 110 | 188 -- 213 | -12.0605 | 3.67613 | 5.31318 |
| LinCd | CD630_n00980 | 8 -- 94 | 308 -- 402 | -12.0416 | 4.33624 | 5.31318 |
| CD630_21090 | CD630_n00980 | 131 -- 188 | 112 -- 161 | -11.9649 | 4.15803 | 5.31318 |
| metQ | CD630_n00980 | 87 -- 116 | 183 -- 213 | -11.9553 | 3.87190 | 5.31318 |
| CD630_14250 | CD630_n00980 | 184 -- 199 | 197 -- 213 | -11.946 | 4.19604 | 5.31318 |
| CD630_34030 | CD630_n00980 | 38 -- 77 | 376 -- 404 | -11.9207 | 3.05766 | 5.31318 |
| CD630_18650 | CD630_n00980 | 167 -- 191 | 297 -- 320 | -11.8965 | 3.75892 | 5.31318 |
| CD630_07870 | CD630_n00980 | 170 -- 179 | 377 -- 386 | -11.844 | 3.22829 | 5.31318 |
| CD630_35260 | CD630_n00980 | 69 -- 130 | 109 -- 164 | -11.8353 | 5.77277 | 5.31318 |
| CD630_23820 | CD630_n00980 | 88 -- 162 | 300 -- 361 | -11.7476 | 3.18419 | 5.31318 |
| CD630_33600 | CD630_n00980 | 139 -- 189 | 272 -- 325 | -11.7169 | 3.73196 | 5.31318 |
| bglA | CD630_n00980 | 59 -- 113 | 184 -- 247 | -11.6455 | 4.50315 | 5.31318 |
| dnaC | CD630_n00980 | 145 -- 175 | 185 -- 213 | -11.6399 | 6.30796 | 5.31318 |
| pgm | CD630_n00980 | 145 -- 176 | 184 -- 217 | -11.6275 | 3.45439 | 5.31318 |
| CD630_23770 | CD630_n00980 | 2 -- 31 | 313 -- 341 | -11.6222 | 3.25181 | 5.31318 |
| CD630_19690 | CD630_n00980 | 115 -- 154 | 114 -- 164 | -11.6162 | 3.85861 | 5.31318 |
| vncR | CD630_n00980 | 138 -- 157 | 190 -- 213 | -11.6078 | 3.93642 | 5.31318 |
| CD630_31010 | CD630_n00980 | 41 -- 98 | 109 -- 164 | -11.6009 | 4.96305 | 5.31318 |
| CD630_15850 | CD630_n00980 | 5 -- 47 | 177 -- 219 | -11.5849 | 4.32968 | 5.31318 |
| CD630_09772 | CD630_n00980 | 39 -- 96 | 111 -- 165 | -11.5683 | 3.66033 | 5.31318 |
| CD630_18311 | CD630_n00980 | 88 -- 113 | 181 -- 210 | -11.5657 | 6.86047 | 5.31318 |
| CD630_23760 | CD630_n00980 | 82 -- 105 | 193 -- 216 | -11.5604 | 4.54307 | 5.31318 |
| CD630_21070 | CD630_n00980 | 185 -- 196 | 351 -- 362 | -11.535 | 5.35019 | 5.31318 |
| CD630_20260 | CD630_n00980 | 86 -- 114 | 183 -- 212 | -11.5295 | 3.84162 | 5.31318 |
| CD630_31461 | CD630_n00980 | 6 -- 25 | 139 -- 156 | -11.5276 | 4.41476 | 5.31318 |
| CD630_36781 | CD630_n00980 | 20 -- 38 | 347 -- 364 | -11.5227 | 3.75946 | 5.31318 |
| CD630_09040 | CD630_n00980 | 91 -- 139 | 111 -- 164 | -11.5059 | 4.11851 | 5.31318 |
| CD630_20670 | CD630_n00980 | 74 -- 112 | 184 -- 226 | -11.4298 | 4.42757 | 5.31318 |
| CD630_15431 | CD630_n00980 | 125 -- 185 | 109 -- 164 | -11.4213 | 5.29780 | 5.31318 |
| CD630_22030 | CD630_n00980 | 7 -- 23 | 51 -- 67 | -11.402 | 3.29934 | 5.31318 |
| CD630_22050 | CD630_n00980 | 5 -- 96 | 114 -- 193 | -11.4008 | 4.87639 | 5.31318 |
| CD630_33560 | CD630_n00980 | 57 -- 94 | 113 -- 149 | -11.3923 | 4.03147 | 5.31318 |
| CD630_20171 | CD630_n00980 | 94 -- 168 | 112 -- 163 | -11.391 | 6.87491 | 5.31318 |
| CD630_17710 | CD630_n00980 | 133 -- 152 | 195 -- 214 | -11.3744 | 4.37688 | 5.31318 |
| CD630_22880 | CD630_n00980 | 175 -- 199 | 132 -- 157 | -11.3654 | 3.62455 | 5.31318 |
| CD630_20660 | CD630_n00980 | 134 -- 167 | 115 -- 157 | -11.3333 | 5.39321 | 5.31318 |
| CD630_09060 | CD630_n00980 | 2 -- 23 | 379 -- 400 | -11.3281 | 4.31834 | 5.31318 |
| CD630_12320 | CD630_n00980 | 37 -- 80 | 315 -- 355 | -11.3199 | 3.28667 | 5.31318 |
| CD630_29960 | CD630_n00980 | 135 -- 201 | 72 -- 145 | -11.316 | 4.51904 | 5.31318 |
| cobC | CD630_n00980 | 164 -- 198 | 111 -- 148 | -11.2858 | 3.15854 | 5.31318 |
| CD630_22850 | CD630_n00980 | 167 -- 201 | 112 -- 156 | -11.2854 | 3.84102 | 5.31318 |
| CD630_28580 | CD630_n00980 | 48 -- 62 | 346 -- 360 | -11.2758 | 4.67130 | 5.31318 |
| CD630_31910 | CD630_n00980 | 86 -- 105 | 193 -- 212 | -11.261 | 3.48314 | 5.31318 |
| CD630_18090 | CD630_n00980 | 10 -- 32 | 191 -- 212 | -11.2492 | 4.23961 | 5.31318 |
| CD630_24590 | CD630_n00980 | 143 -- 150 | 355 -- 362 | -11.2458 | 3.70401 | 5.31318 |
| CD630_16950 | CD630_n00980 | 185 -- 196 | 88 -- 100 | -11.2437 | 4.42569 | 5.31318 |
| CD630_06060 | CD630_n00980 | 102 -- 139 | 184 -- 214 | -11.1239 | 4.94050 | 5.31318 |
| adhE | CD630_n00980 | 35 -- 82 | 315 -- 361 | -11.1225 | 3.06485 | 5.31318 |
| glnA | CD630_n00980 | 151 -- 187 | 115 -- 154 | -11.1194 | 3.87629 | 5.31318 |
| CD630_13400 | CD630_n00980 | 67 -- 118 | 112 -- 157 | -11.1024 | 3.20232 | 5.31318 |
| cprK | CD630_n00980 | 47 -- 56 | 89 -- 98 | -11.0872 | 3.96216 | 5.31318 |
| CD630_32520 | CD630_n00980 | 138 -- 164 | 291 -- 321 | -11.0759 | 4.06098 | 5.31318 |
| CD630_15150 | CD630_n00980 | 8 -- 52 | 110 -- 163 | -11.0672 | 3.35564 | 5.31318 |
| CD630_09970 | CD630_n00980 | 3 -- 39 | 110 -- 147 | -11.0498 | 3.25917 | 5.31318 |
| CD630_06690 | CD630_n00980 | 86 -- 96 | 350 -- 360 | -11.0279 | 5.82709 | 5.31318 |
| CD630_16460 | CD630_n00980 | 15 -- 86 | 172 -- 251 | -10.9693 | 4.64028 | 5.31318 |

| | | | | | | |
|---|---|---|---|---|---|---|
| CD630_04610 | CD630_n00980 | 48 -- 71 | 115 -- 146 | -10.9688 | 3.83025 | 5.31318 |
| flgB | CD630_n00980 | 89 -- 104 | 347 -- 363 | -10.9389 | 3.25736 | 5.31318 |
| opuCA | CD630_n00980 | 87 -- 112 | 182 -- 213 | -10.9324 | 4.07035 | 5.31318 |
| CD630_01560 | CD630_n00980 | 142 -- 151 | 376 -- 385 | -10.9141 | 4.06352 | 5.31318 |
| nusB | CD630_n00980 | 88 -- 157 | 288 -- 360 | -10.9038 | 4.92798 | 5.31318 |
| CD630_19801 | CD630_n00980 | 88 -- 98 | 350 -- 360 | -10.8688 | 5.22035 | 5.31318 |
| CD630_30130 | CD630_n00980 | 1 -- 28 | 182 -- 217 | -10.8571 | 3.28343 | 5.31318 |
| CD630_10830 | CD630_n00980 | 86 -- 95 | 352 -- 362 | -10.849 | 4.82398 | 5.31318 |
| CD630_06150 | CD630_n00980 | 99 -- 128 | 110 -- 131 | -10.8386 | 5.51744 | 5.31318 |
| CD630_13350 | CD630_n00980 | 163 -- 181 | 134 -- 149 | -10.838 | 4.15139 | 5.31318 |
| cdu1 | CD630_n00980 | 87 -- 96 | 352 -- 362 | -10.8279 | 4.46276 | 5.31318 |
| CD630_35780 | CD630_n00980 | 16 -- 67 | 278 -- 337 | -10.8235 | 3.50107 | 5.31318 |
| CD630_24760 | CD630_n00980 | 98 -- 143 | 111 -- 161 | -10.808 | 5.19485 | 5.31318 |
| CD630_01690 | CD630_n00980 | 14 -- 45 | 120 -- 151 | -10.7811 | 4.97234 | 5.31318 |
| CD630_17730 | CD630_n00980 | 76 -- 98 | 111 -- 130 | -10.7437 | 4.73397 | 5.31318 |
| CD630_19401 | CD630_n00980 | 35 -- 82 | 125 -- 172 | -10.7356 | 4.37084 | 5.31318 |
| CD630_08390 | CD630_n00980 | 83 -- 100 | 346 -- 359 | -10.7214 | 5.16000 | 5.31318 |
| CD630_06060 | CD630_n00980 | 41 -- 59 | 349 -- 362 | -10.7199 | 4.94050 | 5.31318 |
| CD630_02430 | CD630_n00980 | 39 -- 84 | 117 -- 145 | -10.6616 | 3.71048 | 5.31318 |
| CD630_22890 | CD630_n00980 | 40 -- 47 | 353 -- 360 | -10.6265 | 3.75591 | 5.31318 |
| CD630_25040 | CD630_n00980 | 144 -- 177 | 26 -- 61 | -10.607 | 3.43898 | 5.31318 |
| CD630_09100 | CD630_n00980 | 2 -- 50 | 27 -- 68 | -10.6054 | 5.92260 | 5.31318 |
| CD630_25610 | CD630_n00980 | 44 -- 94 | 279 -- 320 | -10.5909 | 3.02685 | 5.31318 |
| malY | CD630_n00980 | 77 -- 121 | 316 -- 364 | -10.5868 | 4.46345 | 5.31318 |
| srlEb | CD630_n00980 | 91 -- 111 | 194 -- 213 | -10.548 | 3.50140 | 5.31318 |
| CD630_09760 | CD630_n00980 | 90 -- 125 | 180 -- 213 | -10.5355 | 4.83410 | 5.31318 |
| CD630_28910 | CD630_n00980 | 90 -- 125 | 180 -- 213 | -10.5355 | 4.51664 | 5.31318 |
| CD630_30120 | CD630_n00980 | 43 -- 100 | 111 -- 164 | -10.5294 | 4.04859 | 5.31318 |
| gabT | CD630_n00980 | 36 -- 61 | 218 -- 246 | -10.5131 | 3.67067 | 5.31318 |
| CD630_08940 | CD630_n00980 | 75 -- 91 | 74 -- 90 | -10.5078 | 3.72225 | 5.31318 |
| CD630_10050 | CD630_n00980 | 44 -- 159 | 114 -- 209 | -10.5033 | 3.93750 | 5.31318 |
| CD630_34080 | CD630_n00980 | 168 -- 201 | 107 -- 148 | -10.4943 | 3.20224 | 5.31318 |
| CD630_02830 | CD630_n00980 | 182 -- 198 | 300 -- 317 | -10.4815 | 3.57106 | 5.31318 |
| CD630_08510 | CD630_n00980 | 44 -- 52 | 350 -- 358 | -10.4612 | 5.16329 | 5.31318 |
| CD630_21760 | CD630_n00980 | 64 -- 74 | 89 -- 98 | -10.4599 | 4.43846 | 5.31318 |
| CD630_27310 | CD630_n00980 | 135 -- 177 | 108 -- 148 | -10.4522 | 4.01622 | 5.31318 |
| CD630_18010 | CD630_n00980 | 154 -- 186 | 132 -- 163 | -10.4453 | 4.21967 | 5.31318 |
| cwp29 | CD630_n00980 | 20 -- 56 | 185 -- 225 | -10.4418 | 4.40865 | 5.31318 |
| CD630_14050 | CD630_n00980 | 186 -- 201 | 198 -- 213 | -10.4163 | 3.30894 | 5.31318 |
| CD630_08090 | CD630_n00980 | 143 -- 174 | 109 -- 139 | -10.4078 | 3.57167 | 5.31318 |
| CD630_05230 | CD630_n00980 | 3 -- 15 | 351 -- 363 | -10.4061 | 3.73787 | 5.31318 |
| tcdA | CD630_n00980 | 36 -- 63 | 182 -- 212 | -10.3819 | 4.70593 | 5.31318 |
| CD630_31880 | CD630_n00980 | 15 -- 34 | 22 -- 42 | -10.3237 | 3.97522 | 5.31318 |
| CD630_05150 | CD630_n00980 | 87 -- 104 | 347 -- 364 | -10.3129 | 3.69875 | 5.31318 |
| CD630_06190 | CD630_n00980 | 141 -- 151 | 48 -- 58 | -10.2851 | 3.57064 | 5.31318 |
| CD630_33190 | CD630_n00980 | 115 -- 126 | 184 -- 195 | -10.2718 | 4.13899 | 5.31318 |
| opuCC | CD630_n00980 | 95 -- 144 | 115 -- 164 | -10.2683 | 4.07035 | 5.31318 |
| CD630_06350 | CD630_n00980 | 138 -- 167 | 184 -- 213 | -10.2618 | 4.35983 | 5.31318 |
| CD630_24591 | CD630_n00980 | 88 -- 96 | 352 -- 360 | -10.2536 | 3.54497 | 5.31318 |
| cwp26 | CD630_n00980 | 131 -- 161 | 142 -- 172 | -10.2308 | 4.44377 | 5.31318 |
| CD630_26050 | CD630_n00980 | 173 -- 200 | 183 -- 212 | -10.2244 | 6.99851 | 5.31318 |
| CD630_27200 | CD630_n00980 | 1 -- 36 | 278 -- 302 | -10.2084 | 3.42516 | 5.31318 |
| CD630_06100 | CD630_n00980 | 65 -- 74 | 381 -- 390 | -10.2039 | 4.43918 | 5.31318 |
| CD630_33300 | CD630_n00980 | 190 -- 197 | 356 -- 363 | -10.1866 | 3.79149 | 5.31318 |
| dltB | CD630_n00980 | 7 -- 25 | 380 -- 398 | -10.1797 | 3.81485 | 5.31318 |
| CD630_18400 | CD630_n00980 | 121 -- 158 | 112 -- 149 | -10.1631 | 4.67962 | 5.31318 |
| CD630_15460 | CD630_n00980 | 88 -- 97 | 352 -- 362 | -10.1549 | 6.86296 | 5.31318 |
| CD630_32040 | CD630_n00980 | 24 -- 42 | 257 -- 281 | -10.1279 | 4.93516 | 5.31318 |
| CD630_27120 | CD630_n00980 | 116 -- 161 | 273 -- 321 | -10.1161 | 4.43720 | 5.31318 |
| CD630_28290 | CD630_n00980 | 87 -- 166 | 281 -- 360 | -10.1105 | 5.54192 | 5.31318 |
| CD630_05880 | CD630_n00980 | 91 -- 109 | 197 -- 213 | -10.0578 | 4.28339 | 5.31318 |
| etfA | CD630_n00980 | 124 -- 163 | 179 -- 212 | -10.0345 | 4.80521 | 5.31318 |
| CD630_13360 | CD630_n00980 | 46 -- 99 | 111 -- 158 | -10.0309 | 3.77330 | 5.31318 |
| CD630_18910 | CD630_n00980 | 121 -- 136 | 349 -- 364 | -10.0229 | 3.30688 | 5.31318 |
| CD630_28320 | CD630_n00980 | 1 -- 10 | 349 -- 358 | -10.0012 | 3.99184 | 5.31318 |
| pilC | CD630_n00980 | 87 -- 101 | 308 -- 322 | -9.9873 | 3.54230 | 5.31318 |
| garK | CD630_n00980 | 154 -- 160 | 357 -- 363 | -9.97067 | 3.72836 | 5.31318 |
| cbiQ | CD630_n00980 | 36 -- 58 | 83 -- 100 | -9.94487 | 3.79830 | 5.31318 |
| srlD | CD630_n00980 | 109 -- 148 | 319 -- 357 | -9.92225 | 4.19843 | 5.31318 |
| CD630_15060 | CD630_n00980 | 173 -- 191 | 227 -- 244 | -9.91105 | 3.69396 | 5.31318 |
| CD630_30730 | CD630_n00980 | 186 -- 201 | 382 -- 398 | -9.89251 | 3.13236 | 5.31318 |
| cynT | CD630_n00980 | 146 -- 169 | 108 -- 127 | -9.88715 | 4.21370 | 5.31318 |
| CD630_25320 | CD630_n00980 | 144 -- 173 | 184 -- 213 | -9.88459 | 4.29415 | 5.31318 |
| xylR | CD630_n00980 | 68 -- 92 | 379 -- 403 | -9.86475 | 3.95671 | 5.31318 |
| CD630_18610 | CD630_n00980 | 19 -- 52 | 318 -- 358 | -9.85098 | 4.15582 | 5.31318 |
| CD630_18781 | CD630_n00980 | 109 -- 125 | 196 -- 212 | -9.84787 | 3.77660 | 5.31318 |
| CD630_05950 | CD630_n00980 | 34 -- 69 | 180 -- 212 | -9.84437 | 3.31669 | 5.31318 |
| dapB2 | CD630_n00980 | 139 -- 196 | 299 -- 360 | -9.83576 | 3.27218 | 5.31318 |
| CD630_27280 | CD630_n00980 | 122 -- 141 | 382 -- 401 | -9.83443 | 3.24738 | 5.31318 |
| CD630_15890 | CD630_n00980 | 146 -- 166 | 295 -- 316 | -9.81823 | 3.39962 | 5.31318 |
| CD630_16150 | CD630_n00980 | 81 -- 89 | 378 -- 386 | -9.77006 | 4.20614 | 5.31318 |
| CD630_24920 | CD630_n00980 | 89 -- 150 | 112 -- 164 | -9.75926 | 3.89455 | 5.31318 |
| CD630_21030 | CD630_n00980 | 57 -- 107 | 114 -- 163 | -9.75456 | 3.08733 | 5.31318 |
| CD630_22720 | CD630_n00980 | 122 -- 158 | 333 -- 363 | -9.74845 | 3.81804 | 5.31318 |
| CD630_22510 | CD630_n00980 | 114 -- 127 | 225 -- 238 | -9.74758 | 11.42763 | 5.31318 |
| CD630_25620 | CD630_n00980 | 90 -- 146 | 311 -- 360 | -9.73103 | 3.02685 | 5.31318 |
| cspC | CD630_n00980 | 79 -- 89 | 272 -- 282 | -9.72029 | 5.48848 | 5.31318 |
| CD630_20980 | CD630_n00980 | 101 -- 110 | 87 -- 96 | -9.71741 | 8.12847 | 5.31318 |
| CD630_27230 | CD630_n00980 | 84 -- 92 | 378 -- 386 | -9.70183 | 3.52462 | 5.31318 |
| CD630_15550 | CD630_n00980 | 83 -- 133 | 278 -- 326 | -9.70171 | 3.49019 | 5.31318 |
| CD630_09840 | CD630_n00980 | 162 -- 177 | 44 -- 58 | -9.69133 | 4.50009 | 5.31318 |
| CD630_20460 | CD630_n00980 | 97 -- 147 | 310 -- 360 | -9.68931 | 7.94034 | 5.31318 |
| CD630_33650 | CD630_n00980 | 172 -- 180 | 379 -- 387 | -9.68904 | 4.80565 | 5.31318 |
| terD | CD630_n00980 | 115 -- 127 | 111 -- 123 | -9.67698 | 3.33148 | 5.31318 |
| CD630_11220 | CD630_n00980 | 79 -- 105 | 125 -- 149 | -9.67653 | 3.52327 | 5.31318 |
| CD630_27570 | CD630_n00980 | 128 -- 141 | 113 -- 126 | -9.64587 | 3.46216 | 5.31318 |
| CD630_16450 | CD630_n00980 | 149 -- 163 | 87 -- 100 | -9.64192 | 4.57547 | 5.31318 |
| CD630_22500 | CD630_n00980 | 194 -- 201 | 352 -- 359 | -9.6283 | 4.03688 | 5.31318 |
| CD630_14600 | CD630_n00980 | 65 -- 101 | 126 -- 163 | -9.62582 | 5.90348 | 5.31318 |
| gcvTPA | CD630_n00980 | 170 -- 178 | 89 -- 97 | -9.61805 | 4.77229 | 5.31318 |
| CD630_22710 | CD630_n00980 | 84 -- 115 | 180 -- 213 | -9.59639 | 4.05623 | 5.31318 |
| CD630_14180 | CD630_n00980 | 87 -- 95 | 352 -- 360 | -9.58002 | 3.93696 | 5.31318 |
| CD630_34520 | CD630_n00980 | 77 -- 113 | 114 -- 146 | -9.57564 | 3.83016 | 5.31318 |
| CD630_32990 | CD630_n00980 | 35 -- 98 | 188 -- 261 | -9.54162 | 5.24925 | 5.31318 |
| CD630_04410 | CD630_n00980 | 83 -- 96 | 352 -- 363 | -9.53416 | 3.90451 | 5.31318 |
| CD630_14260 | CD630_n00980 | 39 -- 83 | 197 -- 250 | -9.52346 | 3.22214 | 5.31318 |
| CD630_23070 | CD630_n00980 | 5 -- 16 | 30 -- 41 | -9.52112 | 5.23027 | 5.31318 |
| CD630_08130 | CD630_n00980 | 73 -- 103 | 180 -- 208 | -9.517 | 4.63831 | 5.31318 |
| CD630_09562 | CD630_n00980 | 178 -- 195 | 365 -- 382 | -9.50053 | 3.94307 | 5.31318 |
| CD630_13960 | CD630_n01060 | 20 -- 64 | 65 -- 113 | -17.8196 | 3.30375 | 3.97522 |
| CD630_08950 | CD630_n01060 | 4 -- 57 | 59 -- 113 | -16.5939 | 3.71905 | 3.97522 |
| CD630_07070 | CD630_n01060 | 4 -- 46 | 79 -- 118 | -15.8427 | 3.46414 | 3.97522 |
| oppB | CD630_n01060 | 124 -- 154 | 35 -- 67 | -15.7683 | 2.98328 | 3.97522 |
| CD630_29250 | CD630_n01060 | 4 -- 29 | 95 -- 118 | -14.6139 | 4.61901 | 3.97522 |
| CD630_09410 | CD630_n01060 | 4 -- 29 | 95 -- 118 | -14.6139 | 4.75651 | 3.97522 |
| CD630_19030 | CD630_n01060 | 150 -- 168 | 95 -- 115 | -14.5492 | 3.34240 | 3.97522 |
| CD630_04410 | CD630_n01060 | 52 -- 93 | 63 -- 108 | -14.4268 | 3.90451 | 3.97522 |
| CD630_13950 | CD630_n01060 | 148 -- 197 | 68 -- 117 | -14.2965 | 4.25540 | 3.97522 |
| CD630_08660 | CD630_n01060 | 143 -- 199 | 50 -- 116 | -14.2344 | 3.86467 | 3.97522 |
| CD630_22500 | CD630_n01060 | 138 -- 172 | 39 -- 69 | -13.9604 | 4.03688 | 3.97522 |
| cdtR | CD630_n01060 | 4 -- 42 | 82 -- 112 | -12.9763 | 5.84368 | 3.97522 |
| CD630_20440 | CD630_n01060 | 138 -- 192 | 52 -- 117 | -12.9609 | 3.77388 | 3.97522 |
| CD630_28290 | CD630_n01060 | 15 -- 52 | 79 -- 115 | -12.8549 | 5.54192 | 3.97522 |
| CD630_05830 | CD630_n01060 | 137 -- 184 | 67 -- 119 | -12.7634 | 3.35360 | 3.97522 |
| spaK | CD630_n01060 | 138 -- 193 | 50 -- 98 | -12.6173 | 3.04530 | 3.97522 |
| CD630_05280 | CD630_n01060 | 8 -- 61 | 53 -- 110 | -12.5191 | 3.91665 | 3.97522 |
| CD630_01690 | CD630_n01060 | 153 -- 177 | 92 -- 117 | -12.4555 | 4.97234 | 3.97522 |
| CD630_13020 | CD630_n01060 | 114 -- 147 | 97 -- 115 | -12.3603 | 3.52267 | 3.97522 |
| CD630_01590 | CD630_n01060 | 17 -- 45 | 92 -- 121 | -12.1923 | 3.45609 | 3.97522 |
| CD630_10220 | CD630_n01060 | 79 -- 107 | 60 -- 87 | -12.1577 | 4.08233 | 3.97522 |
| CD630_20170 | CD630_n01060 | 2 -- 25 | 94 -- 114 | -12.0988 | 6.90201 | 3.97522 |
| CD630_06540 | CD630_n01060 | 28 -- 53 | 37 -- 62 | -12.0424 | 3.54673 | 3.97522 |
| CD630_30100 | CD630_n01060 | 148 -- 200 | 66 -- 118 | -12.0352 | 4.13234 | 3.97522 |
| CD630_09100 | CD630_n01060 | 1 -- 86 | 38 -- 115 | -12.034 | 5.92260 | 3.97522 |
| appC | CD630_n01060 | 48 -- 68 | 96 -- 115 | -11.8305 | 4.50438 | 3.97522 |
| CD630_30190 | CD630_n01060 | 118 -- 150 | 69 -- 113 | -11.7723 | 3.27394 | 3.97522 |
| CD630_06920 | CD630_n01060 | 7 -- 59 | 51 -- 119 | -11.7504 | 3.13260 | 3.97522 |
| cwp26 | CD630_n01060 | 132 -- 179 | 58 -- 112 | -11.7295 | 4.44377 | 3.97522 |
| CD630_23930 | CD630_n01060 | 1 -- 24 | 95 -- 113 | -11.5749 | 3.05650 | 3.97522 |
| CD630_36690 | CD630_n01060 | 140 -- 196 | 38 -- 116 | -11.546 | 3.53761 | 3.97522 |
| CD630_13580 | CD630_n01060 | 37 -- 133 | 37 -- 113 | -11.5166 | 3.80715 | 3.97522 |
| CD630_27280 | CD630_n01060 | 5 -- 70 | 34 -- 117 | -11.4449 | 3.24738 | 3.97522 |
| CD630_13790 | CD630_n01060 | 94 -- 135 | 73 -- 113 | -11.3944 | 5.17374 | 3.97522 |
| CD630_07400 | CD630_n01060 | 1 -- 15 | 108 -- 121 | -11.3777 | 4.52273 | 3.97522 |
| CD630_06100 | CD630_n01060 | 136 -- 200 | 48 -- 122 | -11.3553 | 4.43918 | 3.97522 |
| CD630_18010 | CD630_n01060 | 21 -- 49 | 91 -- 122 | -11.2797 | 4.21967 | 3.97522 |
| CD630_21720 | CD630_n01060 | 17 -- 46 | 51 -- 83 | -11.2733 | 4.13030 | 3.97522 |
| CD630_16940 | CD630_n01060 | 10 -- 80 | 43 -- 120 | -11.1301 | 4.14665 | 3.97522 |
| CD630_04341 | CD630_n01060 | 21 -- 50 | 41 -- 68 | -11.1275 | 4.29262 | 3.97522 |
| CD630_34520 | CD630_n01060 | 44 -- 55 | 107 -- 118 | -11.1257 | 3.83016 | 3.97522 |
| CD630_16330 | CD630_n01060 | 164 -- 182 | 97 -- 116 | -11.105 | 4.11180 | 3.97522 |
| xylR | CD630_n01060 | 16 -- 57 | 88 -- 119 | -11.012 | 3.95671 | 3.97522 |
| CD630_21260 | CD630_n01060 | 131 -- 169 | 81 -- 118 | -10.9814 | 4.22255 | 3.97522 |
| CD630_22510 | CD630_n01060 | 114 -- 137 | 96 -- 115 | -10.8351 | 11.42763 | 3.97522 |
| CD630_02790 | CD630_n01060 | 34 -- 55 | 98 -- 119 | -10.7864 | 4.77661 | 3.97522 |
| CD630_28560 | CD630_n01060 | 114 -- 136 | 90 -- 111 | -10.7559 | 3.96803 | 3.97522 |
| CD630_06340 | CD630_n01060 | 5 -- 58 | 40 -- 85 | -10.7536 | 4.35983 | 3.97522 |
| CD630_14620 | CD630_n01060 | 18 -- 47 | 81 -- 119 | -10.7398 | 4.43651 | 3.97522 |
| CD630_24920 | CD630_n01060 | 4 -- 18 | 58 -- 72 | -10.7388 | 3.89455 | 3.97522 |
| CD630_18610 | CD630_n01060 | 138 -- 176 | 67 -- 109 | -10.6029 | 4.15582 | 3.97522 |
| CD630_13530 | CD630_n01060 | 1 -- 36 | 82 -- 120 | -10.5736 | 4.09832 | 3.97522 |
| CD630_26250 | CD630_n01060 | 140 -- 186 | 70 -- 118 | -10.5628 | 5.03379 | 3.97522 |
| CD630_14610 | CD630_n01060 | 152 -- 176 | 96 -- 119 | -10.4575 | 4.89428 | 3.97522 |
| CD630_16160 | CD630_n01060 | 177 -- 197 | 96 -- 116 | -10.4014 | 6.11801 | 3.97522 |
| CD630_23450 | CD630_n01060 | 10 -- 78 | 52 -- 119 | -10.3987 | 3.27218 | 3.97522 |
| CD630_22880 | CD630_n01060 | 34 -- 73 | 89 -- 121 | -10.3706 | 3.62455 | 3.97522 |
| CD630_07470 | CD630_n01060 | 1 -- 38 | 45 -- 82 | -10.3646 | 3.37143 | 3.97522 |
| dnaD | CD630_n01060 | 145 -- 177 | 81 -- 119 | -10.3433 | 3.56078 | 3.97522 |
| CD630_22550 | CD630_n01060 | 20 -- 74 | 40 -- 110 | -10.3196 | 3.62379 | 3.97522 |
| CD630_17720 | CD630_n01060 | 3 -- 35 | 90 -- 122 | -10.315 | 3.22910 | 3.97522 |
| srlD | CD630_n01060 | 167 -- 190 | 59 -- 83 | -10.2655 | 4.19843 | 3.97522 |
| CD630_27240 | CD630_n01060 | 141 -- 163 | 97 -- 117 | -10.1765 | 3.52462 | 3.97522 |
| CD630_15060 | CD630_n01060 | 166 -- 189 | 95 -- 114 | -10.1123 | 3.69396 | 3.97522 |
| CD630_21530 | CD630_n01060 | 148 -- 196 | 39 -- 85 | -10.0501 | 3.39774 | 3.97522 |
| CD630_22050 | CD630_n01060 | 2 -- 28 | 88 -- 116 | -10.0041 | 4.87639 | 3.97522 |
| CD630_10800 | CD630_n01060 | 123 -- 164 | 55 -- 88 | -9.98704 | 5.30319 | 3.97522 |
| CD630_20260 | CD630_n01060 | 7 -- 15 | 77 -- 85 | -9.97574 | 3.84162 | 3.97522 |
| CD630_09750 | CD630_n01060 | 22 -- 33 | 61 -- 72 | -9.96554 | 4.83410 | 3.97522 |
| CD630_28920 | CD630_n01060 | 22 -- 33 | 61 -- 72 | -9.96554 | 4.51664 | 3.97522 |
| CD630_18930 | CD630_n01060 | 9 -- 37 | 89 -- 116 | -9.95702 | 5.32127 | 3.97522 |
| rolA | CD630_n01060 | 17 -- 69 | 20 -- 69 | -9.86645 | 4.56362 | 3.97522 |
| CD630_16640 | CD630_n01060 | 95 -- 124 | 59 -- 83 | -9.82957 | 3.61548 | 3.97522 |
| CD630_36781 | CD630_n01060 | 121 -- 197 | 29 -- 114 | -9.79616 | 3.75946 | 3.97522 |
| CD630_31530 | CD630_n01060 | 9 -- 44 | 51 -- 85 | -9.79251 | 3.49100 | 3.97522 |
| CD630_27360 | CD630_n01060 | 88 -- 139 | 59 -- 114 | -9.78208 | 3.79570 | 3.97522 |
| CD630_17970 | CD630_n01060 | 2 -- 20 | 43 -- 62 | -9.74769 | 3.68802 | 3.97522 |
| CD630_23080 | CD630_n01060 | 24 -- 69 | 81 -- 119 | -9.7335 | 5.23027 | 3.97522 |
| CD630_03920 | CD630_n01060 | 105 -- 159 | 38 -- 113 | -9.72634 | 4.15934 | 3.97522 |
| CD630_21330 | CD630_n01060 | 194 -- 201 | 95 -- 102 | -9.72182 | 4.10111 | 3.97522 |
| CD630_26630 | CD630_n01060 | 8 -- 106 | 5 -- 86 | -9.72105 | 4.61557 | 3.97522 |
| spoVFB | CD630_n01060 | 51 -- 65 | 96 -- 111 | -9.71751 | 4.62758 | 3.97522 |
| CD630_21760 | CD630_n01060 | 177 -- 189 | 60 -- 72 | -9.67866 | 4.43846 | 3.97522 |
| CD630_23070 | CD630_n01060 | 169 -- 199 | 51 -- 71 | -9.65077 | 5.23027 | 3.97522 |
| appF | CD630_n01060 | 54 -- 83 | 53 -- 83 | -9.62352 | 4.25570 | 3.97522 |
| CD630_15200 | CD630_n01060 | 132 -- 150 | 91 -- 109 | -9.59836 | 4.00442 | 3.97522 |
| CD630_19850 | CD630_n01060 | 101 -- 124 | 62 -- 85 | -9.54968 | 6.88486 | 3.97522 |
| CD630_16530 | CD630_n01060 | 116 -- 168 | 58 -- 120 | -9.54908 | 4.37658 | 3.97522 |
| CD630_34030 | CD630_n01060 | 30 -- 56 | 81 -- 116 | -9.5456 | 3.05766 | 3.97522 |
| CD630_30730 | CD630_n01060 | 17 -- 36 | 99 -- 119 | -9.53985 | 3.13236 | 3.97522 |
| CD630_07121 | CD630_n01060 | 2 -- 9 | 97 -- 104 | -9.5309 | 3.26177 | 3.97522 |
| CD630_21090 | CD630_n01060 | 127 -- 148 | 97 -- 118 | -9.50348 | 4.15803 | 3.97522 |
| CD630_07870 | CD630_RNA_15 | 175 -- 197 | 62 -- 83 | -11.9328 | 3.22829 | 4.12920 |
| CD630_09100 | CD630_RNA_15 | 12 -- 26 | 62 -- 76 | -11.3763 | 5.92260 | 4.12920 |
| CD630_09770 | CD630_RNA_15 | 19 -- 35 | 54 -- 70 | -10.8355 | 3.22106 | 4.12920 |
| CD630_21020 | CD630_RNA_15 | 90 -- 108 | 41 -- 60 | -10.6698 | 4.28666 | 4.12920 |
| CD630_30130 | CD630_RNA_15 | 126 -- 137 | 72 -- 83 | -10.1514 | 3.28343 | 4.12920 |
| CD630_24591 | CD630_RNA_15 | 41 -- 48 | 68 -- 75 | -10.0005 | 3.54497 | 4.12920 |
| terD | CD630_RNA_15 | 145 -- 159 | 71 -- 83 | -9.97063 | 3.33148 | 4.12920 |
| garK | CD630_RNA_15 | 13 -- 31 | 44 -- 60 | -9.96532 | 3.72836 | 4.12920 |
| CD630_05821 | CD630_RNA_15 | 102 -- 115 | 39 -- 53 | -9.95863 | 3.75661 | 4.12920 |
| CD630_20260 | CD630_RNA_15 | 1 -- 12 | 39 -- 50 | -9.84439 | 3.84162 | 4.12920 |
| CD630_31382 | CD630_RNA_15 | 174 -- 199 | 1 -- 25 | -9.71376 | 3.49137 | 4.12920 |
| ung | CD630_RNA_15 | 3 -- 13 | 43 -- 53 | -9.60361 | 3.30908 | 4.12920 |
| CD630_08140 | CD630_RNA_15 | 12 -- 19 | 68 -- 75 | -9.50979 | 3.30159 | 4.12920 |
| CD630_03040 | CD630_RNA_5 | 111 -- 149 | 5 -- 34 | -14.4907 | 3.51403 | 4.62291 |
| CD630_06060 | CD630_RNA_5 | 41 -- 60 | 3 -- 23 | -13.8371 | 4.94050 | 4.62291 |
| CD630_23780 | CD630_RNA_5 | 141 -- 175 | 82 -- 104 | -13.677 | 3.91629 | 4.62291 |
| CD630_04770 | CD630_RNA_5 | 36 -- 52 | 3 -- 21 | -12.431 | 3.58478 | 4.62291 |
| cobS | CD630_RNA_5 | 138 -- 153 | 6 -- 21 | -11.9389 | 3.67481 | 4.62291 |
| tcdC | CD630_RNA_5 | 132 -- 158 | 3 -- 32 | -11.8298 | 3.76101 | 4.62291 |
| CD630_21570 | CD630_RNA_5 | 4 -- 19 | 1 -- 16 | -11.7049 | 6.48385 | 4.62291 |
| CD630_27280 | CD630_RNA_5 | 46 -- 68 | 87 -- 105 | -11.4714 | 3.24738 | 4.62291 |
| CD630_14990 | CD630_RNA_5 | 148 -- 167 | 3 -- 23 | -11.4443 | 3.78619 | 4.62291 |
| pilD | CD630_RNA_5 | 3 -- 16 | 8 -- 21 | -11.262 | 4.54123 | 4.62291 |
| CD630_16530 | CD630_RNA_5 | 60 -- 109 | 5 -- 55 | -11.0013 | 4.37658 | 4.62291 |
| CD630_08070 | CD630_RNA_5 | 103 -- 131 | 22 -- 47 | -10.9211 | 4.87819 | 4.62291 |
| CD630_31220 | CD630_RNA_5 | 9 -- 27 | 15 -- 34 | -10.7907 | 3.73779 | 4.62291 |
| iorA | CD630_RNA_5 | 56 -- 107 | 7 -- 60 | -10.7156 | 3.83173 | 4.62291 |
| CD630_09760 | CD630_RNA_5 | 134 -- 150 | 16 -- 33 | -10.586 | 4.83410 | 4.62291 |
| CD630_28910 | CD630_RNA_5 | 134 -- 150 | 16 -- 33 | -10.586 | 4.51664 | 4.62291 |
| CD630_23080 | CD630_RNA_5 | 65 -- 81 | 2 -- 18 | -10.4841 | 5.23027 | 4.62291 |
| CD630_19904 | CD630_RNA_5 | 25 -- 31 | 100 -- 106 | -10.451 | 5.96526 | 4.62291 |
| CD630_16150 | CD630_RNA_5 | 1 -- 21 | 15 -- 30 | -10.3716 | 4.20614 | 4.62291 |
| cdtR | CD630_RNA_5 | 21 -- 33 | 96 -- 107 | -10.1277 | 5.84368 | 4.62291 |
| ppnK | CD630_RNA_5 | 126 -- 154 | 6 -- 43 | -10.0924 | 4.36817 | 4.62291 |
| CD630_32890 | CD630_RNA_5 | 139 -- 155 | 3 -- 20 | -10.0446 | 3.41276 | 4.62291 |
| CD630_18930 | CD630_RNA_5 | 147 -- 166 | 3 -- 23 | -9.99346 | 5.32127 | 4.62291 |
| terD | CD630_RNA_5 | 146 -- 169 | 3 -- 30 | -9.78005 | 3.33148 | 4.62291 |
| CD630_24591 | CD630_s0010 | 13 -- 48 | 1 -- 31 | -13.6811 | 3.54497 | 4.92433 |
| serS1 | CD630_s0010 | 158 -- 171 | 180 -- 193 | -13.4207 | 3.45858 | 4.92433 |
| CD630_19800 | CD630_s0010 | 60 -- 84 | 64 -- 87 | -13.14 | 5.22035 | 4.92433 |
| CD630_17160 | CD630_s0010 | 89 -- 102 | 179 -- 192 | -13.1318 | 3.50228 | 4.92433 |
| CD630_18610 | CD630_s0010 | 79 -- 106 | 166 -- 197 | -13.0547 | 4.15582 | 4.92433 |
| pilC | CD630_s0010 | 125 -- 141 | 180 -- 195 | -12.9801 | 3.54230 | 4.92433 |
| CD630_21240 | CD630_s0010 | 84 -- 132 | 163 -- 196 | -12.4402 | 5.88879 | 4.92433 |
| CD630_09100 | CD630_s0010 | 1 -- 33 | 2 -- 31 | -12.41 | 5.92260 | 4.92433 |
| CD630_23450 | CD630_s0010 | 102 -- 128 | 3 -- 30 | -12.3445 | 3.27218 | 4.92433 |
| CD630_27280 | CD630_s0010 | 153 -- 171 | 1 -- 20 | -12.1814 | 3.24738 | 4.92433 |
| CD630_33580 | CD630_s0010 | 103 -- 129 | 65 -- 87 | -11.8249 | 3.95778 | 4.92433 |
| CD630_18120 | CD630_s0010 | 90 -- 102 | 179 -- 191 | -11.6062 | 3.99713 | 4.92433 |
| CD630_16550 | CD630_s0010 | 127 -- 147 | 16 -- 35 | -11.4563 | 5.61064 | 4.92433 |
| CD630_14620 | CD630_s0010 | 17 -- 42 | 2 -- 27 | -11.4313 | 4.43651 | 4.92433 |
| CD630_22991 | CD630_s0010 | 189 -- 197 | 156 -- 164 | -11.3862 | 6.03728 | 4.92433 |
| CD630_03220 | CD630_s0010 | 107 -- 128 | 60 -- 81 | -11.3428 | 3.36746 | 4.92433 |
| CD630_13370 | CD630_s0010 | 157 -- 177 | 2 -- 22 | -11.2865 | 4.33145 | 4.92433 |
| CD630_30310 | CD630_s0010 | 176 -- 198 | 66 -- 87 | -11.1958 | 4.56059 | 4.92433 |
| CD630_36180 | CD630_s0010 | 176 -- 199 | 167 -- 190 | -11.0303 | 4.83428 | 4.92433 |
| CD630_03400 | CD630_s0010 | 111 -- 131 | 64 -- 86 | -10.9936 | 4.45994 | 4.92433 |
| CD630_06060 | CD630_s0010 | 66 -- 95 | 160 -- 189 | -10.9521 | 4.94050 | 4.92433 |
| CD630_16530 | CD630_s0010 | 133 -- 164 | 73 -- 106 | -10.8073 | 4.37658 | 4.92433 |
| CD630_24580 | CD630_s0010 | 144 -- 159 | 62 -- 77 | -10.6195 | 3.88519 | 4.92433 |
| CD630_08370 | CD630_s0010 | 82 -- 95 | 183 -- 196 | -10.4983 | 3.65639 | 4.92433 |
| opuCA | CD630_s0010 | 84 -- 100 | 178 -- 193 | -10.3908 | 4.07035 | 4.92433 |
| CD630_31490 | CD630_s0010 | 30 -- 48 | 156 -- 174 | -10.3611 | 4.08593 | 4.92433 |
| folC | CD630_s0010 | 112 -- 140 | 63 -- 90 | -10.2668 | 3.85634 | 4.92433 |
| gcp | CD630_s0010 | 24 -- 56 | 54 -- 81 | -10.1962 | 4.92798 | 4.92433 |
| CD630_13530 | CD630_s0010 | 7 -- 31 | 2 -- 26 | -10.1205 | 4.09832 | 4.92433 |
| spaK | CD630_s0010 | 167 -- 181 | 153 -- 167 | -10.1132 | 3.04530 | 4.92433 |
| CD630_06110 | CD630_s0010 | 177 -- 192 | 158 -- 173 | -10.0865 | 5.61380 | 4.92433 |
| CD630_07870 | CD630_s0010 | 184 -- 200 | 6 -- 20 | -10.0259 | 3.22829 | 4.92433 |
| cobQ | CD630_s0010 | 87 -- 100 | 180 -- 193 | -9.98587 | 3.13524 | 4.92433 |
| CD630_20260 | CD630_s0010 | 133 -- 155 | 2 -- 25 | -9.93092 | 3.84162 | 4.92433 |
| CD630_32720 | CD630_s0010 | 122 -- 157 | 94 -- 130 | -9.91184 | 4.74276 | 4.92433 |
| flhB | CD630_s0010 | 188 -- 199 | 20 -- 33 | -9.78949 | 3.55524 | 4.92433 |
| CD630_21620 | CD630_s0010 | 178 -- 189 | 21 -- 32 | -9.76431 | 3.02649 | 4.92433 |
| CD630_27240 | CD630_s0010 | 109 -- 154 | 36 -- 79 | -9.7585 | 3.52462 | 4.92433 |
| CD630_14182 | CD630_s0010 | 189 -- 197 | 156 -- 164 | -9.71847 | 4.82360 | 4.92433 |
| CD630_08140 | CD630_s0010 | 115 -- 145 | 141 -- 165 | -9.70097 | 3.30159 | 4.92433 |
| CD630_25200 | CD630_s0010 | 145 -- 162 | 7 -- 24 | -9.68777 | 3.65158 | 4.92433 |
| kdpD | CD630_s0010 | 181 -- 200 | 18 -- 36 | -9.67078 | 4.27140 | 4.92433 |
| fbp | CD630_s0010 | 92 -- 115 | 166 -- 188 | -9.66228 | 3.80071 | 4.92433 |
| CD630_20541 | CD630_s0010 | 39 -- 52 | 74 -- 88 | -9.60767 | 5.74527 | 4.92433 |
| CD630_21090 | CD630_s0010 | 167 -- 199 | 2 -- 34 | -9.59021 | 4.15803 | 4.92433 |
| CD630_27570 | CD630_s0010 | 1 -- 18 | 1 -- 27 | -9.5894 | 3.46216 | 4.92433 |
| CD630_21490 | CD630_s0010 | 132 -- 200 | 70 -- 113 | -9.57459 | 3.37462 | 4.92433 |
| CD630_36690 | CD630_s0010 | 15 -- 32 | 2 -- 23 | -9.53806 | 3.53761 | 4.92433 |
| CD630_13960 | CD630_s0010 | 48 -- 64 | 16 -- 36 | -9.52588 | 3.30375 | 4.92433 |
| rbsC | CD630_s0010 | 161 -- 177 | 63 -- 79 | -9.51167 | 3.71650 | 4.92433 |
| CD630_21490 | CD630_s0050 | 189 -- 195 | 47 -- 53 | -12.2923 | 3.37462 | 5.03432 |
| CD630_14130 | CD630_s0050 | 143 -- 183 | 12 -- 52 | -11.32 | 4.57970 | 5.03432 |
| CD630_09840 | CD630_s0050 | 162 -- 186 | 36 -- 61 | -11.3105 | 4.50009 | 5.03432 |
| iorA | CD630_s0050 | 55 -- 62 | 47 -- 54 | -11.2574 | 3.83173 | 5.03432 |
| rbsA | CD630_s0050 | 1 -- 21 | 2 -- 19 | -10.7837 | 3.71650 | 5.03432 |
| CD630_28550 | CD630_s0050 | 192 -- 200 | 41 -- 49 | -10.3267 | 3.51196 | 5.03432 |
| CD630_08140 | CD630_s0050 | 3 -- 9 | 12 -- 18 | -10.0307 | 3.30159 | 5.03432 |
| CD630_36040 | CD630_s0050 | 120 -- 126 | 47 -- 53 | -9.83197 | 3.52009 | 5.03432 |
| CD630_30110 | CD630_s0050 | 170 -- 176 | 47 -- 53 | -9.78205 | 3.31041 | 5.03432 |
| CD630_30420 | CD630_s0190 | 120 -- 174 | 50 -- 107 | -18.1492 | 5.27539 | 4.98480 |
| ppnK | CD630_s0190 | 127 -- 192 | 49 -- 122 | -16.9034 | 4.36817 | 4.98480 |
| CD630_06150 | CD630_s0190 | 83 -- 166 | 65 -- 136 | -16.4343 | 5.51744 | 4.98480 |
| CD630_08090 | CD630_s0190 | 128 -- 201 | 63 -- 122 | -15.9049 | 3.57167 | 4.98480 |
| CD630_27280 | CD630_s0190 | 121 -- 196 | 4 -- 74 | -15.794 | 3.24738 | 4.98480 |
| CD630_13020 | CD630_s0190 | 118 -- 145 | 64 -- 92 | -15.4582 | 3.52267 | 4.98480 |
| CD630_06100 | CD630_s0190 | 102 -- 192 | 61 -- 156 | -15.2501 | 4.43918 | 4.98480 |
| CD630_09100 | CD630_s0190 | 8 -- 51 | 1 -- 49 | -15.0277 | 5.92260 | 4.98480 |
| CD630_19440 | CD630_s0190 | 117 -- 185 | 54 -- 134 | -14.9814 | 4.53937 | 4.98480 |
| cwp26 | CD630_s0190 | 126 -- 156 | 53 -- 84 | -14.2629 | 4.44377 | 4.98480 |
| pgm | CD630_s0190 | 87 -- 171 | 50 -- 131 | -14.0989 | 3.45439 | 4.98480 |
| CD630_28870 | CD630_s0190 | 75 -- 101 | 54 -- 81 | -14.0167 | 3.41991 | 4.98480 |
| cobC | CD630_s0190 | 25 -- 54 | 67 -- 88 | -13.6447 | 3.15854 | 4.98480 |
| CD630_01590 | CD630_s0190 | 13 -- 35 | 57 -- 78 | -13.5239 | 3.45609 | 4.98480 |
| CD630_13580 | CD630_s0190 | 112 -- 146 | 70 -- 105 | -13.3161 | 3.80715 | 4.98480 |
| CD630_33682 | CD630_s0190 | 22 -- 45 | 171 -- 192 | -12.9818 | 5.77720 | 4.98480 |
| CD630_23280 | CD630_s0190 | 164 -- 185 | 28 -- 49 | -12.703 | 3.50093 | 4.98480 |
| CD630_24580 | CD630_s0190 | 126 -- 178 | 134 -- 172 | -12.6509 | 3.88519 | 4.98480 |
| CD630_08080 | CD630_s0190 | 2 -- 35 | 37 -- 76 | -12.6245 | 4.87819 | 4.98480 |
| CD630_14240 | CD630_s0190 | 22 -- 45 | 171 -- 192 | -12.554 | 8.60766 | 4.98480 |
| CD630_03170 | CD630_s0190 | 109 -- 178 | 75 -- 151 | -12.2737 | 3.66946 | 4.98480 |
| CD630_08140 | CD630_s0190 | 5 -- 55 | 33 -- 88 | -12.1041 | 3.30159 | 4.98480 |
| CD630_05870 | CD630_s0190 | 120 -- 142 | 75 -- 96 | -11.8873 | 4.28339 | 4.98480 |
| fbp | CD630_s0190 | 92 -- 115 | 176 -- 194 | -11.4958 | 3.80071 | 4.98480 |
| CD630_28560 | CD630_s0190 | 82 -- 93 | 192 -- 203 | -11.4468 | 3.96803 | 4.98480 |
| CD630_21090 | CD630_s0190 | 156 -- 179 | 69 -- 89 | -11.4039 | 4.15803 | 4.98480 |
| CD630_23820 | CD630_s0190 | 167 -- 198 | 61 -- 89 | -11.3651 | 3.18419 | 4.98480 |
| CD630_17630 | CD630_s0190 | 139 -- 169 | 64 -- 87 | -11.2387 | 4.52408 | 4.98480 |
| CD630_18311 | CD630_s0190 | 134 -- 151 | 70 -- 87 | -11.2265 | 6.86047 | 4.98480 |
| CD630_19903 | CD630_s0190 | 21 -- 44 | 171 -- 192 | -11.1952 | 5.79994 | 4.98480 |
| CD630_28290 | CD630_s0190 | 186 -- 196 | 131 -- 141 | -11.0908 | 5.54192 | 4.98480 |
| CD630_05822 | CD630_s0190 | 129 -- 152 | 56 -- 85 | -11.0773 | 4.28113 | 4.98480 |
| CD630_23090 | CD630_s0190 | 25 -- 45 | 171 -- 189 | -11.0634 | 4.25622 | 4.98480 |
| cdtR | CD630_s0190 | 72 -- 124 | 91 -- 145 | -11.0085 | 5.84368 | 4.98480 |
| CD630_22991 | CD630_s0190 | 189 -- 196 | 167 -- 174 | -11.0075 | 6.03728 | 4.98480 |
| CD630_04880 | CD630_s0190 | 147 -- 179 | 61 -- 82 | -10.9671 | 4.62291 | 4.98480 |
| CD630_25300 | CD630_s0190 | 133 -- 156 | 120 -- 148 | -10.893 | 4.49333 | 4.98480 |
| CD630_25620 | CD630_s0190 | 85 -- 101 | 117 -- 134 | -10.8573 | 3.02685 | 4.98480 |
| spaK | CD630_s0190 | 167 -- 200 | 133 -- 177 | -10.807 | 3.04530 | 4.98480 |
| eutP | CD630_s0190 | 162 -- 182 | 101 -- 123 | -10.7921 | 3.82847 | 4.98480 |
| rolA | CD630_s0190 | 1 -- 36 | 13 -- 47 | -10.7791 | 4.56362 | 4.98480 |
| panG | CD630_s0190 | 65 -- 77 | 57 -- 69 | -10.737 | 5.54047 | 4.98480 |
| CD630_16330 | CD630_s0190 | 165 -- 182 | 167 -- 187 | -10.6727 | 4.11180 | 4.98480 |
| CD630_21140 | CD630_s0190 | 14 -- 37 | 60 -- 81 | -10.6719 | 3.33304 | 4.98480 |
| CD630_01690 | CD630_s0190 | 165 -- 182 | 107 -- 124 | -10.6105 | 4.97234 | 4.98480 |
| CD630_18010 | CD630_s0190 | 183 -- 198 | 135 -- 151 | -10.5014 | 4.21967 | 4.98480 |
| rbsA | CD630_s0190 | 129 -- 159 | 79 -- 113 | -10.4784 | 3.71650 | 4.98480 |
| CD630_09970 | CD630_s0190 | 170 -- 185 | 71 -- 86 | -10.4701 | 3.25917 | 4.98480 |
| CD630_26050 | CD630_s0190 | 113 -- 120 | 41 -- 48 | -10.4499 | 6.99851 | 4.98480 |
| CD630_19850 | CD630_s0190 | 110 -- 145 | 34 -- 69 | -10.419 | 6.88486 | 4.98480 |
| CD630_04320 | CD630_s0190 | 169 -- 190 | 165 -- 187 | -10.4123 | 3.68342 | 4.98480 |
| CD630_16550 | CD630_s0190 | 183 -- 198 | 56 -- 71 | -10.3794 | 5.61064 | 4.98480 |
| CD630_05950 | CD630_s0190 | 178 -- 189 | 38 -- 49 | -10.3199 | 3.31669 | 4.98480 |
| CD630_08370 | CD630_s0190 | 188 -- 198 | 164 -- 174 | -10.2981 | 3.65639 | 4.98480 |
| CD630_31500 | CD630_s0190 | 144 -- 191 | 57 -- 113 | -10.2244 | 4.90207 | 4.98480 |
| CD630_13350 | CD630_s0190 | 192 -- 200 | 42 -- 50 | -10.2078 | 4.15139 | 4.98480 |
| CD630_04620 | CD630_s0190 | 5 -- 30 | 67 -- 97 | -10.107 | 3.45413 | 4.98480 |
| CD630_22550 | CD630_s0190 | 124 -- 165 | 29 -- 71 | -10.1062 | 3.62379 | 4.98480 |
| CD630_36170 | CD630_s0190 | 160 -- 168 | 41 -- 49 | -10.0784 | 3.72380 | 4.98480 |
| CD630_14182 | CD630_s0190 | 189 -- 196 | 167 -- 174 | -10.0632 | 4.82360 | 4.98480 |
| CD630_04850 | CD630_s0190 | 153 -- 184 | 60 -- 87 | -10.0168 | 4.40788 | 4.98480 |
| spoVFB | CD630_s0190 | 1 -- 23 | 56 -- 77 | -10.0145 | 4.62758 | 4.98480 |
| CD630_06060 | CD630_s0190 | 76 -- 95 | 170 -- 189 | -9.98804 | 4.94050 | 4.98480 |
| CD630_21730 | CD630_s0190 | 181 -- 192 | 75 -- 86 | -9.98625 | 3.45076 | 4.98480 |
| CD630_26920 | CD630_s0190 | 140 -- 158 | 138 -- 158 | -9.90563 | 4.58060 | 4.98480 |
| CD630_03650 | CD630_s0190 | 69 -- 90 | 52 -- 73 | -9.90074 | 3.93697 | 4.98480 |
| CD630_20650 | CD630_s0190 | 120 -- 126 | 41 -- 47 | -9.89423 | 4.00083 | 4.98480 |
| CD630_10950 | CD630_s0190 | 1 -- 8 | 40 -- 47 | -9.8411 | 3.67675 | 4.98480 |
| CD630_05940 | CD630_s0190 | 56 -- 83 | 33 -- 61 | -9.82824 | 4.18595 | 4.98480 |
| CD630_29840 | CD630_s0190 | 192 -- 201 | 6 -- 15 | -9.81805 | 4.48825 | 4.98480 |
| bltD | CD630_s0190 | 181 -- 198 | 56 -- 73 | -9.74048 | 4.49926 | 4.98480 |
| CD630_28550 | CD630_s0190 | 170 -- 200 | 54 -- 86 | -9.7247 | 3.51196 | 4.98480 |
| CD630_35840 | CD630_s0190 | 83 -- 135 | 63 -- 131 | -9.70456 | 3.14170 | 4.98480 |
| CD630_22030 | CD630_s0190 | 132 -- 200 | 51 -- 122 | -9.68079 | 3.29934 | 4.98480 |
| CD630_36781 | CD630_s0190 | 127 -- 186 | 51 -- 137 | -9.56151 | 3.75946 | 4.98480 |
| CD630_13920 | CD630_s0190 | 174 -- 194 | 102 -- 124 | -9.54409 | 3.25236 | 4.98480 |
| CD630_14150 | CD630_s0190 | 37 -- 50 | 58 -- 72 | -9.53004 | 3.28251 | 4.98480 |
| CD630_27520 | CD630_s0190 | 128 -- 158 | 178 -- 206 | -9.52644 | 4.27693 | 4.98480 |

| | | | | | | |
|---|---|---|---|---|---|---|
| srlEb | CD630_s0190 | 68 -- 173 | 42 -- 140 | -9.50107 | 3.50140 | 4.98480 |
| feoB | CD630_s0210 | 24 -- 40 | 74 -- 90 | -15.4066 | 4.05364 | 4.02122 |
| CD630_19440 | CD630_s0210 | 115 -- 184 | 15 -- 87 | -15.3974 | 4.53937 | 4.02122 |
| CD630_09100 | CD630_s0210 | 1 -- 26 | 34 -- 56 | -14.9944 | 5.92260 | 4.02122 |
| CD630_17360 | CD630_s0210 | 177 -- 198 | 42 -- 61 | -14.7996 | 5.66070 | 4.02122 |
| CD630_03920 | CD630_s0210 | 1 -- 19 | 10 -- 28 | -14.5744 | 4.15934 | 4.02122 |
| CD630_27280 | CD630_s0210 | 147 -- 195 | 3 -- 62 | -14.3493 | 3.24738 | 4.02122 |
| CD630_32040 | CD630_s0210 | 129 -- 192 | 4 -- 61 | -14.1398 | 4.93516 | 4.02122 |
| CD630_32070 | CD630_s0210 | 57 -- 73 | 222 -- 237 | -13.6102 | 3.51964 | 4.02122 |
| CD630_06100 | CD630_s0210 | 95 -- 124 | 30 -- 58 | -13.5068 | 4.43918 | 4.02122 |
| CD630_15190 | CD630_s0210 | 164 -- 185 | 211 -- 231 | -13.1296 | 4.43255 | 4.02122 |
| cspD | CD630_s0210 | 161 -- 185 | 214 -- 237 | -12.8497 | 3.23587 | 4.02122 |
| CD630_08100 | CD630_s0210 | 80 -- 123 | 44 -- 82 | -12.6033 | 4.39181 | 4.02122 |
| CD630_32990 | CD630_s0210 | 171 -- 190 | 40 -- 58 | -12.5338 | 5.24925 | 4.02122 |
| CD630_14260 | CD630_s0210 | 15 -- 45 | 26 -- 57 | -12.4213 | 3.22214 | 4.02122 |
| xdhA | CD630_s0210 | 56 -- 75 | 60 -- 80 | -12.408 | 4.45147 | 4.02122 |
| CD630_05830 | CD630_s0210 | 90 -- 160 | 1 -- 62 | -12.3557 | 3.35360 | 4.02122 |
| spoVFB | CD630_s0210 | 170 -- 190 | 220 -- 239 | -12.3286 | 4.62758 | 4.02122 |
| panB | CD630_s0210 | 97 -- 170 | 5 -- 61 | -12.3128 | 3.40248 | 4.02122 |
| CD630_05211 | CD630_s0210 | 20 -- 28 | 42 -- 50 | -12.31 | 3.97048 | 4.02122 |
| CD630_15900 | CD630_s0210 | 139 -- 189 | 5 -- 60 | -12.299 | 3.17664 | 4.02122 |
| CD630_04850 | CD630_s0210 | 113 -- 133 | 9 -- 28 | -12.0994 | 4.40788 | 4.02122 |
| dltB | CD630_s0210 | 145 -- 180 | 32 -- 60 | -11.9248 | 3.81485 | 4.02122 |
| ppnK | CD630_s0210 | 141 -- 198 | 24 -- 87 | -11.8533 | 4.36817 | 4.02122 |
| CD630_17970 | CD630_s0210 | 91 -- 135 | 116 -- 159 | -11.8378 | 3.68802 | 4.02122 |
| CD630_05820 | CD630_s0210 | 1 -- 22 | 42 -- 63 | -11.7687 | 3.75661 | 4.02122 |
| CD630_21090 | CD630_s0210 | 117 -- 185 | 20 -- 87 | -11.7131 | 4.15803 | 4.02122 |
| CD630_17151 | CD630_s0210 | 130 -- 177 | 19 -- 64 | -11.5785 | 6.43124 | 4.02122 |
| CD630_34520 | CD630_s0210 | 137 -- 201 | 31 -- 88 | -11.3428 | 3.83016 | 4.02122 |
| CD630_06170 | CD630_s0210 | 104 -- 125 | 37 -- 62 | -11.2894 | 4.95366 | 4.02122 |
| uxaA%27 | CD630_s0210 | 137 -- 185 | 17 -- 56 | -11.2203 | 4.26165 | 4.02122 |
| appC | CD630_s0210 | 150 -- 198 | 176 -- 215 | -11.2034 | 4.50438 | 4.02122 |
| CD630_04610 | CD630_s0210 | 149 -- 195 | 131 -- 163 | -11.1995 | 3.83025 | 4.02122 |
| cspC | CD630_s0210 | 157 -- 185 | 214 -- 240 | -11.1846 | 5.48848 | 4.02122 |
| CD630_27210 | CD630_s0210 | 108 -- 149 | 1 -- 57 | -11.146 | 4.18786 | 4.02122 |
| CD630_13880 | CD630_s0210 | 146 -- 162 | 39 -- 60 | -11.0367 | 3.52735 | 4.02122 |
| CD630_14050 | CD630_s0210 | 150 -- 186 | 25 -- 61 | -11.0204 | 3.30894 | 4.02122 |
| CD630_10880 | CD630_s0210 | 144 -- 165 | 220 -- 242 | -11.0201 | 5.05048 | 4.02122 |
| CD630_03160 | CD630_s0210 | 98 -- 118 | 116 -- 135 | -11.0013 | 3.66946 | 4.02122 |
| CD630_19700 | CD630_s0210 | 152 -- 191 | 54 -- 89 | -10.9306 | 3.85861 | 4.02122 |
| flhB | CD630_s0210 | 6 -- 48 | 2 -- 47 | -10.8663 | 3.55524 | 4.02122 |
| CD630_05490 | CD630_s0210 | 55 -- 97 | 17 -- 56 | -10.8525 | 3.89852 | 4.02122 |
| CD630_20650 | CD630_s0210 | 107 -- 123 | 41 -- 60 | -10.7865 | 4.00083 | 4.02122 |
| pilT | CD630_s0210 | 136 -- 151 | 41 -- 57 | -10.7416 | 3.24639 | 4.02122 |
| CD630_31210 | CD630_s0210 | 18 -- 33 | 46 -- 61 | -10.7277 | 4.17693 | 4.02122 |
| CD630_06380 | CD630_s0210 | 119 -- 189 | 7 -- 60 | -10.6732 | 4.32211 | 4.02122 |
| CD630_22660 | CD630_s0210 | 2 -- 22 | 180 -- 200 | -10.6713 | 3.60496 | 4.02122 |
| CD630_13790 | CD630_s0210 | 190 -- 200 | 73 -- 83 | -10.671 | 5.17374 | 4.02122 |
| cprK | CD630_s0210 | 82 -- 122 | 200 -- 234 | -10.664 | 3.96216 | 4.02122 |
| CD630_29960 | CD630_s0210 | 60 -- 116 | 2 -- 60 | -10.647 | 4.51904 | 4.02122 |
| CD630_02420 | CD630_s0210 | 45 -- 56 | 78 -- 89 | -10.6265 | 3.71048 | 4.02122 |
| CD630_05940 | CD630_s0210 | 30 -- 80 | 2 -- 60 | -10.5964 | 4.18595 | 4.02122 |
| CD630_28181 | CD630_s0210 | 145 -- 162 | 222 -- 240 | -10.5436 | 3.91299 | 4.02122 |
| CD630_21260 | CD630_s0210 | 118 -- 150 | 4 -- 49 | -10.5427 | 4.22255 | 4.02122 |
| cobS | CD630_s0210 | 124 -- 138 | 74 -- 89 | -10.5169 | 3.67481 | 4.02122 |
| CD630_05300 | CD630_s0210 | 176 -- 199 | 52 -- 80 | -10.5097 | 4.56885 | 4.02122 |
| adhE | CD630_s0210 | 49 -- 69 | 73 -- 90 | -10.4858 | 3.06485 | 4.02122 |
| CD630_09850 | CD630_s0210 | 168 -- 182 | 46 -- 59 | -10.3183 | 4.18679 | 4.02122 |
| spaK | CD630_s0210 | 153 -- 182 | 144 -- 173 | -10.3124 | 3.04530 | 4.02122 |
| CD630_35780 | CD630_s0210 | 53 -- 66 | 78 -- 90 | -10.3123 | 3.50107 | 4.02122 |
| eutP | CD630_s0210 | 161 -- 178 | 64 -- 82 | -10.2831 | 3.82847 | 4.02122 |
| CD630_34530 | CD630_s0210 | 154 -- 170 | 172 -- 188 | -10.2715 | 3.83016 | 4.02122 |
| CD630_10950 | CD630_s0210 | 85 -- 93 | 221 -- 229 | -10.2689 | 3.67675 | 4.02122 |
| CD630_11220 | CD630_s0210 | 38 -- 98 | 20 -- 80 | -10.2687 | 3.52327 | 4.02122 |
| CD630_28290 | CD630_s0210 | 126 -- 157 | 26 -- 58 | -10.2629 | 5.54192 | 4.02122 |
| CD630_20460 | CD630_s0210 | 139 -- 153 | 137 -- 152 | -10.2563 | 7.94034 | 4.02122 |
| CD630_22991 | CD630_s0210 | 189 -- 197 | 200 -- 208 | -10.2297 | 6.03728 | 4.02122 |
| CD630_29250 | CD630_s0210 | 52 -- 102 | 10 -- 54 | -10.2132 | 4.61901 | 4.02122 |
| CD630_09410 | CD630_s0210 | 52 -- 102 | 10 -- 54 | -10.2132 | 4.75651 | 4.02122 |
| CD630_26020 | CD630_s0210 | 8 -- 32 | 4 -- 29 | -10.1909 | 4.03155 | 4.02122 |
| CD630_30420 | CD630_s0210 | 109 -- 177 | 6 -- 58 | -10.1408 | 5.27539 | 4.02122 |
| CD630_18311 | CD630_s0210 | 121 -- 172 | 1 -- 57 | -10.0956 | 6.86047 | 4.02122 |
| CD630_23080 | CD630_s0210 | 30 -- 53 | 48 -- 67 | -10.0908 | 5.23027 | 4.02122 |
| CD630_20600 | CD630_s0210 | 156 -- 186 | 44 -- 77 | -10.0518 | 4.20093 | 4.02122 |
| CD630_11240 | CD630_s0210 | 7 -- 39 | 122 -- 154 | -10.0177 | 3.20274 | 4.02122 |
| CD630_21240 | CD630_s0210 | 156 -- 197 | 45 -- 82 | -9.96533 | 5.88879 | 4.02122 |
| CD630_31490 | CD630_s0210 | 116 -- 172 | 67 -- 110 | -9.92976 | 4.08593 | 4.02122 |
| CD630_35840 | CD630_s0210 | 182 -- 201 | 48 -- 66 | -9.90677 | 3.14170 | 4.02122 |
| CD630_35430 | CD630_s0210 | 120 -- 164 | 26 -- 59 | -9.90452 | 4.28798 | 4.02122 |
| CD630_27200 | CD630_s0210 | 30 -- 40 | 73 -- 83 | -9.8533 | 3.42516 | 4.02122 |
| CD630_23450 | CD630_s0210 | 184 -- 193 | 41 -- 51 | -9.84509 | 3.27218 | 4.02122 |
| CD630_16530 | CD630_s0210 | 128 -- 169 | 13 -- 59 | -9.84233 | 4.37658 | 4.02122 |
| virS | CD630_s0210 | 177 -- 189 | 46 -- 58 | -9.76883 | 4.74085 | 4.02122 |
| CD630_32890 | CD630_s0210 | 110 -- 136 | 49 -- 75 | -9.75864 | 3.41276 | 4.02122 |
| vncR | CD630_s0210 | 22 -- 44 | 31 -- 49 | -9.71038 | 3.93642 | 4.02122 |
| CD630_25610 | CD630_s0210 | 174 -- 192 | 72 -- 90 | -9.6729 | 3.02685 | 4.02122 |
| CD630_04770 | CD630_s0210 | 26 -- 63 | 26 -- 59 | -9.66958 | 3.58478 | 4.02122 |
| glnA | CD630_s0210 | 171 -- 197 | 136 -- 162 | -9.65703 | 3.87629 | 4.02122 |
| CD630_22880 | CD630_s0210 | 19 -- 31 | 22 -- 34 | -9.6367 | 3.62455 | 4.02122 |
| CD630_32880 | CD630_s0210 | 7 -- 17 | 75 -- 85 | -9.61252 | 3.41276 | 4.02122 |
| CD630_09770 | CD630_s0210 | 145 -- 199 | 11 -- 49 | -9.57327 | 3.22106 | 4.02122 |
| CD630_04620 | CD630_s0210 | 172 -- 183 | 43 -- 54 | -9.5691 | 3.45413 | 4.02122 |
| CD630_08950 | CD630_s0210 | 92 -- 105 | 74 -- 87 | -9.56865 | 3.71905 | 4.02122 |
| CD630_24770 | CD630_s0210 | 173 -- 185 | 206 -- 217 | -9.524 | 3.20798 | 4.02122 |
| CD630_14240 | CD630_s0210 | 40 -- 80 | 21 -- 59 | -9.50444 | 8.60766 | 4.02122 |
| CD630_31220 | CD630_s0220 | 157 -- 188 | 2 -- 36 | -14.0243 | 3.73779 | 3.72587 |
| CD630_19850 | CD630_s0220 | 106 -- 118 | 75 -- 87 | -12.4701 | 6.88486 | 3.72587 |
| cwp26 | CD630_s0220 | 146 -- 186 | 4 -- 37 | -9.92186 | 4.44377 | 3.72587 |
| kdgT1 | CD630_s0250 | 180 -- 201 | 7 -- 27 | -13.0068 | 4.49207 | 3.74616 |
| thiH | CD630_s0250 | 179 -- 189 | 1 -- 11 | -12.4476 | 3.58230 | 3.74616 |
| CD630_31220 | CD630_s0250 | 156 -- 189 | 1 -- 39 | -12.4366 | 3.73779 | 3.74616 |
| CD630_20290 | CD630_s0250 | 63 -- 95 | 74 -- 106 | -11.7791 | 4.38183 | 3.74616 |
| CD630_03220 | CD630_s0250 | 99 -- 111 | 28 -- 41 | -10.8174 | 3.36746 | 3.74616 |
| garK | CD630_s0250 | 138 -- 160 | 77 -- 106 | -10.6522 | 3.72836 | 3.74616 |
| CD630_06642 | CD630_s0250 | 155 -- 173 | 42 -- 60 | -10.1317 | 3.32967 | 3.74616 |
| kdpE | CD630_s0250 | 133 -- 141 | 1 -- 9 | -9.8569 | 4.27140 | 3.74616 |
| CD630_28550 | CD630_s0250 | 190 -- 200 | 41 -- 51 | -9.73947 | 3.51196 | 3.74616 |
| CD630_08100 | CD630_s0250 | 22 -- 35 | 1 -- 14 | -9.53907 | 4.39181 | 3.74616 |
| etfA | CD630_s0250 | 170 -- 181 | 24 -- 35 | -9.51641 | 3.41204 | 3.74616 |
| CD630_33420 | CD630_s0270 | 16 -- 153 | 34 -- 149 | -14.9003 | 3.22095 | 3.63582 |
| CD630_09100 | CD630_s0270 | 3 -- 22 | 4 -- 23 | -14.8163 | 5.92260 | 3.63582 |
| xylR | CD630_s0270 | 140 -- 181 | 6 -- 46 | -14.485 | 3.95671 | 3.63582 |
| CD630_26150 | CD630_s0270 | 133 -- 187 | 9 -- 67 | -14.2402 | 4.06302 | 3.63582 |
| CD630_14340 | CD630_s0270 | 33 -- 79 | 89 -- 147 | -13.8219 | 5.10868 | 3.63582 |
| CD630_27360 | CD630_s0270 | 84 -- 134 | 41 -- 85 | -13.6185 | 3.79570 | 3.63582 |
| CD630_27270 | CD630_s0270 | 90 -- 157 | 8 -- 85 | -13.2714 | 3.50892 | 3.63582 |
| CD630_21730 | CD630_s0270 | 39 -- 135 | 44 -- 150 | -13.2251 | 3.45076 | 3.63582 |
| CD630_22030 | CD630_s0270 | 138 -- 182 | 104 -- 148 | -13.0206 | 3.29934 | 3.63582 |
| CD630_04320 | CD630_s0270 | 160 -- 185 | 207 -- 229 | -12.9978 | 3.68342 | 3.63582 |
| CD630_03170 | CD630_s0270 | 87 -- 107 | 218 -- 239 | -12.7873 | 3.66946 | 3.63582 |
| CD630_21090 | CD630_s0270 | 108 -- 171 | 1 -- 61 | -12.7311 | 4.15803 | 3.63582 |
| pilC | CD630_s0270 | 129 -- 147 | 225 -- 239 | -12.6049 | 3.54230 | 3.63582 |
| CD630_25020 | CD630_s0270 | 10 -- 62 | 6 -- 60 | -12.5785 | 4.17721 | 3.63582 |
| bglA | CD630_s0270 | 178 -- 201 | 215 -- 239 | -12.1615 | 4.86402 | 3.63582 |
| CD630_06380 | CD630_s0270 | 118 -- 165 | 103 -- 147 | -11.9575 | 4.32211 | 3.63582 |
| CD630_34080 | CD630_s0270 | 45 -- 113 | 20 -- 72 | -11.6511 | 3.20224 | 3.63582 |
| CD630_25200 | CD630_s0270 | 179 -- 200 | 112 -- 134 | -11.4811 | 3.65158 | 3.63582 |
| CD630_16950 | CD630_s0270 | 177 -- 199 | 111 -- 131 | -11.177 | 4.42569 | 3.63582 |
| CD630_05030 | CD630_s0270 | 121 -- 153 | 20 -- 49 | -11.1223 | 4.03823 | 3.63582 |
| CD630_16150 | CD630_s0270 | 60 -- 71 | 74 -- 85 | -11.0974 | 4.20614 | 3.63582 |
| CD630_14810 | CD630_s0270 | 127 -- 136 | 213 -- 222 | -11.0648 | 3.35323 | 3.63582 |
| blaI | CD630_s0270 | 40 -- 53 | 62 -- 75 | -11.0548 | 3.73674 | 3.63582 |
| CD630_27280 | CD630_s0270 | 152 -- 197 | 4 -- 44 | -11.0021 | 3.24738 | 3.63582 |
| eutW | CD630_s0270 | 10 -- 22 | 224 -- 237 | -10.9906 | 4.55715 | 3.63582 |
| CD630_10080 | CD630_s0270 | 146 -- 171 | 112 -- 134 | -10.9769 | 3.75625 | 3.63582 |
| CD630_06060 | CD630_s0270 | 77 -- 184 | 127 -- 225 | -10.9693 | 4.94050 | 3.63582 |
| CD630_32060 | CD630_s0270 | 170 -- 189 | 35 -- 50 | -10.9433 | 5.73797 | 3.63582 |
| CD630_11730 | CD630_s0270 | 144 -- 175 | 125 -- 156 | -10.8885 | 4.80521 | 3.63582 |
| hfq | CD630_s0270 | 148 -- 184 | 200 -- 237 | -10.7959 | 5.05609 | 3.63582 |
| chrA | CD630_s0270 | 39 -- 60 | 68 -- 88 | -10.6814 | 5.27255 | 3.63582 |
| CD630_34520 | CD630_s0270 | 139 -- 201 | 33 -- 90 | -10.6768 | 3.83016 | 3.63582 |
| CD630_20980 | CD630_s0270 | 177 -- 201 | 64 -- 90 | -10.66 | 8.12847 | 3.63582 |
| CD630_23820 | CD630_s0270 | 3 -- 43 | 31 -- 78 | -10.6481 | 3.18419 | 3.63582 |
| CD630_08660 | CD630_s0270 | 145 -- 191 | 119 -- 167 | -10.6203 | 3.86467 | 3.63582 |
| CD630_06060 | CD630_s0270 | 106 -- 145 | 58 -- 99 | -10.6135 | 4.94050 | 3.63582 |
| CD630_15890 | CD630_s0270 | 147 -- 170 | 107 -- 127 | -10.5964 | 3.39962 | 3.63582 |
| CD630_19903 | CD630_s0270 | 46 -- 59 | 86 -- 99 | -10.5915 | 5.79994 | 3.63582 |
| CD630_06540 | CD630_s0270 | 82 -- 118 | 200 -- 241 | -10.5212 | 3.54673 | 3.63582 |
| CD630_30420 | CD630_s0270 | 136 -- 200 | 109 -- 178 | -10.4998 | 5.27539 | 3.63582 |
| aroD | CD630_s0270 | 55 -- 75 | 10 -- 28 | -10.4681 | 3.71929 | 3.63582 |
| CD630_05950 | CD630_s0270 | 9 -- 25 | 10 -- 26 | -10.4579 | 3.31669 | 3.63582 |
| CD630_21610 | CD630_s0270 | 139 -- 166 | 50 -- 77 | -10.4182 | 3.16662 | 3.63582 |
| CD630_04540 | CD630_s0270 | 82 -- 109 | 217 -- 240 | -10.4029 | 3.33556 | 3.63582 |
| CD630_05450 | CD630_s0270 | 158 -- 177 | 127 -- 146 | -10.393 | 5.04761 | 3.63582 |
| CD630_14600 | CD630_s0270 | 138 -- 150 | 81 -- 92 | -10.3594 | 5.90348 | 3.63582 |
| CD630_03650 | CD630_s0270 | 174 -- 197 | 45 -- 65 | -10.3576 | 3.93697 | 3.63582 |
| CD630_16160 | CD630_s0270 | 43 -- 66 | 68 -- 89 | -10.3515 | 6.11801 | 3.63582 |
| CD630_09050 | CD630_s0270 | 181 -- 199 | 112 -- 128 | -10.3337 | 3.96216 | 3.63582 |
| CD630_22740 | CD630_s0270 | 78 -- 95 | 220 -- 236 | -10.2984 | 3.08733 | 3.63582 |
| CD630_31461 | CD630_s0270 | 131 -- 166 | 7 -- 41 | -10.2957 | 4.41476 | 3.63582 |
| CD630_22500 | CD630_s0270 | 130 -- 172 | 104 -- 149 | -10.2638 | 4.03688 | 3.63582 |
| CD630_08360 | CD630_s0270 | 45 -- 68 | 91 -- 116 | -10.2181 | 3.19339 | 3.63582 |
| CD630_10960 | CD630_s0270 | 89 -- 107 | 218 -- 237 | -10.2171 | 3.53850 | 3.63582 |
| CD630_05370 | CD630_s0270 | 126 -- 158 | 78 -- 120 | -10.0927 | 3.26947 | 3.63582 |
| CD630_33580 | CD630_s0270 | 178 -- 194 | 58 -- 75 | -10.0439 | 3.95778 | 3.63582 |
| sigK | CD630_s0270 | 107 -- 152 | 109 -- 162 | -10.0416 | 3.08122 | 3.63582 |
| CD630_27610 | CD630_s0270 | 114 -- 124 | 111 -- 121 | -10.032 | 3.96465 | 3.63582 |
| CD630_18120 | CD630_s0270 | 86 -- 100 | 228 -- 242 | -10.0187 | 3.99713 | 3.63582 |
| CD630_24590 | CD630_s0270 | 83 -- 107 | 76 -- 99 | -9.93882 | 3.70401 | 3.63582 |
| cwp26 | CD630_s0270 | 186 -- 199 | 61 -- 74 | -9.93327 | 4.44377 | 3.63582 |
| CD630_34050 | CD630_s0270 | 139 -- 157 | 57 -- 75 | -9.90571 | 4.03863 | 3.63582 |
| CD630_17710 | CD630_s0270 | 64 -- 120 | 15 -- 64 | -9.88438 | 4.37688 | 3.63582 |
| CD630_26750 | CD630_s0270 | 125 -- 152 | 11 -- 34 | -9.81959 | 3.60610 | 3.63582 |
| dnaC | CD630_s0270 | 47 -- 74 | 63 -- 90 | -9.69508 | 6.30796 | 3.63582 |
| CD630_04610 | CD630_s0270 | 144 -- 201 | 1 -- 60 | -9.69446 | 3.83025 | 3.63582 |
| CD630_15460 | CD630_s0270 | 89 -- 106 | 224 -- 240 | -9.6735 | 6.86296 | 3.63582 |
| CD630_22550 | CD630_s0270 | 159 -- 184 | 204 -- 226 | -9.67072 | 3.62379 | 3.63582 |
| CD630_35260 | CD630_s0270 | 66 -- 95 | 80 -- 101 | -9.64525 | 5.77277 | 3.63582 |
| bltD | CD630_s0270 | 188 -- 198 | 221 -- 231 | -9.62257 | 4.49926 | 3.63582 |
| CD630_19700 | CD630_s0270 | 4 -- 54 | 113 -- 160 | -9.61701 | 3.85861 | 3.63582 |
| CD630_06150 | CD630_s0270 | 83 -- 109 | 225 -- 244 | -9.51271 | 5.51744 | 3.63582 |
| serS1 | CD630_s0270 | 159 -- 170 | 228 -- 239 | -9.50573 | 3.45858 | 3.63582 |
| spaK | CD630_s0280 | 132 -- 188 | 71 -- 129 | -17.1279 | 3.04530 | 5.37459 |
| dltB | CD630_s0280 | 139 -- 167 | 45 -- 74 | -16.7015 | 3.81485 | 5.37459 |
| CD630_01560 | CD630_s0280 | 145 -- 185 | 56 -- 94 | -16.3024 | 4.06352 | 5.37459 |
| ldh | CD630_s0280 | 134 -- 164 | 27 -- 56 | -15.8986 | 4.35818 | 5.37459 |
| CD630_22550 | CD630_s0280 | 166 -- 184 | 211 -- 229 | -15.8631 | 3.62379 | 5.37459 |
| CD630_04770 | CD630_s0280 | 39 -- 61 | 134 -- 154 | -15.6192 | 3.58478 | 5.37459 |
| CD630_22710 | CD630_s0280 | 171 -- 184 | 210 -- 223 | -13.4934 | 4.05623 | 5.37459 |
| CD630_31490 | CD630_s0280 | 32 -- 50 | 208 -- 226 | -13.4439 | 4.08593 | 5.37459 |
| CD630_06080 | CD630_s0280 | 19 -- 97 | 20 -- 100 | -13.3973 | 3.82315 | 5.37459 |
| CD630_31530 | CD630_s0280 | 7 -- 41 | 102 -- 136 | -13.2171 | 3.49100 | 5.37459 |
| thiH | CD630_s0280 | 46 -- 74 | 218 -- 242 | -12.8561 | 3.58230 | 5.37459 |
| CD630_23450 | CD630_s0280 | 41 -- 56 | 60 -- 75 | -12.4701 | 3.27218 | 5.37459 |
| CD630_09990 | CD630_s0280 | 119 -- 139 | 8 -- 28 | -12.4149 | 3.86040 | 5.37459 |
| metQ | CD630_s0280 | 122 -- 135 | 38 -- 51 | -12.2629 | 3.87190 | 5.37459 |
| CD630_18311 | CD630_s0280 | 91 -- 102 | 19 -- 30 | -12.0001 | 6.86047 | 5.37459 |
| CD630_17970 | CD630_s0280 | 162 -- 197 | 55 -- 90 | -11.9114 | 3.68802 | 5.37459 |
| CD630_16640 | CD630_s0280 | 91 -- 126 | 55 -- 94 | -11.9053 | 3.61548 | 5.37459 |
| CD630_21090 | CD630_s0280 | 119 -- 160 | 50 -- 87 | -11.8462 | 4.15803 | 5.37459 |
| CD630_13530 | CD630_s0280 | 167 -- 199 | 31 -- 61 | -11.7982 | 4.09832 | 5.37459 |
| CD630_25050 | CD630_s0280 | 98 -- 164 | 34 -- 93 | -11.7981 | 4.33624 | 5.37459 |
| CD630_20450 | CD630_s0280 | 114 -- 136 | 126 -- 151 | -11.6055 | 3.77388 | 5.37459 |
| CD630_28150 | CD630_s0280 | 22 -- 61 | 54 -- 89 | -11.4534 | 4.18447 | 5.37459 |
| pilC | CD630_s0280 | 91 -- 102 | 19 -- 30 | -11.4092 | 3.54230 | 5.37459 |
| feoB | CD630_s0280 | 92 -- 141 | 45 -- 92 | -11.2115 | 4.52257 | 5.37459 |
| CD630_23440 | CD630_s0280 | 168 -- 190 | 215 -- 237 | -11.1796 | 3.34841 | 5.37459 |
| CD630_07840 | CD630_s0280 | 138 -- 181 | 44 -- 87 | -11.177 | 4.26761 | 5.37459 |
| CD630_31381 | CD630_s0280 | 20 -- 60 | 88 -- 128 | -11.1431 | 3.49137 | 5.37459 |
| appF | CD630_s0280 | 53 -- 74 | 63 -- 83 | -11.1371 | 4.25570 | 5.37459 |
| CD630_13560 | CD630_s0280 | 48 -- 92 | 39 -- 77 | -10.9978 | 3.38083 | 5.37459 |
| CD630_06170 | CD630_s0280 | 96 -- 124 | 178 -- 198 | -10.9246 | 4.95366 | 5.37459 |
| CD630_27280 | CD630_s0280 | 183 -- 196 | 179 -- 192 | -10.8961 | 3.24738 | 5.37459 |
| CD630_04320 | CD630_s0280 | 184 -- 196 | 218 -- 230 | -10.8697 | 3.68342 | 5.37459 |
| CD630_10790 | CD630_s0280 | 189 -- 201 | 58 -- 70 | -10.8594 | 4.45284 | 5.37459 |
| pmi | CD630_s0280 | 140 -- 161 | 210 -- 230 | -10.8115 | 3.86845 | 5.37459 |
| CD630_02720 | CD630_s0280 | 101 -- 151 | 179 -- 223 | -10.8026 | 3.38938 | 5.37459 |
| CD630_05840 | CD630_s0280 | 122 -- 159 | 42 -- 79 | -10.7989 | 4.28779 | 5.37459 |
| CD630_15890 | CD630_s0280 | 2 -- 40 | 39 -- 73 | -10.7985 | 3.39962 | 5.37459 |
| CD630_18970 | CD630_s0280 | 97 -- 129 | 78 -- 109 | -10.7226 | 3.57243 | 5.37459 |
| CD630_19800 | CD630_s0280 | 11 -- 33 | 177 -- 198 | -10.6981 | 5.22035 | 5.37459 |
| bcp | CD630_s0280 | 96 -- 129 | 83 -- 117 | -10.6379 | 3.16670 | 5.37459 |
| CD630_10950 | CD630_s0280 | 133 -- 188 | 47 -- 112 | -10.5753 | 3.67675 | 5.37459 |
| CD630_06050 | CD630_s0280 | 107 -- 144 | 56 -- 91 | -10.5198 | 3.59332 | 5.37459 |
| CD630_20650 | CD630_s0280 | 86 -- 128 | 45 -- 85 | -10.4421 | 4.00083 | 5.37459 |
| CD630_16330 | CD630_s0280 | 165 -- 182 | 212 -- 232 | -10.4127 | 4.11180 | 5.37459 |
| CD630_19690 | CD630_s0280 | 41 -- 79 | 57 -- 92 | -10.3824 | 3.85861 | 5.37459 |
| CD630_30130 | CD630_s0280 | 18 -- 96 | 55 -- 113 | -10.3661 | 3.28343 | 5.37459 |
| CD630_04610 | CD630_s0280 | 48 -- 56 | 209 -- 217 | -10.3617 | 3.83025 | 5.37459 |
| CD630_19840 | CD630_s0280 | 25 -- 64 | 53 -- 86 | -10.277 | 3.44724 | 5.37459 |
| CD630_21910 | CD630_s0280 | 179 -- 196 | 6 -- 23 | -10.2691 | 4.66600 | 5.37459 |
| CD630_05830 | CD630_s0280 | 11 -- 46 | 65 -- 96 | -10.2366 | 3.35360 | 5.37459 |
| CD630_18960 | CD630_s0280 | 16 -- 34 | 11 -- 29 | -10.2307 | 4.37822 | 5.37459 |
| CD630_05950 | CD630_s0280 | 175 -- 189 | 181 -- 195 | -10.049 | 3.31669 | 5.37459 |
| CD630_36180 | CD630_s0280 | 170 -- 201 | 221 -- 249 | -10.0395 | 4.83428 | 5.37459 |
| CD630_06190 | CD630_s0280 | 103 -- 133 | 54 -- 85 | -10.0388 | 3.57064 | 5.37459 |
| CD630_06340 | CD630_s0280 | 19 -- 49 | 40 -- 70 | -10.0207 | 4.35983 | 5.37459 |
| CD630_06110 | CD630_s0280 | 178 -- 191 | 213 -- 226 | -10.0093 | 5.61380 | 5.37459 |
| appC | CD630_s0280 | 153 -- 171 | 58 -- 77 | -9.95971 | 4.50438 | 5.37459 |
| CD630_21730 | CD630_s0280 | 167 -- 195 | 74 -- 100 | -9.9351 | 3.45076 | 5.37459 |
| CD630_06060 | CD630_s0280 | 55 -- 92 | 127 -- 165 | -9.91721 | 4.94050 | 5.37459 |
| CD630_05821 | CD630_s0280 | 174 -- 199 | 66 -- 92 | -9.87468 | 3.75661 | 5.37459 |
| CD630_18930 | CD630_s0280 | 46 -- 66 | 18 -- 39 | -9.87228 | 5.32127 | 5.37459 |
| sspA | CD630_s0280 | 51 -- 67 | 43 -- 63 | -9.84905 | 3.62689 | 5.37459 |
| CD630_27210 | CD630_s0280 | 116 -- 145 | 65 -- 92 | -9.82961 | 4.18786 | 5.37459 |
| CD630_25300 | CD630_s0280 | 61 -- 92 | 88 -- 118 | -9.74033 | 4.49333 | 5.37459 |
| CD630_13350 | CD630_s0280 | 88 -- 125 | 56 -- 87 | -9.72506 | 4.15139 | 5.37459 |
| CD630_25430 | CD630_s0280 | 137 -- 154 | 140 -- 156 | -9.69729 | 4.00527 | 5.37459 |
| CD630_36781 | CD630_s0280 | 161 -- 186 | 89 -- 116 | -9.65988 | 3.75946 | 5.37459 |
| CD630_03920 | CD630_s0280 | 19 -- 42 | 126 -- 150 | -9.65084 | 4.15934 | 5.37459 |
| CD630_21770 | CD630_s0280 | 142 -- 153 | 15 -- 26 | -9.62583 | 4.02587 | 5.37459 |
| CD630_06100 | CD630_s0280 | 165 -- 189 | 127 -- 150 | -9.61094 | 4.43918 | 5.37459 |
| CD630_11220 | CD630_s0280 | 48 -- 106 | 54 -- 116 | -9.57465 | 3.52327 | 5.37459 |
| CD630_36190 | CD630_s0280 | 152 -- 180 | 213 -- 240 | -9.54618 | 4.63449 | 5.37459 |
| CD630_05870 | CD630_s0281 | 14 -- 32 | 160 -- 178 | -17.4366 | 4.28339 | 8.30299 |
| CD630_28280 | CD630_s0281 | 12 -- 53 | 9 -- 51 | -16.3338 | 4.59833 | 8.30299 |
| gatC | CD630_s0281 | 43 -- 58 | 180 -- 195 | -16.2731 | 4.25143 | 8.30299 |
| CD630_32520 | CD630_s0281 | 163 -- 194 | 10 -- 41 | -15.6256 | 4.06098 | 8.30299 |
| CD630_08080 | CD630_s0281 | 27 -- 68 | 275 -- 300 | -15.4339 | 4.87819 | 8.30299 |
| CD630_18910 | CD630_s0281 | 40 -- 74 | 11 -- 45 | -14.4707 | 3.30688 | 8.30299 |
| CD630_06960 | CD630_s0281 | 148 -- 177 | 12 -- 48 | -14.0672 | 3.16317 | 8.30299 |
| bcp | CD630_s0281 | 120 -- 143 | 161 -- 180 | -14.0428 | 3.16670 | 8.30299 |
| CD630_22850 | CD630_s0281 | 8 -- 25 | 188 -- 205 | -13.9087 | 3.84102 | 8.30299 |
| pilT | CD630_s0281 | 127 -- 144 | 167 -- 186 | -13.8323 | 3.24639 | 8.30299 |
| CD630_25320 | CD630_s0281 | 117 -- 146 | 156 -- 191 | -13.6233 | 4.29415 | 8.30299 |
| CD630_14150 | CD630_s0281 | 35 -- 99 | 94 -- 171 | -13.5183 | 3.28251 | 8.30299 |
| eutP | CD630_s0281 | 160 -- 181 | 162 -- 182 | -13.4527 | 3.82847 | 8.30299 |
| CD630_19904 | CD630_s0281 | 24 -- 33 | 39 -- 48 | -13.4176 | 5.96526 | 8.30299 |
| CD630_33560 | CD630_s0281 | 94 -- 128 | 8 -- 44 | -13.3686 | 4.03147 | 8.30299 |
| CD630_14630 | CD630_s0281 | 140 -- 169 | 95 -- 110 | -13.0114 | 3.34485 | 8.30299 |
| CD630_24830 | CD630_s0281 | 138 -- 198 | 158 -- 216 | -12.9145 | 4.98239 | 8.30299 |
| eutW | CD630_s0281 | 11 -- 31 | 162 -- 180 | -12.7503 | 4.55715 | 8.30299 |
| CD630_30420 | CD630_s0281 | 38 -- 97 | 3 -- 47 | -12.6963 | 5.27539 | 8.30299 |
| CD630_19800 | CD630_s0281 | 24 -- 44 | 107 -- 126 | -12.3821 | 5.22035 | 8.30299 |
| cdu1 | CD630_s0281 | 147 -- 185 | 162 -- 200 | -12.3507 | 4.46276 | 8.30299 |
| CD630_14420 | CD630_s0281 | 2 -- 32 | 174 -- 204 | -12.2726 | 3.99497 | 8.30299 |
| CD630_29270 | CD630_s0281 | 7 -- 26 | 167 -- 186 | -12.2289 | 3.70174 | 8.30299 |
| CD630_05830 | CD630_s0281 | 114 -- 186 | 17 -- 82 | -12.2175 | 3.35360 | 8.30299 |
| CD630_24050 | CD630_s0281 | 2 -- 46 | 11 -- 54 | -12.1977 | 3.70192 | 8.30299 |
| CD630_09570 | CD630_s0281 | 23 -- 52 | 72 -- 104 | -12.0503 | 5.31213 | 8.30299 |
| CD630_29070 | CD630_s0281 | 23 -- 52 | 72 -- 104 | -12.0503 | 3.89477 | 8.30299 |
| CD630_05820 | CD630_s0281 | 82 -- 107 | 89 -- 112 | -12.0303 | 3.75661 | 8.30299 |
| CD630_06170 | CD630_s0281 | 108 -- 127 | 115 -- 134 | -11.9615 | 4.95366 | 8.30299 |
| CD630_07470 | CD630_s0281 | 23 -- 60 | 6 -- 47 | -11.8464 | 3.37143 | 8.30299 |
| dnaD | CD630_s0281 | 147 -- 196 | 167 -- 196 | -11.8158 | 3.56078 | 8.30299 |
| CD630_06060 | CD630_s0281 | 142 -- 178 | 8 -- 46 | -11.7554 | 4.94050 | 8.30299 |
| CD630_19030 | CD630_s0281 | 2 -- 33 | 23 -- 56 | -11.723 | 3.34240 | 8.30299 |
| CD630_28181 | CD630_s0281 | 125 -- 163 | 156 -- 192 | -11.5695 | 3.91299 | 8.30299 |
| CD630_06100 | CD630_s0281 | 110 -- 150 | 269 -- 304 | -11.5328 | 4.43918 | 8.30299 |
| CD630_13020 | CD630_s0281 | 142 -- 157 | 162 -- 179 | -11.4405 | 3.52267 | 8.30299 |
| CD630_33300 | CD630_s0281 | 165 -- 178 | 152 -- 165 | -11.3532 | 3.79149 | 8.30299 |
| CD630_20260 | CD630_s0281 | 110 -- 149 | 165 -- 204 | -11.3465 | 3.84162 | 8.30299 |
| CD630_25050 | CD630_s0281 | 13 -- 41 | 21 -- 50 | -11.3167 | 4.33624 | 8.30299 |
| CD630_19850 | CD630_s0281 | 134 -- 159 | 277 -- 300 | -11.2796 | 6.88486 | 8.30299 |
| CD630_19840 | CD630_s0281 | 129 -- 161 | 177 -- 204 | -11.2096 | 3.44724 | 8.30299 |
| CD630_14250 | CD630_s0281 | 175 -- 201 | 163 -- 187 | -11.1725 | 4.19604 | 8.30299 |
| CD630_22510 | CD630_s0281 | 111 -- 139 | 272 -- 298 | -11.1594 | 11.42763 | 8.30299 |
| CD630_06140 | CD630_s0281 | 54 -- 90 | 156 -- 195 | -11.1515 | 4.12519 | 8.30299 |
| CD630_08130 | CD630_s0281 | 168 -- 191 | 77 -- 100 | -11.1424 | 4.63831 | 8.30299 |
| cstA | CD630_s0281 | 42 -- 91 | 156 -- 211 | -11.1336 | 3.53584 | 8.30299 |
| opuCA | CD630_s0281 | 99 -- 135 | 157 -- 193 | -11.1322 | 4.07035 | 8.30299 |
| uxaA%27 | CD630_s0281 | 4 -- 22 | 36 -- 53 | -11.1172 | 4.26165 | 8.30299 |
| CD630_06690 | CD630_s0281 | 90 -- 103 | 87 -- 100 | -11.1092 | 5.82709 | 8.30299 |
| CD630_20600 | CD630_s0281 | 168 -- 199 | 169 -- 193 | -11.0317 | 4.20093 | 8.30299 |
| mobA | CD630_s0281 | 138 -- 177 | 155 -- 193 | -10.9484 | 4.25703 | 8.30299 |
| appF | CD630_s0281 | 42 -- 92 | 157 -- 205 | -10.8943 | 4.25570 | 8.30299 |
| CD630_06642 | CD630_s0281 | 151 -- 168 | 164 -- 182 | -10.8104 | 3.32967 | 8.30299 |
| CD630_21030 | CD630_s0281 | 35 -- 73 | 9 -- 47 | -10.7673 | 3.08733 | 8.30299 |
| feoB | CD630_s0281 | 4 -- 33 | 154 -- 180 | -10.741 | 4.52257 | 8.30299 |
| CD630_27600 | CD630_s0281 | 123 -- 135 | 7 -- 19 | -10.7137 | 3.48384 | 8.30299 |
| CD630_23770 | CD630_s0281 | 32 -- 52 | 162 -- 182 | -10.6825 | 3.25181 | 8.30299 |
| CD630_21730 | CD630_s0281 | 158 -- 175 | 167 -- 182 | -10.6424 | 3.45076 | 8.30299 |
| cdu2 | CD630_s0281 | 57 -- 94 | 155 -- 193 | -10.6098 | 2.97335 | 8.30299 |
| sigK | CD630_s0281 | 53 -- 66 | 167 -- 180 | -10.5909 | 3.08122 | 8.30299 |
| CD630_23080 | CD630_s0281 | 22 -- 67 | 110 -- 153 | -10.5379 | 5.23027 | 8.30299 |
| cobQ | CD630_s0281 | 184 -- 200 | 165 -- 182 | -10.5335 | 3.13524 | 8.30299 |
| eutV | CD630_s0281 | 146 -- 179 | 154 -- 188 | -10.4867 | 4.55715 | 8.30299 |
| CD630_08390 | CD630_s0281 | 81 -- 94 | 92 -- 106 | -10.4811 | 5.16000 | 8.30299 |
| CD630_22880 | CD630_s0281 | 49 -- 77 | 270 -- 299 | -10.4711 | 3.62455 | 8.30299 |
| CD630_24760 | CD630_s0281 | 104 -- 123 | 161 -- 180 | -10.4638 | 5.19485 | 8.30299 |
| CD630_05160 | CD630_s0281 | 162 -- 186 | 157 -- 180 | -10.4189 | 5.03573 | 8.30299 |
| CD630_31461 | CD630_s0281 | 44 -- 63 | 162 -- 182 | -10.3708 | 4.41476 | 8.30299 |
| CD630_24590 | CD630_s0281 | 121 -- 150 | 155 -- 189 | -10.3655 | 3.70401 | 8.30299 |
| srlD | CD630_s0281 | 34 -- 60 | 164 -- 191 | -10.3605 | 4.19843 | 8.30299 |
| CD630_14990 | CD630_s0281 | 3 -- 35 | 157 -- 189 | -10.3084 | 3.78619 | 8.30299 |
| CD630_23820 | CD630_s0281 | 92 -- 104 | 97 -- 110 | -10.3084 | 3.18419 | 8.30299 |
| CD630_08940 | CD630_s0281 | 158 -- 177 | 161 -- 182 | -10.3081 | 3.72225 | 8.30299 |
| CD630_08140 | CD630_s0281 | 6 -- 52 | 273 -- 300 | -10.2851 | 3.30159 | 8.30299 |
| dltB | CD630_s0281 | 5 -- 19 | 121 -- 136 | -10.2593 | 3.81485 | 8.30299 |
| CD630_21750 | CD630_s0281 | 121 -- 138 | 193 -- 210 | -10.2537 | 3.70626 | 8.30299 |
| CD630_07480 | CD630_s0281 | 149 -- 194 | 12 -- 57 | -10.2228 | 5.18321 | 8.30299 |
| CD630_16550 | CD630_s0281 | 187 -- 201 | 166 -- 179 | -10.2014 | 5.61064 | 8.30299 |
| gcp | CD630_s0281 | 51 -- 70 | 38 -- 56 | -10.1959 | 4.92798 | 8.30299 |
| CD630_13380 | CD630_s0281 | 53 -- 85 | 156 -- 191 | -10.1632 | 4.74408 | 8.30299 |
| CD630_09100 | CD630_s0281 | 6 -- 42 | 273 -- 301 | -10.1547 | 5.92260 | 8.30299 |
| CD630_15431 | CD630_s0281 | 8 -- 27 | 90 -- 107 | -10.1515 | 5.29780 | 8.30299 |
| cwp17 | CD630_s0281 | 114 -- 133 | 156 -- 175 | -10.1074 | 3.43363 | 8.30299 |
| CD630_09970 | CD630_s0281 | 16 -- 60 | 152 -- 191 | -10.0991 | 3.25917 | 8.30299 |
| CD630_24220 | CD630_s0281 | 82 -- 95 | 97 -- 109 | -10.088 | 4.18037 | 8.30299 |
| CD630_32890 | CD630_s0281 | 34 -- 54 | 177 -- 198 | -10.0851 | 3.41276 | 8.30299 |
| CD630_10050 | CD630_s0281 | 174 -- 199 | 160 -- 182 | -10.0059 | 3.93750 | 8.30299 |
| CD630_17720 | CD630_s0281 | 11 -- 41 | 21 -- 50 | -9.99952 | 3.22910 | 8.30299 |
| CD630_04620 | CD630_s0281 | 155 -- 182 | 165 -- 198 | -9.99485 | 3.45413 | 8.30299 |
| CD630_13530 | CD630_s0281 | 1 -- 24 | 113 -- 135 | -9.99183 | 4.09832 | 8.30299 |
| nusB | CD630_s0281 | 110 -- 120 | 155 -- 165 | -9.97229 | 4.92798 | 8.30299 |
| CD630_09060 | CD630_s0281 | 53 -- 72 | 160 -- 180 | -9.94141 | 4.31834 | 8.30299 |
| CD630_13400 | CD630_s0281 | 189 -- 198 | 36 -- 45 | -9.93739 | 3.20232 | 8.30299 |
| cdtR | CD630_s0281 | 94 -- 143 | 36 -- 87 | -9.89613 | 5.84368 | 8.30299 |
| CD630_01560 | CD630_s0281 | 109 -- 144 | 158 -- 191 | -9.87898 | 4.06352 | 8.30299 |
| CD630_18960 | CD630_s0281 | 61 -- 95 | 161 -- 188 | -9.83193 | 4.37822 | 8.30299 |
| CD630_18440 | CD630_s0281 | 50 -- 75 | 167 -- 194 | -9.82511 | 4.07960 | 8.30299 |
| CD630_03410 | CD630_s0281 | 49 -- 88 | 19 -- 55 | -9.82122 | 4.45994 | 8.30299 |
| kdpD | CD630_s0281 | 13 -- 38 | 273 -- 297 | -9.79057 | 4.27140 | 8.30299 |
| CD630_04341 | CD630_s0281 | 37 -- 59 | 97 -- 116 | -9.77356 | 4.29262 | 8.30299 |
| CD630_34520 | CD630_s0281 | 116 -- 133 | 167 -- 185 | -9.76856 | 3.83016 | 8.30299 |
| cheW | CD630_s0281 | 118 -- 143 | 170 -- 194 | -9.76412 | 3.72371 | 8.30299 |
| CD630_14010 | CD630_s0281 | 17 -- 27 | 167 -- 177 | -9.75571 | 3.69983 | 8.30299 |
| CD630_26630 | CD630_s0281 | 69 -- 93 | 157 -- 192 | -9.75325 | 4.61557 | 8.30299 |
| CD630_14600 | CD630_s0281 | 173 -- 198 | 185 -- 209 | -9.7364 | 5.90348 | 8.30299 |
| CD630_04280 | CD630_s0281 | 84 -- 96 | 96 -- 109 | -9.73216 | 3.61457 | 8.30299 |
| CD630_30190 | CD630_s0281 | 171 -- 182 | 121 -- 132 | -9.72114 | 3.27394 | 8.30299 |
| CD630_05490 | CD630_s0281 | 65 -- 76 | 36 -- 46 | -9.7102 | 3.89852 | 8.30299 |
| CD630_16450 | CD630_s0281 | 65 -- 115 | 85 -- 128 | -9.7034 | 4.57547 | 8.30299 |
| cbiO | CD630_s0281 | 146 -- 164 | 159 -- 177 | -9.64662 | 3.79830 | 8.30299 |
| CD630_28560 | CD630_s0281 | 28 -- 47 | 109 -- 129 | -9.61636 | 3.96803 | 8.30299 |
| CD630_05280 | CD630_s0281 | 161 -- 174 | 157 -- 170 | -9.59845 | 3.91665 | 8.30299 |
| CD630_03180 | CD630_s0281 | 189 -- 201 | 168 -- 180 | -9.59 | 3.73138 | 8.30299 |
| bglA | CD630_s0281 | 74 -- 90 | 161 -- 177 | -9.55452 | 4.86402 | 8.30299 |
| CD630_06410 | CD630_s0281 | 1 -- 36 | 21 -- 49 | -9.55333 | 4.14404 | 8.30299 |
| CD630_22710 | CD630_s0281 | 119 -- 144 | 160 -- 182 | -9.52092 | 4.05623 | 8.30299 |
| ldh | CD630_s0281 | 146 -- 191 | 150 -- 201 | -9.51627 | 4.35818 | 8.30299 |
| CD630_18950 | CD630_s0281 | 2 -- 30 | 8 -- 37 | -9.51433 | 3.40610 | 8.30299 |
| CD630_17281 | CD630_s0281 | 11 -- 34 | 181 -- 205 | -9.50715 | 5.91544 | 8.30299 |
| CD630_15740 | CD630_s0281 | 27 -- 70 | 36 -- 75 | -9.50433 | 4.43222 | 8.30299 |
| CD630_19401 | CD630_s0300 | 27 -- 127 | 6 -- 104 | -17.6705 | 4.37084 | 4.17452 |

| | | | | | | | |
|---|---|---|---|---|---|---|---|
| CD630_16640 | CD630_s0300 | 80 -- 116 | 126 -- 161 | -15.0996 | | 3.61548 | 4.17452 |
| CD630_27280 | CD630_s0300 | 7 -- 40 | 128 -- 166 | -14.6776 | | 3.24738 | 4.17452 |
| thiH | CD630_s0300 | 42 -- 73 | 200 -- 228 | -13.8771 | | 3.58230 | 4.17452 |
| CD630_14130 | CD630_s0300 | 169 -- 198 | 39 -- 66 | -13.5254 | | 4.57970 | 4.17452 |
| feoB | CD630_s0300 | 35 -- 86 | 48 -- 107 | -13.4141 | | 4.05364 | 4.17452 |
| etfA | CD630_s0300 | 45 -- 166 | 41 -- 146 | -13.2066 | | 3.41204 | 4.17452 |
| CD630_35780 | CD630_s0300 | 57 -- 68 | 83 -- 94 | -13.0701 | | 3.50107 | 4.17452 |
| CD630_19700 | CD630_s0300 | 170 -- 187 | 91 -- 107 | -12.8336 | | 3.85861 | 4.17452 |
| CD630_09970 | CD630_s0300 | 87 -- 102 | 212 -- 227 | -12.7886 | | 3.25917 | 4.17452 |
| CD630_28181 | CD630_s0300 | 146 -- 163 | 215 -- 232 | -12.7415 | | 3.91299 | 4.17452 |
| CD630_24050 | CD630_s0300 | 126 -- 140 | 33 -- 47 | -12.7169 | | 3.70192 | 4.17452 |
| CD630_34080 | CD630_s0300 | 126 -- 171 | 55 -- 105 | -12.6367 | | 3.20224 | 4.17452 |
| CD630_27210 | CD630_s0300 | 79 -- 99 | 209 -- 229 | -12.5058 | | 4.18786 | 4.17452 |
| CD630_27220 | CD630_s0300 | 110 -- 142 | 137 -- 170 | -12.3328 | | 4.18786 | 4.17452 |
| CD630_10000 | CD630_s0300 | 1 -- 24 | 40 -- 64 | -12.199 | | 3.86040 | 4.17452 |
| CD630_27570 | CD630_s0300 | 89 -- 126 | 82 -- 113 | -12.1901 | | 3.46216 | 4.17452 |
| CD630_15740 | CD630_s0300 | 11 -- 48 | 134 -- 167 | -12.0978 | | 4.43222 | 4.17452 |
| CD630_03170 | CD630_s0300 | 7 -- 37 | 126 -- 158 | -12.0337 | | 3.66946 | 4.17452 |
| CD630_32520 | CD630_s0300 | 128 -- 186 | 125 -- 180 | -11.9568 | | 4.06098 | 4.17452 |
| CD630_22030 | CD630_s0300 | 5 -- 33 | 80 -- 102 | -11.801 | | 3.29934 | 4.17452 |
| CD630_15890 | CD630_s0300 | 24 -- 53 | 128 -- 156 | -11.7888 | | 3.39962 | 4.17452 |
| CD630_19800 | CD630_s0300 | 14 -- 62 | 48 -- 87 | -11.7619 | | 5.22035 | 4.17452 |
| cbiQ | CD630_s0300 | 159 -- 187 | 29 -- 62 | -11.6618 | | 3.79830 | 4.17452 |
| appC | CD630_s0300 | 36 -- 60 | 125 -- 148 | -11.4002 | | 4.50438 | 4.17452 |
| CD630_28290 | CD630_s0300 | 185 -- 195 | 69 -- 79 | -11.3779 | | 5.54192 | 4.17452 |
| CD630_36190 | CD630_s0300 | 79 -- 93 | 218 -- 232 | -11.3599 | | 4.63449 | 4.17452 |
| CD630_03410 | CD630_s0300 | 99 -- 178 | 105 -- 175 | -11.2462 | | 4.45994 | 4.17452 |
| dnaC | CD630_s0300 | 128 -- 157 | 105 -- 132 | -11.2436 | | 6.30796 | 4.17452 |
| CD630_25150 | CD630_s0300 | 76 -- 93 | 218 -- 235 | -11.2387 | | 3.79339 | 4.17452 |
| CD630_31580 | CD630_s0300 | 186 -- 200 | 47 -- 62 | -11.2234 | | 3.51756 | 4.17452 |
| xdhA | CD630_s0300 | 12 -- 45 | 105 -- 142 | -11.1441 | | 4.45147 | 4.17452 |
| CD630_05150 | CD630_s0300 | 80 -- 98 | 216 -- 233 | -11.0855 | | 3.69875 | 4.17452 |
| CD630_29750 | CD630_s0300 | 20 -- 38 | 209 -- 228 | -11.0796 | | 4.59146 | 4.17452 |
| garK | CD630_s0300 | 13 -- 28 | 212 -- 225 | -11.0729 | | 3.72836 | 4.17452 |
| CD630_17170 | CD630_s0300 | 82 -- 98 | 213 -- 230 | -11.0174 | | 4.10230 | 4.17452 |
| CD630_17151 | CD630_s0300 | 171 -- 197 | 214 -- 232 | -11.0055 | | 6.43124 | 4.17452 |
| appB | CD630_s0300 | 174 -- 195 | 86 -- 107 | -10.9794 | | 6.44103 | 4.17452 |
| CD630_31530 | CD630_s0300 | 144 -- 167 | 29 -- 50 | -10.9356 | | 3.49100 | 4.17452 |
| CD630_06540 | CD630_s0300 | 86 -- 118 | 189 -- 225 | -10.9179 | | 3.54673 | 4.17452 |
| nusB | CD630_s0300 | 36 -- 63 | 21 -- 50 | -10.9151 | | 4.92798 | 4.17452 |
| CD630_13020 | CD630_s0300 | 113 -- 162 | 118 -- 163 | -10.9 | | 3.52267 | 4.17452 |
| CD630_33480 | CD630_s0300 | 86 -- 112 | 208 -- 225 | -10.7822 | | 3.67613 | 4.17452 |
| CD630_14920 | CD630_s0300 | 102 -- 179 | 22 -- 88 | -10.7778 | | 4.35225 | 4.17452 |
| CD630_27230 | CD630_s0300 | 133 -- 142 | 81 -- 90 | -10.7726 | | 3.52462 | 4.17452 |
| CD630_21020 | CD630_s0300 | 100 -- 155 | 126 -- 174 | -10.7652 | | 4.28666 | 4.17452 |
| CD630_34560 | CD630_s0300 | 72 -- 99 | 207 -- 234 | -10.7502 | | 4.02604 | 4.17452 |
| CD630_08460 | CD630_s0300 | 147 -- 161 | 14 -- 28 | -10.7393 | | 4.15083 | 4.17452 |
| CD630_10950 | CD630_s0300 | 74 -- 94 | 213 -- 234 | -10.7233 | | 3.67675 | 4.17452 |
| CD630_04850 | CD630_s0300 | 128 -- 197 | 118 -- 166 | -10.7203 | | 4.40788 | 4.17452 |
| CD630_31210 | CD630_s0300 | 104 -- 119 | 148 -- 163 | -10.699 | | 4.17693 | 4.17452 |
| CD630_21090 | CD630_s0300 | 147 -- 171 | 30 -- 55 | -10.6821 | | 4.15803 | 4.17452 |
| CD630_06560 | CD630_s0300 | 1 -- 47 | 111 -- 153 | -10.6214 | | 4.02532 | 4.17452 |
| CD630_13350 | CD630_s0300 | 153 -- 170 | 1 -- 17 | -10.6112 | | 4.15139 | 4.17452 |
| CD630_25610 | CD630_s0300 | 9 -- 61 | 111 -- 150 | -10.5935 | | 3.02685 | 4.17452 |
| CD630_07840 | CD630_s0300 | 169 -- 178 | 44 -- 53 | -10.5145 | | 4.26761 | 4.17452 |
| CD630_05030 | CD630_s0300 | 32 -- 39 | 194 -- 201 | -10.4648 | | 4.03823 | 4.17452 |
| malY | CD630_s0300 | 75 -- 118 | 124 -- 174 | -10.3773 | | 4.46345 | 4.17452 |
| ogt1 | CD630_s0300 | 83 -- 102 | 212 -- 230 | -10.3712 | | 3.16772 | 4.17452 |
| cprK | CD630_s0300 | 141 -- 159 | 128 -- 146 | -10.2719 | | 3.96216 | 4.17452 |
| CD630_18311 | CD630_s0300 | 165 -- 186 | 142 -- 163 | -10.2658 | | 6.86047 | 4.17452 |
| tcdA | CD630_s0300 | 88 -- 111 | 65 -- 89 | -10.2447 | | 4.70593 | 4.17452 |
| CD630_19440 | CD630_s0300 | 163 -- 191 | 136 -- 168 | -10.2315 | | 4.53937 | 4.17452 |
| CD630_20650 | CD630_s0300 | 101 -- 129 | 109 -- 141 | -10.2236 | | 4.00083 | 4.17452 |
| CD630_28560 | CD630_s0300 | 30 -- 56 | 50 -- 77 | -10.1859 | | 3.96803 | 4.17452 |
| CD630_06060 | CD630_s0300 | 129 -- 162 | 16 -- 46 | -10.1673 | | 4.94050 | 4.17452 |
| cwp17 | CD630_s0300 | 126 -- 152 | 22 -- 47 | -10.1659 | | 3.43363 | 4.17452 |
| CD630_18440 | CD630_s0300 | 8 -- 20 | 40 -- 51 | -10.1639 | | 4.07960 | 4.17452 |
| CD630_06170 | CD630_s0300 | 151 -- 164 | 147 -- 162 | -10.158 | | 4.95366 | 4.17452 |
| CD630_27360 | CD630_s0300 | 185 -- 192 | 44 -- 51 | -10.1454 | | 3.79570 | 4.17452 |
| CD630_06340 | CD630_s0300 | 127 -- 153 | 137 -- 164 | -10.1021 | | 4.35983 | 4.17452 |
| CD630_32890 | CD630_s0300 | 136 -- 164 | 142 -- 168 | -10.0493 | | 3.41276 | 4.17452 |
| CD630_20960 | CD630_s0300 | 55 -- 81 | 208 -- 232 | -10.0492 | | 5.65940 | 4.17452 |
| CD630_31480 | CD630_s0300 | 146 -- 181 | 131 -- 170 | -10.0361 | | 4.88897 | 4.17452 |
| CD630_08080 | CD630_s0300 | 149 -- 168 | 48 -- 78 | -10.0331 | | 4.87819 | 4.17452 |
| CD630_31970 | CD630_s0300 | 115 -- 129 | 92 -- 107 | -10.0209 | | 3.24511 | 4.17452 |
| CD630_05230 | CD630_s0300 | 1 -- 55 | 40 -- 88 | -10.019 | | 3.73787 | 4.17452 |
| CD630_14340 | CD630_s0300 | 71 -- 93 | 64 -- 87 | -9.93793 | | 5.10868 | 4.17452 |
| CD630_32040 | CD630_s0300 | 156 -- 169 | 99 -- 112 | -9.86277 | | 4.93516 | 4.17452 |
| CD630_02420 | CD630_s0300 | 129 -- 156 | 85 -- 119 | -9.85996 | | 3.71048 | 4.17452 |
| CD630_06100 | CD630_s0300 | 142 -- 196 | 1 -- 70 | -9.78325 | | 4.43918 | 4.17452 |
| spaK | CD630_s0300 | 109 -- 118 | 69 -- 78 | -9.73515 | | 3.04530 | 4.17452 |
| pilC | CD630_s0300 | 131 -- 139 | 216 -- 225 | -9.68049 | | 3.54230 | 4.17452 |
| panB | CD630_s0300 | 148 -- 180 | 132 -- 170 | -9.61097 | | 3.40248 | 4.17452 |
| CD630_05211 | CD630_s0300 | 8 -- 20 | 4 -- 16 | -9.60644 | | 3.97048 | 4.17452 |
| pilW | CD630_s0300 | 179 -- 186 | 48 -- 55 | -9.60101 | | 4.96316 | 4.17452 |
| CD630_23770 | CD630_s0300 | 61 -- 119 | 10 -- 77 | -9.59991 | | 3.25181 | 4.17452 |
| CD630_13960 | CD630_s0300 | 81 -- 102 | 209 -- 230 | -9.59599 | | 3.30375 | 4.17452 |
| CD630_24920 | CD630_s0300 | 2 -- 17 | 127 -- 142 | -9.5952 | | 3.89455 | 4.17452 |
| CD630_29960 | CD630_s0300 | 49 -- 63 | 126 -- 141 | -9.58182 | | 4.51904 | 4.17452 |
| CD630_04280 | CD630_s0300 | 128 -- 167 | 16 -- 49 | -9.56769 | | 3.61457 | 4.17452 |
| CD630_04341 | CD630_s0300 | 124 -- 139 | 209 -- 223 | -9.52106 | | 4.29262 | 4.17452 |
| CD630_35780 | CD630_s0310 | 23 -- 68 | 83 -- 130 | -16.2772 | | 3.50107 | 4.17452 |
| CD630_05030 | CD630_s0310 | 32 -- 43 | 188 -- 199 | -16.2488 | | 4.03823 | 4.17452 |
| CD630_02420 | CD630_s0310 | 14 -- 55 | 86 -- 126 | -15.6639 | | 3.71048 | 4.17452 |
| CD630_14620 | CD630_s0310 | 150 -- 183 | 85 -- 114 | -15.5235 | | 4.43651 | 4.17452 |
| CD630_16950 | CD630_s0310 | 167 -- 199 | 81 -- 112 | -14.6684 | | 4.42569 | 4.17452 |
| uxaA%27 | CD630_s0310 | 150 -- 197 | 53 -- 103 | -14.2987 | | 4.26165 | 4.17452 |
| CD630_31500 | CD630_s0310 | 154 -- 198 | 65 -- 111 | -12.9738 | | 4.90207 | 4.17452 |
| CD630_21730 | CD630_s0310 | 116 -- 198 | 75 -- 168 | -12.6805 | | 3.45076 | 4.17452 |
| CD630_08780 | CD630_s0310 | 129 -- 189 | 63 -- 112 | -12.4681 | | 7.76690 | 4.17452 |
| CD630_23450 | CD630_s0310 | 151 -- 189 | 65 -- 102 | -12.2944 | | 3.27218 | 4.17452 |
| CD630_36690 | CD630_s0310 | 146 -- 169 | 192 -- 211 | -12.0645 | | 3.53761 | 4.17452 |
| CD630_06050 | CD630_s0310 | 114 -- 140 | 118 -- 147 | -11.8468 | | 3.59332 | 4.17452 |
| CD630_21720 | CD630_s0310 | 138 -- 176 | 55 -- 101 | -11.5758 | | 4.13030 | 4.17452 |
| CD630_06170 | CD630_s0310 | 142 -- 192 | 124 -- 164 | -11.3766 | | 4.95366 | 4.17452 |
| CD630_22880 | CD630_s0310 | 14 -- 41 | 7 -- 39 | -11.2282 | | 3.62455 | 4.17452 |
| CD630_29250 | CD630_s0310 | 183 -- 196 | 158 -- 171 | -11.1332 | | 4.61901 | 4.17452 |
| CD630_09410 | CD630_s0310 | 183 -- 196 | 158 -- 171 | -11.1332 | | 4.75651 | 4.17452 |
| CD630_18120 | CD630_s0310 | 48 -- 64 | 157 -- 173 | -11.0668 | | 3.99713 | 4.17452 |
| CD630_14810 | CD630_s0310 | 1 -- 24 | 89 -- 114 | -11.0131 | | 3.35323 | 4.17452 |
| CD630_22030 | CD630_s0310 | 5 -- 33 | 80 -- 102 | -10.9941 | | 3.29934 | 4.17452 |
| CD630_27230 | CD630_s0310 | 133 -- 142 | 81 -- 90 | -10.9882 | | 3.52462 | 4.17452 |
| CD630_14340 | CD630_s0310 | 71 -- 86 | 71 -- 87 | -10.9188 | | 5.10868 | 4.17452 |
| CD630_03400 | CD630_s0310 | 1 -- 40 | 6 -- 38 | -10.8843 | | 4.45994 | 4.17452 |
| dltB | CD630_s0310 | 150 -- 178 | 84 -- 108 | -10.791 | | 3.81485 | 4.17452 |
| CD630_10880 | CD630_s0310 | 159 -- 181 | 117 -- 138 | -10.7522 | | 5.05048 | 4.17452 |
| CD630_07400 | CD630_s0310 | 44 -- 98 | 101 -- 157 | -10.6859 | | 4.52273 | 4.17452 |
| kdgT1 | CD630_s0310 | 186 -- 200 | 87 -- 100 | -10.6137 | | 4.49207 | 4.17452 |
| CD630_06100 | CD630_s0310 | 169 -- 196 | 1 -- 40 | -10.4735 | | 4.43918 | 4.17452 |
| CD630_06110 | CD630_s0310 | 177 -- 192 | 191 -- 206 | -10.4356 | | 5.61380 | 4.17452 |
| CD630_08080 | CD630_s0310 | 57 -- 68 | 7 -- 18 | -10.3745 | | 4.87819 | 4.17452 |
| CD630_19903 | CD630_s0310 | 166 -- 192 | 91 -- 114 | -10.245 | | 5.79994 | 4.17452 |
| CD630_31461 | CD630_s0310 | 1 -- 19 | 104 -- 124 | -10.2192 | | 4.41476 | 4.17452 |
| CD630_14240 | CD630_s0310 | 166 -- 192 | 91 -- 114 | -10.1636 | | 8.60766 | 4.17452 |
| CD630_03170 | CD630_s0310 | 7 -- 21 | 141 -- 154 | -10.1519 | | 3.66946 | 4.17452 |
| CD630_13350 | CD630_s0310 | 163 -- 170 | 44 -- 51 | -10.0336 | | 4.15139 | 4.17452 |
| CD630_09770 | CD630_s0310 | 193 -- 201 | 85 -- 93 | -10.0183 | | 3.22106 | 4.17452 |
| CD630_13380 | CD630_s0310 | 122 -- 198 | 32 -- 99 | -9.94245 | | 4.74408 | 4.17452 |
| CD630_18960 | CD630_s0310 | 52 -- 92 | 121 -- 169 | -9.76066 | | 4.37822 | 4.17452 |
| CD630_28580 | CD630_s0310 | 156 -- 175 | 38 -- 56 | -9.74526 | | 4.67130 | 4.17452 |
| spaK | CD630_s0310 | 17 -- 55 | 84 -- 109 | -9.74423 | | 3.04530 | 4.17452 |
| opuCA | CD630_s0310 | 171 -- 195 | 78 -- 102 | -9.74251 | | 4.07035 | 4.17452 |
| CD630_13020 | CD630_s0310 | 121 -- 155 | 86 -- 128 | -9.74183 | | 3.52267 | 4.17452 |
| ldh | CD630_s0310 | 131 -- 171 | 72 -- 109 | -9.72244 | | 4.35818 | 4.17452 |
| CD630_15360 | CD630_s0310 | 163 -- 190 | 67 -- 94 | -9.71975 | | 4.47208 | 4.17452 |
| CD630_19401 | CD630_s0310 | 37 -- 127 | 6 -- 93 | -9.7119 | | 4.37084 | 4.17452 |
| CD630_10050 | CD630_s0310 | 158 -- 182 | 82 -- 112 | -9.70531 | | 3.93750 | 4.17452 |
| CD630_08130 | CD630_s0310 | 111 -- 155 | 2 -- 62 | -9.65382 | | 4.63831 | 4.17452 |
| CD630_32880 | CD630_s0310 | 184 -- 193 | 81 -- 90 | -9.55531 | | 3.41276 | 4.17452 |
| CD630_13930 | CD630_s0320 | 134 -- 168 | 15 -- 38 | -13.2013 | | 4.09246 | 8.15991 |
| etfA | CD630_s0320 | 81 -- 90 | 41 -- 50 | -12.5591 | | 4.80521 | 8.15991 |
| CD630_30420 | CD630_s0320 | 138 -- 159 | 5 -- 24 | -12.5157 | | 5.27539 | 8.15991 |
| CD630_13530 | CD630_s0320 | 160 -- 194 | 15 -- 42 | -12.1975 | | 4.09832 | 8.15991 |
| CD630_24590 | CD630_s0320 | 176 -- 192 | 44 -- 59 | -11.8521 | | 3.70401 | 8.15991 |
| CD630_21760 | CD630_s0320 | 117 -- 133 | 78 -- 94 | -11.7807 | | 4.43846 | 8.15991 |
| kdgT1 | CD630_s0320 | 182 -- 200 | 19 -- 37 | -11.6806 | | 4.49207 | 8.15991 |
| appC | CD630_s0320 | 156 -- 169 | 24 -- 37 | -11.5751 | | 4.50438 | 8.15991 |
| spoVFB | CD630_s0320 | 1 -- 23 | 8 -- 31 | -10.8822 | | 4.62758 | 8.15991 |
| CD630_15550 | CD630_s0320 | 126 -- 133 | 17 -- 24 | -10.8398 | | 3.49019 | 8.15991 |
| CD630_36180 | CD630_s0320 | 18 -- 27 | 10 -- 19 | -10.804 | | 4.83428 | 8.15991 |
| CD630_18090 | CD630_s0320 | 140 -- 157 | 21 -- 38 | -10.7403 | | 4.23961 | 8.15991 |
| appB | CD630_s0320 | 185 -- 201 | 12 -- 28 | -10.7323 | | 6.44103 | 8.15991 |
| dnaC | CD630_s0320 | 32 -- 50 | 22 -- 37 | -10.7169 | | 6.30796 | 8.15991 |
| CD630_03650 | CD630_s0320 | 150 -- 164 | 7 -- 20 | -10.6923 | | 3.93697 | 8.15991 |
| CD630_21070 | CD630_s0320 | 37 -- 52 | 22 -- 38 | -10.6305 | | 5.35019 | 8.15991 |
| opuCC | CD630_s0320 | 103 -- 112 | 16 -- 25 | -10.5432 | | 4.07035 | 8.15991 |
| CD630_09030 | CD630_s0320 | 128 -- 152 | 16 -- 38 | -10.2878 | | 4.21282 | 8.15991 |
| CD630_25200 | CD630_s0320 | 124 -- 134 | 44 -- 54 | -10.1923 | | 3.65158 | 8.15991 |
| cdtR | CD630_s0320 | 22 -- 39 | 1 -- 16 | -10.1164 | | 5.84368 | 8.15991 |
| CD630_08370 | CD630_s0320 | 191 -- 201 | 8 -- 18 | -10.0865 | | 3.65639 | 8.15991 |
| CD630_22030 | CD630_s0320 | 5 -- 20 | 13 -- 27 | -10.0764 | | 3.29934 | 8.15991 |
| oppB | CD630_s0320 | 89 -- 97 | 37 -- 45 | -10.0669 | | 2.98328 | 8.15991 |
| CD630_09771 | CD630_s0320 | 107 -- 120 | 79 -- 92 | -9.99337 | | 4.95362 | 8.15991 |
| CD630_18090 | CD630_s0320 | 5 -- 37 | 28 -- 57 | -9.7279 | | 4.23961 | 8.15991 |
| CD630_14630 | CD630_s0320 | 186 -- 195 | 43 -- 52 | -9.63557 | | 3.34485 | 8.15991 |
| vncR | CD630_s0320 | 154 -- 171 | 18 -- 35 | -9.59579 | | 3.93642 | 8.15991 |
| CD630_08140 | CD630_s0340 | 4 -- 18 | 2 -- 15 | -18.3607 | | 3.30159 | 5.61064 |
| CD630_22550 | CD630_s0340 | 166 -- 188 | 1 -- 21 | -14.4554 | | 3.62379 | 5.61064 |
| CD630_09100 | CD630_s0340 | 14 -- 49 | 40 -- 75 | -14.0533 | | 5.92260 | 5.61064 |
| rolA | CD630_s0340 | 172 -- 186 | 40 -- 53 | -13.3295 | | 4.56362 | 5.61064 |
| CD630_18910 | CD630_s0340 | 27 -- 35 | 5 -- 13 | -12.7153 | | 3.30688 | 5.61064 |
| CD630_23780 | CD630_s0340 | 9 -- 77 | 4 -- 75 | -12.4909 | | 3.91629 | 5.61064 |
| agrB | CD630_s0340 | 124 -- 180 | 1 -- 62 | -12.2455 | | 3.92598 | 5.61064 |
| CD630_04540 | CD630_s0340 | 63 -- 133 | 39 -- 114 | -12.0992 | | 3.33556 | 5.61064 |
| CD630_04850 | CD630_s0340 | 62 -- 171 | 5 -- 108 | -12.0568 | | 4.40788 | 5.61064 |
| CD630_05230 | CD630_s0340 | 148 -- 175 | 144 -- 175 | -11.9953 | | 3.73787 | 5.61064 |
| CD630_07480 | CD630_s0340 | 151 -- 182 | 41 -- 77 | -11.8716 | | 5.18321 | 5.61064 |
| CD630_21090 | CD630_s0340 | 157 -- 169 | 2 -- 14 | -11.8317 | | 4.15803 | 5.61064 |
| CD630_06642 | CD630_s0340 | 128 -- 142 | 135 -- 149 | -11.6144 | | 3.32967 | 5.61064 |
| feoB | CD630_s0340 | 24 -- 44 | 152 -- 171 | -11.5295 | | 4.05364 | 5.61064 |
| CD630_22991 | CD630_s0340 | 193 -- 199 | 7 -- 13 | -11.2516 | | 6.03728 | 5.61064 |
| CD630_08370 | CD630_s0340 | 189 -- 199 | 1 -- 11 | -11.2509 | | 3.65639 | 5.61064 |
| CD630_16150 | CD630_s0340 | 168 -- 192 | 13 -- 41 | -11.1459 | | 4.20614 | 5.61064 |
| CD630_13950 | CD630_s0340 | 137 -- 193 | 36 -- 80 | -11.0344 | | 4.25540 | 5.61064 |
| CD630_26050 | CD630_s0340 | 180 -- 199 | 68 -- 82 | -11.0332 | | 6.99851 | 5.61064 |
| CD630_08950 | CD630_s0340 | 8 -- 26 | 2 -- 15 | -11.0304 | | 3.71905 | 5.61064 |
| CD630_27280 | CD630_s0340 | 107 -- 197 | 3 -- 77 | -11.0191 | | 3.24738 | 5.61064 |
| CD630_14630 | CD630_s0340 | 139 -- 165 | 144 -- 175 | -10.9593 | | 3.34485 | 5.61064 |
| CD630_06100 | CD630_s0340 | 154 -- 192 | 40 -- 78 | -10.8335 | | 4.43918 | 5.61064 |
| CD630_06190 | CD630_s0340 | 130 -- 201 | 7 -- 78 | -10.7493 | | 3.57064 | 5.61064 |
| CD630_24050 | CD630_s0340 | 112 -- 159 | 29 -- 78 | -10.7389 | | 3.70192 | 5.61064 |
| CD630_10080 | CD630_s0340 | 10 -- 39 | 143 -- 166 | -10.6322 | | 3.75625 | 5.61064 |
| CD630_14260 | CD630_s0340 | 74 -- 125 | 25 -- 77 | -10.502 | | 3.22214 | 5.61064 |
| CD630_25040 | CD630_s0340 | 166 -- 200 | 142 -- 172 | -10.4494 | | 3.43898 | 5.61064 |
| CD630_04660 | CD630_s0340 | 145 -- 168 | 145 -- 168 | -10.3419 | | 4.32791 | 5.61064 |
| CD630_28290 | CD630_s0340 | 39 -- 73 | 29 -- 67 | -10.2506 | | 5.54192 | 5.61064 |
| xdhA | CD630_s0340 | 138 -- 165 | 147 -- 175 | -10.2222 | | 4.45147 | 5.61064 |
| CD630_05900 | CD630_s0340 | 100 -- 123 | 145 -- 164 | -10.2204 | | 3.21934 | 5.61064 |
| CD630_17511 | CD630_s0340 | 101 -- 122 | 150 -- 170 | -10.2199 | | 9.12322 | 5.61064 |
| CD630_33600 | CD630_s0340 | 180 -- 192 | 164 -- 176 | -10.1995 | | 3.73196 | 5.61064 |
| CD630_06170 | CD630_s0340 | 1 -- 28 | 55 -- 80 | -10.1975 | | 4.95366 | 5.61064 |
| CD630_32110 | CD630_s0340 | 117 -- 139 | 151 -- 176 | -10.1469 | | 4.45291 | 5.61064 |
| CD630_01700 | CD630_s0340 | 11 -- 24 | 64 -- 77 | -10.131 | | 4.97234 | 5.61064 |
| dpaA | CD630_s0340 | 136 -- 157 | 133 -- 154 | -10.1281 | | 4.65364 | 5.61064 |
| CD630_21240 | CD630_s0340 | 103 -- 125 | 64 -- 87 | -10.0755 | | 5.88879 | 5.61064 |
| CD630_19780 | CD630_s0340 | 99 -- 121 | 151 -- 175 | -10.034 | | 3.64830 | 5.61064 |
| argS | CD630_s0340 | 54 -- 80 | 143 -- 168 | -9.93587 | | 4.96036 | 5.61064 |
| CD630_05822 | CD630_s0340 | 143 -- 174 | 140 -- 171 | -9.87263 | | 4.28113 | 5.61064 |
| spoVFB | CD630_s0340 | 38 -- 70 | 23 -- 59 | -9.86928 | | 4.62758 | 5.61064 |
| CD630_21260 | CD630_s0340 | 121 -- 154 | 41 -- 69 | -9.85893 | | 4.22255 | 5.61064 |
| CD630_15080 | CD630_s0340 | 38 -- 87 | 26 -- 78 | -9.84374 | | 4.03033 | 5.61064 |
| CD630_18930 | CD630_s0340 | 134 -- 165 | 147 -- 177 | -9.80693 | | 5.32127 | 5.61064 |
| CD630_04610 | CD630_s0340 | 147 -- 183 | 40 -- 73 | -9.79959 | | 3.83025 | 5.61064 |
| CD630_13370 | CD630_s0340 | 154 -- 187 | 40 -- 77 | -9.712 | | 4.33145 | 5.61064 |
| cdtR | CD630_s0340 | 3 -- 10 | 4 -- 11 | -9.70961 | | 5.84368 | 5.61064 |
| CD630_17770 | CD630_s0340 | 130 -- 139 | 4 -- 13 | -9.65665 | | 5.06490 | 5.61064 |
| CD630_19840 | CD630_s0340 | 176 -- 189 | 40 -- 54 | -9.64265 | | 3.44724 | 5.61064 |
| CD630_03170 | CD630_s0340 | 127 -- 178 | 53 -- 106 | -9.64141 | | 3.66946 | 5.61064 |
| CD630_08660 | CD630_s0340 | 158 -- 177 | 5 -- 20 | -9.63897 | | 3.86467 | 5.61064 |
| CD630_15431 | CD630_s0340 | 139 -- 199 | 22 -- 82 | -9.62663 | | 5.29780 | 5.61064 |
| CD630_03040 | CD630_s0340 | 180 -- 198 | 147 -- 166 | -9.61625 | | 3.51403 | 5.61064 |
| CD630_31530 | CD630_s0340 | 144 -- 168 | 145 -- 168 | -9.54943 | | 3.49100 | 5.61064 |
| CD630_35260 | CD630_s0340 | 75 -- 84 | 43 -- 52 | -9.54282 | | 5.77277 | 5.61064 |
| CD630_13530 | CD630_s0340 | 12 -- 22 | 2 -- 12 | -9.53586 | | 4.09832 | 5.61064 |
| CD630_08100 | CD630_s0340 | 157 -- 175 | 13 -- 31 | -9.53386 | | 4.39181 | 5.61064 |
| CD630_08130 | CD630_s0340 | 9 -- 40 | 140 -- 175 | -9.52788 | | 4.63831 | 5.61064 |
| CD630_13020 | CD630_s0340 | 100 -- 141 | 55 -- 91 | -9.52524 | | 3.52267 | 5.61064 |
| cspC | CD630_s0370 | 63 -- 93 | 58 -- 87 | -13.8231 | | 5.48848 | 10.21495 |
| oppB | CD630_s0370 | 88 -- 104 | 54 -- 70 | -12.5514 | | 2.98328 | 10.21495 |
| cspD | CD630_s0370 | 63 -- 93 | 58 -- 87 | -12.445 | | 3.23587 | 10.21495 |
| CD630_08460 | CD630_s0370 | 127 -- 161 | 67 -- 106 | -11.5972 | | 4.15083 | 10.21495 |
| xdhA | CD630_s0370 | 11 -- 27 | 68 -- 86 | -11.5971 | | 4.45147 | 10.21495 |
| CD630_26920 | CD630_s0370 | 141 -- 156 | 38 -- 55 | -11.4066 | | 4.58060 | 10.21495 |
| appC | CD630_s0370 | 162 -- 169 | 31 -- 38 | -11.1138 | | 4.50438 | 10.21495 |
| CD630_13530 | CD630_s0370 | 176 -- 183 | 31 -- 38 | -10.9543 | | 4.09832 | 10.21495 |
| CD630_05480 | CD630_s0370 | 9 -- 18 | 30 -- 39 | -10.9424 | | 4.27854 | 10.21495 |
| CD630_25620 | CD630_s0370 | 187 -- 198 | 42 -- 53 | -10.9186 | | 3.02685 | 10.21495 |
| CD630_27040 | CD630_s0370 | 176 -- 195 | 3 -- 22 | -10.9065 | | 3.51812 | 10.21495 |
| CD630_31480 | CD630_s0370 | 165 -- 191 | 77 -- 102 | -10.2852 | | 4.88897 | 10.21495 |
| CD630_20960 | CD630_s0370 | 189 -- 199 | 30 -- 40 | -10.1824 | | 5.65940 | 10.21495 |
| CD630_36250 | CD630_s0370 | 40 -- 54 | 97 -- 111 | -10.1384 | | 4.13224 | 10.21495 |
| CD630_09100 | CD630_s0370 | 8 -- 50 | 4 -- 49 | -10.0093 | | 5.92260 | 10.21495 |
| CD630_14050 | CD630_s0370 | 165 -- 174 | 32 -- 41 | -9.8516 | | 3.30894 | 10.21495 |
| CD630_06060 | CD630_s0370 | 152 -- 184 | 43 -- 79 | -9.75871 | | 4.94050 | 10.21495 |
| CD630_05940 | CD630_s0370 | 189 -- 199 | 31 -- 41 | -9.68102 | | 4.18595 | 10.21495 |
| CD630_16160 | CD630_s0390 | 37 -- 65 | 8 -- 42 | -16.1324 | | 6.11801 | 4.73397 |
| CD630_19030 | CD630_s0390 | 99 -- 146 | 52 -- 93 | -15.8518 | | 3.34240 | 4.73397 |
| CD630_03220 | CD630_s0390 | 99 -- 109 | 135 -- 145 | -14.9677 | | 3.36746 | 4.73397 |
| CD630_21490 | CD630_s0390 | 173 -- 199 | 132 -- 161 | -14.5664 | | 3.37462 | 4.73397 |
| CD630_21540 | CD630_s0390 | 81 -- 128 | 41 -- 89 | -14.3238 | | 4.33090 | 4.73397 |
| CD630_36180 | CD630_s0390 | 167 -- 198 | 10 -- 41 | -14.2971 | | 4.83428 | 4.73397 |
| CD630_21720 | CD630_s0390 | 163 -- 193 | 56 -- 88 | -14.1994 | | 4.13030 | 4.73397 |
| CD630_27280 | CD630_s0390 | 150 -- 196 | 183 -- 230 | -13.9845 | | 3.24738 | 4.73397 |
| CD630_25300 | CD630_s0390 | 134 -- 163 | 59 -- 94 | -13.966 | | 4.49333 | 4.73397 |
| CD630_10220 | CD630_s0390 | 60 -- 129 | 33 -- 100 | -13.957 | | 4.08233 | 4.73397 |
| ssuC | CD630_s0390 | 65 -- 77 | 134 -- 145 | -13.8524 | | 3.45491 | 4.73397 |
| CD630_06970 | CD630_s0390 | 121 -- 143 | 124 -- 146 | -12.9954 | | 3.82312 | 4.73397 |
| CD630_26630 | CD630_s0390 | 176 -- 200 | 69 -- 93 | -12.8256 | | 4.61557 | 4.73397 |
| CD630_25200 | CD630_s0390 | 148 -- 200 | 41 -- 92 | -12.5856 | | 3.65158 | 4.73397 |
| CD630_25310 | CD630_s0390 | 21 -- 43 | 125 -- 143 | -12.2244 | | 3.99861 | 4.73397 |
| CD630_23450 | CD630_s0390 | 46 -- 127 | 4 -- 63 | -12.1003 | | 3.27218 | 4.73397 |
| CD630_28150 | CD630_s0390 | 36 -- 71 | 58 -- 93 | -12.0538 | | 4.18447 | 4.73397 |
| gcp | CD630_s0390 | 124 -- 157 | 3 -- 46 | -11.9915 | | 4.92798 | 4.73397 |
| dpaA | CD630_s0390 | 134 -- 163 | 65 -- 97 | -11.9611 | | 4.65364 | 4.73397 |
| CD630_18940 | CD630_s0390 | 149 -- 188 | 52 -- 92 | -11.8572 | | 4.09988 | 4.73397 |
| CD630_36190 | CD630_s0390 | 149 -- 177 | 124 -- 151 | -11.7764 | | 4.63449 | 4.73397 |
| CD630_28181 | CD630_s0390 | 149 -- 168 | 131 -- 150 | -11.4992 | | 3.91299 | 4.73397 |
| CD630_06170 | CD630_s0390 | 135 -- 154 | 1 -- 21 | -11.4857 | | 4.95366 | 4.73397 |
| CD630_05830 | CD630_s0390 | 148 -- 169 | 146 -- 168 | -11.4757 | | 3.35360 | 4.73397 |
| CD630_15270 | CD630_s0390 | 86 -- 131 | 43 -- 88 | -11.3686 | | 4.66015 | 4.73397 |
| CD630_07470 | CD630_s0390 | 155 -- 175 | 70 -- 88 | -11.3565 | | 3.37143 | 4.73397 |
| CD630_02420 | CD630_s0390 | 1 -- 53 | 53 -- 98 | -11.3062 | | 3.71048 | 4.73397 |
| CD630_19850 | CD630_s0390 | 93 -- 124 | 56 -- 90 | -11.2265 | | 6.88486 | 4.73397 |
| CD630_02790 | CD630_s0390 | 76 -- 96 | 137 -- 156 | -11.2167 | | 4.77661 | 4.73397 |
| cdtR | CD630_s0390 | 57 -- 77 | 213 -- 233 | -11.1856 | | 5.84368 | 4.73397 |
| CD630_07870 | CD630_s0390 | 176 -- 200 | 6 -- 27 | -11.0771 | | 3.22829 | 4.73397 |
| CD630_31490 | CD630_s0390 | 152 -- 181 | 30 -- 57 | -11.0451 | | 4.08593 | 4.73397 |
| CD630_16330 | CD630_s0390 | 122 -- 156 | 51 -- 87 | -11.0146 | | 4.11180 | 4.73397 |
| xylR | CD630_s0390 | 118 -- 155 | 10 -- 50 | -11.0009 | | 3.95671 | 4.73397 |
| CD630_08370 | CD630_s0390 | 142 -- 160 | 65 -- 86 | -10.975 | | 3.65639 | 4.73397 |
| CD630_03650 | CD630_s0390 | 111 -- 129 | 34 -- 55 | -10.9551 | | 3.93697 | 4.73397 |
| CD630_05900 | CD630_s0390 | 38 -- 77 | 54 -- 97 | -10.939 | | 3.21934 | 4.73397 |
| CD630_27240 | CD630_s0390 | 131 -- 175 | 146 -- 195 | -10.9387 | | 3.52462 | 4.73397 |
| CD630_14130 | CD630_s0390 | 138 -- 189 | 114 -- 157 | -10.9338 | | 4.57970 | 4.73397 |
| CD630_30540 | CD630_s0390 | 173 -- 194 | 58 -- 79 | -10.9335 | | 3.63804 | 4.73397 |
| CD630_26250 | CD630_s0390 | 125 -- 153 | 210 -- 238 | -10.8495 | | 5.03379 | 4.73397 |
| CD630_30420 | CD630_s0390 | 138 -- 192 | 35 -- 88 | -10.8342 | | 5.27539 | 4.73397 |
| cheW | CD630_s0390 | 1 -- 14 | 141 -- 154 | -10.7439 | | 3.72371 | 4.73397 |
| CD630_09100 | CD630_s0390 | 3 -- 64 | 4 -- 67 | -10.7323 | | 5.92260 | 4.73397 |
| CD630_32700 | CD630_s0390 | 128 -- 163 | 135 -- 177 | -10.7127 | | 4.74276 | 4.73397 |
| CD630_05230 | CD630_s0390 | 127 -- 144 | 23 -- 40 | -10.6877 | | 3.73787 | 4.73397 |
| CD630_18680 | CD630_s0390 | 144 -- 159 | 185 -- 200 | -10.6638 | | 3.04315 | 4.73397 |
| CD630_23930 | CD630_s0390 | 160 -- 200 | 41 -- 86 | -10.6125 | | 3.05650 | 4.73397 |
| CD630_14630 | CD630_s0390 | 129 -- 157 | 66 -- 90 | -10.5285 | | 3.34485 | 4.73397 |
| CD630_15111 | CD630_s0390 | 97 -- 118 | 63 -- 83 | -10.5165 | | 4.18664 | 4.73397 |
| CD630_17970 | CD630_s0390 | 144 -- 184 | 54 -- 91 | -10.5067 | | 3.68802 | 4.73397 |
| ribD | CD630_s0390 | 40 -- 66 | 209 -- 239 | -10.4154 | | 6.65922 | 4.73397 |
| CD630_20171 | CD630_s0390 | 81 -- 124 | 211 -- 254 | -10.3223 | | 6.87491 | 4.73397 |
| CD630_11220 | CD630_s0390 | 176 -- 187 | 79 -- 89 | -10.2493 | | 3.52327 | 4.73397 |
| etfA | CD630_s0390 | 141 -- 168 | 126 -- 153 | -10.2428 | | 3.41204 | 4.73397 |
| dapB2 | CD630_s0390 | 182 -- 196 | 223 -- 239 | -10.2253 | | 3.27218 | 4.73397 |
| CD630_08940 | CD630_s0390 | 68 -- 88 | 70 -- 89 | -10.2155 | | 3.72225 | 4.73397 |
| pilC | CD630_s0390 | 130 -- 138 | 240 -- 248 | -10.165 | | 3.54230 | 4.73397 |
| CD630_20280 | CD630_s0390 | 109 -- 145 | 136 -- 172 | -10.159 | | 3.46343 | 4.73397 |
| ogt1 | CD630_s0390 | 128 -- 144 | 32 -- 48 | -10.1558 | | 3.16772 | 4.73397 |
| CD630_23820 | CD630_s0390 | 27 -- 43 | 58 -- 74 | -10.1417 | | 3.18419 | 4.73397 |
| CD630_18090 | CD630_s0390 | 18 -- 39 | 212 -- 235 | -10.1325 | | 4.23961 | 4.73397 |
| bioB | CD630_s0390 | 112 -- 127 | 13 -- 28 | -10.0896 | | 3.59022 | 4.73397 |
| CD630_07960 | CD630_s0390 | 179 -- 196 | 130 -- 148 | -10.0679 | | 3.22537 | 4.73397 |
| CD630_32890 | CD630_s0390 | 103 -- 135 | 4 -- 39 | -10.0646 | | 3.41276 | 4.73397 |
| CD630_10080 | CD630_s0390 | 153 -- 179 | 17 -- 44 | -10.058 | | 3.75625 | 4.73397 |
| cprK | CD630_s0390 | 83 -- 112 | 223 -- 250 | -10.0402 | | 3.96216 | 4.73397 |
| CD630_14700 | CD630_s0390 | 181 -- 197 | 70 -- 86 | -10.0241 | | 3.98234 | 4.73397 |
| thiH | CD630_s0390 | 46 -- 73 | 223 -- 246 | -10.0156 | | 3.58230 | 4.73397 |
| tcdC | CD630_s0390 | 149 -- 157 | 34 -- 42 | -10.0148 | | 3.76101 | 4.73397 |
| CD630_06080 | CD630_s0390 | 178 -- 201 | 230 -- 252 | -9.85869 | | 3.82315 | 4.73397 |
| adhE | CD630_s0390 | 162 -- 176 | 83 -- 99 | -9.8531 | | 3.06485 | 4.73397 |
| bglA | CD630_s0390 | 169 -- 195 | 57 -- 86 | -9.8515 | | 4.86402 | 4.73397 |
| CD630_13620 | CD630_s0390 | 177 -- 200 | 130 -- 150 | -9.82272 | | 4.77627 | 4.73397 |
| flhB | CD630_s0390 | 33 -- 53 | 214 -- 233 | -9.80947 | | 3.55524 | 4.73397 |
| CD630_05030 | CD630_s0390 | 32 -- 39 | 217 -- 224 | -9.76826 | | 4.03823 | 4.73397 |
| CD630_22950 | CD630_s0390 | 122 -- 141 | 81 -- 99 | -9.75344 | | 3.96883 | 4.73397 |
| CD630_25320 | CD630_s0390 | 121 -- 162 | 19 -- 59 | -9.73633 | | 4.29415 | 4.73397 |
| CD630_04880 | CD630_s0390 | 123 -- 161 | 45 -- 91 | -9.73271 | | 4.62291 | 4.73397 |
| pilD | CD630_s0390 | 128 -- 162 | 120 -- 155 | -9.70585 | | 4.54123 | 4.73397 |
| CD630_08360 | CD630_s0390 | 46 -- 60 | 66 -- 81 | -9.67692 | | 3.19339 | 4.73397 |
| CD630_06110 | CD630_s0390 | 178 -- 191 | 217 -- 230 | -9.67026 | | 5.61380 | 4.73397 |
| srlEb | CD630_s0390 | 121 -- 153 | 57 -- 85 | -9.64067 | | 3.50140 | 4.73397 |
| CD630_17511 | CD630_s0390 | 89 -- 112 | 217 -- 240 | -9.61606 | | 9.12322 | 4.73397 |
| CD630_32720 | CD630_s0390 | 89 -- 99 | 230 -- 240 | -9.60575 | | 4.74276 | 4.73397 |
| CD630_06060 | CD630_s0390 | 71 -- 92 | 57 -- 77 | -9.58856 | | 4.94050 | 4.73397 |
| CD630_24050 | CD630_s0390 | 129 -- 159 | 9 -- 43 | -9.55873 | | 3.70192 | 4.73397 |
| ung | CD630_s0390 | 140 -- 184 | 2 -- 41 | -9.53721 | | 3.30908 | 4.73397 |
| CD630_26750 | CD630_s0390 | 115 -- 162 | 44 -- 89 | -9.52931 | | 3.60610 | 4.73397 |
| CD630_12650 | CD630_s0390 | 140 -- 177 | 64 -- 97 | -9.52536 | | 6.31827 | 4.73397 |
| CD630_32070 | CD630_s0390 | 147 -- 159 | 146 -- 158 | -9.52382 | | 3.51964 | 4.73397 |
| CD630_27040 | CD630_s0390 | 185 -- 199 | 185 -- 199 | -9.51841 | | 3.51812 | 4.73397 |
| terD | CD630_s0390 | 121 -- 138 | 230 -- 246 | -9.51642 | | 3.33148 | 4.73397 |
| CD630_01640 | CD630_s0390 | 144 -- 166 | 15 -- 32 | -9.5111 | | 4.55381 | 4.73397 |
| CD630_21030 | CD630_s0390 | 179 -- 195 | 13 -- 29 | -9.50077 | | 3.08733 | 4.73397 |
| CD630_29250 | CD630_s0390 | 87 -- 116 | 58 -- 85 | -9.50026 | | 4.61901 | 4.73397 |
| CD630_09410 | CD630_s0390 | 87 -- 116 | 58 -- 85 | -9.50026 | | 4.75651 | 4.73397 |
| CD630_04850 | CD630_s0400 | 137 -- 198 | 27 -- 76 | -16.4074 | | 4.40788 | 4.51086 |
| CD630_16160 | CD630_s0400 | 42 -- 66 | 53 -- 77 | -16.0881 | | 6.11801 | 4.51086 |
| CD630_13580 | CD630_s0400 | 134 -- 161 | 53 -- 82 | -15.1603 | | 3.80715 | 4.51086 |
| spaK | CD630_s0400 | 167 -- 200 | 131 -- 174 | -15.0371 | | 3.04530 | 4.51086 |
| CD630_08100 | CD630_s0400 | 14 -- 36 | 22 -- 43 | -15.0311 | | 4.39181 | 4.51086 |
| CD630_18230 | CD630_s0400 | 113 -- 147 | 13 -- 45 | -14.9621 | | 3.00184 | 4.51086 |
| CD630_08130 | CD630_s0400 | 117 -- 154 | 4 -- 38 | -14.6287 | | 4.63831 | 4.51086 |
| CD630_21090 | CD630_s0400 | 139 -- 199 | 5 -- 58 | -14.2459 | | 4.15803 | 4.51086 |
| CD630_06150 | CD630_s0400 | 83 -- 160 | 37 -- 134 | -14.0636 | | 5.51744 | 4.51086 |
| cwp29 | CD630_s0400 | 88 -- 184 | 22 -- 129 | -14.0405 | | 4.40865 | 4.51086 |
| CD630_08080 | CD630_s0400 | 28 -- 69 | 6 -- 48 | -13.9851 | | 4.87819 | 4.51086 |
| CD630_14260 | CD630_s0400 | 35 -- 102 | 23 -- 75 | -13.7922 | | 3.22214 | 4.51086 |
| CD630_14620 | CD630_s0400 | 38 -- 96 | 12 -- 76 | -13.7723 | | 4.43651 | 4.51086 |
| CD630_08140 | CD630_s0400 | 5 -- 51 | 6 -- 48 | -13.6598 | | 3.30159 | 4.51086 |
| cwp26 | CD630_s0400 | 143 -- 189 | 23 -- 67 | -13.1932 | | 4.44377 | 4.51086 |
| CD630_06100 | CD630_s0400 | 106 -- 144 | 7 -- 47 | -13.1803 | | 4.43918 | 4.51086 |
| CD630_10080 | CD630_s0400 | 9 -- 21 | 63 -- 75 | -13.1283 | | 3.75625 | 4.51086 |
| CD630_23450 | CD630_s0400 | 90 -- 146 | 20 -- 71 | -12.9323 | | 3.27218 | 4.51086 |
| CD630_30190 | CD630_s0400 | 35 -- 72 | 20 -- 58 | -12.8611 | | 3.27394 | 4.51086 |
| CD630_22550 | CD630_s0400 | 25 -- 59 | 113 -- 148 | -12.79 | | 3.62379 | 4.51086 |
| CD630_16940 | CD630_s0400 | 85 -- 161 | 54 -- 129 | -12.2935 | | 4.14665 | 4.51086 |
| CD630_15060 | CD630_s0400 | 170 -- 196 | 20 -- 49 | -12.2854 | | 3.69396 | 4.51086 |
| CD630_17900 | CD630_s0400 | 55 -- 133 | 8 -- 72 | -12.2656 | | 3.52167 | 4.51086 |
| CD630_13530 | CD630_s0400 | 12 -- 44 | 18 -- 49 | -12.2523 | | 4.09832 | 4.51086 |
| CD630_30420 | CD630_s0400 | 140 -- 174 | 50 -- 87 | -12.2407 | | 5.27539 | 4.51086 |
| appC | CD630_s0400 | 34 -- 166 | 6 -- 119 | -12.194 | | 4.50438 | 4.51086 |
| CD630_06560 | CD630_s0400 | 113 -- 153 | 3 -- 43 | -12.1217 | | 4.02532 | 4.51086 |
| CD630_07770 | CD630_s0400 | 138 -- 154 | 20 -- 38 | -12.0302 | | 3.31158 | 4.51086 |
| CD630_21110 | CD630_s0400 | 121 -- 166 | 6 -- 56 | -11.9763 | | 3.89630 | 4.51086 |
| CD630_21260 | CD630_s0400 | 131 -- 140 | 2 -- 11 | -11.9581 | | 4.22255 | 4.51086 |
| CD630_02720 | CD630_s0400 | 163 -- 177 | 27 -- 40 | -11.9094 | | 3.38938 | 4.51086 |
| CD630_26691 | CD630_s0400 | 5 -- 78 | 63 -- 141 | -11.8388 | | 3.96230 | 4.51086 |
| CD630_30100 | CD630_s0400 | 150 -- 185 | 40 -- 78 | -11.8328 | | 4.13234 | 4.51086 |

| | | | | | |
|---|---|---|---|---|---|
| CD630_19850 | CD630_s0400 | 78 – 97 | 8 – 28 | -11.6912 | 6.88486 | 4.51086 |
| CD630_21770 | CD630_s0400 | 110 – 153 | 35 – 77 | -11.6523 | 4.02587 | 4.51086 |
| CD630_17511 | CD630_s0400 | 1 – 60 | 1 – 67 | -11.5282 | 9.12322 | 4.51086 |
| CD630_15460 | CD630_s0400 | 94 – 107 | 178 – 191 | -11.5213 | 6.86296 | 4.51086 |
| CD630_22991 | CD630_s0400 | 189 – 197 | 163 – 171 | -11.475 | 6.03728 | 4.51086 |
| CD630_01640 | CD630_s0400 | 186 – 199 | 45 – 58 | -11.3923 | 4.55381 | 4.51086 |
| CD630_27280 | CD630_s0400 | 165 – 197 | 38 – 76 | -11.3821 | 3.24738 | 4.51086 |
| cdtR | CD630_s0400 | 2 – 16 | 31 – 46 | -11.2844 | 5.84368 | 4.51086 |
| CD630_15270 | CD630_s0400 | 142 – 191 | 8 – 55 | -11.1057 | 4.66015 | 4.51086 |
| dltB | CD630_s0400 | 16 – 86 | 8 – 87 | -10.936 | 3.81485 | 4.51086 |
| CD630_28290 | CD630_s0400 | 20 – 93 | 13 – 75 | -10.8199 | 5.54192 | 4.51086 |
| CD630_16330 | CD630_s0400 | 166 – 183 | 163 – 183 | -10.8038 | 4.11180 | 4.51086 |
| dnaD | CD630_s0400 | 4 – 38 | 44 – 77 | -10.7874 | 3.56078 | 4.51086 |
| CD630_19401 | CD630_s0400 | 31 – 127 | 6 – 83 | -10.7812 | 4.37084 | 4.51086 |
| dnaC | CD630_s0400 | 124 – 179 | 20 – 68 | -10.7493 | 6.30796 | 4.51086 |
| CD630_27110 | CD630_s0400 | 112 – 186 | 20 – 73 | -10.7237 | 3.78231 | 4.51086 |
| xdhA | CD630_s0400 | 111 – 148 | 178 – 205 | -10.6633 | 4.45147 | 4.51086 |
| CD630_05900 | CD630_s0400 | 7 – 47 | 6 – 56 | -10.6611 | 3.21934 | 4.51086 |
| CD630_03170 | CD630_s0400 | 148 – 178 | 31 – 65 | -10.6496 | 3.66946 | 4.51086 |
| srlEb | CD630_s0400 | 187 – 197 | 2 – 12 | -10.5986 | 3.50140 | 4.51086 |
| CD630_01700 | CD630_s0400 | 17 – 39 | 60 – 78 | -10.5807 | 4.97234 | 4.51086 |
| CD630_25050 | CD630_s0400 | 120 – 171 | 21 – 73 | -10.478 | 4.33624 | 4.51086 |
| CD630_28870 | CD630_s0400 | 73 – 86 | 70 – 83 | -10.4452 | 3.41991 | 4.51086 |
| CD630_13960 | CD630_s0400 | 190 – 201 | 20 – 31 | -10.4317 | 3.30375 | 4.51086 |
| CD630_16530 | CD630_s0400 | 107 – 154 | 34 – 74 | -10.4088 | 4.37658 | 4.51086 |
| CD630_07470 | CD630_s0400 | 25 – 60 | 4 – 33 | -10.393 | 3.37143 | 4.51086 |
| CD630_24760 | CD630_s0400 | 139 – 166 | 4 – 30 | -10.3032 | 5.19485 | 4.51086 |
| CD630_22510 | CD630_s0400 | 119 – 157 | 43 – 78 | -10.2747 | 11.42763 | 4.51086 |
| CD630_09040 | CD630_s0400 | 76 – 127 | 24 – 69 | -10.26 | 4.11851 | 4.51086 |
| CD630_03650 | CD630_s0400 | 31 – 80 | 2 – 48 | -10.233 | 3.93697 | 4.51086 |
| CD630_13880 | CD630_s0400 | 154 – 162 | 38 – 46 | -10.1982 | 3.52735 | 4.51086 |
| CD630_31461 | CD630_s0400 | 132 – 142 | 44 – 54 | -10.1178 | 4.41476 | 4.51086 |
| CD630_33610 | CD630_s0400 | 172 – 192 | 137 – 156 | -10.1162 | 3.79322 | 4.51086 |
| CD630_21540 | CD630_s0400 | 35 – 59 | 24 – 47 | -10.0655 | 4.33090 | 4.51086 |
| CD630_29270 | CD630_s0400 | 58 – 92 | 14 – 58 | -10.0471 | 3.70174 | 4.51086 |
| CD630_21070 | CD630_s0400 | 58 – 92 | 14 – 48 | -10.0445 | 5.35019 | 4.51086 |
| CD630_29840 | CD630_s0400 | 159 – 195 | 34 – 68 | -10.0273 | 4.48825 | 4.51086 |
| CD630_05290 | CD630_s0400 | 2 – 111 | 53 – 173 | -10.0254 | 3.97604 | 4.51086 |
| CD630_23861 | CD630_s0400 | 10 – 31 | 10 – 30 | -10.022 | 5.83068 | 4.51086 |
| CD630_18781 | CD630_s0400 | 138 – 174 | 5 – 54 | -9.99607 | 3.77660 | 4.51086 |
| cls | CD630_s0400 | 37 – 57 | 167 – 187 | -9.99364 | 4.46177 | 4.51086 |
| CD630_36690 | CD630_s0400 | 146 – 196 | 5 – 49 | -9.96982 | 3.53761 | 4.51086 |
| CD630_28150 | CD630_s0400 | 9 – 21 | 59 – 71 | -9.9323 | 4.18447 | 4.51086 |
| opuCC | CD630_s0400 | 115 – 198 | 2 – 78 | -9.9128 | 4.07035 | 4.51086 |
| CD630_13790 | CD630_s0400 | 124 – 158 | 35 – 66 | -9.89629 | 5.17374 | 4.51086 |
| CD630_25310 | CD630_s0400 | 165 – 180 | 56 – 72 | -9.87859 | 3.99861 | 4.51086 |
| CD630_14182 | CD630_s0400 | 189 – 197 | 163 – 171 | -9.86522 | 4.82360 | 4.51086 |
| CD630_33420 | CD630_s0400 | 13 – 50 | 147 – 191 | -9.86134 | 3.22095 | 4.51086 |
| CD630_19904 | CD630_s0400 | 12 – 39 | 20 – 47 | -9.85216 | 5.96526 | 4.51086 |
| serS1 | CD630_s0400 | 153 – 170 | 184 – 204 | -9.79036 | 3.45858 | 4.51086 |
| CD630_18090 | CD630_s0400 | 185 – 194 | 42 – 51 | -9.78516 | 4.23961 | 4.51086 |
| CD630_04320 | CD630_s0400 | 170 – 185 | 167 – 183 | -9.76566 | 3.68342 | 4.51086 |
| CD630_05830 | CD630_s0400 | 108 – 160 | 28 – 74 | -9.74903 | 3.35360 | 4.51086 |
| CD630_04880 | CD630_s0400 | 115 – 181 | 5 – 64 | -9.74834 | 4.62291 | 4.51086 |
| CD630_24590 | CD630_s0400 | 173 – 196 | 22 – 43 | -9.73805 | 3.70401 | 4.51086 |
| CD630_27040 | CD630_s0400 | 112 – 160 | 29 – 72 | -9.71348 | 3.51812 | 4.51086 |
| CD630_01570 | CD630_s0400 | 2 – 56 | 21 – 82 | -9.70119 | 6.08699 | 4.51086 |
| CD630_33682 | CD630_s0400 | 22 – 45 | 168 – 189 | -9.69427 | 5.77720 | 4.51086 |
| CD630_19680 | CD630_s0400 | 37 – 96 | 121 – 186 | -9.67122 | 3.58080 | 4.51086 |
| CD630_01690 | CD630_s0400 | 168 – 182 | 105 – 119 | -9.65289 | 4.97234 | 4.51086 |
| CD630_06920 | CD630_s0400 | 141 – 185 | 40 – 87 | -9.63381 | 3.13260 | 4.51086 |
| kdgT1 | CD630_s0400 | 35 – 63 | 6 – 33 | -9.62469 | 4.49207 | 4.51086 |
| CD630_29760 | CD630_s0400 | 157 – 185 | 121 – 150 | -9.61557 | 3.51074 | 4.51086 |
| CD630_18120 | CD630_s0400 | 179 – 199 | 182 – 201 | -9.61372 | 3.99713 | 4.51086 |
| CD630_16550 | CD630_s0400 | 16 – 28 | 165 – 176 | -9.59666 | 5.61064 | 4.51086 |
| CD630_36180 | CD630_s0400 | 12 – 33 | 64 – 87 | -9.58973 | 4.83428 | 4.51086 |
| CD630_05590 | CD630_s0400 | 192 – 201 | 38 – 47 | -9.58644 | 3.39670 | 4.51086 |
| CD630_06690 | CD630_s0400 | 161 – 191 | 47 – 77 | -9.57773 | 5.82709 | 4.51086 |
| CD630_10000 | CD630_s0400 | 146 – 176 | 4 – 32 | -9.55962 | 3.86040 | 4.51086 |
| feoB | CD630_s0400 | 119 – 149 | 19 – 49 | -9.55145 | 4.05364 | 4.51086 |
| CD630_18780 | CD630_s0400 | 184 – 197 | 13 – 27 | -9.5127 | 4.71613 | 4.51086 |
| CD630_03160 | CD630_s0400 | 49 – 116 | 7 – 73 | -9.50252 | 3.66946 | 4.51086 |
| CD630_05030 | CD630_s0480 | 32 – 67 | 151 – 191 | -14.3806 | 4.03823 | 4.87999 |
| CD630_27220 | CD630_s0480 | 131 – 186 | 24 – 83 | -14.1765 | 4.18786 | 4.87999 |
| bglA | CD630_s0480 | 91 – 127 | 43 – 85 | -14.0264 | 4.50315 | 4.87999 |
| CD630_27280 | CD630_s0480 | 45 – 68 | 68 – 91 | -13.7738 | 3.24738 | 4.87999 |
| CD630_31690 | CD630_s0480 | 139 – 196 | 153 – 204 | -13.5071 | 3.71193 | 4.87999 |
| CD630_10220 | CD630_s0480 | 92 – 141 | 39 – 85 | -13.4931 | 4.08233 | 4.87999 |
| CD630_05940 | CD630_s0480 | 64 – 104 | 43 – 81 | -13.1338 | 4.18595 | 4.87999 |
| CD630_15360 | CD630_s0480 | 154 – 187 | 66 – 99 | -13.0406 | 4.47208 | 4.87999 |
| CD630_20980 | CD630_s0480 | 89 – 110 | 41 – 61 | -12.8981 | 8.12847 | 4.87999 |
| nusB | CD630_s0480 | 36 – 66 | 17 – 48 | -12.6333 | 4.92798 | 4.87999 |
| CD630_35780 | CD630_s0480 | 57 – 68 | 81 – 92 | -12.5879 | 3.50107 | 4.87999 |
| serS1 | CD630_s0480 | 156 – 168 | 208 – 220 | -12.4892 | 3.45858 | 4.87999 |
| dltB | CD630_s0480 | 3 – 77 | 14 – 71 | -12.0947 | 3.81485 | 4.87999 |
| CD630_04880 | CD630_s0480 | 84 – 132 | 188 – 220 | -11.7629 | 4.62291 | 4.87999 |
| CD630_06060 | CD630_s0480 | 123 – 164 | 42 – 86 | -11.6972 | 4.94050 | 4.87999 |
| CD630_19401 | CD630_s0480 | 24 – 127 | 5 – 99 | -11.6796 | 4.37084 | 4.87999 |
| CD630_28181 | CD630_s0480 | 151 – 190 | 19 – 64 | -11.6011 | 3.91299 | 4.87999 |
| CD630_27230 | CD630_s0480 | 133 – 142 | 79 – 88 | -11.5736 | 3.52462 | 4.87999 |
| CD630_18650 | CD630_s0480 | 138 – 192 | 43 – 91 | -11.3825 | 3.75892 | 4.87999 |
| CD630_05150 | CD630_s0480 | 82 – 103 | 201 – 221 | -11.3022 | 3.69875 | 4.87999 |
| chrA | CD630_s0480 | 87 – 154 | 39 – 96 | -11.2863 | 5.27255 | 4.87999 |
| CD630_14340 | CD630_s0480 | 71 – 94 | 61 – 85 | -11.2587 | 5.10868 | 4.87999 |
| CD630_21490 | CD630_s0480 | 33 – 82 | 41 – 89 | -11.1165 | 3.37462 | 4.87999 |
| CD630_21540 | CD630_s0480 | 77 – 169 | 13 – 95 | -11.023 | 4.33090 | 4.87999 |
| CD630_21240 | CD630_s0480 | 88 – 112 | 46 – 61 | -11.015 | 5.88879 | 4.87999 |
| tcdC | CD630_s0480 | 51 – 135 | 15 – 81 | -11.0073 | 3.76101 | 4.87999 |
| CD630_22950 | CD630_s0480 | 4 – 51 | 85 – 131 | -10.9301 | 3.96883 | 4.87999 |
| CD630_24920 | CD630_s0480 | 1 – 25 | 105 – 136 | -10.8015 | 3.89455 | 4.87999 |
| CD630_22030 | CD630_s0480 | 3 – 37 | 185 – 212 | -10.7514 | 3.29934 | 4.87999 |
| CD630_04770 | CD630_s0480 | 32 – 68 | 9 – 44 | -10.6853 | 3.58478 | 4.87999 |
| ssuC | CD630_s0480 | 64 – 98 | 76 – 110 | -10.6495 | 3.45491 | 4.87999 |
| cobQ | CD630_s0480 | 86 – 103 | 201 – 219 | -10.6136 | 3.13524 | 4.87999 |
| cbpA | CD630_s0480 | 84 – 96 | 205 – 217 | -10.5784 | 3.03040 | 4.87999 |
| CD630_04610 | CD630_s0480 | 39 – 51 | 161 – 173 | -10.5386 | 3.83025 | 4.87999 |
| CD630_19680 | CD630_s0480 | 63 – 95 | 55 – 86 | -10.4833 | 3.58080 | 4.87999 |
| CD630_14990 | CD630_s0480 | 1 – 16 | 141 – 158 | -10.4781 | 3.78619 | 4.87999 |
| CD630_15160 | CD630_s0480 | 4 – 37 | 156 – 184 | -10.476 | 4.32318 | 4.87999 |
| srlD | CD630_s0480 | 94 – 146 | 81 – 141 | -10.4465 | 4.19843 | 4.87999 |
| CD630_05300 | CD630_s0480 | 3 – 31 | 52 – 82 | -10.2626 | 4.56885 | 4.87999 |
| CD630_31210 | CD630_s0480 | 82 – 97 | 51 – 66 | -10.2408 | 4.17693 | 4.87999 |
| CD630_16940 | CD630_s0480 | 85 – 185 | 138 – 217 | -10.1701 | 4.14665 | 4.87999 |
| CD630_31220 | CD630_s0480 | 124 – 143 | 128 – 150 | -10.1664 | 3.73779 | 4.87999 |
| CD630_26691 | CD630_s0480 | 4 – 35 | 44 – 74 | -10.1381 | 3.96230 | 4.87999 |
| bglA | CD630_s0480 | 46 – 95 | 105 – 155 | -10.1181 | 4.86402 | 4.87999 |
| CD630_32880 | CD630_s0480 | 184 – 193 | 79 – 88 | -10.1169 | 3.41276 | 4.87999 |
| CD630_18950 | CD630_s0480 | 180 – 188 | 41 – 49 | -10.1079 | 3.40610 | 4.87999 |
| CD630_06170 | CD630_s0480 | 8 – 28 | 101 – 122 | -10.0811 | 4.95366 | 4.87999 |
| CD630_27630 | CD630_s0480 | 152 – 176 | 22 – 47 | -10.0693 | 3.54199 | 4.87999 |
| opuCA | CD630_s0480 | 163 – 195 | 36 – 68 | -10.0423 | 4.07035 | 4.87999 |
| CD630_06200 | CD630_s0480 | 168 – 197 | 143 – 172 | -10.0256 | 3.57064 | 4.87999 |
| CD630_27610 | CD630_s0480 | 10 – 27 | 41 – 59 | -10.0091 | 3.96465 | 4.87999 |
| flhB | CD630_s0480 | 159 – 167 | 41 – 49 | -9.92907 | 3.55524 | 4.87999 |
| CD630_09840 | CD630_s0480 | 20 – 31 | 41 – 52 | -9.89908 | 4.50009 | 4.87999 |
| CD630_08070 | CD630_s0480 | 73 – 89 | 107 – 123 | -9.74398 | 4.87819 | 4.87999 |
| CD630_02830 | CD630_s0480 | 161 – 181 | 148 – 169 | -9.68126 | 3.57106 | 4.87999 |
| CD630_28560 | CD630_s0480 | 83 – 89 | 43 – 49 | -9.66578 | 3.96803 | 4.87999 |
| spaK | CD630_s0480 | 7 – 24 | 184 – 201 | -9.60689 | 3.04530 | 4.87999 |
| oppB | CD630_s0480 | 89 – 105 | 199 – 215 | -9.58221 | 2.98328 | 4.87999 |
| CD630_19800 | CD630_s0480 | 14 – 71 | 39 – 85 | -9.55058 | 5.22035 | 4.87999 |
| CD630_18680 | CD630_s0640 | 66 – 94 | 4 – 30 | -14.352 | 3.04315 | 5.24925 |
| CD630_08460 | CD630_s0640 | 131 – 155 | 8 – 32 | -13.4383 | 4.15083 | 5.24925 |
| CD630_13620 | CD630_s0640 | 177 – 198 | 50 – 74 | -11.6224 | 4.77627 | 5.24925 |
| bglA | CD630_s0640 | 108 – 135 | 1 – 31 | -11.0332 | 4.50315 | 5.24925 |
| folC | CD630_s0640 | 88 – 116 | 4 – 30 | -10.8233 | 3.85634 | 5.24925 |
| CD630_19780 | CD630_s0640 | 8 – 26 | 1 – 21 | -10.7494 | 3.64830 | 5.24925 |
| CD630_02420 | CD630_s0640 | 82 – 96 | 1 – 15 | -10.6155 | 3.71048 | 5.24925 |
| CD630_15360 | CD630_s0640 | 65 – 97 | 1 – 29 | -10.5439 | 4.47208 | 5.24925 |
| CD630_05030 | CD630_s0640 | 36 – 57 | 1 – 21 | -10.5246 | 4.03823 | 5.24925 |
| terD | CD630_s0640 | 185 – 201 | 64 – 80 | -10.5095 | 3.33148 | 5.24925 |
| CD630_06080 | CD630_s0640 | 178 – 188 | 1 – 11 | -10.4231 | 3.82315 | 5.24925 |
| CD630_17170 | CD630_s0640 | 84 – 95 | 2 – 13 | -10.3472 | 4.10230 | 5.24925 |
| CD630_11730 | CD630_s0640 | 148 – 160 | 2 – 14 | -10.3406 | 4.80521 | 5.24925 |
| aroD | CD630_s0640 | 87 – 100 | 1 – 14 | -10.2794 | 3.71929 | 5.24925 |
| kdpD | CD630_s0640 | 149 – 158 | 1 – 10 | -10.215 | 4.27140 | 5.24925 |
| def1 | CD630_s0640 | 86 – 117 | 1 – 32 | -10.0852 | 3.99720 | 5.24925 |
| CD630_23450 | CD630_s0640 | 180 – 194 | 98 – 113 | -9.92479 | 3.27218 | 5.24925 |
| CD630_21020 | CD630_s0640 | 87 – 96 | 1 – 10 | -9.87013 | 4.28666 | 5.24925 |
| CD630_18090 | CD630_s0640 | 54 – 75 | 1 – 24 | -9.79196 | 4.23961 | 5.24925 |
| panB | CD630_s0640 | 137 – 148 | 2 – 13 | -9.764 | 3.40248 | 5.24925 |
| bioB | CD630_s0640 | 97 – 109 | 66 – 78 | -9.68685 | 3.59022 | 5.24925 |
| CD630_26680 | CD630_s0641 | 23 – 146 | 77 – 170 | -15.5957 | 3.53956 | 6.68432 |
| gcvTPA | CD630_s0641 | 139 – 180 | 217 – 253 | -15.4589 | 4.77229 | 6.68432 |
| CD630_05821 | CD630_s0641 | 78 – 114 | 253 – 286 | -15.3556 | 3.75661 | 6.68432 |
| CD630_21620 | CD630_s0641 | 5 – 38 | 211 – 242 | -15.2363 | 3.02649 | 6.68432 |
| CD630_14420 | CD630_s0641 | 87 – 100 | 370 – 383 | -15.1257 | 3.99497 | 6.68432 |
| cprK | CD630_s0641 | 95 – 160 | 120 – 168 | -15.0519 | 3.96216 | 6.68432 |
| CD630_20660 | CD630_s0641 | 128 – 162 | 220 – 251 | -14.9291 | 5.39321 | 6.68432 |
| cdu2 | CD630_s0641 | 86 – 118 | 129 – 160 | -14.4605 | 2.97335 | 6.68432 |
| CD630_25610 | CD630_s0641 | 58 – 123 | 110 – 169 | -14.3849 | 3.02685 | 6.68432 |
| CD630_02420 | CD630_s0641 | 6 – 40 | 148 – 181 | -14.3738 | 3.71048 | 6.68432 |
| CD630_27110 | CD630_s0641 | 46 – 60 | 183 – 197 | -13.9335 | 3.78231 | 6.68432 |
| CD630_23770 | CD630_s0641 | 103 – 133 | 219 – 250 | -13.8627 | 3.25181 | 6.68432 |
| CD630_13620 | CD630_s0641 | 148 – 197 | 201 – 252 | -13.738 | 4.77627 | 6.68432 |
| appF | CD630_s0641 | 45 – 192 | 15 – 128 | -13.6925 | 4.25570 | 6.68432 |
| CD630_36690 | CD630_s0641 | 117 – 169 | 54 – 97 | -13.6826 | 3.53761 | 6.68432 |
| dnaD | CD630_s0641 | 15 – 35 | 87 – 108 | -13.6133 | 3.56078 | 6.68432 |
| CD630_17360 | CD630_s0641 | 5 – 32 | 102 – 130 | -13.5117 | 5.66070 | 6.68432 |
| CD630_05211 | CD630_s0641 | 119 – 148 | 292 – 319 | -13.4189 | 3.97048 | 6.68432 |
| CD630_30140 | CD630_s0641 | 3 – 22 | 220 – 242 | -13.3893 | 3.28343 | 6.68432 |
| panB | CD630_s0641 | 146 – 174 | 220 – 252 | -13.2271 | 3.40248 | 6.68432 |
| CD630_34560 | CD630_s0641 | 26 – 65 | 224 – 253 | -13.0178 | 4.02604 | 6.68432 |
| CD630_06050 | CD630_s0641 | 39 – 84 | 35 – 77 | -12.9742 | 3.59332 | 6.68432 |
| CD630_23760 | CD630_s0641 | 145 – 193 | 86 – 132 | -12.8229 | 4.54307 | 6.68432 |
| dnaC | CD630_s0641 | 86 – 151 | 217 – 281 | -12.8125 | 6.30796 | 6.68432 |
| pgm | CD630_s0641 | 147 – 165 | 208 – 226 | -12.787 | 3.45439 | 6.68432 |
| cdtR | CD630_s0641 | 3 – 106 | 38 – 124 | -12.7834 | 5.84368 | 6.68432 |
| CD630_14710 | CD630_s0641 | 51 – 97 | 389 – 437 | -12.6828 | 3.65171 | 6.68432 |
| CD630_22550 | CD630_s0641 | 137 – 165 | 225 – 255 | -12.5767 | 3.62379 | 6.68432 |
| CD630_26020 | CD630_s0641 | 4 – 49 | 110 – 159 | -12.427 | 4.03155 | 6.68432 |
| CD630_06060 | CD630_s0641 | 115 – 158 | 265 – 299 | -12.4185 | 4.94050 | 6.68432 |
| CD630_06340 | CD630_s0641 | 19 – 36 | 75 – 92 | -12.3755 | 4.35983 | 6.68432 |
| CD630_14190 | CD630_s0641 | 147 – 172 | 109 – 142 | -12.3311 | 3.78562 | 6.68432 |
| CD630_06060 | CD630_s0641 | 112 – 147 | 292 – 323 | -12.3291 | 4.94050 | 6.68432 |
| opuCC | CD630_s0641 | 92 – 117 | 124 – 157 | -12.2701 | 4.07035 | 6.68432 |
| CD630_22890 | CD630_s0641 | 2 – 50 | 210 – 256 | -12.1205 | 3.75591 | 6.68432 |
| CD630_32890 | CD630_s0641 | 161 – 186 | 217 – 242 | -12.0622 | 3.41276 | 6.68432 |
| bglA | CD630_s0641 | 152 – 195 | 254 – 290 | -12.0024 | 4.86402 | 6.68432 |
| CD630_18010 | CD630_s0641 | 22 – 61 | 394 – 438 | -11.9985 | 4.21967 | 6.68432 |
| appB | CD630_s0641 | 157 – 174 | 282 – 299 | -11.9336 | 6.44103 | 6.68432 |
| CD630_05490 | CD630_s0641 | 161 – 184 | 219 – 242 | -11.9248 | 3.89852 | 6.68432 |
| CD630_24760 | CD630_s0641 | 86 – 139 | 209 – 256 | -11.9173 | 5.19485 | 6.68432 |
| CD630_16530 | CD630_s0641 | 23 – 169 | 33 – 178 | -11.8573 | 4.37658 | 6.68432 |
| CD630_03180 | CD630_s0641 | 136 – 151 | 393 – 408 | -11.8391 | 3.73138 | 6.68432 |
| CD630_18660 | CD630_s0641 | 126 – 148 | 335 – 354 | -11.8366 | 3.50371 | 6.68432 |
| xdhA | CD630_s0641 | 25 – 69 | 259 – 304 | -11.7012 | 4.45147 | 6.68432 |
| CD630_29650 | CD630_s0641 | 86 – 183 | 66 – 157 | -11.6969 | 3.67128 | 6.68432 |
| CD630_08510 | CD630_s0641 | 58 – 87 | 271 – 302 | -11.6905 | 5.16329 | 6.68432 |
| CD630_24590 | CD630_s0641 | 103 – 136 | 135 – 170 | -11.6253 | 3.70401 | 6.68432 |
| CD630_20170 | CD630_s0641 | 102 – 121 | 390 – 410 | -11.6209 | 6.90201 | 6.68432 |
| CD630_22070 | CD630_s0641 | 90 – 161 | 236 – 304 | -11.6045 | 4.09651 | 6.68432 |
| CD630_13350 | CD630_s0641 | 167 – 195 | 226 – 246 | -11.5834 | 4.15139 | 6.68432 |
| cbiO | CD630_s0641 | 9 – 41 | 220 – 247 | -11.5569 | 3.79830 | 6.68432 |
| CD630_04610 | CD630_s0641 | 149 – 196 | 129 – 179 | -11.5331 | 3.83025 | 6.68432 |
| CD630_28320 | CD630_s0641 | 88 – 99 | 145 – 156 | -11.526 | 3.99184 | 6.68432 |
| CD630_28550 | CD630_s0641 | 78 – 109 | 135 – 160 | -11.478 | 3.51196 | 6.68432 |
| CD630_36781 | CD630_s0641 | 3 – 26 | 151 – 173 | -11.4604 | 3.75946 | 6.68432 |
| CD630_06110 | CD630_s0641 | 94 – 130 | 219 – 253 | -11.3717 | 5.61380 | 6.68432 |
| CD630_26150 | CD630_s0641 | 152 – 190 | 222 – 258 | -11.3702 | 4.06302 | 6.68432 |
| CD630_21770 | CD630_s0641 | 104 – 133 | 225 – 252 | -11.3612 | 4.02587 | 6.68432 |
| CD630_31500 | CD630_s0641 | 146 – 178 | 293 – 325 | -11.3484 | 4.90207 | 6.68432 |
| ppnK | CD630_s0641 | 153 – 197 | 218 – 254 | -11.3306 | 4.36817 | 6.68432 |
| tcdC | CD630_s0641 | 112 – 160 | 203 – 252 | -11.2052 | 3.76101 | 6.68432 |
| CD630_03930 | CD630_s0641 | 133 – 169 | 98 – 132 | -11.0883 | 4.15934 | 6.68432 |
| CD630_25070 | CD630_s0641 | 135 – 170 | 86 – 119 | -11.0871 | 3.45164 | 6.68432 |
| CD630_21090 | CD630_s0641 | 151 – 172 | 415 – 436 | -11.0359 | 4.15803 | 6.68432 |
| CD630_14150 | CD630_s0641 | 22 – 50 | 171 – 197 | -11.0154 | 3.28251 | 6.68432 |
| CD630_22720 | CD630_s0641 | 137 – 157 | 364 – 382 | -10.9891 | 3.81804 | 6.68432 |
| CD630_34530 | CD630_s0641 | 179 – 192 | 271 – 283 | -10.9489 | 3.83016 | 6.68432 |
| CD630_01590 | CD630_s0641 | 18 – 30 | 234 – 246 | -10.9346 | 3.45609 | 6.68432 |
| CD630_23780 | CD630_s0641 | 146 – 185 | 36 – 78 | -10.9168 | 3.91629 | 6.68432 |
| CD630_18960 | CD630_s0641 | 171 – 193 | 53 – 79 | -10.9068 | 4.37822 | 6.68432 |
| CD630_30730 | CD630_s0641 | 73 – 174 | 68 – 170 | -10.8736 | 3.13236 | 6.68432 |
| rbsC | CD630_s0641 | 61 – 106 | 259 – 303 | -10.845 | 3.71650 | 6.68432 |
| CD630_27120 | CD630_s0641 | 131 – 157 | 208 – 233 | -10.7791 | 4.43720 | 6.68432 |
| uxaA | CD630_s0641 | 72 – 94 | 377 – 400 | -10.7589 | 4.26165 | 6.68432 |
| CD630_07840 | CD630_s0641 | 144 – 165 | 99 – 118 | -10.7444 | 4.26761 | 6.68432 |
| CD630_27570 | CD630_s0641 | 122 – 171 | 203 – 256 | -10.6998 | 3.46216 | 6.68432 |
| CD630_16640 | CD630_s0641 | 1 – 136 | 60 – 180 | -10.6872 | 3.61548 | 6.68432 |
| cwp26 | CD630_s0641 | 136 – 149 | 63 – 76 | -10.6445 | 4.44377 | 6.68432 |
| CD630_22710 | CD630_s0641 | 2 – 16 | 154 – 168 | -10.6377 | 4.05623 | 6.68432 |
| CD630_27280 | CD630_s0641 | 10 – 35 | 34 – 58 | -10.625 | 3.24738 | 6.68432 |
| CD630_33580 | CD630_s0641 | 69 – 102 | 147 – 178 | -10.6149 | 3.95778 | 6.68432 |
| CD630_33570 | CD630_s0641 | 134 – 175 | 214 – 250 | -10.611 | 3.95778 | 6.68432 |
| chrA | CD630_s0641 | 138 – 156 | 152 – 171 | -10.6009 | 5.27255 | 6.68432 |
| CD630_06170 | CD630_s0641 | 150 – 165 | 176 – 190 | -10.5897 | 4.95366 | 6.68432 |
| CD630_30310 | CD630_s0641 | 49 – 78 | 217 – 245 | -10.5857 | 4.56059 | 6.68432 |
| bglA | CD630_s0641 | 63 – 100 | 209 – 250 | -10.5784 | 4.50315 | 6.68432 |
| CD630_27040 | CD630_s0641 | 47 – 82 | 95 – 125 | -10.5713 | 3.51812 | 6.68432 |
| srlD | CD630_s0641 | 113 – 147 | 127 – 178 | -10.4754 | 4.19843 | 6.68432 |
| garK | CD630_s0641 | 1 – 25 | 153 – 173 | -10.4711 | 3.72836 | 6.68432 |
| CD630_18120 | CD630_s0641 | 89 – 98 | 374 – 383 | -10.4417 | 3.99713 | 6.68432 |
| CD630_26250 | CD630_s0641 | 122 – 167 | 93 – 140 | -10.4414 | 5.03379 | 6.68432 |
| CD630_09100 | CD630_s0641 | 33 – 71 | 398 – 436 | -10.405 | 5.92260 | 6.68432 |
| CD630_15460 | CD630_s0641 | 137 – 200 | 71 – 141 | -10.3647 | 6.86296 | 6.68432 |
| CD630_20260 | CD630_s0641 | 137 – 146 | 215 – 224 | -10.3553 | 3.84162 | 6.68432 |
| srlEb | CD630_s0641 | 126 – 153 | 241 – 267 | -10.3542 | 3.50140 | 6.68432 |
| CD630_31220 | CD630_s0641 | 151 – 179 | 345 – 367 | -10.3533 | 3.73779 | 6.68432 |
| CD630_27270 | CD630_s0641 | 142 – 173 | 220 – 250 | -10.3347 | 3.50892 | 6.68432 |
| chrA | CD630_s0641 | 97 – 110 | 300 – 314 | -10.3205 | 5.27255 | 6.68432 |
| CD630_07870 | CD630_s0641 | 170 – 201 | 221 – 254 | -10.3121 | 3.22829 | 6.68432 |
| CD630_06100 | CD630_s0641 | 116 – 127 | 183 – 194 | -10.3032 | 4.43918 | 6.68432 |
| CD630_18950 | CD630_s0641 | 137 – 169 | 208 – 242 | -10.2808 | 3.40610 | 6.68432 |
| scpB | CD630_s0641 | 130 – 186 | 241 – 302 | -10.2686 | 3.33560 | 6.68432 |
| CD630_21760 | CD630_s0641 | 170 – 189 | 226 – 244 | -10.2583 | 4.43846 | 6.68432 |
| CD630_08070 | CD630_s0641 | 155 – 177 | 387 – 409 | -10.2448 | 4.87819 | 6.68432 |
| CD630_11241 | CD630_s0641 | 163 – 182 | 367 – 383 | -10.2396 | 4.85723 | 6.68432 |
| CD630_05940 | CD630_s0641 | 162 – 190 | 118 – 146 | -10.2326 | 4.18595 | 6.68432 |
| nrdD | CD630_s0641 | 147 – 169 | 201 – 223 | -10.2084 | 3.28290 | 6.68432 |
| CD630_19690 | CD630_s0641 | 33 – 43 | 218 – 228 | -10.1997 | 3.85861 | 6.68432 |
| CD630_13020 | CD630_s0641 | 118 – 133 | 175 – 191 | -10.1497 | 3.52267 | 6.68432 |
| CD630_34080 | CD630_s0641 | 88 – 112 | 389 – 412 | -10.1462 | 3.20224 | 6.68432 |
| CD630_19942 | CD630_s0641 | 119 – 130 | 215 – 227 | -10.1383 | 4.24574 | 6.68432 |
| CD630_28280 | CD630_s0641 | 38 – 80 | 258 – 293 | -10.1369 | 4.59833 | 6.68432 |
| CD630_04880 | CD630_s0641 | 9 – 19 | 336 – 346 | -10.0918 | 4.62291 | 6.68432 |
| CD630_21110 | CD630_s0641 | 123 – 158 | 212 – 249 | -10.0752 | 3.89630 | 6.68432 |
| CD630_22880 | CD630_s0641 | 174 – 196 | 212 – 235 | -10.0601 | 3.62455 | 6.68432 |
| moaA | CD630_s0641 | 86 – 113 | 135 – 156 | -10.0493 | 3.83799 | 6.68432 |
| CD630_13920 | CD630_s0641 | 172 – 200 | 340 – 364 | -10.0315 | 3.25236 | 6.68432 |
| CD630_36180 | CD630_s0641 | 168 – 200 | 177 – 205 | -10.0252 | 4.83428 | 6.68432 |
| CD630_07480 | CD630_s0641 | 79 – 110 | 255 – 284 | -9.99 | 5.18321 | 6.68432 |
| CD630_31480 | CD630_s0641 | 162 – 192 | 113 – 141 | -9.98871 | 4.88897 | 6.68432 |
| CD630_31960 | CD630_s0641 | 5 – 41 | 265 – 304 | -9.98251 | 4.93757 | 6.68432 |
| CD630_04341 | CD630_s0641 | 127 – 174 | 211 – 252 | -9.95723 | 4.29262 | 6.68432 |
| metQ | CD630_s0641 | 28 – 85 | 95 – 156 | -9.95168 | 3.87190 | 6.68432 |
| CD630_08140 | CD630_s0641 | 1 – 11 | 184 – 194 | -9.9495 | 3.30159 | 6.68432 |
| CD630_06200 | CD630_s0641 | 69 – 98 | 269 – 302 | -9.93141 | 3.57064 | 6.68432 |
| appC | CD630_s0641 | 44 – 66 | 225 – 245 | -9.91711 | 4.50438 | 6.68432 |
| CD630_19700 | CD630_s0641 | 165 – 175 | 297 – 308 | -9.87938 | 3.85861 | 6.68432 |
| CD630_32990 | CD630_s0641 | 169 – 200 | 224 – 245 | -9.87531 | 5.24925 | 6.68432 |
| CD630_08950 | CD630_s0641 | 8 – 21 | 225 – 237 | -9.86368 | 3.71905 | 6.68432 |
| CD630_05300 | CD630_s0641 | 145 – 175 | 67 – 99 | -9.86233 | 4.56885 | 6.68432 |
| CD630_20171 | CD630_s0641 | 33 – 55 | 393 – 415 | -9.85079 | 6.87491 | 6.68432 |
| CD630_33480 | CD630_s0641 | 113 – 139 | 270 – 301 | -9.83305 | 3.67613 | 6.68432 |
| bcp | CD630_s0641 | 138 – 159 | 391 – 410 | -9.82967 | 3.16670 | 6.68432 |
| CD630_26920 | CD630_s0641 | 161 – 174 | 396 – 409 | -9.82193 | 4.58060 | 6.68432 |
| CD630_29750 | CD630_s0641 | 39 – 83 | 205 – 249 | -9.79651 | 4.59146 | 6.68432 |
| CD630_18940 | CD630_s0641 | 52 – 98 | 52 – 96 | -9.76844 | 4.09988 | 6.68432 |
| CD630_09060 | CD630_s0641 | 88 – 108 | 257 – 276 | -9.75509 | 4.31834 | 6.68432 |
| CD630_05460 | CD630_s0641 | 3 – 35 | 6 – 37 | -9.74561 | 3.98753 | 6.68432 |
| cbiQ | CD630_s0641 | 20 – 44 | 173 – 195 | -9.73814 | 3.79830 | 6.68432 |
| CD630_13790 | CD630_s0641 | 93 – 106 | 38 – 51 | -9.73527 | 5.17374 | 6.68432 |
| pilC | CD630_s0641 | 38 – 48 | 337 – 347 | -9.7198 | 3.54230 | 6.68432 |
| CD630_10220 | CD630_s0641 | 193 – 201 | 34 – 42 | -9.71465 | 4.08233 | 6.68432 |
| CD630_28260 | CD630_s0641 | 173 – 198 | 299 – 326 | -9.68851 | 3.51613 | 6.68432 |
| CD630_03040 | CD630_s0641 | 67 – 92 | 217 – 244 | -9.68321 | 3.51403 | 6.68432 |
| CD630_15080 | CD630_s0641 | 14 – 88 | 81 – 168 | -9.66413 | 4.03033 | 6.68432 |
| cls | CD630_s0641 | 177 – 189 | 234 – 246 | -9.65052 | 4.46177 | 6.68432 |
| etfA | CD630_s0641 | 45 – 75 | 222 – 256 | -9.63815 | 3.41204 | 6.68432 |
| CD630_03920 | CD630_s0641 | 97 – 138 | 66 – 111 | -9.62617 | 4.15934 | 6.68432 |
| CD630_32060 | CD630_s0641 | 39 – 49 | 390 – 400 | -9.59227 | 5.73797 | 6.68432 |
| CD630_27200 | CD630_s0641 | 27 – 40 | 177 – 191 | -9.58994 | 3.42516 | 6.68432 |
| cobQ | CD630_s0641 | 88 – 103 | 369 – 383 | -9.58755 | 3.13524 | 6.68432 |
| CD630_17170 | CD630_s0641 | 168 – 177 | 149 – 158 | -9.58421 | 4.10230 | 6.68432 |
| ribD | CD630_s0641 | 170 – 186 | 365 – 380 | -9.5783 | 6.65922 | 6.68432 |
| CD630_21230 | CD630_s0641 | 143 – 160 | 297 – 313 | -9.55361 | 3.89213 | 6.68432 |
| CD630_17720 | CD630_s0641 | 2 – 42 | 63 – 101 | -9.52375 | 3.22910 | 6.68432 |
| CD630_32520 | CD630_s0641 | 132 – 168 | 337 – 372 | -9.51209 | 4.06098 | 6.68432 |
| CD630_33420 | CD630_SQ1005 | 16 – 153 | 9 – 124 | -15.4762 | 3.22095 | 3.63582 |
| CD630_27360 | CD630_SQ1005 | 84 – 134 | 16 – 60 | -14.1649 | 3.79570 | 3.63582 |
| CD630_21730 | CD630_SQ1005 | 39 – 135 | 19 – 125 | -13.7451 | 3.45076 | 3.63582 |
| CD630_14340 | CD630_SQ1005 | 33 – 79 | 64 – 122 | -13.2333 | 5.10868 | 3.63582 |
| CD630_04320 | CD630_SQ1005 | 160 – 185 | 182 – 204 | -13.0132 | 3.68342 | 3.63582 |
| CD630_03170 | CD630_SQ1005 | 87 – 107 | 193 – 214 | -12.808 | 3.66946 | 3.63582 |
| CD630_14600 | CD630_SQ1005 | 138 – 198 | 15 – 67 | -12.7419 | 5.90348 | 3.63582 |
| CD630_22030 | CD630_SQ1005 | 138 – 182 | 79 – 123 | -12.6717 | 3.29934 | 3.63582 |
| pilC | CD630_SQ1005 | 129 – 147 | 200 – 214 | -12.6258 | 3.54230 | 3.63582 |
| bglA | CD630_SQ1005 | 178 – 201 | 190 – 214 | -12.1825 | 4.86402 | 3.63582 |
| CD630_23820 | CD630_SQ1005 | 3 – 43 | 6 – 53 | -11.782 | 3.18419 | 3.63582 |
| CD630_06380 | CD630_SQ1005 | 118 – 165 | 78 – 122 | -11.6161 | 4.32211 | 3.63582 |
| CD630_34520 | CD630_SQ1005 | 139 – 200 | 7 – 65 | -11.4199 | 3.83016 | 3.63582 |
| blaI | CD630_SQ1005 | 40 – 53 | 37 – 50 | -11.3625 | 3.73674 | 3.63582 |
| CD630_14810 | CD630_SQ1005 | 127 – 136 | 188 – 197 | -11.0784 | 3.35323 | 3.63582 |
| CD630_16150 | CD630_SQ1005 | 60 – 71 | 49 – 60 | -11.0494 | 4.20614 | 3.63582 |
| eutW | CD630_SQ1005 | 10 – 22 | 199 – 212 | -11.0098 | 4.55715 | 3.63582 |
| CD630_06060 | CD630_SQ1005 | 106 – 145 | 33 – 74 | -10.9657 | 4.94050 | 3.63582 |
| CD630_06060 | CD630_SQ1005 | 77 – 184 | 102 – 200 | -10.9583 | 4.94050 | 3.63582 |
| CD630_20980 | CD630_SQ1005 | 177 – 201 | 39 – 65 | -10.8853 | 8.12847 | 3.63582 |
| CD630_03650 | CD630_SQ1005 | 174 – 197 | 20 – 40 | -10.8698 | 3.93697 | 3.63582 |
| chrA | CD630_SQ1005 | 39 – 60 | 43 – 63 | -10.8675 | 5.27255 | 3.63582 |
| CD630_21610 | CD630_SQ1005 | 139 – 166 | 25 – 52 | -10.8645 | 3.16662 | 3.63582 |
| CD630_27270 | CD630_SQ1005 | 90 – 109 | 38 – 60 | -10.8559 | 3.50892 | 3.63582 |
| CD630_25200 | CD630_SQ1005 | 179 – 200 | 87 – 109 | -10.831 | 3.65158 | 3.63582 |
| hfq | CD630_SQ1005 | 148 – 184 | 175 – 212 | -10.8152 | 5.05609 | 3.63582 |
| CD630_32060 | CD630_SQ1005 | 161 – 189 | 10 – 34 | -10.7985 | 5.73797 | 3.63582 |
| CD630_19903 | CD630_SQ1005 | 46 – 59 | 61 – 74 | -10.6823 | 5.79994 | 3.63582 |
| CD630_05030 | CD630_SQ1005 | 121 – 136 | 6 – 24 | -10.5912 | 4.03823 | 3.63582 |
| CD630_16950 | CD630_SQ1005 | 177 – 199 | 86 – 106 | -10.5712 | 4.42569 | 3.63582 |
| CD630_06540 | CD630_SQ1005 | 82 – 118 | 175 – 216 | -10.5435 | 3.54673 | 3.63582 |
| CD630_16160 | CD630_SQ1005 | 43 – 66 | 43 – 64 | -10.5333 | 6.11801 | 3.63582 |
| CD630_30420 | CD630_SQ1005 | 136 – 200 | 84 – 153 | -10.4998 | 5.27539 | 3.63582 |
| CD630_04540 | CD630_SQ1005 | 82 – 109 | 192 – 215 | -10.4245 | 3.33556 | 3.63582 |
| pilT | CD630_SQ1005 | 122 – 180 | 5 – 60 | -10.4243 | 3.24639 | 3.63582 |
| CD630_33580 | CD630_SQ1005 | 178 – 194 | 33 – 50 | -10.3841 | 3.95778 | 3.63582 |
| CD630_10080 | CD630_SQ1005 | 146 – 171 | 87 – 109 | -10.3684 | 3.75625 | 3.63582 |
| CD630_05450 | CD630_SQ1005 | 158 – 177 | 102 – 121 | -10.3522 | 5.04761 | 3.63582 |
| CD630_15890 | CD630_SQ1005 | 147 – 170 | 82 – 102 | -10.3172 | 3.39962 | 3.63582 |
| CD630_22740 | CD630_SQ1005 | 78 – 95 | 195 – 211 | -10.3172 | 3.08733 | 3.63582 |
| CD630_21570 | CD630_SQ1005 | 126 – 173 | 10 – 49 | -10.3165 | 6.48385 | 3.63582 |
| CD630_08360 | CD630_SQ1005 | 45 – 68 | 66 – 91 | -10.2587 | 3.19339 | 3.63582 |
| CD630_34050 | CD630_SQ1005 | 139 – 157 | 32 – 50 | -10.2581 | 4.03863 | 3.63582 |
| cwp26 | CD630_SQ1005 | 186 – 199 | 36 – 49 | -10.2485 | 4.44377 | 3.63582 |
| ppnK | CD630_SQ1005 | 126 – 165 | 8 – 50 | -10.2424 | 4.36817 | 3.63582 |
| CD630_10960 | CD630_SQ1005 | 89 – 107 | 193 – 212 | -10.2363 | 3.53850 | 3.63582 |
| CD630_17770 | CD630_SQ1005 | 129 – 142 | 1 – 13 | -10.2029 | 5.06490 | 3.63582 |
| CD630_05370 | CD630_SQ1005 | 126 – 158 | 53 – 95 | -10.0933 | 3.26947 | 3.63582 |
| CD630_24590 | CD630_SQ1005 | 83 – 107 | 51 – 74 | -10.0432 | 3.70401 | 3.63582 |
| CD630_18120 | CD630_SQ1005 | 86 – 100 | 203 – 217 | -10.042 | 3.99713 | 3.63582 |
| CD630_19440 | CD630_SQ1005 | 144 – 186 | 2 – 46 | -10.03 | 4.53937 | 3.63582 |
| CD630_22500 | CD630_SQ1005 | 130 – 172 | 79 – 124 | -9.96521 | 4.03688 | 3.63582 |
| dnaC | CD630_SQ1005 | 47 – 74 | 38 – 65 | -9.92697 | 6.30796 | 3.63582 |
| CD630_09050 | CD630_SQ1005 | 181 – 199 | 87 – 103 | -9.841 | 3.96216 | 3.63582 |
| CD630_27610 | CD630_SQ1005 | 114 – 124 | 86 – 96 | -9.79365 | 3.96465 | 3.63582 |
| CD630_35260 | CD630_SQ1005 | 66 – 95 | 55 – 76 | -9.76747 | 5.77277 | 3.63582 |
| CD630_15460 | CD630_SQ1005 | 89 – 106 | 199 – 215 | -9.69447 | 6.86296 | 3.63582 |
| CD630_22550 | CD630_SQ1005 | 159 – 184 | 179 – 201 | -9.68499 | 3.62379 | 3.63582 |
| bltD | CD630_SQ1005 | 188 – 198 | 196 – 206 | -9.63908 | 4.49926 | 3.63582 |
| CD630_34080 | CD630_SQ1005 | 45 – 75 | 21 – 47 | -9.57283 | 3.20224 | 3.63582 |
| CD630_09100 | CD630_SQ1005 | 1 – 28 | 20 – 46 | -9.53852 | 5.92260 | 3.63582 |
| CD630_06150 | CD630_SQ1005 | 83 – 109 | 200 – 219 | -9.53703 | 5.51744 | 3.63582 |
| serS1 | CD630_SQ1005 | 159 – 170 | 203 – 214 | -9.52644 | 3.45858 | 3.63582 |
| pmi | CD630_SQ1005 | 187 – 201 | 15 – 27 | -9.52555 | 3.86845 | 3.63582 |
| bioB | CD630_SQ1038 | 103 – 140 | 130 – 165 | -18.7435 | 3.59022 | 3.49019 |
| CD630_24770 | CD630_SQ1038 | 164 – 185 | 3 – 27 | -15.5838 | 3.20798 | 3.49019 |

| | | | | | | |
|---|---|---|---|---|---|---|
| xdhA | CD630_SQ1038 | 121–148 | 144–173 | -15.2262 | 4.45147 | 3.49019 |
| panG | CD630_SQ1038 | 48–85 | 118–155 | -15.1716 | 5.54047 | 3.49019 |
| CD630_27200 | CD630_SQ1038 | 1–16 | 112–128 | -14.4274 | 3.42516 | 3.49019 |
| CD630_21140 | CD630_SQ1038 | 11–42 | 92–119 | -14.193 | 3.33304 | 3.49019 |
| CD630_08360 | CD630_SQ1038 | 39–60 | 134–155 | -13.9198 | 3.19339 | 3.49019 |
| CD630_18920 | CD630_SQ1038 | 114–163 | 111–149 | -13.8452 | 4.43558 | 3.49019 |
| CD630_26050 | CD630_SQ1038 | 128–151 | 3–30 | -13.6394 | 6.99851 | 3.49019 |
| CD630_13020 | CD630_SQ1038 | 115–139 | 71–93 | -13.6009 | 3.52267 | 3.49019 |
| CD630_28290 | CD630_SQ1038 | 129–157 | 117–151 | -13.1116 | 5.54192 | 3.49019 |
| CD630_34520 | CD630_SQ1038 | 161–200 | 111–167 | -13.0848 | 3.83016 | 3.49019 |
| thiH | CD630_SQ1038 | 88–102 | 7–21 | -12.9342 | 3.58230 | 3.49019 |
| CD630_05830 | CD630_SQ1038 | 74–169 | 74–175 | -12.6847 | 3.35360 | 3.49019 |
| CD630_32690 | CD630_SQ1038 | 6–29 | 1–21 | -12.1776 | 7.32676 | 3.49019 |
| cwp17 | CD630_SQ1038 | 49–75 | 148–173 | -12.1439 | 3.43363 | 3.49019 |
| CD630_19850 | CD630_SQ1038 | 9–33 | 192–219 | -12.1164 | 6.88486 | 3.49019 |
| CD630_17292 | CD630_SQ1038 | 128–190 | 145–201 | -12.0127 | 6.25799 | 3.49019 |
| CD630_03400 | CD630_SQ1038 | 69–88 | 193–212 | -11.5979 | 4.45994 | 3.49019 |
| CD630_27770 | CD630_SQ1038 | 92–114 | 139–161 | -11.3912 | 4.08588 | 3.49019 |
| CD630_02720 | CD630_SQ1038 | 141–176 | 50–85 | -11.3845 | 3.38938 | 3.49019 |
| iorA | CD630_SQ1038 | 2–38 | 194–231 | -11.1684 | 3.83173 | 3.49019 |
| hfq | CD630_SQ1038 | 159–194 | 114–150 | -11.1136 | 5.05609 | 3.49019 |
| CD630_36040 | CD630_SQ1038 | 51–123 | 118–182 | -11.0004 | 3.52009 | 3.49019 |
| eutV | CD630_SQ1038 | 143–177 | 119–155 | -10.9998 | 4.55715 | 3.49019 |
| CD630_21620 | CD630_SQ1038 | 96–121 | 4–28 | -10.9738 | 3.02649 | 3.49019 |
| CD630_27230 | CD630_SQ1038 | 91–135 | 143–189 | -10.9183 | 3.52462 | 3.49019 |
| CD630_07430 | CD630_SQ1038 | 152–195 | 88–149 | -10.7355 | 4.65568 | 3.49019 |
| CD630_04850 | CD630_SQ1038 | 144–170 | 2–28 | -10.728 | 4.40788 | 3.49019 |
| CD630_19700 | CD630_SQ1038 | 42–57 | 147–162 | -10.6822 | 3.85861 | 3.49019 |
| CD630_06340 | CD630_SQ1038 | 140–179 | 112–149 | -10.6367 | 4.35983 | 3.49019 |
| CD630_06060 | CD630_SQ1038 | 57–94 | 81–130 | -10.6289 | 4.94050 | 3.49019 |
| CD630_33560 | CD630_SQ1038 | 66–118 | 186–233 | -10.6203 | 4.03147 | 3.49019 |
| kdpE | CD630_SQ1038 | 57–94 | 117–155 | -10.5841 | 4.27140 | 3.49019 |
| CD630_08140 | CD630_SQ1038 | 119–190 | 81–164 | -10.5344 | 3.30159 | 3.49019 |
| CD630_27110 | CD630_SQ1038 | 9–55 | 131–176 | -10.4751 | 3.78231 | 3.49019 |
| eutW | CD630_SQ1038 | 9–22 | 15–28 | -10.425 | 4.55715 | 3.49019 |
| CD630_15190 | CD630_SQ1038 | 167–179 | 12–24 | -10.3804 | 4.43255 | 3.49019 |
| CD630_10050 | CD630_SQ1038 | 43–62 | 144–162 | -10.3766 | 3.93750 | 3.49019 |
| CD630_02790 | CD630_SQ1038 | 97–111 | 14–27 | -10.3624 | 4.77661 | 3.49019 |
| moaA | CD630_SQ1038 | 70–114 | 197–236 | -10.2854 | 3.83799 | 3.49019 |
| CD630_17720 | CD630_SQ1038 | 150–166 | 6–23 | -10.2388 | 3.22910 | 3.49019 |
| CD630_05460 | CD630_SQ1038 | 90–133 | 143–189 | -10.1694 | 3.98753 | 3.49019 |
| CD630_19903 | CD630_SQ1038 | 106–122 | 146–162 | -9.99636 | 5.79994 | 3.49019 |
| CD630_03040 | CD630_SQ1038 | 10–26 | 137–153 | -9.97963 | 3.51403 | 3.49019 |
| CD630_15740 | CD630_SQ1038 | 43–92 | 118–165 | -9.95541 | 4.43222 | 3.49019 |
| CD630_16640 | CD630_SQ1038 | 18–31 | 4–17 | -9.94577 | 3.61548 | 3.49019 |
| CD630_08100 | CD630_SQ1038 | 186–201 | 8–23 | -9.9202 | 4.39181 | 3.49019 |
| gcp | CD630_SQ1038 | 20–37 | 1–23 | -9.91854 | 4.92798 | 3.49019 |
| CD630_04341 | CD630_SQ1038 | 167–183 | 145–161 | -9.87469 | 4.29262 | 3.49019 |
| CD630_18090 | CD630_SQ1038 | 14–35 | 133–155 | -9.85684 | 4.23961 | 3.49019 |
| CD630_13790 | CD630_SQ1038 | 167–192 | 131–152 | -9.84887 | 5.17374 | 3.49019 |
| CD630_09760 | CD630_SQ1038 | 109–149 | 116–159 | -9.8471 | 4.83410 | 3.49019 |
| CD630_28910 | CD630_SQ1038 | 109–149 | 116–159 | -9.8471 | 4.51664 | 3.49019 |
| cstA | CD630_SQ1038 | 80–121 | 122–168 | -9.83811 | 3.53584 | 3.49019 |
| CD630_04280 | CD630_SQ1038 | 79–109 | 134–162 | -9.81576 | 3.61457 | 3.49019 |
| CD630_18120 | CD630_SQ1038 | 50–64 | 224–238 | -9.80114 | 3.99713 | 3.49019 |
| CD630_02830 | CD630_SQ1038 | 39–75 | 70–106 | -9.77753 | 3.57106 | 3.49019 |
| metQ | CD630_SQ1038 | 50–130 | 73–170 | -9.76956 | 3.87190 | 3.49019 |
| CD630_24580 | CD630_SQ1038 | 123–135 | 107–119 | -9.76009 | 3.88519 | 3.49019 |
| CD630_32880 | CD630_SQ1038 | 5–29 | 61–84 | -9.75559 | 3.41276 | 3.49019 |
| CD630_14990 | CD630_SQ1038 | 164–181 | 112–130 | -9.65304 | 3.78619 | 3.49019 |
| CD630_22710 | CD630_SQ1038 | 61–103 | 120–161 | -9.61374 | 4.05623 | 3.49019 |
| argS | CD630_SQ1038 | 71–91 | 61–84 | -9.58286 | 4.96036 | 3.49019 |
| CD630_33682 | CD630_SQ1038 | 106–122 | 146–162 | -9.54012 | 5.77720 | 3.49019 |
| CD630_19690 | CD630_SQ1641 | 145–175 | 1–30 | -18.4343 | 3.85861 | 8.65138 |
| CD630_14630 | CD630_SQ1641 | 116–167 | 2–44 | -18.0134 | 3.34485 | 8.65138 |
| CD630_32520 | CD630_SQ1641 | 117–201 | 3–97 | -17.5205 | 4.06098 | 8.65138 |
| CD630_14182 | CD630_SQ1641 | 174–197 | 215–238 | -17.3551 | 4.82360 | 8.65138 |
| CD630_08360 | CD630_SQ1641 | 186–201 | 16–32 | -16.4191 | 3.19339 | 8.65138 |
| CD630_32880 | CD630_SQ1641 | 145–178 | 15–46 | -15.3212 | 3.41276 | 8.65138 |
| CD630_06060 | CD630_SQ1641 | 166–200 | 2–39 | -14.8332 | 4.94050 | 8.65138 |
| CD630_09970 | CD630_SQ1641 | 78–95 | 17–34 | -14.6428 | 3.25917 | 8.65138 |
| CD630_19903 | CD630_SQ1641 | 170–199 | 18–43 | -14.4068 | 5.79994 | 8.65138 |
| dnaC | CD630_SQ1641 | 178–200 | 66–88 | -13.9382 | 6.30796 | 8.65138 |
| CD630_28280 | CD630_SQ1641 | 49–91 | 11–55 | -13.8675 | 4.59833 | 8.65138 |
| CD630_08390 | CD630_SQ1641 | 75–105 | 1–32 | -13.8627 | 5.16000 | 8.65138 |
| opuCC | CD630_SQ1641 | 164–185 | 7–30 | -13.7909 | 4.07035 | 8.65138 |
| CD630_29650 | CD630_SQ1641 | 164–189 | 2–28 | -13.523 | 3.67128 | 8.65138 |
| CD630_36250 | CD630_SQ1641 | 1–24 | 22–44 | -13.4012 | 4.13224 | 8.65138 |
| CD630_11241 | CD630_SQ1641 | 163–198 | 172–208 | -13.2964 | 4.85723 | 8.65138 |
| CD630_23770 | CD630_SQ1641 | 108–133 | 15–45 | -13.2685 | 3.25181 | 8.65138 |
| pilC | CD630_SQ1641 | 57–96 | 2–42 | -13.2681 | 3.54230 | 8.65138 |
| CD630_33682 | CD630_SQ1641 | 170–198 | 2–45 | -13.2304 | 5.77720 | 8.65138 |
| def1 | CD630_SQ1641 | 111–136 | 15–42 | -13.0259 | 3.99720 | 8.65138 |
| CD630_19700 | CD630_SQ1641 | 19–45 | 5–32 | -12.9979 | 3.85861 | 8.65138 |
| CD630_23780 | CD630_SQ1641 | 89–146 | 152–210 | -12.9551 | 3.91629 | 8.65138 |
| CD630_09060 | CD630_SQ1641 | 94–124 | 5–39 | -12.9089 | 4.31834 | 8.65138 |
| CD630_14240 | CD630_SQ1641 | 170–198 | 2–45 | -12.8396 | 8.60766 | 8.65138 |
| CD630_09562 | CD630_SQ1641 | 178–199 | 213–234 | -12.637 | 3.94307 | 8.65138 |
| CD630_24770 | CD630_SQ1641 | 35–52 | 26–43 | -12.5498 | 3.20798 | 8.65138 |
| CD630_31210 | CD630_SQ1641 | 154–180 | 9–31 | -12.4897 | 4.17693 | 8.65138 |
| CD630_31580 | CD630_SQ1641 | 67–106 | 4–42 | -12.4569 | 3.51756 | 8.65138 |
| CD630_15190 | CD630_SQ1641 | 115–142 | 1–30 | -12.4434 | 4.43255 | 8.65138 |
| CD630_21720 | CD630_SQ1641 | 155–193 | 157–191 | -12.3441 | 4.13030 | 8.65138 |
| CD630_23090 | CD630_SQ1641 | 170–199 | 18–45 | -12.3402 | 4.25622 | 8.65138 |
| CD630_30420 | CD630_SQ1641 | 147–187 | 84–117 | -12.2694 | 5.27539 | 8.65138 |
| bglA | CD630_SQ1641 | 154–178 | 1–24 | -12.259 | 4.86402 | 8.65138 |
| CD630_05950 | CD630_SQ1641 | 121–163 | 9–46 | -12.175 | 3.31669 | 8.65138 |
| CD630_33600 | CD630_SQ1641 | 54–145 | 15–99 | -12.1501 | 3.73196 | 8.65138 |
| CD630_16450 | CD630_SQ1641 | 79–105 | 1–27 | -12.0787 | 4.57547 | 8.65138 |
| CD630_24220 | CD630_SQ1641 | 16–98 | 13–103 | -12.042 | 4.18037 | 8.65138 |
| CD630_08130 | CD630_SQ1641 | 52–86 | 5–32 | -12.003 | 4.63831 | 8.65138 |
| gcp | CD630_SQ1641 | 43–67 | 7–32 | -12.0029 | 4.92798 | 8.65138 |
| tcdC | CD630_SQ1641 | 116–142 | 1–26 | -12.002 | 3.76101 | 8.65138 |
| CD630_24830 | CD630_SQ1641 | 95–115 | 80–100 | -11.9975 | 4.98239 | 8.65138 |
| CD630_05000 | CD630_SQ1641 | 139–156 | 13–30 | -11.9907 | 4.02442 | 8.65138 |
| cobS | CD630_SQ1641 | 166–178 | 12–24 | -11.8729 | 3.67481 | 8.65138 |
| gcvTPA | CD630_SQ1641 | 133–179 | 1–33 | -11.7423 | 4.77229 | 8.65138 |
| CD630_15060 | CD630_SQ1641 | 173–198 | 24–44 | -11.7152 | 3.69396 | 8.65138 |
| CD630_23930 | CD630_SQ1641 | 110–130 | 9–29 | -11.6598 | 3.05650 | 8.65138 |
| kdgT1 | CD630_SQ1641 | 72–103 | 3–33 | -11.6012 | 4.49207 | 8.65138 |
| CD630_05220 | CD630_SQ1641 | 163–192 | 16–46 | -11.5688 | 3.65057 | 8.65138 |
| CD630_09050 | CD630_SQ1641 | 90–118 | 15–42 | -11.5043 | 3.96216 | 8.65138 |
| iorA | CD630_SQ1641 | 91–116 | 19–42 | -11.4744 | 3.83173 | 8.65138 |
| CD630_25430 | CD630_SQ1641 | 36–71 | 73–109 | -11.3615 | 4.00527 | 8.65138 |
| etfA | CD630_SQ1641 | 115–169 | 1–57 | -11.3588 | 3.41204 | 8.65138 |
| CD630_30140 | CD630_SQ1641 | 8–33 | 3–30 | -11.3286 | 3.28343 | 8.65138 |
| CD630_28181 | CD630_SQ1641 | 31–79 | 48–94 | -11.3268 | 3.91299 | 8.65138 |
| CD630_17900 | CD630_SQ1641 | 184–201 | 13–30 | -11.2528 | 3.52167 | 8.65138 |
| cdu2 | CD630_SQ1641 | 47–84 | 1–30 | -11.1811 | 2.97335 | 8.65138 |
| CD630_05230 | CD630_SQ1641 | 170–182 | 194–206 | -11.1328 | 3.73787 | 8.65138 |
| xdhA | CD630_SQ1641 | 166–194 | 3–32 | -11.0862 | 4.45147 | 8.65138 |
| CD630_31220 | CD630_SQ1641 | 179–193 | 5–23 | -11.0511 | 3.73779 | 8.65138 |
| CD630_36170 | CD630_SQ1641 | 184–201 | 13–33 | -11.0508 | 3.72380 | 8.65138 |
| CD630_30540 | CD630_SQ1641 | 146–186 | 5–42 | -11.0441 | 3.63804 | 8.65138 |
| CD630_26050 | CD630_SQ1641 | 159–192 | 71–102 | -11.0067 | 6.99851 | 8.65138 |
| CD630_32060 | CD630_SQ1641 | 162–201 | 74–105 | -11.0032 | 5.73797 | 8.65138 |
| CD630_33580 | CD630_SQ1641 | 93–115 | 1–23 | -10.9888 | 3.95778 | 8.65138 |
| eutV | CD630_SQ1641 | 124–150 | 1–24 | -10.9694 | 4.55715 | 8.65138 |
| CD630_03220 | CD630_SQ1641 | 68–118 | 1–53 | -10.9284 | 3.36746 | 8.65138 |
| aroD | CD630_SQ1641 | 160–182 | 78–100 | -10.9223 | 3.71929 | 8.65138 |
| CD630_02720 | CD630_SQ1641 | 1–48 | 158–203 | -10.8937 | 3.38938 | 8.65138 |
| etfA | CD630_SQ1641 | 66–120 | 165–208 | -10.8545 | 4.80521 | 8.65138 |
| glgP | CD630_SQ1641 | 2–30 | 1–28 | -10.8181 | 3.28530 | 8.65138 |
| CD630_01640 | CD630_SQ1641 | 84–104 | 2–22 | -10.7507 | 4.55381 | 8.65138 |
| CD630_08100 | CD630_SQ1641 | 181–197 | 3–20 | -10.7418 | 4.39181 | 8.65138 |
| CD630_06690 | CD630_SQ1641 | 25–64 | 4–42 | -10.6838 | 5.82709 | 8.65138 |
| CD630_06200 | CD630_SQ1641 | 182–199 | 1–20 | -10.6699 | 3.57064 | 8.65138 |
| CD630_10960 | CD630_SQ1641 | 6–30 | 84–109 | -10.6679 | 3.53850 | 8.65138 |
| CD630_03920 | CD630_SQ1641 | 100–135 | 11–45 | -10.6578 | 4.15934 | 8.65138 |
| CD630_08510 | CD630_SQ1641 | 44–54 | 197–207 | -10.6416 | 5.16329 | 8.65138 |
| CD630_03410 | CD630_SQ1641 | 166–178 | 9–21 | -10.6221 | 4.45994 | 8.65138 |
| CD630_28150 | CD630_SQ1641 | 29–64 | 1–44 | -10.6006 | 4.18447 | 8.65138 |
| CD630_18230 | CD630_SQ1641 | 157–197 | 83–125 | -10.5846 | 3.00184 | 8.65138 |
| CD630_13930 | CD630_SQ1641 | 1–18 | 84–101 | -10.5253 | 4.09246 | 8.65138 |
| garK | CD630_SQ1641 | 2–20 | 17–34 | -10.4897 | 3.72836 | 8.65138 |
| CD630_10080 | CD630_SQ1641 | 75–117 | 2–32 | -10.4841 | 3.75625 | 8.65138 |
| CD630_15461 | CD630_SQ1641 | 90–117 | 16–42 | -10.4754 | 9.13442 | 8.65138 |
| mobA | CD630_SQ1641 | 3–22 | 1–22 | -10.4645 | 4.25703 | 8.65138 |
| CD630_17710 | CD630_SQ1641 | 170–196 | 4–30 | -10.4198 | 4.37688 | 8.65138 |
| CD630_21330 | CD630_SQ1641 | 94–109 | 72–87 | -10.4049 | 4.10111 | 8.65138 |
| CD630_31461 | CD630_SQ1641 | 84–100 | 15–32 | -10.3979 | 4.41476 | 8.65138 |
| CD630_13600 | CD630_SQ1641 | 80–95 | 17–33 | -10.3768 | 4.45481 | 8.65138 |
| CD630_18920 | CD630_SQ1641 | 115–127 | 18–30 | -10.3664 | 4.43558 | 8.65138 |
| CD630_15431 | CD630_SQ1641 | 1–28 | 7–33 | -10.3487 | 5.29780 | 8.65138 |
| CD630_17151 | CD630_SQ1641 | 92–109 | 69–86 | -10.3331 | 6.43124 | 8.65138 |
| CD630_21700 | CD630_SQ1641 | 28–56 | 91–111 | -10.3009 | 3.96650 | 8.65138 |
| CD630_14360 | CD630_SQ1641 | 162–200 | 2–37 | -10.291 | 3.02079 | 8.65138 |
| CD630_27610 | CD630_SQ1641 | 30–53 | 3–32 | -10.2761 | 3.96465 | 8.65138 |
| CD630_11240 | CD630_SQ1641 | 4–36 | 1–32 | -10.2661 | 3.20274 | 8.65138 |
| CD630_27520 | CD630_SQ1641 | 85–142 | 144–199 | -10.266 | 4.27693 | 8.65138 |
| CD630_28260 | CD630_SQ1641 | 179–199 | 1–19 | -10.2411 | 3.51613 | 8.65138 |
| eutW | CD630_SQ1641 | 44–70 | 19–46 | -10.2399 | 4.55715 | 8.65138 |
| CD630_06110 | CD630_SQ1641 | 111–130 | 15–35 | -10.2159 | 5.61380 | 8.65138 |
| CD630_13620 | CD630_SQ1641 | 175–184 | 15–24 | -10.2101 | 4.77627 | 8.65138 |
| CD630_20260 | CD630_SQ1641 | 99–142 | 1–46 | -10.1918 | 3.84162 | 8.65138 |
| CD630_32110 | CD630_SQ1641 | 66–97 | 13–45 | -10.1601 | 4.45291 | 8.65138 |
| CD630_08110 | CD630_SQ1641 | 105–136 | 28–60 | -10.1326 | 4.39181 | 8.65138 |
| CD630_06050 | CD630_SQ1641 | 65–94 | 2–33 | -10.0896 | 3.59332 | 8.65138 |
| cynT | CD630_SQ1641 | 113–163 | 40–86 | -10.0865 | 4.21370 | 8.65138 |
| CD630_14700 | CD630_SQ1641 | 170–190 | 1–23 | -10.0789 | 3.98234 | 8.65138 |
| CD630_21570 | CD630_SQ1641 | 32–65 | 76–105 | -10.0788 | 6.48385 | 8.65138 |
| rolB | CD630_SQ1641 | 125–133 | 15–23 | -10.0451 | 3.60270 | 8.65138 |
| glgB | CD630_SQ1641 | 84–112 | 15–45 | -10.0368 | 4.48273 | 8.65138 |
| CD630_31010 | CD630_SQ1641 | 131–141 | 18–28 | -10.0109 | 4.96305 | 8.65138 |
| CD630_08370 | CD630_SQ1641 | 139–175 | 4–30 | -10.0007 | 3.65639 | 8.65138 |
| CD630_22740 | CD630_SQ1641 | 12–22 | 24–34 | -9.98405 | 3.08733 | 8.65138 |
| CD630_20980 | CD630_SQ1641 | 92–108 | 6–23 | -9.95076 | 8.12847 | 8.65138 |
| CD630_22070 | CD630_SQ1641 | 115–170 | 152–206 | -9.94564 | 4.09651 | 8.65138 |
| CD630_31530 | CD630_SQ1641 | 98–117 | 80–99 | -9.93866 | 3.49100 | 8.65138 |
| CD630_35780 | CD630_SQ1641 | 179–201 | 22–42 | -9.93396 | 3.50107 | 8.65138 |
| CD630_29840 | CD630_SQ1641 | 3–23 | 1–20 | -9.93264 | 4.48825 | 8.65138 |
| pgm | CD630_SQ1641 | 148–168 | 5–24 | -9.90694 | 3.45439 | 8.65138 |
| CD630_24590 | CD630_SQ1641 | 166–180 | 152–166 | -9.90371 | 3.70401 | 8.65138 |
| CD630_18910 | CD630_SQ1641 | 45–73 | 171–198 | -9.88349 | 3.30688 | 8.65138 |
| CD630_21620 | CD630_SQ1641 | 129–156 | 4–31 | -9.85982 | 3.02649 | 8.65138 |
| CD630_27120 | CD630_SQ1641 | 124–145 | 15–32 | -9.85967 | 4.43720 | 8.65138 |
| CD630_21030 | CD630_SQ1641 | 64–95 | 1–33 | -9.85887 | 3.08733 | 8.65138 |
| CD630_22660 | CD630_SQ1641 | 142–165 | 77–99 | -9.84437 | 3.60496 | 8.65138 |
| ldh | CD630_SQ1641 | 88–100 | 196–208 | -9.83863 | 4.35818 | 8.65138 |
| dapB2 | CD630_SQ1641 | 134–156 | 5–28 | -9.82621 | 3.27218 | 8.65138 |
| CD630_33570 | CD630_SQ1641 | 3–31 | 9–36 | -9.8071 | 3.95778 | 8.65138 |
| CD630_15360 | CD630_SQ1641 | 7–105 | 1–103 | -9.80553 | 4.47208 | 8.65138 |
| bltD | CD630_SQ1641 | 36–59 | 18–42 | -9.79379 | 4.49926 | 8.65138 |
| CD630_28550 | CD630_SQ1641 | 87–119 | 4–32 | -9.79136 | 3.51196 | 8.65138 |
| CD630_19800 | CD630_SQ1641 | 41–105 | 1–47 | -9.78824 | 5.22035 | 8.65138 |
| CD630_21730 | CD630_SQ1641 | 170–192 | 9–32 | -9.7542 | 3.45076 | 8.65138 |
| CD630_31381 | CD630_SQ1641 | 93–107 | 16–28 | -9.74077 | 3.49137 | 8.65138 |
| cheW | CD630_SQ1641 | 101–148 | 1–44 | -9.70892 | 3.72371 | 8.65138 |
| CD630_15270 | CD630_SQ1641 | 181–195 | 151–166 | -9.67649 | 4.66015 | 8.65138 |
| CD630_25320 | CD630_SQ1641 | 35–148 | 1–102 | -9.67446 | 4.29415 | 8.65138 |
| chrA | CD630_SQ1641 | 142–158 | 14–31 | -9.65385 | 5.27255 | 8.65138 |
| CD630_14150 | CD630_SQ1641 | 29–63 | 12–37 | -9.65383 | 3.28251 | 8.65138 |
| CD630_09760 | CD630_SQ1641 | 116–152 | 6–44 | -9.63639 | 4.83410 | 8.65138 |
| CD630_28910 | CD630_SQ1641 | 116–152 | 6–44 | -9.63639 | 4.51664 | 8.65138 |
| pmi | CD630_SQ1641 | 140–181 | 1–46 | -9.61002 | 3.86845 | 8.65138 |
| ogt1 | CD630_SQ1641 | 30–44 | 18–31 | -9.6096 | 3.16772 | 8.65138 |
| CD630_08460 | CD630_SQ1641 | 147–199 | 1–44 | -9.60008 | 4.15083 | 8.65138 |
| CD630_04320 | CD630_SQ1641 | 76–114 | 15–55 | -9.59996 | 3.68342 | 8.65138 |
| cobC | CD630_SQ1641 | 83–110 | 4–32 | -9.58456 | 3.15854 | 8.65138 |
| CD630_21760 | CD630_SQ1641 | 28–52 | 7–32 | -9.58317 | 4.43846 | 8.65138 |
| CD630_05490 | CD630_SQ1641 | 120–143 | 11–34 | -9.58232 | 3.89852 | 8.65138 |
| CD630_18100 | CD630_SQ1641 | 116–130 | 84–99 | -9.51422 | 5.71658 | 8.65138 |
| CD630_22991 | CD630_SQ1642 | 1–138 | 1–138 | -164.152 | 6.03728 | 8.65138 |
| ssuC | CD630_SQ1642 | 55–102 | 139–183 | -19.6844 | 3.45491 | 8.65138 |
| CD630_09970 | CD630_SQ1642 | 6–98 | 112–202 | -19.6833 | 3.25917 | 8.65138 |
| CD630_26630 | CD630_SQ1642 | 2–84 | 91–174 | -19.5296 | 4.61557 | 8.65138 |
| CD630_29750 | CD630_SQ1642 | 63–139 | 120–181 | -19.4154 | 4.59146 | 8.65138 |
| CD630_10830 | CD630_SQ1642 | 62–184 | 75–189 | -19.3975 | 4.82398 | 8.65138 |
| argS | CD630_SQ1642 | 140–176 | 162–198 | -19.34 | 4.96036 | 8.65138 |
| aroD | CD630_SQ1642 | 49–110 | 147–199 | -19.2972 | 3.71929 | 8.65138 |
| CD630_17170 | CD630_SQ1642 | 74–173 | 91–195 | -18.9122 | 4.10230 | 8.65138 |
| CD630_05230 | CD630_SQ1642 | 3–55 | 146–198 | -18.8433 | 3.73787 | 8.65138 |
| CD630_17770 | CD630_SQ1642 | 163–191 | 163–190 | -18.8392 | 5.06490 | 8.65138 |
| CD630_23280 | CD630_SQ1642 | 53–151 | 116–211 | -18.8372 | 3.50093 | 8.65138 |
| CD630_33600 | CD630_SQ1642 | 9–149 | 52–186 | -18.8052 | 3.73196 | 8.65138 |
| CD630_11241 | CD630_SQ1642 | 143–184 | 140–181 | -18.7568 | 4.85723 | 8.65138 |
| CD630_18950 | CD630_SQ1642 | 114–165 | 147–197 | -18.7467 | 3.40610 | 8.65138 |
| CD630_33420 | CD630_SQ1642 | 6–109 | 74–178 | -18.6944 | 3.22095 | 8.65138 |
| CD630_36170 | CD630_SQ1642 | 148–197 | 153–198 | -18.678 | 3.72380 | 8.65138 |
| CD630_06060 | CD630_SQ1642 | 108–152 | 174–220 | -18.4729 | 4.94050 | 8.65138 |
| CD630_19942 | CD630_SQ1642 | 67–118 | 117–185 | -18.4333 | 4.24574 | 8.65138 |
| CD630_05880 | CD630_SQ1642 | 91–162 | 114–195 | -18.377 | 4.28339 | 8.65138 |
| CD630_04940 | CD630_SQ1642 | 55–156 | 111–198 | -18.3438 | 4.20948 | 8.65138 |
| CD630_17151 | CD630_SQ1642 | 158–199 | 144–192 | -18.3001 | 6.43124 | 8.65138 |
| cspD | CD630_SQ1642 | 80–180 | 81–170 | -18.2352 | 3.23587 | 8.65138 |
| CD630_27600 | CD630_SQ1642 | 57–136 | 101–185 | -18.21 | 3.48384 | 8.65138 |
| CD630_28181 | CD630_SQ1642 | 35–78 | 150–201 | -18.1796 | 3.91299 | 8.65138 |
| CD630_22710 | CD630_SQ1642 | 3–129 | 116–223 | -18.0986 | 4.05623 | 8.65138 |
| cdu2 | CD630_SQ1642 | 57–104 | 152–199 | -18.0539 | 2.97335 | 8.65138 |
| CD630_09771 | CD630_SQ1642 | 1–138 | 1–139 | -18.0063 | 4.95362 | 8.65138 |
| srlD | CD630_SQ1642 | 88–136 | 117–164 | -17.8715 | 4.19843 | 8.65138 |
| CD630_12320 | CD630_SQ1642 | 42–114 | 122–200 | -17.8682 | 3.28667 | 8.65138 |
| pilT | CD630_SQ1642 | 59–129 | 120–197 | -17.8618 | 3.24639 | 8.65138 |
| iorA | CD630_SQ1642 | 9–91 | 118–200 | -17.7529 | 3.83173 | 8.65138 |
| CD630_22200 | CD630_SQ1642 | 84–106 | 37–61 | -17.7254 | 4.18288 | 8.65138 |
| srlEb | CD630_SQ1642 | 58–182 | 112–239 | -17.7218 | 3.50140 | 8.65138 |
| CD630_19790 | CD630_SQ1642 | 22–118 | 72–154 | -17.649 | 3.79593 | 8.65138 |
| cwp17 | CD630_SQ1642 | 53–110 | 147–197 | -17.5842 | 3.43363 | 8.65138 |
| CD630_08510 | CD630_SQ1642 | 65–110 | 142–191 | -17.3723 | 5.16329 | 8.65138 |
| CD630_05280 | CD630_SQ1642 | 146–182 | 160–196 | -17.2718 | 3.91665 | 8.65138 |
| argG | CD630_SQ1642 | 50–136 | 117–196 | -17.1313 | 4.80300 | 8.65138 |
| CD630_20670 | CD630_SQ1642 | 54–157 | 96–193 | -17.0809 | 4.42757 | 8.65138 |
| mobA | CD630_SQ1642 | 125–193 | 117–187 | -17.0508 | 4.25703 | 8.65138 |
| CD630_18090 | CD630_SQ1642 | 11–113 | 92–198 | -17.0298 | 4.23961 | 8.65138 |
| CD630_20171 | CD630_SQ1642 | 140–186 | 165–201 | -17.0068 | 6.87491 | 8.65138 |
| bcp | CD630_SQ1642 | 138–199 | 123–191 | -16.9476 | 3.16670 | 8.65138 |
| CD630_02420 | CD630_SQ1642 | 1–103 | 85–191 | -16.8433 | 3.71048 | 8.65138 |
| malY | CD630_SQ1642 | 161–186 | 161–192 | -16.8062 | 4.46345 | 8.65138 |
| CD630_08360 | CD630_SQ1642 | 30–73 | 155–197 | -16.7277 | 3.19339 | 8.65138 |
| CD630_14360 | CD630_SQ1642 | 58–113 | 150–193 | -16.7234 | 3.02079 | 8.65138 |
| CD630_32720 | CD630_SQ1642 | 35–95 | 144–213 | -16.7067 | 4.74276 | 8.65138 |
| fliI | CD630_SQ1642 | 24–81 | 117–164 | -16.7032 | 3.03666 | 8.65138 |
| CD630_31910 | CD630_SQ1642 | 9–91 | 92–174 | -16.6543 | 3.48314 | 8.65138 |
| bglA | CD630_SQ1642 | 150–198 | 148–194 | -16.6349 | 4.86402 | 8.65138 |
| CD630_04880 | CD630_SQ1642 | 49–129 | 121–212 | -16.6323 | 4.62291 | 8.65138 |
| CD630_22890 | CD630_SQ1642 | 30–141 | 71–202 | -16.6019 | 3.75591 | 8.65138 |
| CD630_17710 | CD630_SQ1642 | 18–154 | 38–169 | -16.547 | 4.37688 | 8.65138 |
| CD630_22720 | CD630_SQ1642 | 91–167 | 143–212 | -16.4491 | 3.81804 | 8.65138 |
| garK | CD630_SQ1642 | 13–42 | 170–198 | -16.4045 | 3.72836 | 8.65138 |
| CD630_20260 | CD630_SQ1642 | 119–144 | 174–198 | -16.3998 | 3.84162 | 8.65138 |
| folC | CD630_SQ1642 | 70–115 | 147–198 | -16.3493 | 3.85634 | 8.65138 |
| CD630_20960 | CD630_SQ1642 | 91–173 | 95–212 | -16.3051 | 5.65940 | 8.65138 |
| xdhA | CD630_SQ1642 | 85–129 | 147–189 | -16.2303 | 4.45147 | 8.65138 |
| CD630_07760 | CD630_SQ1642 | 70–128 | 119–188 | -16.2187 | 3.64186 | 8.65138 |
| CD630_19700 | CD630_SQ1642 | 40–97 | 140–193 | -16.1966 | 3.85861 | 8.65138 |
| gcvTPA | CD630_SQ1642 | 30–102 | 139–199 | -16.1554 | 4.77229 | 8.65138 |
| etfA | CD630_SQ1642 | 21–168 | 75–197 | -16.1528 | 3.41204 | 8.65138 |
| cspC | CD630_SQ1642 | 87–178 | 113–213 | -16.0667 | 5.48848 | 8.65138 |
| CD630_10880 | CD630_SQ1642 | 102–198 | 106–199 | -16.0636 | 5.05048 | 8.65138 |
| CD630_15360 | CD630_SQ1642 | 63–109 | 111–166 | -16.0498 | 4.47208 | 8.65138 |
| CD630_35260 | CD630_SQ1642 | 112–154 | 154–197 | -15.9909 | 5.77277 | 8.65138 |
| dnaC | CD630_SQ1642 | 89–146 | 148–214 | -15.9434 | 6.30796 | 8.65138 |
| CD630_32070 | CD630_SQ1642 | 36–92 | 140–202 | -15.9405 | 3.51964 | 8.65138 |
| CD630_18960 | CD630_SQ1642 | 30–70 | 163–202 | -15.9296 | 4.37822 | 8.65138 |
| CD630_13880 | CD630_SQ1642 | 44–75 | 168–199 | -15.8067 | 3.52735 | 8.65138 |
| eutV | CD630_SQ1642 | 21–114 | 74–188 | -15.8009 | 4.55715 | 8.65138 |
| cheW | CD630_SQ1642 | 6–101 | 78–190 | -15.7872 | 3.72371 | 8.65138 |
| CD630_19680 | CD630_SQ1642 | 40–146 | 113–223 | -15.7719 | 3.58080 | 8.65138 |
| CD630_02430 | CD630_SQ1642 | 69–118 | 117–169 | -15.7521 | 3.71048 | 8.65138 |
| CD630_31000 | CD630_SQ1642 | 52–143 | 87–180 | -15.6863 | 3.70946 | 8.65138 |
| CD630_17160 | CD630_SQ1642 | 3–119 | 72–189 | -15.6597 | 3.50228 | 8.65138 |
| cwp29 | CD630_SQ1642 | 11–60 | 143–190 | -15.644 | 4.40865 | 8.65138 |
| rolB | CD630_SQ1642 | 127–201 | 117–199 | -15.5601 | 3.60270 | 8.65138 |
| CD630_03040 | CD630_SQ1642 | 87–127 | 158–198 | -15.5498 | 3.51403 | 8.65138 |
| CD630_22550 | CD630_SQ1642 | 90–149 | 141–198 | -15.5484 | 3.62379 | 8.65138 |
| CD630_15461 | CD630_SQ1642 | 6–106 | 82–192 | -15.5354 | 9.13442 | 8.65138 |
| CD630_13920 | CD630_SQ1642 | 83–196 | 75–217 | -15.5044 | 3.25236 | 8.65138 |
| CD630_04410 | CD630_SQ1642 | 121–201 | 105–184 | -15.498 | 3.90451 | 8.65138 |
| CD630_36250 | CD630_SQ1642 | 77–201 | 90–196 | -15.4809 | 4.13224 | 8.65138 |
| CD630_32520 | CD630_SQ1642 | 86–201 | 115–216 | -15.4559 | 4.06098 | 8.65138 |
| CD630_14990 | CD630_SQ1642 | 12–78 | 147–194 | -15.4075 | 3.78619 | 8.65138 |
| CD630_14410 | CD630_SQ1642 | 33–53 | 146–164 | -15.2837 | 3.39433 | 8.65138 |
| scpB | CD630_SQ1642 | 80–118 | 164–198 | -15.2695 | 3.33560 | 8.65138 |
| spoVFB | CD630_SQ1642 | 66–196 | 75–192 | -15.2574 | 4.62758 | 8.65138 |
| CD630_23780 | CD630_SQ1642 | 83–118 | 152–184 | -15.2237 | 3.91629 | 8.65138 |
| CD630_21730 | CD630_SQ1642 | 133–190 | 144–197 | -15.195 | 3.45076 | 8.65138 |
| CD630_16460 | CD630_SQ1642 | 133–179 | 147–190 | -15.1892 | 4.64028 | 8.65138 |
| CD630_32700 | CD630_SQ1642 | 59–126 | 116–167 | -15.1801 | 4.74276 | 8.65138 |
| CD630_14250 | CD630_SQ1642 | 161–196 | 176–215 | -15.1073 | 4.19604 | 8.65138 |
| CD630_21330 | CD630_SQ1642 | 133–152 | 160–178 | -15.07 | 4.10111 | 8.65138 |
| CD630_28260 | CD630_SQ1642 | 7–108 | 80–180 | -14.9778 | 3.51613 | 8.65138 |
| cynT | CD630_SQ1642 | 111–161 | 153–200 | -14.9363 | 4.21370 | 8.65138 |
| CD630_09570 | CD630_SQ1642 | 2–44 | 174–214 | -14.9317 | 5.31213 | 8.65138 |
| CD630_29070 | CD630_SQ1642 | 2–44 | 174–214 | -14.9317 | 3.89477 | 8.65138 |
| CD630_03160 | CD630_SQ1642 | 82–105 | 153–178 | -14.8374 | 3.66946 | 8.65138 |
| CD630_21910 | CD630_SQ1642 | 72–103 | 153–195 | -14.8172 | 4.66600 | 8.65138 |
| CD630_20600 | CD630_SQ1642 | 76–123 | 140–188 | -14.765 | 4.20093 | 8.65138 |
| CD630_17292 | CD630_SQ1642 | 164–187 | 175–198 | -14.7396 | 6.25799 | 8.65138 |
| cbiQ | CD630_SQ1642 | 130–176 | 160–194 | -14.7354 | 3.79830 | 8.65138 |
| CD630_14010 | CD630_SQ1642 | 161–189 | 170–198 | -14.7227 | 3.69983 | 8.65138 |
| CD630_01590 | CD630_SQ1642 | 70–136 | 116–187 | -14.7113 | 3.45609 | 8.65138 |
| CD630_18120 | CD630_SQ1642 | 87–114 | 158–200 | -14.6728 | 3.99713 | 8.65138 |
| CD630_34560 | CD630_SQ1642 | 102–151 | 146–199 | -14.6642 | 4.02604 | 8.65138 |
| CD630_05950 | CD630_SQ1642 | 35–93 | 144–198 | -14.6478 | 3.31669 | 8.65138 |
| CD630_14050 | CD630_SQ1642 | 65–132 | 150–200 | -14.6328 | 3.30894 | 8.65138 |
| etfA | CD630_SQ1642 | 61–126 | 120–195 | -14.6259 | 4.80521 | 8.65138 |
| CD630_07770 | CD630_SQ1642 | 68–122 | 144–190 | -14.6126 | 3.31158 | 8.65138 |
| appF | CD630_SQ1642 | 40–180 | 54–186 | -14.5924 | 4.25570 | 8.65138 |
| CD630_28550 | CD630_SQ1642 | 79–115 | 155–199 | -14.592 | 3.51196 | 8.65138 |
| CD630_16150 | CD630_SQ1642 | 84–114 | 167–198 | -14.5787 | 4.20614 | 8.65138 |
| CD630_08460 | CD630_SQ1642 | 93–197 | 115–212 | -14.5776 | 4.15083 | 8.65138 |
| rbsC | CD630_SQ1642 | 30–72 | 151–194 | -14.5738 | 3.71650 | 8.65138 |
| CD630_09990 | CD630_SQ1642 | 1–138 | 89–194 | -14.5198 | 3.86040 | 8.65138 |
| cobC | CD630_SQ1642 | 79–109 | 148–184 | -14.4932 | 3.15854 | 8.65138 |
| glgP | CD630_SQ1642 | 28–81 | 140–189 | -14.4303 | 3.28530 | 8.65138 |
| ppnK | CD630_SQ1642 | 75–115 | 142–181 | -14.402 | 4.36817 | 8.65138 |
| CD630_04341 | CD630_SQ1642 | 159–192 | 140–171 | -14.3892 | 4.29262 | 8.65138 |
| pilD | CD630_SQ1642 | 19–92 | 90–163 | -14.3861 | 4.54123 | 8.65138 |
| CD630_17970 | CD630_SQ1642 | 49–127 | 89–161 | -14.3306 | 3.68802 | 8.65138 |
| pilC | CD630_SQ1642 | 11–45 | 157–190 | -14.3193 | 3.54230 | 8.65138 |
| CD630_02790 | CD630_SQ1642 | 75–103 | 142–175 | -14.3151 | 4.77661 | 8.65138 |
| CD630_14630 | CD630_SQ1642 | 68–104 | 152–194 | -14.3068 | 3.34485 | 8.65138 |
| CD630_19690 | CD630_SQ1642 | 66–153 | 117–193 | -14.3029 | 3.85861 | 8.65138 |
| CD630_32060 | CD630_SQ1642 | 28–62 | 121–158 | -14.2882 | 5.73797 | 8.65138 |
| vncR | CD630_SQ1642 | 131–157 | 41–63 | -14.2608 | 3.93642 | 8.65138 |
| cobQ | CD630_SQ1642 | 88–173 | 113–198 | -14.2497 | 3.13524 | 8.65138 |
| CD630_21620 | CD630_SQ1642 | 70–105 | 147–191 | -14.2147 | 3.02649 | 8.65138 |
| CD630_30140 | CD630_SQ1642 | 80–119 | 163–199 | -14.2063 | 3.28343 | 8.65138 |
| CD630_36190 | CD630_SQ1642 | 75–93 | 157–177 | -14.2025 | 4.63449 | 8.65138 |
| CD630_15550 | CD630_SQ1642 | 92–123 | 157–190 | -14.1933 | 3.49019 | 8.65138 |
| CD630_15850 | CD630_SQ1642 | 6–53 | 157–201 | -14.1576 | 4.32968 | 8.65138 |
| metQ | CD630_SQ1642 | 52–181 | 74–190 | -14.053 | 3.87190 | 8.65138 |
| CD630_20170 | CD630_SQ1642 | 102–151 | 164–199 | -14.0448 | 6.90201 | 8.65138 |
| CD630_08100 | CD630_SQ1642 | 160–200 | 150–193 | -14.0345 | 4.39181 | 8.65138 |
| kdpE | CD630_SQ1642 | 56–134 | 115–198 | -13.9808 | 4.27140 | 8.65138 |
| cwp26 | CD630_SQ1642 | 52–88 | 148–178 | -13.9792 | 4.44377 | 8.65138 |
| CD630_31580 | CD630_SQ1642 | 4–90 | 91–181 | -13.9761 | 3.51756 | 8.65138 |
| CD630_14700 | CD630_SQ1642 | 108–188 | 115–190 | -13.9278 | 3.98234 | 8.65138 |
| CD630_27570 | CD630_SQ1642 | 89–125 | 115–166 | -13.8823 | 3.46216 | 8.65138 |
| CD630_20450 | CD630_SQ1642 | 55–129 | 111–185 | -13.8821 | 3.77388 | 8.65138 |
| bglA | CD630_SQ1642 | 85–131 | 159–199 | -13.8652 | 4.50315 | 8.65138 |
| CD630_34520 | CD630_SQ1642 | 141–195 | 146–203 | -13.8418 | 3.83016 | 8.65138 |
| glnA | CD630_SQ1642 | 86–154 | 144–199 | -13.8381 | 3.87629 | 8.65138 |
| CD630_10220 | CD630_SQ1642 | 88–124 | 159–198 | -13.8121 | 4.08233 | 8.65138 |
| CD630_31220 | CD630_SQ1642 | 62–113 | 74–128 | -13.8095 | 3.73779 | 8.65138 |
| CD630_25320 | CD630_SQ1642 | 74–106 | 146–192 | -13.7986 | 4.29415 | 8.65138 |
| CD630_05490 | CD630_SQ1642 | 71–174 | 81–182 | -13.7827 | 3.89852 | 8.65138 |
| CD630_23000 | CD630_SQ1642 | 49–124 | 113–200 | -13.7548 | 3.54130 | 8.65138 |
| CD630_27520 | CD630_SQ1642 | 101–147 | 157–200 | -13.736 | 4.27693 | 8.65138 |
| pgm | CD630_SQ1642 | 120–175 | 91–180 | -13.5616 | 3.45439 | 8.65138 |
| acoL | CD630_SQ1642 | 77–133 | 120–190 | -13.5127 | 3.32169 | 8.65138 |
| CD630_29500 | CD630_SQ1642 | 126–176 | 152–202 | -13.4652 | 3.70644 | 8.65138 |
| CD630_09772 | CD630_SQ1642 | 70–110 | 148–192 | -13.4456 | 3.66033 | 8.65138 |
| ogt1 | CD630_SQ1642 | 1–100 | 71–173 | -13.4061 | 3.16772 | 8.65138 |
| CD630_18400 | CD630_SQ1642 | 108–123 | 163–178 | -13.3848 | 4.67962 | 8.65138 |

| | | | | | | |
|---|---|---|---|---|---|---|
| CD630_16160 | CD630_SQ1642 | 65 -- 110 | 145 -- 186 | -13.3842 | 6.11801 | 8.65138 |
| CD630_21790 | CD630_SQ1642 | 54 -- 102 | 146 -- 198 | -13.3814 | 4.93762 | 8.65138 |
| CD630_19580 | CD630_SQ1642 | 63 -- 98 | 155 -- 190 | -13.3486 | 4.51902 | 8.65138 |
| CD630_21770 | CD630_SQ1642 | 46 -- 93 | 157 -- 200 | -13.3227 | 4.02587 | 8.65138 |
| CD630_02740 | CD630_SQ1642 | 158 -- 175 | 151 -- 169 | -13.2952 | 5.02277 | 8.65138 |
| CD630_31490 | CD630_SQ1642 | 73 -- 146 | 147 -- 210 | -13.2727 | 4.08593 | 8.65138 |
| CD630_13560 | CD630_SQ1642 | 2 -- 26 | 173 -- 198 | -13.2658 | 3.38083 | 8.65138 |
| CD630_17380 | CD630_SQ1642 | 35 -- 88 | 116 -- 178 | -13.2576 | 5.03207 | 8.65138 |
| CD630_25300 | CD630_SQ1642 | 52 -- 78 | 73 -- 100 | -13.244 | 4.49333 | 8.65138 |
| kdpD | CD630_SQ1642 | 131 -- 194 | 142 -- 198 | -13.2417 | 4.27140 | 8.65138 |
| CD630_07480 | CD630_SQ1642 | 87 -- 112 | 147 -- 171 | -13.2171 | 5.18321 | 8.65138 |
| CD630_32710 | CD630_SQ1642 | 113 -- 156 | 157 -- 195 | -13.1626 | 4.74276 | 8.65138 |
| CD630_10950 | CD630_SQ1642 | 78 -- 111 | 154 -- 197 | -13.1534 | 3.67675 | 8.65138 |
| CD630_13400 | CD630_SQ1642 | 90 -- 120 | 181 -- 212 | -13.1416 | 3.20232 | 8.65138 |
| CD630_08370 | CD630_SQ1642 | 123 -- 170 | 156 -- 198 | -13.1061 | 3.65639 | 8.65138 |
| CD630_30540 | CD630_SQ1642 | 162 -- 187 | 175 -- 200 | -13.0954 | 3.63804 | 8.65138 |
| CD630_29760 | CD630_SQ1642 | 141 -- 185 | 168 -- 197 | -13.0892 | 3.51074 | 8.65138 |
| CD630_06080 | CD630_SQ1642 | 38 -- 112 | 140 -- 214 | -13.0832 | 3.82315 | 8.65138 |
| CD630_16450 | CD630_SQ1642 | 80 -- 117 | 149 -- 186 | -13.0533 | 4.57547 | 8.65138 |
| CD630_06340 | CD630_SQ1642 | 88 -- 113 | 140 -- 164 | -13.0359 | 4.35983 | 8.65138 |
| rolA | CD630_SQ1642 | 32 -- 76 | 157 -- 199 | -13.0288 | 4.56362 | 8.65138 |
| CD630_23820 | CD630_SQ1642 | 31 -- 95 | 94 -- 153 | -12.9562 | 3.18419 | 8.65138 |
| cdu1 | CD630_SQ1642 | 1 -- 120 | 70 -- 179 | -12.9516 | 4.46276 | 8.65138 |
| CD630_08660 | CD630_SQ1642 | 62 -- 90 | 153 -- 191 | -12.9485 | 3.86467 | 8.65138 |
| oppB | CD630_SQ1642 | 90 -- 112 | 74 -- 99 | -12.945 | 2.98328 | 8.65138 |
| CD630_09040 | CD630_SQ1642 | 122 -- 185 | 120 -- 180 | -12.9333 | 4.11851 | 8.65138 |
| CD630_23080 | CD630_SQ1642 | 157 -- 194 | 157 -- 190 | -12.9269 | 5.23027 | 8.65138 |
| glgB | CD630_SQ1642 | 86 -- 111 | 170 -- 199 | -12.9115 | 4.48273 | 8.65138 |
| thiH | CD630_SQ1642 | 166 -- 199 | 173 -- 202 | -12.9025 | 3.58230 | 8.65138 |
| CD630_14920 | CD630_SQ1642 | 91 -- 139 | 146 -- 199 | -12.8828 | 4.35225 | 8.65138 |
| CD630_04320 | CD630_SQ1642 | 2 -- 41 | 148 -- 190 | -12.8802 | 3.68342 | 8.65138 |
| CD630_31382 | CD630_SQ1642 | 5 -- 15 | 113 -- 123 | -12.8757 | 3.49137 | 8.65138 |
| CD630_06050 | CD630_SQ1642 | 113 -- 201 | 148 -- 220 | -12.8629 | 3.59332 | 8.65138 |
| CD630_14930 | CD630_SQ1642 | 170 -- 191 | 177 -- 197 | -12.8455 | 3.31335 | 8.65138 |
| CD630_20460 | CD630_SQ1642 | 16 -- 105 | 92 -- 192 | -12.832 | 7.94034 | 8.65138 |
| CD630_03420 | CD630_SQ1642 | 115 -- 194 | 118 -- 196 | -12.7975 | 4.71868 | 8.65138 |
| CD630_30110 | CD630_SQ1642 | 84 -- 121 | 153 -- 198 | -12.7973 | 3.31041 | 8.65138 |
| ung | CD630_SQ1642 | 78 -- 134 | 143 -- 201 | -12.7831 | 3.30908 | 8.65138 |
| CD630_03220 | CD630_SQ1642 | 41 -- 136 | 113 -- 199 | -12.7381 | 3.36746 | 8.65138 |
| CD630_25620 | CD630_SQ1642 | 127 -- 155 | 170 -- 199 | -12.6978 | 3.02685 | 8.65138 |
| adhE | CD630_SQ1642 | 73 -- 96 | 170 -- 193 | -12.6837 | 3.06485 | 8.65138 |
| CD630_18680 | CD630_SQ1642 | 47 -- 121 | 75 -- 128 | -12.6706 | 3.04315 | 8.65138 |
| CD630_18910 | CD630_SQ1642 | 165 -- 193 | 38 -- 65 | -12.6703 | 3.30688 | 8.65138 |
| tcdC | CD630_SQ1642 | 112 -- 162 | 147 -- 197 | -12.6236 | 3.76101 | 8.65138 |
| CD630_04770 | CD630_SQ1642 | 31 -- 69 | 142 -- 175 | -12.5482 | 3.58478 | 8.65138 |
| serS1 | CD630_SQ1642 | 125 -- 172 | 157 -- 198 | -12.5419 | 3.45858 | 8.65138 |
| CD630_13780 | CD630_SQ1642 | 86 -- 143 | 122 -- 182 | -12.535 | 3.52274 | 8.65138 |
| CD630_19670 | CD630_SQ1642 | 150 -- 176 | 158 -- 183 | -12.4422 | 3.20718 | 8.65138 |
| CD630_26150 | CD630_SQ1642 | 163 -- 171 | 87 -- 95 | -12.3657 | 4.06302 | 8.65138 |
| CD630_08090 | CD630_SQ1642 | 142 -- 191 | 147 -- 199 | -12.3529 | 3.57167 | 8.65138 |
| CD630_21230 | CD630_SQ1642 | 61 -- 110 | 155 -- 200 | -12.3461 | 3.89213 | 8.65138 |
| nusB | CD630_SQ1642 | 89 -- 152 | 134 -- 190 | -12.3076 | 4.92798 | 8.65138 |
| CD630_18650 | CD630_SQ1642 | 82 -- 131 | 145 -- 198 | -12.2701 | 3.75892 | 8.65138 |
| CD630_27770 | CD630_SQ1642 | 84 -- 107 | 174 -- 197 | -12.2538 | 4.08588 | 8.65138 |
| CD630_22050 | CD630_SQ1642 | 81 -- 103 | 153 -- 180 | -12.2013 | 4.87639 | 8.65138 |
| opuCA | CD630_SQ1642 | 126 -- 140 | 183 -- 197 | -12.1982 | 4.07035 | 8.65138 |
| CD630_28320 | CD630_SQ1642 | 47 -- 93 | 116 -- 171 | -12.182 | 3.99184 | 8.65138 |
| CD630_23440 | CD630_SQ1642 | 2 -- 19 | 154 -- 171 | -12.1393 | 3.34841 | 8.65138 |
| CD630_08130 | CD630_SQ1642 | 73 -- 103 | 153 -- 183 | -12.1365 | 4.63831 | 8.65138 |
| CD630_29250 | CD630_SQ1642 | 149 -- 163 | 187 -- 201 | -12.0952 | 4.61901 | 8.65138 |
| CD630_09410 | CD630_SQ1642 | 149 -- 163 | 187 -- 201 | -12.0952 | 4.75651 | 8.65138 |
| CD630_33480 | CD630_SQ1642 | 71 -- 111 | 143 -- 188 | -12.0876 | 3.67613 | 8.65138 |
| CD630_09760 | CD630_SQ1642 | 93 -- 145 | 153 -- 198 | -12.0537 | 4.83410 | 8.65138 |
| CD630_28910 | CD630_SQ1642 | 93 -- 145 | 153 -- 198 | -12.0537 | 4.51664 | 8.65138 |
| CD630_30310 | CD630_SQ1642 | 183 -- 201 | 157 -- 175 | -12.0507 | 4.56059 | 8.65138 |
| CD630_15200 | CD630_SQ1642 | 66 -- 97 | 90 -- 127 | -12.0147 | 4.00442 | 8.65138 |
| CD630_01570 | CD630_SQ1642 | 150 -- 165 | 173 -- 188 | -12.0047 | 6.08699 | 8.65138 |
| CD630_21140 | CD630_SQ1642 | 76 -- 98 | 166 -- 187 | -11.9742 | 3.33304 | 8.65138 |
| pilW | CD630_SQ1642 | 38 -- 99 | 155 -- 197 | -11.9583 | 4.96316 | 8.65138 |
| CD630_21570 | CD630_SQ1642 | 86 -- 109 | 74 -- 99 | -11.7965 | 6.48385 | 8.65138 |
| CD630_05211 | CD630_SQ1642 | 88 -- 152 | 137 -- 199 | -11.7952 | 3.97048 | 8.65138 |
| CD630_18100 | CD630_SQ1642 | 127 -- 164 | 153 -- 190 | -11.7935 | 5.71658 | 8.65138 |
| CD630_32050 | CD630_SQ1642 | 43 -- 58 | 180 -- 195 | -11.755 | 5.73797 | 8.65138 |
| CD630_10000 | CD630_SQ1642 | 77 -- 160 | 155 -- 239 | -11.732 | 3.86040 | 8.65138 |
| CD630_05590 | CD630_SQ1642 | 64 -- 121 | 137 -- 190 | -11.7298 | 3.39670 | 8.65138 |
| CD630_16550 | CD630_SQ1642 | 76 -- 107 | 147 -- 183 | -11.7244 | 5.61064 | 8.65138 |
| CD630_15060 | CD630_SQ1642 | 72 -- 108 | 158 -- 192 | -11.7217 | 3.69396 | 8.65138 |
| CD630_27230 | CD630_SQ1642 | 129 -- 147 | 1 -- 20 | -11.713 | 3.52462 | 8.65138 |
| CD630_28580 | CD630_SQ1642 | 4 -- 28 | 134 -- 159 | -11.6954 | 4.67130 | 8.65138 |
| CD630_14240 | CD630_SQ1642 | 68 -- 128 | 83 -- 139 | -11.6942 | 8.60766 | 8.65138 |
| CD630_30360 | CD630_SQ1642 | 43 -- 129 | 112 -- 198 | -11.6688 | 4.26509 | 8.65138 |
| CD630_06560 | CD630_SQ1642 | 63 -- 116 | 144 -- 197 | -11.6557 | 4.02532 | 8.65138 |
| panG | CD630_SQ1642 | 9 -- 33 | 165 -- 188 | -11.5773 | 5.54047 | 8.65138 |
| CD630_33570 | CD630_SQ1642 | 132 -- 172 | 161 -- 200 | -11.5332 | 3.95778 | 8.65138 |
| CD630_29270 | CD630_SQ1642 | 43 -- 103 | 153 -- 198 | -11.4986 | 3.70174 | 8.65138 |
| uxaA | CD630_SQ1642 | 75 -- 94 | 157 -- 183 | -11.475 | 4.26165 | 8.65138 |
| CD630_22030 | CD630_SQ1642 | 66 -- 104 | 152 -- 188 | -11.4551 | 3.29934 | 8.65138 |
| CD630_18930 | CD630_SQ1642 | 188 -- 201 | 188 -- 201 | -11.4366 | 5.32127 | 8.65138 |
| bltD | CD630_SQ1642 | 56 -- 144 | 120 -- 199 | -11.3615 | 4.49926 | 8.65138 |
| CD630_21110 | CD630_SQ1642 | 89 -- 118 | 168 -- 197 | -11.3454 | 3.89630 | 8.65138 |
| CD630_21700 | CD630_SQ1642 | 164 -- 194 | 146 -- 177 | -11.3223 | 3.96650 | 8.65138 |
| pmi | CD630_SQ1642 | 65 -- 103 | 142 -- 185 | -11.2712 | 3.86845 | 8.65138 |
| CD630_13360 | CD630_SQ1642 | 88 -- 141 | 159 -- 199 | -11.2709 | 3.77330 | 8.65138 |
| CD630_20660 | CD630_SQ1642 | 115 -- 159 | 148 -- 191 | -11.2521 | 5.39321 | 8.65138 |
| CD630_14420 | CD630_SQ1642 | 143 -- 177 | 163 -- 200 | -11.2502 | 3.99497 | 8.65138 |
| CD630_14710 | CD630_SQ1642 | 77 -- 108 | 157 -- 188 | -11.2237 | 3.65171 | 8.65138 |
| CD630_31210 | CD630_SQ1642 | 42 -- 105 | 133 -- 194 | -11.1909 | 4.17693 | 8.65138 |
| tcdA | CD630_SQ1642 | 9 -- 60 | 147 -- 197 | -11.1849 | 4.70593 | 8.65138 |
| CD630_35840 | CD630_SQ1642 | 62 -- 92 | 166 -- 199 | -11.1833 | 3.14170 | 8.65138 |
| nrdD | CD630_SQ1642 | 51 -- 91 | 118 -- 169 | -11.1777 | 3.28290 | 8.65138 |
| ldh | CD630_SQ1642 | 182 -- 191 | 85 -- 94 | -11.1675 | 4.35818 | 8.65138 |
| CD630_07430 | CD630_SQ1642 | 111 -- 128 | 148 -- 164 | -11.1305 | 4.65568 | 8.65138 |
| CD630_25040 | CD630_SQ1642 | 7 -- 55 | 152 -- 197 | -11.1301 | 3.43898 | 8.65138 |
| CD630_27240 | CD630_SQ1642 | 3 -- 29 | 144 -- 183 | -11.0858 | 3.52462 | 8.65138 |
| CD630_30420 | CD630_SQ1642 | 59 -- 93 | 118 -- 156 | -11.0761 | 5.27539 | 8.65138 |
| panB | CD630_SQ1642 | 25 -- 72 | 147 -- 198 | -11.0606 | 3.40248 | 8.65138 |
| CD630_17730 | CD630_SQ1642 | 85 -- 107 | 174 -- 196 | -11.0486 | 4.73397 | 8.65138 |
| CD630_18940 | CD630_SQ1642 | 117 -- 166 | 139 -- 203 | -11.0335 | 4.09988 | 8.65138 |
| CD630_19800 | CD630_SQ1642 | 102 -- 137 | 166 -- 199 | -11.0292 | 5.22035 | 8.65138 |
| CD630_06140 | CD630_SQ1642 | 85 -- 123 | 90 -- 126 | -11.023 | 4.12519 | 8.65138 |
| CD630_14182 | CD630_SQ1642 | 15 -- 133 | 6 -- 127 | -100.288 | 4.82360 | 8.65138 |
| CD630_18780 | CD630_SQ1642 | 138 -- 181 | 119 -- 164 | -10.9714 | 4.71613 | 8.65138 |
| scrR | CD630_SQ1642 | 9 -- 33 | 155 -- 180 | -10.9323 | 3.49166 | 8.65138 |
| CD630_19921 | CD630_SQ1642 | 43 -- 104 | 139 -- 199 | -10.9274 | 5.46345 | 8.65138 |
| CD630_31880 | CD630_SQ1642 | 125 -- 170 | 153 -- 198 | -10.9148 | 3.97522 | 8.65138 |
| CD630_19780 | CD630_SQ1642 | 15 -- 32 | 148 -- 166 | -10.9116 | 3.64830 | 8.65138 |
| CD630_20650 | CD630_SQ1642 | 152 -- 160 | 87 -- 95 | -10.9019 | 4.00083 | 8.65138 |
| CD630_01560 | CD630_SQ1642 | 35 -- 91 | 157 -- 199 | -10.8825 | 4.06352 | 8.65138 |
| CD630_27210 | CD630_SQ1642 | 21 -- 44 | 168 -- 191 | -10.8776 | 4.18786 | 8.65138 |
| rbsA | CD630_SQ1642 | 46 -- 74 | 157 -- 186 | -10.8632 | 3.71650 | 8.65138 |
| CD630_15900 | CD630_SQ1642 | 166 -- 175 | 86 -- 95 | -10.8563 | 3.17664 | 8.65138 |
| CD630_25150 | CD630_SQ1642 | 102 -- 148 | 155 -- 199 | -10.828 | 3.79339 | 8.65138 |
| flgB | CD630_SQ1642 | 66 -- 103 | 153 -- 195 | -10.8052 | 3.25736 | 8.65138 |
| CD630_23930 | CD630_SQ1642 | 115 -- 156 | 150 -- 199 | -10.8012 | 3.05650 | 8.65138 |
| CD630_06350 | CD630_SQ1642 | 82 -- 124 | 154 -- 197 | -10.7839 | 4.35983 | 8.65138 |
| CD630_17960 | CD630_SQ1642 | 80 -- 106 | 146 -- 172 | -10.7635 | 3.71404 | 8.65138 |
| dltA | CD630_SQ1642 | 108 -- 166 | 147 -- 203 | -10.7491 | 3.85429 | 8.65138 |
| CD630_18311 | CD630_SQ1642 | 82 -- 108 | 173 -- 196 | -10.7101 | 6.86047 | 8.65138 |
| CD630_06540 | CD630_SQ1642 | 70 -- 199 | 93 -- 209 | -10.7006 | 3.54673 | 8.65138 |
| CD630_27120 | CD630_SQ1642 | 134 -- 154 | 185 -- 204 | -10.658 | 4.43720 | 8.65138 |
| CD630_28290 | CD630_SQ1642 | 85 -- 95 | 114 -- 124 | -10.6545 | 5.54192 | 8.65138 |
| CD630_24770 | CD630_SQ1642 | 36 -- 105 | 138 -- 211 | -10.6516 | 3.20798 | 8.65138 |
| CD630_11220 | CD630_SQ1642 | 88 -- 128 | 11 -- 52 | -10.6346 | 3.52327 | 8.65138 |
| CD630_27610 | CD630_SQ1642 | 82 -- 126 | 57 -- 102 | -10.5932 | 3.96465 | 8.65138 |
| CD630_09050 | CD630_SQ1642 | 67 -- 103 | 164 -- 195 | -10.5905 | 3.96216 | 8.65138 |
| CD630_06060 | CD630_SQ1642 | 54 -- 112 | 146 -- 201 | -10.5858 | 4.94050 | 8.65138 |
| CD630_12110 | CD630_SQ1642 | 125 -- 164 | 170 -- 204 | -10.5682 | 4.36078 | 8.65138 |
| CD630_08390 | CD630_SQ1642 | 75 -- 114 | 8 -- 44 | -10.5624 | 5.16000 | 8.65138 |
| CD630_20290 | CD630_SQ1642 | 47 -- 90 | 91 -- 147 | -10.5558 | 4.38183 | 8.65138 |
| CD630_34530 | CD630_SQ1642 | 37 -- 50 | 177 -- 190 | -10.55 | 3.83016 | 8.65138 |
| CD630_36690 | CD630_SQ1642 | 85 -- 140 | 136 -- 198 | -10.5075 | 3.53761 | 8.65138 |
| CD630_06970 | CD630_SQ1642 | 87 -- 133 | 159 -- 198 | -10.5072 | 3.82312 | 8.65138 |
| CD630_24920 | CD630_SQ1642 | 19 -- 58 | 140 -- 172 | -10.5023 | 3.89455 | 8.65138 |
| CD630_05150 | CD630_SQ1642 | 81 -- 115 | 155 -- 185 | -10.4851 | 3.69875 | 8.65138 |
| CD630_26050 | CD630_SQ1642 | 170 -- 176 | 91 -- 97 | -10.4814 | 6.99851 | 8.65138 |
| gatC | CD630_SQ1642 | 58 -- 110 | 156 -- 201 | -10.4796 | 4.25143 | 8.65138 |
| moeA | CD630_SQ1642 | 102 -- 144 | 152 -- 199 | -10.4727 | 3.26118 | 8.65138 |
| cbpA | CD630_SQ1642 | 86 -- 105 | 38 -- 54 | -10.4662 | 3.03040 | 8.65138 |
| CD630_28150 | CD630_SQ1642 | 154 -- 194 | 147 -- 179 | -10.4541 | 4.18447 | 8.65138 |
| CD630_24830 | CD630_SQ1642 | 171 -- 178 | 87 -- 94 | -10.4388 | 4.98239 | 8.65138 |
| CD630_30100 | CD630_SQ1642 | 74 -- 90 | 173 -- 189 | -10.4008 | 4.13234 | 8.65138 |
| CD630_21030 | CD630_SQ1642 | 78 -- 108 | 174 -- 199 | -10.3791 | 3.08733 | 8.65138 |
| CD630_08070 | CD630_SQ1642 | 155 -- 172 | 144 -- 162 | -10.3773 | 4.87819 | 8.65138 |
| CD630_27310 | CD630_SQ1642 | 78 -- 107 | 174 -- 198 | -10.3584 | 4.01622 | 8.65138 |
| virS | CD630_SQ1642 | 57 -- 106 | 132 -- 195 | -10.3579 | 4.74085 | 8.65138 |
| CD630_21490 | CD630_SQ1642 | 75 -- 83 | 4 -- 12 | -10.3448 | 3.37462 | 8.65138 |
| CD630_20980 | CD630_SQ1642 | 94 -- 109 | 6 -- 21 | -10.3368 | 8.12847 | 8.65138 |
| CD630_21760 | CD630_SQ1642 | 3 -- 40 | 155 -- 190 | -10.3343 | 4.43846 | 8.65138 |
| CD630_15740 | CD630_SQ1642 | 65 -- 104 | 150 -- 187 | -10.3287 | 4.43222 | 8.65138 |
| CD630_26691 | CD630_SQ1642 | 168 -- 175 | 90 -- 97 | -10.3165 | 3.96230 | 8.65138 |
| CD630_22950 | CD630_SQ1642 | 38 -- 113 | 91 -- 169 | -10.2853 | 3.96883 | 8.65138 |
| CD630_01690 | CD630_SQ1642 | 62 -- 85 | 150 -- 169 | -10.2801 | 4.97234 | 8.65138 |
| CD630_31500 | CD630_SQ1642 | 69 -- 104 | 152 -- 202 | -10.2552 | 4.90207 | 8.65138 |
| CD630_18610 | CD630_SQ1642 | 14 -- 93 | 115 -- 183 | -10.2386 | 4.15582 | 8.65138 |
| ribD | CD630_SQ1642 | 88 -- 142 | 153 -- 212 | -10.2179 | 6.65922 | 8.65138 |
| CD630_31381 | CD630_SQ1642 | 160 -- 195 | 162 -- 198 | -10.2075 | 3.49137 | 8.65138 |
| CD630_13930 | CD630_SQ1642 | 121 -- 143 | 165 -- 187 | -10.2034 | 4.09246 | 8.65138 |
| CD630_34030 | CD630_SQ1642 | 62 -- 78 | 154 -- 171 | -10.2034 | 3.05766 | 8.65138 |
| CD630_14810 | CD630_SQ1642 | 138 -- 154 | 111 -- 127 | -10.1916 | 3.35323 | 8.65138 |
| gabT | CD630_SQ1642 | 108 -- 156 | 152 -- 199 | -10.1674 | 3.67067 | 8.65138 |
| CD630_21720 | CD630_SQ1642 | 89 -- 140 | 160 -- 212 | -10.1643 | 4.13030 | 8.65138 |
| CD630_19904 | CD630_SQ1642 | 13 -- 32 | 63 -- 83 | -10.1471 | 5.96526 | 8.65138 |
| CD630_13600 | CD630_SQ1642 | 120 -- 174 | 149 -- 198 | -10.1406 | 4.45481 | 8.65138 |
| CD630_18781 | CD630_SQ1642 | 51 -- 112 | 147 -- 203 | -10.1007 | 3.77660 | 8.65138 |
| CD630_24580 | CD630_SQ1642 | 85 -- 108 | 31 -- 56 | -10.05 | 3.88519 | 8.65138 |
| CD630_25200 | CD630_SQ1642 | 57 -- 97 | 151 -- 209 | -10.0213 | 3.65158 | 8.65138 |
| CD630_25190 | CD630_SQ1642 | 167 -- 199 | 172 -- 198 | -10.0108 | 3.19663 | 8.65138 |
| CD630_31960 | CD630_SQ1642 | 72 -- 84 | 188 -- 200 | -9.97076 | 4.93757 | 8.65138 |
| CD630_23760 | CD630_SQ1642 | 40 -- 56 | 173 -- 189 | -9.96739 | 4.54307 | 8.65138 |
| opuCC | CD630_SQ1642 | 93 -- 120 | 25 -- 52 | -9.96124 | 4.07035 | 8.65138 |
| CD630_26680 | CD630_SQ1642 | 49 -- 95 | 154 -- 213 | -9.92182 | 3.53956 | 8.65138 |
| CD630_05480 | CD630_SQ1642 | 167 -- 190 | 72 -- 99 | -9.92176 | 4.27854 | 8.65138 |
| CD630_05870 | CD630_SQ1642 | 143 -- 197 | 144 -- 198 | -9.89384 | 4.28339 | 8.65138 |
| CD630_30130 | CD630_SQ1642 | 3 -- 28 | 171 -- 200 | -9.87372 | 3.28343 | 8.65138 |
| CD630_05840 | CD630_SQ1642 | 88 -- 98 | 113 -- 123 | -9.87138 | 4.28779 | 8.65138 |
| eutP | CD630_SQ1642 | 95 -- 128 | 157 -- 186 | -9.84677 | 3.82847 | 8.65138 |
| CD630_26020 | CD630_SQ1642 | 5 -- 42 | 150 -- 180 | -9.81907 | 4.03155 | 8.65138 |
| CD630_12650 | CD630_SQ1642 | 99 -- 143 | 91 -- 144 | -9.77594 | 6.31827 | 8.65138 |
| hfq | CD630_SQ1642 | 16 -- 57 | 168 -- 202 | -9.76394 | 5.05609 | 8.65138 |
| CD630_15270 | CD630_SQ1642 | 89 -- 119 | 73 -- 121 | -9.75078 | 4.66015 | 8.65138 |
| CD630_15150 | CD630_SQ1642 | 1 -- 28 | 142 -- 169 | -9.73989 | 3.35564 | 8.65138 |
| CD630_02720 | CD630_SQ1642 | 121 -- 148 | 158 -- 187 | -9.72641 | 3.38938 | 8.65138 |
| CD630_18410 | CD630_SQ1642 | 43 -- 65 | 178 -- 200 | -9.72622 | 3.59739 | 8.65138 |
| CD630_21020 | CD630_SQ1642 | 85 -- 105 | 34 -- 53 | -9.69868 | 4.28666 | 8.65138 |
| gcp | CD630_SQ1642 | 95 -- 130 | 168 -- 203 | -9.67903 | 4.92798 | 8.65138 |
| CD630_21460 | CD630_SQ1642 | 114 -- 158 | 157 -- 197 | -9.67008 | 4.83005 | 8.65138 |
| CD630_03920 | CD630_SQ1642 | 82 -- 103 | 85 -- 106 | -9.64728 | 4.15934 | 8.65138 |
| CD630_07840 | CD630_SQ1642 | 3 -- 25 | 171 -- 196 | -9.62921 | 4.26761 | 8.65138 |
| CD630_05370 | CD630_SQ1642 | 88 -- 110 | 155 -- 178 | -9.62544 | 3.26947 | 8.65138 |
| CD630_14110 | CD630_SQ1642 | 141 -- 196 | 139 -- 200 | -9.6107 | 3.39550 | 8.65138 |
| CD630_05450 | CD630_SQ1642 | 86 -- 102 | 39 -- 54 | -9.58282 | 5.04761 | 8.65138 |
| CD630_05290 | CD630_SQ1642 | 68 -- 104 | 215 -- 249 | -9.56127 | 3.97604 | 8.65138 |
| CD630_19801 | CD630_SQ1642 | 144 -- 173 | 150 -- 183 | -9.55733 | 5.22035 | 8.65138 |
| CD630_33560 | CD630_SQ1642 | 66 -- 117 | 148 -- 197 | -9.55698 | 4.03147 | 8.65138 |
| spaK | CD630_SQ1642 | 72 -- 93 | 162 -- 185 | -9.54404 | 3.04530 | 8.65138 |
| CD630_14210 | CD630_SQ1642 | 71 -- 98 | 155 -- 183 | -9.5369 | 3.82355 | 8.65138 |
| CD630_07470 | CD630_SQ1642 | 84 -- 115 | 169 -- 200 | -9.52927 | 3.37143 | 8.65138 |
| agrB | CD630_SQ1642 | 91 -- 151 | 130 -- 189 | -9.50693 | 3.92598 | 8.65138 |
| CD630_32880 | CD630_SQ1642 | 90 -- 116 | 160 -- 188 | -9.50493 | 3.41276 | 8.65138 |
| CD630_32520 | CD630_SQ1656 | 152 -- 198 | 8 -- 58 | -18.2755 | 4.06098 | 4.25622 |
| CD630_17360 | CD630_SQ1656 | 98 -- 140 | 5 -- 42 | -17.2864 | 5.66070 | 4.25622 |
| CD630_22740 | CD630_SQ1656 | 163 -- 194 | 7 -- 37 | -15.8997 | 3.08733 | 4.25622 |
| tcdC | CD630_SQ1656 | 117 -- 157 | 4 -- 39 | -15.4546 | 3.76101 | 4.25622 |
| CD630_19790 | CD630_SQ1656 | 49 -- 117 | 6 -- 84 | -15.3615 | 3.79593 | 4.25622 |
| CD630_09771 | CD630_SQ1656 | 155 -- 187 | 1 -- 31 | -15.2466 | 4.95362 | 4.25622 |
| CD630_21330 | CD630_SQ1656 | 48 -- 93 | 8 -- 64 | -14.8038 | 4.10111 | 4.25622 |
| CD630_23820 | CD630_SQ1656 | 152 -- 172 | 62 -- 83 | -14.7541 | 3.18419 | 4.25622 |
| CD630_09060 | CD630_SQ1656 | 6 -- 135 | 9 -- 120 | -14.6726 | 4.31834 | 4.25622 |
| CD630_30140 | CD630_SQ1656 | 67 -- 117 | 24 -- 75 | -14.6414 | 3.28343 | 4.25622 |
| CD630_14182 | CD630_SQ1656 | 126 -- 196 | 1 -- 74 | -14.6145 | 4.82360 | 4.25622 |
| CD630_24760 | CD630_SQ1656 | 98 -- 139 | 7 -- 42 | -14.5509 | 5.19485 | 4.25622 |
| etfA | CD630_SQ1656 | 9 -- 62 | 47 -- 102 | -14.5339 | 3.41204 | 4.25622 |
| CD630_28890 | CD630_SQ1656 | 158 -- 187 | 1 -- 28 | -14.3149 | 7.52469 | 4.25622 |
| CD630_18960 | CD630_SQ1656 | 40 -- 96 | 16 -- 67 | -14.2988 | 4.37822 | 4.25622 |
| CD630_24591 | CD630_SQ1656 | 14 -- 48 | 4 -- 38 | -14.2367 | 3.54497 | 4.25622 |
| mobA | CD630_SQ1656 | 118 -- 150 | 16 -- 44 | -13.9372 | 4.25703 | 4.25622 |
| CD630_08370 | CD630_SQ1656 | 145 -- 195 | 1 -- 37 | -13.9153 | 3.65639 | 4.25622 |
| CD630_05821 | CD630_SQ1656 | 96 -- 115 | 5 -- 25 | -13.6974 | 3.75661 | 4.25622 |
| CD630_19700 | CD630_SQ1656 | 9 -- 52 | 10 -- 53 | -13.6793 | 3.85861 | 4.25622 |
| CD630_20450 | CD630_SQ1656 | 128 -- 165 | 192 -- 229 | -13.6544 | 3.77388 | 4.25622 |
| CD630_21260 | CD630_SQ1656 | 138 -- 152 | 150 -- 164 | -13.3402 | 4.22255 | 4.25622 |
| CD630_22991 | CD630_SQ1656 | 152 -- 196 | 1 -- 47 | -13.1986 | 6.03728 | 4.25622 |
| tcdA | CD630_SQ1656 | 58 -- 166 | 11 -- 124 | -13.1452 | 4.70593 | 4.25622 |
| CD630_31960 | CD630_SQ1656 | 62 -- 90 | 2 -- 36 | -13.1218 | 4.93757 | 4.25622 |
| CD630_14050 | CD630_SQ1656 | 53 -- 137 | 6 -- 80 | -12.9636 | 3.30894 | 4.25622 |
| CD630_20260 | CD630_SQ1656 | 99 -- 131 | 9 -- 41 | -12.8005 | 3.84162 | 4.25622 |
| CD630_18230 | CD630_SQ1656 | 1 -- 36 | 7 -- 38 | -12.7393 | 3.00184 | 4.25622 |
| CD630_34560 | CD630_SQ1656 | 71 -- 153 | 7 -- 80 | -12.7154 | 4.02604 | 4.25622 |
| CD630_17770 | CD630_SQ1656 | 88 -- 133 | 4 -- 49 | -12.6576 | 5.06490 | 4.25622 |
| CD630_16450 | CD630_SQ1656 | 4 -- 23 | 13 -- 32 | -12.6493 | 4.57547 | 4.25622 |
| CD630_26050 | CD630_SQ1656 | 125 -- 157 | 16 -- 49 | -12.6402 | 6.99851 | 4.25622 |
| CD630_02420 | CD630_SQ1656 | 9 -- 50 | 154 -- 193 | -12.542 | 3.71048 | 4.25622 |
| CD630_20980 | CD630_SQ1656 | 100 -- 156 | 9 -- 67 | -12.5075 | 8.12847 | 4.25622 |
| CD630_36781 | CD630_SQ1656 | 17 -- 47 | 12 -- 41 | -12.424 | 3.75946 | 4.25622 |
| CD630_32500 | CD630_SQ1656 | 91 -- 131 | 5 -- 52 | -12.3846 | 6.17288 | 4.25622 |
| cdu1 | CD630_SQ1656 | 126 -- 154 | 63 -- 91 | -12.3782 | 4.46276 | 4.25622 |
| spoVFB | CD630_SQ1656 | 166 -- 201 | 7 -- 47 | -12.3205 | 4.62758 | 4.25622 |
| CD630_04620 | CD630_SQ1656 | 68 -- 98 | 7 -- 37 | -12.3151 | 3.45413 | 4.25622 |
| CD630_21700 | CD630_SQ1656 | 89 -- 122 | 7 -- 37 | -12.2679 | 3.96650 | 4.25622 |
| CD630_23280 | CD630_SQ1656 | 134 -- 152 | 209 -- 227 | -12.1604 | 3.50093 | 4.25622 |
| ppnK | CD630_SQ1656 | 97 -- 148 | 2 -- 43 | -12.1427 | 4.36817 | 4.25622 |
| CD630_11730 | CD630_SQ1656 | 21 -- 43 | 9 -- 31 | -12.1423 | 4.80521 | 4.25622 |
| CD630_25150 | CD630_SQ1656 | 135 -- 157 | 1 -- 26 | -12.1012 | 3.79339 | 4.25622 |
| CD630_25190 | CD630_SQ1656 | 155 -- 191 | 183 -- 229 | -12.101 | 3.19663 | 4.25622 |
| CD630_27120 | CD630_SQ1656 | 124 -- 157 | 6 -- 43 | -12.0587 | 4.43720 | 4.25622 |
| CD630_25070 | CD630_SQ1656 | 130 -- 164 | 4 -- 36 | -12.0502 | 3.45164 | 4.25622 |
| CD630_04341 | CD630_SQ1656 | 37 -- 81 | 7 -- 47 | -12.0285 | 4.29262 | 4.25622 |
| CD630_22710 | CD630_SQ1656 | 6 -- 94 | 7 -- 87 | -12.008 | 4.05623 | 4.25622 |
| dltA | CD630_SQ1656 | 98 -- 125 | 9 -- 42 | -11.9623 | 3.85429 | 4.25622 |
| cspD | CD630_SQ1656 | 98 -- 162 | 35 -- 103 | -11.9128 | 3.23587 | 4.25622 |
| CD630_19000 | CD630_SQ1656 | 80 -- 121 | 4 -- 43 | -11.8753 | 4.18443 | 4.25622 |
| CD630_14420 | CD630_SQ1656 | 33 -- 120 | 13 -- 110 | -11.8564 | 3.99497 | 4.25622 |
| CD630_17170 | CD630_SQ1656 | 100 -- 175 | 164 -- 230 | -11.8379 | 4.10230 | 4.25622 |
| CD630_33650 | CD630_SQ1656 | 98 -- 118 | 90 -- 111 | -11.8188 | 4.80565 | 4.25622 |
| sspA | CD630_SQ1656 | 79 -- 109 | 195 -- 224 | -11.8118 | 3.62689 | 4.25622 |
| CD630_16150 | CD630_SQ1656 | 139 -- 166 | 8 -- 36 | -11.8042 | 4.20614 | 4.25622 |
| CD630_10050 | CD630_SQ1656 | 140 -- 163 | 209 -- 230 | -11.7864 | 3.93750 | 4.25622 |
| CD630_06050 | CD630_SQ1656 | 102 -- 139 | 19 -- 53 | -11.7233 | 3.59332 | 4.25622 |
| CD630_07760 | CD630_SQ1656 | 178 -- 200 | 9 -- 31 | -11.6891 | 3.64186 | 4.25622 |
| CD630_13920 | CD630_SQ1656 | 115 -- 147 | 4 -- 41 | -11.6411 | 3.25236 | 4.25622 |
| CD630_18970 | CD630_SQ1656 | 11 -- 46 | 11 -- 41 | -11.5401 | 3.57243 | 4.25622 |
| CD630_27570 | CD630_SQ1656 | 77 -- 115 | 81 -- 118 | -11.5121 | 3.46216 | 4.25622 |
| CD630_23070 | CD630_SQ1656 | 158 -- 196 | 179 -- 213 | -11.4298 | 5.23027 | 4.25622 |
| CD630_13370 | CD630_SQ1656 | 185 -- 198 | 11 -- 24 | -11.4026 | 4.33145 | 4.25622 |
| CD630_20541 | CD630_SQ1656 | 3 -- 44 | 5 -- 53 | -11.3717 | 5.74527 | 4.25622 |
| bltD | CD630_SQ1656 | 175 -- 193 | 7 -- 33 | -11.3584 | 4.49926 | 4.25622 |
| CD630_08130 | CD630_SQ1656 | 79 -- 112 | 8 -- 37 | -11.3164 | 4.63831 | 4.25622 |
| ssuC | CD630_SQ1656 | 36 -- 127 | 5 -- 82 | -11.275 | 3.45491 | 4.25622 |
| CD630_18940 | CD630_SQ1656 | 48 -- 98 | 4 -- 52 | -11.2627 | 4.09988 | 4.25622 |
| CD630_21760 | CD630_SQ1656 | 52 -- 87 | 182 -- 214 | -11.2417 | 4.43846 | 4.25622 |
| CD630_18610 | CD630_SQ1656 | 17 -- 38 | 156 -- 175 | -11.2363 | 4.15582 | 4.25622 |
| CD630_14130 | CD630_SQ1656 | 146 -- 178 | 16 -- 45 | -11.2263 | 4.57970 | 4.25622 |
| CD630_32070 | CD630_SQ1656 | 62 -- 99 | 6 -- 38 | -11.1792 | 3.51964 | 4.25622 |
| xdhA | CD630_SQ1656 | 49 -- 99 | 10 -- 43 | -11.1742 | 4.45147 | 4.25622 |
| CD630_10960 | CD630_SQ1656 | 7 -- 35 | 207 -- 235 | -11.1318 | 3.53850 | 4.25622 |
| folC | CD630_SQ1656 | 89 -- 117 | 7 -- 37 | -11.0174 | 3.85634 | 4.25622 |
| glgB | CD630_SQ1656 | 78 -- 114 | 9 -- 47 | -10.948 | 4.48273 | 4.25622 |
| CD630_09040 | CD630_SQ1656 | 12 -- 65 | 65 -- 120 | -10.9021 | 4.11851 | 4.25622 |
| CD630_27310 | CD630_SQ1656 | 71 -- 97 | 208 -- 234 | -10.8462 | 4.01622 | 4.25622 |
| CD630_13790 | CD630_SQ1656 | 166 -- 200 | 7 -- 40 | -10.8331 | 5.17374 | 4.25622 |
| CD630_31010 | CD630_SQ1656 | 100 -- 145 | 4 -- 40 | -10.8198 | 4.96305 | 4.25622 |
| gcvTPA | CD630_SQ1656 | 65 -- 107 | 33 -- 98 | -10.747 | 4.77229 | 4.25622 |
| pilW | CD630_SQ1656 | 62 -- 116 | 8 -- 49 | -10.7415 | 4.96316 | 4.25622 |
| CD630_31461 | CD630_SQ1656 | 156 -- 187 | 7 -- 41 | -10.7378 | 4.41476 | 4.25622 |
| CD630_13600 | CD630_SQ1656 | 133 -- 182 | 161 -- 201 | -10.7296 | 4.45481 | 4.25622 |
| CD630_16330 | CD630_SQ1656 | 148 -- 201 | 2 -- 38 | -10.722 | 4.11180 | 4.25622 |
| CD630_10880 | CD630_SQ1656 | 155 -- 197 | 179 -- 227 | -10.7077 | 5.05048 | 4.25622 |
| opuCA | CD630_SQ1656 | 89 -- 115 | 10 -- 37 | -10.6822 | 4.07035 | 4.25622 |
| CD630_35790 | CD630_SQ1656 | 65 -- 123 | 7 -- 58 | -10.6516 | 6.97665 | 4.25622 |
| glnA | CD630_SQ1656 | 144 -- 172 | 7 -- 39 | -10.6371 | 3.87629 | 4.25622 |
| CD630_09050 | CD630_SQ1656 | 93 -- 130 | 7 -- 47 | -10.621 | 3.96216 | 4.25622 |
| CD630_31490 | CD630_SQ1656 | 99 -- 132 | 4 -- 41 | -10.6073 | 4.08593 | 4.25622 |
| ldh | CD630_SQ1656 | 157 -- 171 | 115 -- 129 | -10.5965 | 4.35818 | 4.25622 |
| CD630_09990 | CD630_SQ1656 | 23 -- 110 | 7 -- 84 | -10.5382 | 3.86040 | 4.25622 |
| xylR | CD630_SQ1656 | 121 -- 175 | 4 -- 78 | -10.5296 | 3.95671 | 4.25622 |
| CD630_08780 | CD630_SQ1656 | 141 -- 174 | 180 -- 210 | -10.5082 | 7.76690 | 4.25622 |
| CD630_09562 | CD630_SQ1656 | 175 -- 195 | 2 -- 24 | -10.4905 | 3.94307 | 4.25622 |
| CD630_22660 | CD630_SQ1656 | 156 -- 201 | 183 -- 229 | -10.4878 | 3.60496 | 4.25622 |
| CD630_14210 | CD630_SQ1656 | 115 -- 147 | 7 -- 45 | -10.4778 | 3.82355 | 4.25622 |
| CD630_06340 | CD630_SQ1656 | 162 -- 174 | 223 -- 235 | -10.4481 | 4.35983 | 4.25622 |
| CD630_16670 | CD630_SQ1656 | 62 -- 99 | 9 -- 44 | -10.4333 | 3.33553 | 4.25622 |
| CD630_13780 | CD630_SQ1656 | 120 -- 150 | 4 -- 37 | -10.4133 | 3.52274 | 4.25622 |
| CD630_36690 | CD630_SQ1656 | 94 -- 123 | 207 -- 236 | -10.3809 | 3.53761 | 4.25622 |
| aroD | CD630_SQ1656 | 166 -- 186 | 9 -- 29 | -10.3414 | 3.71929 | 4.25622 |
| CD630_06190 | CD630_SQ1656 | 141 -- 151 | 154 -- 164 | -10.3383 | 3.57064 | 4.25622 |
| CD630_18950 | CD630_SQ1656 | 150 -- 169 | 6 -- 23 | -10.3325 | 3.40610 | 4.25622 |
| cbiQ | CD630_SQ1656 | 2 -- 19 | 5 -- 22 | -10.2725 | 3.79830 | 4.25622 |
| CD630_24590 | CD630_SQ1656 | 1 -- 13 | 223 -- 235 | -10.2371 | 3.70401 | 4.25622 |
| CD630_05220 | CD630_SQ1656 | 163 -- 201 | 6 -- 41 | -10.2302 | 3.65057 | 4.25622 |
| CD630_19780 | CD630_SQ1656 | 73 -- 97 | 33 -- 57 | -10.2188 | 3.64830 | 4.25622 |
| CD630_05370 | CD630_SQ1656 | 91 -- 105 | 35 -- 49 | -10.18 | 3.26947 | 4.25622 |
| CD630_28181 | CD630_SQ1656 | 37 -- 82 | 192 -- 228 | -10.1462 | 3.91299 | 4.25622 |
| CD630_08140 | CD630_SQ1656 | 100 -- 129 | 14 -- 40 | -10.1125 | 3.30159 | 4.25622 |
| CD630_08460 | CD630_SQ1656 | 95 -- 127 | 10 -- 41 | -10.0767 | 4.15083 | 4.25622 |
| CD630_05900 | CD630_SQ1656 | 107 -- 152 | 8 -- 52 | -10.0665 | 3.21934 | 4.25622 |
| CD630_03180 | CD630_SQ1656 | 95 -- 107 | 33 -- 45 | -10.0658 | 3.73138 | 4.25622 |
| CD630_27280 | CD630_SQ1656 | 146 -- 189 | 154 -- 200 | -10.0646 | 3.24738 | 4.25622 |
| CD630_16550 | CD630_SQ1656 | 94 -- 124 | 207 -- 236 | -10.0602 | 5.61064 | 4.25622 |
| CD630_31580 | CD630_SQ1656 | 172 -- 196 | 3 -- 25 | -10.0419 | 3.51756 | 4.25622 |
| dapB2 | CD630_SQ1656 | 170 -- 190 | 81 -- 101 | -10.0277 | 3.27218 | 4.25622 |
| appC | CD630_SQ1656 | 39 -- 58 | 178 -- 193 | -10.0163 | 4.50438 | 4.25622 |
| srlEb | CD630_SQ1656 | 60 -- 92 | 8 -- 43 | -9.99531 | 3.50140 | 4.25622 |
| CD630_05870 | CD630_SQ1656 | 165 -- 198 | 20 -- 54 | -9.99281 | 4.28339 | 4.25622 |
| rbsA | CD630_SQ1656 | 60 -- 75 | 15 -- 30 | -9.99027 | 3.71650 | 4.25622 |
| CD630_33570 | CD630_SQ1656 | 2 -- 37 | 90 -- 118 | -9.97723 | 3.95778 | 4.25622 |
| def1 | CD630_SQ1656 | 23 -- 65 | 8 -- 39 | -9.90124 | 3.99720 | 4.25622 |
| bglA | CD630_SQ1656 | 151 -- 190 | 46 -- 84 | -9.88274 | 4.86402 | 4.25622 |
| CD630_14700 | CD630_SQ1656 | 109 -- 130 | 2 -- 21 | -9.87513 | 3.98234 | 4.25622 |
| CD630_17511 | CD630_SQ1656 | 11 -- 34 | 159 -- 184 | -9.86211 | 9.12322 | 4.25622 |
| CD630_16530 | CD630_SQ1656 | 29 -- 120 | 23 -- 117 | -9.837 | 4.37658 | 4.25622 |
| CD630_36250 | CD630_SQ1656 | 58 -- 88 | 8 -- 39 | -9.80219 | 4.13224 | 4.25622 |
| CD630_22050 | CD630_SQ1656 | 77 -- 107 | 9 -- 45 | -9.79661 | 4.87639 | 4.25622 |
| glgP | CD630_SQ1656 | 12 -- 43 | 9 -- 39 | -9.78636 | 3.28530 | 4.25622 |
| CD630_14190 | CD630_SQ1656 | 117 -- 127 | 4 -- 13 | -9.78265 | 3.78562 | 4.25622 |
| CD630_21620 | CD630_SQ1656 | 139 -- 183 | 19 -- 61 | -9.76876 | 3.02649 | 4.25622 |
| CD630_32050 | CD630_SQ1656 | 176 -- 193 | 6 -- 21 | -9.73385 | 5.73797 | 4.25622 |
| CD630_20960 | CD630_SQ1656 | 90 -- 117 | 8 -- 36 | -9.71825 | 5.65940 | 4.25622 |
| CD630_05950 | CD630_SQ1656 | 43 -- 71 | 9 -- 37 | -9.67617 | 3.31669 | 4.25622 |
| blaI | CD630_SQ1656 | 142 -- 169 | 3 -- 40 | -9.66113 | 3.73674 | 4.25622 |
| CD630_15360 | CD630_SQ1656 | 24 -- 37 | 148 -- 161 | -9.65745 | 4.47208 | 4.25622 |
| CD630_05290 | CD630_SQ1656 | 17 -- 64 | 5 -- 46 | -9.64487 | 3.97604 | 4.25622 |
| CD630_21570 | CD630_SQ1656 | 136 -- 154 | 6 -- 25 | -9.62181 | 6.48385 | 4.25622 |
| CD630_02740 | CD630_SQ1656 | 156 -- 189 | 197 -- 229 | -9.6142 | 5.02277 | 4.25622 |
| CD630_26250 | CD630_SQ1656 | 170 -- 177 | 4 -- 11 | -9.5953 | 5.03379 | 4.25622 |
| agrB | CD630_SQ1656 | 100 -- 128 | 12 -- 36 | -9.55794 | 3.92598 | 4.25622 |
| CD630_03420 | CD630_SQ1656 | 126 -- 172 | 8 -- 41 | -9.55713 | 4.71868 | 4.25622 |
| CD630_08360 | CD630_SQ1656 | 7 -- 28 | 7 -- 31 | -9.54462 | 3.19339 | 4.25622 |
| CD630_19700 | CD630_SQ173 | 55 -- 80 | 42 -- 67 | -16.0976 | 3.85861 | 6.59405 |
| CD630_23760 | CD630_SQ173 | 53 -- 69 | 196 -- 212 | -15.9332 | 4.54307 | 6.59405 |
| CD630_30100 | CD630_SQ173 | 175 -- 197 | 55 -- 84 | -14.271 | 4.13234 | 6.59405 |
| CD630_24050 | CD630_SQ173 | 111 -- 140 | 157 -- 179 | -13.8642 | 3.70192 | 6.59405 |
| CD630_21700 | CD630_SQ173 | 40 -- 68 | 40 -- 67 | -12.9783 | 3.96650 | 6.59405 |
| CD630_04410 | CD630_SQ173 | 60 -- 102 | 137 -- 191 | -12.9749 | 3.90451 | 6.59405 |
| CD630_23070 | CD630_SQ173 | 172 -- 187 | 143 -- 157 | -12.85 | 5.23027 | 6.59405 |
| CD630_21260 | CD630_SQ173 | 138 -- 152 | 104 -- 118 | -12.5902 | 4.22255 | 6.59405 |
| CD630_17511 | CD630_SQ173 | 7 -- 27 | 62 -- 85 | -12.5102 | 9.12322 | 6.59405 |
| CD630_25190 | CD630_SQ173 | 91 -- 113 | 41 -- 62 | -12.1279 | 3.19663 | 6.59405 |
| CD630_25620 | CD630_SQ173 | 34 -- 72 | 26 -- 63 | -12.1086 | 3.02685 | 6.59405 |
| CD630_21720 | CD630_SQ173 | 17 -- 45 | 55 -- 81 | -12.0897 | 4.13030 | 6.59405 |
| CD630_15900 | CD630_SQ173 | 107 -- 173 | 161 -- 209 | -12.0185 | 3.17664 | 6.59405 |
| cwp17 | CD630_SQ173 | 131 -- 150 | 151 -- 168 | -12.008 | 3.43363 | 6.59405 |
| dapB2 | CD630_SQ173 | 145 -- 199 | 6 -- 73 | -11.7585 | 3.27218 | 6.59405 |
| CD630_15270 | CD630_SQ173 | 144 -- 200 | 55 -- 111 | -11.7456 | 4.66015 | 6.59405 |
| CD630_05000 | CD630_SQ173 | 123 -- 156 | 40 -- 83 | -11.7388 | 4.02442 | 6.59405 |
| CD630_09040 | CD630_SQ173 | 91 -- 113 | 41 -- 62 | -11.2175 | 4.11851 | 6.59405 |

| | | | | | | |
|---|---|---|---|---|---|---|
| agrB | CD630_SQ173 | 134 – 165 | 43 – 72 | -11.1462 | 3.92598 | 6.59405 |
| nrdD | CD630_SQ173 | 69 – 108 | 144 – 181 | -11.1281 | 3.28290 | 6.59405 |
| CD630_33570 | CD630_SQ173 | 24 – 55 | 31 – 67 | -11.0085 | 3.95778 | 6.59405 |
| CD630_22550 | CD630_SQ173 | 37 – 79 | 29 – 70 | -10.9704 | 3.62379 | 6.59405 |
| CD630_18311 | CD630_SQ173 | 136 – 185 | 146 – 200 | -10.9472 | 6.86047 | 6.59405 |
| CD630_14130 | CD630_SQ173 | 144 – 201 | 143 – 205 | -10.9004 | 4.57970 | 6.59405 |
| CD630_30120 | CD630_SQ173 | 53 – 91 | 42 – 78 | -10.8693 | 4.04859 | 6.59405 |
| CD630_21540 | CD630_SQ173 | 127 – 156 | 152 – 184 | -10.7379 | 4.33090 | 6.59405 |
| CD630_08080 | CD630_SQ173 | 159 – 176 | 51 – 68 | -10.4243 | 4.87819 | 6.59405 |
| CD630_09772 | CD630_SQ173 | 5 – 45 | 18 – 57 | -10.4103 | 3.66033 | 6.59405 |
| CD630_14420 | CD630_SQ173 | 69 – 97 | 147 – 174 | -10.3989 | 3.99497 | 6.59405 |
| CD630_25310 | CD630_SQ173 | 20 – 34 | 162 – 177 | -10.3474 | 3.99861 | 6.59405 |
| CD630_14920 | CD630_SQ173 | 31 – 49 | 33 – 51 | -10.2432 | 4.35225 | 6.59405 |
| CD630_20280 | CD630_SQ173 | 117 – 132 | 144 – 158 | -10.2168 | 3.46343 | 6.59405 |
| CD630_35260 | CD630_SQ173 | 129 – 175 | 3 – 63 | -10.1183 | 5.77277 | 6.59405 |
| CD630_23450 | CD630_SQ173 | 191 – 199 | 225 – 233 | -9.95014 | 3.27218 | 6.59405 |
| CD630_08460 | CD630_SQ173 | 122 – 163 | 18 – 58 | -9.94887 | 4.15083 | 6.59405 |
| CD630_26680 | CD630_SQ173 | 47 – 61 | 12 – 26 | -9.93346 | 3.53956 | 6.59405 |
| glgP | CD630_SQ173 | 32 – 48 | 41 – 58 | -9.91013 | 3.28530 | 6.59405 |
| CD630_15360 | CD630_SQ173 | 24 – 37 | 102 – 115 | -9.86436 | 4.47208 | 6.59405 |
| CD630_14360 | CD630_SQ173 | 92 – 109 | 45 – 62 | -9.72536 | 3.02079 | 6.59405 |
| CD630_09570 | CD630_SQ173 | 3 – 19 | 8 – 24 | -9.69577 | 5.31213 | 6.59405 |
| CD630_29070 | CD630_SQ173 | 3 – 19 | 8 – 24 | -9.69577 | 3.89477 | 6.59405 |
| CD630_03420 | CD630_SQ173 | 37 – 77 | 32 – 69 | -9.54756 | 4.71868 | 6.59405 |
| CD630_06560 | CD630_SQ173 | 1 – 23 | 49 – 69 | -9.54662 | 4.02532 | 6.59405 |
| feoB | CD630_SQ173 | 185 – 197 | 142 – 155 | -9.50445 | 4.05364 | 6.59405 |
| pgm | CD630_SQ2429 | 63 – 112 | 58 – 100 | -17.2024 | 3.45439 | 5.77720 |
| CD630_17630 | CD630_SQ2429 | 109 – 163 | 59 – 110 | -16.1649 | 4.52408 | 5.77720 |
| CD630_23450 | CD630_SQ2429 | 10 – 67 | 55 – 107 | -15.3495 | 3.27218 | 5.77720 |
| CD630_20920 | CD630_SQ2429 | 33 – 89 | 50 – 108 | -14.3453 | 4.13812 | 5.77720 |
| CD630_30540 | CD630_SQ2429 | 124 – 150 | 2 – 27 | -13.4058 | 3.63804 | 5.77720 |
| CD630_23070 | CD630_SQ2429 | 51 – 186 | 116 – 230 | -13.3272 | 5.23027 | 5.77720 |
| CD630_10000 | CD630_SQ2429 | 107 – 165 | 22 – 66 | -13.2448 | 3.86040 | 5.77720 |
| CD630_05460 | CD630_SQ2429 | 73 – 113 | 192 – 233 | -13.1703 | 3.98753 | 5.77720 |
| CD630_33570 | CD630_SQ2429 | 2 – 37 | 82 – 111 | -12.8439 | 3.95778 | 5.77720 |
| CD630_05950 | CD630_SQ2429 | 43 – 64 | 3 – 27 | -12.6271 | 3.31669 | 5.77720 |
| CD630_21260 | CD630_SQ2429 | 138 – 152 | 143 – 157 | -12.4886 | 4.22255 | 5.77720 |
| CD630_24830 | CD630_SQ2429 | 6 – 28 | 203 – 225 | -12.3366 | 4.98239 | 5.77720 |
| CD630_10960 | CD630_SQ2429 | 92 – 138 | 43 – 87 | -11.6892 | 3.53850 | 5.77720 |
| CD630_07470 | CD630_SQ2429 | 24 – 106 | 24 – 106 | -11.5732 | 3.37143 | 5.77720 |
| CD630_21770 | CD630_SQ2429 | 1 – 37 | 80 – 122 | -11.4732 | 4.02587 | 5.77720 |
| CD630_32990 | CD630_SQ2429 | 41 – 69 | 86 – 122 | -11.3456 | 5.24925 | 5.77720 |
| bglA | CD630_SQ2429 | 152 – 201 | 28 – 76 | -11.3017 | 4.86402 | 5.77720 |
| CD630_15460 | CD630_SQ2429 | 168 – 184 | 195 – 212 | -11.2977 | 6.86296 | 5.77720 |
| CD630_36781 | CD630_SQ2429 | 12 – 28 | 20 – 36 | -11.2356 | 3.75946 | 5.77720 |
| CD630_17720 | CD630_SQ2429 | 9 – 28 | 57 – 76 | -11.1052 | 3.22910 | 5.77720 |
| CD630_03920 | CD630_SQ2429 | 58 – 77 | 6 – 26 | -11.0864 | 4.15934 | 5.77720 |
| CD630_08360 | CD630_SQ2429 | 155 – 199 | 78 – 104 | -11.0302 | 3.19339 | 5.77720 |
| pilD | CD630_SQ2429 | 4 – 71 | 172 – 236 | -10.949 | 4.54123 | 5.77720 |
| CD630_18610 | CD630_SQ2429 | 17 – 38 | 149 – 168 | -10.9345 | 4.15582 | 5.77720 |
| CD630_20600 | CD630_SQ2429 | 179 – 193 | 3 – 17 | -10.9345 | 4.20093 | 5.77720 |
| glgP | CD630_SQ2429 | 83 – 129 | 196 – 236 | -10.8983 | 3.28530 | 5.77720 |
| etfA | CD630_SQ2429 | 2 – 27 | 75 – 101 | -10.8406 | 3.41204 | 5.77720 |
| CD630_04880 | CD630_SQ2429 | 78 – 116 | 16 – 57 | -10.7789 | 4.62291 | 5.77720 |
| CD630_36170 | CD630_SQ2429 | 188 – 200 | 24 – 36 | -10.6983 | 3.72380 | 5.77720 |
| CD630_10800 | CD630_SQ2429 | 2 – 69 | 38 – 108 | -10.6522 | 5.30319 | 5.77720 |
| CD630_06190 | CD630_SQ2429 | 141 – 151 | 147 – 157 | -10.6297 | 3.57064 | 5.77720 |
| CD630_20450 | CD630_SQ2429 | 128 – 165 | 192 – 228 | -10.4974 | 3.77388 | 5.77720 |
| CD630_34520 | CD630_SQ2429 | 186 – 199 | 73 – 87 | -10.4618 | 3.83016 | 5.77720 |
| CD630_09970 | CD630_SQ2429 | 35 – 81 | 55 – 101 | -10.4563 | 3.25917 | 5.77720 |
| CD630_20670 | CD630_SQ2429 | 179 – 197 | 177 – 194 | -10.3685 | 4.42757 | 5.77720 |
| CD630_13020 | CD630_SQ2429 | 93 – 123 | 206 – 236 | -10.3604 | 3.52267 | 5.77720 |
| CD630_20260 | CD630_SQ2429 | 64 – 127 | 1 – 66 | -10.2379 | 3.84162 | 5.77720 |
| ldh | CD630_SQ2429 | 157 – 171 | 108 – 122 | -10.2012 | 4.35818 | 5.77720 |
| CD630_25190 | CD630_SQ2429 | 174 – 181 | 192 – 199 | -10.1528 | 3.19663 | 5.77720 |
| xdhA | CD630_SQ2429 | 72 – 131 | 4 – 67 | -10.1426 | 4.45147 | 5.77720 |
| CD630_02420 | CD630_SQ2429 | 9 – 50 | 147 – 193 | -10.097 | 3.71048 | 5.77720 |
| CD630_06380 | CD630_SQ2429 | 136 – 157 | 199 – 221 | -10.0726 | 4.32211 | 5.77720 |
| CD630_08950 | CD630_SQ2429 | 133 – 184 | 60 – 115 | -10.0506 | 3.71905 | 5.77720 |
| CD630_36250 | CD630_SQ2429 | 154 – 174 | 5 – 27 | -10.0498 | 4.13224 | 5.77720 |
| spaK | CD630_SQ2429 | 139 – 187 | 55 – 110 | -9.97521 | 3.04530 | 5.77720 |
| CD630_19942 | CD630_SQ2429 | 93 – 119 | 3 – 27 | -9.96438 | 4.24574 | 5.77720 |
| CD630_08110 | CD630_SQ2429 | 123 – 144 | 91 – 112 | -9.93826 | 4.39181 | 5.77720 |
| CD630_18960 | CD630_SQ2429 | 12 – 33 | 209 – 229 | -9.90073 | 4.37822 | 5.77720 |
| CD630_15360 | CD630_SQ2429 | 24 – 37 | 141 – 154 | -9.8464 | 4.47208 | 5.77720 |
| CD630_07400 | CD630_SQ2429 | 1 – 13 | 97 – 109 | -9.82922 | 4.52273 | 5.77720 |
| CD630_29760 | CD630_SQ2429 | 69 – 97 | 1 – 30 | -9.74769 | 3.51074 | 5.77720 |
| CD630_30140 | CD630_SQ2429 | 67 – 85 | 48 – 67 | -9.73917 | 3.28343 | 5.77720 |
| CD630_13370 | CD630_SQ2429 | 154 – 165 | 203 – 214 | -9.72185 | 4.33145 | 5.77720 |
| CD630_28290 | CD630_SQ2429 | 34 – 52 | 56 – 73 | -9.72017 | 5.54192 | 5.77720 |
| CD630_15431 | CD630_SQ2429 | 125 – 156 | 4 – 33 | -9.64164 | 5.29780 | 5.77720 |
| CD630_25040 | CD630_SQ2429 | 144 – 177 | 125 – 160 | -9.59662 | 3.43898 | 5.77720 |
| CD630_10970 | CD630_SQ2429 | 97 – 125 | 4 – 33 | -9.58795 | 3.53850 | 5.77720 |
| CD630_34530 | CD630_SQ2429 | 29 – 54 | 209 – 235 | -9.55766 | 3.83016 | 5.77720 |
| CD630_16530 | CD630_SQ327 | 55 – 145 | 77 – 164 | -14.3797 | 4.37658 | 4.62291 |
| chrA | CD630_SQ327 | 7 – 59 | 69 – 121 | -14.3194 | 5.27255 | 4.62291 |
| CD630_08070 | CD630_SQ327 | 103 – 131 | 128 – 153 | -14.2198 | 4.87819 | 4.62291 |
| CD630_23780 | CD630_SQ327 | 131 – 175 | 188 – 220 | -13.8319 | 3.91629 | 4.62291 |
| CD630_27120 | CD630_SQ327 | 42 – 107 | 52 – 127 | -13.3788 | 4.43720 | 4.62291 |
| CD630_03040 | CD630_SQ327 | 111 – 149 | 111 – 140 | -13.2115 | 3.51403 | 4.62291 |
| CD630_21090 | CD630_SQ327 | 102 – 190 | 69 – 166 | -13.0753 | 4.15803 | 4.62291 |
| CD630_04610 | CD630_SQ327 | 145 – 193 | 67 – 114 | -13.0598 | 3.83025 | 4.62291 |
| CD630_06060 | CD630_SQ327 | 41 – 60 | 109 – 129 | -12.9525 | 4.94050 | 4.62291 |
| CD630_06380 | CD630_SQ327 | 111 – 160 | 42 – 88 | -12.9099 | 4.32211 | 4.62291 |
| CD630_34560 | CD630_SQ327 | 4 – 91 | 58 – 124 | -12.8871 | 4.02604 | 4.62291 |
| CD630_18930 | CD630_SQ327 | 50 – 77 | 62 – 91 | -12.5998 | 5.32127 | 4.62291 |
| CD630_32110 | CD630_SQ327 | 11 – 119 | 25 – 139 | -12.4901 | 4.45291 | 4.62291 |
| CD630_15270 | CD630_SQ327 | 175 – 201 | 13 – 38 | -12.4294 | 4.66015 | 4.62291 |
| bglA | CD630_SQ327 | 49 – 86 | 97 – 127 | -12.3027 | 4.86402 | 4.62291 |
| agrB | CD630_SQ327 | 131 – 153 | 79 – 100 | -12.3013 | 3.92598 | 4.62291 |
| CD630_21110 | CD630_SQ327 | 43 – 93 | 58 – 103 | -12.0024 | 3.89630 | 4.62291 |
| cobS | CD630_SQ327 | 138 – 153 | 112 – 127 | -11.8919 | 3.67481 | 4.62291 |
| CD630_04770 | CD630_SQ327 | 36 – 52 | 109 – 127 | -11.8599 | 3.58478 | 4.62291 |
| CD630_21570 | CD630_SQ327 | 4 – 20 | 106 – 122 | -11.7995 | 6.48385 | 4.62291 |
| CD630_28870 | CD630_SQ327 | 160 – 178 | 207 – 224 | -11.6724 | 3.41991 | 4.62291 |
| CD630_21260 | CD630_SQ327 | 115 – 141 | 191 – 218 | -11.6508 | 4.22255 | 4.62291 |
| sspA | CD630_SQ327 | 50 – 64 | 71 – 85 | -11.6425 | 3.62689 | 4.62291 |
| pilD | CD630_SQ327 | 3 – 16 | 114 – 127 | -11.4885 | 4.54123 | 4.62291 |
| CD630_08660 | CD630_SQ327 | 148 – 200 | 88 – 143 | -11.4808 | 3.86467 | 4.62291 |
| CD630_27110 | CD630_SQ327 | 34 – 57 | 191 – 218 | -11.3055 | 3.78231 | 4.62291 |
| iorA | CD630_SQ327 | 138 – 162 | 105 – 129 | -11.1305 | 3.83173 | 4.62291 |
| CD630_07430 | CD630_SQ327 | 34 – 82 | 77 – 122 | -11.115 | 4.65568 | 4.62291 |
| CD630_06690 | CD630_SQ327 | 166 – 197 | 189 – 224 | -11.0532 | 5.82709 | 4.62291 |
| CD630_22500 | CD630_SQ327 | 137 – 179 | 130 – 175 | -11.0288 | 4.03688 | 4.62291 |
| CD630_27040 | CD630_SQ327 | 35 – 59 | 79 – 103 | -10.9885 | 3.51812 | 4.62291 |
| CD630_18781 | CD630_SQ327 | 10 – 86 | 26 – 88 | -10.9148 | 3.77660 | 4.62291 |
| cdtR | CD630_SQ327 | 20 – 33 | 202 – 214 | -10.9132 | 5.84368 | 4.62291 |
| CD630_13370 | CD630_SQ327 | 150 – 162 | 208 – 220 | -10.8969 | 4.33145 | 4.62291 |
| dltB | CD630_SQ327 | 129 – 155 | 70 – 95 | -10.8773 | 3.81485 | 4.62291 |
| CD630_31220 | CD630_SQ327 | 9 – 27 | 121 – 140 | -10.8668 | 3.73779 | 4.62291 |
| ppnK | CD630_SQ327 | 126 – 154 | 112 – 149 | -10.8583 | 4.36817 | 4.62291 |
| CD630_04880 | CD630_SQ327 | 39 – 53 | 202 – 216 | -10.7894 | 4.62291 | 4.62291 |
| CD630_19904 | CD630_SQ327 | 162 – 180 | 147 – 165 | -10.758 | 5.96526 | 4.62291 |
| CD630_09760 | CD630_SQ327 | 134 – 150 | 122 – 139 | -10.7262 | 4.83410 | 4.62291 |
| CD630_28910 | CD630_SQ327 | 134 – 150 | 122 – 139 | -10.7262 | 4.51664 | 4.62291 |
| nusB | CD630_SQ327 | 142 – 175 | 3 – 41 | -10.7131 | 4.92798 | 4.62291 |
| CD630_17900 | CD630_SQ327 | 172 – 201 | 122 – 149 | -10.7091 | 3.52167 | 4.62291 |
| tcdC | CD630_SQ327 | 132 – 158 | 109 – 138 | -10.7046 | 3.76101 | 4.62291 |
| CD630_02420 | CD630_SQ327 | 155 – 181 | 133 – 153 | -10.5846 | 3.71048 | 4.62291 |
| CD630_14990 | CD630_SQ327 | 148 – 167 | 109 – 129 | -10.5806 | 3.78619 | 4.62291 |
| CD630_20660 | CD630_SQ327 | 125 – 141 | 166 – 182 | -10.5096 | 5.39321 | 4.62291 |
| CD630_27280 | CD630_SQ327 | 32 – 68 | 193 – 224 | -10.5024 | 3.24738 | 4.62291 |
| CD630_16150 | CD630_SQ327 | 1 – 21 | 121 – 136 | -10.4158 | 4.20614 | 4.62291 |
| CD630_19030 | CD630_SQ327 | 165 – 182 | 139 – 157 | -10.2893 | 3.34240 | 4.62291 |
| CD630_06080 | CD630_SQ327 | 157 – 199 | 134 – 179 | -10.2038 | 3.82315 | 4.62291 |
| cls | CD630_SQ327 | 148 – 173 | 55 – 81 | -10.1858 | 4.46177 | 4.62291 |
| CD630_14130 | CD630_SQ327 | 143 – 171 | 137 – 165 | -10.1218 | 4.57970 | 4.62291 |
| eutP | CD630_SQ327 | 160 – 172 | 139 – 151 | -10.0235 | 3.82847 | 4.62291 |
| CD630_05830 | CD630_SQ327 | 133 – 170 | 130 – 175 | -9.88698 | 3.35360 | 4.62291 |
| CD630_02720 | CD630_SQ327 | 144 – 201 | 63 – 118 | -9.80063 | 3.38938 | 4.62291 |
| CD630_34520 | CD630_SQ327 | 9 – 60 | 84 – 126 | -9.74279 | 3.83016 | 4.62291 |
| CD630_31880 | CD630_SQ327 | 11 – 39 | 88 – 115 | -9.73384 | 3.97522 | 4.62291 |
| CD630_10950 | CD630_SQ327 | 108 – 148 | 1 – 39 | -9.63741 | 3.67675 | 4.62291 |
| CD630_31490 | CD630_SQ327 | 154 – 162 | 149 – 157 | -9.63111 | 4.08593 | 4.62291 |
| CD630_29840 | CD630_SQ327 | 5 – 24 | 29 – 48 | -9.58387 | 4.48825 | 4.62291 |
| CD630_29250 | CD630_SQ327 | 33 – 84 | 95 – 144 | -9.57904 | 4.61901 | 4.62291 |
| CD630_09410 | CD630_SQ327 | 33 – 84 | 95 – 144 | -9.57904 | 4.75651 | 4.62291 |
| CD630_30190 | CD630_SQ327 | 120 – 150 | 90 – 124 | -9.56939 | 3.27394 | 4.62291 |
| CD630_16750 | CD630_SQ327 | 2 – 36 | 76 – 115 | -9.54826 | 5.04051 | 4.62291 |
| CD630_13020 | CD630_SQ327 | 117 – 147 | 121 – 159 | -9.50412 | 3.52267 | 4.62291 |
| dltD | CD630_SQ367 | 110 – 161 | 1 – 48 | -18.054 | 4.40777 | 4.08290 |
| cwp17 | CD630_SQ367 | 10 – 37 | 172 – 203 | -17.9008 | 3.43363 | 4.08290 |
| CD630_09100 | CD630_SQ367 | 19 – 68 | 138 – 184 | -16.7028 | 5.92260 | 4.08290 |
| CD630_14810 | CD630_SQ367 | 110 – 146 | 2 – 49 | -16.2161 | 3.35323 | 4.08290 |
| CD630_19401 | CD630_SQ367 | 74 – 162 | 127 – 205 | -15.9657 | 4.37084 | 4.08290 |
| CD630_17511 | CD630_SQ367 | 4 – 28 | 129 – 156 | -15.9365 | 9.12322 | 4.08290 |
| CD630_22050 | CD630_SQ367 | 1 – 45 | 121 – 164 | -15.9151 | 4.87639 | 4.08290 |
| CD630_16530 | CD630_SQ367 | 17 – 72 | 132 – 173 | -15.4982 | 4.37658 | 4.08290 |
| CD630_15900 | CD630_SQ367 | 138 – 190 | 108 – 165 | -15.3857 | 3.17664 | 4.08290 |
| CD630_21260 | CD630_SQ367 | 117 – 165 | 140 – 188 | -15.1613 | 4.22255 | 4.08290 |
| CD630_07070 | CD630_SQ367 | 117 – 160 | 145 – 186 | -14.7321 | 3.46414 | 4.08290 |
| cobS | CD630_SQ367 | 85 – 141 | 127 – 193 | -14.4172 | 3.67481 | 4.08290 |
| CD630_31500 | CD630_SQ367 | 128 – 165 | 3 – 33 | -14.2401 | 4.90207 | 4.08290 |
| CD630_26250 | CD630_SQ367 | 111 – 153 | 138 – 183 | -14.233 | 5.03379 | 4.08290 |
| CD630_22880 | CD630_SQ367 | 43 – 85 | 151 – 195 | -14.1046 | 3.62455 | 4.08290 |
| CD630_23450 | CD630_SQ367 | 1 – 67 | 107 – 168 | -14.0699 | 3.27218 | 4.08290 |
| CD630_18311 | CD630_SQ367 | 133 – 185 | 126 – 179 | -14.0262 | 6.86047 | 4.08290 |
| CD630_31690 | CD630_SQ367 | 153 – 196 | 41 – 86 | -13.9999 | 3.71193 | 4.08290 |
| CD630_13530 | CD630_SQ367 | 1 – 31 | 137 – 166 | -13.9786 | 4.09832 | 4.08290 |
| CD630_06060 | CD630_SQ367 | 164 – 182 | 67 – 85 | -13.9019 | 4.94050 | 4.08290 |
| CD630_24830 | CD630_SQ367 | 152 – 179 | 68 – 93 | -13.788 | 4.98239 | 4.08290 |
| CD630_21090 | CD630_SQ367 | 159 – 172 | 143 – 156 | -13.771 | 4.15803 | 4.08290 |
| CD630_14610 | CD630_SQ367 | 143 – 183 | 140 – 179 | -13.6698 | 4.89428 | 4.08290 |
| CD630_21610 | CD630_SQ367 | 128 – 168 | 147 – 187 | -13.611 | 3.16662 | 4.08290 |
| CD630_27280 | CD630_SQ367 | 144 – 196 | 147 – 181 | -13.6089 | 3.24738 | 4.08290 |
| CD630_21570 | CD630_SQ367 | 13 – 53 | 71 – 119 | -13.4517 | 6.48385 | 4.08290 |
| CD630_28550 | CD630_SQ367 | 174 – 199 | 162 – 194 | -13.4249 | 3.51196 | 4.08290 |
| cobQ | CD630_SQ367 | 77 – 105 | 2 – 30 | -13.274 | 3.13524 | 4.08290 |
| cspC | CD630_SQ367 | 123 – 150 | 165 – 193 | -13.2077 | 5.48848 | 4.08290 |
| CD630_33570 | CD630_SQ367 | 41 – 74 | 8 – 46 | -13.1948 | 3.95778 | 4.08290 |
| CD630_27110 | CD630_SQ367 | 1 – 12 | 75 – 86 | -13.0912 | 3.78231 | 4.08290 |
| CD630_25200 | CD630_SQ367 | 20 – 67 | 128 – 179 | -13.041 | 3.65158 | 4.08290 |
| CD630_16940 | CD630_SQ367 | 86 – 133 | 166 – 217 | -12.9918 | 4.14665 | 4.08290 |
| CD630_02420 | CD630_SQ367 | 1 – 30 | 188 – 218 | -12.9303 | 3.71048 | 4.08290 |
| CD630_03400 | CD630_SQ367 | 26 – 58 | 143 – 174 | -12.9049 | 4.45994 | 4.08290 |
| fbp | CD630_SQ367 | 129 – 152 | 12 – 37 | -12.8628 | 3.80071 | 4.08290 |
| vncR | CD630_SQ367 | 59 – 91 | 11 – 45 | -12.805 | 3.93642 | 4.08290 |
| oppB | CD630_SQ367 | 34 – 105 | 121 – 193 | -12.761 | 2.98328 | 4.08290 |
| cspD | CD630_SQ367 | 123 – 153 | 163 – 193 | -12.6907 | 3.23587 | 4.08290 |
| CD630_18100 | CD630_SQ367 | 7 – 44 | 170 – 199 | -12.6481 | 5.71658 | 4.08290 |
| CD630_27570 | CD630_SQ367 | 6 – 21 | 77 – 93 | -12.5879 | 3.46216 | 4.08290 |
| CD630_10220 | CD630_SQ367 | 125 – 164 | 65 – 120 | -12.4802 | 4.08233 | 4.08290 |
| CD630_17720 | CD630_SQ367 | 2 – 41 | 138 – 179 | -12.4777 | 3.22910 | 4.08290 |
| CD630_04850 | CD630_SQ367 | 157 – 197 | 152 – 188 | -12.3903 | 4.40788 | 4.08290 |
| CD630_31960 | CD630_SQ367 | 57 – 73 | 118 – 134 | -12.3068 | 4.93757 | 4.08290 |
| CD630_29760 | CD630_SQ367 | 117 – 164 | 162 – 206 | -12.3028 | 3.51074 | 4.08290 |
| CD630_20650 | CD630_SQ367 | 33 – 43 | 168 – 178 | -12.2963 | 4.00083 | 4.08290 |
| CD630_04610 | CD630_SQ367 | 161 – 200 | 147 – 185 | -12.2259 | 3.83025 | 4.08290 |
| CD630_30420 | CD630_SQ367 | 146 – 174 | 126 – 159 | -12.0611 | 5.27539 | 4.08290 |
| CD630_27600 | CD630_SQ367 | 7 – 43 | 149 – 185 | -12.0364 | 3.48384 | 4.08290 |
| CD630_16460 | CD630_SQ367 | 122 – 161 | 2 – 46 | -12.0318 | 4.64028 | 4.08290 |
| CD630_13790 | CD630_SQ367 | 107 – 155 | 79 – 128 | -11.9897 | 5.17374 | 4.08290 |
| CD630_13960 | CD630_SQ367 | 142 – 155 | 192 – 205 | -11.98 | 3.30375 | 4.08290 |
| CD630_29750 | CD630_SQ367 | 62 – 94 | 11 – 47 | -11.9234 | 4.59146 | 4.08290 |
| ogt1 | CD630_SQ367 | 147 – 186 | 168 – 208 | -11.7349 | 3.16772 | 4.08290 |
| CD630_01690 | CD630_SQ367 | 186 – 201 | 149 – 164 | -11.6223 | 4.97234 | 4.08290 |
| CD630_24050 | CD630_SQ367 | 5 – 73 | 111 – 179 | -11.5956 | 3.70192 | 4.08290 |
| CD630_09040 | CD630_SQ367 | 1 – 32 | 125 – 155 | -11.5816 | 4.11851 | 4.08290 |
| CD630_29250 | CD630_SQ367 | 169 – 196 | 163 – 191 | -11.555 | 4.61901 | 4.08290 |
| CD630_09410 | CD630_SQ367 | 169 – 196 | 163 – 191 | -11.555 | 4.75651 | 4.08290 |
| CD630_04880 | CD630_SQ367 | 107 – 149 | 2 – 47 | -11.5217 | 4.62291 | 4.08290 |
| CD630_03040 | CD630_SQ367 | 161 – 196 | 17 – 49 | -11.5155 | 3.51403 | 4.08290 |
| CD630_13950 | CD630_SQ367 | 137 – 201 | 81 – 174 | -11.4903 | 4.25540 | 4.08290 |
| CD630_07480 | CD630_SQ367 | 150 – 193 | 15 – 47 | -11.3935 | 5.18321 | 4.08290 |
| CD630_13020 | CD630_SQ367 | 55 – 145 | 127 – 204 | -11.3769 | 3.52267 | 4.08290 |
| CD630_08130 | CD630_SQ367 | 177 – 191 | 160 – 174 | -11.3573 | 4.63831 | 4.08290 |
| CD630_27240 | CD630_SQ367 | 140 – 160 | 63 – 87 | -11.3067 | 3.52462 | 4.08290 |
| CD630_10080 | CD630_SQ367 | 100 – 156 | 3 – 49 | -11.2967 | 3.75625 | 4.08290 |
| CD630_18410 | CD630_SQ367 | 117 – 175 | 148 – 221 | -11.2753 | 3.59739 | 4.08290 |
| panB | CD630_SQ367 | 161 – 187 | 63 – 85 | -11.2393 | 3.40248 | 4.08290 |
| CD630_25430 | CD630_SQ367 | 61 – 168 | 106 – 216 | -11.2255 | 4.00527 | 4.08290 |
| chrA | CD630_SQ367 | 38 – 120 | 133 – 205 | -11.22 | 5.27255 | 4.08290 |
| CD630_18960 | CD630_SQ367 | 52 – 94 | 2 – 46 | -11.2178 | 4.37822 | 4.08290 |
| CD630_08510 | CD630_SQ367 | 11 – 48 | 11 – 47 | -11.1346 | 5.16329 | 4.08290 |
| CD630_28580 | CD630_SQ367 | 149 – 191 | 149 – 188 | -11.1178 | 4.67130 | 4.08290 |
| opuCC | CD630_SQ367 | 116 – 198 | 149 – 233 | -11.1123 | 4.07035 | 4.08290 |
| uxaA | CD630_SQ367 | 170 – 185 | 68 – 82 | -11.107 | 4.26165 | 4.08290 |
| CD630_21490 | CD630_SQ367 | 150 – 201 | 163 – 215 | -11.0936 | 3.37462 | 4.08290 |
| CD630_26920 | CD630_SQ367 | 50 – 99 | 142 – 190 | -11.0343 | 4.58060 | 4.08290 |
| CD630_06170 | CD630_SQ367 | 136 – 156 | 145 – 164 | -11.01 | 4.95366 | 4.08290 |
| CD630_35780 | CD630_SQ367 | 53 – 67 | 1 – 24 | -10.9823 | 3.50107 | 4.08290 |
| CD630_25620 | CD630_SQ367 | 79 – 95 | 1 – 24 | -10.8681 | 3.02685 | 4.08290 |
| cdtR | CD630_SQ367 | 56 – 75 | 142 – 165 | -10.8044 | 5.84368 | 4.08290 |
| CD630_25310 | CD630_SQ367 | 13 – 30 | 3 – 21 | -10.7951 | 3.99861 | 4.08290 |
| CD630_17360 | CD630_SQ367 | 5 – 24 | 67 – 85 | -10.6828 | 5.66070 | 4.08290 |
| CD630_14620 | CD630_SQ367 | 13 – 49 | 140 – 185 | -10.6491 | 4.43651 | 4.08290 |
| CD630_06380 | CD630_SQ367 | 142 – 184 | 145 – 194 | -10.6352 | 4.32211 | 4.08290 |
| CD630_10800 | CD630_SQ367 | 20 – 161 | 100 – 227 | -10.5731 | 5.30319 | 4.08290 |
| CD630_04320 | CD630_SQ367 | 159 – 171 | 4 – 16 | -10.57 | 3.68342 | 4.08290 |
| CD630_08390 | CD630_SQ367 | 70 – 115 | 178 – 229 | -10.5111 | 5.16000 | 4.08290 |
| CD630_26150 | CD630_SQ367 | 19 – 31 | 76 – 88 | -10.4242 | 4.06302 | 4.08290 |
| CD630_31381 | CD630_SQ367 | 42 – 105 | 167 – 227 | -10.4219 | 3.49137 | 4.08290 |
| tcdC | CD630_SQ367 | 27 – 62 | 131 – 164 | -10.3939 | 3.76101 | 4.08290 |
| CD630_08080 | CD630_SQ367 | 100 – 156 | 64 – 107 | -10.3583 | 4.87819 | 4.08290 |
| CD630_21030 | CD630_SQ367 | 157 – 195 | 149 – 191 | -10.346 | 3.08733 | 4.08290 |
| CD630_21760 | CD630_SQ367 | 39 – 51 | 171 – 195 | -10.3349 | 4.43846 | 4.08290 |
| CD630_16540 | CD630_SQ367 | 184 – 197 | 87 – 100 | -10.3241 | 4.37658 | 4.08290 |
| CD630_24590 | CD630_SQ367 | 170 – 190 | 10 – 32 | -10.2535 | 3.70401 | 4.08290 |
| CD630_17170 | CD630_SQ367 | 86 – 126 | 10 – 47 | -10.2229 | 4.10230 | 4.08290 |
| CD630_05830 | CD630_SQ367 | 106 – 141 | 70 – 113 | -10.2063 | 3.35360 | 4.08290 |
| CD630_22950 | CD630_SQ367 | 1 – 16 | 132 – 147 | -10.1684 | 3.96883 | 4.08290 |
| CD630_26680 | CD630_SQ367 | 186 – 196 | 189 – 199 | -10.1073 | 3.53956 | 4.08290 |
| CD630_05300 | CD630_SQ367 | 118 – 149 | 14 – 43 | -10.1013 | 4.56885 | 4.08290 |
| CD630_34030 | CD630_SQ367 | 32 – 76 | 12 – 45 | -10.0882 | 3.05766 | 4.08290 |
| CD630_06060 | CD630_SQ367 | 147 – 181 | 2 – 34 | -10.0847 | 4.94050 | 4.08290 |
| CD630_25150 | CD630_SQ367 | 16 – 58 | 160 – 205 | -10.069 | 3.79339 | 4.08290 |
| CD630_16160 | CD630_SQ367 | 41 – 64 | 162 – 185 | -10.0466 | 6.11801 | 4.08290 |
| CD630_25020 | CD630_SQ367 | 4 – 69 | 31 – 98 | -10.0464 | 4.17721 | 4.08290 |
| CD630_14150 | CD630_SQ367 | 28 – 51 | 120 – 141 | -10.0077 | 3.28251 | 4.08290 |
| rolA | CD630_SQ367 | 153 – 173 | 17 – 38 | -9.99247 | 4.56362 | 4.08290 |
| CD630_35430 | CD630_SQ367 | 115 – 124 | 144 – 153 | -9.99157 | 4.28798 | 4.08290 |
| flhB | CD630_SQ367 | 33 – 53 | 148 – 173 | -9.98488 | 3.55524 | 4.08290 |
| CD630_32520 | CD630_SQ367 | 156 – 198 | 2 – 48 | -9.95984 | 4.06098 | 4.08290 |
| CD630_20920 | CD630_SQ367 | 78 – 123 | 67 – 113 | -9.95949 | 4.13812 | 4.08290 |
| CD630_19790 | CD630_SQ367 | 128 – 153 | 172 – 193 | -9.91372 | 3.79593 | 4.08290 |
| CD630_14010 | CD630_SQ367 | 121 – 148 | 127 – 153 | -9.87638 | 3.69983 | 4.08290 |
| CD630_15270 | CD630_SQ367 | 141 – 158 | 4 – 23 | -9.86652 | 4.66015 | 4.08290 |
| CD630_06690 | CD630_SQ367 | 159 – 189 | 16 – 44 | -9.83648 | 5.82709 | 4.08290 |
| CD630_17281 | CD630_SQ367 | 2 – 30 | 3 – 32 | -9.83189 | 5.91544 | 4.08290 |
| ung | CD630_SQ367 | 119 – 186 | 145 – 198 | -9.82042 | 3.30908 | 4.08290 |
| CD630_23780 | CD630_SQ367 | 63 – 108 | 2 – 48 | -9.79941 | 3.91629 | 4.08290 |
| LinCd | CD630_SQ367 | 76 – 100 | 195 – 220 | -9.79397 | 4.33624 | 4.08290 |
| CD630_32710 | CD630_SQ367 | 13 – 37 | 20 – 43 | -9.77887 | 4.74276 | 4.08290 |
| CD630_06100 | CD630_SQ367 | 188 – 201 | 140 – 153 | -9.75702 | 4.43918 | 4.08290 |
| CD630_13580 | CD630_SQ367 | 114 – 165 | 64 – 112 | -9.69902 | 3.80715 | 4.08290 |
| CD630_13350 | CD630_SQ367 | 15 – 62 | 108 – 155 | -9.69411 | 4.15139 | 4.08290 |
| def1 | CD630_SQ367 | 139 – 155 | 243 – 259 | -9.66866 | 3.99720 | 4.08290 |
| srlD | CD630_SQ367 | 133 – 146 | 11 – 24 | -9.63352 | 4.19843 | 4.08290 |
| CD630_28280 | CD630_SQ367 | 11 – 106 | 8 – 89 | -9.62907 | 4.59833 | 4.08290 |
| CD630_18650 | CD630_SQ367 | 49 – 84 | 127 – 165 | -9.62813 | 3.75892 | 4.08290 |
| ldh | CD630_SQ367 | 150 – 179 | 170 – 210 | -9.62058 | 4.35818 | 4.08290 |
| CD630_22500 | CD630_SQ367 | 44 – 57 | 125 – 138 | -9.56519 | 4.03688 | 4.08290 |
| CD630_30100 | CD630_SQ367 | 31 – 56 | 128 – 153 | -9.53762 | 4.13234 | 4.08290 |
| CD630_07400 | CD630_SQ367 | 159 – 173 | 144 – 158 | -9.53473 | 4.52273 | 4.08290 |
| CD630_17970 | CD630_SQ367 | 79 – 97 | 184 – 201 | -9.51617 | 3.68802 | 4.08290 |
| CD630_14190 | CD630_SQ367 | 139 – 156 | 16 – 34 | -9.50607 | 3.78562 | 4.08290 |
| CD630_15360 | CD630_SQ408 | 3 – 64 | 24 – 89 | -18.8638 | 4.47208 | 7.27863 |
| CD630_01590 | CD630_SQ408 | 105 – 192 | 73 – 166 | -15.0713 | 3.45609 | 7.27863 |
| bcp | CD630_SQ408 | 128 – 195 | 21 – 87 | -13.8719 | 3.16670 | 7.27863 |
| CD630_19401 | CD630_SQ408 | 102 – 142 | 145 – 187 | -13.4265 | 4.37084 | 7.27863 |
| CD630_14190 | CD630_SQ408 | 134 – 157 | 94 – 114 | -13.1343 | 3.78562 | 7.27863 |
| CD630_18940 | CD630_SQ408 | 68 – 95 | 113 – 143 | -13.1137 | 4.09988 | 7.27863 |
| CD630_06170 | CD630_SQ408 | 101 – 116 | 100 – 116 | -12.8047 | 4.95366 | 7.27863 |
| CD630_31880 | CD630_SQ408 | 122 – 186 | 41 – 114 | -12.7021 | 3.97522 | 7.27863 |
| ssuC | CD630_SQ408 | 153 – 174 | 177 – 198 | -12.529 | 3.45491 | 7.27863 |
| spoVFB | CD630_SQ408 | 167 – 192 | 169 – 191 | -11.9732 | 4.62758 | 7.27863 |
| CD630_15160 | CD630_SQ408 | 112 – 157 | 72 – 114 | -11.7004 | 4.32318 | 7.27863 |
| CD630_27570 | CD630_SQ408 | 77 – 125 | 155 – 208 | -11.5479 | 3.46216 | 7.27863 |
| cwp17 | CD630_SQ408 | 86 – 132 | 98 – 142 | -11.4983 | 3.43363 | 7.27863 |
| CD630_18610 | CD630_SQ408 | 96 – 176 | 49 – 123 | -11.4406 | 4.15582 | 7.27863 |
| CD630_21750 | CD630_SQ408 | 159 – 186 | 95 – 116 | -11.4282 | 3.70626 | 7.27863 |
| chrA | CD630_SQ408 | 58 – 87 | 145 – 169 | -11.3975 | 5.27255 | 7.27863 |
| virS | CD630_SQ408 | 145 – 193 | 97 – 146 | -11.3271 | 4.74085 | 7.27863 |
| CD630_25020 | CD630_SQ408 | 54 – 84 | 85 – 114 | -11.2743 | 4.17721 | 7.27863 |
| CD630_18400 | CD630_SQ408 | 183 – 199 | 134 – 150 | -11.2719 | 4.67962 | 7.27863 |
| CD630_18680 | CD630_SQ408 | 134 – 166 | 153 – 186 | -11.2213 | 3.04315 | 7.27863 |
| rolB | CD630_SQ408 | 106 – 131 | 5 – 29 | -11.1354 | 3.60270 | 7.27863 |
| CD630_20670 | CD630_SQ408 | 51 – 63 | 177 – 189 | -11.1168 | 4.42757 | 7.27863 |
| CD630_31010 | CD630_SQ408 | 104 – 146 | 1 – 38 | -11.0486 | 4.96305 | 7.27863 |
| cbiQ | CD630_SQ408 | 110 – 175 | 61 – 126 | -11.0338 | 3.79830 | 7.27863 |
| CD630_15890 | CD630_SQ408 | 154 – 176 | 155 – 180 | -11.0195 | 3.39962 | 7.27863 |
| pilC | CD630_SQ408 | 11 – 39 | 155 – 183 | -11.0151 | 3.54230 | 7.27863 |
| CD630_29250 | CD630_SQ408 | 122 – 140 | 18 – 36 | -10.9035 | 4.61901 | 7.27863 |
| CD630_09410 | CD630_SQ408 | 122 – 140 | 18 – 36 | -10.9035 | 4.75651 | 7.27863 |
| CD630_26250 | CD630_SQ408 | 136 – 166 | 140 – 169 | -10.8287 | 5.03379 | 7.27863 |
| CD630_20960 | CD630_SQ408 | 73 – 114 | 137 – 174 | -10.8201 | 5.65940 | 7.27863 |
| CD630_18910 | CD630_SQ408 | 180 – 197 | 170 – 188 | -10.7483 | 3.30688 | 7.27863 |
| kdgT1 | CD630_SQ408 | 46 – 73 | 1 – 26 | -10.6844 | 4.49207 | 7.27863 |
| CD630_21730 | CD630_SQ408 | 10 – 45 | 51 – 85 | -10.6371 | 3.45076 | 7.27863 |
| CD630_24050 | CD630_SQ408 | 4 – 44 | 174 – 219 | -10.6149 | 3.70192 | 7.27863 |
| CD630_11241 | CD630_SQ408 | 25 – 50 | 60 – 86 | -10.5602 | 4.85723 | 7.27863 |
| cobQ | CD630_SQ408 | 37 – 82 | 133 – 177 | -10.5286 | 3.13524 | 7.27863 |
| CD630_28280 | CD630_SQ408 | 125 – 154 | 91 – 116 | -10.515 | 4.59833 | 7.27863 |
| CD630_06140 | CD630_SQ408 | 129 – 170 | 62 – 104 | -10.5065 | 4.12519 | 7.27863 |
| CD630_16520 | CD630_SQ408 | 49 – 114 | 59 – 129 | -10.4305 | 4.28395 | 7.27863 |
| CD630_09570 | CD630_SQ408 | 116 – 152 | 173 – 209 | -10.4258 | 5.31213 | 7.27863 |
| CD630_29070 | CD630_SQ408 | 116 – 152 | 173 – 209 | -10.4258 | 3.89477 | 7.27863 |
| CD630_13930 | CD630_SQ408 | 91 – 109 | 68 – 87 | -10.2863 | 4.09246 | 7.27863 |
| CD630_18781 | CD630_SQ408 | 50 – 124 | 23 – 96 | -10.2627 | 3.77660 | 7.27863 |
| CD630_23450 | CD630_SQ408 | 22 – 64 | 77 – 107 | -10.2033 | 3.27218 | 7.27863 |
| CD630_27280 | CD630_SQ408 | 12 – 31 | 6 – 25 | -10.1921 | 3.24738 | 7.27863 |
| CD630_10960 | CD630_SQ408 | 129 – 163 | 1 – 35 | -10.1909 | 3.53850 | 7.27863 |
| CD630_31220 | CD630_SQ408 | 57 – 74 | 105 – 123 | -10.1767 | 3.73779 | 7.27863 |
| CD630_10970 | CD630_SQ408 | 31 – 48 | 98 – 116 | -10.1711 | 3.53850 | 7.27863 |
| CD630_04341 | CD630_SQ408 | 172 – 189 | 107 – 123 | -10.1683 | 4.29262 | 7.27863 |
| CD630_22660 | CD630_SQ408 | 180 – 193 | 177 – 190 | -10.1456 | 3.60496 | 7.27863 |
| CD630_27060 | CD630_SQ408 | 91 – 123 | 155 – 186 | -10.0789 | 4.06049 | 7.27863 |
| CD630_09772 | CD630_SQ408 | 2 – 44 | 70 – 109 | -9.97222 | 3.66033 | 7.27863 |
| cbpA | CD630_SQ408 | 79 – 131 | 75 – 128 | -9.96328 | 3.03040 | 7.27863 |
| CD630_27230 | CD630_SQ408 | 86 – 146 | 63 – 122 | -9.92566 | 3.52462 | 7.27863 |
| gcp | CD630_SQ408 | 125 – 155 | 2 – 31 | -9.91155 | 4.92798 | 7.27863 |
| CD630_25610 | CD630_SQ408 | 177 – 198 | 154 – 174 | -9.86128 | 3.02685 | 7.27863 |
| dapB2 | CD630_SQ408 | 169 – 198 | 154 – 183 | -9.85807 | 3.27218 | 7.27863 |
| CD630_26691 | CD630_SQ408 | 22 – 38 | 10 – 25 | -9.8569 | 3.96230 | 7.27863 |
| CD630_15740 | CD630_SQ408 | 148 – 159 | 59 – 70 | -9.83423 | 4.43222 | 7.27863 |
| CD630_19700 | CD630_SQ408 | 171 – 189 | 65 – 81 | -9.83224 | 3.85861 | 7.27863 |
| CD630_05820 | CD630_SQ408 | 180 – 195 | 98 – 112 | -9.81961 | 3.75661 | 7.27863 |
| CD630_32710 | CD630_SQ408 | 134 – 148 | 20 – 34 | -9.80256 | 4.74276 | 7.27863 |
| CD630_09770 | CD630_SQ408 | 22 – 44 | 95 – 117 | -9.79751 | 3.22106 | 7.27863 |
| CD630_25040 | CD630_SQ408 | 1 – 61 | 157 – 209 | -9.77855 | 3.43898 | 7.27863 |
| CD630_36170 | CD630_SQ408 | 157 – 166 | 76 – 85 | -9.73269 | 3.72380 | 7.27863 |
| CD630_27210 | CD630_SQ408 | 184 – 194 | 75 – 85 | -9.70292 | 4.18786 | 7.27863 |
| CD630_19850 | CD630_SQ408 | 107 – 120 | 56 – 69 | -9.65847 | 6.88486 | 7.27863 |
| CD630_24830 | CD630_SQ408 | 92 – 104 | 73 – 85 | -9.64148 | 4.98239 | 7.27863 |
| CD630_01690 | CD630_SQ408 | 58 – 84 | 53 – 81 | -9.58334 | 4.97234 | 7.27863 |
| CD630_25310 | CD630_SQ408 | 154 – 201 | 72 – 111 | -9.56478 | 3.99861 | 7.27863 |
| CD630_21090 | CD630_SQ408 | 125 – 151 | 3 – 24 | -9.51803 | 4.15803 | 7.27863 |
| CD630_19790 | RCd1 | 71 – 99 | 141 – 171 | -16.8509 | 3.79593 | 4.09988 |
| glgP | RCd1 | 12 – 45 | 143 – 171 | -13.162 | 3.28530 | 4.09988 |
| appC | RCd1 | 110 – 125 | 144 – 159 | -13.07 | 4.50438 | 4.09988 |
| CD630_06920 | RCd1 | 86 – 151 | 29 – 94 | -12.6503 | 3.13260 | 4.09988 |
| CD630_15461 | RCd1 | 182 – 198 | 146 – 162 | -12.4833 | 9.13442 | 4.09988 |
| CD630_15431 | RCd1 | 138 – 160 | 141 – 162 | -12.4482 | 5.29780 | 4.09988 |
| cbpA | RCd1 | 143 – 158 | 144 – 159 | -11.8965 | 3.03040 | 4.09988 |
| CD630_07760 | RCd1 | 182 – 197 | 142 – 158 | -11.6129 | 3.64186 | 4.09988 |
| cobQ | RCd1 | 102 – 159 | 5 – 49 | -11.5813 | 3.13524 | 4.09988 |
| CD630_28320 | RCd1 | 83 – 93 | 89 – 99 | -11.3862 | 3.99184 | 4.09988 |
| gcvTPA | RCd1 | 174 – 189 | 39 – 51 | -11.2382 | 4.77229 | 4.09988 |
| CD630_21570 | RCd1 | 171 – 183 | 142 – 154 | -11.0661 | 6.48385 | 4.09988 |
| CD630_33300 | RCd1 | 90 – 106 | 78 – 94 | -10.9905 | 3.79149 | 4.09988 |
| CD630_21460 | RCd1 | 133 – 163 | 142 – 174 | -10.9643 | 4.83005 | 4.09988 |
| CD630_19430 | RCd1 | 143 – 160 | 141 – 159 | -10.8599 | 3.13922 | 4.09988 |
| CD630_20460 | RCd1 | 155 – 185 | 141 – 169 | -10.7207 | 7.94034 | 4.09988 |
| CD630_10830 | RCd1 | 164 – 191 | 140 – 171 | -10.6727 | 4.82398 | 4.09988 |
| CD630_32710 | RCd1 | 135 – 161 | 142 – 169 | -10.6542 | 4.74276 | 4.09988 |

| | | | | | | |
|---|---|---|---|---|---|---|
| CD630_04540 | RCd1 | 3 – 13 | 2 – 12 | -10.6536 | 3.33556 | 4.09988 |
| mobA | RCd1 | 182 – 192 | 90 – 101 | -10.5761 | 4.25703 | 4.09988 |
| CD630_21540 | RCd1 | 62 – 80 | 142 – 160 | -10.5269 | 4.33090 | 4.09988 |
| CD630_32990 | RCd1 | 78 – 109 | 142 – 177 | -10.5151 | 5.24925 | 4.09988 |
| CD630_18680 | RCd1 | 97 – 137 | 140 – 171 | -10.4874 | 3.04315 | 4.09988 |
| dapB2 | RCd1 | 172 – 199 | 86 – 115 | -10.4757 | 3.27218 | 4.09988 |
| chrA | RCd1 | 31 – 72 | 149 – 182 | -10.2198 | 5.27255 | 4.09988 |
| ldh | RCd1 | 125 – 156 | 151 – 177 | -10.2055 | 4.35818 | 4.09988 |
| CD630_03220 | RCd1 | 152 – 191 | 142 – 175 | -10.0005 | 3.36746 | 4.09988 |
| CD630_19904 | RCd1 | 3 – 37 | 143 – 179 | -9.91737 | 5.96526 | 4.09988 |
| CD630_31530 | RCd1 | 168 – 189 | 146 – 164 | -9.89191 | 3.49100 | 4.09988 |
| CD630_21230 | RCd1 | 172 – 178 | 42 – 48 | -9.88299 | 3.89213 | 4.09988 |
| CD630_08660 | RCd1 | 155 – 192 | 151 – 185 | -9.86811 | 3.86467 | 4.09988 |
| cspC | RCd1 | 78 – 109 | 141 – 171 | -9.85752 | 5.48848 | 4.09988 |
| CD630_09100 | RCd1 | 11 – 75 | 3 – 44 | -9.85361 | 5.92260 | 4.09988 |
| cdu2 | RCd1 | 21 – 37 | 140 – 156 | -9.81338 | 2.97335 | 4.09988 |
| CD630_14140 | RCd1 | 148 – 168 | 144 – 164 | -9.68329 | 3.16551 | 4.09988 |
| CD630_02420 | RCd1 | 20 – 37 | 92 – 109 | -9.63622 | 3.71048 | 4.09988 |
| CD630_18970 | RCd1 | 21 – 48 | 140 – 167 | -9.59753 | 3.57243 | 4.09988 |
| CD630_21030 | RCd1 | 30 – 61 | 155 – 186 | -9.5636 | 3.08733 | 4.09988 |
| CD630_07121 | RCd1 | 11 – 29 | 155 – 173 | -9.54091 | 3.26177 | 4.09988 |
| CD630_22200 | RCd5 | 102 – 173 | 109 – 179 | -19.6136 | 4.18288 | 6.52781 |
| CD630_21770 | RCd5 | 21 – 161 | 283 – 415 | -19.1695 | 4.02587 | 6.52781 |
| CD630_05160 | RCd5 | 50 – 196 | 274 – 418 | -18.5026 | 5.03573 | 6.52781 |
| CD630_24590 | RCd5 | 4 – 148 | 284 – 428 | -18.1816 | 3.70401 | 6.52781 |
| feoB | RCd5 | 135 – 197 | 82 – 138 | -17.3971 | 4.05364 | 6.52781 |
| CD630_09100 | RCd5 | 3 – 85 | 213 – 285 | -17.1181 | 5.92260 | 6.52781 |
| CD630_09760 | RCd5 | 101 – 149 | 371 – 425 | -16.8006 | 4.83410 | 6.52781 |
| CD630_28910 | RCd5 | 101 – 149 | 371 – 425 | -16.8006 | 4.51664 | 6.52781 |
| CD630_20450 | RCd5 | 118 – 165 | 87 – 137 | -16.4579 | 3.77388 | 6.52781 |
| CD630_20460 | RCd5 | 29 – 102 | 350 – 424 | -16.3442 | 7.94034 | 6.52781 |
| CD630_21090 | RCd5 | 146 – 171 | 266 – 293 | -16.2774 | 4.15803 | 6.52781 |
| CD630_06140 | RCd5 | 16 – 92 | 107 – 181 | -16.0356 | 4.12519 | 6.52781 |
| nusB | RCd5 | 138 – 159 | 401 – 422 | -15.8899 | 4.92798 | 6.52781 |
| CD630_05870 | RCd5 | 160 – 196 | 349 – 393 | -15.7945 | 4.28339 | 6.52781 |
| nrdD | RCd5 | 73 – 106 | 390 – 419 | -15.7894 | 3.28290 | 6.52781 |
| CD630_05280 | RCd5 | 8 – 58 | 246 – 302 | -15.7554 | 3.91665 | 6.52781 |
| CD630_23930 | RCd5 | 45 – 107 | 349 – 435 | -15.6914 | 3.05650 | 6.52781 |
| cobS | RCd5 | 39 – 64 | 391 – 414 | -15.5982 | 3.67481 | 6.52781 |
| def1 | RCd5 | 111 – 188 | 353 – 423 | -15.2916 | 3.99720 | 6.52781 |
| CD630_02430 | RCd5 | 31 – 62 | 403 – 438 | -15.2123 | 3.71048 | 6.52781 |
| CD630_32710 | RCd5 | 110 – 152 | 351 – 385 | -15.1039 | 4.74276 | 6.52781 |
| CD630_23280 | RCd5 | 54 – 155 | 334 – 425 | -14.8928 | 3.50093 | 6.52781 |
| cobQ | RCd5 | 36 – 106 | 351 – 429 | -14.7344 | 3.13524 | 6.52781 |
| CD630_19580 | RCd5 | 68 – 95 | 339 – 366 | -14.7236 | 4.51902 | 6.52781 |
| CD630_23440 | RCd5 | 56 – 106 | 379 – 425 | -14.6681 | 3.34841 | 6.52781 |
| CD630_21260 | RCd5 | 138 – 163 | 32 – 59 | -14.6657 | 4.22255 | 6.52781 |
| CD630_21620 | RCd5 | 11 – 38 | 111 – 137 | -14.5028 | 3.02649 | 6.52781 |
| CD630_05290 | RCd5 | 1 – 48 | 269 – 315 | -14.4938 | 3.97604 | 6.52781 |
| thiH | RCd5 | 171 – 200 | 351 – 375 | -14.4799 | 3.58230 | 6.52781 |
| CD630_21570 | RCd5 | 157 – 186 | 111 – 137 | -14.4523 | 6.48385 | 6.52781 |
| dapB2 | RCd5 | 92 – 191 | 348 – 423 | -14.4226 | 3.27218 | 6.52781 |
| CD630_10220 | RCd5 | 113 – 171 | 181 – 237 | -14.4178 | 4.08233 | 6.52781 |
| CD630_31461 | RCd5 | 38 – 58 | 275 – 295 | -14.3623 | 4.41476 | 6.52781 |
| CD630_08080 | RCd5 | 3 – 56 | 114 – 167 | -14.3304 | 4.87819 | 6.52781 |
| CD630_24591 | RCd5 | 39 – 65 | 404 – 431 | -13.946 | 3.54497 | 6.52781 |
| kdpD | RCd5 | 152 – 201 | 88 – 134 | -13.9186 | 4.27140 | 6.52781 |
| metQ | RCd5 | 29 – 143 | 286 – 422 | -13.8159 | 3.87190 | 6.52781 |
| CD630_19700 | RCd5 | 1 – 81 | 334 – 421 | -13.5539 | 3.85861 | 6.52781 |
| glnA | RCd5 | 163 – 187 | 109 – 137 | -13.5433 | 3.87629 | 6.52781 |
| CD630_32720 | RCd5 | 64 – 94 | 349 – 378 | -13.4809 | 4.74276 | 6.52781 |
| rolB | RCd5 | 129 – 201 | 354 – 425 | -13.3581 | 3.60270 | 6.52781 |
| CD630_04850 | RCd5 | 142 – 196 | 258 – 312 | -13.2413 | 4.40788 | 6.52781 |
| folC | RCd5 | 113 – 160 | 247 – 296 | -13.2383 | 3.85634 | 6.52781 |
| CD630_12650 | RCd5 | 97 – 133 | 168 – 205 | -13.0995 | 6.31827 | 6.52781 |
| CD630_09772 | RCd5 | 56 – 88 | 407 – 435 | -13.0299 | 3.66033 | 6.52781 |
| CD630_36690 | RCd5 | 121 – 151 | 157 – 192 | -13.001 | 3.53761 | 6.52781 |
| CD630_18660 | RCd5 | 3 – 63 | 358 – 420 | -12.9568 | 3.50371 | 6.52781 |
| etfA | RCd5 | 9 – 53 | 354 – 386 | -12.8869 | 3.41204 | 6.52781 |
| CD630_13880 | RCd5 | 51 – 71 | 405 – 425 | -12.8218 | 3.52735 | 6.52781 |
| pilD | RCd5 | 3 – 25 | 109 – 132 | -12.7992 | 4.54123 | 6.52781 |
| CD630_28320 | RCd5 | 84 – 193 | 87 – 195 | -12.7801 | 3.99184 | 6.52781 |
| CD630_01560 | RCd5 | 79 – 147 | 111 – 194 | -12.772 | 4.06352 | 6.52781 |
| CD630_22070 | RCd5 | 100 – 141 | 389 – 426 | -12.7653 | 4.09651 | 6.52781 |
| CD630_27570 | RCd5 | 108 – 138 | 352 – 397 | -12.7056 | 3.46216 | 6.52781 |
| CD630_14110 | RCd5 | 17 – 57 | 144 – 179 | -12.702 | 3.39550 | 6.52781 |
| rbsA | RCd5 | 1 – 73 | 353 – 439 | -12.6623 | 3.71650 | 6.52781 |
| oppB | RCd5 | 117 – 153 | 387 – 424 | -12.6545 | 2.98328 | 6.52781 |
| CD630_20280 | RCd5 | 55 – 132 | 84 – 171 | -12.6511 | 3.46343 | 6.52781 |
| CD630_02790 | RCd5 | 82 – 165 | 286 – 359 | -12.6491 | 4.77661 | 6.52781 |
| CD630_18230 | RCd5 | 120 – 147 | 405 – 439 | -12.6489 | 3.00184 | 6.52781 |
| CD630_32520 | RCd5 | 73 – 102 | 395 – 423 | -12.6323 | 4.06098 | 6.52781 |
| CD630_17151 | RCd5 | 145 – 196 | 110 – 151 | -12.587 | 6.43124 | 6.52781 |
| cbpA | RCd5 | 163 – 199 | 274 – 312 | -12.5695 | 3.03040 | 6.52781 |
| CD630_15160 | RCd5 | 157 – 198 | 266 – 293 | -12.5633 | 4.32318 | 6.52781 |
| CD630_30140 | RCd5 | 66 – 112 | 106 – 149 | -12.5142 | 3.28343 | 6.52781 |
| CD630_15460 | RCd5 | 146 – 180 | 50 – 79 | -12.4712 | 6.86296 | 6.52781 |
| CD630_18610 | RCd5 | 185 – 200 | 263 – 278 | -12.4608 | 4.15582 | 6.52781 |
| CD630_28330 | RCd5 | 14 – 51 | 404 – 440 | -12.3486 | 3.29122 | 6.52781 |
| CD630_31530 | RCd5 | 118 – 144 | 109 – 137 | -12.2866 | 3.49100 | 6.52781 |
| CD630_25150 | RCd5 | 44 – 122 | 372 – 439 | -12.2509 | 3.79339 | 6.52781 |
| CD630_17511 | RCd5 | 11 – 34 | 54 – 79 | -12.2384 | 9.12322 | 6.52781 |
| CD630_14410 | RCd5 | 94 – 112 | 404 – 423 | -12.1998 | 3.39433 | 6.52781 |
| CD630_10970 | RCd5 | 175 – 194 | 110 – 131 | -12.1561 | 3.53850 | 6.52781 |
| CD630_21330 | RCd5 | 1 – 97 | 106 – 178 | -12.1444 | 4.10111 | 6.52781 |
| CD630_07121 | RCd5 | 24 – 54 | 157 – 190 | -12.1395 | 3.26177 | 6.52781 |
| bcp | RCd5 | 77 – 91 | 109 – 123 | -12.1063 | 3.16670 | 6.52781 |
| malY | RCd5 | 48 – 83 | 352 – 391 | -12.0765 | 4.46345 | 6.52781 |
| CD630_04410 | RCd5 | 141 – 158 | 406 – 424 | -12.0172 | 3.90451 | 6.52781 |
| ldh | RCd5 | 117 – 165 | 378 – 423 | -11.9918 | 4.35818 | 6.52781 |
| CD630_19921 | RCd5 | 34 – 65 | 404 – 434 | -11.9898 | 5.46345 | 6.52781 |
| CD630_14420 | RCd5 | 145 – 162 | 405 – 421 | -11.9867 | 3.99497 | 6.52781 |
| CD630_05830 | RCd5 | 112 – 144 | 30 – 71 | -11.9755 | 3.35360 | 6.52781 |
| CD630_06340 | RCd5 | 143 – 190 | 392 – 437 | -11.9722 | 4.35983 | 6.52781 |
| CD630_15850 | RCd5 | 13 – 39 | 391 – 423 | -11.941 | 4.32968 | 6.52781 |
| CD630_08360 | RCd5 | 161 – 187 | 395 – 425 | -11.9092 | 3.19339 | 6.52781 |
| CD630_17720 | RCd5 | 147 – 156 | 405 – 414 | -11.9008 | 3.22910 | 6.52781 |
| CD630_14250 | RCd5 | 163 – 201 | 356 – 394 | -11.8378 | 4.19604 | 6.52781 |
| CD630_27770 | RCd5 | 78 – 102 | 333 – 364 | -11.8225 | 4.08588 | 6.52781 |
| CD630_21110 | RCd5 | 119 – 154 | 126 – 163 | -11.8217 | 3.89630 | 6.52781 |
| CD630_04341 | RCd5 | 48 – 71 | 366 – 387 | -11.8197 | 4.29262 | 6.52781 |
| CD630_34520 | RCd5 | 123 – 182 | 360 – 423 | -11.8155 | 3.83016 | 6.52781 |
| sspA | RCd5 | 64 – 109 | 90 – 130 | -11.786 | 3.62689 | 6.52781 |
| CD630_25020 | RCd5 | 10 – 66 | 120 – 173 | -11.7794 | 4.17721 | 6.52781 |
| CD630_17710 | RCd5 | 58 – 141 | 351 – 425 | -11.7668 | 4.37688 | 6.52781 |
| CD630_13020 | RCd5 | 115 – 162 | 278 – 323 | -11.7541 | 3.52267 | 6.52781 |
| CD630_08130 | RCd5 | 47 – 112 | 353 – 423 | -11.7511 | 4.63831 | 6.52781 |
| gcp | RCd5 | 10 – 38 | 213 – 237 | -11.6922 | 4.92798 | 6.52781 |
| CD630_03040 | RCd5 | 88 – 105 | 407 – 423 | -11.6326 | 3.51403 | 6.52781 |
| CD630_03650 | RCd5 | 150 – 181 | 31 – 62 | -11.5862 | 3.93697 | 6.52781 |
| CD630_13960 | RCd5 | 99 – 162 | 360 – 427 | -11.5692 | 3.30375 | 6.52781 |
| CD630_35430 | RCd5 | 163 – 182 | 115 – 134 | -11.56 | 4.28798 | 6.52781 |
| CD630_26050 | RCd5 | 33 – 52 | 404 – 422 | -11.5557 | 6.99851 | 6.52781 |
| CD630_16540 | RCd5 | 167 – 190 | 399 – 422 | -11.5386 | 4.37658 | 6.52781 |
| CD630_08390 | RCd5 | 61 – 99 | 392 – 430 | -11.5107 | 5.16000 | 6.52781 |
| CD630_36781 | RCd5 | 145 – 198 | 115 – 174 | -11.4974 | 3.75946 | 6.52781 |
| CD630_17170 | RCd5 | 123 – 147 | 213 – 239 | -11.4772 | 4.10230 | 6.52781 |
| CD630_25320 | RCd5 | 132 – 162 | 405 – 439 | -11.4008 | 4.29415 | 6.52781 |
| CD630_07470 | RCd5 | 42 – 63 | 395 – 414 | -11.3923 | 3.37143 | 6.52781 |
| CD630_27360 | RCd5 | 58 – 77 | 68 – 87 | -11.3772 | 3.79570 | 6.52781 |
| CD630_06060 | RCd5 | 13 – 44 | 220 – 266 | -11.3604 | 4.94050 | 6.52781 |
| CD630_13530 | RCd5 | 156 – 194 | 281 – 321 | -11.3127 | 4.09832 | 6.52781 |
| CD630_20260 | RCd5 | 7 – 43 | 265 – 295 | -11.2952 | 3.84162 | 6.52781 |
| CD630_21490 | RCd5 | 163 – 193 | 379 – 415 | -11.2835 | 3.37462 | 6.52781 |
| CD630_21910 | RCd5 | 76 – 95 | 351 – 369 | -11.2775 | 4.66600 | 6.52781 |
| CD630_14130 | RCd5 | 58 – 122 | 350 – 423 | -11.263 | 4.57970 | 6.52781 |
| CD630_14930 | RCd5 | 55 – 140 | 325 – 395 | -11.2453 | 3.31335 | 6.52781 |
| bglA | RCd5 | 88 – 112 | 351 – 375 | -11.2286 | 4.50315 | 6.52781 |
| glgP | RCd5 | 81 – 129 | 91 – 133 | -11.2222 | 3.28530 | 6.52781 |
| CD630_27310 | RCd5 | 89 – 105 | 407 – 423 | -11.2145 | 4.01622 | 6.52781 |
| CD630_24770 | RCd5 | 168 – 178 | 401 – 411 | -11.2063 | 3.20798 | 6.52781 |
| rolA | RCd5 | 23 – 59 | 354 – 394 | -11.1891 | 4.56362 | 6.52781 |
| adhE | RCd5 | 54 – 95 | 104 – 139 | -11.1858 | 3.06485 | 6.52781 |
| CD630_05230 | RCd5 | 4 – 53 | 352 – 416 | -11.1817 | 3.73787 | 6.52781 |
| CD630_31880 | RCd5 | 2 – 14 | 215 – 227 | -11.1772 | 3.97522 | 6.52781 |
| CD630_33600 | RCd5 | 139 – 171 | 354 – 386 | -11.1725 | 3.73196 | 6.52781 |
| CD630_21020 | RCd5 | 21 – 80 | 193 – 248 | -11.1668 | 4.28666 | 6.52781 |
| CD630_31210 | RCd5 | 139 – 178 | 205 – 240 | -11.158 | 4.17693 | 6.52781 |
| CD630_16330 | RCd5 | 154 – 198 | 124 – 162 | -11.1498 | 4.11180 | 6.52781 |
| CD630_06110 | RCd5 | 168 – 188 | 404 – 423 | -11.1223 | 5.61380 | 6.52781 |
| CD630_07960 | RCd5 | 180 – 201 | 274 – 295 | -11.1193 | 3.22537 | 6.52781 |
| CD630_27600 | RCd5 | 31 – 66 | 402 – 440 | -11.1145 | 3.48384 | 6.52781 |
| CD630_30310 | RCd5 | 142 – 167 | 114 – 140 | -11.1123 | 4.56059 | 6.52781 |
| CD630_15360 | RCd5 | 86 – 121 | 391 – 425 | -11.1061 | 4.47208 | 6.52781 |
| CD630_22050 | RCd5 | 170 – 194 | 299 – 325 | -11.0835 | 4.87639 | 6.52781 |
| dltB | RCd5 | 6 – 30 | 124 – 148 | -11.0654 | 3.81485 | 6.52781 |
| CD630_21700 | RCd5 | 20 – 41 | 119 – 145 | -11.0577 | 3.96650 | 6.52781 |
| CD630_21540 | RCd5 | 47 – 70 | 215 – 238 | -11.0441 | 4.33090 | 6.52781 |
| CD630_08140 | RCd5 | 9 – 124 | 36 – 143 | -11.0416 | 3.30159 | 6.52781 |
| CD630_06560 | RCd5 | 4 – 19 | 290 – 306 | -11.04 | 4.02532 | 6.52781 |
| pilC | RCd5 | 175 – 184 | 404 – 413 | -11.0222 | 3.54230 | 6.52781 |
| srlD | RCd5 | 84 – 106 | 334 – 358 | -10.9999 | 4.19843 | 6.52781 |
| CD630_27280 | RCd5 | 45 – 190 | 258 – 406 | -10.9969 | 3.24738 | 6.52781 |
| CD630_34530 | RCd5 | 86 – 159 | 352 – 415 | -10.953 | 3.83016 | 6.52781 |
| CD630_23080 | RCd5 | 1 – 88 | 257 – 347 | -10.9006 | 5.23027 | 6.52781 |
| CD630_13580 | RCd5 | 110 – 141 | 387 – 423 | -10.8611 | 3.80715 | 6.52781 |
| CD630_32070 | RCd5 | 50 – 79 | 104 – 134 | -10.8565 | 3.51964 | 6.52781 |
| CD630_20670 | RCd5 | 179 – 194 | 74 – 89 | -10.8089 | 4.42757 | 6.52781 |
| CD630_25040 | RCd5 | 144 – 177 | 27 – 62 | -10.8006 | 3.43898 | 6.52781 |
| CD630_18920 | RCd5 | 96 – 154 | 370 – 430 | -10.7897 | 4.43558 | 6.52781 |
| CD630_08950 | RCd5 | 156 – 191 | 276 – 313 | -10.7832 | 3.71905 | 6.52781 |
| CD630_08460 | RCd5 | 89 – 136 | 351 – 424 | -10.7827 | 4.15083 | 6.52781 |
| iorA | RCd5 | 53 – 92 | 351 – 397 | -10.7508 | 3.83173 | 6.52781 |
| CD630_06080 | RCd5 | 7 – 29 | 106 – 130 | -10.7059 | 3.82315 | 6.52781 |
| cspC | RCd5 | 91 – 141 | 352 – 393 | -10.6803 | 5.48848 | 6.52781 |
| CD630_29500 | RCd5 | 43 – 60 | 171 – 187 | -10.6636 | 3.70644 | 6.52781 |
| CD630_23780 | RCd5 | 82 – 152 | 346 – 404 | -10.6625 | 3.91629 | 6.52781 |
| CD630_22950 | RCd5 | 3 – 25 | 245 – 269 | -10.6522 | 3.96883 | 6.52781 |
| CD630_02720 | RCd5 | 101 – 148 | 355 – 411 | -10.6505 | 3.38938 | 6.52781 |
| CD630_33420 | RCd5 | 11 – 76 | 324 – 422 | -10.6415 | 3.22095 | 6.52781 |
| CD630_08370 | RCd5 | 2 – 26 | 110 – 135 | -10.6261 | 3.65639 | 6.52781 |
| CD630_24220 | RCd5 | 126 – 169 | 211 – 255 | -10.5927 | 4.18037 | 6.52781 |
| CD630_33570 | RCd5 | 150 – 162 | 227 – 239 | -10.5897 | 3.95778 | 6.52781 |
| CD630_04610 | RCd5 | 18 – 45 | 354 – 377 | -10.5784 | 3.83025 | 6.52781 |
| cspD | RCd5 | 89 – 169 | 335 – 409 | -10.5604 | 3.23587 | 6.52781 |
| CD630_31910 | RCd5 | 88 – 155 | 361 – 423 | -10.5053 | 3.48314 | 6.52781 |
| CD630_32050 | RCd5 | 176 – 193 | 404 – 423 | -10.4939 | 5.73797 | 6.52781 |
| CD630_31490 | RCd5 | 67 – 129 | 360 – 417 | -10.453 | 4.08593 | 6.52781 |
| CD630_19850 | RCd5 | 79 – 99 | 401 – 421 | -10.4351 | 6.88486 | 6.52781 |
| CD630_06050 | RCd5 | 87 – 170 | 353 – 424 | -10.4212 | 3.59332 | 6.52781 |
| CD630_11240 | RCd5 | 4 – 45 | 372 – 418 | -10.4177 | 3.20274 | 6.52781 |
| xdhA | RCd5 | 3 – 148 | 102 – 231 | -10.4071 | 4.45147 | 6.52781 |
| CD630_18120 | RCd5 | 6 – 62 | 115 – 164 | -10.3962 | 3.99713 | 6.52781 |
| CD630_27270 | RCd5 | 93 – 105 | 227 – 239 | -10.3748 | 3.50892 | 6.52781 |
| CD630_19690 | RCd5 | 112 – 182 | 355 – 423 | -10.3642 | 3.85861 | 6.52781 |
| CD630_22550 | RCd5 | 141 – 193 | 260 – 327 | -10.3464 | 3.62379 | 6.52781 |
| CD630_10880 | RCd5 | 167 – 181 | 404 – 422 | -10.3339 | 5.05048 | 6.52781 |
| CD630_25050 | RCd5 | 153 – 164 | 346 – 357 | -10.3254 | 4.33624 | 6.52781 |
| CD630_14050 | RCd5 | 66 – 164 | 337 – 424 | -10.311 | 3.30894 | 6.52781 |
| agrB | RCd5 | 129 – 172 | 404 – 439 | -10.2907 | 3.92598 | 6.52781 |
| CD630_28150 | RCd5 | 152 – 201 | 347 – 400 | -10.2893 | 4.18447 | 6.52781 |
| CD630_23770 | RCd5 | 42 – 144 | 352 – 427 | -10.251 | 3.25181 | 6.52781 |
| CD630_26750 | RCd5 | 22 – 61 | 111 – 149 | -10.2397 | 3.60610 | 6.52781 |
| oppC | RCd5 | 164 – 182 | 111 – 131 | -10.2387 | 3.01695 | 6.52781 |
| CD630_06200 | RCd5 | 44 – 66 | 286 – 313 | -10.2357 | 3.57064 | 6.52781 |
| CD630_27230 | RCd5 | 90 – 121 | 353 – 375 | -10.2298 | 3.52462 | 6.52781 |
| CD630_22660 | RCd5 | 134 – 201 | 78 – 137 | -10.2274 | 3.60496 | 6.52781 |
| CD630_15111 | RCd5 | 37 – 177 | 166 – 297 | -10.2144 | 4.18664 | 6.52781 |
| CD630_15431 | RCd5 | 139 – 153 | 284 – 298 | -10.2117 | 5.29780 | 6.52781 |
| CD630_10000 | RCd5 | 100 – 107 | 405 – 412 | -10.1968 | 3.86040 | 6.52781 |
| CD630_04550 | RCd5 | 95 – 131 | 212 – 247 | -10.1874 | 3.05227 | 6.52781 |
| CD630_10050 | RCd5 | 176 – 198 | 354 – 373 | -10.1853 | 3.93750 | 6.52781 |
| CD630_06170 | RCd5 | 117 – 136 | 113 – 140 | -10.1807 | 4.95366 | 6.52781 |
| CD630_06190 | RCd5 | 141 – 151 | 49 – 59 | -10.166 | 3.57064 | 6.52781 |
| CD630_25190 | RCd5 | 174 – 181 | 87 – 94 | -10.1564 | 3.19663 | 6.52781 |
| CD630_17730 | RCd5 | 76 – 97 | 104 – 124 | -10.1393 | 4.73397 | 6.52781 |
| CD630_27120 | RCd5 | 3 – 36 | 226 – 262 | -10.1229 | 4.43720 | 6.52781 |
| cls | RCd5 | 45 – 77 | 395 – 428 | -10.1206 | 4.46177 | 6.52781 |
| CD630_18960 | RCd5 | 4 – 28 | 109 – 134 | -10.1036 | 4.37822 | 6.52781 |
| rbsC | RCd5 | 34 – 51 | 335 – 352 | -10.1018 | 3.71650 | 6.52781 |
| CD630_34050 | RCd5 | 8 – 20 | 5 – 17 | -10.0935 | 4.03863 | 6.52781 |
| CD630_27610 | RCd5 | 15 – 28 | 234 – 248 | -10.0861 | 3.96465 | 6.52781 |
| CD630_08660 | RCd5 | 148 – 186 | 213 – 251 | -10.0721 | 3.86467 | 6.52781 |
| CD630_11241 | RCd5 | 159 – 184 | 235 – 259 | -10.0618 | 4.85723 | 6.52781 |
| CD630_23070 | RCd5 | 172 – 187 | 83 – 97 | -10.0583 | 5.23027 | 6.52781 |
| CD630_28580 | RCd5 | 127 – 147 | 395 – 415 | -10.0574 | 4.67130 | 6.52781 |
| CD630_14610 | RCd5 | 146 – 177 | 258 – 298 | -10.0534 | 4.89428 | 6.52781 |
| fliI | RCd5 | 51 – 107 | 197 – 245 | -10.0497 | 3.03666 | 6.52781 |
| CD630_06060 | RCd5 | 162 – 184 | 404 – 425 | -10.0435 | 4.94050 | 6.52781 |
| CD630_04940 | RCd5 | 99 – 123 | 405 – 427 | -10.0311 | 4.20948 | 6.52781 |
| CD630_05950 | RCd5 | 19 – 50 | 112 – 142 | -10.0295 | 3.31669 | 6.52781 |
| CD630_07760 | RCd5 | 2 – 14 | 405 – 417 | -10.0239 | 3.64186 | 6.52781 |
| CD630_25070 | RCd5 | 106 – 131 | 111 – 135 | -10.0238 | 3.45164 | 6.52781 |
| CD630_19904 | RCd5 | 25 – 37 | 127 – 141 | -10.0162 | 5.96526 | 6.52781 |
| CD630_30100 | RCd5 | 176 – 186 | 14 – 24 | -9.9901 | 4.13234 | 6.52781 |
| pilT | RCd5 | 84 – 141 | 356 – 426 | -9.97142 | 3.24639 | 6.52781 |
| CD630_30730 | RCd5 | 118 – 130 | 265 – 277 | -9.942 | 3.13236 | 6.52781 |
| CD630_19670 | RCd5 | 72 – 87 | 351 – 367 | -9.94146 | 3.20718 | 6.52781 |
| opuCC | RCd5 | 184 – 197 | 169 – 183 | -9.9252 | 4.07035 | 6.52781 |
| vncR | RCd5 | 110 – 143 | 406 – 440 | -9.92161 | 3.93642 | 6.52781 |
| CD630_18100 | RCd5 | 128 – 152 | 399 – 423 | -9.89591 | 5.71658 | 6.52781 |
| CD630_10830 | RCd5 | 61 – 93 | 352 – 383 | -9.88694 | 4.82398 | 6.52781 |
| CD630_22500 | RCd5 | 149 – 195 | 95 – 148 | -9.88528 | 4.03688 | 6.52781 |
| CD630_20171 | RCd5 | 143 – 178 | 379 – 422 | -9.8695 | 6.87491 | 6.52781 |
| dnaC | RCd5 | 47 – 123 | 354 – 430 | -9.85273 | 6.30796 | 6.52781 |
| CD630_06970 | RCd5 | 110 – 152 | 211 – 256 | -9.83701 | 3.82312 | 6.52781 |
| CD630_31960 | RCd5 | 31 – 52 | 371 – 393 | -9.83273 | 4.93757 | 6.52781 |
| CD630_07870 | RCd5 | 184 – 199 | 158 – 174 | -9.82911 | 3.22829 | 6.52781 |
| CD630_05900 | RCd5 | 19 – 50 | 400 – 439 | -9.81236 | 3.21934 | 6.52781 |
| CD630_28181 | RCd5 | 14 – 76 | 87 – 137 | -9.81231 | 3.91299 | 6.52781 |
| CD630_05220 | RCd5 | 67 – 79 | 335 – 347 | -9.79007 | 3.65057 | 6.52781 |
| CD630_19401 | RCd5 | 26 – 107 | 92 – 172 | -9.77057 | 4.37084 | 6.52781 |
| CD630_06410 | RCd5 | 86 – 130 | 247 – 292 | -9.76527 | 4.14404 | 6.52781 |
| CD630_09970 | RCd5 | 8 – 29 | 113 – 136 | -9.76199 | 3.25917 | 6.52781 |
| eutW | RCd5 | 13 – 20 | 405 – 412 | -9.76003 | 4.55715 | 6.52781 |
| gcvTPA | RCd5 | 26 – 109 | 335 – 428 | -9.75968 | 4.77229 | 6.52781 |
| CD630_13790 | RCd5 | 102 – 132 | 399 – 435 | -9.75608 | 5.17374 | 6.52781 |
| CD630_29960 | RCd5 | 137 – 168 | 114 – 137 | -9.7354 | 4.51904 | 6.52781 |
| cwp26 | RCd5 | 138 – 160 | 213 – 236 | -9.73218 | 4.44377 | 6.52781 |
| CD630_17770 | RCd5 | 76 – 104 | 336 – 364 | -9.73126 | 5.06490 | 6.52781 |
| CD630_24830 | RCd5 | 139 – 171 | 257 – 285 | -9.72123 | 4.98239 | 6.52781 |
| CD630_26680 | RCd5 | 126 – 160 | 372 – 414 | -9.70221 | 3.53956 | 6.52781 |
| virS | RCd5 | 89 – 123 | 397 – 423 | -9.69906 | 4.74085 | 6.52781 |
| CD630_02740 | RCd5 | 88 – 150 | 286 – 355 | -9.6913 | 5.02277 | 6.52781 |
| CD630_14990 | RCd5 | 120 – 151 | 220 – 266 | -9.68788 | 3.78619 | 6.52781 |
| CD630_32690 | RCd5 | 86 – 148 | 104 – 159 | -9.65859 | 7.32676 | 6.52781 |
| CD630_32890 | RCd5 | 188 – 200 | 189 – 201 | -9.64358 | 3.41276 | 6.52781 |
| cdtR | RCd5 | 120 – 137 | 397 – 414 | -9.63231 | 5.84368 | 6.52781 |
| CD630_06060 | RCd5 | 174 – 193 | 402 – 421 | -9.62771 | 4.94050 | 6.52781 |
| ssuC | RCd5 | 54 – 68 | 350 – 364 | -9.62723 | 3.45491 | 6.52781 |
| pgm | RCd5 | 118 – 140 | 349 – 379 | -9.61648 | 3.45439 | 6.52781 |
| CD630_21070 | RCd5 | 21 – 54 | 401 – 440 | -9.60373 | 5.35019 | 6.52781 |
| CD630_26150 | RCd5 | 42 – 90 | 117 – 164 | -9.60062 | 4.06302 | 6.52781 |
| appC | RCd5 | 39 – 58 | 73 – 88 | -9.58936 | 4.50438 | 6.52781 |
| CD630_33560 | RCd5 | 72 – 94 | 405 – 431 | -9.58736 | 4.03147 | 6.52781 |
| CD630_14630 | RCd5 | 79 – 91 | 352 – 364 | -9.58404 | 3.34485 | 6.52781 |
| CD630_20541 | RCd5 | 89 – 113 | 350 – 374 | -9.56404 | 5.74527 | 6.52781 |
| CD630_21730 | RCd5 | 131 – 166 | 109 – 136 | -9.56137 | 3.45076 | 6.52781 |
| dpaA | RCd5 | 163 – 188 | 344 – 364 | -9.55908 | 4.65364 | 6.52781 |
| cobC | RCd5 | 169 – 195 | 351 – 378 | -9.54129 | 3.15854 | 6.52781 |
| CD630_36040 | RCd8 | 18 – 54 | 49 – 84 | -16.2788 | 3.52009 | 6.97567 |
| CD630_32060 | RCd8 | 163 – 194 | 55 – 86 | -15.5556 | 5.73797 | 6.97567 |
| CD630_08460 | RCd8 | 134 – 157 | 5 – 26 | -15.2979 | 4.15083 | 6.97567 |
| oppB | RCd8 | 80 – 103 | 177 – 201 | -14.9823 | 2.98328 | 6.97567 |
| CD630_07770 | RCd8 | 82 – 109 | 55 – 82 | -14.8075 | 3.31158 | 6.97567 |
| CD630_05030 | RCd8 | 148 – 169 | 186 – 208 | -14.5382 | 4.03823 | 6.97567 |
| CD630_27570 | RCd8 | 178 – 196 | 55 – 73 | -14.4303 | 3.46216 | 6.97567 |
| CD630_16940 | RCd8 | 130 – 179 | 52 – 109 | -13.8937 | 4.14665 | 6.97567 |
| fliI | RCd8 | 179 – 199 | 50 – 72 | -13.0968 | 3.03666 | 6.97567 |
| CD630_14990 | RCd8 | 177 – 198 | 48 – 72 | -12.9522 | 3.78619 | 6.97567 |
| CD630_05821 | RCd8 | 180 – 197 | 150 – 167 | -12.612 | 3.75661 | 6.97567 |
| CD630_07480 | RCd8 | 1 – 98 | 63 – 156 | -12.5404 | 5.18321 | 6.97567 |
| CD630_19680 | RCd8 | 39 – 64 | 50 – 71 | -12.4309 | 3.58080 | 6.97567 |
| CD630_17170 | RCd8 | 118 – 129 | 217 – 228 | -12.3342 | 4.10230 | 6.97567 |
| CD630_20450 | RCd8 | 146 – 186 | 57 – 84 | -12.302 | 3.77388 | 6.97567 |
| CD630_03420 | RCd8 | 105 – 123 | 51 – 69 | -12.2936 | 4.71868 | 6.97567 |
| CD630_10960 | RCd8 | 6 – 40 | 55 – 94 | -12.2833 | 3.53850 | 6.97567 |
| CD630_14810 | RCd8 | 125 – 165 | 92 – 132 | -12.192 | 3.35323 | 6.97567 |
| CD630_21720 | RCd8 | 155 – 173 | 52 – 72 | -12.091 | 4.13030 | 6.97567 |
| CD630_24590 | RCd8 | 134 – 201 | 144 – 202 | -12.0226 | 3.70401 | 6.97567 |
| CD630_02430 | RCd8 | 68 – 88 | 52 – 73 | -11.9822 | 3.71048 | 6.97567 |
| CD630_23760 | RCd8 | 179 – 194 | 67 – 82 | -11.9672 | 4.54307 | 6.97567 |
| CD630_10220 | RCd8 | 116 – 147 | 4 – 30 | -11.8219 | 4.08233 | 6.97567 |
| CD630_09760 | RCd8 | 74 – 123 | 146 – 202 | -11.7665 | 4.83410 | 6.97567 |
| CD630_28910 | RCd8 | 74 – 123 | 146 – 202 | -11.7665 | 4.51664 | 6.97567 |
| adhE | RCd8 | 98 – 119 | 63 – 87 | -11.6955 | 3.06485 | 6.97567 |
| CD630_16530 | RCd8 | 33 – 44 | 11 – 22 | -11.5975 | 4.37658 | 6.97567 |
| CD630_28890 | RCd8 | 31 – 53 | 188 – 204 | -11.5764 | 7.52469 | 6.97567 |
| CD630_26050 | RCd8 | 166 – 186 | 49 – 70 | -11.5642 | 6.99851 | 6.97567 |
| CD630_28181 | RCd8 | 31 – 65 | 47 – 79 | -11.4232 | 3.91299 | 6.97567 |
| CD630_19903 | RCd8 | 23 – 71 | 86 – 139 | -11.3583 | 5.79994 | 6.97567 |
| dltB | RCd8 | 14 – 73 | 108 – 164 | -11.3574 | 3.81485 | 6.97567 |
| CD630_05450 | RCd8 | 4 – 20 | 56 – 73 | -11.1065 | 5.04761 | 6.97567 |
| CD630_24580 | RCd8 | 85 – 94 | 218 – 227 | -11.0866 | 3.88519 | 6.97567 |
| CD630_19800 | RCd8 | 119 – 141 | 49 – 69 | -11.0245 | 5.22035 | 6.97567 |
| CD630_24920 | RCd8 | 86 – 97 | 217 – 228 | -10.839 | 3.89455 | 6.97567 |
| CD630_25430 | RCd8 | 36 – 75 | 55 – 94 | -10.7781 | 4.00527 | 6.97567 |
| CD630_29760 | RCd8 | 113 – 144 | 56 – 88 | -10.7669 | 3.51074 | 6.97567 |
| panG | RCd8 | 31 – 68 | 52 – 88 | -10.6831 | 5.54047 | 6.97567 |
| CD630_31690 | RCd8 | 4 – 25 | 49 – 71 | -10.5625 | 3.71193 | 6.97567 |
| bglA | RCd8 | 94 – 111 | 148 – 166 | -10.5431 | 4.50315 | 6.97567 |
| CD630_09771 | RCd8 | 167 – 190 | 200 – 228 | -10.5262 | 4.95362 | 6.97567 |
| CD630_22050 | RCd8 | 190 – 200 | 72 – 82 | -10.4688 | 4.87639 | 6.97567 |
| CD630_09850 | RCd8 | 169 – 188 | 72 – 94 | -10.4584 | 4.18679 | 6.97567 |
| CD630_30360 | RCd8 | 79 – 108 | 127 – 154 | -10.4427 | 4.26509 | 6.97567 |
| CD630_15740 | RCd8 | 136 – 156 | 65 – 87 | -10.4091 | 4.43222 | 6.97567 |
| CD630_29650 | RCd8 | 34 – 49 | 67 – 82 | -10.3276 | 3.67128 | 6.97567 |
| CD630_05230 | RCd8 | 3 – 20 | 99 – 118 | -10.2962 | 3.73787 | 6.97567 |
| CD630_15900 | RCd8 | 1 – 15 | 15 – 29 | -10.2485 | 3.17664 | 6.97567 |
| CD630_27520 | RCd8 | 130 – 152 | 49 – 70 | -10.2469 | 4.27693 | 6.97567 |
| CD630_17900 | RCd8 | 190 – 198 | 65 – 73 | -10.2304 | 3.52167 | 6.97567 |
| nusB | RCd8 | 2 – 11 | 192 – 201 | -10.2041 | 4.92798 | 6.97567 |
| CD630_25200 | RCd8 | 128 – 142 | 58 – 72 | -10.2002 | 3.65158 | 6.97567 |
| CD630_12650 | RCd8 | 116 – 125 | 218 – 227 | -10.1902 | 6.31827 | 6.97567 |
| CD630_14700 | RCd8 | 60 – 69 | 63 – 72 | -10.1723 | 3.98234 | 6.97567 |
| CD630_20650 | RCd8 | 8 – 29 | 52 – 74 | -10.1713 | 4.00083 | 6.97567 |
| CD630_30190 | RCd8 | 132 – 147 | 3 – 19 | -10.1562 | 3.27394 | 6.97567 |
| CD630_25150 | RCd8 | 168 – 192 | 53 – 79 | -10.1306 | 3.79339 | 6.97567 |
| pmi | RCd8 | 79 – 97 | 65 – 84 | -10.0993 | 3.86845 | 6.97567 |
| CD630_16950 | RCd8 | 185 – 199 | 187 – 201 | -10.0966 | 4.42569 | 6.97567 |
| CD630_27110 | RCd8 | 158 – 167 | 198 – 207 | -10.0861 | 3.78231 | 6.97567 |
| rbsC | RCd8 | 46 – 71 | 49 – 72 | -10.0774 | 3.71650 | 6.97567 |
| CD630_20460 | RCd8 | 159 – 176 | 52 – 68 | -10.0526 | 7.94034 | 6.97567 |
| CD630_09100 | RCd8 | 165 – 184 | 50 – 69 | -10.0467 | 5.92260 | 6.97567 |
| CD630_10950 | RCd8 | 91 – 110 | 53 – 72 | -10.0186 | 3.67675 | 6.97567 |
| CD630_15200 | RCd8 | 93 – 101 | 9 – 17 | -10.0031 | 4.00442 | 6.97567 |
| CD630_13790 | RCd8 | 158 – 182 | 54 – 80 | -9.94943 | 5.17374 | 6.97567 |
| CD630_28320 | RCd8 | 157 – 185 | 42 – 70 | -9.93848 | 3.99184 | 6.97567 |
| CD630_09060 | RCd8 | 4 – 19 | 14 – 29 | -9.93697 | 4.31834 | 6.97567 |
| bltD | RCd8 | 89 – 97 | 197 – 205 | -9.92955 | 4.49926 | 6.97567 |
| CD630_19700 | RCd8 | 168 – 180 | 55 – 68 | -9.90628 | 3.85861 | 6.97567 |
| cwp17 | RCd8 | 4 – 51 | 54 – 88 | -9.86145 | 3.43363 | 6.97567 |
| tcdA | RCd8 | 50 – 84 | 50 – 73 | -9.85237 | 4.70593 | 6.97567 |
| sigK | RCd8 | 118 – 126 | 65 – 73 | -9.82434 | 3.08122 | 6.97567 |
| CD630_01700 | RCd8 | 138 – 148 | 64 – 74 | -9.81464 | 4.97234 | 6.97567 |
| CD630_13560 | RCd8 | 190 – 200 | 74 – 84 | -9.78794 | 3.38083 | 6.97567 |
| CD630_22660 | RCd8 | 142 – 187 | 50 – 84 | -9.76088 | 3.60496 | 6.97567 |
| CD630_06060 | RCd8 | 63 – 80 | 150 – 167 | -9.74468 | 4.94050 | 6.97567 |
| CD630_06340 | RCd8 | 2 – 12 | 216 – 226 | -9.73032 | 4.35983 | 6.97567 |
| CD630_13400 | RCd8 | 148 – 167 | 55 – 73 | -9.72545 | 3.20232 | 6.97567 |
| CD630_05211 | RCd8 | 24 – 43 | 141 – 160 | -9.67905 | 3.97048 | 6.97567 |
| CD630_14240 | RCd8 | 23 – 114 | 52 – 140 | -9.65433 | 8.60766 | 6.97567 |
| CD630_21770 | RCd8 | 26 – 41 | 59 – 73 | -9.61351 | 4.02587 | 6.97567 |
| CD630_33420 | RCd8 | 149 – 172 | 67 – 93 | -9.59777 | 3.22095 | 6.97567 |
| CD630_22510 | RCd8 | 175 – 196 | 48 – 70 | -9.59044 | 11.42763 | 6.97567 |
| dapB2 | RCd8 | 189 – 198 | 11 – 20 | -9.57887 | 3.27218 | 6.97567 |
| CD630_14930 | RCd8 | 9 – 26 | 77 – 95 | -9.57796 | 3.31335 | 6.97567 |
| argG | RCd8 | 71 – 104 | 52 – 87 | -9.56177 | 4.80300 | 6.97567 |
| CD630_30130 | RCd8 | 49 – 67 | 50 – 72 | -9.53467 | 3.28343 | 6.97567 |
| CD630_18960 | RCd8 | 12 – 33 | 48 – 70 | -9.53241 | 4.37822 | 6.97567 |
| cheW | RCd8 | 24 – 56 | 133 – 166 | -9.5071 | 3.72371 | 6.97567 |
| CD630_09562 | RCd9 | 52 – 201 | 115 – 264 | -179.115 | 3.94307 | 8.96925 |
| CD630_15550 | RCd9 | 84 – 135 | 181 – 236 | -19.3702 | 3.49019 | 8.96925 |
| CD630_05880 | RCd9 | 89 – 173 | 148 – 229 | -19.0404 | 4.28339 | 8.96925 |
| CD630_29250 | RCd9 | 80 – 108 | 208 – 237 | -18.7095 | 4.61901 | 8.96925 |

| | | | | | | |
|---|---|---|---|---|---|---|
| CD630_09410 | RCd9 | 80 – 108 | 208 – 237 | -18.7095 | 4.75651 | 8.96925 |
| flgB | RCd9 | 88 – 107 | 209 – 229 | -18.5675 | 3.25736 | 8.96925 |
| CD630_08940 | RCd9 | 66 – 94 | 264 – 299 | -18.4558 | 3.72225 | 8.96925 |
| CD630_04341 | RCd9 | 159 – 182 | 209 – 232 | -18.2093 | 4.29262 | 8.96925 |
| CD630_18440 | RCd9 | 93 – 200 | 185 – 298 | -17.3486 | 4.07960 | 8.96925 |
| CD630_14920 | RCd9 | 89 – 114 | 211 – 235 | -16.9107 | 4.35225 | 8.96925 |
| cbpA | RCd9 | 85 – 97 | 218 – 231 | -16.9032 | 3.03040 | 8.96925 |
| CD630_06920 | RCd9 | 81 – 157 | 159 – 230 | -16.4272 | 3.13260 | 8.96925 |
| bglA | RCd9 | 159 – 186 | 89 – 115 | -15.9701 | 4.86402 | 8.96925 |
| CD630_19690 | RCd9 | 107 – 129 | 205 – 230 | -15.936 | 3.85861 | 8.96925 |
| fliI | RCd9 | 39 – 110 | 135 – 193 | -15.8409 | 3.03666 | 8.96925 |
| CD630_17970 | RCd9 | 114 – 135 | 253 – 275 | -15.7487 | 3.68802 | 8.96925 |
| srlD | RCd9 | 126 – 150 | 209 – 230 | -15.667 | 4.19843 | 8.96925 |
| CD630_32520 | RCd9 | 94 – 191 | 168 – 241 | -15.6446 | 4.06098 | 8.96925 |
| CD630_16450 | RCd9 | 79 – 105 | 211 – 236 | -15.6406 | 4.57547 | 8.96925 |
| CD630_13930 | RCd9 | 87 – 109 | 207 – 232 | -15.512 | 4.09246 | 8.96925 |
| CD630_25190 | RCd9 | 155 – 191 | 245 – 268 | -15.4627 | 3.19663 | 8.96925 |
| terD | RCd9 | 181 – 198 | 208 – 229 | -15.4115 | 3.33148 | 8.96925 |
| CD630_09040 | RCd9 | 155 – 191 | 245 – 268 | -15.4113 | 4.11851 | 8.96925 |
| CD630_17170 | RCd9 | 159 – 176 | 215 – 232 | -15.3619 | 4.10230 | 8.96925 |
| CD630_05230 | RCd9 | 3 – 56 | 249 – 298 | -15.3377 | 3.73787 | 8.96925 |
| serS1 | RCd9 | 155 – 196 | 186 – 232 | -15.1554 | 3.45858 | 8.96925 |
| CD630_15890 | RCd9 | 86 – 163 | 153 – 230 | -14.9858 | 3.39962 | 8.96925 |
| CD630_04940 | RCd9 | 140 – 156 | 283 – 298 | -14.686 | 4.20948 | 8.96925 |
| CD630_20670 | RCd9 | 87 – 164 | 139 – 226 | -14.6256 | 4.42757 | 8.96925 |
| CD630_08390 | RCd9 | 82 – 105 | 247 – 267 | -14.5917 | 5.16000 | 8.96925 |
| CD630_29750 | RCd9 | 62 – 97 | 215 – 236 | -14.5639 | 4.59146 | 8.96925 |
| CD630_10800 | RCd9 | 84 – 105 | 211 – 232 | -14.5316 | 5.30319 | 8.96925 |
| CD630_23440 | RCd9 | 55 – 86 | 210 – 235 | -14.5301 | 3.34841 | 8.96925 |
| CD630_31910 | RCd9 | 85 – 95 | 215 – 226 | -14.4216 | 3.48314 | 8.96925 |
| CD630_33420 | RCd9 | 6 – 43 | 263 – 298 | -14.3776 | 3.22095 | 8.96925 |
| CD630_15850 | RCd9 | 125 – 162 | 205 – 228 | -14.2512 | 4.32968 | 8.96925 |
| srlEb | RCd9 | 44 – 96 | 217 – 254 | -14.1511 | 3.50140 | 8.96925 |
| CD630_17730 | RCd9 | 87 – 105 | 211 – 226 | -14.1504 | 4.73397 | 8.96925 |
| CD630_30420 | RCd9 | 88 – 172 | 155 – 229 | -14.1424 | 5.27539 | 8.96925 |
| CD630_27040 | RCd9 | 66 – 100 | 129 – 164 | -14.1312 | 3.51812 | 8.96925 |
| cobQ | RCd9 | 86 – 107 | 211 – 230 | -14.1023 | 3.13524 | 8.96925 |
| ung | RCd9 | 80 – 111 | 205 – 233 | -13.9514 | 3.30908 | 8.96925 |
| CD630_19680 | RCd9 | 33 – 117 | 187 – 281 | -13.8985 | 3.58080 | 8.96925 |
| CD630_13960 | RCd9 | 138 – 162 | 245 – 268 | -13.8878 | 3.30375 | 8.96925 |
| CD630_07480 | RCd9 | 86 – 111 | 211 – 231 | -13.8813 | 5.18321 | 8.96925 |
| CD630_06060 | RCd9 | 84 – 105 | 211 – 235 | -13.8197 | 4.94050 | 8.96925 |
| CD630_33300 | RCd9 | 183 – 196 | 219 – 234 | -13.7929 | 3.79149 | 8.96925 |
| CD630_15360 | RCd9 | 94 – 126 | 268 – 299 | -13.5963 | 4.47208 | 8.96925 |
| CD630_22880 | RCd9 | 43 – 105 | 220 – 267 | -13.5789 | 3.62455 | 8.96925 |
| CD630_18910 | RCd9 | 79 – 108 | 184 – 209 | -13.5742 | 3.30688 | 8.96925 |
| cspD | RCd9 | 80 – 169 | 130 – 229 | -13.554 | 3.23587 | 8.96925 |
| CD630_24590 | RCd9 | 90 – 128 | 263 – 298 | -13.5438 | 3.70401 | 8.96925 |
| CD630_08130 | RCd9 | 28 – 108 | 208 – 298 | -13.4968 | 4.63831 | 8.96925 |
| blaI | RCd9 | 87 – 105 | 211 – 231 | -13.4697 | 3.73674 | 8.96925 |
| CD630_23820 | RCd9 | 69 – 95 | 266 – 297 | -13.4393 | 3.18419 | 8.96925 |
| oppB | RCd9 | 88 – 137 | 168 – 229 | -13.42 | 2.98328 | 8.96925 |
| CD630_35840 | RCd9 | 77 – 92 | 222 – 236 | -13.3933 | 3.14170 | 8.96925 |
| CD630_23280 | RCd9 | 103 – 123 | 208 – 230 | -13.3912 | 3.50093 | 8.96925 |
| pgm | RCd9 | 84 – 104 | 253 – 273 | -13.3896 | 3.45439 | 8.96925 |
| moaA | RCd9 | 87 – 109 | 207 – 227 | -13.3696 | 3.83799 | 8.96925 |
| metQ | RCd9 | 86 – 137 | 181 – 226 | -13.3365 | 3.87190 | 8.96925 |
| CD630_16150 | RCd9 | 33 – 113 | 212 – 295 | -13.2955 | 4.20614 | 8.96925 |
| CD630_34030 | RCd9 | 86 – 108 | 208 – 230 | -13.1952 | 3.05766 | 8.96925 |
| CD630_18090 | RCd9 | 82 – 113 | 264 – 297 | -13.1742 | 4.23961 | 8.96925 |
| nrdD | RCd9 | 48 – 62 | 88 – 101 | -13.1671 | 3.28290 | 8.96925 |
| CD630_17160 | RCd9 | 86 – 107 | 217 – 235 | -13.1433 | 3.50228 | 8.96925 |
| uxaA | RCd9 | 74 – 145 | 163 – 228 | -13.1178 | 4.26165 | 8.96925 |
| CD630_02420 | RCd9 | 23 – 46 | 241 – 267 | -13.0969 | 3.71048 | 8.96925 |
| CD630_26050 | RCd9 | 127 – 166 | 2 – 49 | -13.0956 | 6.99851 | 8.96925 |
| CD630_22720 | RCd9 | 121 – 145 | 207 – 226 | -13.0643 | 3.81804 | 8.96925 |
| dapB2 | RCd9 | 150 – 197 | 254 – 299 | -12.9707 | 3.27218 | 8.96925 |
| CD630_14810 | RCd9 | 140 – 154 | 96 – 112 | -12.9702 | 3.35323 | 8.96925 |
| cdu2 | RCd9 | 82 – 108 | 207 – 232 | -12.9673 | 2.97335 | 8.96925 |
| CD630_33610 | RCd9 | 27 – 36 | 223 – 232 | -12.9475 | 3.79322 | 8.96925 |
| CD630_06060 | RCd9 | 29 – 40 | 217 – 227 | -12.9276 | 4.94050 | 8.96925 |
| CD630_30120 | RCd9 | 129 – 142 | 247 – 260 | -12.918 | 4.04859 | 8.96925 |
| CD630_14410 | RCd9 | 86 – 108 | 208 – 233 | -12.8971 | 3.39433 | 8.96925 |
| CD630_15200 | RCd9 | 66 – 92 | 268 – 299 | -12.8963 | 4.00442 | 8.96925 |
| CD630_24591 | RCd9 | 85 – 107 | 218 – 232 | -12.8916 | 3.54497 | 8.96925 |
| pilT | RCd9 | 179 – 198 | 97 – 115 | -12.8589 | 3.24639 | 8.96925 |
| CD630_22510 | RCd9 | 83 – 105 | 211 – 234 | -12.8104 | 11.42763 | 8.96925 |
| CD630_10000 | RCd9 | 142 – 163 | 263 – 280 | -12.8052 | 3.86040 | 8.96925 |
| CD630_31382 | RCd9 | 5 – 15 | 285 – 295 | -12.7906 | 3.49137 | 8.96925 |
| CD630_13920 | RCd9 | 117 – 141 | 208 – 234 | -12.7826 | 3.25236 | 8.96925 |
| CD630_20541 | RCd9 | 38 – 60 | 94 – 114 | -12.7617 | 5.74527 | 8.96925 |
| CD630_03650 | RCd9 | 83 – 109 | 207 – 233 | -12.7598 | 3.93697 | 8.96925 |
| CD630_27570 | RCd9 | 101 – 130 | 218 – 234 | -12.7516 | 3.46216 | 8.96925 |
| garK | RCd9 | 19 – 43 | 130 – 152 | -12.7162 | 3.72836 | 8.96925 |
| CD630_35260 | RCd9 | 90 – 108 | 207 – 227 | -12.6586 | 5.77277 | 8.96925 |
| CD630_16550 | RCd9 | 82 – 109 | 207 – 231 | -12.6577 | 5.61064 | 8.96925 |
| CD630_05450 | RCd9 | 80 – 105 | 211 – 236 | -12.6536 | 5.04761 | 8.96925 |
| CD630_08460 | RCd9 | 89 – 148 | 171 – 233 | -12.6257 | 4.15083 | 8.96925 |
| gcvTPA | RCd9 | 87 – 108 | 208 – 230 | -12.5255 | 4.77229 | 8.96925 |
| CD630_10970 | RCd9 | 84 – 105 | 211 – 232 | -12.5036 | 3.53850 | 8.96925 |
| CD630_14250 | RCd9 | 83 – 191 | 129 – 231 | -12.4879 | 4.19604 | 8.96925 |
| CD630_25310 | RCd9 | 85 – 105 | 211 – 232 | -12.4701 | 3.99861 | 8.96925 |
| CD630_14050 | RCd9 | 19 – 44 | 90 – 113 | -12.4441 | 3.30894 | 8.96925 |
| CD630_06690 | RCd9 | 83 – 127 | 252 – 295 | -12.4065 | 5.82709 | 8.96925 |
| CD630_06340 | RCd9 | 4 – 48 | 163 – 197 | -12.3985 | 4.35983 | 8.96925 |
| CD630_20920 | RCd9 | 50 – 151 | 207 – 294 | -12.3801 | 4.13812 | 8.96925 |
| vncR | RCd9 | 131 – 156 | 209 – 234 | -12.3365 | 3.93642 | 8.96925 |
| CD630_24920 | RCd9 | 85 – 125 | 191 – 230 | -12.2677 | 3.89455 | 8.96925 |
| CD630_05280 | RCd9 | 87 – 108 | 208 – 229 | -12.2656 | 3.91665 | 8.96925 |
| CD630_28290 | RCd9 | 86 – 94 | 222 – 230 | -12.2614 | 5.54192 | 8.96925 |
| CD630_17360 | RCd9 | 87 – 139 | 180 – 232 | -12.2522 | 5.66070 | 8.96925 |
| glnA | RCd9 | 87 – 107 | 209 – 229 | -12.196 | 3.87629 | 8.96925 |
| CD630_20960 | RCd9 | 81 – 95 | 222 – 236 | -12.1774 | 5.65940 | 8.96925 |
| CD630_28870 | RCd9 | 60 – 77 | 216 – 230 | -12.1413 | 3.41991 | 8.96925 |
| CD630_18660 | RCd9 | 31 – 55 | 131 – 162 | -12.0723 | 3.50371 | 8.96925 |
| CD630_05460 | RCd9 | 82 – 94 | 222 – 235 | -12.0453 | 3.98753 | 8.96925 |
| CD630_04610 | RCd9 | 85 – 185 | 137 – 232 | -12.0084 | 3.83025 | 8.96925 |
| CD630_28580 | RCd9 | 105 – 187 | 129 – 222 | -12.0057 | 4.67130 | 8.96925 |
| CD630_28320 | RCd9 | 79 – 94 | 215 – 231 | -11.9686 | 3.99184 | 8.96925 |
| CD630_06140 | RCd9 | 85 – 120 | 269 – 298 | -11.9639 | 4.12519 | 8.96925 |
| CD630_31000 | RCd9 | 61 – 128 | 167 – 229 | -11.9268 | 3.70946 | 8.96925 |
| CD630_21620 | RCd9 | 146 – 192 | 187 – 237 | -11.9095 | 3.02649 | 8.96925 |
| acoL | RCd9 | 171 – 200 | 263 – 298 | -11.9012 | 3.32169 | 8.96925 |
| CD630_14360 | RCd9 | 88 – 104 | 248 – 266 | -11.8893 | 3.02079 | 8.96925 |
| CD630_07770 | RCd9 | 87 – 105 | 211 – 230 | -11.8695 | 3.31158 | 8.96925 |
| CD630_28181 | RCd9 | 38 – 77 | 182 – 228 | -11.8451 | 3.91299 | 8.96925 |
| CD630_08100 | RCd9 | 182 – 198 | 99 – 115 | -11.8185 | 4.39181 | 8.96925 |
| CD630_04410 | RCd9 | 84 – 95 | 248 – 259 | -11.817 | 3.90451 | 8.96925 |
| CD630_05030 | RCd9 | 152 – 169 | 95 – 115 | -11.8039 | 4.03823 | 8.96925 |
| CD630_15060 | RCd9 | 85 – 117 | 200 – 231 | -11.7799 | 3.69396 | 8.96925 |
| oppC | RCd9 | 153 – 164 | 287 – 299 | -11.7725 | 3.01695 | 8.96925 |
| pmi | RCd9 | 163 – 192 | 25 – 55 | -11.7381 | 3.86845 | 8.96925 |
| CD630_31960 | RCd9 | 26 – 61 | 197 – 230 | -11.7001 | 4.93757 | 8.96925 |
| CD630_18610 | RCd9 | 81 – 98 | 217 – 234 | -11.6941 | 4.15582 | 8.96925 |
| CD630_05820 | RCd9 | 81 – 104 | 212 – 235 | -11.6789 | 3.75661 | 8.96925 |
| CD630_32070 | RCd9 | 46 – 81 | 211 – 235 | -11.6648 | 3.51964 | 8.96925 |
| CD630_31580 | RCd9 | 85 – 92 | 217 – 224 | -11.6515 | 3.51756 | 8.96925 |
| CD630_05490 | RCd9 | 128 – 165 | 253 – 297 | -11.6352 | 3.89852 | 8.96925 |
| ldh | RCd9 | 146 – 192 | 148 – 188 | -11.5886 | 4.35818 | 8.96925 |
| CD630_29270 | RCd9 | 48 – 59 | 287 – 298 | -11.5416 | 3.70174 | 8.96925 |
| CD630_11220 | RCd9 | 87 – 109 | 207 – 229 | -11.5329 | 3.52327 | 8.96925 |
| bglA | RCd9 | 84 – 105 | 211 – 234 | -11.5223 | 4.50315 | 8.96925 |
| CD630_16160 | RCd9 | 45 – 92 | 217 – 256 | -11.5142 | 6.11801 | 8.96925 |
| CD630_13560 | RCd9 | 19 – 26 | 225 – 232 | -11.513 | 3.38083 | 8.96925 |
| CD630_27060 | RCd9 | 84 – 113 | 213 – 234 | -11.5057 | 4.06049 | 8.96925 |
| CD630_20450 | RCd9 | 65 – 92 | 268 – 295 | -11.4912 | 3.77388 | 8.96925 |
| CD630_19000 | RCd9 | 85 – 107 | 218 – 233 | -11.4656 | 4.18443 | 8.96925 |
| CD630_32060 | RCd9 | 2 – 49 | 247 – 296 | -11.3738 | 5.73797 | 8.96925 |
| CD630_18100 | RCd9 | 78 – 131 | 18 – 65 | -11.3645 | 5.71658 | 8.96925 |
| CD630_36170 | RCd9 | 94 – 116 | 209 – 231 | -11.3496 | 3.72380 | 8.96925 |
| CD630_36781 | RCd9 | 5 – 24 | 268 – 288 | -11.3454 | 3.75946 | 8.96925 |
| CD630_10220 | RCd9 | 87 – 174 | 163 – 229 | -11.3425 | 4.08233 | 8.96925 |
| CD630_21020 | RCd9 | 85 – 105 | 211 – 232 | -11.3343 | 4.28666 | 8.96925 |
| CD630_01690 | RCd9 | 65 – 74 | 217 – 227 | -11.2999 | 4.97234 | 8.96925 |
| CD630_06160 | RCd9 | 78 – 91 | 223 – 235 | -11.25 | 5.51744 | 8.96925 |
| CD630_25610 | RCd9 | 22 – 93 | 134 – 192 | -11.2347 | 3.02685 | 8.96925 |
| CD630_17960 | RCd9 | 81 – 109 | 207 – 230 | -11.2338 | 3.71404 | 8.96925 |
| CD630_15111 | RCd9 | 135 – 161 | 167 – 193 | -11.2225 | 4.18664 | 8.96925 |
| CD630_19401 | RCd9 | 34 – 60 | 158 – 182 | -11.1923 | 4.37084 | 8.96925 |
| CD630_14010 | RCd9 | 89 – 111 | 205 – 228 | -11.1896 | 3.69983 | 8.96925 |
| CD630_26691 | RCd9 | 23 – 35 | 254 – 266 | -11.1474 | 3.96230 | 8.96925 |
| CD630_12110 | RCd9 | 53 – 61 | 222 – 230 | -11.1313 | 4.36078 | 8.96925 |
| malY | RCd9 | 76 – 106 | 206 – 234 | -11.1248 | 4.46345 | 8.96925 |
| CD630_18120 | RCd9 | 184 – 198 | 99 – 112 | -11.1155 | 3.99713 | 8.96925 |
| CD630_02740 | RCd9 | 83 – 106 | 210 – 232 | -11.1152 | 5.02277 | 8.96925 |
| CD630_20280 | RCd9 | 85 – 132 | 184 – 231 | -11.1147 | 3.46343 | 8.96925 |
| CD630_03920 | RCd9 | 82 – 95 | 264 – 278 | -11.1057 | 4.15934 | 8.96925 |
| CD630_14630 | RCd9 | 86 – 93 | 225 – 232 | -11.0898 | 3.34485 | 8.96925 |
| CD630_27240 | RCd9 | 75 – 132 | 175 – 238 | -11.0742 | 3.52462 | 8.96925 |
| CD630_20980 | RCd9 | 94 – 147 | 134 – 198 | -11.065 | 8.12847 | 8.96925 |
| CD630_33580 | RCd9 | 91 – 99 | 222 – 230 | -11.0366 | 3.95778 | 8.96925 |
| spoVFB | RCd9 | 166 – 181 | 263 – 277 | -11.0338 | 4.62758 | 8.96925 |
| CD630_21790 | RCd9 | 138 – 162 | 208 – 232 | -10.9927 | 4.93762 | 8.96925 |
| CD630_26150 | RCd9 | 163 – 178 | 255 – 267 | -10.9759 | 4.06302 | 8.96925 |
| CD630_21240 | RCd9 | 87 – 168 | 159 – 229 | -10.9704 | 5.88879 | 8.96925 |
| CD630_27270 | RCd9 | 89 – 106 | 210 – 228 | -10.938 | 3.50892 | 8.96925 |
| CD630_18410 | RCd9 | 85 – 152 | 161 – 229 | -10.9122 | 3.59739 | 8.96925 |
| uxaA%27 | RCd9 | 86 – 107 | 209 – 230 | -10.911 | 4.26165 | 8.96925 |
| CD630_26750 | RCd9 | 86 – 107 | 209 – 232 | -10.9027 | 3.60610 | 8.96925 |
| CD630_07760 | RCd9 | 77 – 109 | 206 – 235 | -10.902 | 3.64186 | 8.96925 |
| CD630_06050 | RCd9 | 45 – 91 | 252 – 296 | -10.8965 | 3.59332 | 8.96925 |
| CD630_19030 | RCd9 | 103 – 114 | 286 – 297 | -10.891 | 3.34240 | 8.96925 |
| CD630_04540 | RCd9 | 84 – 95 | 217 – 231 | -10.8728 | 3.33556 | 8.96925 |
| CD630_09990 | RCd9 | 90 – 154 | 160 – 225 | -10.8568 | 3.86040 | 8.96925 |
| CD630_32700 | RCd9 | 116 – 126 | 288 – 298 | -10.8497 | 4.74276 | 8.96925 |
| argS | RCd9 | 78 – 93 | 263 – 280 | -10.8218 | 4.96036 | 8.96925 |
| mobA | RCd9 | 145 – 186 | 183 – 231 | -10.8194 | 4.25703 | 8.96925 |
| CD630_33480 | RCd9 | 85 – 107 | 209 – 231 | -10.8073 | 3.67613 | 8.96925 |
| CD630_28260 | RCd9 | 61 – 89 | 91 – 115 | -10.8048 | 3.51613 | 8.96925 |
| CD630_05870 | RCd9 | 87 – 173 | 134 – 230 | -10.7953 | 4.28339 | 8.96925 |
| CD630_24580 | RCd9 | 85 – 108 | 208 – 228 | -10.7846 | 3.88519 | 8.96925 |
| CD630_01590 | RCd9 | 90 – 185 | 131 – 230 | -10.7759 | 3.45609 | 8.96925 |
| CD630_17151 | RCd9 | 154 – 187 | 87 – 119 | -10.7512 | 6.43124 | 8.96925 |
| CD630_17292 | RCd9 | 83 – 96 | 209 – 222 | -10.7494 | 6.25799 | 8.96925 |
| CD630_14240 | RCd9 | 70 – 94 | 215 – 234 | -10.731 | 8.60766 | 8.96925 |
| CD630_19790 | RCd9 | 115 – 146 | 138 – 176 | -10.7199 | 3.79593 | 8.96925 |
| CD630_05290 | RCd9 | 66 – 93 | 266 – 295 | -10.7197 | 3.97604 | 8.96925 |
| CD630_18680 | RCd9 | 62 – 91 | 146 – 190 | -10.7139 | 3.04315 | 8.96925 |
| CD630_36190 | RCd9 | 78 – 105 | 211 – 235 | -10.7049 | 4.63449 | 8.96925 |
| CD630_06110 | RCd9 | 84 – 92 | 225 – 233 | -10.7031 | 5.61380 | 8.96925 |
| CD630_20440 | RCd9 | 167 – 191 | 149 – 179 | -10.6981 | 3.77388 | 8.96925 |
| CD630_01640 | RCd9 | 83 – 104 | 212 – 235 | -10.6946 | 4.55381 | 8.96925 |
| CD630_24220 | RCd9 | 146 – 184 | 161 – 194 | -10.6811 | 4.18037 | 8.96925 |
| CD630_06080 | RCd9 | 168 – 201 | 82 – 117 | -10.678 | 3.82315 | 8.96925 |
| glgB | RCd9 | 69 – 94 | 264 – 291 | -10.6726 | 4.48273 | 8.96925 |
| cwp26 | RCd9 | 147 – 158 | 170 – 181 | -10.6711 | 4.44377 | 8.96925 |
| CD630_23770 | RCd9 | 59 – 92 | 263 – 288 | -10.6297 | 3.25181 | 8.96925 |
| cheW | RCd9 | 86 – 95 | 223 – 232 | -10.6141 | 3.72371 | 8.96925 |
| CD630_14190 | RCd9 | 86 – 94 | 224 – 232 | -10.6048 | 3.78562 | 8.96925 |
| CD630_02790 | RCd9 | 84 – 105 | 211 – 231 | -10.6003 | 4.77661 | 8.96925 |
| tcdA | RCd9 | 48 – 64 | 97 – 116 | -10.5963 | 4.70593 | 8.96925 |
| CD630_08950 | RCd9 | 159 – 200 | 129 – 169 | -10.5951 | 3.71905 | 8.96925 |
| CD630_07840 | RCd9 | 87 – 94 | 225 – 232 | -10.5789 | 4.26761 | 8.96925 |
| CD630_22740 | RCd9 | 79 – 96 | 94 – 113 | -10.5687 | 3.08733 | 8.96925 |
| eutV | RCd9 | 68 – 152 | 209 – 296 | -10.5681 | 4.55715 | 8.96925 |
| CD630_09970 | RCd9 | 85 – 98 | 284 – 296 | -10.5636 | 3.25917 | 8.96925 |
| tcdC | RCd9 | 136 – 153 | 217 – 236 | -10.5515 | 3.76101 | 8.96925 |
| CD630_22070 | RCd9 | 7 – 117 | 187 – 293 | -10.5418 | 4.09651 | 8.96925 |
| CD630_25050 | RCd9 | 83 – 113 | 207 – 234 | -10.5329 | 4.33624 | 8.96925 |
| CD630_17511 | RCd9 | 8 – 35 | 137 – 163 | -10.5092 | 9.12322 | 8.96925 |
| CD630_17770 | RCd9 | 129 – 138 | 23 – 32 | -10.4984 | 5.06490 | 8.96925 |
| CD630_33600 | RCd9 | 21 – 32 | 98 – 112 | -10.4955 | 3.73196 | 8.96925 |
| CD630_28550 | RCd9 | 88 – 111 | 208 – 223 | -10.4862 | 3.51196 | 8.96925 |
| CD630_33190 | RCd9 | 6 – 19 | 140 – 153 | -10.4763 | 4.13899 | 8.96925 |
| ribD | RCd9 | 85 – 92 | 222 – 229 | -10.4497 | 6.65922 | 8.96925 |
| flhB | RCd9 | 185 – 197 | 21 – 34 | -10.4326 | 3.55524 | 8.96925 |
| ogt1 | RCd9 | 88 – 108 | 209 – 230 | -10.4077 | 3.16772 | 8.96925 |
| CD630_27280 | RCd9 | 38 – 178 | 158 – 281 | -10.4023 | 3.24738 | 8.96925 |
| CD630_11730 | RCd9 | 146 – 181 | 84 – 119 | -10.3999 | 4.80521 | 8.96925 |
| CD630_23070 | RCd9 | 171 – 185 | 173 – 187 | -10.3778 | 5.23027 | 8.96925 |
| CD630_25020 | RCd9 | 172 – 198 | 265 – 294 | -10.36 | 4.17721 | 8.96925 |
| CD630_13600 | RCd9 | 129 – 178 | 180 – 232 | -10.3566 | 4.45481 | 8.96925 |
| CD630_05590 | RCd9 | 90 – 118 | 209 – 229 | -10.3275 | 3.39670 | 8.96925 |
| kdpE | RCd9 | 184 – 197 | 224 – 236 | -10.3245 | 4.27140 | 8.96925 |
| CD630_30360 | RCd9 | 89 – 122 | 15 – 55 | -10.32 | 4.26509 | 8.96925 |
| CD630_22710 | RCd9 | 118 – 129 | 288 – 298 | -10.2816 | 4.05623 | 8.96925 |
| appF | RCd9 | 85 – 92 | 224 – 231 | -10.2698 | 4.25570 | 8.96925 |
| gatC | RCd9 | 85 – 108 | 208 – 231 | -10.2673 | 4.25143 | 8.96925 |
| CD630_15900 | RCd9 | 166 – 176 | 257 – 267 | -10.2641 | 3.17664 | 8.96925 |
| CD630_10950 | RCd9 | 84 – 107 | 90 – 114 | -10.2322 | 3.67675 | 8.96925 |
| CD630_09750 | RCd9 | 120 – 143 | 206 – 226 | -10.2143 | 4.83410 | 8.96925 |
| CD630_28920 | RCd9 | 120 – 143 | 206 – 226 | -10.2143 | 4.51664 | 8.96925 |
| CD630_17710 | RCd9 | 56 – 94 | 264 – 296 | -10.2108 | 4.37688 | 8.96925 |
| CD630_04320 | RCd9 | 161 – 192 | 159 – 185 | -10.2078 | 3.68342 | 8.96925 |
| CD630_18950 | RCd9 | 181 – 189 | 181 – 189 | -10.2 | 3.40610 | 8.96925 |
| CD630_08510 | RCd9 | 76 – 90 | 100 – 114 | -10.1958 | 5.16329 | 8.96925 |
| CD630_06540 | RCd9 | 83 – 91 | 225 – 233 | -10.1838 | 3.54673 | 8.96925 |
| opuCA | RCd9 | 85 – 109 | 268 – 295 | -10.1825 | 4.07035 | 8.96925 |
| CD630_12320 | RCd9 | 43 – 93 | 219 – 267 | -10.1503 | 3.28667 | 8.96925 |
| CD630_19670 | RCd9 | 79 – 91 | 223 – 237 | -10.1102 | 3.20718 | 8.96925 |
| CD630_22890 | RCd9 | 36 – 46 | 222 – 232 | -10.1029 | 3.75591 | 8.96925 |
| CD630_15740 | RCd9 | 38 – 94 | 224 – 259 | -10.0995 | 4.43222 | 8.96925 |
| CD630_14710 | RCd9 | 87 – 97 | 219 – 229 | -10.0795 | 3.65171 | 8.96925 |
| CD630_05000 | RCd9 | 85 – 95 | 222 – 232 | -10.0718 | 4.02442 | 8.96925 |
| CD630_18930 | RCd9 | 65 – 125 | 214 – 262 | -10.0692 | 5.32127 | 8.96925 |
| CD630_21090 | RCd9 | 16 – 40 | 192 – 215 | -10.0626 | 4.15803 | 8.96925 |
| CD630_23000 | RCd9 | 111 – 124 | 285 – 299 | -10.0602 | 3.54130 | 8.96925 |
| dpaA | RCd9 | 81 – 92 | 224 – 236 | -10.0447 | 4.65364 | 8.96925 |
| CD630_04280 | RCd9 | 81 – 108 | 208 – 231 | -10.0288 | 3.61457 | 8.96925 |
| cwp29 | RCd9 | 28 – 35 | 224 – 231 | -10.0227 | 4.40865 | 8.96925 |
| CD630_21070 | RCd9 | 88 – 95 | 222 – 229 | -10.0146 | 5.35019 | 8.96925 |
| CD630_31490 | RCd9 | 90 – 106 | 210 – 229 | -10.0093 | 4.08593 | 8.96925 |
| CD630_19800 | RCd9 | 61 – 86 | 93 – 115 | -9.9795 | 5.22035 | 8.96925 |
| CD630_21490 | RCd9 | 190 – 196 | 264 – 270 | -9.95545 | 3.37462 | 8.96925 |
| CD630_19840 | RCd9 | 90 – 96 | 223 – 229 | -9.95131 | 3.44724 | 8.96925 |
| CD630_25320 | RCd9 | 87 – 108 | 208 – 229 | -9.94446 | 4.29415 | 8.96925 |
| scrR | RCd9 | 121 – 155 | 132 – 169 | -9.92307 | 3.49166 | 8.96925 |
| feoB | RCd9 | 28 – 50 | 9 – 32 | -9.89291 | 4.05364 | 8.96925 |
| CD630_27220 | RCd9 | 87 – 108 | 208 – 231 | -9.87409 | 4.18786 | 8.96925 |
| CD630_31220 | RCd9 | 127 – 156 | 266 – 297 | -9.85495 | 3.73779 | 8.96925 |
| opuCC | RCd9 | 88 – 112 | 206 – 232 | -9.8368 | 4.07035 | 8.96925 |
| CD630_02830 | RCd9 | 89 – 105 | 207 – 222 | -9.83438 | 3.57106 | 8.96925 |
| CD630_25200 | RCd9 | 74 – 95 | 213 – 233 | -9.83229 | 3.65158 | 8.96925 |
| CD630_15190 | RCd9 | 87 – 93 | 223 – 229 | -9.8195 | 4.43255 | 8.96925 |
| CD630_05840 | RCd9 | 88 – 98 | 285 – 295 | -9.78006 | 4.28779 | 8.96925 |
| CD630_36040 | RCd9 | 84 – 127 | 264 – 295 | -9.77523 | 3.52009 | 8.96925 |
| CD630_21910 | RCd9 | 147 – 197 | 179 – 232 | -9.77416 | 4.66600 | 8.96925 |
| CD630_25040 | RCd9 | 88 – 105 | 211 – 228 | -9.7587 | 3.43898 | 8.96925 |
| aroD | RCd9 | 3 – 32 | 138 – 170 | -9.75157 | 3.71929 | 8.96925 |
| iorA | RCd9 | 151 – 161 | 179 – 189 | -9.71552 | 3.83173 | 8.96925 |
| etfA | RCd9 | 75 – 96 | 215 – 231 | -9.70263 | 4.80521 | 8.96925 |
| CD630_21230 | RCd9 | 84 – 102 | 207 – 223 | -9.66857 | 3.89213 | 8.96925 |
| CD630_22200 | RCd9 | 84 – 101 | 209 – 225 | -9.66717 | 4.18288 | 8.96925 |
| CD630_19921 | RCd9 | 60 – 110 | 165 – 220 | -9.66068 | 5.46345 | 8.96925 |
| CD630_22050 | RCd9 | 89 – 95 | 224 – 230 | -9.65696 | 4.87639 | 8.96925 |
| rolB | RCd9 | 85 – 109 | 207 – 232 | -9.65036 | 3.60270 | 8.96925 |
| CD630_14420 | RCd9 | 87 – 106 | 210 – 228 | -9.64692 | 3.99497 | 8.96925 |
| CD630_01570 | RCd9 | 111 – 120 | 110 – 119 | -9.62554 | 6.08699 | 8.96925 |
| gcp | RCd9 | 86 – 95 | 214 – 223 | -9.60618 | 4.92798 | 8.96925 |
| CD630_10050 | RCd9 | 7 – 27 | 271 – 298 | -9.60585 | 3.93750 | 8.96925 |
| CD630_03040 | RCd9 | 83 – 92 | 289 – 298 | -9.58274 | 3.51403 | 8.96925 |
| CD630_30100 | RCd9 | 85 – 91 | 224 – 230 | -9.54152 | 4.13234 | 8.96925 |
| CD630_16670 | RCd9 | 123 – 131 | 106 – 114 | -9.53691 | 3.33553 | 8.96925 |
| CD630_16940 | RCd9 | 86 – 107 | 209 – 231 | -9.53468 | 4.14665 | 8.96925 |
| CD630_04770 | RCd9 | 85 – 93 | 225 – 233 | -9.52858 | 3.58478 | 8.96925 |
| CD630_15270 | RCd9 | 89 – 100 | 282 – 293 | -9.51087 | 4.66015 | 8.96925 |
| CD630_19904 | RCd9 | 151 – 186 | 228 – 267 | -9.50721 | 5.96526 | 8.96925 |
| CD630_06970 | RCd9 | 122 – 132 | 218 – 228 | -9.49992 | 3.82312 | 8.96925 |

**Table S9. IntaRNA prediction for ncRNA-mRNA interactions**

Strand "-"

| | | | | | RIPseq log p-value | |
|---|---|---|---|---|---|---|
| **CDS** | **ncRNA** | **interaction CDS** | **interaction ncRNA** | **interaction energy (kcal/mol)** | **cds logpVal** | **ncRNA logpVal** |
| CD630_05460 | CD630_Cdi1_10 | 51 -- 113 | 93 -- 154 | -16.6616 | 3.98753 | 8.60766 |
| CD630_21700 | CD630_Cdi1_10 | 22 -- 41 | 125 -- 149 | -15.4903 | 3.96650 | 8.60766 |
| CD630_20450 | CD630_Cdi1_10 | 113 -- 165 | 93 -- 148 | -14.9417 | 3.77388 | 8.60766 |
| CD630_06150 | CD630_Cdi1_10 | 148 -- 175 | 129 -- 154 | -14.125 | 5.51744 | 8.60766 |
| CD630_14600 | CD630_Cdi1_10 | 71 -- 101 | 126 -- 159 | -14.0099 | 5.90348 | 8.60766 |
| CD630_31461 | CD630_Cdi1_10 | 134 -- 190 | 97 -- 151 | -13.3271 | 4.41476 | 8.60766 |
| sspA | CD630_Cdi1_10 | 50 -- 109 | 96 -- 156 | -12.4702 | 3.62689 | 8.60766 |
| CD630_01570 | CD630_Cdi1_10 | 126 -- 167 | 115 -- 156 | -12.2132 | 6.08699 | 8.60766 |
| CD630_08080 | CD630_Cdi1_10 | 26 -- 52 | 123 -- 151 | -12.0032 | 4.87819 | 8.60766 |
| CD630_05160 | CD630_Cdi1_10 | 141 -- 199 | 110 -- 159 | -11.9239 | 5.03573 | 8.60766 |
| CD630_31530 | CD630_Cdi1_10 | 118 -- 146 | 114 -- 143 | -11.8714 | 3.49100 | 8.60766 |
| feoB | CD630_Cdi1_10 | 134 -- 197 | 88 -- 145 | -11.7921 | 4.05364 | 8.60766 |
| CD630_13020 | CD630_Cdi1_10 | 69 -- 123 | 105 -- 158 | -11.7827 | 3.52267 | 8.60766 |
| CD630_17281 | CD630_Cdi1_10 | 50 -- 94 | 114 -- 158 | -11.735 | 5.91544 | 8.60766 |
| CD630_21260 | CD630_Cdi1_10 | 140 -- 152 | 44 -- 56 | -11.6639 | 4.22255 | 8.60766 |
| CD630_21090 | CD630_Cdi1_10 | 161 -- 185 | 129 -- 152 | -11.5015 | 4.15803 | 8.60766 |
| CD630_05830 | CD630_Cdi1_10 | 47 -- 94 | 110 -- 156 | -11.4229 | 3.35360 | 8.60766 |
| ppnK | CD630_Cdi1_10 | 142 -- 162 | 133 -- 148 | -11.3709 | 4.36817 | 8.60766 |
| CD630_19401 | CD630_Cdi1_10 | 50 -- 82 | 125 -- 157 | -11.2814 | 4.37084 | 8.60766 |
| CD630_06140 | CD630_Cdi1_10 | 51 -- 94 | 112 -- 149 | -11.1065 | 4.12519 | 8.60766 |
| CD630_30110 | CD630_Cdi1_10 | 45 -- 98 | 110 -- 158 | -11.0214 | 3.31041 | 8.60766 |
| CD630_06340 | CD630_Cdi1_10 | 142 -- 172 | 125 -- 152 | -10.9153 | 4.35983 | 8.60766 |
| gabT | CD630_Cdi1_10 | 151 -- 174 | 134 -- 158 | -10.7702 | 3.67067 | 8.60766 |
| CD630_20170 | CD630_Cdi1_10 | 129 -- 164 | 114 -- 148 | -10.6829 | 6.90201 | 8.60766 |
| CD630_06170 | CD630_Cdi1_10 | 66 -- 108 | 96 -- 138 | -10.6777 | 4.95366 | 8.60766 |
| CD630_04620 | CD630_Cdi1_10 | 17 -- 50 | 123 -- 150 | -10.6292 | 3.45413 | 8.60766 |
| ldh | CD630_Cdi1_10 | 157 -- 171 | 9 -- 23 | -10.5959 | 4.35818 | 8.60766 |
| CD630_20460 | CD630_Cdi1_10 | 144 -- 185 | 111 -- 151 | -10.5878 | 7.94034 | 8.60766 |
| CD630_19904 | CD630_Cdi1_10 | 20 -- 36 | 134 -- 151 | -10.5873 | 5.96526 | 8.60766 |
| dnaD | CD630_Cdi1_10 | 143 -- 151 | 143 -- 151 | -10.5137 | 3.56078 | 8.60766 |
| cstA | CD630_Cdi1_10 | 17 -- 51 | 125 -- 156 | -10.5023 | 3.53584 | 8.60766 |
| CD630_20600 | CD630_Cdi1_10 | 150 -- 179 | 123 -- 152 | -10.4984 | 4.20093 | 8.60766 |
| CD630_14140 | CD630_Cdi1_10 | 150 -- 172 | 130 -- 154 | -10.4938 | 3.16551 | 8.60766 |
| CD630_13530 | CD630_Cdi1_10 | 161 -- 181 | 129 -- 154 | -10.4921 | 4.09832 | 8.60766 |
| CD630_22550 | CD630_Cdi1_10 | 26 -- 51 | 125 -- 150 | -10.4504 | 3.62379 | 8.60766 |
| CD630_34050 | CD630_Cdi1_10 | 8 -- 20 | 4 -- 16 | -10.4362 | 4.03863 | 8.60766 |
| CD630_25190 | CD630_Cdi1_10 | 174 -- 181 | 93 -- 100 | -10.4303 | 3.19663 | 8.60766 |
| CD630_20670 | CD630_Cdi1_10 | 179 -- 197 | 78 -- 95 | -10.4264 | 4.42757 | 8.60766 |
| cdtR | CD630_Cdi1_10 | 2 -- 41 | 127 -- 151 | -10.3703 | 5.84368 | 8.60766 |
| CD630_16540 | CD630_Cdi1_10 | 161 -- 182 | 124 -- 145 | -10.3089 | 4.37658 | 8.60766 |
| CD630_30310 | CD630_Cdi1_10 | 129 -- 154 | 134 -- 158 | -10.3081 | 4.56059 | 8.60766 |
| CD630_04880 | CD630_Cdi1_10 | 149 -- 175 | 125 -- 151 | -10.292 | 4.62291 | 8.60766 |
| CD630_09772 | CD630_Cdi1_10 | 53 -- 96 | 112 -- 158 | -10.2645 | 3.66033 | 8.60766 |
| CD630_16330 | CD630_Cdi1_10 | 146 -- 166 | 132 -- 151 | -10.2543 | 4.11180 | 8.60766 |
| CD630_13620 | CD630_Cdi1_10 | 133 -- 164 | 124 -- 155 | -10.2157 | 4.77627 | 8.60766 |
| CD630_02720 | CD630_Cdi1_10 | 141 -- 168 | 127 -- 152 | -10.1722 | 3.38938 | 8.60766 |
| CD630_23760 | CD630_Cdi1_10 | 45 -- 62 | 111 -- 128 | -10.1684 | 4.54307 | 8.60766 |
| CD630_15360 | CD630_Cdi1_10 | 24 -- 37 | 42 -- 55 | -10.1303 | 4.47208 | 8.60766 |
| CD630_10880 | CD630_Cdi1_10 | 5 -- 25 | 137 -- 158 | -10.0336 | 5.05048 | 8.60766 |
| CD630_18311 | CD630_Cdi1_10 | 54 -- 75 | 110 -- 131 | -10.022 | 6.86047 | 8.60766 |
| CD630_32990 | CD630_Cdi1_10 | 79 -- 123 | 108 -- 150 | -9.99476 | 5.24925 | 8.60766 |
| tcdA | CD630_Cdi1_10 | 139 -- 174 | 123 -- 158 | -9.95836 | 4.70593 | 8.60766 |
| CD630_02420 | CD630_Cdi1_10 | 9 -- 50 | 48 -- 94 | -9.89244 | 3.71048 | 8.60766 |
| CD630_34030 | CD630_Cdi1_10 | 42 -- 66 | 134 -- 157 | -9.88448 | 3.05766 | 8.60766 |
| CD630_03220 | CD630_Cdi1_10 | 150 -- 196 | 107 -- 154 | -9.86948 | 3.36746 | 8.60766 |
| glnA | CD630_Cdi1_10 | 151 -- 172 | 134 -- 154 | -9.86101 | 3.87629 | 8.60766 |
| CD630_29960 | CD630_Cdi1_10 | 45 -- 63 | 140 -- 158 | -9.85151 | 4.51904 | 8.60766 |
| pmi | CD630_Cdi1_10 | 50 -- 95 | 113 -- 159 | -9.81621 | 3.86845 | 8.60766 |
| mobA | CD630_Cdi1_10 | 137 -- 192 | 94 -- 151 | -9.78079 | 4.25703 | 8.60766 |
| CD630_30120 | CD630_Cdi1_10 | 45 -- 89 | 113 -- 156 | -9.7463 | 4.04859 | 8.60766 |
| CD630_23070 | CD630_Cdi1_10 | 172 -- 187 | 89 -- 103 | -9.74447 | 5.23027 | 8.60766 |
| CD630_09100 | CD630_Cdi1_10 | 13 -- 36 | 127 -- 150 | -9.73639 | 5.92260 | 8.60766 |
| CD630_03170 | CD630_Cdi1_10 | 133 -- 151 | 132 -- 151 | -9.73512 | 3.66946 | 8.60766 |
| moeA | CD630_Cdi1_10 | 49 -- 81 | 134 -- 158 | -9.73278 | 3.26118 | 8.60766 |
| CD630_21110 | CD630_Cdi1_10 | 127 -- 157 | 125 -- 157 | -9.69025 | 3.89630 | 8.60766 |
| CD630_05490 | CD630_Cdi1_10 | 164 -- 194 | 80 -- 118 | -9.67055 | 3.89852 | 8.60766 |
| CD630_07480 | CD630_Cdi1_10 | 156 -- 164 | 139 -- 147 | -9.66146 | 5.18321 | 8.60766 |
| CD630_13370 | CD630_Cdi1_10 | 156 -- 181 | 124 -- 149 | -9.63754 | 4.33145 | 8.60766 |
| appC | CD630_Cdi1_10 | 49 -- 72 | 137 -- 157 | -9.61539 | 4.50438 | 8.60766 |
| CD630_14010 | CD630_Cdi1_10 | 39 -- 54 | 127 -- 146 | -9.51623 | 3.69983 | 8.60766 |
| CD630_09771 | CD630_Cdi1_5 | 52 -- 201 | 343 -- 492 | -191.788 | 4.95362 | 4.83391 |
| CD630_28890 | CD630_Cdi1_5 | 52 -- 201 | 343 -- 492 | -191.788 | 7.52469 | 4.83391 |
| CD630_17170 | CD630_Cdi1_5 | 158 -- 177 | 440 -- 459 | -19.1929 | 4.10230 | 4.83391 |
| srlD | CD630_Cdi1_5 | 125 -- 149 | 434 -- 458 | -18.7704 | 4.19843 | 4.83391 |
| cheW | CD630_Cdi1_5 | 85 -- 105 | 439 -- 460 | -18.5927 | 3.72371 | 4.83391 |
| CD630_30140 | CD630_Cdi1_5 | 84 -- 110 | 435 -- 458 | -18.4877 | 3.28343 | 4.83391 |
| CD630_14410 | CD630_Cdi1_5 | 89 -- 109 | 435 -- 457 | -18.3862 | 3.39433 | 4.83391 |
| CD630_13920 | CD630_Cdi1_5 | 119 -- 139 | 438 -- 458 | -18.2646 | 3.25236 | 4.83391 |
| CD630_09562 | CD630_Cdi1_5 | 46 -- 195 | 358 -- 496 | -18.2463 | 3.94307 | 4.83391 |
| CD630_21490 | CD630_Cdi1_5 | 90 -- 175 | 402 -- 454 | -17.834 | 3.37462 | 4.83391 |
| CD630_10220 | CD630_Cdi1_5 | 87 -- 157 | 387 -- 456 | -17.6773 | 4.08233 | 4.83391 |
| CD630_19680 | CD630_Cdi1_5 | 88 -- 136 | 390 -- 454 | -17.445 | 3.58080 | 4.83391 |
| metQ | CD630_Cdi1_5 | 85 -- 200 | 349 -- 463 | -17.4277 | 3.87190 | 4.83391 |
| CD630_21610 | CD630_Cdi1_5 | 88 -- 123 | 418 -- 456 | -17.1855 | 3.16662 | 4.83391 |
| tcdA | CD630_Cdi1_5 | 36 -- 94 | 422 -- 463 | -16.729 | 4.70593 | 4.83391 |
| CD630_35260 | CD630_Cdi1_5 | 90 -- 104 | 440 -- 454 | -16.6565 | 5.77277 | 4.83391 |
| CD630_06970 | CD630_Cdi1_5 | 84 -- 128 | 420 -- 460 | -16.635 | 3.82312 | 4.83391 |
| CD630_21750 | CD630_Cdi1_5 | 59 -- 106 | 270 -- 323 | -16.5731 | 3.70626 | 4.83391 |
| CD630_27570 | CD630_Cdi1_5 | 102 -- 140 | 422 -- 464 | -16.4989 | 3.46216 | 4.83391 |
| CD630_21570 | CD630_Cdi1_5 | 151 -- 186 | 110 -- 141 | -16.4684 | 6.48385 | 4.83391 |
| CD630_20450 | CD630_Cdi1_5 | 111 -- 165 | 85 -- 142 | -16.396 | 3.77388 | 4.83391 |
| CD630_18120 | CD630_Cdi1_5 | 78 -- 109 | 435 -- 464 | -16.3428 | 3.99713 | 4.83391 |
| CD630_14700 | CD630_Cdi1_5 | 76 -- 157 | 106 -- 200 | -16.3357 | 3.98234 | 4.83391 |
| CD630_09570 | CD630_Cdi1_5 | 21 -- 52 | 426 -- 457 | -16.3185 | 5.31213 | 4.83391 |
| CD630_29070 | CD630_Cdi1_5 | 21 -- 52 | 426 -- 457 | -16.3185 | 3.89477 | 4.83391 |
| CD630_09990 | CD630_Cdi1_5 | 90 -- 174 | 372 -- 456 | -16.2983 | 3.86040 | 4.83391 |
| CD630_23780 | CD630_Cdi1_5 | 90 -- 122 | 316 -- 353 | -16.2513 | 3.91629 | 4.83391 |
| kdpE | CD630_Cdi1_5 | 91 -- 106 | 438 -- 455 | -16.2079 | 4.27140 | 4.83391 |
| CD630_26050 | CD630_Cdi1_5 | 169 -- 179 | 443 -- 453 | -16.1339 | 6.99851 | 4.83391 |
| CD630_11241 | CD630_Cdi1_5 | 164 -- 193 | 178 -- 206 | -16.1288 | 4.85723 | 4.83391 |
| CD630_04341 | CD630_Cdi1_5 | 159 -- 188 | 426 -- 454 | -16.1187 | 4.29262 | 4.83391 |
| xdhA | CD630_Cdi1_5 | 47 -- 101 | 105 -- 157 | -16.07 | 4.45147 | 4.83391 |
| CD630_18910 | CD630_Cdi1_5 | 118 -- 138 | 428 -- 453 | -16.0571 | 3.30688 | 4.83391 |
| CD630_06140 | CD630_Cdi1_5 | 86 -- 123 | 421 -- 453 | -16.0409 | 4.12519 | 4.83391 |
| CD630_05211 | CD630_Cdi1_5 | 112 -- 153 | 212 -- 260 | -15.9862 | 3.97048 | 4.83391 |
| CD630_17281 | CD630_Cdi1_5 | 50 -- 94 | 106 -- 151 | -15.9585 | 5.91544 | 4.83391 |
| CD630_26250 | CD630_Cdi1_5 | 86 -- 157 | 377 -- 457 | -15.9459 | 5.03379 | 4.83391 |
| oppC | CD630_Cdi1_5 | 143 -- 193 | 99 -- 150 | -15.8425 | 3.01695 | 4.83391 |
| CD630_02790 | CD630_Cdi1_5 | 127 -- 171 | 320 -- 356 | -15.8191 | 4.77661 | 4.83391 |
| CD630_18680 | CD630_Cdi1_5 | 86 -- 111 | 425 -- 451 | -15.8026 | 3.04315 | 4.83391 |
| CD630_15360 | CD630_Cdi1_5 | 117 -- 176 | 269 -- 319 | -15.6381 | 4.47208 | 4.83391 |
| cbpA | CD630_Cdi1_5 | 84 -- 95 | 451 -- 462 | -15.5714 | 3.03040 | 4.83391 |
| CD630_32710 | CD630_Cdi1_5 | 91 -- 185 | 373 -- 452 | -15.5189 | 4.74276 | 4.83391 |
| CD630_19780 | CD630_Cdi1_5 | 14 -- 32 | 436 -- 454 | -15.3999 | 3.64830 | 4.83391 |
| CD630_31530 | CD630_Cdi1_5 | 107 -- 146 | 106 -- 144 | -15.3625 | 3.49100 | 4.83391 |
| cbiQ | CD630_Cdi1_5 | 1 -- 89 | 371 -- 451 | -15.3452 | 3.79830 | 4.83391 |
| CD630_06060 | CD630_Cdi1_5 | 29 -- 39 | 444 -- 454 | -15.3369 | 4.94050 | 4.83391 |
| CD630_13960 | CD630_Cdi1_5 | 117 -- 175 | 183 -- 246 | -15.323 | 3.30375 | 4.83391 |
| CD630_05830 | CD630_Cdi1_5 | 47 -- 94 | 102 -- 149 | -15.3102 | 3.35360 | 4.83391 |
| CD630_24590 | CD630_Cdi1_5 | 123 -- 190 | 419 -- 464 | -15.2494 | 3.70401 | 4.83391 |
| terD | CD630_Cdi1_5 | 87 -- 141 | 411 -- 457 | -15.2137 | 3.33148 | 4.83391 |
| bglA | CD630_Cdi1_5 | 155 -- 190 | 438 -- 458 | -15.1268 | 4.86402 | 4.83391 |
| CD630_07840 | CD630_Cdi1_5 | 109 -- 163 | 105 -- 153 | -15.1266 | 4.26761 | 4.83391 |
| CD630_19690 | CD630_Cdi1_5 | 109 -- 126 | 436 -- 449 | -15.1038 | 3.85861 | 4.83391 |
| rbsC | CD630_Cdi1_5 | 34 -- 63 | 435 -- 459 | -15.0494 | 3.71650 | 4.83391 |
| CD630_06080 | CD630_Cdi1_5 | 156 -- 182 | 177 -- 205 | -15.0335 | 3.82315 | 4.83391 |
| CD630_14250 | CD630_Cdi1_5 | 83 -- 104 | 440 -- 458 | -15.0112 | 4.19604 | 4.83391 |
| CD630_18410 | CD630_Cdi1_5 | 85 -- 104 | 436 -- 456 | -14.8044 | 3.59739 | 4.83391 |
| CD630_10950 | CD630_Cdi1_5 | 85 -- 130 | 429 -- 459 | -14.8037 | 3.67675 | 4.83391 |
| CD630_21020 | CD630_Cdi1_5 | 85 -- 108 | 438 -- 459 | -14.7555 | 4.28666 | 4.83391 |
| gabT | CD630_Cdi1_5 | 90 -- 123 | 425 -- 454 | -14.7506 | 3.67067 | 4.83391 |
| cprK | CD630_Cdi1_5 | 84 -- 112 | 419 -- 450 | -14.6514 | 3.96216 | 4.83391 |
| CD630_26750 | CD630_Cdi1_5 | 85 -- 104 | 440 -- 460 | -14.635 | 3.60610 | 4.83391 |
| CD630_14920 | CD630_Cdi1_5 | 95 -- 113 | 440 -- 458 | -14.5499 | 4.35225 | 4.83391 |
| CD630_32700 | CD630_Cdi1_5 | 180 -- 194 | 192 -- 206 | -14.5344 | 4.74276 | 4.83391 |
| CD630_14420 | CD630_Cdi1_5 | 75 -- 97 | 438 -- 460 | -14.5038 | 3.99497 | 4.83391 |
| CD630_05480 | CD630_Cdi1_5 | 118 -- 199 | 371 -- 447 | -14.4989 | 4.27854 | 4.83391 |
| CD630_28550 | CD630_Cdi1_5 | 78 -- 109 | 421 -- 451 | -14.4961 | 3.51196 | 4.83391 |
| CD630_19580 | CD630_Cdi1_5 | 52 -- 65 | 422 -- 435 | -14.4408 | 4.51902 | 4.83391 |
| CD630_07480 | CD630_Cdi1_5 | 87 -- 106 | 438 -- 461 | -14.4247 | 5.18321 | 4.83391 |
| CD630_10050 | CD630_Cdi1_5 | 152 -- 200 | 420 -- 463 | -14.4129 | 3.93750 | 4.83391 |
| CD630_21620 | CD630_Cdi1_5 | 4 -- 38 | 110 -- 141 | -14.412 | 3.02649 | 4.83391 |
| CD630_31000 | CD630_Cdi1_5 | 101 -- 146 | 426 -- 451 | -14.3751 | 3.70946 | 4.83391 |
| CD630_05821 | CD630_Cdi1_5 | 80 -- 103 | 332 -- 357 | -14.2986 | 3.75661 | 4.83391 |
| CD630_29250 | CD630_Cdi1_5 | 87 -- 106 | 438 -- 457 | -14.174 | 4.61901 | 4.83391 |
| CD630_09410 | CD630_Cdi1_5 | 87 -- 106 | 438 -- 457 | -14.174 | 4.75651 | 4.83391 |
| CD630_16550 | CD630_Cdi1_5 | 79 -- 108 | 435 -- 465 | -14.1557 | 5.61064 | 4.83391 |
| CD630_16150 | CD630_Cdi1_5 | 83 -- 105 | 439 -- 460 | -14.1328 | 4.20614 | 4.83391 |
| pilD | CD630_Cdi1_5 | 3 -- 31 | 104 -- 131 | -14.109 | 4.54123 | 4.83391 |
| CD630_14630 | CD630_Cdi1_5 | 139 -- 176 | 422 -- 453 | -14.0951 | 3.34485 | 4.83391 |
| CD630_17292 | CD630_Cdi1_5 | 82 -- 105 | 423 -- 449 | -14.0565 | 6.25799 | 4.83391 |
| acoL | CD630_Cdi1_5 | 104 -- 135 | 348 -- 379 | -14.0258 | 3.32169 | 4.83391 |
| CD630_16670 | CD630_Cdi1_5 | 122 -- 137 | 342 -- 358 | -14.0058 | 3.33553 | 4.83391 |
| CD630_20290 | CD630_Cdi1_5 | 80 -- 180 | 372 -- 465 | -13.9027 | 4.38183 | 4.83391 |
| CD630_01640 | CD630_Cdi1_5 | 76 -- 104 | 440 -- 469 | -13.8612 | 4.55381 | 4.83391 |
| CD630_35430 | CD630_Cdi1_5 | 142 -- 182 | 114 -- 150 | -13.8354 | 4.28798 | 4.83391 |
| CD630_21260 | CD630_Cdi1_5 | 138 -- 152 | 37 -- 51 | -13.8314 | 4.22255 | 4.83391 |
| CD630_17160 | CD630_Cdi1_5 | 88 -- 105 | 435 -- 450 | -13.8088 | 3.50228 | 4.83391 |
| CD630_21460 | CD630_Cdi1_5 | 72 -- 79 | 349 -- 356 | -13.8003 | 4.83005 | 4.83391 |
| dapB2 | CD630_Cdi1_5 | 130 -- 162 | 419 -- 450 | -13.7464 | 3.27218 | 4.83391 |
| CD630_05290 | CD630_Cdi1_5 | 19 -- 63 | 106 -- 154 | -13.7324 | 3.97604 | 4.83391 |
| CD630_33480 | CD630_Cdi1_5 | 86 -- 106 | 438 -- 457 | -13.6884 | 3.67613 | 4.83391 |
| CD630_27240 | CD630_Cdi1_5 | 2 -- 11 | 438 -- 447 | -13.6696 | 3.52462 | 4.83391 |
| CD630_04280 | CD630_Cdi1_5 | 82 -- 109 | 435 -- 457 | -13.6628 | 3.61457 | 4.83391 |
| CD630_26680 | CD630_Cdi1_5 | 75 -- 104 | 187 -- 216 | -13.617 | 3.53956 | 4.83391 |
| flgB | CD630_Cdi1_5 | 89 -- 101 | 437 -- 449 | -13.6106 | 3.25736 | 4.83391 |
| CD630_28320 | CD630_Cdi1_5 | 78 -- 93 | 450 -- 464 | -13.5796 | 3.99184 | 4.83391 |
| CD630_23440 | CD630_Cdi1_5 | 1 -- 13 | 439 -- 452 | -13.5687 | 3.34841 | 4.83391 |
| CD630_15080 | CD630_Cdi1_5 | 85 -- 104 | 440 -- 459 | -13.557 | 4.03033 | 4.83391 |
| CD630_04540 | CD630_Cdi1_5 | 82 -- 93 | 453 -- 464 | -13.5557 | 3.33556 | 4.83391 |
| CD630_36250 | CD630_Cdi1_5 | 170 -- 199 | 179 -- 206 | -13.5125 | 4.13224 | 4.83391 |
| CD630_22890 | CD630_Cdi1_5 | 31 -- 48 | 435 -- 451 | -13.4676 | 3.75591 | 4.83391 |
| feoB | CD630_Cdi1_5 | 131 -- 197 | 80 -- 141 | -13.454 | 4.05364 | 4.83391 |
| thiH | CD630_Cdi1_5 | 169 -- 196 | 264 -- 288 | -13.4322 | 3.58230 | 4.83391 |
| CD630_28560 | CD630_Cdi1_5 | 64 -- 92 | 422 -- 448 | -13.4048 | 3.96803 | 4.83391 |
| CD630_30310 | CD630_Cdi1_5 | 70 -- 93 | 422 -- 444 | -13.3965 | 4.56059 | 4.83391 |
| cdu2 | CD630_Cdi1_5 | 82 -- 139 | 420 -- 463 | -13.3911 | 2.97335 | 4.83391 |
| CD630_05490 | CD630_Cdi1_5 | 155 -- 184 | 423 -- 449 | -13.389 | 3.89852 | 4.83391 |
| ldh | CD630_Cdi1_5 | 39 -- 97 | 343 -- 408 | -13.3818 | 4.35818 | 4.83391 |
| CD630_11240 | CD630_Cdi1_5 | 83 -- 108 | 436 -- 459 | -13.3655 | 3.20274 | 4.83391 |
| CD630_23000 | CD630_Cdi1_5 | 22 -- 33 | 422 -- 433 | -13.3026 | 3.54130 | 4.83391 |
| CD630_03420 | CD630_Cdi1_5 | 138 -- 175 | 106 -- 147 | -13.263 | 4.71868 | 4.83391 |
| CD630_19790 | CD630_Cdi1_5 | 85 -- 125 | 174 -- 209 | -13.1999 | 3.79593 | 4.83391 |
| CD630_14190 | CD630_Cdi1_5 | 82 -- 106 | 438 -- 463 | -13.1964 | 3.78562 | 4.83391 |
| CD630_22200 | CD630_Cdi1_5 | 141 -- 173 | 108 -- 141 | -13.0444 | 4.18288 | 4.83391 |
| CD630_24770 | CD630_Cdi1_5 | 87 -- 100 | 342 -- 356 | -13.0367 | 3.20798 | 4.83391 |
| CD630_15850 | CD630_Cdi1_5 | 12 -- 39 | 438 -- 463 | -13.0213 | 4.32968 | 4.83391 |
| CD630_16450 | CD630_Cdi1_5 | 88 -- 112 | 431 -- 455 | -12.987 | 4.57547 | 4.83391 |
| cspD | CD630_Cdi1_5 | 160 -- 184 | 427 -- 455 | -12.9794 | 3.23587 | 4.83391 |
| CD630_13360 | CD630_Cdi1_5 | 89 -- 139 | 423 -- 455 | -12.9689 | 3.77330 | 4.83391 |
| CD630_21770 | CD630_Cdi1_5 | 87 -- 196 | 104 -- 205 | -12.9463 | 4.02587 | 4.83391 |
| CD630_08360 | CD630_Cdi1_5 | 161 -- 176 | 436 -- 451 | -12.9092 | 3.19339 | 4.83391 |
| CD630_12320 | CD630_Cdi1_5 | 37 -- 85 | 303 -- 348 | -12.8762 | 3.28667 | 4.83391 |
| scrR | CD630_Cdi1_5 | 134 -- 148 | 344 -- 358 | -12.8178 | 3.49166 | 4.83391 |
| bcp | CD630_Cdi1_5 | 44 -- 97 | 102 -- 157 | -12.8175 | 3.16670 | 4.83391 |
| eutV | CD630_Cdi1_5 | 90 -- 109 | 443 -- 461 | -12.7888 | 4.55715 | 4.83391 |
| CD630_31490 | CD630_Cdi1_5 | 90 -- 122 | 422 -- 456 | -12.783 | 4.08593 | 4.83391 |
| sspA | CD630_Cdi1_5 | 53 -- 109 | 88 -- 149 | -12.7506 | 3.62689 | 4.83391 |
| CD630_25190 | CD630_Cdi1_5 | 91 -- 139 | 104 -- 157 | -12.7137 | 3.19663 | 4.83391 |
| CD630_14150 | CD630_Cdi1_5 | 134 -- 184 | 276 -- 320 | -12.7038 | 3.28251 | 4.83391 |
| CD630_03220 | CD630_Cdi1_5 | 29 -- 70 | 421 -- 452 | -12.6957 | 3.36746 | 4.83391 |
| CD630_19000 | CD630_Cdi1_5 | 83 -- 116 | 177 -- 209 | -12.6884 | 4.18443 | 4.83391 |
| CD630_33300 | CD630_Cdi1_5 | 87 -- 105 | 437 -- 455 | -12.645 | 3.79149 | 4.83391 |
| serS1 | CD630_Cdi1_5 | 153 -- 169 | 435 -- 450 | -12.6403 | 3.45858 | 4.83391 |
| CD630_20170 | CD630_Cdi1_5 | 123 -- 164 | 106 -- 142 | -12.6396 | 6.90201 | 4.83391 |
| etfA | CD630_Cdi1_5 | 24 -- 66 | 114 -- 157 | -12.634 | 3.41204 | 4.83391 |
| CD630_34530 | CD630_Cdi1_5 | 2 -- 50 | 108 -- 158 | -12.6264 | 3.83016 | 4.83391 |
| CD630_05280 | CD630_Cdi1_5 | 153 -- 186 | 173 -- 206 | -12.62 | 3.91665 | 4.83391 |
| CD630_18960 | CD630_Cdi1_5 | 135 -- 153 | 136 -- 154 | -12.5983 | 4.37822 | 4.83391 |
| oppB | CD630_Cdi1_5 | 88 -- 99 | 446 -- 456 | -12.5835 | 2.98328 | 4.83391 |
| CD630_29270 | CD630_Cdi1_5 | 26 -- 57 | 445 -- 471 | -12.556 | 3.70174 | 4.83391 |
| CD630_28181 | CD630_Cdi1_5 | 8 -- 78 | 82 -- 143 | -12.5499 | 3.91299 | 4.83391 |
| CD630_06200 | CD630_Cdi1_5 | 118 -- 150 | 104 -- 144 | -12.5442 | 3.57064 | 4.83391 |
| CD630_17960 | CD630_Cdi1_5 | 126 -- 147 | 445 -- 463 | -12.5128 | 3.71404 | 4.83391 |
| CD630_11220 | CD630_Cdi1_5 | 136 -- 160 | 438 -- 461 | -12.5081 | 3.52327 | 4.83391 |
| CD630_03040 | CD630_Cdi1_5 | 51 -- 95 | 105 -- 152 | -12.4872 | 3.51403 | 4.83391 |
| CD630_14810 | CD630_Cdi1_5 | 144 -- 165 | 428 -- 449 | -12.4786 | 3.35323 | 4.83391 |
| CD630_05590 | CD630_Cdi1_5 | 90 -- 135 | 420 -- 460 | -12.4599 | 3.39670 | 4.83391 |
| CD630_30540 | CD630_Cdi1_5 | 80 -- 124 | 422 -- 456 | -12.4388 | 3.63804 | 4.83391 |
| ung | CD630_Cdi1_5 | 1 -- 38 | 102 -- 141 | -12.3865 | 3.30908 | 4.83391 |
| CD630_02420 | CD630_Cdi1_5 | 18 -- 40 | 439 -- 459 | -12.3819 | 3.71048 | 4.83391 |
| CD630_27110 | CD630_Cdi1_5 | 133 -- 178 | 99 -- 144 | -12.3697 | 3.78231 | 4.83391 |
| CD630_28150 | CD630_Cdi1_5 | 25 -- 36 | 342 -- 353 | -12.365 | 4.18447 | 4.83391 |
| CD630_31580 | CD630_Cdi1_5 | 185 -- 195 | 485 -- 496 | -12.3491 | 3.51756 | 4.83391 |
| CD630_08100 | CD630_Cdi1_5 | 180 -- 198 | 438 -- 453 | -12.2799 | 4.39181 | 4.83391 |
| gcvTPA | CD630_Cdi1_5 | 87 -- 111 | 433 -- 457 | -12.2584 | 4.77229 | 4.83391 |
| CD630_15740 | CD630_Cdi1_5 | 48 -- 93 | 422 -- 450 | -12.2355 | 4.43222 | 4.83391 |
| CD630_20171 | CD630_Cdi1_5 | 151 -- 179 | 420 -- 447 | -12.1995 | 6.87491 | 4.83391 |
| CD630_23280 | CD630_Cdi1_5 | 99 -- 134 | 422 -- 461 | -12.1974 | 3.50093 | 4.83391 |
| CD630_20670 | CD630_Cdi1_5 | 74 -- 112 | 177 -- 219 | -12.1971 | 4.42757 | 4.83391 |
| CD630_31910 | CD630_Cdi1_5 | 84 -- 121 | 423 -- 457 | -12.1832 | 3.48314 | 4.83391 |
| CD630_14260 | CD630_Cdi1_5 | 96 -- 109 | 438 -- 451 | -12.1822 | 3.22214 | 4.83391 |
| CD630_27200 | CD630_Cdi1_5 | 1 -- 36 | 271 -- 295 | -12.139 | 3.42516 | 4.83391 |
| CD630_18950 | CD630_Cdi1_5 | 180 -- 194 | 79 -- 92 | -12.134 | 3.40610 | 4.83391 |
| CD630_30100 | CD630_Cdi1_5 | 54 -- 80 | 175 -- 202 | -12.0949 | 4.13234 | 4.83391 |

| | | | | | | |
|---|---|---|---|---|---|---|
| CD630_01590 | CD630_Cdi1_5 | 155 -- 201 | 177 -- 225 | -12.0931 | 3.45609 | 4.83391 |
| CD630_28290 | CD630_Cdi1_5 | 86 -- 94 | 449 -- 457 | -12.0919 | 5.54192 | 4.83391 |
| CD630_16160 | CD630_Cdi1_5 | 87 -- 111 | 432 -- 459 | -12.0844 | 6.11801 | 4.83391 |
| CD630_22050 | CD630_Cdi1_5 | 82 -- 104 | 440 -- 464 | -12.0701 | 4.87639 | 4.83391 |
| ogt1 | CD630_Cdi1_5 | 63 -- 93 | 422 -- 446 | -12.0277 | 3.16772 | 4.83391 |
| CD630_16460 | CD630_Cdi1_5 | 15 -- 86 | 165 -- 244 | -12.0086 | 4.64028 | 4.83391 |
| CD630_21090 | CD630_Cdi1_5 | 131 -- 188 | 105 -- 154 | -11.9649 | 4.15803 | 4.83391 |
| CD630_22550 | CD630_Cdi1_5 | 84 -- 104 | 440 -- 464 | -11.9647 | 3.62379 | 4.83391 |
| bglA | CD630_Cdi1_5 | 86 -- 112 | 423 -- 458 | -11.9203 | 4.50315 | 4.83391 |
| CD630_14360 | CD630_Cdi1_5 | 86 -- 106 | 438 -- 457 | -11.9153 | 3.02079 | 4.83391 |
| CD630_17730 | CD630_Cdi1_5 | 87 -- 109 | 435 -- 453 | -11.858 | 4.73397 | 4.83391 |
| CD630_21540 | CD630_Cdi1_5 | 77 -- 106 | 438 -- 463 | -11.8159 | 4.33090 | 4.83391 |
| CD630_23820 | CD630_Cdi1_5 | 88 -- 106 | 338 -- 354 | -11.776 | 3.18419 | 4.83391 |
| CD630_14340 | CD630_Cdi1_5 | 98 -- 111 | 433 -- 446 | -11.7328 | 5.10868 | 4.83391 |
| CD630_13780 | CD630_Cdi1_5 | 130 -- 176 | 174 -- 209 | -11.7237 | 3.52274 | 4.83391 |
| CD630_32050 | CD630_Cdi1_5 | 176 -- 199 | 184 -- 206 | -11.716 | 5.73797 | 4.83391 |
| CD630_14130 | CD630_Cdi1_5 | 163 -- 190 | 423 -- 450 | -11.7074 | 4.57970 | 4.83391 |
| CD630_10000 | CD630_Cdi1_5 | 52 -- 88 | 422 -- 448 | -11.6834 | 3.86040 | 4.83391 |
| CD630_29650 | CD630_Cdi1_5 | 87 -- 111 | 422 -- 453 | -11.6834 | 3.67128 | 4.83391 |
| CD630_15270 | CD630_Cdi1_5 | 89 -- 111 | 433 -- 455 | -11.6832 | 4.66015 | 4.83391 |
| CD630_15550 | CD630_Cdi1_5 | 88 -- 118 | 426 -- 457 | -11.678 | 3.49019 | 4.83391 |
| CD630_20460 | CD630_Cdi1_5 | 98 -- 112 | 441 -- 454 | -11.6743 | 7.94034 | 4.83391 |
| CD630_21910 | CD630_Cdi1_5 | 85 -- 96 | 453 -- 464 | -11.6713 | 4.66600 | 4.83391 |
| CD630_28580 | CD630_Cdi1_5 | 48 -- 64 | 337 -- 356 | -11.653 | 4.67130 | 4.83391 |
| CD630_07870 | CD630_Cdi1_5 | 90 -- 109 | 435 -- 456 | -11.641 | 3.22829 | 4.83391 |
| CD630_31010 | CD630_Cdi1_5 | 41 -- 98 | 102 -- 157 | -11.6009 | 4.96305 | 4.83391 |
| CD630_28260 | CD630_Cdi1_5 | 66 -- 104 | 419 -- 462 | -11.5845 | 3.51613 | 4.83391 |
| CD630_09772 | CD630_Cdi1_5 | 39 -- 96 | 104 -- 158 | -11.5683 | 3.66033 | 4.83391 |
| CD630_31461 | CD630_Cdi1_5 | 6 -- 25 | 132 -- 149 | -11.5253 | 4.41476 | 4.83391 |
| cwp29 | CD630_Cdi1_5 | 89 -- 157 | 396 -- 453 | -11.5236 | 4.40865 | 4.83391 |
| CD630_09040 | CD630_Cdi1_5 | 91 -- 139 | 104 -- 157 | -11.5059 | 4.11851 | 4.83391 |
| CD630_22030 | CD630_Cdi1_5 | 7 -- 23 | 44 -- 60 | -11.4811 | 3.29934 | 4.83391 |
| uxaA | CD630_Cdi1_5 | 72 -- 91 | 422 -- 451 | -11.4688 | 4.26165 | 4.83391 |
| CD630_18610 | CD630_Cdi1_5 | 15 -- 25 | 435 -- 445 | -11.4671 | 4.15582 | 4.83391 |
| garK | CD630_Cdi1_5 | 123 -- 178 | 422 -- 470 | -11.4461 | 3.72836 | 4.83391 |
| CD630_05870 | CD630_Cdi1_5 | 87 -- 106 | 438 -- 457 | -11.4358 | 4.28339 | 4.83391 |
| CD630_33560 | CD630_Cdi1_5 | 57 -- 94 | 106 -- 142 | -11.4321 | 4.03147 | 4.83391 |
| CD630_15431 | CD630_Cdi1_5 | 125 -- 185 | 102 -- 157 | -11.4213 | 5.29780 | 4.83391 |
| CD630_34030 | CD630_Cdi1_5 | 42 -- 66 | 127 -- 154 | -11.4158 | 3.05766 | 4.83391 |
| CD630_17710 | CD630_Cdi1_5 | 87 -- 108 | 436 -- 457 | -11.4129 | 4.37688 | 4.83391 |
| CD630_27120 | CD630_Cdi1_5 | 113 -- 161 | 266 -- 316 | -11.4069 | 4.43720 | 4.83391 |
| CD630_10880 | CD630_Cdi1_5 | 86 -- 105 | 423 -- 450 | -11.4053 | 5.05048 | 4.83391 |
| dnaC | CD630_Cdi1_5 | 145 -- 175 | 178 -- 206 | -11.4036 | 6.30796 | 4.83391 |
| CD630_29960 | CD630_Cdi1_5 | 135 -- 201 | 65 -- 138 | -11.3727 | 4.51904 | 4.83391 |
| CD630_18311 | CD630_Cdi1_5 | 88 -- 113 | 174 -- 203 | -11.3695 | 6.86047 | 4.83391 |
| ribD | CD630_Cdi1_5 | 85 -- 109 | 431 -- 456 | -11.367 | 6.65922 | 4.83391 |
| CD630_22880 | CD630_Cdi1_5 | 175 -- 199 | 125 -- 150 | -11.3654 | 3.62455 | 4.83391 |
| cspC | CD630_Cdi1_5 | 160 -- 184 | 427 -- 455 | -11.3468 | 5.48848 | 4.83391 |
| CD630_32890 | CD630_Cdi1_5 | 85 -- 118 | 423 -- 458 | -11.3375 | 3.41276 | 4.83391 |
| CD630_20660 | CD630_Cdi1_5 | 134 -- 167 | 108 -- 150 | -11.3333 | 5.39321 | 4.83391 |
| cobC | CD630_Cdi1_5 | 164 -- 198 | 104 -- 141 | -11.3263 | 3.15854 | 4.83391 |
| CD630_17970 | CD630_Cdi1_5 | 164 -- 174 | 436 -- 446 | -11.3201 | 3.68802 | 4.83391 |
| CD630_24760 | CD630_Cdi1_5 | 75 -- 139 | 363 -- 423 | -11.3172 | 5.19485 | 4.83391 |
| CD630_20260 | CD630_Cdi1_5 | 86 -- 114 | 176 -- 205 | -11.3113 | 3.84162 | 4.83391 |
| agrB | CD630_Cdi1_5 | 17 -- 61 | 371 -- 411 | -11.3087 | 3.92598 | 4.83391 |
| glgB | CD630_Cdi1_5 | 87 -- 112 | 435 -- 452 | -11.3037 | 4.48273 | 4.83391 |
| CD630_22850 | CD630_Cdi1_5 | 167 -- 201 | 105 -- 149 | -11.2843 | 3.84102 | 4.83391 |
| vncR | CD630_Cdi1_5 | 138 -- 157 | 183 -- 206 | -11.282 | 3.93642 | 4.83391 |
| CD630_32990 | CD630_Cdi1_5 | 35 -- 68 | 211 -- 254 | -11.2777 | 5.24925 | 4.83391 |
| CD630_17151 | CD630_Cdi1_5 | 166 -- 197 | 421 -- 447 | -11.2695 | 6.43124 | 4.83391 |
| CD630_06340 | CD630_Cdi1_5 | 107 -- 135 | 271 -- 300 | -11.2686 | 4.35983 | 4.83391 |
| CD630_34080 | CD630_Cdi1_5 | 83 -- 106 | 423 -- 447 | -11.2174 | 3.20224 | 4.83391 |
| CD630_31220 | CD630_Cdi1_5 | 85 -- 105 | 423 -- 447 | -11.2148 | 3.73779 | 4.83391 |
| CD630_11730 | CD630_Cdi1_5 | 150 -- 179 | 420 -- 451 | -11.2098 | 4.80521 | 4.83391 |
| CD630_16950 | CD630_Cdi1_5 | 185 -- 196 | 81 -- 93 | -11.203 | 4.42569 | 4.83391 |
| CD630_12110 | CD630_Cdi1_5 | 53 -- 61 | 449 -- 457 | -11.2018 | 4.36078 | 4.83391 |
| CD630_18650 | CD630_Cdi1_5 | 88 -- 113 | 423 -- 458 | -11.1843 | 3.75892 | 4.83391 |
| srlEb | CD630_Cdi1_5 | 79 -- 97 | 440 -- 457 | -11.1599 | 3.50140 | 4.83391 |
| glnA | CD630_Cdi1_5 | 151 -- 187 | 108 -- 147 | -11.1302 | 3.87629 | 4.83391 |
| CD630_36040 | CD630_Cdi1_5 | 64 -- 125 | 420 -- 461 | -11.1036 | 3.52009 | 4.83391 |
| CD630_13400 | CD630_Cdi1_5 | 67 -- 118 | 105 -- 150 | -11.1024 | 3.20232 | 4.83391 |
| CD630_33580 | CD630_Cdi1_5 | 91 -- 99 | 449 -- 457 | -11.0948 | 3.95778 | 4.83391 |
| ssuC | CD630_Cdi1_5 | 61 -- 95 | 421 -- 463 | -11.091 | 3.45491 | 4.83391 |
| CD630_09970 | CD630_Cdi1_5 | 3 -- 39 | 103 -- 140 | -11.0906 | 3.25917 | 4.83391 |
| CD630_15150 | CD630_Cdi1_5 | 8 -- 52 | 103 -- 156 | -11.0672 | 3.35564 | 4.83391 |
| CD630_32690 | CD630_Cdi1_5 | 3 -- 34 | 426 -- 458 | -11.0371 | 7.32676 | 4.83391 |
| CD630_04610 | CD630_Cdi1_5 | 48 -- 71 | 108 -- 139 | -11.0077 | 3.83025 | 4.83391 |
| CD630_10960 | CD630_Cdi1_5 | 87 -- 131 | 423 -- 456 | -10.9334 | 3.53850 | 4.83391 |
| CD630_21080 | CD630_Cdi1_5 | 89 -- 105 | 435 -- 450 | -10.931 | 3.99742 | 4.83391 |
| CD630_14710 | CD630_Cdi1_5 | 87 -- 97 | 445 -- 456 | -10.9277 | 3.65171 | 4.83391 |
| tcdC | CD630_Cdi1_5 | 112 -- 158 | 274 -- 314 | -10.9238 | 3.76101 | 4.83391 |
| CD630_24580 | CD630_Cdi1_5 | 85 -- 141 | 403 -- 455 | -10.908 | 3.88519 | 4.83391 |
| spoVFB | CD630_Cdi1_5 | 83 -- 129 | 421 -- 464 | -10.8813 | 4.62758 | 4.83391 |
| CD630_06150 | CD630_Cdi1_5 | 99 -- 128 | 103 -- 124 | -10.8707 | 5.51744 | 4.83391 |
| CD630_18090 | CD630_Cdi1_5 | 10 -- 32 | 184 -- 205 | -10.8666 | 4.23961 | 4.83391 |
| CD630_13350 | CD630_Cdi1_5 | 163 -- 181 | 127 -- 142 | -10.865 | 4.15139 | 4.83391 |
| CD630_13880 | CD630_Cdi1_5 | 41 -- 55 | 439 -- 453 | -10.8599 | 3.52735 | 4.83391 |
| CD630_15900 | CD630_Cdi1_5 | 84 -- 128 | 423 -- 463 | -10.8485 | 3.17664 | 4.83391 |
| CD630_04660 | CD630_Cdi1_5 | 91 -- 105 | 435 -- 449 | -10.8452 | 4.32791 | 4.83391 |
| CD630_01690 | CD630_Cdi1_5 | 14 -- 45 | 113 -- 144 | -10.8061 | 4.97234 | 4.83391 |
| CD630_21070 | CD630_Cdi1_5 | 185 -- 196 | 344 -- 355 | -10.7953 | 5.35019 | 4.83391 |
| glgP | CD630_Cdi1_5 | 71 -- 115 | 420 -- 456 | -10.7737 | 3.28530 | 4.83391 |
| CD630_09100 | CD630_Cdi1_5 | 2 -- 50 | 20 -- 61 | -10.7656 | 5.92260 | 4.83391 |
| CD630_32520 | CD630_Cdi1_5 | 135 -- 161 | 332 -- 355 | -10.7391 | 4.06098 | 4.83391 |
| CD630_19401 | CD630_Cdi1_5 | 35 -- 82 | 118 -- 165 | -10.7356 | 4.37084 | 4.83391 |
| opuCA | CD630_Cdi1_5 | 87 -- 112 | 175 -- 206 | -10.7343 | 4.07035 | 4.83391 |
| CD630_23760 | CD630_Cdi1_5 | 82 -- 105 | 186 -- 209 | -10.7317 | 4.54307 | 4.83391 |
| pgm | CD630_Cdi1_5 | 167 -- 177 | 444 -- 454 | -10.7273 | 3.45439 | 4.83391 |
| CD630_02740 | CD630_Cdi1_5 | 88 -- 114 | 426 -- 456 | -10.7227 | 5.02277 | 4.83391 |
| pilT | CD630_Cdi1_5 | 8 -- 26 | 297 -- 316 | -10.7175 | 3.24639 | 4.83391 |
| CD630_05820 | CD630_Cdi1_5 | 87 -- 93 | 449 -- 455 | -10.7086 | 3.75661 | 4.83391 |
| CD630_02430 | CD630_Cdi1_5 | 39 -- 84 | 110 -- 138 | -10.6998 | 3.71048 | 4.83391 |
| uxaA%27 | CD630_Cdi1_5 | 86 -- 108 | 438 -- 457 | -10.6601 | 4.26165 | 4.83391 |
| appF | CD630_Cdi1_5 | 84 -- 129 | 421 -- 463 | -10.6545 | 4.25570 | 4.83391 |
| CD630_25040 | CD630_Cdi1_5 | 144 -- 177 | 19 -- 54 | -10.6491 | 3.43898 | 4.83391 |
| CD630_08390 | CD630_Cdi1_5 | 81 -- 101 | 423 -- 443 | -10.616 | 5.16000 | 4.83391 |
| folC | CD630_Cdi1_5 | 87 -- 125 | 421 -- 458 | -10.5653 | 3.85634 | 4.83391 |
| CD630_25620 | CD630_Cdi1_5 | 88 -- 107 | 437 -- 457 | -10.5636 | 3.02685 | 4.83391 |
| CD630_33420 | CD630_Cdi1_5 | 88 -- 105 | 435 -- 451 | -10.5599 | 3.22095 | 4.83391 |
| CD630_34560 | CD630_Cdi1_5 | 84 -- 115 | 432 -- 461 | -10.5498 | 4.02604 | 4.83391 |
| CD630_31880 | CD630_Cdi1_5 | 15 -- 34 | 15 -- 35 | -10.547 | 3.97522 | 4.83391 |
| nusB | CD630_Cdi1_5 | 88 -- 157 | 281 -- 353 | -10.5366 | 4.92798 | 4.83391 |
| CD630_30120 | CD630_Cdi1_5 | 43 -- 100 | 104 -- 157 | -10.5294 | 4.04859 | 4.83391 |
| CD630_23770 | CD630_Cdi1_5 | 52 -- 93 | 108 -- 151 | -10.5083 | 3.25181 | 4.83391 |
| CD630_19030 | CD630_Cdi1_5 | 89 -- 115 | 418 -- 440 | -10.4993 | 3.34240 | 4.83391 |
| CD630_27310 | CD630_Cdi1_5 | 135 -- 177 | 101 -- 141 | -10.4939 | 4.01622 | 4.83391 |
| CD630_27610 | CD630_Cdi1_5 | 5 -- 52 | 271 -- 316 | -10.4844 | 3.96465 | 4.83391 |
| CD630_04880 | CD630_Cdi1_5 | 125 -- 152 | 336 -- 355 | -10.4703 | 4.62291 | 4.83391 |
| CD630_08940 | CD630_Cdi1_5 | 75 -- 91 | 67 -- 83 | -10.47 | 3.72225 | 4.83391 |
| CD630_08090 | CD630_Cdi1_5 | 143 -- 174 | 102 -- 132 | -10.4487 | 3.57167 | 4.83391 |
| CD630_18010 | CD630_Cdi1_5 | 154 -- 186 | 125 -- 156 | -10.4453 | 4.21967 | 4.83391 |
| blaI | CD630_Cdi1_5 | 87 -- 104 | 440 -- 458 | -10.4283 | 3.73674 | 4.83391 |
| CD630_16530 | CD630_Cdi1_5 | 87 -- 166 | 387 -- 460 | -10.4261 | 4.37658 | 4.83391 |
| CD630_21760 | CD630_Cdi1_5 | 64 -- 74 | 82 -- 91 | -10.4184 | 4.43846 | 4.83391 |
| CD630_06920 | CD630_Cdi1_5 | 83 -- 125 | 420 -- 458 | -10.4146 | 3.13260 | 4.83391 |
| cdtR | CD630_Cdi1_5 | 76 -- 92 | 422 -- 438 | -10.4081 | 5.84368 | 4.83391 |
| CD630_25310 | CD630_Cdi1_5 | 160 -- 176 | 294 -- 310 | -10.3925 | 3.99861 | 4.83391 |
| CD630_09760 | CD630_Cdi1_5 | 90 -- 125 | 173 -- 206 | -10.3721 | 4.83410 | 4.83391 |
| CD630_28910 | CD630_Cdi1_5 | 90 -- 125 | 173 -- 206 | -10.3721 | 4.51664 | 4.83391 |
| CD630_09850 | CD630_Cdi1_5 | 87 -- 104 | 440 -- 457 | -10.3662 | 4.18679 | 4.83391 |
| CD630_03930 | CD630_Cdi1_5 | 87 -- 113 | 440 -- 461 | -10.3194 | 4.15934 | 4.83391 |
| CD630_15060 | CD630_Cdi1_5 | 173 -- 191 | 220 -- 237 | -10.2936 | 3.69396 | 4.83391 |
| CD630_33190 | CD630_Cdi1_5 | 115 -- 126 | 177 -- 188 | -10.2735 | 4.13899 | 4.83391 |
| opuCC | CD630_Cdi1_5 | 95 -- 144 | 108 -- 157 | -10.2683 | 4.07035 | 4.83391 |
| CD630_25070 | CD630_Cdi1_5 | 24 -- 46 | 425 -- 446 | -10.249 | 3.45164 | 4.83391 |
| CD630_17360 | CD630_Cdi1_5 | 85 -- 122 | 422 -- 457 | -10.2472 | 5.66070 | 4.83391 |
| CD630_06690 | CD630_Cdi1_5 | 86 -- 96 | 343 -- 353 | -10.243 | 5.82709 | 4.83391 |
| CD630_28280 | CD630_Cdi1_5 | 97 -- 105 | 418 -- 426 | -10.2427 | 4.59833 | 4.83391 |
| CD630_27270 | CD630_Cdi1_5 | 89 -- 118 | 435 -- 455 | -10.2354 | 3.50892 | 4.83391 |
| cwp26 | CD630_Cdi1_5 | 131 -- 161 | 135 -- 165 | -10.2308 | 4.44377 | 4.83391 |
| CD630_10830 | CD630_Cdi1_5 | 86 -- 95 | 345 -- 355 | -10.213 | 4.82398 | 4.83391 |
| CD630_13380 | CD630_Cdi1_5 | 177 -- 186 | 436 -- 445 | -10.2127 | 4.74408 | 4.83391 |
| CD630_06560 | CD630_Cdi1_5 | 137 -- 187 | 390 -- 452 | -10.2123 | 4.02532 | 4.83391 |
| adhE | CD630_Cdi1_5 | 36 -- 54 | 438 -- 456 | -10.2052 | 3.06485 | 4.83391 |
| CD630_18400 | CD630_Cdi1_5 | 121 -- 158 | 105 -- 142 | -10.2032 | 4.67962 | 4.83391 |
| pilW | CD630_Cdi1_5 | 88 -- 118 | 423 -- 452 | -10.1987 | 4.96316 | 4.83391 |
| cdu1 | CD630_Cdi1_5 | 87 -- 96 | 345 -- 355 | -10.192 | 4.46276 | 4.83391 |
| CD630_06110 | CD630_Cdi1_5 | 124 -- 138 | 304 -- 318 | -10.1733 | 5.61380 | 4.83391 |
| CD630_06190 | CD630_Cdi1_5 | 141 -- 151 | 41 -- 51 | -10.1713 | 3.57064 | 4.83391 |
| CD630_27220 | CD630_Cdi1_5 | 86 -- 112 | 439 -- 463 | -10.1647 | 4.18786 | 4.83391 |
| CD630_18920 | CD630_Cdi1_5 | 170 -- 194 | 438 -- 459 | -10.1564 | 4.43558 | 4.83391 |
| CD630_03650 | CD630_Cdi1_5 | 85 -- 94 | 420 -- 429 | -10.1251 | 3.93697 | 4.83391 |
| CD630_08510 | CD630_Cdi1_5 | 42 -- 76 | 171 -- 208 | -10.1198 | 5.16329 | 4.83391 |
| iorA | CD630_Cdi1_5 | 57 -- 98 | 431 -- 453 | -10.1168 | 3.83173 | 4.83391 |
| CD630_05450 | CD630_Cdi1_5 | 85 -- 92 | 451 -- 458 | -10.0847 | 5.04761 | 4.83391 |
| CD630_19801 | CD630_Cdi1_5 | 88 -- 98 | 343 -- 353 | -10.0835 | 5.22035 | 4.83391 |
| CD630_05150 | CD630_Cdi1_5 | 87 -- 109 | 423 -- 450 | -10.0772 | 3.69875 | 4.83391 |
| CD630_35840 | CD630_Cdi1_5 | 80 -- 92 | 452 -- 464 | -10.072 | 3.14170 | 4.83391 |
| CD630_27360 | CD630_Cdi1_5 | 184 -- 196 | 395 -- 407 | -10.0665 | 3.79570 | 4.83391 |
| feoB | CD630_Cdi1_5 | 85 -- 124 | 423 -- 462 | -10.0663 | 4.52257 | 4.83391 |
| CD630_06350 | CD630_Cdi1_5 | 138 -- 167 | 177 -- 206 | -10.0352 | 4.35983 | 4.83391 |
| CD630_14050 | CD630_Cdi1_5 | 186 -- 201 | 191 -- 206 | -10.0271 | 3.30894 | 4.83391 |
| CD630_03160 | CD630_Cdi1_5 | 95 -- 123 | 268 -- 295 | -10.0235 | 3.66946 | 4.83391 |
| CD630_01700 | CD630_Cdi1_5 | 85 -- 98 | 442 -- 457 | -10.0149 | 4.97234 | 4.83391 |
| CD630_32040 | CD630_Cdi1_5 | 24 -- 42 | 250 -- 274 | -10.0114 | 4.93516 | 4.83391 |
| CD630_06060 | CD630_Cdi1_5 | 41 -- 59 | 342 -- 355 | -9.9773 | 4.94050 | 4.83391 |
| CD630_16520 | CD630_Cdi1_5 | 85 -- 104 | 424 -- 451 | -9.9667 | 4.28395 | 4.83391 |
| CD630_06050 | CD630_Cdi1_5 | 74 -- 90 | 439 -- 453 | -9.96635 | 3.59332 | 4.83391 |
| pilC | CD630_Cdi1_5 | 129 -- 139 | 453 -- 463 | -9.95027 | 3.54230 | 4.83391 |
| CD630_36190 | CD630_Cdi1_5 | 73 -- 128 | 440 -- 465 | -9.93539 | 4.63449 | 4.83391 |
| CD630_05230 | CD630_Cdi1_5 | 82 -- 109 | 420 -- 450 | -9.93412 | 3.73787 | 4.83391 |
| cynT | CD630_Cdi1_5 | 146 -- 169 | 101 -- 120 | -9.92724 | 4.21370 | 4.83391 |
| CD630_09060 | CD630_Cdi1_5 | 2 -- 23 | 372 -- 393 | -9.92078 | 4.31834 | 4.83391 |
| CD630_22510 | CD630_Cdi1_5 | 114 -- 127 | 218 -- 231 | -9.88346 | 11.42763 | 4.83391 |
| CD630_13600 | CD630_Cdi1_5 | 129 -- 176 | 437 -- 463 | -9.87758 | 4.45481 | 4.83391 |
| etfA | CD630_Cdi1_5 | 124 -- 163 | 172 -- 205 | -9.8759 | 4.80521 | 4.83391 |
| CD630_29500 | CD630_Cdi1_5 | 47 -- 69 | 196 -- 219 | -9.8505 | 3.70644 | 4.83391 |
| CD630_06060 | CD630_Cdi1_5 | 87 -- 94 | 451 -- 458 | -9.84039 | 4.94050 | 4.83391 |
| rolA | CD630_Cdi1_5 | 33 -- 56 | 300 -- 323 | -9.81817 | 4.56362 | 4.83391 |
| cls | CD630_Cdi1_5 | 86 -- 93 | 451 -- 458 | -9.80556 | 4.46177 | 4.83391 |
| CD630_36180 | CD630_Cdi1_5 | 176 -- 183 | 451 -- 458 | -9.80013 | 4.83428 | 4.83391 |
| CD630_32880 | CD630_Cdi1_5 | 172 -- 190 | 271 -- 291 | -9.77799 | 3.41276 | 4.83391 |
| CD630_01560 | CD630_Cdi1_5 | 142 -- 151 | 369 -- 378 | -9.76101 | 4.06352 | 4.83391 |
| CD630_29760 | CD630_Cdi1_5 | 86 -- 93 | 451 -- 458 | -9.76066 | 3.51074 | 4.83391 |
| CD630_24920 | CD630_Cdi1_5 | 89 -- 150 | 105 -- 157 | -9.75926 | 3.89455 | 4.83391 |
| CD630_21030 | CD630_Cdi1_5 | 57 -- 107 | 107 -- 156 | -9.75456 | 3.08733 | 4.83391 |
| CD630_36690 | CD630_Cdi1_5 | 111 -- 133 | 292 -- 314 | -9.75431 | 3.53761 | 4.83391 |
| argG | CD630_Cdi1_5 | 87 -- 134 | 422 -- 459 | -9.74716 | 4.80300 | 4.83391 |
| CD630_19904 | CD630_Cdi1_5 | 151 -- 171 | 424 -- 450 | -9.74159 | 5.96526 | 4.83391 |
| CD630_18970 | CD630_Cdi1_5 | 145 -- 185 | 276 -- 314 | -9.72584 | 3.57243 | 4.83391 |
| xylR | CD630_Cdi1_5 | 126 -- 139 | 435 -- 448 | -9.72334 | 3.95671 | 4.83391 |
| bioB | CD630_Cdi1_5 | 131 -- 186 | 264 -- 317 | -9.69772 | 3.59022 | 4.83391 |
| CD630_23070 | CD630_Cdi1_5 | 5 -- 16 | 23 -- 34 | -9.6858 | 5.23027 | 4.83391 |
| CD630_20980 | CD630_Cdi1_5 | 101 -- 110 | 80 -- 89 | -9.67716 | 8.12847 | 4.83391 |
| CD630_05950 | CD630_Cdi1_5 | 34 -- 69 | 173 -- 205 | -9.66918 | 3.31669 | 4.83391 |
| CD630_25320 | CD630_Cdi1_5 | 144 -- 173 | 177 -- 206 | -9.65909 | 4.29415 | 4.83391 |
| CD630_27280 | CD630_Cdi1_5 | 146 -- 156 | 82 -- 93 | -9.62835 | 3.24738 | 4.83391 |
| CD630_14600 | CD630_Cdi1_5 | 65 -- 101 | 119 -- 156 | -9.62582 | 5.90348 | 4.83391 |
| CD630_22070 | CD630_Cdi1_5 | 42 -- 53 | 438 -- 450 | -9.62028 | 4.09651 | 4.83391 |
| CD630_04550 | CD630_Cdi1_5 | 88 -- 123 | 421 -- 457 | -9.61782 | 3.05227 | 4.83391 |
| CD630_34520 | CD630_Cdi1_5 | 77 -- 113 | 107 -- 139 | -9.61507 | 3.83016 | 4.83391 |
| CD630_24591 | CD630_Cdi1_5 | 88 -- 96 | 345 -- 353 | -9.61384 | 3.54497 | 4.83391 |
| CD630_30130 | CD630_Cdi1_5 | 2 -- 28 | 175 -- 209 | -9.59855 | 3.28343 | 4.83391 |
| aroD | CD630_Cdi1_5 | 137 -- 176 | 423 -- 455 | -9.59202 | 3.71929 | 4.83391 |
| CD630_31500 | CD630_Cdi1_5 | 19 -- 37 | 300 -- 317 | -9.58198 | 4.90207 | 4.83391 |
| CD630_09840 | CD630_Cdi1_5 | 162 -- 177 | 37 -- 51 | -9.5399 | 4.50009 | 4.83391 |
| CD630_07760 | CD630_Cdi1_5 | 80 -- 94 | 422 -- 446 | -9.53579 | 3.64186 | 4.83391 |
| CD630_19840 | CD630_Cdi1_5 | 90 -- 96 | 450 -- 456 | -9.52671 | 3.44724 | 4.83391 |
| CD630_15460 | CD630_Cdi1_5 | 88 -- 97 | 345 -- 355 | -9.5186 | 6.86296 | 4.83391 |
| CD630_13020 | CD630_Cdi1_5 | 175 -- 200 | 426 -- 453 | -9.50919 | 3.52267 | 4.83391 |
| CD630_36690 | CD630_Cdi2_2 | 148 -- 169 | 64 -- 85 | -18.7481 | 3.53761 | 6.72203 |
| CD630_36690 | CD630_Cdi2_2 | 148 -- 169 | 64 -- 85 | -18.7481 | 3.53761 | 6.72203 |
| CD630_30100 | CD630_Cdi2_2 | 125 -- 182 | 36 -- 84 | -17.9463 | 4.13234 | 6.72203 |
| CD630_30100 | CD630_Cdi2_2 | 125 -- 182 | 36 -- 84 | -17.9463 | 4.13234 | 6.72203 |
| cwp26 | CD630_Cdi2_2 | 142 -- 179 | 37 -- 85 | -15.5178 | 4.44377 | 6.72203 |
| cwp26 | CD630_Cdi2_2 | 142 -- 179 | 37 -- 85 | -15.5178 | 4.44377 | 6.72203 |
| CD630_27280 | CD630_Cdi2_2 | 131 -- 196 | 17 -- 86 | -14.4628 | 3.24738 | 6.72203 |
| CD630_27280 | CD630_Cdi2_2 | 131 -- 196 | 17 -- 86 | -14.4628 | 3.24738 | 6.72203 |
| CD630_13380 | CD630_Cdi2_2 | 118 -- 154 | 54 -- 85 | -14.2087 | 4.74408 | 6.72203 |
| CD630_13380 | CD630_Cdi2_2 | 118 -- 154 | 54 -- 85 | -14.2087 | 4.74408 | 6.72203 |
| CD630_19430 | CD630_Cdi2_2 | 160 -- 197 | 15 -- 47 | -13.4728 | 3.13922 | 6.72203 |
| CD630_19430 | CD630_Cdi2_2 | 160 -- 197 | 15 -- 47 | -13.4728 | 3.13922 | 6.72203 |
| CD630_13020 | CD630_Cdi2_2 | 115 -- 139 | 31 -- 53 | -13.1883 | 3.52267 | 6.72203 |
| CD630_13020 | CD630_Cdi2_2 | 115 -- 139 | 31 -- 53 | -13.1883 | 3.52267 | 6.72203 |
| CD630_36170 | CD630_Cdi2_2 | 157 -- 188 | 22 -- 44 | -13.1883 | 3.72380 | 6.72203 |
| CD630_36170 | CD630_Cdi2_2 | 157 -- 188 | 22 -- 44 | -13.1883 | 3.72380 | 6.72203 |
| CD630_27200 | CD630_Cdi2_2 | 9 -- 20 | 33 -- 44 | -13.0564 | 3.42516 | 6.72203 |
| CD630_27200 | CD630_Cdi2_2 | 9 -- 20 | 33 -- 44 | -13.0564 | 3.42516 | 6.72203 |
| CD630_33560 | CD630_Cdi2_2 | 124 -- 136 | 33 -- 45 | -12.9024 | 4.03147 | 6.72203 |
| CD630_33560 | CD630_Cdi2_2 | 124 -- 136 | 33 -- 45 | -12.9024 | 4.03147 | 6.72203 |
| cheW | CD630_Cdi2_2 | 106 -- 121 | 36 -- 50 | -12.3557 | 3.72371 | 6.72203 |
| cheW | CD630_Cdi2_2 | 106 -- 121 | 36 -- 50 | -12.3557 | 3.72371 | 6.72203 |
| CD630_21090 | CD630_Cdi2_2 | 130 -- 184 | 11 -- 86 | -12.1164 | 4.15803 | 6.72203 |
| CD630_21090 | CD630_Cdi2_2 | 130 -- 184 | 11 -- 86 | -12.1164 | 4.15803 | 6.72203 |
| CD630_17360 | CD630_Cdi2_2 | 5 -- 19 | 48 -- 62 | -12.0025 | 5.66070 | 6.72203 |
| CD630_17360 | CD630_Cdi2_2 | 5 -- 19 | 48 -- 62 | -12.0025 | 5.66070 | 6.72203 |
| CD630_25610 | CD630_Cdi2_2 | 85 -- 149 | 15 -- 80 | -11.6166 | 3.02685 | 6.72203 |
| CD630_25610 | CD630_Cdi2_2 | 85 -- 149 | 15 -- 80 | -11.6166 | 3.02685 | 6.72203 |

| | | | | | | |
|---|---|---|---|---|---|---|
| CD630_19801 | CD630_Cdi2_2 | 115 – 163 | 22 – 79 | -11.5944 | 5.22035 | 6.72203 |
| CD630_19801 | CD630_Cdi2_2 | 115 – 163 | 22 – 79 | -11.5944 | 5.22035 | 6.72203 |
| CD630_19700 | CD630_Cdi2_2 | 153 – 164 | 33 – 45 | -11.2821 | 3.85861 | 6.72203 |
| CD630_19700 | CD630_Cdi2_2 | 153 – 164 | 33 – 45 | -11.2821 | 3.85861 | 6.72203 |
| CD630_15890 | CD630_Cdi2_2 | 149 – 169 | 67 – 84 | -10.945 | 3.39962 | 6.72203 |
| CD630_15890 | CD630_Cdi2_2 | 149 – 169 | 67 – 84 | -10.945 | 3.39962 | 6.72203 |
| CD630_01570 | CD630_Cdi2_2 | 123 – 169 | 27 – 67 | -10.8673 | 6.08699 | 6.72203 |
| CD630_01570 | CD630_Cdi2_2 | 123 – 169 | 27 – 67 | -10.8673 | 6.08699 | 6.72203 |
| CD630_03170 | CD630_Cdi2_2 | 180 – 201 | 61 – 80 | -10.5021 | 3.66946 | 6.72203 |
| CD630_03170 | CD630_Cdi2_2 | 180 – 201 | 61 – 80 | -10.5021 | 3.66946 | 6.72203 |
| CD630_23080 | CD630_Cdi2_2 | 101 – 108 | 35 – 42 | -10.4033 | 5.23027 | 6.72203 |
| CD630_23080 | CD630_Cdi2_2 | 101 – 108 | 35 – 42 | -10.4033 | 5.23027 | 6.72203 |
| CD630_22500 | CD630_Cdi2_2 | 129 – 161 | 44 – 70 | -10.3726 | 4.03688 | 6.72203 |
| CD630_22500 | CD630_Cdi2_2 | 129 – 161 | 44 – 70 | -10.3726 | 4.03688 | 6.72203 |
| CD630_05840 | CD630_Cdi2_2 | 92 – 121 | 44 – 72 | -10.335 | 4.28779 | 6.72203 |
| CD630_05840 | CD630_Cdi2_2 | 92 – 121 | 44 – 72 | -10.335 | 4.28779 | 6.72203 |
| CD630_15270 | CD630_Cdi2_2 | 117 – 192 | 19 – 82 | -10.0887 | 4.66015 | 6.72203 |
| CD630_15270 | CD630_Cdi2_2 | 117 – 192 | 19 – 82 | -10.0887 | 4.66015 | 6.72203 |
| CD630_04320 | CD630_Cdi2_2 | 143 – 154 | 73 – 84 | -10.0522 | 3.68342 | 6.72203 |
| CD630_04320 | CD630_Cdi2_2 | 143 – 154 | 73 – 84 | -10.0522 | 3.68342 | 6.72203 |
| pilD | CD630_Cdi2_2 | 127 – 142 | 70 – 85 | -10.0367 | 4.54123 | 6.72203 |
| pilD | CD630_Cdi2_2 | 127 – 142 | 70 – 85 | -10.0367 | 4.54123 | 6.72203 |
| spaK | CD630_Cdi2_2 | 7 – 44 | 44 – 82 | -10.0192 | 3.04530 | 6.72203 |
| spaK | CD630_Cdi2_2 | 7 – 44 | 44 – 82 | -10.0192 | 3.04530 | 6.72203 |
| kdgT1 | CD630_Cdi2_2 | 9 – 45 | 48 – 84 | -9.99424 | 4.49207 | 6.72203 |
| kdgT1 | CD630_Cdi2_2 | 9 – 45 | 48 – 84 | -9.99424 | 4.49207 | 6.72203 |
| moeA | CD630_Cdi2_2 | 115 – 121 | 38 – 44 | -9.90446 | 3.26118 | 6.72203 |
| moeA | CD630_Cdi2_2 | 115 – 121 | 38 – 44 | -9.90446 | 3.26118 | 6.72203 |
| CD630_23090 | CD630_Cdi2_2 | 26 – 43 | 67 – 87 | -9.88607 | 4.25622 | 6.72203 |
| CD630_23090 | CD630_Cdi2_2 | 26 – 43 | 67 – 87 | -9.88607 | 4.25622 | 6.72203 |
| CD630_06100 | CD630_Cdi2_2 | 21 – 75 | 5 – 50 | -9.87297 | 4.43918 | 6.72203 |
| CD630_06100 | CD630_Cdi2_2 | 21 – 75 | 5 – 50 | -9.87297 | 4.43918 | 6.72203 |
| CD630_17511 | CD630_Cdi2_2 | 4 – 35 | 8 – 47 | -9.69584 | 9.12322 | 6.72203 |
| CD630_17511 | CD630_Cdi2_2 | 4 – 35 | 8 – 47 | -9.69584 | 9.12322 | 6.72203 |
| CD630_29270 | CD630_Cdi2_2 | 6 – 16 | 35 – 45 | -9.6519 | 3.70174 | 6.72203 |
| CD630_29270 | CD630_Cdi2_2 | 6 – 16 | 35 – 45 | -9.6519 | 3.70174 | 6.72203 |
| CD630_16460 | CD630_Cdi2_2 | 132 – 146 | 32 – 45 | -9.63679 | 4.64028 | 6.72203 |
| CD630_16460 | CD630_Cdi2_2 | 132 – 146 | 32 – 45 | -9.63679 | 4.64028 | 6.72203 |
| CD630_18090 | CD630_Cdi2_2 | 7 – 34 | 64 – 84 | -9.62756 | 4.23961 | 6.72203 |
| CD630_18090 | CD630_Cdi2_2 | 7 – 34 | 64 – 84 | -9.62756 | 4.23961 | 6.72203 |
| CD630_36180 | CD630_Cdi2_2 | 109 – 121 | 51 – 63 | -9.58045 | 4.83428 | 6.72203 |
| CD630_36180 | CD630_Cdi2_2 | 109 – 121 | 51 – 63 | -9.58045 | 4.83428 | 6.72203 |
| CD630_14050 | CD630_Cdi2_2 | 154 – 171 | 45 – 63 | -9.55557 | 3.30894 | 6.72203 |
| CD630_14050 | CD630_Cdi2_2 | 154 – 171 | 45 – 63 | -9.55557 | 3.30894 | 6.72203 |
| CD630_34530 | CD630_Cdi2_2 | 65 – 77 | 31 – 42 | -9.50867 | 3.83016 | 6.72203 |
| CD630_34530 | CD630_Cdi2_2 | 65 – 77 | 31 – 42 | -9.50867 | 3.83016 | 6.72203 |
| CD630_09100 | CD630_Cdi2_3 | 34 – 84 | 5 – 63 | -18.9059 | 5.92260 | 3.99184 |
| CD630_24220 | CD630_Cdi2_3 | 7 – 99 | 9 – 106 | -18.5673 | 4.18037 | 3.99184 |
| CD630_08140 | CD630_Cdi2_3 | 4 – 25 | 31 – 61 | -16.6019 | 3.30159 | 3.99184 |
| CD630_36690 | CD630_Cdi2_3 | 148 – 177 | 182 – 212 | -16.2001 | 3.53761 | 3.99184 |
| CD630_19942 | CD630_Cdi2_3 | 108 – 136 | 215 – 244 | -15.6501 | 4.24574 | 3.99184 |
| CD630_23770 | CD630_Cdi2_3 | 41 – 78 | 32 – 66 | -15.0047 | 3.25181 | 3.99184 |
| CD630_16540 | CD630_Cdi2_3 | 21 – 37 | 45 – 61 | -14.6984 | 4.37658 | 3.99184 |
| CD630_08950 | CD630_Cdi2_3 | 5 – 41 | 20 – 64 | -14.6962 | 3.71905 | 3.99184 |
| CD630_25200 | CD630_Cdi2_3 | 24 – 79 | 109 – 160 | -14.67 | 3.65158 | 3.99184 |
| CD630_23760 | CD630_Cdi2_3 | 148 – 188 | 28 – 62 | -14.428 | 4.54307 | 3.99184 |
| CD630_08660 | CD630_Cdi2_3 | 158 – 199 | 29 – 66 | -14.3946 | 3.86467 | 3.99184 |
| opuCC | CD630_Cdi2_3 | 172 – 198 | 39 – 64 | -14.3917 | 4.07035 | 3.99184 |
| CD630_13560 | CD630_Cdi2_3 | 162 – 193 | 150 – 175 | -14.1534 | 3.38083 | 3.99184 |
| CD630_17170 | CD630_Cdi2_3 | 139 – 189 | 154 – 203 | -14.1368 | 4.10230 | 3.99184 |
| CD630_05830 | CD630_Cdi2_3 | 117 – 151 | 19 – 61 | -14.0577 | 3.35360 | 3.99184 |
| CD630_19680 | CD630_Cdi2_3 | 34 – 78 | 142 – 186 | -13.7427 | 3.58080 | 3.99184 |
| CD630_16530 | CD630_Cdi2_3 | 117 – 164 | 11 – 67 | -13.6731 | 4.37658 | 3.99184 |
| CD630_06100 | CD630_Cdi2_3 | 175 – 201 | 33 – 60 | -13.5504 | 4.43918 | 3.99184 |
| CD630_32990 | CD630_Cdi2_3 | 33 – 71 | 22 – 62 | -13.4621 | 5.24925 | 3.99184 |
| CD630_23450 | CD630_Cdi2_3 | 1 – 58 | 5 – 61 | -13.3834 | 3.27218 | 3.99184 |
| CD630_05480 | CD630_Cdi2_3 | 16 – 55 | 9 – 61 | -13.3714 | 4.27854 | 3.99184 |
| CD630_18610 | CD630_Cdi2_3 | 6 – 54 | 151 – 196 | -13.2061 | 4.15582 | 3.99184 |
| chrA | CD630_Cdi2_3 | 92 – 137 | 15 – 61 | -13.0715 | 5.27255 | 3.99184 |
| CD630_16160 | CD630_Cdi2_3 | 176 – 191 | 32 – 47 | -13.0322 | 6.11801 | 3.99184 |
| CD630_26920 | CD630_Cdi2_3 | 20 – 46 | 31 – 60 | -13.0086 | 4.58060 | 3.99184 |
| CD630_06140 | CD630_Cdi2_3 | 1 – 38 | 16 – 56 | -12.9705 | 4.12519 | 3.99184 |
| CD630_06920 | CD630_Cdi2_3 | 44 – 92 | 15 – 66 | -12.8748 | 3.13260 | 3.99184 |
| CD630_21760 | CD630_Cdi2_3 | 173 – 200 | 33 – 60 | -12.7183 | 4.43846 | 3.99184 |
| CD630_04550 | CD630_Cdi2_3 | 61 – 72 | 177 – 188 | -12.6907 | 3.05227 | 3.99184 |
| CD630_06190 | CD630_Cdi2_3 | 37 – 105 | 154 – 203 | -12.6753 | 3.57064 | 3.99184 |
| CD630_19430 | CD630_Cdi2_3 | 170 – 193 | 145 – 165 | -12.6138 | 3.13922 | 3.99184 |
| CD630_06642 | CD630_Cdi2_3 | 135 – 184 | 153 – 198 | -12.5126 | 3.32967 | 3.99184 |
| CD630_36180 | CD630_Cdi2_3 | 7 – 32 | 130 – 153 | -12.307 | 4.83428 | 3.99184 |
| CD630_26250 | CD630_Cdi2_3 | 138 – 188 | 12 – 62 | -12.2905 | 5.03379 | 3.99184 |
| CD630_05370 | CD630_Cdi2_3 | 1 – 11 | 12 – 22 | -12.2204 | 3.26947 | 3.99184 |
| scpB | CD630_Cdi2_3 | 122 – 157 | 13 – 56 | -12.21 | 3.33560 | 3.99184 |
| CD630_28870 | CD630_Cdi2_3 | 90 – 115 | 215 – 241 | -12.1527 | 3.41991 | 3.99184 |
| CD630_24050 | CD630_Cdi2_3 | 134 – 162 | 14 – 42 | -12.0506 | 3.70192 | 3.99184 |
| CD630_22550 | CD630_Cdi2_3 | 162 – 201 | 23 – 62 | -12.0452 | 3.62379 | 3.99184 |
| CD630_20280 | CD630_Cdi2_3 | 21 – 122 | 2 – 97 | -12.0221 | 3.46343 | 3.99184 |
| CD630_28290 | CD630_Cdi2_3 | 33 – 61 | 38 – 68 | -11.9743 | 5.54192 | 3.99184 |
| CD630_30190 | CD630_Cdi2_3 | 123 – 163 | 20 – 57 | -11.971 | 3.27394 | 3.99184 |
| CD630_07070 | CD630_Cdi2_3 | 14 – 47 | 24 – 60 | -11.9562 | 3.46414 | 3.99184 |
| CD630_21230 | CD630_Cdi2_3 | 171 – 179 | 163 – 171 | -11.8824 | 3.89213 | 3.99184 |
| CD630_14210 | CD630_Cdi2_3 | 11 – 90 | 18 – 107 | -11.8411 | 3.82355 | 3.99184 |
| rbsA | CD630_Cdi2_3 | 5 – 20 | 13 – 28 | -11.8096 | 3.71650 | 3.99184 |
| CD630_18090 | CD630_Cdi2_3 | 176 – 199 | 30 – 61 | -11.724 | 4.23961 | 3.99184 |
| serS1 | CD630_Cdi2_3 | 133 – 158 | 223 – 248 | -11.703 | 3.45858 | 3.99184 |
| CD630_32720 | CD630_Cdi2_3 | 129 – 153 | 142 – 166 | -11.6575 | 4.74276 | 3.99184 |
| CD630_18930 | CD630_Cdi2_3 | 150 – 159 | 17 – 26 | -11.6185 | 5.32127 | 3.99184 |
| CD630_03930 | CD630_Cdi2_3 | 166 – 195 | 148 – 177 | -11.5665 | 4.15934 | 3.99184 |
| CD630_09770 | CD630_Cdi2_3 | 3 – 38 | 7 – 42 | -11.5205 | 3.22106 | 3.99184 |
| oppB | CD630_Cdi2_3 | 146 – 177 | 31 – 63 | -11.4979 | 2.98328 | 3.99184 |
| CD630_14150 | CD630_Cdi2_3 | 43 – 103 | 18 – 107 | -11.4242 | 3.28251 | 3.99184 |
| bglA | CD630_Cdi2_3 | 54 – 94 | 11 – 61 | -11.3855 | 4.86402 | 3.99184 |
| ogt1 | CD630_Cdi2_3 | 126 – 163 | 128 – 166 | -11.3761 | 3.16772 | 3.99184 |
| CD630_05450 | CD630_Cdi2_3 | 159 – 171 | 163 – 175 | -11.3742 | 5.04761 | 3.99184 |
| appC | CD630_Cdi2_3 | 38 – 51 | 49 – 62 | -11.3271 | 4.50438 | 3.99184 |
| CD630_19401 | CD630_Cdi2_3 | 34 – 57 | 28 – 50 | -11.2862 | 4.37084 | 3.99184 |
| CD630_06170 | CD630_Cdi2_3 | 120 – 165 | 23 – 60 | -11.2839 | 4.95366 | 3.99184 |
| CD630_04850 | CD630_Cdi2_3 | 146 – 187 | 22 – 66 | -11.2692 | 4.40788 | 3.99184 |
| CD630_17511 | CD630_Cdi2_3 | 186 – 199 | 52 – 66 | -11.1488 | 9.12322 | 3.99184 |
| CD630_36781 | CD630_Cdi2_3 | 161 – 192 | 161 – 190 | -11.1441 | 3.75946 | 3.99184 |
| dapB2 | CD630_Cdi2_3 | 117 – 193 | 101 – 184 | -11.1412 | 3.27218 | 3.99184 |
| CD630_18400 | CD630_Cdi2_3 | 167 – 201 | 24 – 61 | -11.1217 | 4.67962 | 3.99184 |
| CD630_21240 | CD630_Cdi2_3 | 99 – 142 | 22 – 65 | -11.1202 | 5.88879 | 3.99184 |
| kdgT1 | CD630_Cdi2_3 | 167 – 197 | 163 – 195 | -11.0367 | 4.49207 | 3.99184 |
| CD630_20260 | CD630_Cdi2_3 | 73 – 88 | 235 – 250 | -11.0136 | 3.84162 | 3.99184 |
| cdtR | CD630_Cdi2_3 | 3 – 31 | 18 – 45 | -10.9973 | 5.84368 | 3.99184 |
| CD630_14190 | CD630_Cdi2_3 | 153 – 201 | 17 – 61 | -10.9921 | 3.78562 | 3.99184 |
| CD630_29650 | CD630_Cdi2_3 | 162 – 201 | 138 – 188 | -10.9213 | 3.67128 | 3.99184 |
| CD630_14620 | CD630_Cdi2_3 | 15 – 49 | 19 – 57 | -10.9045 | 4.43651 | 3.99184 |
| CD630_21070 | CD630_Cdi2_3 | 43 – 55 | 175 – 187 | -10.9045 | 5.35019 | 3.99184 |
| CD630_19850 | CD630_Cdi2_3 | 118 – 150 | 28 – 62 | -10.8615 | 6.88486 | 3.99184 |
| CD630_24830 | CD630_Cdi2_3 | 1 – 40 | 25 – 62 | -10.8569 | 4.98239 | 3.99184 |
| CD630_30100 | CD630_Cdi2_3 | 125 – 198 | 147 – 211 | -10.8288 | 4.13234 | 3.99184 |
| chrA | CD630_Cdi2_3 | 12 – 48 | 15 – 55 | -10.7907 | 5.27255 | 3.99184 |
| CD630_18910 | CD630_Cdi2_3 | 7 – 56 | 5 – 52 | -10.7862 | 3.30688 | 3.99184 |
| CD630_26150 | CD630_Cdi2_3 | 25 – 51 | 169 – 195 | -10.7846 | 4.06302 | 3.99184 |
| agrB | CD630_Cdi2_3 | 165 – 185 | 38 – 63 | -10.7126 | 3.92598 | 3.99184 |
| CD630_06160 | CD630_Cdi2_3 | 119 – 175 | 7 – 61 | -10.6854 | 5.51744 | 3.99184 |
| CD630_18920 | CD630_Cdi2_3 | 120 – 141 | 1 – 20 | -10.6596 | 4.43558 | 3.99184 |
| CD630_31530 | CD630_Cdi2_3 | 39 – 77 | 11 – 56 | -10.6595 | 3.49100 | 3.99184 |
| CD630_34560 | CD630_Cdi2_3 | 16 – 96 | 5 – 108 | -10.6331 | 4.02604 | 3.99184 |
| CD630_21540 | CD630_Cdi2_3 | 46 – 95 | 13 – 63 | -10.5684 | 4.33090 | 3.99184 |
| CD630_02720 | CD630_Cdi2_3 | 166 – 190 | 41 – 63 | -10.4698 | 3.38938 | 3.99184 |
| CD630_27270 | CD630_Cdi2_3 | 145 – 175 | 156 – 190 | -10.4675 | 3.50892 | 3.99184 |
| CD630_07770 | CD630_Cdi2_3 | 140 – 148 | 53 – 61 | -10.4336 | 3.31158 | 3.99184 |
| CD630_26020 | CD630_Cdi2_3 | 141 – 160 | 44 – 61 | -10.4174 | 4.03155 | 3.99184 |
| CD630_31970 | CD630_Cdi2_3 | 64 – 120 | 139 – 188 | -10.4142 | 3.24511 | 3.99184 |
| CD630_27060 | CD630_Cdi2_3 | 111 – 152 | 11 – 63 | -10.3671 | 4.06049 | 3.99184 |
| CD630_21080 | CD630_Cdi2_3 | 159 – 189 | 133 – 167 | -10.344 | 3.99742 | 3.99184 |
| appB | CD630_Cdi2_3 | 158 – 180 | 36 – 58 | -10.3264 | 6.44103 | 3.99184 |
| CD630_07870 | CD630_Cdi2_3 | 185 – 201 | 47 – 62 | -10.3119 | 3.22829 | 3.99184 |
| CD630_17160 | CD630_Cdi2_3 | 57 – 96 | 11 – 56 | -10.1426 | 3.50228 | 3.99184 |
| dnaC | CD630_Cdi2_3 | 51 – 64 | 4 – 18 | -10.105 | 6.30796 | 3.99184 |
| CD630_15200 | CD630_Cdi2_3 | 131 – 147 | 38 – 61 | -10.0965 | 4.00442 | 3.99184 |
| CD630_28560 | CD630_Cdi2_3 | 38 – 59 | 170 – 189 | -10.0936 | 3.96803 | 3.99184 |
| CD630_01560 | CD630_Cdi2_3 | 45 – 62 | 143 – 160 | -10.0526 | 4.06352 | 3.99184 |
| CD630_35430 | CD630_Cdi2_3 | 7 – 30 | 42 – 59 | -10.0285 | 4.28798 | 3.99184 |
| CD630_26691 | CD630_Cdi2_3 | 5 – 27 | 10 – 32 | -9.98614 | 3.96230 | 3.99184 |
| CD630_36170 | CD630_Cdi2_3 | 157 – 170 | 158 – 169 | -9.95729 | 3.72380 | 3.99184 |
| CD630_25020 | CD630_Cdi2_3 | 4 – 53 | 55 – 103 | -9.91523 | 4.17721 | 3.99184 |
| CD630_10960 | CD630_Cdi2_3 | 151 – 165 | 185 – 198 | -9.88274 | 3.53850 | 3.99184 |
| CD630_22880 | CD630_Cdi2_3 | 38 – 75 | 19 – 59 | -9.87402 | 3.62455 | 3.99184 |
| CD630_31210 | CD630_Cdi2_3 | 43 – 72 | 130 – 159 | -9.87116 | 4.17693 | 3.99184 |
| CD630_19840 | CD630_Cdi2_3 | 187 – 199 | 153 – 165 | -9.86147 | 3.44724 | 3.99184 |
| CD630_19670 | CD630_Cdi2_3 | 28 – 76 | 14 – 61 | -9.85642 | 3.20718 | 3.99184 |
| CD630_34530 | CD630_Cdi2_3 | 66 – 79 | 155 – 167 | -9.84752 | 3.83016 | 3.99184 |
| CD630_32880 | CD630_Cdi2_3 | 43 – 62 | 153 – 171 | -9.83667 | 3.41276 | 3.99184 |
| CD630_13880 | CD630_Cdi2_3 | 144 – 162 | 38 – 55 | -9.79644 | 3.52735 | 3.99184 |
| dltB | CD630_Cdi2_3 | 21 – 41 | 136 – 156 | -9.77852 | 3.81485 | 3.99184 |
| pilD | CD630_Cdi2_3 | 127 – 174 | 134 – 180 | -9.77031 | 4.54123 | 3.99184 |
| CD630_14600 | CD630_Cdi2_3 | 137 – 179 | 17 – 64 | -9.75911 | 5.90348 | 3.99184 |
| CD630_24920 | CD630_Cdi2_3 | 4 – 24 | 16 – 48 | -9.7453 | 3.89455 | 3.99184 |
| CD630_17350 | CD630_Cdi2_3 | 171 – 193 | 40 – 63 | -9.72959 | 3.35334 | 3.99184 |
| CD630_16670 | CD630_Cdi2_3 | 135 – 156 | 8 – 28 | -9.72617 | 3.33553 | 3.99184 |
| CD630_14250 | CD630_Cdi2_3 | 87 – 100 | 158 – 171 | -9.71665 | 4.19604 | 3.99184 |
| CD630_03220 | CD630_Cdi2_3 | 93 – 114 | 154 – 177 | -9.71551 | 3.36746 | 3.99184 |
| CD630_31000 | CD630_Cdi2_3 | 32 – 83 | 1 – 48 | -9.71243 | 3.70946 | 3.99184 |
| CD630_04540 | CD630_Cdi2_3 | 120 – 156 | 29 – 63 | -9.67237 | 3.33556 | 3.99184 |
| CD630_13380 | CD630_Cdi2_3 | 101 – 152 | 183 – 229 | -9.66858 | 4.74408 | 3.99184 |
| CD630_13530 | CD630_Cdi2_3 | 1 – 33 | 24 – 57 | -9.66433 | 4.09832 | 3.99184 |
| CD630_18311 | CD630_Cdi2_3 | 121 – 157 | 18 – 53 | -9.66374 | 6.86047 | 3.99184 |
| appF | CD630_Cdi2_3 | 54 – 104 | 4 – 63 | -9.66235 | 4.25570 | 3.99184 |
| CD630_29250 | CD630_Cdi2_3 | 8 – 29 | 38 – 64 | -9.64163 | 4.61901 | 3.99184 |
| CD630_09410 | CD630_Cdi2_3 | 8 – 29 | 38 – 64 | -9.64163 | 4.75651 | 3.99184 |
| CD630_25050 | CD630_Cdi2_3 | 168 – 180 | 139 – 151 | -9.63073 | 4.33624 | 3.99184 |
| spaK | CD630_Cdi2_3 | 17 – 46 | 141 – 168 | -9.60886 | 3.04530 | 3.99184 |
| CD630_19000 | CD630_Cdi2_3 | 56 – 98 | 8 – 53 | -9.59594 | 4.18443 | 3.99184 |
| CD630_14140 | CD630_Cdi2_3 | 116 – 163 | 17 – 64 | -9.5695 | 3.16551 | 3.99184 |
| CD630_27200 | CD630_Cdi2_3 | 8 – 20 | 159 – 170 | -9.56858 | 3.42516 | 3.99184 |
| CD630_22660 | CD630_Cdi2_3 | 25 – 63 | 12 – 60 | -9.52727 | 3.60496 | 3.99184 |
| CD630_06060 | CD630_Cdi2_3 | 168 – 191 | 32 – 59 | -9.51172 | 4.94050 | 3.99184 |
| CD630_08510 | CD630_Cdi2_3 | 47 – 56 | 179 – 188 | -9.51122 | 5.16329 | 3.99184 |
| CD630_18090 | CD630_Cdi2_3 | 149 – 170 | 41 – 61 | -9.50161 | 4.23961 | 3.99184 |
| bcp | CD630_n00080 | 111 – 162 | 63 – 101 | -19.67 | 3.16670 | 4.59330 |
| etfA | CD630_n00080 | 156 – 180 | 75 – 99 | -19.4739 | 3.41204 | 4.59330 |
| CD630_17160 | CD630_n00080 | 61 – 104 | 64 – 102 | -19.1611 | 3.50228 | 4.59330 |
| CD630_03220 | CD630_n00080 | 94 – 132 | 75 – 103 | -19.0808 | 3.36746 | 4.59330 |
| CD630_24590 | CD630_n00080 | 83 – 158 | 10 – 82 | -18.9306 | 3.70401 | 4.59330 |
| CD630_10050 | CD630_n00080 | 90 – 172 | 1 – 78 | -18.9118 | 3.93750 | 4.59330 |
| CD630_32710 | CD630_n00080 | 140 – 159 | 72 – 91 | -18.8583 | 4.74276 | 4.59330 |
| srlD | CD630_n00080 | 110 – 146 | 58 – 97 | -18.7863 | 4.19843 | 4.59330 |
| argS | CD630_n00080 | 163 – 188 | 76 – 100 | -18.6988 | 4.96036 | 4.59330 |
| CD630_21020 | CD630_n00080 | 86 – 104 | 64 – 82 | -18.611 | 4.28666 | 4.59330 |
| cobQ | CD630_n00080 | 87 – 153 | 6 – 80 | -18.5354 | 3.13524 | 4.59330 |
| CD630_14130 | CD630_n00080 | 142 – 195 | 64 – 100 | -18.519 | 4.57970 | 4.59330 |
| CD630_15360 | CD630_n00080 | 154 – 173 | 79 – 98 | -18.4929 | 4.47208 | 4.59330 |
| CD630_06060 | CD630_n00080 | 15 – 44 | 72 – 98 | -18.4652 | 4.94050 | 4.59330 |
| rbsC | CD630_n00080 | 31 – 61 | 61 – 92 | -18.4415 | 3.71650 | 4.59330 |
| CD630_29840 | CD630_n00080 | 31 – 70 | 69 – 100 | -18.3341 | 4.48825 | 4.59330 |
| CD630_28320 | CD630_n00080 | 81 – 94 | 86 – 100 | -18.2457 | 3.99184 | 4.59330 |
| CD630_23760 | CD630_n00080 | 55 – 126 | 42 – 100 | -18.1592 | 4.54307 | 4.59330 |
| CD630_22720 | CD630_n00080 | 89 – 157 | 50 – 92 | -18.1087 | 3.81804 | 4.59330 |
| CD630_33682 | CD630_n00080 | 82 – 147 | 27 – 93 | -18.077 | 5.77720 | 4.59330 |
| CD630_07480 | CD630_n00080 | 80 – 110 | 64 – 99 | -18.0426 | 5.18321 | 4.59330 |
| dapB2 | CD630_n00080 | 139 – 198 | 57 – 99 | -18.034 | 3.27218 | 4.59330 |
| eutV | CD630_n00080 | 87 – 172 | 8 – 95 | -18.0178 | 4.55715 | 4.59330 |
| CD630_05490 | CD630_n00080 | 162 – 188 | 65 – 102 | -17.9937 | 3.89852 | 4.59330 |
| cspD | CD630_n00080 | 79 – 173 | 10 – 101 | -17.5878 | 3.23587 | 4.59330 |
| CD630_08100 | CD630_n00080 | 127 – 200 | 4 – 78 | -17.5738 | 4.39181 | 4.59330 |
| CD630_20670 | CD630_n00080 | 81 – 158 | 1 – 87 | -17.5724 | 4.42757 | 4.59330 |
| CD630_19904 | CD630_n00080 | 139 – 187 | 48 – 92 | -17.5181 | 5.96526 | 4.59330 |
| nusB | CD630_n00080 | 115 – 142 | 77 – 101 | -17.4139 | 4.92798 | 4.59330 |
| dnaC | CD630_n00080 | 140 – 161 | 65 – 86 | -17.3735 | 6.30796 | 4.59330 |
| CD630_05590 | CD630_n00080 | 94 – 143 | 49 – 100 | -17.3288 | 3.39670 | 4.59330 |
| CD630_05230 | CD630_n00080 | 2 – 54 | 2 – 51 | -17.3139 | 3.73787 | 4.59330 |
| CD630_23820 | CD630_n00080 | 86 – 121 | 59 – 103 | -17.1993 | 3.18419 | 4.59330 |
| CD630_04341 | CD630_n00080 | 123 – 143 | 50 – 72 | -16.799 | 4.29262 | 4.59330 |
| CD630_28260 | CD630_n00080 | 38 – 111 | 4 – 78 | -16.7602 | 3.51613 | 4.59330 |
| mobA | CD630_n00080 | 172 – 199 | 72 – 100 | -16.6838 | 4.25703 | 4.59330 |
| CD630_14050 | CD630_n00080 | 13 – 42 | 64 – 95 | -16.6658 | 3.30894 | 4.59330 |
| CD630_32070 | CD630_n00080 | 51 – 95 | 50 – 92 | -16.6496 | 3.51964 | 4.59330 |
| etfA | CD630_n00080 | 24 – 90 | 49 – 100 | -16.5714 | 4.80521 | 4.59330 |
| pmi | CD630_n00080 | 143 – 165 | 78 – 101 | -16.5339 | 3.86845 | 4.59330 |
| CD630_25610 | CD630_n00080 | 29 – 134 | 1 – 98 | -16.5121 | 3.02685 | 4.59330 |
| bglA | CD630_n00080 | 131 – 183 | 49 – 100 | -16.4387 | 4.86402 | 4.59330 |
| CD630_21720 | CD630_n00080 | 136 – 182 | 65 – 100 | -16.3669 | 4.13030 | 4.59330 |
| CD630_10970 | CD630_n00080 | 36 – 111 | 20 – 100 | -16.3218 | 3.53850 | 4.59330 |
| CD630_15900 | CD630_n00080 | 142 – 170 | 79 – 103 | -16.2381 | 3.17664 | 4.59330 |
| CD630_32690 | CD630_n00080 | 103 – 142 | 64 – 99 | -16.0708 | 7.32676 | 4.59330 |
| CD630_21620 | CD630_n00080 | 70 – 161 | 1 – 96 | -15.9917 | 3.02649 | 4.59330 |
| CD630_24760 | CD630_n00080 | 79 – 160 | 54 – 102 | -15.973 | 5.19485 | 4.59330 |
| CD630_30140 | CD630_n00080 | 160 – 198 | 69 – 98 | -15.9334 | 3.28343 | 4.59330 |
| CD630_11730 | CD630_n00080 | 141 – 179 | 65 – 93 | -15.8427 | 4.80521 | 4.59330 |
| CD630_02740 | CD630_n00080 | 140 – 167 | 67 – 95 | -15.6937 | 5.02277 | 4.59330 |
| CD630_35260 | CD630_n00080 | 67 – 103 | 65 – 96 | -15.6181 | 5.77277 | 4.59330 |
| CD630_28550 | CD630_n00080 | 170 – 201 | 57 – 103 | -15.6127 | 3.51196 | 4.59330 |
| CD630_17292 | CD630_n00080 | 80 – 118 | 68 – 101 | -15.5476 | 6.25799 | 4.59330 |
| CD630_06080 | CD630_n00080 | 151 – 197 | 61 – 100 | -15.43 | 3.82315 | 4.59330 |
| CD630_18910 | CD630_n00080 | 162 – 200 | 43 – 96 | -15.3048 | 3.30688 | 4.59330 |

| | | | | | | |
|---|---|---|---|---|---|---|
| CD630_13920 | CD630_n00080 | 86 -- 123 | 49 -- 92 | -15.2836 | 3.25236 | 4.59330 |
| CD630_11241 | CD630_n00080 | 159 -- 198 | 55 -- 96 | -15.2419 | 4.85723 | 4.59330 |
| CD630_09840 | CD630_n00080 | 78 -- 116 | 42 -- 87 | -15.1263 | 4.50009 | 4.59330 |
| CD630_23280 | CD630_n00080 | 131 -- 150 | 75 -- 95 | -15.0298 | 3.50093 | 4.59330 |
| CD630_23090 | CD630_n00080 | 81 -- 126 | 65 -- 103 | -14.8878 | 4.25622 | 4.59330 |
| CD630_36180 | CD630_n00080 | 164 -- 185 | 73 -- 94 | -14.865 | 4.83428 | 4.59330 |
| CD630_23440 | CD630_n00080 | 181 -- 198 | 84 -- 101 | -14.7868 | 3.34841 | 4.59330 |
| CD630_17960 | CD630_n00080 | 114 -- 149 | 62 -- 96 | -14.7779 | 3.71404 | 4.59330 |
| glgP | CD630_n00080 | 71 -- 126 | 59 -- 100 | -14.7167 | 3.28530 | 4.59330 |
| CD630_02420 | CD630_n00080 | 1 -- 31 | 60 -- 83 | -14.707 | 3.71048 | 4.59330 |
| CD630_22850 | CD630_n00080 | 160 -- 190 | 77 -- 100 | -14.7049 | 3.84102 | 4.59330 |
| CD630_20980 | CD630_n00080 | 125 -- 150 | 75 -- 100 | -14.6934 | 8.12847 | 4.59330 |
| CD630_36781 | CD630_n00080 | 177 -- 201 | 70 -- 96 | -14.6656 | 3.75946 | 4.59330 |
| CD630_19801 | CD630_n00080 | 119 -- 167 | 50 -- 97 | -14.6359 | 5.22035 | 4.59330 |
| iorA | CD630_n00080 | 81 -- 155 | 31 -- 97 | -14.6165 | 3.83173 | 4.59330 |
| CD630_06110 | CD630_n00080 | 168 -- 184 | 60 -- 78 | -14.5043 | 5.61380 | 4.59330 |
| CD630_32520 | CD630_n00080 | 88 -- 140 | 50 -- 92 | -14.4832 | 4.06098 | 4.59330 |
| moaA | CD630_n00080 | 86 -- 120 | 67 -- 100 | -14.4747 | 3.83799 | 4.59330 |
| CD630_20541 | CD630_n00080 | 43 -- 56 | 86 -- 102 | -14.4238 | 5.74527 | 4.59330 |
| folC | CD630_n00080 | 85 -- 113 | 50 -- 93 | -14.3808 | 3.85634 | 4.59330 |
| bglA | CD630_n00080 | 92 -- 133 | 64 -- 95 | -14.3432 | 4.50315 | 4.59330 |
| CD630_13600 | CD630_n00080 | 119 -- 145 | 65 -- 92 | -14.3402 | 4.45481 | 4.59330 |
| CD630_18680 | CD630_n00080 | 62 -- 104 | 64 -- 101 | -14.3285 | 3.04315 | 4.59330 |
| dltA | CD630_n00080 | 138 -- 185 | 50 -- 96 | -14.3036 | 3.85429 | 4.59330 |
| CD630_09040 | CD630_n00080 | 133 -- 180 | 64 -- 100 | -14.29 | 4.11851 | 4.59330 |
| xdhA | CD630_n00080 | 46 -- 144 | 3 -- 100 | -14.1914 | 4.45147 | 4.59330 |
| CD630_36690 | CD630_n00080 | 155 -- 188 | 65 -- 97 | -14.188 | 3.53761 | 4.59330 |
| CD630_25620 | CD630_n00080 | 158 -- 195 | 60 -- 97 | -14.1359 | 3.02685 | 4.59330 |
| CD630_33600 | CD630_n00080 | 147 -- 194 | 55 -- 100 | -14.1114 | 3.73196 | 4.59330 |
| cdu2 | CD630_n00080 | 139 -- 190 | 65 -- 98 | -14.0589 | 2.97335 | 4.59330 |
| pgm | CD630_n00080 | 82 -- 193 | 12 -- 96 | -13.9864 | 3.45439 | 4.59330 |
| CD630_32880 | CD630_n00080 | 24 -- 104 | 9 -- 95 | -13.9552 | 3.41276 | 4.59330 |
| CD630_19780 | CD630_n00080 | 4 -- 27 | 71 -- 91 | -13.8886 | 3.64830 | 4.59330 |
| CD630_06970 | CD630_n00080 | 122 -- 152 | 59 -- 99 | -13.8614 | 3.82312 | 4.59330 |
| CD630_15150 | CD630_n00080 | 72 -- 93 | 75 -- 100 | -13.8487 | 3.35564 | 4.59330 |
| CD630_16460 | CD630_n00080 | 158 -- 182 | 77 -- 100 | -13.8185 | 4.64028 | 4.59330 |
| CD630_16450 | CD630_n00080 | 159 -- 184 | 68 -- 99 | -13.8127 | 4.57547 | 4.59330 |
| CD630_05870 | CD630_n00080 | 161 -- 193 | 49 -- 88 | -13.8121 | 4.28339 | 4.59330 |
| CD630_25310 | CD630_n00080 | 14 -- 45 | 64 -- 95 | -13.7279 | 3.99861 | 4.59330 |
| garK | CD630_n00080 | 150 -- 191 | 46 -- 93 | -13.6619 | 3.72836 | 4.59330 |
| CD630_31010 | CD630_n00080 | 63 -- 107 | 49 -- 96 | -13.6595 | 4.96305 | 4.59330 |
| CD630_13560 | CD630_n00080 | 167 -- 190 | 68 -- 90 | -13.6579 | 3.38083 | 4.59330 |
| CD630_06690 | CD630_n00080 | 84 -- 128 | 33 -- 82 | -13.6328 | 5.82709 | 4.59330 |
| CD630_17970 | CD630_n00080 | 77 -- 143 | 20 -- 78 | -13.5728 | 3.68802 | 4.59330 |
| CD630_10960 | CD630_n00080 | 26 -- 58 | 73 -- 101 | -13.5452 | 3.53850 | 4.59330 |
| CD630_19680 | CD630_n00080 | 89 -- 111 | 57 -- 82 | -13.4395 | 3.58080 | 4.59330 |
| CD630_16160 | CD630_n00080 | 30 -- 53 | 57 -- 78 | -13.4393 | 6.11801 | 4.59330 |
| CD630_19921 | CD630_n00080 | 149 -- 172 | 73 -- 97 | -13.4211 | 5.46345 | 4.59330 |
| terD | CD630_n00080 | 88 -- 161 | 4 -- 80 | -13.4155 | 3.33148 | 4.59330 |
| uxaA | CD630_n00080 | 68 -- 123 | 50 -- 98 | -13.3695 | 4.26165 | 4.59330 |
| CD630_14360 | CD630_n00080 | 22 -- 126 | 15 -- 103 | -13.3378 | 3.02079 | 4.59330 |
| CD630_09990 | CD630_n00080 | 132 -- 174 | 19 -- 62 | -13.324 | 3.86040 | 4.59330 |
| CD630_18930 | CD630_n00080 | 191 -- 201 | 69 -- 79 | -13.3226 | 5.32127 | 4.59330 |
| CD630_01590 | CD630_n00080 | 169 -- 201 | 67 -- 103 | -13.3002 | 3.45609 | 4.59330 |
| CD630_11220 | CD630_n00080 | 137 -- 159 | 69 -- 90 | -13.2823 | 3.52327 | 4.59330 |
| CD630_29270 | CD630_n00080 | 4 -- 94 | 1 -- 95 | -13.2554 | 3.70174 | 4.59330 |
| CD630_32700 | CD630_n00080 | 156 -- 185 | 73 -- 101 | -13.2401 | 4.74276 | 4.59330 |
| CD630_33570 | CD630_n00080 | 142 -- 175 | 64 -- 98 | -13.2394 | 3.95778 | 4.59330 |
| CD630_03160 | CD630_n00080 | 70 -- 95 | 57 -- 88 | -13.2328 | 3.66946 | 4.59330 |
| acoL | CD630_n00080 | 105 -- 201 | 2 -- 96 | -13.1701 | 3.32169 | 4.59330 |
| CD630_30540 | CD630_n00080 | 152 -- 176 | 66 -- 94 | -13.1559 | 3.63804 | 4.59330 |
| CD630_26150 | CD630_n00080 | 84 -- 105 | 51 -- 73 | -13.1419 | 4.06302 | 4.59330 |
| CD630_21490 | CD630_n00080 | 33 -- 142 | 1 -- 103 | -13.0949 | 3.37462 | 4.59330 |
| CD630_06350 | CD630_n00080 | 86 -- 115 | 68 -- 95 | -13.0869 | 4.35983 | 4.59330 |
| CD630_27520 | CD630_n00080 | 131 -- 161 | 60 -- 94 | -13.0592 | 4.27693 | 4.59330 |
| CD630_09030 | CD630_n00080 | 127 -- 159 | 49 -- 82 | -13.0312 | 4.21282 | 4.59330 |
| CD630_07770 | CD630_n00080 | 86 -- 122 | 55 -- 96 | -13.0042 | 3.31158 | 4.59330 |
| CD630_06060 | CD630_n00080 | 67 -- 91 | 61 -- 92 | -12.9116 | 4.94050 | 4.59330 |
| CD630_16530 | CD630_n00080 | 155 -- 173 | 50 -- 67 | -12.8746 | 4.37658 | 4.59330 |
| CD630_31220 | CD630_n00080 | 62 -- 104 | 64 -- 97 | -12.8278 | 3.73779 | 4.59330 |
| rolB | CD630_n00080 | 111 -- 157 | 50 -- 96 | -12.8094 | 3.60270 | 4.59330 |
| serS1 | CD630_n00080 | 122 -- 173 | 50 -- 92 | -12.7963 | 3.45858 | 4.59330 |
| vncR | CD630_n00080 | 137 -- 152 | 65 -- 82 | -12.766 | 3.93642 | 4.59330 |
| CD630_18940 | CD630_n00080 | 141 -- 172 | 59 -- 96 | -12.6412 | 4.09988 | 4.59330 |
| CD630_10220 | CD630_n00080 | 84 -- 125 | 66 -- 103 | -12.634 | 4.08233 | 4.59330 |
| glgB | CD630_n00080 | 86 -- 110 | 58 -- 83 | -12.6168 | 4.48273 | 4.59330 |
| CD630_29750 | CD630_n00080 | 25 -- 41 | 51 -- 65 | -12.5973 | 4.59146 | 4.59330 |
| cprK | CD630_n00080 | 50 -- 149 | 1 -- 66 | -12.5773 | 3.96216 | 4.59330 |
| CD630_14920 | CD630_n00080 | 88 -- 114 | 51 -- 75 | -12.5218 | 4.35225 | 4.59330 |
| CD630_08460 | CD630_n00080 | 15 -- 37 | 1 -- 24 | -12.4862 | 4.15083 | 4.59330 |
| CD630_27310 | CD630_n00080 | 59 -- 103 | 65 -- 103 | -12.4273 | 4.01622 | 4.59330 |
| CD630_35840 | CD630_n00080 | 63 -- 133 | 20 -- 99 | -12.4028 | 3.14170 | 4.59330 |
| spoVFB | CD630_n00080 | 171 -- 200 | 73 -- 101 | -12.3927 | 4.62758 | 4.59330 |
| CD630_15080 | CD630_n00080 | 74 -- 103 | 65 -- 94 | -12.3846 | 4.03033 | 4.59330 |
| CD630_21700 | CD630_n00080 | 179 -- 195 | 65 -- 79 | -12.3442 | 3.96650 | 4.59330 |
| feoB | CD630_n00080 | 11 -- 46 | 73 -- 103 | -12.3238 | 4.05364 | 4.59330 |
| CD630_30420 | CD630_n00080 | 167 -- 200 | 61 -- 94 | -12.31 | 5.27539 | 4.59330 |
| kdpD | CD630_n00080 | 69 -- 91 | 1 -- 20 | -12.3038 | 4.27140 | 4.59330 |
| CD630_27270 | CD630_n00080 | 146 -- 184 | 69 -- 102 | -12.2763 | 3.50892 | 4.59330 |
| CD630_30120 | CD630_n00080 | 133 -- 167 | 65 -- 102 | -12.2745 | 4.04859 | 4.59330 |
| CD630_32890 | CD630_n00080 | 152 -- 189 | 72 -- 103 | -12.2742 | 3.41276 | 4.59330 |
| CD630_02430 | CD630_n00080 | 26 -- 119 | 20 -- 100 | -12.2574 | 3.71048 | 4.59330 |
| ogt1 | CD630_n00080 | 38 -- 72 | 58 -- 89 | -12.2285 | 3.16772 | 4.59330 |
| CD630_22880 | CD630_n00080 | 103 -- 117 | 77 -- 90 | -12.2252 | 3.62455 | 4.59330 |
| CD630_23770 | CD630_n00080 | 77 -- 112 | 64 -- 96 | -12.2113 | 3.25181 | 4.59330 |
| CD630_21790 | CD630_n00080 | 136 -- 169 | 51 -- 78 | -12.1936 | 4.93762 | 4.59330 |
| CD630_21760 | CD630_n00080 | 22 -- 41 | 64 -- 79 | -12.1815 | 4.43846 | 4.59330 |
| CD630_27770 | CD630_n00080 | 35 -- 55 | 51 -- 78 | -12.1458 | 4.08588 | 4.59330 |
| CD630_22890 | CD630_n00080 | 34 -- 46 | 65 -- 78 | -12.1429 | 3.75591 | 4.59330 |
| CD630_18920 | CD630_n00080 | 170 -- 197 | 71 -- 99 | -12.1352 | 4.43558 | 4.59330 |
| CD630_20171 | CD630_n00080 | 88 -- 147 | 2 -- 71 | -12.0334 | 6.87491 | 4.59330 |
| CD630_10880 | CD630_n00080 | 132 -- 165 | 62 -- 98 | -12.0156 | 5.05048 | 4.59330 |
| CD630_10000 | CD630_n00080 | 139 -- 165 | 66 -- 92 | -12.0124 | 3.86040 | 4.59330 |
| CD630_08360 | CD630_n00080 | 26 -- 61 | 50 -- 95 | -12.0039 | 3.19339 | 4.59330 |
| CD630_19790 | CD630_n00080 | 22 -- 122 | 1 -- 103 | -11.9901 | 3.79593 | 4.59330 |
| CD630_15890 | CD630_n00080 | 30 -- 45 | 82 -- 97 | -11.9823 | 3.39962 | 4.59330 |
| fliI | CD630_n00080 | 42 -- 99 | 50 -- 96 | -11.9741 | 3.03666 | 4.59330 |
| CD630_22070 | CD630_n00080 | 25 -- 129 | 1 -- 97 | -11.9507 | 4.09651 | 4.59330 |
| CD630_31970 | CD630_n00080 | 119 -- 132 | 90 -- 103 | -11.9498 | 3.24511 | 4.59330 |
| CD630_14700 | CD630_n00080 | 58 -- 88 | 68 -- 100 | -11.9031 | 3.98234 | 4.59330 |
| CD630_18440 | CD630_n00080 | 163 -- 200 | 61 -- 100 | -11.887 | 4.07960 | 4.59330 |
| CD630_05300 | CD630_n00080 | 87 -- 108 | 66 -- 84 | -11.8827 | 4.56885 | 4.59330 |
| tcdA | CD630_n00080 | 36 -- 54 | 62 -- 80 | -11.8624 | 4.70593 | 4.59330 |
| CD630_19700 | CD630_n00080 | 11 -- 31 | 68 -- 97 | -11.862 | 3.85861 | 4.59330 |
| CD630_23080 | CD630_n00080 | 165 -- 194 | 73 -- 96 | -11.8259 | 5.23027 | 4.59330 |
| CD630_26750 | CD630_n00080 | 81 -- 105 | 69 -- 103 | -11.8148 | 3.60610 | 4.59330 |
| CD630_05880 | CD630_n00080 | 90 -- 151 | 12 -- 77 | -11.8096 | 4.28339 | 4.59330 |
| CD630_05950 | CD630_n00080 | 179 -- 199 | 71 -- 97 | -11.7645 | 3.31669 | 4.59330 |
| opuCC | CD630_n00080 | 87 -- 105 | 56 -- 77 | -11.7402 | 4.07035 | 4.59330 |
| CD630_15550 | CD630_n00080 | 88 -- 115 | 51 -- 81 | -11.732 | 3.49019 | 4.59330 |
| CD630_18781 | CD630_n00080 | 174 -- 200 | 64 -- 93 | -11.6894 | 3.77660 | 4.59330 |
| thiH | CD630_n00080 | 174 -- 200 | 64 -- 90 | -11.6709 | 3.58230 | 4.59330 |
| CD630_14810 | CD630_n00080 | 138 -- 154 | 63 -- 96 | -11.6227 | 3.35323 | 4.59330 |
| CD630_17770 | CD630_n00080 | 162 -- 199 | 62 -- 95 | -11.5774 | 5.06490 | 4.59330 |
| CD630_19800 | CD630_n00080 | 61 -- 90 | 66 -- 97 | -11.5736 | 5.22035 | 4.59330 |
| CD630_27060 | CD630_n00080 | 86 -- 155 | 44 -- 97 | -11.5688 | 4.06049 | 4.59330 |
| CD630_14630 | CD630_n00080 | 122 -- 196 | 2 -- 95 | -11.5668 | 3.34485 | 4.59330 |
| CD630_06920 | CD630_n00080 | 121 -- 136 | 86 -- 101 | -11.5316 | 3.13260 | 4.59330 |
| sigK | CD630_n00080 | 119 -- 129 | 87 -- 98 | -11.4819 | 3.08122 | 4.59330 |
| CD630_14340 | CD630_n00080 | 70 -- 108 | 60 -- 100 | -11.4649 | 5.10868 | 4.59330 |
| CD630_28330 | CD630_n00080 | 179 -- 195 | 68 -- 91 | -11.4497 | 3.29122 | 4.59330 |
| CD630_21530 | CD630_n00080 | 108 -- 190 | 33 -- 100 | -11.4272 | 3.39774 | 4.59330 |
| CD630_08510 | CD630_n00080 | 44 -- 90 | 49 -- 91 | -11.4077 | 5.16329 | 4.59330 |
| CD630_30130 | CD630_n00080 | 6 -- 58 | 41 -- 100 | -11.4066 | 3.28343 | 4.59330 |
| CD630_27230 | CD630_n00080 | 193 -- 200 | 93 -- 100 | -11.3625 | 3.52462 | 4.59330 |
| CD630_08940 | CD630_n00080 | 152 -- 179 | 73 -- 97 | -11.3561 | 3.72225 | 4.59330 |
| opuCA | CD630_n00080 | 86 -- 119 | 71 -- 100 | -11.3476 | 4.07035 | 4.59330 |
| CD630_08390 | CD630_n00080 | 46 -- 100 | 49 -- 101 | -11.3229 | 5.16000 | 4.59330 |
| CD630_27570 | CD630_n00080 | 122 -- 164 | 50 -- 95 | -11.3166 | 3.46216 | 4.59330 |
| CD630_26050 | CD630_n00080 | 82 -- 124 | 42 -- 80 | -11.3126 | 6.99851 | 4.59330 |
| CD630_19942 | CD630_n00080 | 163 -- 199 | 65 -- 100 | -11.3041 | 4.24574 | 4.59330 |
| CD630_19903 | CD630_n00080 | 80 -- 147 | 27 -- 103 | -11.2821 | 5.79994 | 4.59330 |
| CD630_29650 | CD630_n00080 | 86 -- 115 | 72 -- 101 | -11.2758 | 3.67128 | 4.59330 |
| CD630_13780 | CD630_n00080 | 166 -- 185 | 58 -- 80 | -11.2629 | 3.52274 | 4.59330 |
| glnA | CD630_n00080 | 88 -- 104 | 64 -- 79 | -11.262 | 3.87629 | 4.59330 |
| CD630_08950 | CD630_n00080 | 40 -- 74 | 13 -- 45 | -11.2233 | 3.71905 | 4.59330 |
| CD630_31461 | CD630_n00080 | 54 -- 85 | 63 -- 98 | -11.1574 | 4.41476 | 4.59330 |
| CD630_19580 | CD630_n00080 | 174 -- 197 | 1 -- 24 | -11.152 | 4.51902 | 4.59330 |
| CD630_05820 | CD630_n00080 | 146 -- 165 | 49 -- 70 | -11.1309 | 3.75661 | 4.59330 |
| pilC | CD630_n00080 | 13 -- 62 | 71 -- 100 | -11.0931 | 3.54230 | 4.59330 |
| CD630_27220 | CD630_n00080 | 154 -- 197 | 50 -- 101 | -11.092 | 4.18786 | 4.59330 |
| chrA | CD630_n00080 | 141 -- 170 | 53 -- 93 | -11.0651 | 5.27255 | 4.59330 |
| malY | CD630_n00080 | 55 -- 94 | 49 -- 82 | -11.052 | 4.46345 | 4.59330 |
| CD630_26920 | CD630_n00080 | 137 -- 174 | 69 -- 103 | -11.0482 | 4.58060 | 4.59330 |
| CD630_32060 | CD630_n00080 | 37 -- 58 | 52 -- 75 | -11.0237 | 5.73797 | 4.59330 |
| oppB | CD630_n00080 | 87 -- 110 | 49 -- 80 | -11.0215 | 2.98328 | 4.59330 |
| CD630_22710 | CD630_n00080 | 171 -- 199 | 69 -- 97 | -11.0023 | 4.05623 | 4.59330 |
| pilT | CD630_n00080 | 78 -- 91 | 67 -- 80 | -10.9853 | 3.24639 | 4.59330 |
| CD630_33420 | CD630_n00080 | 11 -- 35 | 2 -- 23 | -10.9782 | 3.22095 | 4.59330 |
| CD630_30620 | CD630_n00080 | 12 -- 29 | 1 -- 21 | -10.9777 | 4.15443 | 4.59330 |
| oppC | CD630_n00080 | 152 -- 184 | 73 -- 97 | -10.9513 | 3.01695 | 4.59330 |
| gcvTPA | CD630_n00080 | 84 -- 115 | 67 -- 103 | -10.9444 | 4.77229 | 4.59330 |
| CD630_17710 | CD630_n00080 | 87 -- 113 | 50 -- 81 | -10.9365 | 4.37688 | 4.59330 |
| CD630_20660 | CD630_n00080 | 84 -- 101 | 42 -- 66 | -10.9301 | 5.39321 | 4.59330 |
| dltB | CD630_n00080 | 162 -- 199 | 50 -- 96 | -10.915 | 3.81485 | 4.59330 |
| pilD | CD630_n00080 | 45 -- 95 | 51 -- 96 | -10.9109 | 4.54123 | 4.59330 |
| cbpA | CD630_n00080 | 54 -- 118 | 12 -- 71 | -10.8421 | 3.03040 | 4.59330 |
| cheW | CD630_n00080 | 82 -- 106 | 78 -- 103 | -10.838 | 3.72371 | 4.59330 |
| CD630_18410 | CD630_n00080 | 66 -- 95 | 58 -- 101 | -10.7996 | 3.59739 | 4.59330 |
| CD630_22740 | CD630_n00080 | 79 -- 114 | 64 -- 98 | -10.7769 | 3.08733 | 4.59330 |
| CD630_31580 | CD630_n00080 | 87 -- 104 | 64 -- 81 | -10.7698 | 3.51756 | 4.59330 |
| CD630_03040 | CD630_n00080 | 117 -- 134 | 80 -- 96 | -10.7682 | 3.51403 | 4.59330 |
| CD630_18660 | CD630_n00080 | 131 -- 161 | 49 -- 77 | -10.7594 | 3.50371 | 4.59330 |
| CD630_06560 | CD630_n00080 | 45 -- 67 | 65 -- 88 | -10.7507 | 4.02532 | 4.59330 |
| CD630_16670 | CD630_n00080 | 108 -- 157 | 50 -- 95 | -10.7344 | 3.33553 | 4.59330 |
| CD630_12650 | CD630_n00080 | 84 -- 167 | 13 -- 92 | -10.7279 | 6.31827 | 4.59330 |
| CD630_30100 | CD630_n00080 | 54 -- 66 | 65 -- 77 | -10.7273 | 4.13234 | 4.59330 |
| CD630_23000 | CD630_n00080 | 154 -- 168 | 12 -- 26 | -10.7268 | 3.54130 | 4.59330 |
| CD630_24920 | CD630_n00080 | 143 -- 154 | 71 -- 82 | -10.7042 | 3.89455 | 4.59330 |
| CD630_25190 | CD630_n00080 | 155 -- 171 | 77 -- 94 | -10.6834 | 3.19663 | 4.59330 |
| CD630_19401 | CD630_n00080 | 22 -- 66 | 31 -- 73 | -10.6805 | 4.37084 | 4.59330 |
| CD630_22030 | CD630_n00080 | 82 -- 149 | 7 -- 53 | -10.6759 | 3.29934 | 4.59330 |
| CD630_04410 | CD630_n00080 | 87 -- 166 | 43 -- 102 | -10.6115 | 3.90451 | 4.59330 |
| CD630_22660 | CD630_n00080 | 182 -- 194 | 7 -- 19 | -10.5959 | 3.60496 | 4.59330 |
| CD630_18610 | CD630_n00080 | 87 -- 134 | 34 -- 82 | -10.5645 | 4.15582 | 4.59330 |
| CD630_03420 | CD630_n00080 | 84 -- 148 | 42 -- 97 | -10.4973 | 4.71868 | 4.59330 |
| CD630_32110 | CD630_n00080 | 1 -- 22 | 9 -- 33 | -10.4327 | 4.45291 | 4.59330 |
| CD630_15431 | CD630_n00080 | 105 -- 122 | 65 -- 81 | -10.4027 | 5.29780 | 4.59330 |
| spaK | CD630_n00080 | 79 -- 103 | 65 -- 97 | -10.3781 | 3.04530 | 4.59330 |
| CD630_16150 | CD630_n00080 | 82 -- 109 | 72 -- 96 | -10.3574 | 4.20614 | 4.59330 |
| CD630_08090 | CD630_n00080 | 162 -- 198 | 64 -- 96 | -10.3527 | 3.57167 | 4.59330 |
| CD630_05900 | CD630_n00080 | 32 -- 124 | 12 -- 98 | -10.3434 | 3.21934 | 4.59330 |
| pilW | CD630_n00080 | 88 -- 117 | 49 -- 82 | -10.3238 | 4.96316 | 4.59330 |
| CD630_13960 | CD630_n00080 | 126 -- 158 | 65 -- 93 | -10.3024 | 3.30375 | 4.59330 |
| ribD | CD630_n00080 | 166 -- 177 | 64 -- 75 | -10.297 | 6.65922 | 4.59330 |
| CD630_13380 | CD630_n00080 | 137 -- 149 | 80 -- 98 | -10.2926 | 4.74408 | 4.59330 |
| CD630_24220 | CD630_n00080 | 125 -- 140 | 59 -- 74 | -10.2413 | 4.18037 | 4.59330 |
| CD630_14410 | CD630_n00080 | 90 -- 117 | 51 -- 80 | -10.2362 | 3.39433 | 4.59330 |
| CD630_36170 | CD630_n00080 | 147 -- 192 | 41 -- 80 | -10.2247 | 3.72380 | 4.59330 |
| aroD | CD630_n00080 | 90 -- 106 | 67 -- 83 | -10.1581 | 3.71929 | 4.59330 |
| CD630_36190 | CD630_n00080 | 78 -- 103 | 65 -- 90 | -10.1538 | 4.63449 | 4.59330 |
| CD630_15190 | CD630_n00080 | 169 -- 196 | 17 -- 38 | -10.1183 | 4.43255 | 4.59330 |
| cwp17 | CD630_n00080 | 90 -- 112 | 72 -- 95 | -10.1077 | 3.43363 | 4.59330 |
| argG | CD630_n00080 | 105 -- 136 | 74 -- 100 | -10.0848 | 4.80300 | 4.59330 |
| CD630_26691 | CD630_n00080 | 89 -- 172 | 16 -- 95 | -10.0836 | 3.96230 | 4.59330 |
| CD630_31382 | CD630_n00080 | 95 -- 121 | 54 -- 78 | -10.0788 | 3.49137 | 4.59330 |
| CD630_13930 | CD630_n00080 | 91 -- 108 | 76 -- 92 | -10.0768 | 4.09246 | 4.59330 |
| CD630_25070 | CD630_n00080 | 178 -- 188 | 60 -- 70 | -10.0521 | 3.45164 | 4.59330 |
| CD630_06642 | CD630_n00080 | 107 -- 142 | 42 -- 76 | -10.0453 | 3.32967 | 4.59330 |
| CD630_21730 | CD630_n00080 | 177 -- 201 | 78 -- 97 | -10.0411 | 3.45076 | 4.59330 |
| CD630_04320 | CD630_n00080 | 2 -- 40 | 50 -- 82 | -10.041 | 3.68342 | 4.59330 |
| CD630_05821 | CD630_n00080 | 50 -- 87 | 33 -- 82 | -10.0379 | 3.75661 | 4.59330 |
| CD630_09570 | CD630_n00080 | 19 -- 29 | 65 -- 76 | -9.99872 | 5.31213 | 4.59330 |
| CD630_29070 | CD630_n00080 | 19 -- 29 | 65 -- 76 | -9.99872 | 3.89477 | 4.59330 |
| bltD | CD630_n00080 | 87 -- 95 | 92 -- 100 | -9.94974 | 4.49926 | 4.59330 |
| CD630_03920 | CD630_n00080 | 22 -- 39 | 82 -- 103 | -9.90309 | 4.15934 | 4.59330 |
| CD630_22500 | CD630_n00080 | 85 -- 99 | 54 -- 70 | -9.90189 | 4.03688 | 4.59330 |
| CD630_32050 | CD630_n00080 | 156 -- 197 | 68 -- 97 | -9.90005 | 5.73797 | 4.59330 |
| CD630_13360 | CD630_n00080 | 173 -- 193 | 77 -- 97 | -9.87288 | 3.77330 | 4.59330 |
| CD630_05460 | CD630_n00080 | 173 -- 184 | 92 -- 103 | -9.86916 | 3.98753 | 4.59330 |
| CD630_07121 | CD630_n00080 | 83 -- 95 | 66 -- 79 | -9.81766 | 3.26177 | 4.59330 |
| tcdC | CD630_n00080 | 122 -- 153 | 67 -- 98 | -9.80612 | 3.76101 | 4.59330 |
| CD630_17151 | CD630_n00080 | 158 -- 194 | 41 -- 94 | -9.78195 | 6.43124 | 4.59330 |
| CD630_06200 | CD630_n00080 | 88 -- 108 | 66 -- 93 | -9.76167 | 3.57064 | 4.59330 |
| CD630_08130 | CD630_n00080 | 73 -- 96 | 82 -- 99 | -9.7556 | 4.63831 | 4.59330 |
| CD630_19850 | CD630_n00080 | 88 -- 128 | 52 -- 92 | -9.74354 | 6.88486 | 4.59330 |
| CD630_18970 | CD630_n00080 | 129 -- 173 | 65 -- 96 | -9.70037 | 3.57243 | 4.59330 |
| CD630_13350 | CD630_n00080 | 163 -- 180 | 78 -- 96 | -9.68529 | 4.15139 | 4.59330 |
| CD630_23930 | CD630_n00080 | 73 -- 109 | 54 -- 88 | -9.62323 | 3.05650 | 4.59330 |
| CD630_14260 | CD630_n00080 | 128 -- 137 | 1 -- 10 | -9.61147 | 3.22214 | 4.59330 |
| CD630_26050 | CD630_n00080 | 156 -- 200 | 18 -- 61 | -9.57409 | 6.99851 | 4.59330 |
| CD630_34530 | CD630_n00080 | 119 -- 163 | 45 -- 94 | -9.5669 | 3.83016 | 4.59330 |
| CD630_21910 | CD630_n00080 | 168 -- 199 | 2 -- 28 | -9.56352 | 4.66600 | 4.59330 |
| gcp | CD630_n00080 | 75 -- 98 | 54 -- 75 | -9.56212 | 4.92798 | 4.59330 |
| CD630_28560 | CD630_n00080 | 152 -- 166 | 36 -- 50 | -9.53798 | 3.96803 | 4.59330 |
| CD630_14990 | CD630_n00080 | 10 -- 36 | 74 -- 97 | -9.5257 | 3.78619 | 4.59330 |
| CD630_31381 | CD630_n00080 | 2 -- 11 | 51 -- 60 | -9.514 | 3.49137 | 4.59330 |
| CD630_20280 | CD630_n00080 | 119 -- 137 | 70 -- 89 | -9.51385 | 3.46343 | 4.59330 |

| | | | | | | |
|---|---|---|---|---|---|---|
| cobQ | CD630_n00150 | 96 -- 153 | 1 -- 63 | -11.3119 | 3.13524 | 8.81823 |
| CD630_27570 | CD630_n00150 | 173 -- 200 | 55 -- 83 | -11.1035 | 3.46216 | 8.81823 |
| CD630_27520 | CD630_n00150 | 93 -- 114 | 25 -- 47 | -10.4231 | 4.27693 | 8.81823 |
| CD630_25610 | CD630_n00150 | 90 -- 99 | 26 -- 35 | -10.1356 | 3.02685 | 8.81823 |
| CD630_13380 | CD630_n00150 | 128 -- 140 | 26 -- 38 | -9.86463 | 4.74408 | 8.81823 |
| CD630_19790 | CD630_n00150 | 86 -- 97 | 26 -- 37 | -9.83832 | 3.79593 | 8.81823 |
| CD630_20171 | CD630_n00150 | 79 -- 92 | 30 -- 43 | -9.54102 | 6.87491 | 8.81823 |
| CD630_03220 | CD630_n00290 | 47 -- 110 | 10 -- 71 | -18.6243 | 3.36746 | 3.85358 |
| ung | CD630_n00290 | 61 -- 129 | 5 -- 74 | -15.1559 | 3.30908 | 3.85358 |
| CD630_11730 | CD630_n00290 | 91 -- 116 | 23 -- 46 | -14.6669 | 4.80521 | 3.85358 |
| CD630_14130 | CD630_n00290 | 87 -- 155 | 11 -- 56 | -14.4794 | 4.57970 | 3.85358 |
| CD630_05590 | CD630_n00290 | 101 -- 116 | 45 -- 59 | -14.1427 | 3.39670 | 3.85358 |
| cdtR | CD630_n00290 | 81 -- 114 | 29 -- 64 | -13.6562 | 5.84368 | 3.85358 |
| oppB | CD630_n00290 | 87 -- 119 | 30 -- 55 | -13.2635 | 2.98328 | 3.85358 |
| CD630_19580 | CD630_n00290 | 46 -- 86 | 17 -- 71 | -13.0214 | 4.51902 | 3.85358 |
| xdhA | CD630_n00290 | 46 -- 58 | 43 -- 54 | -12.7388 | 4.45147 | 3.85358 |
| CD630_08090 | CD630_n00290 | 85 -- 95 | 123 -- 134 | -12.6288 | 3.57167 | 3.85358 |
| CD630_14050 | CD630_n00290 | 135 -- 188 | 51 -- 96 | -12.5892 | 3.30894 | 3.85358 |
| CD630_29650 | CD630_n00290 | 21 -- 90 | 20 -- 76 | -12.4924 | 3.67128 | 3.85358 |
| spaK | CD630_n00290 | 51 -- 61 | 44 -- 54 | -12.3668 | 3.04530 | 3.85358 |
| CD630_33682 | CD630_n00290 | 76 -- 93 | 4 -- 20 | -12.3563 | 5.77720 | 3.85358 |
| CD630_04410 | CD630_n00290 | 153 -- 163 | 49 -- 59 | -12.315 | 3.90451 | 3.85358 |
| tcdC | CD630_n00290 | 187 -- 197 | 46 -- 56 | -11.8693 | 3.76101 | 3.85358 |
| CD630_24591 | CD630_n00290 | 28 -- 115 | 34 -- 104 | -11.8547 | 3.54497 | 3.85358 |
| CD630_08100 | CD630_n00290 | 181 -- 199 | 34 -- 52 | -11.803 | 4.39181 | 3.85358 |
| bglA | CD630_n00290 | 50 -- 118 | 37 -- 79 | -11.8016 | 4.50315 | 3.85358 |
| CD630_28320 | CD630_n00290 | 91 -- 110 | 114 -- 132 | -11.5466 | 3.99184 | 3.85358 |
| CD630_28870 | CD630_n00290 | 7 -- 62 | 20 -- 84 | -11.4988 | 3.41991 | 3.85358 |
| CD630_19903 | CD630_n00290 | 76 -- 93 | 4 -- 20 | -11.4412 | 5.79994 | 3.85358 |
| CD630_18090 | CD630_n00290 | 16 -- 33 | 26 -- 43 | -11.3749 | 4.23961 | 3.85358 |
| CD630_35790 | CD630_n00290 | 152 -- 165 | 120 -- 134 | -11.3226 | 6.97665 | 3.85358 |
| CD630_09840 | CD630_n00290 | 74 -- 99 | 39 -- 67 | -11.2589 | 4.50009 | 3.85358 |
| folC | CD630_n00290 | 85 -- 92 | 8 -- 15 | -11.2369 | 3.85634 | 3.85358 |
| mobA | CD630_n00290 | 123 -- 193 | 8 -- 80 | -11.1366 | 4.25703 | 3.85358 |
| CD630_20541 | CD630_n00290 | 104 -- 115 | 48 -- 59 | -11.1031 | 5.74527 | 3.85358 |
| CD630_02420 | CD630_n00290 | 33 -- 44 | 123 -- 134 | -11.0649 | 3.71048 | 3.85358 |
| CD630_16460 | CD630_n00290 | 127 -- 161 | 11 -- 48 | -11.0211 | 4.64028 | 3.85358 |
| CD630_26680 | CD630_n00290 | 50 -- 95 | 35 -- 71 | -11.0031 | 3.53956 | 3.85358 |
| flgB | CD630_n00290 | 68 -- 97 | 49 -- 81 | -10.8863 | 3.25736 | 3.85358 |
| CD630_21750 | CD630_n00290 | 83 -- 92 | 8 -- 17 | -10.879 | 3.70626 | 3.85358 |
| CD630_32720 | CD630_n00290 | 130 -- 145 | 48 -- 63 | -10.7148 | 4.74276 | 3.85358 |
| CD630_14700 | CD630_n00290 | 56 -- 67 | 45 -- 56 | -10.7123 | 3.98234 | 3.85358 |
| CD630_23440 | CD630_n00290 | 182 -- 194 | 42 -- 54 | -10.6403 | 3.34841 | 3.85358 |
| CD630_17160 | CD630_n00290 | 87 -- 123 | 27 -- 59 | -10.5558 | 3.50228 | 3.85358 |
| CD630_21790 | CD630_n00290 | 135 -- 154 | 29 -- 47 | -10.509 | 4.93762 | 3.85358 |
| CD630_06340 | CD630_n00290 | 24 -- 110 | 33 -- 109 | -10.4195 | 4.35983 | 3.85358 |
| cbpA | CD630_n00290 | 67 -- 96 | 8 -- 38 | -10.4162 | 3.03040 | 3.85358 |
| CD630_19904 | CD630_n00290 | 160 -- 173 | 125 -- 135 | -10.3665 | 5.96526 | 3.85358 |
| CD630_32690 | CD630_n00290 | 67 -- 113 | 36 -- 78 | -10.3212 | 7.32676 | 3.85358 |
| CD630_08370 | CD630_n00290 | 82 -- 91 | 51 -- 60 | -10.291 | 3.65639 | 3.85358 |
| CD630_36250 | CD630_n00290 | 187 -- 200 | 43 -- 60 | -10.2297 | 4.13224 | 3.85358 |
| CD630_31010 | CD630_n00290 | 185 -- 196 | 122 -- 133 | -10.2037 | 4.96305 | 3.85358 |
| CD630_14240 | CD630_n00290 | 67 -- 100 | 33 -- 62 | -10.1582 | 8.60766 | 3.85358 |
| CD630_18660 | CD630_n00290 | 8 -- 17 | 51 -- 60 | -10.1259 | 3.50371 | 3.85358 |
| CD630_20670 | CD630_n00290 | 60 -- 99 | 37 -- 82 | -10.1178 | 4.42757 | 3.85358 |
| CD630_34050 | CD630_n00290 | 140 -- 161 | 24 -- 44 | -10.0603 | 4.03863 | 3.85358 |
| CD630_35260 | CD630_n00290 | 72 -- 108 | 4 -- 40 | -9.99076 | 5.77277 | 3.85358 |
| CD630_15850 | CD630_n00290 | 123 -- 131 | 51 -- 59 | -9.96879 | 4.32968 | 3.85358 |
| CD630_08390 | CD630_n00290 | 47 -- 104 | 5 -- 54 | -9.95443 | 5.16000 | 3.85358 |
| CD630_14340 | CD630_n00290 | 70 -- 80 | 124 -- 134 | -9.94163 | 5.10868 | 3.85358 |
| CD630_10220 | CD630_n00290 | 88 -- 96 | 49 -- 57 | -9.94162 | 4.08233 | 3.85358 |
| CD630_28330 | CD630_n00290 | 84 -- 96 | 5 -- 17 | -9.92332 | 3.29122 | 3.85358 |
| CD630_33300 | CD630_n00290 | 122 -- 142 | 43 -- 60 | -9.83128 | 3.79149 | 3.85358 |
| malY | CD630_n00290 | 75 -- 110 | 24 -- 60 | -9.82518 | 4.46345 | 3.85358 |
| CD630_27210 | CD630_n00290 | 88 -- 109 | 25 -- 56 | -9.81609 | 4.18786 | 3.85358 |
| CD630_21610 | CD630_n00290 | 10 -- 93 | 9 -- 101 | -9.73363 | 3.16662 | 3.85358 |
| cdu2 | CD630_n00290 | 180 -- 190 | 40 -- 50 | -9.72799 | 2.97335 | 3.85358 |
| CD630_22070 | CD630_n00290 | 85 -- 98 | 2 -- 15 | -9.72023 | 4.09651 | 3.85358 |
| CD630_05220 | CD630_n00290 | 183 -- 195 | 42 -- 59 | -9.71524 | 3.65057 | 3.85358 |
| cstA | CD630_n00290 | 103 -- 114 | 121 -- 132 | -9.71263 | 3.53584 | 3.85358 |
| CD630_08070 | CD630_n00290 | 82 -- 96 | 34 -- 50 | -9.70104 | 4.87819 | 3.85358 |
| cprK | CD630_n00290 | 24 -- 95 | 11 -- 78 | -9.68136 | 3.96216 | 3.85358 |
| CD630_18650 | CD630_n00290 | 90 -- 99 | 5 -- 14 | -9.63849 | 3.75892 | 3.85358 |
| CD630_17710 | CD630_n00290 | 85 -- 93 | 51 -- 59 | -9.638 | 4.37688 | 3.85358 |
| CD630_34080 | CD630_n00290 | 102 -- 111 | 38 -- 47 | -9.57521 | 3.20224 | 3.85358 |
| CD630_12650 | CD630_n00290 | 111 -- 141 | 51 -- 83 | -9.56831 | 6.31827 | 3.85358 |
| CD630_23090 | CD630_n00290 | 77 -- 93 | 5 -- 20 | -9.55073 | 4.25622 | 3.85358 |
| CD630_21140 | CD630_n00290 | 14 -- 24 | 118 -- 128 | -9.53431 | 3.33304 | 3.85358 |
| gcvTPA | CD630_n00330 | 67 -- 122 | 29 -- 72 | -19.4118 | 4.77229 | 6.46464 |
| oppB | CD630_n00330 | 88 -- 113 | 36 -- 53 | -18.7325 | 2.98328 | 6.46464 |
| CD630_34030 | CD630_n00330 | 73 -- 107 | 30 -- 66 | -18.714 | 3.05766 | 6.46464 |
| CD630_35260 | CD630_n00330 | 66 -- 100 | 35 -- 55 | -18.5825 | 5.77277 | 6.46464 |
| cbpA | CD630_n00330 | 77 -- 96 | 38 -- 60 | -17.546 | 3.03040 | 6.46464 |
| CD630_29750 | CD630_n00330 | 69 -- 95 | 39 -- 66 | -17.1346 | 4.59146 | 6.46464 |
| CD630_19580 | CD630_n00330 | 61 -- 93 | 32 -- 52 | -16.6279 | 4.51902 | 6.46464 |
| CD630_18910 | CD630_n00330 | 120 -- 189 | 36 -- 111 | -16.4037 | 3.30688 | 6.46464 |
| CD630_08100 | CD630_n00330 | 32 -- 50 | 161 -- 179 | -16.3008 | 4.39181 | 6.46464 |
| CD630_25310 | CD630_n00330 | 6 -- 96 | 36 -- 103 | -15.8762 | 3.99861 | 6.46464 |
| CD630_28181 | CD630_n00330 | 63 -- 78 | 165 -- 182 | -15.5053 | 3.91299 | 6.46464 |
| CD630_07480 | CD630_n00330 | 80 -- 108 | 29 -- 66 | -15.094 | 5.18321 | 6.46464 |
| CD630_11241 | CD630_n00330 | 166 -- 192 | 78 -- 109 | -14.1071 | 4.85723 | 6.46464 |
| CD630_21490 | CD630_n00330 | 190 -- 197 | 37 -- 44 | -14.0646 | 3.37462 | 6.46464 |
| CD630_34530 | CD630_n00330 | 38 -- 98 | 45 -- 108 | -13.8823 | 3.83016 | 6.46464 |
| CD630_28320 | CD630_n00330 | 78 -- 96 | 34 -- 54 | -13.8583 | 3.99184 | 6.46464 |
| CD630_33600 | CD630_n00330 | 20 -- 33 | 162 -- 175 | -13.6886 | 3.73196 | 6.46464 |
| CD630_05230 | CD630_n00330 | 10 -- 63 | 142 -- 180 | -13.6709 | 3.73787 | 6.46464 |
| CD630_27630 | CD630_n00330 | 163 -- 177 | 159 -- 173 | -13.661 | 3.54199 | 6.46464 |
| CD630_23820 | CD630_n00330 | 169 -- 185 | 165 -- 180 | -13.653 | 3.18419 | 6.46464 |
| CD630_01560 | CD630_n00330 | 146 -- 165 | 37 -- 53 | -13.5928 | 4.06352 | 6.46464 |
| CD630_14050 | CD630_n00330 | 23 -- 35 | 39 -- 50 | -13.59 | 3.30894 | 6.46464 |
| ssuC | CD630_n00330 | 62 -- 75 | 42 -- 53 | -13.5601 | 3.45491 | 6.46464 |
| uxaA | CD630_n00330 | 187 -- 201 | 168 -- 180 | -13.5329 | 4.26165 | 6.46464 |
| CD630_25620 | CD630_n00330 | 155 -- 168 | 164 -- 177 | -13.3959 | 3.02685 | 6.46464 |
| CD630_15111 | CD630_n00330 | 83 -- 116 | 33 -- 58 | -13.3724 | 4.18664 | 6.46464 |
| nusB | CD630_n00330 | 110 -- 132 | 29 -- 53 | -13.2695 | 4.92798 | 6.46464 |
| CD630_04770 | CD630_n00330 | 71 -- 97 | 152 -- 179 | -13.1747 | 3.58478 | 6.46464 |
| CD630_24591 | CD630_n00330 | 51 -- 94 | 40 -- 86 | -13.1583 | 3.54497 | 6.46464 |
| CD630_16460 | CD630_n00330 | 90 -- 176 | 26 -- 110 | -13.146 | 4.64028 | 6.46464 |
| CD630_28330 | CD630_n00330 | 38 -- 96 | 35 -- 88 | -13.0847 | 3.29122 | 6.46464 |
| CD630_26920 | CD630_n00330 | 171 -- 185 | 169 -- 180 | -13.0708 | 4.58060 | 6.46464 |
| CD630_01700 | CD630_n00330 | 77 -- 87 | 168 -- 178 | -12.9749 | 4.97234 | 6.46464 |
| CD630_06690 | CD630_n00330 | 99 -- 117 | 166 -- 183 | -12.9018 | 5.82709 | 6.46464 |
| dnaC | CD630_n00330 | 82 -- 99 | 159 -- 175 | -12.8958 | 6.30796 | 6.46464 |
| argS | CD630_n00330 | 55 -- 93 | 32 -- 69 | -12.8927 | 4.96036 | 6.46464 |
| bglA | CD630_n00330 | 153 -- 182 | 34 -- 55 | -12.8251 | 4.86402 | 6.46464 |
| etfA | CD630_n00330 | 149 -- 167 | 32 -- 50 | -12.5079 | 3.41204 | 6.46464 |
| CD630_06970 | CD630_n00330 | 164 -- 181 | 166 -- 184 | -12.388 | 3.82312 | 6.46464 |
| CD630_30620 | CD630_n00330 | 2 -- 94 | 35 -- 114 | -12.2871 | 4.15443 | 6.46464 |
| CD630_18120 | CD630_n00330 | 81 -- 109 | 32 -- 66 | -12.2822 | 3.99713 | 6.46464 |
| CD630_20460 | CD630_n00330 | 98 -- 108 | 35 -- 45 | -12.2616 | 7.94034 | 6.46464 |
| def1 | CD630_n00330 | 74 -- 112 | 33 -- 66 | -12.2381 | 3.99720 | 6.46464 |
| CD630_31220 | CD630_n00330 | 89 -- 97 | 37 -- 45 | -11.9932 | 3.73779 | 6.46464 |
| CD630_33300 | CD630_n00330 | 127 -- 142 | 42 -- 57 | -11.9873 | 3.79149 | 6.46464 |
| CD630_14700 | CD630_n00330 | 55 -- 67 | 43 -- 54 | -11.9613 | 3.98234 | 6.46464 |
| CD630_21530 | CD630_n00330 | 56 -- 72 | 168 -- 183 | -11.9084 | 3.39774 | 6.46464 |
| kdpE | CD630_n00330 | 55 -- 72 | 92 -- 109 | -11.7659 | 4.27140 | 6.46464 |
| CD630_21620 | CD630_n00330 | 86 -- 161 | 32 -- 112 | -11.7533 | 3.02649 | 6.46464 |
| CD630_14130 | CD630_n00330 | 101 -- 195 | 43 -- 111 | -11.71 | 4.57970 | 6.46464 |
| CD630_28560 | CD630_n00330 | 152 -- 170 | 35 -- 49 | -11.6617 | 3.96803 | 6.46464 |
| CD630_05220 | CD630_n00330 | 69 -- 81 | 98 -- 110 | -11.6051 | 3.65057 | 6.46464 |
| CD630_09850 | CD630_n00330 | 121 -- 132 | 162 -- 173 | -11.5791 | 4.18679 | 6.46464 |
| metQ | CD630_n00330 | 80 -- 94 | 38 -- 57 | -11.5603 | 3.87190 | 6.46464 |
| CD630_32710 | CD630_n00330 | 146 -- 157 | 32 -- 48 | -11.533 | 4.74276 | 6.46464 |
| CD630_06060 | CD630_n00330 | 23 -- 41 | 33 -- 56 | -11.4801 | 4.94050 | 6.46464 |
| bglA | CD630_n00330 | 63 -- 134 | 42 -- 108 | -11.4545 | 4.50315 | 6.46464 |
| CD630_13600 | CD630_n00330 | 137 -- 155 | 165 -- 182 | -11.4027 | 4.45481 | 6.46464 |
| CD630_18090 | CD630_n00330 | 18 -- 32 | 166 -- 180 | -11.3967 | 4.23961 | 6.46464 |
| CD630_06140 | CD630_n00330 | 31 -- 93 | 43 -- 114 | -11.3891 | 4.12519 | 6.46464 |
| CD630_16950 | CD630_n00330 | 185 -- 200 | 163 -- 177 | -11.2467 | 4.42569 | 6.46464 |
| CD630_02420 | CD630_n00330 | 24 -- 41 | 33 -- 50 | -11.2249 | 3.71048 | 6.46464 |
| pgm | CD630_n00330 | 56 -- 106 | 37 -- 91 | -11.1954 | 3.45439 | 6.46464 |
| cbiO | CD630_n00330 | 176 -- 186 | 48 -- 58 | -11.1819 | 3.79830 | 6.46464 |
| fbp | CD630_n00330 | 144 -- 153 | 166 -- 175 | -11.1641 | 3.80071 | 6.46464 |
| CD630_21030 | CD630_n00330 | 46 -- 59 | 164 -- 177 | -11.1514 | 3.08733 | 6.46464 |
| CD630_15740 | CD630_n00330 | 72 -- 94 | 154 -- 181 | -11.1142 | 4.43222 | 6.46464 |
| tcdC | CD630_n00330 | 186 -- 193 | 47 -- 54 | -11.1099 | 3.76101 | 6.46464 |
| iorA | CD630_n00330 | 57 -- 63 | 38 -- 44 | -11.0338 | 3.83173 | 6.46464 |
| CD630_32880 | CD630_n00330 | 184 -- 195 | 162 -- 173 | -11.0224 | 3.41276 | 6.46464 |
| CD630_08370 | CD630_n00330 | 1 -- 27 | 47 -- 71 | -11.0141 | 3.65639 | 6.46464 |
| CD630_14340 | CD630_n00330 | 53 -- 72 | 32 -- 49 | -11.0071 | 5.10868 | 6.46464 |
| CD630_11240 | CD630_n00330 | 22 -- 91 | 35 -- 103 | -10.9995 | 3.20274 | 6.46464 |
| CD630_05880 | CD630_n00330 | 94 -- 104 | 101 -- 111 | -10.9945 | 4.28339 | 6.46464 |
| ldh | CD630_n00330 | 29 -- 90 | 38 -- 86 | -10.987 | 4.35818 | 6.46464 |
| terD | CD630_n00330 | 184 -- 198 | 97 -- 111 | -10.9869 | 3.33148 | 6.46464 |
| xdhA | CD630_n00330 | 56 -- 66 | 168 -- 179 | -10.9538 | 4.45147 | 6.46464 |
| CD630_04320 | CD630_n00330 | 16 -- 27 | 100 -- 111 | -10.9442 | 3.68342 | 6.46464 |
| CD630_14810 | CD630_n00330 | 90 -- 154 | 41 -- 106 | -10.8727 | 3.35323 | 6.46464 |
| CD630_03220 | CD630_n00330 | 98 -- 112 | 30 -- 45 | -10.723 | 3.36746 | 6.46464 |
| CD630_31010 | CD630_n00330 | 178 -- 188 | 41 -- 51 | -10.7159 | 4.96305 | 6.46464 |
| CD630_10050 | CD630_n00330 | 151 -- 167 | 40 -- 54 | -10.7002 | 3.93750 | 6.46464 |
| CD630_21140 | CD630_n00330 | 11 -- 18 | 171 -- 178 | -10.6946 | 3.33304 | 6.46464 |
| CD630_29270 | CD630_n00330 | 52 -- 68 | 164 -- 179 | -10.6682 | 3.70174 | 6.46464 |
| CD630_08780 | CD630_n00330 | 78 -- 89 | 48 -- 59 | -10.6666 | 7.76690 | 6.46464 |
| CD630_27110 | CD630_n00330 | 191 -- 199 | 167 -- 175 | -10.6587 | 3.78231 | 6.46464 |
| CD630_20170 | CD630_n00330 | 102 -- 114 | 168 -- 180 | -10.6437 | 6.90201 | 6.46464 |
| CD630_14360 | CD630_n00330 | 188 -- 199 | 44 -- 56 | -10.6434 | 3.02079 | 6.46464 |
| CD630_23760 | CD630_n00330 | 56 -- 62 | 173 -- 179 | -10.633 | 4.54307 | 6.46464 |
| CD630_08940 | CD630_n00330 | 41 -- 95 | 37 -- 72 | -10.6282 | 3.72225 | 6.46464 |
| CD630_26750 | CD630_n00330 | 89 -- 107 | 36 -- 53 | -10.6195 | 3.60610 | 6.46464 |
| cwp17 | CD630_n00330 | 19 -- 25 | 44 -- 50 | -10.6175 | 3.43363 | 6.46464 |
| cynT | CD630_n00330 | 75 -- 91 | 40 -- 60 | -10.6125 | 4.21370 | 6.46464 |
| CD630_33682 | CD630_n00330 | 64 -- 92 | 35 -- 59 | -10.5926 | 5.77720 | 6.46464 |
| CD630_36040 | CD630_n00330 | 121 -- 127 | 38 -- 44 | -10.5872 | 3.52009 | 6.46464 |
| CD630_05940 | CD630_n00330 | 85 -- 111 | 32 -- 53 | -10.5709 | 4.18595 | 6.46464 |
| CD630_21910 | CD630_n00330 | 85 -- 95 | 42 -- 54 | -10.5658 | 4.66600 | 6.46464 |
| CD630_21610 | CD630_n00330 | 81 -- 94 | 47 -- 60 | -10.511 | 3.16662 | 6.46464 |
| CD630_16450 | CD630_n00330 | 158 -- 165 | 173 -- 180 | -10.4412 | 4.57547 | 6.46464 |
| chrA | CD630_n00330 | 147 -- 159 | 36 -- 47 | -10.4214 | 5.27255 | 6.46464 |
| dltD | CD630_n00330 | 158 -- 164 | 44 -- 50 | -10.4117 | 4.40777 | 6.46464 |
| srlD | CD630_n00330 | 131 -- 146 | 166 -- 179 | -10.4042 | 4.19843 | 6.46464 |
| CD630_27230 | CD630_n00330 | 194 -- 200 | 173 -- 179 | -10.4014 | 3.52462 | 6.46464 |
| pilC | CD630_n00330 | 48 -- 68 | 157 -- 177 | -10.3941 | 3.54230 | 6.46464 |
| CD630_17160 | CD630_n00330 | 88 -- 111 | 33 -- 54 | -10.3855 | 3.50228 | 6.46464 |
| CD630_23861 | CD630_n00330 | 152 -- 165 | 165 -- 178 | -10.384 | 5.83068 | 6.46464 |
| feoB | CD630_n00330 | 24 -- 31 | 171 -- 178 | -10.3829 | 4.05364 | 6.46464 |
| CD630_21540 | CD630_n00330 | 80 -- 94 | 162 -- 175 | -10.3514 | 4.33090 | 6.46464 |
| CD630_05870 | CD630_n00330 | 189 -- 197 | 45 -- 53 | -10.3369 | 4.28339 | 6.46464 |
| CD630_30140 | CD630_n00330 | 10 -- 22 | 101 -- 113 | -10.3279 | 3.28343 | 6.46464 |
| CD630_17281 | CD630_n00330 | 72 -- 93 | 38 -- 59 | -10.3054 | 5.91544 | 6.46464 |
| eutV | CD630_n00330 | 147 -- 179 | 144 -- 180 | -10.3013 | 4.55715 | 6.46464 |
| cspC | CD630_n00330 | 81 -- 95 | 29 -- 43 | -10.2785 | 5.48848 | 6.46464 |
| CD630_06060 | CD630_n00330 | 132 -- 162 | 158 -- 179 | -10.2625 | 4.94050 | 6.46464 |
| CD630_21700 | CD630_n00330 | 188 -- 201 | 168 -- 181 | -10.2407 | 3.96650 | 6.46464 |
| vncR | CD630_n00330 | 82 -- 92 | 165 -- 175 | -10.2164 | 3.93642 | 6.46464 |
| CD630_26630 | CD630_n00330 | 189 -- 198 | 168 -- 177 | -10.2036 | 4.61557 | 6.46464 |
| CD630_09760 | CD630_n00330 | 72 -- 83 | 169 -- 181 | -10.1971 | 4.83410 | 6.46464 |
| CD630_28910 | CD630_n00330 | 72 -- 83 | 169 -- 181 | -10.1971 | 4.51664 | 6.46464 |
| CD630_04410 | CD630_n00330 | 83 -- 94 | 40 -- 49 | -10.1492 | 3.90451 | 6.46464 |
| CD630_13780 | CD630_n00330 | 182 -- 197 | 166 -- 180 | -10.1224 | 3.52274 | 6.46464 |
| CD630_15461 | CD630_n00330 | 24 -- 92 | 48 -- 109 | -10.1161 | 9.13442 | 6.46464 |
| CD630_19680 | CD630_n00330 | 137 -- 151 | 30 -- 43 | -10.1067 | 3.58080 | 6.46464 |
| CD630_22070 | CD630_n00330 | 42 -- 95 | 35 -- 110 | -10.0464 | 4.09651 | 6.46464 |
| CD630_17151 | CD630_n00330 | 173 -- 200 | 44 -- 65 | -10.0332 | 6.43124 | 6.46464 |
| rolB | CD630_n00330 | 120 -- 133 | 100 -- 113 | -9.97085 | 3.60270 | 6.46464 |
| CD630_20920 | CD630_n00330 | 57 -- 141 | 33 -- 104 | -9.96714 | 4.13812 | 6.46464 |
| cheW | CD630_n00330 | 176 -- 197 | 45 -- 69 | -9.95807 | 3.72371 | 6.46464 |
| cspD | CD630_n00330 | 158 -- 169 | 100 -- 111 | -9.94929 | 3.23587 | 6.46464 |
| CD630_06642 | CD630_n00330 | 193 -- 201 | 39 -- 47 | -9.93908 | 3.32967 | 6.46464 |
| CD630_24760 | CD630_n00330 | 69 -- 87 | 39 -- 66 | -9.92496 | 5.19485 | 6.46464 |
| pilT | CD630_n00330 | 194 -- 200 | 37 -- 43 | -9.88991 | 3.24639 | 6.46464 |
| cobQ | CD630_n00330 | 125 -- 143 | 165 -- 179 | -9.85193 | 3.13524 | 6.46464 |
| CD630_16940 | CD630_n00330 | 147 -- 158 | 166 -- 178 | -9.82722 | 4.14665 | 6.46464 |
| CD630_31210 | CD630_n00330 | 84 -- 90 | 42 -- 48 | -9.81834 | 4.17693 | 6.46464 |
| CD630_27520 | CD630_n00330 | 85 -- 97 | 166 -- 180 | -9.80152 | 4.27693 | 6.46464 |
| CD630_21790 | CD630_n00330 | 156 -- 164 | 172 -- 180 | -9.79961 | 4.93762 | 6.46464 |
| CD630_25040 | CD630_n00330 | 167 -- 198 | 145 -- 178 | -9.7908 | 3.43898 | 6.46464 |
| CD630_32070 | CD630_n00330 | 72 -- 81 | 100 -- 109 | -9.78986 | 3.51964 | 6.46464 |
| CD630_32700 | CD630_n00330 | 192 -- 199 | 171 -- 178 | -9.78003 | 4.74276 | 6.46464 |
| cdu2 | CD630_n00330 | 32 -- 98 | 38 -- 71 | -9.75609 | 2.97335 | 6.46464 |
| cdtR | CD630_n00330 | 78 -- 107 | 36 -- 60 | -9.75397 | 5.84368 | 6.46464 |
| CD630_20650 | CD630_n00330 | 9 -- 17 | 33 -- 41 | -9.68197 | 4.00083 | 6.46464 |
| CD630_13380 | CD630_n00330 | 170 -- 187 | 96 -- 113 | -9.66941 | 4.74408 | 6.46464 |
| bltD | CD630_n00330 | 53 -- 93 | 43 -- 92 | -9.65969 | 4.49926 | 6.46464 |
| CD630_20670 | CD630_n00330 | 83 -- 110 | 33 -- 57 | -9.65725 | 4.42757 | 6.46464 |
| CD630_30360 | CD630_n00330 | 50 -- 57 | 175 -- 182 | -9.60446 | 4.26509 | 6.46464 |
| glnA | CD630_n00330 | 34 -- 43 | 100 -- 109 | -9.59502 | 3.87629 | 6.46464 |
| CD630_21070 | CD630_n00330 | 41 -- 49 | 169 -- 177 | -9.573 | 5.35019 | 6.46464 |
| CD630_14240 | CD630_n00330 | 74 -- 129 | 29 -- 66 | -9.57051 | 8.60766 | 6.46464 |
| CD630_15150 | CD630_n00330 | 46 -- 97 | 34 -- 102 | -9.56571 | 3.35564 | 6.46464 |
| CD630_06050 | CD630_n00330 | 75 -- 81 | 37 -- 43 | -9.56211 | 3.59332 | 6.46464 |
| CD630_08460 | CD630_n00330 | 94 -- 160 | 43 -- 109 | -9.55029 | 4.15083 | 6.46464 |
| srlEb | CD630_n00330 | 67 -- 107 | 143 -- 177 | -9.53119 | 3.50140 | 6.46464 |
| CD630_05230 | CD630_n00340 | 3 -- 41 | 97 -- 138 | -17.662 | 3.73787 | 5.83478 |
| CD630_18910 | CD630_n00340 | 36 -- 73 | 31 -- 63 | -14.7728 | 3.30688 | 5.83478 |
| CD630_19680 | CD630_n00340 | 102 -- 153 | 96 -- 149 | -13.6481 | 3.58080 | 5.83478 |
| etfA | CD630_n00340 | 149 -- 171 | 126 -- 151 | -12.5886 | 3.41204 | 5.83478 |
| CD630_31530 | CD630_n00340 | 17 -- 31 | 32 -- 46 | -12.4571 | 3.49100 | 5.83478 |
| CD630_18610 | CD630_n00340 | 19 -- 53 | 103 -- 142 | -12.2411 | 4.15582 | 5.83478 |

| | | | | | | |
|---|---|---|---|---|---|---|
| CD630_31220 | CD630_n00340 | 25 – 50 | 35 – 58 | -12.2035 | 3.73779 | 5.83478 |
| CD630_27240 | CD630_n00340 | 7 – 14 | 124 – 131 | -11.6164 | 3.52462 | 5.83478 |
| CD630_25190 | CD630_n00340 | 50 – 59 | 121 – 130 | -11.5358 | 3.19663 | 5.83478 |
| CD630_06540 | CD630_n00340 | 43 – 73 | 36 – 66 | -11.4428 | 3.54673 | 5.83478 |
| CD630_23760 | CD630_n00340 | 56 – 70 | 115 – 130 | -11.3862 | 4.54307 | 5.83478 |
| rolA | CD630_n00340 | 19 – 38 | 125 – 140 | -11.3848 | 4.56362 | 5.83478 |
| CD630_08780 | CD630_n00340 | 8 – 19 | 34 – 45 | -11.3362 | 7.76690 | 5.83478 |
| CD630_21490 | CD630_n00340 | 89 – 108 | 125 – 144 | -11.3032 | 3.37462 | 5.83478 |
| CD630_13960 | CD630_n00340 | 31 – 63 | 27 – 58 | -11.0913 | 3.30375 | 5.83478 |
| CD630_18680 | CD630_n00340 | 84 – 138 | 97 – 150 | -11.0105 | 3.04315 | 5.83478 |
| CD630_32700 | CD630_n00340 | 60 – 90 | 96 – 128 | -11.0015 | 4.74276 | 5.83478 |
| CD630_14630 | CD630_n00340 | 137 – 173 | 100 – 143 | -10.9523 | 3.34485 | 5.83478 |
| CD630_30310 | CD630_n00340 | 40 – 59 | 36 – 54 | -10.8108 | 4.56059 | 5.83478 |
| vncR | CD630_n00340 | 172 – 186 | 28 – 42 | -10.7725 | 3.93642 | 5.83478 |
| CD630_23780 | CD630_n00340 | 55 – 95 | 20 – 55 | -10.7283 | 3.91629 | 5.83478 |
| CD630_31000 | CD630_n00340 | 117 – 124 | 124 – 131 | -10.3231 | 3.70946 | 5.83478 |
| CD630_21240 | CD630_n00340 | 91 – 126 | 28 – 65 | -10.2856 | 5.88879 | 5.83478 |
| CD630_21570 | CD630_n00340 | 40 – 87 | 1 – 62 | -10.1733 | 6.48385 | 5.83478 |
| CD630_23820 | CD630_n00340 | 90 – 106 | 127 – 143 | -10.1716 | 3.18419 | 5.83478 |
| CD630_21030 | CD630_n00340 | 46 – 90 | 9 – 55 | -10.1414 | 3.08733 | 5.83478 |
| CD630_20980 | CD630_n00340 | 126 – 132 | 124 – 130 | -10.1245 | 8.12847 | 5.83478 |
| CD630_16640 | CD630_n00340 | 102 – 121 | 30 – 49 | -10.1152 | 3.61548 | 5.83478 |
| CD630_07470 | CD630_n00340 | 20 – 47 | 35 – 64 | -10.0139 | 3.37143 | 5.83478 |
| CD630_33650 | CD630_n00340 | 54 – 104 | 1 – 58 | -10.0019 | 4.80565 | 5.83478 |
| CD630_23770 | CD630_n00340 | 60 – 87 | 101 – 128 | -9.98296 | 3.25181 | 5.83478 |
| CD630_27230 | CD630_n00340 | 194 – 201 | 123 – 130 | -9.76134 | 3.52462 | 5.83478 |
| CD630_29840 | CD630_n00340 | 31 – 51 | 124 – 143 | -9.75231 | 4.48825 | 5.83478 |
| CD630_04550 | CD630_n00340 | 5 – 20 | 44 – 59 | -9.72823 | 3.05227 | 5.83478 |
| CD630_10880 | CD630_n00340 | 11 – 31 | 98 – 116 | -9.65722 | 5.05048 | 5.83478 |
| CD630_02420 | CD630_n00340 | 7 – 31 | 9 – 40 | -9.58642 | 3.71048 | 5.83478 |
| pgm | CD630_n00340 | 83 – 116 | 104 – 138 | -9.54988 | 3.45439 | 5.83478 |
| CD630_07121 | CD630_n00340 | 10 – 54 | 26 – 58 | -9.5326 | 3.26177 | 5.83478 |
| CD630_26050 | CD630_n00370 | 127 – 166 | 2 – 49 | -12.7899 | 6.99851 | 7.05302 |
| CD630_22991 | CD630_n00370 | 191 – 201 | 23 – 33 | -12.1795 | 6.03728 | 7.05302 |
| CD630_05230 | CD630_n00370 | 4 – 20 | 6 – 24 | -11.6463 | 3.73787 | 7.05302 |
| nrdD | CD630_n00370 | 48 – 62 | 88 – 101 | -11.1197 | 3.28290 | 7.05302 |
| CD630_06540 | CD630_n00370 | 93 – 119 | 41 – 65 | -10.9267 | 3.54673 | 7.05302 |
| oppB | CD630_n00370 | 90 – 103 | 86 – 101 | -10.1761 | 2.98328 | 7.05302 |
| CD630_25310 | CD630_n00370 | 13 – 35 | 49 – 65 | -9.99031 | 3.99861 | 7.05302 |
| cheW | CD630_n00370 | 133 – 178 | 52 – 95 | -9.81537 | 3.72371 | 7.05302 |
| gcvTPA | CD630_n00370 | 172 – 181 | 90 – 99 | -9.7664 | 4.77229 | 7.05302 |
| CD630_36781 | CD630_n00370 | 175 – 192 | 44 – 62 | -9.63568 | 3.75946 | 7.05302 |
| CD630_21750 | CD630_n00390 | 59 – 106 | 270 – 323 | -16.7288 | 3.70626 | 4.83391 |
| CD630_21570 | CD630_n00390 | 151 – 186 | 110 – 141 | -16.4684 | 6.48385 | 4.83391 |
| CD630_02790 | CD630_n00390 | 127 – 171 | 320 – 356 | -16.4101 | 4.77661 | 4.83391 |
| CD630_20450 | CD630_n00390 | 111 – 165 | 85 – 142 | -16.396 | 3.77388 | 4.83391 |
| CD630_14700 | CD630_n00390 | 76 – 157 | 106 – 200 | -16.3357 | 3.98234 | 4.83391 |
| CD630_11241 | CD630_n00390 | 164 – 193 | 178 – 206 | -16.1288 | 4.85723 | 4.83391 |
| xdhA | CD630_n00390 | 47 – 101 | 105 – 157 | -16.07 | 4.45147 | 4.83391 |
| CD630_05211 | CD630_n00390 | 112 – 153 | 212 – 260 | -15.9862 | 3.97048 | 4.83391 |
| CD630_17281 | CD630_n00390 | 50 – 94 | 106 – 151 | -15.9585 | 5.91544 | 4.83391 |
| oppC | CD630_n00390 | 143 – 193 | 99 – 150 | -15.8425 | 3.01695 | 4.83391 |
| CD630_15360 | CD630_n00390 | 117 – 176 | 269 – 319 | -15.7781 | 4.47208 | 4.83391 |
| CD630_31530 | CD630_n00390 | 107 – 146 | 106 – 144 | -15.3625 | 3.49100 | 4.83391 |
| CD630_13960 | CD630_n00390 | 117 – 175 | 183 – 246 | -15.323 | 3.30375 | 4.83391 |
| CD630_05830 | CD630_n00390 | 47 – 94 | 102 – 149 | -15.3102 | 3.35360 | 4.83391 |
| CD630_07840 | CD630_n00390 | 109 – 163 | 105 – 153 | -15.1266 | 4.26761 | 4.83391 |
| CD630_06080 | CD630_n00390 | 156 – 182 | 177 – 205 | -15.0335 | 3.82315 | 4.83391 |
| CD630_16670 | CD630_n00390 | 122 – 137 | 342 – 358 | -14.758 | 3.33553 | 4.83391 |
| CD630_32700 | CD630_n00390 | 180 – 194 | 192 – 206 | -14.5344 | 4.74276 | 4.83391 |
| CD630_21460 | CD630_n00390 | 72 – 79 | 349 – 356 | -14.4496 | 4.83005 | 4.83391 |
| CD630_21620 | CD630_n00390 | 4 – 38 | 110 – 141 | -14.412 | 3.02649 | 4.83391 |
| mobA | CD630_n00390 | 181 – 193 | 346 – 359 | -14.153 | 4.25703 | 4.83391 |
| CD630_32710 | CD630_n00390 | 150 – 188 | 174 – 206 | -14.1222 | 4.74276 | 4.83391 |
| pilD | CD630_n00390 | 3 – 31 | 104 – 131 | -14.109 | 4.54123 | 4.83391 |
| CD630_05821 | CD630_n00390 | 79 – 101 | 340 – 358 | -13.8995 | 3.75661 | 4.83391 |
| CD630_35430 | CD630_n00390 | 142 – 182 | 114 – 150 | -13.8354 | 4.28798 | 4.83391 |
| CD630_21260 | CD630_n00390 | 138 – 152 | 37 – 51 | -13.8314 | 4.22255 | 4.83391 |
| CD630_05290 | CD630_n00390 | 19 – 63 | 106 – 154 | -13.7324 | 3.97604 | 4.83391 |
| CD630_24770 | CD630_n00390 | 87 – 100 | 342 – 356 | -13.6982 | 3.20798 | 4.83391 |
| CD630_26680 | CD630_n00390 | 75 – 104 | 187 – 216 | -13.617 | 3.53956 | 4.83391 |
| scrR | CD630_n00390 | 134 – 148 | 344 – 358 | -13.5751 | 3.49166 | 4.83391 |
| CD630_36250 | CD630_n00390 | 170 – 199 | 179 – 206 | -13.5125 | 4.13224 | 4.83391 |
| feoB | CD630_n00390 | 131 – 197 | 80 – 141 | -13.454 | 4.05364 | 4.83391 |
| CD630_12320 | CD630_n00390 | 37 – 85 | 303 – 348 | -13.3256 | 3.28667 | 4.83391 |
| CD630_03420 | CD630_n00390 | 138 – 175 | 106 – 147 | -13.263 | 4.71868 | 4.83391 |
| CD630_19790 | CD630_n00390 | 85 – 125 | 174 – 209 | -13.1999 | 3.79593 | 4.83391 |
| CD630_29250 | CD630_n00390 | 102 – 159 | 106 – 158 | -13.0495 | 4.61901 | 4.83391 |
| CD630_09410 | CD630_n00390 | 102 – 159 | 106 – 158 | -13.0495 | 4.75651 | 4.83391 |
| CD630_22200 | CD630_n00390 | 141 – 173 | 108 – 141 | -13.0444 | 4.18288 | 4.83391 |
| CD630_28150 | CD630_n00390 | 25 – 36 | 342 – 353 | -13.0333 | 4.18447 | 4.83391 |
| CD630_21770 | CD630_n00390 | 87 – 196 | 104 – 205 | -12.9463 | 4.02587 | 4.83391 |
| cdu2 | CD630_n00390 | 66 – 105 | 171 – 206 | -12.942 | 2.97335 | 4.83391 |
| CD630_14150 | CD630_n00390 | 134 – 184 | 276 – 320 | -12.8821 | 3.28251 | 4.83391 |
| acoL | CD630_n00390 | 104 – 135 | 348 – 379 | -12.8395 | 3.32169 | 4.83391 |
| bcp | CD630_n00390 | 44 – 97 | 102 – 157 | -12.8175 | 3.16670 | 4.83391 |
| sspA | CD630_n00390 | 53 – 109 | 88 – 149 | -12.7506 | 3.62689 | 4.83391 |
| terD | CD630_n00390 | 128 – 152 | 304 – 329 | -12.7403 | 3.33148 | 4.83391 |
| CD630_25190 | CD630_n00390 | 91 – 139 | 104 – 157 | -12.7137 | 3.19663 | 4.83391 |
| CD630_19000 | CD630_n00390 | 83 – 116 | 177 – 209 | -12.6884 | 4.18443 | 4.83391 |
| CD630_20170 | CD630_n00390 | 123 – 164 | 106 – 142 | -12.6396 | 6.90201 | 4.83391 |
| etfA | CD630_n00390 | 24 – 66 | 114 – 157 | -12.634 | 3.41204 | 4.83391 |
| CD630_30140 | CD630_n00390 | 70 – 110 | 107 – 151 | -12.629 | 3.28343 | 4.83391 |
| CD630_34530 | CD630_n00390 | 2 – 50 | 108 – 158 | -12.6264 | 3.83016 | 4.83391 |
| CD630_05280 | CD630_n00390 | 153 – 186 | 173 – 206 | -12.62 | 3.91665 | 4.83391 |
| CD630_18960 | CD630_n00390 | 135 – 153 | 136 – 154 | -12.5983 | 4.37822 | 4.83391 |
| CD630_28181 | CD630_n00390 | 8 – 78 | 82 – 143 | -12.5499 | 3.91299 | 4.83391 |
| CD630_06200 | CD630_n00390 | 118 – 150 | 104 – 144 | -12.5442 | 3.57064 | 4.83391 |
| thiH | CD630_n00390 | 186 – 196 | 264 – 274 | -12.494 | 3.58230 | 4.83391 |
| CD630_03040 | CD630_n00390 | 51 – 95 | 105 – 152 | -12.4872 | 3.51403 | 4.83391 |
| CD630_23780 | CD630_n00390 | 90 – 101 | 342 – 353 | -12.4225 | 3.91629 | 4.83391 |
| ung | CD630_n00390 | 1 – 38 | 102 – 141 | -12.3865 | 3.30908 | 4.83391 |
| CD630_27110 | CD630_n00390 | 133 – 178 | 99 – 144 | -12.3697 | 3.78231 | 4.83391 |
| CD630_28550 | CD630_n00390 | 90 – 100 | 343 – 353 | -12.2128 | 3.51196 | 4.83391 |
| CD630_20670 | CD630_n00390 | 74 – 112 | 177 – 219 | -12.1971 | 4.42757 | 4.83391 |
| CD630_22991 | CD630_n00390 | 178 – 196 | 357 – 375 | -12.1588 | 6.03728 | 4.83391 |
| CD630_18950 | CD630_n00390 | 180 – 194 | 79 – 92 | -12.134 | 3.40610 | 4.83391 |
| CD630_30100 | CD630_n00390 | 54 – 80 | 175 – 202 | -12.0949 | 4.13234 | 4.83391 |
| CD630_01590 | CD630_n00390 | 155 – 201 | 177 – 225 | -12.0931 | 3.45609 | 4.83391 |
| ldh | CD630_n00390 | 130 – 171 | 2 – 47 | -12.0897 | 4.35818 | 4.83391 |
| CD630_16460 | CD630_n00390 | 15 – 86 | 165 – 244 | -12.0086 | 4.64028 | 4.83391 |
| CD630_21090 | CD630_n00390 | 131 – 188 | 105 – 154 | -11.9649 | 4.15803 | 4.83391 |
| CD630_35260 | CD630_n00390 | 69 – 130 | 102 – 157 | -11.8353 | 5.77277 | 4.83391 |
| CD630_33480 | CD630_n00390 | 89 – 110 | 181 – 206 | -11.8234 | 3.67613 | 4.83391 |
| bglA | CD630_n00390 | 59 – 113 | 177 – 240 | -11.8093 | 4.50315 | 4.83391 |
| metQ | CD630_n00390 | 87 – 116 | 176 – 206 | -11.7369 | 3.87190 | 4.83391 |
| CD630_13780 | CD630_n00390 | 130 – 176 | 174 – 209 | -11.7237 | 3.52274 | 4.83391 |
| CD630_32050 | CD630_n00390 | 176 – 199 | 184 – 206 | -11.716 | 5.73797 | 4.83391 |
| CD630_07870 | CD630_n00390 | 170 – 179 | 370 – 379 | -11.6675 | 3.22829 | 4.83391 |
| CD630_19690 | CD630_n00390 | 115 – 154 | 107 – 157 | -11.6162 | 3.85861 | 4.83391 |
| CD630_31010 | CD630_n00390 | 41 – 98 | 102 – 157 | -11.6009 | 4.96305 | 4.83391 |
| CD630_09772 | CD630_n00390 | 39 – 96 | 104 – 158 | -11.5683 | 3.66033 | 4.83391 |
| CD630_31461 | CD630_n00390 | 6 – 25 | 132 – 149 | -11.5253 | 4.41476 | 4.83391 |
| CD630_09040 | CD630_n00390 | 91 – 139 | 104 – 157 | -11.5059 | 4.11851 | 4.83391 |
| CD630_22030 | CD630_n00390 | 7 – 23 | 44 – 60 | -11.4811 | 3.29934 | 4.83391 |
| CD630_21070 | CD630_n00390 | 185 – 196 | 344 – 355 | -11.476 | 5.35019 | 4.83391 |
| CD630_27120 | CD630_n00390 | 113 – 161 | 266 – 316 | -11.4634 | 4.43720 | 4.83391 |
| CD630_15850 | CD630_n00390 | 11 – 47 | 170 – 206 | -11.4535 | 4.32968 | 4.83391 |
| CD630_33560 | CD630_n00390 | 57 – 94 | 106 – 142 | -11.4321 | 4.03147 | 4.83391 |
| CD630_15431 | CD630_n00390 | 125 – 185 | 102 – 157 | -11.4213 | 5.29780 | 4.83391 |
| CD630_34030 | CD630_n00390 | 42 – 66 | 127 – 154 | -11.4158 | 3.05766 | 4.83391 |
| CD630_14250 | CD630_n00390 | 184 – 199 | 190 – 206 | -11.4081 | 4.19604 | 4.83391 |
| dnaC | CD630_n00390 | 145 – 175 | 178 – 206 | -11.4036 | 6.30796 | 4.83391 |
| CD630_22050 | CD630_n00390 | 5 – 96 | 107 – 186 | -11.4008 | 4.87639 | 4.83391 |
| CD630_20171 | CD630_n00390 | 94 – 168 | 105 – 156 | -11.391 | 6.87491 | 4.83391 |
| CD630_29960 | CD630_n00390 | 135 – 201 | 65 – 138 | -11.3727 | 4.51904 | 4.83391 |
| CD630_18311 | CD630_n00390 | 88 – 113 | 174 – 203 | -11.3695 | 6.86047 | 4.83391 |
| CD630_22880 | CD630_n00390 | 175 – 199 | 125 – 150 | -11.3654 | 3.62455 | 4.83391 |
| CD630_20660 | CD630_n00390 | 134 – 167 | 108 – 150 | -11.3333 | 5.39321 | 4.83391 |
| cobC | CD630_n00390 | 164 – 198 | 104 – 141 | -11.3263 | 3.15854 | 4.83391 |
| CD630_20260 | CD630_n00390 | 86 – 114 | 176 – 205 | -11.3113 | 3.84162 | 4.83391 |
| CD630_24590 | CD630_n00390 | 143 – 150 | 348 – 355 | -11.3034 | 3.70401 | 4.83391 |
| CD630_22850 | CD630_n00390 | 167 – 201 | 105 – 149 | -11.2843 | 3.84102 | 4.83391 |
| vncR | CD630_n00390 | 138 – 157 | 183 – 206 | -11.282 | 3.93642 | 4.83391 |
| CD630_32990 | CD630_n00390 | 35 – 68 | 211 – 254 | -11.2777 | 5.24925 | 4.83391 |
| CD630_16950 | CD630_n00390 | 185 – 196 | 81 – 93 | -11.203 | 4.42569 | 4.83391 |
| cwp29 | CD630_n00390 | 20 – 56 | 178 – 218 | -11.1349 | 4.40865 | 4.83391 |
| glnA | CD630_n00390 | 151 – 187 | 108 – 147 | -11.1302 | 3.87629 | 4.83391 |
| CD630_13400 | CD630_n00390 | 67 – 118 | 105 – 150 | -11.1024 | 3.20232 | 4.83391 |
| CD630_09970 | CD630_n00390 | 3 – 39 | 103 – 140 | -11.0906 | 3.25917 | 4.83391 |
| tcdC | CD630_n00390 | 112 – 158 | 274 – 314 | -11.0872 | 3.76101 | 4.83391 |
| CD630_15150 | CD630_n00390 | 8 – 52 | 103 – 156 | -11.0672 | 3.35564 | 4.83391 |
| cprK | CD630_n00390 | 47 – 56 | 82 – 91 | -11.0458 | 3.96216 | 4.83391 |
| CD630_04610 | CD630_n00390 | 48 – 71 | 108 – 139 | -11.0077 | 3.83025 | 4.83391 |
| CD630_15900 | CD630_n00390 | 112 – 127 | 371 – 386 | -10.9889 | 3.17664 | 4.83391 |
| pilT | CD630_n00390 | 8 – 26 | 297 – 316 | -10.9591 | 3.24639 | 4.83391 |
| CD630_06690 | CD630_n00390 | 86 – 96 | 343 – 353 | -10.8997 | 5.82709 | 4.83391 |
| CD630_06060 | CD630_n00390 | 102 – 139 | 177 – 207 | -10.8985 | 4.94050 | 4.83391 |
| CD630_10830 | CD630_n00390 | 86 – 95 | 345 – 355 | -10.8974 | 4.82398 | 4.83391 |
| cdu1 | CD630_n00390 | 87 – 96 | 345 – 355 | -10.8787 | 4.46276 | 4.83391 |
| CD630_06150 | CD630_n00390 | 99 – 128 | 103 – 124 | -10.8707 | 5.51744 | 4.83391 |
| CD630_18090 | CD630_n00390 | 10 – 32 | 184 – 205 | -10.8666 | 4.23961 | 4.83391 |
| CD630_13350 | CD630_n00390 | 163 – 181 | 127 – 142 | -10.865 | 4.15139 | 4.83391 |
| CD630_23440 | CD630_n00390 | 1 – 24 | 169 – 206 | -10.8576 | 3.34841 | 4.83391 |
| CD630_24760 | CD630_n00390 | 98 – 143 | 104 – 154 | -10.808 | 5.19485 | 4.83391 |
| CD630_01690 | CD630_n00390 | 14 – 45 | 113 – 144 | -10.8061 | 4.97234 | 4.83391 |
| CD630_31910 | CD630_n00390 | 86 – 105 | 186 – 205 | -10.793 | 3.48314 | 4.83391 |
| CD630_17710 | CD630_n00390 | 133 – 152 | 188 – 207 | -10.7904 | 4.37688 | 4.83391 |
| CD630_17730 | CD630_n00390 | 76 – 98 | 104 – 123 | -10.7731 | 4.73397 | 4.83391 |
| CD630_09100 | CD630_n00390 | 2 – 50 | 20 – 61 | -10.7656 | 5.92260 | 4.83391 |
| CD630_19801 | CD630_n00390 | 88 – 98 | 343 – 353 | -10.7551 | 5.22035 | 4.83391 |
| CD630_19401 | CD630_n00390 | 35 – 82 | 118 – 165 | -10.7356 | 4.37084 | 4.83391 |
| gabT | CD630_n00390 | 36 – 61 | 211 – 239 | -10.7352 | 3.67067 | 4.83391 |
| opuCA | CD630_n00390 | 87 – 112 | 175 – 206 | -10.7343 | 4.07035 | 4.83391 |
| CD630_23760 | CD630_n00390 | 82 – 105 | 186 – 209 | -10.7317 | 4.54307 | 4.83391 |
| CD630_02430 | CD630_n00390 | 39 – 84 | 110 – 138 | -10.6998 | 3.71048 | 4.83391 |
| CD630_01560 | CD630_n00390 | 142 – 151 | 369 – 378 | -10.6866 | 4.06352 | 4.83391 |
| CD630_30540 | CD630_n00390 | 119 – 136 | 299 – 316 | -10.6845 | 3.63804 | 4.83391 |
| pgm | CD630_n00390 | 146 – 176 | 177 – 209 | -10.6792 | 3.45439 | 4.83391 |
| CD630_21490 | CD630_n00390 | 127 – 193 | 263 – 329 | -10.6757 | 3.37462 | 4.83391 |
| CD630_22890 | CD630_n00390 | 40 – 47 | 346 – 353 | -10.6749 | 3.75591 | 4.83391 |
| CD630_25040 | CD630_n00390 | 144 – 177 | 19 – 54 | -10.6491 | 3.43898 | 4.83391 |
| CD630_06060 | CD630_n00390 | 41 – 59 | 342 – 355 | -10.6488 | 4.94050 | 4.83391 |
| CD630_25310 | CD630_n00390 | 160 – 176 | 294 – 310 | -10.6269 | 3.99861 | 4.83391 |
| CD630_23820 | CD630_n00390 | 88 – 97 | 345 – 354 | -10.6264 | 3.18419 | 4.83391 |
| CD630_27610 | CD630_n00390 | 5 – 52 | 271 – 316 | -10.5555 | 3.96465 | 4.83391 |
| CD630_31880 | CD630_n00390 | 15 – 34 | 15 – 35 | -10.547 | 3.97522 | 4.83391 |
| CD630_34080 | CD630_n00390 | 168 – 201 | 100 – 141 | -10.536 | 3.20224 | 4.83391 |
| CD630_30120 | CD630_n00390 | 43 – 100 | 104 – 157 | -10.5294 | 4.04859 | 4.83391 |
| CD630_23770 | CD630_n00390 | 52 – 93 | 108 – 151 | -10.5083 | 3.25181 | 4.83391 |
| CD630_10050 | CD630_n00390 | 44 – 159 | 107 – 202 | -10.5033 | 3.93750 | 4.83391 |
| CD630_11730 | CD630_n00390 | 36 – 55 | 298 – 317 | -10.5021 | 4.80521 | 4.83391 |
| CD630_27310 | CD630_n00390 | 135 – 177 | 101 – 141 | -10.4939 | 4.01622 | 4.83391 |
| CD630_08940 | CD630_n00390 | 75 – 91 | 67 – 83 | -10.47 | 3.72225 | 4.83391 |
| CD630_06110 | CD630_n00390 | 124 – 138 | 304 – 318 | -10.4566 | 5.61380 | 4.83391 |
| CD630_08090 | CD630_n00390 | 143 – 174 | 102 – 132 | -10.4487 | 3.57167 | 4.83391 |
| CD630_32520 | CD630_n00390 | 153 – 198 | 272 – 323 | -10.4458 | 4.06098 | 4.83391 |
| CD630_18010 | CD630_n00390 | 154 – 186 | 125 – 156 | -10.4453 | 4.21967 | 4.83391 |
| CD630_21760 | CD630_n00390 | 64 – 74 | 82 – 91 | -10.4184 | 4.43846 | 4.83391 |
| CD630_09760 | CD630_n00390 | 90 – 125 | 173 – 206 | -10.3721 | 4.83410 | 4.83391 |
| CD630_28910 | CD630_n00390 | 90 – 125 | 173 – 206 | -10.3721 | 4.51664 | 4.83391 |
| CD630_08510 | CD630_n00390 | 44 – 52 | 343 – 351 | -10.3331 | 5.16329 | 4.83391 |
| CD630_08390 | CD630_n00390 | 83 – 92 | 343 – 352 | -10.3259 | 5.16000 | 4.83391 |
| CD630_05230 | CD630_n00390 | 3 – 15 | 344 – 356 | -10.3237 | 3.73787 | 4.83391 |
| CD630_24591 | CD630_n00390 | 88 – 96 | 345 – 353 | -10.3068 | 3.54497 | 4.83391 |
| CD630_15060 | CD630_n00390 | 173 – 191 | 220 – 237 | -10.2936 | 3.69396 | 4.83391 |
| CD630_33190 | CD630_n00390 | 115 – 126 | 177 – 188 | -10.2735 | 4.13899 | 4.83391 |
| opuCC | CD630_n00390 | 95 – 144 | 108 – 157 | -10.2683 | 4.07035 | 4.83391 |
| CD630_33300 | CD630_n00390 | 190 – 197 | 349 – 356 | -10.2451 | 3.79149 | 4.83391 |
| cwp26 | CD630_n00390 | 131 – 161 | 135 – 165 | -10.2308 | 4.44377 | 4.83391 |
| CD630_06340 | CD630_n00390 | 3 – 45 | 263 – 323 | -10.22 | 4.35983 | 4.83391 |
| CD630_15460 | CD630_n00390 | 88 – 97 | 345 – 355 | -10.205 | 6.86296 | 4.83391 |
| CD630_18400 | CD630_n00390 | 121 – 158 | 105 – 142 | -10.2032 | 4.67962 | 4.83391 |
| tcdA | CD630_n00390 | 36 – 63 | 175 – 205 | -10.1776 | 4.70593 | 4.83391 |
| CD630_06190 | CD630_n00390 | 141 – 151 | 41 – 51 | -10.1713 | 3.57064 | 4.83391 |
| nusB | CD630_n00390 | 88 – 157 | 281 – 353 | -10.1645 | 4.92798 | 4.83391 |
| rolA | CD630_n00390 | 33 – 56 | 300 – 323 | -10.1413 | 4.56362 | 4.83391 |
| CD630_06350 | CD630_n00390 | 138 – 167 | 177 – 206 | -10.0352 | 4.35983 | 4.83391 |
| CD630_13360 | CD630_n00390 | 46 – 99 | 104 – 151 | -10.0309 | 3.77330 | 4.83391 |
| CD630_14050 | CD630_n00390 | 186 – 201 | 191 – 206 | -10.0271 | 3.30894 | 4.83391 |
| CD630_32040 | CD630_n00390 | 24 – 42 | 250 – 274 | -10.0208 | 4.93516 | 4.83391 |
| CD630_18910 | CD630_n00390 | 121 – 136 | 342 – 357 | -10.0137 | 3.30688 | 4.83391 |
| garK | CD630_n00390 | 154 – 160 | 350 – 356 | -10.0123 | 3.72836 | 4.83391 |
| CD630_26050 | CD630_n00390 | 173 – 200 | 176 – 205 | -10.0063 | 6.99851 | 4.83391 |
| CD630_36690 | CD630_n00390 | 111 – 133 | 292 – 314 | -9.99923 | 3.53761 | 4.83391 |
| srlEb | CD630_n00390 | 91 – 111 | 187 – 206 | -9.9874 | 3.50140 | 4.83391 |
| cynT | CD630_n00390 | 146 – 169 | 101 – 120 | -9.92724 | 4.21370 | 4.83391 |
| cbiQ | CD630_n00390 | 36 – 58 | 76 – 93 | -9.91539 | 3.79830 | 4.83391 |
| CD630_28320 | CD630_n00390 | 1 – 10 | 342 – 351 | -9.89861 | 3.99184 | 4.83391 |
| cspC | CD630_n00390 | 79 – 89 | 265 – 275 | -9.89641 | 5.48848 | 4.83391 |
| CD630_18970 | CD630_n00390 | 145 – 185 | 276 – 314 | -9.89341 | 3.57243 | 4.83391 |
| CD630_22510 | CD630_n00390 | 114 – 127 | 218 – 231 | -9.88346 | 11.42763 | 4.83391 |
| etfA | CD630_n00390 | 124 – 163 | 172 – 205 | -9.8759 | 4.80521 | 4.83391 |
| CD630_29500 | CD630_n00390 | 47 – 69 | 196 – 219 | -9.8505 | 3.70644 | 4.83391 |
| CD630_31500 | CD630_n00390 | 19 – 37 | 300 – 317 | -9.83562 | 4.90207 | 4.83391 |
| pilC | CD630_n00390 | 3 – 25 | 300 – 322 | -9.80312 | 3.54230 | 4.83391 |
| CD630_06100 | CD630_n00390 | 65 – 74 | 374 – 383 | -9.7632 | 4.43918 | 4.83391 |
| bioB | CD630_n00390 | 131 – 186 | 264 – 317 | -9.75972 | 3.59022 | 4.83391 |

| | | | | | | |
|---|---|---|---|---|---|---|
| CD630_24920 | CD630_n00390 | 89 -- 150 | 105 -- 157 | -9.75926 | 3.89455 | 4.83391 |
| CD630_21030 | CD630_n00390 | 57 -- 107 | 107 -- 156 | -9.75456 | 3.08733 | 4.83391 |
| CD630_11220 | CD630_n00390 | 79 -- 105 | 118 -- 142 | -9.70302 | 3.52327 | 4.83391 |
| srlD | CD630_n00390 | 101 -- 152 | 182 -- 237 | -9.69707 | 4.19843 | 4.83391 |
| CD630_23070 | CD630_n00390 | 5 -- 16 | 23 -- 34 | -9.6858 | 5.23027 | 4.83391 |
| CD630_20980 | CD630_n00390 | 101 -- 110 | 80 -- 89 | -9.67716 | 8.12847 | 4.83391 |
| CD630_27570 | CD630_n00390 | 128 -- 141 | 106 -- 119 | -9.67132 | 3.46216 | 4.83391 |
| CD630_22500 | CD630_n00390 | 194 -- 201 | 345 -- 352 | -9.67012 | 4.03688 | 4.83391 |
| CD630_05950 | CD630_n00390 | 34 -- 69 | 173 -- 205 | -9.66918 | 3.31669 | 4.83391 |
| CD630_27200 | CD630_n00390 | 1 -- 36 | 271 -- 295 | -9.6675 | 3.42516 | 4.83391 |
| CD630_08100 | CD630_n00390 | 102 -- 165 | 264 -- 331 | -9.66342 | 4.39181 | 4.83391 |
| CD630_25320 | CD630_n00390 | 144 -- 173 | 177 -- 206 | -9.65909 | 4.29415 | 4.83391 |
| CD630_31220 | CD630_n00390 | 5 -- 21 | 293 -- 309 | -9.63797 | 3.73779 | 4.83391 |
| CD630_27280 | CD630_n00390 | 146 -- 156 | 82 -- 93 | -9.62835 | 3.24738 | 4.83391 |
| CD630_14600 | CD630_n00390 | 65 -- 101 | 119 -- 156 | -9.62582 | 5.90348 | 4.83391 |
| CD630_14180 | CD630_n00390 | 87 -- 95 | 345 -- 353 | -9.62082 | 3.93696 | 4.83391 |
| CD630_34520 | CD630_n00390 | 77 -- 113 | 107 -- 139 | -9.61507 | 3.83016 | 4.83391 |
| CD630_16450 | CD630_n00390 | 149 -- 163 | 80 -- 93 | -9.60238 | 4.57547 | 4.83391 |
| CD630_30130 | CD630_n00390 | 2 -- 28 | 175 -- 209 | -9.59855 | 3.28343 | 4.83391 |
| CD630_28580 | CD630_n00390 | 48 -- 61 | 340 -- 353 | -9.59421 | 4.67130 | 4.83391 |
| CD630_09060 | CD630_n00390 | 99 -- 134 | 193 -- 220 | -9.59296 | 4.31834 | 4.83391 |
| CD630_04410 | CD630_n00390 | 83 -- 96 | 345 -- 356 | -9.59023 | 3.90451 | 4.83391 |
| CD630_16150 | CD630_n00390 | 81 -- 89 | 371 -- 379 | -9.57856 | 4.20614 | 4.83391 |
| gcvTPA | CD630_n00390 | 170 -- 178 | 82 -- 90 | -9.5768 | 4.77229 | 4.83391 |
| CD630_09840 | CD630_n00390 | 162 -- 177 | 37 -- 51 | -9.5399 | 4.50009 | 4.83391 |
| adhE | CD630_n00390 | 51 -- 80 | 110 -- 142 | -9.5252 | 3.06485 | 4.83391 |
| CD630_27230 | CD630_n00390 | 84 -- 92 | 371 -- 379 | -9.51497 | 3.52462 | 4.83391 |
| CD630_05880 | CD630_n00390 | 91 -- 109 | 190 -- 206 | -9.5118 | 4.28339 | 4.83391 |
| CD630_13530 | CD630_n00470 | 4 -- 104 | 9 -- 90 | -16.7239 | 4.09832 | 3.52267 |
| CD630_28290 | CD630_n00470 | 28 -- 57 | 1 -- 29 | -15.7331 | 5.54192 | 3.52267 |
| CD630_08950 | CD630_n00470 | 8 -- 65 | 22 -- 81 | -14.8255 | 3.71905 | 3.52267 |
| CD630_05830 | CD630_n00470 | 134 -- 187 | 47 -- 97 | -14.7005 | 3.35360 | 3.52267 |
| dltB | CD630_n00470 | 133 -- 176 | 10 -- 53 | -14.5465 | 3.81485 | 3.52267 |
| CD630_14610 | CD630_n00470 | 138 -- 171 | 38 -- 72 | -14.3954 | 4.89428 | 3.52267 |
| CD630_15160 | CD630_n00470 | 3 -- 19 | 11 -- 27 | -13.9498 | 4.32318 | 3.52267 |
| CD630_06540 | CD630_n00470 | 46 -- 118 | 8 -- 82 | -13.3539 | 3.54673 | 3.52267 |
| CD630_04610 | CD630_n00470 | 157 -- 200 | 49 -- 85 | -13.1934 | 3.83025 | 3.52267 |
| CD630_32060 | CD630_n00470 | 162 -- 201 | 18 -- 62 | -13.0403 | 5.73797 | 3.52267 |
| CD630_03930 | CD630_n00470 | 136 -- 156 | 9 -- 28 | -12.9102 | 4.15934 | 3.52267 |
| cdtR | CD630_n00470 | 29 -- 80 | 3 -- 57 | -12.7595 | 5.84368 | 3.52267 |
| CD630_24050 | CD630_n00470 | 133 -- 160 | 52 -- 79 | -12.4058 | 3.70192 | 3.52267 |
| CD630_05940 | CD630_n00470 | 133 -- 199 | 9 -- 85 | -12.2749 | 4.18595 | 3.52267 |
| CD630_13350 | CD630_n00470 | 119 -- 146 | 1 -- 30 | -12.1212 | 4.15139 | 3.52267 |
| CD630_26250 | CD630_n00470 | 65 -- 167 | 8 -- 89 | -11.686 | 5.03379 | 3.52267 |
| CD630_20660 | CD630_n00470 | 132 -- 149 | 10 -- 27 | -11.6728 | 5.39321 | 3.52267 |
| CD630_09100 | CD630_n00470 | 15 -- 41 | 50 -- 81 | -11.5452 | 5.92260 | 3.52267 |
| CD630_20670 | CD630_n00470 | 150 -- 199 | 33 -- 86 | -11.2941 | 4.42757 | 3.52267 |
| blaI | CD630_n00470 | 160 -- 169 | 50 -- 59 | -11.0971 | 3.73674 | 3.52267 |
| CD630_12320 | CD630_n00470 | 166 -- 174 | 51 -- 59 | -10.8456 | 3.28667 | 3.52267 |
| CD630_27630 | CD630_n00470 | 140 -- 169 | 9 -- 41 | -10.6572 | 3.54199 | 3.52267 |
| CD630_07960 | CD630_n00470 | 23 -- 44 | 1 -- 24 | -10.6244 | 3.22537 | 3.52267 |
| CD630_14410 | CD630_n00470 | 6 -- 43 | 26 -- 62 | -10.5794 | 3.39433 | 3.52267 |
| CD630_27280 | CD630_n00470 | 170 -- 180 | 51 -- 62 | -10.4053 | 3.24738 | 3.52267 |
| CD630_23780 | CD630_n00470 | 140 -- 201 | 15 -- 74 | -10.2906 | 3.91629 | 3.52267 |
| oppB | CD630_n00470 | 138 -- 175 | 47 -- 90 | -10.2446 | 2.98328 | 3.52267 |
| CD630_18781 | CD630_n00470 | 85 -- 104 | 21 -- 40 | -10.2306 | 3.77660 | 3.52267 |
| CD630_34080 | CD630_n00470 | 153 -- 201 | 25 -- 56 | -10.0913 | 3.20224 | 3.52267 |
| CD630_14210 | CD630_n00470 | 45 -- 59 | 2 -- 16 | -9.92791 | 3.82355 | 3.52267 |
| CD630_13400 | CD630_n00470 | 82 -- 151 | 8 -- 69 | -9.85407 | 3.20232 | 3.52267 |
| CD630_33650 | CD630_n00470 | 4 -- 16 | 74 -- 86 | -9.81093 | 4.80565 | 3.52267 |
| CD630_14710 | CD630_n00470 | 58 -- 75 | 47 -- 71 | -9.69052 | 3.65171 | 3.52267 |
| CD630_32500 | CD630_n00470 | 1 -- 39 | 11 -- 41 | -9.64825 | 6.17288 | 3.52267 |
| CD630_06140 | CD630_n00470 | 8 -- 34 | 68 -- 93 | -9.58691 | 4.12519 | 3.52267 |
| CD630_14140 | CD630_n00470 | 115 -- 163 | 4 -- 59 | -9.55997 | 3.16551 | 3.52267 |
| CD630_24770 | CD630_n00500 | 87 -- 145 | 50 -- 101 | -18.924 | 3.20798 | 9.35359 |
| CD630_09771 | CD630_n00500 | 188 -- 201 | 88 -- 101 | -17.6587 | 4.95362 | 9.35359 |
| CD630_28890 | CD630_n00500 | 188 -- 201 | 88 -- 101 | -17.6587 | 7.52469 | 9.35359 |
| CD630_23780 | CD630_n00500 | 90 -- 131 | 52 -- 98 | -16.524 | 3.91629 | 9.35359 |
| CD630_36250 | CD630_n00500 | 53 -- 117 | 45 -- 100 | -15.0215 | 4.13224 | 9.35359 |
| CD630_16670 | CD630_n00500 | 124 -- 138 | 86 -- 101 | -14.8545 | 3.33553 | 9.35359 |
| CD630_06060 | CD630_n00500 | 41 -- 61 | 85 -- 100 | -14.6211 | 4.94050 | 9.35359 |
| srlD | CD630_n00500 | 109 -- 150 | 53 -- 95 | -14.3763 | 4.19843 | 9.35359 |
| CD630_21460 | CD630_n00500 | 72 -- 79 | 94 -- 101 | -14.323 | 4.83005 | 9.35359 |
| CD630_14990 | CD630_n00500 | 148 -- 198 | 47 -- 100 | -13.7836 | 3.78619 | 9.35359 |
| CD630_02790 | CD630_n00500 | 127 -- 135 | 93 -- 101 | -13.2426 | 4.77661 | 9.35359 |
| fliI | CD630_n00500 | 33 -- 52 | 41 -- 60 | -13.1933 | 3.03666 | 9.35359 |
| CD630_28150 | CD630_n00500 | 25 -- 43 | 77 -- 98 | -13.0949 | 4.18447 | 9.35359 |
| scrR | CD630_n00500 | 136 -- 148 | 89 -- 101 | -13.0722 | 3.49166 | 9.35359 |
| ldh | CD630_n00500 | 85 -- 97 | 88 -- 100 | -12.8627 | 4.35818 | 9.35359 |
| CD630_28550 | CD630_n00500 | 90 -- 100 | 88 -- 98 | -12.7823 | 3.51196 | 9.35359 |
| CD630_23440 | CD630_n00500 | 62 -- 89 | 74 -- 98 | -12.4058 | 3.34841 | 9.35359 |
| CD630_13920 | CD630_n00500 | 135 -- 152 | 6 -- 22 | -12.4043 | 3.25236 | 9.35359 |
| tcdA | CD630_n00500 | 33 -- 53 | 44 -- 65 | -12.3199 | 4.70593 | 9.35359 |
| mobA | CD630_n00500 | 184 -- 193 | 91 -- 101 | -12.2932 | 4.25703 | 9.35359 |
| CD630_05230 | CD630_n00500 | 3 -- 40 | 43 -- 101 | -12.2563 | 3.73787 | 9.35359 |
| CD630_21070 | CD630_n00500 | 185 -- 196 | 89 -- 100 | -12.2496 | 5.35019 | 9.35359 |
| CD630_24590 | CD630_n00500 | 143 -- 150 | 93 -- 100 | -12.0806 | 3.70401 | 9.35359 |
| oppB | CD630_n00500 | 44 -- 76 | 1 -- 34 | -12.0005 | 2.98328 | 9.35359 |
| CD630_06960 | CD630_n00500 | 65 -- 142 | 15 -- 99 | -11.7204 | 3.16317 | 9.35359 |
| CD630_11220 | CD630_n00500 | 86 -- 98 | 86 -- 99 | -11.6353 | 3.52327 | 9.35359 |
| cdu1 | CD630_n00500 | 87 -- 96 | 90 -- 100 | -11.6039 | 4.46276 | 9.35359 |
| CD630_10830 | CD630_n00500 | 86 -- 95 | 90 -- 100 | -11.5592 | 4.82398 | 9.35359 |
| CD630_19690 | CD630_n00500 | 117 -- 129 | 88 -- 100 | -11.4671 | 3.85861 | 9.35359 |
| CD630_18910 | CD630_n00500 | 122 -- 137 | 86 -- 101 | -11.4063 | 3.30688 | 9.35359 |
| CD630_32060 | CD630_n00500 | 46 -- 59 | 8 -- 22 | -11.3969 | 5.73797 | 9.35359 |
| CD630_06690 | CD630_n00500 | 86 -- 96 | 88 -- 98 | -11.3736 | 5.82709 | 9.35359 |
| CD630_19801 | CD630_n00500 | 88 -- 98 | 88 -- 98 | -11.3509 | 5.22035 | 9.35359 |
| CD630_23820 | CD630_n00500 | 88 -- 97 | 90 -- 99 | -11.178 | 3.18419 | 9.35359 |
| CD630_08510 | CD630_n00500 | 44 -- 55 | 84 -- 96 | -11.1579 | 5.16329 | 9.35359 |
| CD630_22890 | CD630_n00500 | 40 -- 47 | 91 -- 98 | -11.0874 | 3.75591 | 9.35359 |
| CD630_19942 | CD630_n00500 | 86 -- 129 | 48 -- 100 | -11.0746 | 4.24574 | 9.35359 |
| ogt1 | CD630_n00500 | 2 -- 24 | 42 -- 64 | -11.0403 | 3.16772 | 9.35359 |
| nusB | CD630_n00500 | 88 -- 140 | 45 -- 98 | -11.0131 | 4.92798 | 9.35359 |
| CD630_28181 | CD630_n00500 | 148 -- 190 | 14 -- 52 | -10.9877 | 3.91299 | 9.35359 |
| CD630_08390 | CD630_n00500 | 80 -- 92 | 88 -- 99 | -10.9654 | 5.16000 | 9.35359 |
| CD630_15460 | CD630_n00500 | 88 -- 97 | 90 -- 100 | -10.8733 | 6.86296 | 9.35359 |
| CD630_24591 | CD630_n00500 | 88 -- 96 | 90 -- 98 | -10.8119 | 3.54497 | 9.35359 |
| CD630_06110 | CD630_n00500 | 171 -- 188 | 5 -- 21 | -10.7525 | 5.61380 | 9.35359 |
| acoL | CD630_n00500 | 128 -- 135 | 93 -- 100 | -10.732 | 3.32169 | 9.35359 |
| hfq | CD630_n00500 | 51 -- 66 | 85 -- 101 | -10.7268 | 5.05609 | 9.35359 |
| LinCd | CD630_n00500 | 42 -- 94 | 46 -- 99 | -10.7002 | 4.33624 | 9.35359 |
| CD630_16940 | CD630_n00500 | 130 -- 144 | 2 -- 16 | -10.469 | 4.14665 | 9.35359 |
| def1 | CD630_n00500 | 86 -- 164 | 45 -- 100 | -10.4168 | 3.99720 | 9.35359 |
| CD630_18010 | CD630_n00500 | 94 -- 102 | 90 -- 98 | -10.4083 | 4.21967 | 9.35359 |
| cdu2 | CD630_n00500 | 63 -- 80 | 45 -- 62 | -10.3602 | 2.97335 | 9.35359 |
| CD630_22070 | CD630_n00500 | 44 -- 59 | 42 -- 59 | -10.3195 | 4.09651 | 9.35359 |
| garK | CD630_n00500 | 154 -- 160 | 95 -- 101 | -10.3014 | 3.72836 | 9.35359 |
| CD630_04410 | CD630_n00500 | 83 -- 143 | 45 -- 101 | -10.2579 | 3.90451 | 9.35359 |
| CD630_33300 | CD630_n00500 | 190 -- 197 | 94 -- 101 | -10.2406 | 3.79149 | 9.35359 |
| CD630_04770 | CD630_n00500 | 33 -- 54 | 84 -- 100 | -10.2137 | 3.58478 | 9.35359 |
| CD630_27120 | CD630_n00500 | 189 -- 201 | 87 -- 99 | -10.1942 | 4.43720 | 9.35359 |
| CD630_28320 | CD630_n00500 | 1 -- 10 | 87 -- 96 | -10.1732 | 3.99184 | 9.35359 |
| CD630_25020 | CD630_n00500 | 87 -- 116 | 68 -- 98 | -10.132 | 4.17721 | 9.35359 |
| CD630_05030 | CD630_n00500 | 131 -- 155 | 9 -- 32 | -10.116 | 4.03823 | 9.35359 |
| CD630_13380 | CD630_n00500 | 81 -- 91 | 89 -- 99 | -10.0872 | 4.74408 | 9.35359 |
| CD630_22500 | CD630_n00500 | 194 -- 201 | 90 -- 97 | -10.087 | 4.03688 | 9.35359 |
| srlEb | CD630_n00500 | 85 -- 94 | 91 -- 100 | -10.0786 | 3.50140 | 9.35359 |
| CD630_14180 | CD630_n00500 | 87 -- 95 | 90 -- 98 | -10.0219 | 3.93696 | 9.35359 |
| CD630_04341 | CD630_n00500 | 124 -- 181 | 50 -- 101 | -9.94753 | 4.29262 | 9.35359 |
| CD630_32040 | CD630_n00500 | 19 -- 42 | 1 -- 24 | -9.92422 | 4.93516 | 9.35359 |
| CD630_25310 | CD630_n00500 | 163 -- 175 | 40 -- 52 | -9.92027 | 3.99861 | 9.35359 |
| metQ | CD630_n00500 | 194 -- 200 | 94 -- 100 | -9.83968 | 3.87190 | 9.35359 |
| glgP | CD630_n00500 | 109 -- 115 | 92 -- 98 | -9.81127 | 3.28530 | 9.35359 |
| kdgT1 | CD630_n00500 | 87 -- 128 | 56 -- 98 | -9.71162 | 4.49207 | 9.35359 |
| bglA | CD630_n00500 | 173 -- 194 | 46 -- 67 | -9.68694 | 4.86402 | 9.35359 |
| CD630_14250 | CD630_n00500 | 185 -- 197 | 50 -- 62 | -9.63597 | 4.19604 | 9.35359 |
| CD630_22660 | CD630_n00500 | 128 -- 155 | 45 -- 70 | -9.51991 | 3.60496 | 9.35359 |
| CD630_26150 | CD630_n00590 | 138 -- 171 | 44 -- 75 | -18.4126 | 4.06302 | 4.28395 |
| CD630_06170 | CD630_n00590 | 102 -- 155 | 29 -- 74 | -16.1021 | 4.95366 | 4.28395 |
| CD630_05370 | CD630_n00590 | 119 -- 162 | 23 -- 73 | -15.8282 | 3.26947 | 4.28395 |
| CD630_20960 | CD630_n00590 | 39 -- 105 | 31 -- 101 | -15.2739 | 5.65940 | 4.28395 |
| cdtR | CD630_n00590 | 6 -- 76 | 2 -- 77 | -14.4355 | 5.84368 | 4.28395 |
| CD630_36250 | CD630_n00590 | 127 -- 201 | 16 -- 93 | -14.303 | 4.13224 | 4.28395 |
| CD630_33570 | CD630_n00590 | 107 -- 164 | 27 -- 90 | -13.9969 | 3.95778 | 4.28395 |
| CD630_25150 | CD630_n00590 | 14 -- 53 | 63 -- 100 | -13.764 | 3.79339 | 4.28395 |
| CD630_34050 | CD630_n00590 | 143 -- 171 | 69 -- 99 | -13.7496 | 4.03863 | 4.28395 |
| CD630_14010 | CD630_n00590 | 129 -- 198 | 3 -- 79 | -12.9126 | 3.69983 | 4.28395 |
| CD630_19903 | CD630_n00590 | 29 -- 73 | 26 -- 74 | -12.5452 | 5.79994 | 4.28395 |
| CD630_27570 | CD630_n00590 | 2 -- 23 | 52 -- 74 | -12.3307 | 3.46216 | 4.28395 |
| CD630_31490 | CD630_n00590 | 130 -- 163 | 67 -- 100 | -12.101 | 4.08593 | 4.28395 |
| CD630_04770 | CD630_n00590 | 41 -- 67 | 44 -- 76 | -11.8217 | 3.58478 | 4.28395 |
| CD630_18910 | CD630_n00590 | 15 -- 51 | 46 -- 83 | -11.7816 | 3.30688 | 4.28395 |
| CD630_27280 | CD630_n00590 | 130 -- 196 | 31 -- 87 | -11.6939 | 3.24738 | 4.28395 |
| CD630_06060 | CD630_n00590 | 18 -- 39 | 26 -- 47 | -11.6488 | 4.94050 | 4.28395 |
| opuCC | CD630_n00590 | 178 -- 201 | 59 -- 80 | -11.6061 | 4.07035 | 4.28395 |
| CD630_01590 | CD630_n00590 | 8 -- 31 | 22 -- 41 | -11.6026 | 3.45609 | 4.28395 |
| CD630_15060 | CD630_n00590 | 148 -- 193 | 25 -- 88 | -11.4602 | 3.69396 | 4.28395 |
| CD630_09100 | CD630_n00590 | 11 -- 56 | 21 -- 70 | -11.3599 | 5.92260 | 4.28395 |
| CD630_16540 | CD630_n00590 | 128 -- 154 | 48 -- 71 | -11.3565 | 4.37658 | 4.28395 |
| CD630_11240 | CD630_n00590 | 128 -- 183 | 24 -- 76 | -11.2597 | 3.20274 | 4.28395 |
| spaK | CD630_n00590 | 140 -- 201 | 29 -- 87 | -11.2436 | 3.04530 | 4.28395 |
| feoB | CD630_n00590 | 8 -- 41 | 22 -- 57 | -11.2103 | 4.52257 | 4.28395 |
| CD630_05030 | CD630_n00590 | 1 -- 59 | 4 -- 73 | -11.1412 | 4.03823 | 4.28395 |
| CD630_18090 | CD630_n00590 | 5 -- 38 | 20 -- 45 | -11.0762 | 4.23961 | 4.28395 |
| CD630_29650 | CD630_n00590 | 138 -- 181 | 36 -- 81 | -10.9037 | 3.67128 | 4.28395 |
| CD630_13350 | CD630_n00590 | 148 -- 193 | 45 -- 101 | -10.8601 | 4.15139 | 4.28395 |
| CD630_06150 | CD630_n00590 | 8 -- 22 | 56 -- 70 | -10.8482 | 5.51744 | 4.28395 |
| CD630_19840 | CD630_n00590 | 173 -- 189 | 56 -- 73 | -10.7862 | 3.44724 | 4.28395 |
| CD630_28290 | CD630_n00590 | 128 -- 187 | 22 -- 75 | -10.7674 | 5.54192 | 4.28395 |
| CD630_06100 | CD630_n00590 | 122 -- 191 | 23 -- 80 | -10.7164 | 4.43918 | 4.28395 |
| CD630_09770 | CD630_n00590 | 3 -- 22 | 22 -- 40 | -10.7152 | 3.22106 | 4.28395 |
| CD630_07400 | CD630_n00590 | 124 -- 178 | 25 -- 77 | -10.7119 | 4.52273 | 4.28395 |
| CD630_27230 | CD630_n00590 | 133 -- 145 | 88 -- 100 | -10.6648 | 3.52462 | 4.28395 |
| CD630_14110 | CD630_n00590 | 115 -- 129 | 25 -- 39 | -10.6563 | 3.39550 | 4.28395 |
| CD630_08110 | CD630_n00590 | 126 -- 139 | 68 -- 81 | -10.5745 | 4.39181 | 4.28395 |
| CD630_18010 | CD630_n00590 | 174 -- 198 | 57 -- 82 | -10.522 | 4.21967 | 4.28395 |
| dnaD | CD630_n00590 | 29 -- 39 | 67 -- 77 | -10.5017 | 3.56078 | 4.28395 |
| CD630_32520 | CD630_n00590 | 143 -- 156 | 16 -- 29 | -10.4442 | 4.06098 | 4.28395 |
| CD630_08080 | CD630_n00590 | 8 -- 68 | 30 -- 80 | -10.4171 | 4.87819 | 4.28395 |
| CD630_27220 | CD630_n00590 | 131 -- 148 | 56 -- 73 | -10.3906 | 4.18786 | 4.28395 |
| CD630_21700 | CD630_n00590 | 1 -- 28 | 55 -- 79 | -10.2827 | 3.96650 | 4.28395 |
| CD630_15160 | CD630_n00590 | 168 -- 191 | 12 -- 35 | -10.2456 | 4.32318 | 4.28395 |
| CD630_03160 | CD630_n00590 | 17 -- 45 | 13 -- 38 | -10.1983 | 3.66946 | 4.28395 |
| CD630_19030 | CD630_n00590 | 134 -- 173 | 24 -- 80 | -10.165 | 3.34240 | 4.28395 |
| CD630_31960 | CD630_n00590 | 10 -- 35 | 14 -- 41 | -10.0546 | 4.93757 | 4.28395 |
| CD630_07770 | CD630_n00590 | 137 -- 151 | 58 -- 73 | -10.0389 | 3.31158 | 4.28395 |
| CD630_25050 | CD630_n00590 | 19 -- 53 | 65 -- 101 | -9.94645 | 4.33624 | 4.28395 |
| CD630_30620 | CD630_n00590 | 10 -- 72 | 25 -- 99 | -9.93245 | 4.15443 | 4.28395 |
| CD630_21090 | CD630_n00590 | 111 -- 162 | 29 -- 77 | -9.92016 | 4.15803 | 4.28395 |
| CD630_05822 | CD630_n00590 | 188 -- 201 | 21 -- 35 | -9.89869 | 4.28113 | 4.28395 |
| CD630_04610 | CD630_n00590 | 146 -- 182 | 67 -- 101 | -9.78981 | 3.83025 | 4.28395 |
| CD630_18610 | CD630_n00590 | 188 -- 201 | 26 -- 39 | -9.65455 | 4.15582 | 4.28395 |
| CD630_17511 | CD630_n00590 | 4 -- 61 | 29 -- 80 | -9.6285 | 9.12322 | 4.28395 |
| CD630_20260 | CD630_n00590 | 10 -- 52 | 61 -- 100 | -9.52751 | 3.84162 | 4.28395 |
| CD630_25040 | CD630_n00590 | 111 -- 188 | 15 -- 97 | -9.50221 | 3.43898 | 4.28395 |
| CD630_06170 | CD630_n00650 | 99 -- 125 | 45 -- 72 | -15.9644 | 4.95366 | 5.32127 |
| CD630_25020 | CD630_n00650 | 10 -- 68 | 54 -- 97 | -14.1844 | 4.17721 | 5.32127 |
| CD630_26150 | CD630_n00650 | 130 -- 157 | 51 -- 78 | -13.8939 | 4.06302 | 5.32127 |
| CD630_07770 | CD630_n00650 | 134 -- 145 | 48 -- 59 | -13.5894 | 3.31158 | 5.32127 |
| CD630_05490 | CD630_n00650 | 34 -- 57 | 45 -- 68 | -13.0319 | 3.89852 | 5.32127 |
| CD630_15200 | CD630_n00650 | 120 -- 143 | 41 -- 67 | -12.6272 | 4.00442 | 5.32127 |
| oppB | CD630_n00650 | 89 -- 151 | 1 -- 57 | -12.4532 | 2.98328 | 5.32127 |
| CD630_22050 | CD630_n00650 | 105 -- 152 | 38 -- 83 | -12.1179 | 4.87639 | 5.32127 |
| CD630_22070 | CD630_n00650 | 39 -- 79 | 54 -- 93 | -11.9734 | 4.09651 | 5.32127 |
| CD630_27360 | CD630_n00650 | 172 -- 195 | 49 -- 75 | -11.6807 | 3.79570 | 5.32127 |
| CD630_07121 | CD630_n00650 | 120 -- 172 | 47 -- 97 | -11.6294 | 3.26177 | 5.32127 |
| CD630_13930 | CD630_n00650 | 121 -- 156 | 49 -- 83 | -11.5641 | 4.09246 | 5.32127 |
| CD630_05480 | CD630_n00650 | 158 -- 180 | 43 -- 65 | -11.4411 | 4.27854 | 5.32127 |
| CD630_05900 | CD630_n00650 | 1 -- 23 | 51 -- 73 | -11.3544 | 3.21934 | 5.32127 |
| CD630_09100 | CD630_n00650 | 31 -- 87 | 30 -- 95 | -11.2011 | 5.92260 | 5.32127 |
| CD630_18910 | CD630_n00650 | 131 -- 144 | 52 -- 66 | -11.1662 | 3.30688 | 5.32127 |
| xylR | CD630_n00650 | 18 -- 51 | 47 -- 89 | -10.9318 | 3.95671 | 5.32127 |
| CD630_32890 | CD630_n00650 | 99 -- 145 | 48 -- 87 | -10.8659 | 3.41276 | 5.32127 |
| CD630_05820 | CD630_n00650 | 169 -- 196 | 51 -- 81 | -10.6361 | 3.75661 | 5.32127 |
| CD630_20290 | CD630_n00650 | 125 -- 151 | 45 -- 68 | -10.6173 | 4.38183 | 5.32127 |
| CD630_24580 | CD630_n00650 | 22 -- 63 | 44 -- 79 | -10.5374 | 3.88519 | 5.32127 |
| CD630_08950 | CD630_n00650 | 146 -- 196 | 52 -- 98 | -10.5292 | 3.71905 | 5.32127 |
| CD630_23440 | CD630_n00650 | 96 -- 109 | 77 -- 90 | -10.386 | 3.34841 | 5.32127 |
| CD630_25150 | CD630_n00650 | 1 -- 63 | 41 -- 93 | -10.329 | 3.79339 | 5.32127 |
| CD630_28320 | CD630_n00650 | 107 -- 142 | 52 -- 95 | -10.1577 | 3.99184 | 5.32127 |
| CD630_10790 | CD630_n00650 | 51 -- 76 | 49 -- 78 | -10.0811 | 4.45284 | 5.32127 |
| CD630_03420 | CD630_n00650 | 51 -- 82 | 36 -- 68 | -10.035 | 4.71868 | 5.32127 |
| ldh | CD630_n00650 | 34 -- 55 | 54 -- 74 | -9.98135 | 4.35818 | 5.32127 |
| dltA | CD630_n00650 | 131 -- 150 | 52 -- 74 | -9.96107 | 3.85429 | 5.32127 |
| CD630_04410 | CD630_n00650 | 38 -- 58 | 42 -- 60 | -9.83256 | 3.90451 | 5.32127 |
| panG | CD630_n00650 | 163 -- 190 | 54 -- 81 | -9.76924 | 5.54047 | 5.32127 |
| sigK | CD630_n00650 | 97 -- 130 | 51 -- 86 | -9.75637 | 3.08122 | 5.32127 |
| CD630_22660 | CD630_n00650 | 113 -- 148 | 64 -- 97 | -9.66876 | 3.60496 | 5.32127 |
| chrA | CD630_n00650 | 1 -- 95 | 1 -- 84 | -9.64812 | 5.27255 | 5.32127 |
| CD630_25070 | CD630_n00650 | 132 -- 173 | 41 -- 76 | -9.64614 | 3.45164 | 5.32127 |
| cbpA | CD630_n00650 | 145 -- 182 | 35 -- 76 | -9.60381 | 3.03040 | 5.32127 |
| spaK | CD630_n00650 | 150 -- 160 | 51 -- 61 | -9.54587 | 3.04530 | 5.32127 |
| CD630_22500 | CD630_n00690 | 115 -- 156 | 1 -- 39 | -18.0222 | 4.03688 | 5.96526 |
| CD630_14340 | CD630_n00690 | 84 -- 167 | 125 -- 208 | -17.5478 | 5.10868 | 5.96526 |
| CD630_27570 | CD630_n00690 | 107 -- 183 | 153 -- 225 | -17.2621 | 3.46216 | 5.96526 |
| CD630_28260 | CD630_n00690 | 58 -- 190 | 120 -- 256 | -16.9794 | 3.51613 | 5.96526 |
| pilT | CD630_n00690 | 128 -- 168 | 275 -- 313 | -16.8084 | 3.24639 | 5.96526 |
| panG | CD630_n00690 | 14 -- 72 | 158 -- 224 | -16.3558 | 5.54047 | 5.96526 |

| | | | | | | | |
|---|---|---|---|---|---|---|---|
| CD630_33600 | CD630_n00690 | 134 -- 163 | 286 -- 316 | -16.103 | | 3.73196 | 5.96526 |
| CD630_19790 | CD630_n00690 | 49 -- 101 | 276 -- 317 | -16.0935 | | 3.79593 | 5.96526 |
| CD630_06060 | CD630_n00690 | 49 -- 93 | 199 -- 238 | -16.0517 | | 4.94050 | 5.96526 |
| bglA | CD630_n00690 | 53 -- 110 | 275 -- 326 | -15.6789 | | 4.50315 | 5.96526 |
| CD630_06150 | CD630_n00690 | 106 -- 193 | 220 -- 295 | -15.4436 | | 5.51744 | 5.96526 |
| CD630_31910 | CD630_n00690 | 86 -- 193 | 161 -- 251 | -15.2342 | | 3.48314 | 5.96526 |
| CD630_21620 | CD630_n00690 | 117 -- 166 | 280 -- 324 | -15.2295 | | 3.02649 | 5.96526 |
| CD630_18610 | CD630_n00690 | 26 -- 159 | 48 -- 187 | -14.632 | | 4.15582 | 5.96526 |
| CD630_05880 | CD630_n00690 | 90 -- 141 | 180 -- 235 | -14.5868 | | 4.28339 | 5.96526 |
| CD630_23820 | CD630_n00690 | 148 -- 185 | 279 -- 314 | -14.2079 | | 3.18419 | 5.96526 |
| CD630_14110 | CD630_n00690 | 24 -- 49 | 23 -- 44 | -14.1615 | | 3.39550 | 5.96526 |
| folC | CD630_n00690 | 125 -- 188 | 275 -- 329 | -14.0483 | | 3.85634 | 5.96526 |
| cobC | CD630_n00690 | 139 -- 181 | 271 -- 314 | -13.9884 | | 3.15854 | 5.96526 |
| CD630_24920 | CD630_n00690 | 118 -- 156 | 212 -- 248 | -13.9571 | | 3.89455 | 5.96526 |
| CD630_16460 | CD630_n00690 | 134 -- 171 | 208 -- 247 | -13.915 | | 4.64028 | 5.96526 |
| CD630_14250 | CD630_n00690 | 177 -- 198 | 159 -- 181 | -13.8645 | | 4.19604 | 5.96526 |
| CD630_23080 | CD630_n00690 | 151 -- 186 | 275 -- 306 | -13.7628 | | 5.23027 | 5.96526 |
| CD630_22720 | CD630_n00690 | 120 -- 166 | 276 -- 318 | -13.7419 | | 3.81804 | 5.96526 |
| CD630_24050 | CD630_n00690 | 118 -- 135 | 287 -- 305 | -13.7233 | | 3.70192 | 5.96526 |
| CD630_17170 | CD630_n00690 | 99 -- 145 | 274 -- 321 | -13.6907 | | 4.10230 | 5.96526 |
| CD630_21330 | CD630_n00690 | 46 -- 71 | 283 -- 308 | -13.5399 | | 4.10111 | 5.96526 |
| CD630_21540 | CD630_n00690 | 37 -- 97 | 202 -- 262 | -13.51 | | 4.33090 | 5.96526 |
| CD630_25610 | CD630_n00690 | 8 -- 50 | 276 -- 322 | -13.4914 | | 3.02685 | 5.96526 |
| CD630_31530 | CD630_n00690 | 127 -- 183 | 266 -- 313 | -13.4442 | | 3.49100 | 5.96526 |
| CD630_06560 | CD630_n00690 | 4 -- 52 | 275 -- 324 | -13.4244 | | 4.02532 | 5.96526 |
| gcp | CD630_n00690 | 25 -- 58 | 88 -- 116 | -13.2914 | | 4.92798 | 5.96526 |
| CD630_07960 | CD630_n00690 | 34 -- 69 | 279 -- 318 | -13.2208 | | 3.22537 | 5.96526 |
| CD630_03420 | CD630_n00690 | 186 -- 201 | 211 -- 227 | -13.2111 | | 4.71868 | 5.96526 |
| oppC | CD630_n00690 | 98 -- 139 | 109 -- 150 | -13.1493 | | 3.01695 | 5.96526 |
| CD630_06690 | CD630_n00690 | 24 -- 70 | 187 -- 235 | -13.134 | | 5.82709 | 5.96526 |
| srlD | CD630_n00690 | 88 -- 143 | 204 -- 251 | -13.0578 | | 4.19843 | 5.96526 |
| CD630_27520 | CD630_n00690 | 77 -- 141 | 161 -- 218 | -12.9021 | | 4.27693 | 5.96526 |
| xdhA | CD630_n00690 | 99 -- 146 | 280 -- 328 | -12.8828 | | 4.45147 | 5.96526 |
| CD630_32890 | CD630_n00690 | 31 -- 83 | 125 -- 176 | -12.8679 | | 3.41276 | 5.96526 |
| CD630_06970 | CD630_n00690 | 110 -- 154 | 267 -- 322 | -12.8054 | | 3.82312 | 5.96526 |
| CD630_17292 | CD630_n00690 | 143 -- 185 | 275 -- 313 | -12.7519 | | 6.25799 | 5.96526 |
| appF | CD630_n00690 | 42 -- 82 | 84 -- 123 | -12.707 | | 4.25570 | 5.96526 |
| CD630_08460 | CD630_n00690 | 20 -- 52 | 159 -- 187 | -12.6564 | | 4.15083 | 5.96526 |
| CD630_33560 | CD630_n00690 | 73 -- 130 | 127 -- 173 | -12.6421 | | 4.03147 | 5.96526 |
| CD630_07480 | CD630_n00690 | 52 -- 71 | 274 -- 293 | -12.6384 | | 5.18321 | 5.96526 |
| CD630_27310 | CD630_n00690 | 85 -- 156 | 155 -- 226 | -12.6384 | | 4.01622 | 5.96526 |
| CD630_13620 | CD630_n00690 | 136 -- 199 | 124 -- 177 | -12.5787 | | 4.77627 | 5.96526 |
| CD630_14360 | CD630_n00690 | 88 -- 103 | 209 -- 225 | -12.5625 | | 3.02079 | 5.96526 |
| CD630_31490 | CD630_n00690 | 103 -- 116 | 108 -- 121 | -12.5024 | | 4.08593 | 5.96526 |
| CD630_16330 | CD630_n00690 | 129 -- 155 | 40 -- 72 | -12.4865 | | 4.11180 | 5.96526 |
| CD630_21730 | CD630_n00690 | 151 -- 181 | 278 -- 311 | -12.454 | | 3.45076 | 5.96526 |
| CD630_27200 | CD630_n00690 | 2 -- 15 | 207 -- 220 | -12.4177 | | 3.42516 | 5.96526 |
| CD630_14420 | CD630_n00690 | 24 -- 66 | 154 -- 196 | -12.3473 | | 3.99497 | 5.96526 |
| CD630_24590 | CD630_n00690 | 74 -- 158 | 178 -- 248 | -12.3468 | | 3.70401 | 5.96526 |
| CD630_28181 | CD630_n00690 | 43 -- 163 | 107 -- 225 | -12.3175 | | 3.91299 | 5.96526 |
| CD630_13880 | CD630_n00690 | 22 -- 53 | 209 -- 239 | -12.2802 | | 3.52735 | 5.96526 |
| CD630_18970 | CD630_n00690 | 10 -- 32 | 292 -- 313 | -12.2766 | | 3.57243 | 5.96526 |
| def1 | CD630_n00690 | 122 -- 164 | 275 -- 321 | -12.232 | | 3.99720 | 5.96526 |
| CD630_19401 | CD630_n00690 | 49 -- 84 | 25 -- 62 | -12.2315 | | 4.37084 | 5.96526 |
| CD630_31970 | CD630_n00690 | 156 -- 196 | 41 -- 69 | -12.2047 | | 3.24511 | 5.96526 |
| CD630_25020 | CD630_n00690 | 20 -- 47 | 60 -- 86 | -12.2033 | | 4.17721 | 5.96526 |
| CD630_20170 | CD630_n00690 | 163 -- 196 | 293 -- 328 | -12.1757 | | 6.90201 | 5.96526 |
| CD630_07770 | CD630_n00690 | 82 -- 199 | 211 -- 324 | -12.1723 | | 3.31158 | 5.96526 |
| argS | CD630_n00690 | 171 -- 190 | 217 -- 236 | -12.1484 | | 4.96036 | 5.96526 |
| CD630_02420 | CD630_n00690 | 1 -- 24 | 91 -- 115 | -12.1074 | | 3.71048 | 5.96526 |
| CD630_29960 | CD630_n00690 | 168 -- 197 | 282 -- 310 | -12.0864 | | 4.51904 | 5.96526 |
| CD630_22950 | CD630_n00690 | 103 -- 136 | 200 -- 232 | -11.9986 | | 3.96883 | 5.96526 |
| CD630_19440 | CD630_n00690 | 140 -- 178 | 6 -- 46 | -11.9172 | | 4.53937 | 5.96526 |
| CD630_21720 | CD630_n00690 | 155 -- 197 | 195 -- 247 | -11.8779 | | 4.13030 | 5.96526 |
| CD630_09850 | CD630_n00690 | 150 -- 201 | 2 -- 59 | -11.819 | | 4.18679 | 5.96526 |
| cbiQ | CD630_n00690 | 88 -- 168 | 155 -- 225 | -11.7999 | | 3.79830 | 5.96526 |
| CD630_32520 | CD630_n00690 | 154 -- 194 | 275 -- 317 | -11.7883 | | 4.06098 | 5.96526 |
| nusB | CD630_n00690 | 126 -- 201 | 166 -- 235 | -11.7765 | | 4.92798 | 5.96526 |
| CD630_19904 | CD630_n00690 | 140 -- 172 | 281 -- 313 | -11.7672 | | 5.96526 | 5.96526 |
| CD630_32710 | CD630_n00690 | 135 -- 150 | 275 -- 292 | -11.6575 | | 4.74276 | 5.96526 |
| CD630_31690 | CD630_n00690 | 117 -- 134 | 19 -- 35 | -11.6447 | | 3.71193 | 5.96526 |
| CD630_30110 | CD630_n00690 | 101 -- 122 | 307 -- 328 | -11.6322 | | 3.31041 | 5.96526 |
| CD630_32110 | CD630_n00690 | 79 -- 139 | 160 -- 221 | -11.5522 | | 4.45291 | 5.96526 |
| CD630_09050 | CD630_n00690 | 96 -- 149 | 128 -- 183 | -11.5464 | | 3.96216 | 5.96526 |
| CD630_21260 | CD630_n00690 | 48 -- 190 | 20 -- 153 | -11.5441 | | 4.22255 | 5.96526 |
| CD630_05820 | CD630_n00690 | 179 -- 196 | 289 -- 307 | -11.5205 | | 3.75661 | 5.96526 |
| CD630_32070 | CD630_n00690 | 38 -- 70 | 279 -- 315 | -11.5162 | | 3.51964 | 5.96526 |
| CD630_20290 | CD630_n00690 | 135 -- 174 | 42 -- 75 | -11.5007 | | 4.38183 | 5.96526 |
| dltB | CD630_n00690 | 8 -- 65 | 126 -- 180 | -11.453 | | 3.81485 | 5.96526 |
| CD630_10220 | CD630_n00690 | 121 -- 166 | 274 -- 308 | -11.4301 | | 4.08233 | 5.96526 |
| feoB | CD630_n00690 | 144 -- 185 | 276 -- 321 | -11.425 | | 4.05364 | 5.96526 |
| CD630_33570 | CD630_n00690 | 140 -- 174 | 209 -- 244 | -11.4087 | | 3.95778 | 5.96526 |
| CD630_15850 | CD630_n00690 | 154 -- 197 | 182 -- 221 | -11.4049 | | 4.32968 | 5.96526 |
| CD630_32700 | CD630_n00690 | 155 -- 200 | 205 -- 253 | -11.3815 | | 4.74276 | 5.96526 |
| CD630_27280 | CD630_n00690 | 130 -- 165 | 3 -- 37 | -11.3728 | | 3.24738 | 5.96526 |
| CD630_31000 | CD630_n00690 | 105 -- 153 | 187 -- 225 | -11.2618 | | 3.70946 | 5.96526 |
| CD630_21140 | CD630_n00690 | 26 -- 171 | 48 -- 180 | -11.2049 | | 3.33304 | 5.96526 |
| CD630_12650 | CD630_n00690 | 106 -- 128 | 99 -- 123 | -11.1894 | | 6.31827 | 5.96526 |
| CD630_10800 | CD630_n00690 | 72 -- 124 | 273 -- 311 | -11.1767 | | 5.30319 | 5.96526 |
| CD630_19800 | CD630_n00690 | 100 -- 152 | 184 -- 247 | -11.1625 | | 5.22035 | 5.96526 |
| glgB | CD630_n00690 | 163 -- 185 | 160 -- 182 | -11.0995 | | 4.48273 | 5.96526 |
| etfA | CD630_n00690 | 54 -- 130 | 161 -- 245 | -11.0928 | | 4.80521 | 5.96526 |
| CD630_31480 | CD630_n00690 | 166 -- 186 | 280 -- 301 | -11.0638 | | 4.88897 | 5.96526 |
| CD630_06100 | CD630_n00690 | 144 -- 171 | 61 -- 89 | -11.0456 | | 4.43918 | 5.96526 |
| CD630_25320 | CD630_n00690 | 147 -- 169 | 280 -- 308 | -11.0361 | | 4.29415 | 5.96526 |
| appC | CD630_n00690 | 32 -- 76 | 161 -- 208 | -11.0289 | | 4.50438 | 5.96526 |
| CD630_20450 | CD630_n00690 | 146 -- 182 | 276 -- 310 | -11.0277 | | 3.77388 | 5.96526 |
| CD630_27610 | CD630_n00690 | 13 -- 47 | 158 -- 190 | -11.0146 | | 3.96465 | 5.96526 |
| CD630_20670 | CD630_n00690 | 50 -- 124 | 199 -- 255 | -10.9997 | | 4.42757 | 5.96526 |
| CD630_30190 | CD630_n00690 | 97 -- 124 | 20 -- 46 | -10.9988 | | 3.27394 | 5.96526 |
| CD630_11220 | CD630_n00690 | 174 -- 196 | 23 -- 46 | -10.959 | | 3.52327 | 5.96526 |
| CD630_06960 | CD630_n00690 | 31 -- 48 | 160 -- 176 | -10.9363 | | 3.16317 | 5.96526 |
| nrdD | CD630_n00690 | 6 -- 71 | 163 -- 228 | -10.9055 | | 3.28290 | 5.96526 |
| ssuC | CD630_n00690 | 47 -- 74 | 290 -- 313 | -10.8983 | | 3.45491 | 5.96526 |
| cobQ | CD630_n00690 | 171 -- 194 | 273 -- 294 | -10.8974 | | 3.13524 | 5.96526 |
| CD630_33650 | CD630_n00690 | 184 -- 201 | 30 -- 46 | -10.8919 | | 4.80565 | 5.96526 |
| CD630_05000 | CD630_n00690 | 98 -- 167 | 125 -- 190 | -10.8554 | | 4.02442 | 5.96526 |
| rbsC | CD630_n00690 | 55 -- 92 | 221 -- 253 | -10.8301 | | 3.71650 | 5.96526 |
| CD630_09040 | CD630_n00690 | 152 -- 186 | 289 -- 322 | -10.822 | | 4.11851 | 5.96526 |
| dpaA | CD630_n00690 | 19 -- 54 | 189 -- 219 | -10.7296 | | 4.65364 | 5.96526 |
| CD630_22070 | CD630_n00690 | 23 -- 55 | 279 -- 312 | -10.7122 | | 4.09651 | 5.96526 |
| CD630_18100 | CD630_n00690 | 81 -- 154 | 156 -- 229 | -10.7075 | | 5.71658 | 5.96526 |
| CD630_32060 | CD630_n00690 | 159 -- 181 | 274 -- 292 | -10.6998 | | 5.73797 | 5.96526 |
| CD630_36781 | CD630_n00690 | 153 -- 170 | 17 -- 37 | -10.6967 | | 3.75946 | 5.96526 |
| CD630_15460 | CD630_n00690 | 119 -- 143 | 277 -- 301 | -10.6903 | | 6.86296 | 5.96526 |
| malY | CD630_n00690 | 52 -- 74 | 281 -- 303 | -10.6826 | | 4.46345 | 5.96526 |
| CD630_21750 | CD630_n00690 | 49 -- 97 | 285 -- 328 | -10.6551 | | 3.70626 | 5.96526 |
| CD630_21700 | CD630_n00690 | 37 -- 58 | 145 -- 165 | -10.6263 | | 3.96650 | 5.96526 |
| CD630_18660 | CD630_n00690 | 125 -- 139 | 38 -- 52 | -10.5915 | | 3.50371 | 5.96526 |
| CD630_22880 | CD630_n00690 | 74 -- 113 | 275 -- 312 | -10.5862 | | 3.62455 | 5.96526 |
| CD630_02720 | CD630_n00690 | 137 -- 168 | 284 -- 313 | -10.573 | | 3.38938 | 5.96526 |
| opuCC | CD630_n00690 | 161 -- 193 | 41 -- 70 | -10.5425 | | 4.07035 | 5.96526 |
| moaA | CD630_n00690 | 98 -- 114 | 280 -- 295 | -10.5148 | | 3.83799 | 5.96526 |
| CD630_31010 | CD630_n00690 | 131 -- 152 | 75 -- 97 | -10.4893 | | 4.96305 | 5.96526 |
| cprK | CD630_n00690 | 162 -- 201 | 48 -- 88 | -10.4629 | | 3.96216 | 5.96526 |
| CD630_34030 | CD630_n00690 | 75 -- 108 | 279 -- 312 | -10.453 | | 3.05766 | 5.96526 |
| spaK | CD630_n00690 | 87 -- 103 | 209 -- 225 | -10.4438 | | 3.04530 | 5.96526 |
| CD630_09100 | CD630_n00690 | 6 -- 33 | 42 -- 70 | -10.4434 | | 5.92260 | 5.96526 |
| CD630_15360 | CD630_n00690 | 89 -- 121 | 274 -- 301 | -10.4335 | | 4.47208 | 5.96526 |
| cbiO | CD630_n00690 | 84 -- 118 | 160 -- 191 | -10.4273 | | 3.79830 | 5.96526 |
| CD630_27220 | CD630_n00690 | 110 -- 140 | 145 -- 176 | -10.4065 | | 4.18786 | 5.96526 |
| CD630_17960 | CD630_n00690 | 131 -- 152 | 226 -- 247 | -10.3928 | | 3.71404 | 5.96526 |
| CD630_32990 | CD630_n00690 | 51 -- 104 | 283 -- 332 | -10.3672 | | 5.24925 | 5.96526 |
| feoB | CD630_n00690 | 21 -- 35 | 227 -- 241 | -10.3552 | | 4.52257 | 5.96526 |
| CD630_31461 | CD630_n00690 | 151 -- 190 | 278 -- 321 | -10.3526 | | 4.41476 | 5.96526 |
| CD630_14190 | CD630_n00690 | 99 -- 107 | 280 -- 288 | -10.3264 | | 3.78562 | 5.96526 |
| CD630_04610 | CD630_n00690 | 104 -- 115 | 165 -- 176 | -10.3116 | | 3.83025 | 5.96526 |
| CD630_25150 | CD630_n00690 | 176 -- 201 | 11 -- 31 | -10.3094 | | 3.79339 | 5.96526 |
| CD630_04550 | CD630_n00690 | 28 -- 45 | 165 -- 182 | -10.2662 | | 3.05227 | 5.96526 |
| CD630_25310 | CD630_n00690 | 8 -- 45 | 275 -- 313 | -10.2655 | | 3.99861 | 5.96526 |
| eutW | CD630_n00690 | 58 -- 85 | 162 -- 192 | -10.2404 | | 4.55715 | 5.96526 |
| CD630_13530 | CD630_n00690 | 167 -- 197 | 37 -- 67 | -10.2345 | | 4.09832 | 5.96526 |
| rolB | CD630_n00690 | 147 -- 191 | 276 -- 324 | -10.2326 | | 3.60270 | 5.96526 |
| CD630_15740 | CD630_n00690 | 48 -- 73 | 159 -- 182 | -10.2169 | | 4.43222 | 5.96526 |
| CD630_18930 | CD630_n00690 | 116 -- 158 | 185 -- 226 | -10.2119 | | 5.32127 | 5.96526 |
| CD630_27630 | CD630_n00690 | 83 -- 101 | 225 -- 242 | -10.1796 | | 3.54199 | 5.96526 |
| CD630_06200 | CD630_n00690 | 99 -- 130 | 281 -- 312 | -10.1603 | | 3.57064 | 5.96526 |
| CD630_06080 | CD630_n00690 | 123 -- 155 | 289 -- 319 | -10.1344 | | 3.82315 | 5.96526 |
| CD630_23000 | CD630_n00690 | 161 -- 190 | 286 -- 319 | -10.1274 | | 3.54130 | 5.96526 |
| CD630_19921 | CD630_n00690 | 89 -- 111 | 201 -- 224 | -10.1264 | | 5.46345 | 5.96526 |
| CD630_18680 | CD630_n00690 | 118 -- 149 | 211 -- 242 | -10.1205 | | 3.04315 | 5.96526 |
| CD630_14700 | CD630_n00690 | 170 -- 194 | 157 -- 181 | -10.1172 | | 3.98234 | 5.96526 |
| CD630_18440 | CD630_n00690 | 156 -- 201 | 180 -- 227 | -10.1059 | | 4.07960 | 5.96526 |
| CD630_13560 | CD630_n00690 | 191 -- 201 | 25 -- 35 | -10.0785 | | 3.38083 | 5.96526 |
| bioB | CD630_n00690 | 115 -- 148 | 152 -- 181 | -10.0498 | | 3.59022 | 5.96526 |
| CD630_15111 | CD630_n00690 | 29 -- 43 | 279 -- 293 | -10.0148 | | 4.18664 | 5.96526 |
| CD630_19680 | CD630_n00690 | 37 -- 151 | 100 -- 221 | -10.0026 | | 3.58080 | 5.96526 |
| vncR | CD630_n00690 | 137 -- 152 | 209 -- 225 | -10.0022 | | 3.93642 | 5.96526 |
| CD630_20960 | CD630_n00690 | 169 -- 183 | 220 -- 235 | -9.98938 | | 5.65940 | 5.96526 |
| CD630_28550 | CD630_n00690 | 172 -- 199 | 270 -- 302 | -9.98652 | | 3.51196 | 5.96526 |
| cwp26 | CD630_n00690 | 136 -- 154 | 20 -- 37 | -9.97132 | | 4.44377 | 5.96526 |
| CD630_26691 | CD630_n00690 | 91 -- 117 | 200 -- 225 | -9.96856 | | 3.96230 | 5.96526 |
| CD630_09770 | CD630_n00690 | 127 -- 157 | 280 -- 301 | -9.96808 | | 3.22106 | 5.96526 |
| CD630_17770 | CD630_n00690 | 113 -- 133 | 124 -- 142 | -9.96237 | | 5.06490 | 5.96526 |
| CD630_18920 | CD630_n00690 | 170 -- 180 | 290 -- 300 | -9.95095 | | 4.43558 | 5.96526 |
| adhE | CD630_n00690 | 162 -- 170 | 219 -- 227 | -9.94636 | | 3.06485 | 5.96526 |
| CD630_20980 | CD630_n00690 | 139 -- 155 | 211 -- 229 | -9.93773 | | 8.12847 | 5.96526 |
| CD630_06170 | CD630_n00690 | 65 -- 81 | 161 -- 177 | -9.93492 | | 4.95366 | 5.96526 |
| ppnK | CD630_n00690 | 146 -- 167 | 283 -- 304 | -9.90302 | | 4.36817 | 5.96526 |
| CD630_13600 | CD630_n00690 | 154 -- 181 | 149 -- 176 | -9.90121 | | 4.45481 | 5.96526 |
| CD630_27210 | CD630_n00690 | 10 -- 53 | 286 -- 319 | -9.87652 | | 4.18786 | 5.96526 |
| CD630_08360 | CD630_n00690 | 165 -- 187 | 278 -- 301 | -9.8504 | | 3.19339 | 5.96526 |
| CD630_07840 | CD630_n00690 | 42 -- 97 | 279 -- 329 | -9.79715 | | 4.26761 | 5.96526 |
| CD630_04850 | CD630_n00690 | 124 -- 199 | 1 -- 75 | -9.78659 | | 4.40788 | 5.96526 |
| CD630_03410 | CD630_n00690 | 30 -- 76 | 135 -- 182 | -9.78211 | | 4.45994 | 5.96526 |
| pilC | CD630_n00690 | 16 -- 55 | 290 -- 321 | -9.78198 | | 3.54230 | 5.96526 |
| ldh | CD630_n00690 | 131 -- 152 | 273 -- 294 | -9.7665 | | 4.35818 | 5.96526 |
| CD630_30140 | CD630_n00690 | 92 -- 125 | 276 -- 313 | -9.75111 | | 3.28343 | 5.96526 |
| CD630_28280 | CD630_n00690 | 73 -- 82 | 278 -- 287 | -9.73979 | | 4.59833 | 5.96526 |
| terD | CD630_n00690 | 9 -- 33 | 129 -- 162 | -9.71819 | | 3.33148 | 5.96526 |
| CD630_33580 | CD630_n00690 | 176 -- 199 | 280 -- 295 | -9.70724 | | 3.95778 | 5.96526 |
| CD630_03920 | CD630_n00690 | 144 -- 168 | 181 -- 203 | -9.68594 | | 4.15934 | 5.96526 |
| CD630_18910 | CD630_n00690 | 93 -- 139 | 194 -- 249 | -9.68522 | | 3.30688 | 5.96526 |
| CD630_31960 | CD630_n00690 | 6 -- 60 | 125 -- 176 | -9.65169 | | 4.93757 | 5.96526 |
| panB | CD630_n00690 | 4 -- 35 | 278 -- 313 | -9.59901 | | 3.40248 | 5.96526 |
| CD630_35260 | CD630_n00690 | 26 -- 38 | 163 -- 175 | -9.59423 | | 5.77277 | 5.96526 |
| CD630_24770 | CD630_n00690 | 159 -- 174 | 281 -- 296 | -9.5813 | | 3.20798 | 5.96526 |
| iorA | CD630_n00690 | 99 -- 134 | 276 -- 312 | -9.56686 | | 3.83173 | 5.96526 |
| pilD | CD630_n00690 | 1 -- 48 | 274 -- 310 | -9.56577 | | 4.54123 | 5.96526 |
| CD630_10960 | CD630_n00690 | 12 -- 41 | 281 -- 311 | -9.53881 | | 3.53850 | 5.96526 |
| CD630_02430 | CD630_n00690 | 54 -- 70 | 162 -- 178 | -9.5346 | | 3.71048 | 5.96526 |
| CD630_28330 | CD630_n00690 | 48 -- 61 | 280 -- 295 | -9.5152 | | 3.29122 | 5.96526 |
| CD630_32070 | CD630_n00790 | 52 -- 83 | 204 -- 242 | -19.4584 | | 3.51964 | 5.51210 |
| srlD | CD630_n00790 | 77 -- 106 | 200 -- 228 | -19.3783 | | 4.19843 | 5.51210 |
| CD630_18100 | CD630_n00790 | 123 -- 165 | 207 -- 241 | -19.1924 | | 5.71658 | 5.51210 |
| CD630_27570 | CD630_n00790 | 102 -- 183 | 153 -- 243 | -19.113 | | 3.46216 | 5.51210 |
| CD630_31580 | CD630_n00790 | 127 -- 167 | 472 -- 510 | -18.9419 | | 3.51756 | 5.51210 |
| bcp | CD630_n00790 | 89 -- 161 | 203 -- 274 | -18.6833 | | 3.16670 | 5.51210 |
| CD630_05880 | CD630_n00790 | 75 -- 181 | 179 -- 290 | -18.4858 | | 4.28339 | 5.51210 |
| CD630_32520 | CD630_n00790 | 122 -- 199 | 211 -- 285 | -18.3522 | | 4.06098 | 5.51210 |
| CD630_04620 | CD630_n00790 | 105 -- 145 | 340 -- 376 | -18.2932 | | 3.45413 | 5.51210 |
| CD630_21790 | CD630_n00790 | 136 -- 198 | 214 -- 274 | -18.1492 | | 4.93762 | 5.51210 |
| CD630_10880 | CD630_n00790 | 82 -- 187 | 214 -- 296 | -18.1437 | | 5.05048 | 5.51210 |
| etfA | CD630_n00790 | 33 -- 164 | 99 -- 243 | -17.9995 | | 3.41204 | 5.51210 |
| CD630_18920 | CD630_n00790 | 148 -- 189 | 475 -- 513 | -17.949 | | 4.43558 | 5.51210 |
| cobQ | CD630_n00790 | 86 -- 107 | 200 -- 218 | -17.9362 | | 3.13524 | 5.51210 |
| CD630_07840 | CD630_n00790 | 42 -- 106 | 205 -- 263 | -17.8773 | | 4.26761 | 5.51210 |
| oppC | CD630_n00790 | 66 -- 137 | 109 -- 175 | -17.8624 | | 3.01695 | 5.51210 |
| CD630_12320 | CD630_n00790 | 69 -- 132 | 991 -- 1041 | -17.7558 | | 3.28667 | 5.51210 |
| CD630_14410 | CD630_n00790 | 49 -- 159 | 126 -- 237 | -17.5118 | | 3.39433 | 5.51210 |
| CD630_23440 | CD630_n00790 | 1 -- 23 | 191 -- 216 | -17.5073 | | 3.34841 | 5.51210 |
| CD630_17960 | CD630_n00790 | 82 -- 148 | 209 -- 289 | -17.4152 | | 3.71404 | 5.51210 |
| CD630_19000 | CD630_n00790 | 46 -- 124 | 999 -- 1069 | -17.3725 | | 4.18443 | 5.51210 |
| CD630_05870 | CD630_n00790 | 158 -- 194 | 201 -- 237 | -17.3366 | | 4.28339 | 5.51210 |
| iorA | CD630_n00790 | 48 -- 102 | 193 -- 241 | -17.0839 | | 3.83173 | 5.51210 |
| CD630_18680 | CD630_n00790 | 88 -- 109 | 198 -- 216 | -17.0141 | | 3.04315 | 5.51210 |
| CD630_16550 | CD630_n00790 | 82 -- 102 | 193 -- 219 | -16.9907 | | 5.61064 | 5.51210 |
| CD630_19780 | CD630_n00790 | 17 -- 31 | 204 -- 218 | -16.9898 | | 3.64830 | 5.51210 |
| CD630_07760 | CD630_n00790 | 40 -- 104 | 207 -- 266 | -16.9827 | | 3.64186 | 5.51210 |
| cobS | CD630_n00790 | 55 -- 114 | 195 -- 254 | -16.8926 | | 3.67481 | 5.51210 |
| CD630_01590 | CD630_n00790 | 167 -- 201 | 200 -- 230 | -16.8391 | | 3.45609 | 5.51210 |
| CD630_19942 | CD630_n00790 | 76 -- 135 | 214 -- 284 | -16.777 | | 4.24574 | 5.51210 |
| mobA | CD630_n00790 | 129 -- 198 | 205 -- 290 | -16.7627 | | 4.25703 | 5.51210 |
| CD630_25150 | CD630_n00790 | 147 -- 201 | 11 -- 47 | -16.7304 | | 3.79339 | 5.51210 |
| CD630_32060 | CD630_n00790 | 28 -- 56 | 464 -- 490 | -16.694 | | 5.73797 | 5.51210 |
| srlEb | CD630_n00790 | 24 -- 99 | 205 -- 272 | -16.6258 | | 3.50140 | 5.51210 |
| CD630_10050 | CD630_n00790 | 153 -- 201 | 200 -- 242 | -16.6164 | | 3.93750 | 5.51210 |
| CD630_24590 | CD630_n00790 | 113 -- 157 | 972 -- 1014 | -16.5652 | | 3.70401 | 5.51210 |
| fliI | CD630_n00790 | 179 -- 201 | 997 -- 1019 | -16.4743 | | 3.03666 | 5.51210 |
| xdhA | CD630_n00790 | 105 -- 158 | 995 -- 1047 | -16.3538 | | 4.45147 | 5.51210 |
| CD630_17160 | CD630_n00790 | 89 -- 108 | 200 -- 216 | -16.3159 | | 3.50228 | 5.51210 |
| CD630_06340 | CD630_n00790 | 26 -- 123 | 271 -- 356 | -16.2826 | | 4.35983 | 5.51210 |
| panB | CD630_n00790 | 41 -- 187 | 114 -- 260 | -16.2071 | | 3.40248 | 5.51210 |
| CD630_19401 | CD630_n00790 | 34 -- 114 | 855 -- 938 | -15.988 | | 4.37084 | 5.51210 |
| CD630_21620 | CD630_n00790 | 93 -- 183 | 174 -- 236 | -15.9432 | | 3.02649 | 5.51210 |
| CD630_06160 | CD630_n00790 | 164 -- 190 | 139 -- 166 | -15.5582 | | 5.51744 | 5.51210 |
| CD630_22070 | CD630_n00790 | 35 -- 80 | 530 -- 569 | -15.5366 | | 4.09651 | 5.51210 |
| CD630_22880 | CD630_n00790 | 30 -- 108 | 102 -- 183 | -15.5222 | | 3.62455 | 5.51210 |

| | | | | | | |
|---|---|---|---|---|---|---|
| CD630_36250 | CD630_n00790 | 53 -- 201 | 104 -- 239 | -15.3967 | 4.13224 | 5.51210 |
| CD630_18120 | CD630_n00790 | 63 -- 111 | 200 -- 244 | -15.3792 | 3.99713 | 5.51210 |
| CD630_02430 | CD630_n00790 | 17 -- 137 | 781 -- 912 | -15.2168 | 3.71048 | 5.51210 |
| CD630_28181 | CD630_n00790 | 4 -- 76 | 644 -- 712 | -15.159 | 3.91299 | 5.51210 |
| CD630_08940 | CD630_n00790 | 50 -- 94 | 211 -- 256 | -15.1552 | 3.72225 | 5.51210 |
| rbsC | CD630_n00790 | 157 -- 190 | 801 -- 837 | -15.1518 | 3.71650 | 5.51210 |
| CD630_17170 | CD630_n00790 | 67 -- 109 | 197 -- 239 | -15.1377 | 4.10230 | 5.51210 |
| CD630_04341 | CD630_n00790 | 120 -- 192 | 185 -- 278 | -15.1215 | 4.29262 | 5.51210 |
| CD630_31220 | CD630_n00790 | 86 -- 188 | 205 -- 289 | -15.0625 | 3.73779 | 5.51210 |
| CD630_13920 | CD630_n00790 | 188 -- 201 | 209 -- 222 | -15.0434 | 3.25236 | 5.51210 |
| CD630_30120 | CD630_n00790 | 113 -- 149 | 873 -- 918 | -15.0334 | 4.04859 | 5.51210 |
| cdu2 | CD630_n00790 | 40 -- 88 | 202 -- 243 | -15.016 | 2.97335 | 5.51210 |
| CD630_13380 | CD630_n00790 | 109 -- 149 | 202 -- 242 | -14.9496 | 4.74408 | 5.51210 |
| CD630_35260 | CD630_n00790 | 65 -- 97 | 477 -- 498 | -14.9065 | 5.77277 | 5.51210 |
| CD630_16670 | CD630_n00790 | 71 -- 110 | 994 -- 1034 | -14.8738 | 3.33553 | 5.51210 |
| acoL | CD630_n00790 | 142 -- 201 | 212 -- 274 | -14.8432 | 3.32169 | 5.51210 |
| glgB | CD630_n00790 | 159 -- 201 | 206 -- 247 | -14.8268 | 4.48273 | 5.51210 |
| CD630_32710 | CD630_n00790 | 114 -- 159 | 209 -- 247 | -14.8115 | 4.74276 | 5.51210 |
| CD630_18610 | CD630_n00790 | 30 -- 92 | 199 -- 258 | -14.7736 | 4.15582 | 5.51210 |
| CD630_31910 | CD630_n00790 | 48 -- 107 | 200 -- 253 | -14.7364 | 3.48314 | 5.51210 |
| CD630_30140 | CD630_n00790 | 43 -- 95 | 208 -- 252 | -14.6878 | 3.28343 | 5.51210 |
| def1 | CD630_n00790 | 68 -- 156 | 162 -- 247 | -14.625 | 3.99720 | 5.51210 |
| CD630_29760 | CD630_n00790 | 80 -- 163 | 195 -- 278 | -14.5284 | 3.51074 | 5.51210 |
| CD630_09100 | CD630_n00790 | 4 -- 78 | 586 -- 646 | -14.5283 | 5.92260 | 5.51210 |
| cwp29 | CD630_n00790 | 10 -- 43 | 205 -- 240 | -14.4557 | 4.40865 | 5.51210 |
| CD630_07470 | CD630_n00790 | 43 -- 57 | 1060 -- 1074 | -14.4555 | 3.37143 | 5.51210 |
| CD630_20600 | CD630_n00790 | 76 -- 106 | 205 -- 242 | -14.455 | 4.20093 | 5.51210 |
| nrdD | CD630_n00790 | 57 -- 89 | 996 -- 1020 | -14.4392 | 3.28290 | 5.51210 |
| CD630_14920 | CD630_n00790 | 41 -- 115 | 205 -- 280 | -14.4318 | 4.35225 | 5.51210 |
| CD630_32690 | CD630_n00790 | 81 -- 125 | 998 -- 1041 | -14.418 | 7.32676 | 5.51210 |
| CD630_14260 | CD630_n00790 | 27 -- 69 | 883 -- 919 | -14.3871 | 3.22214 | 5.51210 |
| CD630_08130 | CD630_n00790 | 33 -- 90 | 203 -- 255 | -14.3803 | 4.63831 | 5.51210 |
| CD630_05150 | CD630_n00790 | 80 -- 109 | 202 -- 228 | -14.3035 | 3.69875 | 5.51210 |
| CD630_06060 | CD630_n00790 | 26 -- 46 | 214 -- 239 | -14.2411 | 4.94050 | 5.51210 |
| CD630_09570 | CD630_n00790 | 16 -- 28 | 199 -- 211 | -14.239 | 5.31213 | 5.51210 |
| CD630_29070 | CD630_n00790 | 16 -- 28 | 199 -- 211 | -14.239 | 3.89477 | 5.51210 |
| CD630_25610 | CD630_n00790 | 76 -- 137 | 212 -- 290 | -14.2272 | 3.02685 | 5.51210 |
| CD630_05460 | CD630_n00790 | 18 -- 161 | 214 -- 362 | -14.203 | 3.98753 | 5.51210 |
| CD630_09850 | CD630_n00790 | 167 -- 201 | 2 -- 42 | -14.1929 | 4.18679 | 5.51210 |
| CD630_17360 | CD630_n00790 | 14 -- 137 | 216 -- 337 | -14.1767 | 5.66070 | 5.51210 |
| nusB | CD630_n00790 | 44 -- 115 | 199 -- 282 | -14.1694 | 4.92798 | 5.51210 |
| CD630_14180 | CD630_n00790 | 174 -- 196 | 398 -- 420 | -14.1157 | 3.93696 | 5.51210 |
| CD630_21760 | CD630_n00790 | 50 -- 94 | 994 -- 1036 | -14.0885 | 4.43846 | 5.51210 |
| CD630_28320 | CD630_n00790 | 59 -- 126 | 160 -- 239 | -14.0836 | 3.99184 | 5.51210 |
| CD630_23080 | CD630_n00790 | 166 -- 194 | 210 -- 240 | -14.0688 | 5.23027 | 5.51210 |
| argS | CD630_n00790 | 9 -- 93 | 201 -- 287 | -14.0661 | 4.96036 | 5.51210 |
| CD630_27770 | CD630_n00790 | 78 -- 95 | 210 -- 225 | -14.03 | 4.08588 | 5.51210 |
| CD630_17710 | CD630_n00790 | 75 -- 138 | 730 -- 797 | -13.9676 | 4.37688 | 5.51210 |
| CD630_17292 | CD630_n00790 | 81 -- 200 | 84 -- 217 | -13.9523 | 6.25799 | 5.51210 |
| CD630_20171 | CD630_n00790 | 86 -- 165 | 200 -- 290 | -13.9285 | 6.87491 | 5.51210 |
| CD630_03220 | CD630_n00790 | 86 -- 119 | 205 -- 253 | -13.9274 | 3.36746 | 5.51210 |
| CD630_15160 | CD630_n00790 | 4 -- 44 | 877 -- 919 | -13.9243 | 4.32318 | 5.51210 |
| CD630_06050 | CD630_n00790 | 120 -- 169 | 1006 -- 1047 | -13.8799 | 3.59332 | 5.51210 |
| CD630_19690 | CD630_n00790 | 96 -- 130 | 196 -- 227 | -13.8663 | 3.85861 | 5.51210 |
| metQ | CD630_n00790 | 67 -- 101 | 1007 -- 1041 | -13.8553 | 3.87190 | 5.51210 |
| CD630_04280 | CD630_n00790 | 53 -- 88 | 109 -- 145 | -13.843 | 3.61457 | 5.51210 |
| CD630_28550 | CD630_n00790 | 68 -- 108 | 997 -- 1041 | -13.8274 | 3.51196 | 5.51210 |
| ssuC | CD630_n00790 | 55 -- 102 | 193 -- 242 | -13.7906 | 3.45491 | 5.51210 |
| CD630_14210 | CD630_n00790 | 4 -- 28 | 139 -- 163 | -13.7635 | 3.82355 | 5.51210 |
| CD630_28260 | CD630_n00790 | 45 -- 101 | 202 -- 274 | -13.7453 | 3.51613 | 5.51210 |
| dnaC | CD630_n00790 | 109 -- 156 | 207 -- 241 | -13.7385 | 6.30796 | 5.51210 |
| CD630_29650 | CD630_n00790 | 63 -- 129 | 192 -- 246 | -13.7102 | 3.67128 | 5.51210 |
| CD630_08390 | CD630_n00790 | 40 -- 104 | 231 -- 292 | -13.6949 | 5.16000 | 5.51210 |
| CD630_21490 | CD630_n00790 | 72 -- 110 | 211 -- 249 | -13.6908 | 3.37462 | 5.51210 |
| CD630_15080 | CD630_n00790 | 168 -- 197 | 468 -- 496 | -13.6658 | 4.03033 | 5.51210 |
| CD630_10220 | CD630_n00790 | 132 -- 166 | 846 -- 881 | -13.5986 | 4.08233 | 5.51210 |
| CD630_26920 | CD630_n00790 | 124 -- 166 | 268 -- 323 | -13.5795 | 4.58060 | 5.51210 |
| CD630_07770 | CD630_n00790 | 90 -- 121 | 198 -- 218 | -13.5666 | 3.31158 | 5.51210 |
| CD630_29250 | CD630_n00790 | 170 -- 199 | 18 -- 48 | -13.5334 | 4.61901 | 5.51210 |
| CD630_09410 | CD630_n00790 | 170 -- 199 | 18 -- 48 | -13.5334 | 4.75651 | 5.51210 |
| malY | CD630_n00790 | 61 -- 80 | 477 -- 496 | -13.5208 | 4.46345 | 5.51210 |
| CD630_23770 | CD630_n00790 | 59 -- 121 | 193 -- 254 | -13.4471 | 3.25181 | 5.51210 |
| CD630_13600 | CD630_n00790 | 75 -- 181 | 149 -- 248 | -13.4353 | 4.45481 | 5.51210 |
| CD630_28560 | CD630_n00790 | 109 -- 200 | 235 -- 339 | -13.3726 | 3.96803 | 5.51210 |
| CD630_27220 | CD630_n00790 | 46 -- 101 | 207 -- 262 | -13.352 | 4.18786 | 5.51210 |
| CD630_36180 | CD630_n00790 | 169 -- 184 | 266 -- 282 | -13.342 | 4.83428 | 5.51210 |
| CD630_15360 | CD630_n00790 | 113 -- 164 | 731 -- 784 | -13.3342 | 4.47208 | 5.51210 |
| CD630_19700 | CD630_n00790 | 35 -- 91 | 997 -- 1039 | -13.3168 | 3.85861 | 5.51210 |
| CD630_10080 | CD630_n00790 | 32 -- 106 | 187 -- 244 | -13.3161 | 3.75625 | 5.51210 |
| bglA | CD630_n00790 | 152 -- 178 | 477 -- 502 | -13.3008 | 4.86402 | 5.51210 |
| CD630_20670 | CD630_n00790 | 13 -- 158 | 214 -- 354 | -13.254 | 4.42757 | 5.51210 |
| appF | CD630_n00790 | 110 -- 180 | 202 -- 272 | -13.233 | 4.25570 | 5.51210 |
| CD630_23450 | CD630_n00790 | 147 -- 189 | 540 -- 574 | -13.2246 | 3.27218 | 5.51210 |
| CD630_33570 | CD630_n00790 | 131 -- 172 | 731 -- 776 | -13.2077 | 3.95778 | 5.51210 |
| CD630_16520 | CD630_n00790 | 86 -- 186 | 174 -- 275 | -13.1948 | 4.28395 | 5.51210 |
| pgm | CD630_n00790 | 100 -- 177 | 211 -- 319 | -13.1755 | 3.45439 | 5.51210 |
| CD630_32050 | CD630_n00790 | 156 -- 196 | 217 -- 262 | -13.0793 | 5.73797 | 5.51210 |
| CD630_14810 | CD630_n00790 | 119 -- 153 | 207 -- 239 | -13.0769 | 3.35323 | 5.51210 |
| CD630_02420 | CD630_n00790 | 89 -- 145 | 214 -- 274 | -13.0588 | 3.71048 | 5.51210 |
| CD630_21730 | CD630_n00790 | 56 -- 196 | 151 -- 252 | -13.0535 | 3.45076 | 5.51210 |
| CD630_09970 | CD630_n00790 | 170 -- 190 | 538 -- 556 | -13.0067 | 3.25917 | 5.51210 |
| garK | CD630_n00790 | 4 -- 26 | 467 -- 489 | -13.0001 | 3.72836 | 5.51210 |
| CD630_18930 | CD630_n00790 | 87 -- 121 | 995 -- 1023 | -12.9895 | 5.32127 | 5.51210 |
| serS1 | CD630_n00790 | 128 -- 170 | 210 -- 247 | -12.9768 | 3.45858 | 5.51210 |
| CD630_21460 | CD630_n00790 | 121 -- 195 | 962 -- 1032 | -12.9638 | 4.83005 | 5.51210 |
| CD630_25320 | CD630_n00790 | 135 -- 194 | 977 -- 1036 | -12.9444 | 4.29415 | 5.51210 |
| CD630_18910 | CD630_n00790 | 169 -- 201 | 244 -- 274 | -12.9209 | 3.30688 | 5.51210 |
| CD630_18311 | CD630_n00790 | 88 -- 109 | 202 -- 216 | -12.9071 | 6.86047 | 5.51210 |
| CD630_03170 | CD630_n00790 | 163 -- 181 | 396 -- 418 | -12.903 | 3.66946 | 5.51210 |
| CD630_09040 | CD630_n00790 | 154 -- 186 | 199 -- 228 | -12.8803 | 4.11851 | 5.51210 |
| CD630_09772 | CD630_n00790 | 146 -- 166 | 341 -- 361 | -12.8614 | 3.66033 | 5.51210 |
| CD630_06970 | CD630_n00790 | 92 -- 141 | 204 -- 251 | -12.8384 | 3.82312 | 5.51210 |
| CD630_21330 | CD630_n00790 | 92 -- 151 | 214 -- 274 | -12.8162 | 4.10111 | 5.51210 |
| CD630_07400 | CD630_n00790 | 6 -- 64 | 125 -- 179 | -12.7839 | 4.52273 | 5.51210 |
| CD630_24050 | CD630_n00790 | 111 -- 131 | 113 -- 133 | -12.7377 | 3.70192 | 5.51210 |
| CD630_19440 | CD630_n00790 | 144 -- 187 | 377 -- 410 | -12.7364 | 4.53937 | 5.51210 |
| CD630_11220 | CD630_n00790 | 138 -- 192 | 997 -- 1043 | -12.722 | 3.52327 | 5.51210 |
| CD630_30360 | CD630_n00790 | 50 -- 116 | 1058 -- 1121 | -12.7136 | 4.26509 | 5.51210 |
| CD630_06150 | CD630_n00790 | 88 -- 113 | 198 -- 218 | -12.6987 | 5.51744 | 5.51210 |
| CD630_08510 | CD630_n00790 | 60 -- 96 | 209 -- 247 | -12.6804 | 5.16329 | 5.51210 |
| CD630_15550 | CD630_n00790 | 98 -- 116 | 1005 -- 1025 | -12.6709 | 3.49019 | 5.51210 |
| CD630_21910 | CD630_n00790 | 139 -- 187 | 221 -- 289 | -12.659 | 4.66600 | 5.51210 |
| spoVFB | CD630_n00790 | 102 -- 200 | 201 -- 308 | -12.6578 | 4.62758 | 5.51210 |
| CD630_06080 | CD630_n00790 | 103 -- 185 | 201 -- 274 | -12.6529 | 3.82315 | 5.51210 |
| CD630_01560 | CD630_n00790 | 85 -- 147 | 210 -- 274 | -12.6509 | 4.06352 | 5.51210 |
| ribD | CD630_n00790 | 99 -- 176 | 210 -- 302 | -12.6469 | 6.65922 | 5.51210 |
| CD630_26150 | CD630_n00790 | 163 -- 196 | 20 -- 61 | -12.6212 | 4.06302 | 5.51210 |
| CD630_21240 | CD630_n00790 | 111 -- 135 | 112 -- 133 | -12.6194 | 5.88879 | 5.51210 |
| CD630_30730 | CD630_n00790 | 8 -- 39 | 129 -- 155 | -12.6172 | 3.13236 | 5.51210 |
| cls | CD630_n00790 | 168 -- 196 | 629 -- 659 | -12.5983 | 4.46177 | 5.51210 |
| CD630_28870 | CD630_n00790 | 37 -- 139 | 482 -- 598 | -12.5849 | 3.41991 | 5.51210 |
| CD630_24770 | CD630_n00790 | 127 -- 181 | 213 -- 277 | -12.583 | 3.20798 | 5.51210 |
| aroD | CD630_n00790 | 92 -- 109 | 202 -- 218 | -12.5828 | 3.71929 | 5.51210 |
| CD630_27630 | CD630_n00790 | 41 -- 78 | 233 -- 265 | -12.555 | 3.54199 | 5.51210 |
| CD630_03420 | CD630_n00790 | 183 -- 200 | 216 -- 232 | -12.5542 | 4.71868 | 5.51210 |
| CD630_09840 | CD630_n00790 | 15 -- 55 | 549 -- 595 | -12.5538 | 4.50009 | 5.51210 |
| bltD | CD630_n00790 | 14 -- 95 | 100 -- 185 | -12.4906 | 4.49926 | 5.51210 |
| blaI | CD630_n00790 | 79 -- 146 | 224 -- 292 | -12.4813 | 3.73674 | 5.51210 |
| CD630_06110 | CD630_n00790 | 84 -- 94 | 199 -- 209 | -12.4319 | 5.61380 | 5.51210 |
| CD630_27110 | CD630_n00790 | 119 -- 155 | 2 -- 43 | -12.4306 | 3.78231 | 5.51210 |
| CD630_25200 | CD630_n00790 | 81 -- 107 | 475 -- 500 | -12.4055 | 3.65158 | 5.51210 |
| CD630_11730 | CD630_n00790 | 57 -- 101 | 205 -- 248 | -12.4033 | 4.80521 | 5.51210 |
| CD630_06920 | CD630_n00790 | 51 -- 199 | 207 -- 340 | -12.3879 | 3.13260 | 5.51210 |
| CD630_09760 | CD630_n00790 | 77 -- 112 | 199 -- 231 | -12.3718 | 4.83410 | 5.51210 |
| CD630_28910 | CD630_n00790 | 77 -- 112 | 199 -- 231 | -12.3718 | 4.51664 | 5.51210 |
| CD630_05820 | CD630_n00790 | 22 -- 109 | 195 -- 283 | -12.3689 | 3.75661 | 5.51210 |
| CD630_01570 | CD630_n00790 | 128 -- 148 | 131 -- 150 | -12.3429 | 6.08699 | 5.51210 |
| CD630_19680 | CD630_n00790 | 8 -- 110 | 201 -- 279 | -12.3349 | 3.58080 | 5.51210 |
| CD630_08090 | CD630_n00790 | 87 -- 150 | 727 -- 782 | -12.332 | 3.57167 | 5.51210 |
| CD630_05950 | CD630_n00790 | 76 -- 112 | 469 -- 501 | -12.3265 | 3.31669 | 5.51210 |
| CD630_03040 | CD630_n00790 | 116 -- 147 | 1000 -- 1027 | -12.3233 | 3.51403 | 5.51210 |
| pilD | CD630_n00790 | 2 -- 42 | 999 -- 1035 | -12.3208 | 4.54123 | 5.51210 |
| pilC | CD630_n00790 | 16 -- 51 | 998 -- 1047 | -12.3043 | 3.54230 | 5.51210 |
| CD630_30310 | CD630_n00790 | 58 -- 100 | 194 -- 229 | -12.3013 | 4.56059 | 5.51210 |
| CD630_04940 | CD630_n00790 | 61 -- 92 | 105 -- 133 | -12.2988 | 4.20948 | 5.51210 |
| CD630_20920 | CD630_n00790 | 110 -- 170 | 998 -- 1065 | -12.2947 | 4.13812 | 5.51210 |
| cheW | CD630_n00790 | 11 -- 109 | 197 -- 316 | -12.2794 | 3.72371 | 5.51210 |
| etfA | CD630_n00790 | 22 -- 37 | 479 -- 495 | -12.2423 | 4.80521 | 5.51210 |
| cspD | CD630_n00790 | 116 -- 155 | 259 -- 291 | -12.237 | 3.23587 | 5.51210 |
| appC | CD630_n00790 | 63 -- 121 | 119 -- 174 | -12.2218 | 4.50438 | 5.51210 |
| agrB | CD630_n00790 | 168 -- 185 | 629 -- 646 | -12.2206 | 3.92598 | 5.51210 |
| CD630_08360 | CD630_n00790 | 35 -- 65 | 265 -- 300 | -12.2109 | 3.19339 | 5.51210 |
| dapB2 | CD630_n00790 | 182 -- 199 | 95 -- 111 | -12.1979 | 3.27218 | 5.51210 |
| CD630_02740 | CD630_n00790 | 75 -- 96 | 197 -- 221 | -12.1799 | 5.02277 | 5.51210 |
| CD630_14250 | CD630_n00790 | 160 -- 186 | 204 -- 229 | -12.1606 | 4.19604 | 5.51210 |
| CD630_27060 | CD630_n00790 | 58 -- 134 | 994 -- 1045 | -12.1258 | 4.06049 | 5.51210 |
| kdgT1 | CD630_n00790 | 32 -- 122 | 110 -- 175 | -12.1252 | 4.49207 | 5.51210 |
| CD630_21020 | CD630_n00790 | 87 -- 105 | 202 -- 218 | -12.1152 | 4.28666 | 5.51210 |
| CD630_03930 | CD630_n00790 | 142 -- 172 | 915 -- 939 | -12.1081 | 4.15934 | 5.51210 |
| CD630_15850 | CD630_n00790 | 121 -- 133 | 200 -- 212 | -12.0977 | 4.32968 | 5.51210 |
| CD630_22890 | CD630_n00790 | 30 -- 45 | 549 -- 565 | -12.0937 | 3.75591 | 5.51210 |
| CD630_19800 | CD630_n00790 | 10 -- 90 | 199 -- 272 | -12.0771 | 5.22035 | 5.51210 |
| CD630_14050 | CD630_n00790 | 104 -- 199 | 202 -- 312 | -12.0701 | 3.30894 | 5.51210 |
| moaA | CD630_n00790 | 4 -- 21 | 211 -- 225 | -12.0694 | 3.83799 | 5.51210 |
| CD630_08100 | CD630_n00790 | 180 -- 197 | 479 -- 495 | -12.0574 | 4.39181 | 5.51210 |
| CD630_13560 | CD630_n00790 | 19 -- 27 | 200 -- 208 | -12.0571 | 3.38083 | 5.51210 |
| CD630_06200 | CD630_n00790 | 87 -- 147 | 202 -- 276 | -12.0345 | 3.57064 | 5.51210 |
| CD630_06690 | CD630_n00790 | 170 -- 186 | 997 -- 1013 | -12.0323 | 5.82709 | 5.51210 |
| CD630_26250 | CD630_n00790 | 137 -- 184 | 381 -- 421 | -11.9981 | 5.03379 | 5.51210 |
| CD630_23090 | CD630_n00790 | 17 -- 36 | 221 -- 239 | -11.9791 | 4.25622 | 5.51210 |
| CD630_22200 | CD630_n00790 | 87 -- 99 | 206 -- 218 | -11.9783 | 4.18288 | 5.51210 |
| htpG | CD630_n00790 | 31 -- 137 | 977 -- 1068 | -11.9722 | 3.45147 | 5.51210 |
| cspC | CD630_n00790 | 116 -- 174 | 207 -- 291 | -11.9686 | 5.48848 | 5.51210 |
| CD630_29270 | CD630_n00790 | 37 -- 52 | 201 -- 219 | -11.9651 | 3.70174 | 5.51210 |
| kdpD | CD630_n00790 | 146 -- 200 | 219 -- 273 | -11.9519 | 4.27140 | 5.51210 |
| CD630_09770 | CD630_n00790 | 1 -- 42 | 467 -- 502 | -11.9144 | 3.22106 | 5.51210 |
| CD630_27230 | CD630_n00790 | 93 -- 151 | 798 -- 852 | -11.9054 | 3.52462 | 5.51210 |
| CD630_31461 | CD630_n00790 | 141 -- 201 | 124 -- 171 | -11.8957 | 4.41476 | 5.51210 |
| CD630_06060 | CD630_n00790 | 102 -- 127 | 334 -- 363 | -11.8749 | 4.94050 | 5.51210 |
| CD630_20960 | CD630_n00790 | 40 -- 183 | 202 -- 330 | -11.862 | 5.65940 | 5.51210 |
| adhE | CD630_n00790 | 157 -- 196 | 780 -- 823 | -11.8564 | 3.06485 | 5.51210 |
| CD630_29750 | CD630_n00790 | 71 -- 94 | 111 -- 135 | -11.8496 | 4.59146 | 5.51210 |
| CD630_32700 | CD630_n00790 | 43 -- 183 | 200 -- 313 | -11.8484 | 4.74276 | 5.51210 |
| CD630_30130 | CD630_n00790 | 7 -- 109 | 202 -- 274 | -11.8272 | 3.28343 | 5.51210 |
| CD630_17770 | CD630_n00790 | 169 -- 199 | 279 -- 313 | -11.8107 | 5.06490 | 5.51210 |
| CD630_18950 | CD630_n00790 | 88 -- 167 | 204 -- 274 | -11.8057 | 3.40610 | 5.51210 |
| CD630_12110 | CD630_n00790 | 3 -- 12 | 280 -- 289 | -11.7873 | 4.36078 | 5.51210 |
| CD630_14630 | CD630_n00790 | 130 -- 177 | 994 -- 1031 | -11.7863 | 3.34485 | 5.51210 |
| CD630_25040 | CD630_n00790 | 97 -- 171 | 739 -- 829 | -11.7838 | 3.43898 | 5.51210 |
| CD630_30420 | CD630_n00790 | 166 -- 199 | 737 -- 765 | -11.7503 | 5.27539 | 5.51210 |
| CD630_27200 | CD630_n00790 | 110 -- 137 | 981 -- 1001 | -11.7474 | 3.42516 | 5.51210 |
| CD630_27240 | CD630_n00790 | 158 -- 176 | 128 -- 147 | -11.733 | 3.52462 | 5.51210 |
| CD630_25050 | CD630_n00790 | 83 -- 94 | 199 -- 210 | -11.7317 | 4.33624 | 5.51210 |
| CD630_18660 | CD630_n00790 | 125 -- 136 | 403 -- 415 | -11.7236 | 3.50371 | 5.51210 |
| CD630_04410 | CD630_n00790 | 151 -- 171 | 337 -- 356 | -11.7231 | 3.90451 | 5.51210 |
| CD630_31880 | CD630_n00790 | 110 -- 125 | 1058 -- 1073 | -11.7162 | 3.97522 | 5.51210 |
| CD630_14990 | CD630_n00790 | 133 -- 153 | 214 -- 239 | -11.7042 | 3.78619 | 5.51210 |
| CD630_28330 | CD630_n00790 | 66 -- 107 | 221 -- 260 | -11.701 | 3.29122 | 5.51210 |
| tcdC | CD630_n00790 | 134 -- 151 | 266 -- 288 | -11.6954 | 3.76101 | 5.51210 |
| CD630_03920 | CD630_n00790 | 151 -- 201 | 221 -- 263 | -11.6855 | 4.15934 | 5.51210 |
| CD630_10790 | CD630_n00790 | 183 -- 201 | 532 -- 546 | -11.6499 | 4.45284 | 5.51210 |
| CD630_27600 | CD630_n00790 | 68 -- 131 | 724 -- 788 | -11.6205 | 3.48384 | 5.51210 |
| CD630_11240 | CD630_n00790 | 68 -- 99 | 1005 -- 1033 | -11.6122 | 3.20274 | 5.51210 |
| CD630_10800 | CD630_n00790 | 41 -- 64 | 469 -- 490 | -11.611 | 5.30319 | 5.51210 |
| CD630_30540 | CD630_n00790 | 167 -- 183 | 467 -- 483 | -11.6096 | 3.63804 | 5.51210 |
| cwp17 | CD630_n00790 | 57 -- 115 | 194 -- 247 | -11.6011 | 3.43363 | 5.51210 |
| CD630_18960 | CD630_n00790 | 19 -- 53 | 207 -- 240 | -11.5984 | 4.37822 | 5.51210 |
| CD630_05300 | CD630_n00790 | 117 -- 147 | 36 -- 68 | -11.5772 | 4.56885 | 5.51210 |
| CD630_04320 | CD630_n00790 | 74 -- 91 | 731 -- 747 | -11.5603 | 3.68342 | 5.51210 |
| CD630_16540 | CD630_n00790 | 86 -- 148 | 154 -- 204 | -11.5587 | 4.37658 | 5.51210 |
| CD630_07121 | CD630_n00790 | 29 -- 78 | 598 -- 655 | -11.5482 | 3.26177 | 5.51210 |
| CD630_19790 | CD630_n00790 | 26 -- 95 | 200 -- 274 | -11.538 | 3.79593 | 5.51210 |
| cprK | CD630_n00790 | 83 -- 147 | 217 -- 276 | -11.5365 | 3.96216 | 5.51210 |
| CD630_05370 | CD630_n00790 | 126 -- 137 | 412 -- 423 | -11.523 | 3.26947 | 5.51210 |
| pilT | CD630_n00790 | 150 -- 199 | 211 -- 262 | -11.5104 | 3.24639 | 5.51210 |
| CD630_05590 | CD630_n00790 | 92 -- 111 | 199 -- 218 | -11.4971 | 3.39670 | 5.51210 |
| bglA | CD630_n00790 | 170 -- 188 | 261 -- 285 | -11.4893 | 4.50315 | 5.51210 |
| CD630_15461 | CD630_n00790 | 3 -- 29 | 268 -- 292 | -11.4887 | 9.13442 | 5.51210 |
| CD630_29960 | CD630_n00790 | 140 -- 193 | 196 -- 269 | -11.4685 | 4.51904 | 5.51210 |
| CD630_13360 | CD630_n00790 | 159 -- 193 | 119 -- 158 | -11.4601 | 3.77330 | 5.51210 |
| CD630_10970 | CD630_n00790 | 183 -- 199 | 475 -- 489 | -11.4313 | 3.53850 | 5.51210 |
| CD630_24591 | CD630_n00790 | 30 -- 48 | 630 -- 644 | -11.4244 | 3.54497 | 5.51210 |
| CD630_22660 | CD630_n00790 | 185 -- 195 | 281 -- 291 | -11.4239 | 3.60496 | 5.51210 |
| cbiQ | CD630_n00790 | 145 -- 176 | 213 -- 251 | -11.3954 | 3.79830 | 5.51210 |
| CD630_14610 | CD630_n00790 | 139 -- 164 | 339 -- 365 | -11.387 | 4.89428 | 5.51210 |
| CD630_26750 | CD630_n00790 | 93 -- 177 | 367 -- 446 | -11.3636 | 3.60610 | 5.51210 |
| CD630_06350 | CD630_n00790 | 75 -- 118 | 268 -- 313 | -11.3589 | 4.35983 | 5.51210 |
| CD630_28290 | CD630_n00790 | 159 -- 178 | 630 -- 648 | -11.3502 | 5.54192 | 5.51210 |
| CD630_23820 | CD630_n00790 | 26 -- 38 | 1015 -- 1027 | -11.3413 | 3.18419 | 5.51210 |
| CD630_31480 | CD630_n00790 | 71 -- 95 | 212 -- 241 | -11.3404 | 4.88897 | 5.51210 |
| cobC | CD630_n00790 | 167 -- 198 | 1001 -- 1030 | -11.3388 | 3.15854 | 5.51210 |
| CD630_21230 | CD630_n00790 | 77 -- 113 | 1001 -- 1028 | -11.3306 | 3.89213 | 5.51210 |
| CD630_02790 | CD630_n00790 | 86 -- 162 | 730 -- 812 | -11.3067 | 4.77661 | 5.51210 |
| CD630_33300 | CD630_n00790 | 130 -- 147 | 554 -- 570 | -11.2915 | 3.79149 | 5.51210 |
| CD630_14360 | CD630_n00790 | 86 -- 101 | 202 -- 218 | -11.2822 | 3.02079 | 5.51210 |
| CD630_21540 | CD630_n00790 | 69 -- 120 | 631 -- 687 | -11.2753 | 4.33090 | 5.51210 |
| CD630_23280 | CD630_n00790 | 113 -- 184 | 960 -- 1025 | -11.2681 | 3.50093 | 5.51210 |
| CD630_04610 | CD630_n00790 | 147 -- 199 | 5 -- 42 | -11.25 | 3.83025 | 5.51210 |

| | | | | | | |
|---|---|---|---|---|---|---|
| CD630_13780 | CD630_n00790 | 113 -- 180 | 203 -- 274 | -11.247 | 3.52274 | 5.51210 |
| CD630_05830 | CD630_n00790 | 8 -- 28 | 222 -- 242 | -11.242 | 3.35360 | 5.51210 |
| CD630_10950 | CD630_n00790 | 187 -- 195 | 410 -- 418 | -11.2187 | 3.67675 | 5.51210 |
| CD630_09060 | CD630_n00790 | 11 -- 91 | 201 -- 279 | -11.1986 | 4.31834 | 5.51210 |
| CD630_32880 | CD630_n00790 | 179 -- 201 | 872 -- 893 | -11.1896 | 3.41276 | 5.51210 |
| cbpA | CD630_n00790 | 84 -- 96 | 200 -- 218 | -11.1814 | 3.03040 | 5.51210 |
| glnA | CD630_n00790 | 66 -- 108 | 998 -- 1035 | -11.1549 | 3.87629 | 5.51210 |
| CD630_32040 | CD630_n00790 | 1 -- 36 | 466 -- 501 | -11.147 | 4.93516 | 5.51210 |
| CD630_13530 | CD630_n00790 | 111 -- 124 | 1056 -- 1069 | -11.1467 | 4.09832 | 5.51210 |
| CD630_31000 | CD630_n00790 | 43 -- 109 | 731 -- 796 | -11.1346 | 3.70946 | 5.51210 |
| CD630_28580 | CD630_n00790 | 152 -- 191 | 6 -- 44 | -11.1274 | 4.67130 | 5.51210 |
| vncR | CD630_n00790 | 104 -- 131 | 349 -- 374 | -11.1263 | 3.93642 | 5.51210 |
| flgB | CD630_n00790 | 122 -- 141 | 1000 -- 1025 | -11.1085 | 3.25736 | 5.51210 |
| CD630_06170 | CD630_n00790 | 14 -- 154 | 958 -- 1067 | -11.1055 | 4.95366 | 5.51210 |
| CD630_05030 | CD630_n00790 | 133 -- 155 | 1019 -- 1037 | -11.1039 | 4.03823 | 5.51210 |
| glgP | CD630_n00790 | 9 -- 38 | 1000 -- 1025 | -11.0815 | 3.28530 | 5.51210 |
| CD630_14130 | CD630_n00790 | 97 -- 150 | 991 -- 1026 | -11.0698 | 4.57970 | 5.51210 |
| eutV | CD630_n00790 | 130 -- 178 | 1060 -- 1119 | -11.064 | 4.55715 | 5.51210 |
| CD630_14930 | CD630_n00790 | 5 -- 43 | 905 -- 941 | -11.0606 | 3.31335 | 5.51210 |
| CD630_13370 | CD630_n00790 | 18 -- 63 | 730 -- 774 | -11.0592 | 4.33145 | 5.51210 |
| rbsA | CD630_n00790 | 132 -- 151 | 397 -- 417 | -11.0462 | 3.71650 | 5.51210 |
| CD630_33480 | CD630_n00790 | 37 -- 75 | 118 -- 157 | -11.0135 | 3.67613 | 5.51210 |
| CD630_10830 | CD630_n00790 | 150 -- 200 | 998 -- 1040 | -10.9974 | 4.82398 | 5.51210 |
| CD630_21770 | CD630_n00790 | 148 -- 171 | 493 -- 515 | -10.9628 | 4.02587 | 5.51210 |
| CD630_04880 | CD630_n00790 | 147 -- 159 | 540 -- 552 | -10.9529 | 4.62291 | 5.51210 |
| oppB | CD630_n00790 | 53 -- 99 | 552 -- 598 | -10.9421 | 2.98328 | 5.51210 |
| CD630_05220 | CD630_n00790 | 181 -- 200 | 266 -- 286 | -10.9368 | 3.65057 | 5.51210 |
| CD630_25620 | CD630_n00790 | 44 -- 94 | 212 -- 248 | -10.9352 | 3.02685 | 5.51210 |
| CD630_18650 | CD630_n00790 | 88 -- 97 | 199 -- 208 | -10.9282 | 3.75892 | 5.51210 |
| CD630_17730 | CD630_n00790 | 81 -- 107 | 204 -- 224 | -10.9187 | 4.73397 | 5.51210 |
| CD630_04660 | CD630_n00790 | 88 -- 101 | 206 -- 219 | -10.9123 | 4.32791 | 5.51210 |
| terD | CD630_n00790 | 9 -- 31 | 137 -- 162 | -10.9005 | 3.33148 | 5.51210 |
| CD630_27520 | CD630_n00790 | 84 -- 109 | 202 -- 227 | -10.8841 | 4.27693 | 5.51210 |
| CD630_09990 | CD630_n00790 | 91 -- 106 | 205 -- 219 | -10.8613 | 3.86040 | 5.51210 |
| CD630_26020 | CD630_n00790 | 39 -- 114 | 1000 -- 1061 | -10.8461 | 4.03155 | 5.51210 |
| CD630_22850 | CD630_n00790 | 148 -- 196 | 4 -- 46 | -10.846 | 3.84102 | 5.51210 |
| uxaA%27 | CD630_n00790 | 48 -- 93 | 199 -- 245 | -10.8177 | 4.26165 | 5.51210 |
| CD630_19921 | CD630_n00790 | 46 -- 66 | 219 -- 237 | -10.8147 | 5.46345 | 5.51210 |
| pmi | CD630_n00790 | 41 -- 95 | 993 -- 1040 | -10.8015 | 3.86845 | 5.51210 |
| CD630_35780 | CD630_n00790 | 29 -- 66 | 996 -- 1027 | -10.801 | 3.50107 | 5.51210 |
| CD630_31970 | CD630_n00790 | 111 -- 132 | 551 -- 571 | -10.7715 | 3.24511 | 5.51210 |
| CD630_20450 | CD630_n00790 | 103 -- 136 | 85 -- 106 | -10.759 | 3.77388 | 5.51210 |
| CD630_26691 | CD630_n00790 | 61 -- 99 | 879 -- 913 | -10.7507 | 3.96230 | 5.51210 |
| CD630_33650 | CD630_n00790 | 150 -- 199 | 624 -- 673 | -10.7437 | 4.80565 | 5.51210 |
| CD630_06540 | CD630_n00790 | 164 -- 195 | 390 -- 422 | -10.7056 | 3.54673 | 5.51210 |
| CD630_30100 | CD630_n00790 | 37 -- 66 | 1014 -- 1044 | -10.6945 | 4.13234 | 5.51210 |
| CD630_21030 | CD630_n00790 | 89 -- 106 | 200 -- 217 | -10.6786 | 3.08733 | 5.51210 |
| CD630_17970 | CD630_n00790 | 114 -- 171 | 1018 -- 1078 | -10.6629 | 3.68802 | 5.51210 |
| CD630_20980 | CD630_n00790 | 87 -- 156 | 206 -- 272 | -10.6626 | 8.12847 | 5.51210 |
| CD630_18090 | CD630_n00790 | 179 -- 200 | 995 -- 1019 | -10.6542 | 4.23961 | 5.51210 |
| CD630_33600 | CD630_n00790 | 150 -- 193 | 277 -- 320 | -10.6504 | 3.73196 | 5.51210 |
| CD630_22991 | CD630_n00790 | 25 -- 46 | 1019 -- 1040 | -10.6467 | 6.03728 | 5.51210 |
| CD630_06560 | CD630_n00790 | 68 -- 139 | 976 -- 1045 | -10.6356 | 4.02532 | 5.51210 |
| spaK | CD630_n00790 | 77 -- 98 | 463 -- 487 | -10.6232 | 3.04530 | 5.51210 |
| CD630_29840 | CD630_n00790 | 20 -- 49 | 1004 -- 1035 | -10.622 | 4.48825 | 5.51210 |
| CD630_08460 | CD630_n00790 | 90 -- 141 | 727 -- 773 | -10.6166 | 4.15083 | 5.51210 |
| CD630_08070 | CD630_n00790 | 75 -- 105 | 477 -- 509 | -10.6098 | 4.87819 | 5.51210 |
| CD630_08140 | CD630_n00790 | 150 -- 197 | 883 -- 924 | -10.6034 | 3.30159 | 5.51210 |
| CD630_14190 | CD630_n00790 | 86 -- 136 | 154 -- 208 | -10.6019 | 3.78562 | 5.51210 |
| CD630_21140 | CD630_n00790 | 28 -- 42 | 999 -- 1013 | -10.5986 | 3.33304 | 5.51210 |
| CD630_25020 | CD630_n00790 | 117 -- 158 | 99 -- 133 | -10.5907 | 4.17721 | 5.51210 |
| CD630_07480 | CD630_n00790 | 80 -- 111 | 202 -- 232 | -10.5654 | 5.18321 | 5.51210 |
| CD630_31381 | CD630_n00790 | 158 -- 172 | 204 -- 218 | -10.5549 | 3.49137 | 5.51210 |
| CD630_22050 | CD630_n00790 | 89 -- 106 | 205 -- 218 | -10.5436 | 4.87639 | 5.51210 |
| CD630_24760 | CD630_n00790 | 104 -- 134 | 204 -- 237 | -10.5205 | 5.19485 | 5.51210 |
| CD630_22030 | CD630_n00790 | 86 -- 123 | 102 -- 139 | -10.4621 | 3.29934 | 5.51210 |
| CD630_11241 | CD630_n00790 | 149 -- 177 | 202 -- 232 | -10.4141 | 4.85723 | 5.51210 |
| CD630_05900 | CD630_n00790 | 8 -- 43 | 206 -- 233 | -10.4054 | 3.21934 | 5.51210 |
| CD630_17151 | CD630_n00790 | 18 -- 53 | 723 -- 756 | -10.3968 | 6.43124 | 5.51210 |
| CD630_33580 | CD630_n00790 | 70 -- 106 | 801 -- 841 | -10.3752 | 3.95778 | 5.51210 |
| CD630_22720 | CD630_n00790 | 79 -- 165 | 198 -- 284 | -10.3745 | 3.81804 | 5.51210 |
| CD630_19580 | CD630_n00790 | 172 -- 190 | 347 -- 362 | -10.3604 | 4.51902 | 5.51210 |
| CD630_33610 | CD630_n00790 | 27 -- 34 | 201 -- 208 | -10.359 | 3.79322 | 5.51210 |
| CD630_05490 | CD630_n00790 | 154 -- 178 | 194 -- 217 | -10.3425 | 3.89852 | 5.51210 |
| CD630_10000 | CD630_n00790 | 7 -- 29 | 565 -- 595 | -10.3374 | 3.86040 | 5.51210 |
| CD630_05230 | CD630_n00790 | 148 -- 181 | 216 -- 251 | -10.325 | 3.73787 | 5.51210 |
| CD630_14420 | CD630_n00790 | 17 -- 30 | 1000 -- 1013 | -10.3206 | 3.99497 | 5.51210 |
| CD630_22550 | CD630_n00790 | 28 -- 84 | 126 -- 173 | -10.3114 | 3.62379 | 5.51210 |
| CD630_26050 | CD630_n00790 | 48 -- 195 | 110 -- 256 | -10.308 | 6.99851 | 5.51210 |
| CD630_34530 | CD630_n00790 | 2 -- 22 | 1009 -- 1035 | -10.3012 | 3.83016 | 5.51210 |
| CD630_20541 | CD630_n00790 | 24 -- 54 | 215 -- 249 | -10.2636 | 5.74527 | 5.51210 |
| CD630_16750 | CD630_n00790 | 50 -- 64 | 17 -- 32 | -10.2618 | 5.04051 | 5.51210 |
| CD630_01690 | CD630_n00790 | 167 -- 189 | 125 -- 147 | -10.2345 | 4.97234 | 5.51210 |
| CD630_23930 | CD630_n00790 | 130 -- 194 | 978 -- 1040 | -10.2243 | 3.05650 | 5.51210 |
| kdpE | CD630_n00790 | 87 -- 99 | 266 -- 278 | -10.2194 | 4.27140 | 5.51210 |
| CD630_04770 | CD630_n00790 | 69 -- 94 | 200 -- 218 | -10.2153 | 3.58478 | 5.51210 |
| CD630_35840 | CD630_n00790 | 57 -- 65 | 489 -- 497 | -10.208 | 3.14170 | 5.51210 |
| CD630_16940 | CD630_n00790 | 82 -- 95 | 208 -- 221 | -10.2066 | 4.14665 | 5.51210 |
| gcvTPA | CD630_n00790 | 31 -- 113 | 202 -- 273 | -10.2044 | 4.77229 | 5.51210 |
| sspA | CD630_n00790 | 101 -- 130 | 993 -- 1022 | -10.1928 | 3.62689 | 5.51210 |
| CD630_17630 | CD630_n00790 | 190 -- 199 | 475 -- 484 | -10.1758 | 4.52408 | 5.51210 |
| cdtR | CD630_n00790 | 3 -- 10 | 629 -- 636 | -10.1753 | 5.84368 | 5.51210 |
| CD630_16460 | CD630_n00790 | 128 -- 162 | 995 -- 1021 | -10.169 | 4.64028 | 5.51210 |
| CD630_14182 | CD630_n00790 | 28 -- 49 | 1019 -- 1040 | -10.1654 | 4.82360 | 5.51210 |
| uxaA | CD630_n00790 | 82 -- 121 | 196 -- 237 | -10.1649 | 4.26165 | 5.51210 |
| CD630_33560 | CD630_n00790 | 75 -- 106 | 205 -- 245 | -10.1648 | 4.03147 | 5.51210 |
| CD630_19670 | CD630_n00790 | 114 -- 125 | 476 -- 487 | -10.1645 | 3.20718 | 5.51210 |
| CD630_18090 | CD630_n00790 | 78 -- 104 | 184 -- 212 | -10.1137 | 4.23961 | 5.51210 |
| rolB | CD630_n00790 | 76 -- 98 | 803 -- 823 | -10.1026 | 3.60270 | 5.51210 |
| CD630_17380 | CD630_n00790 | 73 -- 93 | 995 -- 1015 | -10.0894 | 5.03207 | 5.51210 |
| CD630_26050 | CD630_n00790 | 133 -- 148 | 483 -- 498 | -10.0674 | 6.99851 | 5.51210 |
| CD630_27120 | CD630_n00790 | 130 -- 165 | 469 -- 505 | -10.0474 | 4.43720 | 5.51210 |
| opuCC | CD630_n00790 | 78 -- 111 | 104 -- 133 | -10.033 | 4.07035 | 5.51210 |
| CD630_16450 | CD630_n00790 | 80 -- 93 | 199 -- 211 | -10.0171 | 4.57547 | 5.51210 |
| CD630_19904 | CD630_n00790 | 141 -- 186 | 204 -- 251 | -10.016 | 5.96526 | 5.51210 |
| CD630_04850 | CD630_n00790 | 51 -- 130 | 953 -- 1030 | -10.0112 | 4.40788 | 5.51210 |
| CD630_36690 | CD630_n00790 | 85 -- 93 | 200 -- 208 | -10.0013 | 3.53761 | 5.51210 |
| CD630_20280 | CD630_n00790 | 119 -- 170 | 617 -- 677 | -9.99406 | 3.46343 | 5.51210 |
| CD630_15890 | CD630_n00790 | 4 -- 36 | 998 -- 1031 | -9.96206 | 3.39962 | 5.51210 |
| CD630_22510 | CD630_n00790 | 163 -- 184 | 388 -- 409 | -9.95729 | 11.42763 | 5.51210 |
| CD630_31490 | CD630_n00790 | 103 -- 114 | 110 -- 121 | -9.8999 | 4.08593 | 5.51210 |
| appB | CD630_n00790 | 140 -- 174 | 885 -- 919 | -9.89952 | 6.44103 | 5.51210 |
| CD630_36781 | CD630_n00790 | 24 -- 63 | 214 -- 251 | -9.87643 | 3.75946 | 5.51210 |
| ung | CD630_n00790 | 100 -- 127 | 557 -- 586 | -9.84422 | 3.30908 | 5.51210 |
| CD630_33420 | CD630_n00790 | 12 -- 46 | 473 -- 507 | -9.84243 | 3.22095 | 5.51210 |
| CD630_27280 | CD630_n00790 | 159 -- 190 | 374 -- 417 | -9.83562 | 3.24738 | 5.51210 |
| CD630_06100 | CD630_n00790 | 58 -- 77 | 142 -- 163 | -9.8341 | 4.43918 | 5.51210 |
| CD630_16150 | CD630_n00790 | 85 -- 106 | 205 -- 219 | -9.82147 | 4.20614 | 5.51210 |
| CD630_14240 | CD630_n00790 | 107 -- 120 | 202 -- 214 | -9.81186 | 8.60766 | 5.51210 |
| CD630_20260 | CD630_n00790 | 110 -- 119 | 1061 -- 1070 | -9.80905 | 3.84162 | 5.51210 |
| CD630_32990 | CD630_n00790 | 39 -- 67 | 911 -- 942 | -9.80853 | 5.24925 | 5.51210 |
| rolA | CD630_n00790 | 33 -- 76 | 210 -- 241 | -9.80081 | 4.56362 | 5.51210 |
| CD630_09030 | CD630_n00790 | 129 -- 158 | 222 -- 274 | -9.78024 | 4.21282 | 5.51210 |
| CD630_14600 | CD630_n00790 | 20 -- 50 | 317 -- 346 | -9.75298 | 5.90348 | 5.51210 |
| CD630_05940 | CD630_n00790 | 83 -- 93 | 199 -- 209 | -9.7481 | 4.18595 | 5.51210 |
| dltB | CD630_n00790 | 39 -- 49 | 401 -- 411 | -9.73051 | 3.81485 | 5.51210 |
| CD630_13960 | CD630_n00790 | 145 -- 189 | 999 -- 1028 | -9.7097 | 3.30375 | 5.51210 |
| CD630_34560 | CD630_n00790 | 83 -- 94 | 209 -- 220 | -9.69674 | 4.02604 | 5.51210 |
| CD630_16330 | CD630_n00790 | 141 -- 185 | 139 -- 181 | -9.69164 | 4.11180 | 5.51210 |
| CD630_04540 | CD630_n00790 | 79 -- 93 | 202 -- 216 | -9.67634 | 3.33556 | 5.51210 |
| CD630_16530 | CD630_n00790 | 116 -- 175 | 701 -- 758 | -9.65395 | 4.37658 | 5.51210 |
| CD630_28150 | CD630_n00790 | 151 -- 166 | 668 -- 683 | -9.62605 | 4.18447 | 5.51210 |
| CD630_16950 | CD630_n00790 | 128 -- 149 | 733 -- 754 | -9.58257 | 4.42569 | 5.51210 |
| CD630_05000 | CD630_n00790 | 145 -- 154 | 211 -- 220 | -9.57647 | 4.02442 | 5.51210 |
| dnaD | CD630_n00790 | 134 -- 144 | 1063 -- 1073 | -9.55794 | 3.56078 | 5.51210 |
| virS | CD630_n00790 | 151 -- 169 | 865 -- 884 | -9.54337 | 4.74085 | 5.51210 |
| CD630_23000 | CD630_n00790 | 149 -- 185 | 998 -- 1034 | -9.54159 | 3.54130 | 5.51210 |
| CD630_07960 | CD630_n00790 | 176 -- 191 | 202 -- 214 | -9.52571 | 3.22537 | 5.51210 |
| CD630_21080 | CD630_n00790 | 90 -- 112 | 196 -- 216 | -9.5162 | 3.99742 | 5.51210 |
| CD630_30190 | CD630_n00790 | 165 -- 200 | 766 -- 806 | -9.50848 | 3.27394 | 5.51210 |
| pilT | CD630_n00910 | 158 -- 189 | 2 -- 38 | -10.0749 | 3.24639 | 4.43720 |
| CD630_27280 | CD630_n00930 | 51 -- 188 | 120 -- 266 | -17.0479 | 3.24738 | 5.79130 |
| CD630_14150 | CD630_n00930 | 40 -- 59 | 24 -- 43 | -16.6298 | 3.28251 | 5.79130 |
| CD630_31461 | CD630_n00930 | 128 -- 193 | 115 -- 162 | -14.8107 | 4.41476 | 5.79130 |
| CD630_06050 | CD630_n00930 | 132 -- 177 | 23 -- 64 | -14.7568 | 3.59332 | 5.79130 |
| CD630_30420 | CD630_n00930 | 57 -- 120 | 4 -- 68 | -14.6788 | 5.27539 | 5.79130 |
| CD630_06100 | CD630_n00930 | 92 -- 164 | 116 -- 175 | -14.62 | 4.43918 | 5.79130 |
| oppB | CD630_n00930 | 90 -- 147 | 8 -- 48 | -14.3218 | 2.98328 | 5.79130 |
| CD630_28181 | CD630_n00930 | 32 -- 70 | 27 -- 54 | -14.1597 | 3.91299 | 5.79130 |
| CD630_09840 | CD630_n00930 | 79 -- 128 | 12 -- 57 | -14.0316 | 4.50009 | 5.79130 |
| argS | CD630_n00930 | 52 -- 61 | 335 -- 344 | -13.9706 | 4.96036 | 5.79130 |
| ribD | CD630_n00930 | 165 -- 181 | 332 -- 349 | -13.5904 | 6.65922 | 5.79130 |
| scpB | CD630_n00930 | 5 -- 29 | 46 -- 70 | -13.5843 | 3.33560 | 5.79130 |
| flhB | CD630_n00930 | 21 -- 32 | 21 -- 32 | -13.5637 | 3.55524 | 5.79130 |
| CD630_02790 | CD630_n00930 | 186 -- 201 | 29 -- 43 | -13.5397 | 4.77661 | 5.79130 |
| CD630_13780 | CD630_n00930 | 164 -- 193 | 102 -- 133 | -13.53 | 3.52274 | 5.79130 |
| CD630_04540 | CD630_n00930 | 70 -- 140 | 104 -- 180 | -13.5091 | 3.33556 | 5.79130 |
| CD630_21720 | CD630_n00930 | 165 -- 190 | 311 -- 345 | -13.497 | 4.13030 | 5.79130 |
| CD630_22500 | CD630_n00930 | 11 -- 157 | 120 -- 247 | -13.4844 | 4.03688 | 5.79130 |
| CD630_26750 | CD630_n00930 | 68 -- 132 | 9 -- 68 | -13.3221 | 3.60610 | 5.79130 |
| CD630_20650 | CD630_n00930 | 145 -- 165 | 19 -- 39 | -13.1267 | 4.00083 | 5.79130 |
| CD630_14630 | CD630_n00930 | 59 -- 115 | 18 -- 72 | -13.1192 | 3.34485 | 5.79130 |
| gatC | CD630_n00930 | 75 -- 117 | 8 -- 61 | -13.1027 | 4.25143 | 5.79130 |
| CD630_05900 | CD630_n00930 | 21 -- 47 | 330 -- 357 | -12.9753 | 3.21934 | 5.79130 |
| CD630_02420 | CD630_n00930 | 9 -- 38 | 27 -- 57 | -12.9721 | 3.71048 | 5.79130 |
| CD630_03410 | CD630_n00930 | 38 -- 104 | 147 -- 238 | -12.8879 | 4.45994 | 5.79130 |
| CD630_10080 | CD630_n00930 | 143 -- 174 | 22 -- 60 | -12.8567 | 3.75625 | 5.79130 |
| CD630_30100 | CD630_n00930 | 39 -- 76 | 29 -- 63 | -12.8054 | 4.13234 | 5.79130 |
| CD630_28320 | CD630_n00930 | 88 -- 100 | 332 -- 344 | -12.78 | 3.99184 | 5.79130 |
| CD630_15111 | CD630_n00930 | 182 -- 195 | 47 -- 60 | -12.7167 | 4.18664 | 5.79130 |
| CD630_21620 | CD630_n00930 | 148 -- 159 | 336 -- 347 | -12.711 | 3.02649 | 5.79130 |
| CD630_21030 | CD630_n00930 | 165 -- 193 | 4 -- 29 | -12.7 | 3.08733 | 5.79130 |
| CD630_14250 | CD630_n00930 | 163 -- 199 | 6 -- 43 | -12.6403 | 4.19604 | 5.79130 |
| CD630_06690 | CD630_n00930 | 88 -- 106 | 187 -- 207 | -12.6179 | 5.82709 | 5.79130 |
| CD630_10800 | CD630_n00930 | 153 -- 164 | 22 -- 33 | -12.404 | 5.30319 | 5.79130 |
| argG | CD630_n00930 | 71 -- 89 | 183 -- 202 | -12.3887 | 4.80300 | 5.79130 |
| ssuC | CD630_n00930 | 131 -- 159 | 1 -- 31 | -12.3142 | 3.45491 | 5.79130 |
| etfA | CD630_n00930 | 139 -- 183 | 155 -- 204 | -12.3082 | 3.41204 | 5.79130 |
| CD630_14240 | CD630_n00930 | 103 -- 145 | 4 -- 46 | -12.2989 | 8.60766 | 5.79130 |
| CD630_21080 | CD630_n00930 | 45 -- 117 | 5 -- 68 | -12.2059 | 3.99742 | 5.79130 |
| CD630_19401 | CD630_n00930 | 13 -- 67 | 2 -- 70 | -12.1893 | 4.37084 | 5.79130 |
| iorA | CD630_n00930 | 28 -- 124 | 154 -- 235 | -12.0794 | 3.83173 | 5.79130 |
| CD630_23090 | CD630_n00930 | 98 -- 145 | 4 -- 51 | -11.9601 | 4.25622 | 5.79130 |
| opuCA | CD630_n00930 | 103 -- 114 | 336 -- 347 | -11.9261 | 4.07035 | 5.79130 |
| CD630_20290 | CD630_n00930 | 51 -- 125 | 2 -- 66 | -11.8883 | 4.38183 | 5.79130 |
| CD630_28550 | CD630_n00930 | 92 -- 109 | 182 -- 207 | -11.8687 | 3.51196 | 5.79130 |
| CD630_08130 | CD630_n00930 | 170 -- 182 | 35 -- 47 | -11.8248 | 4.63831 | 5.79130 |
| CD630_31210 | CD630_n00930 | 2 -- 26 | 2 -- 29 | -11.7948 | 4.17693 | 5.79130 |
| CD630_09100 | CD630_n00930 | 36 -- 84 | 112 -- 155 | -11.7504 | 5.92260 | 5.79130 |
| CD630_34520 | CD630_n00930 | 138 -- 167 | 185 -- 215 | -11.4641 | 3.83016 | 5.79130 |
| CD630_31960 | CD630_n00930 | 34 -- 67 | 102 -- 135 | -11.4525 | 4.93757 | 5.79130 |
| CD630_13560 | CD630_n00930 | 186 -- 197 | 22 -- 32 | -11.4523 | 3.38083 | 5.79130 |
| CD630_25430 | CD630_n00930 | 94 -- 156 | 131 -- 192 | -11.4444 | 4.00527 | 5.79130 |
| gcvTPA | CD630_n00930 | 155 -- 179 | 183 -- 216 | -11.4441 | 4.77229 | 5.79130 |
| CD630_16530 | CD630_n00930 | 96 -- 148 | 7 -- 68 | -11.4355 | 4.37658 | 5.79130 |
| CD630_14130 | CD630_n00930 | 164 -- 175 | 29 -- 41 | -11.3704 | 4.57970 | 5.79130 |
| CD630_20171 | CD630_n00930 | 168 -- 180 | 202 -- 214 | -11.3696 | 6.87491 | 5.79130 |
| CD630_19903 | CD630_n00930 | 107 -- 145 | 4 -- 41 | -11.3572 | 5.79994 | 5.79130 |
| CD630_15740 | CD630_n00930 | 43 -- 95 | 29 -- 73 | -11.275 | 4.43222 | 5.79130 |
| CD630_31382 | CD630_n00930 | 171 -- 189 | 152 -- 181 | -11.247 | 3.49137 | 5.79130 |
| CD630_17281 | CD630_n00930 | 22 -- 42 | 21 -- 38 | -11.2438 | 5.91544 | 5.79130 |
| CD630_23440 | CD630_n00930 | 156 -- 190 | 24 -- 53 | -11.205 | 3.34841 | 5.79130 |
| CD630_04610 | CD630_n00930 | 158 -- 199 | 104 -- 158 | -11.2035 | 3.83025 | 5.79130 |
| CD630_25310 | CD630_n00930 | 25 -- 75 | 3 -- 63 | -11.1802 | 3.99861 | 5.79130 |
| CD630_13020 | CD630_n00930 | 117 -- 168 | 17 -- 70 | -11.1426 | 3.52267 | 5.79130 |
| CD630_31910 | CD630_n00930 | 116 -- 139 | 18 -- 44 | -11.1401 | 3.48314 | 5.79130 |
| eutW | CD630_n00930 | 182 -- 200 | 206 -- 224 | -11.0623 | 4.55715 | 5.79130 |
| CD630_32700 | CD630_n00930 | 107 -- 153 | 1 -- 47 | -11.0549 | 4.74276 | 5.79130 |
| CD630_22510 | CD630_n00930 | 121 -- 153 | 5 -- 38 | -11.0284 | 11.42763 | 5.79130 |
| CD630_08510 | CD630_n00930 | 35 -- 49 | 202 -- 216 | -11.0182 | 5.16329 | 5.79130 |
| opuCC | CD630_n00930 | 88 -- 108 | 330 -- 349 | -10.9567 | 4.07035 | 5.79130 |
| CD630_33682 | CD630_n00930 | 103 -- 145 | 4 -- 46 | -10.9435 | 5.77720 | 5.79130 |
| CD630_23280 | CD630_n00930 | 62 -- 129 | 20 -- 66 | -10.925 | 3.50093 | 5.79130 |
| CD630_06060 | CD630_n00930 | 48 -- 87 | 194 -- 245 | -10.9074 | 4.94050 | 5.79130 |
| CD630_21140 | CD630_n00930 | 4 -- 57 | 304 -- 348 | -10.9056 | 3.33304 | 5.79130 |
| CD630_15900 | CD630_n00930 | 163 -- 186 | 10 -- 35 | -10.8874 | 3.17664 | 5.79130 |
| cspD | CD630_n00930 | 162 -- 177 | 335 -- 349 | -10.8565 | 3.23587 | 5.79130 |
| CD630_32520 | CD630_n00930 | 135 -- 170 | 311 -- 346 | -10.7465 | 4.06098 | 5.79130 |
| CD630_15360 | CD630_n00930 | 81 -- 97 | 335 -- 351 | -10.744 | 4.47208 | 5.79130 |
| pgm | CD630_n00930 | 85 -- 106 | 28 -- 49 | -10.7358 | 3.45439 | 5.79130 |
| CD630_21760 | CD630_n00930 | 91 -- 135 | 172 -- 216 | -10.7026 | 4.43846 | 5.79130 |
| CD630_06170 | CD630_n00930 | 114 -- 149 | 208 -- 251 | -10.7022 | 4.95366 | 5.79130 |
| CD630_25150 | CD630_n00930 | 6 -- 45 | 173 -- 221 | -10.6062 | 3.79339 | 5.79130 |
| CD630_08090 | CD630_n00930 | 55 -- 95 | 177 -- 217 | -10.5995 | 3.57167 | 5.79130 |
| CD630_29960 | CD630_n00930 | 16 -- 60 | 294 -- 338 | -10.5495 | 4.51904 | 5.79130 |
| CD630_31010 | CD630_n00930 | 170 -- 196 | 150 -- 175 | -10.5345 | 4.96305 | 5.79130 |
| CD630_05211 | CD630_n00930 | 155 -- 189 | 5 -- 41 | -10.5248 | 3.97048 | 5.79130 |
| CD630_23820 | CD630_n00930 | 32 -- 44 | 30 -- 43 | -10.5232 | 3.18419 | 5.79130 |
| CD630_26050 | CD630_n00930 | 116 -- 153 | 2 -- 31 | -10.5086 | 6.99851 | 5.79130 |
| CD630_32040 | CD630_n00930 | 17 -- 43 | 13 -- 36 | -10.5018 | 4.93516 | 5.79130 |
| CD630_08460 | CD630_n00930 | 98 -- 162 | 156 -- 208 | -10.434 | 4.15083 | 5.79130 |
| CD630_09760 | CD630_n00930 | 76 -- 112 | 21 -- 54 | -10.4319 | 4.83410 | 5.79130 |
| CD630_28910 | CD630_n00930 | 76 -- 112 | 21 -- 54 | -10.4319 | 4.51664 | 5.79130 |
| CD630_23000 | CD630_n00930 | 89 -- 115 | 22 -- 46 | -10.3925 | 3.54130 | 5.79130 |
| CD630_21770 | CD630_n00930 | 106 -- 142 | 24 -- 60 | -10.384 | 4.02587 | 5.79130 |
| uxaA | CD630_n00930 | 172 -- 196 | 104 -- 128 | -10.3763 | 4.26165 | 5.79130 |
| CD630_14920 | CD630_n00930 | 160 -- 190 | 164 -- 186 | -10.3756 | 4.35225 | 5.79130 |
| rbsC | CD630_n00930 | 157 -- 185 | 203 -- 231 | -10.3505 | 3.71650 | 5.79130 |

| | | | | | | |
|---|---|---|---|---|---|---|
| CD630_15080 | CD630_n00930 | 10 -- 49 | 6 -- 44 | -10.3404 | 4.03033 | 5.79130 |
| CD630_06970 | CD630_n00930 | 119 -- 139 | 29 -- 51 | -10.3392 | 3.82312 | 5.79130 |
| CD630_32060 | CD630_n00930 | 37 -- 48 | 337 -- 348 | -10.3259 | 5.73797 | 5.79130 |
| CD630_36250 | CD630_n00930 | 182 -- 199 | 183 -- 216 | -10.3225 | 4.13224 | 5.79130 |
| CD630_13600 | CD630_n00930 | 126 -- 143 | 28 -- 50 | -10.3195 | 4.45481 | 5.79130 |
| CD630_22050 | CD630_n00930 | 145 -- 166 | 1 -- 25 | -10.2967 | 4.87639 | 5.79130 |
| CD630_14410 | CD630_n00930 | 49 -- 61 | 331 -- 344 | -10.2961 | 3.39433 | 5.79130 |
| CD630_03180 | CD630_n00930 | 126 -- 156 | 78 -- 110 | -10.254 | 3.73138 | 5.79130 |
| CD630_19850 | CD630_n00930 | 111 -- 127 | 22 -- 43 | -10.248 | 6.88486 | 5.79130 |
| nusB | CD630_n00930 | 70 -- 119 | 183 -- 226 | -10.2471 | 4.92798 | 5.79130 |
| CD630_21530 | CD630_n00930 | 54 -- 80 | 19 -- 47 | -10.2465 | 3.39774 | 5.79130 |
| CD630_34030 | CD630_n00930 | 18 -- 54 | 3 -- 39 | -10.2391 | 3.05766 | 5.79130 |
| CD630_09570 | CD630_n00930 | 2 -- 40 | 304 -- 340 | -10.2354 | 5.31213 | 5.79130 |
| CD630_29070 | CD630_n00930 | 2 -- 40 | 304 -- 340 | -10.2354 | 3.89477 | 5.79130 |
| CD630_17160 | CD630_n00930 | 92 -- 124 | 5 -- 42 | -10.2342 | 3.50228 | 5.79130 |
| CD630_04410 | CD630_n00930 | 58 -- 91 | 183 -- 215 | -10.2188 | 3.90451 | 5.79130 |
| CD630_01570 | CD630_n00930 | 3 -- 28 | 111 -- 136 | -10.2107 | 6.08699 | 5.79130 |
| malY | CD630_n00930 | 4 -- 15 | 21 -- 33 | -10.1931 | 4.46345 | 5.79130 |
| CD630_16950 | CD630_n00930 | 169 -- 199 | 201 -- 235 | -10.1892 | 4.42569 | 5.79130 |
| CD630_32070 | CD630_n00930 | 52 -- 81 | 29 -- 60 | -10.1848 | 3.51964 | 5.79130 |
| CD630_11220 | CD630_n00930 | 90 -- 107 | 27 -- 42 | -10.1822 | 3.52327 | 5.79130 |
| CD630_05840 | CD630_n00930 | 91 -- 134 | 3 -- 42 | -10.1674 | 4.28779 | 5.79130 |
| rolA | CD630_n00930 | 36 -- 50 | 337 -- 350 | -10.1606 | 4.56362 | 5.79130 |
| CD630_10220 | CD630_n00930 | 64 -- 129 | 150 -- 207 | -10.1396 | 4.08233 | 5.79130 |
| CD630_27630 | CD630_n00930 | 11 -- 25 | 107 -- 122 | -10.1218 | 3.54199 | 5.79130 |
| CD630_18930 | CD630_n00930 | 80 -- 116 | 105 -- 136 | -10.1079 | 5.32127 | 5.79130 |
| dltA | CD630_n00930 | 161 -- 176 | 332 -- 347 | -10.1054 | 3.85429 | 5.79130 |
| CD630_08390 | CD630_n00930 | 84 -- 100 | 183 -- 207 | -10.0977 | 5.16000 | 5.79130 |
| CD630_36170 | CD630_n00930 | 29 -- 38 | 154 -- 163 | -10.0573 | 3.72380 | 5.79130 |
| CD630_24220 | CD630_n00930 | 128 -- 143 | 23 -- 41 | -10.0543 | 4.18037 | 5.79130 |
| CD630_13400 | CD630_n00930 | 114 -- 163 | 180 -- 228 | -10.0525 | 3.20232 | 5.79130 |
| CD630_08940 | CD630_n00930 | 161 -- 185 | 52 -- 74 | -10.0512 | 3.72225 | 5.79130 |
| CD630_04280 | CD630_n00930 | 97 -- 108 | 202 -- 214 | -10.0099 | 3.61457 | 5.79130 |
| CD630_15150 | CD630_n00930 | 89 -- 98 | 335 -- 344 | -9.99893 | 3.35564 | 5.79130 |
| CD630_23780 | CD630_n00930 | 88 -- 108 | 183 -- 208 | -9.9983 | 3.91629 | 5.79130 |
| CD630_05450 | CD630_n00930 | 1 -- 30 | 292 -- 318 | -9.9442 | 5.04761 | 5.79130 |
| dltD | CD630_n00930 | 151 -- 167 | 22 -- 38 | -9.94317 | 4.40777 | 5.79130 |
| spoVFB | CD630_n00930 | 112 -- 137 | 35 -- 60 | -9.93793 | 4.62758 | 5.79130 |
| CD630_05880 | CD630_n00930 | 91 -- 108 | 334 -- 349 | -9.92563 | 4.28339 | 5.79130 |
| CD630_21610 | CD630_n00930 | 162 -- 173 | 325 -- 336 | -9.92347 | 3.16662 | 5.79130 |
| CD630_07470 | CD630_n00930 | 130 -- 159 | 5 -- 28 | -9.91508 | 3.37143 | 5.79130 |
| CD630_19690 | CD630_n00930 | 120 -- 136 | 195 -- 208 | -9.89321 | 3.85861 | 5.79130 |
| CD630_03420 | CD630_n00930 | 5 -- 20 | 116 -- 131 | -9.89052 | 4.71868 | 5.79130 |
| CD630_08660 | CD630_n00930 | 142 -- 183 | 1 -- 39 | -9.8781 | 3.86467 | 5.79130 |
| CD630_16450 | CD630_n00930 | 146 -- 183 | 29 -- 63 | -9.84443 | 4.57547 | 5.79130 |
| vncR | CD630_n00930 | 114 -- 144 | 183 -- 212 | -9.83242 | 3.93642 | 5.79130 |
| cdu1 | CD630_n00930 | 142 -- 196 | 5 -- 60 | -9.83027 | 4.46276 | 5.79130 |
| CD630_19840 | CD630_n00930 | 50 -- 62 | 52 -- 64 | -9.82298 | 3.44724 | 5.79130 |
| CD630_31000 | CD630_n00930 | 55 -- 95 | 296 -- 350 | -9.82232 | 3.70946 | 5.79130 |
| CD630_19700 | CD630_n00930 | 170 -- 191 | 3 -- 30 | -9.80369 | 3.85861 | 5.79130 |
| CD630_19904 | CD630_n00930 | 142 -- 169 | 24 -- 63 | -9.79057 | 5.96526 | 5.79130 |
| sspA | CD630_n00930 | 44 -- 63 | 114 -- 129 | -9.77778 | 3.62689 | 5.79130 |
| ldh | CD630_n00930 | 144 -- 168 | 41 -- 65 | -9.77705 | 4.35818 | 5.79130 |
| appF | CD630_n00930 | 19 -- 30 | 50 -- 61 | -9.77088 | 4.25570 | 5.79130 |
| panG | CD630_n00930 | 41 -- 54 | 337 -- 350 | -9.76998 | 5.54047 | 5.79130 |
| CD630_16330 | CD630_n00930 | 105 -- 156 | 104 -- 153 | -9.76528 | 4.11180 | 5.79130 |
| srlD | CD630_n00930 | 98 -- 128 | 316 -- 349 | -9.76426 | 4.19843 | 5.79130 |
| CD630_31580 | CD630_n00930 | 185 -- 196 | 331 -- 343 | -9.75951 | 3.51756 | 5.79130 |
| CD630_05820 | CD630_n00930 | 96 -- 119 | 174 -- 204 | -9.75028 | 3.75661 | 5.79130 |
| CD630_22710 | CD630_n00930 | 190 -- 198 | 336 -- 344 | -9.73668 | 4.05623 | 5.79130 |
| acoL | CD630_n00930 | 89 -- 119 | 310 -- 344 | -9.72166 | 3.32169 | 5.79130 |
| CD630_23770 | CD630_n00930 | 72 -- 130 | 1 -- 69 | -9.71201 | 3.25181 | 5.79130 |
| CD630_18100 | CD630_n00930 | 130 -- 192 | 2 -- 63 | -9.70821 | 5.71658 | 5.79130 |
| CD630_14050 | CD630_n00930 | 111 -- 179 | 106 -- 176 | -9.70419 | 3.30894 | 5.79130 |
| CD630_21700 | CD630_n00930 | 92 -- 117 | 188 -- 206 | -9.6981 | 3.96650 | 5.79130 |
| CD630_35780 | CD630_n00930 | 30 -- 63 | 331 -- 350 | -9.67702 | 3.50107 | 5.79130 |
| CD630_14930 | CD630_n00930 | 11 -- 28 | 20 -- 39 | -9.67148 | 3.31335 | 5.79130 |
| CD630_11240 | CD630_n00930 | 30 -- 45 | 308 -- 323 | -9.66764 | 3.20274 | 5.79130 |
| CD630_14360 | CD630_n00930 | 58 -- 92 | 316 -- 347 | -9.64725 | 3.02079 | 5.79130 |
| CD630_06560 | CD630_n00930 | 139 -- 166 | 45 -- 75 | -9.62739 | 4.02532 | 5.79130 |
| CD630_02720 | CD630_n00930 | 139 -- 153 | 52 -- 66 | -9.60841 | 3.38938 | 5.79130 |
| CD630_20960 | CD630_n00930 | 127 -- 142 | 20 -- 36 | -9.60213 | 5.65940 | 5.79130 |
| CD630_03400 | CD630_n00930 | 85 -- 116 | 3 -- 35 | -9.51252 | 4.45994 | 5.79130 |
| CD630_16670 | CD630_n00990 | 106 -- 157 | 396 -- 433 | -19.9028 | 3.33553 | 3.72051 |
| CD630_07760 | CD630_n00990 | 73 -- 109 | 960 -- 1004 | -19.3754 | 3.64186 | 3.72051 |
| nusB | CD630_n00990 | 123 -- 186 | 928 -- 996 | -19.0833 | 4.92798 | 3.72051 |
| CD630_21620 | CD630_n00990 | 50 -- 189 | 961 -- 1081 | -18.9529 | 3.02649 | 3.72051 |
| CD630_31000 | CD630_n00990 | 101 -- 152 | 948 -- 986 | -18.9114 | 3.70946 | 3.72051 |
| terD | CD630_n00990 | 1 -- 65 | 848 -- 911 | -18.8389 | 3.33148 | 3.72051 |
| CD630_18610 | CD630_n00990 | 14 -- 155 | 878 -- 1002 | -18.7996 | 4.15582 | 3.72051 |
| moeA | CD630_n00990 | 49 -- 123 | 948 -- 1022 | -18.7461 | 3.26118 | 3.72051 |
| CD630_24590 | CD630_n00990 | 2 -- 144 | 915 -- 1063 | -18.6448 | 3.70401 | 3.72051 |
| CD630_19942 | CD630_n00990 | 83 -- 145 | 917 -- 989 | -18.6053 | 4.24574 | 3.72051 |
| gcvTPA | CD630_n00990 | 37 -- 120 | 392 -- 464 | -18.5996 | 4.77229 | 3.72051 |
| CD630_03170 | CD630_n00990 | 119 -- 192 | 844 -- 909 | -18.5954 | 3.66946 | 3.72051 |
| CD630_32710 | CD630_n00990 | 124 -- 167 | 383 -- 441 | -18.4994 | 4.74276 | 3.72051 |
| CD630_25620 | CD630_n00990 | 89 -- 201 | 877 -- 984 | -18.4939 | 3.02685 | 3.72051 |
| CD630_15080 | CD630_n00990 | 22 -- 111 | 388 -- 476 | -18.3041 | 4.03033 | 3.72051 |
| CD630_02430 | CD630_n00990 | 67 -- 141 | 400 -- 469 | -18.1958 | 3.71048 | 3.72051 |
| CD630_22500 | CD630_n00990 | 115 -- 150 | 130 -- 163 | -18.0623 | 4.03688 | 3.72051 |
| CD630_10880 | CD630_n00990 | 148 -- 200 | 375 -- 434 | -18.0472 | 5.05048 | 3.72051 |
| CD630_19401 | CD630_n00990 | 21 -- 64 | 964 -- 1005 | -17.9061 | 4.37084 | 3.72051 |
| CD630_16450 | CD630_n00990 | 20 -- 110 | 391 -- 480 | -17.5987 | 4.57547 | 3.72051 |
| CD630_34030 | CD630_n00990 | 23 -- 78 | 974 -- 1024 | -17.53 | 3.05766 | 3.72051 |
| CD630_22070 | CD630_n00990 | 85 -- 170 | 459 -- 534 | -17.4329 | 4.09651 | 3.72051 |
| srlEb | CD630_n00990 | 65 -- 97 | 394 -- 428 | -17.4218 | 3.50140 | 3.72051 |
| CD630_13370 | CD630_n00990 | 103 -- 168 | 244 -- 299 | -17.4026 | 4.33145 | 3.72051 |
| CD630_31490 | CD630_n00990 | 59 -- 125 | 401 -- 474 | -17.3589 | 4.08593 | 3.72051 |
| appF | CD630_n00990 | 52 -- 122 | 385 -- 472 | -17.3484 | 4.25570 | 3.72051 |
| CD630_03040 | CD630_n00990 | 84 -- 127 | 978 -- 1013 | -17.0506 | 3.51403 | 3.72051 |
| CD630_28181 | CD630_n00990 | 4 -- 76 | 369 -- 439 | -16.9484 | 3.91299 | 3.72051 |
| CD630_17170 | CD630_n00990 | 115 -- 150 | 961 -- 996 | -16.9198 | 4.10230 | 3.72051 |
| garK | CD630_n00990 | 174 -- 191 | 390 -- 408 | -16.8243 | 3.72836 | 3.72051 |
| CD630_08080 | CD630_n00990 | 2 -- 65 | 716 -- 780 | -16.785 | 4.87819 | 3.72051 |
| CD630_11241 | CD630_n00990 | 158 -- 195 | 412 -- 444 | -16.7205 | 4.85723 | 3.72051 |
| CD630_14110 | CD630_n00990 | 5 -- 41 | 246 -- 286 | -16.6892 | 3.39550 | 3.72051 |
| CD630_20171 | CD630_n00990 | 71 -- 154 | 405 -- 499 | -16.6464 | 6.87491 | 3.72051 |
| CD630_04850 | CD630_n00990 | 165 -- 198 | 731 -- 765 | -16.5754 | 4.40788 | 3.72051 |
| CD630_06540 | CD630_n00990 | 127 -- 198 | 115 -- 183 | -16.4936 | 3.54673 | 3.72051 |
| CD630_20450 | CD630_n00990 | 110 -- 165 | 922 -- 986 | -16.4224 | 3.77388 | 3.72051 |
| opuCC | CD630_n00990 | 102 -- 198 | 664 -- 788 | -16.3896 | 4.07035 | 3.72051 |
| spaK | CD630_n00990 | 72 -- 110 | 960 -- 1004 | -16.3061 | 3.04530 | 3.72051 |
| CD630_25610 | CD630_n00990 | 111 -- 153 | 964 -- 1006 | -16.2571 | 3.02685 | 3.72051 |
| CD630_08940 | CD630_n00990 | 45 -- 123 | 960 -- 1022 | -16.2351 | 3.72225 | 3.72051 |
| CD630_19700 | CD630_n00990 | 48 -- 95 | 524 -- 566 | -16.2311 | 3.85861 | 3.72051 |
| CD630_27280 | CD630_n00990 | 136 -- 197 | 755 -- 812 | -16.1861 | 3.24738 | 3.72051 |
| CD630_21260 | CD630_n00990 | 121 -- 143 | 747 -- 771 | -16.1852 | 4.22255 | 3.72051 |
| CD630_34520 | CD630_n00990 | 88 -- 161 | 924 -- 982 | -16.096 | 3.83016 | 3.72051 |
| ung | CD630_n00990 | 100 -- 141 | 394 -- 445 | -16.0365 | 3.30908 | 3.72051 |
| CD630_19690 | CD630_n00990 | 29 -- 130 | 386 -- 482 | -16.0333 | 3.85861 | 3.72051 |
| CD630_15900 | CD630_n00990 | 140 -- 190 | 717 -- 768 | -15.9338 | 3.17664 | 3.72051 |
| CD630_06060 | CD630_n00990 | 129 -- 144 | 827 -- 842 | -15.9265 | 4.94050 | 3.72051 |
| glgB | CD630_n00990 | 100 -- 172 | 933 -- 1007 | -15.8817 | 4.48273 | 3.72051 |
| CD630_19790 | CD630_n00990 | 21 -- 157 | 460 -- 596 | -15.8416 | 3.79593 | 3.72051 |
| CD630_29500 | CD630_n00990 | 47 -- 139 | 384 -- 483 | -15.7942 | 3.70644 | 3.72051 |
| CD630_21570 | CD630_n00990 | 39 -- 108 | 394 -- 454 | -15.6988 | 6.48385 | 3.72051 |
| CD630_04410 | CD630_n00990 | 142 -- 201 | 917 -- 982 | -15.6522 | 3.90451 | 3.72051 |
| CD630_36690 | CD630_n00990 | 140 -- 197 | 265 -- 329 | -15.6462 | 3.53761 | 3.72051 |
| CD630_29650 | CD630_n00990 | 34 -- 129 | 382 -- 482 | -15.6092 | 3.67128 | 3.72051 |
| oppB | CD630_n00990 | 46 -- 102 | 275 -- 331 | -15.6075 | 2.98328 | 3.72051 |
| etfA | CD630_n00990 | 146 -- 183 | 523 -- 563 | -15.5282 | 3.41204 | 3.72051 |
| cbiO | CD630_n00990 | 91 -- 107 | 964 -- 986 | -15.4871 | 3.79830 | 3.72051 |
| oppC | CD630_n00990 | 106 -- 156 | 961 -- 1018 | -15.4297 | 3.01695 | 3.72051 |
| CD630_23000 | CD630_n00990 | 1 -- 53 | 540 -- 592 | -15.4268 | 3.54130 | 3.72051 |
| CD630_18781 | CD630_n00990 | 115 -- 158 | 18 -- 70 | -15.4215 | 3.77660 | 3.72051 |
| CD630_26630 | CD630_n00990 | 58 -- 193 | 154 -- 273 | -15.4139 | 4.61557 | 3.72051 |
| CD630_19921 | CD630_n00990 | 145 -- 198 | 948 -- 1008 | -15.3643 | 5.46345 | 3.72051 |
| CD630_10970 | CD630_n00990 | 3 -- 146 | 828 -- 969 | -15.3411 | 3.53850 | 3.72051 |
| rolB | CD630_n00990 | 120 -- 157 | 38 -- 79 | -15.3303 | 3.60270 | 3.72051 |
| CD630_14920 | CD630_n00990 | 87 -- 107 | 980 -- 1002 | -15.2906 | 4.35225 | 3.72051 |
| CD630_30730 | CD630_n00990 | 8 -- 36 | 254 -- 279 | -15.2852 | 3.13236 | 3.72051 |
| CD630_21090 | CD630_n00990 | 111 -- 183 | 846 -- 903 | -15.2555 | 4.15803 | 3.72051 |
| CD630_32690 | CD630_n00990 | 122 -- 159 | 963 -- 1004 | -15.2403 | 7.32676 | 3.72051 |
| CD630_05820 | CD630_n00990 | 65 -- 93 | 401 -- 436 | -15.175 | 3.75661 | 3.72051 |
| CD630_13960 | CD630_n00990 | 130 -- 156 | 530 -- 554 | -15.1683 | 3.30375 | 3.72051 |
| CD630_18970 | CD630_n00990 | 86 -- 158 | 948 -- 1011 | -15.1481 | 3.57243 | 3.72051 |
| CD630_08140 | CD630_n00990 | 6 -- 55 | 860 -- 901 | -15.1188 | 3.30159 | 3.72051 |
| CD630_18100 | CD630_n00990 | 11 -- 148 | 913 -- 1057 | -15.0943 | 5.71658 | 3.72051 |
| CD630_18930 | CD630_n00990 | 13 -- 53 | 763 -- 802 | -15.0032 | 5.32127 | 3.72051 |
| CD630_06340 | CD630_n00990 | 68 -- 109 | 963 -- 1006 | -14.9575 | 4.35983 | 3.72051 |
| CD630_17511 | CD630_n00990 | 3 -- 60 | 248 -- 304 | -14.9488 | 9.12322 | 3.72051 |
| CD630_19850 | CD630_n00990 | 90 -- 115 | 530 -- 554 | -14.918 | 6.88486 | 3.72051 |
| CD630_28280 | CD630_n00990 | 21 -- 116 | 707 -- 810 | -14.9116 | 4.59833 | 3.72051 |
| CD630_28550 | CD630_n00990 | 68 -- 106 | 965 -- 1009 | -14.8849 | 3.51196 | 3.72051 |
| mobA | CD630_n00990 | 141 -- 193 | 55 -- 99 | -14.8679 | 4.25703 | 3.72051 |
| CD630_09760 | CD630_n00990 | 111 -- 144 | 964 -- 1004 | -14.7982 | 4.83410 | 3.72051 |
| CD630_28910 | CD630_n00990 | 111 -- 144 | 964 -- 1004 | -14.7982 | 4.51664 | 3.72051 |
| CD630_32990 | CD630_n00990 | 22 -- 54 | 783 -- 817 | -14.7589 | 5.24925 | 3.72051 |
| cobQ | CD630_n00990 | 88 -- 110 | 977 -- 997 | -14.7101 | 3.13524 | 3.72051 |
| CD630_08100 | CD630_n00990 | 87 -- 135 | 923 -- 984 | -14.7014 | 4.39181 | 3.72051 |
| bglA | CD630_n00990 | 159 -- 185 | 56 -- 77 | -14.6719 | 4.86402 | 3.72051 |
| dltB | CD630_n00990 | 1 -- 34 | 161 -- 199 | -14.637 | 3.81485 | 3.72051 |
| CD630_04610 | CD630_n00990 | 143 -- 201 | 752 -- 815 | -14.5914 | 3.83025 | 3.72051 |
| CD630_19030 | CD630_n00990 | 109 -- 198 | 726 -- 809 | -14.5726 | 3.34240 | 3.72051 |
| CD630_05950 | CD630_n00990 | 29 -- 63 | 394 -- 433 | -14.5698 | 3.31669 | 3.72051 |
| chrA | CD630_n00990 | 1 -- 52 | 847 -- 901 | -14.5431 | 5.27255 | 3.72051 |
| CD630_27570 | CD630_n00990 | 84 -- 194 | 896 -- 1015 | -14.5067 | 3.46216 | 3.72051 |
| CD630_06160 | CD630_n00990 | 164 -- 199 | 254 -- 290 | -14.487 | 5.51744 | 3.72051 |
| CD630_21760 | CD630_n00990 | 173 -- 199 | 246 -- 274 | -14.4617 | 4.43846 | 3.72051 |
| CD630_01590 | CD630_n00990 | 2 -- 28 | 744 -- 766 | -14.4573 | 3.45609 | 3.72051 |
| CD630_27110 | CD630_n00990 | 174 -- 199 | 243 -- 270 | -14.442 | 3.78231 | 3.72051 |
| CD630_28290 | CD630_n00990 | 25 -- 74 | 849 -- 891 | -14.4328 | 5.54192 | 3.72051 |
| CD630_21730 | CD630_n00990 | 166 -- 193 | 238 -- 265 | -14.4144 | 3.45076 | 3.72051 |
| CD630_15360 | CD630_n00990 | 36 -- 106 | 392 -- 459 | -14.3487 | 4.47208 | 3.72051 |
| CD630_06080 | CD630_n00990 | 45 -- 104 | 374 -- 435 | -14.3418 | 3.82315 | 3.72051 |
| CD630_28150 | CD630_n00990 | 21 -- 53 | 398 -- 436 | -14.3192 | 4.18447 | 3.72051 |
| acoL | CD630_n00990 | 106 -- 138 | 31 -- 64 | -14.3154 | 3.32169 | 3.72051 |
| CD630_16330 | CD630_n00990 | 149 -- 200 | 247 -- 299 | -14.3084 | 4.11180 | 3.72051 |
| CD630_30310 | CD630_n00990 | 155 -- 188 | 32 -- 64 | -14.2962 | 4.56059 | 3.72051 |
| CD630_32700 | CD630_n00990 | 102 -- 119 | 526 -- 543 | -14.2673 | 4.74276 | 3.72051 |
| CD630_21240 | CD630_n00990 | 121 -- 152 | 788 -- 814 | -14.2657 | 5.88879 | 3.72051 |
| CD630_15200 | CD630_n00990 | 8 -- 138 | 683 -- 800 | -14.2517 | 4.00442 | 3.72051 |
| CD630_06970 | CD630_n00990 | 83 -- 151 | 918 -- 985 | -14.239 | 3.82312 | 3.72051 |
| kdpD | CD630_n00990 | 141 -- 199 | 960 -- 1002 | -14.2351 | 4.27140 | 3.72051 |
| moaA | CD630_n00990 | 51 -- 120 | 528 -- 576 | -14.2338 | 3.83799 | 3.72051 |
| CD630_21140 | CD630_n00990 | 128 -- 195 | 905 -- 971 | -14.1954 | 3.33304 | 3.72051 |
| fbp | CD630_n00990 | 172 -- 189 | 798 -- 818 | -14.1864 | 3.80071 | 3.72051 |
| CD630_14360 | CD630_n00990 | 40 -- 116 | 954 -- 1018 | -14.1674 | 3.02079 | 3.72051 |
| CD630_27610 | CD630_n00990 | 11 -- 29 | 462 -- 480 | -14.0759 | 3.96465 | 3.72051 |
| CD630_01700 | CD630_n00990 | 1 -- 50 | 246 -- 295 | -14.0537 | 4.97234 | 3.72051 |
| pilC | CD630_n00990 | 11 -- 110 | 391 -- 471 | -14.0361 | 3.54230 | 3.72051 |
| CD630_22550 | CD630_n00990 | 125 -- 196 | 238 -- 327 | -14.0346 | 3.62379 | 3.72051 |
| CD630_31010 | CD630_n00990 | 133 -- 158 | 945 -- 971 | -14.0279 | 4.96305 | 3.72051 |
| CD630_20960 | CD630_n00990 | 50 -- 111 | 392 -- 446 | -14.0124 | 5.65940 | 3.72051 |
| CD630_17710 | CD630_n00990 | 86 -- 109 | 961 -- 987 | -14.0018 | 4.37688 | 3.72051 |
| eutW | CD630_n00990 | 129 -- 144 | 252 -- 267 | -13.9663 | 4.55715 | 3.72051 |
| CD630_17360 | CD630_n00990 | 89 -- 123 | 403 -- 428 | -13.964 | 5.66070 | 3.72051 |
| CD630_26920 | CD630_n00990 | 33 -- 57 | 245 -- 269 | -13.9528 | 4.58060 | 3.72051 |
| CD630_13880 | CD630_n00990 | 111 -- 158 | 229 -- 270 | -13.9478 | 3.52735 | 3.72051 |
| CD630_19840 | CD630_n00990 | 133 -- 178 | 669 -- 719 | -13.9389 | 3.44724 | 3.72051 |
| CD630_05000 | CD630_n00990 | 110 -- 159 | 922 -- 967 | -13.9297 | 4.02442 | 3.72051 |
| cdu2 | CD630_n00990 | 36 -- 162 | 393 -- 505 | -13.9047 | 2.97335 | 3.72051 |
| CD630_22740 | CD630_n00990 | 62 -- 119 | 461 -- 512 | -13.9012 | 3.08733 | 3.72051 |
| CD630_09772 | CD630_n00990 | 69 -- 92 | 991 -- 1014 | -13.8969 | 3.66033 | 3.72051 |
| CD630_32520 | CD630_n00990 | 166 -- 201 | 975 -- 1008 | -13.8898 | 4.06098 | 3.72051 |
| CD630_30120 | CD630_n00990 | 70 -- 134 | 947 -- 1007 | -13.8806 | 4.04859 | 3.72051 |
| CD630_14340 | CD630_n00990 | 44 -- 103 | 33 -- 86 | -13.8679 | 5.10868 | 3.72051 |
| cobC | CD630_n00990 | 165 -- 195 | 990 -- 1018 | -13.8566 | 3.15854 | 3.72051 |
| CD630_06050 | CD630_n00990 | 89 -- 166 | 384 -- 471 | -13.8301 | 3.59332 | 3.72051 |
| CD630_05450 | CD630_n00990 | 83 -- 103 | 968 -- 988 | -13.822 | 5.04761 | 3.72051 |
| CD630_35260 | CD630_n00990 | 171 -- 188 | 525 -- 543 | -13.7314 | 5.77277 | 3.72051 |
| CD630_13950 | CD630_n00990 | 149 -- 201 | 749 -- 799 | -13.7019 | 4.25540 | 3.72051 |
| bcp | CD630_n00990 | 89 -- 151 | 330 -- 398 | -13.701 | 3.16670 | 3.72051 |
| uxaA%27 | CD630_n00990 | 88 -- 160 | 890 -- 982 | -13.6809 | 4.26165 | 3.72051 |
| CD630_14630 | CD630_n00990 | 55 -- 134 | 952 -- 1022 | -13.6768 | 3.34485 | 3.72051 |
| CD630_05460 | CD630_n00990 | 51 -- 93 | 989 -- 1029 | -13.6761 | 3.98753 | 3.72051 |
| CD630_22030 | CD630_n00990 | 156 -- 198 | 923 -- 981 | -13.6365 | 3.29934 | 3.72051 |
| CD630_14930 | CD630_n00990 | 2 -- 43 | 698 -- 730 | -13.6042 | 3.31335 | 3.72051 |
| CD630_13530 | CD630_n00990 | 5 -- 31 | 765 -- 790 | -13.6027 | 4.09832 | 3.72051 |
| CD630_16150 | CD630_n00990 | 5 -- 75 | 469 -- 541 | -13.5995 | 4.20614 | 3.72051 |
| cbpA | CD630_n00990 | 75 -- 107 | 961 -- 1002 | -13.5839 | 3.03040 | 3.72051 |
| xdhA | CD630_n00990 | 99 -- 150 | 948 -- 1015 | -13.5696 | 4.45147 | 3.72051 |
| CD630_05821 | CD630_n00990 | 176 -- 199 | 881 -- 904 | -13.5593 | 3.75661 | 3.72051 |
| CD630_14130 | CD630_n00990 | 89 -- 138 | 961 -- 1011 | -13.5105 | 4.57970 | 3.72051 |
| CD630_23820 | CD630_n00990 | 44 -- 109 | 962 -- 1015 | -13.4666 | 3.18419 | 3.72051 |
| CD630_23780 | CD630_n00990 | 5 -- 54 | 712 -- 761 | -13.4381 | 3.91629 | 3.72051 |
| CD630_32040 | CD630_n00990 | 22 -- 41 | 387 -- 407 | -13.4371 | 4.93516 | 3.72051 |
| CD630_25320 | CD630_n00990 | 142 -- 153 | 974 -- 985 | -13.4154 | 4.29415 | 3.72051 |
| appC | CD630_n00990 | 43 -- 76 | 285 -- 320 | -13.4061 | 4.50438 | 3.72051 |
| CD630_20980 | CD630_n00990 | 78 -- 152 | 401 -- 499 | -13.3979 | 8.12847 | 3.72051 |
| CD630_27230 | CD630_n00990 | 6 -- 26 | 947 -- 968 | -13.3958 | 3.52462 | 3.72051 |
| dapB2 | CD630_n00990 | 136 -- 160 | 974 -- 1000 | -13.3876 | 3.27218 | 3.72051 |
| CD630_13350 | CD630_n00990 | 158 -- 199 | 313 -- 354 | -13.3843 | 4.15139 | 3.72051 |
| CD630_27210 | CD630_n00990 | 118 -- 150 | 574 -- 607 | -13.3746 | 4.18786 | 3.72051 |
| CD630_27310 | CD630_n00990 | 7 -- 62 | 48 -- 102 | -13.3624 | 4.01622 | 3.72051 |
| CD630_10950 | CD630_n00990 | 30 -- 106 | 394 -- 469 | -13.3552 | 3.67675 | 3.72051 |
| CD630_24580 | CD630_n00990 | 11 -- 46 | 229 -- 266 | -13.3238 | 3.88519 | 3.72051 |
| scpB | CD630_n00990 | 92 -- 151 | 951 -- 1007 | -13.3052 | 3.33560 | 3.72051 |

| | | | | | | | |
|---|---|---|---|---|---|---|---|
| def1 | CD630_n00990 | 86 – 188 | 927 – 986 | -13.3035 | | 3.99720 | 3.72051 |
| CD630_31210 | CD630_n00990 | 3 – 69 | 708 – 767 | -13.2859 | | 4.17693 | 3.72051 |
| CD630_14260 | CD630_n00990 | 22 – 68 | 226 – 259 | -13.2774 | | 3.22214 | 3.72051 |
| CD630_27630 | CD630_n00990 | 1 – 73 | 693 – 764 | -13.2625 | | 3.54199 | 3.72051 |
| CD630_23080 | CD630_n00990 | 1 – 56 | 251 – 303 | -13.2518 | | 5.23027 | 3.72051 |
| argG | CD630_n00990 | 70 – 107 | 394 – 425 | -13.2492 | | 4.80300 | 3.72051 |
| cwp29 | CD630_n00990 | 89 – 137 | 937 – 984 | -13.2463 | | 4.40865 | 3.72051 |
| CD630_28320 | CD630_n00990 | 80 – 95 | 52 – 67 | -13.2259 | | 3.99184 | 3.72051 |
| CD630_06140 | CD630_n00990 | 81 – 124 | 33 – 70 | -13.1491 | | 4.12519 | 3.72051 |
| CD630_21610 | CD630_n00990 | 156 – 171 | 773 – 788 | -13.137 | | 3.16662 | 3.72051 |
| CD630_06380 | CD630_n00990 | 147 – 201 | 732 – 785 | -13.1264 | | 4.32211 | 3.72051 |
| CD630_27270 | CD630_n00990 | 158 – 174 | 966 – 986 | -13.1192 | | 3.50892 | 3.72051 |
| CD630_25300 | CD630_n00990 | 73 – 154 | 224 – 300 | -13.1104 | | 4.49333 | 3.72051 |
| CD630_16460 | CD630_n00990 | 91 – 160 | 916 – 982 | -13.0767 | | 4.64028 | 3.72051 |
| CD630_23930 | CD630_n00990 | 71 – 200 | 878 – 1022 | -13.0615 | | 3.05650 | 3.72051 |
| CD630_13780 | CD630_n00990 | 85 – 130 | 385 – 434 | -13.0479 | | 3.52274 | 3.72051 |
| CD630_07840 | CD630_n00990 | 114 – 154 | 521 – 560 | -13.0162 | | 4.26761 | 3.72051 |
| dnaD | CD630_n00990 | 114 – 172 | 252 – 300 | -13.0132 | | 3.56078 | 3.72051 |
| CD630_34530 | CD630_n00990 | 65 – 121 | 963 – 1020 | -12.9866 | | 3.83016 | 3.72051 |
| CD630_21770 | CD630_n00990 | 52 – 112 | 959 – 1022 | -12.8947 | | 4.02587 | 3.72051 |
| CD630_30130 | CD630_n00990 | 31 – 76 | 963 – 1011 | -12.8796 | | 3.28343 | 3.72051 |
| CD630_15850 | CD630_n00990 | 121 – 197 | 942 – 1001 | -12.8389 | | 4.32968 | 3.72051 |
| CD630_24760 | CD630_n00990 | 55 – 161 | 388 – 496 | -12.8313 | | 5.19485 | 3.72051 |
| CD630_10830 | CD630_n00990 | 66 – 104 | 967 – 1005 | -12.8219 | | 4.82398 | 3.72051 |
| CD630_13600 | CD630_n00990 | 35 – 165 | 394 – 522 | -12.8055 | | 4.45481 | 3.72051 |
| CD630_21700 | CD630_n00990 | 121 – 139 | 244 – 263 | -12.8023 | | 3.96650 | 3.72051 |
| CD630_19430 | CD630_n00990 | 88 – 116 | 955 – 984 | -12.7372 | | 3.13922 | 3.72051 |
| malY | CD630_n00990 | 4 – 18 | 460 – 473 | -12.7149 | | 4.46345 | 3.72051 |
| CD630_16750 | CD630_n00990 | 4 – 36 | 720 – 755 | -12.7133 | | 5.04051 | 3.72051 |
| CD630_18650 | CD630_n00990 | 29 – 98 | 401 – 478 | -12.7091 | | 3.75892 | 3.72051 |
| CD630_31580 | CD630_n00990 | 85 – 114 | 962 – 986 | -12.702 | | 3.51756 | 3.72051 |
| CD630_30140 | CD630_n00990 | 15 – 62 | 927 – 971 | -12.6995 | | 3.28343 | 3.72051 |
| srlD | CD630_n00990 | 88 – 104 | 967 – 984 | -12.6558 | | 4.19843 | 3.72051 |
| CD630_05230 | CD630_n00990 | 127 – 169 | 284 – 333 | -12.6443 | | 3.73787 | 3.72051 |
| eutV | CD630_n00990 | 102 – 135 | 960 – 985 | -12.6313 | | 4.55715 | 3.72051 |
| CD630_06060 | CD630_n00990 | 172 – 193 | 956 – 985 | -12.5974 | | 4.94050 | 3.72051 |
| ssuC | CD630_n00990 | 47 – 83 | 974 – 1007 | -12.5856 | | 3.45491 | 3.72051 |
| CD630_06100 | CD630_n00990 | 137 – 201 | 749 – 790 | -12.5806 | | 4.43918 | 3.72051 |
| CD630_09750 | CD630_n00990 | 89 – 128 | 976 – 1010 | -12.5647 | | 4.83410 | 3.72051 |
| CD630_28920 | CD630_n00990 | 89 – 128 | 976 – 1010 | -12.5647 | | 4.51664 | 3.72051 |
| CD630_33600 | CD630_n00990 | 21 – 31 | 31 – 41 | -12.5436 | | 3.73196 | 3.72051 |
| CD630_25050 | CD630_n00990 | 1 – 67 | 704 – 765 | -12.5075 | | 4.33624 | 3.72051 |
| CD630_32050 | CD630_n00990 | 132 – 177 | 816 – 854 | -12.4965 | | 5.73797 | 3.72051 |
| CD630_13020 | CD630_n00990 | 117 – 168 | 319 – 349 | -12.4818 | | 3.52267 | 3.72051 |
| CD630_25310 | CD630_n00990 | 22 – 31 | 530 – 539 | -12.4706 | | 3.99861 | 3.72051 |
| CD630_25020 | CD630_n00990 | 12 – 53 | 733 – 775 | -12.4192 | | 4.17721 | 3.72051 |
| CD630_04550 | CD630_n00990 | 94 – 136 | 267 – 305 | -12.4017 | | 3.05227 | 3.72051 |
| CD630_18090 | CD630_n00990 | 81 – 168 | 426 – 539 | -12.3592 | | 4.23961 | 3.72051 |
| CD630_19000 | CD630_n00990 | 54 – 136 | 393 – 462 | -12.3513 | | 4.18443 | 3.72051 |
| CD630_05830 | CD630_n00990 | 134 – 183 | 757 – 811 | -12.3496 | | 3.35360 | 3.72051 |
| CD630_04940 | CD630_n00990 | 89 – 100 | 973 – 984 | -12.3351 | | 4.20948 | 3.72051 |
| CD630_05490 | CD630_n00990 | 137 – 178 | 384 – 421 | -12.322 | | 3.89852 | 3.72051 |
| CD630_06170 | CD630_n00990 | 102 – 168 | 638 – 718 | -12.3126 | | 4.95366 | 3.72051 |
| CD630_04280 | CD630_n00990 | 156 – 169 | 405 – 417 | -12.3011 | | 3.61457 | 3.72051 |
| CD630_19680 | CD630_n00990 | 39 – 97 | 424 – 484 | -12.3011 | | 3.58080 | 3.72051 |
| argS | CD630_n00990 | 176 – 195 | 346 – 363 | -12.2762 | | 4.96036 | 3.72051 |
| CD630_05822 | CD630_n00990 | 172 – 201 | 891 – 923 | -12.2754 | | 4.28113 | 3.72051 |
| pgm | CD630_n00990 | 16 – 43 | 382 – 405 | -12.264 | | 3.45439 | 3.72051 |
| pilT | CD630_n00990 | 102 – 180 | 688 – 766 | -12.2523 | | 3.24639 | 3.72051 |
| CD630_07870 | CD630_n00990 | 159 – 200 | 847 – 898 | -12.2502 | | 3.22829 | 3.72051 |
| CD630_23770 | CD630_n00990 | 59 – 201 | 370 – 511 | -12.2349 | | 3.25181 | 3.72051 |
| CD630_31500 | CD630_n00990 | 135 – 168 | 894 – 932 | -12.2327 | | 4.90207 | 3.72051 |
| CD630_06110 | CD630_n00990 | 170 – 186 | 981 – 997 | -12.2267 | | 5.61380 | 3.72051 |
| CD630_33420 | CD630_n00990 | 142 – 159 | 795 – 812 | -12.1676 | | 3.22095 | 3.72051 |
| panG | CD630_n00990 | 161 – 199 | 832 – 865 | -12.1658 | | 5.54047 | 3.72051 |
| CD630_32070 | CD630_n00990 | 158 – 179 | 799 – 818 | -12.1466 | | 3.51964 | 3.72051 |
| CD630_15550 | CD630_n00990 | 85 – 139 | 949 – 1000 | -12.1381 | | 3.49019 | 3.72051 |
| CD630_05880 | CD630_n00990 | 104 – 174 | 961 – 1044 | -12.138 | | 4.28339 | 3.72051 |
| CD630_20170 | CD630_n00990 | 99 – 119 | 401 – 421 | -12.1366 | | 6.90201 | 3.72051 |
| CD630_22050 | CD630_n00990 | 134 – 191 | 848 – 909 | -12.1354 | | 4.87639 | 3.72051 |
| CD630_09040 | CD630_n00990 | 23 – 53 | 924 – 949 | -12.1326 | | 4.11851 | 3.72051 |
| CD630_30420 | CD630_n00990 | 137 – 175 | 747 – 785 | -12.0895 | | 5.27539 | 3.72051 |
| CD630_19580 | CD630_n00990 | 63 – 95 | 977 – 1011 | -12.0892 | | 4.51902 | 3.72051 |
| kdgT1 | CD630_n00990 | 67 – 111 | 958 – 1020 | -12.0629 | | 4.49207 | 3.72051 |
| CD630_18010 | CD630_n00990 | 175 – 201 | 761 – 786 | -12.0301 | | 4.21967 | 3.72051 |
| CD630_05870 | CD630_n00990 | 158 – 174 | 964 – 980 | -12.0106 | | 4.28339 | 3.72051 |
| CD630_10080 | CD630_n00990 | 9 – 51 | 706 – 754 | -12.0073 | | 3.75625 | 3.72051 |
| cspD | CD630_n00990 | 86 – 168 | 32 – 112 | -11.9795 | | 3.23587 | 3.72051 |
| CD630_16520 | CD630_n00990 | 148 – 193 | 926 – 982 | -11.9742 | | 4.28395 | 3.72051 |
| CD630_13560 | CD630_n00990 | 131 – 179 | 923 – 971 | -11.9709 | | 3.38083 | 3.72051 |
| hfq | CD630_n00990 | 150 – 197 | 929 – 982 | -11.962 | | 5.05609 | 3.72051 |
| CD630_08130 | CD630_n00990 | 13 – 96 | 386 – 470 | -11.9518 | | 4.63831 | 3.72051 |
| CD630_17292 | CD630_n00990 | 93 – 149 | 962 – 1007 | -11.9503 | | 6.25799 | 3.72051 |
| CD630_23861 | CD630_n00990 | 17 – 41 | 248 – 267 | -11.9495 | | 5.83068 | 3.72051 |
| CD630_14710 | CD630_n00990 | 60 – 106 | 823 – 871 | -11.9468 | | 3.65171 | 3.72051 |
| CD630_15060 | CD630_n00990 | 170 – 196 | 255 – 275 | -11.9458 | | 3.69396 | 3.72051 |
| CD630_23440 | CD630_n00990 | 53 – 80 | 964 – 1002 | -11.9383 | | 3.34841 | 3.72051 |
| cspC | CD630_n00990 | 160 – 177 | 964 – 982 | -11.9041 | | 5.48848 | 3.72051 |
| CD630_32720 | CD630_n00990 | 113 – 151 | 519 – 565 | -11.903 | | 4.74276 | 3.72051 |
| CD630_12320 | CD630_n00990 | 50 – 105 | 966 – 1015 | -11.9018 | | 3.28667 | 3.72051 |
| CD630_10050 | CD630_n00990 | 154 – 163 | 975 – 984 | -11.8978 | | 3.93750 | 3.72051 |
| CD630_22880 | CD630_n00990 | 39 – 64 | 244 – 268 | -11.8845 | | 3.62455 | 3.72051 |
| CD630_10000 | CD630_n00990 | 82 – 113 | 816 – 844 | -11.8843 | | 3.86040 | 3.72051 |
| CD630_31220 | CD630_n00990 | 165 – 195 | 324 – 350 | -11.8649 | | 3.73779 | 3.72051 |
| CD630_14240 | CD630_n00990 | 70 – 133 | 948 – 999 | -11.8608 | | 8.60766 | 3.72051 |
| CD630_02790 | CD630_n00990 | 15 – 135 | 34 – 156 | -11.8442 | | 4.77661 | 3.72051 |
| CD630_09840 | CD630_n00990 | 105 – 130 | 248 – 272 | -11.827 | | 4.50009 | 3.72051 |
| iorA | CD630_n00990 | 2 – 22 | 815 – 835 | -11.8259 | | 3.83173 | 3.72051 |
| CD630_03160 | CD630_n00990 | 67 – 80 | 528 – 541 | -11.8188 | | 3.66946 | 3.72051 |
| uxaA | CD630_n00990 | 74 – 98 | 390 – 407 | -11.796 | | 4.26165 | 3.72051 |
| CD630_02740 | CD630_n00990 | 140 – 162 | 964 – 983 | -11.7888 | | 5.02277 | 3.72051 |
| CD630_08370 | CD630_n00990 | 112 – 155 | 395 – 436 | -11.7868 | | 3.65639 | 3.72051 |
| CD630_18960 | CD630_n00990 | 11 – 55 | 975 – 1006 | -11.775 | | 4.37822 | 3.72051 |
| CD630_18410 | CD630_n00990 | 123 – 172 | 247 – 292 | -11.7644 | | 3.59739 | 3.72051 |
| CD630_16550 | CD630_n00990 | 80 – 107 | 394 – 434 | -11.7541 | | 5.61064 | 3.72051 |
| CD630_20670 | CD630_n00990 | 151 – 159 | 979 – 987 | -11.7506 | | 4.42757 | 3.72051 |
| CD630_10960 | CD630_n00990 | 11 – 40 | 395 – 423 | -11.7336 | | 3.53850 | 3.72051 |
| CD630_28260 | CD630_n00990 | 39 – 108 | 398 – 471 | -11.7211 | | 3.51613 | 3.72051 |
| CD630_19780 | CD630_n00990 | 2 – 18 | 466 – 482 | -11.7195 | | 3.64830 | 3.72051 |
| CD630_24050 | CD630_n00990 | 5 – 52 | 707 – 763 | -11.7158 | | 3.70192 | 3.72051 |
| CD630_14810 | CD630_n00990 | 54 – 65 | 403 – 414 | -11.7024 | | 3.35323 | 3.72051 |
| CD630_31461 | CD630_n00990 | 141 – 201 | 250 – 295 | -11.5869 | | 4.41476 | 3.72051 |
| thiH | CD630_n00990 | 68 – 99 | 984 – 1016 | -11.573 | | 3.58230 | 3.72051 |
| CD630_02420 | CD630_n00990 | 3 – 35 | 393 – 428 | -11.5637 | | 3.71048 | 3.72051 |
| CD630_30540 | CD630_n00990 | 123 – 156 | 382 – 408 | -11.5598 | | 3.63804 | 3.72051 |
| CD630_21490 | CD630_n00990 | 149 – 169 | 791 – 811 | -11.551 | | 3.37462 | 3.72051 |
| cwp26 | CD630_n00990 | 70 – 88 | 967 – 985 | -11.5349 | | 4.44377 | 3.72051 |
| CD630_31690 | CD630_n00990 | 112 – 160 | 252 – 298 | -11.5343 | | 3.71193 | 3.72051 |
| CD630_27200 | CD630_n00990 | 122 – 137 | 312 – 327 | -11.5262 | | 3.42516 | 3.72051 |
| CD630_08070 | CD630_n00990 | 86 – 125 | 952 – 983 | -11.5261 | | 4.87819 | 3.72051 |
| CD630_16160 | CD630_n00990 | 66 – 111 | 959 – 1004 | -11.5237 | | 6.11801 | 3.72051 |
| htpG | CD630_n00990 | 46 – 93 | 990 – 1022 | -11.5019 | | 3.45147 | 3.72051 |
| cdu1 | CD630_n00990 | 3 – 35 | 532 – 565 | -11.5003 | | 4.46276 | 3.72051 |
| CD630_13790 | CD630_n00990 | 78 – 118 | 977 – 1004 | -11.4958 | | 5.17374 | 3.72051 |
| CD630_03400 | CD630_n00990 | 17 – 46 | 36 – 67 | -11.4951 | | 4.45994 | 3.72051 |
| CD630_28870 | CD630_n00990 | 32 – 73 | 552 – 594 | -11.4836 | | 3.41991 | 3.72051 |
| CD630_20600 | CD630_n00990 | 79 – 124 | 951 – 1004 | -11.4664 | | 4.20093 | 3.72051 |
| CD630_13580 | CD630_n00990 | 153 – 174 | 69 – 90 | -11.4638 | | 3.80715 | 3.72051 |
| CD630_08460 | CD630_n00990 | 148 – 166 | 522 – 543 | -11.4482 | | 4.15083 | 3.72051 |
| CD630_12110 | CD630_n00990 | 167 – 191 | 735 – 758 | -11.4476 | | 4.36078 | 3.72051 |
| CD630_19904 | CD630_n00990 | 154 – 200 | 316 – 366 | -11.4443 | | 5.96526 | 3.72051 |
| CD630_32060 | CD630_n00990 | 161 – 189 | 691 – 721 | -11.4404 | | 5.73797 | 3.72051 |
| CD630_03220 | CD630_n00990 | 86 – 135 | 392 – 444 | -11.4121 | | 3.36746 | 3.72051 |
| metQ | CD630_n00990 | 88 – 104 | 967 – 984 | -11.4097 | | 3.87190 | 3.72051 |
| CD630_23280 | CD630_n00990 | 71 – 140 | 961 – 1022 | -11.4051 | | 3.50093 | 3.72051 |
| CD630_01690 | CD630_n00990 | 151 – 171 | 850 – 869 | -11.3978 | | 4.97234 | 3.72051 |
| panB | CD630_n00990 | 153 – 187 | 238 – 270 | -11.3821 | | 3.40248 | 3.72051 |
| CD630_04880 | CD630_n00990 | 122 – 160 | 883 – 923 | -11.3812 | | 4.62291 | 3.72051 |
| CD630_29760 | CD630_n00990 | 134 – 150 | 991 – 1007 | -11.3698 | | 3.51074 | 3.72051 |
| CD630_04540 | CD630_n00990 | 80 – 94 | 56 – 70 | -11.3355 | | 3.33556 | 3.72051 |
| CD630_26150 | CD630_n00990 | 19 – 83 | 712 – 787 | -11.3253 | | 4.06302 | 3.72051 |
| CD630_36250 | CD630_n00990 | 53 – 152 | 917 – 1015 | -11.3107 | | 4.13224 | 3.72051 |
| CD630_05900 | CD630_n00990 | 116 – 174 | 866 – 910 | -11.3058 | | 3.21934 | 3.72051 |
| CD630_13920 | CD630_n00990 | 25 – 49 | 245 – 268 | -11.2962 | | 3.25236 | 3.72051 |
| CD630_07430 | CD630_n00990 | 3 – 22 | 256 – 275 | -11.2936 | | 4.65568 | 3.72051 |
| CD630_29840 | CD630_n00990 | 25 – 38 | 31 – 44 | -11.2922 | | 4.48825 | 3.72051 |
| dpaA | CD630_n00990 | 40 – 54 | 230 – 245 | -11.2918 | | 4.65364 | 3.72051 |
| CD630_20460 | CD630_n00990 | 77 – 100 | 990 – 1013 | -11.2905 | | 7.94034 | 3.72051 |
| CD630_13400 | CD630_n00990 | 149 – 180 | 885 – 923 | -11.2647 | | 3.20232 | 3.72051 |
| CD630_25040 | CD630_n00990 | 1 – 53 | 926 – 976 | -11.2618 | | 3.43898 | 3.72051 |
| CD630_15740 | CD630_n00990 | 71 – 97 | 392 – 419 | -11.2574 | | 4.43222 | 3.72051 |
| CD630_17281 | CD630_n00990 | 74 – 151 | 278 – 352 | -11.2397 | | 5.91544 | 3.72051 |
| CD630_29250 | CD630_n00990 | 81 – 135 | 361 – 420 | -11.2373 | | 4.61901 | 3.72051 |
| CD630_09410 | CD630_n00990 | 81 – 135 | 361 – 420 | -11.2373 | | 4.75651 | 3.72051 |
| CD630_02830 | CD630_n00990 | 155 – 174 | 824 – 844 | -11.2338 | | 3.57106 | 3.72051 |
| CD630_18311 | CD630_n00990 | 96 – 181 | 881 – 954 | -11.2285 | | 6.86047 | 3.72051 |
| CD630_32880 | CD630_n00990 | 106 – 196 | 233 – 315 | -11.2145 | | 3.41276 | 3.72051 |
| cls | CD630_n00990 | 168 – 201 | 675 – 713 | -11.1989 | | 4.46177 | 3.72051 |
| CD630_21070 | CD630_n00990 | 169 – 197 | 29 – 66 | -11.1941 | | 5.35019 | 3.72051 |
| CD630_27060 | CD630_n00990 | 102 – 161 | 926 – 985 | -11.1905 | | 4.06049 | 3.72051 |
| CD630_08110 | CD630_n00990 | 106 – 116 | 459 – 469 | -11.1853 | | 4.39181 | 3.72051 |
| CD630_04660 | CD630_n00990 | 73 – 100 | 991 – 1018 | -11.1763 | | 4.32791 | 3.72051 |
| CD630_26680 | CD630_n00990 | 70 – 98 | 397 – 428 | -11.161 | | 3.53956 | 3.72051 |
| CD630_03930 | CD630_n00990 | 138 – 166 | 739 – 766 | -11.1564 | | 4.15934 | 3.72051 |
| CD630_18920 | CD630_n00990 | 116 – 189 | 394 – 483 | -11.1294 | | 4.43558 | 3.72051 |
| CD630_34050 | CD630_n00990 | 80 – 151 | 918 – 996 | -11.097 | | 4.03863 | 3.72051 |
| CD630_27240 | CD630_n00990 | 147 – 170 | 253 – 275 | -11.0936 | | 3.52462 | 3.72051 |
| CD630_14610 | CD630_n00990 | 152 – 183 | 684 – 716 | -11.0541 | | 4.89428 | 3.72051 |
| CD630_25150 | CD630_n00990 | 99 – 157 | 687 – 763 | -11.0372 | | 3.79339 | 3.72051 |
| CD630_26050 | CD630_n00990 | 68 – 135 | 941 – 1007 | -11.0341 | | 6.99851 | 3.72051 |
| CD630_06150 | CD630_n00990 | 144 – 177 | 244 – 275 | -11.0258 | | 5.51744 | 3.72051 |
| CD630_18440 | CD630_n00990 | 105 – 118 | 82 – 95 | -11.0182 | | 4.07960 | 3.72051 |
| CD630_26050 | CD630_n00990 | 171 – 188 | 967 – 986 | -10.998 | | 6.99851 | 3.72051 |
| CD630_33580 | CD630_n00990 | 174 – 201 | 711 – 748 | -10.9928 | | 3.95778 | 3.72051 |
| CD630_18120 | CD630_n00990 | 164 – 200 | 52 – 95 | -10.9484 | | 3.99713 | 3.72051 |
| CD630_30110 | CD630_n00990 | 137 – 180 | 28 – 69 | -10.9408 | | 3.31041 | 3.72051 |
| CD630_23760 | CD630_n00990 | 151 – 200 | 304 – 352 | -10.9193 | | 4.54307 | 3.72051 |
| pilW | CD630_n00990 | 90 – 104 | 967 – 984 | -10.9144 | | 4.96316 | 3.72051 |
| CD630_08360 | CD630_n00990 | 30 – 66 | 401 – 433 | -10.911 | | 3.19339 | 3.72051 |
| CD630_33570 | CD630_n00990 | 132 – 168 | 790 – 828 | -10.8859 | | 3.95778 | 3.72051 |
| pilD | CD630_n00990 | 35 – 77 | 916 – 956 | -10.8826 | | 4.54123 | 3.72051 |
| CD630_07121 | CD630_n00990 | 44 – 77 | 243 – 273 | -10.8798 | | 3.26177 | 3.72051 |
| CD630_09770 | CD630_n00990 | 123 – 144 | 977 – 996 | -10.8757 | | 3.22106 | 3.72051 |
| CD630_08390 | CD630_n00990 | 87 – 116 | 517 – 543 | -10.8707 | | 5.16000 | 3.72051 |
| CD630_24591 | CD630_n00990 | 59 – 107 | 979 – 1022 | -10.8667 | | 3.54497 | 3.72051 |
| CD630_09030 | CD630_n00990 | 94 – 136 | 975 – 1026 | -10.8541 | | 4.21282 | 3.72051 |
| CD630_06642 | CD630_n00990 | 94 – 123 | 71 – 98 | -10.8442 | | 3.32967 | 3.72051 |
| vncR | CD630_n00990 | 126 – 178 | 950 – 1004 | -10.8432 | | 3.93642 | 3.72051 |
| CD630_31960 | CD630_n00990 | 125 – 146 | 882 – 901 | -10.8368 | | 4.93757 | 3.72051 |
| CD630_05370 | CD630_n00990 | 72 – 107 | 977 – 1007 | -10.8275 | | 3.26947 | 3.72051 |
| CD630_06350 | CD630_n00990 | 139 – 157 | 967 – 984 | -10.8248 | | 4.35983 | 3.72051 |
| CD630_17160 | CD630_n00990 | 85 – 108 | 977 – 1001 | -10.8198 | | 3.50228 | 3.72051 |
| CD630_22991 | CD630_n00990 | 13 – 46 | 32 – 64 | -10.819 | | 6.03728 | 3.72051 |
| CD630_14182 | CD630_n00990 | 16 – 49 | 32 – 64 | -10.7642 | | 4.82360 | 3.72051 |
| CD630_05840 | CD630_n00990 | 52 – 138 | 683 – 779 | -10.7518 | | 4.28779 | 3.72051 |
| CD630_11220 | CD630_n00990 | 138 – 168 | 913 – 945 | -10.6826 | | 3.52327 | 3.72051 |
| serS1 | CD630_n00990 | 138 – 158 | 55 – 74 | -10.6824 | | 3.45858 | 3.72051 |
| CD630_03180 | CD630_n00990 | 81 – 95 | 54 – 68 | -10.667 | | 3.73138 | 3.72051 |
| CD630_15111 | CD630_n00990 | 20 – 45 | 994 – 1021 | -10.6619 | | 4.18664 | 3.72051 |
| CD630_20650 | CD630_n00990 | 26 – 122 | 948 – 1070 | -10.6583 | | 4.00083 | 3.72051 |
| CD630_02720 | CD630_n00990 | 138 – 176 | 856 – 890 | -10.6499 | | 3.38938 | 3.72051 |
| cdtR | CD630_n00990 | 4 – 73 | 748 – 814 | -10.6481 | | 5.84368 | 3.72051 |
| CD630_13380 | CD630_n00990 | 20 – 84 | 929 – 980 | -10.6468 | | 4.74408 | 3.72051 |
| CD630_21110 | CD630_n00990 | 13 – 32 | 168 – 189 | -10.645 | | 3.89630 | 3.72051 |
| CD630_08660 | CD630_n00990 | 52 – 81 | 927 – 957 | -10.6422 | | 3.86467 | 3.72051 |
| CD630_23070 | CD630_n00990 | 176 – 196 | 951 – 971 | -10.6344 | | 5.23027 | 3.72051 |
| rbsC | CD630_n00990 | 160 – 190 | 960 – 1001 | -10.6316 | | 3.71650 | 3.72051 |
| CD630_22890 | CD630_n00990 | 87 – 120 | 31 – 61 | -10.6274 | | 3.75591 | 3.72051 |
| CD630_05220 | CD630_n00990 | 160 – 197 | 977 – 1023 | -10.5906 | | 3.65057 | 3.72051 |
| CD630_22720 | CD630_n00990 | 92 – 147 | 962 – 1007 | -10.582 | | 3.81804 | 3.72051 |
| CD630_24920 | CD630_n00990 | 86 – 96 | 523 – 533 | -10.5695 | | 3.89455 | 3.72051 |
| feoB | CD630_n00990 | 144 – 181 | 984 – 1015 | -10.5691 | | 4.05364 | 3.72051 |
| CD630_33480 | CD630_n00990 | 15 – 33 | 258 – 274 | -10.5633 | | 3.67613 | 3.72051 |
| CD630_03650 | CD630_n00990 | 177 – 197 | 746 – 776 | -10.5234 | | 3.93697 | 3.72051 |
| CD630_01640 | CD630_n00990 | 18 – 56 | 783 – 821 | -10.515 | | 4.55381 | 3.72051 |
| cheW | CD630_n00990 | 71 – 105 | 393 – 427 | -10.5144 | | 3.72371 | 3.72051 |
| CD630_27770 | CD630_n00990 | 87 – 111 | 958 – 984 | -10.5077 | | 4.08588 | 3.72051 |
| CD630_17730 | CD630_n00990 | 77 – 98 | 52 – 69 | -10.4874 | | 4.73397 | 3.72051 |
| CD630_26750 | CD630_n00990 | 166 – 193 | 339 – 370 | -10.4821 | | 3.60610 | 3.72051 |
| CD630_05290 | CD630_n00990 | 6 – 39 | 295 – 325 | -10.4817 | | 3.97604 | 3.72051 |
| CD630_24830 | CD630_n00990 | 21 – 39 | 755 – 774 | -10.4765 | | 4.98239 | 3.72051 |
| CD630_27520 | CD630_n00990 | 111 – 147 | 977 – 1015 | -10.4764 | | 4.27693 | 3.72051 |
| CD630_36190 | CD630_n00990 | 76 – 104 | 967 – 1002 | -10.4681 | | 4.63449 | 3.72051 |
| CD630_27040 | CD630_n00990 | 37 – 69 | 248 – 276 | -10.4565 | | 3.51812 | 3.72051 |
| CD630_07480 | CD630_n00990 | 5 – 48 | 728 – 773 | -10.4513 | | 5.18321 | 3.72051 |
| sspA | CD630_n00990 | 87 – 161 | 894 – 982 | -10.4434 | | 3.62689 | 3.72051 |
| CD630_19440 | CD630_n00990 | 82 – 124 | 822 – 874 | -10.4257 | | 4.53937 | 3.72051 |
| CD630_25200 | CD630_n00990 | 168 – 198 | 753 – 791 | -10.4251 | | 3.65158 | 3.72051 |
| CD630_09850 | CD630_n00990 | 1 – 10 | 975 – 984 | -10.4184 | | 4.18679 | 3.72051 |
| CD630_04770 | CD630_n00990 | 85 – 105 | 414 – 434 | -10.415 | | 3.58478 | 3.72051 |
| CD630_36170 | CD630_n00990 | 148 – 165 | 960 – 981 | -10.4018 | | 3.72380 | 3.72051 |
| CD630_09570 | CD630_n00990 | 16 – 26 | 990 – 1000 | -10.3924 | | 5.31213 | 3.72051 |
| CD630_29070 | CD630_n00990 | 16 – 26 | 990 – 1000 | -10.3924 | | 3.89477 | 3.72051 |
| tcdC | CD630_n00990 | 114 – 151 | 405 – 433 | -10.3616 | | 3.76101 | 3.72051 |
| CD630_33560 | CD630_n00990 | 91 – 128 | 252 – 299 | -10.3573 | | 4.03147 | 3.72051 |
| CD630_34560 | CD630_n00990 | 85 – 104 | 967 – 984 | -10.3304 | | 4.02604 | 3.72051 |
| CD630_21910 | CD630_n00990 | 106 – 142 | 243 – 275 | -10.3289 | | 4.66600 | 3.72051 |
| CD630_11240 | CD630_n00990 | 2 – 39 | 230 – 265 | -10.3286 | | 3.20274 | 3.72051 |

| | | | | | | |
|---|---|---|---|---|---|---|
| CD630_05590 | CD630_n00990 | 65 -- 114 | 388 -- 433 | -10.3019 | 3.39670 | 3.72051 |
| rbsA | CD630_n00990 | 45 -- 74 | 411 -- 443 | -10.2959 | 3.71650 | 3.72051 |
| gcp | CD630_n00990 | 87 -- 145 | 919 -- 982 | -10.2563 | 4.92798 | 3.72051 |
| CD630_18910 | CD630_n00990 | 166 -- 192 | 974 -- 1000 | -10.2546 | 3.30688 | 3.72051 |
| CD630_06560 | CD630_n00990 | 120 -- 167 | 287 -- 326 | -10.2531 | 4.02532 | 3.72051 |
| CD630_05150 | CD630_n00990 | 79 -- 112 | 977 -- 1006 | -10.2518 | 3.69875 | 3.72051 |
| CD630_17970 | CD630_n00990 | 50 -- 123 | 466 -- 531 | -10.2461 | 3.68802 | 3.72051 |
| CD630_16530 | CD630_n00990 | 20 -- 43 | 235 -- 258 | -10.2457 | 4.37658 | 3.72051 |
| gabT | CD630_n00990 | 90 -- 116 | 36 -- 61 | -10.2455 | 3.67067 | 3.72051 |
| CD630_14420 | CD630_n00990 | 90 -- 147 | 414 -- 471 | -10.2409 | 3.99497 | 3.72051 |
| CD630_14700 | CD630_n00990 | 138 -- 155 | 991 -- 1008 | -10.23 | 3.98234 | 3.72051 |
| CD630_25190 | CD630_n00990 | 128 -- 141 | 977 -- 996 | -10.2298 | 3.19663 | 3.72051 |
| CD630_20290 | CD630_n00990 | 119 -- 175 | 230 -- 297 | -10.2154 | 4.38183 | 3.72051 |
| CD630_19801 | CD630_n00990 | 125 -- 177 | 964 -- 1020 | -10.2134 | 5.22035 | 3.72051 |
| CD630_18940 | CD630_n00990 | 89 -- 126 | 822 -- 875 | -10.2083 | 4.09988 | 3.72051 |
| CD630_03420 | CD630_n00990 | 81 -- 136 | 962 -- 1018 | -10.2026 | 4.71868 | 3.72051 |
| spoVFB | CD630_n00990 | 179 -- 200 | 404 -- 428 | -10.1828 | 4.62758 | 3.72051 |
| chrA | CD630_n00990 | 133 -- 154 | 465 -- 481 | -10.1732 | 5.27255 | 3.72051 |
| CD630_03920 | CD630_n00990 | 151 -- 168 | 570 -- 586 | -10.1596 | 4.15934 | 3.72051 |
| CD630_13360 | CD630_n00990 | 165 -- 189 | 250 -- 273 | -10.1483 | 3.77330 | 3.72051 |
| CD630_28580 | CD630_n00990 | 95 -- 110 | 34 -- 49 | -10.1215 | 4.67130 | 3.72051 |
| CD630_18680 | CD630_n00990 | 41 -- 153 | 359 -- 494 | -10.1179 | 3.04315 | 3.72051 |
| CD630_26250 | CD630_n00990 | 139 -- 178 | 292 -- 332 | -10.1156 | 5.03379 | 3.72051 |
| CD630_26020 | CD630_n00990 | 5 -- 56 | 948 -- 999 | -10.0875 | 4.03155 | 3.72051 |
| cprK | CD630_n00990 | 162 -- 201 | 172 -- 212 | -10.0832 | 3.96216 | 3.72051 |
| flgB | CD630_n00990 | 77 -- 185 | 869 -- 1007 | -10.0825 | 3.25736 | 3.72051 |
| CD630_29960 | CD630_n00990 | 1 -- 15 | 878 -- 892 | -10.08 | 4.51904 | 3.72051 |
| CD630_31970 | CD630_n00990 | 159 -- 184 | 741 -- 764 | -10.0785 | 3.24511 | 3.72051 |
| CD630_20920 | CD630_n00990 | 15 -- 74 | 816 -- 873 | -10.0641 | 4.13812 | 3.72051 |
| CD630_27220 | CD630_n00990 | 40 -- 118 | 379 -- 460 | -10.0624 | 4.18786 | 3.72051 |
| CD630_09562 | CD630_n00990 | 18 -- 51 | 32 -- 64 | -10.0471 | 3.94307 | 3.72051 |
| CD630_14150 | CD630_n00990 | 27 -- 71 | 852 -- 897 | -10.0354 | 3.28251 | 3.72051 |
| appB | CD630_n00990 | 140 -- 173 | 717 -- 766 | -10.0105 | 6.44103 | 3.72051 |
| ldh | CD630_n00990 | 180 -- 191 | 984 -- 995 | -10.0094 | 4.35818 | 3.72051 |
| CD630_29270 | CD630_n00990 | 103 -- 175 | 914 -- 982 | -9.99597 | 3.70174 | 3.72051 |
| CD630_21020 | CD630_n00990 | 12 -- 28 | 203 -- 219 | -9.98656 | 4.28666 | 3.72051 |
| CD630_21720 | CD630_n00990 | 155 -- 195 | 28 -- 75 | -9.98503 | 4.13030 | 3.72051 |
| CD630_06060 | CD630_n00990 | 169 -- 186 | 251 -- 268 | -9.98449 | 4.94050 | 3.72051 |
| CD630_36781 | CD630_n00990 | 153 -- 170 | 141 -- 161 | -9.96726 | 3.75946 | 3.72051 |
| dltD | CD630_n00990 | 127 -- 177 | 245 -- 297 | -9.9497 | 4.40777 | 3.72051 |
| glgP | CD630_n00990 | 178 -- 199 | 244 -- 267 | -9.93762 | 3.28530 | 3.72051 |
| CD630_26691 | CD630_n00990 | 1 -- 42 | 937 -- 981 | -9.92325 | 3.96230 | 3.72051 |
| bglA | CD630_n00990 | 92 -- 135 | 31 -- 75 | -9.9226 | 4.50315 | 3.72051 |
| CD630_14250 | CD630_n00990 | 10 -- 46 | 311 -- 344 | -9.91303 | 4.19604 | 3.72051 |
| cobS | CD630_n00990 | 157 -- 173 | 965 -- 987 | -9.90017 | 3.67481 | 3.72051 |
| agrB | CD630_n00990 | 128 -- 146 | 238 -- 257 | -9.89921 | 3.92598 | 3.72051 |
| CD630_22660 | CD630_n00990 | 9 -- 40 | 723 -- 755 | -9.8971 | 3.60496 | 3.72051 |
| eutP | CD630_n00990 | 79 -- 126 | 978 -- 1018 | -9.89618 | 3.82847 | 3.72051 |
| CD630_22710 | CD630_n00990 | 132 -- 197 | 52 -- 117 | -9.89612 | 4.05623 | 3.72051 |
| CD630_21790 | CD630_n00990 | 142 -- 176 | 945 -- 982 | -9.89193 | 4.93762 | 3.72051 |
| CD630_05160 | CD630_n00990 | 4 -- 13 | 666 -- 675 | -9.89042 | 5.03573 | 3.72051 |
| LinCd | CD630_n00990 | 1 -- 36 | 251 -- 275 | -9.83909 | 4.33624 | 3.72051 |
| CD630_17630 | CD630_n00990 | 8 -- 40 | 733 -- 767 | -9.82905 | 4.52408 | 3.72051 |
| CD630_08780 | CD630_n00990 | 161 -- 189 | 241 -- 269 | -9.80724 | 7.76690 | 3.72051 |
| CD630_10220 | CD630_n00990 | 85 -- 174 | 455 -- 535 | -9.79135 | 4.08233 | 3.72051 |
| CD630_09970 | CD630_n00990 | 167 -- 190 | 241 -- 265 | -9.77934 | 3.25917 | 3.72051 |
| CD630_06920 | CD630_n00990 | 40 -- 65 | 786 -- 813 | -9.7575 | 3.13260 | 3.72051 |
| CD630_14190 | CD630_n00990 | 89 -- 171 | 928 -- 984 | -9.75617 | 3.78562 | 3.72051 |
| CD630_19903 | CD630_n00990 | 170 -- 199 | 939 -- 966 | -9.74477 | 5.79994 | 3.72051 |
| etfA | CD630_n00990 | 75 -- 96 | 975 -- 995 | -9.72673 | 4.80521 | 3.72051 |
| CD630_31480 | CD630_n00990 | 9 -- 22 | 706 -- 719 | -9.7266 | 4.88897 | 3.72051 |
| adhE | CD630_n00990 | 160 -- 194 | 257 -- 305 | -9.7265 | 3.06485 | 3.72051 |
| CD630_05211 | CD630_n00990 | 89 -- 104 | 967 -- 983 | -9.72504 | 3.97048 | 3.72051 |
| CD630_14210 | CD630_n00990 | 4 -- 39 | 252 -- 287 | -9.71386 | 3.82355 | 3.72051 |
| CD630_15431 | CD630_n00990 | 140 -- 172 | 38 -- 76 | -9.69432 | 5.29780 | 3.72051 |
| CD630_31382 | CD630_n00990 | 1 -- 20 | 521 -- 539 | -9.67634 | 3.49137 | 3.72051 |
| CD630_33682 | CD630_n00990 | 70 -- 112 | 966 -- 999 | -9.66633 | 5.77720 | 3.72051 |
| CD630_13620 | CD630_n00990 | 32 -- 71 | 830 -- 872 | -9.64396 | 4.77627 | 3.72051 |
| CD630_21540 | CD630_n00990 | 31 -- 45 | 251 -- 266 | -9.62739 | 4.33090 | 3.72051 |
| CD630_21750 | CD630_n00990 | 49 -- 92 | 978 -- 1022 | -9.6051 | 3.70626 | 3.72051 |
| CD630_34080 | CD630_n00990 | 22 -- 78 | 851 -- 911 | -9.58773 | 3.20224 | 3.72051 |
| CD630_17380 | CD630_n00990 | 184 -- 196 | 519 -- 531 | -9.5845 | 5.03207 | 3.72051 |
| CD630_32500 | CD630_n00990 | 53 -- 101 | 971 -- 1012 | -9.55775 | 6.17288 | 3.72051 |
| CD630_21460 | CD630_n00990 | 180 -- 201 | 989 -- 1011 | -9.51771 | 4.83005 | 3.72051 |
| CD630_17151 | CD630_n00990 | 132 -- 191 | 816 -- 871 | -9.51133 | 6.43124 | 3.72051 |
| CD630_22850 | CD630_n00990 | 16 -- 57 | 718 -- 769 | -9.50565 | 3.84102 | 3.72051 |
| CD630_14010 | CD630_n00990 | 40 -- 86 | 722 -- 767 | -9.50385 | 3.69983 | 3.72051 |
| CD630_07070 | CD630_n00990 | 32 -- 47 | 882 -- 896 | -9.50138 | 3.46414 | 3.72051 |
| CD630_19790 | CD630_n01010 | 49 -- 93 | 839 -- 884 | -19.7527 | 3.79593 | 4.59146 |
| cspD | CD630_n01010 | 122 -- 183 | 368 -- 425 | -19.6734 | 3.23587 | 4.59146 |
| cobQ | CD630_n01010 | 51 -- 107 | 850 -- 910 | -19.6043 | 3.13524 | 4.59146 |
| CD630_03220 | CD630_n01010 | 40 -- 135 | 777 -- 877 | -19.5777 | 3.36746 | 4.59146 |
| CD630_09760 | CD630_n01010 | 6 -- 152 | 803 -- 940 | -19.5486 | 4.83410 | 4.59146 |
| CD630_28910 | CD630_n01010 | 6 -- 152 | 803 -- 940 | -19.5486 | 4.51664 | 4.59146 |
| CD630_32700 | CD630_n01010 | 59 -- 201 | 1062 -- 1206 | -19.1162 | 4.74276 | 4.59146 |
| argS | CD630_n01010 | 64 -- 105 | 1100 -- 1143 | -18.8316 | 4.96036 | 4.59146 |
| CD630_08360 | CD630_n01010 | 30 -- 171 | 793 -- 941 | -18.8018 | 3.19339 | 4.59146 |
| CD630_02790 | CD630_n01010 | 85 -- 199 | 3 -- 104 | -18.7661 | 4.77661 | 4.59146 |
| CD630_22890 | CD630_n01010 | 59 -- 142 | 789 -- 881 | -18.7455 | 3.75591 | 4.59146 |
| cdu2 | CD630_n01010 | 72 -- 109 | 828 -- 870 | -18.5464 | 2.97335 | 4.59146 |
| CD630_08390 | CD630_n01010 | 5 -- 107 | 793 -- 902 | -18.3239 | 5.16000 | 4.59146 |
| etfA | CD630_n01010 | 64 -- 95 | 1388 -- 1420 | -18.2495 | 4.80521 | 4.59146 |
| CD630_18970 | CD630_n01010 | 10 -- 158 | 1085 -- 1208 | -18.2076 | 3.57243 | 4.59146 |
| CD630_15850 | CD630_n01010 | 155 -- 197 | 1013 -- 1048 | -18.1659 | 4.32968 | 4.59146 |
| CD630_27280 | CD630_n01010 | 4 -- 73 | 1258 -- 1326 | -18.1137 | 3.24738 | 4.59146 |
| CD630_11730 | CD630_n01010 | 131 -- 192 | 840 -- 910 | -18.074 | 4.80521 | 4.59146 |
| glgB | CD630_n01010 | 139 -- 201 | 382 -- 448 | -18.0532 | 4.48273 | 4.59146 |
| CD630_10800 | CD630_n01010 | 86 -- 194 | 99 -- 199 | -18.0466 | 5.30319 | 4.59146 |
| gabT | CD630_n01010 | 29 -- 140 | 38 -- 128 | -17.9802 | 3.67067 | 4.59146 |
| CD630_06340 | CD630_n01010 | 74 -- 108 | 315 -- 343 | -17.9445 | 4.35983 | 4.59146 |
| vncR | CD630_n01010 | 31 -- 147 | 793 -- 909 | -17.8771 | 3.93642 | 4.59146 |
| CD630_32060 | CD630_n01010 | 5 -- 74 | 48 -- 119 | -17.8257 | 5.73797 | 4.59146 |
| CD630_28550 | CD630_n01010 | 69 -- 105 | 833 -- 876 | -17.777 | 3.51196 | 4.59146 |
| nusB | CD630_n01010 | 114 -- 194 | 328 -- 417 | -17.75 | 4.92798 | 4.59146 |
| CD630_08100 | CD630_n01010 | 56 -- 144 | 802 -- 882 | -17.7143 | 4.39181 | 4.59146 |
| bglA | CD630_n01010 | 57 -- 110 | 752 -- 803 | -17.7133 | 4.50315 | 4.59146 |
| CD630_06080 | CD630_n01010 | 39 -- 160 | 794 -- 896 | -17.6828 | 3.82315 | 4.59146 |
| ung | CD630_n01010 | 119 -- 166 | 819 -- 857 | -17.6463 | 3.30908 | 4.59146 |
| cprK | CD630_n01010 | 1 -- 112 | 315 -- 416 | -17.5507 | 3.96216 | 4.59146 |
| CD630_04620 | CD630_n01010 | 42 -- 188 | 776 -- 895 | -17.5464 | 3.45413 | 4.59146 |
| aroD | CD630_n01010 | 123 -- 194 | 863 -- 932 | -17.4421 | 3.71929 | 4.59146 |
| xdhA | CD630_n01010 | 48 -- 110 | 1099 -- 1147 | -17.3175 | 4.45147 | 4.59146 |
| CD630_11241 | CD630_n01010 | 160 -- 182 | 835 -- 857 | -17.3154 | 4.85723 | 4.59146 |
| CD630_05950 | CD630_n01010 | 29 -- 70 | 834 -- 875 | -17.0669 | 3.31669 | 4.59146 |
| CD630_06060 | CD630_n01010 | 95 -- 130 | 39 -- 71 | -16.9457 | 4.94050 | 4.59146 |
| pilD | CD630_n01010 | 35 -- 91 | 841 -- 895 | -16.9212 | 4.54123 | 4.59146 |
| CD630_20920 | CD630_n01010 | 114 -- 139 | 1389 -- 1414 | -16.9024 | 4.13812 | 4.59146 |
| CD630_19700 | CD630_n01010 | 151 -- 197 | 1116 -- 1159 | -16.8673 | 3.85861 | 4.59146 |
| CD630_18650 | CD630_n01010 | 76 -- 100 | 846 -- 868 | -16.8656 | 3.75892 | 4.59146 |
| tcdC | CD630_n01010 | 118 -- 155 | 186 -- 223 | -16.7448 | 3.76101 | 4.59146 |
| CD630_04540 | CD630_n01010 | 131 -- 180 | 1088 -- 1125 | -16.7291 | 3.33556 | 4.59146 |
| CD630_20450 | CD630_n01010 | 69 -- 100 | 1387 -- 1418 | -16.6997 | 3.77388 | 4.59146 |
| CD630_17292 | CD630_n01010 | 79 -- 147 | 777 -- 856 | -16.696 | 6.25799 | 4.59146 |
| CD630_17970 | CD630_n01010 | 100 -- 183 | 1314 -- 1415 | -16.5641 | 3.68802 | 4.59146 |
| bglA | CD630_n01010 | 151 -- 190 | 845 -- 884 | -16.5317 | 4.86402 | 4.59146 |
| CD630_35840 | CD630_n01010 | 11 -- 85 | 38 -- 124 | -16.5263 | 3.14170 | 4.59146 |
| CD630_32520 | CD630_n01010 | 150 -- 193 | 169 -- 209 | -16.4465 | 4.06098 | 4.59146 |
| glgP | CD630_n01010 | 31 -- 130 | 39 -- 127 | -16.357 | 3.28530 | 4.59146 |
| CD630_11220 | CD630_n01010 | 138 -- 192 | 1098 -- 1147 | -16.2774 | 3.52327 | 4.59146 |
| CD630_32070 | CD630_n01010 | 52 -- 85 | 839 -- 875 | -16.1538 | 3.51964 | 4.59146 |
| CD630_14630 | CD630_n01010 | 43 -- 149 | 795 -- 904 | -16.0938 | 3.34485 | 4.59146 |
| spaK | CD630_n01010 | 109 -- 129 | 4 -- 24 | -16.0732 | 3.04530 | 4.59146 |
| CD630_31580 | CD630_n01010 | 131 -- 192 | 58 -- 119 | -15.9483 | 3.51756 | 4.59146 |
| blaI | CD630_n01010 | 10 -- 116 | 822 -- 939 | -15.9302 | 3.73674 | 4.59146 |
| moaA | CD630_n01010 | 1 -- 71 | 796 -- 861 | -15.9003 | 3.83799 | 4.59146 |
| CD630_32050 | CD630_n01010 | 166 -- 196 | 175 -- 208 | -15.8762 | 5.73797 | 4.59146 |
| CD630_14420 | CD630_n01010 | 40 -- 181 | 793 -- 910 | -15.8561 | 3.99497 | 4.59146 |
| CD630_24590 | CD630_n01010 | 110 -- 198 | 37 -- 140 | -15.8482 | 3.70401 | 4.59146 |
| cspC | CD630_n01010 | 81 -- 173 | 798 -- 884 | -15.8349 | 5.48848 | 4.59146 |
| CD630_22050 | CD630_n01010 | 84 -- 121 | 1100 -- 1138 | -15.8215 | 4.87639 | 4.59146 |
| CD630_23780 | CD630_n01010 | 32 -- 96 | 324 -- 387 | -15.8196 | 3.91629 | 4.59146 |
| CD630_18120 | CD630_n01010 | 61 -- 112 | 839 -- 889 | -15.8186 | 3.99713 | 4.59146 |
| mobA | CD630_n01010 | 119 -- 188 | 38 -- 115 | -15.8174 | 4.25703 | 4.59146 |
| CD630_08510 | CD630_n01010 | 27 -- 175 | 126 -- 268 | -15.7812 | 5.16329 | 4.59146 |
| CD630_02420 | CD630_n01010 | 112 -- 144 | 979 -- 1008 | -15.7594 | 3.71048 | 4.59146 |
| CD630_10880 | CD630_n01010 | 167 -- 200 | 1089 -- 1130 | -15.7349 | 5.05048 | 4.59146 |
| CD630_04341 | CD630_n01010 | 160 -- 194 | 824 -- 863 | -15.7223 | 4.29262 | 4.59146 |
| pilT | CD630_n01010 | 135 -- 200 | 845 -- 910 | -15.7223 | 3.24639 | 4.59146 |
| CD630_06970 | CD630_n01010 | 39 -- 173 | 1043 -- 1190 | -15.713 | 3.82312 | 4.59146 |
| CD630_16460 | CD630_n01010 | 98 -- 162 | 604 -- 650 | -15.7095 | 4.64028 | 4.59146 |
| CD630_14190 | CD630_n01010 | 139 -- 189 | 1109 -- 1158 | -15.6977 | 3.78562 | 4.59146 |
| CD630_15740 | CD630_n01010 | 25 -- 118 | 35 -- 124 | -15.671 | 4.43222 | 4.59146 |
| CD630_15200 | CD630_n01010 | 5 -- 112 | 821 -- 930 | -15.6677 | 4.00442 | 4.59146 |
| CD630_31910 | CD630_n01010 | 86 -- 182 | 806 -- 884 | -15.6452 | 3.48314 | 4.59146 |
| hfq | CD630_n01010 | 145 -- 195 | 410 -- 463 | -15.6342 | 5.05609 | 4.59146 |
| CD630_16150 | CD630_n01010 | 133 -- 196 | 820 -- 875 | -15.6284 | 4.20614 | 4.59146 |
| CD630_08940 | CD630_n01010 | 74 -- 95 | 1390 -- 1418 | -15.613 | 3.72225 | 4.59146 |
| CD630_18610 | CD630_n01010 | 7 -- 43 | 39 -- 82 | -15.5887 | 4.15582 | 4.59146 |
| CD630_17151 | CD630_n01010 | 52 -- 201 | 796 -- 945 | -15.5865 | 6.43124 | 4.59146 |
| eutV | CD630_n01010 | 15 -- 161 | 798 -- 936 | -15.5851 | 4.55715 | 4.59146 |
| CD630_31880 | CD630_n01010 | 133 -- 187 | 819 -- 875 | -15.5521 | 3.97522 | 4.59146 |
| CD630_32690 | CD630_n01010 | 68 -- 150 | 796 -- 880 | -15.5295 | 7.32676 | 4.59146 |
| metQ | CD630_n01010 | 48 -- 148 | 39 -- 154 | -15.5067 | 3.87190 | 4.59146 |
| CD630_31000 | CD630_n01010 | 55 -- 124 | 55 -- 144 | -15.4821 | 3.70946 | 4.59146 |
| CD630_10220 | CD630_n01010 | 127 -- 169 | 613 -- 656 | -15.464 | 4.08233 | 4.59146 |
| CD630_33600 | CD630_n01010 | 125 -- 172 | 839 -- 893 | -15.4572 | 3.73196 | 4.59146 |
| CD630_16670 | CD630_n01010 | 96 -- 200 | 1029 -- 1138 | -15.415 | 3.33553 | 4.59146 |
| CD630_32880 | CD630_n01010 | 90 -- 180 | 793 -- 875 | -15.3639 | 3.41276 | 4.59146 |
| CD630_28260 | CD630_n01010 | 50 -- 100 | 852 -- 902 | -15.2955 | 3.51613 | 4.59146 |
| gatC | CD630_n01010 | 40 -- 148 | 1067 -- 1184 | -15.2742 | 4.25143 | 4.59146 |
| sspA | CD630_n01010 | 90 -- 160 | 1108 -- 1167 | -15.2639 | 3.62689 | 4.59146 |
| CD630_22200 | CD630_n01010 | 79 -- 122 | 1359 -- 1412 | -15.2408 | 4.18288 | 4.59146 |
| CD630_04410 | CD630_n01010 | 128 -- 185 | 802 -- 877 | -15.2126 | 3.90451 | 4.59146 |
| pgm | CD630_n01010 | 84 -- 176 | 794 -- 861 | -15.204 | 3.45439 | 4.59146 |
| moeA | CD630_n01010 | 49 -- 123 | 822 -- 895 | -15.1843 | 3.26118 | 4.59146 |
| eutW | CD630_n01010 | 75 -- 169 | 335 -- 432 | -15.1655 | 4.55715 | 4.59146 |
| CD630_24591 | CD630_n01010 | 7 -- 95 | 847 -- 943 | -15.1151 | 3.54497 | 4.59146 |
| rolA | CD630_n01010 | 14 -- 50 | 396 -- 433 | -15.0499 | 4.56362 | 4.59146 |
| CD630_21570 | CD630_n01010 | 30 -- 173 | 309 -- 444 | -15.0258 | 6.48385 | 4.59146 |
| CD630_08460 | CD630_n01010 | 22 -- 113 | 834 -- 914 | -15.0077 | 4.15083 | 4.59146 |
| flgB | CD630_n01010 | 62 -- 115 | 822 -- 874 | -15.0062 | 3.25736 | 4.59146 |
| CD630_10050 | CD630_n01010 | 160 -- 188 | 791 -- 819 | -15.0056 | 3.93750 | 4.59146 |
| CD630_29760 | CD630_n01010 | 109 -- 176 | 818 -- 874 | -14.9905 | 3.51074 | 4.59146 |
| CD630_17160 | CD630_n01010 | 87 -- 109 | 839 -- 861 | -14.9175 | 3.50228 | 4.59146 |
| CD630_15360 | CD630_n01010 | 114 -- 156 | 842 -- 885 | -14.8646 | 4.47208 | 4.59146 |
| CD630_27570 | CD630_n01010 | 99 -- 127 | 1391 -- 1416 | -14.8357 | 3.46216 | 4.59146 |
| CD630_21140 | CD630_n01010 | 122 -- 194 | 38 -- 122 | -14.7649 | 3.33304 | 4.59146 |
| CD630_16520 | CD630_n01010 | 45 -- 191 | 802 -- 912 | -14.7512 | 4.28395 | 4.59146 |
| CD630_07770 | CD630_n01010 | 29 -- 132 | 1085 -- 1166 | -14.75 | 3.31158 | 4.59146 |
| CD630_01690 | CD630_n01010 | 164 -- 190 | 424 -- 448 | -14.6861 | 4.97234 | 4.59146 |
| CD630_14130 | CD630_n01010 | 81 -- 183 | 773 -- 875 | -14.6736 | 4.57970 | 4.59146 |
| CD630_27520 | CD630_n01010 | 90 -- 183 | 601 -- 720 | -14.6146 | 4.27693 | 4.59146 |
| CD630_19680 | CD630_n01010 | 136 -- 145 | 1388 -- 1397 | -14.5536 | 3.58080 | 4.59146 |
| ssuC | CD630_n01010 | 65 -- 81 | 845 -- 863 | -14.5331 | 3.45491 | 4.59146 |
| dnaC | CD630_n01010 | 88 -- 154 | 793 -- 857 | -14.5261 | 6.30796 | 4.59146 |
| CD630_30120 | CD630_n01010 | 68 -- 144 | 794 -- 871 | -14.503 | 4.04859 | 4.59146 |
| CD630_08370 | CD630_n01010 | 122 -- 155 | 841 -- 875 | -14.47 | 3.65639 | 4.59146 |
| CD630_14600 | CD630_n01010 | 31 -- 84 | 752 -- 802 | -14.4558 | 5.90348 | 4.59146 |
| CD630_14410 | CD630_n01010 | 82 -- 114 | 176 -- 212 | -14.4406 | 3.39433 | 4.59146 |
| oppC | CD630_n01010 | 160 -- 191 | 240 -- 271 | -14.4393 | 3.01695 | 4.59146 |
| CD630_33570 | CD630_n01010 | 93 -- 180 | 308 -- 394 | -14.4237 | 3.95778 | 4.59146 |
| CD630_07480 | CD630_n01010 | 84 -- 111 | 846 -- 872 | -14.3894 | 5.18321 | 4.59146 |
| CD630_09040 | CD630_n01010 | 116 -- 195 | 407 -- 468 | -14.3891 | 4.11851 | 4.59146 |
| CD630_22660 | CD630_n01010 | 17 -- 62 | 136 -- 182 | -14.3673 | 3.60496 | 4.59146 |
| LinCd | CD630_n01010 | 6 -- 71 | 74 -- 129 | -14.3518 | 4.33624 | 4.59146 |
| CD630_27770 | CD630_n01010 | 155 -- 188 | 1203 -- 1235 | -14.3485 | 4.08588 | 4.59146 |
| htpG | CD630_n01010 | 30 -- 98 | 846 -- 911 | -14.3456 | 3.45147 | 4.59146 |
| CD630_15890 | CD630_n01010 | 148 -- 180 | 1392 -- 1417 | -14.3215 | 3.39962 | 4.59146 |
| CD630_19780 | CD630_n01010 | 14 -- 34 | 837 -- 857 | -14.3155 | 3.64830 | 4.59146 |
| CD630_14700 | CD630_n01010 | 50 -- 159 | 315 -- 460 | -14.3047 | 3.98234 | 4.59146 |
| CD630_35260 | CD630_n01010 | 29 -- 104 | 843 -- 919 | -14.2643 | 5.77277 | 4.59146 |
| iorA | CD630_n01010 | 99 -- 155 | 177 -- 221 | -14.1978 | 3.83173 | 4.59146 |
| CD630_18910 | CD630_n01010 | 106 -- 198 | 361 -- 472 | -14.1585 | 3.30688 | 4.59146 |
| CD630_25610 | CD630_n01010 | 90 -- 154 | 265 -- 320 | -14.142 | 3.02685 | 4.59146 |
| CD630_34530 | CD630_n01010 | 1 -- 146 | 75 -- 219 | -14.1264 | 3.83016 | 4.59146 |
| CD630_21750 | CD630_n01010 | 41 -- 119 | 1069 -- 1165 | -14.0617 | 3.70626 | 4.59146 |
| CD630_18680 | CD630_n01010 | 86 -- 141 | 794 -- 861 | -14.0269 | 3.04315 | 4.59146 |
| CD630_20460 | CD630_n01010 | 69 -- 109 | 842 -- 880 | -14.0162 | 7.94034 | 4.59146 |
| pilW | CD630_n01010 | 61 -- 110 | 837 -- 902 | -14.0073 | 4.96316 | 4.59146 |
| CD630_13400 | CD630_n01010 | 107 -- 160 | 796 -- 848 | -13.9909 | 3.20232 | 4.59146 |
| CD630_21910 | CD630_n01010 | 71 -- 105 | 842 -- 876 | -13.9761 | 4.66600 | 4.59146 |
| CD630_31382 | CD630_n01010 | 171 -- 192 | 1234 -- 1255 | -13.9532 | 3.49137 | 4.59146 |
| CD630_27610 | CD630_n01010 | 15 -- 35 | 607 -- 627 | -13.9419 | 3.96465 | 4.59146 |
| CD630_20670 | CD630_n01010 | 36 -- 106 | 832 -- 902 | -13.9394 | 4.42757 | 4.59146 |
| CD630_06060 | CD630_n01010 | 137 -- 158 | 1107 -- 1128 | -13.9373 | 4.94050 | 4.59146 |
| CD630_13380 | CD630_n01010 | 12 -- 98 | 834 -- 924 | -13.9306 | 4.74408 | 4.59146 |
| CD630_23280 | CD630_n01010 | 53 -- 120 | 389 -- 450 | -13.9278 | 3.50093 | 4.59146 |
| CD630_07760 | CD630_n01010 | 28 -- 101 | 843 -- 915 | -13.9262 | 3.64186 | 4.59146 |
| CD630_21540 | CD630_n01010 | 22 -- 94 | 644 -- 718 | -13.89 | 4.33090 | 4.59146 |
| CD630_18230 | CD630_n01010 | 6 -- 36 | 323 -- 351 | -13.8695 | 3.00184 | 4.59146 |
| CD630_18090 | CD630_n01010 | 18 -- 91 | 61 -- 129 | -13.856 | 4.23961 | 4.59146 |
| CD630_23450 | CD630_n01010 | 39 -- 72 | 1282 -- 1317 | -13.8146 | 3.27218 | 4.59146 |
| scpB | CD630_n01010 | 113 -- 162 | 311 -- 354 | -13.8145 | 3.33560 | 4.59146 |
| CD630_05450 | CD630_n01010 | 84 -- 98 | 1386 -- 1399 | -13.8044 | 5.04761 | 4.59146 |
| CD630_34520 | CD630_n01010 | 76 -- 197 | 779 -- 891 | -13.7657 | 3.83016 | 4.59146 |
| CD630_02430 | CD630_n01010 | 54 -- 133 | 79 -- 153 | -13.7602 | 3.71048 | 4.59146 |
| CD630_19903 | CD630_n01010 | 69 -- 146 | 1344 -- 1415 | -13.7598 | 5.79994 | 4.59146 |
| CD630_06050 | CD630_n01010 | 47 -- 81 | 1390 -- 1418 | -13.7541 | 3.59332 | 4.59146 |

| | | | | | | | |
|---|---|---|---|---|---|---|---|
| bcp | CD630_n01010 | 39 -- 136 | 1091 -- 1179 | -13.7403 | | 3.16670 | 4.59146 |
| CD630_18440 | CD630_n01010 | 150 -- 179 | 841 -- 871 | -13.7307 | | 4.07960 | 4.59146 |
| CD630_09770 | CD630_n01010 | 6 -- 34 | 452 -- 490 | -13.719 | | 3.22106 | 4.59146 |
| gcvTPA | CD630_n01010 | 140 -- 178 | 291 -- 330 | -13.7047 | | 4.77229 | 4.59146 |
| CD630_04940 | CD630_n01010 | 38 -- 92 | 59 -- 114 | -13.6668 | | 4.20948 | 4.59146 |
| CD630_12320 | CD630_n01010 | 44 -- 62 | 846 -- 862 | -13.655 | | 3.28667 | 4.59146 |
| CD630_20660 | CD630_n01010 | 186 -- 201 | 598 -- 613 | -13.6508 | | 5.39321 | 4.59146 |
| serS1 | CD630_n01010 | 105 -- 146 | 380 -- 422 | -13.6468 | | 3.45858 | 4.59146 |
| appC | CD630_n01010 | 39 -- 74 | 138 -- 180 | -13.6376 | | 4.50438 | 4.59146 |
| sigK | CD630_n01010 | 93 -- 153 | 1273 -- 1336 | -13.6212 | | 3.08122 | 4.59146 |
| CD630_11240 | CD630_n01010 | 135 -- 170 | 851 -- 875 | -13.5734 | | 3.20274 | 4.59146 |
| CD630_25320 | CD630_n01010 | 78 -- 147 | 800 -- 875 | -13.5658 | | 4.29415 | 4.59146 |
| CD630_28320 | CD630_n01010 | 1 -- 141 | 797 -- 915 | -13.5177 | | 3.99184 | 4.59146 |
| CD630_20600 | CD630_n01010 | 79 -- 193 | 82 -- 219 | -13.5047 | | 4.20093 | 4.59146 |
| srlD | CD630_n01010 | 88 -- 140 | 792 -- 854 | -13.497 | | 4.19843 | 4.59146 |
| CD630_03040 | CD630_n01010 | 75 -- 98 | 442 -- 465 | -13.4929 | | 3.51403 | 4.59146 |
| CD630_26630 | CD630_n01010 | 55 -- 99 | 1113 -- 1167 | -13.4861 | | 4.61557 | 4.59146 |
| CD630_01590 | CD630_n01010 | 27 -- 171 | 774 -- 921 | -13.4529 | | 3.45609 | 4.59146 |
| CD630_04850 | CD630_n01010 | 152 -- 188 | 1016 -- 1057 | -13.4334 | | 4.40788 | 4.59146 |
| CD630_30310 | CD630_n01010 | 88 -- 97 | 973 -- 982 | -13.4279 | | 4.56059 | 4.59146 |
| CD630_27110 | CD630_n01010 | 20 -- 118 | 292 -- 399 | -13.3992 | | 3.78231 | 4.59146 |
| rbsC | CD630_n01010 | 8 -- 62 | 1367 -- 1419 | -13.388 | | 3.71650 | 4.59146 |
| CD630_22950 | CD630_n01010 | 126 -- 174 | 100 -- 160 | -13.3825 | | 3.96883 | 4.59146 |
| CD630_19904 | CD630_n01010 | 137 -- 171 | 1098 -- 1130 | -13.381 | | 5.96526 | 4.59146 |
| CD630_28181 | CD630_n01010 | 51 -- 79 | 850 -- 877 | -13.3566 | | 3.91299 | 4.59146 |
| cbpA | CD630_n01010 | 85 -- 95 | 976 -- 986 | -13.3196 | | 3.03040 | 4.59146 |
| CD630_14360 | CD630_n01010 | 87 -- 113 | 834 -- 863 | -13.2993 | | 3.02079 | 4.59146 |
| CD630_30420 | CD630_n01010 | 140 -- 172 | 291 -- 323 | -13.2892 | | 5.27539 | 4.59146 |
| CD630_05160 | CD630_n01010 | 177 -- 201 | 389 -- 417 | -13.2843 | | 5.03573 | 4.59146 |
| CD630_06160 | CD630_n01010 | 145 -- 180 | 1033 -- 1066 | -13.2753 | | 5.51744 | 4.59146 |
| CD630_21490 | CD630_n01010 | 175 -- 197 | 1390 -- 1410 | -13.2678 | | 3.37462 | 4.59146 |
| dnaD | CD630_n01010 | 3 -- 100 | 846 -- 938 | -13.266 | | 3.56078 | 4.59146 |
| def1 | CD630_n01010 | 80 -- 139 | 792 -- 863 | -13.2587 | | 3.99720 | 4.59146 |
| CD630_13350 | CD630_n01010 | 179 -- 199 | 888 -- 910 | -13.2526 | | 4.15139 | 4.59146 |
| pmi | CD630_n01010 | 132 -- 150 | 847 -- 864 | -13.2189 | | 3.86845 | 4.59146 |
| CD630_24760 | CD630_n01010 | 61 -- 156 | 841 -- 937 | -13.2178 | | 5.19485 | 4.59146 |
| CD630_09970 | CD630_n01010 | 3 -- 62 | 68 -- 126 | -13.217 | | 3.25917 | 4.59146 |
| scrR | CD630_n01010 | 79 -- 102 | 438 -- 461 | -13.2127 | | 3.49166 | 4.59146 |
| CD630_14340 | CD630_n01010 | 102 -- 195 | 34 -- 129 | -13.197 | | 5.10868 | 4.59146 |
| CD630_08130 | CD630_n01010 | 176 -- 195 | 1397 -- 1418 | -13.1912 | | 4.63831 | 4.59146 |
| CD630_32990 | CD630_n01010 | 41 -- 70 | 1348 -- 1376 | -13.1792 | | 5.24925 | 4.59146 |
| fliI | CD630_n01010 | 82 -- 185 | 795 -- 891 | -13.1651 | | 3.03666 | 4.59146 |
| CD630_27310 | CD630_n01010 | 29 -- 105 | 210 -- 290 | -13.1234 | | 4.01622 | 4.59146 |
| panG | CD630_n01010 | 9 -- 66 | 336 -- 410 | -13.0999 | | 5.54047 | 4.59146 |
| CD630_21760 | CD630_n01010 | 179 -- 192 | 1053 -- 1066 | -13.0664 | | 4.43846 | 4.59146 |
| CD630_10000 | CD630_n01010 | 134 -- 167 | 847 -- 880 | -13.0578 | | 3.86040 | 4.59146 |
| feoB | CD630_n01010 | 4 -- 20 | 797 -- 813 | -13.0505 | | 4.05364 | 4.59146 |
| CD630_03400 | CD630_n01010 | 97 -- 122 | 853 -- 872 | -13.0365 | | 4.45994 | 4.59146 |
| CD630_30100 | CD630_n01010 | 3 -- 83 | 381 -- 458 | -13.0206 | | 4.13234 | 4.59146 |
| CD630_05230 | CD630_n01010 | 27 -- 54 | 840 -- 866 | -13.0204 | | 3.73787 | 4.59146 |
| CD630_04320 | CD630_n01010 | 82 -- 108 | 1388 -- 1418 | -13.0042 | | 3.68342 | 4.59146 |
| CD630_06060 | CD630_n01010 | 67 -- 96 | 839 -- 866 | -12.9693 | | 4.94050 | 4.59146 |
| CD630_23770 | CD630_n01010 | 113 -- 194 | 1081 -- 1163 | -12.9679 | | 3.25181 | 4.59146 |
| CD630_18930 | CD630_n01010 | 111 -- 200 | 137 -- 215 | -12.9379 | | 5.32127 | 4.59146 |
| CD630_22500 | CD630_n01010 | 5 -- 40 | 1315 -- 1352 | -12.9305 | | 4.03688 | 4.59146 |
| CD630_34560 | CD630_n01010 | 10 -- 25 | 1339 -- 1354 | -12.9201 | | 4.02604 | 4.59146 |
| spoVFB | CD630_n01010 | 176 -- 198 | 1395 -- 1420 | -12.9149 | | 4.62758 | 4.59146 |
| malY | CD630_n01010 | 166 -- 187 | 1390 -- 1418 | -12.8835 | | 4.46345 | 4.59146 |
| CD630_25620 | CD630_n01010 | 110 -- 142 | 378 -- 405 | -12.825 | | 3.02685 | 4.59146 |
| CD630_27270 | CD630_n01010 | 160 -- 174 | 964 -- 981 | -12.8087 | | 3.50892 | 4.59146 |
| CD630_05590 | CD630_n01010 | 67 -- 117 | 833 -- 880 | -12.8074 | | 3.39670 | 4.59146 |
| CD630_28560 | CD630_n01010 | 152 -- 200 | 1131 -- 1183 | -12.806 | | 3.96803 | 4.59146 |
| CD630_24050 | CD630_n01010 | 103 -- 116 | 391 -- 404 | -12.7763 | | 3.70192 | 4.59146 |
| CD630_02720 | CD630_n01010 | 105 -- 141 | 1258 -- 1294 | -12.7748 | | 3.38938 | 4.59146 |
| CD630_08660 | CD630_n01010 | 52 -- 82 | 1128 -- 1159 | -12.7648 | | 3.86467 | 4.59146 |
| CD630_14260 | CD630_n01010 | 22 -- 72 | 192 -- 237 | -12.7036 | | 3.22214 | 4.59146 |
| CD630_16550 | CD630_n01010 | 73 -- 108 | 839 -- 872 | -12.7035 | | 5.61064 | 4.59146 |
| nrdD | CD630_n01010 | 12 -- 26 | 970 -- 983 | -12.6693 | | 3.28290 | 4.59146 |
| CD630_07121 | CD630_n01010 | 77 -- 98 | 1385 -- 1404 | -12.6583 | | 3.26177 | 4.59146 |
| CD630_25190 | CD630_n01010 | 132 -- 138 | 976 -- 982 | -12.653 | | 3.19663 | 4.59146 |
| CD630_17710 | CD630_n01010 | 55 -- 165 | 34 -- 124 | -12.6423 | | 4.37688 | 4.59146 |
| CD630_28580 | CD630_n01010 | 34 -- 54 | 1391 -- 1411 | -12.6322 | | 4.67130 | 4.59146 |
| CD630_28280 | CD630_n01010 | 14 -- 61 | 79 -- 112 | -12.6292 | | 4.59833 | 4.59146 |
| CD630_26050 | CD630_n01010 | 158 -- 175 | 1391 -- 1407 | -12.6146 | | 6.99851 | 4.59146 |
| terD | CD630_n01010 | 17 -- 65 | 1281 -- 1323 | -12.6113 | | 3.33148 | 4.59146 |
| CD630_08950 | CD630_n01010 | 34 -- 64 | 1085 -- 1116 | -12.5938 | | 3.71905 | 4.59146 |
| CD630_18400 | CD630_n01010 | 109 -- 196 | 349 -- 460 | -12.5803 | | 4.67962 | 4.59146 |
| CD630_13920 | CD630_n01010 | 132 -- 158 | 39 -- 65 | -12.5668 | | 3.25236 | 4.59146 |
| CD630_06350 | CD630_n01010 | 65 -- 104 | 843 -- 883 | -12.5532 | | 4.35983 | 4.59146 |
| CD630_29840 | CD630_n01010 | 21 -- 69 | 38 -- 85 | -12.5307 | | 4.48825 | 4.59146 |
| CD630_19430 | CD630_n01010 | 175 -- 200 | 982 -- 1013 | -12.5207 | | 3.13922 | 4.59146 |
| CD630_20171 | CD630_n01010 | 146 -- 183 | 840 -- 880 | -12.5003 | | 6.87491 | 4.59146 |
| dapB2 | CD630_n01010 | 169 -- 197 | 1393 -- 1418 | -12.485 | | 3.27218 | 4.59146 |
| CD630_05211 | CD630_n01010 | 110 -- 188 | 1307 -- 1387 | -12.485 | | 3.97048 | 4.59146 |
| cheW | CD630_n01010 | 4 -- 98 | 1120 -- 1204 | -12.4604 | | 3.72371 | 4.59146 |
| CD630_29750 | CD630_n01010 | 77 -- 115 | 1369 -- 1407 | -12.4475 | | 4.59146 | 4.59146 |
| CD630_22710 | CD630_n01010 | 57 -- 199 | 795 -- 943 | -12.4219 | | 4.05623 | 4.59146 |
| CD630_21460 | CD630_n01010 | 125 -- 194 | 639 -- 716 | -12.421 | | 4.83005 | 4.59146 |
| pilC | CD630_n01010 | 7 -- 56 | 388 -- 416 | -12.4135 | | 3.54230 | 4.59146 |
| CD630_05880 | CD630_n01010 | 65 -- 105 | 842 -- 882 | -12.389 | | 4.28339 | 4.59146 |
| CD630_26750 | CD630_n01010 | 84 -- 97 | 973 -- 986 | -12.3863 | | 3.60610 | 4.59146 |
| CD630_20170 | CD630_n01010 | 99 -- 125 | 853 -- 879 | -12.3842 | | 6.90201 | 4.59146 |
| CD630_05900 | CD630_n01010 | 83 -- 99 | 844 -- 861 | -12.361 | | 3.21934 | 4.59146 |
| CD630_14240 | CD630_n01010 | 69 -- 146 | 1344 -- 1415 | -12.3592 | | 8.60766 | 4.59146 |
| panB | CD630_n01010 | 81 -- 171 | 819 -- 921 | -12.3551 | | 3.40248 | 4.59146 |
| CD630_36250 | CD630_n01010 | 79 -- 109 | 839 -- 863 | -12.3366 | | 4.13224 | 4.59146 |
| CD630_19000 | CD630_n01010 | 127 -- 175 | 1104 -- 1136 | -12.3335 | | 4.18443 | 4.59146 |
| CD630_07840 | CD630_n01010 | 142 -- 178 | 62 -- 103 | -12.3115 | | 4.26761 | 4.59146 |
| CD630_02740 | CD630_n01010 | 38 -- 105 | 842 -- 896 | -12.3103 | | 5.02277 | 4.59146 |
| CD630_17511 | CD630_n01010 | 4 -- 30 | 705 -- 731 | -12.2949 | | 9.12322 | 4.59146 |
| CD630_17960 | CD630_n01010 | 157 -- 171 | 794 -- 808 | -12.2838 | | 3.71404 | 4.59146 |
| thiH | CD630_n01010 | 175 -- 199 | 449 -- 471 | -12.2576 | | 3.58230 | 4.59146 |
| CD630_10830 | CD630_n01010 | 139 -- 182 | 38 -- 83 | -12.2357 | | 4.82398 | 4.59146 |
| CD630_05460 | CD630_n01010 | 49 -- 119 | 1103 -- 1167 | -12.2078 | | 3.98753 | 4.59146 |
| CD630_36190 | CD630_n01010 | 78 -- 93 | 1391 -- 1407 | -12.2052 | | 4.63449 | 4.59146 |
| CD630_05821 | CD630_n01010 | 51 -- 71 | 37 -- 61 | -12.2022 | | 3.75661 | 4.59146 |
| CD630_19580 | CD630_n01010 | 60 -- 95 | 839 -- 864 | -12.1933 | | 4.51902 | 4.59146 |
| CD630_17730 | CD630_n01010 | 169 -- 197 | 159 -- 187 | -12.1517 | | 4.73397 | 4.59146 |
| flhB | CD630_n01010 | 9 -- 45 | 289 -- 337 | -12.1421 | | 3.55524 | 4.59146 |
| CD630_36170 | CD630_n01010 | 14 -- 39 | 397 -- 420 | -12.1317 | | 3.72380 | 4.59146 |
| CD630_13620 | CD630_n01010 | 5 -- 64 | 802 -- 854 | -12.1108 | | 4.77627 | 4.59146 |
| cobC | CD630_n01010 | 27 -- 65 | 616 -- 656 | -12.1084 | | 3.15854 | 4.59146 |
| CD630_31530 | CD630_n01010 | 127 -- 148 | 1122 -- 1143 | -12.0709 | | 3.49100 | 4.59146 |
| CD630_25310 | CD630_n01010 | 83 -- 149 | 794 -- 864 | -12.0643 | | 3.99861 | 4.59146 |
| cstA | CD630_n01010 | 64 -- 98 | 845 -- 878 | -12.0514 | | 3.53584 | 4.59146 |
| CD630_18940 | CD630_n01010 | 86 -- 140 | 794 -- 843 | -12.0478 | | 4.09988 | 4.59146 |
| CD630_36781 | CD630_n01010 | 40 -- 189 | 78 -- 218 | -12.0318 | | 3.75946 | 4.59146 |
| fbp | CD630_n01010 | 96 -- 120 | 325 -- 348 | -12.022 | | 3.80071 | 4.59146 |
| CD630_02830 | CD630_n01010 | 2 -- 74 | 37 -- 107 | -12.0183 | | 3.57106 | 4.59146 |
| CD630_08080 | CD630_n01010 | 166 -- 197 | 129 -- 156 | -11.9944 | | 4.87819 | 4.59146 |
| CD630_33610 | CD630_n01010 | 154 -- 200 | 1115 -- 1167 | -11.9842 | | 3.79322 | 4.59146 |
| CD630_06642 | CD630_n01010 | 2 -- 43 | 1033 -- 1061 | -11.97 | | 3.32967 | 4.59146 |
| CD630_31500 | CD630_n01010 | 29 -- 54 | 322 -- 348 | -11.9522 | | 4.90207 | 4.59146 |
| CD630_31010 | CD630_n01010 | 83 -- 114 | 173 -- 202 | -11.941 | | 4.96305 | 4.59146 |
| CD630_10970 | CD630_n01010 | 177 -- 199 | 976 -- 998 | -11.9277 | | 3.53850 | 4.59146 |
| CD630_05870 | CD630_n01010 | 168 -- 199 | 967 -- 1009 | -11.8955 | | 4.28339 | 4.59146 |
| CD630_21030 | CD630_n01010 | 169 -- 200 | 859 -- 892 | -11.8706 | | 3.08733 | 4.59146 |
| cwp26 | CD630_n01010 | 134 -- 158 | 1213 -- 1237 | -11.8637 | | 4.44377 | 4.59146 |
| CD630_06110 | CD630_n01010 | 54 -- 89 | 1127 -- 1160 | -11.86 | | 5.61380 | 4.59146 |
| CD630_17900 | CD630_n01010 | 137 -- 200 | 838 -- 910 | -11.8585 | | 3.52167 | 4.59146 |
| CD630_33682 | CD630_n01010 | 69 -- 146 | 1344 -- 1415 | -11.8568 | | 5.77720 | 4.59146 |
| CD630_33300 | CD630_n01010 | 185 -- 198 | 1124 -- 1137 | -11.8511 | | 3.79149 | 4.59146 |
| CD630_25040 | CD630_n01010 | 141 -- 200 | 38 -- 98 | -11.8504 | | 3.43898 | 4.59146 |
| CD630_13880 | CD630_n01010 | 136 -- 158 | 1113 -- 1132 | -11.8284 | | 3.52735 | 4.59146 |
| CD630_16940 | CD630_n01010 | 17 -- 161 | 313 -- 446 | -11.8098 | | 4.14665 | 4.59146 |
| CD630_28290 | CD630_n01010 | 184 -- 196 | 14 -- 26 | -11.8048 | | 5.54192 | 4.59146 |
| CD630_07470 | CD630_n01010 | 140 -- 180 | 1126 -- 1165 | -11.7961 | | 3.37143 | 4.59146 |
| CD630_14990 | CD630_n01010 | 1 -- 44 | 832 -- 880 | -11.7806 | | 3.78619 | 4.59146 |
| CD630_18100 | CD630_n01010 | 179 -- 195 | 1115 -- 1130 | -11.779 | | 5.71658 | 4.59146 |
| CD630_06540 | CD630_n01010 | 31 -- 120 | 748 -- 871 | -11.7342 | | 3.54673 | 4.59146 |
| dltD | CD630_n01010 | 37 -- 118 | 632 -- 699 | -11.7136 | | 4.40777 | 4.59146 |
| CD630_25300 | CD630_n01010 | 138 -- 152 | 643 -- 657 | -11.7082 | | 4.49333 | 4.59146 |
| CD630_31210 | CD630_n01010 | 39 -- 91 | 850 -- 905 | -11.7056 | | 4.17693 | 4.59146 |
| cwp29 | CD630_n01010 | 27 -- 106 | 328 -- 416 | -11.6945 | | 4.40865 | 4.59146 |
| CD630_31220 | CD630_n01010 | 43 -- 115 | 836 -- 895 | -11.6914 | | 3.73779 | 4.59146 |
| opuCA | CD630_n01010 | 4 -- 97 | 965 -- 1039 | -11.6822 | | 4.07035 | 4.59146 |
| CD630_17360 | CD630_n01010 | 6 -- 37 | 1150 -- 1184 | -11.6817 | | 5.66070 | 4.59146 |
| CD630_30540 | CD630_n01010 | 73 -- 96 | 836 -- 870 | -11.6465 | | 3.63804 | 4.59146 |
| CD630_22030 | CD630_n01010 | 84 -- 122 | 57 -- 104 | -11.6431 | | 3.29934 | 4.59146 |
| CD630_22850 | CD630_n01010 | 82 -- 93 | 1393 -- 1404 | -11.639 | | 3.84102 | 4.59146 |
| CD630_05840 | CD630_n01010 | 106 -- 167 | 628 -- 696 | -11.6337 | | 4.28779 | 4.59146 |
| CD630_21720 | CD630_n01010 | 155 -- 172 | 798 -- 815 | -11.6319 | | 4.13030 | 4.59146 |
| CD630_19840 | CD630_n01010 | 122 -- 187 | 568 -- 644 | -11.6285 | | 3.44724 | 4.59146 |
| CD630_31461 | CD630_n01010 | 17 -- 99 | 834 -- 924 | -11.6229 | | 4.41476 | 4.59146 |
| CD630_33560 | CD630_n01010 | 160 -- 185 | 598 -- 623 | -11.5977 | | 4.03147 | 4.59146 |
| CD630_13600 | CD630_n01010 | 85 -- 97 | 1387 -- 1400 | -11.5929 | | 4.45481 | 4.59146 |
| CD630_12650 | CD630_n01010 | 106 -- 150 | 309 -- 341 | -11.588 | | 6.31827 | 4.59146 |
| CD630_15550 | CD630_n01010 | 49 -- 93 | 839 -- 888 | -11.5665 | | 3.49019 | 4.59146 |
| argG | CD630_n01010 | 59 -- 108 | 839 -- 878 | -11.5633 | | 4.80300 | 4.59146 |
| CD630_05220 | CD630_n01010 | 102 -- 134 | 1389 -- 1419 | -11.5578 | | 3.65057 | 4.59146 |
| CD630_29250 | CD630_n01010 | 70 -- 83 | 1292 -- 1305 | -11.5105 | | 4.61901 | 4.59146 |
| CD630_09410 | CD630_n01010 | 70 -- 83 | 1292 -- 1305 | -11.5105 | | 4.75651 | 4.59146 |
| glnA | CD630_n01010 | 131 -- 192 | 69 -- 119 | -11.5061 | | 3.87629 | 4.59146 |
| CD630_27240 | CD630_n01010 | 3 -- 15 | 54 -- 65 | -11.4774 | | 3.52462 | 4.59146 |
| CD630_34030 | CD630_n01010 | 14 -- 72 | 850 -- 903 | -11.4766 | | 3.05766 | 4.59146 |
| CD630_06190 | CD630_n01010 | 68 -- 95 | 1122 -- 1155 | -11.4504 | | 3.57064 | 4.59146 |
| CD630_30620 | CD630_n01010 | 2 -- 63 | 519 -- 569 | -11.4137 | | 4.15443 | 4.59146 |
| dltB | CD630_n01010 | 136 -- 168 | 66 -- 98 | -11.4127 | | 3.81485 | 4.59146 |
| CD630_06170 | CD630_n01010 | 109 -- 145 | 105 -- 155 | -11.4094 | | 4.95366 | 4.59146 |
| CD630_06100 | CD630_n01010 | 60 -- 164 | 68 -- 164 | -11.407 | | 4.43918 | 4.59146 |
| CD630_21700 | CD630_n01010 | 179 -- 201 | 852 -- 874 | -11.3765 | | 3.96650 | 4.59146 |
| CD630_14810 | CD630_n01010 | 117 -- 157 | 197 -- 239 | -11.3726 | | 3.35323 | 4.59146 |
| CD630_09840 | CD630_n01010 | 150 -- 157 | 976 -- 983 | -11.3305 | | 4.50009 | 4.59146 |
| CD630_32720 | CD630_n01010 | 34 -- 95 | 1122 -- 1169 | -11.3214 | | 4.74276 | 4.59146 |
| CD630_25020 | CD630_n01010 | 127 -- 200 | 51 -- 129 | -11.3172 | | 4.17721 | 4.59146 |
| CD630_08140 | CD630_n01010 | 6 -- 30 | 1215 -- 1236 | -11.3019 | | 3.30159 | 4.59146 |
| CD630_05300 | CD630_n01010 | 172 -- 199 | 1117 -- 1141 | -11.2911 | | 4.56885 | 4.59146 |
| CD630_05000 | CD630_n01010 | 124 -- 152 | 1125 -- 1155 | -11.2883 | | 4.02442 | 4.59146 |
| appF | CD630_n01010 | 56 -- 105 | 1117 -- 1173 | -11.2768 | | 4.25570 | 4.59146 |
| CD630_08780 | CD630_n01010 | 145 -- 174 | 1016 -- 1045 | -11.2546 | | 7.76690 | 4.59146 |
| CD630_21730 | CD630_n01010 | 158 -- 196 | 578 -- 615 | -11.2347 | | 3.45076 | 4.59146 |
| CD630_20650 | CD630_n01010 | 52 -- 129 | 800 -- 882 | -11.2341 | | 4.00083 | 4.59146 |
| CD630_23930 | CD630_n01010 | 73 -- 171 | 383 -- 470 | -11.2023 | | 3.05650 | 4.59146 |
| CD630_14250 | CD630_n01010 | 148 -- 195 | 853 -- 892 | -11.1862 | | 4.19604 | 4.59146 |
| oppB | CD630_n01010 | 97 -- 147 | 98 -- 138 | -11.1585 | | 2.98328 | 4.59146 |
| CD630_25200 | CD630_n01010 | 62 -- 96 | 1388 -- 1421 | -11.1297 | | 3.65158 | 4.59146 |
| CD630_31970 | CD630_n01010 | 118 -- 134 | 792 -- 807 | -11.1296 | | 3.24511 | 4.59146 |
| CD630_08090 | CD630_n01010 | 36 -- 56 | 837 -- 862 | -11.1249 | | 3.57167 | 4.59146 |
| CD630_29270 | CD630_n01010 | 2 -- 116 | 785 -- 875 | -11.1181 | | 3.70174 | 4.59146 |
| CD630_15190 | CD630_n01010 | 9 -- 157 | 253 -- 394 | -11.1165 | | 4.43255 | 4.59146 |
| CD630_34050 | CD630_n01010 | 29 -- 45 | 309 -- 327 | -11.1147 | | 4.03863 | 4.59146 |
| CD630_25430 | CD630_n01010 | 139 -- 163 | 1019 -- 1044 | -11.093 | | 4.00527 | 4.59146 |
| CD630_05820 | CD630_n01010 | 68 -- 123 | 821 -- 874 | -11.0857 | | 3.75661 | 4.59146 |
| CD630_21020 | CD630_n01010 | 85 -- 110 | 1113 -- 1138 | -11.0855 | | 4.28666 | 4.59146 |
| CD630_18960 | CD630_n01010 | 25 -- 51 | 326 -- 349 | -11.0815 | | 4.37822 | 4.59146 |
| CD630_14610 | CD630_n01010 | 146 -- 176 | 1280 -- 1312 | -11.0776 | | 4.89428 | 4.59146 |
| chrA | CD630_n01010 | 148 -- 167 | 1387 -- 1411 | -11.0731 | | 5.27255 | 4.59146 |
| CD630_01700 | CD630_n01010 | 74 -- 95 | 846 -- 865 | -11.0651 | | 4.97234 | 4.59146 |
| CD630_34080 | CD630_n01010 | 47 -- 115 | 1101 -- 1168 | -11.0644 | | 3.20224 | 4.59146 |
| acoL | CD630_n01010 | 84 -- 133 | 1097 -- 1137 | -11.0593 | | 3.32169 | 4.59146 |
| CD630_18660 | CD630_n01010 | 14 -- 29 | 1104 -- 1119 | -11.0326 | | 3.50371 | 4.59146 |
| CD630_13960 | CD630_n01010 | 153 -- 177 | 834 -- 853 | -11.027 | | 3.30375 | 4.59146 |
| CD630_05490 | CD630_n01010 | 114 -- 132 | 5 -- 22 | -11.0209 | | 3.89852 | 4.59146 |
| CD630_27220 | CD630_n01010 | 1 -- 16 | 330 -- 345 | -11.0162 | | 4.18786 | 4.59146 |
| CD630_33420 | CD630_n01010 | 6 -- 36 | 833 -- 863 | -11.0111 | | 3.22095 | 4.59146 |
| CD630_21240 | CD630_n01010 | 88 -- 94 | 976 -- 982 | -10.9742 | | 5.88879 | 4.59146 |
| cobS | CD630_n01010 | 86 -- 100 | 46 -- 61 | -10.9557 | | 3.67481 | 4.59146 |
| CD630_26150 | CD630_n01010 | 159 -- 195 | 1303 -- 1334 | -10.9462 | | 4.06302 | 4.59146 |
| CD630_09990 | CD630_n01010 | 19 -- 47 | 1104 -- 1133 | -10.9409 | | 3.86040 | 4.59146 |
| CD630_32890 | CD630_n01010 | 60 -- 85 | 525 -- 552 | -10.9383 | | 3.41276 | 4.59146 |
| CD630_15080 | CD630_n01010 | 80 -- 106 | 841 -- 868 | -10.9178 | | 4.03033 | 4.59146 |
| CD630_19921 | CD630_n01010 | 62 -- 98 | 322 -- 358 | -10.9165 | | 5.46345 | 4.59146 |
| CD630_03420 | CD630_n01010 | 106 -- 140 | 1387 -- 1419 | -10.9159 | | 4.71868 | 4.59146 |
| CD630_04880 | CD630_n01010 | 89 -- 111 | 832 -- 854 | -10.9141 | | 4.62291 | 4.59146 |
| CD630_05030 | CD630_n01010 | 136 -- 168 | 52 -- 85 | -10.9115 | | 4.03823 | 4.59146 |
| CD630_14920 | CD630_n01010 | 74 -- 107 | 976 -- 1009 | -10.905 | | 4.35225 | 4.59146 |
| CD630_18090 | CD630_n01010 | 109 -- 199 | 1097 -- 1190 | -10.8951 | | 4.23961 | 4.59146 |
| CD630_06560 | CD630_n01010 | 39 -- 178 | 751 -- 878 | -10.879 | | 4.02532 | 4.59146 |
| uxaA | CD630_n01010 | 186 -- 199 | 667 -- 680 | -10.8781 | | 4.26165 | 4.59146 |
| CD630_26050 | CD630_n01010 | 57 -- 78 | 859 -- 879 | -10.8767 | | 6.99851 | 4.59146 |
| CD630_13560 | CD630_n01010 | 5 -- 23 | 1127 -- 1152 | -10.8369 | | 3.38083 | 4.59146 |
| CD630_15461 | CD630_n01010 | 89 -- 115 | 833 -- 854 | -10.8061 | | 9.13442 | 4.59146 |
| CD630_27120 | CD630_n01010 | 5 -- 44 | 318 -- 356 | -10.802 | | 4.43720 | 4.59146 |
| CD630_09060 | CD630_n01010 | 12 -- 58 | 1328 -- 1366 | -10.7954 | | 4.31834 | 4.59146 |
| ogt1 | CD630_n01010 | 63 -- 95 | 1391 -- 1417 | -10.7817 | | 3.16772 | 4.59146 |
| rolB | CD630_n01010 | 141 -- 201 | 840 -- 916 | -10.7732 | | 3.60270 | 4.59146 |
| CD630_23080 | CD630_n01010 | 157 -- 190 | 840 -- 875 | -10.7469 | | 5.23027 | 4.59146 |
| CD630_14150 | CD630_n01010 | 144 -- 198 | 1095 -- 1133 | -10.7323 | | 3.28251 | 4.59146 |
| CD630_16450 | CD630_n01010 | 86 -- 109 | 838 -- 861 | -10.7319 | | 4.57547 | 4.59146 |
| CD630_01640 | CD630_n01010 | 164 -- 192 | 319 -- 346 | -10.7188 | | 4.55381 | 4.59146 |
| CD630_14110 | CD630_n01010 | 2 -- 29 | 886 -- 913 | -10.7064 | | 3.39550 | 4.59146 |
| cynT | CD630_n01010 | 113 -- 162 | 834 -- 874 | -10.6931 | | 4.21370 | 4.59146 |
| CD630_06920 | CD630_n01010 | 53 -- 139 | 46 -- 136 | -10.6886 | | 3.13260 | 4.59146 |
| CD630_23861 | CD630_n01010 | 154 -- 167 | 1390 -- 1404 | -10.6879 | | 5.83068 | 4.59146 |
| CD630_03930 | CD630_n01010 | 110 -- 184 | 1259 -- 1336 | -10.6823 | | 4.15934 | 4.59146 |
| CD630_27200 | CD630_n01010 | 1 -- 15 | 1387 -- 1408 | -10.6785 | | 3.42516 | 4.59146 |
| CD630_04280 | CD630_n01010 | 103 -- 124 | 388 -- 409 | -10.6749 | | 3.61457 | 4.59146 |
| CD630_04610 | CD630_n01010 | 145 -- 190 | 1315 -- 1367 | -10.6681 | | 3.83025 | 4.59146 |

| | | | | | |
|---|---|---|---|---|---|
| CD630_13780 | CD630_n01010 | 152 -- 186 | 182 -- 216 | -10.6674 | 3.52274 | 4.59146 |
| CD630_29500 | CD630_n01010 | 115 -- 155 | 378 -- 421 | -10.6636 | 3.70644 | 4.59146 |
| CD630_28330 | CD630_n01010 | 178 -- 198 | 186 -- 208 | -10.6622 | 3.29122 | 4.59146 |
| CD630_15150 | CD630_n01010 | 75 -- 104 | 182 -- 207 | -10.6554 | 3.35564 | 4.59146 |
| garK | CD630_n01010 | 152 -- 201 | 1101 -- 1138 | -10.6532 | 3.72836 | 4.59146 |
| CD630_05290 | CD630_n01010 | 59 -- 114 | 309 -- 349 | -10.647 | 3.97604 | 4.59146 |
| CD630_23000 | CD630_n01010 | 94 -- 113 | 834 -- 852 | -10.632 | 3.54130 | 4.59146 |
| CD630_18311 | CD630_n01010 | 85 -- 113 | 836 -- 857 | -10.6141 | 6.86047 | 4.59146 |
| CD630_18410 | CD630_n01010 | 31 -- 110 | 38 -- 101 | -10.6018 | 3.59739 | 4.59146 |
| CD630_19690 | CD630_n01010 | 38 -- 126 | 60 -- 153 | -10.5977 | 3.85861 | 4.59146 |
| CD630_16640 | CD630_n01010 | 90 -- 111 | 1110 -- 1130 | -10.5852 | 3.61548 | 4.59146 |
| CD630_19800 | CD630_n01010 | 21 -- 48 | 312 -- 341 | -10.5847 | 5.22035 | 4.59146 |
| CD630_21090 | CD630_n01010 | 124 -- 142 | 1345 -- 1362 | -10.5772 | 4.15803 | 4.59146 |
| CD630_22740 | CD630_n01010 | 55 -- 194 | 277 -- 421 | -10.5734 | 3.08733 | 4.59146 |
| CD630_10080 | CD630_n01010 | 37 -- 165 | 596 -- 698 | -10.5712 | 3.75625 | 4.59146 |
| CD630_26050 | CD630_n01010 | 77 -- 152 | 1060 -- 1140 | -10.5696 | 6.99851 | 4.59146 |
| CD630_15431 | CD630_n01010 | 170 -- 201 | 843 -- 875 | -10.5671 | 5.29780 | 4.59146 |
| CD630_22880 | CD630_n01010 | 74 -- 117 | 839 -- 879 | -10.5543 | 3.62455 | 4.59146 |
| CD630_32040 | CD630_n01010 | 15 -- 32 | 1303 -- 1320 | -10.5527 | 4.93516 | 4.59146 |
| CD630_21770 | CD630_n01010 | 5 -- 105 | 842 -- 941 | -10.5435 | 4.02587 | 4.59146 |
| CD630_07400 | CD630_n01010 | 148 -- 187 | 625 -- 654 | -10.5219 | 4.52273 | 4.59146 |
| CD630_27210 | CD630_n01010 | 30 -- 119 | 101 -- 198 | -10.487 | 4.18786 | 4.59146 |
| CD630_20980 | CD630_n01010 | 126 -- 133 | 54 -- 61 | -10.4759 | 8.12847 | 4.59146 |
| CD630_07870 | CD630_n01010 | 182 -- 196 | 1056 -- 1071 | -10.4303 | 3.22829 | 4.59146 |
| CD630_27060 | CD630_n01010 | 94 -- 159 | 800 -- 852 | -10.4197 | 4.06049 | 4.59146 |
| CD630_25150 | CD630_n01010 | 67 -- 142 | 604 -- 683 | -10.4132 | 3.79339 | 4.59146 |
| bltD | CD630_n01010 | 136 -- 154 | 1401 -- 1420 | -10.3968 | 4.49926 | 4.59146 |
| CD630_16530 | CD630_n01010 | 157 -- 167 | 1387 -- 1396 | -10.3894 | 4.37658 | 4.59146 |
| CD630_01560 | CD630_n01010 | 148 -- 165 | 1390 -- 1407 | -10.3782 | 4.06352 | 4.59146 |
| CD630_26250 | CD630_n01010 | 161 -- 187 | 132 -- 163 | -10.3724 | 5.03379 | 4.59146 |
| CD630_23440 | CD630_n01010 | 182 -- 199 | 1111 -- 1128 | -10.3528 | 3.34841 | 4.59146 |
| CD630_19030 | CD630_n01010 | 1 -- 40 | 935 -- 971 | -10.3251 | 3.34240 | 4.59146 |
| CD630_27230 | CD630_n01010 | 5 -- 43 | 398 -- 451 | -10.3148 | 3.52462 | 4.59146 |
| opuCC | CD630_n01010 | 178 -- 187 | 1 -- 10 | -10.3017 | 4.07035 | 4.59146 |
| CD630_19801 | CD630_n01010 | 160 -- 189 | 1300 -- 1338 | -10.2875 | 5.22035 | 4.59146 |
| CD630_13930 | CD630_n01010 | 4 -- 30 | 398 -- 421 | -10.2775 | 4.09246 | 4.59146 |
| CD630_33190 | CD630_n01010 | 56 -- 120 | 836 -- 895 | -10.241 | 4.13899 | 4.59146 |
| CD630_05280 | CD630_n01010 | 35 -- 50 | 57 -- 73 | -10.216 | 3.91665 | 4.59146 |
| CD630_26020 | CD630_n01010 | 93 -- 100 | 974 -- 981 | -10.2111 | 4.03155 | 4.59146 |
| CD630_05940 | CD630_n01010 | 87 -- 93 | 975 -- 981 | -10.2106 | 4.18595 | 4.59146 |
| CD630_08110 | CD630_n01010 | 81 -- 162 | 1334 -- 1406 | -10.1872 | 4.39181 | 4.59146 |
| kdgT1 | CD630_n01010 | 110 -- 126 | 326 -- 342 | -10.1769 | 4.49207 | 4.59146 |
| CD630_12110 | CD630_n01010 | 141 -- 177 | 236 -- 275 | -10.1715 | 4.36078 | 4.59146 |
| CD630_22550 | CD630_n01010 | 180 -- 201 | 889 -- 909 | -10.1502 | 3.62379 | 4.59146 |
| CD630_03170 | CD630_n01010 | 102 -- 175 | 1291 -- 1378 | -10.143 | 3.66946 | 4.59146 |
| CD630_28890 | CD630_n01010 | 25 -- 54 | 452 -- 479 | -10.1418 | 7.52469 | 4.59146 |
| CD630_13580 | CD630_n01010 | 103 -- 147 | 1035 -- 1077 | -10.1357 | 3.80715 | 4.59146 |
| CD630_10960 | CD630_n01010 | 17 -- 30 | 204 -- 217 | -10.1015 | 3.53850 | 4.59146 |
| tcdA | CD630_n01010 | 40 -- 64 | 852 -- 875 | -10.0812 | 4.70593 | 4.59146 |
| CD630_36180 | CD630_n01010 | 91 -- 116 | 309 -- 331 | -10.0748 | 4.83428 | 4.59146 |
| CD630_30110 | CD630_n01010 | 88 -- 114 | 389 -- 415 | -10.0708 | 3.31041 | 4.59146 |
| CD630_21330 | CD630_n01010 | 86 -- 106 | 1119 -- 1140 | -10.0706 | 4.10111 | 4.59146 |
| CD630_16160 | CD630_n01010 | 42 -- 109 | 838 -- 907 | -10.0649 | 6.11801 | 4.59146 |
| CD630_05370 | CD630_n01010 | 168 -- 200 | 647 -- 680 | -10.0545 | 3.26947 | 4.59146 |
| CD630_16750 | CD630_n01010 | 135 -- 154 | 1121 -- 1140 | -10.0473 | 5.04051 | 4.59146 |
| CD630_20960 | CD630_n01010 | 69 -- 107 | 312 -- 350 | -10.0444 | 5.65940 | 4.59146 |
| CD630_20541 | CD630_n01010 | 79 -- 97 | 1110 -- 1129 | -10.0306 | 5.74527 | 4.59146 |
| CD630_03160 | CD630_n01010 | 44 -- 74 | 557 -- 591 | -10.0265 | 3.66946 | 4.59146 |
| CD630_21260 | CD630_n01010 | 55 -- 97 | 981 -- 1014 | -10.0263 | 4.22255 | 4.59146 |
| CD630_14010 | CD630_n01010 | 191 -- 201 | 1340 -- 1350 | -10.0161 | 3.69983 | 4.59146 |
| cdu1 | CD630_n01010 | 129 -- 155 | 269 -- 296 | -10.0044 | 4.46276 | 4.59146 |
| CD630_21790 | CD630_n01010 | 137 -- 164 | 1387 -- 1418 | -9.99059 | 4.93762 | 4.59146 |
| CD630_15160 | CD630_n01010 | 178 -- 189 | 133 -- 144 | -9.9774 | 4.32318 | 4.59146 |
| cbiQ | CD630_n01010 | 113 -- 178 | 1114 -- 1166 | -9.97678 | 3.79830 | 4.59146 |
| CD630_07960 | CD630_n01010 | 177 -- 198 | 1084 -- 1111 | -9.94413 | 3.22537 | 4.59146 |
| CD630_13360 | CD630_n01010 | 87 -- 105 | 842 -- 861 | -9.94124 | 3.77330 | 4.59146 |
| CD630_06200 | CD630_n01010 | 91 -- 153 | 39 -- 104 | -9.9206 | 3.57064 | 4.59146 |
| CD630_13790 | CD630_n01010 | 170 -- 194 | 883 -- 910 | -9.91179 | 5.17374 | 4.59146 |
| CD630_27360 | CD630_n01010 | 58 -- 186 | 1002 -- 1145 | -9.90755 | 3.79570 | 4.59146 |
| CD630_10950 | CD630_n01010 | 135 -- 162 | 252 -- 284 | -9.89783 | 3.67675 | 4.59146 |
| CD630_27600 | CD630_n01010 | 4 -- 33 | 1020 -- 1047 | -9.88077 | 3.48384 | 4.59146 |
| CD630_23760 | CD630_n01010 | 185 -- 199 | 1374 -- 1388 | -9.87269 | 4.54307 | 4.59146 |
| CD630_17380 | CD630_n01010 | 115 -- 130 | 1395 -- 1412 | -9.85409 | 5.03207 | 4.59146 |
| cbiO | CD630_n01010 | 93 -- 99 | 975 -- 981 | -9.84895 | 3.79830 | 4.59146 |
| CD630_31490 | CD630_n01010 | 123 -- 135 | 141 -- 153 | -9.84794 | 4.08593 | 4.59146 |
| CD630_23820 | CD630_n01010 | 88 -- 104 | 834 -- 855 | -9.82329 | 3.18419 | 4.59146 |
| CD630_14050 | CD630_n01010 | 70 -- 104 | 834 -- 870 | -9.82182 | 3.30894 | 4.59146 |
| CD630_29650 | CD630_n01010 | 163 -- 176 | 384 -- 397 | -9.7924 | 3.67128 | 4.59146 |
| CD630_20260 | CD630_n01010 | 7 -- 34 | 538 -- 564 | -9.75992 | 3.84162 | 4.59146 |
| CD630_14930 | CD630_n01010 | 1 -- 25 | 361 -- 385 | -9.75792 | 3.31335 | 4.59146 |
| CD630_24770 | CD630_n01010 | 57 -- 100 | 972 -- 1009 | -9.75511 | 3.20798 | 4.59146 |
| CD630_18920 | CD630_n01010 | 152 -- 189 | 976 -- 1009 | -9.75422 | 4.43558 | 4.59146 |
| ppnK | CD630_n01010 | 81 -- 96 | 981 -- 994 | -9.74886 | 4.36817 | 4.59146 |
| CD630_06380 | CD630_n01010 | 134 -- 189 | 1346 -- 1406 | -9.72951 | 4.32211 | 4.59146 |
| CD630_22720 | CD630_n01010 | 126 -- 161 | 840 -- 875 | -9.72597 | 3.81804 | 4.59146 |
| CD630_26680 | CD630_n01010 | 21 -- 34 | 981 -- 994 | -9.71316 | 3.53956 | 4.59146 |
| CD630_09100 | CD630_n01010 | 35 -- 63 | 896 -- 922 | -9.67689 | 5.92260 | 4.59146 |
| CD630_06690 | CD630_n01010 | 89 -- 106 | 701 -- 716 | -9.66944 | 5.82709 | 4.59146 |
| eutP | CD630_n01010 | 100 -- 111 | 836 -- 847 | -9.66026 | 3.82847 | 4.59146 |
| CD630_04660 | CD630_n01010 | 99 -- 171 | 61 -- 128 | -9.65939 | 4.32791 | 4.59146 |
| CD630_08070 | CD630_n01010 | 89 -- 105 | 389 -- 405 | -9.6395 | 4.87819 | 4.59146 |
| CD630_31690 | CD630_n01010 | 156 -- 195 | 123 -- 165 | -9.62215 | 3.71193 | 4.59146 |
| CD630_21230 | CD630_n01010 | 84 -- 105 | 842 -- 864 | -9.61846 | 3.89213 | 4.59146 |
| CD630_21610 | CD630_n01010 | 52 -- 95 | 848 -- 879 | -9.61524 | 3.16662 | 4.59146 |
| CD630_09570 | CD630_n01010 | 24 -- 38 | 391 -- 405 | -9.60523 | 5.31213 | 4.59146 |
| CD630_29070 | CD630_n01010 | 24 -- 38 | 391 -- 405 | -9.60523 | 3.89477 | 4.59146 |
| CD630_36690 | CD630_n01010 | 120 -- 165 | 1274 -- 1305 | -9.58175 | 3.53761 | 4.59146 |
| ldh | CD630_n01010 | 118 -- 144 | 57 -- 86 | -9.57416 | 4.35818 | 4.59146 |
| CD630_14140 | CD630_n01010 | 118 -- 144 | 29 -- 52 | -9.57331 | 3.16551 | 4.59146 |
| CD630_07070 | CD630_n01010 | 11 -- 27 | 1216 -- 1233 | -9.55623 | 3.46414 | 4.59146 |
| CD630_19440 | CD630_n01010 | 107 -- 133 | 1104 -- 1138 | -9.5491 | 4.53937 | 4.59146 |
| CD630_21110 | CD630_n01010 | 90 -- 106 | 788 -- 804 | -9.50095 | 3.89630 | 4.59146 |
| CD630_25200 | CD630_RNA_7 | 158 -- 196 | 48 -- 95 | -12.6231 | 3.65158 | 7.42886 |
| CD630_32040 | CD630_RNA_7 | 19 -- 32 | 94 -- 107 | -12.3139 | 4.93516 | 7.42886 |
| CD630_02830 | CD630_RNA_7 | 81 -- 126 | 84 -- 113 | -12.1088 | 3.57106 | 7.42886 |
| CD630_29760 | CD630_RNA_7 | 63 -- 96 | 83 -- 113 | -11.6247 | 3.51074 | 7.42886 |
| CD630_18930 | CD630_RNA_7 | 9 -- 18 | 50 -- 59 | -11.1218 | 5.32127 | 7.42886 |
| CD630_09760 | CD630_RNA_7 | 78 -- 108 | 82 -- 113 | -10.9484 | 4.83410 | 7.42886 |
| CD630_28910 | CD630_RNA_7 | 78 -- 108 | 82 -- 113 | -10.9484 | 4.51664 | 7.42886 |
| mobA | CD630_RNA_7 | 138 -- 149 | 100 -- 111 | -10.9447 | 4.25703 | 7.42886 |
| cdtR | CD630_RNA_7 | 179 -- 188 | 95 -- 104 | -10.9337 | 5.84368 | 7.42886 |
| virS | CD630_RNA_7 | 3 -- 31 | 23 -- 52 | -10.4515 | 4.74085 | 7.42886 |
| CD630_07070 | CD630_RNA_7 | 31 -- 45 | 50 -- 63 | -10.3943 | 3.46414 | 7.42886 |
| CD630_06690 | CD630_RNA_7 | 25 -- 33 | 95 -- 103 | -10.3261 | 5.82709 | 7.42886 |
| CD630_27520 | CD630_RNA_7 | 85 -- 110 | 84 -- 105 | -10.2358 | 4.27693 | 7.42886 |
| CD630_09100 | CD630_RNA_7 | 2 -- 89 | 32 -- 105 | -10.1049 | 5.92260 | 7.42886 |
| CD630_04880 | CD630_RNA_7 | 129 -- 147 | 95 -- 113 | -9.84526 | 4.62291 | 7.42886 |
| CD630_16450 | CD630_RNA_7 | 102 -- 114 | 94 -- 105 | -9.83473 | 4.57547 | 7.42886 |
| CD630_23780 | CD630_RNA_7 | 164 -- 187 | 39 -- 62 | -9.80031 | 3.91629 | 7.42886 |
| appF | CD630_RNA_7 | 131 -- 139 | 96 -- 104 | -9.75033 | 4.25570 | 7.42886 |
| CD630_32520 | CD630_RNA_7 | 87 -- 96 | 96 -- 105 | -9.748 | 4.06098 | 7.42886 |
| CD630_13880 | CD630_RNA_7 | 23 -- 41 | 92 -- 107 | -9.72233 | 3.52735 | 7.42886 |
| CD630_06540 | CD630_RNA_7 | 168 -- 183 | 80 -- 97 | -9.68915 | 3.54673 | 7.42886 |
| CD630_08370 | CD630_RNA_7 | 186 -- 194 | 16 -- 24 | -9.67139 | 3.65639 | 7.42886 |
| bglA | CD630_RNA_7 | 153 -- 186 | 83 -- 106 | -9.64489 | 4.86402 | 7.42886 |
| CD630_05280 | CD630_RNA_7 | 174 -- 183 | 83 -- 92 | -9.64079 | 3.91665 | 7.42886 |
| CD630_26920 | CD630_RNA_7 | 85 -- 98 | 96 -- 108 | -9.62358 | 4.58060 | 7.42886 |
| CD630_17770 | CD630_RNA_7 | 131 -- 139 | 51 -- 59 | -9.50699 | 5.06490 | 7.42886 |
| CD630_05230 | CD630_s0330 | 69 -- 143 | 2 -- 94 | -19.088 | 3.73787 | 4.37658 |
| CD630_14130 | CD630_s0330 | 140 -- 191 | 1 -- 44 | -14.0021 | 4.57970 | 4.37658 |
| CD630_28181 | CD630_s0330 | 153 -- 189 | 2 -- 33 | -13.1949 | 3.91299 | 4.37658 |
| CD630_19850 | CD630_s0330 | 106 -- 118 | 75 -- 87 | -12.7362 | 6.88486 | 4.37658 |
| CD630_05300 | CD630_s0330 | 170 -- 199 | 7 -- 37 | -12.7203 | 4.56885 | 4.37658 |
| CD630_06110 | CD630_s0330 | 175 -- 189 | 26 -- 40 | -12.5562 | 5.61380 | 4.37658 |
| CD630_14182 | CD630_s0330 | 189 -- 198 | 8 -- 17 | -12.3966 | 4.82360 | 4.37658 |
| CD630_14250 | CD630_s0330 | 151 -- 169 | 27 -- 45 | -11.7494 | 4.19604 | 4.37658 |
| CD630_33600 | CD630_s0330 | 51 -- 66 | 76 -- 91 | -11.322 | 3.73196 | 4.37658 |
| CD630_28560 | CD630_s0330 | 152 -- 162 | 30 -- 40 | -11.2161 | 3.96803 | 4.37658 |
| CD630_28550 | CD630_s0330 | 190 -- 200 | 41 -- 51 | -10.9085 | 3.51196 | 4.37658 |
| CD630_36190 | CD630_s0330 | 79 -- 112 | 7 -- 39 | -10.8586 | 4.63449 | 4.37658 |
| CD630_20650 | CD630_s0330 | 7 -- 17 | 2 -- 12 | -10.7886 | 4.00083 | 4.37658 |
| CD630_09771 | CD630_s0330 | 180 -- 189 | 8 -- 17 | -10.6055 | 4.95362 | 4.37658 |
| CD630_31010 | CD630_s0330 | 183 -- 191 | 25 -- 33 | -10.4947 | 4.96305 | 4.37658 |
| spaK | CD630_s0330 | 150 -- 178 | 9 -- 36 | -10.258 | 3.04530 | 4.37658 |
| CD630_25300 | CD630_s0330 | 134 -- 155 | 4 -- 25 | -10.1722 | 4.49333 | 4.37658 |
| CD630_28890 | CD630_s0330 | 180 -- 189 | 8 -- 17 | -10.1654 | 7.52469 | 4.37658 |
| CD630_16450 | CD630_s0330 | 131 -- 161 | 3 -- 42 | -10.1158 | 4.57547 | 4.37658 |
| CD630_05870 | CD630_s0330 | 168 -- 181 | 78 -- 91 | -10.0519 | 4.28339 | 4.37658 |
| flgB | CD630_s0330 | 185 -- 197 | 77 -- 89 | -10.0234 | 3.25736 | 4.37658 |
| CD630_33580 | CD630_s0330 | 104 -- 114 | 4 -- 14 | -9.94153 | 3.95778 | 4.37658 |
| CD630_06060 | CD630_s0330 | 137 -- 151 | 26 -- 40 | -9.93531 | 4.94050 | 4.37658 |
| CD630_30100 | CD630_s0330 | 137 -- 161 | 13 -- 40 | -9.93002 | 4.13234 | 4.37658 |
| CD630_04850 | CD630_s0330 | 147 -- 171 | 10 -- 36 | -9.88221 | 4.40788 | 4.37658 |
| dltA | CD630_s0330 | 148 -- 170 | 53 -- 74 | -9.85996 | 3.85429 | 4.37658 |
| CD630_17160 | CD630_s0330 | 65 -- 95 | 2 -- 30 | -9.77206 | 3.50228 | 4.37658 |
| CD630_19780 | CD630_s0330 | 8 -- 21 | 1 -- 18 | -9.73546 | 3.64830 | 4.37658 |
| CD630_06380 | CD630_s0330 | 183 -- 194 | 78 -- 88 | -9.70599 | 4.32211 | 4.37658 |
| CD630_35260 | CD630_s0330 | 66 -- 100 | 4 -- 36 | -9.67581 | 5.77277 | 4.37658 |
| CD630_18910 | CD630_s0360 | 28 -- 53 | 2 -- 23 | -15.0362 | 3.30688 | 6.65922 |
| CD630_18781 | CD630_s0360 | 138 -- 159 | 101 -- 122 | -12.2843 | 3.77660 | 6.65922 |
| CD630_03400 | CD630_s0360 | 169 -- 188 | 100 -- 118 | -11.6926 | 4.45994 | 6.65922 |
| CD630_01590 | CD630_s0360 | 12 -- 35 | 94 -- 117 | -11.6199 | 3.45609 | 6.65922 |
| CD630_27280 | CD630_s0360 | 191 -- 199 | 108 -- 116 | -11.0088 | 3.24738 | 6.65922 |
| gcvTPA | CD630_s0360 | 168 -- 181 | 25 -- 38 | -10.704 | 4.77229 | 6.65922 |
| CD630_22991 | CD630_s0360 | 191 -- 201 | 11 -- 21 | -10.6072 | 6.03728 | 6.65922 |
| CD630_08140 | CD630_s0360 | 1 -- 15 | 10 -- 23 | -10.4891 | 3.30159 | 6.65922 |
| CD630_13790 | CD630_s0360 | 1 -- 13 | 97 -- 109 | -10.0494 | 5.17374 | 6.65922 |
| CD630_14150 | CD630_s0360 | 28 -- 49 | 96 -- 118 | -9.75455 | 3.28251 | 6.65922 |
| CD630_23450 | CD630_s0360 | 181 -- 194 | 104 -- 115 | -9.6991 | 3.27218 | 6.65922 |
| CD630_08460 | CD630_s0360 | 134 -- 160 | 1 -- 29 | -9.66026 | 4.15083 | 6.65922 |
| CD630_02430 | CD630_s0360 | 27 -- 48 | 6 -- 29 | -9.59237 | 3.71048 | 6.65922 |
| CD630_08080 | CD630_s0360 | 6 -- 28 | 100 -- 119 | -9.5387 | 4.87819 | 6.65922 |
| dnaD | CD630_s0450 | 104 -- 162 | 28 -- 90 | -17.8068 | 3.56078 | 5.67265 |
| CD630_09770 | CD630_s0450 | 160 -- 201 | 52 -- 93 | -17.037 | 3.22106 | 5.67265 |
| CD630_27280 | CD630_s0450 | 132 -- 172 | 3 -- 44 | -15.6173 | 3.24738 | 5.67265 |
| flhB | CD630_s0450 | 151 -- 193 | 5 -- 46 | -15.1524 | 3.55524 | 5.67265 |
| CD630_03930 | CD630_s0450 | 136 -- 153 | 99 -- 115 | -14.1866 | 4.15934 | 5.67265 |
| rbsC | CD630_s0450 | 168 -- 192 | 180 -- 209 | -14.067 | 3.71650 | 5.67265 |
| agrB | CD630_s0450 | 3 -- 34 | 15 -- 46 | -14.041 | 3.92598 | 5.67265 |
| CD630_21090 | CD630_s0450 | 147 -- 175 | 24 -- 48 | -13.5049 | 4.15803 | 5.67265 |
| CD630_26250 | CD630_s0450 | 131 -- 179 | 45 -- 89 | -13.4168 | 5.03379 | 5.67265 |
| CD630_08370 | CD630_s0450 | 188 -- 198 | 4 -- 14 | -13.3309 | 3.65639 | 5.67265 |
| cdtR | CD630_s0450 | 2 -- 49 | 5 -- 46 | -13.2727 | 5.84368 | 5.67265 |
| CD630_26150 | CD630_s0450 | 20 -- 41 | 65 -- 90 | -13.0529 | 4.06302 | 5.67265 |
| CD630_06170 | CD630_s0450 | 190 -- 201 | 177 -- 188 | -12.6367 | 4.95366 | 5.67265 |
| CD630_23070 | CD630_s0450 | 154 -- 190 | 4 -- 41 | -12.2534 | 5.23027 | 5.67265 |
| CD630_05490 | CD630_s0450 | 66 -- 82 | 135 -- 151 | -12.2386 | 3.89852 | 5.67265 |
| CD630_09850 | CD630_s0450 | 163 -- 180 | 172 -- 188 | -12.0908 | 4.18679 | 5.67265 |
| CD630_17360 | CD630_s0450 | 177 -- 192 | 10 -- 26 | -11.9368 | 5.66070 | 5.67265 |
| CD630_06100 | CD630_s0450 | 108 -- 124 | 135 -- 153 | -11.9304 | 4.43918 | 5.67265 |
| CD630_06150 | CD630_s0450 | 137 -- 156 | 27 -- 46 | -11.8601 | 5.51744 | 5.67265 |
| CD630_04880 | CD630_s0450 | 118 -- 160 | 22 -- 64 | -11.843 | 4.62291 | 5.67265 |
| CD630_36781 | CD630_s0450 | 162 -- 187 | 128 -- 153 | -11.8008 | 3.75946 | 5.67265 |
| CD630_25300 | CD630_s0450 | 87 -- 114 | 56 -- 83 | -11.7203 | 4.49333 | 5.67265 |
| CD630_21140 | CD630_s0450 | 14 -- 36 | 61 -- 82 | -11.7022 | 3.33304 | 5.67265 |
| CD630_21910 | CD630_s0450 | 112 -- 137 | 135 -- 160 | -11.6637 | 4.66600 | 5.67265 |
| CD630_04850 | CD630_s0450 | 152 -- 171 | 169 -- 187 | -11.6503 | 4.40788 | 5.67265 |
| CD630_03650 | CD630_s0450 | 151 -- 181 | 5 -- 37 | -11.6423 | 3.93697 | 5.67265 |
| CD630_13530 | CD630_s0450 | 11 -- 27 | 134 -- 150 | -11.5733 | 4.09832 | 5.67265 |
| CD630_14610 | CD630_s0450 | 129 -- 164 | 5 -- 44 | -11.5509 | 4.89428 | 5.67265 |
| CD630_23861 | CD630_s0450 | 8 -- 37 | 170 -- 192 | -11.5148 | 5.83068 | 5.67265 |
| CD630_21020 | CD630_s0450 | 85 -- 161 | 140 -- 206 | -11.4801 | 4.28666 | 5.67265 |
| tcdA | CD630_s0450 | 149 -- 170 | 187 -- 205 | -11.4475 | 4.70593 | 5.67265 |
| CD630_14240 | CD630_s0450 | 105 -- 146 | 170 -- 209 | -11.4365 | 8.60766 | 5.67265 |
| CD630_05150 | CD630_s0450 | 181 -- 200 | 31 -- 49 | -11.4293 | 3.69875 | 5.67265 |
| CD630_13020 | CD630_s0450 | 118 -- 133 | 78 -- 93 | -11.3861 | 3.52267 | 5.67265 |
| CD630_21080 | CD630_s0450 | 147 -- 194 | 67 -- 116 | -11.2402 | 3.99742 | 5.67265 |
| CD630_27230 | CD630_s0450 | 7 -- 26 | 171 -- 189 | -11.182 | 3.52462 | 5.67265 |
| CD630_22200 | CD630_s0450 | 74 -- 108 | 182 -- 209 | -11.1816 | 4.18288 | 5.67265 |
| CD630_13580 | CD630_s0450 | 136 -- 167 | 13 -- 43 | -11.1714 | 3.80715 | 5.67265 |
| CD630_22950 | CD630_s0450 | 5 -- 29 | 24 -- 45 | -11.1288 | 3.96883 | 5.67265 |
| CD630_23450 | CD630_s0450 | 8 -- 53 | 6 -- 46 | -11.0032 | 3.27218 | 5.67265 |
| cwp29 | CD630_s0450 | 125 -- 169 | 34 -- 86 | -10.9602 | 4.40865 | 5.67265 |
| CD630_05830 | CD630_s0450 | 146 -- 162 | 26 -- 41 | -10.9208 | 3.35360 | 5.67265 |
| CD630_08660 | CD630_s0450 | 48 -- 62 | 100 -- 114 | -10.8751 | 3.86467 | 5.67265 |
| CD630_02720 | CD630_s0450 | 163 -- 176 | 28 -- 40 | -10.8643 | 3.38938 | 5.67265 |
| CD630_33420 | CD630_s0450 | 139 -- 171 | 10 -- 44 | -10.6585 | 3.22095 | 5.67265 |
| CD630_27600 | CD630_s0450 | 173 -- 201 | 129 -- 158 | -10.642 | 3.48384 | 5.67265 |
| CD630_21070 | CD630_s0450 | 45 -- 60 | 22 -- 40 | -10.578 | 5.35019 | 5.67265 |
| CD630_22991 | CD630_s0450 | 189 -- 197 | 168 -- 176 | -10.5285 | 6.03728 | 5.67265 |
| CD630_16160 | CD630_s0450 | 166 -- 197 | 7 -- 47 | -10.513 | 6.11801 | 5.67265 |
| CD630_16330 | CD630_s0450 | 160 -- 183 | 168 -- 194 | -10.3914 | 4.11180 | 5.67265 |
| chrA | CD630_s0450 | 176 -- 191 | 35 -- 48 | -10.3779 | 5.27255 | 5.67265 |
| CD630_30420 | CD630_s0450 | 34 -- 72 | 4 -- 40 | -10.3765 | 5.27539 | 5.67265 |
| opuCC | CD630_s0450 | 81 -- 137 | 102 -- 146 | -10.3725 | 4.07035 | 5.67265 |
| CD630_05030 | CD630_s0450 | 132 -- 161 | 66 -- 88 | -10.2539 | 4.03823 | 5.67265 |
| CD630_17720 | CD630_s0450 | 2 -- 18 | 27 -- 43 | -10.166 | 3.22910 | 5.67265 |
| spaK | CD630_s0450 | 155 -- 200 | 135 -- 187 | -10.1353 | 3.04530 | 5.67265 |
| CD630_31010 | CD630_s0450 | 140 -- 157 | 170 -- 187 | -10.1306 | 4.96305 | 5.67265 |
| CD630_33650 | CD630_s0450 | 6 -- 34 | 66 -- 96 | -10.1232 | 4.80565 | 5.67265 |
| blaI | CD630_s0450 | 164 -- 175 | 24 -- 36 | -10.1088 | 3.73674 | 5.67265 |
| CD630_09050 | CD630_s0450 | 176 -- 193 | 169 -- 187 | -10.108 | 3.96216 | 5.67265 |
| CD630_01700 | CD630_s0450 | 1 -- 13 | 63 -- 75 | -10.0725 | 4.97234 | 5.67265 |
| CD630_15060 | CD630_s0450 | 4 -- 21 | 26 -- 43 | -10.0277 | 3.69396 | 5.67265 |
| CD630_07121 | CD630_s0450 | 120 -- 140 | 25 -- 47 | -10.0129 | 3.26177 | 5.67265 |
| CD630_14150 | CD630_s0450 | 41 -- 70 | 56 -- 83 | -9.99854 | 3.28251 | 5.67265 |
| dpaA | CD630_s0450 | 134 -- 197 | 143 -- 207 | -9.91789 | 4.65364 | 5.67265 |
| CD630_34560 | CD630_s0450 | 17 -- 32 | 65 -- 81 | -9.90091 | 4.02604 | 5.67265 |

| | | | | | | |
|---|---|---|---|---|---|---|
| CD630_22550 | CD630_s0450 | 133 – 177 | 67 – 118 | -9.88398 | 3.62379 | 5.67265 |
| CD630_20960 | CD630_s0450 | 157 – 199 | 34 – 76 | -9.81314 | 5.65940 | 5.67265 |
| CD630_34030 | CD630_s0450 | 28 – 42 | 101 – 115 | -9.80572 | 3.05766 | 5.67265 |
| CD630_04770 | CD630_s0450 | 39 – 68 | 120 – 150 | -9.78525 | 3.58478 | 5.67265 |
| CD630_08140 | CD630_s0450 | 1 – 12 | 27 – 39 | -9.7564 | 3.30159 | 5.67265 |
| kdgT1 | CD630_s0450 | 180 – 199 | 31 – 47 | -9.75537 | 4.49207 | 5.67265 |
| CD630_04620 | CD630_s0450 | 129 – 193 | 56 – 114 | -9.74669 | 3.45413 | 5.67265 |
| CD630_01690 | CD630_s0450 | 168 – 189 | 100 – 122 | -9.74468 | 4.97234 | 5.67265 |
| CD630_27310 | CD630_s0450 | 176 – 189 | 182 – 195 | -9.67377 | 4.01622 | 5.67265 |
| ppnK | CD630_s0450 | 79 – 102 | 186 – 208 | -9.67242 | 4.36817 | 5.67265 |
| CD630_07870 | CD630_s0450 | 161 – 200 | 9 – 46 | -9.65388 | 3.22829 | 5.67265 |
| CD630_05822 | CD630_s0450 | 129 – 137 | 31 – 39 | -9.65343 | 4.28113 | 5.67265 |
| panB | CD630_s0450 | 157 – 180 | 170 – 190 | -9.64423 | 3.40248 | 5.67265 |
| CD630_10790 | CD630_s0450 | 187 – 197 | 140 – 150 | -9.61478 | 4.45284 | 5.67265 |
| eutW | CD630_s0450 | 12 – 27 | 183 – 197 | -9.60393 | 4.55715 | 5.67265 |
| CD630_32890 | CD630_s0450 | 151 – 163 | 34 – 46 | -9.60101 | 3.41276 | 5.67265 |
| CD630_22500 | CD630_s0450 | 132 – 145 | 138 – 151 | -9.59621 | 4.03688 | 5.67265 |
| dnaC | CD630_s0450 | 125 – 157 | 154 – 187 | -9.56944 | 6.30796 | 5.67265 |
| CD630_04341 | CD630_s0450 | 158 – 195 | 182 – 207 | -9.50455 | 4.29262 | 5.67265 |
| CD630_31220 | CD630_s0460 | 163 – 181 | 1 – 19 | -17.6458 | 3.73779 | 5.74527 |
| CD630_08140 | CD630_s0460 | 4 – 15 | 4 – 15 | -15.8097 | 3.30159 | 5.74527 |
| CD630_21090 | CD630_s0460 | 142 – 165 | 6 – 22 | -13.9849 | 4.15803 | 5.74527 |
| spoVFB | CD630_s0460 | 2 – 15 | 1 – 14 | -12.1226 | 4.62758 | 5.74527 |
| CD630_05460 | CD630_s0460 | 113 – 180 | 1 – 66 | -11.8624 | 3.98753 | 5.74527 |
| CD630_26020 | CD630_s0460 | 108 – 158 | 27 – 66 | -11.76 | 4.03155 | 5.74527 |
| rolB | CD630_s0460 | 67 – 151 | 11 – 80 | -11.6657 | 3.60270 | 5.74527 |
| CD630_05830 | CD630_s0460 | 122 – 137 | 151 – 165 | -11.4825 | 3.35360 | 5.74527 |
| CD630_18910 | CD630_s0460 | 27 – 36 | 4 – 13 | -11.462 | 3.30688 | 5.74527 |
| CD630_31461 | CD630_s0460 | 7 – 23 | 12 – 29 | -11.3517 | 4.41476 | 5.74527 |
| CD630_01560 | CD630_s0460 | 178 – 193 | 7 – 22 | -11.3158 | 4.06352 | 5.74527 |
| dltB | CD630_s0460 | 131 – 165 | 43 – 74 | -11.2982 | 3.81485 | 5.74527 |
| cprK | CD630_s0460 | 79 – 112 | 1 – 31 | -10.9904 | 3.96216 | 5.74527 |
| CD630_06970 | CD630_s0460 | 144 – 157 | 7 – 20 | -10.9399 | 3.82312 | 5.74527 |
| CD630_03650 | CD630_s0460 | 151 – 188 | 4 – 45 | -10.8873 | 3.93697 | 5.74527 |
| CD630_01570 | CD630_s0460 | 109 – 129 | 150 – 168 | -10.8386 | 6.08699 | 5.74527 |
| CD630_04341 | CD630_s0460 | 177 – 195 | 6 – 26 | -10.8109 | 4.29262 | 5.74527 |
| CD630_14410 | CD630_s0460 | 102 – 124 | 4 – 26 | -10.7456 | 3.39433 | 5.74527 |
| CD630_27280 | CD630_s0460 | 135 – 198 | 6 – 69 | -10.6634 | 3.24738 | 5.74527 |
| CD630_05950 | CD630_s0460 | 177 – 192 | 7 – 23 | -10.6117 | 3.31669 | 5.74527 |
| CD630_05870 | CD630_s0460 | 1 – 43 | 28 – 75 | -10.5414 | 4.28339 | 5.74527 |
| cwp26 | CD630_s0460 | 125 – 142 | 6 – 23 | -10.3383 | 4.44377 | 5.74527 |
| CD630_19430 | CD630_s0460 | 122 – 139 | 4 – 21 | -10.3192 | 3.13922 | 5.74527 |
| CD630_09100 | CD630_s0460 | 12 – 54 | 5 – 41 | -10.2374 | 5.92260 | 5.74527 |
| CD630_21790 | CD630_s0460 | 178 – 192 | 4 – 19 | -10.2078 | 4.93762 | 5.74527 |
| CD630_34520 | CD630_s0460 | 126 – 154 | 28 – 56 | -10.1859 | 3.83016 | 5.74527 |
| CD630_19700 | CD630_s0460 | 168 – 196 | 33 – 63 | -10.1733 | 3.85861 | 5.74527 |
| CD630_17511 | CD630_s0460 | 100 – 114 | 8 – 22 | -10.0799 | 9.12322 | 5.74527 |
| CD630_07770 | CD630_s0460 | 134 – 147 | 8 – 21 | -10.0649 | 3.31158 | 5.74527 |
| CD630_06110 | CD630_s0460 | 181 – 197 | 7 – 23 | -10.0498 | 5.61380 | 5.74527 |
| gcp | CD630_s0460 | 131 – 155 | 6 – 28 | -10.0348 | 4.92798 | 5.74527 |
| CD630_28890 | CD630_s0460 | 24 – 53 | 140 – 169 | -9.9663 | 7.52469 | 5.74527 |
| ribD | CD630_s0460 | 149 – 178 | 1 – 31 | -9.88794 | 6.65922 | 5.74527 |
| chrA | CD630_s0460 | 175 – 191 | 1 – 18 | -9.88793 | 5.27255 | 5.74527 |
| CD630_33300 | CD630_s0460 | 133 – 150 | 151 – 168 | -9.88564 | 3.79149 | 5.74527 |
| CD630_22991 | CD630_s0460 | 188 – 199 | 7 – 19 | -9.83601 | 6.03728 | 5.74527 |
| CD630_06100 | CD630_s0460 | 164 – 197 | 36 – 75 | -9.82918 | 4.43918 | 5.74527 |
| CD630_04850 | CD630_s0460 | 150 – 171 | 148 – 169 | -9.52138 | 4.40788 | 5.74527 |
| dpaA | CD630_s0470 | 148 – 200 | 43 – 88 | -18.0297 | 4.65364 | 5.35019 |
| CD630_01700 | CD630_s0470 | 90 – 115 | 30 – 56 | -16.72 | 4.97234 | 5.35019 |
| CD630_05230 | CD630_s0470 | 26 – 57 | 69 – 99 | -15.5456 | 3.73787 | 5.35019 |
| CD630_14620 | CD630_s0470 | 11 – 64 | 3 – 51 | -15.4521 | 4.43651 | 5.35019 |
| CD630_06050 | CD630_s0470 | 90 – 173 | 1 – 91 | -15.366 | 3.59332 | 5.35019 |
| CD630_17170 | CD630_s0470 | 167 – 184 | 35 – 56 | -12.411 | 4.10230 | 5.35019 |
| CD630_15461 | CD630_s0470 | 152 – 190 | 18 – 55 | -12.4056 | 9.13442 | 5.35019 |
| CD630_03170 | CD630_s0470 | 120 – 168 | 31 – 76 | -12.3731 | 3.66946 | 5.35019 |
| cheW | CD630_s0470 | 75 – 125 | 34 – 88 | -12.3115 | 3.72371 | 5.35019 |
| CD630_18680 | CD630_s0470 | 46 – 92 | 27 – 91 | -12.0738 | 3.04315 | 5.35019 |
| CD630_25300 | CD630_s0470 | 101 – 121 | 2 – 22 | -11.9685 | 4.49333 | 5.35019 |
| CD630_31970 | CD630_s0470 | 153 – 170 | 1 – 17 | -11.7747 | 3.24511 | 5.35019 |
| CD630_24590 | CD630_s0470 | 113 – 199 | 10 – 88 | -11.7633 | 3.70401 | 5.35019 |
| CD630_05460 | CD630_s0470 | 1 – 25 | 1 – 25 | -11.6875 | 3.98753 | 5.35019 |
| moaA | CD630_s0470 | 50 – 110 | 36 – 91 | -11.5918 | 3.83799 | 5.35019 |
| CD630_19942 | CD630_s0470 | 165 – 181 | 4 – 22 | -11.493 | 4.24574 | 5.35019 |
| cwp17 | CD630_s0470 | 70 – 119 | 4 – 55 | -11.2206 | 3.43363 | 5.35019 |
| garK | CD630_s0470 | 19 – 34 | 3 – 18 | -11.0387 | 3.72836 | 5.35019 |
| CD630_09990 | CD630_s0470 | 142 – 160 | 50 – 67 | -11.0245 | 3.86040 | 5.35019 |
| CD630_10220 | CD630_s0470 | 119 – 173 | 9 – 54 | -11.0165 | 4.08233 | 5.35019 |
| CD630_08360 | CD630_s0470 | 141 – 172 | 1 – 32 | -10.7978 | 3.19339 | 5.35019 |
| CD630_25620 | CD630_s0470 | 183 – 199 | 38 – 54 | -10.6999 | 3.02685 | 5.35019 |
| CD630_25610 | CD630_s0470 | 93 – 158 | 3 – 53 | -10.6329 | 3.02685 | 5.35019 |
| CD630_18910 | CD630_s0470 | 178 – 197 | 75 – 95 | -10.6253 | 3.30688 | 5.35019 |
| CD630_25050 | CD630_s0470 | 43 – 76 | 2 – 32 | -10.588 | 4.33624 | 5.35019 |
| CD630_03930 | CD630_s0470 | 132 – 149 | 30 – 51 | -10.4928 | 4.15934 | 5.35019 |
| CD630_17380 | CD630_s0470 | 165 – 192 | 7 – 31 | -10.4453 | 5.03207 | 5.35019 |
| CD630_22850 | CD630_s0470 | 137 – 192 | 3 – 54 | -10.3808 | 3.84102 | 5.35019 |
| feoB | CD630_s0470 | 26 – 40 | 39 – 54 | -10.2699 | 4.52257 | 5.35019 |
| bltD | CD630_s0470 | 171 – 199 | 3 – 33 | -10.0428 | 4.49926 | 5.35019 |
| CD630_15431 | CD630_s0470 | 140 – 191 | 30 – 89 | -10.041 | 5.29780 | 5.35019 |
| CD630_36170 | CD630_s0470 | 1 – 18 | 2 – 20 | -10.0064 | 3.72380 | 5.35019 |
| CD630_19700 | CD630_s0470 | 4 – 48 | 48 – 85 | -9.99386 | 3.85861 | 5.35019 |
| CD630_19790 | CD630_s0470 | 61 – 104 | 42 – 83 | -9.9586 | 3.79593 | 5.35019 |
| cbiO | CD630_s0470 | 19 – 29 | 41 – 51 | -9.88038 | 3.79830 | 5.35019 |
| etfA | CD630_s0470 | 147 – 160 | 53 – 66 | -9.87361 | 3.41204 | 5.35019 |
| CD630_31530 | CD630_s0470 | 10 – 41 | 2 – 34 | -9.87241 | 3.49100 | 5.35019 |
| aroD | CD630_s0470 | 143 – 201 | 3 – 56 | -9.82546 | 3.71929 | 5.35019 |
| CD630_13350 | CD630_s0470 | 95 – 122 | 3 – 28 | -9.82329 | 4.15139 | 5.35019 |
| CD630_08460 | CD630_s0470 | 14 – 43 | 1 – 30 | -9.7874 | 4.15083 | 5.35019 |
| dltA | CD630_s0470 | 135 – 148 | 3 – 17 | -9.74539 | 3.85429 | 5.35019 |
| xdhA | CD630_s0470 | 39 – 103 | 1 – 73 | -9.68775 | 4.45147 | 5.35019 |
| CD630_14420 | CD630_s0470 | 17 – 28 | 1 – 12 | -9.68454 | 3.99497 | 5.35019 |
| CD630_05030 | CD630_s0470 | 1 – 33 | 11 – 45 | -9.62737 | 4.03823 | 5.35019 |
| cstA | CD630_s0470 | 121 – 170 | 49 – 94 | -9.60536 | 3.53584 | 5.35019 |
| CD630_26050 | CD630_s0470 | 156 – 189 | 41 – 75 | -9.60238 | 6.99851 | 5.35019 |
| CD630_06060 | CD630_s0470 | 92 – 141 | 3 – 53 | -9.59886 | 4.94050 | 5.35019 |
| bglA | CD630_s0470 | 152 – 198 | 28 – 70 | -9.59827 | 4.86402 | 5.35019 |
| CD630_22660 | CD630_s0470 | 150 – 197 | 50 – 99 | -9.57319 | 3.60496 | 5.35019 |
| CD630_24591 | CD630_s0470 | 89 – 108 | 48 – 66 | -9.53788 | 3.54497 | 5.35019 |
| CD630_34530 | CD630_s0470 | 131 – 179 | 1 – 48 | -9.50229 | 3.83016 | 5.35019 |
| CD630_27280 | CD630_s0500 | 181 – 198 | 85 – 103 | -13.5536 | 3.24738 | 3.25870 |
| bglA | CD630_s0500 | 148 – 201 | 27 – 79 | -13.1804 | 4.86402 | 3.25870 |
| CD630_03400 | CD630_s0500 | 164 – 178 | 89 – 103 | -12.1563 | 4.45994 | 3.25870 |
| CD630_29840 | CD630_s0500 | 178 – 195 | 84 – 101 | -11.3764 | 4.48825 | 3.25870 |
| CD630_32690 | CD630_s0500 | 18 – 46 | 21 – 57 | -11.3582 | 7.32676 | 3.25870 |
| CD630_07400 | CD630_s0500 | 164 – 173 | 87 – 96 | -11.3537 | 4.52273 | 3.25870 |
| ppnK | CD630_s0500 | 54 – 102 | 11 – 55 | -11.3523 | 4.36817 | 3.25870 |
| CD630_04610 | CD630_s0500 | 188 – 201 | 90 – 103 | -11.1088 | 3.83025 | 3.25870 |
| CD630_19680 | CD630_s0500 | 123 – 147 | 11 – 35 | -10.8508 | 3.58080 | 3.25870 |
| CD630_20450 | CD630_s0500 | 36 – 57 | 38 – 55 | -10.6735 | 3.77388 | 3.25870 |
| CD630_36690 | CD630_s0500 | 141 – 156 | 88 – 102 | -10.6326 | 3.53761 | 3.25870 |
| CD630_16950 | CD630_s0500 | 179 – 199 | 64 – 87 | -10.5036 | 4.42569 | 3.25870 |
| CD630_13020 | CD630_s0500 | 113 – 123 | 93 – 103 | -10.3534 | 3.52267 | 3.25870 |
| eutV | CD630_s0500 | 87 – 107 | 43 – 63 | -10.2016 | 4.55715 | 3.25870 |
| CD630_02420 | CD630_s0500 | 91 – 142 | 16 – 55 | -10.1936 | 3.71048 | 3.25870 |
| CD630_28870 | CD630_s0500 | 170 – 182 | 88 – 100 | -10.0484 | 3.41991 | 3.25870 |
| CD630_21610 | CD630_s0500 | 160 – 173 | 89 – 102 | -10.0175 | 3.16662 | 3.25870 |
| CD630_15060 | CD630_s0500 | 180 – 195 | 88 – 102 | -9.73043 | 3.69396 | 3.25870 |
| CD630_30110 | CD630_s0500 | 136 – 166 | 16 – 58 | -9.58941 | 3.31041 | 3.25870 |
| CD630_06060 | CD630_s0500 | 77 – 152 | 17 – 72 | -9.56611 | 4.94050 | 3.25870 |
| CD630_08080 | CD630_s0510 | 18 – 69 | 6 – 62 | -16.4607 | 4.87819 | 3.37631 |
| CD630_22550 | CD630_s0510 | 24 – 59 | 117 – 154 | -16.3113 | 3.62379 | 3.37631 |
| CD630_06100 | CD630_s0510 | 66 – 163 | 6 – 91 | -15.9102 | 4.43918 | 3.37631 |
| cwp29 | CD630_s0510 | 120 – 165 | 13 – 62 | -15.5533 | 4.40865 | 3.37631 |
| CD630_03170 | CD630_s0510 | 149 – 178 | 6 – 40 | -15.4822 | 3.66946 | 3.37631 |
| CD630_06060 | CD630_s0510 | 124 – 186 | 3 – 68 | -15.4331 | 4.94050 | 3.37631 |
| CD630_01690 | CD630_s0510 | 168 – 186 | 105 – 123 | -15.2984 | 4.97234 | 3.37631 |
| CD630_21090 | CD630_s0510 | 156 – 172 | 6 – 23 | -15.2403 | 4.15803 | 3.37631 |
| CD630_04880 | CD630_s0510 | 136 – 191 | 8 – 70 | -15.1713 | 4.62291 | 3.37631 |
| CD630_07430 | CD630_s0510 | 82 – 110 | 172 – 203 | -15.0451 | 4.65568 | 3.37631 |
| CD630_28290 | CD630_s0510 | 27 – 67 | 10 – 50 | -14.8211 | 5.54192 | 3.37631 |
| CD630_10950 | CD630_s0510 | 119 – 193 | 29 – 110 | -14.7012 | 3.67675 | 3.37631 |
| CD630_12110 | CD630_s0510 | 103 – 135 | 26 – 54 | -14.5434 | 4.36078 | 3.37631 |
| CD630_22950 | CD630_s0510 | 6 – 50 | 19 – 62 | -14.5332 | 3.96883 | 3.37631 |
| CD630_30360 | CD630_s0510 | 98 – 156 | 8 – 61 | -14.4762 | 4.26509 | 3.37631 |
| spaK | CD630_s0510 | 148 – 200 | 5 – 62 | -14.1484 | 3.04530 | 3.37631 |
| cdtR | CD630_s0510 | 4 – 62 | 6 – 62 | -14.0147 | 5.84368 | 3.37631 |
| CD630_14130 | CD630_s0510 | 135 – 162 | 45 – 75 | -13.6835 | 4.57970 | 3.37631 |
| dnaD | CD630_s0510 | 131 – 151 | 39 – 57 | -13.5983 | 3.56078 | 3.37631 |
| CD630_18930 | CD630_s0510 | 154 – 182 | 33 – 60 | -13.4873 | 5.32127 | 3.37631 |
| CD630_36781 | CD630_s0510 | 163 – 187 | 128 – 152 | -13.4751 | 3.75946 | 3.37631 |
| CD630_05211 | CD630_s0510 | 20 – 28 | 42 – 50 | -13.1007 | 3.97048 | 3.37631 |
| cwp26 | CD630_s0510 | 123 – 171 | 10 – 59 | -13.0293 | 4.44377 | 3.37631 |
| CD630_04850 | CD630_s0510 | 128 – 178 | 109 – 149 | -13.0144 | 4.40788 | 3.37631 |
| CD630_23450 | CD630_s0510 | 183 – 193 | 41 – 52 | -12.9004 | 3.27218 | 3.37631 |
| CD630_15160 | CD630_s0510 | 3 – 17 | 101 – 115 | -12.7632 | 4.32318 | 3.37631 |
| CD630_29250 | CD630_s0510 | 10 – 39 | 31 – 62 | -12.4166 | 4.61901 | 3.37631 |
| CD630_09410 | CD630_s0510 | 10 – 39 | 31 – 62 | -12.4166 | 4.75651 | 3.37631 |
| CD630_16940 | CD630_s0510 | 85 – 144 | 76 – 133 | -12.4082 | 4.14665 | 3.37631 |
| CD630_21760 | CD630_s0510 | 93 – 200 | 16 – 122 | -12.3709 | 4.43846 | 3.37631 |
| CD630_15111 | CD630_s0510 | 19 – 56 | 9 – 51 | -12.3444 | 4.18664 | 3.37631 |
| CD630_18781 | CD630_s0510 | 137 – 199 | 66 – 127 | -12.2648 | 3.77660 | 3.37631 |
| CD630_15080 | CD630_s0510 | 8 – 24 | 43 – 59 | -12.1654 | 4.03033 | 3.37631 |
| CD630_06170 | CD630_s0510 | 114 – 154 | 3 – 49 | -12.1603 | 4.95366 | 3.37631 |
| CD630_25050 | CD630_s0510 | 96 – 139 | 36 – 71 | -12.0938 | 4.33624 | 3.37631 |
| CD630_36690 | CD630_s0510 | 4 – 48 | 8 – 47 | -11.9054 | 3.53761 | 3.37631 |
| CD630_13530 | CD630_s0510 | 5 – 22 | 4 – 20 | -11.7673 | 4.09832 | 3.37631 |
| CD630_13950 | CD630_s0510 | 134 – 191 | 5 – 57 | -11.6981 | 4.25540 | 3.37631 |
| CD630_21260 | CD630_s0510 | 122 – 140 | 6 – 25 | -11.6807 | 4.22255 | 3.37631 |
| ldh | CD630_s0510 | 123 – 179 | 8 – 62 | -11.6749 | 4.35818 | 3.37631 |
| CD630_05830 | CD630_s0510 | 112 – 175 | 1 – 59 | -11.6622 | 3.35360 | 3.37631 |
| rbsC | CD630_s0510 | 170 – 182 | 193 – 205 | -11.6046 | 3.71650 | 3.37631 |
| CD630_32990 | CD630_s0510 | 78 – 160 | 6 – 81 | -11.5365 | 5.24925 | 3.37631 |
| CD630_19440 | CD630_s0510 | 163 – 182 | 29 – 54 | -11.5323 | 4.53937 | 3.37631 |
| tcdA | CD630_s0510 | 149 – 159 | 193 – 203 | -11.48 | 4.70593 | 3.37631 |
| CD630_18910 | CD630_s0510 | 24 – 63 | 12 – 49 | -11.2755 | 3.30688 | 3.37631 |
| CD630_03180 | CD630_s0510 | 6 – 64 | 2 – 55 | -11.1374 | 3.73138 | 3.37631 |
| CD630_30100 | CD630_s0510 | 144 – 179 | 31 – 62 | -11.0888 | 4.13234 | 3.37631 |
| panB | CD630_s0510 | 82 – 186 | 31 – 138 | -11.0333 | 3.40248 | 3.37631 |
| CD630_18680 | CD630_s0510 | 48 – 92 | 26 – 75 | -11.0212 | 3.04315 | 3.37631 |
| CD630_17360 | CD630_s0510 | 189 – 198 | 42 – 51 | -10.8073 | 5.66070 | 3.37631 |
| folC | CD630_s0510 | 16 – 159 | 24 – 142 | -10.7871 | 3.85634 | 3.37631 |
| CD630_16160 | CD630_s0510 | 170 – 186 | 135 – 150 | -10.7643 | 6.11801 | 3.37631 |
| CD630_22991 | CD630_s0510 | 189 – 197 | 168 – 176 | -10.757 | 6.03728 | 3.37631 |
| CD630_34520 | CD630_s0510 | 136 – 150 | 25 – 39 | -10.7202 | 3.83016 | 3.37631 |
| CD630_08370 | CD630_s0510 | 189 – 198 | 4 – 13 | -10.7 | 3.65639 | 3.37631 |
| CD630_26050 | CD630_s0510 | 174 – 199 | 38 – 61 | -10.697 | 6.99851 | 3.37631 |
| CD630_27630 | CD630_s0510 | 110 – 195 | 8 – 93 | -10.6729 | 3.54199 | 3.37631 |
| CD630_34050 | CD630_s0510 | 2 – 45 | 19 – 62 | -10.6166 | 4.03863 | 3.37631 |
| CD630_14180 | CD630_s0510 | 165 – 195 | 19 – 48 | -10.6093 | 3.93696 | 3.37631 |
| CD630_14182 | CD630_s0510 | 189 – 198 | 167 – 176 | -10.604 | 4.82360 | 3.37631 |
| CD630_07480 | CD630_s0510 | 157 – 166 | 34 – 43 | -10.5757 | 5.18321 | 3.37631 |
| CD630_13790 | CD630_s0510 | 79 – 113 | 167 – 208 | -10.561 | 5.17374 | 3.37631 |
| CD630_19904 | CD630_s0510 | 9 – 32 | 36 – 51 | -10.5445 | 5.96526 | 3.37631 |
| CD630_19800 | CD630_s0510 | 3 – 31 | 38 – 61 | -10.5432 | 5.22035 | 3.37631 |
| CD630_13880 | CD630_s0510 | 154 – 175 | 26 – 47 | -10.4408 | 3.52735 | 3.37631 |
| CD630_07121 | CD630_s0510 | 119 – 135 | 3 – 17 | -10.3958 | 3.26177 | 3.37631 |
| CD630_23780 | CD630_s0510 | 148 – 170 | 4 – 24 | -10.3358 | 3.91629 | 3.37631 |
| gcp | CD630_s0510 | 136 – 151 | 108 – 123 | -10.2987 | 4.92798 | 3.37631 |
| CD630_27240 | CD630_s0510 | 159 – 173 | 110 – 123 | -10.265 | 3.52462 | 3.37631 |
| CD630_07840 | CD630_s0510 | 146 – 178 | 31 – 61 | -10.2579 | 4.26761 | 3.37631 |
| CD630_31530 | CD630_s0510 | 12 – 56 | 33 – 76 | -10.2238 | 3.49100 | 3.37631 |
| CD630_17511 | CD630_s0510 | 2 – 60 | 3 – 62 | -10.2187 | 9.12322 | 3.37631 |
| CD630_36180 | CD630_s0510 | 164 – 200 | 8 – 32 | -10.207 | 4.83428 | 3.37631 |
| CD630_32060 | CD630_s0510 | 1 – 11 | 199 – 209 | -10.1303 | 5.73797 | 3.37631 |
| CD630_29760 | CD630_s0510 | 177 – 199 | 112 – 133 | -10.1236 | 3.51074 | 3.37631 |
| CD630_31880 | CD630_s0510 | 22 – 44 | 69 – 91 | -10.105 | 3.97522 | 3.37631 |
| CD630_21490 | CD630_s0510 | 127 – 180 | 166 – 204 | -10.0948 | 3.37462 | 3.37631 |
| adhE | CD630_s0510 | 37 – 59 | 47 – 69 | -10.0696 | 3.06485 | 3.37631 |
| CD630_03400 | CD630_s0510 | 2 – 40 | 6 – 43 | -10.0192 | 4.45994 | 3.37631 |
| CD630_32050 | CD630_s0510 | 119 – 141 | 188 – 209 | -9.95471 | 5.73797 | 3.37631 |
| kdpD | CD630_s0510 | 151 – 201 | 167 – 207 | -9.9401 | 4.27140 | 3.37631 |
| CD630_02720 | CD630_s0510 | 139 – 187 | 5 – 50 | -9.92918 | 3.38938 | 3.37631 |
| CD630_04320 | CD630_s0510 | 170 – 185 | 172 – 187 | -9.92014 | 3.68342 | 3.37631 |
| feoB | CD630_s0510 | 115 – 163 | 8 – 61 | -9.88213 | 4.52257 | 3.37631 |
| glgB | CD630_s0510 | 143 – 161 | 104 – 123 | -9.88113 | 4.48273 | 3.37631 |
| CD630_26150 | CD630_s0510 | 123 – 150 | 33 – 55 | -9.87392 | 4.06302 | 3.37631 |
| terD | CD630_s0510 | 118 – 171 | 166 – 207 | -9.87182 | 3.33148 | 3.37631 |
| CD630_27110 | CD630_s0510 | 2 – 15 | 136 – 149 | -9.79077 | 3.78231 | 3.37631 |
| CD630_06110 | CD630_s0510 | 145 – 154 | 104 – 113 | -9.77104 | 5.61380 | 3.37631 |
| CD630_13620 | CD630_s0510 | 114 – 160 | 5 – 55 | -9.75944 | 4.77627 | 3.37631 |
| CD630_20171 | CD630_s0510 | 42 – 59 | 20 – 37 | -9.74819 | 6.87491 | 3.37631 |
| CD630_16550 | CD630_s0510 | 16 – 28 | 170 – 181 | -9.73068 | 5.61064 | 3.37631 |
| CD630_03650 | CD630_s0510 | 151 – 195 | 5 – 54 | -9.72253 | 3.93697 | 3.37631 |
| malY | CD630_s0510 | 6 – 95 | 52 – 131 | -9.71951 | 4.46345 | 3.37631 |
| CD630_13370 | CD630_s0510 | 158 – 177 | 5 – 22 | -9.68019 | 4.33145 | 3.37631 |
| feoB | CD630_s0510 | 104 – 139 | 17 – 55 | -9.65762 | 4.05364 | 3.37631 |
| CD630_06060 | CD630_s0510 | 24 – 50 | 182 – 209 | -9.64329 | 4.94050 | 3.37631 |
| CD630_07400 | CD630_s0510 | 112 – 121 | 8 – 17 | -9.63361 | 4.52273 | 3.37631 |
| CD630_14260 | CD630_s0510 | 23 – 29 | 44 – 50 | -9.55596 | 3.22214 | 3.37631 |
| CD630_18010 | CD630_s0510 | 153 – 193 | 5 – 56 | -9.55529 | 4.21967 | 3.37631 |
| CD630_14110 | CD630_s0510 | 1 – 25 | 10 – 33 | -9.55032 | 3.39550 | 3.37631 |
| CD630_08110 | CD630_s0510 | 121 – 134 | 17 – 30 | -9.54657 | 4.39181 | 3.37631 |
| CD630_29840 | CD630_s0510 | 61 – 94 | 127 – 161 | -9.53262 | 4.48825 | 3.37631 |
| CD630_33610 | CD630_s0510 | 181 – 195 | 139 – 153 | -9.52249 | 3.79322 | 3.37631 |
| CD630_06100 | CD630_s0590 | 104 – 154 | 28 – 82 | -16.7104 | 4.43918 | 8.13716 |
| chrA | CD630_s0590 | 151 – 166 | 134 – 149 | -15.0242 | 5.27255 | 8.13716 |
| CD630_21090 | CD630_s0590 | 157 – 170 | 1 – 14 | -14.5063 | 4.15803 | 8.13716 |
| CD630_21140 | CD630_s0590 | 7 – 20 | 165 – 178 | -14.3359 | 3.33304 | 8.13716 |
| CD630_08140 | CD630_s0590 | 4 – 17 | 1 – 15 | -14.1812 | 3.30159 | 8.13716 |

| | | | | | | |
|---|---|---|---|---|---|---|
| CD630_13580 | CD630_s0590 | 1 – 25 | 29 – 54 | -13.8292 | 3.80715 | 8.13716 |
| CD630_06060 | CD630_s0590 | 174 – 190 | 134 – 153 | -12.7931 | 4.94050 | 8.13716 |
| CD630_17350 | CD630_s0590 | 183 – 198 | 1 – 16 | -12.7352 | 3.35334 | 8.13716 |
| CD630_22070 | CD630_s0590 | 89 – 141 | 111 – 153 | -12.3119 | 4.09651 | 8.13716 |
| CD630_09100 | CD630_s0590 | 43 – 54 | 5 – 16 | -12.218 | 5.92260 | 8.13716 |
| CD630_14410 | CD630_s0590 | 160 – 178 | 136 – 157 | -12.1986 | 3.39433 | 8.13716 |
| opuCA | CD630_s0590 | 88 – 107 | 135 – 153 | -12.1312 | 4.07035 | 8.13716 |
| CD630_06920 | CD630_s0590 | 117 – 144 | 71 – 99 | -11.8332 | 3.13260 | 8.13716 |
| CD630_22991 | CD630_s0590 | 193 – 199 | 7 – 13 | -11.7308 | 6.03728 | 8.13716 |
| CD630_15461 | CD630_s0590 | 90 – 118 | 137 – 153 | -11.7203 | 9.13442 | 8.13716 |
| CD630_33560 | CD630_s0590 | 172 – 194 | 75 – 96 | -11.5715 | 4.03147 | 8.13716 |
| CD630_01640 | CD630_s0590 | 145 – 159 | 143 – 157 | -11.4616 | 4.55381 | 8.13716 |
| CD630_01690 | CD630_s0590 | 190 – 201 | 5 – 16 | -11.4296 | 4.97234 | 8.13716 |
| CD630_04770 | CD630_s0590 | 39 – 55 | 1 – 15 | -11.3962 | 3.58478 | 8.13716 |
| feoB | CD630_s0590 | 22 – 32 | 166 – 176 | -11.3897 | 4.05364 | 8.13716 |
| CD630_22720 | CD630_s0590 | 124 – 154 | 128 – 154 | -11.3648 | 3.81804 | 8.13716 |
| CD630_06060 | CD630_s0590 | 83 – 145 | 83 – 152 | -11.2703 | 4.94050 | 8.13716 |
| CD630_19580 | CD630_s0590 | 180 – 192 | 132 – 144 | -11.1506 | 4.51902 | 8.13716 |
| CD630_07770 | CD630_s0590 | 137 – 152 | 4 – 18 | -10.9016 | 3.31158 | 8.13716 |
| cdtR | CD630_s0590 | 3 – 14 | 1 – 11 | -10.8392 | 5.84368 | 8.13716 |
| CD630_19440 | CD630_s0590 | 141 – 154 | 3 – 16 | -10.7781 | 4.53937 | 8.13716 |
| CD630_22550 | CD630_s0590 | 151 – 189 | 36 – 76 | -10.7765 | 3.62379 | 8.13716 |
| CD630_21260 | CD630_s0590 | 41 – 56 | 1 – 17 | -10.7087 | 4.22255 | 8.13716 |
| iorA | CD630_s0590 | 110 – 119 | 135 – 144 | -10.7023 | 3.83173 | 8.13716 |
| CD630_25050 | CD630_s0590 | 89 – 107 | 135 – 153 | -10.5869 | 4.33624 | 8.13716 |
| CD630_11220 | CD630_s0590 | 184 – 191 | 135 – 142 | -10.4555 | 3.52327 | 8.13716 |
| CD630_16460 | CD630_s0590 | 113 – 133 | 79 – 97 | -10.403 | 4.64028 | 8.13716 |
| CD630_06340 | CD630_s0590 | 116 – 137 | 78 – 101 | -10.3074 | 4.35983 | 8.13716 |
| kdpD | CD630_s0590 | 1 – 21 | 1 – 16 | -10.254 | 4.27140 | 8.13716 |
| CD630_27280 | CD630_s0590 | 187 – 194 | 5 – 12 | -10.2274 | 3.24738 | 8.13716 |
| CD630_20660 | CD630_s0590 | 28 – 38 | 3 – 13 | -10.1698 | 5.39321 | 8.13716 |
| CD630_22850 | CD630_s0590 | 178 – 186 | 74 – 82 | -10.0939 | 3.84102 | 8.13716 |
| CD630_08370 | CD630_s0590 | 89 – 107 | 135 – 153 | -10.0838 | 3.65639 | 8.13716 |
| CD630_27110 | CD630_s0590 | 51 – 59 | 8 – 16 | -10.0799 | 3.78231 | 8.13716 |
| pilC | CD630_s0590 | 69 – 82 | 137 – 150 | -10.0703 | 3.54230 | 8.13716 |
| panG | CD630_s0590 | 163 – 177 | 1 – 15 | -9.91042 | 5.54047 | 8.13716 |
| CD630_13020 | CD630_s0590 | 118 – 126 | 7 – 15 | -9.91015 | 3.52267 | 8.13716 |
| CD630_05830 | CD630_s0590 | 153 – 161 | 7 – 15 | -9.90703 | 3.35360 | 8.13716 |
| CD630_28550 | CD630_s0590 | 90 – 107 | 135 – 153 | -9.90397 | 3.51196 | 8.13716 |
| rolA | CD630_s0590 | 176 – 189 | 2 – 15 | -9.89301 | 4.56362 | 8.13716 |
| CD630_31381 | CD630_s0590 | 100 – 133 | 115 – 145 | -9.8393 | 3.49137 | 8.13716 |
| scrR | CD630_s0590 | 152 – 162 | 126 – 136 | -9.75525 | 3.49166 | 8.13716 |
| CD630_19430 | CD630_s0590 | 89 – 117 | 126 – 153 | -9.61822 | 3.13922 | 8.13716 |
| CD630_18910 | CD630_s0590 | 27 – 38 | 2 – 13 | -9.54669 | 3.30688 | 8.13716 |
| CD630_31530 | CD630_s0591 | 161 – 187 | 29 – 54 | -14.0192 | 3.49100 | 6.84967 |
| CD630_06170 | CD630_s0591 | 117 – 128 | 119 – 131 | -13.3001 | 4.95366 | 6.84967 |
| CD630_09100 | CD630_s0591 | 92 – 106 | 119 – 133 | -12.3231 | 5.92260 | 6.84967 |
| CD630_04550 | CD630_s0591 | 1 – 12 | 121 – 132 | -10.8895 | 3.05227 | 6.84967 |
| CD630_19904 | CD630_s0591 | 24 – 30 | 126 – 132 | -10.5942 | 5.96526 | 6.84967 |
| CD630_04620 | CD630_s0591 | 156 – 166 | 121 – 131 | -10.3928 | 3.45413 | 6.84967 |
| CD630_31690 | CD630_s0591 | 108 – 126 | 117 – 134 | -9.95583 | 3.71193 | 6.84967 |
| CD630_21260 | CD630_s0591 | 45 – 55 | 120 – 130 | -9.62821 | 4.22255 | 6.84967 |
| CD630_29250 | CD630_s0591 | 176 – 191 | 116 – 131 | -9.61415 | 4.61901 | 6.84967 |
| CD630_09410 | CD630_s0591 | 176 – 191 | 116 – 131 | -9.61415 | 4.75651 | 6.84967 |
| CD630_21090 | CD630_s0591 | 164 – 172 | 124 – 132 | -9.57889 | 4.15803 | 6.84967 |
| CD630_27570 | CD630_s0591 | 10 – 24 | 119 – 135 | -9.57713 | 3.46216 | 6.84967 |
| cbpA | CD630_s0591 | 158 – 179 | 32 – 54 | -9.50974 | 3.03040 | 6.84967 |
| CD630_30120 | CD630_s0631 | 10 – 37 | 80 – 105 | -18.2768 | 4.04859 | 4.95025 |
| CD630_21260 | CD630_s0631 | 150 – 180 | 51 – 83 | -16.038 | 4.22255 | 4.95025 |
| CD630_08140 | CD630_s0631 | 11 – 44 | 311 – 340 | -14.7348 | 3.30159 | 4.95025 |
| CD630_03400 | CD630_s0631 | 52 – 83 | 313 – 339 | -14.643 | 4.45994 | 4.95025 |
| CD630_17511 | CD630_s0631 | 6 – 37 | 3 – 37 | -14.6345 | 9.12322 | 4.95025 |
| CD630_09760 | CD630_s0631 | 47 – 104 | 150 – 202 | -14.4964 | 4.83410 | 4.95025 |
| CD630_28910 | CD630_s0631 | 47 – 104 | 150 – 202 | -14.4964 | 4.51664 | 4.95025 |
| pilC | CD630_s0631 | 128 – 146 | 252 – 267 | -14.3407 | 3.54230 | 4.95025 |
| CD630_18400 | CD630_s0631 | 183 – 201 | 31 – 50 | -14.1878 | 4.67962 | 4.95025 |
| CD630_05820 | CD630_s0631 | 112 – 124 | 329 – 342 | -14.1077 | 3.75661 | 4.95025 |
| CD630_19903 | CD630_s0631 | 53 – 116 | 288 – 341 | -13.8611 | 5.79994 | 4.95025 |
| CD630_21090 | CD630_s0631 | 112 – 183 | 9 – 72 | -13.8274 | 4.15803 | 4.95025 |
| rbsC | CD630_s0631 | 56 – 92 | 159 – 195 | -13.5226 | 3.71650 | 4.95025 |
| CD630_15270 | CD630_s0631 | 145 – 157 | 324 – 336 | -13.3791 | 4.66015 | 4.95025 |
| CD630_06060 | CD630_s0631 | 128 – 180 | 61 – 113 | -13.2615 | 4.94050 | 4.95025 |
| kdgT1 | CD630_s0631 | 116 – 148 | 63 – 96 | -13.1683 | 4.49207 | 4.95025 |
| CD630_13350 | CD630_s0631 | 168 – 200 | 61 – 85 | -13.1658 | 4.15139 | 4.95025 |
| CD630_03220 | CD630_s0631 | 87 – 135 | 70 – 116 | -13.0356 | 3.36746 | 4.95025 |
| CD630_09100 | CD630_s0631 | 3 – 55 | 7 – 75 | -13.0283 | 5.92260 | 4.95025 |
| CD630_02790 | CD630_s0631 | 21 – 50 | 312 – 339 | -12.9571 | 4.77661 | 4.95025 |
| CD630_33610 | CD630_s0631 | 116 – 192 | 24 – 99 | -12.8538 | 3.79322 | 4.95025 |
| CD630_04850 | CD630_s0631 | 148 – 195 | 21 – 69 | -12.7486 | 4.40788 | 4.95025 |
| dnaD | CD630_s0631 | 4 – 34 | 18 – 47 | -12.4722 | 3.56078 | 4.95025 |
| CD630_10800 | CD630_s0631 | 107 – 139 | 236 – 269 | -12.4088 | 5.30319 | 4.95025 |
| CD630_13530 | CD630_s0631 | 14 – 36 | 11 – 33 | -12.3689 | 4.09832 | 4.95025 |
| CD630_30540 | CD630_s0631 | 121 – 163 | 238 – 265 | -11.9172 | 3.63804 | 4.95025 |
| spoVFB | CD630_s0631 | 14 – 21 | 329 – 336 | -11.9044 | 4.62758 | 4.95025 |
| CD630_35840 | CD630_s0631 | 178 – 191 | 329 – 342 | -11.8873 | 3.14170 | 4.95025 |
| CD630_06050 | CD630_s0631 | 107 – 137 | 193 – 228 | -11.8624 | 3.59332 | 4.95025 |
| cobQ | CD630_s0631 | 87 – 102 | 252 – 267 | -11.8417 | 3.13524 | 4.95025 |
| CD630_33682 | CD630_s0631 | 58 – 116 | 288 – 335 | -11.814 | 5.77720 | 4.95025 |
| terD | CD630_s0631 | 117 – 136 | 248 – 267 | -11.8042 | 3.33148 | 4.95025 |
| CD630_35260 | CD630_s0631 | 179 – 200 | 24 – 44 | -11.7878 | 5.77277 | 4.95025 |
| CD630_22660 | CD630_s0631 | 151 – 192 | 58 – 98 | -11.5916 | 3.60496 | 4.95025 |
| gcvTPA | CD630_s0631 | 42 – 78 | 86 – 115 | -11.5837 | 4.77229 | 4.95025 |
| CD630_18230 | CD630_s0631 | 166 – 186 | 315 – 339 | -11.4801 | 3.00184 | 4.95025 |
| CD630_14240 | CD630_s0631 | 54 – 116 | 288 – 338 | -11.4769 | 8.60766 | 4.95025 |
| CD630_18610 | CD630_s0631 | 82 – 106 | 246 – 268 | -11.475 | 4.15582 | 4.95025 |
| CD630_21730 | CD630_s0631 | 181 – 197 | 94 – 109 | -11.47 | 3.45076 | 4.95025 |
| CD630_10220 | CD630_s0631 | 119 – 166 | 60 – 116 | -11.4216 | 4.08233 | 4.95025 |
| def1 | CD630_s0631 | 64 – 107 | 62 – 115 | -11.3248 | 3.99720 | 4.95025 |
| CD630_23090 | CD630_s0631 | 33 – 40 | 60 – 67 | -11.2537 | 4.25622 | 4.95025 |
| serS1 | CD630_s0631 | 158 – 171 | 254 – 267 | -11.1889 | 3.45858 | 4.95025 |
| CD630_06380 | CD630_s0631 | 179 – 199 | 70 – 88 | -11.1791 | 4.32211 | 4.95025 |
| CD630_14010 | CD630_s0631 | 37 – 64 | 2 – 29 | -11.173 | 3.69983 | 4.95025 |
| CD630_21700 | CD630_s0631 | 1 – 49 | 2 – 50 | -11.1227 | 3.96650 | 4.95025 |
| CD630_03420 | CD630_s0631 | 94 – 110 | 313 – 329 | -11.0368 | 4.71868 | 4.95025 |
| adhE | CD630_s0631 | 162 – 196 | 92 – 122 | -11.0245 | 3.06485 | 4.95025 |
| feoB | CD630_s0631 | 5 – 62 | 80 – 137 | -11.001 | 4.52257 | 4.95025 |
| CD630_05150 | CD630_s0631 | 156 – 166 | 324 – 334 | -10.9987 | 3.69875 | 4.95025 |
| LinCd | CD630_s0631 | 102 – 119 | 98 – 116 | -10.939 | 4.33624 | 4.95025 |
| CD630_27280 | CD630_s0631 | 141 – 200 | 12 – 73 | -10.9205 | 3.24738 | 4.95025 |
| mobA | CD630_s0631 | 5 – 40 | 56 – 94 | -10.8969 | 4.25703 | 4.95025 |
| CD630_15360 | CD630_s0631 | 159 – 178 | 98 – 116 | -10.8566 | 4.47208 | 4.95025 |
| CD630_14620 | CD630_s0631 | 157 – 189 | 87 – 115 | -10.8353 | 4.43651 | 4.95025 |
| CD630_13360 | CD630_s0631 | 171 – 200 | 312 – 338 | -10.7917 | 3.77330 | 4.95025 |
| eutV | CD630_s0631 | 159 – 184 | 57 – 84 | -10.7662 | 4.55715 | 4.95025 |
| CD630_19850 | CD630_s0631 | 134 – 151 | 322 – 339 | -10.7552 | 6.88486 | 4.95025 |
| etfA | CD630_s0631 | 44 – 65 | 312 – 341 | -10.7438 | 4.80521 | 4.95025 |
| CD630_28560 | CD630_s0631 | 82 – 92 | 57 – 68 | -10.7236 | 3.96803 | 4.95025 |
| CD630_06410 | CD630_s0631 | 171 – 187 | 88 – 105 | -10.5936 | 4.14404 | 4.95025 |
| CD630_13950 | CD630_s0631 | 186 – 198 | 323 – 335 | -10.5789 | 4.25540 | 4.95025 |
| CD630_04610 | CD630_s0631 | 146 – 160 | 327 – 340 | -10.5406 | 3.83025 | 4.95025 |
| CD630_08110 | CD630_s0631 | 163 – 192 | 19 – 49 | -10.5356 | 4.39181 | 4.95025 |
| CD630_18100 | CD630_s0631 | 181 – 198 | 99 – 115 | -10.4911 | 5.71658 | 4.95025 |
| CD630_06690 | CD630_s0631 | 178 – 200 | 43 – 69 | -10.4572 | 5.82709 | 4.95025 |
| CD630_13960 | CD630_s0631 | 51 – 63 | 328 – 340 | -10.3464 | 3.30375 | 4.95025 |
| CD630_27360 | CD630_s0631 | 184 – 191 | 61 – 68 | -10.3039 | 3.79570 | 4.95025 |
| CD630_28290 | CD630_s0631 | 38 – 149 | 10 – 111 | -10.2577 | 5.54192 | 4.95025 |
| CD630_33420 | CD630_s0631 | 157 – 168 | 58 – 69 | -10.2281 | 3.22095 | 4.95025 |
| CD630_18950 | CD630_s0631 | 179 – 188 | 59 – 68 | -10.2069 | 3.40610 | 4.95025 |
| CD630_31220 | CD630_s0631 | 119 – 168 | 99 – 164 | -10.1668 | 3.73779 | 4.95025 |
| bcp | CD630_s0631 | 91 – 123 | 102 – 137 | -10.1622 | 3.16670 | 4.95025 |
| CD630_27040 | CD630_s0631 | 79 – 101 | 287 – 309 | -10.1433 | 3.51812 | 4.95025 |
| CD630_33560 | CD630_s0631 | 181 – 198 | 159 – 175 | -10.1419 | 4.03147 | 4.95025 |
| CD630_26050 | CD630_s0631 | 191 – 200 | 329 – 338 | -10.1405 | 6.99851 | 4.95025 |
| CD630_27220 | CD630_s0631 | 109 – 134 | 90 – 115 | -10.1259 | 4.18786 | 4.95025 |
| CD630_04540 | CD630_s0631 | 83 – 99 | 248 – 266 | -10.1124 | 3.33556 | 4.95025 |
| CD630_12320 | CD630_s0631 | 170 – 190 | 98 – 111 | -10.0778 | 3.28667 | 4.95025 |
| cbiQ | CD630_s0631 | 74 – 96 | 251 – 272 | -10.0074 | 3.79830 | 4.95025 |
| CD630_09850 | CD630_s0631 | 167 – 194 | 316 – 339 | -9.96702 | 4.18679 | 4.95025 |
| CD630_33300 | CD630_s0631 | 131 – 147 | 248 – 263 | -9.93961 | 3.79149 | 4.95025 |
| CD630_10050 | CD630_s0631 | 13 – 52 | 289 – 341 | -9.9393 | 3.93750 | 4.95025 |
| CD630_05230 | CD630_s0631 | 1 – 20 | 250 – 268 | -9.93583 | 3.73787 | 4.95025 |
| CD630_05211 | CD630_s0631 | 114 – 131 | 319 – 336 | -9.91638 | 3.97048 | 4.95025 |
| CD630_06060 | CD630_s0631 | 164 – 187 | 6 – 31 | -9.89604 | 4.94050 | 4.95025 |
| panB | CD630_s0631 | 159 – 181 | 13 – 33 | -9.85052 | 3.40248 | 4.95025 |
| moaA | CD630_s0631 | 14 – 39 | 239 – 261 | -9.84985 | 3.83799 | 4.95025 |
| CD630_21610 | CD630_s0631 | 158 – 168 | 330 – 340 | -9.83595 | 3.16662 | 4.95025 |
| CD630_15890 | CD630_s0631 | 30 – 73 | 24 – 68 | -9.81983 | 3.39962 | 4.95025 |
| CD630_15160 | CD630_s0631 | 100 – 150 | 35 – 81 | -9.81603 | 4.32318 | 4.95025 |
| CD630_18010 | CD630_s0631 | 173 – 196 | 4 – 28 | -9.81437 | 4.21967 | 4.95025 |
| CD630_32890 | CD630_s0631 | 158 – 173 | 326 – 339 | -9.80534 | 3.41276 | 4.95025 |
| CD630_29760 | CD630_s0631 | 138 – 167 | 239 – 268 | -9.80231 | 3.51074 | 4.95025 |
| CD630_19680 | CD630_s0631 | 147 – 174 | 71 – 95 | -9.78525 | 3.58080 | 4.95025 |
| CD630_18780 | CD630_s0631 | 92 – 154 | 57 – 115 | -9.7636 | 4.71613 | 4.95025 |
| CD630_05000 | CD630_s0631 | 149 – 179 | 47 – 78 | -9.75363 | 4.02442 | 4.95025 |
| CD630_08130 | CD630_s0631 | 119 – 154 | 45 – 68 | -9.75187 | 4.63831 | 4.95025 |
| CD630_27230 | CD630_s0631 | 54 – 66 | 59 – 71 | -9.73918 | 3.52462 | 4.95025 |
| cls | CD630_s0631 | 190 – 201 | 293 – 304 | -9.72153 | 4.46177 | 4.95025 |
| CD630_31490 | CD630_s0631 | 41 – 60 | 224 – 243 | -9.71715 | 4.08593 | 4.95025 |
| CD630_10880 | CD630_s0631 | 160 – 188 | 56 – 84 | -9.71116 | 5.05048 | 4.95025 |
| CD630_28580 | CD630_s0631 | 151 – 173 | 317 – 337 | -9.70786 | 4.67130 | 4.95025 |
| CD630_22200 | CD630_s0631 | 102 – 136 | 305 – 337 | -9.70505 | 4.18288 | 4.95025 |
| CD630_06642 | CD630_s0631 | 35 – 48 | 326 – 338 | -9.68255 | 3.32967 | 4.95025 |
| CD630_30140 | CD630_s0631 | 50 – 120 | 64 – 123 | -9.67652 | 3.28343 | 4.95025 |
| CD630_25040 | CD630_s0631 | 53 – 73 | 326 – 342 | -9.67591 | 3.43898 | 4.95025 |
| CD630_31382 | CD630_s0631 | 171 – 185 | 327 – 342 | -9.67146 | 3.49137 | 4.95025 |
| CD630_36781 | CD630_s0631 | 140 – 157 | 321 – 335 | -9.66461 | 3.75946 | 4.95025 |
| CD630_05830 | CD630_s0631 | 114 – 129 | 123 – 138 | -9.64652 | 3.35360 | 4.95025 |
| CD630_36180 | CD630_s0631 | 169 – 181 | 158 – 170 | -9.61278 | 4.83428 | 4.95025 |
| CD630_28181 | CD630_s0631 | 44 – 77 | 247 – 267 | -9.60002 | 3.91299 | 4.95025 |
| CD630_26250 | CD630_s0631 | 134 – 162 | 171 – 199 | -9.52896 | 5.03379 | 4.95025 |
| spaK | CD630_s0631 | 155 – 199 | 14 – 68 | -9.52578 | 3.04530 | 4.95025 |
| CD630_24770 | CD630_s0631 | 156 – 194 | 39 – 82 | -9.52547 | 3.20798 | 4.95025 |
| folC | CD630_s0631 | 145 – 182 | 60 – 98 | -9.52484 | 3.85634 | 4.95025 |
| CD630_07870 | CD630_s0631 | 179 – 197 | 129 – 148 | -9.50699 | 3.22829 | 4.95025 |
| CD630_06100 | CD630_s0631 | 185 – 201 | 321 – 338 | -9.50195 | 4.43918 | 4.95025 |
| CD630_28870 | CD630_s0631 | 173 – 198 | 306 – 340 | -9.50136 | 3.41991 | 4.95025 |
| CD630_05950 | CD630_s0642 | 21 – 49 | 310 – 338 | -17.4948 | 3.31669 | 6.00600 |
| mobA | CD630_s0642 | 174 – 201 | 18 – 46 | -17.4553 | 4.25703 | 6.00600 |
| cbpA | CD630_s0642 | 78 – 129 | 111 – 161 | -16.9675 | 3.03040 | 6.00600 |
| CD630_17730 | CD630_s0642 | 162 – 181 | 147 – 167 | -15.8827 | 4.73397 | 6.00600 |
| fliI | CD630_s0642 | 177 – 199 | 134 – 154 | -15.7602 | 3.03666 | 6.00600 |
| CD630_31382 | CD630_s0642 | 11 – 55 | 10 – 48 | -15.7481 | 3.49137 | 6.00600 |
| CD630_06080 | CD630_s0642 | 126 – 175 | 309 – 354 | -15.5432 | 3.82315 | 6.00600 |
| CD630_10830 | CD630_s0642 | 120 – 201 | 231 – 317 | -15.3628 | 4.82398 | 6.00600 |
| CD630_12650 | CD630_s0642 | 102 – 124 | 236 – 258 | -15.2301 | 6.31827 | 6.00600 |
| CD630_07400 | CD630_s0642 | 152 – 174 | 10 – 33 | -15.2153 | 4.52273 | 6.00600 |
| CD630_21750 | CD630_s0642 | 151 – 193 | 7 – 54 | -15.1339 | 3.70626 | 6.00600 |
| appF | CD630_s0642 | 70 – 94 | 146 – 170 | -14.8509 | 4.25570 | 6.00600 |
| dapB2 | CD630_s0642 | 146 – 201 | 234 – 288 | -14.6419 | 3.27218 | 6.00600 |
| CD630_15200 | CD630_s0642 | 35 – 96 | 148 – 203 | -14.6365 | 4.00442 | 6.00600 |
| CD630_04850 | CD630_s0642 | 144 – 174 | 12 – 43 | -14.5508 | 4.40788 | 6.00600 |
| CD630_33560 | CD630_s0642 | 181 – 201 | 20 – 41 | -14.4523 | 4.03147 | 6.00600 |
| CD630_26050 | CD630_s0642 | 158 – 195 | 133 – 165 | -14.3398 | 6.99851 | 6.00600 |
| CD630_31000 | CD630_s0642 | 77 – 153 | 211 – 280 | -14.2341 | 3.70946 | 6.00600 |
| CD630_29750 | CD630_s0642 | 59 – 95 | 232 – 267 | -14.1264 | 4.59146 | 6.00600 |
| CD630_14920 | CD630_s0642 | 55 – 105 | 310 – 355 | -14.0533 | 4.35225 | 6.00600 |
| CD630_06060 | CD630_s0642 | 112 – 152 | 231 – 266 | -14.0366 | 4.94050 | 6.00600 |
| CD630_21240 | CD630_s0642 | 70 – 100 | 17 – 48 | -13.8741 | 5.88879 | 6.00600 |
| CD630_08090 | CD630_s0642 | 77 – 165 | 90 – 161 | -13.8239 | 3.57167 | 6.00600 |
| cdtR | CD630_s0642 | 15 – 55 | 388 – 420 | -13.7029 | 5.84368 | 6.00600 |
| pilT | CD630_s0642 | 129 – 157 | 20 – 48 | -13.6197 | 3.24639 | 6.00600 |
| CD630_09990 | CD630_s0642 | 120 – 159 | 147 – 192 | -13.5876 | 3.86040 | 6.00600 |
| CD630_02420 | CD630_s0642 | 120 – 168 | 119 – 180 | -13.5749 | 3.71048 | 6.00600 |
| CD630_05160 | CD630_s0642 | 150 – 194 | 10 – 46 | -13.5631 | 5.03573 | 6.00600 |
| CD630_11730 | CD630_s0642 | 147 – 163 | 301 – 317 | -13.4835 | 4.80521 | 6.00600 |
| CD630_21110 | CD630_s0642 | 149 – 159 | 145 – 155 | -13.4129 | 3.89630 | 6.00600 |
| garK | CD630_s0642 | 134 – 159 | 147 – 168 | -13.1795 | 3.72836 | 6.00600 |
| htpG | CD630_s0642 | 164 – 186 | 20 – 59 | -13.1683 | 3.45147 | 6.00600 |
| metQ | CD630_s0642 | 81 – 98 | 143 – 160 | -13.1444 | 3.87190 | 6.00600 |
| spoVFB | CD630_s0642 | 165 – 181 | 146 – 162 | -13.0982 | 4.62758 | 6.00600 |
| dpaA | CD630_s0642 | 117 – 147 | 13 – 46 | -13.0852 | 4.65364 | 6.00600 |
| cls | CD630_s0642 | 146 – 194 | 11 – 57 | -12.9865 | 4.46177 | 6.00600 |
| CD630_25020 | CD630_s0642 | 178 – 197 | 255 – 275 | -12.9591 | 4.17721 | 6.00600 |
| CD630_28320 | CD630_s0642 | 149 – 200 | 232 – 272 | -12.9198 | 3.99184 | 6.00600 |
| CD630_19030 | CD630_s0642 | 117 – 139 | 14 – 35 | -12.8548 | 3.34240 | 6.00600 |
| CD630_15740 | CD630_s0642 | 23 – 62 | 18 – 54 | -12.8043 | 4.43222 | 6.00600 |
| acoL | CD630_s0642 | 120 – 164 | 291 – 336 | -12.6771 | 3.32169 | 6.00600 |
| CD630_21540 | CD630_s0642 | 1 – 47 | 12 – 59 | -12.5988 | 4.33090 | 6.00600 |
| etfA | CD630_s0642 | 77 – 97 | 143 – 165 | -12.5912 | 4.80521 | 6.00600 |
| CD630_18781 | CD630_s0642 | 9 – 45 | 221 – 247 | -12.5459 | 3.77660 | 6.00600 |
| CD630_03410 | CD630_s0642 | 142 – 163 | 315 – 339 | -12.5115 | 4.45994 | 6.00600 |
| CD630_13360 | CD630_s0642 | 46 – 62 | 32 – 48 | -12.5091 | 3.77330 | 6.00600 |
| CD630_29760 | CD630_s0642 | 81 – 98 | 143 – 160 | -12.4419 | 3.51074 | 6.00600 |
| CD630_18930 | CD630_s0642 | 11 – 158 | 81 – 229 | -12.4395 | 5.32127 | 6.00600 |
| oppC | CD630_s0642 | 110 – 159 | 231 – 284 | -12.4178 | 3.01695 | 6.00600 |
| ssuC | CD630_s0642 | 28 – 74 | 231 – 284 | -12.4072 | 3.45491 | 6.00600 |
| CD630_21490 | CD630_s0642 | 97 – 119 | 133 – 165 | -12.3903 | 3.37462 | 6.00600 |
| CD630_35780 | CD630_s0642 | 148 – 195 | 92 – 143 | -12.3413 | 3.50107 | 6.00600 |
| CD630_07430 | CD630_s0642 | 38 – 72 | 9 – 42 | -12.3087 | 4.65568 | 6.00600 |
| CD630_13580 | CD630_s0642 | 24 – 42 | 30 – 48 | -12.2988 | 3.80715 | 6.00600 |
| CD630_31910 | CD630_s0642 | 16 – 65 | 231 – 281 | -12.2275 | 3.48314 | 6.00600 |
| CD630_25300 | CD630_s0642 | 102 – 128 | 13 – 37 | -12.213 | 4.49333 | 6.00600 |
| CD630_02740 | CD630_s0642 | 134 – 178 | 292 – 341 | -12.2037 | 5.02277 | 6.00600 |
| dltB | CD630_s0642 | 159 – 191 | 23 – 55 | -12.1015 | 3.81485 | 6.00600 |
| CD630_13930 | CD630_s0642 | 91 – 128 | 119 – 155 | -12.0821 | 4.09246 | 6.00600 |
| sigK | CD630_s0642 | 51 – 95 | 188 – 229 | -12.0507 | 3.08122 | 6.00600 |
| cdu2 | CD630_s0642 | 83 – 98 | 141 – 155 | -12.0406 | 2.97335 | 6.00600 |
| cspC | CD630_s0642 | 161 – 170 | 146 – 155 | -12.0233 | 5.48848 | 6.00600 |

| | | | | | | |
|---|---|---|---|---|---|---|
| CD630_27120 | CD630_s0642 | 109 – 154 | 231 – 278 | -12.0108 | 4.43720 | 6.00600 |
| bglA | CD630_s0642 | 154 – 184 | 141 – 172 | -11.9846 | 4.86402 | 6.00600 |
| CD630_18120 | CD630_s0642 | 43 – 96 | 230 – 278 | -11.9143 | 3.99713 | 6.00600 |
| CD630_13620 | CD630_s0642 | 178 – 198 | 222 – 248 | -11.9088 | 4.77627 | 6.00600 |
| CD630_13880 | CD630_s0642 | 193 – 201 | 17 – 25 | -11.8776 | 3.52735 | 6.00600 |
| CD630_09040 | CD630_s0642 | 152 – 200 | 317 – 357 | -11.8321 | 4.11851 | 6.00600 |
| xdhA | CD630_s0642 | 30 – 77 | 118 – 161 | -11.8187 | 4.45147 | 6.00600 |
| CD630_06050 | CD630_s0642 | 79 – 130 | 226 – 278 | -11.7641 | 3.59332 | 6.00600 |
| CD630_15431 | CD630_s0642 | 143 – 179 | 240 – 275 | -11.726 | 5.29780 | 6.00600 |
| CD630_15270 | CD630_s0642 | 166 – 199 | 7 – 41 | -11.7107 | 4.66015 | 6.00600 |
| CD630_19700 | CD630_s0642 | 152 – 185 | 197 – 235 | -11.6652 | 3.85861 | 6.00600 |
| CD630_31970 | CD630_s0642 | 121 – 134 | 229 – 243 | -11.5982 | 3.24511 | 6.00600 |
| CD630_06970 | CD630_s0642 | 64 – 91 | 23 – 52 | -11.5815 | 3.82312 | 6.00600 |
| folC | CD630_s0642 | 161 – 172 | 301 – 312 | -11.5246 | 3.85634 | 6.00600 |
| CD630_30130 | CD630_s0642 | 42 – 68 | 136 – 159 | -11.5201 | 3.28343 | 6.00600 |
| CD630_18610 | CD630_s0642 | 157 – 196 | 14 – 55 | -11.5056 | 4.15582 | 6.00600 |
| CD630_22070 | CD630_s0642 | 107 – 146 | 133 – 171 | -11.5013 | 4.09651 | 6.00600 |
| CD630_10800 | CD630_s0642 | 114 – 156 | 13 – 58 | -11.495 | 5.30319 | 6.00600 |
| CD630_23820 | CD630_s0642 | 148 – 187 | 134 – 172 | -11.4924 | 3.18419 | 6.00600 |
| CD630_06060 | CD630_s0642 | 132 – 139 | 146 – 153 | -11.4902 | 4.94050 | 6.00600 |
| CD630_34530 | CD630_s0642 | 158 – 182 | 33 – 58 | -11.4738 | 3.83016 | 6.00600 |
| CD630_17960 | CD630_s0642 | 137 – 146 | 146 – 155 | -11.46 | 3.71404 | 6.00600 |
| chrA | CD630_s0642 | 48 – 93 | 150 – 192 | -11.4502 | 5.27255 | 6.00600 |
| CD630_32880 | CD630_s0642 | 5 – 37 | 315 – 339 | -11.3677 | 3.41276 | 6.00600 |
| CD630_05880 | CD630_s0642 | 39 – 102 | 230 – 285 | -11.3203 | 4.28339 | 6.00600 |
| CD630_05150 | CD630_s0642 | 151 – 190 | 10 – 45 | -11.3007 | 3.69875 | 6.00600 |
| CD630_27110 | CD630_s0642 | 148 – 174 | 317 – 339 | -11.2977 | 3.78231 | 6.00600 |
| CD630_30100 | CD630_s0642 | 132 – 162 | 278 – 308 | -11.2876 | 4.13234 | 6.00600 |
| oppB | CD630_s0642 | 169 – 192 | 17 – 40 | -11.2821 | 2.98328 | 6.00600 |
| CD630_27230 | CD630_s0642 | 92 – 147 | 234 – 283 | -11.2788 | 3.52462 | 6.00600 |
| CD630_21910 | CD630_s0642 | 59 – 95 | 146 – 180 | -11.259 | 4.66600 | 6.00600 |
| CD630_25620 | CD630_s0642 | 7 – 51 | 241 – 283 | -11.2112 | 3.02685 | 6.00600 |
| CD630_09770 | CD630_s0642 | 145 – 169 | 13 – 35 | -11.2034 | 3.22106 | 6.00600 |
| CD630_29960 | CD630_s0642 | 181 – 189 | 147 – 155 | -11.1246 | 4.51904 | 6.00600 |
| CD630_27570 | CD630_s0642 | 110 – 128 | 146 – 167 | -11.0762 | 3.46216 | 6.00600 |
| CD630_24920 | CD630_s0642 | 117 – 155 | 136 – 173 | -11.0501 | 3.89455 | 6.00600 |
| kdgT1 | CD630_s0642 | 68 – 86 | 321 – 340 | -11.0453 | 4.49207 | 6.00600 |
| CD630_12320 | CD630_s0642 | 135 – 174 | 17 – 55 | -11.0046 | 3.28667 | 6.00600 |
| CD630_28550 | CD630_s0642 | 86 – 119 | 140 – 171 | -10.9924 | 3.51196 | 6.00600 |
| CD630_26750 | CD630_s0642 | 164 – 179 | 17 – 35 | -10.9836 | 3.60610 | 6.00600 |
| cprK | CD630_s0642 | 84 – 97 | 144 – 156 | -10.9811 | 3.96216 | 6.00600 |
| CD630_03220 | CD630_s0642 | 157 – 176 | 297 – 313 | -10.9701 | 3.36746 | 6.00600 |
| CD630_06340 | CD630_s0642 | 106 – 137 | 12 – 49 | -10.9507 | 4.35983 | 6.00600 |
| uxaA | CD630_s0642 | 172 – 200 | 133 – 161 | -10.9275 | 4.26165 | 6.00600 |
| CD630_29840 | CD630_s0642 | 27 – 42 | 144 – 158 | -10.9111 | 4.48825 | 6.00600 |
| CD630_20960 | CD630_s0642 | 73 – 112 | 149 – 191 | -10.8384 | 5.65940 | 6.00600 |
| CD630_21090 | CD630_s0642 | 124 – 177 | 3 – 54 | -10.8043 | 4.15803 | 6.00600 |
| CD630_23000 | CD630_s0642 | 41 – 65 | 231 – 263 | -10.7964 | 3.54130 | 6.00600 |
| CD630_36781 | CD630_s0642 | 176 – 194 | 141 – 160 | -10.7949 | 3.75946 | 6.00600 |
| CD630_14620 | CD630_s0642 | 23 – 63 | 40 – 82 | -10.7873 | 4.43651 | 6.00600 |
| nusB | CD630_s0642 | 116 – 137 | 133 – 154 | -10.7702 | 4.92798 | 6.00600 |
| pilC | CD630_s0642 | 90 – 146 | 224 – 268 | -10.7568 | 3.54230 | 6.00600 |
| CD630_16530 | CD630_s0642 | 92 – 169 | 17 – 92 | -10.7479 | 4.37658 | 6.00600 |
| ldh | CD630_s0642 | 68 – 94 | 143 – 171 | -10.7426 | 4.35818 | 6.00600 |
| CD630_06920 | CD630_s0642 | 188 – 196 | 147 – 155 | -10.7119 | 3.13260 | 6.00600 |
| CD630_15890 | CD630_s0642 | 150 – 180 | 148 – 183 | -10.6547 | 3.39962 | 6.00600 |
| CD630_03920 | CD630_s0642 | 24 – 38 | 18 – 32 | -10.5882 | 4.15934 | 6.00600 |
| CD630_08950 | CD630_s0642 | 42 – 76 | 188 – 234 | -10.5858 | 3.71905 | 6.00600 |
| CD630_05030 | CD630_s0642 | 130 – 165 | 19 – 42 | -10.5304 | 4.03823 | 6.00600 |
| CD630_10220 | CD630_s0642 | 82 – 98 | 147 – 162 | -10.5234 | 4.08233 | 6.00600 |
| CD630_22550 | CD630_s0642 | 31 – 74 | 2 – 53 | -10.4919 | 3.62379 | 6.00600 |
| CD630_25320 | CD630_s0642 | 161 – 177 | 317 – 333 | -10.4825 | 4.29415 | 6.00600 |
| CD630_14210 | CD630_s0642 | 136 – 148 | 144 – 155 | -10.4579 | 3.82355 | 6.00600 |
| argS | CD630_s0642 | 52 – 67 | 134 – 149 | -10.457 | 4.96036 | 6.00600 |
| srlEb | CD630_s0642 | 69 – 98 | 133 – 165 | -10.4419 | 3.50140 | 6.00600 |
| CD630_21020 | CD630_s0642 | 86 – 104 | 136 – 155 | -10.4305 | 4.28666 | 6.00600 |
| CD630_15360 | CD630_s0642 | 141 – 169 | 317 – 345 | -10.4072 | 4.47208 | 6.00600 |
| CD630_18660 | CD630_s0642 | 126 – 142 | 302 – 316 | -10.4022 | 3.50371 | 6.00600 |
| CD630_17710 | CD630_s0642 | 40 – 55 | 387 – 402 | -10.3887 | 4.37688 | 6.00600 |
| CD630_28260 | CD630_s0642 | 83 – 101 | 141 – 159 | -10.3751 | 3.51613 | 6.00600 |
| CD630_06060 | CD630_s0642 | 33 – 45 | 147 – 160 | -10.3589 | 4.94050 | 6.00600 |
| CD630_05230 | CD630_s0642 | 1 – 30 | 3 – 31 | -10.3483 | 3.73787 | 6.00600 |
| CD630_24590 | CD630_s0642 | 179 – 189 | 144 – 153 | -10.3443 | 3.70401 | 6.00600 |
| CD630_05460 | CD630_s0642 | 102 – 117 | 23 – 37 | -10.3402 | 3.98753 | 6.00600 |
| CD630_21140 | CD630_s0642 | 173 – 197 | 133 – 155 | -10.3378 | 3.33304 | 6.00600 |
| CD630_27220 | CD630_s0642 | 141 – 186 | 6 – 44 | -10.3368 | 4.18786 | 6.00600 |
| CD630_27520 | CD630_s0642 | 84 – 98 | 143 – 157 | -10.3136 | 4.27693 | 6.00600 |
| CD630_07760 | CD630_s0642 | 45 – 91 | 232 – 284 | -10.27 | 3.64186 | 6.00600 |
| feoB | CD630_s0642 | 1 – 39 | 133 – 161 | -10.2624 | 4.05364 | 6.00600 |
| CD630_09750 | CD630_s0642 | 11 – 39 | 167 – 191 | -10.2252 | 4.83410 | 6.00600 |
| CD630_28920 | CD630_s0642 | 11 – 39 | 167 – 191 | -10.2252 | 4.51664 | 6.00600 |
| moaA | CD630_s0642 | 54 – 111 | 6 – 48 | -10.2061 | 3.83799 | 6.00600 |
| CD630_25190 | CD630_s0642 | 154 – 200 | 317 – 355 | -10.1989 | 3.19663 | 6.00600 |
| CD630_18090 | CD630_s0642 | 174 – 194 | 17 – 34 | -10.1988 | 4.23961 | 6.00600 |
| CD630_18650 | CD630_s0642 | 11 – 23 | 317 – 329 | -10.192 | 3.75892 | 6.00600 |
| CD630_29650 | CD630_s0642 | 151 – 192 | 20 – 54 | -10.1901 | 3.67128 | 6.00600 |
| CD630_14190 | CD630_s0642 | 14 – 32 | 200 – 218 | -10.1851 | 3.78562 | 6.00600 |
| CD630_03930 | CD630_s0642 | 137 – 184 | 33 – 89 | -10.179 | 4.15934 | 6.00600 |
| CD630_05220 | CD630_s0642 | 169 – 201 | 22 – 54 | -10.1728 | 3.65057 | 6.00600 |
| CD630_32050 | CD630_s0642 | 87 – 129 | 21 – 61 | -10.1646 | 5.73797 | 6.00600 |
| CD630_05830 | CD630_s0642 | 31 – 63 | 19 – 54 | -10.1608 | 3.35360 | 6.00600 |
| CD630_13350 | CD630_s0642 | 165 – 183 | 317 – 339 | -10.1587 | 4.15139 | 6.00600 |
| CD630_31530 | CD630_s0642 | 1 – 37 | 241 – 279 | -10.1496 | 3.49100 | 6.00600 |
| CD630_16750 | CD630_s0642 | 164 – 201 | 10 – 38 | -10.1437 | 5.04051 | 6.00600 |
| CD630_25200 | CD630_s0642 | 2 – 10 | 149 – 157 | -10.1432 | 3.65158 | 6.00600 |
| CD630_05820 | CD630_s0642 | 150 – 197 | 26 – 63 | -10.1374 | 3.75661 | 6.00600 |
| CD630_01590 | CD630_s0642 | 154 – 200 | 146 – 193 | -10.1135 | 3.45609 | 6.00600 |
| CD630_19790 | CD630_s0642 | 50 – 97 | 139 – 178 | -10.1121 | 3.79593 | 6.00600 |
| bglA | CD630_s0642 | 89 – 97 | 147 – 155 | -10.1062 | 4.50315 | 6.00600 |
| bcp | CD630_s0642 | 182 – 197 | 232 – 247 | -10.102 | 3.16670 | 6.00600 |
| CD630_13950 | CD630_s0642 | 24 – 42 | 318 – 337 | -10.0993 | 4.25540 | 6.00600 |
| CD630_13780 | CD630_s0642 | 124 – 171 | 302 – 344 | -10.0942 | 3.52274 | 6.00600 |
| CD630_18960 | CD630_s0642 | 28 – 53 | 147 – 171 | -10.0923 | 4.37822 | 6.00600 |
| CD630_15461 | CD630_s0642 | 1 – 25 | 28 – 49 | -10.0848 | 9.13442 | 6.00600 |
| CD630_33600 | CD630_s0642 | 7 – 39 | 136 – 167 | -10.082 | 3.73196 | 6.00600 |
| CD630_36250 | CD630_s0642 | 53 – 61 | 143 – 151 | -10.0535 | 4.13224 | 6.00600 |
| CD630_36690 | CD630_s0642 | 120 – 158 | 148 – 176 | -10.0519 | 3.53761 | 6.00600 |
| CD630_05450 | CD630_s0642 | 142 – 180 | 147 – 174 | -10.047 | 5.04761 | 6.00600 |
| CD630_32710 | CD630_s0642 | 131 – 152 | 225 – 247 | -10.0373 | 4.74276 | 6.00600 |
| CD630_05900 | CD630_s0642 | 10 – 37 | 147 – 165 | -10.02 | 3.21934 | 6.00600 |
| CD630_03170 | CD630_s0642 | 150 – 173 | 157 – 184 | -10.008 | 3.66946 | 6.00600 |
| CD630_24770 | CD630_s0642 | 47 – 94 | 232 – 275 | -9.98762 | 3.20798 | 6.00600 |
| iorA | CD630_s0642 | 59 – 74 | 298 – 314 | -9.9452 | 3.83173 | 6.00600 |
| CD630_05870 | CD630_s0642 | 161 – 201 | 228 – 274 | -9.93477 | 4.28339 | 6.00600 |
| CD630_26630 | CD630_s0642 | 80 – 102 | 135 – 160 | -9.91944 | 4.61557 | 6.00600 |
| CD630_28181 | CD630_s0642 | 149 – 172 | 10 – 35 | -9.91827 | 3.91299 | 6.00600 |
| CD630_14700 | CD630_s0642 | 22 – 68 | 230 – 274 | -9.91589 | 3.98234 | 6.00600 |
| CD630_17281 | CD630_s0642 | 14 – 42 | 45 – 73 | -9.91301 | 5.91544 | 6.00600 |
| CD630_16670 | CD630_s0642 | 98 – 139 | 12 – 42 | -9.8972 | 3.33553 | 6.00600 |
| cdu1 | CD630_s0642 | 79 – 95 | 143 – 157 | -9.88978 | 4.46276 | 6.00600 |
| CD630_15150 | CD630_s0642 | 59 – 108 | 133 – 171 | -9.87954 | 3.35564 | 6.00600 |
| CD630_31220 | CD630_s0642 | 48 – 72 | 313 – 337 | -9.87767 | 3.73779 | 6.00600 |
| CD630_32520 | CD630_s0642 | 132 – 176 | 228 – 277 | -9.86621 | 4.06098 | 6.00600 |
| CD630_14990 | CD630_s0642 | 119 – 164 | 308 – 347 | -9.86486 | 3.78619 | 6.00600 |
| CD630_17170 | CD630_s0642 | 163 – 173 | 146 – 156 | -9.84248 | 4.10230 | 6.00600 |
| CD630_31960 | CD630_s0642 | 17 – 34 | 144 – 161 | -9.84062 | 4.93757 | 6.00600 |
| CD630_23760 | CD630_s0642 | 54 – 61 | 148 – 155 | -9.82717 | 4.54307 | 6.00600 |
| CD630_35260 | CD630_s0642 | 137 – 148 | 136 – 147 | -9.81637 | 5.77277 | 6.00600 |
| CD630_21570 | CD630_s0642 | 79 – 118 | 132 – 161 | -9.81528 | 6.48385 | 6.00600 |
| CD630_06160 | CD630_s0642 | 184 – 198 | 323 – 339 | -9.79982 | 5.51744 | 6.00600 |
| CD630_23861 | CD630_s0642 | 9 – 42 | 10 – 48 | -9.79705 | 5.83068 | 6.00600 |
| CD630_36170 | CD630_s0642 | 33 – 69 | 19 – 58 | -9.79415 | 3.72380 | 6.00600 |
| CD630_27280 | CD630_s0642 | 171 – 191 | 212 – 230 | -9.78982 | 3.24738 | 6.00600 |
| CD630_15850 | CD630_s0642 | 56 – 83 | 6 – 38 | -9.76092 | 4.32968 | 6.00600 |
| CD630_20670 | CD630_s0642 | 151 – 194 | 12 – 62 | -9.76062 | 4.42757 | 6.00600 |
| terD | CD630_s0642 | 104 – 132 | 145 – 171 | -9.72837 | 3.33148 | 6.00600 |
| CD630_09570 | CD630_s0642 | 138 – 164 | 140 – 166 | -9.72626 | 5.31213 | 6.00600 |
| CD630_29070 | CD630_s0642 | 138 – 164 | 140 – 166 | -9.72626 | 3.89477 | 6.00600 |
| CD630_31461 | CD630_s0642 | 121 – 141 | 12 – 32 | -9.67931 | 4.41476 | 6.00600 |
| CD630_21760 | CD630_s0642 | 49 – 84 | 19 – 54 | -9.66208 | 4.43846 | 6.00600 |
| CD630_30190 | CD630_s0642 | 91 – 98 | 147 – 154 | -9.64405 | 3.27394 | 6.00600 |
| CD630_22740 | CD630_s0642 | 65 – 94 | 140 – 172 | -9.64356 | 3.08733 | 6.00600 |
| CD630_09030 | CD630_s0642 | 45 – 66 | 386 – 409 | -9.63415 | 4.21282 | 6.00600 |
| CD630_10000 | CD630_s0642 | 78 – 86 | 28 – 36 | -9.63389 | 3.86040 | 6.00600 |
| CD630_01560 | CD630_s0642 | 135 – 167 | 307 – 333 | -9.62996 | 4.06352 | 6.00600 |
| pmi | CD630_s0642 | 1 – 37 | 4 – 37 | -9.60934 | 3.86845 | 6.00600 |
| CD630_19401 | CD630_s0642 | 33 – 46 | 152 – 165 | -9.58178 | 4.37084 | 6.00600 |
| CD630_04341 | CD630_s0642 | 24 – 75 | 141 – 180 | -9.53077 | 4.29262 | 6.00600 |
| CD630_17511 | CD630_s0660 | 6 – 60 | 3 – 59 | -16.7174 | 9.12322 | 4.92199 |
| CD630_05460 | CD630_s0660 | 88 – 109 | 208 – 232 | -15.731 | 3.98753 | 4.92199 |
| CD630_04341 | CD630_s0660 | 51 – 75 | 223 – 244 | -15.1336 | 4.29262 | 4.92199 |
| CD630_05030 | CD630_s0660 | 132 – 177 | 46 – 89 | -14.6819 | 4.03823 | 4.92199 |
| CD630_20450 | CD630_s0660 | 102 – 141 | 21 – 62 | -14.396 | 3.77388 | 4.92199 |
| CD630_21620 | CD630_s0660 | 4 – 43 | 19 – 55 | -14.1366 | 3.02649 | 4.92199 |
| CD630_36180 | CD630_s0660 | 169 – 200 | 8 – 39 | -14.0148 | 4.83428 | 4.92199 |
| CD630_23820 | CD630_s0660 | 160 – 200 | 15 – 56 | -13.8586 | 3.18419 | 4.92199 |
| serS1 | CD630_s0660 | 161 – 168 | 225 – 232 | -13.5702 | 3.45858 | 4.92199 |
| CD630_18100 | CD630_s0660 | 87 – 122 | 207 – 232 | -13.0939 | 5.71658 | 4.92199 |
| CD630_25320 | CD630_s0660 | 125 – 177 | 20 – 87 | -13.0252 | 4.29415 | 4.92199 |
| CD630_21240 | CD630_s0660 | 83 – 94 | 225 – 236 | -12.7946 | 5.88879 | 4.92199 |
| CD630_06060 | CD630_s0660 | 138 – 150 | 227 – 242 | -12.7764 | 4.94050 | 4.92199 |
| CD630_09100 | CD630_s0660 | 5 – 56 | 1 – 48 | -12.6597 | 5.92260 | 4.92199 |
| CD630_23861 | CD630_s0660 | 83 – 96 | 222 – 236 | -12.546 | 5.83068 | 4.92199 |
| CD630_08390 | CD630_s0660 | 70 – 89 | 225 – 239 | -12.3817 | 5.16000 | 4.92199 |
| CD630_31461 | CD630_s0660 | 133 – 142 | 46 – 55 | -12.2187 | 4.41476 | 4.92199 |
| CD630_19700 | CD630_s0660 | 81 – 90 | 228 – 237 | -12.1359 | 3.85861 | 4.92199 |
| CD630_08110 | CD630_s0660 | 88 – 110 | 207 – 228 | -12.0401 | 4.39181 | 4.92199 |
| CD630_16460 | CD630_s0660 | 1 – 69 | 10 – 76 | -11.9292 | 4.64028 | 4.92199 |
| CD630_13780 | CD630_s0660 | 85 – 117 | 140 – 175 | -11.8709 | 3.52274 | 4.92199 |
| CD630_08090 | CD630_s0660 | 84 – 92 | 224 – 232 | -11.8557 | 3.57167 | 4.92199 |
| CD630_21570 | CD630_s0660 | 84 – 97 | 220 – 232 | -11.816 | 6.48385 | 4.92199 |
| CD630_34520 | CD630_s0660 | 125 – 166 | 80 – 128 | -11.4656 | 3.83016 | 4.92199 |
| kdpD | CD630_s0660 | 109 – 139 | 19 – 48 | -11.3792 | 4.27140 | 4.92199 |
| CD630_32890 | CD630_s0660 | 158 – 176 | 107 – 128 | -11.3673 | 3.41276 | 4.92199 |
| gcp | CD630_s0660 | 132 – 151 | 46 – 63 | -11.3268 | 4.92798 | 4.92199 |
| oppB | CD630_s0660 | 89 – 101 | 221 – 232 | -11.3196 | 2.98328 | 4.92199 |
| CD630_14260 | CD630_s0660 | 81 – 110 | 19 – 46 | -11.2704 | 3.22214 | 4.92199 |
| CD630_06190 | CD630_s0660 | 13 – 40 | 19 – 48 | -11.1235 | 3.57064 | 4.92199 |
| CD630_06170 | CD630_s0660 | 137 – 154 | 32 – 49 | -11.019 | 4.95366 | 4.92199 |
| thiH | CD630_s0660 | 189 – 201 | 21 – 33 | -10.9834 | 3.58230 | 4.92199 |
| CD630_24591 | CD630_s0660 | 88 – 95 | 225 – 232 | -10.9593 | 3.54497 | 4.92199 |
| CD630_08940 | CD630_s0660 | 154 – 169 | 76 – 91 | -10.9095 | 3.72225 | 4.92199 |
| CD630_28580 | CD630_s0660 | 39 – 54 | 225 – 239 | -10.8198 | 4.67130 | 4.92199 |
| feoB | CD630_s0660 | 30 – 78 | 10 – 55 | -10.7241 | 4.52257 | 4.92199 |
| CD630_09050 | CD630_s0660 | 177 – 187 | 108 – 118 | -10.7235 | 3.96216 | 4.92199 |
| cstA | CD630_s0660 | 83 – 106 | 211 – 237 | -10.7014 | 3.53584 | 4.92199 |
| CD630_24770 | CD630_s0660 | 158 – 191 | 117 – 147 | -10.6568 | 3.20798 | 4.92199 |
| CD630_31960 | CD630_s0660 | 25 – 71 | 40 – 86 | -10.6202 | 4.93757 | 4.92199 |
| cbpA | CD630_s0660 | 85 – 103 | 214 – 232 | -10.5588 | 3.03040 | 4.92199 |
| eutV | CD630_s0660 | 98 – 106 | 229 – 237 | -10.5433 | 4.55715 | 4.92199 |
| CD630_05590 | CD630_s0660 | 93 – 122 | 207 – 232 | -10.5081 | 3.39670 | 4.92199 |
| CD630_10880 | CD630_s0660 | 129 – 160 | 22 – 58 | -10.4979 | 5.05048 | 4.92199 |
| dapB2 | CD630_s0660 | 117 – 156 | 107 – 147 | -10.4963 | 3.27218 | 4.92199 |
| flgB | CD630_s0660 | 91 – 99 | 224 – 232 | -10.4127 | 3.25736 | 4.92199 |
| CD630_31490 | CD630_s0660 | 128 – 162 | 80 – 117 | -10.3936 | 4.08593 | 4.92199 |
| CD630_03410 | CD630_s0660 | 121 – 157 | 33 – 71 | -10.3817 | 4.45994 | 4.92199 |
| gcvTPA | CD630_s0660 | 144 – 190 | 15 – 55 | -10.2788 | 4.77229 | 4.92199 |
| CD630_17720 | CD630_s0660 | 8 – 31 | 33 – 55 | -10.2624 | 3.22910 | 4.92199 |
| opuCA | CD630_s0660 | 169 – 191 | 141 – 170 | -10.2029 | 4.07035 | 4.92199 |
| spaK | CD630_s0660 | 160 – 200 | 32 – 74 | -10.1857 | 3.04530 | 4.92199 |
| CD630_05900 | CD630_s0660 | 4 – 47 | 17 – 60 | -10.1725 | 3.21934 | 4.92199 |
| CD630_13790 | CD630_s0660 | 89 – 113 | 144 – 174 | -10.1524 | 5.17374 | 4.92199 |
| CD630_08780 | CD630_s0660 | 83 – 114 | 207 – 235 | -10.1251 | 7.76690 | 4.92199 |
| CD630_33610 | CD630_s0660 | 134 – 192 | 32 – 86 | -10.0405 | 3.79322 | 4.92199 |
| CD630_18311 | CD630_s0660 | 164 – 185 | 36 – 55 | -9.96416 | 6.86047 | 4.92199 |
| CD630_31580 | CD630_s0660 | 143 – 154 | 20 – 31 | -9.95939 | 3.51756 | 4.92199 |
| CD630_05821 | CD630_s0660 | 10 – 30 | 106 – 128 | -9.94333 | 3.75661 | 4.92199 |
| CD630_17730 | CD630_s0660 | 67 – 94 | 223 – 243 | -9.93692 | 4.73397 | 4.92199 |
| CD630_07760 | CD630_s0660 | 78 – 111 | 207 – 244 | -9.9171 | 3.64186 | 4.92199 |
| CD630_30140 | CD630_s0660 | 90 – 102 | 162 – 174 | -9.89869 | 3.28343 | 4.92199 |
| CD630_09030 | CD630_s0660 | 35 – 48 | 210 – 223 | -9.83692 | 4.21282 | 4.92199 |
| CD630_32700 | CD630_s0660 | 101 – 126 | 222 – 244 | -9.8321 | 4.74276 | 4.92199 |
| CD630_22550 | CD630_s0660 | 171 – 182 | 206 – 217 | -9.81634 | 3.62379 | 4.92199 |
| dltB | CD630_s0660 | 16 – 36 | 69 – 89 | -9.78322 | 3.81485 | 4.92199 |
| CD630_10050 | CD630_s0660 | 139 – 183 | 140 – 178 | -9.77162 | 3.93750 | 4.92199 |
| eutW | CD630_s0660 | 1 – 29 | 54 – 85 | -9.76254 | 4.55715 | 4.92199 |
| CD630_14710 | CD630_s0660 | 86 – 97 | 223 – 233 | -9.74902 | 3.65171 | 4.92199 |
| CD630_20460 | CD630_s0660 | 150 – 170 | 72 – 89 | -9.73604 | 7.94034 | 4.92199 |
| CD630_18930 | CD630_s0660 | 146 – 167 | 134 – 154 | -9.71593 | 5.32127 | 4.92199 |
| CD630_13020 | CD630_s0660 | 117 – 160 | 34 – 80 | -9.71168 | 3.52267 | 4.92199 |
| CD630_20170 | CD630_s0660 | 187 – 198 | 51 – 62 | -9.69613 | 6.90201 | 4.92199 |
| CD630_06050 | CD630_s0660 | 48 – 59 | 20 – 31 | -9.64766 | 3.59332 | 4.92199 |
| CD630_28330 | CD630_s0660 | 81 – 112 | 207 – 239 | -9.62688 | 3.29122 | 4.92199 |
| iorA | CD630_s0660 | 58 – 69 | 134 – 145 | -9.60096 | 3.83173 | 4.92199 |
| CD630_20670 | CD630_s0660 | 78 – 102 | 219 – 238 | -9.59039 | 4.42757 | 4.92199 |
| CD630_14130 | CD630_s0660 | 140 – 180 | 186 – 228 | -9.56084 | 4.57970 | 4.92199 |
| CD630_04610 | CD630_s0660 | 108 – 121 | 77 – 89 | -9.5455 | 3.83025 | 4.92199 |
| CD630_05830 | CD630_s0660 | 26 – 66 | 42 – 85 | -9.52953 | 3.35360 | 4.92199 |
| CD630_21700 | CD630_s0660 | 31 – 47 | 37 – 54 | -9.52826 | 3.96650 | 4.92199 |
| CD630_05822 | CD630_s0660 | 126 – 139 | 72 – 84 | -9.52799 | 4.28113 | 4.92199 |
| CD630_24590 | CD630_s0660 | 119 – 129 | 228 – 237 | -9.51934 | 3.70401 | 4.92199 |
| CD630_06080 | CD630_s0660 | 21 – 31 | 21 – 31 | -9.51195 | 3.82315 | 4.92199 |
| CD630_19401 | CD630_s0670 | 7 – 45 | 79 – 105 | -18.9929 | 4.37084 | 4.56362 |
| CD630_04880 | CD630_s0670 | 47 – 91 | 63 – 105 | -15.0125 | 4.62291 | 4.56362 |
| CD630_25320 | CD630_s0670 | 126 – 168 | 43 – 82 | -14.6976 | 4.29415 | 4.56362 |
| CD630_32520 | CD630_s0670 | 3 – 33 | 157 – 192 | -14.4753 | 4.06098 | 4.56362 |
| CD630_08100 | CD630_s0670 | 146 – 162 | 39 – 55 | -14.3317 | 4.39181 | 4.56362 |
| CD630_21260 | CD630_s0670 | 128 – 143 | 28 – 43 | -13.0738 | 4.22255 | 4.56362 |

| | | | | | | |
|---|---|---|---|---|---|---|
| rbsA | CD630_s0670 | 168 – 192 | 90 – 113 | -13.0414 | 3.71650 | 4.56362 |
| CD630_09100 | CD630_s0670 | 39 – 83 | 154 – 193 | -13.037 | 5.92260 | 4.56362 |
| CD630_10880 | CD630_s0670 | 141 – 183 | 40 – 76 | -12.5713 | 5.05048 | 4.56362 |
| CD630_18090 | CD630_s0670 | 185 – 196 | 176 – 187 | -12.554 | 4.23961 | 4.56362 |
| acoL | CD630_s0670 | 127 – 162 | 50 – 84 | -12.3297 | 3.32169 | 4.56362 |
| CD630_16330 | CD630_s0670 | 44 – 65 | 92 – 113 | -12.2619 | 4.11180 | 4.56362 |
| oppB | CD630_s0670 | 88 – 119 | 45 – 68 | -12.2584 | 2.98328 | 4.56362 |
| CD630_24590 | CD630_s0670 | 142 – 167 | 52 – 84 | -12.2301 | 3.70401 | 4.56362 |
| CD630_02430 | CD630_s0670 | 68 – 80 | 49 – 61 | -12.2207 | 3.71048 | 4.56362 |
| CD630_19680 | CD630_s0670 | 141 – 176 | 48 – 90 | -12.0548 | 3.58080 | 4.56362 |
| CD630_16670 | CD630_s0670 | 120 – 129 | 79 – 88 | -11.7615 | 3.33553 | 4.56362 |
| CD630_07480 | CD630_s0670 | 89 – 106 | 43 – 60 | -11.5678 | 5.18321 | 4.56362 |
| CD630_27570 | CD630_s0670 | 171 – 198 | 43 – 68 | -11.5633 | 3.46216 | 4.56362 |
| CD630_31010 | CD630_s0670 | 146 – 196 | 35 – 101 | -11.5275 | 4.96305 | 4.56362 |
| xdhA | CD630_s0670 | 16 – 71 | 41 – 100 | -11.4442 | 4.45147 | 4.56362 |
| CD630_13790 | CD630_s0670 | 164 – 185 | 165 – 186 | -11.1161 | 5.17374 | 4.56362 |
| CD630_24220 | CD630_s0670 | 164 – 179 | 48 – 62 | -11.0652 | 4.18037 | 4.56362 |
| CD630_27240 | CD630_s0670 | 142 – 161 | 176 – 193 | -11.0498 | 3.52462 | 4.56362 |
| opuCC | CD630_s0670 | 184 – 200 | 150 – 166 | -10.88 | 4.07035 | 4.56362 |
| CD630_17511 | CD630_s0670 | 189 – 197 | 181 – 189 | -10.829 | 9.12322 | 4.56362 |
| CD630_28560 | CD630_s0670 | 113 – 130 | 173 – 190 | -10.7979 | 3.96803 | 4.56362 |
| CD630_21530 | CD630_s0670 | 62 – 113 | 60 – 111 | -10.7482 | 3.39774 | 4.56362 |
| CD630_05821 | CD630_s0670 | 78 – 86 | 79 – 87 | -10.7269 | 3.75661 | 4.56362 |
| CD630_21090 | CD630_s0670 | 145 – 172 | 22 – 45 | -10.6902 | 4.15803 | 4.56362 |
| CD630_29250 | CD630_s0670 | 176 – 194 | 12 – 31 | -10.6757 | 4.61901 | 4.56362 |
| CD630_09410 | CD630_s0670 | 176 – 194 | 12 – 31 | -10.6757 | 4.75651 | 4.56362 |
| CD630_32690 | CD630_s0670 | 88 – 111 | 45 – 67 | -10.6456 | 7.32676 | 4.56362 |
| bcp | CD630_s0670 | 103 – 118 | 49 – 63 | -10.5909 | 3.16670 | 4.56362 |
| CD630_32720 | CD630_s0670 | 182 – 200 | 176 – 190 | -10.5592 | 4.74276 | 4.56362 |
| CD630_05211 | CD630_s0670 | 182 – 200 | 42 – 63 | -10.5413 | 3.97048 | 4.56362 |
| CD630_14630 | CD630_s0670 | 188 – 200 | 48 – 60 | -10.4655 | 3.34485 | 4.56362 |
| CD630_20170 | CD630_s0670 | 185 – 198 | 37 – 49 | -10.4544 | 6.90201 | 4.56362 |
| CD630_18910 | CD630_s0670 | 175 – 198 | 48 – 68 | -10.4083 | 3.30688 | 4.56362 |
| CD630_06170 | CD630_s0670 | 64 – 79 | 177 – 192 | -10.3351 | 4.95366 | 4.56362 |
| CD630_18010 | CD630_s0670 | 174 – 186 | 24 – 36 | -10.3228 | 4.21967 | 4.56362 |
| CD630_06100 | CD630_s0670 | 91 – 124 | 23 – 60 | -10.2736 | 4.43918 | 4.56362 |
| CD630_15060 | CD630_s0670 | 4 – 22 | 89 – 108 | -10.23 | 3.69396 | 4.56362 |
| thiH | CD630_s0670 | 189 – 199 | 63 – 73 | -10.0799 | 3.58230 | 4.56362 |
| CD630_20670 | CD630_s0670 | 110 – 136 | 43 – 68 | -10.0294 | 4.42757 | 4.56362 |
| CD630_08660 | CD630_s0670 | 170 – 200 | 165 – 192 | -9.91771 | 3.86467 | 4.56362 |
| CD630_21760 | CD630_s0670 | 183 – 199 | 151 – 167 | -9.86749 | 4.43846 | 4.56362 |
| CD630_04341 | CD630_s0670 | 46 – 59 | 40 – 53 | -9.8674 | 4.29262 | 4.56362 |
| srlD | CD630_s0670 | 108 – 133 | 78 – 103 | -9.841 | 4.19843 | 4.56362 |
| CD630_31690 | CD630_s0670 | 108 – 121 | 182 – 194 | -9.83947 | 3.71193 | 4.56362 |
| CD630_14130 | CD630_s0670 | 114 – 158 | 41 – 70 | -9.82343 | 4.57970 | 4.56362 |
| gatC | CD630_s0670 | 40 – 65 | 168 – 191 | -9.78478 | 4.25143 | 4.56362 |
| CD630_18950 | CD630_s0670 | 184 – 201 | 43 – 61 | -9.76053 | 3.40610 | 4.56362 |
| CD630_34520 | CD630_s0670 | 7 – 49 | 151 – 192 | -9.73313 | 3.83016 | 4.56362 |
| CD630_14050 | CD630_s0670 | 17 – 37 | 47 – 66 | -9.69036 | 3.30894 | 4.56362 |
| CD630_28290 | CD630_s0670 | 173 – 196 | 28 – 52 | -9.68036 | 5.54192 | 4.56362 |
| flhB | CD630_s0670 | 179 – 194 | 90 – 106 | -9.60446 | 3.55524 | 4.56362 |
| CD630_30100 | CD630_s0670 | 137 – 159 | 30 – 60 | -9.55227 | 4.13234 | 4.56362 |
| CD630_05000 | CD630_s0670 | 181 – 196 | 146 – 161 | -9.54921 | 4.02442 | 4.56362 |
| CD630_31960 | CD630_s0670 | 91 – 138 | 26 – 66 | -9.50924 | 4.93757 | 4.56362 |
| terD | CD630_s0670 | 121 – 136 | 39 – 55 | -9.50471 | 3.33148 | 4.56362 |
| mobA | CD630_SQ1002 | 147 – 195 | 164 – 212 | -18.7143 | 4.25703 | 4.43255 |
| CD630_34030 | CD630_SQ1002 | 53 – 122 | 139 – 208 | -18.4806 | 3.05766 | 4.43255 |
| CD630_23280 | CD630_SQ1002 | 59 – 114 | 86 – 144 | -17.6769 | 3.50093 | 4.43255 |
| CD630_20670 | CD630_SQ1002 | 54 – 123 | 90 – 165 | -17.6599 | 4.42757 | 4.43255 |
| CD630_10000 | CD630_SQ1002 | 27 – 93 | 87 – 154 | -17.0332 | 3.86040 | 4.43255 |
| CD630_31910 | CD630_SQ1002 | 86 – 129 | 131 – 171 | -16.5536 | 3.48314 | 4.43255 |
| CD630_26750 | CD630_SQ1002 | 45 – 91 | 164 – 204 | -16.1949 | 3.60610 | 4.43255 |
| CD630_10970 | CD630_SQ1002 | 53 – 114 | 143 – 200 | -16.0464 | 3.53850 | 4.43255 |
| CD630_08130 | CD630_SQ1002 | 3 – 138 | 85 – 231 | -15.8443 | 4.63831 | 4.43255 |
| CD630_32110 | CD630_SQ1002 | 92 – 138 | 87 – 136 | -15.5977 | 4.45291 | 4.43255 |
| folC | CD630_SQ1002 | 87 – 131 | 126 – 171 | -15.1383 | 3.85634 | 4.43255 |
| CD630_18120 | CD630_SQ1002 | 156 – 191 | 84 – 110 | -15.0453 | 3.99713 | 4.43255 |
| moeA | CD630_SQ1002 | 85 – 174 | 87 – 173 | -14.9548 | 3.26118 | 4.43255 |
| CD630_30540 | CD630_SQ1002 | 170 – 188 | 158 – 172 | -14.7691 | 3.63804 | 4.43255 |
| CD630_08510 | CD630_SQ1002 | 41 – 85 | 84 – 121 | -14.7133 | 5.16329 | 4.43255 |
| CD630_28181 | CD630_SQ1002 | 152 – 164 | 160 – 173 | -14.5896 | 3.91299 | 4.43255 |
| CD630_06050 | CD630_SQ1002 | 116 – 189 | 116 – 191 | -14.503 | 3.59332 | 4.43255 |
| CD630_32880 | CD630_SQ1002 | 113 – 200 | 84 – 155 | -14.297 | 3.41276 | 4.43255 |
| CD630_24590 | CD630_SQ1002 | 80 – 194 | 86 – 225 | -14.2945 | 3.70401 | 4.43255 |
| CD630_32070 | CD630_SQ1002 | 36 – 73 | 85 – 122 | -14.2729 | 3.51964 | 4.43255 |
| cobC | CD630_SQ1002 | 139 – 157 | 94 – 112 | -14.17 | 3.15854 | 4.43255 |
| CD630_04341 | CD630_SQ1002 | 66 – 74 | 164 – 172 | -13.8677 | 4.29262 | 4.43255 |
| tcdC | CD630_SQ1002 | 128 – 154 | 90 – 118 | -13.833 | 3.76101 | 4.43255 |
| CD630_27600 | CD630_SQ1002 | 90 – 129 | 129 – 171 | -13.6607 | 3.48384 | 4.43255 |
| CD630_20170 | CD630_SQ1002 | 103 – 152 | 124 – 169 | -13.6438 | 6.90201 | 4.43255 |
| pilC | CD630_SQ1002 | 122 – 148 | 88 – 110 | -13.4812 | 3.54230 | 4.43255 |
| cbpA | CD630_SQ1002 | 86 – 98 | 158 – 171 | -13.344 | 3.03040 | 4.43255 |
| glnA | CD630_SQ1002 | 87 – 105 | 153 – 170 | -13.1585 | 3.87629 | 4.43255 |
| CD630_32690 | CD630_SQ1002 | 134 – 158 | 206 – 230 | -13.0817 | 7.32676 | 4.43255 |
| etfA | CD630_SQ1002 | 74 – 96 | 162 – 197 | -12.7978 | 4.80521 | 4.43255 |
| CD630_34520 | CD630_SQ1002 | 4 – 18 | 109 – 123 | -12.7169 | 3.83016 | 4.43255 |
| def1 | CD630_SQ1002 | 120 – 154 | 84 – 118 | -12.5787 | 3.99720 | 4.43255 |
| CD630_17730 | CD630_SQ1002 | 55 – 95 | 93 – 130 | -12.5492 | 4.73397 | 4.43255 |
| CD630_30130 | CD630_SQ1002 | 85 – 150 | 89 – 147 | -12.466 | 3.28343 | 4.43255 |
| CD630_25050 | CD630_SQ1002 | 92 – 120 | 83 – 111 | -12.4658 | 4.33624 | 4.43255 |
| flgB | CD630_SQ1002 | 148 – 168 | 87 – 104 | -12.4602 | 3.25736 | 4.43255 |
| CD630_19580 | CD630_SQ1002 | 52 – 72 | 160 – 183 | -12.2971 | 4.51902 | 4.43255 |
| CD630_28330 | CD630_SQ1002 | 152 – 198 | 83 – 134 | -12.267 | 3.29122 | 4.43255 |
| CD630_09060 | CD630_SQ1002 | 104 – 119 | 128 – 143 | -12.2652 | 4.31834 | 4.43255 |
| CD630_26630 | CD630_SQ1002 | 77 – 98 | 89 – 111 | -12.2557 | 4.61557 | 4.43255 |
| CD630_27210 | CD630_SQ1002 | 82 – 122 | 128 – 172 | -12.2486 | 4.18786 | 4.43255 |
| pilW | CD630_SQ1002 | 28 – 97 | 87 – 165 | -12.2406 | 4.96316 | 4.43255 |
| CD630_19903 | CD630_SQ1002 | 159 – 195 | 94 – 130 | -12.1111 | 5.79994 | 4.43255 |
| malY | CD630_SQ1002 | 56 – 126 | 94 – 168 | -12.0756 | 4.46345 | 4.43255 |
| CD630_19790 | CD630_SQ1002 | 63 – 122 | 92 – 157 | -12.0742 | 3.79593 | 4.43255 |
| CD630_17380 | CD630_SQ1002 | 59 – 93 | 88 – 122 | -12.0608 | 5.03207 | 4.43255 |
| garK | CD630_SQ1002 | 15 – 101 | 89 – 168 | -11.9619 | 3.72836 | 4.43255 |
| nusB | CD630_SQ1002 | 92 – 121 | 89 – 118 | -11.9065 | 4.92798 | 4.43255 |
| vncR | CD630_SQ1002 | 115 – 146 | 87 – 122 | -11.8315 | 3.93642 | 4.43255 |
| aroD | CD630_SQ1002 | 110 – 148 | 160 – 196 | -11.8206 | 3.71929 | 4.43255 |
| CD630_16940 | CD630_SQ1002 | 71 – 136 | 116 – 188 | -11.7349 | 4.14665 | 4.43255 |
| CD630_20920 | CD630_SQ1002 | 153 – 174 | 160 – 174 | -11.734 | 4.13812 | 4.43255 |
| CD630_10880 | CD630_SQ1002 | 151 – 181 | 83 – 111 | -11.7321 | 5.05048 | 4.43255 |
| CD630_32520 | CD630_SQ1002 | 121 – 197 | 99 – 166 | -11.5373 | 4.06098 | 4.43255 |
| CD630_19000 | CD630_SQ1002 | 43 – 119 | 89 – 153 | -11.5218 | 4.18443 | 4.43255 |
| CD630_06140 | CD630_SQ1002 | 86 – 93 | 164 – 171 | -11.4519 | 4.12519 | 4.43255 |
| CD630_09040 | CD630_SQ1002 | 116 – 133 | 218 – 235 | -11.4299 | 4.11851 | 4.43255 |
| CD630_14810 | CD630_SQ1002 | 144 – 153 | 161 – 170 | -11.41 | 3.35323 | 4.43255 |
| cspD | CD630_SQ1002 | 160 – 169 | 160 – 169 | -11.4003 | 3.23587 | 4.43255 |
| CD630_23440 | CD630_SQ1002 | 1 – 12 | 160 – 171 | -11.3055 | 3.34841 | 4.43255 |
| CD630_21570 | CD630_SQ1002 | 73 – 102 | 138 – 170 | -11.2158 | 6.48385 | 4.43255 |
| CD630_08360 | CD630_SQ1002 | 47 – 137 | 89 – 172 | -11.185 | 3.19339 | 4.43255 |
| CD630_18090 | CD630_SQ1002 | 80 – 98 | 179 – 200 | -11.1734 | 4.23961 | 4.43255 |
| cobQ | CD630_SQ1002 | 84 – 109 | 165 – 198 | -11.1444 | 3.13524 | 4.43255 |
| CD630_18950 | CD630_SQ1002 | 87 – 159 | 84 – 155 | -11.0507 | 3.40610 | 4.43255 |
| eutV | CD630_SQ1002 | 102 – 109 | 165 – 172 | -10.953 | 4.55715 | 4.43255 |
| cprK | CD630_SQ1002 | 48 – 97 | 162 – 198 | -10.9478 | 3.96216 | 4.43255 |
| CD630_33190 | CD630_SQ1002 | 7 – 19 | 59 – 71 | -10.9369 | 4.13899 | 4.43255 |
| CD630_18230 | CD630_SQ1002 | 1 – 40 | 86 – 119 | -10.919 | 3.00184 | 4.43255 |
| CD630_25620 | CD630_SQ1002 | 8 – 51 | 86 – 129 | -10.9093 | 3.02685 | 4.43255 |
| CD630_21620 | CD630_SQ1002 | 98 – 166 | 89 – 160 | -10.9057 | 3.02649 | 4.43255 |
| xdhA | CD630_SQ1002 | 42 – 161 | 90 – 200 | -10.8961 | 4.45147 | 4.43255 |
| CD630_15900 | CD630_SQ1002 | 157 – 176 | 127 – 146 | -10.8606 | 3.17664 | 4.43255 |
| CD630_29750 | CD630_SQ1002 | 63 – 93 | 182 – 210 | -10.8155 | 4.59146 | 4.43255 |
| CD630_33600 | CD630_SQ1002 | 135 – 159 | 87 – 113 | -10.7845 | 3.73196 | 4.43255 |
| bglA | CD630_SQ1002 | 54 – 93 | 165 – 207 | -10.6763 | 4.86402 | 4.43255 |
| cspC | CD630_SQ1002 | 160 – 169 | 160 – 169 | -10.6749 | 5.48848 | 4.43255 |
| CD630_09850 | CD630_SQ1002 | 2 – 10 | 162 – 170 | -10.6736 | 4.18679 | 4.43255 |
| CD630_14360 | CD630_SQ1002 | 194 – 200 | 165 – 171 | -10.6577 | 3.02079 | 4.43255 |
| metQ | CD630_SQ1002 | 89 – 97 | 162 – 170 | -10.6145 | 3.87190 | 4.43255 |
| CD630_20171 | CD630_SQ1002 | 85 – 118 | 138 – 173 | -10.5906 | 6.87491 | 4.43255 |
| CD630_14920 | CD630_SQ1002 | 102 – 114 | 160 – 172 | -10.5848 | 4.35225 | 4.43255 |
| dnaC | CD630_SQ1002 | 64 – 84 | 84 – 105 | -10.5827 | 6.30796 | 4.43255 |
| CD630_27610 | CD630_SQ1002 | 173 – 195 | 88 – 111 | -10.5557 | 3.96465 | 4.43255 |
| CD630_02790 | CD630_SQ1002 | 126 – 155 | 190 – 216 | -10.515 | 4.77661 | 4.43255 |
| oppC | CD630_SQ1002 | 72 – 161 | 85 – 156 | -10.505 | 3.01695 | 4.43255 |
| CD630_06350 | CD630_SQ1002 | 140 – 163 | 147 – 170 | -10.5024 | 4.35983 | 4.43255 |
| CD630_08950 | CD630_SQ1002 | 30 – 53 | 187 – 209 | -10.5015 | 3.71905 | 4.43255 |
| CD630_27770 | CD630_SQ1002 | 88 – 98 | 162 – 171 | -10.4936 | 4.08588 | 4.43255 |
| CD630_27570 | CD630_SQ1002 | 59 – 127 | 87 – 155 | -10.4732 | 3.46216 | 4.43255 |
| CD630_34560 | CD630_SQ1002 | 103 – 135 | 131 – 168 | -10.4537 | 4.02604 | 4.43255 |
| CD630_20260 | CD630_SQ1002 | 86 – 116 | 143 – 171 | -10.4084 | 3.84162 | 4.43255 |
| CD630_04540 | CD630_SQ1002 | 73 – 114 | 192 – 231 | -10.4076 | 3.33556 | 4.43255 |
| CD630_13620 | CD630_SQ1002 | 174 – 196 | 90 – 113 | -10.3672 | 4.77627 | 4.43255 |
| CD630_07470 | CD630_SQ1002 | 163 – 198 | 94 – 129 | -10.3203 | 3.37143 | 4.43255 |
| CD630_33570 | CD630_SQ1002 | 86 – 103 | 155 – 171 | -10.308 | 3.95778 | 4.43255 |
| bglA | CD630_SQ1002 | 110 – 133 | 164 – 190 | -10.2947 | 4.50315 | 4.43255 |
| bltD | CD630_SQ1002 | 174 – 197 | 1 – 23 | -10.2863 | 4.49926 | 4.43255 |
| CD630_17770 | CD630_SQ1002 | 51 – 69 | 183 – 200 | -10.2602 | 5.06490 | 4.43255 |
| CD630_05000 | CD630_SQ1002 | 183 – 200 | 96 – 113 | -10.2407 | 4.02442 | 4.43255 |
| dltB | CD630_SQ1002 | 61 – 80 | 91 – 110 | -10.2117 | 3.81485 | 4.43255 |
| CD630_11240 | CD630_SQ1002 | 82 – 133 | 127 – 172 | -10.1801 | 3.20274 | 4.43255 |
| acoL | CD630_SQ1002 | 73 – 114 | 192 – 230 | -10.1564 | 3.32169 | 4.43255 |
| CD630_16460 | CD630_SQ1002 | 158 – 164 | 165 – 171 | -10.1502 | 4.64028 | 4.43255 |
| CD630_31580 | CD630_SQ1002 | 75 – 200 | 83 – 181 | -10.1486 | 3.51756 | 4.43255 |
| CD630_06920 | CD630_SQ1002 | 171 – 198 | 85 – 107 | -10.1283 | 3.13260 | 4.43255 |
| CD630_19700 | CD630_SQ1002 | 26 – 93 | 89 – 157 | -10.1246 | 3.85861 | 4.43255 |
| spoVFB | CD630_SQ1002 | 166 – 201 | 90 – 133 | -10.0885 | 4.62758 | 4.43255 |
| CD630_32700 | CD630_SQ1002 | 74 – 90 | 91 – 107 | -10.0852 | 4.74276 | 4.43255 |
| CD630_14050 | CD630_SQ1002 | 75 – 137 | 83 – 157 | -10.083 | 3.30894 | 4.43255 |
| rolB | CD630_SQ1002 | 125 – 137 | 99 – 111 | -10.0777 | 3.60270 | 4.43255 |
| srlD | CD630_SQ1002 | 89 – 95 | 165 – 171 | -10.0312 | 4.19843 | 4.43255 |
| CD630_12650 | CD630_SQ1002 | 111 – 129 | 155 – 173 | -10.0012 | 6.31827 | 4.43255 |
| CD630_17160 | CD630_SQ1002 | 154 – 166 | 186 – 198 | -9.9951 | 3.50228 | 4.43255 |
| oppB | CD630_SQ1002 | 88 – 98 | 188 – 198 | -9.96524 | 2.98328 | 4.43255 |
| CD630_19780 | CD630_SQ1002 | 16 – 23 | 165 – 172 | -9.91317 | 3.64830 | 4.43255 |
| CD630_25320 | CD630_SQ1002 | 146 – 168 | 84 – 105 | -9.90024 | 4.29415 | 4.43255 |
| CD630_09050 | CD630_SQ1002 | 44 – 65 | 4 – 26 | -9.88858 | 3.96216 | 4.43255 |
| CD630_07770 | CD630_SQ1002 | 84 – 97 | 184 – 199 | -9.88564 | 3.31158 | 4.43255 |
| CD630_08100 | CD630_SQ1002 | 181 – 195 | 160 – 171 | -9.8064 | 4.39181 | 4.43255 |
| CD630_31220 | CD630_SQ1002 | 88 – 165 | 90 – 168 | -9.79763 | 3.73779 | 4.43255 |
| CD630_33420 | CD630_SQ1002 | 6 – 47 | 107 – 155 | -9.74549 | 3.22095 | 4.43255 |
| fliI | CD630_SQ1002 | 176 – 192 | 152 – 168 | -9.71711 | 3.03666 | 4.43255 |
| CD630_23760 | CD630_SQ1002 | 54 – 87 | 188 – 212 | -9.70987 | 4.54307 | 4.43255 |
| argS | CD630_SQ1002 | 52 – 108 | 86 – 150 | -9.684 | 4.96036 | 4.43255 |
| CD630_17292 | CD630_SQ1002 | 82 – 114 | 141 – 170 | -9.65442 | 6.25799 | 4.43255 |
| CD630_20600 | CD630_SQ1002 | 19 – 78 | 100 – 147 | -9.64814 | 4.20093 | 4.43255 |
| CD630_08780 | CD630_SQ1002 | 150 – 170 | 124 – 144 | -9.61519 | 7.76690 | 4.43255 |
| CD630_22030 | CD630_SQ1002 | 113 – 136 | 83 – 105 | -9.57271 | 3.29934 | 4.43255 |
| bcp | CD630_SQ1002 | 89 – 184 | 84 – 169 | -9.56466 | 3.16670 | 4.43255 |
| CD630_25190 | CD630_SQ1002 | 134 – 140 | 165 – 171 | -9.53501 | 3.19663 | 4.43255 |
| CD630_19680 | CD630_SQ1002 | 90 – 96 | 165 – 171 | -9.51382 | 3.58080 | 4.43255 |
| CD630_17970 | CD630_SQ808 | 57 – 92 | 39 – 72 | -14.192 | 3.68802 | 6.85680 |
| CD630_18970 | CD630_SQ808 | 101 – 132 | 50 – 80 | -13.7438 | 3.57243 | 6.85680 |
| CD630_33600 | CD630_SQ808 | 138 – 186 | 29 – 80 | -13.5032 | 3.73196 | 6.85680 |
| CD630_15850 | CD630_SQ808 | 88 – 112 | 36 – 61 | -13.2565 | 4.32968 | 6.85680 |
| CD630_10800 | CD630_SQ808 | 27 – 57 | 28 – 57 | -13.2501 | 5.30319 | 6.85680 |
| cheW | CD630_SQ808 | 113 – 183 | 26 – 78 | -13.0134 | 3.72371 | 6.85680 |
| CD630_19903 | CD630_SQ808 | 35 – 71 | 150 – 191 | -12.806 | 5.79994 | 6.85680 |
| CD630_19690 | CD630_SQ808 | 75 – 157 | 70 – 148 | -12.6537 | 3.85861 | 6.85680 |
| CD630_01590 | CD630_SQ808 | 150 – 192 | 33 – 75 | -12.526 | 3.45609 | 6.85680 |
| CD630_02430 | CD630_SQ808 | 90 – 139 | 31 – 75 | -12.459 | 3.71048 | 6.85680 |
| CD630_16940 | CD630_SQ808 | 130 – 166 | 138 – 173 | -12.4166 | 4.14665 | 6.85680 |
| CD630_13400 | CD630_SQ808 | 50 – 104 | 29 – 78 | -12.3884 | 3.20232 | 6.85680 |
| CD630_32520 | CD630_SQ808 | 131 – 161 | 36 – 67 | -12.2376 | 4.06098 | 6.85680 |
| CD630_15190 | CD630_SQ808 | 132 – 198 | 134 – 205 | -12.2299 | 4.43255 | 6.85680 |
| CD630_18440 | CD630_SQ808 | 58 – 126 | 24 – 79 | -12.0942 | 4.07960 | 6.85680 |
| CD630_09970 | CD630_SQ808 | 155 – 184 | 50 – 77 | -12.0936 | 3.25917 | 6.85680 |
| CD630_32720 | CD630_SQ808 | 76 – 106 | 42 – 78 | -11.9208 | 4.74276 | 6.85680 |
| cdu2 | CD630_SQ808 | 27 – 45 | 123 – 143 | -11.8767 | 2.97335 | 6.85680 |
| CD630_14130 | CD630_SQ808 | 146 – 173 | 36 – 68 | -11.8616 | 4.57970 | 6.85680 |
| opuCC | CD630_SQ808 | 18 – 55 | 40 – 79 | -11.7985 | 4.07035 | 6.85680 |
| CD630_13920 | CD630_SQ808 | 116 – 142 | 24 – 48 | -11.7732 | 3.25236 | 6.85680 |
| CD630_31910 | CD630_SQ808 | 149 – 201 | 24 – 78 | -11.702 | 3.48314 | 6.85680 |
| CD630_24590 | CD630_SQ808 | 90 – 146 | 25 – 78 | -11.6082 | 3.70401 | 6.85680 |
| CD630_10050 | CD630_SQ808 | 174 – 183 | 95 – 104 | -11.3519 | 3.93750 | 6.85680 |
| CD630_21770 | CD630_SQ808 | 5 – 15 | 51 – 61 | -11.305 | 4.02587 | 6.85680 |
| CD630_28550 | CD630_SQ808 | 76 – 108 | 194 – 225 | -11.3045 | 3.51196 | 6.85680 |
| spoVFB | CD630_SQ808 | 52 – 86 | 125 – 162 | -11.2928 | 4.62758 | 6.85680 |
| CD630_22510 | CD630_SQ808 | 126 – 154 | 141 – 168 | -11.2659 | 11.42763 | 6.85680 |
| CD630_08140 | CD630_SQ808 | 135 – 166 | 122 – 151 | -11.246 | 3.30159 | 6.85680 |
| ldh | CD630_SQ808 | 160 – 200 | 190 – 226 | -11.2359 | 4.35818 | 6.85680 |
| CD630_18230 | CD630_SQ808 | 157 – 195 | 136 – 174 | -11.2153 | 3.00184 | 6.85680 |
| CD630_18910 | CD630_SQ808 | 53 – 83 | 121 – 151 | -11.208 | 3.30688 | 6.85680 |
| srlEb | CD630_SQ808 | 102 – 137 | 24 – 53 | -11.1644 | 3.50140 | 6.85680 |
| CD630_04660 | CD630_SQ808 | 14 – 41 | 26 – 59 | -11.142 | 4.32791 | 6.85680 |
| CD630_05230 | CD630_SQ808 | 4 – 20 | 163 – 181 | -11.1262 | 3.73787 | 6.85680 |
| CD630_04550 | CD630_SQ808 | 178 – 191 | 48 – 61 | -11.1063 | 3.05227 | 6.85680 |
| CD630_06060 | CD630_SQ808 | 100 – 146 | 34 – 78 | -11.0597 | 4.94050 | 6.85680 |
| cdu1 | CD630_SQ808 | 126 – 153 | 36 – 62 | -10.9426 | 4.46276 | 6.85680 |
| CD630_32050 | CD630_SQ808 | 73 – 86 | 209 – 222 | -10.8389 | 5.73797 | 6.85680 |
| CD630_36250 | CD630_SQ808 | 53 – 62 | 95 – 104 | -10.8196 | 4.13224 | 6.85680 |
| CD630_03920 | CD630_SQ808 | 72 – 110 | 37 – 76 | -10.7394 | 4.15934 | 6.85680 |
| nusB | CD630_SQ808 | 129 – 156 | 122 – 148 | -10.7354 | 4.92798 | 6.85680 |
| CD630_14700 | CD630_SQ808 | 28 – 150 | 29 – 131 | -10.5092 | 3.98234 | 6.85680 |
| CD630_30140 | CD630_SQ808 | 7 – 33 | 30 – 52 | -10.4888 | 3.28343 | 6.85680 |
| bglA | CD630_SQ808 | 59 – 99 | 30 – 78 | -10.4842 | 4.50315 | 6.85680 |
| CD630_24770 | CD630_SQ808 | 161 – 187 | 35 – 63 | -10.3917 | 3.20798 | 6.85680 |
| fbp | CD630_SQ808 | 99 – 114 | 34 – 49 | -10.3836 | 3.80071 | 6.85680 |
| CD630_23450 | CD630_SQ808 | 45 – 84 | 29 – 58 | -10.2869 | 3.27218 | 6.85680 |
| CD630_27240 | CD630_SQ808 | 4 – 13 | 25 – 34 | -10.1671 | 3.52462 | 6.85680 |
| CD630_33570 | CD630_SQ808 | 144 – 164 | 30 – 52 | -10.1622 | 3.95778 | 6.85680 |
| CD630_36190 | CD630_SQ808 | 74 – 110 | 36 – 76 | -10.13 | 4.63449 | 6.85680 |

| | | | | | | | |
|---|---|---|---|---|---|---|---|
| CD630_06200 | CD630_SQ808 | 172 -- 194 | 205 -- 225 | -10.1251 | | 3.57064 | 6.85680 |
| CD630_13790 | CD630_SQ808 | 156 -- 176 | 37 -- 59 | -10.0727 | | 5.17374 | 6.85680 |
| CD630_15740 | CD630_SQ808 | 5 -- 153 | 54 -- 195 | -10.0545 | | 4.43222 | 6.85680 |
| CD630_16160 | CD630_SQ808 | 29 -- 75 | 172 -- 220 | -10.0178 | | 6.11801 | 6.85680 |
| CD630_03040 | CD630_SQ808 | 175 -- 198 | 120 -- 144 | -9.97387 | | 3.51403 | 6.85680 |
| CD630_21490 | CD630_SQ808 | 75 -- 109 | 24 -- 59 | -9.93748 | | 3.37462 | 6.85680 |
| CD630_23280 | CD630_SQ808 | 177 -- 201 | 27 -- 52 | -9.9058 | | 3.50093 | 6.85680 |
| CD630_34560 | CD630_SQ808 | 80 -- 106 | 42 -- 67 | -9.86884 | | 4.02604 | 6.85680 |
| CD630_22720 | CD630_SQ808 | 2 -- 26 | 124 -- 148 | -9.86296 | | 3.81804 | 6.85680 |
| CD630_11241 | CD630_SQ808 | 161 -- 191 | 36 -- 63 | -9.8477 | | 4.85723 | 6.85680 |
| CD630_18930 | CD630_SQ808 | 140 -- 169 | 31 -- 60 | -9.82618 | | 5.32127 | 6.85680 |
| fliI | CD630_SQ808 | 180 -- 189 | 95 -- 104 | -9.81266 | | 3.03666 | 6.85680 |
| gcp | CD630_SQ808 | 8 -- 28 | 25 -- 44 | -9.80271 | | 4.92798 | 6.85680 |
| CD630_06110 | CD630_SQ808 | 169 -- 189 | 55 -- 75 | -9.79826 | | 5.61380 | 6.85680 |
| CD630_31220 | CD630_SQ808 | 123 -- 142 | 206 -- 225 | -9.77934 | | 3.73779 | 6.85680 |
| CD630_09760 | CD630_SQ808 | 110 -- 158 | 30 -- 78 | -9.77925 | | 4.83410 | 6.85680 |
| CD630_28910 | CD630_SQ808 | 110 -- 158 | 30 -- 78 | -9.77925 | | 4.51664 | 6.85680 |
| CD630_17281 | CD630_SQ808 | 144 -- 154 | 94 -- 104 | -9.75403 | | 5.91544 | 6.85680 |
| glgB | CD630_SQ808 | 70 -- 106 | 42 -- 78 | -9.68349 | | 4.48273 | 6.85680 |
| scpB | CD630_SQ808 | 113 -- 158 | 27 -- 78 | -9.6367 | | 3.33560 | 6.85680 |
| garK | CD630_SQ808 | 1 -- 27 | 97 -- 121 | -9.63495 | | 3.72836 | 6.85680 |
| CD630_31000 | CD630_SQ808 | 145 -- 155 | 139 -- 149 | -9.62956 | | 3.70946 | 6.85680 |
| CD630_07430 | CD630_SQ808 | 97 -- 114 | 138 -- 155 | -9.61425 | | 4.65568 | 6.85680 |
| CD630_34520 | CD630_SQ808 | 138 -- 175 | 28 -- 52 | -9.5751 | | 3.83016 | 6.85680 |
| CD630_18680 | CD630_SQ808 | 82 -- 139 | 32 -- 91 | -9.5701 | | 3.04315 | 6.85680 |
| CD630_32110 | CD630_SQ808 | 2 -- 24 | 124 -- 147 | -9.56402 | | 4.45291 | 6.85680 |
| CD630_21700 | CD630_SQ808 | 28 -- 56 | 140 -- 160 | -9.5163 | | 3.96650 | 6.85680 |
| CD630_06050 | RCd2 | 61 -- 110 | 18 -- 70 | -17.3671 | | 3.59332 | 3.80984 |
| CD630_05880 | RCd2 | 62 -- 110 | 16 -- 70 | -15.989 | | 4.28339 | 3.80984 |
| pilT | RCd2 | 83 -- 140 | 16 -- 65 | -15.5369 | | 3.24639 | 3.80984 |
| CD630_27280 | RCd2 | 59 -- 173 | 8 -- 113 | -15.4335 | | 3.24738 | 3.80984 |
| dnaD | RCd2 | 48 -- 98 | 31 -- 81 | -14.8699 | | 3.56078 | 3.80984 |
| CD630_23820 | RCd2 | 141 -- 178 | 18 -- 57 | -14.7067 | | 3.18419 | 3.80984 |
| CD630_21260 | RCd2 | 59 -- 106 | 19 -- 71 | -14.3396 | | 4.22255 | 3.80984 |
| CD630_36180 | RCd2 | 95 -- 112 | 17 -- 34 | -14.286 | | 4.83428 | 3.80984 |
| CD630_24770 | RCd2 | 121 -- 170 | 46 -- 96 | -14.2571 | | 3.20798 | 3.80984 |
| malY | RCd2 | 3 -- 13 | 19 -- 29 | -14.1745 | | 4.46345 | 3.80984 |
| glgP | RCd2 | 170 -- 201 | 168 -- 198 | -14.1436 | | 3.28530 | 3.80984 |
| CD630_33650 | RCd2 | 153 -- 183 | 167 -- 201 | -14.0517 | | 4.80565 | 3.80984 |
| CD630_24590 | RCd2 | 89 -- 130 | 24 -- 57 | -14.0146 | | 3.70401 | 3.80984 |
| etfA | RCd2 | 126 -- 170 | 19 -- 65 | -13.9407 | | 3.41204 | 3.80984 |
| CD630_27120 | RCd2 | 110 -- 160 | 15 -- 68 | -13.2701 | | 4.43720 | 3.80984 |
| CD630_31490 | RCd2 | 43 -- 126 | 21 -- 108 | -12.7603 | | 4.08593 | 3.80984 |
| CD630_28181 | RCd2 | 9 -- 76 | 15 -- 77 | -12.6936 | | 3.91299 | 3.80984 |
| CD630_33570 | RCd2 | 150 -- 178 | 18 -- 46 | -12.6357 | | 3.95778 | 3.80984 |
| moaA | RCd2 | 51 -- 113 | 19 -- 70 | -12.5576 | | 3.83799 | 3.80984 |
| CD630_14360 | RCd2 | 16 -- 106 | 19 -- 94 | -12.5565 | | 3.02079 | 3.80984 |
| CD630_21730 | RCd2 | 13 -- 27 | 173 -- 187 | -12.4716 | | 3.45076 | 3.80984 |
| CD630_21910 | RCd2 | 60 -- 98 | 31 -- 69 | -12.4637 | | 4.66600 | 3.80984 |
| bglA | RCd2 | 157 -- 190 | 19 -- 57 | -12.4537 | | 4.86402 | 3.80984 |
| CD630_05821 | RCd2 | 90 -- 106 | 19 -- 37 | -12.4532 | | 3.75661 | 3.80984 |
| CD630_32060 | RCd2 | 7 -- 54 | 14 -- 76 | -12.34 | | 5.73797 | 3.80984 |
| CD630_05280 | RCd2 | 161 -- 187 | 162 -- 186 | -12.2117 | | 3.91665 | 3.80984 |
| CD630_34530 | RCd2 | 3 -- 20 | 124 -- 141 | -12.1305 | | 3.83016 | 3.80984 |
| CD630_14050 | RCd2 | 30 -- 43 | 24 -- 38 | -12.0314 | | 3.30894 | 3.80984 |
| CD630_32990 | RCd2 | 60 -- 87 | 169 -- 199 | -12.0302 | | 5.24925 | 3.80984 |
| CD630_36170 | RCd2 | 2 -- 40 | 30 -- 81 | -11.8484 | | 3.72380 | 3.80984 |
| CD630_22660 | RCd2 | 111 -- 146 | 163 -- 199 | -11.7358 | | 3.60496 | 3.80984 |
| CD630_06340 | RCd2 | 97 -- 134 | 25 -- 55 | -11.7316 | | 4.35983 | 3.80984 |
| CD630_17970 | RCd2 | 62 -- 108 | 19 -- 69 | -11.7214 | | 3.68802 | 3.80984 |
| CD630_33420 | RCd2 | 15 -- 52 | 16 -- 65 | -11.6856 | | 3.22095 | 3.80984 |
| CD630_14140 | RCd2 | 90 -- 108 | 21 -- 39 | -11.6055 | | 3.16551 | 3.80984 |
| opuCC | RCd2 | 177 -- 196 | 173 -- 196 | -11.5985 | | 4.07035 | 3.80984 |
| panG | RCd2 | 49 -- 71 | 18 -- 38 | -11.5756 | | 5.54047 | 3.80984 |
| CD630_06080 | RCd2 | 119 -- 201 | 13 -- 107 | -11.3417 | | 3.82315 | 3.80984 |
| CD630_29750 | RCd2 | 66 -- 82 | 87 -- 103 | -11.3398 | | 4.59146 | 3.80984 |
| CD630_04410 | RCd2 | 72 -- 103 | 26 -- 67 | -11.3307 | | 3.90451 | 3.80984 |
| CD630_16950 | RCd2 | 174 -- 184 | 68 -- 78 | -11.2838 | | 4.42569 | 3.80984 |
| CD630_16450 | RCd2 | 107 -- 162 | 19 -- 61 | -11.2543 | | 4.57547 | 3.80984 |
| CD630_01590 | RCd2 | 11 -- 29 | 176 -- 195 | -11.1597 | | 3.45609 | 3.80984 |
| CD630_14710 | RCd2 | 57 -- 80 | 169 -- 196 | -11.1375 | | 3.65171 | 3.80984 |
| CD630_32520 | RCd2 | 168 -- 201 | 165 -- 196 | -11.1274 | | 4.06098 | 3.80984 |
| CD630_29250 | RCd2 | 156 -- 195 | 70 -- 103 | -11.0924 | | 4.61901 | 3.80984 |
| CD630_09410 | RCd2 | 156 -- 195 | 70 -- 103 | -11.0924 | | 4.75651 | 3.80984 |
| CD630_18950 | RCd2 | 90 -- 117 | 167 -- 192 | -11.0813 | | 3.40610 | 3.80984 |
| CD630_32720 | RCd2 | 44 -- 100 | 12 -- 67 | -11.026 | | 4.74276 | 3.80984 |
| CD630_02790 | RCd2 | 30 -- 63 | 165 -- 195 | -10.8596 | | 4.77661 | 3.80984 |
| gcp | RCd2 | 138 -- 159 | 174 -- 194 | -10.8539 | | 4.92798 | 3.80984 |
| CD630_20920 | RCd2 | 38 -- 59 | 44 -- 65 | -10.8538 | | 4.13812 | 3.80984 |
| CD630_19680 | RCd2 | 37 -- 86 | 15 -- 53 | -10.8534 | | 3.58080 | 3.80984 |
| def1 | RCd2 | 151 -- 186 | 20 -- 57 | -10.8318 | | 3.99720 | 3.80984 |
| CD630_21030 | RCd2 | 49 -- 79 | 29 -- 61 | -10.7843 | | 3.08733 | 3.80984 |
| CD630_16160 | RCd2 | 176 -- 199 | 174 -- 196 | -10.7729 | | 6.11801 | 3.80984 |
| cobC | RCd2 | 92 -- 114 | 15 -- 37 | -10.6863 | | 3.15854 | 3.80984 |
| CD630_32040 | RCd2 | 157 -- 169 | 24 -- 38 | -10.6267 | | 4.93516 | 3.80984 |
| argS | RCd2 | 46 -- 61 | 19 -- 33 | -10.6111 | | 4.96036 | 3.80984 |
| CD630_06100 | RCd2 | 128 -- 156 | 171 -- 201 | -10.6016 | | 4.43918 | 3.80984 |
| CD630_17720 | RCd2 | 152 -- 165 | 128 -- 141 | -10.5937 | | 3.22910 | 3.80984 |
| CD630_17292 | RCd2 | 14 -- 25 | 22 -- 33 | -10.5772 | | 6.25799 | 3.80984 |
| CD630_15890 | RCd2 | 147 -- 167 | 14 -- 35 | -10.5392 | | 3.39962 | 3.80984 |
| CD630_23770 | RCd2 | 62 -- 91 | 168 -- 201 | -10.5381 | | 3.25181 | 3.80984 |
| terD | RCd2 | 50 -- 68 | 171 -- 188 | -10.5131 | | 3.33148 | 3.80984 |
| ldh | RCd2 | 132 -- 150 | 25 -- 39 | -10.4565 | | 4.35818 | 3.80984 |
| CD630_08080 | RCd2 | 66 -- 93 | 163 -- 195 | -10.4452 | | 4.87819 | 3.80984 |
| CD630_02720 | RCd2 | 160 -- 181 | 59 -- 76 | -10.4334 | | 3.38938 | 3.80984 |
| CD630_19401 | RCd2 | 20 -- 77 | 6 -- 80 | -10.4028 | | 4.37084 | 3.80984 |
| CD630_29760 | RCd2 | 113 -- 154 | 15 -- 69 | -10.4004 | | 3.51074 | 3.80984 |
| CD630_16330 | RCd2 | 105 -- 128 | 168 -- 189 | -10.3918 | | 4.11180 | 3.80984 |
| CD630_22070 | RCd2 | 55 -- 84 | 170 -- 201 | -10.3563 | | 4.09651 | 3.80984 |
| CD630_27570 | RCd2 | 90 -- 115 | 18 -- 39 | -10.3331 | | 3.46216 | 3.80984 |
| CD630_08390 | RCd2 | 84 -- 100 | 24 -- 38 | -10.3318 | | 5.16000 | 3.80984 |
| cheW | RCd2 | 54 -- 105 | 20 -- 71 | -10.2908 | | 3.72371 | 3.80984 |
| rolA | RCd2 | 147 -- 157 | 65 -- 75 | -10.1936 | | 4.56362 | 3.80984 |
| CD630_19921 | RCd2 | 63 -- 111 | 21 -- 68 | -10.1372 | | 5.46345 | 3.80984 |
| CD630_13580 | RCd2 | 134 -- 183 | 47 -- 93 | -10.1244 | | 3.80715 | 3.80984 |
| CD630_21020 | RCd2 | 40 -- 108 | 19 -- 68 | -10.1178 | | 4.28666 | 3.80984 |
| CD630_22880 | RCd2 | 175 -- 201 | 19 -- 45 | -10.1152 | | 3.62455 | 3.80984 |
| CD630_20670 | RCd2 | 45 -- 89 | 24 -- 65 | -10.1071 | | 4.42757 | 3.80984 |
| cwp26 | RCd2 | 67 -- 76 | 33 -- 42 | -10.0893 | | 4.44377 | 3.80984 |
| CD630_16150 | RCd2 | 146 -- 177 | 7 -- 34 | -10.0855 | | 4.20614 | 3.80984 |
| CD630_08510 | RCd2 | 40 -- 85 | 19 -- 61 | -10.0176 | | 5.16329 | 3.80984 |
| CD630_33682 | RCd2 | 81 -- 123 | 21 -- 59 | -9.98891 | | 5.77720 | 3.80984 |
| CD630_18090 | RCd2 | 120 -- 142 | 19 -- 42 | -9.98527 | | 4.23961 | 3.80984 |
| CD630_22550 | RCd2 | 30 -- 86 | 16 -- 73 | -9.97337 | | 3.62379 | 3.80984 |
| CD630_20960 | RCd2 | 108 -- 152 | 19 -- 57 | -9.97189 | | 5.65940 | 3.80984 |
| CD630_28550 | RCd2 | 77 -- 116 | 16 -- 49 | -9.8734 | | 3.51196 | 3.80984 |
| CD630_05900 | RCd2 | 4 -- 18 | 171 -- 186 | -9.84328 | | 3.21934 | 3.80984 |
| CD630_26250 | RCd2 | 167 -- 188 | 60 -- 79 | -9.8372 | | 5.03379 | 3.80984 |
| CD630_10080 | RCd2 | 182 -- 197 | 124 -- 139 | -9.80326 | | 3.75625 | 3.80984 |
| moeA | RCd2 | 33 -- 93 | 19 -- 84 | -9.76591 | | 3.26118 | 3.80984 |
| CD630_10220 | RCd2 | 64 -- 96 | 29 -- 57 | -9.73551 | | 4.08233 | 3.80984 |
| CD630_05840 | RCd2 | 122 -- 148 | 170 -- 196 | -9.72174 | | 4.28779 | 3.80984 |
| CD630_19030 | RCd2 | 11 -- 36 | 51 -- 74 | -9.7069 | | 3.34240 | 3.80984 |
| CD630_24580 | RCd2 | 95 -- 108 | 21 -- 35 | -9.69693 | | 3.88519 | 3.80984 |
| cbiQ | RCd2 | 147 -- 169 | 159 -- 182 | -9.6732 | | 3.79830 | 3.80984 |
| CD630_14340 | RCd2 | 54 -- 111 | 14 -- 65 | -9.64811 | | 5.10868 | 3.80984 |
| cdu2 | RCd2 | 49 -- 95 | 17 -- 51 | -9.64205 | | 2.97335 | 3.80984 |
| CD630_31461 | RCd2 | 155 -- 185 | 169 -- 201 | -9.61917 | | 4.41476 | 3.80984 |
| CD630_04540 | RCd2 | 25 -- 84 | 21 -- 84 | -9.61049 | | 3.33556 | 3.80984 |
| feoB | RCd2 | 1 -- 22 | 56 -- 80 | -9.55832 | | 4.52257 | 3.80984 |
| CD630_13020 | RCd2 | 119 -- 148 | 70 -- 89 | -9.55196 | | 3.52267 | 3.80984 |

**Table S10. IntaRNA prediction for ncRNA-ncRNA interactions**

| | | | | | RIPseq log p-value | |
|---|---|---|---|---|---|---|
| **ncRNA1** | **ncRNA2** | **interaction ncRNA1** | **interaction ncRNA2** | **interaction energy (kcal/mol)** | **ncRNA1 logpVal** | **ncRNA2 logpVal** |
| CD630_s0470 | CD630_n01010 | 8 -- 99 | 1275 -- 1353 | -19.842 | 5.35019 | 4.59146 |
| CD630_n01010 | CD630_s0470 | 1275 -- 1353 | 8 -- 99 | -19.842 | 4.59146 | 5.35019 |
| CD630_n00600 | CD630_s0450 | 797 -- 836 | 15 -- 49 | -19.7696 | 5.31146 | 5.67265 |
| CD630_s0450 | CD630_n00600 | 15 -- 49 | 797 -- 836 | -19.7696 | 5.67265 | 5.31146 |
| CD630_n01010 | CD630_n00380 | 309 -- 458 | 158 -- 306 | -19.7007 | 4.59146 | 4.27916 |
| CD630_n01010 | CD630_n00990 | 309 -- 458 | 247 -- 395 | -19.7007 | 4.59146 | 3.72051 |
| CD630_s0400 | CD630_n00080 | 21 -- 78 | 2 -- 60 | -19.4845 | 4.51086 | 4.59330 |
| CD630_n00080 | CD630_s0400 | 2 -- 60 | 21 -- 78 | -19.4845 | 4.59330 | 4.51086 |
| CD630_n00380 | CD630_n01010 | 20 -- 57 | 62 -- 98 | -19.4462 | 4.27916 | 4.59146 |
| CD630_Cdi1_3 | CD630_n00440 | 468 -- 496 | 56 -- 87 | -19.3477 | 6.59405 | 4.85723 |
| CD630_n00440 | CD630_Cdi1_3 | 56 -- 87 | 468 -- 496 | -19.3477 | 4.85723 | 6.59405 |
| CD630_n00080 | CD630_s0450 | 60 -- 100 | 7 -- 48 | -19.192 | 4.59330 | 5.67265 |
| CD630_s0450 | CD630_n00080 | 7 -- 48 | 60 -- 100 | -19.192 | 5.67265 | 4.59330 |
| CD630_s0642 | CD630_s0642 | 29 -- 177 | 25 -- 173 | -19.1117 | 6.00600 | 6.00600 |
| RCd8 | CD630_n00690 | 6 -- 29 | 2 -- 25 | -19.0198 | 6.97567 | 5.96526 |
| CD630_n00690 | RCd8 | 2 -- 25 | 6 -- 29 | -19.0198 | 5.96526 | 6.97567 |
| CD630_n00380 | CD630_n00380 | 725 -- 752 | 706 -- 734 | -18.8887 | 4.27916 | 4.27916 |
| CD630_n00380 | CD630_n00990 | 725 -- 752 | 795 -- 823 | -18.8887 | 4.27916 | 3.72051 |
| CD630_n00990 | CD630_n00380 | 814 -- 841 | 706 -- 734 | -18.8887 | 3.72051 | 4.27916 |
| CD630_n00990 | CD630_n00990 | 814 -- 841 | 795 -- 823 | -18.8887 | 3.72051 | 3.72051 |
| CD630_n00990 | CD630_n01010 | 109 -- 146 | 62 -- 98 | -18.7669 | 3.72051 | 4.59146 |
| CD630_n00690 | CD630_n01010 | 7 -- 54 | 35 -- 77 | -18.5332 | 5.96526 | 4.59146 |
| CD630_n01010 | CD630_n00690 | 35 -- 77 | 7 -- 54 | -18.5332 | 4.59146 | 5.96526 |
| CD630_n00510 | CD630_n00790 | 321 -- 347 | 465 -- 490 | -18.4504 | 3.92762 | 5.51210 |
| CD630_n00790 | CD630_n00510 | 465 -- 493 | 319 -- 347 | -18.4066 | 5.51210 | 3.92762 |
| CD630_n00080 | CD630_s0510 | 57 -- 92 | 5 -- 49 | -17.9709 | 4.59330 | 3.37631 |
| CD630_s0510 | CD630_n00080 | 5 -- 49 | 57 -- 92 | -17.9709 | 3.37631 | 4.59330 |
| CD630_n00380 | CD630_SQ1642 | 643 -- 691 | 147 -- 192 | -17.8859 | 4.27916 | 8.65138 |
| CD630_n00990 | CD630_SQ1642 | 732 -- 780 | 147 -- 192 | -17.8859 | 3.72051 | 8.65138 |
| CD630_SQ1642 | CD630_n00380 | 147 -- 177 | 662 -- 691 | -17.8617 | 8.65138 | 4.27916 |
| CD630_SQ1642 | CD630_n00990 | 147 -- 177 | 751 -- 780 | -17.8617 | 8.65138 | 3.72051 |
| CD630_SQ1641 | RCd9 | 106 -- 154 | 254 -- 298 | -17.7432 | 8.65138 | 8.96925 |
| RCd9 | CD630_SQ1641 | 254 -- 298 | 106 -- 154 | -17.7432 | 8.96925 | 8.65138 |
| RCd8 | CD630_n00790 | 6 -- 29 | 2 -- 25 | -17.2847 | 6.97567 | 5.51210 |
| CD630_n00790 | RCd8 | 2 -- 25 | 6 -- 29 | -17.2847 | 5.51210 | 6.97567 |
| CD630_n00600 | CD630_n00380 | 337 -- 365 | 790 -- 818 | -17.2738 | 5.31146 | 4.27916 |
| CD630_n00600 | CD630_n00990 | 337 -- 365 | 879 -- 907 | -17.2738 | 5.31146 | 3.72051 |
| CD630_Cdi1_3 | CD630_Cdi1_5 | 485 -- 497 | 437 -- 449 | -17.202 | 6.59405 | 4.83391 |
| CD630_Cdi1_5 | CD630_Cdi1_3 | 437 -- 449 | 485 -- 497 | -17.202 | 4.83391 | 6.59405 |
| CD630_Cdi1_3 | CD630_Cdi1_4 | 485 -- 497 | 444 -- 456 | -17.166 | 6.59405 | 5.31318 |
| CD630_Cdi1_4 | CD630_Cdi1_3 | 444 -- 456 | 485 -- 497 | -17.166 | 5.31318 | 6.59405 |
| CD630_s0340 | CD630_n00380 | 53 -- 78 | 309 -- 339 | -17.1279 | 5.61064 | 4.27916 |
| CD630_s0340 | CD630_n00990 | 53 -- 78 | 398 -- 428 | -17.1279 | 5.61064 | 3.72051 |
| CD630_n00560 | CD630_n00330 | 276 -- 286 | 168 -- 178 | -17.1092 | 5.82053 | 6.46464 |
| CD630_n00330 | CD630_n00560 | 168 -- 178 | 276 -- 286 | -17.1092 | 6.46464 | 5.82053 |
| CD630_s0642 | CD630_n00790 | 191 -- 225 | 104 -- 134 | -17.0208 | 6.00600 | 5.51210 |
| CD630_n00380 | CD630_s0340 | 309 -- 353 | 42 -- 78 | -16.9751 | 4.27916 | 5.61064 |
| CD630_n00990 | CD630_s0340 | 398 -- 442 | 42 -- 78 | -16.9751 | 3.72051 | 5.61064 |
| CD630_SQ1642 | CD630_n00460 | 5 -- 41 | 863 -- 912 | -16.9712 | 8.65138 | 5.55447 |
| RCd5 | CD630_n00290 | 111 -- 194 | 23 -- 104 | -16.8234 | 6.52781 | 3.85358 |
| CD630_n00680 | CD630_n00290 | 111 -- 194 | 23 -- 104 | -16.8234 | 6.52781 | 3.85358 |
| CD630_n00290 | RCd5 | 23 -- 104 | 111 -- 194 | -16.8234 | 3.85358 | 6.52781 |
| CD630_n00290 | CD630_n00680 | 23 -- 104 | 111 -- 194 | -16.8234 | 3.85358 | 6.52781 |
| CD630_n01010 | CD630_n01010 | 281 -- 403 | 1053 -- 1161 | -16.7626 | 4.59146 | 4.59146 |
| CD630_n01010 | CD630_n00510 | 50 -- 73 | 10 -- 43 | -16.7516 | 4.59146 | 3.92762 |
| CD630_s0641 | CD630_n01010 | 92 -- 134 | 814 -- 859 | -16.7068 | 6.68432 | 4.59146 |
| CD630_n01010 | CD630_s0641 | 814 -- 859 | 92 -- 134 | -16.7068 | 4.59146 | 6.68432 |
| CD630_SQ367 | CD630_SQ1642 | 67 -- 191 | 78 -- 201 | -16.6649 | 4.08290 | 8.65138 |
| CD630_n00600 | CD630_n00470 | 514 -- 565 | 37 -- 97 | -16.6609 | 5.31146 | 3.52267 |
| CD630_n00470 | CD630_n00600 | 37 -- 97 | 514 -- 565 | -16.6609 | 3.52267 | 5.31146 |
| CD630_n01010 | CD630_n00600 | 35 -- 78 | 6 -- 54 | -16.5719 | 4.59146 | 5.31146 |
| CD630_s0390 | CD630_s0670 | 214 -- 251 | 26 -- 67 | -16.5575 | 4.73397 | 4.56362 |
| CD630_Cdi2_3 | CD630_n00500 | 34 -- 55 | 42 -- 63 | -16.5042 | 3.99184 | 9.35359 |
| CD630_n00500 | CD630_Cdi2_3 | 42 -- 63 | 34 -- 55 | -16.5041 | 9.35359 | 3.99184 |
| CD630_n00080 | CD630_s0460 | 52 -- 102 | 3 -- 60 | -16.4597 | 4.59330 | 5.74527 |
| RCd5 | CD630_n00380 | 335 -- 430 | 615 -- 701 | -16.4183 | 6.52781 | 4.27916 |
| RCd5 | CD630_n00990 | 335 -- 430 | 704 -- 790 | -16.4183 | 6.52781 | 3.72051 |
| CD630_n00380 | RCd5 | 615 -- 701 | 335 -- 430 | -16.4182 | 4.27916 | 6.52781 |
| CD630_n00990 | RCd5 | 704 -- 790 | 335 -- 430 | -16.4182 | 3.72051 | 6.52781 |
| CD630_n01060 | CD630_SQ1642 | 89 -- 115 | 163 -- 183 | -16.281 | 3.97522 | 8.65138 |
| CD630_s0631 | CD630_s0642 | 315 -- 342 | 21 -- 50 | -16.247 | 4.95025 | 6.00600 |
| CD630_s0642 | CD630_s0631 | 21 -- 50 | 315 -- 342 | -16.247 | 6.00600 | 4.95025 |
| CD630_Cdi1_8 | CD630_SQ1641 | 230 -- 270 | 5 -- 43 | -16.222 | 5.79994 | 8.65138 |
| CD630_n00380 | CD630_s0641 | 200 -- 343 | 74 -- 209 | -16.177 | 4.27916 | 6.68432 |
| CD630_n00990 | CD630_s0641 | 289 -- 432 | 74 -- 209 | -16.177 | 3.72051 | 6.68432 |
| CD630_s0190 | CD630_s0010 | 78 -- 109 | 58 -- 87 | -16.1513 | 4.98480 | 4.92433 |
| CD630_s0010 | CD630_s0190 | 58 -- 88 | 77 -- 109 | -16.1484 | 4.92433 | 4.98480 |
| CD630_n00460 | CD630_s0370 | 290 -- 374 | 4 -- 97 | -16.0927 | 5.55447 | 10.21495 |
| CD630_SQ1642 | CD630_n01060 | 163 -- 194 | 76 -- 115 | -16.0757 | 8.65138 | 3.97522 |
| CD630_n00560 | CD630_s0400 | 409 -- 470 | 1 -- 54 | -16.0313 | 5.82053 | 4.51086 |
| CD630_n00510 | CD630_n01010 | 10 -- 23 | 60 -- 73 | -16.0303 | 3.92762 | 4.59146 |
| CD630_s0641 | CD630_s0641 | 253 -- 291 | 215 -- 248 | -16.0157 | 6.68432 | 6.68432 |
| CD630_s0641 | CD630_SQ1642 | 32 -- 175 | 57 -- 195 | -16.0093 | 6.68432 | 8.65138 |
| CD630_n00380 | CD630_s0510 | 304 -- 339 | 6 -- 52 | -15.9836 | 4.27916 | 3.37631 |
| CD630_s0510 | CD630_n00380 | 6 -- 52 | 304 -- 339 | -15.9836 | 3.37631 | 4.27916 |
| CD630_s0510 | CD630_n00990 | 6 -- 52 | 393 -- 428 | -15.9836 | 3.37631 | 3.72051 |
| CD630_n00990 | CD630_s0510 | 393 -- 428 | 6 -- 52 | -15.9836 | 3.72051 | 3.37631 |
| CD630_s0340 | CD630_n00790 | 20 -- 74 | 199 -- 248 | -15.9413 | 5.61064 | 5.51210 |
| CD630_n00600 | CD630_s0340 | 105 -- 139 | 132 -- 170 | -15.9157 | 5.31146 | 5.61064 |
| CD630_n00460 | CD630_SQ1642 | 863 -- 895 | 10 -- 41 | -15.9056 | 5.55447 | 8.65138 |
| CD630_SQ1641 | CD630_Cdi1_8 | 5 -- 40 | 233 -- 270 | -15.8979 | 8.65138 | 5.79994 |
| CD630_n00980 | CD630_s0642 | 320 -- 360 | 16 -- 53 | -15.8959 | 5.31318 | 6.00600 |
| CD630_SQ808 | CD630_n00790 | 24 -- 55 | 377 -- 405 | -15.8907 | 6.85680 | 5.51210 |
| CD630_n00790 | CD630_SQ808 | 377 -- 405 | 24 -- 55 | -15.8907 | 5.51210 | 6.85680 |
| CD630_s0642 | CD630_n00980 | 16 -- 43 | 330 -- 360 | -15.8783 | 6.00600 | 5.31318 |
| CD630_s0400 | CD630_n00560 | 1 -- 68 | 393 -- 470 | -15.863 | 4.51086 | 5.82053 |
| CD630_s0450 | CD630_n00460 | 3 -- 48 | 286 -- 344 | -15.8107 | 5.67265 | 5.55447 |
| CD630_n01010 | CD630_s0210 | 36 -- 90 | 8 -- 61 | -15.7751 | 4.59146 | 4.02122 |
| CD630_s0210 | CD630_n00460 | 8 -- 51 | 205 -- 246 | -15.7571 | 4.02122 | 5.55447 |
| CD630_n00460 | CD630_s0210 | 205 -- 246 | 8 -- 51 | -15.7571 | 5.55447 | 4.02122 |
| CD630_SQ1642 | CD630_SQ367 | 78 -- 195 | 78 -- 191 | -15.674 | 8.65138 | 4.08290 |
| CD630_n00600 | CD630_s0510 | 795 -- 843 | 9 -- 53 | -15.6666 | 5.31146 | 3.37631 |
| CD630_s0510 | CD630_n00600 | 9 -- 53 | 795 -- 843 | -15.6666 | 3.37631 | 5.31146 |
| CD630_s0340 | CD630_n00600 | 132 -- 169 | 106 -- 139 | -15.5592 | 5.61064 | 5.31146 |
| CD630_SQ327 | CD630_n01010 | 54 -- 123 | 851 -- 912 | -15.529 | 4.62291 | 4.59146 |
| CD630_n01010 | CD630_SQ327 | 851 -- 912 | 54 -- 123 | -15.529 | 4.59146 | 4.62291 |
| CD630_Cdi2_2 | CD630_n00380 | 11 -- 61 | 301 -- 344 | -15.529 | 6.72203 | 4.27916 |
| CD630_Cdi2_2 | CD630_n00990 | 11 -- 61 | 390 -- 433 | -15.529 | 6.72203 | 3.72051 |
| CD630_Cdi2_2 | CD630_n00380 | 11 -- 61 | 301 -- 344 | -15.529 | 6.72203 | 4.27916 |
| CD630_Cdi2_2 | CD630_n00990 | 11 -- 61 | 390 -- 433 | -15.529 | 6.72203 | 3.72051 |
| CD630_n00560 | CD630_s0390 | 468 -- 513 | 16 -- 64 | -15.5279 | 5.82053 | 4.73397 |
| CD630_s0390 | CD630_n00560 | 16 -- 64 | 468 -- 513 | -15.5279 | 4.73397 | 5.82053 |
| CD630_n00080 | CD630_n00590 | 9 -- 90 | 23 -- 99 | -15.527 | 4.59330 | 4.28395 |
| CD630_n00600 | CD630_n01010 | 981 -- 1069 | 330 -- 419 | -15.5205 | 5.31146 | 4.59146 |
| CD630_s0210 | CD630_n01010 | 8 -- 60 | 37 -- 90 | -15.4908 | 4.02122 | 4.59146 |
| CD630_n00560 | RCd8 | 1 -- 23 | 6 -- 28 | -15.4561 | 5.82053 | 6.97567 |
| RCd8 | CD630_n00560 | 6 -- 28 | 1 -- 23 | -15.4561 | 6.97567 | 5.82053 |
| CD630_n00680 | RCd5 | 255 -- 386 | 300 -- 431 | -15.4253 | 6.52781 | 6.52781 |
| CD630_Cdi1_4 | CD630_s0642 | 330 -- 360 | 16 -- 43 | -15.3871 | 5.31318 | 6.00600 |
| CD630_s0642 | CD630_Cdi1_4 | 16 -- 43 | 330 -- 360 | -15.3871 | 6.00600 | 5.31318 |
| CD630_s0641 | CD630_n00380 | 74 -- 207 | 202 -- 343 | -15.3614 | 6.68432 | 4.27916 |
| CD630_s0641 | CD630_n00990 | 74 -- 207 | 291 -- 432 | -15.3614 | 6.68432 | 3.72051 |
| CD630_s0210 | CD630_n00690 | 13 -- 87 | 159 -- 242 | -15.3588 | 4.02122 | 5.96526 |
| CD630_n00080 | CD630_n00380 | 48 -- 89 | 761 -- 807 | -15.3584 | 4.59330 | 4.27916 |
| CD630_n00080 | CD630_n00990 | 48 -- 89 | 850 -- 896 | -15.3584 | 4.59330 | 3.72051 |
| CD630_n00460 | CD630_n00380 | 690 -- 744 | 754 -- 814 | -15.3362 | 5.55447 | 4.27916 |
| CD630_n00460 | CD630_n00990 | 690 -- 744 | 843 -- 903 | -15.3362 | 5.55447 | 3.72051 |
| RCd5 | CD630_n00680 | 300 -- 426 | 263 -- 386 | -15.3238 | 6.52781 | 6.52781 |
| CD630_n00460 | CD630_s0450 | 286 -- 326 | 13 -- 48 | -15.3052 | 5.55447 | 5.67265 |
| CD630_s0190 | CD630_n00690 | 74 -- 109 | 202 -- 235 | -15.2444 | 4.98480 | 5.96526 |
| CD630_n00690 | CD630_s0190 | 202 -- 235 | 74 -- 109 | -15.2444 | 5.96526 | 4.98480 |
| CD630_n00470 | CD630_n00790 | 21 -- 86 | 193 -- 255 | -15.2269 | 3.52267 | 5.51210 |
| CD630_n00790 | CD630_n00470 | 193 -- 255 | 21 -- 86 | -15.2269 | 5.51210 | 3.52267 |
| CD630_Cdi1_4 | CD630_Cdi2_2 | 136 -- 164 | 20 -- 49 | -15.1773 | 5.31318 | 6.72203 |
| CD630_Cdi1_4 | CD630_Cdi2_2 | 136 -- 164 | 20 -- 49 | -15.1773 | 5.31318 | 6.72203 |
| CD630_n00980 | CD630_Cdi2_2 | 136 -- 164 | 20 -- 49 | -15.1773 | 5.31318 | 6.72203 |
| CD630_n00980 | CD630_Cdi2_2 | 136 -- 164 | 20 -- 49 | -15.1773 | 5.31318 | 6.72203 |
| CD630_Cdi1_5 | CD630_Cdi2_2 | 129 -- 157 | 20 -- 49 | -15.1773 | 4.83391 | 6.72203 |
| CD630_Cdi1_5 | CD630_Cdi2_2 | 129 -- 157 | 20 -- 49 | -15.1773 | 4.83391 | 6.72203 |
| CD630_n00390 | CD630_Cdi2_2 | 129 -- 157 | 20 -- 49 | -15.1773 | 4.83391 | 6.72203 |
| CD630_n00390 | CD630_Cdi2_2 | 129 -- 157 | 20 -- 49 | -15.1773 | 4.83391 | 6.72203 |
| CD630_Cdi2_2 | CD630_Cdi1_4 | 20 -- 49 | 136 -- 164 | -15.1773 | 6.72203 | 5.31318 |
| CD630_Cdi2_2 | CD630_n00980 | 20 -- 49 | 136 -- 164 | -15.1773 | 6.72203 | 5.31318 |
| CD630_Cdi2_2 | CD630_Cdi1_5 | 20 -- 49 | 129 -- 157 | -15.1773 | 6.72203 | 4.83391 |
| CD630_Cdi2_2 | CD630_n00390 | 20 -- 49 | 129 -- 157 | -15.1773 | 6.72203 | 4.83391 |
| CD630_Cdi2_2 | CD630_Cdi1_4 | 20 -- 49 | 136 -- 164 | -15.1773 | 6.72203 | 5.31318 |
| CD630_Cdi2_2 | CD630_n00980 | 20 -- 49 | 136 -- 164 | -15.1773 | 6.72203 | 5.31318 |

| | | | | | | |
|---|---|---|---|---|---|---|
| CD630_Cdi2_2 | CD630_Cdi1_5 | 20 -- 49 | 129 -- 157 | -15.1773 | 6.72203 | 4.83391 |
| CD630_Cdi2_2 | CD630_n00390 | 20 -- 49 | 129 -- 157 | -15.1773 | 6.72203 | 4.83391 |
| CD630_n00600 | CD630_s0390 | 793 -- 818 | 32 -- 60 | -15.1709 | 5.31146 | 4.73397 |
| CD630_s0390 | CD630_n00600 | 32 -- 60 | 793 -- 818 | -15.1709 | 4.73397 | 5.31146 |
| CD630_s0590 | CD630_n01010 | 37 -- 108 | 50 -- 129 | -15.1555 | 8.13716 | 4.59146 |
| CD630_n00590 | CD630_n00080 | 23 -- 76 | 42 -- 90 | -15.1288 | 4.28395 | 4.59330 |
| CD630_n00680 | CD630_n00380 | 353 -- 388 | 656 -- 689 | -15.1127 | 6.52781 | 4.27916 |
| CD630_n00680 | CD630_n00990 | 353 -- 388 | 745 -- 778 | -15.1127 | 6.52781 | 3.72051 |
| CD630_SQ1642 | CD630_s0510 | 158 -- 178 | 5 -- 25 | -15.0931 | 8.65138 | 3.37631 |
| CD630_s0510 | CD630_SQ1642 | 5 -- 25 | 158 -- 178 | -15.0931 | 3.37631 | 8.65138 |
| RCd5 | CD630_SQ1641 | 234 -- 261 | 5 -- 32 | -15.0769 | 6.52781 | 8.65138 |
| CD630_n00680 | CD630_SQ1641 | 234 -- 261 | 5 -- 32 | -15.0769 | 6.52781 | 8.65138 |
| CD630_SQ1641 | RCd5 | 5 -- 32 | 234 -- 261 | -15.0769 | 8.65138 | 6.52781 |
| CD630_SQ1641 | CD630_n00680 | 5 -- 32 | 234 -- 261 | -15.0769 | 8.65138 | 6.52781 |
| CD630_n00380 | CD630_n00080 | 761 -- 808 | 47 -- 89 | -15.0535 | 4.27916 | 4.59330 |
| CD630_n00990 | CD630_n00080 | 850 -- 897 | 47 -- 89 | -15.0535 | 3.72051 | 4.59330 |
| CD630_n00790 | CD630_s0642 | 104 -- 143 | 190 -- 225 | -15.0522 | 5.51210 | 6.00600 |
| CD630_n00380 | CD630_Cdi1_4 | 155 -- 190 | 182 -- 216 | -14.9903 | 4.27916 | 5.31318 |
| CD630_n00380 | CD630_n00980 | 155 -- 190 | 182 -- 216 | -14.9903 | 4.27916 | 5.31318 |
| CD630_n00990 | CD630_Cdi1_4 | 244 -- 279 | 182 -- 216 | -14.9903 | 3.72051 | 5.31318 |
| CD630_n00990 | CD630_n00980 | 244 -- 279 | 182 -- 216 | -14.9903 | 3.72051 | 5.31318 |
| CD630_n00380 | CD630_n00560 | 671 -- 698 | 460 -- 486 | -14.972 | 4.27916 | 5.82053 |
| CD630_n00560 | CD630_n00380 | 460 -- 486 | 671 -- 698 | -14.972 | 5.82053 | 4.27916 |
| CD630_n00560 | CD630_n00990 | 460 -- 486 | 760 -- 787 | -14.972 | 5.82053 | 3.72051 |
| CD630_n00990 | CD630_n00560 | 760 -- 787 | 460 -- 486 | -14.972 | 3.72051 | 5.82053 |
| RCd8 | CD630_n00460 | 68 -- 206 | 647 -- 785 | -14.9677 | 6.97567 | 5.55447 |
| CD630_n01010 | CD630_s0590 | 50 -- 130 | 36 -- 108 | -14.9287 | 4.59146 | 8.13716 |
| CD630_n00080 | CD630_s0590 | 10 -- 98 | 7 -- 80 | -14.8986 | 4.59330 | 8.13716 |
| CD630_s0590 | CD630_n00080 | 7 -- 80 | 10 -- 98 | -14.8986 | 8.13716 | 4.59330 |
| CD630_SQ1642 | CD630_n00220 | 134 -- 198 | 5 -- 82 | -14.8834 | 8.65138 | 4.94050 |
| CD630_Cdi1_3 | CD630_n01010 | 458 -- 497 | 844 -- 874 | -14.8573 | 6.59405 | 4.59146 |
| CD630_n01010 | CD630_Cdi1_3 | 844 -- 874 | 458 -- 497 | -14.8573 | 4.59146 | 6.59405 |
| CD630_n01010 | CD630_n00560 | 254 -- 337 | 24 -- 85 | -14.8519 | 4.59146 | 5.82053 |
| CD630_n00600 | CD630_n00620 | 792 -- 838 | 55 -- 99 | -14.7638 | 5.31146 | 5.82838 |
| CD630_n00620 | CD630_n00600 | 55 -- 99 | 792 -- 838 | -14.7638 | 5.82838 | 5.31146 |
| CD630_s0010 | CD630_SQ1641 | 58 -- 84 | 5 -- 29 | -14.7487 | 4.92433 | 8.65138 |
| CD630_SQ1641 | CD630_s0010 | 5 -- 29 | 58 -- 84 | -14.7487 | 8.65138 | 4.92433 |
| CD630_n00380 | CD630_Cdi2_2 | 301 -- 331 | 33 -- 61 | -14.707 | 4.27916 | 6.72203 |
| CD630_n00380 | CD630_Cdi2_2 | 301 -- 331 | 33 -- 61 | -14.707 | 4.27916 | 6.72203 |
| CD630_n00990 | CD630_Cdi2_2 | 390 -- 420 | 33 -- 61 | -14.707 | 3.72051 | 6.72203 |
| CD630_n00990 | CD630_Cdi2_2 | 390 -- 420 | 33 -- 61 | -14.707 | 3.72051 | 6.72203 |
| CD630_n00600 | CD630_n00600 | 513 -- 606 | 1009 -- 1111 | -14.703 | 5.31146 | 5.31146 |
| CD630_n00380 | CD630_n00460 | 754 -- 847 | 659 -- 744 | -14.67 | 4.27916 | 5.55447 |
| CD630_n00990 | CD630_n00460 | 843 -- 936 | 659 -- 744 | -14.67 | 3.72051 | 5.55447 |
| CD630_n00560 | CD630_n01010 | 5 -- 52 | 35 -- 77 | -14.6656 | 5.82053 | 4.59146 |
| RCd9 | CD630_s0660 | 93 -- 115 | 106 -- 128 | -14.6588 | 8.96925 | 4.92199 |
| CD630_s0660 | RCd9 | 106 -- 128 | 93 -- 115 | -14.6588 | 4.92199 | 8.96925 |
| CD630_n00080 | CD630_s0010 | 63 -- 93 | 18 -- 47 | -14.6376 | 4.59330 | 4.92433 |
| CD630_n00650 | CD630_n00930 | 53 -- 97 | 27 -- 67 | -14.6308 | 5.32127 | 5.79130 |
| CD630_n00930 | CD630_n00650 | 27 -- 67 | 53 -- 97 | -14.6308 | 5.79130 | 5.32127 |
| CD630_n00380 | CD630_n00680 | 656 -- 701 | 335 -- 388 | -14.6146 | 4.27916 | 6.52781 |
| CD630_n00990 | CD630_n00680 | 745 -- 790 | 335 -- 388 | -14.6146 | 3.72051 | 6.52781 |
| CD630_s0460 | CD630_n00080 | 3 -- 16 | 86 -- 102 | -14.6104 | 5.74527 | 4.59330 |
| CD630_n00460 | RCd5 | 205 -- 231 | 122 -- 148 | -14.5423 | 5.55447 | 6.52781 |
| CD630_n00460 | CD630_n00680 | 205 -- 231 | 122 -- 148 | -14.5423 | 5.55447 | 6.52781 |
| RCd5 | CD630_n00460 | 122 -- 148 | 205 -- 231 | -14.5423 | 6.52781 | 5.55447 |
| CD630_n00680 | CD630_n00460 | 122 -- 148 | 205 -- 231 | -14.5423 | 6.52781 | 5.55447 |
| CD630_n00790 | CD630_s0340 | 199 -- 248 | 20 -- 74 | -14.5413 | 5.51210 | 5.61064 |
| CD630_n00390 | CD630_s0642 | 290 -- 353 | 16 -- 74 | -14.5267 | 4.83391 | 6.00600 |
| CD630_s0642 | CD630_n00390 | 16 -- 74 | 290 -- 353 | -14.5267 | 6.00600 | 4.83391 |
| CD630_s0190 | CD630_n00560 | 67 -- 89 | 465 -- 487 | -14.52 | 4.98480 | 5.82053 |
| CD630_n00560 | CD630_s0190 | 465 -- 487 | 67 -- 89 | -14.52 | 5.82053 | 4.98480 |
| CD630_s0190 | CD630_s0210 | 75 -- 158 | 9 -- 90 | -14.504 | 4.98480 | 4.02122 |
| CD630_s0010 | CD630_n00080 | 18 -- 48 | 58 -- 93 | -14.4958 | 4.92433 | 4.59330 |
| CD630_s0370 | CD630_n00460 | 4 -- 40 | 339 -- 374 | -14.4865 | 10.21495 | 5.55447 |
| CD630_Cdi1_4 | CD630_Cdi2_3 | 141 -- 164 | 146 -- 170 | -14.4757 | 5.31318 | 3.99184 |
| CD630_n00980 | CD630_Cdi2_3 | 141 -- 164 | 146 -- 170 | -14.4757 | 5.31318 | 3.99184 |
| CD630_Cdi1_5 | CD630_Cdi2_3 | 134 -- 157 | 146 -- 170 | -14.4757 | 4.83391 | 3.99184 |
| CD630_n00390 | CD630_Cdi2_3 | 134 -- 157 | 146 -- 170 | -14.4757 | 4.83391 | 3.99184 |
| CD630_Cdi2_3 | CD630_Cdi1_4 | 146 -- 170 | 141 -- 164 | -14.4757 | 3.99184 | 5.31318 |
| CD630_Cdi2_3 | CD630_n00980 | 146 -- 170 | 141 -- 164 | -14.4757 | 3.99184 | 5.31318 |
| CD630_Cdi2_3 | CD630_Cdi1_5 | 146 -- 170 | 134 -- 157 | -14.4757 | 3.99184 | 4.83391 |
| CD630_Cdi2_3 | CD630_n00390 | 146 -- 170 | 134 -- 157 | -14.4757 | 3.99184 | 4.83391 |
| CD630_n00380 | CD630_n00620 | 514 -- 649 | 32 -- 179 | -14.4694 | 4.27916 | 5.82838 |
| CD630_n00990 | CD630_n00620 | 603 -- 738 | 32 -- 179 | -14.4694 | 3.72051 | 5.82838 |
| CD630_s0670 | CD630_s0390 | 26 -- 54 | 220 -- 251 | -14.4285 | 4.56362 | 4.73397 |
| CD630_n00220 | CD630_SQ1642 | 5 -- 45 | 162 -- 198 | -14.4074 | 4.94050 | 8.65138 |
| CD630_s0390 | CD630_n00080 | 186 -- 223 | 68 -- 100 | -14.4041 | 4.73397 | 4.59330 |
| CD630_n00600 | CD630_s0210 | 797 -- 818 | 30 -- 51 | -14.4038 | 5.31146 | 4.02122 |
| CD630_n00080 | CD630_n00460 | 71 -- 101 | 37 -- 77 | -14.4003 | 4.59330 | 5.55447 |
| CD630_n00560 | CD630_n00600 | 54 -- 85 | 676 -- 708 | -14.3318 | 5.82053 | 5.31146 |
| CD630_n00600 | CD630_n00560 | 676 -- 708 | 54 -- 85 | -14.3318 | 5.31146 | 5.82053 |
| CD630_s0280 | CD630_Cdi1_4 | 49 -- 91 | 277 -- 322 | -14.3276 | 5.37459 | 5.31318 |
| CD630_Cdi1_5 | CD630_s0631 | 104 -- 138 | 303 -- 343 | -14.3264 | 4.83391 | 4.95025 |
| CD630_n00390 | CD630_s0631 | 104 -- 138 | 303 -- 343 | -14.3264 | 4.83391 | 4.95025 |
| CD630_s0631 | CD630_Cdi1_5 | 303 -- 343 | 104 -- 138 | -14.3264 | 4.95025 | 4.83391 |
| CD630_s0631 | CD630_n00390 | 303 -- 343 | 104 -- 138 | -14.3264 | 4.95025 | 4.83391 |
| CD630_n00470 | CD630_SQ1002 | 58 -- 81 | 84 -- 107 | -14.3099 | 3.52267 | 4.43255 |
| CD630_SQ1002 | CD630_n00470 | 84 -- 107 | 58 -- 81 | -14.3099 | 4.43255 | 3.52267 |
| CD630_n00600 | CD630_Cdi1_5 | 786 -- 834 | 15 -- 63 | -14.3082 | 5.31146 | 4.83391 |
| CD630_n00600 | CD630_n00390 | 786 -- 834 | 15 -- 63 | -14.3082 | 5.31146 | 4.83391 |
| CD630_Cdi1_5 | CD630_n00600 | 15 -- 63 | 786 -- 834 | -14.3082 | 4.83391 | 5.31146 |
| CD630_n00390 | CD630_n00600 | 15 -- 63 | 786 -- 834 | -14.3082 | 4.83391 | 5.31146 |
| CD630_SQ1642 | CD630_n00510 | 174 -- 202 | 937 -- 966 | -14.3003 | 8.65138 | 3.92762 |
| CD630_Cdi1_4 | CD630_s0631 | 111 -- 145 | 303 -- 343 | -14.2862 | 5.31318 | 4.95025 |
| CD630_n00980 | CD630_s0631 | 111 -- 145 | 303 -- 343 | -14.2862 | 5.31318 | 4.95025 |
| CD630_s0631 | CD630_Cdi1_4 | 303 -- 343 | 111 -- 145 | -14.2862 | 4.95025 | 5.31318 |
| CD630_s0631 | CD630_n00980 | 303 -- 343 | 111 -- 145 | -14.2862 | 4.95025 | 5.31318 |
| CD630_n00690 | CD630_s0210 | 159 -- 230 | 25 -- 87 | -14.2773 | 5.96526 | 4.02122 |
| CD630_n00380 | CD630_Cdi1_5 | 155 -- 190 | 175 -- 209 | -14.2529 | 4.27916 | 4.83391 |
| CD630_n00380 | CD630_n00390 | 155 -- 190 | 175 -- 209 | -14.2529 | 4.27916 | 4.83391 |
| CD630_n00990 | CD630_Cdi1_5 | 244 -- 279 | 175 -- 209 | -14.2529 | 3.72051 | 4.83391 |
| CD630_n00990 | CD630_n00390 | 244 -- 279 | 175 -- 209 | -14.2529 | 3.72051 | 4.83391 |
| CD630_Cdi1_5 | CD630_n00460 | 399 -- 443 | 866 -- 914 | -14.2527 | 4.83391 | 5.55447 |
| CD630_n00690 | CD630_s0450 | 290 -- 328 | 13 -- 49 | -14.2314 | 5.96526 | 5.67265 |
| CD630_s0450 | CD630_n00690 | 13 -- 49 | 290 -- 328 | -14.2314 | 5.67265 | 5.96526 |
| CD630_n00330 | CD630_s0450 | 33 -- 52 | 134 -- 154 | -14.1937 | 6.46464 | 5.67265 |
| CD630_s0450 | CD630_n00330 | 134 -- 154 | 33 -- 52 | -14.1937 | 5.67265 | 6.46464 |
| CD630_n00620 | CD630_n00380 | 37 -- 169 | 319 -- 439 | -14.1669 | 5.82838 | 4.27916 |
| CD630_n00620 | CD630_n00990 | 37 -- 169 | 408 -- 528 | -14.1669 | 5.82838 | 3.72051 |
| CD630_s0210 | CD630_n00600 | 76 -- 86 | 676 -- 686 | -14.1591 | 4.02122 | 5.31146 |
| RCd9 | CD630_s0591 | 224 -- 233 | 122 -- 131 | -14.1558 | 8.96925 | 6.84967 |
| CD630_s0591 | RCd9 | 122 -- 131 | 224 -- 233 | -14.1558 | 6.84967 | 8.96925 |
| CD630_SQ1642 | CD630_s0631 | 118 -- 199 | 13 -- 110 | -14.1456 | 8.65138 | 4.95025 |
| RCd2 | CD630_n00380 | 24 -- 59 | 156 -- 186 | -14.1409 | 3.80984 | 4.27916 |
| RCd2 | CD630_n00990 | 24 -- 59 | 245 -- 275 | -14.1409 | 3.80984 | 3.72051 |
| CD630_n01010 | CD630_s0450 | 795 -- 843 | 6 -- 50 | -14.1349 | 4.59146 | 5.67265 |
| CD630_s0280 | CD630_n00980 | 49 -- 91 | 277 -- 322 | -14.1269 | 5.37459 | 5.31318 |
| CD630_n00510 | RCd8 | 8 -- 24 | 6 -- 22 | -14.1152 | 3.92762 | 6.97567 |
| RCd8 | CD630_n00510 | 6 -- 22 | 8 -- 24 | -14.1152 | 6.97567 | 3.92762 |
| CD630_Cdi1_5 | CD630_s0642 | 290 -- 353 | 16 -- 74 | -14.112 | 4.83391 | 6.00600 |
| CD630_s0642 | CD630_Cdi1_5 | 16 -- 74 | 290 -- 353 | -14.112 | 6.00600 | 4.83391 |
| CD630_n00510 | CD630_SQ1642 | 895 -- 955 | 146 -- 214 | -14.1016 | 3.92762 | 8.65138 |
| CD630_s0340 | CD630_n00080 | 5 -- 17 | 85 -- 100 | -14.0969 | 5.61064 | 4.59330 |
| CD630_n00600 | RCd5 | 797 -- 836 | 7 -- 59 | -14.0931 | 5.31146 | 6.52781 |
| CD630_n00600 | CD630_n00680 | 797 -- 836 | 7 -- 59 | -14.0931 | 5.31146 | 6.52781 |
| RCd5 | CD630_n00600 | 7 -- 59 | 797 -- 836 | -14.0931 | 6.52781 | 5.31146 |
| CD630_n00680 | CD630_n00600 | 7 -- 59 | 797 -- 836 | -14.0931 | 6.52781 | 5.31146 |
| CD630_n00600 | CD630_Cdi1_4 | 786 -- 834 | 22 -- 70 | -14.08 | 5.31146 | 5.31318 |
| CD630_n00600 | CD630_n00980 | 786 -- 834 | 22 -- 70 | -14.08 | 5.31146 | 5.31318 |
| CD630_Cdi1_4 | CD630_n00600 | 22 -- 70 | 786 -- 834 | -14.08 | 5.31318 | 5.31146 |
| CD630_n00980 | CD630_n00600 | 22 -- 70 | 786 -- 834 | -14.08 | 5.31318 | 5.31146 |
| CD630_n00460 | CD630_n00080 | 37 -- 71 | 79 -- 101 | -14.0561 | 5.55447 | 4.59330 |
| CD630_s0281 | RCd5 | 162 -- 202 | 215 -- 262 | -14.0455 | 8.30299 | 6.52781 |
| CD630_s0281 | CD630_n00680 | 162 -- 202 | 215 -- 262 | -14.0455 | 8.30299 | 6.52781 |
| RCd5 | CD630_s0281 | 215 -- 262 | 162 -- 202 | -14.0455 | 6.52781 | 8.30299 |
| CD630_n00680 | CD630_s0281 | 215 -- 262 | 162 -- 202 | -14.0455 | 6.52781 | 8.30299 |
| CD630_n00470 | CD630_n00560 | 21 -- 86 | 454 -- 515 | -14.0446 | 3.52267 | 5.82053 |
| RCd8 | CD630_n01010 | 144 -- 168 | 1260 -- 1285 | -14.0424 | 6.97567 | 4.59146 |
| CD630_n01010 | RCd8 | 1260 -- 1285 | 144 -- 168 | -14.0424 | 4.59146 | 6.97567 |
| CD630_n00380 | RCd2 | 164 -- 204 | 3 -- 49 | -14.0267 | 4.27916 | 3.80984 |
| CD630_n00990 | RCd2 | 253 -- 293 | 3 -- 49 | -14.0267 | 3.72051 | 3.80984 |
| CD630_s0450 | CD630_n01010 | 6 -- 45 | 797 -- 843 | -14.0169 | 5.67265 | 4.59146 |
| CD630_n00440 | CD630_Cdi2_3 | 216 -- 239 | 21 -- 49 | -14.001 | 4.85723 | 3.99184 |
| CD630_Cdi2_3 | CD630_n00440 | 21 -- 49 | 216 -- 239 | -14.001 | 3.99184 | 4.85723 |
| CD630_s0190 | CD630_n01010 | 40 -- 86 | 318 -- 358 | -14.0009 | 4.98480 | 4.59146 |
| CD630_n00080 | CD630_s0340 | 85 -- 101 | 1 -- 17 | -13.991 | 4.59330 | 5.61064 |
| CD630_s0642 | CD630_n00080 | 9 -- 72 | 15 -- 84 | -13.9822 | 6.00600 | 4.59330 |
| CD630_Cdi1_8 | CD630_n01010 | 129 -- 206 | 1344 -- 1415 | -13.9642 | 5.79994 | 4.59146 |
| CD630_n01010 | CD630_Cdi1_8 | 1344 -- 1415 | 129 -- 206 | -13.9642 | 4.59146 | 5.79994 |

| | | | | | | |
|---|---|---|---|---|---|---|
| CD630_n00460 | CD630_Cdi1_5 | 333 -- 373 | 25 -- 64 | -13.9568 | 5.55447 | 4.83391 |
| CD630_n00460 | CD630_n00390 | 333 -- 373 | 25 -- 64 | -13.9568 | 5.55447 | 4.83391 |
| CD630_n00390 | CD630_n00460 | 25 -- 64 | 333 -- 373 | -13.9568 | 4.83391 | 5.55447 |
| CD630_Cdi1_4 | CD630_n00460 | 406 -- 450 | 866 -- 914 | -13.9219 | 5.31318 | 5.55447 |
| CD630_n00560 | CD630_n00470 | 454 -- 499 | 45 -- 86 | -13.8834 | 5.82053 | 3.52267 |
| CD630_n00690 | CD630_Cdi1_8 | 185 -- 328 | 184 -- 310 | -13.8742 | 5.96526 | 5.79994 |
| CD630_n00460 | CD630_Cdi1_4 | 333 -- 373 | 32 -- 71 | -13.8215 | 5.55447 | 5.31318 |
| CD630_n00460 | CD630_n00980 | 333 -- 373 | 32 -- 71 | -13.8215 | 5.55447 | 5.31318 |
| CD630_n00980 | CD630_n00460 | 32 -- 71 | 333 -- 373 | -13.8215 | 5.31318 | 5.55447 |
| CD630_s0631 | CD630_SQ1642 | 13 -- 72 | 148 -- 199 | -13.7898 | 4.95025 | 8.65138 |
| CD630_s0510 | CD630_n00790 | 29 -- 67 | 83 -- 115 | -13.7882 | 3.37631 | 5.51210 |
| CD630_s0210 | CD630_n00560 | 7 -- 31 | 397 -- 421 | -13.7792 | 4.02122 | 5.82053 |
| CD630_n00560 | CD630_s0210 | 397 -- 421 | 7 -- 31 | -13.7792 | 5.82053 | 4.02122 |
| CD630_s0210 | CD630_s0281 | 42 -- 57 | 91 -- 107 | -13.7624 | 4.02122 | 8.30299 |
| CD630_s0281 | CD630_s0210 | 91 -- 107 | 42 -- 57 | -13.7624 | 8.30299 | 4.02122 |
| CD630_n01010 | CD630_s0190 | 318 -- 341 | 60 -- 86 | -13.7404 | 4.59146 | 4.98480 |
| CD630_n00790 | CD630_n00440 | 918 -- 1012 | 157 -- 237 | -13.7267 | 5.51210 | 4.85723 |
| CD630_s0190 | CD630_n00080 | 40 -- 48 | 57 -- 65 | -13.7125 | 4.98480 | 4.59330 |
| CD630_s0270 | CD630_n00080 | 11 -- 65 | 61 -- 101 | -13.7103 | 3.63582 | 4.59330 |
| CD630_n00080 | CD630_s0270 | 61 -- 101 | 11 -- 65 | -13.7103 | 4.59330 | 3.63582 |
| CD630_s0400 | CD630_n00790 | 2 -- 51 | 204 -- 273 | -13.7086 | 4.51086 | 5.51210 |
| CD630_n00790 | CD630_s0400 | 204 -- 273 | 2 -- 51 | -13.7086 | 5.51210 | 4.51086 |
| CD630_s0270 | CD630_s0281 | 112 -- 138 | 167 -- 193 | -13.7073 | 3.63582 | 8.30299 |
| CD630_s0320 | CD630_s0330 | 30 -- 62 | 4 -- 32 | -13.6966 | 8.15991 | 4.37658 |
| CD630_n01060 | CD630_n01010 | 32 -- 119 | 837 -- 919 | -13.6924 | 3.97522 | 4.59146 |
| CD630_n00790 | CD630_s0631 | 202 -- 232 | 8 -- 53 | -13.6897 | 5.51210 | 4.95025 |
| CD630_s0631 | CD630_n00790 | 8 -- 53 | 202 -- 232 | -13.6897 | 4.95025 | 5.51210 |
| CD630_n00220 | CD630_n00440 | 70 -- 100 | 207 -- 240 | -13.6716 | 4.94050 | 4.85723 |
| CD630_n00440 | CD630_n00220 | 207 -- 240 | 70 -- 100 | -13.6716 | 4.85723 | 4.94050 |
| CD630_n00080 | CD630_s0642 | 15 -- 83 | 10 -- 72 | -13.6627 | 4.59330 | 6.00600 |
| CD630_s0660 | CD630_s0370 | 10 -- 71 | 4 -- 55 | -13.6389 | 4.92199 | 10.21495 |
| CD630_s0280 | CD630_s0280 | 214 -- 223 | 214 -- 223 | -13.6375 | 5.37459 | 5.37459 |
| CD630_s0480 | CD630_n00150 | 3 -- 40 | 27 -- 64 | -13.6357 | 4.87999 | 8.81823 |
| CD630_n00150 | CD630_s0480 | 27 -- 64 | 3 -- 40 | -13.6357 | 8.81823 | 4.87999 |
| CD630_SQ1642 | CD630_n00370 | 86 -- 123 | 1 -- 41 | -13.6146 | 8.65138 | 7.05302 |
| CD630_n00370 | CD630_SQ1642 | 1 -- 41 | 86 -- 123 | -13.6146 | 7.05302 | 8.65138 |
| CD630_s0190 | CD630_n00600 | 74 -- 150 | 534 -- 614 | -13.612 | 4.98480 | 5.31146 |
| CD630_n00790 | CD630_s0450 | 97 -- 119 | 24 -- 47 | -13.6045 | 5.51210 | 5.67265 |
| CD630_Cdi1_8 | CD630_s0631 | 113 -- 176 | 288 -- 341 | -13.5981 | 5.79994 | 4.95025 |
| CD630_s0631 | CD630_Cdi1_8 | 288 -- 341 | 113 -- 176 | -13.5981 | 4.95025 | 5.79994 |
| CD630_SQ808 | CD630_n00380 | 30 -- 69 | 196 -- 236 | -13.5856 | 6.85680 | 4.27916 |
| CD630_SQ1642 | RCd9 | 86 -- 123 | 1 -- 41 | -13.5826 | 8.65138 | 8.96925 |
| CD630_n00380 | CD630_n00790 | 515 -- 624 | 976 -- 1074 | -13.5754 | 4.27916 | 5.51210 |
| CD630_n00990 | CD630_n00790 | 604 -- 713 | 976 -- 1074 | -13.5754 | 3.72051 | 5.51210 |
| CD630_n00790 | CD630_n00690 | 90 -- 122 | 8 -- 41 | -13.5742 | 5.51210 | 5.96526 |
| CD630_Cdi1_4 | CD630_n00590 | 382 -- 464 | 22 -- 101 | -13.5579 | 5.31318 | 4.28395 |
| CD630_n00850 | CD630_s0631 | 113 -- 175 | 288 -- 342 | -13.5487 | 5.53585 | 4.95025 |
| CD630_Cdi1_3 | CD630_n00290 | 464 -- 493 | 49 -- 81 | -13.547 | 6.59405 | 3.85358 |
| CD630_s0320 | CD630_n00600 | 1 -- 33 | 538 -- 572 | -13.5436 | 8.15991 | 5.31146 |
| CD630_n00600 | CD630_s0320 | 538 -- 572 | 1 -- 33 | -13.5436 | 5.31146 | 8.15991 |
| CD630_n00790 | CD630_s0510 | 82 -- 122 | 22 -- 68 | -13.5392 | 5.51210 | 3.37631 |
| CD630_s0190 | CD630_n00790 | 55 -- 93 | 198 -- 231 | -13.5255 | 4.98480 | 5.51210 |
| CD630_n00790 | CD630_s0190 | 198 -- 231 | 55 -- 93 | -13.5255 | 5.51210 | 4.98480 |
| CD630_Cdi1_3 | CD630_s0660 | 487 -- 495 | 224 -- 232 | -13.5203 | 6.59405 | 4.92199 |
| CD630_s0660 | CD630_Cdi1_3 | 224 -- 232 | 487 -- 495 | -13.5203 | 4.92199 | 6.59405 |
| CD630_n00440 | RCd9 | 185 -- 229 | 140 -- 184 | -13.5068 | 4.85723 | 8.96925 |
| CD630_n00510 | CD630_SQ1002 | 933 -- 956 | 89 -- 110 | -13.4998 | 3.92762 | 4.43255 |
| CD630_n00690 | CD630_n00790 | 8 -- 23 | 107 -- 122 | -13.4968 | 5.96526 | 5.51210 |
| CD630_n00600 | CD630_n00080 | 39 -- 144 | 3 -- 101 | -13.4916 | 5.31146 | 4.59330 |
| CD630_n00080 | CD630_n00600 | 3 -- 101 | 39 -- 144 | -13.4916 | 4.59330 | 5.31146 |
| CD630_s0400 | CD630_n00460 | 21 -- 71 | 181 -- 227 | -13.4913 | 4.51086 | 5.55447 |
| CD630_n00600 | CD630_n00690 | 673 -- 821 | 146 -- 293 | -13.4841 | 5.31146 | 5.96526 |
| CD630_SQ1642 | CD630_n00590 | 97 -- 197 | 1 -- 100 | -13.4675 | 8.65138 | 4.28395 |
| RCd9 | CD630_SQ1642 | 1 -- 48 | 78 -- 123 | -13.4651 | 8.96925 | 8.65138 |
| CD630_Cdi1_5 | CD630_n00590 | 375 -- 457 | 22 -- 101 | -13.4618 | 4.83391 | 4.28395 |
| RCd9 | CD630_n00440 | 140 -- 173 | 195 -- 229 | -13.4499 | 8.96925 | 4.85723 |
| CD630_n00600 | CD630_s0190 | 123 -- 148 | 169 -- 195 | -13.4441 | 5.31146 | 4.98480 |
| CD630_n00790 | CD630_n00620 | 781 -- 836 | 147 -- 203 | -13.4296 | 5.51210 | 5.82838 |
| CD630_n00620 | CD630_SQ1642 | 40 -- 76 | 163 -- 199 | -13.4246 | 5.82838 | 8.65138 |
| CD630_n00560 | CD630_s0300 | 474 -- 498 | 139 -- 167 | -13.4237 | 5.82053 | 4.17452 |
| CD630_s0300 | CD630_n00560 | 139 -- 167 | 474 -- 498 | -13.4237 | 4.17452 | 5.82053 |
| CD630_n00690 | CD630_n00990 | 93 -- 122 | 132 -- 158 | -13.4146 | 5.96526 | 3.72051 |
| CD630_n00990 | CD630_n00690 | 132 -- 158 | 93 -- 122 | -13.4146 | 3.72051 | 5.96526 |
| CD630_n01010 | CD630_n01060 | 837 -- 918 | 33 -- 119 | -13.3989 | 4.59146 | 3.97522 |
| CD630_Cdi1_10 | CD630_n00290 | 125 -- 158 | 33 -- 62 | -13.3776 | 8.60766 | 3.85358 |
| CD630_n00680 | CD630_n00680 | 2 -- 34 | 339 -- 370 | -13.3737 | 6.52781 | 6.52781 |
| CD630_n00460 | CD630_s0300 | 54 -- 74 | 212 -- 230 | -13.3494 | 5.55447 | 4.17452 |
| CD630_s0300 | CD630_n00460 | 212 -- 230 | 54 -- 74 | -13.3494 | 4.17452 | 5.55447 |
| CD630_SQ1642 | CD630_n00620 | 163 -- 192 | 46 -- 76 | -13.3482 | 8.65138 | 5.82838 |
| CD630_n00850 | CD630_n00080 | 163 -- 185 | 65 -- 85 | -13.3324 | 5.53585 | 4.59330 |
| CD630_n00080 | CD630_s0631 | 61 -- 100 | 23 -- 67 | -13.3271 | 4.59330 | 4.95025 |
| CD630_n00240 | CD630_s0631 | 4 -- 23 | 8 -- 26 | -13.32 | 5.82709 | 4.95025 |
| CD630_s0631 | CD630_n00240 | 8 -- 26 | 4 -- 23 | -13.32 | 4.95025 | 5.82709 |
| CD630_s0281 | CD630_SQ1656 | 12 -- 48 | 15 -- 51 | -13.3097 | 8.30299 | 4.25622 |
| CD630_SQ1656 | CD630_s0281 | 15 -- 51 | 12 -- 48 | -13.3097 | 4.25622 | 8.30299 |
| RCd2 | CD630_n00790 | 3 -- 53 | 125 -- 169 | -13.2769 | 3.80984 | 5.51210 |
| CD630_n00290 | CD630_Cdi1_3 | 126 -- 134 | 488 -- 496 | -13.2747 | 3.85358 | 6.59405 |
| RCd5 | CD630_Cdi1_4 | 404 -- 432 | 375 -- 408 | -13.2661 | 6.52781 | 5.31318 |
| CD630_Cdi1_4 | RCd5 | 375 -- 408 | 404 -- 432 | -13.2661 | 5.31318 | 6.52781 |
| CD630_s0360 | CD630_n00330 | 8 -- 21 | 40 -- 56 | -13.254 | 6.65922 | 6.46464 |
| CD630_n00790 | CD630_n00380 | 200 -- 272 | 598 -- 696 | -13.2198 | 5.51210 | 4.27916 |
| CD630_n00790 | CD630_n00990 | 200 -- 272 | 687 -- 785 | -13.2198 | 5.51210 | 3.72051 |
| CD630_n00600 | CD630_s0660 | 603 -- 624 | 104 -- 129 | -13.1987 | 5.31146 | 4.92199 |
| RCd9 | CD630_n00370 | 257 -- 295 | 1 -- 42 | -13.1987 | 8.96925 | 7.05302 |
| CD630_n00370 | RCd9 | 1 -- 42 | 257 -- 295 | -13.1987 | 7.05302 | 8.96925 |
| CD630_s0660 | CD630_n00600 | 104 -- 129 | 603 -- 624 | -13.1987 | 4.92199 | 5.31146 |
| CD630_n00590 | CD630_Cdi1_4 | 22 -- 58 | 430 -- 464 | -13.1928 | 4.28395 | 5.31318 |
| CD630_SQ1642 | RCd5 | 110 -- 170 | 259 -- 326 | -13.1868 | 8.65138 | 6.52781 |
| CD630_n00380 | CD630_s0642 | 1 -- 46 | 229 -- 274 | -13.1866 | 4.27916 | 6.00600 |
| CD630_s0642 | CD630_n00380 | 229 -- 274 | 1 -- 46 | -13.1866 | 6.00600 | 4.27916 |
| CD630_n00080 | CD630_n00850 | 65 -- 84 | 164 -- 185 | -13.1803 | 4.59330 | 5.53585 |
| CD630_n00460 | CD630_s0510 | 205 -- 249 | 9 -- 51 | -13.1778 | 5.55447 | 3.37631 |
| CD630_s0510 | CD630_n00460 | 9 -- 51 | 205 -- 249 | -13.1778 | 3.37631 | 5.55447 |
| RCd9 | RCd9 | 257 -- 295 | 1 -- 42 | -13.1661 | 8.96925 | 8.96925 |
| CD630_n00460 | CD630_n00600 | 60 -- 147 | 350 -- 425 | -13.1607 | 5.55447 | 5.31146 |
| CD630_n00600 | CD630_n00460 | 350 -- 425 | 60 -- 147 | -13.1607 | 5.31146 | 5.55447 |
| RCd9 | CD630_s0460 | 94 -- 112 | 1 -- 20 | -13.1602 | 8.96925 | 5.74527 |
| CD630_s0460 | RCd9 | 1 -- 20 | 94 -- 112 | -13.1602 | 5.74527 | 8.96925 |
| CD630_SQ327 | CD630_SQ1642 | 90 -- 122 | 162 -- 198 | -13.1491 | 4.62291 | 8.65138 |
| CD630_SQ1642 | CD630_SQ327 | 162 -- 198 | 90 -- 122 | -13.1491 | 8.65138 | 4.62291 |
| CD630_s0300 | CD630_s0631 | 110 -- 171 | 52 -- 102 | -13.1436 | 4.17452 | 4.95025 |
| CD630_s0280 | CD630_Cdi1_5 | 68 -- 91 | 270 -- 295 | -13.1358 | 5.37459 | 4.83391 |
| CD630_Cdi1_5 | CD630_s0280 | 270 -- 295 | 68 -- 91 | -13.1358 | 4.83391 | 5.37459 |
| CD630_SQ1656 | CD630_n01010 | 2 -- 39 | 953 -- 982 | -13.1332 | 4.25622 | 4.59146 |
| CD630_n01010 | CD630_SQ1656 | 953 -- 982 | 2 -- 39 | -13.1332 | 4.59146 | 4.25622 |
| CD630_n00440 | CD630_s0400 | 62 -- 98 | 22 -- 48 | -13.1331 | 4.85723 | 4.51086 |
| CD630_Cdi1_5 | CD630_SQ808 | 106 -- 122 | 150 -- 166 | -13.1274 | 4.83391 | 6.85680 |
| CD630_n00390 | CD630_SQ808 | 106 -- 122 | 150 -- 166 | -13.1274 | 4.83391 | 6.85680 |
| CD630_SQ808 | CD630_Cdi1_5 | 150 -- 166 | 106 -- 122 | -13.1274 | 6.85680 | 4.83391 |
| CD630_SQ808 | CD630_n00390 | 150 -- 166 | 106 -- 122 | -13.1274 | 6.85680 | 4.83391 |
| CD630_n00340 | CD630_n00590 | 1 -- 57 | 26 -- 76 | -13.1229 | 5.83478 | 4.28395 |
| CD630_n00330 | CD630_s0360 | 40 -- 51 | 11 -- 21 | -13.1167 | 6.46464 | 6.65922 |
| CD630_n00790 | RCd2 | 98 -- 168 | 4 -- 84 | -13.1147 | 5.51210 | 3.80984 |
| CD630_s0300 | CD630_n01010 | 127 -- 167 | 62 -- 103 | -13.1127 | 4.17452 | 4.59146 |
| CD630_n01010 | CD630_s0300 | 62 -- 103 | 127 -- 167 | -13.1127 | 4.59146 | 4.17452 |
| CD630_n00620 | CD630_n00790 | 32 -- 105 | 229 -- 305 | -13.1124 | 5.82838 | 5.51210 |
| CD630_s0642 | CD630_s0340 | 146 -- 155 | 4 -- 13 | -13.1091 | 6.00600 | 5.61064 |
| CD630_n00560 | CD630_s0670 | 670 -- 710 | 2 -- 43 | -13.1036 | 5.82053 | 4.56362 |
| CD630_s0670 | CD630_n00560 | 2 -- 43 | 670 -- 710 | -13.1036 | 4.56362 | 5.82053 |
| CD630_n00380 | CD630_SQ808 | 196 -- 240 | 32 -- 69 | -13.0974 | 4.27916 | 6.85680 |
| CD630_Cdi1_8 | CD630_n00080 | 284 -- 321 | 63 -- 97 | -13.0971 | 5.79994 | 4.59330 |
| CD630_n00080 | CD630_Cdi1_8 | 63 -- 97 | 284 -- 321 | -13.0971 | 4.59330 | 5.79994 |
| CD630_Cdi1_4 | CD630_SQ808 | 113 -- 129 | 150 -- 166 | -13.0948 | 5.31318 | 6.85680 |
| CD630_n00980 | CD630_SQ808 | 113 -- 129 | 150 -- 166 | -13.0948 | 5.31318 | 6.85680 |
| CD630_SQ808 | CD630_Cdi1_4 | 150 -- 166 | 113 -- 129 | -13.0948 | 6.85680 | 5.31318 |
| CD630_SQ808 | CD630_n00980 | 150 -- 166 | 113 -- 129 | -13.0948 | 6.85680 | 5.31318 |
| CD630_s0390 | RCd9 | 206 -- 221 | 221 -- 236 | -13.0943 | 4.73397 | 8.96925 |
| RCd9 | CD630_s0390 | 221 -- 236 | 206 -- 221 | -13.0943 | 8.96925 | 4.73397 |
| CD630_n00510 | CD630_n00560 | 6 -- 23 | 104 -- 121 | -13.0936 | 3.92762 | 5.82053 |
| CD630_n00560 | CD630_n00510 | 104 -- 121 | 6 -- 23 | -13.0936 | 5.82053 | 3.92762 |
| CD630_SQ367 | RCd9 | 62 -- 86 | 264 -- 289 | -13.0824 | 4.08290 | 8.96925 |
| RCd9 | CD630_SQ367 | 264 -- 289 | 62 -- 86 | -13.0824 | 8.96925 | 4.08290 |
| CD630_Cdi2_3 | CD630_n00790 | 20 -- 67 | 999 -- 1040 | -13.0721 | 3.99184 | 5.51210 |
| CD630_Cdi1_5 | CD630_s0660 | 392 -- 429 | 37 -- 74 | -13.0633 | 4.83391 | 4.92199 |
| CD630_s0660 | CD630_Cdi1_5 | 37 -- 74 | 392 -- 429 | -13.0633 | 4.92199 | 4.83391 |
| CD630_n01010 | CD630_s0642 | 353 -- 404 | 158 -- 208 | -13.0483 | 4.59146 | 6.00600 |
| CD630_s0642 | CD630_n01010 | 158 -- 208 | 353 -- 404 | -13.0483 | 6.00600 | 4.59146 |
| CD630_n00460 | CD630_n01010 | 9 -- 22 | 60 -- 73 | -13.0454 | 5.55447 | 4.59146 |
| CD630_n01010 | CD630_n00460 | 60 -- 73 | 9 -- 22 | -13.0454 | 4.59146 | 5.55447 |
| CD630_n00460 | CD630_s0400 | 181 -- 217 | 38 -- 71 | -13.0398 | 5.55447 | 4.51086 |

| | | | | | | |
|---|---|---|---|---|---|---|
| CD630_n00380 | CD630_Cdi2_3 | 289 -- 317 | 137 -- 158 | -13.0374 | 4.27916 | 3.99184 |
| CD630_Cdi2_3 | CD630_n00380 | 137 -- 158 | 289 -- 317 | -13.0374 | 3.99184 | 4.27916 |
| CD630_Cdi2_3 | CD630_n00990 | 137 -- 158 | 378 -- 406 | -13.0374 | 3.99184 | 3.72051 |
| CD630_n00990 | CD630_Cdi2_3 | 378 -- 406 | 137 -- 158 | -13.0374 | 3.72051 | 3.99184 |
| CD630_s0281 | CD630_n00600 | 273 -- 299 | 1120 -- 1146 | -13.0274 | 8.30299 | 5.31146 |
| CD630_n00600 | CD630_s0281 | 1120 -- 1146 | 273 -- 299 | -13.0274 | 5.31146 | 8.30299 |
| CD630_Cdi1_3 | CD630_s0190 | 319 -- 336 | 58 -- 76 | -13.024 | 6.59405 | 4.98480 |
| CD630_s0190 | CD630_Cdi1_3 | 58 -- 76 | 319 -- 336 | -13.024 | 4.98480 | 6.59405 |
| CD630_SQ1642 | CD630_s0641 | 140 -- 192 | 68 -- 123 | -13.0169 | 8.65138 | 6.68432 |
| CD630_s0190 | CD630_SQ1642 | 5 -- 87 | 91 -- 178 | -13.0155 | 4.98480 | 8.65138 |
| CD630_SQ1642 | CD630_s0190 | 91 -- 178 | 5 -- 87 | -13.0155 | 8.65138 | 4.98480 |
| CD630_n00080 | CD630_s0190 | 57 -- 97 | 10 -- 48 | -13.0143 | 4.59330 | 4.98480 |
| CD630_SQ808 | CD630_n00470 | 26 -- 58 | 25 -- 56 | -13.0084 | 6.85680 | 3.52267 |
| CD630_n00470 | CD630_SQ808 | 25 -- 56 | 26 -- 58 | -13.0084 | 3.52267 | 6.85680 |
| CD630_Cdi1_8 | CD630_n00690 | 184 -- 280 | 220 -- 328 | -13.0062 | 5.79994 | 5.96526 |
| CD630_Cdi1_4 | CD630_s0510 | 426 -- 470 | 2 -- 49 | -12.9913 | 5.31318 | 3.37631 |
| CD630_s0510 | CD630_Cdi1_4 | 2 -- 49 | 426 -- 470 | -12.9913 | 3.37631 | 5.31318 |
| CD630_n00600 | CD630_s0631 | 518 -- 548 | 10 -- 41 | -12.9858 | 5.31146 | 4.95025 |
| CD630_n00460 | CD630_s0390 | 210 -- 250 | 17 -- 58 | -12.9674 | 5.55447 | 4.73397 |
| CD630_s0390 | CD630_n00460 | 17 -- 58 | 210 -- 250 | -12.9674 | 4.73397 | 5.55447 |
| CD630_n00290 | CD630_Cdi1_10 | 33 -- 60 | 127 -- 158 | -12.9389 | 3.85358 | 8.60766 |
| CD630_n00590 | CD630_Cdi1_5 | 22 -- 58 | 423 -- 457 | -12.9383 | 4.28395 | 4.83391 |
| CD630_s0590 | CD630_s0642 | 5 -- 13 | 146 -- 154 | -12.9371 | 8.13716 | 6.00600 |
| CD630_s0642 | CD630_s0590 | 146 -- 154 | 5 -- 13 | -12.9371 | 6.00600 | 8.13716 |
| CD630_s0281 | CD630_SQ1642 | 11 -- 49 | 157 -- 199 | -12.9215 | 8.30299 | 8.65138 |
| CD630_SQ1642 | CD630_s0281 | 157 -- 199 | 11 -- 49 | -12.9215 | 8.65138 | 8.30299 |
| CD630_s0330 | CD630_s0320 | 4 -- 28 | 37 -- 62 | -12.9164 | 4.37658 | 8.15991 |
| CD630_n00510 | CD630_s0450 | 759 -- 794 | 13 -- 45 | -12.9085 | 3.92762 | 5.67265 |
| CD630_s0450 | CD630_n00510 | 13 -- 45 | 759 -- 794 | -12.9085 | 5.67265 | 3.92762 |
| CD630_n01060 | CD630_SQ1002 | 97 -- 122 | 147 -- 190 | -12.8909 | 3.97522 | 4.43255 |
| CD630_n01010 | RCd5 | 829 -- 872 | 122 -- 159 | -12.8759 | 4.59146 | 6.52781 |
| CD630_n01010 | CD630_n00680 | 829 -- 872 | 122 -- 159 | -12.8759 | 4.59146 | 6.52781 |
| CD630_Cdi1_11 | CD630_n00290 | 134 -- 151 | 4 -- 20 | -12.8612 | 5.77720 | 3.85358 |
| CD630_n01010 | CD630_n00220 | 1122 -- 1174 | 45 -- 97 | -12.8352 | 4.59146 | 4.94050 |
| CD630_Cdi1_5 | CD630_s0510 | 419 -- 463 | 2 -- 49 | -12.8327 | 4.83391 | 3.37631 |
| CD630_s0510 | CD630_Cdi1_5 | 2 -- 49 | 419 -- 463 | -12.8327 | 3.37631 | 4.83391 |
| CD630_s0310 | CD630_Cdi1_5 | 69 -- 96 | 419 -- 448 | -12.8303 | 4.17452 | 4.83391 |
| CD630_Cdi1_5 | CD630_s0310 | 419 -- 448 | 69 -- 96 | -12.8303 | 4.83391 | 4.17452 |
| CD630_s0281 | CD630_n01010 | 271 -- 298 | 447 -- 463 | -12.8258 | 8.30299 | 4.59146 |
| CD630_n01010 | CD630_s0281 | 447 -- 463 | 271 -- 298 | -12.8258 | 4.59146 | 8.30299 |
| CD630_SQ1642 | CD630_n00680 | 110 -- 170 | 259 -- 326 | -12.8158 | 8.65138 | 6.52781 |
| CD630_n00560 | CD630_n00560 | 148 -- 292 | 158 -- 305 | -12.8048 | 5.82053 | 5.82053 |
| CD630_SQ1005 | CD630_s0281 | 87 -- 113 | 167 -- 193 | -12.8047 | 3.63582 | 8.30299 |
| RCd5 | CD630_SQ808 | 52 -- 76 | 40 -- 64 | -12.7961 | 6.52781 | 6.85680 |
| CD630_n00680 | CD630_SQ808 | 52 -- 76 | 40 -- 64 | -12.7961 | 6.52781 | 6.85680 |
| CD630_SQ808 | RCd5 | 40 -- 64 | 52 -- 76 | -12.7961 | 6.85680 | 6.52781 |
| CD630_SQ808 | CD630_n00680 | 40 -- 64 | 52 -- 76 | -12.7961 | 6.85680 | 6.52781 |
| CD630_Cdi1_3 | CD630_n00500 | 485 -- 496 | 89 -- 100 | -12.7896 | 6.59405 | 9.35359 |
| CD630_n00500 | CD630_Cdi1_3 | 89 -- 100 | 485 -- 496 | -12.7896 | 9.35359 | 6.59405 |
| CD630_n00990 | CD630_s0642 | 709 -- 784 | 146 -- 200 | -12.775 | 3.72051 | 6.00600 |
| CD630_SQ173 | CD630_n00380 | 5 -- 73 | 603 -- 676 | -12.7561 | 6.59405 | 4.27916 |
| CD630_SQ173 | CD630_n00990 | 5 -- 73 | 692 -- 765 | -12.7561 | 6.59405 | 3.72051 |
| CD630_n00380 | CD630_SQ173 | 603 -- 676 | 5 -- 73 | -12.7561 | 4.27916 | 6.59405 |
| CD630_n00990 | CD630_SQ173 | 692 -- 765 | 5 -- 73 | -12.7561 | 3.72051 | 6.59405 |
| CD630_s0320 | CD630_n00080 | 9 -- 35 | 85 -- 102 | -12.7524 | 8.15991 | 4.59330 |
| CD630_n00080 | CD630_s0320 | 85 -- 102 | 9 -- 35 | -12.7524 | 4.59330 | 8.15991 |
| CD630_Cdi1_3 | CD630_n00980 | 485 -- 496 | 351 -- 363 | -12.7512 | 6.59405 | 5.31318 |
| CD630_n00980 | CD630_Cdi1_3 | 351 -- 363 | 485 -- 496 | -12.7512 | 5.31318 | 6.59405 |
| CD630_s0631 | CD630_n00850 | 289 -- 343 | 112 -- 174 | -12.7471 | 4.95025 | 5.53585 |
| CD630_n00560 | CD630_n00690 | 88 -- 117 | 11 -- 41 | -12.7407 | 5.82053 | 5.96526 |
| CD630_Cdi1_3 | RCd9 | 483 -- 497 | 215 -- 230 | -12.7235 | 6.59405 | 8.96925 |
| RCd9 | CD630_Cdi1_3 | 215 -- 230 | 483 -- 497 | -12.7235 | 8.96925 | 6.59405 |
| CD630_n00930 | CD630_n00620 | 2 -- 55 | 127 -- 191 | -12.7227 | 5.79130 | 5.82838 |
| CD630_s0281 | CD630_Cdi2_3 | 155 -- 195 | 12 -- 61 | -12.7224 | 8.30299 | 3.99184 |
| CD630_Cdi2_3 | CD630_s0281 | 12 -- 61 | 155 -- 195 | -12.7224 | 3.99184 | 8.30299 |
| CD630_Cdi1_4 | CD630_s0660 | 399 -- 436 | 37 -- 74 | -12.7122 | 5.31318 | 4.92199 |
| CD630_s0660 | CD630_Cdi1_4 | 37 -- 74 | 399 -- 436 | -12.7122 | 4.92199 | 5.31318 |
| CD630_n00390 | CD630_n00470 | 272 -- 325 | 5 -- 52 | -12.7087 | 4.83391 | 3.52267 |
| CD630_n00240 | CD630_Cdi1_10 | 3 -- 78 | 77 -- 151 | -12.7074 | 5.82709 | 8.60766 |
| CD630_n00600 | CD630_s0641 | 535 -- 565 | 171 -- 194 | -12.7073 | 5.31146 | 6.68432 |
| CD630_s0641 | CD630_n00600 | 171 -- 194 | 535 -- 565 | -12.7073 | 6.68432 | 5.31146 |
| CD630_Cdi1_11 | CD630_n00330 | 122 -- 150 | 35 -- 59 | -12.6825 | 5.77720 | 6.46464 |
| CD630_n00330 | CD630_Cdi1_11 | 35 -- 59 | 122 -- 150 | -12.6825 | 6.46464 | 5.77720 |
| CD630_SQ367 | CD630_n00850 | 67 -- 88 | 382 -- 403 | -12.6773 | 4.08290 | 5.53585 |
| CD630_n00850 | CD630_SQ367 | 382 -- 403 | 67 -- 88 | -12.6773 | 5.53585 | 4.08290 |
| CD630_n01010 | CD630_n00790 | 1028 -- 1175 | 699 -- 821 | -12.6666 | 4.59146 | 5.51210 |
| CD630_n00080 | CD630_s0390 | 68 -- 91 | 199 -- 223 | -12.6572 | 4.59330 | 4.73397 |
| CD630_s0400 | CD630_n00440 | 22 -- 49 | 60 -- 98 | -12.6563 | 4.51086 | 4.85723 |
| CD630_Cdi1_10 | CD630_n00240 | 77 -- 147 | 7 -- 78 | -12.644 | 8.60766 | 5.82709 |
| CD630_Cdi1_3 | CD630_n00790 | 458 -- 495 | 206 -- 241 | -12.6401 | 6.59405 | 5.51210 |
| CD630_n00790 | CD630_Cdi1_3 | 206 -- 241 | 458 -- 495 | -12.6401 | 5.51210 | 6.59405 |
| CD630_n00290 | CD630_s0360 | 115 -- 136 | 9 -- 28 | -12.6361 | 3.85358 | 6.65922 |
| CD630_Cdi1_3 | CD630_n00390 | 485 -- 496 | 344 -- 356 | -12.6332 | 6.59405 | 4.83391 |
| CD630_n00390 | CD630_Cdi1_3 | 344 -- 356 | 485 -- 496 | -12.6332 | 4.83391 | 6.59405 |
| CD630_s0631 | CD630_n00080 | 23 -- 64 | 64 -- 100 | -12.6261 | 4.95025 | 4.59330 |
| CD630_n00460 | CD630_Cdi1_8 | 395 -- 462 | 237 -- 296 | -12.6237 | 5.55447 | 5.79994 |
| CD630_Cdi1_8 | CD630_n00460 | 237 -- 296 | 395 -- 462 | -12.6237 | 5.79994 | 5.55447 |
| CD630_SQ1641 | CD630_n00380 | 15 -- 46 | 486 -- 521 | -12.6209 | 8.65138 | 4.27916 |
| CD630_n00380 | CD630_n00510 | 294 -- 332 | 936 -- 975 | -12.6181 | 4.27916 | 3.92762 |
| CD630_n00510 | CD630_n00380 | 936 -- 975 | 294 -- 332 | -12.6181 | 3.92762 | 4.27916 |
| CD630_n00510 | CD630_n00990 | 936 -- 975 | 383 -- 421 | -12.6181 | 3.92762 | 3.72051 |
| CD630_n00990 | CD630_n00510 | 383 -- 421 | 936 -- 975 | -12.6181 | 3.72051 | 3.92762 |
| CD630_n00790 | CD630_n00790 | 159 -- 212 | 623 -- 690 | -12.6139 | 5.51210 | 5.51210 |
| CD630_s0360 | CD630_n00290 | 9 -- 20 | 125 -- 136 | -12.6128 | 6.65922 | 3.85358 |
| CD630_n00560 | CD630_Cdi1_8 | 332 -- 364 | 226 -- 261 | -12.6126 | 5.82053 | 5.79994 |
| CD630_Cdi1_8 | CD630_n00560 | 226 -- 261 | 332 -- 364 | -12.6126 | 5.79994 | 5.82053 |
| CD630_s0641 | CD630_n00790 | 66 -- 174 | 128 -- 255 | -12.6068 | 6.68432 | 5.51210 |
| CD630_n00790 | CD630_s0641 | 128 -- 255 | 66 -- 174 | -12.6068 | 5.51210 | 6.68432 |
| CD630_n00510 | CD630_Cdi1_4 | 804 -- 897 | 127 -- 212 | -12.603 | 3.92762 | 5.31318 |
| CD630_n00510 | CD630_n00980 | 804 -- 897 | 127 -- 212 | -12.603 | 3.92762 | 5.31318 |
| CD630_n00510 | CD630_Cdi1_5 | 804 -- 897 | 120 -- 205 | -12.603 | 3.92762 | 4.83391 |
| CD630_n00510 | CD630_n00390 | 804 -- 897 | 120 -- 205 | -12.603 | 3.92762 | 4.83391 |
| CD630_Cdi1_4 | CD630_n00510 | 127 -- 212 | 804 -- 897 | -12.603 | 5.31318 | 3.92762 |
| CD630_n00980 | CD630_n00510 | 127 -- 212 | 804 -- 897 | -12.603 | 5.31318 | 3.92762 |
| CD630_Cdi1_5 | CD630_n00510 | 120 -- 205 | 804 -- 897 | -12.603 | 4.83391 | 3.92762 |
| CD630_n00390 | CD630_n00510 | 120 -- 205 | 804 -- 897 | -12.603 | 4.83391 | 3.92762 |
| CD630_SQ1002 | CD630_n00510 | 89 -- 104 | 941 -- 956 | -12.5815 | 4.43255 | 3.92762 |
| CD630_n00380 | CD630_SQ1641 | 486 -- 528 | 7 -- 46 | -12.5677 | 4.27916 | 8.65138 |
| RCd5 | CD630_SQ1642 | 262 -- 325 | 111 -- 168 | -12.5657 | 6.52781 | 8.65138 |
| CD630_Cdi1_5 | CD630_s0210 | 101 -- 131 | 21 -- 49 | -12.5636 | 4.83391 | 4.02122 |
| CD630_n00390 | CD630_s0210 | 101 -- 131 | 21 -- 49 | -12.5636 | 4.83391 | 4.02122 |
| CD630_n01010 | RCd9 | 15 -- 118 | 140 -- 231 | -12.5525 | 4.59146 | 8.96925 |
| CD630_Cdi1_3 | CD630_s0631 | 170 -- 220 | 196 -- 244 | -12.5399 | 6.59405 | 4.95025 |
| CD630_s0631 | CD630_Cdi1_3 | 196 -- 244 | 170 -- 220 | -12.5399 | 4.95025 | 6.59405 |
| CD630_Cdi1_5 | CD630_n00470 | 272 -- 325 | 5 -- 52 | -12.5332 | 4.83391 | 3.52267 |
| CD630_s0340 | CD630_s0642 | 4 -- 24 | 131 -- 155 | -12.5254 | 5.61064 | 6.00600 |
| CD630_Cdi1_4 | CD630_s0210 | 108 -- 138 | 21 -- 49 | -12.5222 | 5.31318 | 4.02122 |
| CD630_n00980 | CD630_s0210 | 108 -- 138 | 21 -- 49 | -12.5222 | 5.31318 | 4.02122 |
| CD630_n00460 | CD630_SQ1641 | 103 -- 134 | 71 -- 120 | -12.5173 | 5.55447 | 8.65138 |
| RCd9 | CD630_Cdi1_10 | 215 -- 234 | 128 -- 152 | -12.5139 | 8.96925 | 8.60766 |
| CD630_Cdi1_10 | RCd9 | 128 -- 152 | 215 -- 234 | -12.5139 | 8.60766 | 8.96925 |
| CD630_SQ1038 | CD630_n00340 | 111 -- 170 | 2 -- 65 | -12.5103 | 3.49019 | 5.83478 |
| CD630_n00340 | CD630_SQ1038 | 2 -- 65 | 111 -- 170 | -12.5103 | 5.83478 | 3.49019 |
| CD630_Cdi1_8 | CD630_SQ808 | 95 -- 131 | 150 -- 191 | -12.5099 | 5.79994 | 6.85680 |
| RCd5 | RCd8 | 270 -- 331 | 55 -- 110 | -12.5097 | 6.52781 | 6.97567 |
| CD630_s0310 | CD630_Cdi1_4 | 69 -- 96 | 426 -- 455 | -12.5024 | 4.17452 | 5.31318 |
| CD630_Cdi1_4 | CD630_s0310 | 426 -- 455 | 69 -- 96 | -12.5024 | 5.31318 | 4.17452 |
| CD630_s0450 | CD630_n00790 | 143 -- 152 | 102 -- 111 | -12.4966 | 5.67265 | 5.51210 |
| CD630_s0280 | CD630_s0510 | 11 -- 53 | 26 -- 75 | -12.4844 | 5.37459 | 3.37631 |
| CD630_s0510 | CD630_s0280 | 26 -- 75 | 11 -- 53 | -12.4844 | 3.37631 | 5.37459 |
| RCd2 | CD630_Cdi2_3 | 19 -- 57 | 22 -- 61 | -12.4789 | 3.80984 | 3.99184 |
| CD630_Cdi2_3 | RCd2 | 22 -- 61 | 19 -- 57 | -12.4789 | 3.99184 | 3.80984 |
| CD630_s0300 | CD630_n00980 | 28 -- 60 | 349 -- 386 | -12.474 | 4.17452 | 5.31318 |
| CD630_n00980 | CD630_s0300 | 349 -- 386 | 28 -- 60 | -12.474 | 5.31318 | 4.17452 |
| CD630_SQ367 | CD630_SQ1002 | 162 -- 185 | 84 -- 108 | -12.4309 | 4.08290 | 4.43255 |
| CD630_SQ1002 | CD630_SQ367 | 84 -- 108 | 162 -- 185 | -12.4309 | 4.43255 | 4.08290 |
| CD630_s0280 | CD630_n00390 | 54 -- 91 | 270 -- 304 | -12.4258 | 5.37459 | 4.83391 |
| CD630_n00390 | CD630_s0280 | 270 -- 304 | 54 -- 91 | -12.4258 | 4.83391 | 5.37459 |
| CD630_Cdi2_3 | CD630_n00080 | 52 -- 63 | 85 -- 97 | -12.4204 | 3.99184 | 4.59330 |
| CD630_n00680 | RCd8 | 270 -- 331 | 55 -- 110 | -12.414 | 6.52781 | 6.97567 |
| CD630_Cdi2_3 | CD630_n01010 | 30 -- 65 | 847 -- 880 | -12.4135 | 3.99184 | 4.59146 |
| CD630_n01010 | CD630_Cdi2_3 | 847 -- 880 | 30 -- 65 | -12.4135 | 4.59146 | 3.99184 |
| CD630_n00460 | RCd8 | 647 -- 673 | 186 -- 206 | -12.4081 | 5.55447 | 6.97567 |
| CD630_n00460 | RCd9 | 644 -- 673 | 95 -- 119 | -12.4068 | 5.55447 | 8.96925 |
| CD630_n00600 | CD630_Cdi2_3 | 802 -- 839 | 30 -- 64 | -12.4023 | 5.31146 | 3.99184 |
| CD630_Cdi2_3 | CD630_n00600 | 30 -- 64 | 802 -- 839 | -12.4023 | 3.99184 | 5.31146 |
| CD630_n00790 | CD630_Cdi2_3 | 1000 -- 1035 | 25 -- 66 | -12.3938 | 5.51210 | 3.99184 |
| CD630_n00600 | CD630_Cdi1_3 | 122 -- 152 | 179 -- 212 | -12.384 | 5.31146 | 6.59405 |
| CD630_n00790 | CD630_n01010 | 699 -- 746 | 1117 -- 1175 | -12.37 | 5.51210 | 4.59146 |
| CD630_SQ1656 | CD630_n01060 | 7 -- 43 | 52 -- 100 | -12.3653 | 4.25622 | 3.97522 |

| | | | | | | |
|---|---|---|---|---|---|---|
| CD630_n01060 | CD630_SQ1656 | 52 -- 100 | 7 -- 43 | -12.3653 | 3.97522 | 4.25622 |
| CD630_s0641 | CD630_SQ2429 | 244 -- 286 | 115 -- 160 | -12.3588 | 6.68432 | 5.77720 |
| CD630_s0300 | CD630_n00390 | 28 -- 60 | 342 -- 379 | -12.3575 | 4.17452 | 4.83391 |
| CD630_n00390 | CD630_s0300 | 342 -- 379 | 28 -- 60 | -12.3575 | 4.83391 | 4.17452 |
| CD630_n00600 | CD630_s0642 | 107 -- 138 | 191 -- 226 | -12.3554 | 5.31146 | 6.00600 |
| CD630_s0642 | CD630_n00600 | 191 -- 226 | 107 -- 138 | -12.3554 | 6.00600 | 5.31146 |
| RCd8 | CD630_s0450 | 189 -- 209 | 167 -- 186 | -12.3506 | 6.97567 | 5.67265 |
| CD630_s0450 | RCd8 | 167 -- 186 | 189 -- 209 | -12.3506 | 5.67265 | 6.97567 |
| CD630_SQ367 | CD630_n00440 | 145 -- 192 | 196 -- 236 | -12.3248 | 4.08290 | 4.85723 |
| CD630_n00440 | CD630_SQ367 | 196 -- 236 | 145 -- 192 | -12.3248 | 4.85723 | 4.08290 |
| CD630_s0210 | CD630_n00790 | 33 -- 51 | 207 -- 224 | -12.3243 | 4.02122 | 5.51210 |
| CD630_n00790 | CD630_s0210 | 207 -- 224 | 33 -- 51 | -12.3243 | 5.51210 | 4.02122 |
| CD630_Cdi1_3 | CD630_n00600 | 179 -- 201 | 129 -- 152 | -12.3185 | 6.59405 | 5.31146 |
| CD630_n00460 | CD630_n00620 | 338 -- 377 | 55 -- 94 | -12.317 | 5.55447 | 5.82838 |
| CD630_n00620 | CD630_n00460 | 55 -- 94 | 338 -- 377 | -12.317 | 5.82838 | 5.55447 |
| CD630_n00590 | CD630_SQ1642 | 1 -- 80 | 117 -- 197 | -12.3092 | 4.28395 | 8.65138 |
| RCd8 | CD630_n00600 | 1 -- 22 | 9 -- 31 | -12.3033 | 6.97567 | 5.31146 |
| CD630_s0280 | CD630_s0330 | 78 -- 88 | 7 -- 17 | -12.3025 | 5.37459 | 4.37658 |
| CD630_s0330 | CD630_s0280 | 7 -- 17 | 78 -- 88 | -12.3025 | 4.37658 | 5.37459 |
| CD630_n00470 | CD630_n00390 | 5 -- 44 | 283 -- 325 | -12.2974 | 3.52267 | 4.83391 |
| CD630_SQ1656 | CD630_s0631 | 141 -- 197 | 60 -- 118 | -12.2931 | 4.25622 | 4.95025 |
| CD630_n00470 | CD630_n01010 | 1 -- 83 | 248 -- 326 | -12.2755 | 3.52267 | 4.59146 |
| CD630_SQ1642 | CD630_n00690 | 82 -- 201 | 37 -- 141 | -12.2748 | 8.65138 | 5.96526 |
| CD630_n00690 | CD630_SQ1642 | 37 -- 141 | 82 -- 201 | -12.2748 | 5.96526 | 8.65138 |
| CD630_n00460 | CD630_s0642 | 557 -- 593 | 190 -- 225 | -12.2741 | 5.55447 | 6.00600 |
| CD630_s0642 | CD630_n00460 | 190 -- 225 | 557 -- 593 | -12.2741 | 6.00600 | 5.55447 |
| CD630_n00290 | CD630_Cdi1_11 | 4 -- 15 | 140 -- 151 | -12.2633 | 3.85358 | 5.77720 |
| CD630_n00600 | CD630_SQ2429 | 797 -- 836 | 105 -- 157 | -12.2614 | 5.31146 | 5.77720 |
| CD630_SQ2429 | CD630_n00600 | 105 -- 157 | 797 -- 836 | -12.2614 | 5.77720 | 5.31146 |
| CD630_n00500 | CD630_s0631 | 95 -- 101 | 334 -- 340 | -12.2459 | 9.35359 | 4.95025 |
| CD630_s0631 | CD630_n00500 | 334 -- 340 | 95 -- 101 | -12.2459 | 4.95025 | 9.35359 |
| CD630_n00440 | CD630_n00990 | 63 -- 75 | 191 -- 203 | -12.2435 | 4.85723 | 3.72051 |
| CD630_n00990 | CD630_n00440 | 191 -- 203 | 63 -- 75 | -12.2435 | 3.72051 | 4.85723 |
| CD630_n00510 | CD630_n00600 | 333 -- 404 | 478 -- 551 | -12.2409 | 3.92762 | 5.31146 |
| CD630_n00380 | CD630_n00690 | 43 -- 69 | 93 -- 122 | -12.2399 | 4.27916 | 5.96526 |
| CD630_n00690 | CD630_n00380 | 93 -- 122 | 43 -- 69 | -12.2399 | 5.96526 | 4.27916 |
| CD630_n00460 | CD630_s0281 | 211 -- 220 | 289 -- 298 | -12.2387 | 5.55447 | 8.30299 |
| CD630_s0281 | CD630_n00460 | 289 -- 298 | 211 -- 220 | -12.2387 | 8.30299 | 5.55447 |
| CD630_n00690 | CD630_n00560 | 11 -- 24 | 104 -- 117 | -12.231 | 5.96526 | 5.82053 |
| CD630_n00620 | CD630_n00930 | 127 -- 176 | 17 -- 55 | -12.2283 | 5.82838 | 5.79130 |
| CD630_SQ1002 | CD630_n01060 | 147 -- 171 | 97 -- 122 | -12.2031 | 4.43255 | 3.97522 |
| CD630_n00460 | CD630_Cdi1_11 | 291 -- 314 | 122 -- 147 | -12.1979 | 5.55447 | 5.77720 |
| CD630_Cdi1_11 | CD630_n00460 | 122 -- 147 | 291 -- 314 | -12.1979 | 5.77720 | 5.55447 |
| CD630_s0010 | CD630_n00330 | 155 -- 163 | 39 -- 47 | -12.1948 | 4.92433 | 6.46464 |
| CD630_n00330 | CD630_s0010 | 39 -- 47 | 155 -- 163 | -12.1948 | 6.46464 | 4.92433 |
| CD630_n00380 | CD630_n00440 | 832 -- 851 | 1 -- 18 | -12.1812 | 4.27916 | 4.85723 |
| CD630_n00440 | CD630_n00380 | 1 -- 18 | 832 -- 851 | -12.1812 | 4.85723 | 4.27916 |
| CD630_n00460 | CD630_s0310 | 98 -- 109 | 196 -- 207 | -12.1772 | 5.55447 | 4.17452 |
| CD630_s0310 | CD630_n00460 | 196 -- 207 | 98 -- 109 | -12.1772 | 4.17452 | 5.55447 |
| CD630_n00600 | CD630_n00930 | 1136 -- 1172 | 5 -- 39 | -12.1769 | 5.31146 | 5.79130 |
| CD630_n00930 | CD630_n00600 | 5 -- 39 | 1136 -- 1172 | -12.1769 | 5.79130 | 5.31146 |
| CD630_SQ1656 | CD630_n00790 | 11 -- 44 | 583 -- 614 | -12.1745 | 4.25622 | 5.51210 |
| CD630_n00680 | CD630_SQ1642 | 259 -- 325 | 111 -- 170 | -12.1712 | 6.52781 | 8.65138 |
| CD630_Cdi1_4 | CD630_n00380 | 111 -- 213 | 600 -- 700 | -12.1257 | 5.31318 | 4.27916 |
| CD630_Cdi1_4 | CD630_n00990 | 111 -- 213 | 689 -- 789 | -12.1257 | 5.31318 | 3.72051 |
| CD630_n00980 | CD630_n00380 | 111 -- 213 | 600 -- 700 | -12.1257 | 5.31318 | 4.27916 |
| CD630_n00980 | CD630_n00990 | 111 -- 213 | 689 -- 789 | -12.1257 | 5.31318 | 3.72051 |
| CD630_Cdi1_5 | CD630_n00380 | 104 -- 206 | 600 -- 700 | -12.1257 | 4.83391 | 4.27916 |
| CD630_Cdi1_5 | CD630_n00990 | 104 -- 206 | 689 -- 789 | -12.1257 | 4.83391 | 3.72051 |
| CD630_n00390 | CD630_n00380 | 104 -- 206 | 600 -- 700 | -12.1257 | 4.83391 | 4.27916 |
| CD630_n00390 | CD630_n00990 | 104 -- 206 | 689 -- 789 | -12.1257 | 4.83391 | 3.72051 |
| CD630_n00600 | CD630_SQ1656 | 1002 -- 1023 | 7 -- 29 | -12.1093 | 5.31146 | 4.25622 |
| CD630_SQ1656 | CD630_n00600 | 7 -- 29 | 1002 -- 1023 | -12.1093 | 4.25622 | 5.31146 |
| CD630_n00440 | CD630_n01060 | 215 -- 240 | 95 -- 121 | -12.1026 | 4.85723 | 3.97522 |
| CD630_n01060 | CD630_n00440 | 95 -- 121 | 215 -- 240 | -12.1026 | 3.97522 | 4.85723 |
| CD630_s0281 | CD630_SQ1005 | 158 -- 210 | 70 -- 120 | -12.0898 | 8.30299 | 3.63582 |
| CD630_SQ1642 | CD630_s0450 | 117 -- 181 | 27 -- 90 | -12.0882 | 8.65138 | 5.67265 |
| CD630_n00220 | CD630_n00240 | 1 -- 19 | 1 -- 19 | -12.0805 | 4.94050 | 5.82709 |
| CD630_n00240 | CD630_n00220 | 1 -- 19 | 1 -- 19 | -12.0805 | 5.82709 | 4.94050 |
| CD630_s0340 | CD630_n01010 | 1 -- 14 | 450 -- 463 | -12.0783 | 5.61064 | 4.59146 |
| CD630_n01010 | CD630_s0340 | 450 -- 463 | 1 -- 14 | -12.0783 | 4.59146 | 5.61064 |
| CD630_n00470 | CD630_Cdi1_5 | 5 -- 44 | 283 -- 325 | -12.0609 | 3.52267 | 4.83391 |
| CD630_Cdi1_4 | CD630_s0280 | 277 -- 302 | 68 -- 91 | -12.0578 | 5.31318 | 5.37459 |
| CD630_s0281 | CD630_s0270 | 158 -- 210 | 95 -- 145 | -12.0525 | 8.30299 | 3.63582 |
| CD630_SQ1005 | CD630_n00690 | 188 -- 204 | 125 -- 141 | -12.0521 | 3.63582 | 5.96526 |
| CD630_n00690 | CD630_SQ1005 | 125 -- 141 | 188 -- 204 | -12.0521 | 5.96526 | 3.63582 |
| CD630_s0641 | CD630_SQ1656 | 244 -- 286 | 122 -- 167 | -12.0519 | 6.68432 | 4.25622 |
| RCd5 | CD630_n01010 | 258 -- 279 | 833 -- 860 | -12.0493 | 6.52781 | 4.59146 |
| CD630_s0270 | CD630_n00690 | 213 -- 229 | 125 -- 141 | -12.0367 | 3.63582 | 5.96526 |
| CD630_n00690 | CD630_s0270 | 125 -- 141 | 213 -- 229 | -12.0367 | 5.96526 | 3.63582 |
| CD630_s0642 | CD630_n00990 | 229 -- 274 | 90 -- 135 | -12.0367 | 6.00600 | 3.72051 |
| CD630_n00510 | CD630_n00510 | 104 -- 122 | 6 -- 24 | -12.0085 | 3.92762 | 3.92762 |
| CD630_s0210 | CD630_Cdi1_4 | 12 -- 59 | 76 -- 118 | -12.0057 | 4.02122 | 5.31318 |
| CD630_s0210 | CD630_n00980 | 12 -- 59 | 76 -- 118 | -12.0057 | 4.02122 | 5.31318 |
| CD630_n01010 | CD630_s0460 | 1086 -- 1128 | 40 -- 82 | -11.9835 | 4.59146 | 5.74527 |
| CD630_Cdi1_8 | CD630_n00290 | 136 -- 153 | 4 -- 20 | -11.9808 | 5.79994 | 3.85358 |
| CD630_n01010 | CD630_s0390 | 1077 -- 1133 | 183 -- 236 | -11.976 | 4.59146 | 4.73397 |
| CD630_n00560 | CD630_n00930 | 657 -- 677 | 332 -- 353 | -11.97 | 5.82053 | 5.79130 |
| CD630_n00930 | CD630_n00560 | 332 -- 353 | 657 -- 677 | -11.97 | 5.79130 | 5.82053 |
| CD630_SQ327 | CD630_n00380 | 86 -- 142 | 877 -- 925 | -11.9688 | 4.62291 | 4.27916 |
| CD630_SQ327 | CD630_n00990 | 86 -- 142 | 966 -- 1014 | -11.9688 | 4.62291 | 3.72051 |
| CD630_n00380 | CD630_SQ327 | 877 -- 925 | 86 -- 142 | -11.9688 | 4.27916 | 4.62291 |
| CD630_n00990 | CD630_SQ327 | 966 -- 1014 | 86 -- 142 | -11.9688 | 3.72051 | 4.62291 |
| CD630_n00680 | CD630_n01010 | 122 -- 148 | 843 -- 872 | -11.9677 | 6.52781 | 4.59146 |
| CD630_s0210 | CD630_Cdi1_5 | 12 -- 59 | 69 -- 111 | -11.9654 | 4.02122 | 4.83391 |
| CD630_s0210 | CD630_n00390 | 12 -- 59 | 69 -- 111 | -11.9654 | 4.02122 | 4.83391 |
| CD630_n00560 | CD630_Cdi2_3 | 445 -- 484 | 24 -- 57 | -11.9653 | 5.82053 | 3.99184 |
| CD630_Cdi2_3 | CD630_n00560 | 24 -- 57 | 445 -- 484 | -11.9653 | 3.99184 | 5.82053 |
| CD630_SQ367 | CD630_n00150 | 159 -- 181 | 42 -- 62 | -11.963 | 4.08290 | 8.81823 |
| CD630_n00150 | CD630_SQ367 | 42 -- 62 | 159 -- 181 | -11.963 | 8.81823 | 4.08290 |
| CD630_RNA_5 | CD630_s0330 | 7 -- 18 | 28 -- 39 | -11.9497 | 4.62291 | 4.37658 |
| CD630_s0330 | CD630_RNA_5 | 28 -- 39 | 7 -- 18 | -11.9497 | 4.37658 | 4.62291 |
| CD630_s0510 | CD630_n01010 | 9 -- 61 | 833 -- 884 | -11.943 | 3.37631 | 4.59146 |
| CD630_n01010 | CD630_s0510 | 833 -- 884 | 9 -- 61 | -11.943 | 4.59146 | 3.37631 |
| CD630_s0510 | CD630_s0660 | 8 -- 58 | 201 -- 242 | -11.9395 | 3.37631 | 4.92199 |
| CD630_s0660 | CD630_s0510 | 201 -- 242 | 8 -- 58 | -11.9395 | 4.92199 | 3.37631 |
| CD630_SQ1642 | CD630_n00500 | 87 -- 123 | 1 -- 40 | -11.9393 | 8.65138 | 9.35359 |
| CD630_n00500 | CD630_SQ1642 | 1 -- 40 | 87 -- 123 | -11.9393 | 9.35359 | 8.65138 |
| CD630_Cdi1_4 | CD630_s0591 | 460 -- 469 | 123 -- 132 | -11.9388 | 5.31318 | 6.84967 |
| CD630_s0591 | CD630_Cdi1_4 | 123 -- 132 | 460 -- 469 | -11.9388 | 6.84967 | 5.31318 |
| CD630_s0641 | CD630_Cdi1_5 | 244 -- 286 | 9 -- 54 | -11.9289 | 6.68432 | 4.83391 |
| CD630_s0641 | CD630_n00390 | 244 -- 286 | 9 -- 54 | -11.9289 | 6.68432 | 4.83391 |
| CD630_n00600 | CD630_Cdi1_10 | 797 -- 836 | 6 -- 58 | -11.9144 | 5.31146 | 8.60766 |
| CD630_SQ1642 | CD630_s0642 | 139 -- 155 | 32 -- 48 | -11.9078 | 8.65138 | 6.00600 |
| CD630_s0642 | CD630_SQ1642 | 32 -- 48 | 139 -- 155 | -11.9078 | 6.00600 | 8.65138 |
| CD630_s0280 | CD630_s0190 | 75 -- 94 | 75 -- 95 | -11.9014 | 5.37459 | 4.98480 |
| CD630_Cdi2_3 | CD630_n00460 | 32 -- 64 | 718 -- 749 | -11.8944 | 3.99184 | 5.55447 |
| RCd9 | CD630_n00500 | 255 -- 295 | 1 -- 46 | -11.8941 | 8.96925 | 9.35359 |
| CD630_n00500 | RCd9 | 1 -- 46 | 255 -- 295 | -11.8941 | 9.35359 | 8.96925 |
| CD630_n00440 | CD630_n00460 | 219 -- 237 | 585 -- 602 | -11.8936 | 4.85723 | 5.55447 |
| CD630_n00460 | CD630_n00440 | 585 -- 602 | 219 -- 237 | -11.8936 | 5.55447 | 4.85723 |
| CD630_n00440 | CD630_n00470 | 13 -- 26 | 1 -- 13 | -11.8907 | 4.85723 | 3.52267 |
| CD630_n00470 | CD630_n00440 | 1 -- 13 | 13 -- 26 | -11.8907 | 3.52267 | 4.85723 |
| CD630_s0641 | CD630_Cdi1_4 | 244 -- 286 | 16 -- 61 | -11.8899 | 6.68432 | 5.31318 |
| CD630_s0641 | CD630_n00980 | 244 -- 286 | 16 -- 61 | -11.8899 | 6.68432 | 5.31318 |
| CD630_SQ1642 | CD630_s0660 | 4 -- 17 | 47 -- 60 | -11.8851 | 8.65138 | 4.92199 |
| CD630_s0660 | CD630_SQ1642 | 47 -- 60 | 4 -- 17 | -11.8851 | 4.92199 | 8.65138 |
| CD630_s0631 | CD630_s0670 | 331 -- 341 | 77 -- 87 | -11.882 | 4.95025 | 4.56362 |
| CD630_s0670 | CD630_s0631 | 77 -- 87 | 331 -- 341 | -11.882 | 4.56362 | 4.95025 |
| CD630_s0190 | CD630_s0641 | 55 -- 84 | 295 -- 323 | -11.8691 | 4.98480 | 6.68432 |
| CD630_s0641 | CD630_s0190 | 295 -- 323 | 55 -- 84 | -11.8691 | 6.68432 | 4.98480 |
| CD630_s0660 | CD630_s0281 | 223 -- 242 | 276 -- 301 | -11.8659 | 4.92199 | 8.30299 |
| CD630_s0281 | CD630_s0660 | 276 -- 301 | 223 -- 242 | -11.8658 | 8.30299 | 4.92199 |
| CD630_SQ1642 | CD630_n00930 | 170 -- 193 | 4 -- 26 | -11.8627 | 8.65138 | 5.79130 |
| CD630_n00930 | CD630_SQ1642 | 4 -- 26 | 170 -- 193 | -11.8627 | 5.79130 | 8.65138 |
| CD630_SQ1641 | CD630_n00930 | 2 -- 34 | 106 -- 134 | -11.8617 | 8.65138 | 5.79130 |
| CD630_n00850 | CD630_SQ808 | 114 -- 130 | 150 -- 166 | -11.8607 | 5.53585 | 6.85680 |
| CD630_SQ808 | CD630_n00850 | 150 -- 166 | 114 -- 130 | -11.8607 | 6.85680 | 5.53585 |
| CD630_s0370 | CD630_s0660 | 4 -- 46 | 19 -- 71 | -11.8463 | 10.21495 | 4.92199 |
| CD630_s0190 | CD630_Cdi1_5 | 70 -- 90 | 349 -- 371 | -11.845 | 4.98480 | 4.83391 |
| CD630_Cdi1_5 | CD630_s0190 | 349 -- 371 | 70 -- 90 | -11.845 | 4.83391 | 4.98480 |
| CD630_s0190 | CD630_n00850 | 171 -- 192 | 80 -- 103 | -11.8311 | 4.98480 | 5.53585 |
| CD630_n00850 | CD630_s0190 | 80 -- 103 | 171 -- 192 | -11.8311 | 5.53585 | 4.98480 |
| CD630_Cdi1_5 | CD630_s0591 | 453 -- 462 | 123 -- 132 | -11.8291 | 4.83391 | 6.84967 |
| CD630_s0591 | CD630_Cdi1_5 | 123 -- 132 | 453 -- 462 | -11.8291 | 6.84967 | 4.83391 |
| RCd5 | CD630_s0631 | 36 -- 92 | 60 -- 118 | -11.8221 | 6.52781 | 4.95025 |
| CD630_n00680 | CD630_s0631 | 36 -- 92 | 60 -- 118 | -11.8221 | 6.52781 | 4.95025 |
| CD630_s0631 | RCd5 | 60 -- 118 | 36 -- 92 | -11.8221 | 4.95025 | 6.52781 |
| CD630_s0631 | CD630_n00680 | 60 -- 118 | 36 -- 92 | -11.8221 | 4.95025 | 6.52781 |
| CD630_n00680 | CD630_s0470 | 240 -- 289 | 44 -- 99 | -11.8053 | 6.52781 | 5.35019 |

| | | | | | | |
|---|---|---|---|---|---|---|
| CD630_s0470 | RCd5 | 44 -- 99 | 240 -- 289 | -11.8053 | 5.35019 | 6.52781 |
| CD630_s0470 | CD630_n00680 | 44 -- 99 | 240 -- 289 | -11.8053 | 5.35019 | 6.52781 |
| RCd5 | CD630_Cdi2_2 | 88 -- 131 | 1 -- 46 | -11.8027 | 6.52781 | 6.72203 |
| RCd5 | CD630_Cdi2_2 | 88 -- 131 | 1 -- 46 | -11.8027 | 6.52781 | 6.72203 |
| CD630_n00680 | CD630_Cdi2_2 | 88 -- 131 | 1 -- 46 | -11.8027 | 6.52781 | 6.72203 |
| CD630_n00680 | CD630_Cdi2_2 | 88 -- 131 | 1 -- 46 | -11.8027 | 6.52781 | 6.72203 |
| CD630_Cdi2_2 | RCd5 | 1 -- 46 | 88 -- 131 | -11.8027 | 6.72203 | 6.52781 |
| CD630_Cdi2_2 | CD630_n00680 | 1 -- 46 | 88 -- 131 | -11.8027 | 6.72203 | 6.52781 |
| CD630_Cdi2_2 | RCd5 | 1 -- 46 | 88 -- 131 | -11.8027 | 6.72203 | 6.52781 |
| CD630_Cdi2_2 | CD630_n00680 | 1 -- 46 | 88 -- 131 | -11.8027 | 6.72203 | 6.52781 |
| RCd5 | CD630_Cdi1_5 | 404 -- 432 | 368 -- 401 | -11.7788 | 6.52781 | 4.83391 |
| CD630_Cdi1_5 | RCd5 | 368 -- 401 | 404 -- 432 | -11.7788 | 4.83391 | 6.52781 |
| CD630_s0631 | CD630_n00600 | 320 -- 343 | 420 -- 443 | -11.7763 | 4.95025 | 5.31146 |
| CD630_n00930 | CD630_SQ1641 | 106 -- 132 | 5 -- 34 | -11.7696 | 5.79130 | 8.65138 |
| CD630_s0340 | CD630_s0390 | 31 -- 116 | 15 -- 106 | -11.7675 | 5.61064 | 4.73397 |
| CD630_n00330 | CD630_Cdi1_10 | 44 -- 58 | 136 -- 150 | -11.7566 | 6.46464 | 8.60766 |
| CD630_Cdi1_10 | CD630_n00330 | 136 -- 150 | 44 -- 58 | -11.7566 | 8.60766 | 6.46464 |
| CD630_Cdi1_3 | CD630_s0450 | 209 -- 266 | 69 -- 124 | -11.7553 | 6.59405 | 5.67265 |
| CD630_s0450 | CD630_Cdi1_3 | 69 -- 124 | 209 -- 266 | -11.7553 | 5.67265 | 6.59405 |
| RCd5 | CD630_n00790 | 282 -- 430 | 911 -- 1039 | -11.7552 | 6.52781 | 5.51210 |
| CD630_n00790 | RCd5 | 911 -- 934 | 407 -- 430 | -11.7423 | 5.51210 | 6.52781 |
| CD630_n00460 | CD630_Cdi1_10 | 291 -- 309 | 127 -- 147 | -11.7407 | 5.55447 | 8.60766 |
| CD630_Cdi1_10 | CD630_n00460 | 127 -- 147 | 291 -- 309 | -11.7407 | 8.60766 | 5.55447 |
| CD630_n00620 | CD630_n01010 | 69 -- 97 | 631 -- 659 | -11.7292 | 5.82838 | 4.59146 |
| CD630_n01010 | CD630_n00620 | 631 -- 659 | 69 -- 97 | -11.7292 | 4.59146 | 5.82838 |
| CD630_s0300 | CD630_SQ1642 | 44 -- 52 | 7 -- 15 | -11.7289 | 4.17452 | 8.65138 |
| CD630_SQ1642 | CD630_s0300 | 7 -- 15 | 44 -- 52 | -11.7289 | 8.65138 | 4.17452 |
| CD630_n00380 | CD630_SQ1002 | 662 -- 703 | 127 -- 168 | -11.7224 | 4.27916 | 4.43255 |
| CD630_SQ1002 | CD630_n00380 | 127 -- 168 | 662 -- 703 | -11.7224 | 4.43255 | 4.27916 |
| CD630_SQ1002 | CD630_n00990 | 127 -- 168 | 751 -- 792 | -11.7224 | 4.43255 | 3.72051 |
| CD630_n00990 | CD630_SQ1002 | 751 -- 792 | 127 -- 168 | -11.7224 | 3.72051 | 4.43255 |
| CD630_SQ367 | CD630_n00690 | 73 -- 115 | 278 -- 322 | -11.7126 | 4.08290 | 5.96526 |
| CD630_n00690 | CD630_SQ367 | 278 -- 322 | 73 -- 115 | -11.7126 | 5.96526 | 4.08290 |
| CD630_s0280 | CD630_n01010 | 209 -- 217 | 1392 -- 1400 | -11.7038 | 5.37459 | 4.59146 |
| CD630_n01010 | CD630_s0280 | 1392 -- 1400 | 209 -- 217 | -11.7038 | 4.59146 | 5.37459 |
| CD630_n00220 | CD630_n01010 | 2 -- 32 | 842 -- 865 | -11.7025 | 4.94050 | 4.59146 |
| CD630_n00240 | RCd9 | 98 -- 121 | 146 -- 169 | -11.7013 | 5.82709 | 8.96925 |
| RCd9 | CD630_n00240 | 146 -- 169 | 98 -- 121 | -11.7013 | 8.96925 | 5.82709 |
| CD630_n00600 | CD630_n00510 | 519 -- 546 | 939 -- 962 | -11.6947 | 5.31146 | 3.92762 |
| CD630_n00460 | CD630_s0480 | 100 -- 152 | 133 -- 197 | -11.6922 | 5.55447 | 4.87999 |
| CD630_s0480 | CD630_n00460 | 133 -- 197 | 100 -- 152 | -11.6922 | 4.87999 | 5.55447 |
| CD630_s0010 | CD630_n00850 | 19 -- 36 | 233 -- 250 | -11.6893 | 4.92433 | 5.53585 |
| CD630_n00850 | CD630_s0010 | 233 -- 250 | 19 -- 36 | -11.6893 | 5.53585 | 4.92433 |
| CD630_s0400 | RCd8 | 162 -- 174 | 197 -- 209 | -11.6781 | 4.51086 | 6.97567 |
| RCd8 | CD630_s0400 | 197 -- 209 | 162 -- 174 | -11.6781 | 6.97567 | 4.51086 |
| CD630_n00560 | CD630_s0450 | 465 -- 485 | 70 -- 90 | -11.6718 | 5.82053 | 5.67265 |
| CD630_s0450 | CD630_n00560 | 70 -- 90 | 465 -- 485 | -11.6718 | 5.67265 | 5.82053 |
| CD630_s0320 | CD630_s0190 | 30 -- 46 | 36 -- 54 | -11.6717 | 8.15991 | 4.98480 |
| CD630_s0390 | CD630_n01010 | 183 -- 235 | 1078 -- 1133 | -11.6713 | 4.73397 | 4.59146 |
| CD630_s0300 | CD630_s0641 | 137 -- 177 | 138 -- 173 | -11.6646 | 4.17452 | 6.68432 |
| CD630_SQ1641 | CD630_Cdi1_5 | 101 -- 117 | 106 -- 122 | -11.6646 | 8.65138 | 4.83391 |
| CD630_SQ1641 | CD630_n00390 | 101 -- 117 | 106 -- 122 | -11.6646 | 8.65138 | 4.83391 |
| CD630_Cdi1_5 | CD630_SQ1641 | 106 -- 122 | 101 -- 117 | -11.6646 | 4.83391 | 8.65138 |
| CD630_n00390 | CD630_SQ1641 | 106 -- 122 | 101 -- 117 | -11.6646 | 4.83391 | 8.65138 |
| CD630_n00510 | CD630_n00690 | 80 -- 118 | 11 -- 47 | -11.6587 | 3.92762 | 5.96526 |
| CD630_Cdi1_4 | CD630_s0390 | 446 -- 461 | 215 -- 230 | -11.6465 | 5.31318 | 4.73397 |
| CD630_n00790 | CD630_s0670 | 210 -- 224 | 24 -- 38 | -11.6417 | 5.51210 | 4.56362 |
| CD630_s0670 | CD630_n00790 | 24 -- 38 | 210 -- 224 | -11.6417 | 4.56362 | 5.51210 |
| CD630_n00680 | CD630_Cdi1_8 | 352 -- 370 | 1 -- 20 | -11.6355 | 6.52781 | 5.79994 |
| CD630_s0010 | RCd8 | 155 -- 167 | 197 -- 209 | -11.6348 | 4.92433 | 6.97567 |
| RCd8 | CD630_s0010 | 197 -- 209 | 155 -- 167 | -11.6348 | 6.97567 | 4.92433 |
| RCd9 | CD630_s0450 | 89 -- 103 | 30 -- 43 | -11.6338 | 8.96925 | 5.67265 |
| CD630_s0450 | RCd9 | 30 -- 43 | 89 -- 103 | -11.6338 | 5.67265 | 8.96925 |
| CD630_SQ1641 | CD630_Cdi1_4 | 101 -- 117 | 113 -- 129 | -11.6311 | 8.65138 | 5.31318 |
| CD630_SQ1641 | CD630_n00980 | 101 -- 117 | 113 -- 129 | -11.6311 | 8.65138 | 5.31318 |
| CD630_Cdi1_4 | CD630_SQ1641 | 113 -- 129 | 101 -- 117 | -11.6311 | 5.31318 | 8.65138 |
| CD630_n00980 | CD630_SQ1641 | 113 -- 129 | 101 -- 117 | -11.6311 | 5.31318 | 8.65138 |
| CD630_s0220 | CD630_n00080 | 7 -- 26 | 84 -- 103 | -11.6302 | 3.72587 | 4.59330 |
| CD630_n00080 | CD630_s0220 | 84 -- 103 | 7 -- 26 | -11.6302 | 4.59330 | 3.72587 |
| CD630_n00380 | CD630_n00930 | 766 -- 777 | 336 -- 347 | -11.6295 | 4.27916 | 5.79130 |
| CD630_n00930 | CD630_n00380 | 336 -- 347 | 766 -- 777 | -11.6295 | 5.79130 | 4.27916 |
| CD630_n00930 | CD630_n00990 | 336 -- 347 | 855 -- 866 | -11.6295 | 5.79130 | 3.72051 |
| CD630_n00990 | CD630_n00930 | 855 -- 866 | 336 -- 347 | -11.6295 | 3.72051 | 5.79130 |
| CD630_s0400 | CD630_Cdi1_5 | 132 -- 145 | 434 -- 448 | -11.6264 | 4.51086 | 4.83391 |
| CD630_Cdi1_5 | CD630_s0400 | 434 -- 448 | 132 -- 145 | -11.6264 | 4.83391 | 4.51086 |
| CD630_SQ1038 | CD630_s0480 | 4 -- 28 | 61 -- 83 | -11.6207 | 3.49019 | 4.87999 |
| CD630_s0480 | CD630_SQ1038 | 61 -- 83 | 4 -- 28 | -11.6207 | 4.87999 | 3.49019 |
| CD630_Cdi1_4 | CD630_s0300 | 349 -- 386 | 28 -- 60 | -11.6124 | 5.31318 | 4.17452 |
| CD630_s0300 | CD630_Cdi1_4 | 28 -- 60 | 349 -- 386 | -11.6123 | 4.17452 | 5.31318 |
| CD630_SQ367 | CD630_s0470 | 1 -- 49 | 6 -- 42 | -11.6107 | 4.08290 | 5.35019 |
| CD630_n00600 | CD630_Cdi1_8 | 797 -- 836 | 6 -- 58 | -11.601 | 5.31146 | 5.79994 |
| CD630_Cdi1_8 | CD630_n00600 | 6 -- 58 | 797 -- 836 | -11.601 | 5.79994 | 5.31146 |
| RCd5 | RCd2 | 247 -- 272 | 167 -- 193 | -11.5927 | 6.52781 | 3.80984 |
| CD630_n00680 | RCd2 | 247 -- 272 | 167 -- 193 | -11.5927 | 6.52781 | 3.80984 |
| RCd2 | RCd5 | 167 -- 193 | 247 -- 272 | -11.5927 | 3.80984 | 6.52781 |
| RCd2 | CD630_n00680 | 167 -- 193 | 247 -- 272 | -11.5927 | 3.80984 | 6.52781 |
| CD630_s0450 | CD630_SQ1642 | 27 -- 90 | 117 -- 181 | -11.5882 | 5.67265 | 8.65138 |
| CD630_n00560 | CD630_SQ808 | 159 -- 194 | 39 -- 69 | -11.5855 | 5.82053 | 6.85680 |
| CD630_s0641 | CD630_s0300 | 138 -- 175 | 138 -- 177 | -11.5855 | 6.68432 | 4.17452 |
| CD630_SQ808 | CD630_n00560 | 39 -- 69 | 159 -- 194 | -11.5855 | 6.85680 | 5.82053 |
| CD630_SQ367 | CD630_s0642 | 111 -- 191 | 119 -- 193 | -11.5807 | 4.08290 | 6.00600 |
| RCd9 | CD630_n00790 | 223 -- 235 | 624 -- 638 | -11.5773 | 8.96925 | 5.51210 |
| CD630_n00440 | CD630_s0510 | 63 -- 73 | 42 -- 52 | -11.5749 | 4.85723 | 3.37631 |
| CD630_s0510 | CD630_n00440 | 42 -- 52 | 63 -- 73 | -11.5749 | 3.37631 | 4.85723 |
| RCd5 | CD630_n00690 | 178 -- 227 | 92 -- 142 | -11.5615 | 6.52781 | 5.96526 |
| CD630_n00680 | CD630_n00690 | 178 -- 227 | 92 -- 142 | -11.5615 | 6.52781 | 5.96526 |
| CD630_n00690 | RCd5 | 92 -- 142 | 178 -- 227 | -11.5615 | 5.96526 | 6.52781 |
| CD630_n00690 | CD630_n00680 | 92 -- 142 | 178 -- 227 | -11.5615 | 5.96526 | 6.52781 |
| CD630_n00460 | CD630_n00790 | 116 -- 148 | 281 -- 312 | -11.557 | 5.55447 | 5.51210 |
| CD630_n00790 | CD630_n00460 | 281 -- 312 | 116 -- 148 | -11.557 | 5.51210 | 5.55447 |
| CD630_SQ1641 | CD630_s0330 | 1 -- 34 | 4 -- 37 | -11.5552 | 8.65138 | 4.37658 |
| CD630_s0330 | CD630_SQ1641 | 4 -- 37 | 1 -- 34 | -11.5552 | 4.37658 | 8.65138 |
| CD630_s0400 | RCd5 | 42 -- 74 | 380 -- 414 | -11.5453 | 4.51086 | 6.52781 |
| RCd5 | CD630_s0400 | 380 -- 414 | 42 -- 74 | -11.5453 | 6.52781 | 4.51086 |
| CD630_s0050 | CD630_s0310 | 47 -- 54 | 189 -- 196 | -11.5451 | 5.03432 | 4.17452 |
| CD630_s0310 | CD630_s0050 | 189 -- 196 | 47 -- 54 | -11.5451 | 4.17452 | 5.03432 |
| CD630_Cdi1_5 | CD630_s0390 | 439 -- 454 | 215 -- 230 | -11.5376 | 4.83391 | 4.73397 |
| CD630_n00470 | CD630_n00080 | 8 -- 87 | 42 -- 95 | -11.5355 | 3.52267 | 4.59330 |
| CD630_n00680 | CD630_Cdi1_10 | 352 -- 370 | 1 -- 20 | -11.5277 | 6.52781 | 8.60766 |
| CD630_s0310 | CD630_n00330 | 191 -- 197 | 38 -- 44 | -11.5182 | 4.17452 | 6.46464 |
| CD630_n00330 | CD630_s0310 | 38 -- 44 | 191 -- 197 | -11.5182 | 6.46464 | 4.17452 |
| CD630_n00510 | CD630_s0510 | 86 -- 114 | 29 -- 63 | -11.5089 | 3.92762 | 3.37631 |
| CD630_s0390 | CD630_n00590 | 135 -- 156 | 56 -- 77 | -11.5044 | 4.73397 | 4.28395 |
| CD630_n00590 | CD630_s0390 | 56 -- 77 | 135 -- 156 | -11.5044 | 4.28395 | 4.73397 |
| CD630_RNA_5 | CD630_Cdi1_4 | 85 -- 106 | 452 -- 466 | -11.5037 | 4.62291 | 5.31318 |
| CD630_Cdi1_4 | CD630_RNA_5 | 452 -- 466 | 85 -- 106 | -11.5037 | 5.31318 | 4.62291 |
| CD630_s0400 | CD630_Cdi1_4 | 132 -- 145 | 441 -- 455 | -11.5015 | 4.51086 | 5.31318 |
| CD630_Cdi1_4 | CD630_s0400 | 441 -- 455 | 132 -- 145 | -11.5015 | 5.31318 | 4.51086 |
| RCd1 | CD630_s0642 | 27 -- 48 | 17 -- 39 | -11.4953 | 4.09988 | 6.00600 |
| CD630_s0642 | RCd1 | 17 -- 39 | 27 -- 48 | -11.4953 | 6.00600 | 4.09988 |
| CD630_n01010 | CD630_n00290 | 120 -- 166 | 22 -- 70 | -11.4944 | 4.59146 | 3.85358 |
| CD630_s0010 | CD630_n00790 | 2 -- 21 | 1002 -- 1019 | -11.491 | 4.92433 | 5.51210 |
| CD630_n00790 | CD630_s0010 | 1002 -- 1019 | 2 -- 21 | -11.491 | 5.51210 | 4.92433 |
| CD630_s0641 | CD630_n00460 | 91 -- 134 | 377 -- 421 | -11.4858 | 6.68432 | 5.55447 |
| CD630_SQ1002 | CD630_s0510 | 153 -- 168 | 6 -- 20 | -11.485 | 4.43255 | 3.37631 |
| CD630_s0510 | CD630_SQ1002 | 6 -- 20 | 153 -- 168 | -11.485 | 3.37631 | 4.43255 |
| CD630_s0480 | CD630_n01010 | 26 -- 39 | 794 -- 807 | -11.4847 | 4.87999 | 4.59146 |
| CD630_n01010 | CD630_s0480 | 794 -- 807 | 26 -- 39 | -11.4847 | 4.59146 | 4.87999 |
| CD630_s0250 | CD630_n00590 | 77 -- 86 | 68 -- 77 | -11.4844 | 3.74616 | 4.28395 |
| CD630_n00590 | CD630_s0250 | 68 -- 77 | 77 -- 86 | -11.4844 | 4.28395 | 3.74616 |
| CD630_s0631 | CD630_SQ1656 | 60 -- 107 | 151 -- 197 | -11.4807 | 4.95025 | 4.25622 |
| CD630_SQ367 | CD630_n00600 | 132 -- 168 | 1136 -- 1170 | -11.4654 | 4.08290 | 5.31146 |
| CD630_n00600 | CD630_SQ367 | 1136 -- 1170 | 132 -- 168 | -11.4654 | 5.31146 | 4.08290 |
| CD630_n00460 | CD630_n00510 | 649 -- 674 | 202 -- 229 | -11.4609 | 5.55447 | 3.92762 |
| CD630_n00510 | CD630_n00460 | 202 -- 229 | 649 -- 674 | -11.4609 | 3.92762 | 5.55447 |
| CD630_s0390 | CD630_Cdi1_4 | 86 -- 124 | 429 -- 461 | -11.4604 | 4.73397 | 5.31318 |
| CD630_SQ327 | CD630_s0330 | 113 -- 124 | 28 -- 39 | -11.4571 | 4.62291 | 4.37658 |
| CD630_s0330 | CD630_SQ327 | 28 -- 39 | 113 -- 124 | -11.4571 | 4.37658 | 4.62291 |
| CD630_n00290 | CD630_Cdi1_8 | 4 -- 15 | 142 -- 153 | -11.4496 | 3.85358 | 5.79994 |
| CD630_n00380 | CD630_SQ1656 | 651 -- 689 | 8 -- 51 | -11.4472 | 4.27916 | 4.25622 |
| CD630_SQ1656 | CD630_n00380 | 8 -- 51 | 651 -- 689 | -11.4472 | 4.25622 | 4.27916 |
| CD630_SQ1656 | CD630_n00990 | 8 -- 51 | 740 -- 778 | -11.4472 | 4.25622 | 3.72051 |
| CD630_n00990 | CD630_SQ1656 | 740 -- 778 | 8 -- 51 | -11.4472 | 3.72051 | 4.25622 |
| CD630_n00380 | CD630_s0670 | 861 -- 905 | 22 -- 64 | -11.4384 | 4.27916 | 4.56362 |
| CD630_n00990 | CD630_s0670 | 950 -- 994 | 22 -- 64 | -11.4384 | 3.72051 | 4.56362 |
| CD630_s0670 | CD630_n00380 | 22 -- 64 | 861 -- 905 | -11.4384 | 4.56362 | 4.27916 |
| CD630_s0670 | CD630_n00990 | 22 -- 64 | 950 -- 994 | -11.4384 | 4.56362 | 3.72051 |
| CD630_s0340 | CD630_n00330 | 3 -- 14 | 43 -- 53 | -11.4347 | 5.61064 | 6.46464 |
| CD630_n00330 | CD630_s0340 | 43 -- 53 | 3 -- 14 | -11.4347 | 6.46464 | 5.61064 |
| CD630_Cdi1_8 | CD630_n00680 | 1 -- 33 | 339 -- 370 | -11.4295 | 5.79994 | 6.52781 |

| | | | | | | |
|---|---|---|---|---|---|---|
| CD630_n00790 | CD630_Cdi1_8 | 273 -- 300 | 287 -- 320 | -11.4223 | 5.51210 | 5.79994 |
| CD630_n00790 | CD630_SQ1656 | 583 -- 632 | 8 -- 44 | -11.4126 | 5.51210 | 4.25622 |
| CD630_s0210 | CD630_s0390 | 25 -- 59 | 67 -- 102 | -11.4101 | 4.02122 | 4.73397 |
| CD630_s0390 | CD630_s0210 | 67 -- 102 | 25 -- 59 | -11.4101 | 4.73397 | 4.02122 |
| CD630_n00600 | CD630_Cdi1_11 | 797 -- 836 | 6 -- 58 | -11.4043 | 5.31146 | 5.77720 |
| CD630_n00590 | CD630_n01010 | 52 -- 77 | 1097 -- 1122 | -11.3879 | 4.28395 | 4.59146 |
| CD630_RNA_5 | CD630_Cdi1_5 | 85 -- 106 | 445 -- 459 | -11.3824 | 4.62291 | 4.83391 |
| CD630_Cdi1_5 | CD630_RNA_5 | 445 -- 459 | 85 -- 106 | -11.3824 | 4.83391 | 4.62291 |
| CD630_Cdi1_10 | CD630_n00680 | 1 -- 33 | 339 -- 370 | -11.3816 | 8.60766 | 6.52781 |
| CD630_n00680 | CD630_SQ2429 | 339 -- 380 | 91 -- 132 | -11.3725 | 6.52781 | 5.77720 |
| CD630_SQ2429 | CD630_n00680 | 91 -- 132 | 339 -- 380 | -11.3725 | 5.77720 | 6.52781 |
| CD630_n00290 | CD630_n01010 | 22 -- 48 | 142 -- 166 | -11.3646 | 3.85358 | 4.59146 |
| CD630_s0300 | CD630_Cdi1_5 | 28 -- 60 | 342 -- 379 | -11.3525 | 4.17452 | 4.83391 |
| CD630_Cdi1_5 | CD630_s0300 | 342 -- 379 | 28 -- 60 | -11.3525 | 4.83391 | 4.17452 |
| CD630_s0190 | CD630_s0280 | 75 -- 94 | 76 -- 94 | -11.3518 | 4.98480 | 5.37459 |
| CD630_n00460 | CD630_n00290 | 165 -- 189 | 22 -- 48 | -11.3477 | 5.55447 | 3.85358 |
| CD630_n00290 | CD630_n00460 | 22 -- 48 | 165 -- 189 | -11.3477 | 3.85358 | 5.55447 |
| CD630_SQ1642 | CD630_n00470 | 140 -- 191 | 11 -- 60 | -11.3239 | 8.65138 | 3.52267 |
| CD630_n00470 | CD630_SQ1642 | 11 -- 60 | 140 -- 191 | -11.3239 | 3.52267 | 8.65138 |
| CD630_s0390 | CD630_Cdi1_5 | 86 -- 124 | 422 -- 454 | -11.3076 | 4.73397 | 4.83391 |
| CD630_n00980 | CD630_s0280 | 277 -- 302 | 68 -- 91 | -11.3065 | 5.31318 | 5.37459 |
| CD630_SQ408 | CD630_s0210 | 148 -- 188 | 43 -- 80 | -11.2983 | 7.27863 | 4.02122 |
| CD630_s0210 | CD630_SQ408 | 43 -- 80 | 148 -- 188 | -11.2983 | 4.02122 | 7.27863 |
| CD630_s0340 | CD630_n00290 | 3 -- 14 | 45 -- 56 | -11.2928 | 5.61064 | 3.85358 |
| CD630_n00290 | CD630_s0340 | 45 -- 56 | 3 -- 14 | -11.2928 | 3.85358 | 5.61064 |
| CD630_s0480 | CD630_Cdi1_5 | 66 -- 90 | 422 -- 447 | -11.2829 | 4.87999 | 4.83391 |
| CD630_Cdi1_5 | CD630_s0480 | 422 -- 447 | 66 -- 90 | -11.2829 | 4.83391 | 4.87999 |
| CD630_Cdi1_3 | CD630_Cdi1_8 | 183 -- 274 | 114 -- 205 | -11.2778 | 6.59405 | 5.79994 |
| CD630_Cdi1_8 | CD630_Cdi1_3 | 114 -- 205 | 183 -- 274 | -11.2778 | 5.79994 | 6.59405 |
| CD630_Cdi1_8 | CD630_n00930 | 131 -- 205 | 4 -- 61 | -11.2777 | 5.79994 | 5.79130 |
| CD630_n00930 | CD630_Cdi1_8 | 4 -- 61 | 131 -- 205 | -11.2777 | 5.79130 | 5.79994 |
| CD630_n00080 | CD630_n00470 | 42 -- 53 | 76 -- 87 | -11.2694 | 4.59330 | 3.52267 |
| CD630_SQ367 | CD630_n00460 | 152 -- 175 | 727 -- 748 | -11.2692 | 4.08290 | 5.55447 |
| CD630_n00460 | CD630_SQ367 | 727 -- 748 | 152 -- 175 | -11.2692 | 5.55447 | 4.08290 |
| CD630_n00440 | CD630_n00600 | 63 -- 75 | 67 -- 79 | -11.2586 | 4.85723 | 5.31146 |
| CD630_s0510 | CD630_s0190 | 130 -- 154 | 119 -- 135 | -11.2571 | 3.37631 | 4.98480 |
| CD630_SQ1005 | CD630_Cdi1_8 | 61 -- 74 | 106 -- 119 | -11.2554 | 3.63582 | 5.79994 |
| CD630_Cdi1_8 | CD630_SQ1005 | 106 -- 119 | 61 -- 74 | -11.2554 | 5.79994 | 3.63582 |
| CD630_n00380 | RCd9 | 157 -- 242 | 145 -- 233 | -11.2539 | 4.27916 | 8.96925 |
| RCd9 | CD630_n00380 | 145 -- 233 | 157 -- 242 | -11.2539 | 8.96925 | 4.27916 |
| RCd9 | CD630_n00990 | 145 -- 233 | 246 -- 331 | -11.2539 | 8.96925 | 3.72051 |
| CD630_n00990 | RCd9 | 246 -- 331 | 145 -- 233 | -11.2539 | 3.72051 | 8.96925 |
| CD630_s0210 | CD630_s0280 | 204 -- 213 | 214 -- 223 | -11.2317 | 4.02122 | 5.37459 |
| CD630_s0280 | CD630_s0210 | 214 -- 223 | 204 -- 213 | -11.2317 | 5.37459 | 4.02122 |
| CD630_n00510 | CD630_s0340 | 104 -- 119 | 155 -- 170 | -11.229 | 3.92762 | 5.61064 |
| CD630_s0340 | CD630_n00510 | 155 -- 170 | 104 -- 119 | -11.229 | 5.61064 | 3.92762 |
| RCd9 | CD630_n01010 | 140 -- 162 | 99 -- 118 | -11.2283 | 8.96925 | 4.59146 |
| CD630_n00680 | CD630_Cdi1_9 | 352 -- 370 | 1 -- 20 | -11.2218 | 6.52781 | 4.25622 |
| RCd5 | CD630_Cdi1_8 | 352 -- 370 | 1 -- 20 | -11.2177 | 6.52781 | 5.79994 |
| CD630_n00380 | RCd1 | 163 -- 177 | 2 -- 16 | -11.2104 | 4.27916 | 4.09988 |
| RCd1 | CD630_n00380 | 2 -- 16 | 163 -- 177 | -11.2104 | 4.09988 | 4.27916 |
| RCd1 | CD630_n00990 | 2 -- 16 | 252 -- 266 | -11.2104 | 4.09988 | 3.72051 |
| CD630_n00990 | RCd1 | 252 -- 266 | 2 -- 16 | -11.2104 | 3.72051 | 4.09988 |
| CD630_s0281 | CD630_s0642 | 163 -- 200 | 143 -- 174 | -11.2071 | 8.30299 | 6.00600 |
| CD630_s0642 | CD630_s0281 | 143 -- 174 | 163 -- 200 | -11.2071 | 6.00600 | 8.30299 |
| CD630_n00590 | CD630_n00930 | 28 -- 72 | 180 -- 234 | -11.2053 | 4.28395 | 5.79130 |
| CD630_n00930 | CD630_n00590 | 180 -- 234 | 28 -- 72 | -11.2053 | 5.79130 | 4.28395 |
| CD630_n00460 | CD630_s0631 | 210 -- 245 | 1 -- 28 | -11.2027 | 5.55447 | 4.95025 |
| CD630_s0631 | CD630_n00460 | 1 -- 28 | 210 -- 245 | -11.2027 | 4.95025 | 5.55447 |
| CD630_s0280 | CD630_n00340 | 19 -- 31 | 28 -- 40 | -11.1985 | 5.37459 | 5.83478 |
| CD630_n00340 | CD630_s0280 | 28 -- 40 | 19 -- 31 | -11.1985 | 5.83478 | 5.37459 |
| CD630_Cdi1_8 | RCd5 | 1 -- 33 | 339 -- 370 | -11.1977 | 5.79994 | 6.52781 |
| CD630_n00680 | CD630_Cdi1_11 | 352 -- 370 | 1 -- 20 | -11.1953 | 6.52781 | 5.77720 |
| CD630_n00690 | CD630_n00510 | 107 -- 118 | 11 -- 22 | -11.1948 | 5.96526 | 3.92762 |
| CD630_Cdi1_3 | CD630_n00330 | 484 -- 495 | 34 -- 45 | -11.1777 | 6.59405 | 6.46464 |
| CD630_n00330 | CD630_Cdi1_3 | 34 -- 45 | 484 -- 495 | -11.1777 | 6.46464 | 6.59405 |
| CD630_s0670 | CD630_n00440 | 157 -- 194 | 196 -- 230 | -11.1768 | 4.56362 | 4.85723 |
| CD630_n00460 | CD630_n00460 | 110 -- 148 | 404 -- 435 | -11.1692 | 5.55447 | 5.55447 |
| CD630_s0270 | CD630_Cdi1_8 | 86 -- 99 | 106 -- 119 | -11.1646 | 3.63582 | 5.79994 |
| CD630_Cdi1_8 | CD630_s0270 | 106 -- 119 | 86 -- 99 | -11.1646 | 5.79994 | 3.63582 |
| CD630_s0480 | CD630_s0670 | 34 -- 65 | 25 -- 55 | -11.1644 | 4.87999 | 4.56362 |
| CD630_s0670 | CD630_s0480 | 25 -- 55 | 34 -- 65 | -11.1644 | 4.56362 | 4.87999 |
| CD630_n00510 | CD630_s0390 | 211 -- 241 | 4 -- 39 | -11.1639 | 3.92762 | 4.73397 |
| CD630_s0390 | CD630_n00510 | 4 -- 39 | 211 -- 241 | -11.1639 | 4.73397 | 3.92762 |
| CD630_n00380 | CD630_n00470 | 301 -- 357 | 21 -- 69 | -11.1571 | 4.27916 | 3.52267 |
| CD630_n00990 | CD630_n00470 | 390 -- 446 | 21 -- 69 | -11.1571 | 3.72051 | 3.52267 |
| CD630_n00470 | CD630_n00380 | 21 -- 69 | 301 -- 357 | -11.157 | 3.52267 | 4.27916 |
| CD630_n00470 | CD630_n00990 | 21 -- 69 | 390 -- 446 | -11.157 | 3.52267 | 3.72051 |
| CD630_s0470 | CD630_s0631 | 3 -- 17 | 326 -- 342 | -11.1563 | 5.35019 | 4.95025 |
| CD630_s0631 | CD630_s0470 | 326 -- 342 | 3 -- 17 | -11.1563 | 4.95025 | 5.35019 |
| CD630_Cdi1_9 | CD630_n00680 | 1 -- 33 | 339 -- 370 | -11.1545 | 4.25622 | 6.52781 |
| RCd5 | CD630_Cdi1_10 | 339 -- 370 | 1 -- 33 | -11.1498 | 6.52781 | 8.60766 |
| CD630_Cdi1_10 | RCd5 | 1 -- 33 | 339 -- 370 | -11.1498 | 8.60766 | 6.52781 |
| CD630_n00440 | CD630_s0670 | 196 -- 226 | 161 -- 194 | -11.1489 | 4.85723 | 4.56362 |
| CD630_n00790 | RCd9 | 624 -- 669 | 187 -- 235 | -11.1325 | 5.51210 | 8.96925 |
| CD630_s0270 | CD630_n01010 | 102 -- 127 | 375 -- 405 | -11.1322 | 3.63582 | 4.59146 |
| CD630_n01010 | CD630_s0270 | 375 -- 405 | 102 -- 127 | -11.1322 | 4.59146 | 3.63582 |
| CD630_Cdi1_11 | CD630_n00680 | 1 -- 33 | 339 -- 370 | -11.1263 | 5.77720 | 6.52781 |
| CD630_n00560 | CD630_SQ1656 | 183 -- 203 | 22 -- 44 | -11.1245 | 5.82053 | 4.25622 |
| CD630_SQ1656 | CD630_n00560 | 22 -- 44 | 183 -- 203 | -11.1245 | 4.25622 | 5.82053 |
| RCd5 | CD630_s0470 | 240 -- 288 | 45 -- 99 | -11.1205 | 6.52781 | 5.35019 |
| CD630_s0050 | CD630_n00440 | 13 -- 24 | 60 -- 73 | -11.1195 | 5.03432 | 4.85723 |
| CD630_n00440 | CD630_s0050 | 60 -- 73 | 13 -- 24 | -11.1195 | 4.85723 | 5.03432 |
| CD630_n00600 | CD630_Cdi1_9 | 797 -- 836 | 6 -- 58 | -11.115 | 5.31146 | 4.25622 |
| CD630_n00510 | CD630_n00290 | 166 -- 190 | 22 -- 48 | -11.0876 | 3.92762 | 3.85358 |
| CD630_n00290 | CD630_n00510 | 22 -- 48 | 166 -- 190 | -11.0876 | 3.85358 | 3.92762 |
| CD630_n00440 | CD630_n00790 | 157 -- 173 | 996 -- 1012 | -11.0786 | 4.85723 | 5.51210 |
| RCd5 | CD630_s0510 | 207 -- 228 | 33 -- 53 | -11.0767 | 6.52781 | 3.37631 |
| CD630_n00680 | CD630_s0510 | 207 -- 228 | 33 -- 53 | -11.0767 | 6.52781 | 3.37631 |
| CD630_s0510 | RCd5 | 33 -- 53 | 207 -- 228 | -11.0767 | 3.37631 | 6.52781 |
| CD630_s0510 | CD630_n00680 | 33 -- 53 | 207 -- 228 | -11.0767 | 3.37631 | 6.52781 |
| CD630_n00500 | CD630_s0500 | 91 -- 99 | 83 -- 91 | -11.0758 | 9.35359 | 3.25870 |
| CD630_s0500 | CD630_n00500 | 83 -- 91 | 91 -- 99 | -11.0758 | 3.25870 | 9.35359 |
| CD630_n00460 | CD630_Cdi2_3 | 350 -- 378 | 30 -- 61 | -11.0666 | 5.55447 | 3.99184 |
| CD630_s0010 | RCd2 | 158 -- 175 | 21 -- 38 | -11.0641 | 4.92433 | 3.80984 |
| RCd2 | CD630_s0010 | 21 -- 38 | 158 -- 175 | -11.0641 | 3.80984 | 4.92433 |
| CD630_n00620 | RCd5 | 43 -- 101 | 371 -- 428 | -11.0525 | 5.82838 | 6.52781 |
| RCd5 | CD630_n00620 | 371 -- 428 | 43 -- 101 | -11.0525 | 6.52781 | 5.82838 |
| CD630_s0390 | CD630_n00330 | 7 -- 20 | 164 -- 177 | -11.0464 | 4.73397 | 6.46464 |
| CD630_n00330 | CD630_s0390 | 164 -- 177 | 7 -- 20 | -11.0464 | 6.46464 | 4.73397 |
| CD630_s0460 | CD630_n00290 | 64 -- 75 | 48 -- 59 | -11.0441 | 5.74527 | 3.85358 |
| CD630_SQ2429 | RCd5 | 59 -- 132 | 339 -- 416 | -11.0343 | 5.77720 | 6.52781 |
| CD630_n00380 | CD630_s0450 | 529 -- 562 | 13 -- 44 | -11.0304 | 4.27916 | 5.67265 |
| CD630_s0450 | CD630_n00380 | 13 -- 44 | 529 -- 562 | -11.0304 | 5.67265 | 4.27916 |
| CD630_s0450 | CD630_n00990 | 13 -- 44 | 618 -- 651 | -11.0304 | 5.67265 | 3.72051 |
| CD630_n00990 | CD630_s0450 | 618 -- 651 | 13 -- 44 | -11.0304 | 3.72051 | 5.67265 |
| CD630_n00440 | CD630_n01010 | 26 -- 43 | 141 -- 157 | -11.0179 | 4.85723 | 4.59146 |
| CD630_n01010 | CD630_n00440 | 141 -- 157 | 26 -- 43 | -11.0179 | 4.59146 | 4.85723 |
| CD630_Cdi1_3 | CD630_Cdi1_10 | 239 -- 275 | 112 -- 151 | -11.011 | 6.59405 | 8.60766 |
| CD630_Cdi1_10 | CD630_Cdi1_3 | 112 -- 151 | 239 -- 275 | -11.011 | 8.60766 | 6.59405 |
| CD630_SQ1642 | CD630_s0500 | 163 -- 176 | 90 -- 102 | -11.006 | 8.65138 | 3.25870 |
| CD630_s0500 | CD630_SQ1642 | 90 -- 102 | 163 -- 176 | -11.006 | 3.25870 | 8.65138 |
| CD630_n00600 | RCd8 | 9 -- 24 | 7 -- 22 | -10.9895 | 5.31146 | 6.97567 |
| CD630_s0642 | CD630_SQ367 | 119 -- 175 | 127 -- 191 | -10.9782 | 6.00600 | 4.08290 |
| CD630_n00240 | CD630_SQ2429 | 60 -- 78 | 176 -- 195 | -10.9729 | 5.82709 | 5.77720 |
| CD630_SQ2429 | CD630_n00240 | 176 -- 195 | 60 -- 78 | -10.9729 | 5.77720 | 5.82709 |
| CD630_n00560 | RCd9 | 157 -- 171 | 250 -- 264 | -10.9721 | 5.82053 | 8.96925 |
| RCd9 | CD630_n00560 | 250 -- 264 | 157 -- 171 | -10.9721 | 8.96925 | 5.82053 |
| CD630_RNA_7 | CD630_s0642 | 94 -- 104 | 17 -- 28 | -10.9676 | 7.42886 | 6.00600 |
| CD630_s0642 | CD630_RNA_7 | 17 -- 28 | 94 -- 104 | -10.9676 | 6.00600 | 7.42886 |
| RCd5 | CD630_SQ2429 | 339 -- 380 | 91 -- 132 | -10.9603 | 6.52781 | 5.77720 |
| CD630_s0480 | CD630_Cdi1_4 | 66 -- 90 | 429 -- 454 | -10.9582 | 4.87999 | 5.31318 |
| CD630_Cdi1_4 | CD630_s0480 | 429 -- 454 | 66 -- 90 | -10.9582 | 5.31318 | 4.87999 |
| CD630_n00340 | CD630_n00650 | 103 -- 152 | 51 -- 101 | -10.956 | 5.83478 | 5.32127 |
| CD630_n00650 | CD630_n00340 | 51 -- 101 | 103 -- 152 | -10.956 | 5.32127 | 5.83478 |
| CD630_SQ2429 | CD630_Cdi2_2 | 193 -- 227 | 9 -- 46 | -10.955 | 5.77720 | 6.72203 |
| CD630_SQ2429 | CD630_Cdi2_2 | 193 -- 227 | 9 -- 46 | -10.955 | 5.77720 | 6.72203 |
| CD630_s0470 | CD630_SQ367 | 6 -- 35 | 20 -- 49 | -10.9461 | 5.35019 | 4.08290 |
| CD630_Cdi1_8 | CD630_n00790 | 226 -- 299 | 864 -- 940 | -10.9403 | 5.79994 | 5.51210 |
| CD630_n00850 | CD630_n00930 | 162 -- 204 | 4 -- 46 | -10.9371 | 5.53585 | 5.79130 |
| CD630_n00930 | CD630_n00850 | 4 -- 46 | 162 -- 204 | -10.9371 | 5.79130 | 5.53585 |
| CD630_Cdi1_8 | CD630_SQ1642 | 277 -- 330 | 151 -- 190 | -10.9331 | 5.79994 | 8.65138 |
| CD630_SQ1642 | CD630_Cdi1_8 | 151 -- 190 | 277 -- 330 | -10.9331 | 8.65138 | 5.79994 |
| CD630_n00240 | CD630_s0400 | 5 -- 23 | 48 -- 78 | -10.9289 | 5.82709 | 4.51086 |
| CD630_s0400 | CD630_n00240 | 48 -- 78 | 5 -- 23 | -10.9289 | 4.51086 | 5.82709 |
| RCd5 | CD630_Cdi1_9 | 339 -- 370 | 1 -- 33 | -10.9225 | 6.52781 | 4.25622 |
| CD630_Cdi1_9 | RCd5 | 1 -- 33 | 339 -- 370 | -10.9225 | 4.25622 | 6.52781 |
| CD630_SQ1002 | CD630_s0631 | 86 -- 107 | 321 -- 340 | -10.9215 | 4.43255 | 4.95025 |
| CD630_s0631 | CD630_SQ1002 | 321 -- 340 | 86 -- 107 | -10.9215 | 4.95025 | 4.43255 |
| RCd9 | CD630_s0510 | 211 -- 232 | 33 -- 52 | -10.9185 | 8.96925 | 3.37631 |

| | | | | | | |
|---|---|---|---|---|---|---|
| CD630_Cdi1_8 | CD630_s0641 | 277 -- 332 | 256 -- 304 | -10.9177 | 5.79994 | 6.68432 |
| CD630_s0641 | CD630_Cdi1_8 | 256 -- 304 | 277 -- 332 | -10.9177 | 6.68432 | 5.79994 |
| CD630_s0400 | CD630_n00330 | 2 -- 49 | 38 -- 108 | -10.917 | 4.51086 | 6.46464 |
| CD630_n00330 | CD630_s0400 | 38 -- 108 | 2 -- 49 | -10.917 | 6.46464 | 4.51086 |
| CD630_n00850 | CD630_SQ1642 | 127 -- 187 | 83 -- 139 | -10.9169 | 5.53585 | 8.65138 |
| CD630_n00690 | CD630_n00690 | 90 -- 122 | 8 -- 41 | -10.9122 | 5.96526 | 5.96526 |
| CD630_n00470 | CD630_Cdi1_3 | 1 -- 68 | 307 -- 371 | -10.9086 | 3.52267 | 6.59405 |
| CD630_n00600 | CD630_n00330 | 13 -- 65 | 32 -- 72 | -10.9074 | 5.31146 | 6.46464 |
| CD630_n00330 | CD630_n00600 | 32 -- 72 | 13 -- 65 | -10.9074 | 6.46464 | 5.31146 |
| CD630_n00240 | CD630_Cdi1_11 | 3 -- 78 | 77 -- 151 | -10.9035 | 5.82709 | 5.77720 |
| CD630_n00560 | CD630_n00790 | 116 -- 189 | 250 -- 312 | -10.9006 | 5.82053 | 5.51210 |
| RCd5 | CD630_Cdi1_11 | 339 -- 370 | 1 -- 33 | -10.8944 | 6.52781 | 5.77720 |
| CD630_Cdi1_11 | RCd5 | 1 -- 33 | 339 -- 370 | -10.8944 | 5.77720 | 6.52781 |
| CD630_s0010 | CD630_Cdi1_5 | 63 -- 87 | 270 -- 295 | -10.8855 | 4.92433 | 4.83391 |
| CD630_Cdi1_5 | CD630_s0010 | 270 -- 295 | 63 -- 87 | -10.8855 | 4.83391 | 4.92433 |
| CD630_Cdi1_8 | CD630_SQ1002 | 219 -- 255 | 94 -- 130 | -10.8825 | 5.79994 | 4.43255 |
| CD630_SQ1002 | CD630_Cdi1_8 | 94 -- 130 | 219 -- 255 | -10.8825 | 4.43255 | 5.79994 |
| CD630_SQ1642 | RCd2 | 164 -- 193 | 173 -- 199 | -10.8772 | 8.65138 | 3.80984 |
| RCd2 | CD630_SQ1642 | 173 -- 199 | 164 -- 193 | -10.8772 | 3.80984 | 8.65138 |
| CD630_n00080 | CD630_n00690 | 77 -- 100 | 41 -- 73 | -10.8705 | 4.59330 | 5.96526 |
| CD630_n00690 | CD630_n00080 | 41 -- 73 | 77 -- 100 | -10.8705 | 5.96526 | 4.59330 |
| CD630_s0210 | CD630_s0370 | 10 -- 49 | 15 -- 55 | -10.8702 | 4.02122 | 10.21495 |
| CD630_Cdi1_3 | CD630_n00680 | 466 -- 490 | 351 -- 369 | -10.8683 | 6.59405 | 6.52781 |
| CD630_n00680 | CD630_Cdi1_3 | 351 -- 369 | 466 -- 490 | -10.8683 | 6.52781 | 6.59405 |
| CD630_n00690 | CD630_RNA_7 | 160 -- 182 | 89 -- 113 | -10.8674 | 5.96526 | 7.42886 |
| CD630_RNA_7 | CD630_n00690 | 89 -- 113 | 160 -- 182 | -10.8674 | 7.42886 | 5.96526 |
| CD630_n00440 | CD630_s0340 | 196 -- 240 | 4 -- 49 | -10.8609 | 4.85723 | 5.61064 |
| CD630_s0340 | CD630_n00440 | 4 -- 49 | 196 -- 240 | -10.8609 | 5.61064 | 4.85723 |
| CD630_s0390 | CD630_n00340 | 84 -- 91 | 125 -- 132 | -10.8561 | 4.73397 | 5.83478 |
| CD630_SQ1005 | CD630_n01010 | 77 -- 102 | 375 -- 405 | -10.8546 | 3.63582 | 4.59146 |
| CD630_n01010 | CD630_SQ1005 | 375 -- 405 | 77 -- 102 | -10.8546 | 4.59146 | 3.63582 |
| CD630_SQ1642 | CD630_Cdi2_3 | 174 -- 197 | 37 -- 60 | -10.8503 | 8.65138 | 3.99184 |
| CD630_Cdi2_3 | CD630_SQ1642 | 37 -- 60 | 174 -- 197 | -10.8503 | 3.99184 | 8.65138 |
| CD630_n00600 | CD630_n00440 | 67 -- 77 | 65 -- 75 | -10.8383 | 5.31146 | 4.85723 |
| CD630_n00650 | CD630_n00980 | 42 -- 80 | 317 -- 355 | -10.8347 | 5.32127 | 5.31318 |
| CD630_n00380 | CD630_Cdi1_10 | 889 -- 910 | 128 -- 147 | -10.8314 | 4.27916 | 8.60766 |
| CD630_Cdi1_10 | CD630_n00380 | 128 -- 147 | 889 -- 910 | -10.8314 | 8.60766 | 4.27916 |
| CD630_Cdi1_10 | CD630_n00990 | 128 -- 147 | 978 -- 999 | -10.8314 | 8.60766 | 3.72051 |
| CD630_n00990 | CD630_Cdi1_10 | 978 -- 999 | 128 -- 147 | -10.8314 | 3.72051 | 8.60766 |
| CD630_Cdi1_3 | CD630_s0641 | 487 -- 496 | 147 -- 156 | -10.8258 | 6.59405 | 6.68432 |
| CD630_s0641 | CD630_Cdi1_3 | 147 -- 156 | 487 -- 496 | -10.8258 | 6.68432 | 6.59405 |
| CD630_SQ1642 | CD630_s0591 | 40 -- 56 | 123 -- 137 | -10.8206 | 8.65138 | 6.84967 |
| CD630_s0591 | CD630_SQ1642 | 123 -- 137 | 40 -- 56 | -10.8206 | 6.84967 | 8.65138 |
| CD630_n00790 | CD630_n00560 | 107 -- 122 | 6 -- 21 | -10.8174 | 5.51210 | 5.82053 |
| CD630_n00460 | CD630_n00470 | 1006 -- 1033 | 56 -- 85 | -10.8169 | 5.55447 | 3.52267 |
| CD630_n00470 | CD630_n00460 | 56 -- 85 | 1006 -- 1033 | -10.8169 | 3.52267 | 5.55447 |
| CD630_n00340 | CD630_s0390 | 125 -- 133 | 83 -- 91 | -10.8115 | 5.83478 | 4.73397 |
| RCd1 | CD630_n01010 | 54 -- 93 | 1085 -- 1125 | -10.8089 | 4.09988 | 4.59146 |
| CD630_s0670 | CD630_n01010 | 150 -- 194 | 845 -- 886 | -10.7931 | 4.56362 | 4.59146 |
| CD630_n00440 | CD630_n00560 | 26 -- 43 | 164 -- 180 | -10.7861 | 4.85723 | 5.82053 |
| CD630_n00560 | CD630_n00440 | 164 -- 180 | 26 -- 43 | -10.7861 | 5.82053 | 4.85723 |
| CD630_Cdi1_11 | CD630_n00240 | 77 -- 96 | 60 -- 78 | -10.7777 | 5.77720 | 5.82709 |
| CD630_SQ367 | CD630_n01010 | 137 -- 172 | 841 -- 875 | -10.7757 | 4.08290 | 4.59146 |
| CD630_n01010 | CD630_SQ367 | 841 -- 875 | 137 -- 172 | -10.7757 | 4.59146 | 4.08290 |
| CD630_SQ808 | CD630_Cdi1_8 | 150 -- 169 | 113 -- 131 | -10.7739 | 6.85680 | 5.79994 |
| CD630_n00560 | CD630_s0631 | 533 -- 545 | 327 -- 336 | -10.7679 | 5.82053 | 4.95025 |
| CD630_s0631 | CD630_n00560 | 327 -- 336 | 533 -- 545 | -10.7679 | 4.95025 | 5.82053 |
| CD630_s0300 | CD630_s0190 | 39 -- 94 | 78 -- 139 | -10.7671 | 4.17452 | 4.98480 |
| CD630_n00370 | CD630_n00930 | 40 -- 67 | 209 -- 234 | -10.7658 | 7.05302 | 5.79130 |
| CD630_n00930 | CD630_n00370 | 209 -- 234 | 40 -- 67 | -10.7658 | 5.79130 | 7.05302 |
| CD630_n01010 | CD630_s0670 | 845 -- 882 | 156 -- 194 | -10.7618 | 4.59146 | 4.56362 |
| CD630_n00600 | CD630_SQ808 | 191 -- 215 | 207 -- 231 | -10.7603 | 5.31146 | 6.85680 |
| CD630_SQ808 | CD630_n00600 | 207 -- 231 | 191 -- 215 | -10.7603 | 6.85680 | 5.31146 |
| CD630_n00240 | CD630_n00380 | 4 -- 31 | 654 -- 691 | -10.7562 | 5.82709 | 4.27916 |
| CD630_n00240 | CD630_n00990 | 4 -- 31 | 743 -- 780 | -10.7562 | 5.82709 | 3.72051 |
| CD630_n00380 | CD630_n00240 | 654 -- 691 | 4 -- 31 | -10.7562 | 4.27916 | 5.82709 |
| CD630_n00990 | CD630_n00240 | 743 -- 780 | 4 -- 31 | -10.7562 | 3.72051 | 5.82709 |
| CD630_n00290 | CD630_n00380 | 10 -- 58 | 157 -- 207 | -10.7502 | 3.85358 | 4.27916 |
| CD630_n00290 | CD630_n00990 | 10 -- 58 | 246 -- 296 | -10.7502 | 3.85358 | 3.72051 |
| CD630_n00790 | CD630_s0591 | 201 -- 209 | 122 -- 130 | -10.7467 | 5.51210 | 6.84967 |
| CD630_s0591 | CD630_n00790 | 122 -- 130 | 201 -- 209 | -10.7467 | 6.84967 | 5.51210 |
| CD630_n00600 | CD630_s0470 | 121 -- 147 | 9 -- 35 | -10.7338 | 5.31146 | 5.35019 |
| CD630_s0470 | CD630_n00600 | 9 -- 35 | 121 -- 147 | -10.7338 | 5.35019 | 5.31146 |
| CD630_s0190 | RCd8 | 167 -- 177 | 197 -- 207 | -10.7327 | 4.98480 | 6.97567 |
| RCd8 | CD630_s0190 | 197 -- 207 | 167 -- 177 | -10.7327 | 6.97567 | 4.98480 |
| CD630_n00290 | CD630_s0510 | 2 -- 44 | 8 -- 55 | -10.7302 | 3.85358 | 3.37631 |
| CD630_SQ173 | CD630_n00510 | 43 -- 69 | 579 -- 604 | -10.7234 | 6.59405 | 3.92762 |
| CD630_n00510 | CD630_SQ173 | 579 -- 604 | 43 -- 69 | -10.7234 | 3.92762 | 6.59405 |
| CD630_n00440 | CD630_s0641 | 64 -- 72 | 97 -- 105 | -10.7206 | 4.85723 | 6.68432 |
| CD630_s0641 | CD630_n00440 | 97 -- 105 | 64 -- 72 | -10.7206 | 6.68432 | 4.85723 |
| CD630_s0281 | CD630_n00790 | 12 -- 45 | 883 -- 912 | -10.7178 | 8.30299 | 5.51210 |
| CD630_n00790 | CD630_s0281 | 883 -- 912 | 12 -- 45 | -10.7178 | 5.51210 | 8.30299 |
| CD630_s0400 | CD630_SQ1642 | 46 -- 153 | 86 -- 199 | -10.6998 | 4.51086 | 8.65138 |
| CD630_s0390 | CD630_Cdi1_8 | 85 -- 98 | 288 -- 301 | -10.6972 | 4.73397 | 5.79994 |
| CD630_Cdi1_8 | CD630_s0390 | 288 -- 301 | 85 -- 98 | -10.6972 | 5.79994 | 4.73397 |
| CD630_n00590 | CD630_n00380 | 53 -- 93 | 426 -- 467 | -10.694 | 4.28395 | 4.27916 |
| CD630_n00590 | CD630_n00990 | 53 -- 93 | 515 -- 556 | -10.694 | 4.28395 | 3.72051 |
| CD630_s0400 | CD630_s0631 | 14 -- 32 | 318 -- 335 | -10.6931 | 4.51086 | 4.95025 |
| CD630_s0631 | CD630_s0400 | 318 -- 335 | 14 -- 32 | -10.6931 | 4.95025 | 4.51086 |
| CD630_n00690 | CD630_n00600 | 11 -- 60 | 84 -- 119 | -10.6879 | 5.96526 | 5.31146 |
| CD630_s0190 | CD630_n00460 | 40 -- 88 | 202 -- 249 | -10.6827 | 4.98480 | 5.55447 |
| CD630_n00460 | CD630_s0190 | 202 -- 249 | 40 -- 88 | -10.6827 | 5.55447 | 4.98480 |
| CD630_n00380 | CD630_RNA_7 | 195 -- 211 | 96 -- 113 | -10.6815 | 4.27916 | 7.42886 |
| CD630_RNA_7 | CD630_n00380 | 96 -- 113 | 195 -- 211 | -10.6815 | 7.42886 | 4.27916 |
| CD630_RNA_7 | CD630_n00990 | 96 -- 113 | 284 -- 300 | -10.6815 | 7.42886 | 3.72051 |
| CD630_n00990 | CD630_RNA_7 | 284 -- 300 | 96 -- 113 | -10.6815 | 3.72051 | 7.42886 |
| CD630_n00560 | CD630_SQ1642 | 317 -- 360 | 142 -- 192 | -10.6769 | 5.82053 | 8.65138 |
| CD630_SQ1642 | CD630_n00560 | 142 -- 192 | 317 -- 360 | -10.6769 | 8.65138 | 5.82053 |
| RCd8 | CD630_Cdi1_5 | 86 -- 102 | 106 -- 122 | -10.6761 | 6.97567 | 4.83391 |
| RCd8 | CD630_n00390 | 86 -- 102 | 106 -- 122 | -10.6761 | 6.97567 | 4.83391 |
| CD630_Cdi1_5 | RCd8 | 106 -- 122 | 86 -- 102 | -10.6761 | 4.83391 | 6.97567 |
| CD630_n00390 | RCd8 | 106 -- 122 | 86 -- 102 | -10.6761 | 4.83391 | 6.97567 |
| CD630_s0641 | CD630_Cdi1_10 | 244 -- 286 | 16 -- 61 | -10.6755 | 6.68432 | 8.60766 |
| CD630_n00620 | RCd9 | 52 -- 96 | 159 -- 197 | -10.6634 | 5.82838 | 8.96925 |
| RCd9 | CD630_n00620 | 159 -- 197 | 52 -- 96 | -10.6634 | 8.96925 | 5.82838 |
| CD630_SQ1642 | CD630_s0400 | 86 -- 99 | 137 -- 153 | -10.663 | 8.65138 | 4.51086 |
| CD630_s0310 | CD630_SQ1642 | 1 -- 12 | 155 -- 166 | -10.6571 | 4.17452 | 8.65138 |
| CD630_SQ1642 | CD630_s0310 | 155 -- 166 | 1 -- 12 | -10.6571 | 8.65138 | 4.17452 |
| CD630_n00460 | RCd1 | 103 -- 148 | 47 -- 90 | -10.6466 | 5.55447 | 4.09988 |
| RCd1 | CD630_n00460 | 47 -- 90 | 103 -- 148 | -10.6466 | 4.09988 | 5.55447 |
| CD630_s0340 | CD630_Cdi1_4 | 1 -- 14 | 458 -- 471 | -10.6443 | 5.61064 | 5.31318 |
| CD630_Cdi1_4 | CD630_s0340 | 458 -- 471 | 1 -- 14 | -10.6443 | 5.31318 | 5.61064 |
| RCd8 | CD630_Cdi1_4 | 86 -- 102 | 113 -- 129 | -10.6431 | 6.97567 | 5.31318 |
| RCd8 | CD630_n00980 | 86 -- 102 | 113 -- 129 | -10.6431 | 6.97567 | 5.31318 |
| CD630_Cdi1_4 | RCd8 | 113 -- 129 | 86 -- 102 | -10.6431 | 5.31318 | 6.97567 |
| CD630_n00980 | RCd8 | 113 -- 129 | 86 -- 102 | -10.6431 | 5.31318 | 6.97567 |
| CD630_n00080 | CD630_Cdi2_3 | 85 -- 93 | 55 -- 63 | -10.6431 | 4.59330 | 3.99184 |
| CD630_Cdi1_3 | RCd5 | 466 -- 490 | 351 -- 369 | -10.6426 | 6.59405 | 6.52781 |
| RCd5 | CD630_Cdi1_3 | 351 -- 369 | 466 -- 490 | -10.6426 | 6.52781 | 6.59405 |
| CD630_n01010 | CD630_s0631 | 794 -- 869 | 1 -- 55 | -10.6404 | 4.59146 | 4.95025 |
| CD630_SQ408 | CD630_n00790 | 74 -- 114 | 870 -- 916 | -10.6374 | 7.27863 | 5.51210 |
| CD630_n00790 | CD630_SQ408 | 870 -- 916 | 74 -- 114 | -10.6374 | 5.51210 | 7.27863 |
| CD630_n00600 | CD630_Cdi2_2 | 796 -- 834 | 11 -- 48 | -10.6349 | 5.31146 | 6.72203 |
| CD630_n00600 | CD630_Cdi2_2 | 796 -- 834 | 11 -- 48 | -10.6349 | 5.31146 | 6.72203 |
| CD630_Cdi2_2 | CD630_n00600 | 11 -- 48 | 796 -- 834 | -10.6349 | 6.72203 | 5.31146 |
| CD630_Cdi2_2 | CD630_n00600 | 11 -- 48 | 796 -- 834 | -10.6349 | 6.72203 | 5.31146 |
| CD630_s0641 | CD630_Cdi2_3 | 177 -- 195 | 7 -- 25 | -10.6283 | 6.68432 | 3.99184 |
| CD630_s0010 | CD630_SQ1642 | 156 -- 189 | 65 -- 100 | -10.6276 | 4.92433 | 8.65138 |
| CD630_SQ1642 | CD630_s0010 | 65 -- 100 | 156 -- 189 | -10.6276 | 8.65138 | 4.92433 |
| CD630_s0510 | RCd9 | 38 -- 51 | 212 -- 225 | -10.6243 | 3.37631 | 8.96925 |
| RCd5 | CD630_n00930 | 401 -- 425 | 324 -- 340 | -10.6196 | 6.52781 | 5.79130 |
| CD630_n00930 | RCd5 | 324 -- 340 | 401 -- 425 | -10.6196 | 5.79130 | 6.52781 |
| CD630_n01010 | CD630_n00080 | 1213 -- 1297 | 1 -- 75 | -10.618 | 4.59146 | 4.59330 |
| RCd9 | CD630_s0642 | 14 -- 33 | 144 -- 161 | -10.6133 | 8.96925 | 6.00600 |
| CD630_n00850 | CD630_SQ1002 | 380 -- 401 | 121 -- 148 | -10.612 | 5.53585 | 4.43255 |
| CD630_SQ1002 | CD630_n00850 | 121 -- 148 | 380 -- 401 | -10.612 | 4.43255 | 5.53585 |
| CD630_n00790 | CD630_s0320 | 154 -- 223 | 7 -- 85 | -10.612 | 5.51210 | 8.15991 |
| CD630_n00380 | CD630_Cdi1_11 | 889 -- 910 | 128 -- 147 | -10.611 | 4.27916 | 5.77720 |
| CD630_Cdi1_11 | CD630_n00380 | 128 -- 147 | 889 -- 910 | -10.611 | 5.77720 | 4.27916 |
| CD630_Cdi1_11 | CD630_n00990 | 128 -- 147 | 978 -- 999 | -10.611 | 5.77720 | 3.72051 |
| CD630_n00990 | CD630_Cdi1_11 | 978 -- 999 | 128 -- 147 | -10.611 | 3.72051 | 5.77720 |
| CD630_SQ173 | CD630_n00460 | 42 -- 93 | 483 -- 533 | -10.6088 | 6.59405 | 5.55447 |
| CD630_n00460 | CD630_SQ173 | 483 -- 533 | 42 -- 93 | -10.6088 | 5.55447 | 6.59405 |
| CD630_Cdi1_5 | CD630_Cdi1_5 | 482 -- 500 | 272 -- 290 | -10.6053 | 4.83391 | 4.83391 |
| CD630_s0641 | CD630_Cdi1_11 | 244 -- 286 | 16 -- 61 | -10.5973 | 6.68432 | 5.77720 |
| CD630_s0660 | CD630_RNA_5 | 218 -- 242 | 5 -- 30 | -10.5944 | 4.92199 | 4.62291 |
| CD630_s0190 | CD630_s0670 | 40 -- 49 | 55 -- 64 | -10.5883 | 4.98480 | 4.56362 |
| CD630_Cdi1_9 | CD630_s0631 | 41 -- 91 | 60 -- 112 | -10.584 | 4.25622 | 4.95025 |
| CD630_s0190 | CD630_s0510 | 72 -- 143 | 125 -- 207 | -10.5836 | 4.98480 | 3.37631 |
| CD630_s0010 | CD630_n01010 | 165 -- 184 | 592 -- 611 | -10.5811 | 4.92433 | 4.59146 |

| | | | | | | |
|---|---|---|---|---|---|---|
| CD630_n01010 | CD630_s0010 | 592 -- 611 | 165 -- 184 | -10.5811 | 4.59146 | 4.92433 |
| CD630_n00460 | CD630_s0641 | 377 -- 443 | 65 -- 134 | -10.5796 | 5.55447 | 6.68432 |
| CD630_s0280 | CD630_s0642 | 31 -- 62 | 293 -- 327 | -10.578 | 5.37459 | 6.00600 |
| CD630_s0642 | CD630_s0280 | 293 -- 327 | 31 -- 62 | -10.578 | 6.00600 | 5.37459 |
| CD630_s0400 | RCd9 | 122 -- 143 | 97 -- 118 | -10.5769 | 4.51086 | 8.96925 |
| CD630_n00560 | CD630_SQ1005 | 123 -- 139 | 188 -- 204 | -10.5766 | 5.82053 | 3.63582 |
| CD630_n00460 | CD630_Cdi2_2 | 556 -- 564 | 33 -- 41 | -10.5747 | 5.55447 | 6.72203 |
| CD630_n00460 | CD630_Cdi2_2 | 556 -- 564 | 33 -- 41 | -10.5747 | 5.55447 | 6.72203 |
| CD630_Cdi2_2 | CD630_n00460 | 33 -- 41 | 556 -- 564 | -10.5747 | 6.72203 | 5.55447 |
| CD630_Cdi2_2 | CD630_n00460 | 33 -- 41 | 556 -- 564 | -10.5747 | 6.72203 | 5.55447 |
| CD630_s0360 | CD630_s0642 | 5 -- 22 | 143 -- 161 | -10.5673 | 6.65922 | 6.00600 |
| CD630_n00560 | CD630_s0270 | 123 -- 139 | 213 -- 229 | -10.5612 | 5.82053 | 3.63582 |
| CD630_n00650 | CD630_n00560 | 51 -- 85 | 456 -- 494 | -10.5591 | 5.32127 | 5.82053 |
| CD630_s0281 | CD630_n00380 | 277 -- 297 | 900 -- 922 | -10.556 | 8.30299 | 4.27916 |
| CD630_s0281 | CD630_n00990 | 277 -- 297 | 989 -- 1011 | -10.556 | 8.30299 | 3.72051 |
| CD630_n00500 | CD630_s0642 | 88 -- 98 | 16 -- 26 | -10.5437 | 9.35359 | 6.00600 |
| CD630_s0642 | CD630_n00500 | 16 -- 26 | 88 -- 98 | -10.5437 | 6.00600 | 9.35359 |
| CD630_s0340 | CD630_Cdi1_5 | 1 -- 14 | 451 -- 464 | -10.5405 | 5.61064 | 4.83391 |
| CD630_Cdi1_5 | CD630_s0340 | 451 -- 464 | 1 -- 14 | -10.5405 | 4.83391 | 5.61064 |
| CD630_Cdi1_10 | CD630_n00600 | 127 -- 154 | 351 -- 378 | -10.5373 | 8.60766 | 5.31146 |
| CD630_n00560 | CD630_RNA_7 | 158 -- 174 | 96 -- 113 | -10.5288 | 5.82053 | 7.42886 |
| CD630_RNA_7 | CD630_n00560 | 96 -- 113 | 158 -- 174 | -10.5288 | 7.42886 | 5.82053 |
| CD630_SQ1002 | CD630_s0450 | 184 -- 227 | 135 -- 188 | -10.5287 | 4.43255 | 5.67265 |
| CD630_s0450 | CD630_SQ1002 | 135 -- 188 | 184 -- 227 | -10.5287 | 5.67265 | 4.43255 |
| CD630_s0400 | CD630_s0320 | 165 -- 202 | 12 -- 45 | -10.5266 | 4.51086 | 8.15991 |
| CD630_n00440 | CD630_Cdi1_4 | 13 -- 38 | 385 -- 410 | -10.5239 | 4.85723 | 5.31318 |
| CD630_Cdi1_4 | CD630_n00440 | 385 -- 410 | 13 -- 38 | -10.5239 | 5.31318 | 4.85723 |
| CD630_n00330 | CD630_n00790 | 89 -- 112 | 452 -- 475 | -10.5159 | 6.46464 | 5.51210 |
| CD630_n00790 | CD630_n00330 | 452 -- 475 | 89 -- 112 | -10.5159 | 5.51210 | 6.46464 |
| CD630_s0320 | CD630_s0400 | 12 -- 54 | 161 -- 202 | -10.5123 | 8.15991 | 4.51086 |
| CD630_s0190 | CD630_s0360 | 120 -- 142 | 1 -- 23 | -10.5099 | 4.98480 | 6.65922 |
| CD630_s0360 | CD630_s0190 | 1 -- 23 | 120 -- 142 | -10.5099 | 6.65922 | 4.98480 |
| CD630_SQ1641 | CD630_n00790 | 15 -- 33 | 913 -- 934 | -10.5082 | 8.65138 | 5.51210 |
| CD630_n00790 | CD630_SQ1641 | 913 -- 934 | 15 -- 33 | -10.5082 | 5.51210 | 8.65138 |
| CD630_s0631 | CD630_n01010 | 320 -- 336 | 601 -- 616 | -10.5049 | 4.95025 | 4.59146 |
| CD630_SQ1656 | CD630_SQ1656 | 223 -- 234 | 223 -- 234 | -10.5025 | 4.25622 | 4.25622 |
| CD630_n00080 | CD630_n01010 | 49 -- 64 | 5 -- 19 | -10.4944 | 4.59330 | 4.59146 |
| CD630_n00620 | CD630_s0400 | 65 -- 121 | 6 -- 57 | -10.4939 | 5.82838 | 4.51086 |
| CD630_s0400 | CD630_n00620 | 6 -- 57 | 65 -- 121 | -10.4939 | 4.51086 | 5.82838 |
| CD630_SQ2429 | CD630_n00790 | 23 -- 65 | 590 -- 627 | -10.4911 | 5.77720 | 5.51210 |
| CD630_n00790 | CD630_SQ2429 | 590 -- 627 | 23 -- 65 | -10.4911 | 5.51210 | 5.77720 |
| CD630_s0670 | CD630_n00330 | 20 -- 36 | 38 -- 55 | -10.4875 | 4.56362 | 6.46464 |
| CD630_n00380 | CD630_s0281 | 900 -- 918 | 269 -- 297 | -10.4826 | 4.27916 | 8.30299 |
| CD630_n00990 | CD630_s0281 | 989 -- 1007 | 269 -- 297 | -10.4826 | 3.72051 | 8.30299 |
| CD630_n00240 | CD630_n01010 | 4 -- 15 | 588 -- 599 | -10.4814 | 5.82709 | 4.59146 |
| CD630_n01010 | CD630_n00240 | 588 -- 599 | 4 -- 15 | -10.4814 | 4.59146 | 5.82709 |
| CD630_SQ367 | CD630_s0641 | 171 -- 193 | 235 -- 256 | -10.4764 | 4.08290 | 6.68432 |
| CD630_n00330 | CD630_s0670 | 38 -- 52 | 22 -- 36 | -10.4705 | 6.46464 | 4.56362 |
| CD630_n00510 | CD630_s0642 | 444 -- 462 | 190 -- 208 | -10.4678 | 3.92762 | 6.00600 |
| CD630_s0642 | CD630_n00510 | 190 -- 208 | 444 -- 462 | -10.4678 | 6.00600 | 3.92762 |
| CD630_s0300 | CD630_n00150 | 4 -- 17 | 51 -- 64 | -10.4669 | 4.17452 | 8.81823 |
| CD630_n00150 | CD630_s0300 | 51 -- 64 | 4 -- 17 | -10.4669 | 8.81823 | 4.17452 |
| CD630_n00560 | CD630_s0510 | 664 -- 710 | 9 -- 58 | -10.4647 | 5.82053 | 3.37631 |
| CD630_s0320 | CD630_s0450 | 17 -- 38 | 183 -- 203 | -10.4616 | 8.15991 | 5.67265 |
| CD630_s0450 | CD630_s0320 | 183 -- 203 | 17 -- 38 | -10.4616 | 5.67265 | 8.15991 |
| CD630_s0460 | CD630_n00930 | 4 -- 19 | 334 -- 350 | -10.4594 | 5.74527 | 5.79130 |
| CD630_s0300 | RCd9 | 2 -- 52 | 253 -- 296 | -10.4588 | 4.17452 | 8.96925 |
| RCd9 | CD630_s0300 | 253 -- 296 | 2 -- 52 | -10.4588 | 8.96925 | 4.17452 |
| CD630_n00390 | CD630_s0500 | 346 -- 354 | 83 -- 91 | -10.4582 | 4.83391 | 3.25870 |
| CD630_s0500 | CD630_n00390 | 83 -- 91 | 346 -- 354 | -10.4582 | 3.25870 | 4.83391 |
| CD630_s0270 | CD630_s0330 | 10 -- 19 | 25 -- 34 | -10.4572 | 3.63582 | 4.37658 |
| CD630_s0330 | CD630_s0270 | 25 -- 34 | 10 -- 19 | -10.4572 | 4.37658 | 3.63582 |
| CD630_Cdi1_10 | CD630_s0210 | 104 -- 138 | 21 -- 54 | -10.4556 | 8.60766 | 4.02122 |
| CD630_n00380 | CD630_Cdi1_3 | 156 -- 187 | 179 -- 216 | -10.4412 | 4.27916 | 6.59405 |
| CD630_n00990 | CD630_Cdi1_3 | 245 -- 276 | 179 -- 216 | -10.4412 | 3.72051 | 6.59405 |
| CD630_SQ327 | CD630_Cdi1_4 | 191 -- 212 | 452 -- 466 | -10.4379 | 4.62291 | 5.31318 |
| CD630_Cdi1_4 | CD630_SQ327 | 452 -- 466 | 191 -- 212 | -10.4379 | 5.31318 | 4.62291 |
| CD630_Cdi1_3 | CD630_s0300 | 158 -- 179 | 157 -- 177 | -10.4369 | 6.59405 | 4.17452 |
| CD630_s0300 | CD630_Cdi1_3 | 157 -- 177 | 158 -- 179 | -10.4369 | 4.17452 | 6.59405 |
| CD630_s0340 | CD630_n00560 | 135 -- 167 | 106 -- 134 | -10.4365 | 5.61064 | 5.82053 |
| CD630_s0281 | CD630_s0510 | 97 -- 107 | 42 -- 51 | -10.4334 | 8.30299 | 3.37631 |
| CD630_s0510 | CD630_s0281 | 42 -- 51 | 97 -- 107 | -10.4334 | 3.37631 | 8.30299 |
| CD630_s0190 | CD630_Cdi1_8 | 171 -- 192 | 81 -- 104 | -10.4216 | 4.98480 | 5.79994 |
| CD630_Cdi1_8 | CD630_s0190 | 81 -- 104 | 171 -- 192 | -10.4216 | 5.79994 | 4.98480 |
| CD630_s0320 | CD630_n00290 | 6 -- 20 | 14 -- 28 | -10.4137 | 8.15991 | 3.85358 |
| CD630_n00290 | CD630_s0320 | 14 -- 28 | 6 -- 20 | -10.4137 | 3.85358 | 8.15991 |
| CD630_n00460 | CD630_n00930 | 409 -- 434 | 109 -- 135 | -10.4106 | 5.55447 | 5.79130 |
| CD630_n00930 | CD630_n00460 | 109 -- 135 | 409 -- 434 | -10.4106 | 5.79130 | 5.55447 |
| CD630_s0642 | CD630_s0310 | 119 -- 179 | 94 -- 150 | -10.4048 | 6.00600 | 4.17452 |
| CD630_Cdi1_10 | CD630_n01010 | 142 -- 148 | 975 -- 981 | -10.4036 | 8.60766 | 4.59146 |
| CD630_n01010 | CD630_Cdi1_10 | 975 -- 981 | 142 -- 148 | -10.4036 | 4.59146 | 8.60766 |
| CD630_n00980 | CD630_s0500 | 353 -- 361 | 83 -- 91 | -10.4015 | 5.31318 | 3.25870 |
| CD630_s0500 | CD630_n00980 | 83 -- 91 | 353 -- 361 | -10.4015 | 3.25870 | 5.31318 |
| CD630_n00930 | CD630_s0281 | 23 -- 65 | 23 -- 54 | -10.399 | 5.79130 | 8.30299 |
| CD630_s0281 | CD630_n00930 | 23 -- 54 | 23 -- 65 | -10.3989 | 8.30299 | 5.79130 |
| CD630_RNA_5 | CD630_s0642 | 81 -- 104 | 310 -- 335 | -10.3988 | 4.62291 | 6.00600 |
| CD630_s0642 | CD630_RNA_5 | 310 -- 335 | 81 -- 104 | -10.3988 | 6.00600 | 4.62291 |
| CD630_s0281 | CD630_RNA_7 | 175 -- 182 | 94 -- 101 | -10.3922 | 8.30299 | 7.42886 |
| CD630_RNA_7 | CD630_s0281 | 94 -- 101 | 175 -- 182 | -10.3922 | 7.42886 | 8.30299 |
| CD630_n00290 | CD630_s0460 | 48 -- 58 | 65 -- 75 | -10.3921 | 3.85358 | 5.74527 |
| CD630_n01010 | RCd1 | 1391 -- 1419 | 8 -- 38 | -10.3866 | 4.59146 | 4.09988 |
| CD630_n00620 | CD630_s0631 | 68 -- 94 | 87 -- 113 | -10.3834 | 5.82838 | 4.95025 |
| CD630_s0631 | CD630_n00620 | 87 -- 113 | 68 -- 94 | -10.3834 | 4.95025 | 5.82838 |
| CD630_SQ1641 | CD630_n00850 | 101 -- 117 | 114 -- 130 | -10.377 | 8.65138 | 5.53585 |
| CD630_n00850 | CD630_SQ1641 | 114 -- 130 | 101 -- 117 | -10.377 | 5.53585 | 8.65138 |
| CD630_Cdi2_3 | CD630_s0642 | 53 -- 62 | 149 -- 158 | -10.3761 | 3.99184 | 6.00600 |
| CD630_s0642 | CD630_Cdi2_3 | 149 -- 158 | 53 -- 62 | -10.3761 | 6.00600 | 3.99184 |
| CD630_n00650 | CD630_n01010 | 46 -- 85 | 1092 -- 1130 | -10.375 | 5.32127 | 4.59146 |
| CD630_n01010 | CD630_n00650 | 1092 -- 1130 | 46 -- 85 | -10.375 | 4.59146 | 5.32127 |
| CD630_n00460 | CD630_s0340 | 103 -- 118 | 155 -- 170 | -10.3747 | 5.55447 | 5.61064 |
| CD630_s0340 | CD630_n00460 | 155 -- 170 | 103 -- 118 | -10.3747 | 5.61064 | 5.55447 |
| CD630_SQ1005 | CD630_n00080 | 6 -- 40 | 61 -- 96 | -10.3639 | 3.63582 | 4.59330 |
| CD630_n00080 | CD630_SQ1005 | 61 -- 96 | 6 -- 40 | -10.3639 | 4.59330 | 3.63582 |
| CD630_s0270 | CD630_Cdi1_4 | 2 -- 20 | 194 -- 213 | -10.3623 | 3.63582 | 5.31318 |
| CD630_s0270 | CD630_n00980 | 2 -- 20 | 194 -- 213 | -10.3623 | 3.63582 | 5.31318 |
| CD630_Cdi1_4 | CD630_s0270 | 194 -- 213 | 2 -- 20 | -10.3623 | 5.31318 | 3.63582 |
| CD630_n00980 | CD630_s0270 | 194 -- 213 | 2 -- 20 | -10.3623 | 5.31318 | 3.63582 |
| CD630_n00930 | CD630_n00510 | 213 -- 234 | 859 -- 881 | -10.353 | 5.79130 | 3.92762 |
| RCd9 | CD630_n00590 | 269 -- 297 | 72 -- 100 | -10.3449 | 8.96925 | 4.28395 |
| CD630_n00590 | RCd9 | 72 -- 100 | 269 -- 297 | -10.3449 | 4.28395 | 8.96925 |
| CD630_s0510 | CD630_n00510 | 29 -- 51 | 93 -- 114 | -10.3431 | 3.37631 | 3.92762 |
| CD630_n00650 | CD630_Cdi1_4 | 42 -- 80 | 317 -- 355 | -10.3404 | 5.32127 | 5.31318 |
| CD630_n00790 | CD630_SQ367 | 221 -- 247 | 126 -- 153 | -10.3372 | 5.51210 | 4.08290 |
| CD630_n00440 | CD630_n00510 | 12 -- 43 | 165 -- 191 | -10.3283 | 4.85723 | 3.92762 |
| CD630_SQ1642 | CD630_s0670 | 5 -- 15 | 39 -- 50 | -10.3268 | 8.65138 | 4.56362 |
| CD630_s0670 | CD630_SQ1642 | 39 -- 50 | 5 -- 15 | -10.3268 | 4.56362 | 8.65138 |
| CD630_Cdi2_3 | CD630_Cdi2_3 | 45 -- 187 | 45 -- 187 | -10.3253 | 3.99184 | 3.99184 |
| CD630_s0591 | CD630_n00690 | 31 -- 55 | 289 -- 315 | -10.3212 | 6.84967 | 5.96526 |
| CD630_SQ327 | CD630_Cdi1_5 | 191 -- 212 | 445 -- 459 | -10.3166 | 4.62291 | 4.83391 |
| CD630_Cdi1_5 | CD630_SQ327 | 445 -- 459 | 191 -- 212 | -10.3166 | 4.83391 | 4.62291 |
| CD630_n00560 | CD630_s0340 | 462 -- 480 | 52 -- 71 | -10.3155 | 5.82053 | 5.61064 |
| CD630_s0400 | CD630_s0642 | 21 -- 40 | 146 -- 164 | -10.3115 | 4.51086 | 6.00600 |
| CD630_s0642 | CD630_s0400 | 146 -- 164 | 21 -- 40 | -10.3115 | 6.00600 | 4.51086 |
| CD630_s0250 | CD630_s0631 | 77 -- 106 | 313 -- 340 | -10.3108 | 3.74616 | 4.95025 |
| CD630_s0631 | CD630_s0250 | 313 -- 340 | 77 -- 106 | -10.3108 | 4.95025 | 3.74616 |
| CD630_s0460 | CD630_n00680 | 28 -- 73 | 350 -- 394 | -10.3069 | 5.74527 | 6.52781 |
| CD630_s0641 | CD630_Cdi1_9 | 244 -- 286 | 16 -- 61 | -10.3013 | 6.68432 | 4.25622 |
| CD630_s0480 | RCd9 | 1 -- 37 | 264 -- 296 | -10.2957 | 4.87999 | 8.96925 |
| RCd9 | CD630_s0480 | 264 -- 296 | 1 -- 37 | -10.2957 | 8.96925 | 4.87999 |
| CD630_SQ367 | CD630_n00790 | 126 -- 157 | 217 -- 247 | -10.29 | 4.08290 | 5.51210 |
| CD630_SQ367 | CD630_n00380 | 74 -- 202 | 799 -- 917 | -10.2876 | 4.08290 | 4.27916 |
| CD630_SQ367 | CD630_n00990 | 74 -- 202 | 888 -- 1006 | -10.2876 | 4.08290 | 3.72051 |
| CD630_n00380 | CD630_SQ367 | 799 -- 917 | 74 -- 202 | -10.2876 | 4.27916 | 4.08290 |
| CD630_n00990 | CD630_SQ367 | 888 -- 1006 | 74 -- 202 | -10.2876 | 3.72051 | 4.08290 |
| RCd8 | CD630_s0631 | 197 -- 209 | 235 -- 247 | -10.285 | 6.97567 | 4.95025 |
| CD630_s0631 | RCd8 | 235 -- 247 | 197 -- 209 | -10.285 | 4.95025 | 6.97567 |
| CD630_n00600 | CD630_SQ1642 | 30 -- 139 | 82 -- 200 | -10.2796 | 5.31146 | 8.65138 |
| CD630_s0460 | CD630_n00790 | 27 -- 66 | 88 -- 133 | -10.2791 | 5.74527 | 5.51210 |
| CD630_Cdi1_3 | CD630_SQ1642 | 462 -- 497 | 155 -- 195 | -10.2765 | 6.59405 | 8.65138 |
| CD630_SQ1642 | CD630_Cdi1_3 | 155 -- 195 | 462 -- 497 | -10.2765 | 8.65138 | 6.59405 |
| CD630_n00690 | CD630_s0470 | 4 -- 60 | 2 -- 54 | -10.2765 | 5.96526 | 5.35019 |
| CD630_s0470 | CD630_n00690 | 2 -- 54 | 4 -- 60 | -10.2765 | 5.35019 | 5.96526 |
| CD630_n00240 | CD630_s0281 | 8 -- 28 | 26 -- 50 | -10.2756 | 5.82709 | 8.30299 |
| CD630_s0281 | CD630_n00240 | 26 -- 50 | 8 -- 28 | -10.2756 | 8.30299 | 5.82709 |
| CD630_s0320 | CD630_n01010 | 20 -- 35 | 1113 -- 1129 | -10.2704 | 8.15991 | 4.59146 |
| CD630_n01010 | CD630_s0320 | 1113 -- 1129 | 20 -- 35 | -10.2704 | 4.59146 | 8.15991 |
| CD630_n00460 | CD630_SQ808 | 876 -- 921 | 32 -- 70 | -10.2698 | 5.55447 | 6.85680 |
| CD630_n00380 | CD630_n00590 | 426 -- 453 | 68 -- 93 | -10.2619 | 4.27916 | 4.28395 |
| CD630_n00990 | CD630_n00590 | 515 -- 542 | 68 -- 93 | -10.2619 | 3.72051 | 4.28395 |
| CD630_n00460 | CD630_s0460 | 100 -- 125 | 143 -- 165 | -10.2592 | 5.55447 | 5.74527 |

| | | | | | | |
|---|---|---|---|---|---|---|
| CD630_SQ1642 | CD630_n00600 | 88 -- 151 | 125 -- 175 | -10.259 | 8.65138 | 5.31146 |
| CD630_n00510 | CD630_s0641 | 100 -- 111 | 428 -- 439 | -10.257 | 3.92762 | 6.68432 |
| CD630_s0641 | CD630_n00510 | 428 -- 439 | 100 -- 111 | -10.257 | 6.68432 | 3.92762 |
| CD630_RNA_5 | CD630_n00380 | 4 -- 36 | 877 -- 904 | -10.2455 | 4.62291 | 4.27916 |
| CD630_RNA_5 | CD630_n00990 | 4 -- 36 | 966 -- 993 | -10.2455 | 4.62291 | 3.72051 |
| CD630_n00380 | CD630_RNA_5 | 877 -- 904 | 4 -- 36 | -10.2455 | 4.27916 | 4.62291 |
| CD630_n00990 | CD630_RNA_5 | 966 -- 993 | 4 -- 36 | -10.2455 | 3.72051 | 4.62291 |
| CD630_SQ1641 | CD630_s0670 | 16 -- 28 | 89 -- 100 | -10.2438 | 8.65138 | 4.56362 |
| CD630_s0670 | CD630_SQ1641 | 89 -- 100 | 16 -- 28 | -10.2438 | 4.56362 | 8.65138 |
| CD630_s0300 | CD630_Cdi1_8 | 14 -- 152 | 40 -- 181 | -10.2382 | 4.17452 | 5.79994 |
| CD630_n00460 | CD630_n01060 | 1004 -- 1028 | 94 -- 116 | -10.2369 | 5.55447 | 3.97522 |
| CD630_n01060 | CD630_n00460 | 94 -- 116 | 1004 -- 1028 | -10.2369 | 3.97522 | 5.55447 |
| CD630_s0250 | CD630_Cdi1_5 | 8 -- 16 | 491 -- 499 | -10.236 | 3.74616 | 4.83391 |
| CD630_Cdi1_5 | CD630_s0250 | 491 -- 499 | 8 -- 16 | -10.236 | 4.83391 | 3.74616 |
| CD630_n00380 | CD630_s0210 | 430 -- 461 | 24 -- 55 | -10.2341 | 4.27916 | 4.02122 |
| CD630_s0210 | CD630_n00380 | 24 -- 55 | 430 -- 461 | -10.2341 | 4.02122 | 4.27916 |
| CD630_s0210 | CD630_n00990 | 24 -- 55 | 519 -- 550 | -10.2341 | 4.02122 | 3.72051 |
| CD630_n00990 | CD630_s0210 | 519 -- 550 | 24 -- 55 | -10.2341 | 3.72051 | 4.02122 |
| CD630_n00590 | CD630_n00330 | 19 -- 42 | 36 -- 55 | -10.2293 | 4.28395 | 6.46464 |
| CD630_n00330 | CD630_n00590 | 36 -- 56 | 18 -- 42 | -10.225 | 6.46464 | 4.28395 |
| CD630_Cdi1_10 | CD630_n00790 | 144 -- 155 | 875 -- 886 | -10.2245 | 8.60766 | 5.51210 |
| CD630_n00790 | CD630_Cdi1_10 | 875 -- 886 | 144 -- 155 | -10.2245 | 5.51210 | 8.60766 |
| CD630_n00680 | CD630_n00590 | 335 -- 364 | 56 -- 87 | -10.2187 | 6.52781 | 4.28395 |
| CD630_n00590 | CD630_n00680 | 56 -- 87 | 335 -- 364 | -10.2187 | 4.28395 | 6.52781 |
| CD630_n00220 | RCd9 | 3 -- 17 | 210 -- 223 | -10.2177 | 4.94050 | 8.96925 |
| RCd9 | CD630_n00220 | 210 -- 223 | 3 -- 17 | -10.2177 | 8.96925 | 4.94050 |
| CD630_Cdi1_11 | CD630_n01010 | 142 -- 148 | 975 -- 981 | -10.2128 | 5.77720 | 4.59146 |
| CD630_n01010 | CD630_Cdi1_11 | 975 -- 981 | 142 -- 148 | -10.2128 | 4.59146 | 5.77720 |
| RCd8 | CD630_s0510 | 197 -- 208 | 168 -- 179 | -10.2102 | 6.97567 | 3.37631 |
| CD630_s0510 | RCd8 | 168 -- 179 | 197 -- 208 | -10.2102 | 3.37631 | 6.97567 |
| CD630_n01060 | CD630_n00290 | 94 -- 116 | 38 -- 56 | -10.2087 | 3.97522 | 3.85358 |
| CD630_n00290 | CD630_n01060 | 38 -- 56 | 94 -- 116 | -10.2087 | 3.85358 | 3.97522 |
| CD630_s0590 | CD630_Cdi1_4 | 38 -- 106 | 406 -- 465 | -10.2037 | 8.13716 | 5.31318 |
| CD630_s0400 | CD630_n01010 | 167 -- 182 | 601 -- 615 | -10.2026 | 4.51086 | 4.59146 |
| CD630_s0210 | CD630_s0190 | 9 -- 101 | 62 -- 158 | -10.1946 | 4.02122 | 4.98480 |
| CD630_Cdi1_3 | CD630_SQ1038 | 160 -- 246 | 53 -- 124 | -10.1923 | 6.59405 | 3.49019 |
| CD630_n00380 | CD630_n00290 | 157 -- 182 | 35 -- 58 | -10.1905 | 4.27916 | 3.85358 |
| CD630_n00990 | CD630_n00290 | 246 -- 271 | 35 -- 58 | -10.1905 | 3.72051 | 3.85358 |
| CD630_s0641 | CD630_s0590 | 151 -- 162 | 3 -- 15 | -10.1894 | 6.68432 | 8.13716 |
| CD630_s0590 | CD630_s0641 | 3 -- 15 | 151 -- 162 | -10.1894 | 8.13716 | 6.68432 |
| CD630_Cdi2_2 | CD630_n00790 | 44 -- 61 | 465 -- 480 | -10.1868 | 6.72203 | 5.51210 |
| CD630_Cdi2_2 | CD630_n00790 | 44 -- 61 | 465 -- 480 | -10.1868 | 6.72203 | 5.51210 |
| CD630_s0190 | CD630_n00380 | 70 -- 91 | 889 -- 910 | -10.185 | 4.98480 | 4.27916 |
| CD630_s0190 | CD630_n00990 | 70 -- 91 | 978 -- 999 | -10.185 | 4.98480 | 3.72051 |
| CD630_n00380 | CD630_s0190 | 889 -- 910 | 70 -- 91 | -10.185 | 4.27916 | 4.98480 |
| CD630_n00990 | CD630_s0190 | 978 -- 999 | 70 -- 91 | -10.185 | 3.72051 | 4.98480 |
| CD630_s0190 | CD630_Cdi1_10 | 171 -- 192 | 80 -- 103 | -10.1845 | 4.98480 | 8.60766 |
| CD630_Cdi1_10 | CD630_s0190 | 80 -- 103 | 171 -- 192 | -10.1845 | 8.60766 | 4.98480 |
| CD630_s0340 | CD630_SQ808 | 54 -- 77 | 38 -- 61 | -10.1607 | 5.61064 | 6.85680 |
| CD630_SQ808 | CD630_s0340 | 38 -- 61 | 54 -- 77 | -10.1607 | 6.85680 | 5.61064 |
| CD630_SQ1002 | CD630_n00790 | 188 -- 199 | 32 -- 43 | -10.1529 | 4.43255 | 5.51210 |
| CD630_n00790 | CD630_SQ1002 | 32 -- 43 | 188 -- 199 | -10.1529 | 5.51210 | 4.43255 |
| CD630_n00850 | RCd9 | 141 -- 179 | 264 -- 296 | -10.1516 | 5.53585 | 8.96925 |
| RCd9 | CD630_n00850 | 264 -- 296 | 141 -- 179 | -10.1516 | 8.96925 | 5.53585 |
| CD630_s0480 | CD630_SQ1656 | 43 -- 52 | 57 -- 66 | -10.1513 | 4.87999 | 4.25622 |
| CD630_SQ1656 | CD630_s0480 | 57 -- 66 | 43 -- 52 | -10.1513 | 4.25622 | 4.87999 |
| CD630_n00590 | CD630_n00340 | 26 -- 73 | 5 -- 57 | -10.1491 | 4.28395 | 5.83478 |
| CD630_n00690 | CD630_s0591 | 289 -- 299 | 45 -- 55 | -10.1383 | 5.96526 | 6.84967 |
| CD630_n00680 | CD630_n00790 | 257 -- 295 | 530 -- 576 | -10.1329 | 6.52781 | 5.51210 |
| CD630_n00790 | CD630_n00680 | 530 -- 576 | 257 -- 295 | -10.1329 | 5.51210 | 6.52781 |
| RCd9 | CD630_n00460 | 93 -- 119 | 99 -- 148 | -10.1308 | 8.96925 | 5.55447 |
| CD630_s0400 | CD630_n00690 | 32 -- 71 | 274 -- 308 | -10.1295 | 4.51086 | 5.96526 |
| RCd8 | CD630_SQ808 | 50 -- 70 | 34 -- 56 | -10.1295 | 6.97567 | 6.85680 |
| CD630_SQ808 | RCd8 | 34 -- 56 | 50 -- 70 | -10.1295 | 6.85680 | 6.97567 |
| CD630_n00690 | CD630_s0400 | 274 -- 308 | 32 -- 71 | -10.1295 | 5.96526 | 4.51086 |
| CD630_n00930 | CD630_n01010 | 6 -- 28 | 859 -- 885 | -10.1281 | 5.79130 | 4.59146 |
| CD630_n01010 | CD630_n00930 | 859 -- 885 | 6 -- 28 | -10.1281 | 4.59146 | 5.79130 |
| CD630_n00380 | CD630_n01060 | 834 -- 891 | 49 -- 115 | -10.1208 | 4.27916 | 3.97522 |
| CD630_n01060 | CD630_n00380 | 49 -- 115 | 834 -- 891 | -10.1208 | 3.97522 | 4.27916 |
| CD630_n01060 | CD630_n00990 | 49 -- 115 | 923 -- 980 | -10.1208 | 3.97522 | 3.72051 |
| CD630_n00990 | CD630_n01060 | 923 -- 980 | 49 -- 115 | -10.1208 | 3.72051 | 3.97522 |
| CD630_s0470 | CD630_s0642 | 3 -- 34 | 33 -- 62 | -10.12 | 5.35019 | 6.00600 |
| CD630_n00460 | CD630_n00340 | 294 -- 315 | 36 -- 58 | -10.1186 | 5.55447 | 5.83478 |
| CD630_n00340 | CD630_n00460 | 36 -- 58 | 294 -- 315 | -10.1186 | 5.83478 | 5.55447 |
| CD630_n00330 | CD630_s0660 | 39 -- 47 | 202 -- 210 | -10.1154 | 6.46464 | 4.92199 |
| CD630_s0660 | CD630_n00330 | 202 -- 210 | 39 -- 47 | -10.1154 | 4.92199 | 6.46464 |
| CD630_SQ173 | CD630_n01010 | 105 -- 133 | 628 -- 654 | -10.1119 | 6.59405 | 4.59146 |
| CD630_n01010 | CD630_SQ173 | 628 -- 654 | 105 -- 133 | -10.1119 | 4.59146 | 6.59405 |
| CD630_s0300 | CD630_n00080 | 32 -- 52 | 60 -- 82 | -10.1087 | 4.17452 | 4.59330 |
| CD630_n00080 | CD630_s0300 | 60 -- 82 | 32 -- 52 | -10.1087 | 4.59330 | 4.17452 |
| CD630_n00470 | CD630_Cdi1_4 | 3 -- 24 | 108 -- 129 | -10.1086 | 3.52267 | 5.31318 |
| CD630_n00470 | CD630_n00980 | 3 -- 24 | 108 -- 129 | -10.1086 | 3.52267 | 5.31318 |
| CD630_SQ1002 | CD630_s0670 | 164 -- 171 | 30 -- 37 | -10.1049 | 4.43255 | 4.56362 |
| CD630_n00690 | CD630_s0510 | 201 -- 251 | 9 -- 51 | -10.1049 | 5.96526 | 3.37631 |
| CD630_s0670 | CD630_SQ1002 | 30 -- 37 | 164 -- 171 | -10.1049 | 4.56362 | 4.43255 |
| CD630_s0270 | RCd2 | 112 -- 127 | 161 -- 177 | -10.1046 | 3.63582 | 3.80984 |
| RCd2 | CD630_s0270 | 161 -- 177 | 112 -- 127 | -10.1046 | 3.80984 | 3.63582 |
| CD630_n00620 | RCd2 | 156 -- 173 | 13 -- 31 | -10.1006 | 5.82838 | 3.80984 |
| RCd2 | CD630_n00620 | 13 -- 31 | 156 -- 173 | -10.1006 | 3.80984 | 5.82838 |
| CD630_n00440 | CD630_n00930 | 213 -- 236 | 5 -- 40 | -10.0955 | 4.85723 | 5.79130 |
| CD630_n00930 | CD630_n00440 | 5 -- 40 | 213 -- 236 | -10.0955 | 5.79130 | 4.85723 |
| CD630_s0190 | CD630_s0370 | 191 -- 201 | 30 -- 40 | -10.0909 | 4.98480 | 10.21495 |
| CD630_s0370 | CD630_s0190 | 30 -- 40 | 191 -- 201 | -10.0909 | 10.21495 | 4.98480 |
| CD630_s0210 | CD630_SQ1642 | 199 -- 207 | 93 -- 101 | -10.0903 | 4.02122 | 8.65138 |
| CD630_SQ1642 | CD630_s0210 | 93 -- 101 | 199 -- 207 | -10.0903 | 8.65138 | 4.02122 |
| CD630_SQ327 | CD630_s0281 | 82 -- 124 | 155 -- 193 | -10.0871 | 4.62291 | 8.30299 |
| CD630_s0010 | CD630_s0460 | 62 -- 86 | 1 -- 21 | -10.0846 | 4.92433 | 5.74527 |
| RCd9 | CD630_s0400 | 135 -- 176 | 35 -- 75 | -10.0841 | 8.96925 | 4.51086 |
| CD630_n00500 | CD630_s0510 | 48 -- 62 | 3 -- 17 | -10.0801 | 9.35359 | 3.37631 |
| CD630_s0510 | CD630_n00500 | 3 -- 17 | 48 -- 62 | -10.0801 | 3.37631 | 9.35359 |
| CD630_n00600 | CD630_n01060 | 1118 -- 1145 | 88 -- 118 | -10.078 | 5.31146 | 3.97522 |
| CD630_n01060 | CD630_n00600 | 88 -- 118 | 1118 -- 1145 | -10.078 | 3.97522 | 5.31146 |
| CD630_Cdi1_9 | CD630_n01010 | 137 -- 146 | 845 -- 854 | -10.0766 | 4.25622 | 4.59146 |
| CD630_n01010 | CD630_Cdi1_9 | 845 -- 854 | 137 -- 146 | -10.0766 | 4.59146 | 4.25622 |
| CD630_s0590 | CD630_n00600 | 23 -- 79 | 791 -- 831 | -10.0737 | 8.13716 | 5.31146 |
| CD630_Cdi1_5 | CD630_s0450 | 434 -- 448 | 136 -- 150 | -10.0716 | 4.83391 | 5.67265 |
| CD630_s0450 | CD630_Cdi1_5 | 136 -- 150 | 434 -- 448 | -10.0716 | 5.67265 | 4.83391 |
| CD630_n00460 | CD630_n00560 | 9 -- 22 | 104 -- 117 | -10.0614 | 5.55447 | 5.82053 |
| CD630_SQ1005 | CD630_n00560 | 188 -- 206 | 123 -- 139 | -10.0603 | 3.63582 | 5.82053 |
| CD630_n00080 | CD630_n00650 | 14 -- 37 | 49 -- 69 | -10.0561 | 4.59330 | 5.32127 |
| CD630_s0250 | CD630_n01010 | 2 -- 16 | 453 -- 466 | -10.0499 | 3.74616 | 4.59146 |
| CD630_n01010 | CD630_s0250 | 453 -- 466 | 2 -- 16 | -10.0499 | 4.59146 | 3.74616 |
| CD630_SQ1656 | CD630_s0641 | 3 -- 61 | 103 -- 156 | -10.0468 | 4.25622 | 6.68432 |
| CD630_s0270 | CD630_n00560 | 213 -- 231 | 123 -- 139 | -10.0442 | 3.63582 | 5.82053 |
| CD630_Cdi1_8 | RCd8 | 60 -- 131 | 86 -- 157 | -10.0403 | 5.79994 | 6.97567 |
| CD630_s0330 | CD630_s0450 | 2 -- 44 | 35 -- 92 | -10.0386 | 4.37658 | 5.67265 |
| CD630_s0450 | CD630_s0330 | 35 -- 92 | 2 -- 44 | -10.0386 | 5.67265 | 4.37658 |
| CD630_s0400 | CD630_n00980 | 38 -- 85 | 280 -- 330 | -10.0367 | 4.51086 | 5.31318 |
| CD630_n00650 | CD630_n00690 | 48 -- 69 | 214 -- 235 | -10.036 | 5.32127 | 5.96526 |
| CD630_n00690 | CD630_n00650 | 214 -- 235 | 48 -- 69 | -10.036 | 5.96526 | 5.32127 |
| RCd1 | CD630_n00790 | 42 -- 83 | 113 -- 157 | -10.0355 | 4.09988 | 5.51210 |
| CD630_n00790 | RCd1 | 113 -- 157 | 42 -- 83 | -10.0355 | 5.51210 | 4.09988 |
| CD630_n00620 | CD630_Cdi1_5 | 11 -- 19 | 422 -- 430 | -10.0326 | 5.82838 | 4.83391 |
| CD630_Cdi1_5 | CD630_n00620 | 422 -- 430 | 11 -- 19 | -10.0326 | 4.83391 | 5.82838 |
| CD630_s0631 | CD630_Cdi1_9 | 60 -- 107 | 45 -- 91 | -10.032 | 4.95025 | 4.25622 |
| CD630_n00930 | CD630_s0642 | 174 -- 199 | 2 -- 28 | -10.0281 | 5.79130 | 6.00600 |
| CD630_s0642 | CD630_n00930 | 2 -- 28 | 174 -- 199 | -10.0281 | 6.00600 | 5.79130 |
| RCd9 | CD630_n00930 | 40 -- 67 | 209 -- 234 | -10.0279 | 8.96925 | 5.79130 |
| CD630_n00510 | CD630_n00620 | 96 -- 115 | 71 -- 90 | -10.0272 | 3.92762 | 5.82838 |
| CD630_n00620 | CD630_n00510 | 71 -- 90 | 96 -- 115 | -10.0272 | 5.82838 | 3.92762 |
| CD630_n00440 | CD630_n00620 | 2 -- 39 | 161 -- 199 | -10.0256 | 4.85723 | 5.82838 |
| CD630_n00620 | CD630_n00440 | 161 -- 199 | 2 -- 39 | -10.0256 | 5.82838 | 4.85723 |
| CD630_s0480 | CD630_n00790 | 8 -- 33 | 883 -- 913 | -10.0242 | 4.87999 | 5.51210 |
| CD630_n00790 | CD630_s0480 | 883 -- 913 | 8 -- 33 | -10.0242 | 5.51210 | 4.87999 |
| CD630_Cdi2_3 | CD630_s0670 | 174 -- 186 | 43 -- 54 | -10.0225 | 3.99184 | 4.56362 |
| CD630_s0670 | CD630_Cdi2_3 | 43 -- 54 | 174 -- 186 | -10.0225 | 4.56362 | 3.99184 |
| CD630_s0190 | CD630_SQ2429 | 171 -- 192 | 179 -- 202 | -10.0217 | 4.98480 | 5.77720 |
| CD630_SQ2429 | CD630_s0190 | 179 -- 202 | 171 -- 192 | -10.0217 | 5.77720 | 4.98480 |
| CD630_n00240 | CD630_Cdi1_8 | 3 -- 76 | 80 -- 153 | -10.0103 | 5.82709 | 5.79994 |
| CD630_Cdi1_3 | CD630_n00850 | 239 -- 275 | 112 -- 152 | -10.0083 | 6.59405 | 5.53585 |
| CD630_n00850 | CD630_Cdi1_3 | 112 -- 152 | 239 -- 275 | -10.0083 | 5.53585 | 6.59405 |
